# The Leipzig Catalogue of Vascular Plants (LCVP) – An improved taxonomic reference list for all known vascular plants

**DOI:** 10.1101/2020.05.08.077149

**Authors:** Martin Freiberg, Marten Winter, Alessandro Gentile, Alexander Zizka, Alexandra Nora Muellner-Riehl, Alexandra Weigelt, Christian Wirth

**Author notes:** corresponding author: Martin Freiberg.

## Abstract

The lack of comprehensive and standardized taxonomic reference information is an impediment for robust plant research, e.g. in systematics, biogeography or macroecology. Here we provide an updated and much improved reference list of 1,315,479 scientific plant taxa names for all described vascular plant taxa names globally. The Leipzig Catalogue of Vascular Plants (LCVP; version 1.0.2) contains 351.176 accepted species (plus 6.160 natural hybrids), within 13.422 genera, 561 families and 84 orders. The LCVP a) contains more information on the taxonomic status of global plant names than any other similar resource and b) significantly improves the reliability of our knowledge by e.g. resolving the taxonomic status of ∼184.000 taxa names compared to The Plant List, the up to date most commonly used plant name resource. We used ∼4500 publications, existing relevant databases and available studies on molecular phylogenetics to construct a robust reference backbone. For easy access and integration into automated data processing pipelines, we provide an ‘R’-package (*lcvplants*) with the LCVP.

## Background and summary

Due to substantial progress in the last decade in improving plant taxonomy with phylogenetic findings, an updated global taxonomic reference list was urgently required. To date, the most commonly used reference list of vascular plant taxa names is *The Plant List* (TPL^1^), hosted by the Royal Botanic Gardens, Kew. TPL contains 1,166,081 vascular plants names, including 308,407 accepted names, 304,419 of them angiosperms. ∼700,000 names of TPL are synonyms or other taxonomic ranks (subspecies, varieties, forms), including 227,025 unresolved names. The here presented Leipzig Catalogue of Vascular Plants (LCVP) updates significantly the global knowledge of plant names not only compared to TPL (see Table 1) and thus is a major improvement for global plant research. It is based on existing databases and an additional 4,500 publications (see Supplementary File 3 and 4 for a list of full literature references ordered by families – as plain text and as bibliography RIS file and File 5 for a list of abbreviated literature references ordered by individual taxa), which helped to clarify the status of plant names (i.e. accepted, synonymy, taxonomic placement; see Methods). In the end, 4059 publications provided relevant and sufficiently robust additional information, e.g. changes in taxa names and/or their status. A guiding principle during the compilation of the LCVP was to avoid polyphyletic genera, which are frequent in TPL, either by splitting genera (e.g. separating *Goeppertia* from *Calathea*) or fusing them (e.g. *Stapelia* and *Duvalia* in *Ceropegia*). However, we did not recombine any species name in the LCVP and in cases of unclear phylogenetic position of genera, we used the conservative (i.e. existing) name.

**Table 1:**
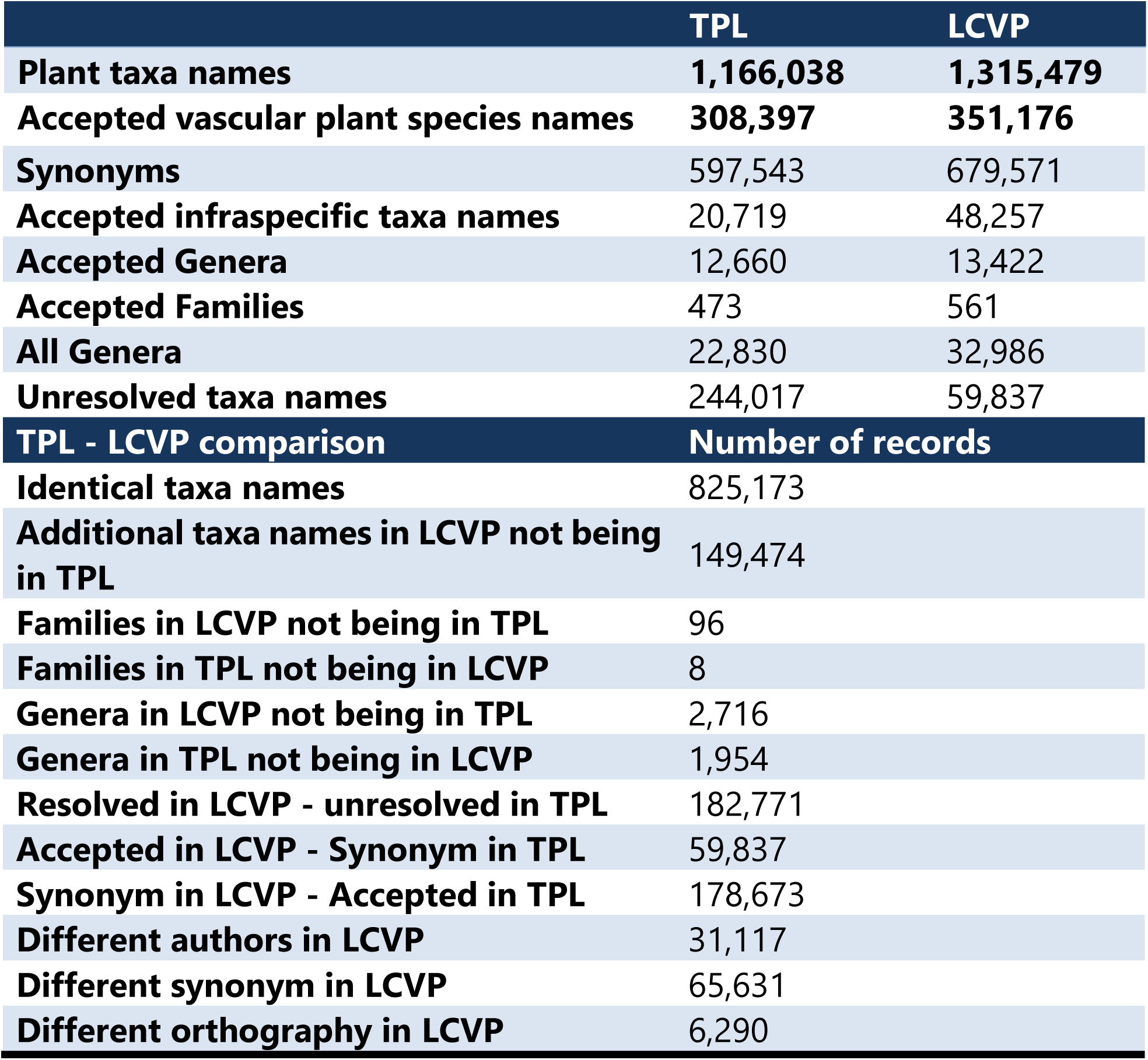
Summary of information content in the LCVP and taxa name differences between LCVP and TPL.

Taxonomists, ecologists and conservation biologists often work with many species (names) and cannot keep pace with the rapid progress in (plant) systematics, boosted by molecular phylogenetic methods^2^. These researchers often rely on taxonomic reference lists as tools to translate taxa names to accepted species names via accepted synonyms.

Comprehensive taxa lists, such as the LCVP, are essential to standardize taxonomic names in databases compiled from various sources, relying on a robust ‘translation’ of species names into one scheme. The TRY database of functional plant traits (TRY^3^; www.try-db.org) is one of the most prominent examples containing trait information for about 150,000 vascular plant species. Other global databases using plant name reference lists focus on plant co-occurrence patterns, such as sPlot containing about 1,1 million vegetation surveys (^4^∼55,000 species), or use any plant species occurrence information, such as the Global Biodiversity Information Facility (GBIF^5^: ∼315,000 vascular plant species; www.gbif.org), the Botanical Information and Ecology Network (BIEN^6^: ∼348,000). The Global Inventory of Floras and Traits (GIFT^7^: ∼268,000; http://gift.uni-goettingen.de/home) or the inventory of the Global Naturalized Alien Flora (GloNAF^8^∼14,000; glonaf.org) focus on plant distribution information from regional floras or floristic inventories.

Generally, such databases were compiled from heterogeneous data sources varying in time of publication and place of origin. The underlying sources may be primary or secondary literature-based using work of scientists with excellent to no plant taxonomic background, thus combining data with various degrees of complexities and uncertainties. The merging of these databases works via species identities and thus depends on the use of accepted species names. These databases typically tap phylogenetic information contained in taxonomic references lists via available tools supporting automated matching and error checking (i.e. taxon scrubbing). There is a variety of R packages (e.g. taxonstand^9^; taxize^10^; RBIEN^11^) or online tools (e.g. Global Name Resolver http://resolver.globalnames.org/ or the Taxonomic Name Resolution Service^12^ http://tnrs.iplantcollaborative.org/TNRSapp.html) supporting researchers to check their taxonomic information (see^13^ for a review on some of those tools). Most of these tools rely on TPL as part of their reference lists, which, however, has not been updated for a decade and originated in a time when phylogenetic information on many genera did not exist.

Global taxonomic name databases are useful in their own right, and jointly create synergies that have transformed ecology into a synthetic and global science, and can help identifying knowledge gaps^14^. For example, functional biogeography combines information on community composition, plant species distribution and functional traits of the component species to make inferences on determinants of global trait distribution^15^. While there is high potential for exciting research using most up-to-date taxonomic information, it can be only as good as the input data and the ability of the user to understand the advantage and shortcoming of the data coming from those resources. For example, missing taxonomic background often leads to neglecting the importance of citing authors of names and inevitably leads to inconsistencies when data from different sources are matched. LCVP shows that when matching plant taxa names without author names, results could have up to 10% mismatches (i.e. ∼10% of all LCVP plant taxa names are the same but ultimately refer to different accepted plant taxa).

## Methods

The creation of the LCVP involved three major steps. (1) We did a thorough search of available and relevant plant taxonomic databases (Supplementary File 1) to collate a raw data table of existing plant names (see Step 1: Producing the raw data table). This table included many contradictory opinions in taxonomic placement of species. (2) Based on additional information in ∼4500 publications and the reliability, timeliness and quality of relevant scientific evidences in this literature we decided for each name, whether that name is in LCVP accepted, synonymous or unresolved (see for more details Step 2: Decision making). Additionally, we harmonized and corrected taxa names orthographically. (3) We implemented the LCVP in an R package (LCVP) which is accessible under a MIT license from GitHub (https://github.com/idiv-biodiversity/LCVP) and will ensure a coherent versioning of the list and future updates. Furthermore, we provide a utility function to use LCVP for taxonomic name resolution (lcvplants), which is also available under the same license from GitHub (https://github.com/idiv-biodiversity/lcvplants).

### Step 1: Producing the raw data table

TPL provided the core of the basic raw data table for published vascular plant names, primarily supplemented by the International Plant Names Index (IPNI^16^, https://www.ipni.org/). IPNI provides a list of published names and their source, but does not provide any information on accepted or synonymous names. We used additional major and minor databases (^17-55^) which we have chosen based on their availability, on our expert judgement on comprehensiveness, and whether they contained information if taxa names are accepted or not (see Supplementary File 1 for a table of used databases). All additional names and potential synonyms found in those databases were incorporated in the raw data table.

### Step 2: Decision making

The extensive raw data table of more than two million entries of plant taxa names contained a high number of orthographic errors, inconsistencies and contradictory opinions concerning the status of the names. A rough guideline for the acceptance of names was a subjective assignment of quality and reliability to the source. Generally, changes were only applied when the authors of the respective publications were clearly suggesting those changes. We ascribed a higher reliability rank (e.g. for conflicting information) usually to the most recent publications. Additionally, when conflicting information appeared we usually used information from publications with a) a more thorough literature section and b) a more comprehensive synonymy history than to those without. A complete synonymy history should include and properly cite not only the latest accepted taxon, but also the depending taxonomic history of all names connected to this taxon (e.g. if it is a recombined taxon) with all homonymic (i.e. species epitheton is the same) and heteronymic (i.e. genus name is the same) synonyms. Since phylogenies based on morphological data alone are prone to homoplasy, only phylogenetic studies that made taxonomical decisions also based on molecular data were taken into account. We did not create new species name combinations. In case of conflicting evidence on the phylogenetic placement or species name, due to e.g. different methods to build phylogenetic trees, species names were marked “comb.ined.” following the basionym author.

The following examples illustrate how we treated name changes: The genus *Dracaena* and *Sansevieria* are closely related^56^, where *Sansevieria* seems to be clearly nested within *Dracaena*, but the differences between both genera are fluently. Lu *et al*.^56^ separated the Hawaiian species of *Dracaena* in a new genus *Chrysodracon*, but did not recombine *Sansevieria* with *Dracaena* yet. The presented argumentation and data in^56^ were thorough and comprehensive and thus we accepted the authors arguments, kept *Sansevieria* and *Dracaena* as distinct genera and separated the Hawaiian species of *Dracaena* in the new genus *Chrysodracon*. In another case Borchsenius et al.^57^ clearly showed that *Calathea* in the traditional description was polyphyletic. In order to keep *Ischnosiphon* and *Monotagma* as distinct genera, being the sister clade to a smaller *Calathea* clade including the type species, the larger clade of *Calathea* was put into the then resurrected genus *Goeppertia*. The argumentation and presentation in^57^ was robustly based on a molecular phylogeny producing well supported clades. As consequence we accepted the recombination of the much larger clade as suggested in^57^.

Supplementary Files 3 and 4 provide a complete list of all ∼4,500 literature references ordered by plant families that we used to decide upon species names to create LCVP. We focused on literature published from 1994 onwards, when molecular phylogenies became widespread (^58,59^). Supplementary File 5 provides a list directly matching >104.000 individual taxa and literature, used to inform the applied name changes for the respective taxa.

We also applied changes to the spelling of species names. Generally, we recommend to check the species names prior to automated list treatments, following the guidelines given in ^60^ and the rules of the current version of the International Code of Nomenclature for algae, fungi, and plants (Shenzhen Code^61^). We followed the Shenzen Code using standardized orthography of epitheta across genera and families, e.g. *warscewiczii* (neither *warscewitzii* nor *warszewiczii*). Only upper cases from ‘A’ to ‘Z’, lower cases from ‘a’ to ‘z’ and the hyphen ‘-’ should be used in the scientific names, special characters are not valid and to be avoided (Isoëtes-> Isoetes, Köberlinia -> Koeberlinia). Authors were given in their short form as provided by IPNI. For further standardization and easier use in automated workflows, we omitted spaces within author names (C. F. W. Meissn., C.F. W. Meissn., C. F.W. Meissn. C. F. W.Meissn. -> C.F.W.Meissn.; Balf. f.-> Balf.f.). We linked names published by two authors with the ‘&’ sign (e.g. *Primula minor* Balf.f. & Kingdon-Ward). Names published by three and more authors were restricted to the first authors followed by ‘& al.’ (e.g. *Limonium irtaense* P.P.Ferrer, A.Navarro, P.Pérez, R.Roselló, Rosselló, M.Rosato & E.Laguna -> *Limonium irtaense* P.P.Ferrer & al.). This refers to the recommendation of the Shenzhen Code, Art. 46 c. We tried to include only natural hybrids (i.e. no cultivars; based on expert judgement of LCVP authors) in the LCVP. Since hybrids were not the focus of the LCVP, we only marked them with ‘_x’, either following the genus name or the epitheton to recognize them as such, but we did not give any parent taxa information.

In most cases, we adopted the names used by the taxonomic expert (i.e. reference author who is usually a person with a publication record within a certain taxonomic group). However, there are many taxa belonging to genera or species which have not been phylogenetically analyzed yet. For those, we adapted the most frequently used taxon name from the recent literature (see Supplementary Files 3 -5). Despite a major effort, there are still names, which we could not resolve.

### Step 3: Implementation in R

Besides providing LCVP as downloadable tables, we also provide the LCVP as an R package (LCVP) for easy integration with analyses pipelines. We also provide a tool for fuzzy-matching-based taxonomic name resolution directly linked with LCVP (lcvplants). This fuzzy-matching algorithm is applied at species, infra-species and authority level of a plant taxon; it uses the ‘max.distance’ argument from the agrep() function’ to assess the comparison between the searched plant name and the closest plant name from the LCVP list (in terms of number of the same character and their order). The taxonomic names resolution is implemented in a user-friendly way, and can be done with few lines of code:

~~~
‘‘‘
# install LCVP and lcvplants from GitHub
install.packages(“devtools”)
library(devtools)
devtools::install_github(“idiv-biodiversity/LCVP”)
devtools::install_github(“idiv-biodiversity/lcvplants”)
# load the package
library(lcvplants)
# run analyses
LCVP(“Hibiscus vitifolius”)
‘‘‘
~~~

We provide a description of the fuzzy matching algorithm and its implementation in Supplementary File 2 and as detailed tutorial on how to use lcvplants online (https://idiv-biodiversity.github.io/lcvplants/).

### Data Records

LCVP contains 1,315,479 vascular plant taxa names. There are 351,176 accepted species names in the LCVP. The accepted species in LCVP belong to 13,422 genera, 561 families, and 84 orders, respectively. LCVP significantly reduced the number of unresolved plant taxa names by ∼184,000 to ∼ 60,000 (5%) taxa (see Table 1). The LCVP version 1.0.2 is available in both Microsoft Excel and text formats in the iDiv data portal (https://idata.idiv.de/ddm/Data/ShowData/1806). A developmental version of the LCVP and the lcvplants package are publicly available via GitHub (https://github.com/idiv-biodiversity/lcvplants). LCVP version 1.0.2 has a DOI (10.25829/idiv.1806-40-3009 via the iDiv data portal). We will constantly curate the LCVP and plan to release a new version once every second to third year.

### Technical Validation

We tested whether all synonyms lead to an accepted name or another synonym. One major issue with TPL is the high amount of unresolved taxa. A link to another name sometimes is another synonym leading to unresolved loops. LCVP only links to accepted names, not to the taxonomic predecessor. If taxon A is synonym to taxon B and it turned out, that taxon B is synonym to taxon C, the accepted name given for taxon A is taxon C, not B. We treated invalid names as synonyms and assigned them to their appropriate accepted name.

Most of the still unresolved species names in LCVP were originally published in the 19th century. There is a high probability that the majority of them are synonyms, e.g. because of historic transfer errors from one publication to the other. An extraordinarily high amount of unresolved taxa can be found in Asteraceae (*Hieracium* 5,800 names out of a total of 19,300 names are unresolved, *Senecio* 685 out of 6,680, *Cirsium* 357 out of 2,170), Rosaceae (*Rubus* 4,040 out of 10,200, *Rosa* 2,300 out of 5,970, *Prunus* 512 out of 2,070, *Potentilla* 728 out of 3,950, *Crataegus* 730 out of 2,700, *Pyrus* 379 out of 1,200), Salicaceae (*Salix* 619 out of 3,800), Araceae (*Anthurium* 585 out of 2,260), and Geraniaceae (*Pelargonium* 963 out of 1,840).

### Comparison to TPL

Due to the improved name resolution and increased name information in general in LCVP compared to TPL, any work flow including taxonomic harmonization of plant names, will very likely yield more robust and reliable results for e.g. species richness pattern, matches between different data sources. For an easier comparison and guidance for users on the differences between LCVP and TPL, LCVP includes information whether taxa name entries are identical, differ in the cross-reference to a synonym, differ only orthographically either by the name or the author, or whether a name is new in the LCVP and not present in TPL. This unique information makes it possible for the users of TPL to update their names according to the LCVP, because all differences are clearly stated in the column ‘status’ of the LCVP.

Kew Gardens’ research effort to standardize plant names recently focuses on their new flagship program, Plants of the World Online (POWO, http://www.plantsoftheworldonline.org/), which includes a not yet published new taxonomic reference backbone (Alan Paton from Kew Gardens, pers. comm.). Given that this is becoming the successor of TPL (see http://www.plantsoftheworldonline.org/about) we also compared the available POWO list with LCVP (POWO access date: November 2018; provided by Kew). With ∼335,000 accepted species names and ∼458,000 taxa of vascular plants marked as synonyms in POWO, LCVP contains also significantly more species name information than POWO (this comparison includes only vascular plants and excludes infra-specific taxa since LCVP covers only vascular plants and POWO does not include taxa below species level).

TPL and POWO cover all plants, LCVP only vascular plants. With the current information we have, LCVP contains more information about vascular plant taxa names (e.g. more resolved taxa, more accepted species, more synonyms) than TPL and POWO. A user is more likely to resolve a given vascular plant taxa name with LCVP than with the given versions of TPL and POWO. LCVP covers also infraspecific taxa names which are not covered in POWO yet. The information in LCVP to which genus a species belongs and/or thus which accepted name should be used, is based on taxonomic, but also on most recent phylogenetic (i.e. mainly genetic) information. TPL was not updated for many years, and is mainly based on taxonomic information (i.e. not molecular phylogenies). With respect to usability of LCVP, we do see advantages compared to POWO, which to our knowledge does not offer an R package nor any other functionality of (half)automatic name checking or any fuzzy name matching functions.

### Code Availability

The LCVP generally consists of (1) the LCVP itself available as R data package (version 1.0.2 as of April 2020) and as tab-delimited textfile file and (2) the R-package lcvplants. The static LCVP file and the lcvplants tools are publicly available either via the iDiv data repository: https://idata.idiv.de/ddm/Data/ShowData/1806 or as developemental versions via GitHub (https://github.com/idiv-biodiversity/LCVP, https://github.com/idiv-biodiversity/lcvplants). We plan to closely collaborate with plant synonymy services and tools like e.g. BIEN, GNR, R packages taxonstand and taxize, to include LCVP as reference option. Requests for integrating LCVP can be made via the projects GitHub (https://github.com/idiv-biodiversity/LCVP/issues)

## Supporting information

Supplementary file 1 - Major sources in compiling the raw-data list

Supplementary file 2 - lcvplants description

Supplementary file 3 - Literature used to compile LCVP ordered by plant families

Supplementary file 4 - Literature used to compile LCVP ordered by plant families 2

Supplementary file 5 - Literature used to compile LCVP ordered by plant families 3

## Acknowledgements

We thank thousands of experts working sometimes their whole life on plant taxonomy and systematics, creating invaluable information and knowledge the LCVP is also based on. Without this knowledge the LCVP would not exist. We also thank Anke Stein, Patrick Weigelt, Aldo Compagnoni, Jitendra Gaikwad, Jens Kattge & Ingolf Kühn for helpful comments on the draft and R package functionalities. MW, AZ & AG thank DFG for funding (via iDiv, FZT 118). ANMR thanks BMBF for funding (Grant no. 16GW0120K).

## Author contributions

MF compiled the LCVP. AG, MW & AZ designed the R packages. AG & AZ implemented the R functions based on discussions with MW & MF. MW & MF compiled the drafts of the data paper. All authors contributed to the writing of the manuscript.

