## Supplementary file 1 - Major sources in compiling the raw-data list for "The Leipzig Catalogue of Vascular Plants (LCVP) – An improved taxonomic reference list for all known vascular plants"

**Supplementary file 1**: Major sources used in compiling the raw-data list

| Source | Reference information if available | License information if available | Copyright information if available |
|---|---|---|---|
| Carnivorous Plant Database (http://www.omnisterra.com/bot/cp_home.cgi) | Carnivorous Plant Database: http://www.omnisterra.com/bot/cp_home.cgi, International Carnivorous Plant Society (2017) | | Dr. Barry Rice, Science Editor of Carnivorous Plant Newsletter |
| Co's Digital Flora of the Phillippines (http://www.philippineplants.org/) | Pelser, P.B., J.F. Barcelona & D.L. Nickrent (eds.). Co's Digital Flora of the Phillippines. 2011 onwards. | | Co's Digital Flora of the Philippines |
| Dipterocarpaceae Data Base (http://rbg-web2.rbge.org.uk/diptero/) | Mark Newman, (Supervisor), Edinburgh; Dipterocarpaceae Data Base at the RBGE PANDORA database system (2015) | | Copyright Royal Botanic Garden Edinburgh 1998 |
| Gentian Research Network (http://gentian.rutgers.edu/classNEW123.htm) | At 23rd March 2020 not available anymore | | |
| Gesneriaceae Research (http://botany.si.edu/gesneriaceae/checklist/result.cfm) | Not existent anymore | | |
| Global Compositae Checklist (https://compositae.landcareresearch.co.nz/) | Flann, C (ed) 2009+ Global Compositae Checklist. Accessed 2016-2019 | Use of these data allowable with due attribution [citation as mentioned] | |
| India Biodiversity Portal (https://indiabiodiversity.org/) | Vattakaven T, George R, Balasubramanian D, Réjou-Méchain M, Muthusankar G, Ramesh B, Prabhakar R (2016) India Biodiversity Portal: An integrated, interactive and participatory biodiversity informatics platform. Biodiversity Data Journal 4: e10279. https://doi.org/10.3897/BDJ.4.e10279 | | |
| International Plant Names Index | IPNI (2010). International Plant Names Index. Published on the Internet http://www.ipni.org, The Royal Botanic Gardens, Kew, Harvard University Herbaria & Libraries and Australian | | |

| | | | |
|---|---|---|---|
| | National Botanic Gardens. [Retrieved 1 April 2010]. | | |
| Parasitic Plant Classification (https://parasiticplants.siu.edu/ListParasites.html) | Daniel L. Nickrent, https://parasiticplants.siu.edu/ListParasites.htm, Department of Plant Biology, Southern Illinois University (2016) | | Copyright © 1997 Daniel L. Nickrent |
| Plants of the World online (http://www.plantsoftheworldonline.org/) | Plants of the World Online. Facilitated by the Royal Botanic Gardens, Kew. Published on the Internet; http://www.plantsoftheworldonline.org/ last time assessed 01.12.2019 | CC - BY | Copyright Board of Trustees of the Royal Botanic Gardens, Kew |
| SysTax (http://www.systax.org/) | SysTax – ein Datenbanksystem für Systematik und Taxonomie © Uni Ulm 2015 | | |
| The African Plant Database (Botanical Garden of Geneva, http://www.ville-ge.ch/musinfo/bd/cjb/africa/recherche.php?langue=an) | © 2012 Conservatoire et Jardin botaniques & South African National Biodiversity Institute | | |
| The BrassiBase (https://brassibase.cos.uni-heidelberg.de/) | Kiefer M, Schmickl R, German DA, Lysak M, Al-Shehbaz IA, Franzke A, Mummenhoff K, Stamatakis A, Koch MA. 2014. BrassiBase: Introdcution to a novel database on Brassicaceae evolution. Plant Cell Physiol., 55(1): e3, doi:10.1093/pcp/pct158.<br><br>Koch MA, German DA, Kiefer M, Franzke A. 2018. Database taxonomics as key to modern plant biology. Trends Plant Sci. 23(1): 4–6. DOI: 10.1016/j.tplants.2017.10.005 | | Usually webpages may be freely distributed and copied. However, it is requested that in any subsequent use of this work, BrassiBase be given appropriate acknowledgment and cited. |
| The Catalogue of Life (http://www.catalogueoflife.org/) | Roskov Y., Ower G., Orrell T., Nicolson D., Bailly N., Kirk P.M., Bourgoin T., DeWalt R.E., Decock W., Nieukerken E. van, Zarucchi J., Penev L., eds. (2019). Species 2000 & ITIS Catalogue of Life, 2019 Annual Checklist. Digital resource at | Use of the content (such as the classification, synonymic species checklist, and scientific names) for publications and databases by individuals and organizations for not-for-profit usage is | This online database is copyrighted by Species 2000 on behalf of the Catalogue of Life partners |

| | | |
|---|---|---|
| | www.catalogueoflife.org/annual-checklist/2019. Species 2000: Naturalis, Leiden, the Netherlands. ISSN 2405-884X<br><br>Hassler M. (2018). World Ferns: Checklist of Ferns and Lycophytes of the World (version Apr 2018). In: Species 2000 & ITIS Catalogue of Life, 2018 Annual Checklist (Roskov Y., Abucay L., Orrell T., Nicolson D., Bailly N., Kirk P.M., Bourgoin T., DeWalt R.E., Decock W., De Wever A., Nieukerken E. van, Zarucchi J., Penev L., eds.). Digital resource at www.catalogueoflife.org/annual-checklist/2018. Species 2000: Naturalis, Leiden, the Netherlands. ISSN 2405-884X.<br><br>Calonje M., Stanberg L. & Stevenson D. (eds). (2018). The World List of Cycads, online edition (version Jun 2015). In: Species 2000 & ITIS Catalogue of Life, 2018 Annual Checklist (Roskov Y., Abucay L., Orrell T., Nicolson D., Bailly N., Kirk P.M., Bourgoin T., DeWalt R.E., Decock W., De Wever A., Nieukerken E. van, Zarucchi J., Penev L., eds.). Digital resource at www.catalogueoflife.org/annual-checklist/2018. Species 2000: Naturalis, Leiden, the Netherlands. ISSN 2405-884X.<br><br>Farjon A., Gardner M. & Thomas P. (2018). Conifer Database (version | encouraged, on condition that full and precise credit is given at three levels on all occasions that records are shown. |

| | |
|---|---|
| | Jan 2014). In: Species 2000 & ITIS Catalogue of Life, 2018 Annual Checklist (Roskov Y., Abucay L., Orrell T., Nicolson D., Bailly N., Kirk P.M., Bourgoin T., DeWalt R.E., Decock W., De Wever A., Nieukerken E. van, Zarucchi J., Penev L., eds.). Digital resource at www.catalogueoflife.org/annual-checklist/2018. Species 2000: Naturalis, Leiden, the Netherlands. ISSN 2405-884X<br><br>Govaerts R. (ed). For a full list of reviewers see: http://apps.kew.org/wcsp/compilersReviewers.do (2018). WCSP: World Checklist of Selected Plant Families (version Aug 2017). In: Species 2000 & ITIS Catalogue of Life, 2018 Annual Checklist (Roskov Y., Abucay L., Orrell T., Nicolson D., Bailly N., Kirk P.M., Bourgoin T., DeWalt R.E., Decock W., De Wever A., Nieukerken E. van, Zarucchi J., Penev L., eds.). Digital resource at www.catalogueoflife.org/annual-checklist/2018. Species 2000: Naturalis, Leiden, the Netherlands. ISSN 2405-884X<br><br>Hassler M. (2018). World Plants: Synonymic Checklists of the Vascular Plants of the World (version Apr 2018). In: Species 2000 & ITIS Catalogue of Life, 2018 Annual Checklist (Roskov Y., Abucay L., Orrell |

| | |
|---|---|
| | T., Nicolson D., Bailly N., Kirk P.M., Bourgoin T., DeWalt R.E., Decock W., De Wever A., Nieukerken E. van, Zarucchi J., Penev L., eds.). Digital resource at www.catalogueoflife.org/annual-checklist/2018. Species 2000: Naturalis, Leiden, the Netherlands. ISSN 2405-884X.<br><br>Rainer H. & Chatrou L.W. (eds) (2018). AnnonBase: Annonaceae GSD (version Jan 2014). In: Species 2000 & ITIS Catalogue of Life, 2018 Annual Checklist (Roskov Y., Abucay L., Orrell T., Nicolson D., Bailly N., Kirk P.M., Bourgoin T., DeWalt R.E., Decock W., De Wever A., Nieukerken E. van, Zarucchi J., Penev L., eds.). Digital resource at www.catalogueoflife.org/annual-checklist/2018. Species 2000: Naturalis, Leiden, the Netherlands. ISSN 2405-884X.<br><br>Warwick S.I., Francis A. & Al-Shehbaz I.A. (2018). Brassicaceae species checklist and database (version 2, Oct 2009). In: Species 2000 & ITIS Catalogue of Life, 2018 Annual Checklist (Roskov Y., Abucay L., Orrell T., Nicolson D., Bailly N., Kirk P.M., Bourgoin T., DeWalt R.E., Decock W., De Wever A., Nieukerken E. van, Zarucchi J., Penev L., eds.). Digital resource at www.catalogueoflife.org/annual-checklist/2018. Species 2000: |

| | |
|---|---|
| | Naturalis, Leiden, the Netherlands. ISSN 2405-884X.<br><br>Culham A. & Yesson C. (2018). Droseraceae Database (version 0.1, Dec 2008). In: Species 2000 & ITIS Catalogue of Life, 2018 Annual Checklist (Roskov Y., Abucay L., Orrell T., Nicolson D., Bailly N., Kirk P.M., Bourgoin T., DeWalt R.E., Decock W., De Wever A., Nieukerken E. van, Zarucchi J., Penev L., eds.). Digital resource at www.catalogueoflife.org/annual-checklist/2018. Species 2000: Naturalis, Leiden, the Netherlands. ISSN 2405-884X.<br><br>Roskov Y., Zarucchi J., Novoselova M. & Bisby F.(†) (eds). (2018). ILDIS World Database of Legumes (version 12, May 2014). In: Species 2000 & ITIS Catalogue of Life, 2018 Annual Checklist (Roskov Y., Abucay L., Orrell T., Nicolson D., Bailly N., Kirk P.M., Bourgoin T., DeWalt R.E., Decock W., De Wever A., Nieukerken E. van, Zarucchi J., Penev L., eds.). Digital resource at www.catalogueoflife.org/annual-checklist/2018. Species 2000: Naturalis, Leiden, the Netherlands. ISSN 2405-884X.<br><br>Maslin B. (2018). WWW: World Wide Wattle (version 2, Jan 2018). In: Species 2000 & ITIS Catalogue of Life, 2018 Annual Checklist (Roskov |

| | |
|---|---|
| | Y., Abucay L., Orrell T., Nicolson D., Bailly N., Kirk P.M., Bourgoin T., DeWalt R.E., Decock W., De Wever A., Nieukerken E. van, Zarucchi J., Penev L., eds.). Digital resource at [www.catalogueoflife.org/annual-checklist/2018](www.catalogueoflife.org/annual-checklist/2018). Species 2000: Naturalis, Leiden, the Netherlands. ISSN 2405-884X<br><br>Aedo C. (2018). RJB Geranium: Geranium Taxonomic Information System (version Sep 2017). In: Species 2000 & ITIS Catalogue of Life, 2018 Annual Checklist (Roskov Y., Abucay L., Orrell T., Nicolson D., Bailly N., Kirk P.M., Bourgoin T., DeWalt R.E., Decock W., De Wever A., Nieukerken E. van, Zarucchi J., Penev L., eds.). Digital resource at [www.catalogueoflife.org/annual-checklist/2018](www.catalogueoflife.org/annual-checklist/2018). Species 2000: Naturalis, Leiden, the Netherlands. ISSN 2405-884X. |
| The Chinese Virtual Herbarium (CVH, [http://www.cvh.ac.cn/](http://www.cvh.ac.cn/)) | Biodiversity occurrence data provided by:Chengdu Institute of Biology,Chinese Academy of Sciences;Wuhan Botanical Garden,Chinese Academy of Sciences;Xishuangbanna Tropical Botanical Garden, Chinese Academy of Sciences;Northwest Institute of Plateau Biology,Chinese Academy of Sciences;Guangxi Institute of Botany, Chinese Academy of Sciences;South China Botanical Garden,Chinese Academy of Sciences;Institute of Applied Ecology, |

| | | |
|---|---|---|
| | Chinese Academy of Sciences;Kunming Institute of Botany, Chinese Academy of Sciences;Lushan Botanical Garden,Chinese Academy of Sciences;Institute of Botany, Jiangsu Province and Chinese Academy of Sciences;Chinese National Herbarium, Institute of Botany, Chinese Academy of Sciences;WUK Herbarium, Northwest Agriculture & Forestry University (Accessed through Chinese Virtual Herbarium(CVH) Data Portal, www.cvh.ac.cn, 2019-04-02) | |
| The Cichorieae Portal (http://cichorieae.e-taxonomy.net/) | Kilian N., Hand R. & Raab-Straube E. von (eds) 2009+ (continuously updated): Cichorieae Systematics Portal. last time accessed 06.12.2019 | CC BY 4.0 |
| The Euro+Med PlantBase (http://ww2.bgbm.org/EuroPlusMed/query.asp) | Euro+Med (2006-): Euro+Med PlantBase - the information resource for Euro-Mediterranean plant diversity. Published on the Internet http://ww2.bgbm.org/EuroPlusMed/. last time accessed 05.06.2019 | |
| The Flora Do Brasil 2020 (http://floradobrasil.jbrj.gov.br/reflora/listaBrasil/ConsultaPublicaUC/ConsultaPublicaUC.do#CondicaoTaxonCP) | Flora do Brasil 2020 under construction. Jardim Botânico do Rio de Janeiro. Available at: < http://floradobrasil.jbrj.gov.br/ >. Accessed on: 03.03.2019 | CC BY-SA 4.0 |
| The Flora Malesiana (https://floramalesiana.org/new/) http://portal.cybertaxonomy.org/flora-malesiana/ | The Flora Malesiana, https://floramalesiana.org/new/ | |
| The Integrated Taxonomic Information System (ITIS, https://www.itis.gov/) | Retrieved [October, 10, 2010], from the Integrated Taxonomic Information System on-line database, http://www.itis.gov. | Information presented on the ITIS website is considered public information and may be distributed or copie |

| | | | |
|---|---|---|---|
| The Melastomataceae database (MEL names, http://www.melastomataceae.net/MELnames/) | Melastomataceae.Net 2007–2020. A Site with Information on the Biodiversity of Melastomataceae: www.melastomataceae.net; assessed: 02.04.2019 | | |
| The Pelargonium Page (http://www.pelargonium.si/index.html) | The Pelargonium Page, Matija Strlic, http://www.pelargonium.si/index.html. assessed: 02.04.2019 | This work is licensed under a Creative Commons Attribution-NonCommercial-ShareAlike 4.0 International License. | |
| The Plants Database of the USDA (https://plants.usda.gov/java/) | USDA, NRCS. 2019. The PLANTS Database (http://plants.usda.gov, 23 March 2019). National Plant Data Team, Greensboro, NC 27401-4901 USA | | |
| The Plant List (TPL) (http://www.theplantlist.org) | The Plant List (2013). Version 1.1. Published on the Internet; http://www.theplantlist.org/ (accessed 1st February 2013). | Use of Content for non-commercial or non-profit use is encouraged | |
| The Smithsonian National Museum of Natural History (https://collections.nmnh.si.edu/search/botany/) | Assessed: 01.08.2019 | CC- 0 | Information provided with the permission of the National Museum of Natural History, Smithsonian Institution, 10th and Constitution Ave. N.W., Washington, DC 20560-0193. (https://collections.nmnh.si.edu/) |
| The Southern African plant names and floristic details (SANBI, http://posa.sanbi.org/sanbi) | South African National Biodiversity Institute. 2016. Botanical Database of Southern Africa (BODATSA) | Creative Commons Attribution-NonCommercial-ShareAlike 4.0 International License | |
| The World Checklist of Selected Plant Families (WCSP, Botanical Gardens Kew, https://wcsp.science.kew.org), | WCSP (2020). 'World Checklist of Selected Plant Families. Facilitated by the Royal Botanic Gardens, Kew. Published on the Internet; http://wcsp.science.kew.org/ Retrieved | CC BY | Board of Trustees of the Royal Botanic Gardens, Kew |
| Grassbase (http://www.kew.org/data/grassbase/index.html) | Clayton, W.D., Vorontsova, M.S., Harman, K.T. and Williamson, H. (2006 onwards). GrassBase - The Online World Grass Flora. | CC BY | Board of Trustees of the Royal Botanic Gardens, Kew |

| | | | |
|---|---|---|---|
| | http://www.kew.org/data/grasses-db.html. [accessed 08 November 2006] | | |
| Palmweb (http://palmweb.org/) | Palmweb [2019]. Palmweb: Palms of the World Online. Published on the internet [http://palmweb.org/]. Acccessed on [05/08/2019]. | CC-BY-NC-SA | Board of Trustees of the Royal Botanic Gardens, Kew |
| TROPICOS (Missouri Botanical Gardens, http://www.tropicos.org/) | Tropicos.org. Missouri Botanical Garden. 01 Aug 2019 <http://www.tropicos.org> | This site is designed for scientific research by individual researchers. Attribution should be given for use of data derived from the site. | |
| World Annonaceae (http://annonaceae.myspecies.info) | World Annonaceae, Thomas L.P. Couvreur, http://annonaceae.myspecies.info, assessed: 01.08.2019 | CC BY | |
