## Supplementary file 2 - lcvplants description for "The Leipzig Catalogue of Vascular Plants (LCVP) – An improved taxonomic reference list for all known vascular plants"

**Supplementary file 2: Leipzig Catalogue of Plants (LCVP) R-Package (lcvplants) description**

### The Leipzig Catalogue of Vascular Plants (LCVP) – An improved taxonomic reference list for all known vascular plants

Martin Freiberg[1*], Marten Winter[2,] Alessandro Gentile[2], Alexander Zizka[2,3], Alexandra Nora Muellner-Riehl[2,4], Alexandra Weigelt[1,2], Christian Wirth[1,2,4,5]

#### Introduction

The Leipzig Catalogue of Vascular Plants is implemented in two R packages: 1) LCVP, a datapackage, which contains the actual list of plant names and 2) lcvplants which contains a fuzzy matching algorithm to use this list for taxonomic name resolution. LCVP development was led by Martin Freiberg at the Botanical Garden of the Leipzig University and lcvplants was developed at the German Centre for Integrative Biodiversity Research (iDiv). Together these packages provide a tool suite to taxonomically and orthographically resolve plant names using the Leipzig Catalogue of Vascular Plants. Here we describe the structure of the data used by both packages and the algorithm used by lcvplants for taxonomic name resolution, a description on how to run the resolution can be found in the vignette "Taxonomic name resolution with lcvplants" available online at: https://idiv-biodiversity.github.io/lcvplants/articles/taxonomic_resolution_using_lcplants.html

#### 1. The data of the Leipzig Catalogue of Vascular Plants (the LCVP package)

LCVP is a data package solely containing two data files: 1) tab_lcvp which is the dataset of plant names and their taxonomic status, and ref_lcvp, which is a list of the literature references used to compile the list. Furthermore LCVP contains an internal table named tab_position, which is used by lcvplants to speed up the name matching The tab_lcvp table contains seven columns:

1. *Input.Taxon*: a list of all vascular plant species names listed in the Leipzig Catalogue of Vascular Plants (LCVP).
2. *Status*: description if a taxon is classified as 'valid', 'synonym', 'unresolved', 'external' or 'blanks'.
   The 'unresolved' rank means that the status of the plant name could be either valid or synonym, but the information available does not allow a definitive decision.
   'External' in an extra rank which lists names outside the scope of this publication but useful to keep on this updated list.
   'Blanks' means that the respective name exists in bibliography but it is neither clear where it came from valid, synonym or unresolved.
3. *PL.comparison*: field provides a direct comparison with 'The Plant List' (TPL; The Plant List http://www.theplantlist.org/ accessed: 1.1. 2013) reporting further information such as 'identical', 'synonym', 'other synonym', 'different authors', 'missing', 'misspelling', 'unresolved'.
4. *PL.Alternative*: possible alternative name from the TPL.
5. *Output.Taxon*: variable contains the list of the accepted plant taxa names according to the LCVP.
6. *Family:* corresponding family name of the Input.Taxon, staying empty if the Status is unresolved.
7. *Order*: corresponding order name of the Input.Taxon, staying empty if the Status is unresolved.

#### 2. The fuzzy matching function and algorithm in lcvplants

The lcvplants package contains the function *LCVP* for a text-based search of plant names on the LCVP using a fuzzy match algorithm. The fuzzy match algorithm can be applied at genus, epithet, infraspecies and authority level of a plant species and it uses the 'max.distance' argument from the agrep() R-function to assess the comparison between the searched plant name and the closest (in terms of number of similar characters) plant name from the LCVP.

For the input data, following the International Code of Nomenclature for algae, fungi, and plants (Shenzhen Code: https://www.iapt-taxon.org/nomen/main.php), genus, epithet, infraspecies rank, infraspecies name and authorities need to be separated by spaces (e.g. *Draba mollissima* var. *kusnezowii* N.Busch). Special characters (such as ü, á, ø, etc.) are allowed for the authority names but not the genus name, epitheton and infraspecies names. Infraspecific names have to be preceded by their rank (e.g. "subsp.", "var.", "forma", "ssp.", "f.", "subvar.", "subf."). The genus name and the epitheton are necessary for a name to be matched; the infraspecific ranks and authority names are optional for better results. If the genus or the epithet names are composed of two words, they have to be separated by a hyphen ("-") but never by a space (e.g. *Hibiscus rosa-sinensis* L not *Hibiscus rosa sinensis* L.). Hybrid names use the characters '_x' in the end of the genus and epithet name (e.g. *Spartocytisus*_x *filipes* Webb & Berthel., *Lycopodium habereri*_x House) annotations in other formats such as 'x' or 'x_' before the names both upper or lower case (e.g. x *Spartocytisus filipes* Webb & Berthel., x_*Spartocytisus filipes* Webb & Berthel., *Lycopodium* x *habereri* House) are also supported and changed automatically into the required format. The commonly used special Unicode Character '×' (U+00D7) for indicating hybrids is not accepted (e.g. *Crassocephalum* ×*picridifolium*).

Through the *LCVP* function the user can search either for a single plant name or for a list of names. There are no limitation on the list length, but we recommend to submit less than 5000 names at a time to ensure a reasonable computation time. See the vignette "Taxonomic resolution using lcvplants" available with lcvplants and at https://idiv-biodiversity.github.io/lcvplants/articles/taxonomic_resolution_using_lcplants.html for a tutorial on how to do the name resolution within few lines of R code.

*LCVP* function, executed as LCVP("*Polygonatum pubescens*", …). has a set of arguments to customize the fuzzy matching

- *genus_search*: is a logical value, FALSE (default). If TRUE, the function will apply the fuzzy match algorithm also for the search of the genus name, otherwise as default the search is applied only to the epithet, the infraspecies and the name author.
- *max.distance*: is an integer value. It represents the maximum distance (number of characters) allowed for a match when comparing the submitted name with the closest name matches in the LCVP.
- *status:* is a logical value, TRUE (default). If FALSE, the function will return not only the valid epithet for a species name but also all the possible synonyms.
- *max.cores*: is an integer value, indicating the number of CPU cores to be used for the parallelization of the plant name search when a list of plant taxa names is submitted. As default, the maximum number of CPU cores available on the working machine menus one is set.
- *genus_tab*: is a logical value, FALSE (default). If TRUE, the function will return the list of plant taxa names belonging to the same genus name submitted by the user.
- *family_tab*: is a logical value, FALSE (default). If TRUE, the function will return the list of plant taxa names belonging to the same family name submitted by the user.
- *order_tab*: is a logical value, FALSE (default). If TRUE, the function will return the list of plant taxa names belonging to the same order name submitted by the user.

- *infraspecies_tab*: is a logical value, FALSE (default). If TRUE, the function will return also all the infraspecies names found for a submitted plant name.
- *encoding*: is a character vector, "UTF-8" (default). This value will allow the user to set the specific codification of the strings.
- *out_path*: is a character vector, which allow the user to define the path where the output file has to be saved. The working directory is set as default.
- *save*: is a logical value. FALSE (default). If TRUE, the function will write the output file as comma-separated format (.csv), saving it into the working directory or in the directory already set through the 'out_path' option.
- *visualize*: is a logical value, TRUE (default). If TRUE the function will visualize the output search on the 'Source Tab' of RStudio. This option has to be turned off (FALSE) if the package is executed in a UNIX environment (from the command line) without having a Graphical User Interface.
- *version*: A character vector indicating the current version of the package (current version is 1.0). A new version is under development allowing the package to connect the web API that is under construction.

The *LCVP* function applies a string comparison between the submitted names from the user with the list of taxa listed in the 'input.taxon' column of the tab_LCVP table, applying a fuzzy match algorithm for possible orthographic errors solving. Taxa are parse one at a time. Following the fuzzy matching, the function performs a further search starting with the genus name, the epithet name and eventually, if inserted from the user, the infraspecies and the authority information. The entire is executed in the following subsequent steps

1. preprocessing of the submitted name. The function takes account of how many terms the submitted taxon is composed. If the number of the term is two, the function will apply the search only for the genus and the epithet term. If the submitted taxon is formed by more than two terms, then the infraspecies name or the authority name or both (if submitted by the user) will be considered in the searching process. The first term is always associated to the genus name and the second term to the epithet name. If the third term is belonging to the infraspecies categories (such as: "subsp.", "var.", "forma", "ssp.", "f.", "subvar.", "subf.") it will be associated to one of those rank, so the function will associate the fourth word to the infraspecies name. If the third term is not in the list of the infraspecies rank, all the words from the third to the end of the submitted name will be associated to the authority description.
2. Once the all components of the species name are parsed and identified name resolutions starts with the genus name. The searching process will begin to select the starting position and the ending position from the tab_position table. First, the first three letters of the submitted genus name (e.g. 'Las', for "*Laserpitium eliasii*") are compared to the records in the list of Triphthongs in tab_position. When matched, the corresponding position information is stored in memory (e.g. 688815). This value will be the starting point from where to start the search on the tab_lcvp; and the following position value (e.g. 691389) will be used as the end point to the search.
3. At this point, the searching process for the genus name of the submitted taxon starts at the first entries in tab_lcvp. If the input genus name does not match any LCVP genus name, an empty row is returned and the message "Genus name not found" is returned in the 'Score' column.
4. If a match for the submitted genus name is found, similar matching will be done to find the correct species epitheton term. If the submitted epitheton cannot be matched in LCVP, an empty row is returned and the message "Epithet name not found" is returned in the 'Score' column.
5. At this point, if the 'Status' option is set as TRUE (default) the function will select the only taxa name with a 'valid' status. If there are no valid name in the LCVP for a specific taxon, the *LCVP* function will return all relevant synonyms.
6. Instead, if the user has activated the option 'infraspecies_tab' (infraspecies_tab = TRUE), this will allow the function to return all the records belonging to the taxa name matched (valid, synonym and subspecies ranks).
7. If an input taxa name matches an LCVP taxa name (i.e. both the genus and the epithet were found), the *LCVP* function proceeds to the next step. If the user has specified also the infraspecies term, the function

will start the searching process, applying the fuzzy match algorithm, to the infraspecies names, similarly to the previous steps. Finally, also for the taxa author (if included in the input data) will be processed with the fuzzy match algorithm.

8. In a final step, the results for all submitted names will be combined into the output table and the results will be returned by the function and printed to screen.

**Description of the output table**

The *LCVP* function returns a data.frame of the submitted and corresponding taxon names with additional information on the taxonomic status. If the option 'save' is turned active (Save = TRUE), the output will additionally be saved in a comma-separated file (.csv) in the working directory or the path specified with the 'out_path' option. The following list describes the format of the output table of the *LCVP* function. If a name could not be resolved, in the "Leipzig Catalogue of Vascular Plants" the output data.frame has empty columns unless for the field of the 'Submitted_Name' and the 'Score' fields (columns).

- *ID*: it is a progressive integer associated to each taxa; the progression of the numbers follows the same order of the submitted names in the input list (introduced by the user). The only cases where it is possible to find the same ID value is when the searching process for one taxon will return more than one records (e.g. synonyms).
- *Submitted_Name*: it is a character string of all the input names submitted by the users.
- *Order*: It is the corresponding taxonomic order name for the matched taxon name.
- *Family*: It is the corresponding taxonomic family name for the matched taxon name.
- *Genus*: It is the corresponding taxonomic genus name for the matched taxon name.
- *Species*: It is the corresponding taxonomic epithet name for the matched taxon name.
- *Infrasp*: It is the corresponding taxonomic infraspecific rank for the matched taxon name.
- *Infraspecies*: It is the corresponding taxonomic infraspecific name for the matched taxon name.
- *Authors*: it is the description of the authorship for the matched taxon name.
- *Status*: it is a character string, which provides a description if a taxon is classified as 'valid', 'synonym', or 'unresolved'.
- *LCVP_Accepted_Taxon*: it is a character string, it provides the accepted (Status = valid) plant taxa name according to the LCVP system.
- *PL_Comparison*: it is a character string, it provides a direct comparison with 'The Plant List' (TPL)
- *PL_Alternative*: it is a character string, it provides an alternative name from the TPL
- *Score*: it is a description of the result for the searching process in the LCVP. In this field different information are stored depending by the results of the searching and solving activities (e.g. 'name found', 'genus name not found', 'epithet name not found', misspelling: epithet', etc.).
- *Insertion*: it is an integer; it describes how many characters the submitted name differs (as inserted types) for the full match with the corresponding name found in the LCVP.
- *Deletion*: it is an integer; it describes how many characters the submitted name differs (as deleted) from the matched name in the LCVP.
- *Substitution*: it is an integer; it describes how many characters the submitted name differs (as substituted) from the matched name in the LCVP.
