## Supplementary file 3 - Literature used to compile LCVP ordered by plant families for "The Leipzig Catalogue of Vascular Plants (LCVP) – An improved taxonomic reference list for all known vascular plants"

### Acanthaceae

**Arolla, Rajender Goud; Cherukupalli, Neeraja; Khareedu, Venkateswara Rao; Vudem, Dashavantha Reddy (2015):**

**DNA barcoding and haplotyping in different Species of *Andrographis*.**

In: Biochemical Systematics and Ecology 62, p. 91–97. DOI: 10.1016/j.bse.2015.08.001.

**Borg, Agneta Julia; McDade, Lucinda A.; Schönenberger, Jürgen (2008):**

**Molecular Phylogenetics and morphological Evolution of Thunbergioideae (Acanthaceae).**

In: Taxon 57 (3), p. 811–822. DOI: 10.1002/tax.573012.

**Carine, Mark A.; Scotland, Robert W. (2002):**

**Classification of Strobilanthinae (Acanthaceae): Trying to Classify the Unclassifiable?**

In: Taxon 51 (2), p. 259–279. DOI: 10.2307/1554926.

**Côrtes, Ana Luiza A.; Daniel, Thomas F.; Rapini, Alessandro (2016):**

**Taxonomic Revision of the Genus *Schaueria* (Acanthaceae).**

In: Plant Systematics and Evolution 302 (7), p. 819–851. DOI: 10.1007/s00606-016-1301-y.

**Côrtes, Ana Luiza A.; Rapini, Alessandro; Daniel, Thomas F. (2015):**

**The Tetramerium Lineage (Acanthaceae: Justicieae) does not support the Pleistocene Arc Hypothesis for South American seasonally dry Forests.**

In: American Journal of Botany 102 (6), p. 992–1007. DOI: 10.3732/ajb.1400558.

**Daniel, Thomas F.; McDade, Lucinda A. (2014):**

**Nelsonioideae (Lamiales: Acanthaceae): Revision of Genera and Catalog of Species.**

In: Aliso 32 (1), p. 1–45. DOI: 10.5642/aliso.20143201.02.

**Ezcurra, Cecilia (2002):**

**El Género *Justicia* (Acanthaceae) en Sudamérica Austral.**

In: Annals of the Missouri Botanical Garden 89, p. 225–280.

**Fisher, Amanda E.; McDade, Lucinda A.; Kiel, Carrie A.; Khoshravesh, Roxanne; Johnson, Melissa A.; Stata, Matt et al. (2015):**

**Evolutionary History of *Blepharis* (Acanthaceae) and the Origin of C 4 Photosynthesis in Section Acanthodium.**

In: International Journal of Plant Sciences 176 (8), p. 770–790. DOI: 10.1086/683011.

**Gnanasekaran, G.; Murthy, G.V.S.; Deng, Yun-Fei (2016):**

**Resurrection of the Genus *Haplanthus* (Acanthaceae: Andrographinae).**

In: Blumea 61 (3), p. 165–169. DOI: 10.3767/000651916X693185.

**Graham, Victoria A. W. (1988):**

**Delimitation and Infra-Generic Classification of *Justicia* (Acanthaceae).**

In: Kew Bulletin 43 (4), p. 551–624.

**Hilsenbeck, Richard A. (1990):**

**Systematics of *Justicia* Sect. Pentaloba (Acanthaceae).**

In: Plant Systematics and Evolution 169, p. 219–235.

**Kiel, Carrie A.; McDade, Lucinda A.; Daniel, Thomas F.; Champluvier, Dominique (2006):**

**Phylogenetic Delimitation of Isoglossinae (Acanthaceae: Justicieae) and Relationships among constituent Genera.**

In: Taxon 55 (3), p. 683–694. DOI: 10.2307/25065644.

**Manktelow, Mariette; McDade, Lucinda A.; Oxelman, Bengt; Furness, Carol A.; Balkwill, Mandy-Jane (2001):**

**The Enigmatic Tribe Whitfieldieae (Acanthaceae): Delimitation and phylogenetic Relationships based on molecular and morphological Data.**

In: Systematic Botany 26 (1), p. 104–119.

**McDade, Lucinda A.; Daniel, Thomas F.; Kiel, Carrie A. (2008):**

**Toward a comprehensive Understanding of phylogenetic Relationships among Lineages of Acanthaceae s.l. (Lamiales).**

In: American Journal of Botany 95 (9), p. 1136–1152. DOI: 10.3732/ajb.0800096.

**McDade, Lucinda A.; Daniel, Thomas F.; Kiel, Carrie A.; Borg, Agneta Julia (2012):**

**Phylogenetic placement, Delimitation, and Relationships among Genera of the enigmatic Nelsonioideae (Lamiales: Acanthaceae).**

In: Taxon 61 (3), p. 637–651. DOI: 10.1002/tax.613012.

**McDade, Lucinda A.; Daniel, Thomas F.; Kiel, Carrie A.; Vollesen, Kaj (2005):**

**Phylogenetic Relationships among Acantheae (Acanthaceae): Major Lineages Present Contrasting Patterns of molecular Evolution and morphological Differentiation.**

In: Systematic Botany 30 (4), p. 834–862.

**McDade, Lucinda A.; Masta, Susan E.; Moody, Michael L.; Waters, Elizabeth (2000):**

**Phylogenetic Relationships among Acanthaceae: Evidence from Two Genomes.**

In: Systematic Botany 25 (1), p. 106–121. DOI: 10.2307/2666677.

**McDade, Lucinda A.; Moody, Michael L. (1999):**

**Phylogenetic Relationships among Acanthaceae: Evidence from noncoding trnL-trnF Chloroplast DNA Sequences.**

In: American Journal of Botany 86 (1), p. 70–80. DOI: 10.2307/2656956.

**Moylan, Elizabeth C.; Bennett, Jonathan R.; Carine, Mark A.; Olmstead, Richard G.; Scotland, Robert W. (2004):**

**Phylogenetic Relationships among *Strobilanthes* s.l. (Acanthaceae): Evidence from ITS nrDNA, trnL-F cpDNA, and Morphology.**

In: American Journal of Botany 91 (5), p. 724–735. DOI: 10.3732/ajb.91.5.724.

**Onjalalaina, Guy Eric; Darbyshire, Iain (2016):**

**An endangered new Species of *Podorungia* (Acanthaceae), with Notes on the Tribe Barlerieae in Madagascar.**

In: Kew Bulletin 71 (3), p. 117. DOI: 10.1007/S12225-016-9657-2.

**Schmidt-Lebuhn, Alexander N. (2005):**

**Evolution of (Acanthaceae) inferred from Morphology, AFLP Data, and ITS rDNA Sequences.**

In: Organisms Diversity and Evolution 5 (1), p. 1–13. DOI: 10.1016/j.ode.2004.04.006.

**Schwarzbach, Andrea E.; McDade, Lucinda A. (2002):**

**Phylogenetic Relationships of the Mangrove Family Avicenniaceae based on Chloroplast and Nuclear Ribosomal DNA Sequences.**

In: Systematic Botany 27 (1), p. 84–98.

**Tripp, Erin A. (2007):**

**Evolutionary Relationships within the Species-Rich Genus *Ruellia* (Acanthaceae).**

In: Systematic Botany 32 (3), p. 628–649.

**TRIPP, ERIN A.; DANIEL, THOMAS F.; FATIMAH, SITI; MCDADE, LUCINDA A. (2013):**
**Phylogenetic Relationships within Ruellieae (Acanthaceae) and a revised Classification.**
In: International Journal of Plant Sciences 174 (1), p. 97–137. DOI: 10.1086/668248.

**TRIPP, ERIN A.; DANIEL, THOMAS F.; LENDEMER, JAMES C.; MCDADE, LUCINDA A. (2009):**
**New molecular and morphological Insights prompt transfer of *Blechum* to *Ruellia* (Acanthaceae).**
In: Taxon 58 (3), p. 893–906. DOI: 10.1002/tax.583017.

**TRIPP, ERIN A.; DARBYSHIRE, IAIN (2017):**
**Phylogenetic Relationships among Old World *Ruellia* L.: A new Classification and Reinstatement of the Genus *Dinteracanthus* Schinz.**
In: Systematic Botany 42 (3), p. 470–483. DOI: 10.1600/036364417X695961.

**TRIPP, ERIN A.; FATIMAH, SITI (2012):**
**Comparative Anatomy, Morphology, and molecular Phylogenetics of the African Genus *Satanocrater* (Acanthaceae).**
In: American Journal of Botany 99 (6), p. 967–982. DOI: 10.3732/ajb.1100354.

**TRIPP, ERIN A.; MCDADE, LUCINDA A. (2012):**
**New synonymies for *Ruellia* (Acanthaceae) of Costa Rica and Notes on other Neotropical Species.**
In: Brittonia 64 (3), p. 305–317. DOI: 10.1007/s12228-012-9244-2.

**WASSHAUSEN, DIETER C. (1992):**
**New Species and new Combinations of *Justicia* (Acanthaceae) from the Venezuelan Guayana.**
In: Novon: A Journal for Botanical Nomenclature 2 (1), p. 62–80.

### Achariaceae

**CHASE, MARK W.; ZMARZTY, SUE; LLEDÓ, M. DOLORES; WURDACK, KENNETH J.J.; SWENSEN, SUSAN M.; FAY, MICHAEL F. (2002):**
**When in Doubt, put It in Flacourtiaceae: A molecular phylogenetic Analysis based on Plastid rbcL DNA Sequences.**
In: Kew Bulletin 57, p. 141–181.

**GROPPO, MILTON; FIASCHI, PEDRO; SALATINO, MARIA LUIZA FARIA; CECCANTINI, GREGÓRIO CARDOSO TÁPIAS; ASSIS RIBEIRO DOS SANTOS, FRANCISCO; VEROLA, CHRISTIANO FRANCO; ANTONELLI, ALEXANDRE (2010):**
**Placement of *Kuhlmanniodendron* Fiaschi & Groppo in Lindackerieae (Achariaceae, Malpighiales) confirmed by Analyses of rbcL Sequences, with Notes on Pollen Morphology and wood Anatomy.**
In: Plant Systematics and Evolution 286 (1-2), p. 27–37. DOI: 10.1007/s00606-010-0276-3.

**WEBBER, BRUCE L.; MILLER, REBECCA E. (2008):**
**Gynocardin from *Baileyoxylon lanceolatum* and a Revision of cyanogenic Glycosides in Achariaceae.**
In: Biochemical Systematics and Ecology 36 (7), p. 545–553. DOI: 10.1016/j.bse.2008.03.011.

### Actinidiaceae

**CHAT, JOËLLE; JÁUREGUI, BLANCA; PETIT, RÉMY J.; NADOT, SOPHIE (2004):**
**Reticulate Evolution in kiwifruit (*Actinidia*, Actinidiaceae) identified by comparing their maternal and paternal Phylogenies.**

In: American Journal of Botany 91 (5), p. 736–747. DOI: 10.3732/ajb.91.5.736.

**Shi, Tao; Huang, Hongwen; Barker, Michael p. (2010):**
**Ancient Genome duplications during the Evolution of Kiwifruit (*Actinidia*) and related Ericales.**
In: Annals of Botany 106 (3), p. 497–504. DOI: 10.1093/aob/mcq129.

**Yao, Xiaohong; Liu, Lei; Yan, Mingke; Li, Dawei; Zhong, Caihong; Huang, Hongwen (2015):**
**Exon primed Intron-crossing (Epic) Markers reveal natural Hybridization and Introgression in *Actinidia* (Actinidiaceae) with sympatric Distribution.**
In: Biochemical Systematics and Ecology 59, p. 246–255. DOI: 10.1016/j.bse.2015.01.023.

### Adoxaceae

**Jacobs, Bart; Huysmans, Suzy; Smets, Erik (2010):**
**Evolution and systematic value of Fruit and Seed Characters in Adoxaceae (Dipsacales).**
In: Taxon 59 (3), p. 850–865.

### Agavaceae

**Althoff, David M.; Segraves, Kari A.; Smith, Christopher I.; Leebens-Mack, James H.; Pellmyr, Olle (2012):**
**Geographic isolation trumps CoEvolution as a Driver of *Yucca* and *Yucca*-moth Diversification.**
In: Molecular Phylogenetics and Evolution 62 (3), p. 898–906. DOI: 10.1016/j.ympev.2011.11.024.

**Archibald, Jenny K.; Kephart, Susan R.; Theiss, Kathryn E.; Petrosky, Anna L.; Culley, Theresa M. (2015):**
**Multilocus phylogenetic Inference in Subfamily Chlorogaloideae and related Genera of Agavaceae - informing questions in Taxonomy at multiple ranks.**
In: Molecular Phylogenetics and Evolution 84, p. 266–283. DOI: 10.1016/j.ympev.2014.12.014.

**Fishbein, Mark; Kephart, Susan R.; Wilder, Mike; Halpin, Kate M.; Datwyler, Shannon L. (2010):**
**Phylogeny of *Camassia* (Agavaceae) inferred from Plastid rpl16 Intron and trnD–trnY–trnE–trnT Intergenic Spacer DNA Sequences: Implications for Species Delimitation .**
In: Systematic Botany 35 (1), p. 77–85. DOI: 10.1600/036364410790862588.

**Good-Avila, Sara V.; Souza, Valeria; Gaut, Brandon S.; Eguiarte, Luis E. (2006):**
**Timing and rate of Speciation in *Agave* (Agavaceae).**
In: Proceedings of the National Academy of Sciences of the United States of America 103 (24), p. 9124–9129. DOI: 10.1073/pnas.0603312103.

**Heyduk, Karolina; McKain, Michael R.; Lalani, Falak; Leebens-Mack, James H. (2016):**
**Evolution of a CAM Anatomy predates the Origins of Crassulacean acid metabolism in the Agavoideae (Asparagaceae).**
In: Molecular Phylogenetics and Evolution 105, p. 102–113. DOI: 10.1016/j.ympev.2016.08.018.

**Lopes, Rosana Conrado; Andreata, Regina Helena Potsch; Cartaxo-Pinto, Simone; Trovó, Marcelo; Gonçalves-Esteves, Vânia (2013):**
**Pollen Morphology and wall Structure of Neotropical Species of *Herreria* and *Clara* (Asparagaceae-Agavoideae) and its taxonomic Implications.**
In: Plant Systematics and Evolution 299 (1), p. 25–34. DOI: 10.1007/s00606-012-0699-0.

**Pellmyr, Olle; Segraves, Kari A.; Althoff, David M.; Balcázar-Lara, Manuel; Leebens-Mack, James H. (2007):**
**The Phylogeny of yuccas.**
In: Molecular Phylogenetics and Evolution 43 (2), p. 493–501. DOI: 10.1016/j.ympev.2006.12.015.

**Pires, J. Chris; Maureira, Ivan J.; Rebman, Jon P.; Salazar, Gerardo A.; Cavrera, Lidia I.; Fay, Michael F.; Chase, Mark W. (2004):**

**Molecular Data confirm the phylogenetic Placement of the enigmatic *Hesperocallis* (Hesperocallidaceae) with *Agave*.**

In: Madroño 51 (3), p. 307–311.

### Aizoaceae

**Bohley, Katharina; Joos, Olga; Hartmann, Heidrun E.K.; Sage, Rowan F.; Liede-Schumann, Sigrid; Kadereit, Gudrun (2015):**

**Phylogeny of Sesuvioideae (Aizoaceae) – Biogeography, Leaf Anatomy and the Evolution of C4 photosynthesis.**

In: Perspectives in Plant Ecology, Evolution and Systematics 17 (2), p. 116–130. DOI: 10.1016/j.ppees.2014.12.003.

**Hassan, N. S.; Thiede, J.; Liede-Schumann, Sigrid (2005):**

**Phylogenetic Analysis of Sesuvioideae (Aizoaceae) inferred from nrDNA Internal Transcribed Spacer (ITS) Sequences and morphological Data.**

In: Plant Systematics and Evolution 255 (3-4), p. 121–143. DOI: 10.1007/s00606-005-0357-x.

**Kellner, A.; Ritz, Christine M.; Schlittenhardt, P.; Hellwig, Frank H. (2011):**

**Genetic differentiation in the Genus *Lithops* L. (Ruschioideae, Aizoaceae) reveals a high level of convergent Evolution and reflects geographic Distribution.**

In: Plant Biology 13 (2), p. 368–380. DOI: 10.1111/j.1438-8677.2010.00354.x.

**Klak, Cornelia; Bruyns, Peter V. (2012):**

**Phylogeny of the Dorotheantheae (Aizoaceae), a Tribe of succulent annuals**

61 (2), p. 293–307.

**Klak, Cornelia; Bruyns, Peter V. (2013):**

**A new infrageneric Classification for *Mesembryanthemum* (Aizoaceae: Mesembryanthemoideae).**

In: Bothalia 43 (2), p. 197–206. DOI: 10.4102/abc.v43i2.95.

**Klak, Cornelia; Bruyns, Peter V.; Hanáček, Pavel (2013):**

**A phylogenetic Hypothesis for the recently diversified Ruschieae (Aizoaceae) in southern Africa.**

In: Molecular Phylogenetics and Evolution 69 (3), p. 1005–1020. DOI: 10.1016/j.ympev.2013.05.030.

**Klak, Cornelia; Bruyns, Peter V.; Hedderson, Terry A. (2007):**

**A Phylogeny and new Classification for Mesembryanthemoideae (Aizoaceae).**

In: Taxon 56 (3), p. 737. DOI: 10.2307/25065858.

**Klak, Cornelia; Hanáček, Pavel; Bruyns, Peter V. (2014):**

**Phylogeny and Taxonomy for *Mesembryanthemum* Subg. Volkeranthus (Aizoaceae-Mesembryanthemoideae).**

In: South African Journal of Botany 95, p. 112–122. DOI: 10.1016/j.sajb.2014.08.006.

**Klak, Cornelia; Hanáček, Pavel; Bruyns, Peter V. (2015):**

**A Phylogeny and revised Classification for the Apatesieae (Aizoaceae: Ruschioideae) with a Comparison of centres of diversity.**

In: Taxon 64 (3), p. 507–522. DOI: 10.12705/643.7.

**Klak, Cornelia; Hanáček, Pavel; Bruyns, Peter V. (2017):**

**Disentangling the Aizooideae: new generic Concepts and a new Subfamily in Aizoaceae.**
In: Taxon 66 (5), p. 1147–1170. DOI: 10.12705/665.9.

KLAK, CORNELIA; HANÁČEK, PAVEL; BRUYNS, PETER V. (2017):
**Out of southern Africa: Origin, Biogeography and age of the Aizooideae (Aizoaceae).**
In: Molecular Phylogenetics and Evolution 109, p. 203–216. DOI: 10.1016/j.ympev.2016.12.016.

KLAK, CORNELIA; HEDDERSON, TERRY A.; LINDER, HANS PETER (2003):
**A molecular Systematic Study of the *Lampranthus* Group (Aizoaceae) based on the Chloroplast TrnL-trnF and Nuclear ITS and 5s Nts Sequence Data.**
In: Systematic Botany 28 (1), p. 70–85.

KLAK, CORNELIA; KHUNOU, ANGELINE; REEVES, GAIL; HEDDERSON, TERRY A. (2003):
**A phylogenetic Hypothesis for the Aizoaceae (Caryophyllales) based on four Plastid DNA Regions.**
In: American Journal of Botany 90 (10), p. 1433–1445. DOI: 10.3732/ajb.90.10.1433.

KLAK, CORNELIA; NOWELL, TRACEY L.; HEDDERSON, TERRY A. J. (2006):
**Phylogeny and Revision of *Brownanthus* and its close allies *Aspazoma* and *Dactylopsis* (Aizoaceae) based on Morphology and four DNA Regions.**
In: Kew Bulletin 61 (3), p. 353–400.

OPEL, MATTHEW R. (2005):
**A morphological Phylogeny of the Genus *Conophytum* N.E.Br. (Aizoaceae).**
In: Haseltonia 11, p. 53–77. DOI: 10.2985/1070-0048(2005)11[53:AMPOTG]2.0.CO;2.

POWELL, ROBYN F.; BOATWRIGHT, JAMES S.; KLAK, CORNELIA; MAGEE, ANTHONY RICHARD (2016):
**Phylogenetic Placement and generic re-Circumscriptions of the multilocular Genera *Arenifera*, *Octopoma* and *Schlechteranthus* (Aizoaceae: Ruschieae): Evidence from anatomical, morphological and Plastid DNA Data.**
In: Taxon 65 (2), p. 249–261. DOI: 10.12705/652.3.

THULIN, MATS; THIEDE, JOACHIM; LIEDE-SCHUMANN, SIGRID (2012):
**Phylogeny and Taxonomy of *Tribulocarpus* (Aizoaceae): A paraphyletic Species and an adaptive shift from zoochorous trample burrs to anemochorous nuts.**
In: Taxon 61 (1), p. 55–66.

VALENTE, LUIS M.; BRITTON, ADAM W.; POWELL, MARTYN P.; PAPADOPULOS, ALEXANDER P. T.; BURGOYNE, PRISCILLA M.; SAVOLAINEN, VINCENT (2014):
**Correlates of hyperdiversity in southern African ice plants (Aizoaceae).**
In: Botanical Journal of the Linnean Society 174 (1), p. 110–129. DOI: 10.1111/boj.12117.

### Alangiaceae

DEWILDE, W.J.J.O.; DUYFJES-DE WILDE, BRIGITTA E.E. (2017):
**Taxonomy of *Alangium* Section Conostigma (Alangiaceae ).**
In: Blumea 62 (1), p. 29–46. DOI: 10.3767/000651917X695164.

FENG, CHUN-MIAO; MANCHESTER, STEVEN R.; XIANG, QIU-YUN (2009):
**Phylogeny and Biogeography of Alangiaceae (Cornales) inferred from DNA Sequences, Morphology, and fossils.**
In: Molecular Phylogenetics and Evolution 51 (2), p. 201–214. DOI: 10.1016/j.ympev.2009.01.017.

WILDE, WILLEM J.J.O.; DUYFJES-DE WILDE, BRIGITTA E.E. (2017):
**The Species of *Alangium* Section Rhytidandra (Alangiaceae ).**

In: Blumea 62 (1), p. 75–83. DOI: 10.3767/000651917X695443.

### Alismataceae

**Arrigo, Nils; Buerki, Sven; Sarr, Anouk; Guadagnuolo, Roberto; Kozlowski, Gregor (2011):**

**Phylogenetics and Phylogeography of the monocot Genus *Baldellia* (Alismataceae): Mediterranean refugia, suture zones and Implications for conservation.**

In: Molecular Phylogenetics and Evolution 58 (1), p. 33–42. DOI: 10.1016/j.ympev.2010.11.009.

**Chen, Ling-Yun; Chen, Jin-Ming; Gituru, Robert Wahiti; Temam, Tamru Demsis; Wang, Qing-Feng (2012):**

**Generic Phylogeny and historical Biogeography of Alismataceae, inferred from multiple DNA Sequences.**

In: Molecular Phylogenetics and Evolution 63 (2), p. 407–416. DOI: 10.1016/j.ympev.2012.01.016.

**Jacobson, A.; Hedrén, Mikael (2007):**

**Phylogenetic Relationships in *Alisma* (Alismataceae) based on RAPDs, and Sequence Data from ITS and trnL.**

In: Plant Systematics and Evolution 265 (1-2), p. 27–44. DOI: 10.1007/s00606-006-0514-x.

**Lehtonen, Samuli (2006):**

**Phylogenetics of *Echinodorus* (Alismataceae) based on morphological Data.**

In: Botanical Journal of the Linnean Society 150 (3), p. 291–305. DOI: 10.1111/j.1095-8339.2006.00478.x.

**Lehtonen, Samuli (2017):**

**Splitting *Caldesia* in favour of *Albidella* (Alismataceae).**

In: Australian Systematic Botany 30 (1), p. 64. DOI: 10.1071/SB16050.

**Lehtonen, Samuli; Myllys, Leena (2008):**

**Cladistic Analysis of *Echinodorus* (Alismataceae): simultaneous Analysis of molecular and morphological Data.**

In: Cladistics 24 (2), p. 218–239. DOI: 10.1111/j.1096-0031.2007.00177.x.

**Liao, Yi-Ying; Gichira, Andrew Wanyoike; Wang, Qing-Feng; Chen, Jin-Ming (2016):**

**Molecular Phylogeography of four endemic *Sagittaria* Species (Alismataceae) in the Sino-Japanese Floristic Region of East Asia.**

In: Botanical Journal of the Linnean Society 180 (1), p. 6–20. DOI: 10.1111/boj.12351.

### Alismatales

**Chen, Ling-Yun; Chen, Jin-Ming; Gituru, Robert Wahiti; Wang, Qing-Feng (2013):**

**Eurasian Origin of Alismatidae inferred from statistical Dispersal-vicariance Analysis.**

In: Molecular Phylogenetics and Evolution 67 (1), p. 38–42. DOI: 10.1016/j.ympev.2013.01.001.

**Petersen, Gitte; Seberg, Ole; Cuenca, Argelia; Stevenson, Dennis Wm.; Thadeo, Marcela; Davis, Jerrold I. et al. (2016):**

**Phylogeny of the Alismatales (Monocotyledons) and the Relationship of *Acorus* (Acorales?).**

In: Cladistics 32 (2), p. 141–159. DOI: 10.1111/cla.12120.

### Alliaceae

**Chase, Mark W.; Reveal, James L.; Fay, Michael F. (2009):**

**A subfamilial Classification for the expanded asparagalean families Amaryllidaceae, Asparagaceae and Xanthorrhoeaceae.**

In: Botanical Journal of the Linnean Society 161 (2), p. 132–136. DOI: 10.1111/j.1095-8339.2009.00999.x.

**Choi, Hyeok Jae; Giussani, Liliana M.; Jang, Chang-Gee; Oh, Byoung Un; Cota-Sánchez, J. Hugo (2012):**

**Systematics of disjunct northeastern Asian and northern North American *Allium* (Amaryllidaceae).**
In: Botany 90 (6), p. 491–508. DOI: 10.1139/B2012-031.

DENIZ, İSMAIL GÖKHAN; GENÇ, İLKER; SARI, DUYGU (2015):
**Morphological and molecular Data reveal a new Species of *Allium* (Amaryllidaceae) from Sw Anatolia, Turkey.**
In: Phytotaxa 212 (4), p. 283. DOI: 10.11646/phytotaxa.212.4.4.

ESCOBAR, INELIA; NOVOA, PATRICIO; RUIZ, EDUARDO; NEGRITTO, MARÍA A.; BAEZA, CARLOS M. (2010):
**Nuevo hallazgo de *Miersia cornuta* Phil. (Gilliesieae-Alliaceae).**
In: Gayana Botánica 67 (1), p. 130–134. DOI: 10.4067/S0717-66432010000100012.

FAY, MICHAEL F.; RUDALL, PAULA J.; CHASE, MARK W. (2006):
**Molecular Studies of Subfamily Gilliesioideae (Alliaceae).**
In: Aliso 22, p. 367–371.

FRIESEN, NIKOLAI; FRITSCH, REINHARD M.; BLATTNER, FRANK R. (2006):
**Phylogeny and new Intrageneric Classification of *Allium* (Alliaceae) based on Nuclear Ribosomal DNA ITS Sequences.**
In: Aliso 22, p. 372–395.

FRIESEN, NIKOLAI; HERDEN, TOBIAS; SCHOENFELDER, PETER (2015):
***Allium canariense* (Amaryllidaceae), a Species endemic to the Canary Islands.**
In: Phytotaxa 221 (1), p. 1. DOI: 10.11646/phytotaxa.221.1.1.

GURUSHIDZE, MAIA; FRITSCH, REINHARD M.; BLATTNER, FRANK R. (2008):
**Phylogenetic Analysis of *Allium* Subg. Melanocrommyum infers cryptic Species and demands a new Sectional Classification.**
In: Molecular Phylogenetics and Evolution 49 (3), p. 997–1007. DOI: 10.1016/j.ympev.2008.09.003.

GURUSHIDZE, MAIA; FRITSCH, REINHARD M.; BLATTNER, FRANK R. (2010):
**Species-level Phylogeny of *Allium* Subgenus Melanocrommyum: Incomplete Lineage sorting, Hybridization and trnF Gene duplication.**
In: Taxon 59 (3), p. 829–840. DOI: 10.1002/tax.593012.

GURUSHIDZE, MAIA; MASHAYEKHI, SAEIDEH; BLATTNER, FRANK R.; FRIESEN, NIKOLAI; FRITSCH, REINHARD M. (2007):
**Phylogenetic Relationships of wild and cultivated Species of *Allium* Section Cepa inferred by nuclear rDNA ITS Sequence Analysis.**
In: Plant Systematics and Evolution 269 (3-4), p. 259–269. DOI: 10.1007/s00606-007-0596-0.

HERDEN, TOBIAS; HANELT, PETER; FRIESEN, NIKOLAI (2016):
**Phylogeny of *Allium* L. Subgenus Anguinum (G. Don. ex W.d.j. Koch) N. Friesen (Amaryllidaceae).**
In: Molecular Phylogenetics and Evolution 95, p. 79–93. DOI: 10.1016/j.ympev.2015.11.004.

HUANG, DE-QING; YANG, JING-TIAN; ZHOU, CHUN-JING; ZHOU, SONG-DONG; HE, XING-JIN (2014):
**Phylogenetic reappraisal of *Allium* Subgenus Cyathophora (Amaryllidaceae) and related taxa, with a proposal of two new Sections.**
In: Journal of Plant Research 127 (2), p. 275–286. DOI: 10.1007/s10265-013-0617-8.

JARA-ARANCIO, PAOLA; ARROYO, MARY T.K.; GUERRERO, PABLO C.; HINOJOSA, LUIS F.; ARANCIO, GINA; MÉNDEZ, MARCO A. (2014):
**Phylogenetic Perspectives on biome shifts in *Leucocoryne* (Alliaceae) in relation to climatic niche Evolution in western South America.**

In: Journal of Biogeography 41 (2), p. 328–338. DOI: 10.1111/jbi.12186.

**Li, Min-Jie; Tan, Jin-Bo; Xie, Deng-Feng; Huang, De-Qing; Gao, Yun-Dong; He, Xing-Jin (2016):**
**Revisiting the evolutionary events in *Allium* Subgenus Cyathophora (Amaryllidaceae): Insights into the effect of the Hengduan Mountains Region (HMR) uplift and Quaternary climatic fluctuations to the environmental Changes in the Qinghai-Tibet Plateau.**
In: Molecular Phylogenetics and Evolution 94 (Pt B), p. 802–813. DOI: 10.1016/j.ympev.2015.10.002.

**Li, Qin-Qin; Zhou, Song-Dong; He, Xing-Jin; Yu, Yang; Zhang, Yu-Cheng; Wei, Xian-Qin (2010):**
**Phylogeny and Biogeography of *Allium* (Amaryllidaceae: Allieae) based on nuclear ribosomal Internal Transcribed Spacer and Chloroplast rps16 Sequences, focusing on the inclusion of Species endemic to China.**
In: Annals of Botany 106 (5), p. 709–733. DOI: 10.1093/aob/mcq177.

**Memariani, Farshid; Joharchi, Mohammad Reza; Arjmandi, Ali Asghar (2012):**
***Allium aladaghense* (Amaryllidaceae, Allieae), a new Species of Section Asteroprason from northeast of Iran.**
In: Phytotaxa 56, p. 28–34.

**Nguyen, Nhu H.; Driscoll, Heather E.; Specht, Chelsea D. (2008):**
**A molecular Phylogeny of the wild onions (*Allium*; Alliaceae) with a focus on the western North American center of diversity.**
In: Molecular Phylogenetics and Evolution 47 (3), p. 1157–1172. DOI: 10.1016/j.ympev.2007.12.006.

**Sassone, Agostina Belen; Belgrano, Manuel J.; Guaglianone, Encarnación Rosa (2015):**
**The Reinstatement of *Latace* Phil. (Amaryllidaceae, Allioideae).**
In: Phytotaxa 239 (3), p. 253. DOI: 10.11646/phytotaxa.239.3.6.

**Sassone, Agostina Belen; Giussani, Liliana M.; Guaglianone, Encarnación Rosa (2014):**
***Beauverdia*, a resurrected Genus of Amaryllidaceae (Allioideae, Gilliesieae).**
In: Systematic Botany 39 (3), p. 767–775. DOI: 10.1600/036364414X681527.

**Sassone, Agostina Belen; Giussani, Liliana M.; Guaglianone, Encarnación Rosa (2013):**
**Multivariate studies of *Ipheion* (Amaryllidaceae, Allioideae) and related Genera.**
In: Plant Systematics and Evolution 299 (8), p. 1561–1575. DOI: 10.1007/s00606-013-0819-5.

**Seregin, Alexey P.; Anačkov, Goran; Friesen, Nikolai (2015):**
**Molecular and morphological Revision of the *Allium saxatile* Group (Amaryllidaceae): geographical isolation as the driving force of underestimated speciation.**
In: Botanical Journal of the Linnean Society 178 (1), p. 67–101. DOI: 10.1111/boj.12269.

**Sinitsyna, Tatiana A.; Herden, Tobias; Friesen, Nikolai (2016):**
**Dated Phylogeny and Biogeography of the Eurasian *Allium* Section Rhizirideum (Amaryllidaceae).**
In: Plant Systematics and Evolution 302 (9), p. 1311–1328. DOI: 10.1007/s00606-016-1333-3.

**Souza, Gustavo; Crosa, Orfeo; Guerra, Marcelo (2015):**
**Karyological, morphological, and phylogenetic Diversification in *Leucocoryne* Lindl. (Allioideae, Amaryllidaceae).**
In: Plant Systematics and Evolution 301 (8), p. 2013–2023. DOI: 10.1007/s00606-015-1216-z.

**Stafford, Gary I.; Wikkelsø, Mette J.; Nancke, Louise; Jäger, Anna K.; Möller, Michael; Rønsted, Nina (2016):**

**The first phylogenetic Hypothesis for the southern African endemic Genus *Tulbaghia* (Amaryllidaceae, Allioideae) based on Plastid and nuclear DNA Sequences.**
In: Botanical Journal of the Linnean Society 181 (2), p. 156–170. DOI: 10.1111/boj.12417.

**WHEELER, ERICA J.; MASHAYEKHI, SAEIDEH; MCNEAL, DALE W.; COLUMBUS, TRAVIS J.; PIRES, J. CHRIS (2013):**
**Molecular Systematics of *Allium* Subgenus Amerallium (Amaryllidaceae) in North America.**
In: American Journal of Botany 100 (4), p. 701–711. DOI: 10.3732/ajb.1200641.

### Alstroemeriaceae

**ALZATE, FERNANDO; MORT, MARK E.; RAMIREZ, MONICA (2008):**
**Phylogenetic Analyses of *Bomarea* (Alstroemeriaceae) based on combined Analyses of nrDNA ITS, psbA-trnH, rpoB-trnC and matK Sequences.**
In: Taxon 57 (3), p. 853–862. DOI: 10.1002/tax.573014.

**ALZATE, FERNANDO; QUIJANO-ABRIL, MARIO ALBERTO; MORRONE, JUAN J. (2008):**
**Panbiogeographical Analysis of the Genus *Bomarea* (Alstroemeriaceae).**
In: Journal of Biogeography 35 (7), p. 1250–1257. DOI: 10.1111/j.1365-2699.2008.01896.x.

**CHACÓN, JULIANA; SOUSA, ARETUZA; BAEZA, CARLOS M.; RENNER, SUSANNE P. (2012):**
**Ribosomal DNA Distribution and a Genus-wide Phylogeny reveal Patterns of chromosomal Evolution in *Alstroemeria* (Alstroemeriaceae).**
In: American Journal of Botany 99 (9), p. 1501–1512. DOI: 10.3732/ajb.1200104.

**SANSO, A. MARIEL; AAGESEN, LONE; XIFREDA, CECILIA C. (2014):**
**Foliar Anatomy and microMorphology of southern South American Alstroemeriaceae: Alstroemerieae, and its systematic Implications in *Alstroemeria*.**
In: Nordic Journal of Botany 32 (6), p. 731–743. DOI: 10.1111/njb.00470.

### Altingiaceae

**ICKERT-BOND, STEFANIE M.; WEN, JUN (2006):**
**Phylogeny and Biogeography of Altingiaceae: Evidence from combined Analysis of five non-coding Chloroplast Regions.**
In: Molecular Phylogenetics and Evolution 39 (2), p. 512–528. DOI: 10.1016/j.ympev.2005.12.003.

**SHI, S.; HUANG, Y.; ZHONG, YANG; DU, Y.; ZHANG, QIANG; CHANG, H.; BOUFFORD, D.E. (2001):**
**Phylogeny of the Altingiaceae based on cpDNA matK, Py-igs and nrDNA ITS Sequences.**
In: Plant Systematics and Evolution 230, p. 13–24.

### Amaranthaceae

**AKHANI, HOSSEIN; EDWARDS, GERALD E.; ROALSON, ERIC H. (2007):**
**Diversification of the Old World Salsoleae s.l. (Chenopodiaceae): molecular phylogenetic Analysis of Nuclear and Chloroplast Data Sets and a revised Classification.**
In: International Journal of Plant Sciences 168 (6), p. 931–956.

**APPLEQUIST, WENDY L.; PRATT, DONALD B. (2005):**
**The Malagasy Endemic *Dendroportulaca* (Portulacaceae) is referable to *Deeringia* (Amaranthaceae): molecular and morphological Evidence.**
In: Taxon 54 (3), p. 681–687. DOI: 10.2307/25065424.

**BRANDT, RONNY; LOMONOSOVA, MARIA; WEISING, KURT; WAGNER, NATASCHA; FREITAG, HELMUT (2015):**

**Phylogeny and Biogeography of *Suaeda* Subg. Brezia (Chenopodiaceae/Amaranthaceae) in the Americas.**
In: Plant Systematics and Evolution 301 (10), p. 2351–2375. DOI: 10.1007/s00606-015-1233-y.

**Brullo, Cristian; Brullo, Salvatore; Gaskin, John F.; Galdo, Gianpietro Giusso Del; Hrusa, G. Frederic; Salmeri, Cristina (2015):**
**A new Species of *Kali* (Salsoloideae, Chenopodiaceae) from Sicily, supported by molecular Analysis.**
In: Phytotaxa 201 (4), p. 256. DOI: 10.11646/phytotaxa.201.4.2.

**Cabrera, Jonathan F.; Jacobs, Surrey W.L.; Kadereit, Gudrun (2009):**
**Phylogeny of the Australian Camphorosmeae (Chenopodiaceae) and the taxonomic Significance of the Fruiting Perianth.**
In: International Journal of Plant Sciences 170 (4), p. 505–521. DOI: 10.1086/597267.

**del-Pino, Ivonne Sánchez; Borsch, Thomas; Motley, Timothy J. (2009):**
**trnL-F and rpl16 Sequence Data and dense Taxon Sampling reveal Monophyly of Unilocular Anthered Gomphrenoideae (Amaranthaceae) and an improved Picture of their internal Relationships.**
In: Systematic Botany 34 (1), p. 57–67. DOI: 10.1600/036364409787602401.

**Fuentes-Bazán, Susy; Mansion, Guilhem; Borsch, Thomas (2012):**
**Towards a Species level Tree of the globally diverse Genus *Chenopodium* (Chenopodiaceae).**
In: Molecular Phylogenetics and Evolution 62 (1), p. 359–374. DOI: 10.1016/j.ympev.2011.10.006.

**Fuentes-Bazán, Susy; Uotila, Pertti; Borsch, Thomas (2012):**
**A novel Phylogeny-based generic Classification for *Chenopodium* sensu lato, and a tribal rearrangement of Chenopodioideae (Chenopodiaceae).**
In: Willdenowia 42 (1), p. 5–24. DOI: 10.3372/wi.42.42101.

**Hammer, Timothy; Davis, Robert; Thiele, Kevin R. (2015):**
**A molecular Famework Phylogeny for *Ptilotus* (Amaranthaceae): Evidence for the rapid Diversification of an arid Australian Genus.**
In: Taxon 64 (2), p. 272–285. DOI: 10.12705/642.6.

**Heklau, Heike; Röser, Martin (2008):**
**Delineation, Taxonomy and phylogenetic Relationships of the Genus *Krascheninnikovia* (Amaranthaceae Subtribe Axyridinae).**
In: Taxon 57 (2), p. 563–576.

**Hohmann, Sandra; Kadereit, Joachim W.; Kadereit, Gudrun (2006):**
**Understanding Mediterranean-Californian disjunctions: molecular Evidence from Chenopodiaceae-Betoideae.**
In: Taxon 55 (1), p. 67–78.

**Kadereit, Gudrun; Ball, Peter; Beer, Svetlana; Mucina, Ladislav; Sokoloff, Dmitry D.; Teege, Patrick et al. (2007):**
**A taxonomic Nightmare Comes True: Phylogeny and Biogeography of Glassworts (*Salicornia* L., Chenopodiaceae).**
In: Taxon 56 (4), p. 1143–1170. DOI: 10.2307/25065909.

**Kadereit, Gudrun; Borsch, Thomas; Weising, Kurt; Freitag, Helmut (2003):**

**Phylogeny of Amaranthaceae and Chenopodiaceae and the Evolution of C4 Photosynthesis.**
In: International Journal of Plant Sciences 164 (6).

**Kadereit, Gudrun; Freitag, Helmut (2011):**
**Molecular Phylogeny of Camphorosmeae (Camphorosmoideae, Chenopodiaceae): Implications for Biogeography, Evolution of C4-photosynthesis and Taxonomy.**
In: Taxon 60 (1), p. 51–78.

**Kadereit, Gudrun; Mavrodiev, Evgeny V.; Zacharias, Elizabeth H.; Sukhorukov, Alexander P. (2010):**
**Molecular Phylogeny of Atripliceae (Chenopodioideae, Chenopodiaceae): Implications for Systematics, Biogeography, flower and Fruit Evolution, and the Origin of C4 photosynthesis.**
In: American Journal of Botany 97 (10), p. 1664–1687. DOI: 10.3732/ajb.1000169.

**Kadereit, Gudrun; Mucina, Ladislav; Freitag, Helmut (2006):**
**Phylogeny of Salicornioideae (Chenopodiaceae): Diversification, Biogeography, and evolutionary Trends in Leaf and Flower Morphology.**
In: Taxon 55 (3), p. 617–642. DOI: 10.2307/25065639.

**Kadereit, Gudrun; Piirainen, Mikko; Lambinon, Jacques; Vanderpoorten, Alain (2012):**
**Cryptic Taxa should have names: Reflections in the glasswort Genus *Salicornia* (Amaranthaceae).**
In: Taxon 61 (6), p. 1227–1239.

**Kapralov, Maxim V.; Akhani, Hossein; Voznesenskaya, Elena V.; Edwards, Gerald E.; Franceschi, Vincent; Roalson, Eric H. (2006):**
**Phylogenetic Relationships in the Salicornioideae / Suaedoideae / Salsoloideae s.l. (Chenopodiaceae) Clade and a Clarification of the phylogenetic Position of *Bienertia* and *Alexandra* using multiple DNA Sequence Datasets.**
In: Systematic Botany 31 (3), p. 571–585.

**Kolano, Bozena; McCann, Jamie; Orzechowska, Maja; Siwinska, Dorota; Temsch, Eva M.; Weiss-Schneeweiss, Hanna (2016):**
**Molecular and cytogenetic Evidence for an allotetraploid Origin of *Chenopodium quinoa* and *C. berlandieri* (Amaranthaceae).**
In: Molecular Phylogenetics and Evolution 100, p. 109–123. DOI: 10.1016/j.ympev.2016.04.009.

**Kolano, Bozena; Siwinska, Dorota; McCann, Jamie; Weiss-Schneeweiss, Hanna (2015):**
**The Evolution of Genome size and rDNA in diploid Species of *Chenopodium* s.l. (Amaranthaceae).**
In: Botanical Journal of the Linnean Society 179 (2), p. 218–235. DOI: 10.1111/boj.12321.

**Lamonico, Duilio; Jarvis, Charles E. (2012):**
**Lectotypification of two Linnaean names in *Chenopodium* L. (Chenopodiaceae).**
In: Taxon 61 (4), p. 864–865.

**Lee, Kok K.; Harrison, Dion K.; Johnston, Margaret E.; Williams, Richard R. (2007):**
**Molecular taxonomic clarification of *Ptilotus exaltatus* and *Ptilotus nobilis* (Amaranthaceae).**
In: Australian Systematic Botany 20 (1), p. 72–81. DOI: 10.1071/SB06010.

**Masson, Rüdiger; Kadereit, Gudrun (2013):**

**Phylogeny of Polycnemoideae (Amaranthaceae): Implications for Biogeography, Character Evolution and Taxonomy.**
In: Taxon 62 (1), p. 100–111.

**MCCAULEY, ROSS A.; BALLARD, HARVEY EUGENE (2007):**
**Systematics of North American *Froelichia* (Amaranthaceae Subfam. Gomphrenoideae) Ii: Phylogeny and biogeographic Speciation Patterns inferred from nrITS Sequence Data.**
In: Brittonia 59 (3), p. 275–289. DOI: 10.1663/0007-196X(2007)59[275:SONAFA]2.0.CO;2.

**MOLERO, J.; MONTSERRAT, J.M. (2006):**
**Quenopodiáceas nuevas o raras para la Flora de Marruecos.**
In: Lagascalia 26, p. 7–24.

**MOSYAKIN, SERGEI L. (2013):**
**New nomenclatural Combinations in *Blitum, Oxybasis, Chenopodiastrum*, and *Lipandra* (Chenopodiaceae).**
In: Phytoneuron 56, p. 1–8.

**MURAKEÖZY, E. P.; AÏNOUCHE, ABDELKADER; MEUDEC, A.; DESLANDES, E.; POUPART, N. (2007):**
**Phylogenetic Relationships and genetic Diversity of the Salicornieae (Chenopodiaceae) native to the Atlantic coasts of France.**
In: Plant Systematics and Evolution 264 (3-4), p. 217–237. DOI: 10.1007/s00606-006-0511-0.

**PIIRAINEN, MIKKO; LIEBISCH, OSKAR; KADEREIT, GUDRUN (2017):**
**Phylogeny, Biogeography, Systematics and Taxonomy of Salicornioideae (Amaranthaceae/Chenopodiaceae) – A cosmopolitan, highly specialized hygrohalophyte Lineage dating back to the Oligocene.**
In: Taxon 66 (1), p. 109–132. DOI: 10.12705/661.6.

**PINO, IVONNE SÁNCHEZ-DEL; MOTLEY, TIMOTHY J. (2010):**
**Evolution of *Tidestromia* (Amaranthaceae) in the deserts of the southwestern United States and Mexico.**
In: Taxon 59 (1), p. 38–48.

**PINO, IVONNE SÁNCHEZ-DEL; MOTLEY, TIMOTHY J.; BORSCH, THOMAS (2012):**
**Molecular Phylogenetics of *Alternanthera* (Gomphrenoideae, Amaranthaceae): resolving a complex taxonomic History caused by different interpretations of morphological Characters in a Lineage with C4 and C3–c4 intermediate Species.**
In: Botanical Journal of the Linnean Society 169, p. 493–517.

**RAAB-STRAUBE, ECKHARD (2013):**
**Euro+Med-Checklist Notulae, 2.**
In: Willdenowia 43 (2), p. 239–249. DOI: 10.3372/wi.43.43202.

**ROMEIRAS, MARIA M.; VIEIRA, ANA; SILVA, DIOGO N.; MOURA, MÓNICA; SANTOS-GUERRA, ARNOLDO; BATISTA, DORA ET AL. (2016):**
**Evolutionary and Biogeographic Insights on the Macaronesian *Beta patellifolia* Species (Amaranthaceae) from a Time-Scaled molecular Phylogeny.**
In: Public Library of Science One 11 (3), e0152456. DOI: 10.1371/journal.pone.0152456.

**SCHÜTZE, P.; FREITAG, HELMUT; WEISING, KURT (2003):**
**An integrated molecular and morphological Study of the Subfamily Suaedoideae Ulbr. (Chenopodiaceae).**

In: Plant Systematics and Evolution 239 (3-4), p. 257–286. DOI: 10.1007/s00606-003-0013-2.

SHEPHERD, KELLY A.; WAYCOTT, MICHELLE; CALLADINE, AINSLEY (2004):
**Radiation of the Australian Salicornioideae (Chenopodiaceae) - based on Evidence from Nuclear and Chloroplast DNA Sequences.**
In: American Journal of Botany 91 (9), p. 1387–1397.

SHEPHERD, KELLY A.; WILSON, PAUL G. (2007):
**Incorporation of the Australian Genera *Halosarcia, Pachycornia, Sclerostegia* and *Tegicornia* into *Tecticornia* (Salicornioideae, Chenopodiaceae).**
In: Australian Systematic Botany 20 (4), p. 319–331. DOI: 10.1071/SB07002.

STEFFEN, SIMONE; BALL, PETER; MUCINA, LADISLAV; KADEREIT, GUDRUN (2015):
**Phylogeny, Biogeography and ecological Diversification of *Sarcocornia* (Salicornioideae, Amaranthaceae).**
In: Annals of Botany 115 (3), p. 353–368. DOI: 10.1093/aob/mcu260.

STETTER, MARKUS G.; SCHMID, KARL J. (2017):
**Analysis of phylogenetic Relationships and Genome size Evolution of the *Amaranthus* Genus using GBS indicates the ancestors of an ancient crop.**
In: Molecular Phylogenetics and Evolution 109, p. 80–92. DOI: 10.1016/j.ympev.2016.12.029.

SUKHORUKOV, ALEXANDER P.; KUSHUNINA, MARIA (2015):
**Corrigenda to "taxonomic Revision of Chenopodiaceae in Nepal" [Phytotaxa 191: 10–44. 2014].**
In: Phytotaxa 226 (3), p. 288. DOI: 10.11646/phytotaxa.226.3.10.

THIV, MIKE; THULIN, MATS; KILIAN, NORBERT; LINDER, HANS PETER (2006):
**Eritreo-Arabian Affinities of the Socotran Flora as Revealed from the molecular Phylogeny of *Aerva* (Amaranthaceae).**
In: Systematic Botany 31 (3), p. 560–570.

THULIN, MATS; RYDBERG, ANDERS; THIEDE, JOACHIM (2010):
**Identity of *Tetragonia pentandra* and Taxonomy and Distribution of *Patellifolia* (Chenopodiaceae).**
In: Willdenowia 40 (1), p. 5–11. DOI: 10.3372/wi.40.40101.

WEN, ZHI-BIN; ZHANG, MING-LI; ZHU, GE-LIN; SANDERSON, STEWART C. (2010):
**Phylogeny of Salsoleae s.l. (Chenopodiaceae) based on DNA Sequence Data from ITS, psbB–psbH, and rbcL, with Emphasis on Taxa of northwestern China.**
In: Plant Systematics and Evolution 288 (1-2), p. 25–42. DOI: 10.1007/s00606-010-0310-5.

XU, F.; SUN, M. (2001):
**Comparative Analysis of phylogenetic Relationships of grain amaranths and their wild relatives (Amaranthus; Amaranthaceae) using internal transcribed spacer, amplified fragment length polymorphism, and double-primer fluorescent intersimple Sequence repeat Markers.**
In: Molecular Phylogenetics and Evolution 21 (3), p. 372–387. DOI: 10.1006/mpev.2001.1016.

ZACHARIAS, ELIZABETH H.; BALDWIN, BRUCE G. (2010):
**A molecular Phylogeny of North American Atripliceae (Chenopodiaceae), with Implications for Floral and Photosynthetic Pathway Evolution.**
In: Systematic Botany 35 (4), p. 839–857. DOI: 10.1600/036364410X539907.

### Amaryllidaceae

**Bay-Smidt, M.G.K.; Jäger, Anna K.; Krydsfeldt, K.; Meerow, Alan W.; Stafford, Gary I.; van Staden, J.; Rønsted, Nina (2011):**

**Phylogenetic selection of target Species in Amaryllidaceae Tribe Haemantheae for acetylcholinesterase inhibition and affinity to the serotonin reuptake transport protein.**

In: South African Journal of Botany 77 (1), p. 175–183. DOI: 10.1016/j.sajb.2010.07.016.

**Bjorå, Charlotte Sletten; Kwembeya, Ezekiel G.; Bogner, Josef; Nordal, Inger (2009):**

**Geophytes diverging in rivers-a Study on the Genus *Crinum*, with two new rheophytic Taxa from Cameroon.**

In: Taxon 58 (2), p. 561–571. DOI: 10.1002/tax.582020.

**Bush, Catherine M.; Rollins, Dollie; Smith, Gerald L. (2010):**

**The Phylogeny of the Southeastern United States *Hymenocallis* (Amaryllidaceae) based on ISSR Fingerprinting and morphological Data.**

In: Castanea 75 (3), p. 368–380. DOI: 10.2179/09-012.1.

**Castro, Olga; Brullo, Salvatore; Colombo, Paolo; Jury, Stephen L.; Luca, Paolo; Di Maio, Antonietta (2012):**

**Phylogenetic and biogeographical Inferences for *Pancratium* (Amaryllidaceae), with an Emphasis on the Mediterranean Species based on Plastid Sequence Data.**

In: Botanical Journal of the Linnean Society 170 (1), p. 12–28. DOI: 10.1111/j.1095-8339.2012.01268.x.

**Gage, Ewan; Wilkin, Paul; Chase, Mark W.; Hawkins, Julie A. (2011):**

**Phylogenetic Systematics of *Sternbergia* (Amaryllidaceae) based on Plastid and ITS Sequence Data.**

In: Botanical Journal of the Linnean Society 166 (2), p. 149–162. DOI: 10.1111/j.1095-8339.2011.01138.x.

**García, Nicolás; Meerow, Alan W.; Soltis, Douglas E.; Soltis, Pamela p. (2014):**

**Testing deep reticulate Evolution in Amaryllidaceae Tribe Hippeastreae (Asparagales) with ITS and Chloroplast Sequence Data.**

In: Systematic Botany 39 (1), p. 75–89. DOI: 10.1600/036364414X678099.

**Graham, Sean W.; Barrett, Spencer C. H. (2004):**

**Phylogenetic Reconstruction of the Evolution of stylar polymorphisms in *Narcissus* (Amaryllidaceae).**

In: American Journal of Botany 91 (7), p. 1007–1021. DOI: 10.3732/ajb.91.7.1007.

**Kwembeya, Ezekiel G.; Bjorå, Charlotte Sletten; Stedje, Brita; Nordal, Inger (2007):**

**Phylogenetic Relationships in the Genus *Crinum* (Amaryllidaceae) with Emphasis on tropical African species: Evidence from trnL-F and nuclear ITS DNA Sequence Data.**

In: Taxon 56 (3), p. 801–810. DOI: 10.2307/25065862.

**Lled, Ma D.; Davis, Aaron P.; Crespo, Manuel B.; Chase, Mark W.; Fay, Michael F. (2004):**

**Phylogenetic Analysis of *Leucojum* and *Galanthus* (Amaryllidaceae) based on Plastid matK and nuclear ribosomal spacer (ITS) DNA Sequences and Morphology.**

In: Plant Systematics and Evolution 246 (3-4), p. 223–243. DOI: 10.1007/s00606-004-0152-0.

**Meerow, Alan W.; Clayton, Jason R. (2004):**

**Generic Relationships among the baccate-fruited Amaryllidaceae (tribe Haemantheae) inferred from Plastid and nuclear non-coding DNA Sequences.**

In: Plant Systematics and Evolution 244 (3-4), p. 141–155. DOI: 10.1007/s00606-003-0085-z.

**Meerow, Alan W.; Francisco-Ortega, Javier; Kuhn, David N.; Schnell, Raymond J. (2006):**

**Phylogenetic Relationships and Biogeography within the Eurasian Clade of Amaryllidaceae based on Plastid ndhF and nrDNA ITS Sequences: Lineage Sorting in a Reticulate Area?**
In: Systematic Botany 31 (1), p. 42–60.

**MEEROW, ALAN W.; GUY, CHARLES L.; LI, QIN-BAO; YANG, SI-LIN (2000):**
**Phylogeny of the American Amaryllidaceae based on nrDNA ITS Sequences.**
In: Systematic Botany 25 (4), p. 708–726. DOI: 10.2307/2666729.

**MEEROW, ALAN W.; JOST, LOU; OLEAS, NORA (2015):**
**Two new Species of endemic Ecuadorean Amaryllidaceae (Asparagales, Amaryllidaceae, Amarylloideae, Eucharideae).**
In: PhytoKeys (48), p. 1–9. DOI: 10.3897/phytokeys.48.4399.

**MEEROW, ALAN W.; LEHMILLER, DAVID J.; CLAYTON, JASON R. (2003):**
**Phylogeny and Biogeography of *Crinum* L. (Amaryllidaceae) inferred from nuclear and limited Plastid non-coding DNA Sequences.**
In: Botanical Journal of the Linnean Society 141 (3), p. 349–363. DOI: 10.1046/j.1095-8339.2003.00142.x.

**MEEROW, ALAN W.; SNUMAN, DEIRDRE (2006):**
**The Never-ending Story: Multigene Approaches to the Phylogeny of Amaryllidaceae.**
In: Aliso 22 (1), p. 355–366. DOI: 10.5642/aliso.20062201.29.

**MUÑOZ, MANUEL; RIEGEL, RICARDO; SEEMANN, PETER; PEÑAILILLO, PATRICIO; SCHIAPPACASSE, FLAVIA; NÚÑEZ, JOSÉ (2011):**
**Relaciones filogenéticas de *Rhodolirium montanum* Phil. y especies afines, basadas en secuencias nucleotídicas de la región ITS y análisis cariotípico.**
In: Gayana Botánica 68 (1), p. 40–48. DOI: 10.4067/S0717-66432011000100005.

**RAN, Y. (2001):**
**Phylogenetic Analysis and Karyotype Evolution in the Genus *Clivia* (Amaryllidaceae).**
In: Annals of Botany 87 (6), p. 823–830. DOI: 10.1006/anbo.2001.1422.

**RØNSTED, NINA; ZUBOV, DIMITRI; BRUUN-LUND, SAM; DAVIS, AARON P. (2013):**
**Snowdrops falling slowly into place: an improved Phylogeny for *Galanthus* (Amaryllidaceae).**
In: Molecular Phylogenetics and Evolution 69 (1), p. 205–217. DOI: 10.1016/j.ympev.2013.05.019.

**SANTOS-GALLY, ROCÍO; VARGAS, PABLO; ARROYO, JUAN (2012):**
**Insights into Neogene Mediterranean Biogeography based on phylogenetic Relationships of mountain and lowland Lineages of *Narcissus* (Amaryllidaceae).**
In: Journal of Biogeography 39 (4), p. 782–798. DOI: 10.1111/j.1365-2699.2011.02526.x.

**SHI, SHUDE; SUN, YAQING; WEI, LEI; LEI, XUEFENG; CAMERON, KENNETH M.; FU, CHENG-XIN (2014):**
**Plastid DNA Sequence Data help to clarify phylogenetic Relationships and reticulate Evolution in *Lycoris* (Amaryllidaceae).**
In: Botanical Journal of the Linnean Society 31, p. 115–126. DOI: 10.1111/boj.12198.

**TAŞCI MARGOZ, NIVART; YÜZBAŞIOĞLU, SIRRI İBRAHIM; ÇELEN, ZEYNEP; EKÍM, TUNA; BILGIN, AYŞE NEŞE (2013):**
**Molecular Phylogeny of *Galanthus* (Amaryllidaceae) of Anatolia inferred from multiple nuclear and Chloroplast DNA Regions.**
In: Turkish Journal of Botany 37, p. 993–1007. DOI: 10.3906/bot-1209-41.

**WEICHHARDT-KULESSA, KATJA; BÖRNER, THOMAS; SCHMITZ, JÜRGEN; MÜLLER-DOBLIES, UTE; MÜLLER-DOBLIES, DIETRICH (2000):**

**Controversial Taxonomy of Strumariinae (Amaryllidaceae) investigated by nuclear rDNA (ITS) Sequences.**
In: Plant Systematics and Evolution 223 (1-2), p. 1–13. DOI: 10.1007/BF00985323.

**ZONNEVELD, BEN J. M. (2008):**
**The systematic value of nuclear DNA content for all Species of *Narcissus* L. (Amaryllidaceae).**
In: Plant Systematics and Evolution 275 (1-2), p. 109–132. DOI: 10.1007/s00606-008-0015-1.

### Anacampserotaceae

**DREHER, HAGEN; RODGERSON, CHRIS; YOUNG, ANDREW (2014):**
**A new Combination and the Description of a new Species in the Section Avonia of the Genus *Anacampseros* (Anacampserotaceae).**
In: Bradleya 32 (32), p. 105–111. DOI: 10.25223/brad.n32.2014.a4.

**GURALNICK, LONNIE J.; CLINE, AMANDA; SMITH, MONICA; SAGE, ROWAN F. (2008):**
**Evolutionary physiology: the extent of C4 and Cam photosynthesis in the Genera *Anacampseros* and *Grahamia* of the Portulacaceae.**
In: Journal of Experimental Botany 59 (7), p. 1735–1742. DOI: 10.1093/jxb/ern081.

**NYFFELER, RETO (2007):**
**The closest relatives of cacti: Insights from phylogenetic Analyses of Chloroplast and mitochondrial Sequences with special Emphasis on Relationships in the Tribe Anacampseroteae.**
In: American Journal of Botany 94 (1), p. 89–101. DOI: 10.3732/ajb.94.1.89.

### Anacardiaceae

**ANDRÉS-HERNÁNDEZ, AGUSTINA ROSA; TERRAZAS, TERESA; SALAZAR, GERARDO A.; OCHOTERENA, HELGA (2014):**
**Phylogenetic Analysis based on structural and combined Analyses of *Rhus* s.s. (Anacardiaceae).**
In: Botanical Journal of the Linnean Society 176, p. 452–468.

**KAFKAS, SALIH (2006):**
**Phylogenetic Analysis of the Genus *Pistacia* by AFLP Markers.**
In: Plant Systematics and Evolution 262 (1-2), p. 113–124. DOI: 10.1007/s00606-006-0460-7.

**KARIMI, HAMID REZA; KAFKAS, SALIH (2011):**
**Genetic Relationships among *Pistacia* Species studied by SAMPL Markers.**
In: Plant Systematics and Evolution 297 (3-4), p. 207–212. DOI: 10.1007/s00606-011-0508-1.

**MILLER, ALLISON J.; YOUNG, DAVID A.; WEN, JUN (2001):**
**Phylogeny and Biogeography of *Rhus* (Anacardiaceae) based on ITS Sequence Data.**
In: International Journal of Plant Sciences 162 (6), p. 1401–1407.

**MITCHELL, JOHN D.; DALY, DOUGLAS C. (2015):**
**A Revision of *Spondias* L. (Anacardiaceae) in the Neotropics.**
In: PhytoKeys (55), p. 1–92. DOI: 10.3897/phytokeys.55.8489.

**NIE, ZE-LONG; SUN, HANG; MENG, YING; WEN, JUN (2009):**
**Phylogenetic Analysis of *Toxicodendron* (Anacardiaceae) and its biogeographic Implications on the Evolution of north temperate and tropical intercontinental disjunctions.**
In: Journal of Systematics and Evolution 47 (5), p. 416–430. DOI: 10.1111/j.1759-6831.2009.00045.x.

**PAN, YUE-ZHI; GONG, XUN; YANG, YANG (2008):**

**Phylogenetic Position of the Genus *Dobinea*: Evidence from nucleotide Sequences of the Chloroplast Gene rbcL and the nuclear ribosomal ITS Region.**
In: Journal of Systematics and Evolution 46 (4), p. 586–594.

PELL, SUSAN K.; MITCHELL, JOHN D.; LOWRY, PORTER PRESCOTT II.; RANDRIANASOLO, ARMAND; URBATSCH, LOWELL E. (2008):
**Phylogenetic Split of Malagasy and African Taxa of *Protorhus* and *Rhus* (Anacardiaceae) based on cpDNA trnL–trnF and nrDNA ETS and ITS Sequence Data.**
In: Systematic Botany 33 (2), p. 375–383.

SILVA, JÉSSICA NAIANA; BEZERRA DA COSTA, ALINE; SILVA, JOSÉ VIEIRA; ALMEIDA, CICERO (2015):
**DNA barcoding and Phylogeny in Neotropical Species of the Genus *Spondias*.**
In: Biochemical Systematics and Ecology 61, p. 240–243. DOI: 10.1016/j.bse.2015.06.005.

TALEBI, MAJID; KAZEMI, MAHBOUBEH; SAYED-TABATABAEI, B. EBRAHIM (2012):
**Molecular Diversity and phylogenetic Relationships of *Pistacia vera*, *Pistacia atlantica* subsp. mutica and *Pistacia khinjuk* using SRAP Markers.**
In: Biochemical Systematics and Ecology 44, p. 179–185. DOI: 10.1016/j.bse.2012.05.013.

WANNAN, B. P. (2006):
**Analysis of Generic Relationships in Anacardiaceae.**
In: Blumea 51 (1), p. 165–195. DOI: 10.3767/000651906X622427.

XIE, LEI; YANG, ZHI-YUN; WEN, JUN; LI, DE-ZHU; YI, TING-SHUANG (2014):
**Biogeographic History of *Pistacia* (Anacardiaceae), emphasizing the Evolution of the Madrean-Tethyan and the eastern Asian-Tethyan disjunctions.**
In: Molecular Phylogenetics and Evolution 77, p. 136–146. DOI: 10.1016/j.ympev.2014.04.006.

YANG, YING-YING; MENG, YING; WEN, JUN; SUN, HANG; NIE, ZE-LONG (2016):
**Phylogenetic Analyses of *Searsia* (Anacardiaceae) from eastern Asia and its biogeographic disjunction with its African relatives.**
In: South African Journal of Botany 106, p. 129–136. DOI: 10.1016/j.sajb.2016.05.021.

YI, TING-SHUANG; MILLER, ALLISON J.; WEN, JUN (2004):
**Phylogenetic and biogeographic Diversification of *Rhus* (Anacardiaceae) in the Northern Hemisphere.**
In: Molecular Phylogenetics and Evolution 33 (3), p. 861–879. DOI: 10.1016/j.ympev.2004.07.006.

YI, TING-SHUANG; MILLER, ALLISON J.; WEN, JUN (2007):
**Phylogeny of *Rhus* (Anacardiaceae) based on Sequences of Nuclear NIA-I3 Intron and Chloroplast trnC-trnD.**
In: Systematic Botany 32 (2), p. 379–391.

YI, TING-SHUANG; WEN, JUN; GOLAN-GOLDHIRSH, AVI; PARFITT, DAN E. (2008):
**Phylogenetics and reticulate Evolution in *Pistacia* (Anacardiaceae).**
In: American Journal of Botany 95 (2), p. 241–251.

### Ancistrocladaceae

MEIMBERG, HARALD; RISCHER, HEIKO; TURINI, FLORIAN G.; CHAMCHUMROON, VORADOL; DREYER, MICHAEL; SOMMARO, MARCELLA ET AL. (2010):
**Evidence for Species differentiation within the *Ancistrocladus tectorius* complex (Ancistrocladaceae) in Southeast Asia: a molecular approach.**
In: Plant Systematics and Evolution 284 (1-2), p. 77–98. DOI: 10.1007/s00606-009-0241-1.

TURINI, FLORIAN G.; STEINERT, CLAUDIA; HEUBL, GÜNTHER; BRINGMANN, GERHARD; LOMBE, B. KIMBADI; MUDOGO, VIRIMA; MEIMBERG, HARALD (2014):

**Microsatellites facilitate Species Delimitation in Congolese *Ancistrocladus* (Ancistrocladaceae), a Genus with pharmacologically potent naphthylisoquinoline alkaloids.**

In: Taxon 63 (2), p. 329–341. DOI: 10.12705/632.36.

### Anemiaceae

LABIAK, PAULO HENRIQUE; MICKEL, JOHN T.; HANKS, JUDITH GARRISON (2015):

**Molecular Phylogeny and Character Evolution of Anemiaceae (Schizaeales).**

In: Taxon 64 (6), p. 1141–1158. DOI: 10.12705/646.3.

SKOG, JUDITH E.; ZIMMER, ELIZABETH A.; MICKEL, JOHN T. (2002):

**Additional Support for two Subgenera of *Anemia* (Schizaeaceae) from Data for the Chloroplast Intergenic Spacer Region trnL-F and Morphology.**

In: American Fern Journal 92 (2), p. 119–130. DOI: 10.1640/0002-8444(2002)092[0119:ASFTSO]2.0.CO;2.

### Angiospermae

ANGIOSPERM PHYLOGENY GROUP III. THIS PAPER WAS COMPILED BY BIRGITTA BREMER, KÅRE BREMER (2009):

**An update of the Angiosperm Phylogeny Group: Classification for the Orders and Families of flowering Plants: APG.**

In: Botanical Journal of the Linnean Society 161, p. 105–121.

BEAULIEU, JEREMY M.; TANK, DAVID C.; DONOGHUE, MICHAEL J. (2013):

**A Southern Hemisphere Origin for Campanulid Angiosperms, with traces of the break-up of Gondwana.**

In: BMC Evolutionary Biology 13, p. 80. DOI: 10.1186/1471-2148-13-80.

BELL, CHARLES D.; SOLTIS, DOUGLAS E.; SOLTIS, PAMELA P. (2010):

**The age and Diversification of the Angiosperms re-revisited.**

In: American Journal of Botany 97 (8), p. 1296–1303. DOI: 10.3732/ajb.0900346.

BREMER, KARE; BACKLUND, A.; SENNBLAD, BENGT; SWENSON, ULF; ANDREASEN, K.; HJERTSON, M. ET AL. (2001):

**A phylogenetic Analysis of 100+ Genera and 50+ families of euasterids based on morphological and molecular Data with Notes on possible higher level morphological synapomorphies.**

In: Plant Systematics and Evolution 229, p. 137–169.

BURLEIGH, JOHN GORDON; BANSAL, MUKUL S.; EULENSTEIN, OLIVER; HARTMANN, STEFANIE; WEHE, ANDRÉ; VISION, TODD J. (2011):

**Genome-scale Phylogenetics: Inferring the plant Tree of life from 18,896 Gene Trees.**

In: Systematic Biology 60 (2), p. 117–125. DOI: 10.1093/sysbio/syq072.

CAI, ZHENGQIU; PEÑAFLOR, CYNTHIA; KUEHL, JENNIFER V.; LEEBENS-MACK, JAMES H.; CARLSON, JOHN E.; DEPAMPHILIS, CLAUDE W. ET AL. (2006):

**Complete Plastid Genome Sequences of *Drimys, Liriodendron*, and *Piper*. Implications for the phylogenetic Relationships of magnoliids.**

In: BMC Evolutionary Biology 6, p. 77. DOI: 10.1186/1471-2148-6-77.

CHASE, MARK W.; REVEAL, JAMES L. (2009):

**A phylogenetic Classification of the land plants to accompany APG III.**

In: Botanical Journal of the Linnean Society (161), p. 122–127.

Drew, Bryan T.; Ruhfel, Bradley R.; Smith, Stephen A.; Moore, Michael J.; Briggs, Barbara G.; Gitzendanner, Matthew A. et al. (2014):
**Another look at the root of the Angiosperms reveals a familiar tale.**
In: Systematic Biology 63 (3), p. 368–382. DOI: 10.1093/sysbio/syt108.

Fiz-Palacios, Omar; Schneider, Harald; Heinrichs, Jochen; Savolainen, Vincent (2011):
**Diversification of land plants: Insights from a Family-level phylogenetic Analysis.**
In: BMC Evolutionary Biology 11, p. 341. DOI: 10.1186/1471-2148-11-341.

Goremykin, Vadim V.; Nikiforova, Svetlana V.; Biggs, Patrick J.; Zhong, Bojian; Delange, Peter; Martin, William et al. (2013):
**The evolutionary root of Flowering Plants.**
In: Systematic Biology 62 (1), p. 50–61. DOI: 10.1093/sysbio/sys070.

Group, Angiosperm Phylogeny (2016):
**An update of the Angiosperm Phylogeny Group An update of the Angiosperm Phylogeny Group Classification for the orders and families of Flowering Plants: APG IV.**
In: Botanical Journal of the Linnean Society 181, p. 1–20.

Hansen, Debra R.; Dastidar, Sayantani G.; Cai, Zhengqiu; Peñaflor, Cynthia; Kuehl, Jennifer V.; Boore, Jeffrey L.; Jansen, Robert K. (2007):
**Phylogenetic and evolutionary Implications of complete Chloroplast Genome Sequences of four early-diverging Angiosperms. *Buxus* (Buxaceae), *Chloranthus* (Chloranthaceae), *Dioscorea* (Dioscoreaceae), and *Illicium* (Schisandraceae).**
In: Molecular Phylogenetics and Evolution 45 (2), p. 547–563. DOI: 10.1016/j.ympev.2007.06.004.

Hilu, Khidir W.; Black, Chelsea; Diouf, Diaga; Burleigh, John Gordon (2008):
**Phylogenetic signal in matK vs. trnK. A case Study in early diverging Eudicots (Angiosperms).**
In: Molecular Phylogenetics and Evolution 48 (3), p. 1120–1130. DOI: 10.1016/j.ympev.2008.05.021.

Kim, Sangtae; Soltis, Douglas E.; Soltis, Pamela S.; Zanis, Michael J.; Suh, Youngbae (2004):
**Phylogenetic Relationships among early-diverging Eudicots based on four Genes. Were the Eudicots ancestrally woody?**
In: Molecular Phylogenetics and Evolution 31 (1), p. 16–30. DOI: 10.1016/j.ympev.2003.07.017.

Magallón, Susana (2010):
**Using fossils to break long branches in molecular dating. A Comparison of relaxed Clocks applied to the Origin of Angiosperms.**
In: Systematic Biology 59 (4), p. 384–399. DOI: 10.1093/sysbio/syq027.

Maia, Vitor H.; Gitzendanner, Matthew A.; Soltis, Pamela S.; Wong, Gane Ka-Shu; Soltis, Douglas E. (2014):
**Angiosperm Phylogeny based on 18s/26s rDNA Sequence Data: Constructing a large Data Set using Next-Generation Sequence Data.**
In: International Journal of Plant Sciences 175 (6), p. 613–650. DOI: 10.1086/676675.

Massoni, Julien; Forest, Félix; Sauquet, Hervé (2014):
**Increased Sampling of both Genes and Taxa improves Resolution of phylogenetic Relationships within Magnoliidae, a large and early-diverging Clade of Angiosperms.**
In: Molecular Phylogenetics and Evolution 70, p. 84–93. DOI: 10.1016/j.ympev.2013.09.010.

Moore, Michael J.; Hassan, Nasr; Gitzendanner, Matthew A.; Bruenn, Riva A.; Croley, Matthew; Vandeventer, Alexia et al. (2011):

**Phylogenetic Analysis of the Plastid inverted repeat for 244 Species. Insights into Deeper-Level Angiosperm Relationships from a long, slowly evolving Sequence Region.**
In: International Journal of Plant Sciences 172 (4), p. 541–558. DOI: 10.1086/658923.

OLMSTEAD, RICHARD G.; KIM, K. J.; JANSEN, ROBERT K.; WAGSTAFF, STEVEN J. (2000):
**The Phylogeny of the Asteridae sensu lato based on Chloroplast ndhF Gene Sequences.**
In: Molecular Phylogenetics and Evolution 16 (1), p. 96–112. DOI: 10.1006/mpev.1999.0769.

OXELMAN, BENGT; YOSHIKAWA, NORI; MCCONAUGHY, BETTY L.; LUO, JIE; DENTON, AMY L.; HALL, BENJAMIN D. (2004):
**Rpb2 Gene Phylogeny in Flowering Plants, with particular Emphasis on Asterids.**
In: Molecular Phylogenetics and Evolution 32 (2), p. 462–479. DOI: 10.1016/j.ympev.2004.01.014.

PIRANI, JOSÉ RUBENS; PRADO, JEFFERSON (2012):
**Embryopsida, a new Name for the Class of Land Plants.**
In: Taxon 61 (5), p. 1096–1098.

REYES, ELISABETH; SAUQUET, HERVÉ; NADOT, SOPHIE (2016):
**Perianth Symmetry Changed at least 199 times in Angiosperm Evolution.**
In: Taxon 65 (5), p. 945–964. DOI: 10.12705/655.1.

SHI, SU-HUA; HUANG, YE-LIN; ZENG, KAI; TAN, FENGXIAO; HE, HANGHANG; HUANG, JIANZI; FU, YUNXIN (2005):
**Molecular phylogenetic Analysis of Mangroves. Independent evolutionary Origins of Vivipary and Salt Secretion.**
In: Molecular Phylogenetics and Evolution 34 (1), p. 159–166. DOI: 10.1016/j.ympev.2004.09.002.

SOLTIS, DOUGLAS E.; SMITH, STEPHEN A.; CELLINESE, NICOLETTA; WURDACK, KENNETH J.J.; TANK, DAVID C.; BROCKINGTON, SAMUEL F. ET AL. (2011):
**Angiosperm Phylogeny. 17 genes, 640 taxa.**
In: American Journal of Botany 98 (4), p. 704–730. DOI: 10.3732/ajb.1000404.

SUN, MIAO; NAEEM, REHAN; SU, JUN-XIA; CAO, ZHI-YONG; BURLEIGH, JOHN GORDON; SOLTIS, PAMELA P. ET AL. (2016):
**Phylogeny of the Rosidae : A dense Taxon Sampling Analysis.**
In: Journal of Systematics and Evolution 54 (4), p. 363–391. DOI: 10.1111/jse.12211.

SUN, MIAO; SOLTIS, DOUGLAS E.; SOLTIS, PAMELA S.; ZHU, XINYU; BURLEIGH, JOHN GORDON; CHEN, ZHI-DUAN (2015):
**Deep phylogenetic Incongruence in the Angiosperm Clade Rosidae.**
In: Molecular Phylogenetics and Evolution 83, p. 156–166. DOI: 10.1016/j.ympev.2014.11.003.

SUN, YANXIA; MOORE, MICHAEL J.; ZHANG, SHOUJUN; SOLTIS, PAMELA S.; SOLTIS, DOUGLAS E.; ZHAO, TINGTING ET AL. (2016):
**Phylogenomic and structural Analyses of 18 complete Plastomes across nearly all Families of early-diverging Eudicots, including an Angiosperm-wide Analysis of Ir Gene Content Evolution.**
In: Molecular Phylogenetics and Evolution 96, p. 93–101. DOI: 10.1016/j.ympev.2015.12.006.

TANK, DAVID C.; DONOGHUE, MICHAEL J. (2010):
**Phylogeny and phylogenetic Nomenclature of the Campanulidae based on an expanded Sample of Genes and Taxa.**
In: Systematic Botany 35 (2), p. 425–441. DOI: 10.1600/036364410791638306.

WINKWORTH, RICHARD C.; LUNDBERG, JOHANNES; DONOGHUE, MICHAEL J. (2008):
**Toward a resolution of Campanulid Phylogeny, with special Reference to the Placement of Dipsacales.**

In: Taxon 57 (1), p. 53–65.

**Zhao, Lei; Li, Xia; Zhang, Ning; Zhang, Shu-Dong; Yi, Ting-Shuang; Ma, Hong et al. (2016):**
**Phylogenomic Analyses of large-scale nuclear Genes provide new Insights into the evolutionary Relationships within the Rosids.**
In: Molecular Phylogenetics and Evolution 105, p. 166–176. DOI: 10.1016/j.ympev.2016.06.007.

**Zhu, Xin-Yu; Chase, Mark W.; Qiu, Yin-Long; Kong, Hong-Zhi; Dilcher, David L.; Li, Jian-Hua; Chen, Zhi-Duan (2007):**
**Mitochondrial matR Sequences help to resolve deep phylogenetic Relationships in Rosids.**
In: BMC Evolutionary Biology 7, p. 217. DOI: 10.1186/1471-2148-7-217.

### Anisophylleaceae

**Chen, Xin; He, Hai; Zhang, Li-Bing (2015):**
**A monograph of the Anisophylleaceae (Cucurbitales) with Description of 18 new Species of *Anisophyllea*.**
In: Phytotaxa 229 (1), p. 1. DOI: 10.11646/phytotaxa.229.1.1.

**Zhang, Li-Bing; Simmons, Mark P.; Renner, Susanne p. (2007):**
**A Phylogeny of Anisophylleaceae based on six nuclear and Plastid Loci: ancient Disjunctions and recent Dispersal between South America, Africa, and Asia.**
In: Molecular Phylogenetics and Evolution 44 (3), p. 1057–1067. DOI: 10.1016/j.ympev.2007.03.002.

### Annonaceae

**Botermans, Marleen; Sosef, Marc p. M.; Chatrou, Lars W.; Couvreur, Thomas L.P. (2011):**
**Revision of the African Genus *Hexalobus* (Annonaceae).**
In: Systematic Botany 36 (1), p. 33–48. DOI: 10.1600/036364411X553108.

**Chaowasku, Tanawat; Johnson, David M.; Ham, Raymond W.J.M. van der; Chatrou, Lars W. (2012):**
**Characterization of *Hubera* (Annonaceae), a new Genus segregated from *Polyalthia* and allied to *Miliusa*.**
In: Phytotaxa 69, p. 33–56.

**Chaowasku, Tanawat; Keßler, Paul J.A.; van der Ham, Raymond W. J. M. (2012):**
**A taxonomic Revision and Pollen Morphology of the Genus *Dendrokingstonia* (Annonaceae).**
In: Botanical Journal of the Linnean Society 168 (1), p. 76–90. DOI: 10.1111/j.1095-8339.2011.01187.x.

**Chaowasku, Tanawat; Thomas, Daniel C.; van der Ham, Raymond W. J. M.; Smets, Erik F.; Mols, Johan B.; Chatrou, Lars W. (2014):**
**A Plastid DNA Phylogeny of Tribe Miliuseae. Insights into Relationships and Character Evolution in one of the most recalcitrant major Clades of Annonaceae.**
In: American Journal of Botany 101 (4), p. 691–709. DOI: 10.3732/ajb.1300403.

**Chatrou, Lars W.; Pirie, Michael D.; Erkens, Roy H.J.; Couvreur, Thomas L.P.; Neubig, Kurt M.; Abbott, Richard J. et al. (2012):**
**A new subfamilial and tribal Classification of the pantropical flowering Plant Family Annonaceae informed by molecular Phylogenetics.**
In: Botanical Journal of the Linnean Society 169 (1), p. 5–40. DOI: 10.1111/j.1095-8339.2012.01235.x.

**Couvreur, T. L. P.; Richardson, James E.; Sosef, Marc p. M.; Erkens, Roy H.J.; Chatrou, Lars W. (2008):**
**Evolution of syncarpy and other morphological Characters in African Annonaceae. A posterior mapping approach.**

In: Molecular Phylogenetics and Evolution 47 (1), p. 302–318. DOI: 10.1016/j.ympev.2008.01.018.

COUVREUR, THOMAS L.P.; VAN DER HAM, RAYMOND W. J. M.; MBELE, YOUSSOUFA M.; MBAGO, FRANK M.; JOHNSON, DAVID M. (2009):
**Molecular and morphological Characterization of a new monotypic Genus of Annonaceae, *Mwasumbia* from Tanzania.**
In: Systematic Botany 34 (2), p. 266–276. DOI: 10.1600/036364409788606398.

ERKENS, ROY H.J.; CHATROU, LARS W.; CHAOWASKU, TANAWAT; WESTRA, LUBBERT Y. TH.; MAAS, JAN W.A.N.; MAAS, PAUL J.M. (2014):
**A decade of uncertainty: Resolving the phylogenetic Position of *Diclinanona* (Annonaceae), including taxonomic Notes and a key to the Species.**
In: Taxon 63 (6), p. 1244–1252. DOI: 10.12705/636.34.

ERKENS, ROY H.J.; CHATROU, LARS W.; KOEK-NOORMAN, JIFKE; MAAS, JAN W.A.N.; MAAS, PAUL J.M. (2007):
**Classification of a large and widespread Genus of Neotropical Trees, *Guatteria* (Annonaceae) and its three satellite Genera *Guatteriella, Guatteriopsis* and *Heteropetalum*.**
In: Taxon 56 (3), p. 757–774.

ERKENS, ROY H.J.; CHATROU, LARS W.; MAAS, JAN W.A.N.; VAN DER NIET, TIMOTHEÜS; SAVOLAINEN, VINCENT (2007):
**A rapid Diversification of Rainforest Trees (*Guatteria*; Annonaceae) following Dispersal from Central into South America.**
In: Molecular Phylogenetics and Evolution 44 (1), p. 399–411. DOI: 10.1016/j.ympev.2007.02.017.

GOTTSBERGER, GERHARD (2016):
**The reproductive biology of the early-divergent Genus *Anaxagorea* (Annonaceae), and its Significance for the evolutionary development of the Family.**
In: Acta Botanica Brasilica 30 (2), p. 313–325. DOI: 10.1590/0102-33062015abb0311.

GUO, XING; HOEKSTRA, PAUL H.; TANG, CHIN CHEUNG; THOMAS, DANIEL C.; WIERINGA, JAN J.; CHATROU, LARS W.; SAUNDERS, RICHARD M.K. (2017):
**Cutting up the climbers: Evidence for extensive Polyphyly in *Friesodielsia* (Annonaceae) necessitates generic Realignment across the Tribe Uvarieae.**
In: Taxon 66 (1), p. 3–19. DOI: 10.12705/661.1.

GUO, XING; TANG, CHIN CHEUNG; THOMAS, DANIEL C.; COUVREUR, THOMAS L.P.; SAUNDERS, RICHARD M.K. (2017):
**A mega-phylogeny of the Annonaceae: taxonomic Placement of five enigmatic Genera and support for a new Tribe, Phoenicantheae.**
In: Scientific Reports 7 (1), p. 7323. DOI: 10.1038/s41598-017-07252-2.

GUO, XING; THOMAS, DANIEL C.; SAUNDERS, RICHARD M.K. (2018):
**Gene Tree discordance and coalescent Methods support ancient intergeneric Hybridisation between *Dasymaschalon* and *Friesodielsia* (Annonaceae).**
In: Molecular Phylogenetics and Evolution 127, p. 14–29. DOI: 10.1016/j.ympev.2018.04.009.

GUO, XING; WANG, JING; XUE, BINE; THOMAS, DANIEL C.; SU, YVONNE C.F.; TAN, YUN-HONG; SAUNDERS, RICHARD M.K. (2014):
**Reassessing the taxonomic Status of two enigmatic *Desmos* Species (Annonaceae). morphological and molecular phylogenetic Support for a new Genus, *Wangia*.**
In: Journal of Systematics and Evolution 52 (1), p. 1–15. DOI: 10.1111/jse.12064.

HOEKSTRA, PAUL H.; WIERINGA, JAN J.; SMETS, ERIK; BRANDÃO, RITA D.; LOPES, JENIFER DE CARVALHO; ERKENS, ROY H.J.; CHATROU, LARS W. (2017):
**Correlated evolutionary Rates across genomic Compartments in Annonaceae.**

In: Molecular Phylogenetics and Evolution 114, p. 63–72. DOI: 10.1016/j.ympev.2017.05.026.

**Li, Pui-Sze; Thomas, Daniel C.; Saunders, Richard M.K. (2015):**
**Phylogenetic Reconstruction, morphological Diversification and Generic Delimitation of *Disepalum* (Annonaceae).**
In: Public Library of Science One 10 (12), e0143481. DOI: 10.1371/journal.pone.0143481.

**Lopes, J. C.; Chatrou, Lars W.; Mello-Silva, Renato; Rudall, Paula J.; Sajo, M. G. (2018):**
**Phylogenomics and Evolution of Floral traits in the Neotropical Tribe Malmeeae (Annonaceae).**
In: Molecular Phylogenetics and Evolution 118, p. 379–391. DOI: 10.1016/j.ympev.2017.10.020.

**Maas, Paul J.M.; Westra, L.Y.T.; Guerrero, p. Arias; Lobão, Adriana Q.; Scharf, Uwe; Zamora, N. A.; Erkens, Roy H.J. (2015):**
**Confronting a morphological nightmare: Revision of the Neotropical Genus *Guatteria* (Annonaceae ).**
In: Blumea 60 (1), p. 1–219. DOI: 10.3767/000651915X690341.

**Maas, Paul J.M.; Westra, Lubbert Y. Th.; Rainer, Heimo; Lobão, Adriana Q.; Erkens, Roy H.J. (2011):**
**An updated index to Genera, Species, and infraspecific Taxa of Neotropical Annonaceae.**
In: Nordic Journal of Botany 29 (3), p. 257–356. DOI: 10.1111/j.1756-1051.2011.01092.x.

**Maas, Roy H. J. Erkens & Paul J. M. (2008):**
**The *Guatteria* Group disentangled: sinling *Guatteriopsis, Guatteriella* and *Heteropetalum* into *Guatteria*.**
In: Rodriguésia 59 (2), p. 401–406.

**Mercer, Elizabeth; Griffin, Brandi; Steele, Joshua; Goodrich, Katherine R.; Bush, Catherine M. (2016):**
**Phylogenetic Relationships of *Asimina* and *Deeringothamnus* (Annonaceae) based on Morphology, Floral scent Chemistry, and Inter-Simple Sequence Repeat Data.**
In: the Journal of the Torrey Botanical Society 143 (1), p. 58–68. DOI: 10.3159/TORREY-D-14-00045.1.

**Mols, Johan B.; Gravendeel, Barbara; Chatrou, Lars W.; Pirie, Michael D.; Bygrave, Paul C.; Chase, Mark W.; Keßler, Paul J.A. (2004):**
**Identifying Clades in Asian Annonaceae. Monophyletic Genera in the polyphyletic Miliuseae.**
In: American Journal of Botany 91 (4), p. 590–600. DOI: 10.3732/ajb.91.4.590.

**Mols, Johan B.; Keßler, Paul J.A.; Rogstad, Steven H.; Saunders, Richard M.K. (2008):**
**Reassignment of six *Polyalthia* Species to the new Genus *Maasia* (Annonaceae): molecular and morphological Congruence.**
In: Systematic Botany 33 (3), p. 490–494.

**Nakkuntod, Maliwan; Su, Yvonne C.F.; Seelanan, Tosak Saunders; Saunders, Richard M.K. (2009):**
**Molecular phylogenetic and morphological Evidence for the congeneric Status of *Goniothalamus* and *Richella* (Annonaceae).**
In: Taxon 58 (1), p. 127–132.

**Ortiz-Rodriguez, Andrés Ernesto; Ornelas, Juan Francisco; Ruiz-Sanchez, Eduardo (2018):**
**A jungle tale: molecular Phylogeny and Divergence time estimates of the *Desmopsis-Stenanona* Clade (Annonaceae) in Mesoamerica.**
In: Molecular Phylogenetics and Evolution 122, p. 80–94. DOI: 10.1016/j.ympev.2018.01.021.

**Pirie, Michael D.; Vargas, Maria Paula Balcázar; Botermans, Marleen; Bakker, Freek T.; Chatrou, Lars W. (2007):**

**Ancient Paralogy in the cpDNA trnL-F Region in Annonaceae. Implications for Plant molecular Systematics.**

In: American Journal of Botany 94 (6), p. 1003–1016. DOI: 10.3732/ajb.94.6.1003.

**Rainer, Heimo (2007):**

**Monographic studies in the Genus *Annona* L. (Annonaceae): Inclusion of the Genus *Rollinia* A.ST.-Hil.**

In: Annalen des Naturhistorischen Museums in Wien, Serie B 108B, p. 191–205.

**Saunders, Richard M.K.; Su, Yvonne C.F.; Xue, Bine (2011):**

**Phylogenetic affinities of *Polyalthia* Species (Annonaceae) with columellar-sulcate Pollen: Enlarging the Madagascan endemic Genus *Fenerivia*.**

In: Taxon 60 (5), p. 1407–1416.

**Saunders, Richard M.K.; Wang, Jing (2011):**

**Five new nomenclatural Combinations in *Dasymaschalon* and *Goniothalamus* (Annonaceae).**

In: Nordic Journal of Botany 29 (6), p. 674–676. DOI: 10.1111/j.1756-1051.2011.01293.x.

**Scharaschkin, Tanya; Doyle, James A. (2005):**

**Phylogeny and Historical Biogeography of *Anaxagorea* (Annonaceae) using Morphology and Non-Coding Chloroplast Sequence Data.**

In: Systematic Botany 30 (4), p. 712–735. DOI: 10.1600/036364405775097888.

**Su, Yvonne C.F.; Chaowasku, Tanawat; Saunders, Richard M.K. (2010):**

**An extended Phylogeny of *Pseuduvaria* (Annonaceae) with Descriptions of three new Species and a Reassessment of the generic Status of *Oreomitra*.**

In: Systematic Botany 35 (1), p. 30–39. DOI: 10.1600/036364410790862533.

**Su, Yvonne C.F.; Mols, Johan B.; Takeuchi, Wayne N.; Keßler, Paul J.A.; Saunders, Richard M.K. (2005):**

**Reassessing the generic Status of *Petalolophus* (Annonaceae): Evidence for the Evolution of a distinct sapromyophilous Lineage within *Pseuduvaria*.**

In: Systematic Botany 30 (3), p. 494–502.

**Su, Yvonne C.F.; Saunders, Richard M.K. (2009):**

**Evolutionary Divergence times in the Annonaceae. Evidence of a late Miocene Origin of *Pseuduvaria* in Sundaland with subsequent Diversification in New Guinea.**

In: BMC Evolutionary Biology 9, p. 153. DOI: 10.1186/1471-2148-9-153.

**Su, Yvonne C.F.; Smith, Gavin J. D.; Saunders, Richard M.K. (2008):**

**Phylogeny of the basal Angiosperm Genus *Pseuduvaria* (Annonaceae) inferred from five Chloroplast DNA Regions, with interpretation of morphological Character Evolution.**

In: Molecular Phylogenetics and Evolution 48 (1), p. 188–206. DOI: 10.1016/j.ympev.2008.03.028.

**Surveswaran, Siddharthan; Wang, Rui-Jiang; Su, Yvonne C.F.; Saunders, Richard M.K. (2010):**

**Generic Delimitation and historical Biogeography in the early-divergent 'ambavioid' Lineage of Annonaceae: *Cananga, Cyathocalyx* and *Drepananthus*.**

In: Taxon 59 (6), p. 1721–1734.

**Tang, Chin Cheung; Thomas, Daniel C.; Saunders, Richard M.K. (2015):**

**Molecular Phylogenetics of the species-rich Angiosperm Genus *Goniothalamus* (Annonaceae) inferred from nine Chloroplast DNA Regions. Synapomorphies and putative correlated evolutionary Changes in Fruit and Seed Morphology.**
In: Molecular Phylogenetics and Evolution 92, p. 124–139. DOI: 10.1016/j.ympev.2015.06.016.

THOMAS, DANIEL C.; CHATROU, LARS W.; STULL, GREGORY W.; JOHNSON, DAVID M.; HARRIS, DAVID J.; THONGPAIROJ, U-SA; SAUNDERS, RICHARD M.K. (2015):
**The historical Origins of palaeotropical intercontinental Disjunctions in the pantropical flowering plant Family Annonaceae.**
In: Perspectives in Plant Ecology, Evolution and Systematics 17 (1), p. 1–16. DOI: 10.1016/j.ppees.2014.11.001.

THOMAS, DANIEL C.; SURVESWARAN, SIDDHARTHAN; XUE, BINE; SANKOWSKY, GARRY; MOLS, JOHAN B. (2012):
**Molecular Phylogenetics and historical Biogeography of the *Meiogyne-Fitzalania* Clade (Annonaceae): Generic Paraphyly and late Miocene-Pliocene Diversification in Australasia and the Pacific.**
In: Taxon 61 (3), p. 559–575.

TURNER, IAN M. (2012):
**A new Combination in *Monocarpia* (Annonaceae).**
In: Edinburgh Journal of Botany 69 (01), p. 95–98. DOI: 10.1017/S0960428611000424.

TURNER, IAN M. (2015):
**A Conspectus of Indo-Burmese Annonaceae.**
In: Nordic Journal of Botany 33 (3), p. 257–299. DOI: 10.1111/njb.00689.

TURNER, IAN M.; UTTERIDGE, TIMOTHY M.A. (2015):
**A new Species and a new Combination in *Meiogyne* (Annonaceae) of New Guinea. Contributions to the Flora of Mt Jaya, XXI.**
In: Kew Bulletin 70 (2). DOI: 10.1007/S12225-015-9577-6.

VAN ZUILEN, CAROLINA M.; KOEK-NOORMAN, JIFKE; MAAS, PAUL J.M. (1995):
**A phylogenetic Analysis of *Duguetia* (Annonaceae) based on morphological Data.**
In: Plant Systematics and Evolution 194, p. 173–188.

WANG, JING; THOMAS, DANIEL C.; SU, YVONNE C.F.; MEINKE, SVENJA; CHATROU, LARS W. (2012):
**A Plastid DNA Phylogeny of *Dasymaschalon* (Annonaceae) and allied Genera: Evidence for generic non-Monophyly and the parallel evolutionary loss of inner petals.**
In: Taxon 61 (3), p. 545–558.

XUE, BINE; SU, YVONNE C.F.; THOMAS, DANIEL C.; SAUNDERS, RICHARD M.K. (2012):
**Pruning the polyphyletic Genus *Polyalthia* (Annonaceae) and resurrecting the Genus *Monoon*.**
In: Taxon 61 (5), p. 1021–1039.

XUE, BINE; TAN, YUN-HONG; YE, XING-ER (2016):
**The identity of *Polyalthia florulenta* (Annonaceae): a second Species of *Wangia* in China.**
In: Phytotaxa 283 (2), p. 163–171. http://www.mapress.com/j/pt/article/download/phytotaxa.283.2.5/9246.

XUE, BINE; THOMAS, DANIEL C.; CHAOWASKU, TANAWAT; JOHNSON, DAVID M.; SAUNDERS, RICHARD M.K. (2014):
**Molecular phylogenetic Support for the taxonomic Merger of *Fitzalania* and *Meiogyne* (Annonaceae). new nomenclatural Combinations under the conserved Name *Meiogyne*.**

In: Systematic Botany 39 (2), p. 396–404. DOI: 10.1600/036364414X680825.

**Zhou, Linlin; Su, Yvonne C.F.; Chalermglin, Piya; Saunders, Richard M.K. (2010):**
**Molecular Phylogenetics of *Uvaria* (Annonaceae). Relationships with *Balonga, Dasoclema* and Australian Species of *Melodorum*.**
In: Botanical Journal of the Linnean Society 163 (1), p. 33–43. DOI: 10.1111/j.1095-8339.2010.01045.x.

**Zhou, Linlin; Su, Yvonne C.F.; Thomas, Daniel C.; Saunders, Richard M.K. (2012):**
**'Out-of-Africa' Dispersal of tropical Floras during the Miocene climatic Pptimum. Evidence from *Uvaria* (Annonaceae).**
In: Journal of Biogeography 39 (2), p. 322–335. DOI: 10.1111/j.1365-2699.2011.02598.x.

### Anthericaceae

**Bjorå, Charlotte Sletten; Hemp, Andreas; Hoell, Gry; Nordal, Inger (2008):**
**A taxonomic and ecological Analysis of two Forest *Chlorophytum* Taxa (Anthericaceae) on Mount Kilimanjaro, Tanzania.**
In: Plant Systematics and Evolution 274 (3-4), p. 243–253. DOI: 10.1007/s00606-008-0032-0.

**Bjorå, Charlotte Sletten; Hoell, Gry; Kativu, Shakkie; Nordal, Inger (2008):**
**New Taxa of *Chlorophytum* (Anthericaceae) from southern tropical Africa with Notes on their sister Group Relationships.**
In: Botanical Journal of the Linnean Society 157 (2), p. 223–238. DOI: 10.1111/j.1095-8339.2008.00811.x.

**Cruden, Robert William (2009):**
**A Synopsis of South American *Echeandia* (Anthericaceae).**
In: Annals of the Missouri Botanical Garden 96 (2), p. 251–267. DOI: 10.3417/2002129.

**Katoch, Meenu; Kumar, Rajinder; Pal, Swadesh; Ahuja, Ashok (2010):**
**Identification of *Chlorophytum* Species (*C. borivilianum, C. arundinaceum, C. laxum, C. capense* and *C. comosum*) using molecular Markers.**
In: Industrial Crops and Products 32 (3), p. 389–393. DOI: 10.1016/j.indcrop.2010.06.001.

**Meerts, Pierre; Bjorå, Charlotte Sletten (2012):**
**Synopsis of the Genus *Chlorophytum* (Asparagaceae) in Central Africa (Democratic Republic of the Congo, Rwanda, Burundi).**
In: Plant Ecology and Evolution 145 (3), p. 373–409. DOI: 10.5091/plecevo.2012.668.

### Apiaceae

**Ajani, Yousef; Ajani, Ahmad; Cordes, Jenny M.; Watson, Mark F.; Downie, Stephen R. (2008):**
**Phylogenetic Analysis of nrDNA ITS Sequences reveals Relationships within five Groups of Iranian Apiaceae Subfamily Apioideae.**
In: Taxon, p. 383–401.

**Ajani, Yousef; Ajani, Ahmad; Cordes, Jenny M.; Watson, Mark F.; Downie, Stephen R. (2008):**
**Phylogenetic Analysis of nrDNA ITS Sequences reveals Relationships within five Groups of Iranian Apiaceae Subfamily Apioideae.**
In: Taxon 57 (2), p. 383–401.

**Aliabadian, Mansour; Kaboli, Mohammad; Prodon, Roger; Nijman, Vincent; Vences, Miguel (2007):**
**Phylogeny of Palaearctic wheatears (genus *Oenanthe*)--congruence between morphometric and molecular Data.**
In: Molecular Phylogenetics and Evolution 42 (3), p. 665–675. DOI: 10.1016/j.ympev.2006.08.018.

**Andersson, Lennart; Kocsis, Marianna; Eriksson, Roger (2006):**

**Relationships of the Genus *Azorella* (Apiaceae) and other hydrocotyloids inferred from Sequence Variation in three Plastid Markers.**
In: Taxon 55 (2), p. 270–280.

ARBIZU, CARLOS; REITSMA, KATHLEEN R.; SIMON, PHILIPP W.; SPOONER, DAVID M. (2014):
**Morphometrics of *Daucus* (Apiaceae): a Counterpart to a phylogenomic Study.**
In: American Journal of Botany 101 (11), p. 2005–2016. DOI: 10.3732/ajb.1400252.

ARBIZU, CARLOS; RUESS, HOLLY; SENALIK, DOUGLAS; SIMON, PHILIPP W.; SPOONER, DAVID M. (2014):
**Phylogenomics of the Carrot Genus (*Daucus*, Apiaceae).**
In: American Journal of Botany 101 (10), p. 1666–1685. DOI: 10.3732/ajb.1400106.

BANASIAK, ŁUKASZ; WOJEWÓDZKA, ANETA; BACZYŃSKI, JAKUB; REDURON, JEAN-PIERRE; PIWCZYŃSKI, MARCIN; KURZYNA-MŁYNIK, RENATA ET AL. (2016):
**Phylogeny of Apiaceae Subtribe Daucinae and the taxonomic Delineation of its Genera.**
In: Taxon 65 (3), p. 563–585. DOI: 10.12705/653.8.

BONE, TIFFANY S.; DOWNIE, STEPHEN R.; AFFOLTER, JAMES M.; SPALIK, KRYSZTOF (2011):
**A phylogenetic and Biogeographic Study of the Genus *Lilaeopsis* (Apiaceae Tribe Oenantheae).**
In: Systematic Botany 36 (3), p. 789–805. DOI: 10.1600/036364411X583745.

BRULLO, CRISTIAN; BRULLO, SALVATORE; DOWNIE, STEPHEN R.; DANDERSON, CLARK A.; DEL GALDO, GIANPIETRO GIUSSO (2013):
***Siculosciadium*, a new monotypic Genus of Apiaceae from Sicily.**
In: Annals of the Missouri Botanical Garden 99 (1), p. 1–18. DOI: 10.3417/2011009.

CALVIÑO, CAROLINA ISABEL; DOWNIE, STEPHEN R. (2007):
**Circumscription and Phylogeny of Apiaceae Subfamily Saniculoideae based on Chloroplast DNA Sequences.**
In: Molecular Phylogenetics and Evolution 44 (1), p. 175–191. DOI: 10.1016/j.ympev.2007.01.002.

CALVIÑO, CAROLINA ISABEL; MARTÍNEZ, SUSANA G.; DOWNIE, STEPHEN R. (2008):
**Morphology and Biogeography of Apiaceae Subfamily Saniculoideae as inferred by phylogenetic Analysis of molecular Data.**
In: American Journal of Botany 95 (2), p. 196–214. DOI: 10.3732/ajb.95.2.196.

CALVIÑO, CAROLINA ISABEL; MARTÍNEZ, SUSANA G.; DOWNIE, STEPHEN R. (2008):
**The evolutionary History of *Eryngium* (Apiaceae, Saniculoideae): rapid Radiations, long distance Dispersals, and Hybridizations.**
In: Molecular Phylogenetics and Evolution 46 (3), p. 1129–1150. DOI: 10.1016/j.ympev.2007.10.021.

CALVIÑO, CAROLINA ISABEL; TILNEY, PATRICIA M.; VAN WYK, BEN-ERIK; DOWNIE, STEPHEN R. (2006):
**A molecular phylogenetic Study of southern African Apiaceae.**
In: American Journal of Botany 93 (12), p. 1828–1847. DOI: 10.3732/ajb.93.12.1828.

CASTRO, OLGA; CENNAMO, PAOLA; LUCA, PAOLO (2009):
**Analysis of the Genus *Petagnaea* Caruel (Apiaceae), using new molecular and Literature Data.**
In: Plant Systematics and Evolution 278 (3-4), p. 239–249. DOI: 10.1007/s00606-008-0133-9.

CHUNG, KUO-FANG (2007):
**Inclusion of the South Pacific Alpine Genus *Oreomyrrhis* (Apiaceae) in *Chaerophyllum* based on nuclear and Chloroplast DNA Sequences.**

In: Systematic Botany 32 (3), p. 671–681. DOI: 10.1600/036364407782250517.

**Chung, Kuo-Fang; Peng, Ching-I.; Downie, Stephen R.; Spalik, Krysztof; Schaal, Barbara A. (2005):**
**Molecular Systematics of the trans-Pacific alpine Genus *Oreomyrrhis* (Apiaceae): phylogenetic affinities and biogeographic Implications.**
In: American Journal of Botany 92 (12), p. 2054–2071. DOI: 10.3732/ajb.92.12.2054.

**Çinbilgel, İlker; Eren, Özkan; Duman, Hayri; Gökceoğlu, Mustafa (2015):**
***Pimpinella ibradiensis* (Apiaceae), an unusual new Species from Turkey.**
In: Phytotaxa 217 (2), p. 164. DOI: 10.11646/phytotaxa.217.2.6.

**Danderson, Clark A.; Downie, Stephen R.; Hermann, Michael (2018):**
**Rampant Polyphyly in the *Arracacia*-clade (Apiaceae) and an assessment of the phylogenetic Utility of 20 noncoding Plastid loci.**
In: Molecular Phylogenetics and Evolution 118, p. 286–305. DOI: 10.1016/j.ympev.2017.10.006.

**Danderson, Clark A.; Hermann, Michael; Downie, Stephen R. (2008):**
**Resolving phylogenetic Relationships within the *Arracacia*-clade (Apiaceae Subfamily Apioideae) using cpDNA Sequence Data.**
University of Illinois Urbana-Champaign.

**Degtjareva, Galina V.; Kljuykov, Eugene V.; Samigullin, Tahir H.; Valiejo-Roman, Carmen M.; Pimenov, Michael G. (2013):**
**ITS Phylogeny of Middle Asian geophilic Umbelliferae-Apioideae Genera with comments on their Morphology and Utility of psbA-trnH Sequences.**
In: Plant Systematics and Evolution 299 (5), p. 985–1010. DOI: 10.1007/s00606-013-0779-9.

**Degtjareva, Galina V.; Kljuykov, Eugene V.; Samigullin, Tahir H.; Valiejo-Roman, Carmen M.; Pimenov, Michael G. (2009):**
**Molecular appraisal of *Bunium* and some related arid and subarid geophilic Apiaceae-Apioideae Taxa of the Ancient Mediterranean.**
In: Botanical Journal of the Linnean Society 160 (2), p. 149–170. DOI: 10.1111/j.1095-8339.2009.00970.x.

**Doğan, Bekir; Duran, Ahmet; Bagci, Yavuz; Dinc, Muhittin; Martin, Esra; Çetin, Özlem; Öztürk, Meryem (2010):**
**Phylogenetic Relationships among the Taxa of the Genus *Johrensia* DC. (Apiaceae) from Turkey based on molecular Method.**
In: Bangladesh Journal of Plant Taxononmy 17 (2), p. 113–120.

**Doğru-Koca, Asli (2016):**
**Phylogeny of the Genus *Tordylium* (Tordylineae, Apioideae, Apiaceae) inferred from morphological Data.**
In: Nordic Journal of Botany 34 (1), p. 111–119. DOI: 10.1111/njb.00991.

**Downie, Stephen R.; Hartman, Ronald L.; Sun, Feng-Jie; Katz-Downie, Deborah p. (2002):**
**Polyphyly of the spring-parsleys (*Cymopterus*): molecular and morphological Evidence suggests complex Relationships among the perennial endemic Genera of western North American Apiaceae.**
In: Canadian Journal of Botany 80 (12), p. 1295–1324. DOI: 10.1139/B02-119.

**Downie, Stephen R.; Katz-Downie, Deborah S.; Spalik, Krysztof (2000):**
**A Phylogeny of Apiaceae Tribe Scandiceae: Evidence from nuclear ribosomal DNA Internal Transcribed Spacer Sequences.**
In: American Journal of Botany 87 (1), p. 76–95. DOI: 10.2307/2656687.

**Downie, Stephen R.; Katz-Downie, Deborah S.; Sun, Feng-Jie; Lee, Chang-Shook (2008):**
**Phylogeny and Biogeography of Apiaceae Tribe Oenantheae inferred from nuclear rDNA ITS and cpDNA psbI – 5′trnK(UUU) Sequences, with Emphasis on the North American Endemics Clade.**
In: Botany 86 (9), p. 1039–1064. DOI: 10.1139/B08-055.

**Downie, Stephen R.; Spalik, Krysztof; Katz-Downie, Deborah S.; Reduron, Jean-Pierre (2010):**
**Major Clades within Apiaceae Subfamily Apioideae as inferred by phylogenetic Analysis of nrDNA ITS Sequences.**
In: Plant Diversity and Evolution 128 (1), p. 111–136. DOI: 10.1127/1869-6155/2010/0128-0005.

**Downie, Stephen R.; Watson, Mark F.; Spalik, Krysztof; Katz-Downie, Deborah p. (2000):**
**Molecular Systematics of Old World Apioideae (Apiaceae): Relationships among some Members of Tribe Peucedaneae sensu lato, the Placement of several Island-endemic Species, and Resolution within the apioid Superclade.**
In: Canadian Journal of Botany 78 (4), p. 506–528. DOI: 10.1139/b00-029.

**Feist, Mary Ann E.; Downie, Stephen R. (2008):**
**A phylogenetic Study of *Oxypolis* and *Ptilimnium* (Apiaceae) based on nuclear rDNA ITS Sequences.**
In: Systematic Botany 33 (2), p. 447–458. DOI: 10.1600/036364408784571509.

**Feist, Mary Ann E.; Downie, Stephen R.; Magee, Anthony Richard; Liu, Mei (2012):**
**Revised generic Delimitations for *Oxypolis* and *Ptilimnium* (Apiaceae) based on Leaf Morphology, comparative Fruit Anatomy, and phylogenetic Analysis of nuclear rDNA ITS and cpDNA trnQ-trnK intergenic Spacer Sequence Data.**
In: Taxon 61 (2), p. 402–418.

**Feng, Tu; Downie, Stephen R.; Yu, Yang; Zhang, Xuemei; Chen, Wei-Wei; He, Xing-Jin; Liu, Shuang (2009):**
**Molecular Systematics of *Angelica* and allied Genera (Apiaceae) from the Hengduan Mountains of China based on nrDNA ITS Sequences: phylogenetic affinities and biogeographic Implications.**
In: Journal of Plant Research 122 (4), p. 403–414. DOI: 10.1007/s10265-009-0238-4.

**Fernández, Martina; Ezcurra, Cecilia; Calviño, Carolina Isabel (2017):**
**Chloroplast and ITS Phylogenies to understand the evolutionary History of southern South American *Azorella, Laretia* and *Mulinum* (Azorelloideae, Apiaceae).**
In: Molecular Phylogenetics and Evolution 108, p. 1–21. DOI: 10.1016/j.ympev.2017.01.016.

**Hand, Ralf (2011):**
**The Euro+Med Treatment of Apiaceae.**
In: Willdenowia 41 (2), p. 245–250. DOI: 10.3372/wi.41.41205.

**Hartman, Ronald L.; Nesom, Guy L. (2012):**
**Taxonomy of the Genus *Vesper* (Apiaceae).**
In: Phytoneuron 94, p. 1–9.

**Henwood, Murray J.; Hart, J. M. (2001):**
**Towards an Understanding of the phylogenetic Relationships of Australian Hydrocotyloideae (Apiaceae).**
In: Edinburgh Journal of Botany 58 (02). DOI: 10.1017/S0960428601000634.

**Jimenez-Mejias, Pedro; Vargas, Pablo (2015):**

**Taxonomy of the Tribe Apieae (Apiaceae) revisited as revealed by molecular Phylogenies and morphological Characters.**
In: Phytotaxa 212 (1), p. 57. DOI: 10.11646/phytotaxa.212.1.2.

KADEREIT, JOACHIM W.; REPPLINGER, MIRIAM; SCHMALZ, NATALIE; UHINK, CHRISTIAN H.; WÖRZ, ARNO (2008):
**The Phylogeny and Biogeography of Apiaceae subf. Saniculoideae Tribe Saniculeae: from South to North and South again.**
In: Taxon 57 (2), p. 365–382.

KURZYNA-MŁYNIK, RENATA; OSKOLSKI, ALEXEI A.; DOWNIE, STEPHEN R.; KOPACZ, RAFAŁ; WOJEWÓDZKA, ANETA; SPALIK, KRYSZTOF (2008):
**Phylogenetic Position of the Genus *Ferula* (Apiaceae) and its Placement in Tribe Scandiceae as inferred from nrDNA ITS Sequence Variation.**
In: Plant Systematics and Evolution 274 (1-2), p. 47–66. DOI: 10.1007/s00606-008-0022-2.

LEE, CHANG-SHOOK; DOWNIE, STEPHEN R. (2006):
**Phylogenetic Relationships within *Cicuta* (Apiaceae Tribe Oenantheae) inferred from nuclear rDNA ITS and cpDNA Sequence Data.**
In: Canadian Journal of Botany 84 (3), p. 453–468. DOI: 10.1139/B06-016.

LIAO, CHEN-YANG; DOWNIE, STEPHEN R.; LI, QIN-QIN; YU, YANG; HE, XING-JIN; ZHOU, BO (2013):
**New Insights into the Phylogeny of *Angelica* and its Allies (Apiaceae) with Emphasis on East Asian Species, inferred from nrDNA, cpDNA, and morphological Evidence.**
In: Systematic Botany 38 (1), p. 266–281. DOI: 10.1600/036364413X662060.

LIAO, CHEN-YANG; DOWNIE, STEPHEN R.; YU, YANG; HE, XING-JIN (2012):
**Historical Biogeography of the *Angelica* Group (Apiaceae Tribe Selineae) inferred from Analyses of nrDNA and cpDNA Sequences.**
In: Journal of Systematics and Evolution 50 (3), p. 206–217. DOI: 10.1111/j.1759-6831.2012.00182.x.

LIU, MEI; PLUNKETT, GREGORY M.; LOWRY, PORTER PRESCOTT; VAN WYK, BEN-ERIK; TILNEY, PATRICIA M.; NICOLAS, ANTOINE N. (2016):
**The phylogenetic Significance of Fruit and Trichome Structures in Apiaceae Subfamily Mackinlayoideae.**
In: Systematic Botany 41 (3), p. 685–699. DOI: 10.1600/036364416X692541.

LOGACHEVA, MARIA D.; VALIEJO-ROMAN, CARMEN M.; DEGTJAREVA, GALINA V.; STRATTON, JENNY M.; DOWNIE, STEPHEN R.; SAMIGULLIN, TAHIR H.; PIMENOV, MICHAEL G. (2010):
**A Comparison of nrDNA ITS and ETS loci for phylogenetic Inference in the Umbelliferae: an example from Tribe Tordylieae.**
In: Molecular Phylogenetics and Evolution 57 (1), p. 471–476. DOI: 10.1016/j.ympev.2010.06.001.

LOGACHEVA, MARIA D.; VALIEJO-ROMAN, CARMEN M.; PIMENOV, MICHAEL G. (2008):
**ITS Phylogeny of West Asian *Heracleum* Species and related Taxa of Umbelliferae–Tordylieae W.D.J.Koch, with Notes on Evolution of their psbA-trnH Sequences.**
In: Plant Systematics and Evolution 270 (3-4), p. 139–157. DOI: 10.1007/s00606-007-0619-x.

LYSKOV, DMITRY; DEGTJAREVA, GALINA V.; SAMIGULLIN, TAHIR H.; PIMENOV, MICHAEL G. (2015):
**Systematic Placement of the Turkish endemic Genus *Ekimia* (Apiaceae) based on morphological and molecular Data.**
In: Turkish Journal of Botany 39, p. 673–680. DOI: 10.3906/bot-1405-111.

MAGEE, ANTHONY RICHARD; CALVIÑO, CAROLINA ISABEL; LIU, MEI; DOWNIE, STEPHEN R.; TILNEY, PATRICIA M. (2010):

**New tribal Delimitations for the early diverging Lineages of Apiaceae Subfamily Apioideae.**
In: Taxon 59 (2), p. 567–580.

MAGEE, ANTHONY RICHARD; CURTIS, ODETTE E.; VAN WYK, BEN-ERIK (2016):
**A refined Circumscription of *Notobubon striatum* and the Resurrection of *Dregea collina* Ecklon & Zeyher (Apiaceae, Apioideae).**
In: Phytotaxa 266 (1), p. 27. DOI: 10.11646/phytotaxa.266.1.4.

MAGEE, ANTHONY RICHARD; VAN WYK, BEN-ERIK; TILNEY, PATRICIA M. (2009):
**A taxonomic Revision of the woody South African Genus *Notobubon* (Apiaceae: Apioideae).**
In: Systematic Botany 34 (1), p. 220–242. DOI: 10.1600/036364409787602294.

MAGEE, ANTHONY RICHARD; VAN WYK, BEN-ERIK; TILNEY, PATRICIA M.; DOWNIE, STEPHEN R. (2009):
**Generic Delimitation s and Relationships of the Cape Genera *Capnophyllum, Dasispermum,* and *Sonderina*, the North African Genera *Krubera* and *Stoibrax*, and a new monotypic Genus of the Subfamily Apioideae (Apiaceae).**
In: Systematic Botany 34 (3), p. 580–594. DOI: 10.1600/036364409789271218.

MITCHELL, ANTHONY D.; WEBB, C. J.; WAGSTAFF, STEVEN J. (1998):
**Phylogenetic Relationships of Species of *Gingidia* and related Genera (Apiaceae, Subfamily Apioideae).**
In: New Zealand Journal of Botany 36 (3), p. 417–424. DOI: 10.1080/0028825X.1998.9512580.

NESOM, GUY L. (2012):
***Villarrealia* (Apiaceae), a new Genus from Northern Mexico.**
In: Phytoneuron 85, p. 1–6.

NEVES, SUSANA S.; WATSON, MARK F. (2004):
**Phylogenetic Relationships in *Bupleurum* (Apiaceae) based on nuclear ribosomal DNA ITS Sequence Data.**
In: Annals of Botany 93 (4), p. 379–398. DOI: 10.1093/aob/mch052.

NICOLAS, ANTOINE N.; PLUNKETT, GREGORY M. (2012):
**Untangling generic limits in *Azorella, Laretia,* and *Mulinum* (Apiaceae: Azorelloideae): Insights from Phylogenetics and Biogeography.**
In: Taxon 61 (4), p. 826–840.

PANAHI, MEHRNOUSH; BANASIAK, ŁUKASZ; PIWCZYŃSKI, MARCIN; PUCHAŁKA, RADOSŁAW; OSKOLSKI, ALEXEI A.; SPALIK, KRYSZTOF (2015):
**Phylogenetic Relationships among *Dorema, Ferula* and *Leutea* (Apiaceae: Scandiceae: Ferulinae) inferred from nrDNA ITS and cpDNA noncoding Sequences.**
In: Taxon 64 (4), p. 770–783. DOI: 10.12705/644.8.

PETERSEN, GITTE; SEBERG, OLE; LARSEN, SIDSEL (2002):
**The phylogenetic and taxonomic Position of *Lilaeopsis* (Apiaceae), with Notes on the applicability of ITS Sequence Data for phylogenetic reconstruction.**
In: Australian Systematic Botany 15 (2), p. 181. DOI: 10.1071/SB01007.

PIMENOV, MICHAEL G.; DEGTJAREVA, GALINA V.; OSTROUMOVA, TATIANA A.; SAMIGULLIN, TAHIR H.; AVERYANOV, LEONID V. (2016):
***Xyloselinum laoticum* (Umbelliferae), a new Species from Laos, and taxonomic Placement of the Genus in the light of nrDNA ITS Sequence Analysis.**

In: Phytotaxa 244 (3), p. 248. DOI: 10.11646/phytotaxa.244.3.2.

PIMENOV, MICHAEL G.; KLJUYKOV, EUGENE V. (2005):
**New West Himalayan Genus of the Umbelliferae, with Notes on Tibetan species, described in *Pachypleurum*.**
In: Feddes Repertorium 116 (1-2), p. 80–91. DOI: 10.1002/fedr.200411060.

PIMENOV, MICHAEL G.; VALIEJO-ROMAN, CARMEN M.; TERENTIEVA, ELENA I.; SAMIGULLIN, TAHIR H.; MOZAFFARIAN, VALIOLLAH (2007):
**Enigmatic Genus *Haussknechtia* (Umbelliferae): systematic Relationships based on molecular and carpological Data.**
In: Nordic Journal of Botany 24 (5), p. 555–565.

PIWCZYŃSKI, MARCIN; PUCHAŁKA, RADOSŁAW; SPALIK, KRYSZTOF (2015):
**The infrageneric Taxonomy of *Chaerophyllum* (Apiaceae) revisited: new Evidence from nuclear ribosomal DNA ITS Sequences and Fruit Anatomy.**
In: Botanical Journal of the Linnean Society 178 (2), p. 298–313. DOI: 10.1111/boj.12282.

PLUNKETT, GREGORY M.; NICOLAS, ANTOINE N. (2017):
**Assessing *Azorella* (Apiaceae) and its allies: Phylogenetics and a new Classification.**
In: Brittonia 69 (1), p. 31–61. DOI: 10.1007/s12228-016-9446-0.

RADFORD, ELIZABETH A.; WATSON, MARK F.; PRESTON, JILLIAN (2001):
**Phylogenetic Relationships of Species of *Aciphylla* (Apiaceae, Subfamily Apioideae) and related Genera using molecular, morphological, and combined Data Sets.**
In: New Zealand Journal of Botany 39 (2), p. 183–208. DOI: 10.1080/0028825X.2001.9512730.

SMITH, JAMES F.; MANSFIELD, DONALD H.; STEVENS, MCKAYLA; SOSA, EDGAR; FEIST, MARY ANN E.; DOWNIE, STEPHEN R. ET AL. (2018):
**Try Tri again? Resolving Species boundaries in the *Lomatium triternatum* (Apiaceae) Complex.**
In: Journal of Systematics and Evolution 56 (3), p. 218–230. DOI: 10.1111/jse.12418.

SPALIK, KRYSZTOF; DOWNIE, STEPHEN R. (2006):
**The evolutionary History of *Sium* sensu lato (Apiaceae): Dispersal, Vicariance, and Domestication as inferred from ITS rDNA Phylogeny.**
In: American Journal of Botany 93 (5), p. 747–761. DOI: 10.3732/ajb.93.5.747.

SPALIK, KRYSZTOF; DOWNIE, STEPHEN R.; WATSON, MARK F. (2009):
**Generic Delimitations within the *Sium* alliance (Apiaceae Tribe Oenantheae) inferred from cpDNA rps16-5′trnK (Uuu) and nrDNA ITS Sequences.**
In: Taxon 58 (3), p. 735–748.

SPALIK, KRYSZTOF; REDURON, JEAN-PIERRE; DOWNIE, STEPHEN R. (2004):
**The phylogenetic Position of *Peucedanum* sensu lato and allied Genera and their Placement in Tribe Selineae (Apiaceae, Subfamily Apioideae).**
In: Plant Systematics and Evolution 243 (3-4), p. 189–210. DOI: 10.1007/s00606-003-0066-2.

SPALIK, KRYSZTOF; WOJEWÓDZKA, ANETA; DOWNIE, STEPHEN R. (2001):
**The Evolution of Fruit in Scandiceae Subtribe Scandicinae (Apiaceae).**
In: Canadian Journal of Botany 79 (11), p. 1358–1374. DOI: 10.1139/cjb-79-11-1358.

SUN, FENG-JIE; DOWNIE, STEPHEN R.; HARTMAN, RONALD L. (2004):
**An ITS-based phylogenetic Analysis of the perennial, endemic Apiaceae Subfamily Apioideae of Western North America.**

In: Systematic Botany 29 (2), p. 419–431. DOI: 10.1600/036364404774195601.

SUN, NA; HE, XING-JIN; ZHOU, SONG-DONG (2008):
**Morphological cladistic Analysis of *Ligusticum* (Umbelliferae) in China.**
In: Nordic Journal of Botany 26 (1-2), p. 118–128. DOI: 10.1111/j.0107-055X.2008.00144.x.

TERENTIEVA, ELENA I.; VALIEJO-ROMAN, CARMEN M.; SAMIGULLIN, TAHIR H.; PIMENOV, MICHAEL G.; TILNEY, PATRICIA M. (2015):
**Molecular phylogenetic and morphological Analyses of the traditional Tribe Coriandreae (Umbelliferae-Apioideae).**
In: Phytotaxa 195 (4), p. 251. DOI: 10.11646/phytotaxa.195.4.1.

VALIEJO-ROMAN, CARMEN M.; SHNEYER, V. S.; SAMIGULLIN, TAHIR H.; TERENTIEVA, ELENA I.; PIMENOV, MICHAEL G. (2006):
**An attempt to clarify taxonomic Relationships in ``Verwandtschaftskreis der Gattung *Ligusticum*'' (Umbelliferae-Apioideae) by molecular Analysis.**
In: Plant Systematics and Evolution 257 (1-2), p. 25–43. DOI: 10.1007/s00606-005-0383-8.

VALIEJO-ROMAN, CARMEN M.; TERENTIEVA, ELENA I.; SAMIGULLIN, TAHIR H.; PIMENOV, MICHAEL G.; GHAHREMANINEJAD, FARROKH; MOZAFFARIAN, V. (2006):
**Molecular Sata (nrITS-Sequencing) reveal Relationships among Iranian endemic Taxa of the Umbelliferae.**
In: Feddes Repertorium 117 (5-6), p. 367–388. DOI: 10.1002/fedr.200611106.

VALIEJO-ROMAN, CARMEN M.; TERENTIEVA, ELENA I.; SAMIGULLIN, TAHIR H.; PIMENOV, MICHAEL G. (2002):
**NrDNA ITS Sequences and affinities of Sino-Himalayan Apioideae (Umbelliferae).**
In: Taxon 51, p. 685–701.

WANG, CHANG-BAO; MA, XIANG-GUANG; HE, XING-JIN (2011):
**A taxonomic re-assessment in the Chinese *Bupleurum* (Apiaceae): Insights from Morphology, nuclear ribosomal internal transcribed Spacer, and Chloroplast (trnH-psbA, matK) Sequences.**
In: Journal of Systematics and Evolution 49 (6), p. 558–589. DOI: 10.1111/j.1759-6831.2011.00157.x.

WANG, QI-ZHI; HE, XING-JIN; ZHOU, SONG-DONG; WU, YUN-KE; YU, YANG; PANG, YUN-LI (2008):
**Phylogenetic Inference of the Genus *Bupleurum* (Apiaceae) in Hengduan Mountains based on Chromosome counts and nuclear ribosomal DNA ITS Sequences.**
In: Journal of Systematics and Evolution 46 (2), p. 142–154.

WANG, ZHI-XIN; DOWNIE, STEPHEN R.; TAN, JIN-BO; LIAO, CHEN-YANG; YU, YANG; HE, XING-JIN (2014):
**Molecular Phylogenetics of *Pimpinella* and allied Genera (Apiaceae), with Emphasis on Chinese native species, inferred from nrDNA ITS and cpDNA Intron Sequence Data.**
In: Nordic Journal of Botany 32 (5), p. 642–657. DOI: 10.1111/j.1756-1051.2013.00343.x.

WEITZEL, CORINNA; RØNSTED, NINA; SPALIK, KRYSZTOF; SIMONSEN, HENRIK TOFT (2014):
**Resurrecting deadly carrots: towards a Revision of *Thapsia* (Apiaceae) based on phylogenetic Analysis of nrITS Sequences and chemical profiles.**
In: Botanical Journal of the Linnean Society 174 (4), p. 620–636. DOI: 10.1111/boj.12144.

WINTER, PIETER J.D.; MAGEE, ANTHONY RICHARD; PHEPHU, NONKULULO; TILNEY, PATRICIA M.; DOWNIE, STEPHEN R.; VAN WYK, BEN-ERIK (2008):
**A new generic Classification for African peucedanoid Species (Apiaceae).**
In: Taxon 57 (2), p. 347–364.

YI, TING-SHUANG; JIN, GUI-HUA; WEN, JUN (2015):

**Chloroplast capture and intra- and inter-continental biogeographic Diversification in the Asian - New World disjunct plant Genus *Osmorhiza* (Apiaceae).**
In: Molecular Phylogenetics and Evolution 85, p. 10–21. DOI: 10.1016/j.ympev.2014.09.028.

**YU, YANG; DOWNIE, STEPHEN R.; HE, XING-JIN; DENG, XIANLAN; YAN, LING (2011):**
**Phylogeny and Biogeography of Chinese *Heracleum* (Apiaceae Tribe Tordylieae) with comments on their Fruit Morphology.**
In: Plant Systematics and Evolution 296 (3-4), p. 179–203. DOI: 10.1007/s00606-011-0486-3.

**YUAN, QING-JUN; ZHANG, BIN; JIANG, DAN; ZHANG, WEN-JING; LIN, TSAI-YUN; WANG, NIAN-HE ET AL. (2015):**
**Identification of Species and Materia Medica within *Angelica* L. (Umbelliferae) based on Phylogeny inferred from DNA barcodes.**
In: molecular Ecology Resources 15 (2), p. 358–371. DOI: 10.1111/1755-0998.12296.

**ZAKHAROVA, EKATERINA A.; DEGTJAREVA, GALINA V.; PIMENOV, MICHAEL G. (2012):**
**Redefined generic Limits of *Carum* (Umbelliferae, Apioideae) and new systematic Placement of some of its Taxa.**
In: Willdenowia 42 (2), p. 149–168. DOI: 10.3372/wi.42.42201.

**ZAKHAROVA, EKATERINA A.; KLJUYKOV, EUGENE V.; DEGTJAREVA, GALINA V.; SAMIGULLIN, TAHIR H.; UKRAINSKAYA, ULIANA A.; DOWNIE, STEPHEN R. (2016):**
**A taxonomic Study of the Genus *Hellenocarum* H.Wolff (Umbelliferae-Apioideae) based on Morphology, Fruit Anatomy, and molecular Data.**
In: Turkish Journal of Botany 40, p. 176–193. DOI: 10.3906/bot-1504-40.

**ZHOU, JING; GONG, XUN; DOWNIE, STEPHEN R.; PENG, HUA (2009):**
**Towards a more robust molecular Phylogeny of Chinese Apiaceae Subfamily Apioideae: additional Evidence from nrDNA ITS and cpDNA Intron (rpl16 and rps16) Sequences.**
In: Molecular Phylogenetics and Evolution 53 (1), p. 56–68. DOI: 10.1016/j.ympev.2009.05.029.

**ZHOU, JING; PENG, HUA; DOWNIE, STEPHEN R.; LIU, ZHEN-WEN; GONG, XUN (2008):**
**A molecular Phylogeny of Chinese Apiaceae Subfamily Apioideae inferred from nuclear ribosomal DNA Internal Transcribed Spacer Sequences.**
In: Taxon 57 (2), p. 402–416.

### Apiales

**CHANDLER, GREGORY T.; PLUNKETT, GREGORY M. (2004):**
**Evolution in Apiales: nuclear and Chloroplast Markers together in (almost) perfect Harmony.**
In: Botanical Journal of the Linnean Society 144 (2), p. 123–147. DOI: 10.1111/j.1095-8339.2003.00247.x.

**DOWNIE, STEPHEN R.; KATZ-DOWNIE, DEBORAH S.; WATSON, MARK F. (2000):**
**A Phylogeny of the flowering plant Family Apiaceae based on Chloroplast DNA rpl16 and rpoC1 Intron Sequences: towards a suprageneric Classification of Subfamily Apioideae.**
In: American Journal of Botany 87 (2), p. 273–292. DOI: 10.2307/2656915.

**PLUNKETT, GREGORY M.; CHANDLER, GREGORY T.; LOWRY, PORTER PRESCOTT II.; PINNEY, STEVEN M.; SPRENKLE, T. P. (2004):**
**Recent Advances in Understanding Apiales and a revised Classification.**
In: South African Journal of Botany 70 (3), p. 371–381.

**PLUNKETT, GREGORY M.; LOWRY, PORTER PRESCOTT (2001):**

**Relationships among "ancient araliads" and their Significance for the Systematics of Apiales.**
In: Molecular Phylogenetics and Evolution 19 (2), p. 259–276. DOI: 10.1006/mpev.2000.0920.

### Apocynaceae

ALVARADO-CÁRDENAS, LEONARDO O.; OCHOTERENA, HELGA (2007):
**A phylogenetic Analysis of the *Cascabela–Thevetia* Species Complex (Plumerieae, Apocynaceae) based on Morphology.**
In: Annals of the Missouri Botanical Garden 94 (2), p. 298–323. DOI: 10.3417/0026-6493(2007)94[298:APAOTC]2.0.CO;2.

BRUYNS, PETER V. (2002):
**Monograph of *Orbea* and *Ballyanthus* (Apocynaceae-Asclepiadoideae-Ceropegieae).**
In: Systematic Botany Monographs 63, p. 1–196.

BRUYNS, PETER V.; FARSI, AMINA AL; TERRY HEDDERSON, A. (2010):
**Phylogenetic Relationships of *Caralluma* R.Br. (Apocynaceae).**
In: Taxon 59 (4), p. 1031–1043.

BRUYNS, PETER V.; KLAK, CORNELIA; HANÁČEK, PAVEL (2014):
**Evolution of the stapeliads (Apocynaceae-Asclepiadoideae) - repeated major Radiation across Africa in an Old World Group.**
In: Molecular Phylogenetics and Evolution 77, p. 251–263. DOI: 10.1016/j.ympev.2014.03.022.

BRUYNS, PETER V.; KLAK, CORNELIA; HANÁČEK, PAVEL (2015):
**Recent Radiation of *Brachystelma* and *Ceropegia* (Apocynaceae) across the Old World against a Background of climatic Change.**
In: Molecular Phylogenetics and Evolution 90, p. 49–66. DOI: 10.1016/j.ympev.2015.04.015.

BRUYNS, PETER V.; KLAK, CORNELIA; HANÁČEK, PAVEL (2017):
**A revised, phylogenetically-based Concept of *Ceropegia* (Apocynaceae).**
In: South African Journal of Botany 112, p. 399–436. DOI: 10.1016/j.sajb.2017.06.021.

BRUYNS, PETER V.; NOWELL, TRACEY L.; HEDDERSON, TERRY A. J. (2005):
**A Revision and phylogenetic Analysis of *Stapeliopsis* (Apocynaceae).**
In: Botanical Journal of the Linnean Society 148, p. 125–155.

ENDRESS, MARY E.; LIEDE-SCHUMANN, SIGRID; MEVE, ULRICH (2014):
**An updated Classification for Apocynaceae.**
In: Phytotaxa 159 (3), p. 175. DOI: 10.11646/phytotaxa.159.3.2.

ENDRESS, MARY E.; VAN DER HAM, RAYMOND W. J. M.; NILSSON, SIWERT; CIVEYREL, LAURE; CHASE, MARK W.; SENNBLAD, BENGT ET AL. (2007):
**A phylogenetic Analysis of Alyxieae (Apocynaceae) based on Rbc L, matK, Trn L Intron, TrnL-f Spacer Sequences, and morphological Characters.**
In: Annals of the Missouri Botanical Garden 94 (1), p. 1–35. DOI: 10.3417/0026-6493(2007)94[1:APAOAA]2.0.CO;2.

FISHBEIN, MARK; CHUBA, DAVID; ELLISON, CHRIS; MASON-GAMER, ROBERTA J.; LYNCH, STEVEN P. (2011):
**Phylogenetic Relationships of *Asclepias* (Apocynaceae) inferred from non-coding Chloroplast DNA Sequences.**
In: Systematic Botany 36 (4), p. 1008–1023. DOI: 10.1600/036364411X605010.

GILBERT, MICHAEL G. (2015):

**_Echidnopsis thulinii_ sp. nov. (Apocynaceae-Asclepiadoideae) with a Review of the infrageneric Classification of the Genus.**
In: Nordic Journal of Botany 33 (6), p. 646–654. DOI: 10.1111/njb.00775.

**Goyder, David J. (2016):**
**The correct name for _Matelea chacoensis_ Goyder (Apocynaceae: Asclepiadoideae).**
In: Kew Bulletin 71 (2), p. 232. DOI: 10.1007/S12225-016-9631-Z.

**Goyder, David J.; Nicholas, Ashley; Liede-Schumann, Sigrid (2007):**
**Phylogenetic Relationships in Subtribe Asclepiadinae (Apocynaceae: Asclepiadoideae) 1.**
In: Annals of the Missouri Botanical Garden 94 (2), p. 423–434. DOI: 10.3417/0026-6493(2007)94[423:PRISAA]2.0.CO;2.

**Hechem, Viviana; Calviño, Carolina Isabel; Ezcurra, Cecilia (2011):**
**Molecular Phylogeny of _Diplolepis_ (Apocynaceae-Asclepiadoideae) and allied Genera, and taxonomic Implications.**
In: Taxon 60 (3), p. 638–648.

**Ionta, Gretchen M.; Judd, Walter p. (2007):**
**Phylogenetic Relationships in Periplocoideae (Apocynaceae S.l.) and Insights into the Origin of Pollinia.**
In: Annals of the Missouri Botanical Garden 94 (2), p. 360–375. DOI: 10.3417/0026-6493(2007)94[360:PRIPAS]2.0.CO;2.

**Joubert, Lize; Klak, Cornelia; Venter, Andor M.; Venter, Hendrik J.T.; Bruyns, Peter V. (2016):**
**A widespread Radiation in the Periplocoideae (Apocynaceae): the case of _Cryptolepis_.**
In: Taxon 65 (3), p. 487–501. DOI: 10.12705/653.4.

**Khanum, Rizwana; Surveswaran, Siddharthan; Meve, Ulrich; Liede-Schumann, Sigrid (2016):**
**_Cynanchum_ (Apocynaceae: Asclepiadoideae): A Pantropical Asclepiadoid Genus revisited.**
In: Taxon 65 (3), p. 467–486. DOI: 10.12705/653.3.

**Konno, Tatiana U. P.; Rapini, Alessandro; Goyder, David J.; Chase, Mark W. (2006):**
**The new Genus _Minaria_ (Asclepiadoideae, Apocynaceae).**
In: Taxon 55 (2), p. 421–430.

**Krings, Alexander; Thomas, David T.; Xiang, Qiu-Yun (2008):**
**On the Generic Circumscription of _Gonolobus_ (Apocynaceae, Asclepiadoideae): Evidence from Molecules and Morphology.**
In: Systematic Botany 33 (2), p. 403–415.

**Kunze, Henning; Wanntorp, Livia (2008):**
**The Gynostegium of _Hoya spartioides_ (Apocynaceae – Asclepiadoideae): A striking Case of Incongruence between molecular and phenotypic Evolution.**
In: Organisms Diversity and Evolution 8 (5), p. 346–357. DOI: 10.1016/j.ode.2008.06.002.

**Lahaye, Renaud; Civeyrel, Laure; Speck, Thomas; Rowe, Nick P. (2005):**
**Evolution of Shrub-like Growth Forms in the Lianoid Subfamily Secamonoideae (Apocynaceae S.l.) of Madagascar: Phylogeny, Biomechanics, and Development.**
In: American Journal of Botany 92 (8), p. 1381–1396.

**Lahaye, Renaud; Klackenberg, Jens; Källersjö, Mari; van Campo, Elise; Civeyrel, Laure (2007):**

**Phylogenetic Relationships between derived Apocynaceae s.l. and within Secamonoideae based on Chloroplast Sequences.**
In: Annals of the Missouri Botanical Garden 94 (2), p. 376–391. DOI: 10.3417/0026-6493(2007)94[376:PRBDAS]2.0.CO;2.

LIEDE, SIGRID (2001):
**Subtribe Astephaninae (Apocynaceae-Asclepiadoideae) Reconsidered: new Evidence based on cpDNA Spacers.**
In: Annals of the Missouri Botanical Garden 88 (4), p. 657–668.

LIEDE, SIGRID; KUNZE, HENNING (2002):
***Cynanchum* and the Cynanchinae (Apocynaceae? Asclepiadoideae): a molecular, anatomical and latex triterpenoid study.**
In: Organisms Diversity and Evolution 2 (3), p. 239–269. DOI: 10.1078/1439-6092-00045.

LIEDE, SIGRID; MEVE, ULRICH; TÄUBER, ANGELIKA (2002):
**What is the Subtribe Glossonematinae (Apocynaceae: Asclepiadoideae)? A phylogenetic Study based on cpDNA spacer.**
In: Botanical Journal of the Linnean Society 139, p. 145–158.

LIEDE, SIGRID; TÄUBER, ANGELIKA (2000):
***Sarcostemma* R.Br. (Apocynaceae - Asclepiadoideae) - a controversial generic Circumscription reconsidered: Evidence from trnL-F spacers.**
In: Plant Systematics and Evolution 225, p. 133–140.

LIEDE-SCHUMANN, SIGRID; DÖTTERL, STEFAN; GEBAUER, MARGIT; MEVE, ULRICH (2013):
**A Rapd Study of the *Sarcostemma* Group of *Cynanchum* (Apocynaceae-Asclepiadoideae-Asclepiadeae).**
In: Organisms Diversity and Evolution 13 (1), p. 15–31. DOI: 10.1007/s13127-012-0099-x.

LIEDE-SCHUMANN, SIGRID; KHANUM, RIZWANA; MUMTAZ, ABDUL SAMAD; GHERGHEL, IULIAN; PAHLEVANI, AMIRHOSSEIN (2016):
**Going west - A subtropical Lineage (*Vincetoxicum*, Apocynaceae: Asclepiadoideae) expanding into Europe.**
In: Molecular Phylogenetics and Evolution 94, p. 436–446. DOI: 10.1016/j.ympev.2015.09.021.

LIEDE-SCHUMANN, SIGRID; KONG, HANGHUI; MEVE, ULRICH; THIV, MIKE (2012):
***Vincetoxicum* and *Tylophora* (Apocynaceae: Asclepiadoideae: Asclepiadeae).Two sides of the same medal: Independent Shifts from tropical to temperate Habitats.**
In: Taxon 61 (4), p. 803–825.

LIEDE-SCHUMANN, SIGRID; MEVE, ULRICH (2013):
**The Orthosiinae revisited (Apocynaceae, Asclepiadoideae, Asclepiadeae) 1.**
In: Annals of the Missouri Botanical Garden 99 (1), p. 44–81. DOI: 10.3417/2010130.

LIEDE-SCHUMANN, SIGRID; RAPINI, ALESSANDRO; GOYDER, DAVID J.; CHASE, MARK W. (2005):
**Phylogenetics of the New World Subtribes of Asclepiadeae (Apocynaceae - Asclepiadoideae): Metastelmatinae, Oxypetalinae, and Gonolobinae.**
In: Systematic Botany 30 (1), p. 184–195. DOI: 10.1600/0363644053661832.

LIVSHULTZ, TATYANA (2010):
**The phylogenetic Position of milkweeds (Apocynaceae subfamilies Secamonoideae and Asclepiadoideae): Evidence from the Nucleus and Chloroplast.**
In: Taxon 59 (4), p. 1016–1030.

**Mangelsdorff, Ralph D.; Meve, Ulrich; Liede-Schumann, Sigrid (2016):**
**Phylogeny and Circumscription of Antillean *Anemotrochus*, gen. nov., and *Tylodontia* (Apocynaceae : Asclepiadoideae : Gonolobinae ).**
In: Willdenowia 46 (3), p. 443–474. DOI: 10.3372/wi.46.46311.

**McDonnell, Angela; Fishbein, Mark (2016):**
***Polystemma canisferum* (Apocynaceae, Asclepiadoideae): a distinctive new gonoloboid milkweed Vine from Sonora, Mexico.**
In: Phytotaxa 246 (1), p. 78. DOI: 10.11646/phytotaxa.246.1.6.

**Meve, Ulrich; Liede, Sigrid (2001):**
**Inclusion of *Tenaris* and *Macropetalum* in *Brachystelma* (Apocynaceae - Asclepioideae - Cecropegieae) inferres from non-cosing nuclear and Chloroplast DNA Sequences.**
In: Plant Systematics and Evolution 228, p. 89–105.

**Meve, Ulrich; Liede, Sigrid (2002):**
**A molecular Phylogeny and generic Rearrangement of the stapelioid Ceropegieae (Apocynaceae-Asclepiadoideae).**
In: Plant Systematics and Evolution 234 (1), p. 171–209. DOI: 10.1007/s00606-002-0220-2.

**Meve, Ulrich; Liede, Sigrid (2004):**
**Generic Delimitations in tuberous Periplocoideae (Apocynaceae) from Africa and Madagascar.**
In: Annals of Botany 93 (4), p. 407–414. DOI: 10.1093/aob/mch057.

**Meve, Ulrich; Liede-Schumann, Sigrid (2007):**
**Ceropegia (Apocyneae, Ceropegieae, Stapeliinae): Paraphyletic but still Taxonomically sound.**
In: Annals of the Missouri Botanical Garden 94 (2), p. 392–406. DOI: 10.3417/0026-6493(2007)94[392:CACSPB]2.0.CO;2.

**Meve, Ulrich; Liede-Schumann, Sigrid (2015):**
**Taxonomy of the Andean Genus *Pentacyphus* (Apocynaceae: Asclepiadeae–Pentacyphinae).**
In: Plant Systematics and Evolution 301 (3), p. 997–1004. DOI: 10.1007/s00606-014-1130-9.

**Morales, J. Francisco; Endress, Mary E.; Liede-Schumann, Sigrid (2017):**
**Sex, drugs and pupusas: Disentangling Relationships in Echiteae (Apocynaceae).**
In: Taxon 66 (3), p. 623–644. DOI: 10.12705/663.7.

**Morillo, Gilberto (2012):**
**Aportes al Conocimiento de las Gonolobinae (Apocynaceae - Asclepiadoideae).**
In: Pittieria 36, p. 13–57.

**Nazar, Nazia; Goyder, David J.; Clarkson, James J.; Mahmood, Tariq; Chase, Mark W. (2013):**
**The Taxonomy and Systematics of Apocynaceae: where we stand in 2012.**
In: Botanical Journal of the Linnean Society 171, p. 482–490.

**Pereira, Fontella J.; Santos, R. G. P.; Goes, M. B. de; Moral, p. A. Cáceres (2014):**
**Notas Taxonómicas sobre *Hemipogon* subgen. *Astephanopsis*, y Descripción de un nuevo Género (Apocynaceae, Asclepiadoideae, Asclepiadeae, Metastelmatinae).**
In: Bonplandia 23 (1), p. 25–31.

**Rapini, Alessandro (2012):**

**Taxonomy "under construction": advances in the Systematics of Apocynaceae, with Emphasis on the Brazilian Asclepiadoideae.**
In: Rodriguésia 63 (1), p. 75–88.

**Rapini, Alessandro; Chase, Mark W.; Goyder, David J.; Griffiths, Jayne (2003):**
**Asclepiadeae Classification: evaluating the phylogenetic Relationships of New World Asclepiadoideae (Apocynaceae).**
In: Taxon 52, p. 33–50.

**Rapini, Alessandro; Chase, Mark W.; Konno, Tatiana U. P. (2006):**
**Phylogenetics of South American Asclepiadoideae (Apocynaceae).**
In: Taxon 55 (1), p. 119–124.

**Ribeiro, Patrícia Luz; Rapini, Alessandro; Silva, Uiara Catharina Soares E.; van den Berg, Cássio (2012):**
**Using multiple analytical methods to improve phylogenetic hypotheses in *Minaria* (Apocynaceae).**
In: Molecular Phylogenetics and Evolution 65 (3), p. 915–925. DOI: 10.1016/j.ympev.2012.08.019.

**Rodda, Michele (2016):**
**Checklist and typification of *Heterostemma* (Apocynaceae, Asclepiadoideae, Ceropegieae).**
In: Phytotaxa 263 (1), p. 1. DOI: 10.11646/phytotaxa.263.1.1.

**Sennblad, Bengt; Bremer, Birgitta (2002):**
**Classification of Apocynaceae s.l. according to a new Approach combining Linnaean and phylogenetic Taxonomy.**
In: Systematic Biology 51 (3), p. 389–409. DOI: 10.1080/10635150290069869.

**Simões, André Olmos; Endress, Mary E.; Conti, Elena (2010):**
**Systematics and Character Evolution of Tabernaemontaneae (Apocynaceae, Rauvolfioideae) based on molecular and morphological Evidence.**
In: Taxon 59 (3), p. 772–790.

**Simões, André Olmos; Endress, Mary E.; van der Niet, Timotheüs; Kinoshita, Luiza S.; Conti, Elena (2006):**
**Is *Mandevilla* (Apocynaceae, Mesechiteae) monophyletic? Evidence from five Plastid DNA Loci and Morphology.**
In: Annals of the Missouri Botanical Garden 93 (4), p. 565–591. DOI: 10.3417/0026-6493(2006)93[565:IMAMME]2.0.CO;2.

**Simões, André Olmos; Kinoshita, Luiza S.; Koch, Ingrid; Silva, Márcio José; Endress, Mary E. (2016):**
**Systematics and Character Evolution of Vinceae (Apocynaceae).**
In: Taxon 65 (1), p. 99–122. DOI: 10.12705/651.7.

**Simões, André Olmos; Livshultz, Tatyana; Conti, Elena; Endress, Mary E. (2007):**
**Phylogeny and Systematics of the Rauvolfioideae (Apocynaceae) based on molecular and morphological Evidence.**
In: Annals of the Missouri Botanical Garden 94 (2), p. 268–297. DOI: 10.3417/0026-6493(2007)94[268:PASOTR]2.0.CO;2.

**Simoes, André; Endress, Mary E.; van der Niet, Timotheüs; Kinoshita, Luiza S.; Conti, Elena (2004):**
**Tribal and Intergeneric Relationships of Mesechiteae (Apocynoideae, Apocynaceae): Evidence from three noncoding Plastid DNA Regions and Morphology.**
In: American Journal of Botany 91 (9), p. 1409–1418.

**Straub, Shannon C. K.; Moore, Michael J.; Soltis, Pamela S.; Soltis, Douglas E.; Liston, Aaron; Livshultz, Tatyana (2014):**
**Phylogenetic Signal Detection from an ancient rapid Radiation: Effects of noise Reduction, long-branch Attraction, and model Selection in Crown Clade Apocynaceae.**
In: Molecular Phylogenetics and Evolution 80, p. 169–185. DOI: 10.1016/j.ympev.2014.07.020.

**Surveswaran, Siddharthan; Kamble, Mayur Y.; Yadav, Shrirang Ramchandra; Sun, Mei (2009):**
**Molecular Phylogeny of *Ceropegia* (Asclepiadoideae, Apocynaceae) from Indian Western Ghats.**
In: Plant Systematics and Evolution 281 (1-4), p. 51–63. DOI: 10.1007/s00606-009-0182-8.

**Surveswaran, Siddharthan; Sun, Mei; Grimm, Guido W.; Liede-Schumann, Sigrid (2014):**
**On the systematic Position of some Asian enigmatic Genera of Asclepiadoideae (Apocynaceae).**
In: Botanical Journal of the Linnean Society 174, p. 601–619.

**Thiv, Mike; Meve, Ulrich (2007):**
**A phylogenetic Study of *Echidnopsis* Hook. f. (Apocynaceae-Asclepiadoideae) - taxonomic Implications and the Colonization of the Socotran Archipelago.**
In: Plant Systematics and Evolution 265 (1-2), p. 71–86. DOI: 10.1007/s00606-007-0516-3.

**Turner, Ian M. (2016):**
**Two new Combinations in tropical Apocynaceae.**
In: Webbia 71 (2), p. 227–228. DOI: 10.1080/00837792.2016.1186336.

**Uemachi, Azusa; Shimomura, Takashi (2013):**
**Molecular Phylogeny of *Tachelospermum* (Apocynaceae) in Japan based on cpDNA and nrDNA nucleotide Sequences.**
In: Acta Phytotaxonomica Geobotanica 64 (1), p. 1–13.

**Ungaretti, Tatiana; Konno, Paleo; Pereira, Jorge Fontella (2004):**
**Some Nomenclatural and taxonomic Notes on Brazilian *Ditassa* (Apocynaceae: Asclepiadoideae).**
In: Kew Bulletin 59 (2), p. 297–300.

**Venter, Hendrik J.T. (2008):**
**Taxonomy of *Chlorocyathus* (Apocynaceae: Periplocoideae).**
In: South African Journal of Botany 74 (2), p. 288–294. DOI: 10.1016/j.sajb.2007.12.004.

**Wanntorp, Livia (2007):**
**Pollinaria of *Hoya* (Marsdenieae, Apocynaceae) - shedding light on molecular Phylogenetics.**
In: Taxon 56 (2), p. 465–478.

**Wanntorp, Livia (2009):**
**Phylogenetic Systematics of *Hoya* (Apocynaceae ).**
In: Blumea 54 (1), p. 228–232. DOI: 10.3767/000651909X476201.

**Wanntorp, Livia; Forster, Paul I. (2007):**
**Phylogenetic Relationships between Hoya and the monotypic Genera *Madangia, Absolmsia,* and *Micholitzia* (Apocynaceae, Marsdenieae): Insights from Flower Morphology.**
In: Annals of the Missouri Botanical Garden 94 (1), p. 36–55. DOI: 10.3417/0026-6493(2007)94[36:PRBHAT]2.0.CO;2.

WANNTORP, LIVIA; GOTTHARDT, KATHERINA; MÜLLNER, ALEXANDRA N. (2011):

**Revisiting the wax plants (Hoya, Marsdenieae, Apocynaceae): phylogenetic Tree using the matK Gene and psbA-trnH intergenic Spacer.**

In: Taxon 60 (1), p. 4–14.

WANNTORP, LIVIA; KOCYAN, ALEXANDER; RENNER, SUSANNE P. (2006):

**Wax plants disentangled: a Phylogeny of *Hoya* (Marsdenieae, Apocynaceae) inferred from nuclear and Chloroplast DNA Sequences.**

In: Molecular Phylogenetics and Evolution 39 (3), p. 722–733. DOI: 10.1016/j.ympev.2006.01.022.

WANNTORP, LIVIA; KOCYAN, ALEXANDER; VAN DONKELAAR, RUURD; RENNER, SUSANNE P. (2006):

**Towards a Monophyletic *Hoya* (Marsdenieae, Apocynaceae): Inferences from the Chloroplast trnL Region and the rbcL-atpB Spacer.**

In: Systematic Botany 31 (3), p. 586–596.

WANNTORP, LIVIA; KUNZE, HENNING (2009):

**Identifying Synapomorphies in the Flowers of *Hoya* and *Dischidia* -Toward phylogenetic Understanding.**

In: International Journal of Plant Sciences 170 (3), p. 331–342. DOI: 10.1086/596329.

WEIDE, JASMIJN C. VAN DER; HAM, RAYMOND W.J.M. VAN DER (2012):

**Pollen Morphology and Phylogeny of the Tribe Tabernaemontaneae (Apocynaceae, Subfamily Rauvolfioideae).**

In: Taxon 61 (1), p. 131–145.

YAMASHIRO, TADASHI; FUKUDA, TATSUYA; YOKOYAMA, JUN U.N.; MAKI, MASAYUKI (2004):

**Molecular Phylogeny of *Vincetoxicum* (Apocynaceae-Asclepiadoideae) based on the nucleotide Sequences of cpDNA and nrDNA.**

In: Molecular Phylogenetics and Evolution 31 (2), p. 689–700. DOI: 10.1016/j.ympev.2003.08.016.

### Apodanthaceae

BELLOT, SIDONIE; RENNER, SUSANNE P. (2014):

**Exploring new dating Approaches for Parasites: the worldwide Apodanthaceae (Cucurbitales) as an example.**

In: Molecular Phylogenetics and Evolution 80, p. 1–10. DOI: 10.1016/j.ympev.2014.07.005.

BLARER, ALBERT; NICKRENT, DANIEL LEE; ENDRESS, PETER K. (2004):

**Comparative Floral Structure and Systematics in Apodanthaceae (Rafflesiales).**

In: Plant Systematics and Evolution 245 (1-2), p. 119–142. DOI: 10.1007/s00606-003-0090-2.

FILIPOWICZ, NATALIA; RENNER, SUSANNE P. (2010):

**The worldwide holoparasitic Apodanthaceae confidently placed in the Cucurbitales by nuclear and mitochondrial Gene Trees.**

In: BMC Evolutionary Biology 10, p. 219. DOI: 10.1186/1471-2148-10-219.

### Aponogetonaceae

CHEN, LING-YUN; GRIMM, GUIDO W.; WANG, QING-FENG; RENNER, SUSANNE P. (2015):

**A Phylogeny and biogeographic Analysis for the Cape-Pondweed Family Aponogetonaceae (Alismatales).**

In: Molecular Phylogenetics and Evolution 82 Pt A, p. 111–117. DOI: 10.1016/j.ympev.2014.10.007.

LES, DONALD H.; MOODY, MICHAEL L.; JACOBS, SURREY W.L. (2005):

**Phylogeny and Systematics of *Aponogeton* (Aponogetonaceae): the Australian Species.**

In: Systematic Botany 30 (3), p. 503–519. DOI: 10.1600/0363644054782215.

### Aquifoliaceae

GOTTLIEB, ALEXANDRA M.; GIBERTI, GUSTAVO C.; POGGIO, LIDIA (2005):
**Molecular Analyses of the Genus *Ilex* (Aquifoliaceae) in Southern South America, Evidence from AFLP and ITS Sequence Data.**
In: American Journal of Botany 92 (2), p. 352–369.

HONG, DE-YUAN (2015):
**A Taxonomical Revision of *Ilex* (Aquifoliaceae) in the Pan-Himalaya and unraveling its Distribution Patterns.**
In: Phytotaxa 230 (2), p. 151. DOI: 10.11646/phytotaxa.230.2.3.

MANEN, JEAN-FRANÇOIS; BARRIERA, GABRIELLE; LOIZEAU, PIERRE-ANDRÉ; NACIRI, YAMAMA (2010):
**The History of extant *Ilex* Species (Aquifoliaceae): Evidence of Hybridization within a Miocene Radiation.**
In: Molecular Phylogenetics and Evolution 57 (3), p. 961–977. DOI: 10.1016/j.ympev.2010.09.006.

MANEN, JEAN-FRANÇOIS; BOULTER, M. C.; NACIRI-GRAVEN, Y. (2002):
**The complex History of the Genus *Ilex* L. (Aquifoliaceae): Evidence from the Comparison of Plastid and nuclear DNA Sequences and from fossil Data.**
In: Plant Systematics and Evolution 235 (1), p. 79–98. DOI: 10.1007/s00606-002-0225-x.

SELBACH-SCHNADELBACH, ALESSANDRA; CAVALLI, SUZANA SMITH; MANEN, JEAN-FRANÇOIS; COELHO, GERALDO CENI; SOUZA-CHIES, TATIANA TEIXEIRA DE (2009):
**New information for Ilex Phylogenetics based on the Plastid psbA-trnH intergenic spacer (Aquifoliaceae).**
In: Botanical Journal of the Linnean Society 159, p. 182–193.

### Araceae

CABRERA, LIDIA I.; SALAZAR, GERARDO A.; CHASE, MARK W.; MAYO, SIMON J.; BOGNER, JOSEF; DÁVILA, PATRICIA (2008):
**Phylogenetic Relationships of aroids and duckweeds (Araceae) inferred from coding and noncoding Plastid DNA.**
In: American Journal of Botany 95 (9), p. 1153–1165. DOI: 10.3732/ajb.0800073.

CARLSEN, MÓNICA M.; CROAT, THOMAS B. (2013):
**A molecular Phylogeny of the species-rich Neotropical Genus *Anthurium* (Araceae) based on combined Chloroplast and Nuclear DNA.**
In: Systematic Botany 38 (3), p. 576–588. DOI: 10.1600/036364413X670287.

CUSIMANO, NATALIE; BARRETT, MATTHEW D.; HETTERSCHEID, WILBERT L.A.; RENNER, SUSANNE P. (2010):
**A Phylogeny of the Areae (Araceae) implies that *Typhonium, Sauromatum*, and the Australian Species of *Typhonium* are distinct Clades.**
In: Taxon 59 (2), p. 439–447.

CUSIMANO, NATALIE; BOGNER, JOSEF; MAYO, SIMON J.; BOYCE, PETER C.; WONG, SIN YENG; HESSE, MICHAEL ET AL. (2011):
**Relationships within the Araceae: Comparison of morphological Patterns with molecular Phylogenies.**
In: American Journal of Botany 98 (4), p. 654–668. DOI: 10.3732/ajb.1000158.

ESPÍNDOLA, ANAHÍ; BUERKI, SVEN; BEDALOV, MARIJA; KÜPFER, PHILIPPE; ALVAREZ, NADIR (2010):

**New Insights into the Phylogenetics and Biogeography of *Arum* (Araceae): unravelling its evolutionary History.**
In: Botanical Journal of the Linnean Society 163 (1), p. 14–32. DOI: 10.1111/j.1095-8339.2010.01049.x.

GAUTHIER, MARIE-PIERRE L.; BARABÉ, DENIS; BRUNEAU, ANNE (2008):
**Molecular Phylogeny of the Genus *Philodendron* (Araceae): Delimitation and infrageneric Classification.**
In: Botanical Journal of the Linnean Society 156 (1), p. 13–27. DOI: 10.1111/j.1095-8339.2007.00733.x.

GONÇALVES, EDUARDO G.; MAYO, SIMON J.; VAN SLUYS, MARIE-ANNE; SALATINO, ANTONIO (2007):
**Combined genotypic-phenotypic Phylogeny of the Tribe Spathicarpeae (Araceae) with Reference to independent Events of Invasion to Andean Regions.**
In: Molecular Phylogenetics and Evolution 43 (3), p. 1023–1039. DOI: 10.1016/j.ympev.2007.01.008.

GROB, G. B.J.; GRAVENDEEL, BARBARA; EURLINGS, MARCEL C.M. (2004):
**Potential phylogenetic utility of the nuclear Floricaula/leafy second Intron: Comparison with three Chloroplast DNA Regions in *Amorphophallus* (Araceae).**
In: Molecular Phylogenetics and Evolution 30 (1), p. 13–23. DOI: 10.1016/S1055-7903(03)00183-0.

GROB, G. B.J.; GRAVENDEEL, BARBARA; EURLINGS, MARCEL C.M.; HETTERSCHEID, WILBERT L.A. (2002):
**Phylogeny of the Tribe Thomsonieae (Araceae) based on Chloroplast matK and trnL Intron Sequences.**
In: Systematic Botany 27 (3), p. 453–467.

HENRIQUEZ, CLAUDIA L.; ARIAS, TATIANA; PIRES, J. CHRIS; CROAT, THOMAS B.; SCHAAL, BARBARA A. (2014):
**Phylogenomics of the Plant Family Araceae.**
In: Molecular Phylogenetics and Evolution 75, p. 91–102. DOI: 10.1016/j.ympev.2014.02.017.

HETTERSCHEID, WILBERT L.A.; CLAUDEL, CYRILLE (2012):
**The End of *Pseudodracontium* N.E.Br.**
In: Aroideana 35, p. 40–46.

LES, DONALD H.; CRAWFORD, DANIEL J.; LANDOLT, ELIAS; GABEL, JOHN D.; KIMBALL, REBECCA T. (2002):
**Phylogeny and Systematics of Lemnaceae, the Duckweed Family.**
In: Systematic Botany 27 (2), p. 221–240.

LIN, JEANINE; STÖKL, JOHANNES; URRU, ISABELLA; KRÜGEL, TAMARA; STENSMYR, MARCUS C. (2010):
**Molecular Phylogeny of the Genus *Arum* (Araceae) inferred from multi-locus Sequence Data and AFLPs.**
In: Taxon 59 (2), p. 405–415.

LOW, SHOOK LING; WONG, SIN YENG; BOYCE, PETER C. (2014):
***Schottarum* (Schismatoglottideae: Araceae) substantiated based on combined nuclear and Plastid DNA Sequences.**
In: Plant Systematics and Evolution 300 (4), p. 607–617. DOI: 10.1007/s00606-013-0906-7.

MAGEE, ANTHONY RICHARD; VAN WYK, BEN-ERIK; TILNEY, PATRICIA M.; DOWNIE, STEPHEN R. (2010):
**Phylogenetic Position of African and Malagasy *Pimpinella* Species and related Genera (Apiaceae, Pimpinelleae).**
In: Plant Systematics and Evolution 288 (3-4), p. 201–211. DOI: 10.1007/s00606-010-0325-y.

MEKKERDCHOO, ORACHORN; BOROMPICHAICHARTKUL, CHALEEDA; PERRIGO, ALLISON; SRZEDNICKI, GEORGE; PRAKITCHAIWATTANA, CHEUNJIT; ANTONELLI, ALEXANDRE (2016):
**Tracing the Evolution and Economic Potential of Konjac Glucomannan in *Amorphophallus* Species (Araceae) using molecular Phylogeny and Rapd Markers.**

In: Phytotaxa 282 (2), p. 81. DOI: 10.11646/phytotaxa.282.2.1.

**Nauheimer, Lars; Boyce, Peter C. (2014):**
***Englerarum* (Araceae, Aroideae): a new Genus supported by Plastid and nuclear Phylogenies.**
In: Plant Systematics and Evolution 300 (4), p. 709–715. DOI: 10.1007/s00606-013-0914-7.

**Nauheimer, Lars; Boyce, Peter C.; Renner, Susanne p. (2012):**
**Giant taro and its relatives: a Phylogeny of the large Genus *Alocasia* (Araceae) sheds Light on Miocene floristic Exchange in the Malesian Region.**
In: Molecular Phylogenetics and Evolution 63 (1), p. 43–51. DOI: 10.1016/j.ympev.2011.12.011.

**Nie, Ze-long; Sun, Hang; Li, Heng; Wen, Jun (2006):**
**Intercontinental Biogeography of Subfamily Orontioideae (Symplocarpus, Lysichiton, and Orontium) of Araceae in Eastern Asia and North America.**
In: Molecular Phylogenetics and Evolution 40 (1), p. 155–165. DOI: 10.1016/j.ympev.2006.03.012.

**Ohi-Toma, Tetsuo; Wu, Sugong; Murata, Hiroko; Murata, Jin (2016):**
**An updated Genus-wide phylogenetic Analysis of *Arisaema* (Araceae) with Reference to Sections.**
In: Botanical Journal of the Linnean Society 182 (1), p. 100–114. DOI: 10.1111/boj.12459.

**Ohi-Toma, Tetsuo; Wu, Sugong; Yadav, Shrirang Ramchandra; Murata, Hiroko; Murata, Jin (2010):**
**Molecular Phylogeny of *Typhonium* sensu lato and its allied Genera in the Tribe Areae of the Subfamily Aroideae (Araceae) based on Sequences of six Chloroplast Regions.**
In: Systematic Botany 35 (2), p. 244–251. DOI: 10.1600/036364410791638315.

**Renner, Susanne S.; Zhang, Li-Bing (2004):**
**Biogeography of the *Pistia* Clade (Araceae): based on Chloroplast and Mitochondrial DNA Sequences and Bayesian Divergence Time Inference.**
In: Systematic Biology 53 (3), p. 422–432. DOI: 10.1080/10635150490445904.

**Renner, Susanne S.; Zhang, Li-Bing; Murata, Jin (2004):**
**A Chloroplast Phylogeny of *Arisaema* (Araceae) illustrates Tertiary floristic links between Asia, North America, and East Africa.**
In: American Journal of Botany 91 (6), p. 881–888. DOI: 10.3732/ajb.91.6.881.

**Rothwell, Gar W.; van Atta, Michelle R.; Ballard, Harvey Eugene; Stockey, Ruth A. (2004):**
**Molecular phylogenetic Relationships among Lemnaceae and Araceae using the Chloroplast trnL–trnF intergenic Spacer.**
In: Molecular Phylogenetics and Evolution 30 (2), p. 378–385. DOI: 10.1016/S1055-7903(03)00205-7.

**Soares, Maria de Lourdes; Mayo, Simon J.; Gribel, Rogério (2013):**
**A Preliminary taxonomic Revision of *Heteropsis* (Araceae).**
In: Systematic Botany 38 (4), p. 925–974. DOI: 10.1600/036364413X674715.

**Tam, Sheh-May; Boyce, Peter C.; Upson, Tim M.; Barabé, Denis; Bruneau, Anne; Forest, Félix; Parker, John p. (2004):**
**Intergeneric and infrafamilial Phylogeny of Subfamily Monsteroideae (Araceae) revealed by Chloroplast trnL-F Sequences.**
In: American Journal of Botany 91 (3), p. 490–498. DOI: 10.3732/ajb.91.3.490.

**Tippery, Nicholas P.; Les, Donald H.; Crawford, Daniel J. (2015):**
**Evaluation of phylogenetic Relationships in Lemnaceae using nuclear ribosomal Data.**

In: Plant Biology 17 Suppl 1, p. 50–58. DOI: 10.1111/plb.12203.

**TRUDGEN, MELINDA S.; BAKER, WILLIAM J. (2008):**
**A Revision of the *Heterospathe elegans* (Arecaceae) complex in New Guinea.**
In: Kew Bulletin 63 (4), p. 639–647. DOI: 10.1007/s12225-008-9063-5.

**VASCONCELOS, SANTELMO; SOARES, MARIA DE LOURDES; SAKURAGUI, CÁSSIA M.; CROAT, THOMAS B.; OLIVEIRA, GUILHERME; BENKO-ISEPPON, ANA MARIA (2018):**
**New Insights on the phylogenetic Relationships among the traditional *Philodendron* subgenera and the other Groups of the *Homalomena* Clade (Araceae).**
In: Molecular Phylogenetics and Evolution 127, p. 168–178. DOI: 10.1016/j.ympev.2018.05.017.

**WONG, SIN YENG; BOYCE, PETER C.; OTHMAN, AHMAD SOFIMAN; PIN, LEAW CHUI (2010):**
**Molecular Phylogeny of Tribe Schismatoglottideae (Araceae) based on two Plastid Markers and Recognition of a new Tribe, Philonotieae, from the neotropics.**
In: Taxon 59 (1), p. 117–124.

**YENG, WONG SIN; MEEROW, ALAN W.; CROAT, THOMAS B. (2016):**
**Resurrection and new Species of the Neotropical Genus *Adelonema* (Araceae: *Philodendron* Clade).**
In: Systematic Botany 41 (1), p. 32–48. DOI: 10.1600/036364416X690732.

### Araliaceae

**ACKERFIELD, JENNIFER; WEN, JUN (2003):**
**Evolution of *Hedera* (the Ivy Genus, Araliceae): Insights form Chloroplast DNA Data.**
In: International Journal of Plant Sciences 164 (4), p. 593–602.

**CHOI, KYOUNG-SU; PARK, SEON-JOO (2012):**
**Molecular phylogenetic Studies of Korean *Hydrocotyle* L.**
In: Korean Journal of Plant Resources 25 (4), p. 490–497. DOI: 10.7732/kjpr.2012.25.4.490.

**COI, HONG-KEUN; WEN, JUN (2000):**
**A phylogenetic Analysis of *Panax* (Araliaceae): integrating cpDNA restriction site and nuclear rDNA ITS Sequence Data.**
In: Plant Systematics and Evolution 224, p. 109–120.

**COSTELLO, ANNEMARIE; MOTLEY, TIMOTHY J. (2007):**
**Phylogenetics of the *Tetraplasandra* Group (Araliaceae) inferred from ITS, 5S-NTS, and Morphology.**
In: Systematic Botany 32 (2), p. 464–477. DOI: 10.1600/036364407781179626.

**FIASCHI, PEDRO; FRODIN, DAVID G. (2014):**
**Two new Combinations in Brazilian *Dendropanax* (Araliaceae).**
In: Phytotaxa 159 (3), p. 236. DOI: 10.11646/phytotaxa.159.3.6.

**GREEN, ADAM F.; RAMSEY, TARA S.; RAMSEY, JUSTIN (2011):**
**Phylogeny and Biogeography of Ivies (*Hedera* spp., Araliaceae), a polyploid Complex of woody Vines.**
In: Systematic Botany 36 (4), p. 1114–1127. DOI: 10.1600/036364411X605100.

**HENWOOD, MURRAY J.; LU-IRVING, PATRICIA; PERKINS, ANDREW J. (2010):**
**Can molecular Systematics provide Insights into aspects of the reproductive biology of *Trachymene* Rudge (Araliaceae)?**
In: Plant Diversity and Evolution 128 (1), p. 85–110. DOI: 10.1127/1869-6155/2010/0128-0004.

**HWANG, HWAN SU; CHOI, YONG EUI (2016):**

**Phylogenetic Analysis of 14 Korean Araliaceae Species using Chloroplast DNA Barcode Analysis.**
In: Journal of Plant Biotechnology 43 (1), p. 82–90. DOI: 10.5010/JPB.2016.43.1.82.

KARUPPUSAMY, S.; ALI, M. AJMAL; RAJASEKARAN, K. M.; LEE, JOONGKU; KIM, SOO-YOUNG; PANDEY, ARUN KUMAR; AL-.HEMAID, FAHAD M. A. (2014):
**A new Species of *Hydrocotyle* L. (Araliaceae) from India.**
In: Bangladesh Journal of Plant Taxononmy 21 (2), p. 167–173.

LEE, CHUNGHEE; WEN, JUN (2004):
**Phylogeny of Panax using Chloroplast trnC-trnD intergenic Region and the utility of trnC-trnD in interspecific studies of plants.**
In: Molecular Phylogenetics and Evolution 31 (3), p. 894–903. DOI: 10.1016/j.ympev.2003.10.009.

LI, RONG; WEN, JUN (2014):
**Phylogeny and Biogeography of Asian *Schefflera* (Araliaceae) based on nuclear and Plastid DNA Sequence Data.**
In: Journal of Systematics and Evolution 52 (4), p. 431–449. DOI: 10.1111/jse.12052.

LI, RONG; WEN, JUN (2016):
**Phylogeny and Diversification of Chinese Araliaceae based on nuclear and Plastid DNA Sequence Data.**
In: Journal of Systematics and Evolution 54 (4), p. 453–467. DOI: 10.1111/jse.12196.

LOWRY, PORTER PRESCOTT; PLUNKETT, GREGORY M. (2010):
**Recircumscription of *Polyscias* (Araliaceae) to include six related Genera, with a new infrageneric Classification and a Synopsis of Species.**
In: Plant Diversity and Evolution 128 (1), p. 55–84. DOI: 10.1127/1869-6155/2010/0128-0003.

LOWRY, PORTER PRESCOTT; PLUNKETT, GREGORY M.; GOSTEL, MORGAN R.; FRODIN, DAVID G. (2017):
**A Synopsis of the Afro-Malagasy Species previously included in *Schefflera* (Araliaceae): Resurrection of the Genera *Astropanax* and *Neocussonia*.**
In: Candollea 72 (2), p. 265–282. DOI: 10.15553/c2017v722a4.

MANZANILLA, V.; KOOL, ANNELEEN; NGUYEN NHAT, L.; NONG VAN, H.; LE THI THU, H.; BOER, HUGO J. (2018):
**Phylogenomics and barcoding of *Panax*: toward the Identification of Ginseng Species.**
In: BMC Evolutionary Biology 18 (1), p. 44. DOI: 10.1186/s12862-018-1160-y.

MITCHELL, ANTHONY D.; LI, RONG; BROWN, JOSEPH W.; SCHÖNBERGER, INES; WEN, JUN (2012):
**Ancient Divergence and Biogeography of *Raukaua* (Araliaceae) and close Relatives in the Southern Hemisphere.**
In: Australian Systematic Botany 25 (6), p. 432. DOI: 10.1071/SB12020.

MITCHELL, ANTHONY D.; WEN, JUN (2005):
**Phylogeny of *Brassaiopsis* (Araliaceae) in Asia based on Nuclear ITS and 5S-NTS DNA Sequences.**
In: Systematic Botany 30 (4), p. 872–886. DOI: 10.1600/036364405775097761.

NICOLAS, ANTOINE N.; PLUNKETT, GREGORY M. (2009):
**The demise of Subfamily Hydrocotyloideae (Apiaceae) and the Re-alignment of its Genera across the entire Order Apiales.**
In: Molecular Phylogenetics and Evolution 53 (1), p. 134–151. DOI: 10.1016/j.ympev.2009.06.010.

PERRIE, LEON R.; SHEPHERD, LARA D. (2009):

**Reconstructing the Species Phylogeny of *Pseudopanax* (Araliaceae), a Genus of Hybridising Trees.**
In: Molecular Phylogenetics and Evolution 52 (3), p. 774–783. DOI: 10.1016/j.ympev.2009.05.030.

**PLUNKETT, GREGORY M.; LOWRY, PORTER PRESCOTT II.; VU, NINH V. (2004):**
**Phylogenetic Relationships among *Polyscias* (Araliaceae) and close Relatives from the Western Indian Ocean Basin.**
In: International Journal of Plant Sciences 165 (5), p. 861–873.

**PLUNKETT, GREGORY M.; WEN, JUN; LOWRY, PORTER PRESCOTT II. (2004):**
**Infrafamilial Classifications and Characters in Araliaceae: Insights from the phylogenetic Analysis of nuclear (ITS) and Plastid (trnL-trnF) Sequence Data.**
In: Plant Systematics and Evolution 245 (1-2). DOI: 10.1007/s00606-003-0101-3.

**TRONCHET, FRÉDÉRIC; PLUNKETT, GREGORY M.; JÉRÉMIE, JOËL; LOWRY, PORTER PRESCOTT (2005):**
**Monophyly and major Clades of *Meryta* (Araliaceae).**
In: Systematic Botany 30 (3), p. 657–670. DOI: 10.1600/0363644054782279.

**VALCÁRCEL, VIRGINIA; FIZ, OMAR; VARGAS, PABLO (2003):**
**Chloroplast and nuclear Evidence for multiple Origins of polyploids and diploids of *Hedera* (Araliaceae) in the Mediterranean basin.**
In: Molecular Phylogenetics and Evolution 27 (1), p. 1–20. DOI: 10.1016/S1055-7903(02)00364-0.

**VALCÁRCEL, VIRGINIA; FIZ-PALACIOS, OMAR; WEN, JUN (2014):**
**The Origin of the early differentiation of ivies (*Hedera* L.) and the Radiation of the Asian Palmate Group (Araliaceae).**
In: Molecular Phylogenetics and Evolution 70, p. 492–503. DOI: 10.1016/j.ympev.2013.10.016.

**VALCÁRCEL, VIRGINIA; VARGAS, PABLO (2010):**
**Quantitative Morphology and Species Delimitation under the General Lineage Concept: Optimization for *Hedera* (Araliaceae).**
In: American Journal of Botany 97 (9), p. 1555–1573. DOI: 10.3732/ajb.1000115.

**VALCÁRCEL, VIRGINIA; VARGAS, PABLO (2013):**
**Phylogenetic Reconstruction of key traits in the Evolution of ivies (*Hedera* L.).**
In: Plant Systematics and Evolution 299 (2), p. 447–458. DOI: 10.1007/s00606-012-0734-1.

**WEN, JUN (2000):**
**Internal Transcribed Space Phylogeny of the Asian and Eastern North American disjunct *Aralia* Sect. *Dimorphanthus* (Araliaceae) and its biogeographic Implications.**
In: International Journal of Plant Sciences 161 (6), p. 959–966.

**WEN, JUN; CHUNGHEE, LEE; LOWRY, PORTER PRESCOTT II.; HIEP, NGUYEN TIEN (2003):**
**Inclusion of the Vietnamese endemic Genus *Grushvitzkya* in *Brassaiopsis* (Araliaceae): Evidence from nuclear ribosomal ITS and Chloroplast ndh F Sequences.**
In: Botanical Journal of the Linnean Society 142, p. 455–463.

**WEN, JUN; PLUNKETT, GREGORY M.; MITCHELL, ANTHONY D.; WAGSTAFF, STEVEN J. (2001):**
**The Evolution of Araliaceae: A phylogenetic Analysis based on ITS Sequences of Nuclear Ribosomal DNA.**
In: Systematic Botany 26 (1), p. 144–167.

**ZUO, YUN-JUAN; WEN, JUN; ZHOU, SHI-LIANG (2017):**

**Intercontinental and intracontinental Biogeography of the eastern Asian - Eastern North American disjunct *Panax* (the Ginseng Genus, Araliaceae), emphasizing its Diversification Processes in Eastern Asia.**
In: Molecular Phylogenetics and Evolution 117, p. 60–74. DOI: 10.1016/j.ympev.2017.06.016.

### Araucariaceae

**Escapa, Ignacio H.; Catalano, Santiago A. (2013):**
**Phylogenetic Analysis of Araucariaceae: Integrating Molecules, Morphology, and Fossils.**
In: International Journal of Plant Sciences 174 (8), p. 1153–1170. DOI: 10.1086/672369.

**Gaudeul, Myriam; Gardner, Martin F.; Thomas, Philip I.; Ennos, Richard A.; Hollingsworth, Peter M. (2014):**
**Evolutionary dynamics of emblematic *Araucaria* Species (Araucariaceae) in New Caledonia: nuclear and Chloroplast Markers suggest recent Diversification, introgression, and a tight Link between Genetics and Geography within Species.**
In: BMC Evolutionary Biology 14, p. 171. DOI: 10.1186/s12862-014-0171-6.

**Gaudeul, Myriam; Rouhan, Germinal; Gardner, Martin F.; Hollingsworth, Peter M. (2012):**
**AFLP Markers provide Insights into the evolutionary Relationships and Diversification of New Caledonian *Araucaria* Species (Araucariaceae).**
In: American Journal of Botany 99 (1), p. 68–81. DOI: 10.3732/ajb.1100321.

**Knapp, Michael; Mudaliar, Ragini; Havell, David; Wagstaff, Steven J.; Lockhart, Peter J. (2007):**
**The drowning of New Zealand and the problem of *Agathis*.**
In: Systematic Biology 56 (5), p. 862–870. DOI: 10.1080/10635150701636412.

**Liu, Nian; Zhu, Yong; Wei, ZongXian; Chen, Jie; Wang, QingBiao; Jian, Shu-Guang et al. (2009):**
**Phylogenetic Relationships and Divergence times of the Family Araucariaceae based on the DNA Sequences of eight Genes.**
In: Science Bulletin 54 (15), p. 2648–2655. DOI: 10.1007/s11434-009-0373-2.

**Stefenon, By V. M.; Gailing, Oliver; Finkeldey, Reiner (2006):**
**Phylogenetic Relationship within Genus *Araucaria* (Araucariaceae) assessed by means of AFLP Fingerprints.**
In: Silvae Genetica 55 (2), p. 45–52.

**Zonneveld, Ben J. M. (2012):**
**Genome sizes of all 19 *Araucaria* Species are correlated with their geographical Distribution.**
In: Plant Systematics and Evolution 298 (7), p. 1249–1255. DOI: 10.1007/s00606-012-0631-7.

### Arecaceae

**Alapetite, Elodie; Baker, William J.; Nadot, Sophie (2014):**
**Evolution of stamen Number in Ptychospermatinae (Arecaceae): Insights from a new molecular Phylogeny of the Subtribe.**
In: Molecular Phylogenetics and Evolution 76, p. 227–240. DOI: 10.1016/j.ympev.2014.02.026.

**Asmussen-Lange, Conny B.; Dransfield, John; Deickmann, Vinnie; Barfod, Anders S.; Pintaud, Jean-Christophe; Baker, William J. (2006):**
**A new Subfamily Classification of the Palm Family (Arecaceae): Evidence from Plastid DNA Phylogeny.**
In: Botanical Journal of the Linnean Society 151 (1), p. 15–38. DOI: 10.1111/j.1095-8339.2006.00521.x.

**Bacon, Christine D.; Baker, William J.; Simmons, Mark P. (2012):**

**Miocene Dispersal drives Island Radiations in the Palm Tribe Trachycarpeae (Arecaceae).**
In: Systematic Biology 61 (3), p. 426–442. DOI: 10.1093/sysbio/syr123.

BACON, CHRISTINE D.; MCKENNA, MILES J.; SIMMONS, MARK P.; WAGNER, WARREN L. (2012):
**Evaluating multiple criteria for Species Delimitation: an empirical example using Hawaiian Palms (Arecaceae: Pritchardia).**
In: BMC Evolutionary Biology 12, p. 23. DOI: 10.1186/1471-2148-12-23.

BAKER, WILLIAM J. (2015):
**A revised Delimitation of the rattan Genus *Calamus* (Arecaceae).**
In: Phytotaxa 197 (2), p. 139. DOI: 10.11646/phytotaxa.197.2.7.

BAKER, WILLIAM J.; DRANSFIELD, JOHN; HEDDERSON, TERRY A. (2000):
**Phylogeny, Character Evolution, and a new Classification of the Calamoid Palms.**
In: Systematic Botany 25 (2), p. 297. DOI: 10.2307/2666644.

BAKER, WILLIAM J.; NORUP, MARIA VIBE; CLARKSON, JAMES J.; COUVREUR, THOMAS L.P.; DOWE, JOHN L.; LEWIS, CARL E. ET AL. (2011):
**Phylogenetic Relationships among Arecoid Palms (Arecaceae: Arecoideae).**
In: Annals of Botany 108 (8), p. 1417–1432. DOI: 10.1093/aob/mcr020.

BAKER, WILLIAM J.; SAVOLAINEN, VINCENT; ASMUSSEN-LANGE, CONNY B.; CHASE, MARK W.; DRANSFIELD, JOHN; FOREST, FÉLIX ET AL. (2009):
**Complete generic-level phylogenetic Analyses of Palms (Arecaceae) with Comparisons of Supertree and Supermatrix Approaches.**
In: Systematic Biology 58 (2), p. 240–256. DOI: 10.1093/sysbio/syp021.

CANO, ÁNGELA; PERRET, MATHIEU; STAUFFER, FRED W. (2013):
**A Revision of the Genus *Trithrinax* (Cryosophileae, Coryphoideae, Arecaceae).**
In: Phytotaxa 136 (1), p. 1. DOI: 10.11646/phytotaxa.136.1.1.

COMER, JASON R.; ZOMLEFER, WENDY B.; BARRETT, CRAIG F.; DAVIS, JERROLD I.; STEVENSON, DENNIS WM.; HEYDUK, KAROLINA; LEEBENS-MACK, JAMES H. (2015):
**Resolving Relationships within the Palm Subfamily Arecoideae (Arecaceae) using Plastid Sequences derived from Next-Generation Sequencing.**
In: American Journal of Botany 102 (6), p. 888–899. DOI: 10.3732/ajb.1500057.

COMER, JASON R.; ZOMLEFER, WENDY B.; BARRETT, CRAIG F.; STEVENSON, DENNIS WM.; HEYDUK, KAROLINA; LEEBENS-MACK, JAMES H. (2016):
**Nuclear phylogenomics of the Palm Subfamily Arecoideae (Arecaceae).**
In: Molecular Phylogenetics and Evolution 97, p. 32–42. DOI: 10.1016/j.ympev.2015.12.015.

CRISP, MICHAEL D.; ISAGI, YUJI; KATO, YOHEI; COOK, LYN G.; BOWMAN, DAVID M. J. P. (2010):
***Livistona* palms in Australia: ancient Relics or opportunistic Immigrants?**
In: Molecular Phylogenetics and Evolution 54 (2), p. 512–523. DOI: 10.1016/j.ympev.2009.09.020.

CUENCA, ARGELIA; ASMUSSEN-LANGE, CONNY B. (2007):
**Phylogeny of the Palm Tribe Chamaedoreeae (Arecaceae) based on Plastid DNA Sequences.**
In: Systematic Botany 32 (2), p. 250–263. DOI: 10.1600/036364407781179644.

CUENCA, ARGELIA; ASMUSSEN-LANGE, CONNY B.; BORCHSENIUS, FINN (2008):
**A dated Phylogeny of the Palm Tribe Chamaedoreeae supports Eocene Dispersal between Africa, North and South America.**
In: Molecular Phylogenetics and Evolution 46 (2), p. 760–775. DOI: 10.1016/j.ympev.2007.10.010.

**Cuenca, Argelia; Dransfield, John; Asmussen-Lange, Conny B. (2009):**
**Phylogeny and Evolution of morphological Characters in Tribe Chamaedoreeae (Arecaceae).**
In: Taxon 58 (4), p. 1092–1108.

**Domenech, Boris; Asmussen-Lange, Conny B.; Baker, William J.; Alapetite, Elodie; Pintaud, Jean-Christophe; Nadot, Sophie (2014):**
**A phylogenetic Analysis of Palm Subtribe Archontophoenicinae (Arecaceae) based on 14 DNA Regions.**
In: Botanical Journal of the Linnean Society 175 (4), p. 469–481. DOI: 10.1111/boj.12179.

**Dransfield, John; Uhl, Natalie W.; Asmussen-Lange, Conny B.; Baker, William J.; Harley, Madeline M.; Lewis, Carl E. (2005):**
**A new phylogenetic Classification of the Palm Family, Arecaceae.**
In: Kew Bulletin 60, p. 559–569.

**Eiserhardt, Wolf L.; Pintaud, Jean-Christophe; Asmussen-Lange, Conny B.; Hahn, William J.; Bernal, Rodrigo; Balslev, Henrik; Borchsenius, Finn (2011):**
**Phylogeny and Divergence times of Bactridinae (Arecaceae, Palmae) based on Plastid and nuclear DNA Sequences.**
In: Taxon 60 (2), p. 485–498.

**Faurby, Søren; Eiserhardt, Wolf L.; Baker, William J.; Svenning, Jens-Christian (2016):**
**An all-Evidence Species-level Supertree for the Palms (Arecaceae).**
In: Molecular Phylogenetics and Evolution 100, p. 57–69. DOI: 10.1016/j.ympev.2016.03.002.

**Hahn, William J. (2002):**
**A molecular phylogenetic Study of the Palmae (Arecaceae) based on atpB, rbcL, and 18s nrDNA Sequences.**
In: Systematic Biology 51 (1), p. 92–112.

**Hahn, William J. (2002):**
**A phylogenetic Analysis of the Arecoid Line of Palms based on Plastid DNA Sequence Data.**
In: Molecular Phylogenetics and Evolution 23 (2), p. 189–204. DOI: 10.1016/S1055-7903(02)00022-2.

**Henderson, Andrew J. (2002):**
**Phenetic and phylogenetic Analysis of *Reinhardtia* (Palmae).**
In: American Journal of Botany 89 (9), p. 1491–1502. DOI: 10.3732/ajb.89.9.1491.

**Henderson, Andrew J. (2005):**
**A multivariate Study of *Calyptrogyne* (Palmae).**
In: Systematic Botany 30 (1), p. 60–83. DOI: 10.1600/0363644053661913.

**Henderson, Andrew J. (2011):**
**A Revision of *Desmoncus* (Arecaceae).**
Auckland, N.Z.: Magnolia Press (Phytotaxa, 35).

**Henderson, Andrew J. (2011):**
**A Revision of *Geonoma* (Arecaceae).**
Auckland, N.Z.: Magnolia Press (Phytotaxa, 17).

**Henderson, Andrew J. (2012):**
**A Revision of *Pholidostachys* (Arecaceae).**
In: Phytotaxa 43, p. 1–48.

**Horn, James W.; Fisher, Jack B.; Tomlinson, P. Barry; Lewis, Carl E.; Laubengayer, Karen (2009):**

**Evolution of Lamina Anatomy in the Palm Family (Arecaceae).**
In: American Journal of Botany 96 (8), p. 1462–1486. DOI: 10.3732/ajb.0800396.

**KEIM, ARY P.R.Y.; DRANSFIELD, JOHN (2012):**
**A monograph of the Genus *Orania* (Arecaceae: Oranieae).**
In: Kew Bulletin 67, p. 127–190.

**LEWIS, CARL E.; DOYLE, J. J. (2002):**
**A phylogenetic Analysis of Tribe Areceae (Arecaceae) using two low-copy nuclear Genes.**
In: Plant Systematics and Evolution 236 (1), p. 1–17. DOI: 10.1007/s00606-002-0205-1.

**LOO, ADRIAN H. B.; DRANSFIELD, JOHN; CHASE, MARK W.; BAKER, WILLIAM J. (2006):**
**Low-copy nuclear DNA, Phylogeny and the Evolution of Dichogamy in the Betel Nut Palms and their Relatives (Arecinae; Arecaceae).**
In: Molecular Phylogenetics and Evolution 39 (3), p. 598–618. DOI: 10.1016/j.ympev.2005.12.006.

**LUDEÑA, BERTHA; CHABRILLANGE, NATHALIE; ABERLENC-BERTOSSI, FRÉDÉRIQUE; ADAM, HÉLÈNE; TREGEAR, JAMES W.; PINTAUD, JEAN-CHRISTOPHE (2011):**
**Phylogenetic utility of the nuclear Genes Agamous 1 and Phytochrome B in Palms (Arecaceae): an Example within Bactridinae.**
In: Annals of Botany 108 (8), p. 1433–1444. DOI: 10.1093/aob/mcr191.

**NOBLICK, LARRY R.; HAHN, WILLIAM J.; GRIFFITH, M. PATRICK (2012):**
**Structural cladistic Study of Cocoseae, Subtribe Attaleinae (Arecaceae): Evaluating taxonomic Limits in Attaleinae and the Neotropical Genus *Syagrus*.**
In: Brittonia 65 (2), p. 232–261.

**NORUP, MARIA VIBE; DRANSFIELD, JOHN; CHASE, MARK W.; BARFOD, ANDERS S.; FERNANDO, EDWINO S.; BAKER, WILLIAM J. (2006):**
**Homoplasious Character Combinations and generic Delimitation : a case Study from the Indo-Pacific Arecoid Palms (Arecaceae: Areceae).**
In: American Journal of Botany 93 (7), p. 1065–1080. DOI: 10.3732/ajb.93.7.1065.

**RIVERA, DIEGO; OBÓN, CONCEPCIÓN; GARCÍA-ARTEAGA, JOAQUÍN; EGEA, TERESA; ALCARAZ, FRANCISCO; LAGUNA, EMILIO ET AL. (2014):**
**Carpological Analysis of *Phoenix* (Arecaceae): contributions to the Taxonomy and evolutionary History of the Genus.**
In: Botanical Journal of the Linnean Society 175 (1), p. 74–122. DOI: 10.1111/boj.12164.

**RONCAL, JULISSA; FRANCISCO-ORTEGA, JAVIER; ASMUSSEN-LANGE, CONNY B.; LEWIS, CARL E. (2005):**
**Molecular Phylogenetics of Tribe Geonomeae (Arecaceae) using Nuclear DNA Sequences of Phosphoribulokinase and RNA Polymerase II.**
In: Systematic Botany 30 (2), p. 275–283. DOI: 10.1600/0363644054223620.

**SANÍN, MARÍA JOSÉ; GALEANO, GLORIA (2011):**
**A Revision of the Andean Wax Palms, *Ceroxylon* (Arecaceae).**
Auckland, N.Z.: Magnolia Press (Phytotaxa, 34).

**SUNDERLAND, TERRY C. H. (2012):**
**A taxonomic Revision of the Rattans of Africa ('Arecaceae' : 'Calamoideae').**
Auckland: Magnolia Press (Phytotaxa, 51).

**THOMAS, MEREDITH M.; GARWOOD, NANCY C.; BAKER, WILLIAM J.; HENDERSON, SALLY A.; RUSSELL, STEPHEN J.; HODEL, DONALD R.; BATEMAN, RICHARD M. (2006):**

**Molecular Phylogeny of the palm Genus *Chamaedorea*, based on the low-copy nuclear Genes Prk and Rpb2.**
In: Molecular Phylogenetics and Evolution 38 (2), p. 398–415. DOI: 10.1016/j.ympev.2005.08.019.

Trénel, Philipp; Gustafsson, Mats H.G.; Baker, William J.; Asmussen-Lange, Conny B.; Dransfield, John; Borchsenius, Finn (2007):
**Mid-Tertiary Dispersal, not Gondwanan Vicariance explains Distribution Patterns in the Wax Palm Subfamily (Ceroxyloideae: Arecaceae).**
In: Molecular Phylogenetics and Evolution 45 (1), p. 272–288. DOI: 10.1016/j.ympev.2007.03.018.

Zona, Scott (2005):
**A Revision of *Ptychococcus* (Arecaceae).**
In: Systematic Botany 30 (3), p. 520–529.

Zona, Scott; Francisco-Ortega, Javier; Jestrow, Brett; Baker, William J.; Lewis, Carl E. (2011):
**Molecular Phylogenetics of the Palm Subtribe Ptychospermatinae (Arecaceae).**
In: American Journal of Botany 98 (10), p. 1716–1726. DOI: 10.3732/ajb.1100218.

### Aristolochiaceae

Berjano, Regina; Roa, Fernando; Talavera, Salvador; Guerra, Marcelo (2009):
**CytoTaxonomy of diploid and polyploid Aristolochia (Aristolochiaceae) Species based on the Distribution of Cma/dapi bands and 5s and 45s rDNA sites.**
In: Plant Systematics and Evolution 280 (3-4), p. 219–227. DOI: 10.1007/s00606-009-0184-6.

Buchwalder, Katja; Samain, Marie-Stéphanie; Sankowsky, Garry; Neinhuis, Christoph; Wanke, Stefan (2014):
**Nomenclatural updates of *Aristolochia* Subgenus *Pararistolochia* (Aristolochiaceae).**
In: Australian Systematic Botany 27 (1), p. 48–55. DOI: 10.1071/SB13042.

Neinhuis, C.; Wanke, Stefan; Hilu, Khidir W.; Müller, Kai F.; Borsch, Thomas (2005):
**Phylogeny of Aristolochiaceae based on Parsimony, Likelihood, and Bayesian Analyses of trnL-trnF Sequences.**
In: Plant Systematics and Evolution 250 (1-2), p. 7–26. DOI: 10.1007/s00606-004-0217-0.

Ohi-Toma, Tetsuo; Sugawara, Takashi; Murata, Hiroko; Wanke, Stefan; Neinhuis, Christoph; Murata, Jin (2006):
**Molecular Phylogeny of *Aristolochia* sensu lato (Aristolochiaceae) based on Sequences of rbcL, matK, and phyA Genes, with Special Reference to Differentiation of Chromosome Numbers.**
In: Systematic Botany 31 (3), p. 481–492.

Wanke, Stefan; González, Favio; Neinhuis, Christoph (2006):
**Systematics of Pipevines: Combining morphological and Fast-Evolving molecular Characters to Investigate the Relationships within Subfamily Aristolochioideae (Aristolochiaceae).**
In: International Journal of Plant Sciences 167 (6), p. 1215–1227.

### Arthropteridaceae

Liu, Hong-Mei; Jiang, Ri-Hong; Guo, Jian; Hovenkamp, Peter H.; Perrie, Leon R.; Shepherd, Lara D. et al. (2013):
**Towards a phylogenetic Classification of the climbing Fern Genus *Arthropteris*.**
In: Taxon 62 (4), p. 688–700. DOI: 10.12705/624.26.

### Asaraceae

**Kelly, Lawrence M. (1998):**
**Phylogenetic Relationships in *Asarum* (Aristolochiaceae) based on Morphology and ITS Sequences.**
In: American Journal of Botany 85 (10), p. 1454–1467. DOI: 10.2307/2446402.

**Sinn, Brandon T.; Kelly, Lawrence M.; Freudenstein, John V. (2015):**
**Phylogenetic Relationships in Asarum: Effect of Data Partitioning and a revised Classification.**
In: American Journal of Botany 102 (5), p. 765–779. DOI: 10.3732/ajb.1400316.

**Sinn, Brandon T.; Kelly, Lawrence M.; Freudenstein, John V. (2015):**
**Putative Floral brood-site mimicry, loss of autonomous selfing, and reduced vegetative Growth are significantly correlated with increased Diversification in *Asarum* (Aristolochiaceae).**
In: Molecular Phylogenetics and Evolution 89, p. 194–204. DOI: 10.1016/j.ympev.2015.04.019.

**Sugawara, Takashi; Fijii, Noriyuki; Senni, Kei; Murata, Jin (2005):**
**Morphological and karyological Characteristics and phylogenetic Relationship of *Asarum cordifolium* C.E.Fisch. (Aristolochiaceae) occurring in Myanmar.**
In: Acta Phytotaxonomica Geobotanica 56 (3), p. 247–255.

**Yamaji, Hiroki; Fukuda, Tatsuya; Yokoyama, Jun U.N.; Pak, Jae-Hong; Zhou, Chang-Zheng; Yang, Chun-shu et al. (2007):**
**Reticulate Evolution and Phylogeography in *Asarum* Sect. *Asiasarum* (Aristolochiaceae) documented in internal transcribed Spacer Sequences (ITS) of nuclear ribosomal DNA.**
In: Molecular Phylogenetics and Evolution 44 (2), p. 863–884. DOI: 10.1016/j.ympev.2007.01.011.

### Asparagaceae

**Fukuda, Tatsuya; Ashizawa, Hiroki; Suzuki, Ryoko; Ochiai, Toshinori; Nakamura, Toru; Kanno, Akira et al. (2005):**
**Molecular Phylogeny of the Genus *Asparagus* (Asparagaceae) inferred from Plastid petB Intron and petD-rpoA intergenic Spacer Sequences.**
In: Plant Species Biology 20 (2), p. 121–132. DOI: 10.1111/j.1442-1984.2005.00131.x.

**Gándara, Etelvina; Specht, Chelsea D.; Sosa, Victoria (2014):**
**Origin and Diversification of the *Milla* Clade (Brodiaeoideae, Asparagaceae): a Neotropical Group of six geophytic Genera.**
In: Molecular Phylogenetics and Evolution 75, p. 118–125. DOI: 10.1016/j.ympev.2014.02.014.

**Kubota, Shosei; Konno, Itaru; Kanno, Akira (2012):**
**Molecular Phylogeny of the Genus *Asparagus* (Asparagaceae) explains interspecific Crossability between the Garden *Asparagus* (*A. officinalis*) and other *Asparagus* Species.**
In: Theoretical and applied genetics 124 (2), p. 345–354. DOI: 10.1007/s00122-011-1709-2.

**Norup, Maria F.; Petersen, Gitte; Burrows, Sandie; Bouchenak-Khelladi, Yanis; Leebens-Mack, Jim H.; Pires, J. Chris et al. (2015):**
**Evolution of *Asparagus* L. (Asparagaceae): Out-of-South-Africa and multiple Origins of sexual Dimorphism.**
In: Molecular Phylogenetics and Evolution 92, p. 25–44. DOI: 10.1016/j.ympev.2015.06.002.

### Asphodelaceae

**Adams, Stuart P.; Leitch, Ilia J.; Bennett, Michael D.; Chase, Mark W.; Leitch, Andrew R. (2000):**
**Ribosomal DNA Evolution and Phylogeny in *Aloe* (Asphodelaceae).**
In: American Journal of Botany 87 (11), p. 1578–1583. DOI: 10.2307/2656733.

**Chase, Mark W. (2000):**
**Phylogenetics of Asphodelaceae (Asparagales): An Analysis of Plastid rbcL and trnL-F DNA Sequences.**
In: Annals of Botany 86 (5), p. 935–951. DOI: 10.1006/anbo.2000.1262.

**Daru, Barnabas H.; Manning, John C.; Boatwright, James S.; Maurin, Olivier; Maclean, Norman; Schaefer, Hanno et al. (2013):**
**Molecular and morphological Analysis of Subfamily Alooideae (Asphodelaceae) and the inclusion of *Chortolirion* in *Aloe*.**
In: Taxon 62 (1), p. 62–76.

**Grace, Olwen M.; Klopper, Ronell R.; Smith, Gideon F.; Crouch, Neil R.; Figueiredo, Estrela; Rønsted, Nina; van Wyk, Abraham E. (2013):**
**A revised generic Classification for *Aloe* (Xanthorrhoeaceae Subfam. Asphodeloideae).**
In: Phytotaxa 76 (1), p. 7. DOI: 10.11646/phytotaxa.76.1.2.

**Klopper, Ronell R.; van Wyk, Abraham E.; Smith, Gideon F. (2010):**
**Phylogenetic Relationships in the Family Asphodelaceae (Asparagales).**
In: Biodiversity and Ecology 6.

**Manning, John C.; Boatwright, James S.; Daru, Barnabas H.; Maurin, Olivier; van der Bank, Michelle (2014):**
**A molecular Phylogeny and Generic Classification of Asphodelaceae Subfamily Alooideae: A Final Resolution of the Prickly Issue of Polyphyly in the Alooids?**
In: Systematic Botany 39 (1), p. 55–74. DOI: 10.1600/036364414X678044.

**Naderi Safar, Kosar; Kazempour-Osaloo, Shahrokh; Assadi, Mostafa; Zarrei, Mehdi; Khoshsokhan Mozaffar, Maryam (2014):**
**Phylogenetic Analysis of *Eremurus, Asphodelus,* and *Asphodeline* (Xanthorrhoeaceae-Asphodeloideae) inferred from Plastid trnL-F and nrDNA ITS Sequences.**
In: Biochemical Systematics and Ecology 56, p. 32–39. DOI: 10.1016/j.bse.2014.04.015.

**Ramdhani, Syd; Barker, Nigel P.; Baijnath, Himansu (2009):**
**Rampant Non-Monophyly of Species in *Kniphofia* Moench (Asphodelaceae) suggests a recent Afromontane Radiation.**
In: Taxon 58 (4), p. 1141–1152. DOI: 10.1002/tax.584008.

**Ramdhani, Syd; Barker, Nigel P.; Cowling, Richard M. (2011):**
**Revisiting Monophyly in *Haworthia* Duval (Asphodelaceae): Incongruence, Hybridization and contemporary speciation.**
In: Taxon 60 (4), p. 1001–1014.

**Treutlein, Jens; Smith, Gideon F.; Wyk, B.-E.; Wink, Michael (2003):**
**Evidence for the Polyphyly of *Haworthia* (Asphodelaceae Subfamily Alooideae; Asparagales) inferred from Nucleotide Sequences of rbcL, matK, ITS1 and Genomic Fingerprinting with LSSR-PCR.**
In: Plant Biology 5 (5), p. 513–521. DOI: 10.1055/s-2003-44793.

### Aspleniaceae

**Heede, Caroline J. van den; Viane, Ronald L. L.; Chase, Mark W. (2003):**

**Phylogenetic Analysis of *Asplenium* Subgenus *Ceterach* (Pteridophyta: Aspleniaceae) based on Plastid and Nuclear Ribosomal ITS DNA Sequences.**
In: American Journal of Botany 90 (3), p. 481–495.

**Ohlsen, Daniel J.; Perrie, Leon R.; Shepherd, Lara D.; Brownsey, Patrick J.; Bayly, Michael J. (2014):**
**Investigation of Species boundaries and Relationships in the *Asplenium paleaceum* complex (Aspleniaceae) using AFLP fingerprinting and Chloroplast and nuclear DNA Sequences.**
In: Australian Systematic Botany 27 (6), p. 378–394. DOI: 10.1071/SB14024.

**Ohlsen, Daniel J.; Perrie, Leon R.; Shepherd, Lara D.; Brownsey, Patrick J.; Bayly, Michael J. (2014):**
**Phylogeny of the Fern Family Aspleniaceae in Australasia and the south-western Pacific.**
In: Australian Systematic Botany 27 (6), p. 355–371. DOI: 10.1071/SB14043.

**Perrie, Leon R.; Brownsey, Patrick J. (2005):**
**Insights into the Biogeography and Polyploid Evolution of New Zealand *Asplenium* from Chloroplast DNA Sequence Data.**
In: American Fern Journal 95 (1), p. 1–21. DOI: 10.1640/0002-8444(2005)095[0001:IITBAP]2.0.CO;2.

**Schneider, Harald; Russell, Steve J.; Cox, Cymon J.; Bakker, Freek T.; Henderson, Sally A.; Rumsey, Fred et al. (2004):**
**Chloroplast Phylogeny of Asplenioid Ferns based on rbcL and trnL-F Spacer Sequences (Polypodiidae, Aspleniaceae) and its Implications for Biogeography.**
In: Systematic Botany 29 (2), p. 260–274. DOI: 10.1600/036364404774195476.

**Schulze, G.; Treutlein, Jens; Wink, Michael (2001):**
**Phylogenetic Relationships between *Asplenium bourgaei* (Boiss.) Milde and *A. jahandiezii* (Litard.) Rouy inferred from morphological Characters and rbcL Sequences.**
In: Plant Biology 3, p. 364–371.

**Shepherd, Lara D.; Holland, Barbara R.; Perrie, Leon R. (2008):**
**Conflict amongst Chloroplast DNA Sequences obscures the Phylogeny of a Group of *Asplenium* Ferns.**
In: Molecular Phylogenetics and Evolution 48 (1), p. 176–187. DOI: 10.1016/j.ympev.2008.02.023.

**Yatabe, Yoko; Masuyama, Shigeo; Darnaedi, Dedy; Murakami, Noriaki (2001):**
**Molecular Systematics of the *Asplenium nidus* Complex from Mt. Halimun National Park, Indonesia: Evidence for reproductive Isolation among three sympatric Rbcl Sequence Types.**
In: American Journal of Botany 88 (8), p. 1517–1522.

### Asteliaceae

**Birch, Joanne L. (2015):**
**A Revision of infrageneric Classification in *Astelia* Banks & Sol. ex R.Br. (Asteliaceae).**
In: PhytoKeys (52), p. 105–132. DOI: 10.3897/phytokeys.52.4768.

**Birch, Joanne L.; Keeley, Sterling C.; Morden, Clifford W. (2012):**
**Molecular Phylogeny and dating of Asteliaceae (Asparagales): *Astelia* s.l. Evolution provides Insight into the Oligocene History of New Zealand.**
In: Molecular Phylogenetics and Evolution 65 (1), p. 102–115. DOI: 10.1016/j.ympev.2012.05.031.

### Asteraceae

**Altinordu, Fahim; Martin, Esra; Hamzaoğlu, Ergin; Çetin, Özlem (2014):**

**New Chromosome counts, karyotype Analyses and asymmetry Indices in some Taxa of Genus *Senecio* L. and related Genera *Tephroseris* (Rchb.) Rchb. and *Turanecio* Hamzaoğlu belong to Tribe Senecioneae (Asteraceae) from Turkey.**

In: Plant Systematics and Evolution 300 (10), p. 2205–2216. DOI: 10.1007/s00606-014-1042-8.

**Anderberg, Arne A. (2005):**

**Evolutionary Relationships in the Asteraceae Tribe Inuleae (incl. Plucheeae) Evidenced by DNA Sequences of ndhF; with Notes on the systematic Positions of some aberrant Genera.**

In: Organisms Diversity and Evolution 5 (2), p. 135–146. DOI: 10.1016/j.ode.2004.10.015.

**Andrés-Sánchez, Santiago; Temsch, Eva M.; Rico, Enrique; Montserrat Martínez-Ortega, Maria (2013):**

**Genome Size in *Filago* L. (Asteraceae, Gnaphalieae) and related Genera: phylogenetic, evolutionary and ecological Implications.**

In: Plant Systematics and Evolution 299 (2), p. 331–345. DOI: 10.1007/s00606-012-0724-3.

**Archibald, Jenny K.; Crawford, Daniel J.; Santos-Guerra, Arnoldo; Mort, Mark E. (2006):**

**The utility of automated Analysis of inter-simple Sequence Repeat (Issr) Loci for resolving Relationships in the Canary Island Species of *Tolpis* (Asteraceae).**

In: American Journal of Botany 93 (8), p. 1154–1162. DOI: 10.3732/ajb.93.8.1154.

**Bain, J. F.; Golden, J. L. (2000):**

**A Phylogeny of *Packera* (Senecioneae; asteraceae) based on Internal Transcribed Spacer Region Sequence Data and a broad Sampling of Outgroups.**

In: Molecular Phylogenetics and Evolution 16 (3), p. 331–338. DOI: 10.1006/mpev.2000.0804.

**Baird, Kristen E.; Funk, Vicki A.; Wen, Jun; Weeks, Andrea (2010):**

**Molecular phylogenetic Analysis of *Leibnitzia* Cass. (Asteraceae: Mutisieae: *Gerbera*-complex), an Asian-North American disjunct Genus.**

In: Journal of Systematics and Evolution 48 (3), p. 161–174. DOI: 10.1111/j.1759-6831.2010.00077.x.

**Baldwin, Bruce G.; Wessa, Bridget L. (2000):**

**Origin and Relationships of the tarweed-silversword Lineage (Compositae-Madiinae).**

In: American Journal of Botany 87 (12), p. 1890–1908. DOI: 10.2307/2656841.

**Barker, Nigel P.; Howis, Seranne; Nordenstam, Bertil; Källersjö, Mari; Eldenäs, Pia K.; Griffioen, C.; Linder, Hans Peter (2009):**

**Nuclear and Chloroplast DNA-based Phylogenies of *Chrysanthemoides* Tourn. ex Medik. (Calenduleae; Asteraceae) reveal extensive incongruence and generic Paraphyly, but support the Recognition of infraspecific Taxa in C. monilifera.**

In: South African Journal of Botany 75 (3), p. 560–572. DOI: 10.1016/j.sajb.2009.05.006.

**Barres, Laia; Sanmartín, Isabel; Anderson, Cajsa Lisa; Susanna, Alfonso; Buerki, Sven; Galbany-Casals, Mercè; Vilatersana, Roser (2013):**

**Reconstructing the Evolution and biogeographic History of Tribe Cardueae (Compositae).**

In: American Journal of Botany 100 (5), p. 867–882. DOI: 10.3732/ajb.1200058.

**Bayer, Randall J.; Cross, E. W. (2003):**

**A reassessment of tribal Affinities of *Cratystylis* and *Haegiela* (Asteraceae) based on three Chloroplast DNA Sequences.**

In: Plant Systematics and Evolution 236 (3-4), p. 207–220. DOI: 10.1007/s00606-002-0233-x.

**Bayer, Randall J.; Puttock, Christopher F.; Kelchner, Scot A. (2000):**
**Phylogeny of South African Gnaphalieae (Asteraceae) based on two noncoding Chloroplast Sequences.**
In: American Journal of Botany 87 (2), p. 259–272. DOI: 10.2307/2656914.

**Bean, A. R. (2013):**
**Reinstatement and Revision of *Sphaeromorphaea* DC. and *Ethuliopsis* F.Muell. (Asteraceae: Plucheinae).**
In: Austrobaileya 9 (1), p. 30–59.

**Beck, James B.; Nesom, Guy L.; Calie, Patrick J.; Baird, Gary I.; Small, Randall L.; Schilling, Edward E. (2004):**
**Is Subtribe Solidagininae (Asteraceae) monophyletic?**
In: Taxon 53 (3), p. 691–698.

**Bengtson, Annika; Anderberg, Arne A.; Karis, Per Ola (2011):**
**Phylogeny and Generic Delimitation of the *Metalasia* Clade (Asteraceae-Gnaphalieae).**
In: International Journal of Plant Sciences 172 (8), p. 1067–1075. DOI: 10.1086/661294.

**Bengtson, Annika; Anderberg, Arne A.; Karis, Per Ola (2014):**
**Phylogeny and Evolution of the South African Genus *Metalasia* (Asteraceae-Gnaphalieae) inferred from molecular and morphological Data.**
In: Botanical Journal of the Linnean Society 174 (2), p. 173–198. DOI: 10.1111/boj.12114.

**Bentley, Joanne; Klaassen, Esmerialda S.; Bergh, Nicola G. (2015):**
***Philyrophyllum* (Asteraceae) transferred from Gnaphalieae to Athroismeae based on phylogenetic Analysis of nuclear and Plastid DNA Sequence Data.**
In: Taxon 64 (5), p. 975–986. DOI: 10.12705/645.7.

**Bergh, Nicola G.; Linder, Hans Peter (2009):**
**Cape Diversification and repeated out-of-southern-Africa Dispersal in paper daisies (Asteraceae-Gnaphalieae).**
In: Molecular Phylogenetics and Evolution 51 (1), p. 5–18. DOI: 10.1016/j.ympev.2008.09.001.

**Bergh, Nicola G.; Trisos, Christopher H.; Verboom, George Anthony (2011):**
**Phylogeny of the "*Ifloga* Clade" (Asteraceae, Gnaphalieae), a Lineage occurring disjointly in the Northern and Southern Hemisphere, and inclusion of *Trichogyne* in Synonymy with *Ifloga*.**
In: Taxon 60 (4), p. 1065–1075.

**Bergh, Nicola G.; Verboom, George Anthony (2011):**
**Anomalous Capitulum Structure and Monoecy may confer Flexibility in sex Allocation and life History Evolution in the *Ifloga* Lineage of Paper Daisies (Compositae: Gnaphalieae).**
In: American Journal of Botany 98 (7), p. 1113–1127. DOI: 10.3732/ajb.1000457.

**Blöch, Cordula; Dickoré, W. Bernhard; Samuel, L.; Stuessy, Tod F. (2010):**
**Molecular Phylogeny of the Edelweiss (*Leontopodium*, Asteraceae – Gnaphalieae).**
In: Edinburgh Journal of Botany 67 (02), p. 235–264. DOI: 10.1017/S0960428610000065.

**Blöch, Cordula; Weiss-Schneeweiss, Hanna; Schneeweiss, Gerald M.; Barfuss, Michael H. J.; Rebernig, Carolin A.; Villaseñor, José Luis; Stuessy, Tod F. (2009):**

**Molecular phylogenetic Analyses of nuclear and Plastid DNA Sequences support dysploid and polyploid Chromosome Number Changes and reticulate Evolution in the Diversification of *Melampodium* (Millerieae, Asteraceae).**
In: Molecular Phylogenetics and Evolution 53 (1), p. 220–233. DOI: 10.1016/j.ympev.2009.02.021.

**BONIFACINO, JOSÉ MAURICIO; FUNK, VICKI A. (2012):**
**Phylogenetics of the *Chiliotrichum* Group (Compositae: Astereae): the Story of the fascinating Radiation in the paleate Astereae Genera from Southern South America.**
In: Taxon 61 (1), p. 180–196.

**BORŠIĆ, IGOR; SUSANNA, ALFONSO; BANCHEVA, SVETLANA; GARCIA-JACAS, NÚRIA (2011):**
***Centaurea* Sect. *Cyanus* : Nuclear Phylogeny, Biogeography, and Life-Form Evolution.**
In: International Journal of Plant Sciences 172 (2), p. 238–249. DOI: 10.1086/657645.

**BOWLES, VICTORIA G.; MAYERHOFER, REINHOLD; DAVIS, COREY; GOOD, ALLEN G.; HALL, JOCELYN C. (2010):**
**A phylogenetic Investigation of Carthamus combining Sequence and microsatellite Data.**
In: Plant Systematics and Evolution 287 (1-2), p. 85–97. DOI: 10.1007/s00606-010-0292-3.

**BREITWIESER, ILSE; GLENNY, DAVID S.; THORNE, ANITA; WAGSTAFF, STEVEN J. (1999):**
**Phylogenetic Relationships in Australasian Gnaphalieae (Compositae) inferred from ITS Sequences.**
In: New Zealand Journal of Botany 37 (3), p. 399–412. DOI: 10.1080/0028825X.1999.9512644.

**BROUILLET, LUC; ANDERBERG, ARNE A.; NESOM, GUY L.; LOWREY, TIMOTHY K.; URBATSCH, LOWELL E. (2009):**
***Welwitschiella* is a member of the African Subtribe Grangeinae (Asteraceae Astereae): a new phylogenetic Position based on ndh and ITS Sequence Data.**
In: Kew Bulletin 64, p. 645–660.

**CALVO, JOEL; ÁLVAREZ, INÉS; AEDO, CARLOS; PELSER, PIETER B. (2013):**
**A phylogenetic Analysis and new Delimitation of *Senecio* Sect. *Crociseris* (Compositae: Senecioneae), with Evidence of intergeneric Hybridization.**
In: Taxon 62 (1), p. 127–140.

**CARIAGA, KATHLEEN A.; PRUSKI, JOHN F.; OVIEDO, RAMONA; ANDERBERG, ARNE A.; LEWIS, CARL E.; FRANCISCO-ORTEGA, JAVIER (2008):**
**Phylogeny and Systematic Position of *Feddea* (Asteraceae: Feddeeae): a Taxonomically enigmatic and critically endangered Genus endemic to Cuba.**
In: Systematic Botany 33 (1), p. 193–202. DOI: 10.1600/036364408783887348.

**CHAN, RAYMUND; BALDWIN, BRUCE G.; ORNDUFF, ROBERT (2001):**
**Goldfields revisited: a molecular phylogenetic Perspective on the Evolution of *Lasthenia* (Compositae: Heliantheae sensu lato).**
In: International Journal of Plant Sciences 162 (6), p. 1347–1360.

**CLEVINGER, JENNIFER A.; PANERO, JOSE L. (2000):**
**Phylogenetic Analysis of *Silphium* and Subtribe Engelmanniinae (Asteraceae: Heliantheae) based on ITS and ETS Sequence Data.**
In: American Journal of Botany 87 (4), p. 565–572. DOI: 10.2307/2656600.

**COLEMAN, MAX; LISTON, AARON; KADEREIT, JOACHIM W.; ABBOTT, RICHARD J. (2003):**
**Repeat intercontinental Dispersal and Pleistocene Speciation in disjunct Mediterranean and desert *Senecio* (Asteraceae).**
In: American Journal of Botany 90 (10), p. 1446–1454. DOI: 10.3732/ajb.90.10.1446.

**Costa, Clayton M. (2014):**
**A molecular Phylogeny of the Golden Asters, Subtribe Chrysopsidinae (Asteraceae: Astereae), based on nuclear ribosomal and Chloroplast Sequence Data.**
Master of Science in Biology. Towson University, Towson, Maryland, USA.

**Crawford, D.J.; Kimball, R. T.; Tadesse, Mesfin (2001):**
**The generic Placement of a morphologically enigmatic Species in Asteraceae: Evidence from ITS Sequences.**
In: Plant Systematics and Evolution 228, p. 63–69.

**Crawford, Daniel J.; Mort, Mark E. (2005):**
**Phylogeny of Eastern North American *Coreopsis* (Asteraceae-Coreopsideae): Insights from nuclear and Plastid Sequences, and comments on Character Evolution.**
In: American Journal of Botany 92 (2), p. 330–336. DOI: 10.3732/ajb.92.2.330.

**Cron, Glynis V.; Balkwill, Kevin; Knox, Eric B. (2008):**
**Phylogenetic Evidence for the generic Circumscription of *Cineraria* L. (Asteraceae-Senecioneae).**
In: Taxon 57 (3), p. 779–798.

**Cross, E. W.; Quinn, Christopher J.; Wagstaff, Steven J. (2002):**
**Molecular Evidence for the Polyphyly of *Olearia* (Astereae: Asteraceae).**
In: Plant Systematics and Evolution 235 (1), p. 99–120. DOI: 10.1007/s00606-002-0198-9.

**Cruz-Mazo, G.; Buide, M. L.; Samuel, L.; Narbona, E. (2009):**
**Molecular Phylogeny of *Scorzoneroides* (Asteraceae): Evolution of heterocarpy and annual Habit in unpredictable Environments.**
In: Molecular Phylogenetics and Evolution 53 (3), p. 835–847. DOI: 10.1016/j.ympev.2009.08.001.

**Denda, T.; Watanabe, K.; Kosuge, Keiko; Yahara, T.; Ito, Motomi (1999):**
**Molecular Phylogeny of *Brachyscome* (Asteraceae).**
In: Plant Systematics and Evolution 217 (217), p. 299–311.

**Devos, Nicolas; Barker, Nigel P.; Nordenstam, Bertil; Mucina, Ladislav (2010):**
**A multi-locus Phylogeny of *Euryops* (Asteraceae, Senecioneae) augments Support for the “Cape to Cairo” Hypothesis of Floral Migrations in Africa.**
In: Taxon 59 (1), p. 57–67.

**Diazgranados, Mauricio (2012):**
**A Nomenclator for the Frailejones (Espeletiinae Cuatrec., Asteraceae).**
In: PhytoKeys (16), p. 1–52. DOI: 10.3897/phytokeys.16.3186.

**Ekenäs, Catarina; Baldwin, Bruce G.; Andreasen, Katarina (2007):**
**A molecular phylogenetic Study of *Arnica* (Asteraceae): Low Chloroplast DNA Variation and problematic subgeneric Classification.**
In: Systematic Botany 32 (4), p. 917–928.

**Ekenäs, Catarina; Heidari, Nahid; Andreasen, Katarina (2012):**
***Arnica* (Asteraceae) Phylogeny revisited using RPB2: complex Patterns and multiple d-Paralogues.**
In: Molecular Phylogenetics and Evolution 64 (2), p. 261–270. DOI: 10.1016/j.ympev.2012.02.032.

**Eldenäs, Pia K.; Källersjö, Mari; Anderberg, Arne A. (1999):**
**Phylogenetic Placement and Circumscription of Tribes Inuleae s. str. and Plucheeae (Asteraceae): Evidence from Sequences of Chloroplast Gene ndhF.**

In: Molecular Phylogenetics and Evolution 13 (1), p. 50–58.

**Eldens, Pia; Anderberg, Arne A.; Kllersj, Mari (1998):**
**Molecular Phylogenetics of the TribeInuleae s. str. (Asteraceae), based on ITS Sequences of nuclear ribosomal DNA.**
In: Plant Systematics and Evolution 210 (3-4), p. 159–173. DOI: 10.1007/BF00985666.

**Englund, Markus; Pornpongrungrueng, Pimwadee; Gustafsson, Mats H.G.; Anderberg, Arne A. (2009):**
**Phylogenetic Relationships and generic Delimitation in Inuleae Subtribe Inulinae (Asteraceae) based on ITS and cpDNA Sequence Data.**
In: Cladistics 25 (4), p. 319–352. DOI: 10.1111/j.1096-0031.2009.00256.x.

**Enke, Neela; Gemeinholzer, Birgit (2008):**
**Babcock revisited: new Insights into generic Delimitation and Character Evolution in *Crepis* L. (Compositae: Cichorieae) from ITS and matK Sequence Data.**
In: Taxon 57 (3), p. 756–768.

**Enke, Neela; Gemeinholzer, Birgit; Zidorn, Christian (2012):**
**Molecular and phytochemical Systematics of the Subtribe Hypochaeridinae (Asteraceae, Cichorieae).**
In: Organisms Diversity and Evolution 12 (1), p. 1–16. DOI: 10.1007/s13127-011-0064-0.

**Esselman, Elizabeth J.; Crawford, Daniel J.; Brauner, Soren; Stuessy, Tod F.; Anderson, Gregory J.; Silva O, Mario (2000):**
**Rapd marker Diversity within and Divergence among Species of *Dendroseris* (Asteraceae: Lactuceae).**
In: American Journal of Botany 87 (4), p. 591–596. DOI: 10.2307/2656603.

**Fehlberg, Shannon D.; Ranker, Tom A. (2007):**
**Phylogeny and Biogeography of *Encelia* (Asteraceae) in the Sonoran and Peninsular Deserts based on multiple DNA Sequences.**
In: Systematic Botany 32 (3), p. 692–699. DOI: 10.1600/036364407782250689.

**Fehrer, Judith; Gemeinholzer, Birgit; Chrtek, Jindrich; Bräutigam, Siegfried (2007):**
**Incongruent Plastid and nuclear DNA Phylogenies reveal ancient intergeneric Hybridization in *Pilosella* hawkweeds (*Hieracium*, Cichorieae, Asteraceae).**
In: Molecular Phylogenetics and Evolution 42 (2), p. 347–361. DOI: 10.1016/j.ympev.2006.07.004.

**Fernández, I. A.; Aguilar, J. F.; Panero, Jose L.; Feliner, G. N. (2001):**
**A phylogenetic Analysis of *Doronicum* (Asteraceae, Senecioneae) based on morphological, nuclear ribosomal (ITS), and Chloroplast (trnL-F) Evidence.**
In: Molecular Phylogenetics and Evolution 20 (1), p. 41–64. DOI: 10.1006/mpev.2001.0954.

**Ferreira, Maria Zita; Zahradníček, Jaroslav; Kadlecová, Jana; Sequeira, Miguel Menezes; Chrtek Jr., Jindřich; Fehrer, Judith (2015):**
**Tracing the evolutionary History of the little-known Mediterranean-Macaronesian Genus *Andryala* (Asteraceae) by multigene Sequencing.**
In: Taxon 64 (3), p. 535–551. DOI: 10.12705/643.10.

**Flagel, Lex E.; Rapp, Ryan A.; Grover, Corrinne E.; Widrlechner, Mark P.; Hawkins, Jennifer; Grafenberg, Jessie L. et al. (2008):**
**Phylogenetic, morphological, and chemotaxonomic Incongruence in the North American endemic Genus *Echinacea*.**
In: American Journal of Botany 95 (6), p. 756–765. DOI: 10.3732/ajb.0800049.

**Flann, Christian; Greuter, Werner; Hind, D. NicholasJ. (2010):**
**Cassini's Compositae Genera: A nomenclatural and taxonomic Assessment.**
In: Taxon 59 (4), p. 1206–1244.

**Font, Mònica; Garcia-Jacas, Núria; Vilatersana, Roser; Roquet, Cristina; Susanna, Alfonso (2009):**
**Evolution and Biogeography of *Centaurea* Section *Acrocentron* inferred from nuclear and Plastid DNA Sequence Analyses.**
In: Annals of Botany 103 (6), p. 985–997. DOI: 10.1093/aob/mcp022.

**Ford, Kerry A.; Ward, Josephine M.; Smissen, Rob D.O.B.; Wagstaff, Steven J.; Breitwieser, Ilse (2007):**
**Phylogeny and Biogeography of *Craspedia* (Asteraceae: Gnaphalieae) based on ITS, ETS and psbA-trnH Sequence Data.**
In: Taxon 56 (3), p. 783–794.

**Francisco-Ortega, Javier; Barber, Janet C.; Santos-Guerra, Arnoldo; Febles-Hernández, Rosa; Jansen, Robert K. (2001):**
**Origin and Evolution of the endemic Genera of Gonosperminae (Asteraceae: Anthemideae) from the Canary Islands: Evidence from nucleotide Sequences of the internal transcribed Spacers of the nuclear ribosomal DNA.**
In: American Journal of Botany 88 (1), p. 161–169. DOI: 10.2307/2657136.

**Freire, Susana Edith (2007):**
**Systematic Revision and Phylogeny of *Ainsliaea* DC. (Asteraceae, Mutisieae) 1.**
In: Annals of the Missouri Botanical Garden 94 (1), p. 79–191. DOI: 10.3417/0026-6493(2007)94[79:SRAPOA]2.0.CO;2.

**Freire, Susana Edith; Chemisquy, Maria Amelia; Anderberg, Arne A.; Beck, Stephan G.; Meneses, Rosa I.; Loeuille, Benoît; Urtubey, Estrella (2015):**
**The *Lucilia* Group (Asteraceae, Gnaphalieae): phylogenetic and taxonomic Considerations based on molecular and morphological Evidence.**
In: Plant Systematics and Evolution 301 (4), p. 1227–1248. DOI: 10.1007/s00606-014-1147-0.

**Funk, Vicki A.; Chan, Raymund (2008):**
**Phylogeny of the Spiny African Daisies (Compositae, Tribe Arctotideae, Subtribe Gorteriinae) based on trnL-F, ndhF, and ITS Sequence Data.**
In: Molecular Phylogenetics and Evolution 48 (1), p. 47–60. DOI: 10.1016/j.ympev.2008.03.035.

**Funk, Vicki A.; Chan, Raymund; Keeley, Sterling C. (2004):**
**Insights into the Evolution of the Tribe Arctoteae (Compositae: Subfamily Cichorioideae s.s.) using trnL-F, ndhF, and ITS.**
In: Taxon 53 (3), p. 637–655.

**Funk, Vicki A.; Kelloff, Carol L.; Chan, Raymund (2012):**
**Phylogeny and Biogeography of the Tribe Liabeae (Compositae Subfamily Cichorioideae).**
In: Taxon 61 (2), p. 437–455.

**Funk, Vicki A.; Koekemoer, Marinda (2011):**
**A Monograph of the small Tribe Platycarpheae (Compositae: Cichorioideae).**
In: Systematic Botany 36 (1), p. 191–208. DOI: 10.1600/036364411X553270.

**Funk, Vicki A.; Sancho, Gisela; Roque, Nádia; Kelloff, Carol L.; Ventosa-Rodríguez, Iralys; Diazgranados, Mauricio et al. (2014):**
**A Phylogeny of the Gochnatieae: Understanding a critically placed Tribe in the Compositae.**

In: Taxon 63 (4), p. 859–882. DOI: 10.12705/634.27.

**FUNSTON, A. MICHELE (2008):**
**Taxonomic Revision of *Roldana* (Asteraceae: Senecioneae), a Genus of the Southwestern U.S.A., Mexico, and Central America.**
In: Annals of the Missouri Botanical Garden 95 (2), p. 282–337. DOI: 10.3417/2003151.

**GALBANY-CASALS, MERCÈ; ANDRÉS-SÁNCHEZ, SANTIAGO; GARCIA-JACAS, NÚRIA; SUSANNA, ALFONSO; RICO, ENRIQUE; MARTÍNEZ-ORTEGA, M. MONTSERRAT (2010):**
**How many of Cassini anagrams should there be? molecular Systematics and phylogenetic Relationships in the *Filago* Group (Asteraceae, Gnaphalieae), with special Focus on the Genus *Filago*.**
In: Taxon 59 (6), p. 1671–1689.

**GALBANY-CASALS, MERCÈ; GARCIA-JACAS, NÚRIA; SÁEZ, LLORENÇ; BENEDÍ, CARLES; SUSANNA, ALFONSO (2009):**
**Phylogeny, Biogeography, and Character Evolution in Mediterranean, Asiatic, and Macaronesian *Helichrysum* (Asteraceae, Gnaphalieae) inferred from Nuclear phylogenetic Analyses.**
In: International Journal of Plant Sciences 170 (3), p. 365–380. DOI: 10.1086/596332.

**GALBANY-CASALS, MERCÈ; UNWIN, MATTHEW M.; GARCIA-JACAS, NÚRIA; SMISSEN, ROB D.O.B.; SUSANNA, ALFONSO; BAYER, RANDALL J. (2014):**
**Phylogenetic Relationships in *Helichrysum* (Compositae: Gnaphalieae) and related Genera: Incongruence between nuclear and Plastid Phylogenies, biogeographic and morphological Patterns, and Implications for generic Delimitation.**
In: Taxon 63 (3), p. 608–624. DOI: 10.12705/633.8.

**GAO, TIAN-GANG; WANG, WEI; BAYER, RANDALL J.; LI, DE-ZHU (2009):**
**Systematic Position of the enigmatic Genus *Sheareria* (Asteraceae) - Evidence from molecular, morphological and cytological Data.**
In: Taxon 58 (3), p. 769–780.

**GARCIA, S.; GARNATJE, TERESA; HIDALGO, ORIANE; MCARTHUR, E. DURANT; SILJAK-YAKOVLEV, SONJA; VALLÈS, JOAN (2007):**
**Extensive ribosomal DNA (18s-5.8s-26s and 5s) Colocalization in the North American endemic sagebrushes (Subgenus *Tridentatae*, *Artemisia*, Asteraceae) revealed by Fish.**
In: Plant Systematics and Evolution 267 (1-4), p. 79–92. DOI: 10.1007/s00606-007-0558-6.

**GARCIA, SÒNIA; MCARTHUR, E. DURANT; PELLICER, JAUME; SANDERSON, STEWART C.; VALLÈS, JOAN; GARNATJE, TERESA (2011):**
**A molecular phylogenetic approach to western North America endemic *Artemisia* and allies (Asteraceae): untangling the sagebrushes.**
In: American Journal of Botany 98 (4), p. 638–653. DOI: 10.3732/ajb.1000386.

**GARCIA-JACAS, NÚRIA (2001):**
**Generic Delimitation and Phylogeny of the Subtribe Centaureinae (Asteraceae): A Combined Nuclear and Chloroplast DNA Analysis.**
In: Annals of Botany 87 (4), p. 503–515. DOI: 10.1006/anbo.2000.1364.

**GARCIA-JACAS, NÚRIA; GARNATJE, TERESA; SUSANNA, ALFONSO; VILATERSANA, ROSER (2002):**
**Tribal and subtribal Delimitation and Phylogeny of the Cardueae (Asteraceae): a combined nuclear and Chloroplast DNA Analysis.**
In: Molecular Phylogenetics and Evolution 22 (1), p. 51–64. DOI: 10.1006/mpev.2001.1038.

**Garcia-Jacas, Núria; Susanna, Alfonso; Mozaffarian, V.; Ilarslan, R. (2000):**

**The natural Delimitation of *Centaurea* (Asteraceae: Cardueae): ITS Sequence Analysis of the *Centaurea jacea* Group.**

In: Plant Systematics and Evolution 223, p. 185–199.

**Garcia-Jacas, Núria; Uysal, Tuna; Romaschenko, Konstantin; Suárez-Santiago, Víctor N.; Ertuğrul, Kuddisi; Susanna, Alfonso (2006):**

***Centaurea* revisited: A molecular survey of the *Jacea* Group.**

In: Annals of Botany 98 (4), p. 741–753. DOI: 10.1093/aob/mcl157.

**Gaskin, John F.; Wilson, L. M. (2007):**

**Phylogenetic Relationships among native and naturalized *Hieracium* (Asteraceae) in Canada and the United States based on Plastid DNA Sequences.**

In: Systematic Botany 32 (2), p. 478–485. DOI: 10.1600/036364407781179752.

**Geleta, Mulatu; Bekele, Endashaw; Dagne, Kifle; Bryngelsson, Tomas (2010):**

**Phylogenetics and taxonomic Delimitation of the Genus *Guizotia* (Asteraceae) based on Sequences derived from various Chloroplast DNA Regions.**

In: Plant Systematics and Evolution 289 (1-2), p. 77–89. DOI: 10.1007/s00606-010-0334-x.

**Geleta, Mulatu; Bryngelsson, Tomas; Bekele, Endashaw; Dagne, Kifle (2007):**

**Comparative Analysis of genetic Relationship and diagnostic Markers of several Taxa of *Guizotia* Cass. (Asteraceae) as revealed by AFLPs and RAPDs.**

In: Plant Systematics and Evolution 265 (3-4), p. 221–239. DOI: 10.1007/s00606-007-0521-6.

**Goertzen, Leslie R.; Cannone, Jamie J.; Gutell, Robin R.; Jansen, Robert K. (2003):**

**ITS secondary Structure derived from comparative Analysis: Implications for Sequence Alignment and Phylogeny of the Asteraceae.**

In: Molecular Phylogenetics and Evolution 29 (2), p. 216–234. DOI: 10.1016/S1055-7903(03)00094-0.

**Greiner, Roland; Vogt, Robert; Oberprieler, Christoph (2012):**

**Phylogenetic studies in the polyploid complex of the Genus *Leucanthemum* Mill. (Compositae, Anthemideae) based on cpDNA Sequence Variation.**

In: Plant Systematics and Evolution 298 (7), p. 1407–1414. DOI: 10.1007/s00606-012-0636-2.

**Grossi, Mariana A.; Gutiérrez, Diego G.; Berrueta, Pedro C.; Martínez, Juan J. (2011):**

***Acanthostyles* (Asteraceae, Eupatorieae): a Revision with a multivariate Analysis.**

In: Australian Systematic Botany 24 (2), p. 87. DOI: 10.1071/SB10038.

**Gruenstaeudl, Michael; Santos-Guerra, Arnoldo; Jansen, Robert K. (2013):**

**Phylogenetic Analyses of *Tolpis* Adans. (Asteraceae) reveal Patterns of adaptive Radiation, multiple Colonization and interspecific Hybridization.**

In: Cladistics 29 (4), p. 416–434. DOI: 10.1111/cla.12005.

**Gruenstaeudl, Michael; Urtubey, Estrella; Jansen, Robert K.; Samuel, Rosabelle; Barfuss, Michael H. J.; Stuessy, Tod F. (2009):**

**Phylogeny of Barnadesioideae (Asteraceae) inferred from DNA Sequence Data and Morphology.**

In: Molecular Phylogenetics and Evolution 51 (3), p. 572–587. DOI: 10.1016/j.ympev.2009.01.023.

**Gültepe, Mutlu; Coşkunçelebi, Kamil; Makbul, Serdar; Sağlam, Coşkun (2015):**

***Tragopogon turcicus* sp. nov. (Asteraceae) from Turkey and its phylogenetic Position.**

In: Nordic Journal of Botany 33 (5), p. 540–547. DOI: 10.1111/njb.00851.

**Guo, Yan-Ping; Ehrendorfer, Friedrich; Samuel, Rosabelle (2004):**

**Phylogeny and Systematics of *Achillea* (Asteraceae-Anthemideae) inferred from nrITS and Plastid trnL-F DNA Sequences.**
In: Taxon 53 (3), p. 657–672.

HEIDEN, GUSTAVO (2013):
**Two new Combinations in *Baccharis* (Asteraceae: Asterae).**
In: Phytoneuron 78, p. 1–2.

HEIDEN, GUSTAVO; PIRANI, JOSÉ RUBENS (2012):
**A Synopsis and Notes for *Baccharis* subgen. *Tarchonanthoides* (Asteraceae: Astereae).**
In: Phytotaxa 60, p. 41–49.

HERSHKOVITZ, MARK A.; ARROYO, MARY T.K.; BELL, CHARLES D.; HINOJOSA, L. FELIPE (2006):
**Phylogeny of *Chaetanthera* (Asteraceae: Mutisieae) reveals both ancient and recent Origins of the high Elevation Lineages.**
In: Molecular Phylogenetics and Evolution 41 (3), p. 594–605. DOI: 10.1016/j.ympev.2006.05.003.

HIDALGO, ORIANE; GARCIA-JACAS, NÚRIA; GARNATJE, TERESA; ROMASCHENKO, KONSTANTIN; SUSANNA, ALFONSO; SILJAK-YAKOVLEV, SONJA (2008):
**Extreme environmental conditions and phylogenetic Inheritance: Systematics of *Myopordon* and *Oligochaeta* (Asteraceae, Cardueae-Centaureinae).**
In: Taxon 57 (3), p. 769–778.

HIDALGO, ORIANE; GARCIA-JACAS, NÚRIA; GARNATJE, TERESA; SUSANNA, ALFONSO (2006):
**Phylogeny of *Rhaponticum* (Asteraceae, Cardueae-Centaureinae) and related Genera inferred from nuclear and Chloroplast DNA Sequence Data: taxonomic and biogeographic Implications.**
In: Annals of Botany 97 (5), p. 705–714. DOI: 10.1093/aob/mcl029.

HILPOLD, ANDREAS; SCHÖNSWETTER, PETER; SUSANNA, ALFONSO; GARCIA-JACAS, NÚRIA; VILATERSANA, ROSER (2011):
**Evolution of the central Mediterranean *Centaurea cineraria* Group (Asteraceae): Evidence for relatively recent, allopatric Diversification following transoceanic Seed Dispersal.**
In: Taxon 60 (2), p. 528–538.

HILPOLD, ANDREAS; VILATERSANA, ROSER; SUSANNA, ALFONSO; MESEGUER, ANDREA SÁNCHEZ; BORŠIĆ, IGOR; CONSTANTINIDIS, THEOPHANIS ET AL. (2014):
**Phylogeny of the *Centaurea* Group (*Centaurea*, Compositae) - Geography is a better Predictor than Morphology.**
In: Molecular Phylogenetics and Evolution 77, p. 195–215. DOI: 10.1016/j.ympev.2014.04.022.

HIMMELREICH, SVEN; BREITWIESER, ILSE; OBERPRIELER, CHRISTOPH (2014):
**Phylogenetic Relationships in the extreme polyploid complex of the New Zealand Genus *Leptinella* (Compositae: Anthemideae) based on AFLP Data.**
In: Taxon 63 (4), p. 883–898. DOI: 10.12705/634.19.

HIMMELREICH, SVEN; BREITWIESER, ILSE; OBERPRIELER, CHRISTOPH (2012):
**Phylogeny, Biogeography, and Evolution of sex Expression in the Southern Hemisphere Genus *Leptinella* (Compositae, Anthemideae).**
In: Molecular Phylogenetics and Evolution 65 (2), p. 464–481. DOI: 10.1016/j.ympev.2012.07.001.

HIMMELREICH, SVEN; KÄLLERSJÖ, MARI; ELDENÄS, PIA K.; OBERPRIELER, CHRISTOPH (2008):

**Phylogeny of Southern Hemisphere Compositae-Anthemideae based on nrDNA ITS and cpDNA ndhF Sequence Information.**
In: Plant Systematics and Evolution 272 (1-4), p. 131–153. DOI: 10.1007/s00606-007-0634-y.

**Howis, Seranne; Barker, Nigel P.; Mucina, Ladislav (2009):**
**Globally grown, but poorly known: Species limits and Biogeography of *Gazania* Gaertn. (Asteraceae) inferred from Chloroplast and nuclear DNA Sequence Data.**
In: Taxon 58 (3), p. 871–882.

**Ito, Motomi; Yahara, Tetsukazu; King, Robert M.; Watanabe, Kuniaki; Oshita, Sanae; Yokoyama, Jun U.N.; Crawford, Daniel J. (2000):**
**Molecular Phylogeny of Eupatorieae (Asteraceae) Estimated from cpDNA RFLP and its Implication for the Polyploid Origin Hypothesis of the Tribe.**
In: Journal of Plant Research 113, p. 91–96.

**Jafari, Farzaneh; Osaloo, Shahrokh Kazempour; Mozaffarian, Valiollah (2015):**
**Molecular Phylogeny of the Tribe Astereae (Asteraceae ) in SW Asia based on nrDNA ITS and cpDNA psbA-trnH Sequences.**
In: Willdenowia 45 (1), p. 77–92. DOI: 10.3372/wi.45.45108.

**Jansen, Robert K.; Holsinger, Kent E.; Michaels, Helen J.; Palmer, Jeffrey D. (1990):**
**Phylogenetic Analysis of Chloroplast DNA Restriction site Data at higher taxonomic Levels: an Exanple from the Asteraceae.**
In: Evolution 44 (8), p. 2089–2105.

**Karaman-Castro, Vesna; Urbatsch, Lowell E. (2009):**
**Phylogeny of *Hinterhubera* Group and related Genera (Hinterhuberinae: Astereae) based on the nrDNA ITS and ETS Sequences.**
In: Systematic Botany 34 (4), p. 805–817. DOI: 10.1600/036364409790139772.

**Karis, Per Ola (2006):**
**Morphological Data indicates two major Clades of the Subtribe Gorteriinae (Asteraceae-Arctotideae).**
In: Cladistics 22, p. 199–221.

**Katinas, Liliana; Crisci, Jorge V.; Jabaily, Rachel Schmidt; Williams, Cody; Walker, Jay B.; Drew, Bryan T. et al. (2008):**
**Evolution of secondary heads in Nassauviinae (Asteraceae, Mutisieae).**
In: American Journal of Botany 95 (2), p. 229–240. DOI: 10.3732/ajb.95.2.229.

**Katinas, Liliana; Sancho, Gisela; Vitali, Maira Soledad (2013):**
**A Revision of *Lophopappus* (Asteraceae, Nassauvieae).**
In: Phytotaxa 103 (1), p. 25. DOI: 10.11646/phytotaxa.103.1.2.

**Keeley, Sterling C.; Forsman, Zac H.; Chan, Raymund (2007):**
**A Phylogeny of the "Evil Tribe" (Vernonieae: Compositae) reveals Old/New World long distance Dispersal: support from separate and combined congruent Datasets (trnL-F, ndhF, ITS).**
In: Molecular Phylogenetics and Evolution 44 (1), p. 89–103. DOI: 10.1016/j.ympev.2006.12.024.

**Kim, H.-G.; Funk, Vicki A.; Vlasak, A.; Zimmer, Elizabeth A. (2003):**
**A Phylogeny of the Munnoziinae (Asteraceae, Liabeae): Circumscription of *Munnozia* and a new Placement of *M. perfoliata*.**
In: Plant Systematics and Evolution 239 (3-4), p. 171–185. DOI: 10.1007/s00606-003-0003-4.

**KIM, SEUNG-CHUL; CHUNGHEE, LEE; MEJÍAS, JOSÉ A. (2007):**

**Phylogenetic Analysis of Chloroplast DNA matK Gene and ITS of nrDNA Sequences reveals Polyphyly of the Genus *Sonchus* and new Relationships among the Subtribe Sonchinae (Asteraceae: Cichorieae).**

In: Molecular Phylogenetics and Evolution 44 (2), p. 578–597. DOI: 10.1016/j.ympev.2007.03.014.

**KIM, SEUNG-CHUL; CRAWFORD, DANIEL J.; JANSEN, ROBERT K.; SANTOS-GUERRA, ARNOLDO (1999):**

**The use of a non-coding Region of Chloroplast DNA in phylogenetic Studies of the Subtribe Sonchinae (Asteraceae: Lactuceae).**

In: Plant Systematics and Evolution 215, p. 85–99.

**KIM, SEUNG-CHUL; LU, CHRISTINA T.; LEPSCHI, BRENDAN J. (2004):**

**Phylogenetic Positions of *Actites megalocarpa* and *Sonchus hydrophilus* (Sonchinae: Asteraceae) based on ITS and Chloroplast non-coding DNA Sequences.**

In: Australian Systematic Botany 17 (1), p. 73. DOI: 10.1071/SB03019.

**KIMBALL, REBECCA T.; CRAWFORD, DANIEL J. (2004):**

**Phylogeny of Coreopsideae (Asteraceae) using ITS Sequences suggests Lability in reproductive Characters.**

In: Molecular Phylogenetics and Evolution 33 (1), p. 127–139. DOI: 10.1016/j.ympev.2004.04.022.

**KIMBALL, REBECCA T.; CRAWFORD, DANIEL J.; SMITH, EDWIN B. (2003):**

**Evolutionary Processes in the Genus *Coreocarpus*: Insights from molecular Phylogenetics.**

In: Evolution 57 (1), p. 52–61.

**KIRSCHNER, JAN; ZÁVESKÁ DRÁBKOVÁ, LENKA; ŠTĚPÁNEK, JAN; UHLEMANN, INGO (2015):**

**Towards a better Understanding of the *Taraxacum* Evolution (Compositae–Cichorieae) on the basis of nrDNA of sexually reproducing Species.**

In: Plant Systematics and Evolution 301 (4), p. 1135–1156. DOI: 10.1007/s00606-014-1139-0.

**KITA, YOKO; FUJIKAWA, KAZUMI; ITO, MOTOMI; OHBA, HIDEAKI; KATO, MASAHIRO (2004):**

**Molecular phylogenetic Analyses and Systematics of the Genus *Saussurea* and related Genera (Asteraceae, Cardueae).**

In: Taxon 53 (3), p. 679–690.

**KOEKEMOER, MARINDA (2016):**

**A monograph of *Stoebe* and *Seriphium* (Gnaphalieae, Asteraceae) in southern Africa and neighbouring Islands.**

In: Phytotaxa 242 (1), p. 1. DOI: 10.11646/phytotaxa.242.1.1.

**KONISHI, N.; WATANABE, K.; KOSUGE, KEIKO (2000):**

**Molecular Systematics of Australian *Podolepis* (Asteraceae: Gnaphalieae): Evidence from DNA Sequences of the nuclear ITS Region and the Chloroplast matK Gene.**

In: Australian Systematic Botany 13 (5), p. 709. DOI: 10.1071/SB99030.

**KONOWALIK, KAMIL; WAGNER, FLORIAN; TOMASELLO, SALVATORE; VOGT, ROBERT; OBERPRIELER, CHRISTOPH (2015):**

**Detecting reticulate Relationships among diploid *Leucanthemum* Mill. (Compositae, Anthemideae) Taxa using multilocus Species Tree Reconstruction Methods and AFLP Fingerprinting.**

In: Molecular Phylogenetics and Evolution 92, p. 308–328. DOI: 10.1016/j.ympev.2015.06.003.

**KOOPMAN, WIM J. M.; GUETTA, ELI; VAN DE WIEL, CLEMENS C. M.; VOSMAN, BEN; VAN DEN BERG, RONALD G. (1998):**

**Phylogenetic Relationships among *Lactuca* (Asteraceae) Species and related Genera based on ITS-1 DNA Sequences.**
In: American Journal of Botany 85 (11), p. 1517–1530. DOI: 10.2307/2446479.

**Koopman, Wim J. M.; Zevenbergen, Martin J.; van den Berg, Ronald G. (2001):**
**Species Relationships in *Lactuca* s.l. (Lactuceae, Asteraceae) inferred from AFLP Fingerprints.**
In: American Journal of Botany 88 (10), p. 1881–1887. DOI: 10.2307/3558364.

**Kornkven, Amy B.; Watson, Linda E.; Estes, James R. (1998):**
**Phylogenetic Analysis of *Artemisia* Section *Tridentatae* (Asteraceae) based on Sequences from the internal transcribed Spacers (ITS) of nuclear ribosomal DNA.**
In: American Journal of Botany 85 (12), p. 1787–1795. DOI: 10.2307/2446513.

**Lee, Chunghee; Kim, Seung-Chul; Lundy, Karen; Santos-Guerra, Arnoldo (2005):**
**Chloroplast DNA Phylogeny of the woody *Sonchus* alliance (Asteraceae: Sonchinae) in the Macaronesian Islands.**
In: American Journal of Botany 92 (12), p. 2072–2085. DOI: 10.3732/ajb.92.12.2072.

**Lee, Joongku; Baldwin, Bruce G.; Gottlieb, L. D. (2002):**
**Phylogeny of *Stephanomeria* and related Genera (Compositae-Lactuceae) based on Analysis of 18s-26s nuclear rDNA ITS and ETS Sequences.**
In: American Journal of Botany 89 (1), p. 160–168. DOI: 10.3732/ajb.89.1.160.

**Lee, Joongku; Baldwin, Bruce G.; Gottlieb, L. D. (2003):**
**Phylogenetic Relationships among the Primarily North American Genera of Cichorieae (Compositae) based on Analysis of 18s–26s Nuclear rDNA ITS and ETS Sequences.**
In: Systematic Botany 28 (3), p. 616–626.

**Li, Wei-Ping; Qian, Feng-Ming; Yang, Xiu-Lin; Chen, San-Mao (2014):**
**Systematic Position of *Cyathocline* Cass. (Asteraceae): Evidences from molecular, cytological and morphological Data.**
In: Plant Systematics and Evolution 300 (4), p. 595–606. DOI: 10.1007/s00606-013-0905-8.

**Li, Wei-Ping; Yang, Fu-Sheng; Jivkova, Todorka; Yin, Gen-Shen (2012):**
**Phylogenetic Relationships and generic Delimitation of Eurasian *Aster* (Asteraceae: Astereae) inferred from ITS, ETS and trnL-F Sequence Data.**
In: Annals of Botany 109 (7), p. 1341–1357. DOI: 10.1093/aob/mcs054.

**Liu, Hong; Trusty, Jennifer; Oviedo, Ramona; Anderberg, Arne A.; Francisco-Ortega, Javier (2004):**
**Molecular Phylogenetics of the Caribbean Genera *Rhodogeron* and *Sachsia* (Asteraceae).**
In: International Journal of Plant Sciences 165 (1), p. 209–217.

**Liu, Jian-Quan; Gao, Tian-Gang; Chen, Zhi-Duan; Lu, An-Ming (2002):**
**Molecular Phylogeny and Biogeography of the Qinghai-Tibet Plateau endemic *Nannoglottis* (Asteraceae).**
In: Molecular Phylogenetics and Evolution 23, p. 307–325.

**Liu, Jian-Quan; Wang, Yu-Jing; Wang, Ai-Lan; Hideaki, Ohba; Abbott, Richard J. (2006):**
**Radiation and Diversification within the *Ligularia-Cremanthodium-Parasenecio* complex (Asteraceae) triggered by Uplift of the Qinghai-Tibetan Plateau.**
In: Molecular Phylogenetics and Evolution 38 (1), p. 31–49. DOI: 10.1016/j.ympev.2005.09.010.

**Liu, Ying; Chen, You-Sheng; Yang, Qin-Er (2013):**
**Generic status, Circumscription, and allopolyploid Origin of *Faberia* (Asteraceae: Cichorieae) as revealed by ITS and Chloroplast DNA Sequence Data.**
In: Taxon 62 (6), p. 1235–1247. DOI: 10.12705/626.14.

**Liu, Ying; Yang, Qin-Er (2011):**
**Cytology and its systematic Implications in *Sinosenecio* (Senecioneae-Asteraceae) and two closely related Genera.**
In: Plant Systematics and Evolution 291 (1-2), p. 7–24. DOI: 10.1007/s00606-010-0365-3.

**Lo Presti, Rosa Maria; Oppolzer, Stephanie; Oberprieler, Christoph (2012):**
**A molecular Phylogeny and a revised Classification of the Mediterranean Genus *Anthemis* s.l. (Compositae, Anthemideae) based on three molecular Markers and micromorphological characters.**
In: Taxon 59 (5), p. 1441–1456.

**Loeuille, Benoît Francis Patrice (2011):**
**Towards a phylogenetic Classification of Lychnophorinae (Asteraceae: Vernonieae).**
Universidade de São Paulo, São Paulo.

**Loeuille, Benoît; Semir, João; Lohmann, Lúcia Garces; Pirani, José Rubens (2015):**
**A phylogenetic Analysis of Lychnophorinae (Asteraceae: Vernonieae) based on molecular and morphological Data.**
In: Systematic Botany 40 (1), p. 299–315. DOI: 10.1600/036364415X686585.

**Loockerman, Dennis J.; Turner, Billie L.; Jansen, Robert K. (2003):**
**Phylogenetic Relationships within the Tageteae (Asteraceae) based on Nuclear Ribosomal ITS and Chloroplast ndhF Gene Sequences.**
In: Systematic Botany 28 (1), p. 191–2007.

**López-Vinyallonga, Sara; López-Pujol, Jordi; Constantinidis, Theophanis; Susanna, Alfonso; Garcia-Jacas, Núria (2015):**
**Mountains and Refuges: Genetic Structure and evolutionary History in closely related, endemic *Centaurea* in continental Greece.**
In: Molecular Phylogenetics and Evolution 92, p. 243–254. DOI: 10.1016/j.ympev.2015.06.018.

**López-Vinyallonga, Sara; Mehregan, Iraj; Garcia-Jacas, Núria; Tscherneva, Olga; Susanna, Alfonso; Kadereit, Joachim W. (2009):**
**Phylogeny and Evolution of the *Arctium-Cousinia* Complex (Compositae, Cardueae-Carduinae).**
In: Taxon 58 (1), p. 153–171.

**López-Vinyallonga, Sara; Romaschenko, Konstantin; Susanna, Alfonso; Garcia-Jacas, Núria (2011):**
**Systematics of the Arctioid Group: Disentangling *Arctium* and *Cousinia* (Cardueae, Carduinae).**
In: Taxon 60 (2), p. 539–554.

**Lowrey, Timothy K.; Quinn, Christopher J.; Taylor, Rachael K.; Chan, Raymund; Kimball, Rebecca T.; Nardi, Jan C. (2001):**
**Molecular and morphological Reassessment of Relationships within the *Vittadinia* Group of Astereae (Asteraceae).**
In: American Journal of Botany 88 (7), p. 1279–1289. DOI: 10.2307/3558339.

**Lowrey, Timothy K.; Whitkus, Richard; Sykes, William R. (2005):**

**A new Species of *Tetramolopium* (Asteraceae) from Mitiaro, Cook Islands: Biogeography, phylogenetic Relationships, and Dispersal.**
In: Systematic Botany 30 (2), p. 448–455.

**LUEBERT, FEDERICO; WEN, JUN; DILLON, MICHAEL O. (2009):**
**Systematic Placement and biogeographical Relationships of the monotypic Genera *Gypothamnium* and *Oxyphyllum* (Asteraceae: Mutisioideae) from the Atacama Desert.**
In: Botanical Journal of the Linnean Society 159, p. 32–51.

**LYU, MING-JU AMY; GOWIK, UDO; KELLY, STEVE; COVSHOFF, SARAH; MALLMANN, JULIA; WESTHOFF, PETER ET AL. (2015):**
**RNA-Seq based Phylogeny recapitulates previous Phylogeny of the Genus *Flaveria* (Asteraceae) with some modifications.**
In: BMC Evolutionary Biology 15, p. 116. DOI: 10.1186/s12862-015-0399-9.

**MAGEE, ANTHONY RICHARD; NICOLAS, ANTOINE N.; TILNEY, PATRICIA M.; PLUNKETT, GREGORY M. (2015):**
**Phylogenetic Relationships and generic Realignments in the early Diverging Subtribe Pentziinae (Asteraceae, Anthemideae).**
In: Botanical Journal of the Linnean Society 178 (4), p. 633–647. DOI: 10.1111/boj.12295.

**MAKBUL, SERDAR; COŞKUNÇELEBI, KAMIL; GÜLTEPE, MUTLU; OKUR, SEDA; GÜZEL, MURAT ERDEM (2012):**
***Scorzonera ahmet-duranii* sp. nov. (Asteraceae) from southwest Anatolia, and its phylogenetic Position.**
In: Nordic Journal of Botany 30 (1), p. 2–11. DOI: 10.1111/j.1756-1051.2011.01235.x.

**MALINSKA, HANA; TATE, JENNIFER A.; MATYASEK, ROMAN; LEITCH, ANDREW R.; SOLTIS, DOUGLAS E.; SOLTIS, PAMELA S.; KOVARIK, ALES (2010):**
**Similar Patterns of rDNA Evolution in synthetic and recently formed natural Populations of *Tragopogon* (Asteraceae) Allotetraploids.**
In: BMC Evolutionary Biology 10, p. 291. DOI: 10.1186/1471-2148-10-291.

**MANDEL, JENNIFER R.; DIKOW, REBECCA B.; FUNK, VICKI A. (2015):**
**Using phylogenomics to resolve Mega-families: An example from Compositae.**
In: Journal of Systematics and Evolution 53 (5), p. 391–402. DOI: 10.1111/jse.12167.

**MARANER, FABRIZIO; SAMUEL, ROSABELLE; STUESSY, TOD F.; CRAWFORD, DANIEL J.; CRISCI, JORGE V.; PANDEY, ARUN KUMAR; MORT, MARK E. (2012):**
**Molecular Phylogeny of *Nassauvia* (Asteraceae, Mutisieae) based on nrDNA ITS Sequences.**
In: Plant Systematics and Evolution 298 (2), p. 399–408. DOI: 10.1007/s00606-011-0553-9.

**MARKOS, STACI; BALDWIN, BRUCE G. (2002):**
**Structure, molecular Evolution, and phylogenetic Utility of the 50 Region of the external transcribed spacer of 18s-26s rDNA in *Lessingia* (Compositae, Astereae).**
In: Molecular Phylogenetics and Evolution 23, p. 214–228.

**MARLOWE, K.; HUFFORD, LARRY (2007):**
**Taxonomy and Biogeography of *Gaillardia* (Asteraceae): A phylogenetic Analysis.**
In: Systematic Botany 32 (1), p. 208–226. DOI: 10.1600/036364407780360229.

**MARTINS, LUDWIG.; HELLWIG, FRANK H. (2005):**
**Phylogenetic Relationships of the enigmatic Species *Serratula chinensis* and *Serratula forrestii* (Asteraceae - Cardueae).**
In: Plant Systematics and Evolution 255 (3-4), p. 215–224. DOI: 10.1007/s00606-005-0342-4.

Martins, Ludwig; Hellwig, Frank H. (2005):
**Systematic Position of the Genera *Serratula* and *Klasea* within Centaureinae (Cardueae, Asteraceae) inferred from ETS and ITS Sequence Data and new Combinations in *Klasea*.**
In: Taxon 54 (3), p. 632–638.

Mason, Chase M.; Donovan, Lisa A. (2015):
**Evolution of the Leaf economics Spectrum in Herbs: Evidence from Environmental Divergences in Leaf Physiology across *Helianthus* (Asteraceae).**
In: Evolution 69 (10), p. 2705–2720. DOI: 10.1111/evo.12768.

Mavrodiev, Evgeny V.; Edwards, Christine E.; Albach, Dirk C.; Gitzendanner, Matthew A.; Soltis, Pamela S.; Soltis, Douglas E. (2004):
**Phylogenetic Relationships in Subtribe Scorzonerinae (Asteraceae: Cichorioideae: Cichorieae) based on ITS Sequence Data.**
In: Taxon 53 (3), p. 699–712.

Mavrodiev, Evgeny V.; Soltis, Pamela S.; Soltis, Douglas E. (2008):
**Putative parentage of six Old World polyploids in *Tragopogon* L. (Asteraceae: Scorzonerinae) based on ITS, ETS, and Plastid Sequence Data.**
In: Taxon 57 (4), p. 1215–1232.

Mavrodiev, Evgeny V.; Tancig, Mark; Sherwood, Anna M.; Gitzendanner, Matthew A.; Rocca, Jennifer; Soltis, Pamela S.; Soltis, Douglas E. (2005):
**Phylogeny of *Tragopogon* L. (Asteraceae) based on internal and external transcribed spacer Sequence Data.**
In: International Journal of Plant Sciences 166 (1), p. 117–133.

McKenzie, Robert J.; Barker, Nigel P. (2008):
**Radiation of southern African daisies: biogeographic Inferences for Subtribe Arctotidinae (Asteraceae, Arctotideae).**
In: Molecular Phylogenetics and Evolution 49 (1), p. 1–16. DOI: 10.1016/j.ympev.2008.07.007.

McKenzie, Robert J.; Muller, Elizabeth M.; Skinner, Amy K. W.; Karis, Per Ola; Barker, Nigel P. (2006):
**Phylogenetic Relationships and generic Delimitation in Subtribe Arctotidinae (Asteraceae: Arctotideae) inferred by DNA Sequence Data from ITS and five Chloroplast Regions.**
In: American Journal of Botany 93 (8), p. 1222–1235. DOI: 10.3732/ajb.93.8.1222.

McKown, Athena D.; Moncalvo, Jean-Marc; Dengler, Nancy G. (2005):
**Phylogeny of *Flaveria* (Asteraceae) and Inference of C4 Photosynthesis Evolution.**
In: American Journal of Botany 92 (11), p. 1911–1928. DOI: 10.3732/ajb.92.11.1911.

Messina, Andre; Walsh, Neville G.; Hoebee, Susan E.; Green, Peter T. (2014):
**A Revision of *Olearia* Section *Asterotriche* (Asteraceae: Asterae).**
In: Australian Systematic Botany 27 (3), p. 199. DOI: 10.1071/SB14012.

Milton, Joseph J. (2009):
**Phylogenetic Analyses and taxonomic Studies of Senecioninae: Southern African *Senecio* Section *Senecio*.**
Dissertation. University of St. Andrews, UK.

Mitsui, Yuki; Chen, Shao-Tien; Zhou, Zhe-Kun; Peng, Ching-I.; Deng, Yun-Fei; Setoguchi, Hiroaki (2008):

**Phylogeny and Biogeography of the Genus *Ainsliaea* (Asteraceae) in the Sino-Japanese Region based on nuclear rDNA and Plastid DNA Sequence Data.**
In: Annals of Botany 101 (1), p. 111–124. DOI: 10.1093/aob/mcm267.

**Mitsui, Yuki; Setoguchi, Hiroaki (2012):**
**Demographic histories of adaptively diverged riparian and non-riparian Species of Ainsliaea (Asteraceae) inferred from coalescent Analyses using multiple nuclear Loci.**
In: BMC Evolutionary Biology 12, p. 254. DOI: 10.1186/1471-2148-12-254.

**Mitsui, Yuki; Setoguchi, Hiroaki (2012):**
**Recent Origin and adaptive Diversification of *Ainsliaea* (Asteraceae) in the Ryukyu Islands: molecular phylogenetic Inference using nuclear microsatellite Markers.**
In: Plant Systematics and Evolution 298 (5), p. 985–996. DOI: 10.1007/s00606-012-0608-6.

**Montes-Moreno, Noemí; Sáez, Llorenç; Benedí, Carles; Susanna, Alfonso; Garcia-Jacas, Núria (2010):**
**Generic delineation, Phylogeny and subtribal affinities of *Phagnalon* and *Aliella* (Compositae, Gnaphalieae) based on nuclear and Chloroplast Sequences.**
In: Taxon 59 (6), p. 1654–1670.

**Moody, Michael L.; Rieseberg, Loren H. (2012):**
**Sorting through the chaff, nDNA Gene Trees for phylogenetic Inference and hybrid Identification of annual Sunflowers (*Helianthus* Sect. *Helianthus*).**
In: Molecular Phylogenetics and Evolution 64 (1), p. 145–155. DOI: 10.1016/j.ympev.2012.03.012.

**Moore, Abigail J.; Bartoli, Adriana; Tortosa, Roberto D.; Baldwin, Bruce G. (2012):**
**Phylogeny, Biogeography, and Chromosome Evolution of the amphitropical Genus *Grindelia* (Asteraceae) inferred from nuclear ribosomal and Chloroplast Sequence Data.**
In: Taxon 61 (1), p. 211–230.

**Moore, Abigail J.; Bohs, Lynn (2003):**
**An ITS Phylogeny of *Balsamorhiza* and *Wyethia* (Asteraceae: Heliantheae).**
In: American Journal of Botany 90 (11), p. 1653–1660. DOI: 10.3732/ajb.90.11.1653.

**Moore, Abigail J.; Bohs, Lynn (2007):**
**Phylogeny of *Balsamorhiza* and *Wyethia* (Asteraceae: Heliantheae) using ITS, ETS, and trnK Sequence Data.**
In: Systematic Botany 32 (3), p. 682–691. DOI: 10.1600/036364407782250661.

**Moore, Michael J.; Francisco-Ortega, Javier; Santos-Guerra, Arnoldo; Jansen, Robert K. (2002):**
**Chloroplast DNA Evidence for the Roles of Island Colonization and Extinction in *Tolpis* (Asteraceae: Lactuceae).**
In: American Journal of Botany 89 (3), p. 518–526. DOI: 10.3732/ajb.89.3.518.

**Moran, Emily; Funk, Vicki A. (2006):**
**A Revision of *Erato* (Compositae: Liabeae).**
In: Systematic Botany 31 (3), p. 597–609. DOI: 10.1600/036364406778388728.

**Morgan, David R.; Korn, Randi-Lynn; Mugleston, Spencer L. (2009):**
**Insights into reticulate Evolution in Machaerantherinae (Asteraceae: Astereae): 5s ribosomal RNA Spacer Variation, estimating Support for Incongruence, and Constructing reticulate Phylogenies.**
In: American Journal of Botany 96 (5), p. 920–932. DOI: 10.3732/ajb.0800308.

**MORT, MARK E.; CRAWFORD, DANIEL J.; SANTOS-GUERRA, ARNOLDO; FRANCISCO-ORTEGA, JAVIER; ESSELMAN, ELIZABETH J.; WOLFE, ANDREA D. (2003):**

**Relationships among the Macaronesian members of *Tolpis* (Asteraceae: Lactuceae) based upon Analyses of inter simple Sequence repeat (ISSR) Markers.**

In: Taxon 52, p. 511–518.

**MORT, MARK E.; RANDLE, CHRISTOPHER P.; KIMBALL, REBECCA T.; TADESSE, MESFIN; CRAWFORD, DANIEL J. (2008):**

**Phylogeny of Coreopsideae (Asteraceae) inferred from nuclear and Plastid DNA Sequences.**

In: Taxon 57 (1), p. 109–120.

**MOURA, MÓNICA; SILVA, LUÍS; DIAS, ELISABETE F.; SCHAEFER, HANNO; CARINE, MARK A. (2015):**

**A Revision of the Genus *Leontodon* (Asteraceae) in the Azores based on morphological and molecular Evidence.**

In: Phytotaxa 210 (1), p. 24. DOI: 10.11646/phytotaxa.210.1.4.

**NIE, ZE-LONG; FUNK, VICKI A.; SUN, HANG; DENG, TAO; MENG, YING; WEN, JUN (2013):**

**Molecular Phylogeny of *Anaphalis* (Asteraceae, Gnaphalieae) with biogeographic Implications in the Northern Hemisphere.**

In: Journal of Plant Research 126 (1), p. 17–32. DOI: 10.1007/s10265-012-0506-6.

**NOMURA, NAOFUMI; TAKASO, TOKUSHIRO; PENG, CHING-I.; KONO, YOSHIKO; OGINUMA, KAZUO; MITSUI, YUKI; SETOGUCHI, HIROAKI (2010):**

**Molecular Phylogeny and habitat Diversification of the Genus *Farfugium* (Asteraceae) based on nuclear rDNA and Plastid DNA.**

In: Annals of Botany 106 (3), p. 467–482. DOI: 10.1093/aob/mcq139.

**NYLINDER, STEPHAN; ANDERBERG, ARNE A. (2015):**

**Phylogeny of the Inuleae (Asteraceae) with special Emphasis on the Inuleae-Plucheinae.**

In: Taxon 64 (1), p. 110–130. DOI: 10.12705/641.22.

**NYLINDER, STEPHAN; CRONHOLM, BODIL; LANGE, PETER J.; WALSH, NEVILLE G.; ANDERBERG, ARNE A. (2013):**

**Species Tree Phylogeny and Character Evolution in the Genus *Centipeda* (Asteraceae): Evidence from DNA Sequences from coding and non-coding loci from the Plastid and nuclear Genomes.**

In: Molecular Phylogenetics and Evolution 68 (2), p. 239–250. DOI: 10.1016/j.ympev.2013.03.020.

**OBERPRIELER, CHRISTOPH (2002):**

**A phylogenetic Analysis of *Chamaemelum* Mill. (Compositae: Anthemideae) and related Genera based upon nrDNA ITS and cpDNA trnL/trnF Igs Sequence Variation.**

In: Botanical Journal of the Linnean Society 138, p. 255–273.

**OBERPRIELER, CHRISTOPH (2005):**

**Temporal and spatial Diversification of Circum-Mediterranean Compositae-Anthemideae.**

In: Taxon 54 (4), p. 951–966.

**OBERPRIELER, CHRISTOPH; VOGT, ROBERT (2000):**

**The Position of *Castrilanthemum* Vogt & Oberprieler and the Phylogeny of Mediterranean Anthemideae (Compositae) as inferred from nrDNA ITS and cpDNA trnL/trnF Igs Sequence Variation.**

In: Plant Systematics and Evolution 225, p. 145–170.

**Oberprieler, Christoph; Vogt, Robert (2006):**

**The taxonomic Position of *Matricaria macrotis* (Compositae-Anthemideae).**

In: Willdenowia 36 (1), p. 329–338. DOI: 10.3372/wi.36.36128.

**Orchard, A. E. (2004):**

**A reassessment of the Genus *Haeckeria* (Asteraceae: Gnaphalieae), with Definition of new Species in *Cassinia*.**

In: Australian Systematic Botany 17 (5), p. 447. DOI: 10.1071/SB04021.

**Orchard, A. E. (2009):**

**A Revision of *Cassinia* (Asteraceae: Gnaphalieae) in Australia. 6. Section *Cassinia*.**

In: Australian Systematic Botany 22 (5), p. 344. DOI: 10.1071/SB09018.

**Ortiz, Santiago; Carbajal, Rodrigo; Serrano, Miguel; Rodríguez-Oubiña, Juan; Iglesias, Isabel (2013):**

**Phylogeny of the African Mutisieae s.l. (Asteraceae) based on ndhF and trnL-F Sequences (cpDNA).**

In: Taxon 62 (3), p. 525–536. DOI: 10.12705/623.11.

**Panero, Jose L.; Freire, Susana Edith; Ariza Espinar, Luis; Crozier, Bonnie S.; Barboza, Gloria Estela; Cantero, Juan J. (2014):**

**Resolution of deep Nodes yields an improved Backbone Phylogeny and a new basal Lineage to Study early Evolution of Asteraceae.**

In: Molecular Phylogenetics and Evolution 80, p. 43–53. DOI: 10.1016/j.ympev.2014.07.012.

**Panero, Jose L.; Funk, Vicki A. (2008):**

**The value of Sampling anomalous Taxa in phylogenetic Studies: major Clades of the Asteraceae revealed.**

In: Molecular Phylogenetics and Evolution 47 (2), p. 757–782. DOI: 10.1016/j.ympev.2008.02.011.

**Panero, Jose L.; Jansen, Robert K.; Clevinger, Jennifer A. (1999):**

**Phylogenetic Relationships of Subtribe Ecliptinae (Asteraceae: Heliantheae) based on Chloroplast DNA restriction site Data.**

In: American Journal of Botany 86 (3), p. 413–427. DOI: 10.2307/2656762.

**Park, Seon-Joo; Korompai, E.J.; Farncisco-Ortega, J.; Santos-Guerra, Arnoldo; Jansen, Robert K. (2001):**

**Phylogenetic Relationships of *Tolpis* (Asteraceae: Lactuceae) based on ndhF Sequence Data.**

In: Plant Systematics and Evolution 226, p. 23–33.

**Peirson, Jess A.; Reznicek, Anton A.; Semple, John C. (2012):**

**Polyploidy, infraspecific cytotype Variation, and Speciation in Goldenrods: the Cytogeography of *Solidago* subSect. *Humiles* (Asteraceae) in North America.**

In: Taxon 61 (1), p. 197–210.

**Pellicer, Jaume; Vallès, Joan; Korobkov, Aleksandr A.; Garnatje, Teresa (2011):**

**Phylogenetic Relationships of *Artemisia* Subg. *Dracunculus* (Asteraceae) based on ribosomal and Chloroplast DNA Sequences.**

In: Taxon 60 (3), p. 691–704.

**Pelser, Pieter B.; Kennedy, Aaron H.; Tepe, Eric J.; Shidler, Jacob B.; Nordenstam, Bertil; Kadereit, Joachim W.; Watson, Linda E. (2010):**

**Patterns and causes of incongruence between Plastid and nuclear Senecioneae (Asteraceae) Phylogenies.**

In: American Journal of Botany 97 (5), p. 856–873. DOI: 10.3732/ajb.0900287.

**Pelser, Pieter B.; Nordenstam, Bertil; Kadereit, Joachim W.; Watson, Linda E. (2007):**
**An ITS Phylogeny of Tribe Senecioneae (Asteraceae) and a new Delimitation of *Senecio* L.**
In: Taxon 56 (4), p. 1077–1104.

**Peng, Yu-lan; Zhang, Yu; Gao, Xin-Fen; Tong, Lin-jing; Li, Liang; Li, Ren-Yuan et al. (2014):**
**A phylogenetic Analysis and new Delimitation of *Crepidiastrum* (Asteraceae, Tribe Cichorieae).**
In: Phytotaxa 159 (4), p. 241. DOI: 10.11646/phytotaxa.159.4.1.

**Plovanich, Anne E.; Panero, Jose L. (2004):**
**A Phylogeny of the ITS and ETS for *Montanoa* (Asteraceae: Heliantheae).**
In: Molecular Phylogenetics and Evolution 31 (3), p. 815–821. DOI: 10.1016/j.ympev.2003.10.021.

**Pornpongrungrueng, Pimwadee; Borchsenius, Finn; Englund, Markus; Anderberg, Arne A.; Gustafsson, Mats H.G. (2007):**
**Phylogenetic Relationships in *Blumea* (Asteraceae: Inuleae) as Evidenced by molecular and morphological Data.**
In: Plant Systematics and Evolution 269 (3-4), p. 223–243. DOI: 10.1007/s00606-007-0581-7.

**Pornpongrungrueng, Pimwadee; Borchsenius, Finn; Gustafsson, Mats H.G. (2009):**
**Relationships within *Blumea* (Inuleae, Asteraceae) and the utility of the 5s-NTS in species-level Phylogeny Reconstruction.**
In: Taxon 58 (4), p. 1181–1193.

**Ranjbar, Massoud; Negaresh, Kazem (2013):**
**A Revision of *Centaurea* Sect. *Phaeopappus* (Asteraceae, Cardueae).**
In: Phytotaxa 123 (1), p. 1. DOI: 10.11646/phytotaxa.123.1.1.

**Rauscher, Jason T. (2002):**
**Molecular Phylogenetics of the *Espeletia* Complex (Asteraceae): Evidence from nrDNA ITS Sequences on the closest Relatives of an Andean adaptive Radiation.**
In: American Journal of Botany 89 (7), p. 1074–1084. DOI: 10.3732/ajb.89.7.1074.

**Repplinger, Miriam; Johannesen, J.; Seitz, A.; Comes, Hans Peter (2007):**
**Morphological and molecular Evidence for Hybridization and introgression in Central European *Arctium* (Asteraceae).**
In: Plant Systematics and Evolution 268 (1-4), p. 75–95. DOI: 10.1007/s00606-007-0547-9.

**Riggins, Chance W.; Seigler, David p. (2012):**
**The Genus *Artemisia* (Asteraceae: Anthemideae) at a continental Crossroads: molecular Insights into Migrations, Disjunctions, and Reticulations among Old and New World Species from a Beringian Perspective.**
In: Molecular Phylogenetics and Evolution 64 (3), p. 471–490. DOI: 10.1016/j.ympev.2012.05.003.

**Rivera, Vanessa Lopes; Panero, Jose L.; Schilling, Edward E.; Crozier, Bonnie S.; Moraes, Marta Dias (2016):**
**Origins and recent Radiation of Brazilian Eupatorieae (Asteraceae) in the eastern Cerrado and Atlantic Forest.**
In: Molecular Phylogenetics and Evolution 97, p. 90–100. DOI: 10.1016/j.ympev.2015.11.013.

**Rivero-Guerra, Aixa O.; Laurin, Michel (2012):**
**Phylogenetic Analysis of the *Santolina rosmarinifolia* Aggregate (Asteraceae: Anthemideae: Santolininae) based on morphological Characteristics.**
In: Nordic Journal of Botany 30 (5), p. 533–545. DOI: 10.1111/j.1756-1051.2011.01382.x.

**Robba, L.; Carine, Mark A.; Russell, Stephen J.; Raimondo, F. M. (2005):**
**The Monophyly and Evolution of *Cynara* L. (Asteraceae) sensu lato: Evidence from the Internal Transcribed Spacer Region of nrDNA.**
In: Plant Systematics and Evolution 253 (1-4), p. 53–64. DOI: 10.1007/s00606-004-0259-3.

**Roberts, Roland P. (2002):**
**Phylogeny of *Ericameria, Chrysothamnus* and related Genera (Asteraceae: Astereae) based on Nuclear Ribosomal DNA Sequence Data.**
Dissertation. Louisiana State University.

**Roberts, Roland P.; Urbatsch, Lowell E. (2003):**
**Molecular Phylogeny of *Ericameria* (Asteraceae, Astereae) based on nuclear ribosomal 3 ETS and ITS Sequence Data.**
In: Taxon 52, p. 209–228.

**Robinson, Harold Ernest (1999):**
**Generic and subtribal Classification of American Vernonieae.**
Washington, D.C.: Smithsonian Institution Press.

**Robinson, Harold Ernest; Funk, Vicki A. (2011):**
**A new Genus, *Nothovernonia*, from tropical Africa (Asteraceae or Compositae, Vernonieae).**
In: PhytoKeys (3), p. 21–34. DOI: 10.3897/phytokeys.3.1131.

**Robinson, Harold Ernest; Skvarla, John J.; Funk, Vicki A. (2016):**
**Vernonieae (Asteraceae) of southern Africa: A generic DisPosition of the Species and a Study of their pollen.**
In: PhytoKeys (60), p. 49–126. DOI: 10.3897/phytokeys.60.6734.

**Roque, Nádia; Afonso Santana, Fernanda (2014):**
**A new Species for a monotypic Genus: *Anteremanthus* (Asteraceae: Vernonieae).**
In: Systematic Botany 39 (2), p. 656–661. DOI: 10.1600/036364414X680771.

**Safer, Stefan; Tremetsberger, Karin; Guo, Yan-Ping; Kohl, Gudrun; Samuel, Mary R.; Stuessy, Tod F.; Stuppner, Hermann (2011):**
**Phylogenetic Relationships in the Genus *Leontopodium* (Asteraceae: Gnaphalieae) based on AFLP Data.**
In: Botanical Journal of the Linnean Society 165 (4), p. 364–377. DOI: 10.1111/j.1095-8339.2011.01117.x.

**Salomón, Luciana; Hernández, Marcelo Paulo; Giugliano, Daniel Alejandro; Freire, Susana Edith (2016):**
**Floral Microcharacters in South American Species of *Senecio* s.str. (Asteraceae) with considerations on the Circumscription of this Genus.**
In: Phytotaxa 244 (1), p. 1. DOI: 10.11646/phytotaxa.244.1.1.

**Samuel, Rosabelle; Gutermann, Walter; Stuessy, Tod F.; Ruas, Claudete de Fátima; Lack, Hans-Walter; Tremetsberger, Karin et al. (2006):**
**Molecular Phylogenetics reveals *Leontodon* (Asteraceae, Lactuceae) to be diphyletic.**
In: American Journal of Botany 93 (8), p. 1193–1205. DOI: 10.3732/ajb.93.8.1193.

**Samuel, Rosabelle; Stuessy, Tod F.; Tremetsberger, Karin; Baeza, Carlos M.; Siljak-Yakovlev, Sonja (2003):**
**Phylogenetic Relationships among Species of *Hypochaeris* (Asteraceae, Cichorieae) based on ITS, Plastid trnL Intron, trnL-F spacer, and matK Sequences.**
In: American Journal of Botany 90 (3), p. 496–507. DOI: 10.3732/ajb.90.3.496.

**Sánchez-Jiménez, Ismael; Lazkov, Georgy A.; Hidalgo, Oriane; Garnatje, Teresa (2010):**

**Molecular Systematics of *Echinops* L. (Asteraceae, Cynareae): A Phylogeny based on ITS and trnL-trnF Sequences with Emphasis on Sectional Delimitation.**
In: Taxon 59 (3), p. 698–708.

**SANCHO, GISELA; KARAMAN-CASTRO, VESNA (2008):**
**A phylogenetic Study in American Podocominae (Asteraceae: Astereae) based on morphological and molecular Data.**
In: Systematic Botany 33 (4), p. 762–775. DOI: 10.1600/036364408786500262.

**SANCHO, GISELA; KATINAS, LILIANA; PLOS, ANABELA (2014):**
**Is Morphology supporting a monophyletic *Proustia* Lag., (Nassauvieae, Asteraceae)?**
In: Plant Systematics and Evolution 300 (10), p. 2265–2276. DOI: 10.1007/s00606-014-1052-6.

**SANTIAGO, LOUIS S.; KIM, SEUNG-CHUL (2009):**
**Correlated Evolution of Leaf Shape and Physiology in the Woody *Sonchus* Alliance (Asteraceae: Sonchinae) in Macaronesia.**
In: International Journal of Plant Sciences 170 (1), p. 83–92. DOI: 10.1086/593044.

**SANZ, MARÍA; VILATERSANA, ROSER; HIDALGO, ORIANE; GARCIA-JACAS, NÚRIA; SUSANNA, ALFONSO; SCHNEEWEISS, GERALD M.; VALLÈS, JOAN (2008):**
**Molecular Phylogeny and Evolution of Floral Characters of *Artemisia* and Allies (Anthemideae, Asteraceae): Evidence from nrDNA ETS and ITS Sequences.**
In: Taxon 57 (1), p. 66–78.

**SCHAEFER, HANNO (2015):**
**On the Origin and systematic Position of the Azorean goldenrod, *Solidago azorica* (Asteraceae).**
In: Phytotaxa 210 (1), p. 47. DOI: 10.11646/phytotaxa.210.1.5.

**SCHILLING, EDWARD E. (2011):**
**Hybrid Genera in Liatrinae (Asteraceae: Eupatorieae).**
In: Molecular Phylogenetics and Evolution 59 (1), p. 158–167. DOI: 10.1016/j.ympev.2011.01.011.

**SCHILLING, EDWARD E.; PANERO, JOSE L. (2010):**
**Transfers to *Simsia* and Description of *Davilanthus*, a new Genus of Asteraceae (Heliantheae).**
In: Brittonia 62 (4), p. 309–320.

**SCHILLING, EDWARD E.; PANERO, JOSE L. (2011):**
**A revised Classification of Subtribe Helianthinae (Asteraceae: Heliantheae) II. Derived Lineages.**
In: Botanical Journal of the Linnean Society 167, p. 311–331.

**SCHILLING, EDWARD E.; PANERO, JOSE L.; COX, PATRICIA B. (1999):**
**Chloroplast DNA restriction site Data support a narrowed interpretation of *Eupatorium* (Asteraceae).**
In: Plant Systematics and Evolution 219, p. 209–223.

**SCHILLING, EDWARD E.; PANERO, JOSE L.; CROZIER, BONNIE S.; SCOTT, RANDALL W.; DÁVILA, PATRICIA (2015):**
**Bricklebush (*Brickellia*) Phylogeny reveals Dimensions of the great Asteraceae Radiation in Mexico.**
In: Molecular Phylogenetics and Evolution 85, p. 161–170. DOI: 10.1016/j.ympev.2015.02.007.

**SCHMIDT, GREGORY J.; SCHILLING, EDWARD E. (2000):**

**Phylogeny and Biogeography of *Eupatorium* (Asteraceae: Eupatorieae) based on nuclear ITS Sequence Data.**
In: American Journal of Botany 87 (5), p. 716–726. DOI: 10.2307/2656858.

**Schmidt-Lebuhn, Alexander N.; Bruhl, Jeremy J.; Telford, Ian R.H.; Wilson, Paul G. (2015):**
**Phylogenetic Relationships of *Coronidium, Xerochrysum* and several neglected Australian Species of "*Helichrysum*" (Asteraceae: Gnaphalieae).**
In: Taxon 64 (1), p. 96–109. DOI: 10.12705/641.5.

**Schmidt-Lebuhn, Alexander N.; Constable, Lee (2013):**
**Phylogenetic Relationships of the Australasian shrubby Everlastings *Ozothamnus* and *Cassinia* (Asteraceae: Asteroideae: Gnaphalieae).**
In: Cladistics 29 (6), p. 574–588. DOI: 10.1111/cla.12007.

**Sehgal, Deepmala; Raina, Soom Nath; Devarumath, Rachhaya M.; Sasakuma, Tetsuo (2009):**
**Nuclear DNA Assay in solving issues related to Ancestry of the domesticated diploid Safflower (*Carthamus tinctorius* L.) and the polyploid (*Carthamus*) taxa, and phylogenetic and genomic Relationships in the Genus *Carthamus* L. (Asteraceae).**
In: Molecular Phylogenetics and Evolution 53 (3), p. 631–644. DOI: 10.1016/j.ympev.2009.07.012.

**Simpson, Beryl B.; Arroyo, Mary T.K.; Sipe, Sandra; Dias de Moraes, Marta; McDill, Joshua R. (2009):**
**Phylogeny and Evolution of *Perezia* (Asteraceae: Mutisieae: Nassauviinae).**
In: Journal of Systematics and Evolution 47 (5), p. 431–443. DOI: 10.1111/j.1759-6831.2009.00039.x.

**Smissen, Rob D.O.B.; Breitwieser, I.; Ward, Josephine M. (2004):**
**Phylogenetic Implications of trans-specific Chloroplast DNA Sequence Polymorphism in New Zealand Gnaphalieae (Asteraceae).**
In: Plant Systematics and Evolution 249 (1-2), p. 37–53. DOI: 10.1007/s00606-004-0209-0.

**Soares, Polyana Noronha (2012):**
**Taxonomia de *Acilepidopsis, Chrysolaena, Echinocoryne, Stenocephalum* e *Vernonanthura* (Vernonieae, Asteraceae) de Minas Gerais, Brasil.**
Dissertation. Universidade Federal de Ueberlandia, Uberlandia, MG, Brasil.

**Sonboli, Ali; Kazempour-Osaloo, Shahrokh; Vallès, Joan; Oberprieler, Christoph (2011):**
**Systematic Status and phylogenetic Relationships of the enigmatic *Tanacetum paradoxum* Bornm. (Asteraceae, Anthemideae): Evidences from nrDNA ITS, micromorphological, and cytological Data.**
In: Plant Systematics and Evolution 292 (1-2), p. 85–93. DOI: 10.1007/s00606-010-0415-x.

**Sonboli, Ali; Oberprieler, Christoph (2012):**
**Insights into the phylogenetic and taxonomic Position of *Tanacetum semenovii* Herder (Compositae, Anthemideae) based on nrDNA ITS Sequences Data.**
In: Biochemical Systematics and Ecology 45, p. 166–170. DOI: 10.1016/j.bse.2012.06.026.

**Sonboli, Ali; Stroka, Kathrin; Kazempour-Osaloo, Shahrokh; Oberprieler, Christoph (2012):**
**Molecular Phylogeny and Taxonomy of *Tanacetum* L. (Compositae, Anthemideae) inferred from nrDNA ITS and cpDNA trnH–psbA Sequence Variation.**
In: Plant Systematics and Evolution 298 (2), p. 431–444. DOI: 10.1007/s00606-011-0556-6.

**Soto-Trejo, Fabiola; Schilling, Edward E.; Solórzano, Sofía; Oyama, Ken; Lira, Rafael; Dávila, Patricia (2015):**
**Phylogenetic Relationships in the Genus *Florestina* (Asteraceae, Bahieae).**
In: Plant Systematics and Evolution 301 (9), p. 2147–2160. DOI: 10.1007/s00606-015-1220-3.

**Stångberg, Frida; Ellis, Allan G.; Anderberg, Arne A. (2013):**
**Evolutionary Relationships in *Gorteria*: A Re-evaluation.**
In: Taxon 62 (3), p. 537–549. DOI: 10.12705/623.3.

**Steffen, Simone (2013):**
**Evolution von Miniaturisierung in arktisch-alpinen Lebensräumen in *Petasites* Mill., *Endocellion* Turcz. ex Herder, *Homogyne* Cass. und *Tussilago* L. (Asteraceae) sowie *Soldanella* L. (Primulaceae).**
Dissertation. Johannes Gutenberg-Universität Mainz, Mainz.

**Suárez-Santiago, Víctor N.; Consuelo Díaz E.; Soltis, Douglas E.; Soltis, Pamela S.; Blanca, Gabriel (2011):**
***Tragopogon lainzii* a new Species of *Tragopogon* (Asteraceae) Segregated from *T. dubius* : Evidence from morphological and molecular Data.**
In: Systematic Botany 36 (2), p. 470–480. DOI: 10.1600/036364411X569651.

**Suárez-Santiago, Víctor N.; Salinas, María J.; Garcia-Jacas, Núria; Soltis, Pamela S.; Soltis, Douglas E.; Blanca, Gabriel (2007):**
**Reticulate Evolution in the *Acrolophus* subgroup (*Centaurea* L., Compositae) from the western Mediterranean: Origin and Diversification of Section *Willkommia* Blanca.**
In: Molecular Phylogenetics and Evolution 43 (1), p. 156–172. DOI: 10.1016/j.ympev.2006.08.006.

**Susanna, Alfonso; Galbany-Casals, Mercè; Romaschenko, Konstantin; Barres, Laia; Martín, Joan; Garcia-Jacas, Núria (2011):**
**Lessons from *Plectocephalus* (Compositae, Cardueae-Centaureinae): ITS Disorientation in annuals and Beringian Dispersal as revealed by molecular Analyses.**
In: Annals of Botany 108 (2), p. 263–277. DOI: 10.1093/aob/mcr138.

**Susanna, Alfonso; Garcia-Jacas, Núria; Hidalgo, Oriane; Vilatersana, Roser; Garnatje, Teresa (2006):**
**The Cardueae (Compositae) Revisited: Insights from ITS, trn L- trn F, and matK Nuclear and Chloroplast DNA Analysis.**
In: Annals of the Missouri Botanical Garden 93 (1), p. 150–171. DOI: 10.3417/0026-6493(2006)93[150:TCCRIF]2.0.CO;2.

**Susanna, Alfonso; Garnatje, Teresa; Garcia-Jacas, Núria (1999):**
**Molecular Phylogeny of *Cheirolophus* (Asteraceae: Cardueae-Centaureinae) based on ITS Sequences of nuclear ribosomal DNA.**
In: Plant Systematics and Evolution 214, p. 147–160.

**Swenson, Ulf; Manns, Ulrika (2003):**
**Phylogeny of *Pericallis* (Asteraceae): a total Evidence Approach Reappraising the double Origin of Woodiness.**
In: Taxon 52, p. 533–546.

**Tadesse, Mesfin; Crawford, Daniel J. (2014):**
**The phytomelanin layer in traditional Members of *Bidens* and *Coreopsis* and Phylogeny of the Coreopsideae (Compositae).**
In: Nordic Journal of Botany 32, p. 80–91.

**Takayama, Koji; López-Sepúlveda, Patricio; Greimler, Josef; Crawford, Daniel J.; Peñailillo, Patricio; Baeza, Marcelo et al. (2015):**

**Relationships and genetic Consequences of contrasting Modes of Speciation among endemic Species of *Robinsonia* (Asteraceae, Senecioneae) of the Juan Fernández Archipelago, Chile, based on AFLPs and SSRs.**
In: the new Phytologist 205 (1), p. 415–428. DOI: 10.1111/nph.13000.

TARIKAHYA-HACIOĞLU, BURCU; KARACAOĞLU, ÇAĞAŞAN; ÖZÜDOĞRU, BARIŞ (2014):
**The Speciation History and Systematics of *Carthamus* (Asteraceae) with special Emphasis on Turkish Species by integrating phylogenetic and ecological Niche modelling Data.**
In: Plant Systematics and Evolution 300 (6), p. 1349–1359. DOI: 10.1007/s00606-013-0966-8.

TERRAB, ANASS; ORTIZ, MARÍA ANGELES; TALAVERA, MARÍA; ARIZA, MARÍA JESÚS; MORIANA, MARÍA DEL CARMEN; GARCÍA-CASTAÑO, JUAN LUIS ET AL. (2009):
**AFLP and breeding system Studies indicate Vicariance Origin for scattered Populations and enigmatic low Fecundity in the Moroccan endemic *Hypochaeris angustifolia* (Asteraceae), sister Taxon to all of the South American *Hypochaeris* Species.**
In: Molecular Phylogenetics and Evolution 53 (1), p. 13–22. DOI: 10.1016/j.ympev.2009.06.008.

TIMME, RUTH E.; SIMPSON, BERYL B.; LINDER, C. RANDAL (2007):
**High-resolution Phylogeny for *Helianthus* (Asteraceae) using the 18s-26s ribosomal DNA external transcribed spacer.**
In: American Journal of Botany 94 (11), p. 1837–1852. DOI: 10.3732/ajb.94.11.1837.

TOMASELLO, SALVATORE; ÁLVAREZ, INÉS; VARGAS, PABLO; OBERPRIELER, CHRISTOPH (2015):
**Is the extremely rare Iberian endemic Plant Species *Castrilanthemum debeauxii* (Compositae, Anthemideae) a 'living fossil'? Evidence from a multi-locus Species Tree Reconstruction.**
In: Molecular Phylogenetics and Evolution 82 Pt A, p. 118–130. DOI: 10.1016/j.ympev.2014.09.007.

TORICES, RUBÉN; ANDERBERG, ARNE A. (2009):
**Phylogenetic Analysis of sexual Systems in Inuleae (Asteraceae).**
In: American Journal of Botany 96 (5), p. 1011–1019. DOI: 10.3732/ajb.0800231.

TORRELL, M.; CERBAH, M.; SILJAK-YAKOVLEV, SONJA; VALLS, J. (2003):
**Molecular cytogenetics of the Genus *Artemisia* (Asteraceae, Anthemideae): Fluorochrome banding and fluorescence in situ Hybridization. I. Subgenus *Seriphidium* and related Taxa.**
In: Plant Systematics and Evolution 239 (1-2), p. 141–153. DOI: 10.1007/s00606-002-0259-0.

TREMETSBERGER, KARIN; STUESSY, TOD F.; KADLEC, GERTRUD; URTUBEY, ESTRELLA; BAEZA, CARLOS M.; BECK, STEPHAN G. ET AL. (2006):
**AFLP Phylogeny of South American Species of *Hypochaeris* (Asteraceae, Lactuceae).**
In: Systematic Botany 31 (3), p. 610–626. DOI: 10.1600/036364406778388520.

TREMETSBERGER, KARIN; WEISS-SCHNEEWEISS, HANNA; STUESSY, TOD F.; SAMUEL, ROSABELLE; KADLEC, GERTRUD; ORTIZ, MARÍA ANGELES; TALAVERA, SALVADOR (2005):
**Nuclear ribosomal DNA and Karyotypes indicate a NW-African Origin of South American *Hypochaeris* (Asteraceae, Cichorieae).**
In: Molecular Phylogenetics and Evolution 35 (1), p. 102–116. DOI: 10.1016/j.ympev.2004.12.022.

URBATSCH, LOWELL E.; ROBERTS, ROLAND P.; KARAMAN, VESNA (2003):
**Phylogenetic Evaluation of *Xylothamia, Gundlachia*, and related Genera (Asteraceae, Astereae) based on ETS and ITS nrDNA Sequence Data.**

In: American Journal of Botany 90 (4), p. 634–649. DOI: 10.3732/ajb.90.4.634.

**Vaezi, Jamil; Brouillet, Luc (2009):**
**Phylogenetic Relationships among diploid Species of *Symphyotrichum* (Asteraceae: Astereae) based on two nuclear Markers, ITS and Gapdh.**
In: Molecular Phylogenetics and Evolution 51 (3), p. 540–553. DOI: 10.1016/j.ympev.2009.03.003.

**Vanijajiva, Ongkarn; Kadereit, Joachim W. (2008):**
**A Revision of *Cissampelopsis* (Asteraceae: Senecioneae).**
In: Kew Bulletin 63, p. 213–226.

**Vanijajiva, Ongkarn; Kadereit, Joachim W. (2011):**
**A Revision of *Gynura* (Asteraceae: Senecioneae).**
In: Journal of Systematics and Evolution 49 (4), p. 285–314. DOI: 10.1111/j.1759-6831.2011.00139.x.

**Vargas, Oscar M.; Madriñán, Santiago (2012):**
**Preliminary Phylogeny of *Diplostephium* (Asteraceae): Speciation Rate and Character Evolution.**
In: Lundellia 15, p. 1–15.

**Vilatersana, Roser; Garcia-Jacas, Núria; Garnatje, Teresa; Molero, Julián; Sonnante, Gabriella; Susanna, Alfonso (2010):**
**Molecular Phylogeny of the Genus *Ptilostemon* (Compositae: Cardueae) and its Relationships with *Cynara* and *Lamyropsis*.**
In: Systematic Botany 35 (4), p. 907–917. DOI: 10.1600/036364410X539952.

**Vilatersana, Roser; Susanna, Alfonso; Garcia-Jacas, Núria; Garnatje, Teresa (2000):**
**Generic Delimitation and Phylogeny of the *Carduncellus-Carthamus* complex (Asteraceae) based on ITS Sequences.**
In: Plant Systematics and Evolution 221, p. 89–105.

**Vitali, Maira Soledad; Sancho, Gisela; Katinas, Liliana (2015):**
**A Revision of *Smallanthus* (Asteraceae, Millerieae), the “yacón” Genus.**
In: Phytotaxa 214 (1), p. 1. DOI: 10.11646/phytotaxa.214.1.1.

**Wagstaff, Steven J.; Breitwieser, I. (2002):**
**Phylogenetic Relationships of New Zealand Asteraceae inferred from ITS Sequences.**
In: Plant Systematics and Evolution 231, p. 203–224.

**Wagstaff, Steven J.; Breitwieser, Ilse (2004):**
**Phylogeny and Classification of *Brachyglottis* (Senecioneae, Asteraceae): An Example of a Rapid Species Radiation in New Zealand.**
In: Systematic Botany 29 (4), p. 1003–1010. DOI: 10.1600/0363644042450991.

**Wagstaff, Steven J.; Breitwieser, Ilse; Ito, Motomi (2011):**
**Evolution and Biogeography of *Pleurophyllum* (Astereae, Asteraceae), a small Genus of Megaherbs endemic to the Subantarctic Islands.**
In: American Journal of Botany 98 (1), p. 62–75. DOI: 10.3732/ajb.1000238.

**Wagstaff, Steven J.; Breitwieser, Ilse; Swenson, Ulf (2006):**
**Origin and Relationships of the austral Genus *Abrotanella* (Asteraceae) inferred from DNA Sequences.**
In: Taxon 55 (1), p. 95–106.

**Wahrmund, Ute; Heklau, Heike; Röser, Martin; Kästner, Arndt; Vitek, Ernst; Ehrendorfer, Friedrich; Hagen, K. Bernhard von (2010):**

**A molecular Phylogeny reveals frequent Changes of growth form in *Carlina* (Asteraceae).**
In: Taxon 59 (2), p. 367–378.

Wang, Guang-yan; Meng, Ying; Deng, T. A.O.; Yang, Yong-Ping (2014):
**Molecular Phylogeny of *Faberia* (Asteraceae: Cichorieae) based on nuclear and Chloroplast Sequences.**
In: Phytotaxa 167 (3), p. 223. DOI: 10.11646/phytotaxa.167.3.1.

Wang, Qiang; Ma, Xin-Tang; Hong, De-Yuan (2014):
**Phylogenetic Analyses reveal three new Genera of the Campanulaceae.**
In: Journal of Systematics and Evolution 52 (5), p. 541–550. DOI: 10.1111/jse.12096.

Wang, Yu-Jin; Raab-Straube, Eckhard; Susanna, Alfonso; Liu, Jian-Quan (2013):
***Shangwua* (Compositae), a new Genus from the Qinghai-Tibetan Plateau and Himalayas.**
In: Taxon 62 (5), p. 984–996. DOI: 10.12705/625.19.

Watanabe, K.; Kosuge, Keiko; Shimamura, R.; Konishi, N.; Taniguchi, K. (2006):
**Molecular Systematics of Australian *Calotis* (Asteraceae: Astereae).**
In: Australian Systematic Botany 19 (2), p. 155. DOI: 10.1071/SB05001.

Watson, Linda E.; Bates, Paul L.; Evans, Timothy M.; Unwin, Matthew M.; Estes, James R. (2002):
**Molecular Phylogeny of Subtribe Artemisiinae (Asteraceae), including *Artemisia* and its allied and segregate Genera.**
In: BMC Evolutionary Biology 2, p. 17.

Watson, Linda E.; Evans, Timothy M.; Boluarte, T. (2000):
**Molecular Phylogeny and Biogeography of Tribe Anthemideae (Asteraceae), based on Chloroplast Gene ndhF.**
In: Molecular Phylogenetics and Evolution 15 (1), p. 59–69. DOI: 10.1006/mpev.1999.0714.

Weiss-Schneeweiss, Hanna; Blöch, Cordula; Turner, Barbara; Villaseñor, José Luis; Stuessy, Tod F.; Schneeweiss, Gerald M. (2012):
**The promiscuous and the chaste: frequent allopolyploid Speciation and its genomic Consequences in American daisies (*Melampodium* Sect. *Melampodium*; Asteraceae).**
In: Evolution 66 (1), p. 211–228. DOI: 10.1111/j.1558-5646.2011.01424.x.

Weiss-Schneeweiss, Hanna; Tremetsberger, Karin; Schneeweiss, Gerald M.; Parker, John S.; Stuessy, Tod F. (2008):
**Karyotype Diversification and Evolution in diploid and polyploid South American Hypochaeris (Asteraceae) inferred from rDNA localization and genetic fingerprint Data.**
In: Annals of Botany 101 (7), p. 909–918. DOI: 10.1093/aob/mcn023.

Yuan, Qian; Bi, Yanchao; Chen, You-Sheng (2015):
**Diplazoptilon (Asteraceae) is merged with *Saussurea* based on Evidence from Morphology and molecular Systematics.**
In: Phytotaxa 236 (1), p. 53. DOI: 10.11646/phytotaxa.236.1.4.

Yu-Jin, Wang; Jian-Quan, Liu (2004):
**A preliminary Investigation on the Phylogeny of *Saussurea* (Asteraceae: Cardueae) based on Chloroplast DNA trnL-F Sequences.**
In: Acta Phytotaxonomica Sinica 42 (2), p. 136–153.

ZHANG, JIAN-WEN; BOUFFORD, DAVID E.; SUN, HANG (2011):
**_Parasyncalathium_ J.W.Zhang, Boufford & H. Sun (Asteraceae, Cichorieae): A new Genus endemic to the Himalaya-Hengduan Mountains.**
In: Taxon 60 (6), p. 1678–1684.

ZHANG, JIAN-WEN; NIE, ZE-LONG; WEN, JUN; SUN, HANG (2011):
**Molecular Phylogeny and Biogeography of three closely related Genera, _Soroseris, Stebbinsia,_ and _Syncalathium_ (Asteraceae, Cichorieae), endemic to the Tibetan Plateau, SW-China.**
In: Taxon 60 (1), p. 15–26.

ZHAO, GUOHUA; CAO, ZHENGFEI; ZHANG, WEI; ZHAO, HONG (2015):
**The sesquiterpenoids and their chemotaxonomic Implications in _Senecio_ L. (Asteraceae).**
In: Biochemical Systematics and Ecology 59, p. 340–347. DOI: 10.1016/j.bse.2015.02.001.

ZHAO, HONG-BO; CHEN, FA-DI; CHEN, SU-MEI; WU, GUO-SHENG; GUO, WEI-MING (2010):
**Molecular Phylogeny of _Chrysanthemum, Ajania_ and its allies (Anthemideae, Asteraceae) as inferred from nuclear ribosomal ITS and Chloroplast trnL-F Igs Sequences.**
In: Plant Systematics and Evolution 284 (3-4), p. 153–169. DOI: 10.1007/s00606-009-0242-0.

### Asterales

KÅREHED, JESPER; LUNDBERG, JOHANNES; BREMER, BIRGITTA; BREMER, KARE (1999):
**Evolution of the Australasian families Alseusomiaceae, Argophyllaceae, and Phellinaceae.**
In: Systematic Botany 24 (4), p. 660–682.

LINDER, C. RANDAL; GOERTZEN, LESLIE R.; HEUVEL, B. VANDEN; FRANCISCO-ORTEGA, JAVIER; JANSEN, ROBERT K. (2000):
**The complete external transcribed spacer of 18s-26s rDNA: amplification and phylogenetic Utility at low taxonomic Levels in Asteraceae and closely allied Families.**
In: Molecular Phylogenetics and Evolution 14 (2), p. 285–303. DOI: 10.1006/mpev.1999.0706.

### Athyriaceae

ADJIEI, BAYU; TAKAMIYA, MASAYUKI; OHTA, MICHIHITO; OHSAWA, TAKESHI A.; WATANO, YASUYUKI (2008):
**Molecular Phylogeny of the Lady Fern Genus _Athyrium_ in Japan based on Chloroplast rbcL and trnL-trnF- Sequences.**
In: Acta Phytotaxonomica Geobotanica 59 (2), p. 79–95.

KUO, LI-YAUNG; EBIHARA, ATSUSHI; SHINOHARA, WATARU; ROUHAN, GERMINAL; WOOD, KENNETH R.; WANG, CHUN-NENG; CHIOU, WEN-LIANG (2016):
**Historical Biogeography of the Fern Genus _Deparia_ (Athyriaceae) and its Relation with Polyploidy.**
In: Molecular Phylogenetics and Evolution 104, p. 123–134. DOI: 10.1016/j.ympev.2016.08.004.

LIU, YEA-CHEN; CHIOU, WEN-LIANG; KATO, MASAHIRO (2011):
**Molecular Phylogeny and Taxonomy of the Fern Genus _Anisocampium_ (Athyriaceae).**
In: Taxon 60 (3), p. 824–830.

WEI, RAN; EBIHARA, ATSUSHI; ZHU, YAN-MEI; ZHAO, CUN-FENG; HENNEQUIN, SABINE; ZHANG, XIAN-CHUN (2018):
**A total-Evidence Phylogeny of the Lady Fern Genus _Athyrium_ Roth (Athyriaceae) with a new infrageneric Classification.**

In: Molecular Phylogenetics and Evolution 119, p. 25–36. DOI: 10.1016/j.ympev.2017.10.019.

**Wei, Ran; Schneider, Harald; Zhang, Xian-Chun (2013):**

**Toward a new Circumscription of the twinsorus-fern Genus *Diplazium* (Athyriaceae): A molecular Phylogeny with morphological Implications and infrageneric Taxonomy.**

In: Taxon 62 (3), p. 441–457.

### Aucubaceae

**Ohi, Tetsuo; Kajita, Tadashi; Murata, Jin (2003):**

**Distinct geographic Structure as Evidenced by Chloroplast DNA haplotypes and ploidy level in Japanese *Aucuba* (Aucubaceae).**

In: American Journal of Botany 90 (11), p. 1645–1652. DOI: 10.3732/ajb.90.11.1645.

### Balanophoraceae

**Su, Huei-Jiun; Hu, Jer-Ming; Anderson, Frank E.; Der, Joshua P.; Nickrent, Daniel Lee (2015):**

**Phylogenetic Relationships of Santalales with Insights into the Origins of holoparasitic Balanophoraceae.**

In: Taxon 64 (3), p. 491–506. DOI: 10.12705/643.2.

**Su, Huei-Jiun; Murata, Jin; Hu, Jer-Ming (2012):**

**Morphology and Phylogenetics of two holoparasitic plants, *Balanophora japonica* and *Balanophora yakushimensis* (Balanophoraceae), and their hosts in Taiwan and Japan.**

In: Journal of Plant Research 125 (3), p. 317–326. DOI: 10.1007/s10265-011-0447-5.

### Balsaminaceae

**Fischer, Eberhard; Rahelivololona, Marie Elisette (2016):**

**New Taxa of *Impatiens* (Balsaminaceae) from Madagascar VIII. *Impatiens max-huberi*, a new Species from Marojejy and Anjanaharibe-Sud.**

In: Phytotaxa 244 (2), p. 191. DOI: 10.11646/phytotaxa.244.2.7.

**Janssens, Steven B.; Fischer, Eberhard; Stévart, Tariq (2010):**

**New Insights into the Origin of two new epiphytic *Impatiens* Species (Balsaminaceae) from West Central Africa based on molecular phylogenetic Analyses.**

In: Taxon 59 (5), p. 1508–1518.

**Janssens, Steven B.; Geuten, Koen; Viaene, Tom; Yuan, Yong-Ming; Song, Yi; Smets, Erik (2007):**

**Phylogenetic utility of the Ap3/def K-domain and its molecular Evolution in *Impatiens* (Balsaminaceae).**

In: Molecular Phylogenetics and Evolution 43 (1), p. 225–239. DOI: 10.1016/j.ympev.2006.11.016.

**Janssens, Steven B.; Geuten, Koen; Yuan, Yong-Ming; Song, Yi; Küpfer, Philippe; Smets, Erik (2006):**

**Phylogenetics of *Impatiens* and *Hydrocera* (Balsaminaceae) using Chloroplast atpB-rbcL Spacer Sequences.**

In: Systematic Botany 31 (1), p. 171–180.

**Janssens, Steven B.; Knox, Eric B.; Huysmans, Suzy; Smets, Erik F.; Merckx, Vincent S.F.T. (2009):**

**Rapid Radiation of *Impatiens* (Balsaminaceae) during Pliocene and Pleistocene: Result of a global Climate Change.**

In: Molecular Phylogenetics and Evolution 52 (3), p. 806–824. DOI: 10.1016/j.ympev.2009.04.013.

**Janssens, Steven B.; Sonké, Bonaventure; Lachenaud, Olivier; Lemaire, Benny; Simo-Droissart, Murielle; Smets, Erik (2015):**

**Morphology, Molecular Phylogenetics and Biogeography of *Impatiens akomensis* (Balsaminaceae), a new Species from Cameroon.**
In: Plant Ecology and Evolution 148 (3), p. 397–408. DOI: 10.5091/plecevo.2015.965.

JANSSENS, STEVEN B.; WILSON, YI SONG; YUAN, YONG-MING; NAGELS, ANNE; SMETS, ERIK F.; HUYSMANS, SUZY (2012):
**A total Evidence approach using palynological Characters to infer the complex evolutionary History of the Asian *Impatiens* (Balsaminaceae).**
In: Taxon 61 (2), p. 355–367.

UTAMI, NANDA; ARDIYANI, MARLINA (2015):
**Phylogenetic Study of Sumatran *Impatiens* (Balsaminaceae) using Nuclear and Plastid DNA Sequences.**
In: Acta Phytotaxonomica Geobotanica 66 (2), p. 81–90.

UTAMI, NANDA; SHIMIZU, TATEMI (2005):
**Seed Morphology and Classification of *Impatiens* (Balsaminaceae).**
In: Blumea 50 (3), p. 447–456. DOI: 10.3767/000651905X622699.

VOLKMAR, UTE; SMETS, ERIK F.; LENZ, HENNING; JANSSENS, STEVEN B. (2014):
**Intron Evolution in a phylogenetic Perspective: Divergent Trends in the two Copies of the duplicated DEF Gene in *Impatiens* L. (Balsaminaceae).**
In: Journal of Systematics and Evolution 52 (2), p. 134–148. DOI: 10.1111/jse.12070.

YU, SHENG-XIANG; JANSSENS, STEVEN B.; ZHU, XIANG-YUN; LIDÉN, MAGNUS; GAO, TIAN-GANG; WANG, WEI (2016):
**Phylogeny of *Impatiens* (Balsaminaceae): integrating molecular and morphological Evidence into a new Classification.**
In: Cladistics 32 (2), p. 179–197. DOI: 10.1111/cla.12119.

YUAN, YONG-MING; SONG, YI; GEUTEN, KOEN; RAHELIVOLOLONA, MARIE ELISETTE; WOHLHAUSER, SÉBASTIEN; FISCHER, EBERHARD ET AL. (2004):
**Phylogeny and Biogeography of Balsaminaceae inferred from ITS Sequences.**
In: Taxon 53 (2), p. 391–403.

ZHANG, JASON GECHEN; ZHANG, LI-BING (2011):
***Impatiens shimianensis* sp. nov. (Balsaminaceae): A new Species from Sichuan, China, based on morphological and molecular Evidence.**
In: Systematic Botany 36 (3), p. 721–729. DOI: 10.1600/036364411X583682.

### Basellaceae

ERIKSSON, ROGER (2007):
**A Synopsis of Basellaceae.**
In: Kew Bulletin 62 (2), p. 297–320.

### Begoniaceae

BRENNAN, ADRIAN CHRISTOPHER; BRIDGETT, STEPHEN; SHAUKAT ALI, MOBINA; HARRISON, NICOLA; MATTHEWS, ANDREW; PELLICER, JAUME ET AL. (2012):
**Genomic Resources for evolutionary Studies in the large, diverse tropical Genus *Begonia*.**
In: Tropical Plant Biology 5 (4), p. 261–276. DOI: 10.1007/s12042-012-9109-6.

CLEMENT, WENDY L.; TEBBITT, MARK C.; FORREST, LAURA L.; BLAIR, JAIME E.; BROUILLET, LUC; ERIKSSON, TORSTEN; SWENSEN, SUSAN M. (2004):
**Phylogenetic Position and Biogeography of *Hillebrandia sandwicensis* (Begoniaceae): a rare Hawaiian relict.**

In: American Journal of Botany 91 (6), p. 905–917. DOI: 10.3732/ajb.91.6.905.

**Lowe-Forrest, Laura; Hughes, Mark; Hollingsworth, Peter M. (2005):**
**A Phylogeny of *Begonia* using Nuclear Ribosomal Sequence Data and morphological Characters.**
In: Systematic Botany 30 (3), p. 671–682.

**Plana, Vanessa; Gascoigne, Angus; Forrest, Laura L.; Harris, David J.; Pennington, R. Toby (2004):**
**Pleistocene and pre-Pleistocene *Begonia* Speciation in Africa.**
In: Molecular Phylogenetics and Evolution 31 (2), p. 449–461. DOI: 10.1016/j.ympev.2003.08.023.

**Tebbitt, Mark C.; Dickson, James H. (2000):**
**Amended Descriptions and Revised Sectional Assignment of Some Asian Begonias (Begoniaceae).**
In: Brittonia 52 (1), p. 112. DOI: 10.2307/2666496.

**Tebbitt, Mark C.; Lowe-Forrest, Laura; Santoriello, Anthony; Clement, Wendy L.; Swensen, Susan M. (2006):**
**Phylogenetic Relationships of Asian *Begonia*, with an Emphasis on the Evolution of Rain-ballist and Animal Dispersal Mechanisms in Sections *Platycentrum, Sphenanthera* and *Leprosae*.**
In: Systematic Botany 31 (2), p. 327–336.

**Thomas, Daniel C.; Hughes, Mark; Phutthai, T.; Rajbhandary, S.; Rubite, R.; Ardi, W. H.; Richardson, James E. (2011):**
**A non-coding Plastid DNA Phylogeny of Asian *Begonia* (Begoniaceae): Evidence for morphological Homoplasy and Sectional Polyphyly.**
In: Molecular Phylogenetics and Evolution 60 (3), p. 428–444. DOI: 10.1016/j.ympev.2011.05.006.

### Berberidaceae

**Adhikari, Bhaskar; Milne, Richard Ian; Pennington, R. Toby; Särkinen, Tiina E.; Pendry, Colin Alistair (2015):**
**Systematics and Biogeography of *Berberis* s.l. inferred from nuclear ITS and Chloroplast ndhF Gene Sequences.**
In: Taxon 64 (1), p. 39–48. DOI: 10.12705/641.21.

**Kim, Young-Dong; Jansen, Robert K. (1998):**
**Chloroplast DNA Restriction Site Variation and Phylogeny of the Berberidaceae.**
In: American Journal of Botany 85 (12), p. 1766–1778.

**Kim, Young-Dong; Kim, Sung-Hee; Kim, Chul Hwan; Jansen, Robert K. (2004):**
**Phylogeny of Berberidaceae based on Sequences of the Chloroplast Gene ndhF.**
In: Biochemical Systematics and Ecology 32 (3), p. 291–301. DOI: 10.1016/j.bse.2003.08.002.

**Kim, Young-Dong; Kim, Sung-Hee; Landrum, Leslie R. (2004):**
**Taxonomic and phytogeographic Implications from ITS Phylogeny in *Berberis* (Berberidaceae).**
In: Journal of Plant Research 117 (3), p. 175–182. DOI: 10.1007/s10265-004-0145-7.

**Li, Yong; Zhai, Sheng-Nan; Qiu, Ying-Xiong; Guo, Yan-Ping; Ge, Xue-Jun; Comes, Hans Peter (2011):**
**Glacial survival east and west of the 'Mekong-Salween Divide' in the Himalaya-Hengduan Mountains Region as revealed by AFLPs and cpDNA Sequence Variation in *Sinopodophyllum hexandrum* (Berberidaceae).**
In: Molecular Phylogenetics and Evolution 59 (2), p. 412–424. DOI: 10.1016/j.ympev.2011.01.009.

**Sheng, Mao-Yin; Wang, Ling-Jiao (2010):**

**Chromosomal localization of 45s and 5s rDNA in 14 Species and the Implications for Genome Evolution of Genus *Epimedium*.**
In: Plant Systematics and Evolution 290 (1-4), p. 65–73. DOI: 10.1007/s00606-010-0349-3.

SUN, YE; FUNG, KWOK-PUI; LEUNG, PING-CHUNG; SHAW, PANG-CHUI (2005):
**A phylogenetic Analysis of *Epimedium* (Berberidaceae) based on nuclear ribosomal DNA Sequences.**
In: Molecular Phylogenetics and Evolution 35 (1), p. 287–291. DOI: 10.1016/j.ympev.2004.12.014.

WANG, WEI; CHEN, ZHI-DUAN; LIU, YANG; LI, RUI-QI; LI, JIAN-HUA (2007):
**Phylogenetic and Biogeographic Diversification of Berberidaceae in the Northern Hemisphere.**
In: Systematic Botany 32 (4), p. 731–742.

ZHANG, MING-LI; UHINK, CHRISTIAN H.; KADEREIT, JOACHIM W. (2007):
**Phylogeny and Biogeography of *Epimedium/Vancouveria* (Berberidaceae): Western North American - East Asian Disjunctions, the Origin of European Mountain Plant Taxa, and East Asian Species Diversity.**
In: Systematic Botany 32 (1), p. 81–92.

### Betulaceae

CHEN, ZHI-DUAN; LI, JIAN-HUA (2004):
**Phylogenetics and Biogeography of *Alnus* (Betulaceae) inferred from Sequences of Nuclear Ribosomal DNA ITS Region.**
In: International Journal of Plant Sciences 165 (2), p. 325–335.

CHEN, ZHI-DUAN; MANCHESTER, STEVEN R.; SUN, HAI-YING (1999):
**Phylogeny and Evolution of the Betulaceae as inferred from DNA Sequences, Morphology, and Paleobotany.**
In: American Journal of Botany 86 (8), p. 1168–1181. DOI: 10.2307/2656981.

FOREST, FÉLIX; BRUNEAU, ANNE (2000):
**Phylogenetic Analysis, Organization, and molecular Evolution of the Nontranscribed Spacer of 5s Ribosomal RNA Genes in *Corylus* (Betulaceae).**
In: International Journal of Plant Sciences 161 (5), p. 793–806.

JÄRVINEN, PIA; PALMÉ, ANNA; ORLANDO MORALES, LUIS; LÄNNENPÄÄ, MIKA; KEINÄNEN, MARKKU; SOPANEN, TUOMAS; LASCOUX, MARTIN (2004):
**Phylogenetic Relationships of *Betula* Species (Betulaceae) based on nuclear Adh and Chloroplast matK Sequences.**
In: American Journal of Botany 91 (11), p. 1834–1845. DOI: 10.3732/ajb.91.11.1834.

LI, JIAN-HUA; SHOUP, SUZANNE; CHEN, ZHI-DUAN (2005):
**Phylogenetics of *Betula* (Betulaceae) inferred from Sequences of nuclear ribosomal DNA.**
In: Rhodora 107 (929), p. 69–86. DOI: 10.3119/04-14.1.

LI, JIAN-HUA; SHOUP, SUZANNE; CHEN, ZHI-DUAN (2007):
**Phylogenetic Relationships of Diploid Species of *Betula* (Betulaceae) inferred from DNA Sequences of Nuclear Nitrate Reductase.**
In: Systematic Botany 32 (2), p. 357–365.

MA, HUI; LU, JING; LIU, BING-BING; DUAN, BING-BING; HE, XIAO-DONG; LIU, JIAN-QUAN (2015):
**Phylotranscriptomic Analyses in plants using Betulaceae as an example.**

In: Journal of Systematics and Evolution 53 (5), p. 403–410. DOI: 10.1111/jse.12178.

**Yoo, Ki-Oug; Wen, Jun (2002):**
**Phylogeny and Biogeography of *Carpinus* and Subfamily Coryloideae (Betulaceae).**
In: International Journal of Plant Sciences 163 (4), p. 641–650.

**Yoo, Ki-Oug; Wen, Jun (2007):**
**Phylogeny of *Carpinus* and Subfamily Coryloideae (Betulaceae) based on Chloroplast and nuclear ribosomal Sequence Data.**
In: Plant Systematics and Evolution 267 (1-4), p. 25–35. DOI: 10.1007/s00606-007-0533-2.

### Biebersteiniaceae

**Müllner, Alexandra N.; Vassiliades, D. D.; Renner, Susanne p. (2007):**
**Placing Biebersteiniaceae, a herbaceous Clade of Sapindales, in a temporal and geographic Context.**
In: Plant Systematics and Evolution 266 (3-4), p. 233–252. DOI: 10.1007/s00606-007-0546-x.

### Bignoniaceae

**Alcantara, Suzana; Lohmann, Lúcia Garces (2010):**
**Evolution of Floral Morphology and Pollination System in Bignonieae (Bignoniaceae).**
In: American Journal of Botany 97 (5), p. 782–796. DOI: 10.3732/ajb.0900182.

**Callmander, Martin W.; Phillipson, Peter B.; Plunkett, Gregory M.; Edwards, Molly B.; Buerki, Sven (2016):**
**Generic Delimitations, Biogeography and Evolution in the Tribe Coleeae (Bignoniaceae), endemic to Madagascar and the smaller Islands of the western Indian Ocean.**
In: Molecular Phylogenetics and Evolution 96, p. 178–186. DOI: 10.1016/j.ympev.2015.11.016.

**Chen, Shaotian; Guan, Kaiyun; Zhou, Zhekun; Olmstead, Richard G.; Cronk, Quentin C.B. (2005):**
**Molecular Phylogeny of *Incarvillea* (Bignoniaceae) based on ITS and trnL-F Sequences.**
In: American Journal of Botany 92 (4), p. 625–633. DOI: 10.3732/ajb.92.4.625.

**Collevatti, Rosane G.; Dornelas, Marcelo C. (2016):**
**Clues to the Evolution of Genome size and Chromosome Number in *Tabebuia* alliance (Bignoniaceae).**
In: Plant Systematics and Evolution 302 (5), p. 601–607. DOI: 10.1007/s00606-016-1280-z.

**Fonseca, Luiz Henrique Martins; Lohmann, Lúcia Garces (2015):**
**Biogeography and Evolution of *Dolichandra* (Bignonieae, Bignoniaceae).**
In: Botanical Journal of the Linnean Society 179, p. 403–420.

**Fonseca, Luiz Henrique Martins; Lohmann, Lúcia Garces (2018):**
**Combining high-throughput Sequencing and targeted Loci Data to infer the Phylogeny of the "*Adenocalymma-Neojobertia*" Clade (Bignonieae, Bignoniaceae).**
In: Molecular Phylogenetics and Evolution 123, p. 1–15. DOI: 10.1016/j.ympev.2018.01.023.

**Gomes, Beatriz M.; Proença, Carolyn E.B. (2010):**
**New Species of *Pleonotoma* (Bignonieae, Bignoniaceae) from Amazonia, Brazil.**
In: Kew Bulletin 65, p. 269–273.

**Grose, Susan O.; Olmstead, Richard G. (2007):**
**Evolution of a Charismatic Neotropical Clade: Molecular Phylogeny of *Tabebuia* s. l., Crescentieae, and Allied Genera (Bignoniaceae).**
In: Systematic Botany 32 (3), p. 650–659.

**Grose, Susan O.; Olmstead, Richard G. (2007):**
**Taxonomic Revisions in the Polyphyletic Genus *Tabebuia* s. l. (Bignoniaceae).**
In: Systematic Botany 32 (3), p. 660–670.

**Kaehler, Miriam; Michelangeli, Fabián Armando; Lohmann, Lúcia Garces (2012):**
**Phylogeny of *Lundia* (Bignoniaceae) based on ndhF and PepC Sequences.**
In: Taxon 61 (2), p. 368–380. DOI: 10.1002/tax.612008.

**Li, Jian-Hua (2008):**
**Phylogeny of *Catalpa* (Bignoniaceae) inferred from Sequences of Chloroplast ndhF and nuclear ribosomal DNA.**
In: Journal of Systematics and Evolution 46 (3), p. 341–348.

**Lohmann, Lúcia Garces (2006):**
**Untangling the Phylogeny of Neotropical Lianas (Bignonieae, Bignoniaceae).**
In: American Journal of Botany 93 (2), p. 304–318.

**Medeiros, Maria Cláudia Melo Pacheco de; Lohmann, Lúcia Garces (2015):**
**Phylogeny and Biogeography of *Tynanthus* Miers (Bignonieae, Bignoniaceae).**
In: Molecular Phylogenetics and Evolution 85, p. 32–40. DOI: 10.1016/j.ympev.2015.01.010.

**Medeiros, Maria Cláudia Melo Pacheco de; Lohmann, Lúcia Garces (2015):**
**Taxonomic Revision of *Tynanthus* (Bignonieae, Bignoniaceae).**
In: Phytotaxa 216 (1), p. 1. DOI: 10.11646/phytotaxa.216.1.1.

**Olmstead, Richard G. (2013):**
**Phylogeny and Biogeography in Solanaceae, Verbenaceae and Bignoniaceae: a Comparison of continental and intercontinental Diversification Patterns.**
In: Botanical Journal of the Linnean Society 171, p. 80–102.

**Olmstead, Richard G.; Zjhra, Michelle L.; Lohmann, Lúcia Garces; Grose, Susan O.; Eckert, Andrew J. (2009):**
**A molecular Phylogeny and Classification of Bignoniaceae.**
In: American Journal of Botany 96 (9), p. 1731–1743. DOI: 10.3732/ajb.0900004.

**Pace, Marcelo R.; Lohmann, Lúcia Garces; Olmstead, Richard G.; Angyalossy, Veronica (2015):**
**Wood Anatomy of major Bignoniaceae Clades.**
In: Plant Systematics and Evolution 301 (3), p. 967–995. DOI: 10.1007/s00606-014-1129-2.

**Pace, Marcelo R.; Zuntini, Alexandre R.; Lohmann, Lúcia Garces; Angyalossy, Veronica (2016):**
**Phylogenetic Relationships of enigmatic *Sphingiphila* (Bignoniaceae) based on molecular and wood anatomical Data.**
In: Taxon 65 (5), p. 1050–1063. DOI: 10.12705/655.7.

**Pool, Amy (2007):**
**A Review of the Genus *Distictis* (Bignoniaceae).**
In: Annals of the Missouri Botanical Garden 94 (4), p. 791–820. DOI: 10.3417/0026-6493(2007)94[791:AROTGD]2.0.CO;2.

**Pool, Amy (2007):**
**A Revision of the Genus *Pithecoctenium* (Bignoniaceae).**
In: Annals of the Missouri Botanical Garden 94 (3), p. 622–642. DOI: 10.3417/0026-6493(2007)94[622:AROTGP]2.0.CO;2.

**Pool, Amy (2008):**
**A Review of the Genus *Pyrostegia* (Bignoniaceae).**
In: Annals of the Missouri Botanical Garden 95 (3), p. 495–510. DOI: 10.3417/2003090.

**Pool, Amy (2009):**
**A Review of the Genus *Distictella* (Bignoniaceae).**
In: Annals of the Missouri Botanical Garden 96 (2), p. 286–323. DOI: 10.3417/2006156.

**Zjhra, Michelle L.; Sytsma, Kenneth J.; Olmstead, Richard G. (2004):**
**Delimitation of Malagasy Tribe Coleeae and Implications for Fruit Evolution in Bignoniaceae inferred from a Chloroplast DNA Phylogeny.**
In: Plant Systematics and Evolution 245 (1-2). DOI: 10.1007/s00606-003-0025-y.


### Blechnaceae

**Gasper, André Luís de; Almeida, Thaís Elias; Oliveira, Dittrich Vinícius Antonio De; Smith, Alan R.; Salino, Alexandre (2017):**
**Molecular Phylogeny of the Fern Family Blechnaceae (Polypodiales) with a revised Genus-level Treatment.**
In: Cladistics 33 (4), p. 429–446. DOI: 10.1111/cla.12173.

**Gasper, André Luís de; Oliveira, Dittrich Vinícius Antonio De; Salino, Alexandre (2016):**
**A Classification for Blechnaceae (Polypodiales: Polypodiopsida): new Genera, resurrected names, and Combinations.**
In: Phytotaxa 275 (3), p. 191. DOI: 10.11646/phytotaxa.275.3.1.

**Perrie, Leon R.; Wilson, Ruby K.; Shepherd, Lara D.; Ohlsen, Daniel J.; Batty, Erin L.; Brownsey, Patrick J.; Bayly, Michael J. (2014):**
**Molecular Phylogenetics and generic Taxonomy of Blechnaceae Ferns.**
In: Taxon 63 (4), p. 745–758. DOI: 10.12705/634.13.


### Bombacaceae

**Baum, David A.; Dewitt Smith, Stacey; Yen, Alan C.; Alverson, William S.; Nyffeler, Reto; Whitlock, Barbara A.; Oldham, Rebecca L. (2004):**
**Phylogenetic Relationships of *Malvatheca* (Bombacoideae and Malvoideae; Malvaceae sensu lato) as inferred from Plastid DNA Sequences.**
In: American Journal of Botany 91 (11), p. 1863–1871. DOI: 10.3732/ajb.91.11.1863.

**Carvalho-Sobrinho, Jefferson Guedes de; Alverson, William S.; Alcantara, Suzana; Queiroz, Luciano Paganucci; Mota, Aline C.; Baum, David A. (2016):**
**Revisiting the Phylogeny of Bombacoideae (Malvaceae): Novel Relationships, morphologically cohesive Clades, and a new tribal Classification based on multilocus phylogenetic Analyses.**
In: Molecular Phylogenetics and Evolution 101, p. 56–74. DOI: 10.1016/j.ympev.2016.05.006.

**Carvalho-Sobrinho, Jefferson Guedes de; Queiroz, Luciano Paganucci (2011):**
**Morphological cladistic Analysis of *Pseudobombax* Dugand (Malvaceae, Bombacoideae) and allied Genera.**
In: Brazilian Journal of Botany 34 (2), p. 197–209. DOI: 10.1590/S0100-84042011000200007.

**Duarte, Marília C.; Esteves, Gerleni Lopes; Salatino, Maria Luiza Faria; Walsh, Karen C.; Baum, David A. (2011):**
**Phylogenetic Analyses of *Eriotheca* and related Genera (Bombacoideae, Malvaceae).**
In: Systematic Botany 36 (3), p. 690–701. DOI: 10.1600/036364411X583655.

**Marinho, Rafaela C.; Mendes-Rodrigues, Clesnan; Balao, Francisco; Ortiz, Pedro L.; Yamagishi-Costa, Júlia; Bonetti, Ana M.; Oliveira, Paulo E. (2014):**
**Do Chromosome Numbers reflect Phylogeny? New counts for Bombacoideae and a Review of Malvaceae s.l.**

In: American Journal of Botany 101 (9), p. 1456–1465. DOI: 10.3732/ajb.1400248.

PETTIGREW FRS, JACK D.; BELL, KAREN L.; BHAGWANDIN, ADHIL; GRINAN, EUNICE; JILLANI, NGALLA; MEYER, JEAN ET AL. (2012):

**Morphology, ploidy and molecular Phylogenetics reveal a new diploid Species from Africa in the baobab Genus *Adansonia* (Malvaceae: Bombacoideae).**

In: Taxon 61 (6), p. 1240–1250. DOI: 10.1002/tax.616006.

RAZANAMARO, ONJA; RASOAMANANA, ELYSÉE; RAKOUTH, BAKOLIMALALA; RANDRIAMALALA, JOSOA RAMAROLANONANA; RABAKONADRIANINA, ELISABETH; CLÉMENT-VIDAL, ANNE ET AL. (2015):

**Chemical characterization of Floral scents in the six endemic baobab Species (*Adansonia* sp.) of Madagascar.**

In: Biochemical Systematics and Ecology 60, p. 238–248. DOI: 10.1016/j.bse.2015.04.005.

### Boraginaceae

CECCHI, LORENZO; COPPI, ANDREA; SELVI, FEDERICO (2009):

***Nonea palmyrensis* (Boraginaceae): Morphology and phylogenetic affinities of a rare endemic of the Syro-Iraqi desert.**

In: Nordic Journal of Botany 27 (5), p. 381–387. DOI: 10.1111/j.1756-1051.2009.00418.x.

CECCHI, LORENZO; COPPI, ANDREA; SELVI, FEDERICO (2011):

**Evolutionary dynamics of serpentine Adaptation in *Onosma* (Boraginaceae) as revealed by ITS Sequence Data.**

In: Plant Systematics and Evolution 297 (3-4), p. 185–199. DOI: 10.1007/s00606-011-0506-3.

CECCHI, LORENZO; SELVI, FEDERICO (2009):

**Phylogenetic Relationships of the monotypic Genera *Halacsya* and *Paramoltkia* and the Origins of serpentine Adaptation in Circum-mediterranean Lithospermeae (Boraginaceae): Insights from ITS and matK DNA Sequences.**

In: Taxon 58 (3), p. 700–714.

CHACÓN, JULIANA; LUEBERT, FEDERICO; HILGER, HARTMUT H.; OVCHINNIKOVA, SVETLANA; SELVI, FEDERICO; CECCHI, LORENZO ET AL. (2016):

**The Borage Family (Boraginaceae s.str.): A revised infrafamilial Classification based on new phylogenetic Evidence, with Emphasis on the Placement of some enigmatic Genera.**

In: Taxon 65 (3), p. 523–546. DOI: 10.12705/653.6.

COHEN, JAMES I. (2011):

**A phylogenetic Analysis of morphological and molecular Characters of *Lithospermum* L. (Boraginaceae) and related taxa: evolutionary Relationships and Character Evolution.**

In: Cladistics 27 (6), p. 559–580. DOI: 10.1111/j.1096-0031.2011.00352.x.

COHEN, JAMES I. (2014):

**A phylogenetic Analysis of morphological and molecular Characters of Boraginaceae: evolutionary Relationships, Taxonomy, and Patterns of Character Evolution.**

In: Cladistics 30 (2), p. 139–169. DOI: 10.1111/cla.12036.

DICKORÉ, W. BERNHARD; HILGER, HARTMUT H. (2015):

***Decalepidanthus* (Boraginaceae) includes and antedates *Pseudomertensia*; a Synopsis of the Genus.**

In: Phytotaxa 226 (2), p. 131. DOI: 10.11646/phytotaxa.226.2.3.

Dimon, Richard J.; Renner, Matt A. M. (2017):
**A new *Cynoglossum* species, and transfers to *Hackelia* in eastern Australian Boraginaceae.**
In: Australian Systematic Botany 30 (2), p. 113. DOI: 10.1071/SB17004.

García-Maroto, Federico; Mañas-Fernández, Aurora; Garrido-Cárdenas, José A.; Alonso, Diego López; Guil-Guerrero, José L.; Guzmán, Beatriz; Vargas, Pablo (2009):
**Delta6-desaturase Sequence Evidence for explosive Pliocene Radiations within the adaptive Radiation of Macaronesian *Echium* (Boraginaceae).**
In: Molecular Phylogenetics and Evolution 52 (3), p. 563–574. DOI: 10.1016/j.ympev.2009.04.009.

Hasenstab-Lehman, Kristen E.; Simpson, Michael G. (2012):
**Cat's Eyes and Popcorn Flowers: phylogenetic Systematics of the Genus *Cryptantha* s. l. (Boraginaceae).**
In: Systematic Botany 37 (3), p. 738–757. DOI: 10.1600/036364412X648706.

Hilger, Hartmut H.; Greuter, Werner; Stier, Victoria (2015):
**Taxa and Names in *Cynoglossum* sensu lato (Boraginaceae, Cynoglosseae): an annotated, synonymic Inventory, with links to the Protologues and Mention of original Material.**
In: Biodiversity Data Journal (3), e4831. DOI: 10.3897/BDJ.3.e4831.

Hilger, Hartmut H.; Selvi, Federico; Papini, Alessio; Bigazzi, Massimo (2004):
**Molecular Systematics of Boraginaceae Tribe Boragineae based on ITS1 and trnL Sequences, with special Reference to *Anchusa* s.l..**
In: Annals of Botany 94 (2), p. 201–212. DOI: 10.1093/aob/mch132.

Huang, Jian-Feng; Zhang, Ming-Li; Cohen, James I. (2013):
**Phylogenetic Analysis of *Lappula* Moench (Boraginaceae) based on molecular and morphological Data.**
In: Plant Systematics and Evolution 299 (5), p. 913–926. DOI: 10.1007/s00606-013-0772-3.

Khoshsokhan Mozaffar, Maryam; Kazempour-Osaloo, Shahrokh; Oskoueiyan, Roghayeh; Naderi Saffar, Kosar; Amirahmadi, Atefe (2013):
**Tribe Eritrichieae (Boraginaceae s.str.) in West Asia: a molecular phylogenetic Perspective.**
In: Plant Systematics and Evolution 299 (1), p. 197–208. DOI: 10.1007/s00606-012-0715-4.

Kolarčik, V.; Zozomová-Lihová, Judita; Mártonfi, P. (2010):
**Systematics and evolutionary History of the *Asterotricha* Group of the Genus *Onosma* (Boraginaceae) in central and southern Europe inferred from AFLP and nrDNA ITS Data.**
In: Plant Systematics and Evolution 290 (1-4), p. 21–45. DOI: 10.1007/s00606-010-0346-6.

König, Julia; Siwakoti, Mohan; Hilger, Hartmut H.; Weigend, Maximilian (2015):
**A Revision of the Genus *Cynoglossum* L. (Boraginaceae Juss.) in Nepal and Notes on the widespread Asian Species.**
In: Phytotaxa 224 (1), p. 1. DOI: 10.11646/phytotaxa.224.1.1.

Långström, Elisabeth; Chase, Mark W. (2002):
**Tribes of Boraginoideae (Boraginaceae) and Placement of *Antiphytum, Echiochilon, Ogastemma* and *Sericostoma*: A phylogenetic Analysis based on atpB Plastid DNA Sequence Data.**
In: Plant Systematics and Evolution 234 (1), p. 137–153. DOI: 10.1007/s00606-002-0195-z.

**Långström, Elisabeth; Oxelman, Bengt (2003):**
**Phylogeny of *Echiochilon* (Echiochileae, Boraginaceae) based on ITS Sequences and Morphology.**
In: Taxon 52 (4), p. 725. DOI: 10.2307/3647347.

**Meudt, Heidi M.; Prebble, Jessica M.; Lehnebach, Carlos A. (2015):**
**Native New Zealand Forget-me-nots (*Myosotis*, Boraginaceae) comprise a Pleistocene Species Radiation with very low genetic Divergence.**
In: Plant Systematics and Evolution 301 (5), p. 1455–1471. DOI: 10.1007/s00606-014-1166-x.

**Moore, Michael J.; Jansen, Robert K. (2006):**
**Molecular Evidence for the Age, Origin, and evolutionary History of the American Desert Plant Genus *Tiquilia* (Boraginaceae).**
In: Molecular Phylogenetics and Evolution 39 (3), p. 668–687. DOI: 10.1016/j.ympev.2006.01.020.

**Moore, Michael J.; Tye, Alan; Jansen, Robert K. (2006):**
**Patterns of long-distance Dispersal in *Tiquilia* Subg. *Tiquilia* (Boraginaceae): Implications for the Origins of amphitropical disjuncts and Galapagos Islands endemics.**
In: American Journal of Botany 93 (8), p. 1163–1177. DOI: 10.3732/ajb.93.8.1163.

**Nazaire, Mare; Wang, Xiao-Quan; Hufford, Larry (2014):**
**Geographic Origins and Patterns of Radiation of *Mertensia* (Boraginaceae).**
In: American Journal of Botany 101 (1), p. 104–118. DOI: 10.3732/ajb.1300320.

**Otero, Ana; Jiménez-Mejías, Pedro; Valcárcel, Virginia; Vargas, Pablo (2014):**
**Molecular Phylogenetics and Morphology support two new Genera (*Memoremea* and *Nihon*) of Boraginaceae s.s.**
In: Phytotaxa 173 (4), p. 241. DOI: 10.11646/phytotaxa.173.4.1.

**Romeiras, Maria M.; Ascensão, Lia; Duarte, Maria Cristina; Diniz, Maria A.; Pais, Maria Salomé (2008):**
**Taxonomy of *Echium* (Boraginaceae) Species from Cape Verde Islands.**
In: Australian Systematic Botany 21 (1), p. 26. DOI: 10.1071/SB07016.

**Romeiras, Maria M.; Paulo, Octávio S.; Duarte, Maria Cristina; Pina-Martins, Francisco; Cotrim, M. Helena; Carine, Mark A.; Pais, Maria Salomé (2011):**
**Origin and Diversification of the Genus *Echium* (Boraginaceae) in the Cape Verde archipelago.**
In: Taxon 60 (5), p. 1375–1385.

**Selvi, Federico; Bigazzi, Massimo; Hilger, Hartmut H.; Papini, Alessio (2006):**
**Molecular Phylogeny, Morphology and taxonomic Re-Circumscription of the Generic Complex *Nonea/Elizaldia/Pulmonaria/Paraskevia* (Boraginaceae-Boragineae).**
In: Taxon 55 (4), p. 907. DOI: 10.2307/25065685.

**Selvi, Federico; Cecchi, Lorenzo; Coppi, Andrea (2009):**
**Phylogeny, Karyotype Evolution and Taxonomy of *Cerinthe* L. (Boraginaceae).**
In: Taxon 58 (4), p. 1307–1325.

**Selvi, Federico; Coppi, Andrea; Cecchi, Lorenzo (2011):**
**High epizoochorous specialization and low DNA Sequence Divergence in Mediterranean *Cynoglossum* (Boraginaceae): Evidence from Fruit traits and ITS Region.**
In: Taxon 60 (4), p. 969–985.

**Selvi, Federico; Papini, Alessio; Hilger, Hartmut H.; Bigazzi, Massimo; Nardi, E. (2004):**
**The phylogenetic Relationships of *Cynoglottis* (Boraginaceae- Boragineae) inferred from ITS, 5.8s and trnL Sequences.**
In: Plant Systematics and Evolution 246 (3-4). DOI: 10.1007/s00606-004-0151-1.

**Sutorý, Karel (2010):**
***Oncaglossum*, a new Genus of the Boraginaceae, Tribe Cynoglosseae, from Mexico.**
In: Novon: A Journal for Botanical Nomenclature 20 (4), p. 463–469. DOI: 10.3417/2008059.

**Sutorý, Karel (2016):**
**New Names in the '*Cynoglossum montanum* Group' (Boraginaceae) in the Mediterranean Area.**
In: Edinburgh Journal of Botany 73 (03), p. 265–275. DOI: 10.1017/S0960428616000123.

**Thomas, Daniel C.; Weigend, Maximilian; Hilger, Hartmut H. (2008):**
**Phylogeny and Systematics of *Lithodora* (Boraginaceae-Lithospermeae) and its affinities to the monotypic Genera *Mairetis, Halacsya* and *Paramoltkia* based on ITS1 and trnLUAA-sequence Data and Morphology.**
In: Taxon 57 (1), p. 79–97.

**Weigend, Maximilian; Gottschling, Marc; Hilger, Hartmut H.; Nürk, Nicolai M. (2010):**
**Five new Species of *Lithospermum* L. (Boraginaceae Tribe Lithospermeae) in Andean South America: Another Radiation in the Amotape-Huancabamba Zone.**
In: Taxon 59 (4), p. 1161–1179.

**Weigend, Maximilian; Gottschling, Marc; Selvi, Federico; Hilger, Hartmut H. (2009):**
**Marbleseeds are gromwells - Systematics and Evolution of *Lithospermum* and allies (Boraginaceae Tribe Lithospermeae) based on molecular and morphological Data.**
In: Molecular Phylogenetics and Evolution 52 (3), p. 755–768. DOI: 10.1016/j.ympev.2009.05.013.

**Weigend, Maximilian; Luebert, Federico; Gottschling, Marc; Couvreur, Thomas L.P.; Hilger, Hartmut H.; Miller, James p. (2014):**
**From Capsules to Nutlets - phylogenetic Relationships in the Boraginales.**
In: Cladistics 30 (5), p. 508–518. DOI: 10.1111/cla.12061.

**Weigend, Maximilian; Luebert, Federico; Selvi, Federico; Brokamp, Grischa; Hilger, Hartmut H. (2013):**
**Multiple Origins for Hound's tongues (*Cynoglossum* L.) and Navel seeds (*Omphalodes* Mill.) - the Phylogeny of the borage Family (Boraginaceae s.str.).**
In: Molecular Phylogenetics and Evolution 68 (3), p. 604–618. DOI: 10.1016/j.ympev.2013.04.009.

**Winkworth, Richard C.; Grau, Jürke; Robertson, Alastair W.; Lockhart, Peter J. (2002):**
**The Origins and Evolution of the Genus *Myosotis* L. (Boraginaceae).**
In: Molecular Phylogenetics and Evolution 24 (2), p. 180–193. DOI: 10.1016/S1055-7903(02)00210-5.

### Boraginaceae; Boraginales

**Luebert, Federico; Cecchi, Lorenzo; Frohlich, Michael W.; Gottschling, Marc; Guilliams, C. Matt; Hasenstab-Lehman, Kristen E. et al. (2016):**
**Familial Classification of the Boraginales.**
In: Taxon 65 (3), p. 502–522. DOI: 10.12705/653.5.

**Miller, James S.; Gottschling, Marc (2017):**
**Generic Transfers in Malagasy Boraginales.**
In: Candollea 72 (2), p. 329–332. DOI: 10.15553/c2017v722a9.

### Borthwickiaceae

SU, JUN-XIA; WANG, WEI; ZHANG, LI-BING; CHEN, ZHI-DUAN (2012):

**Phylogenetic Placement of two enigmatic Genera, *Borthwickia* and *Stixis*, based on molecular and Pollen Data, and the Description of a new Family of Brassicales, Borthwickiaceae.**

In: Taxon 61 (3), p. 601–611.

### Brassicaceae

ALEXANDER, PATRICK J.; WINDHAM, MICHAEL D.; GOVINDARAJULU, RAJANIKANTH; AL-SHEHBAZ, IHSAN A.; BAILEY, C. DONOVAN (2010):

**Molecular Phylogenetics and Taxonomy of the Genus *Thysanocarpus* (Brassicaceae).**

In: Systematic Botany 35 (3), p. 559–577. DOI: 10.1600/036364410792495926.

ALI, TAHIR; SCHMUKER, ANGELIKA; RUNGE, FABIAN; SOLOVYEVA, IRINA; NIGRELLI, LISA; PAULE, JURAJ ET AL. (2016):

**Morphology, Phylogeny, and Taxonomy of *Microthlaspi* (Brassicaceae: Coluteocarpeae) and related Genera.**

In: Taxon 65 (1), p. 79–98. DOI: 10.12705/651.6.

AL-SHEHBAZ, IHSAN A. (2010):

***Arabis mexicana* belongs to *Planodes* (Brassicaceae).**

In: Harvard Papers in Botany 15 (1), p. 137–138. DOI: 10.3100/025.015.0106.

AL-SHEHBAZ, IHSAN A. (2012):

**A generic and tribal Synopsis of the Brassicaceae (Cruciferae).**

In: Taxon 61 (5), p. 931–954.

AL-SHEHBAZ, IHSAN A. (2013):

***Clypeola* is united with *Alyssum* (Brassicaceae).**

In: Harvard Papers in Botany 18 (2), p. 125–128. DOI: 10.3100/025.018.0204.

AL-SHEHBAZ, IHSAN A. (2014):

**A Synopsis of the Genus *Noccaea* (Coluteocarpeae, Brassicaceae).**

In: Harvard Papers in Botany 19 (1), p. 25–51. DOI: 10.3100/hpib.v19iss1.2014.n3.

AL-SHEHBAZ, IHSAN A.; BEILSTEIN, MARK A.; KELLOGG, ELIZABETH A. (2006):

**Systematics and Phylogeny of the Brassicaceae (Cruciferae): an Overview.**

In: Plant Systematics and Evolution 259 (2-4), p. 89–120. DOI: 10.1007/s00606-006-0415-z.

AMARASINGHE, SHANIKA; WATSON-HAIGH, NATHAN S.; GILLIHAM, MATTHEW; ROY, STUART; BAUMANN, UTE (2016):

**The evolutionary Origin of Cipk16: A Gene involved in enhanced Salt Tolerance.**

In: Molecular Phylogenetics and Evolution 100, p. 135–147. DOI: 10.1016/j.ympev.2016.03.031.

ARIAS, TATIANA; PIRES, J. CHRIS (2012):

**A fully resolved Chloroplast Phylogeny of the *Brassica* crops and wild relatives (Brassicaceae: Brassiceae): Novel Clades and potential taxonomic Implications.**

In: Taxon 61 (5), p. 980–988.

BAILEY, C. DONOVAN; KOCH, MARCUS A.; MAYER, MICHAEL S.; MUMMENHOFF, KLAUS; O'KANE, STEVE L.; WARWICK, SUZANNE I. ET AL. (2006):

**Toward a global Phylogeny of the Brassicaceae.**

In: Molecular Biology and Evolution 23 (11), p. 2142–2160. DOI: 10.1093/molbev/msl087.

BECK, JAMES B.; AL-SHEHBAZ, IHSAN A.; O'KANE, STEVE L.; SCHAAL, BARBARA A. (2007):

**Further Insights into the Phylogeny of *Arabidopsis* (Brassicaceae) from nuclear Atmyb2 flanking Sequence.**

In: Molecular Phylogenetics and Evolution 42 (1), p. 122–130. DOI: 10.1016/j.ympev.2006.06.011.

**Beilstein, Mark A.; Al-Shehbaz, Ihsan A.; Kellogg, Elizabeth A. (2006):**
**Brassicaceae Phylogeny and Trichome Evolution.**
In: American Journal of Botany 93 (4), p. 607–619.

**Beilstein, Mark A.; Al-Shehbaz, Ihsan A.; Mathews, Sarah; Kellogg, Elizabeth A. (2008):**
**Brassicaceae Phylogeny inferred from Phytochrome A and ndhF Sequence Data: Tribes and Trichomes revisited.**
In: American Journal of Botany 95 (10), p. 1307–1327. DOI: 10.3732/ajb.0800065.

**Bleeker, Walter; Weber-Sparenberg, C.; Hurka, Herbert (2002):**
**Chloroplast DNA Variation and Biogeography in the Genus *Rorippa* Scop. (Brassicaceae).**
In: Plant Biology 4 (1), p. 104–111. DOI: 10.1055/s-2002-20442.

**Cai, Liming; Ma, Hong (2016):**
**Using nuclear Genes to reconstruct Angiosperm Phylogeny at the Species Level: A case Study with Brassicaceae Species.**
In: Journal of Systematics and Evolution 54 (4), p. 438–452. DOI: 10.1111/jse.12204.

**Cappa, Jennifer J.; Yetter, Crystal; Fakra, Sirine; Cappa, Patrick J.; DeTar, Rachael; Landes, Corbett et al. (2015):**
**Evolution of Selenium Hyperaccumulation in *Stanleya* (Brassicaceae) as inferred from Phylogeny, Physiology and X-ray microprobe Analysis.**
In: the new Phytologist 205 (2), p. 583–595. DOI: 10.1111/nph.13071.

**Carlsen, Tor; Bleeker, Walter; Hurka, Herbert; Elven, Reidar; Brochmann, Christian (2009):**
**Biogeography and Phylogeny of *Cardamine* (Brassicaceae) 1.**
In: Annals of the Missouri Botanical Garden 96 (2), p. 215–236. DOI: 10.3417/2007047.

**Carlsen, Tor; Elven, Reidar; Brochmann, Christian (2010):**
**The evolutionary History of Beringian *Smelowskia* (Brassicaceae) inferred from combined Microsatellite and DNA Sequence Data.**
In: Taxon 59 (2), p. 427–438.

**Chen, Sheng-Yun; Wu, Guili; Chen, Shi-Long; Ren, Jiawen; Qin, Dahe (2010):**
**Molecular Phylogeny and Biogeography of the narrow endemic *Coelonema* and affinitive *Draba* (Brassicaceae) based on two DNA Regions.**
In: Biochemical Systematics and Ecology 38 (4), p. 796–805. DOI: 10.1016/j.bse.2010.08.002.

**Couvreur, Thomas L.P.; Franzke, Andreas; Al-Shehbaz, Ihsan A.; Bakker, Freek T.; Koch, Marcus A.; Mummenhoff, Klaus (2010):**
**Molecular Phylogenetics, temporal Diversification, and Principles of Evolution in the mustard Family (Brassicaceae).**
In: Molecular Biology and Evolution 27 (1), p. 55–71. DOI: 10.1093/molbev/msp202.

**Doğan, Bekir; Çelik, Mustafa; Ünal, Murat; Sefali, Abdurrahman; Martin, Esra; Kaya, Ayla (2016):**
**Study of phylogenetic Relationship of Turkish Species of *Matthiola* (Brassicaceae) based on ISSR amplification.**
In: Turkish Journal of Botany 40, p. 130–136. DOI: 10.3906/bot-1412-34.

**Eschmann-Grupe, G.; Hurka, Herbert; Neuffer, Barbara (2003):**
**Species Relationships within *Diplotaxis* (Brassicaceae) and the phylogenetic Origin of *D. muralis*.**
In: Plant Systematics and Evolution 243 (1-2), p. 13–29. DOI: 10.1007/s00606-003-0047-5.

**Fırat, Mehmet; Özüdoğru, Barış; Tarikahya-Hacıoğlu, Burcu; Bülbül, Ali Savaş; Al-Shehbaz, Ihsan A.; Mummenhoff, Klaus (2014):**
**Phylogenetic Position and taxonomic Assignment of *Thlaspi aghricum* P.H.Davis & K.Tan (Brassicaceae).**
In: Phytotaxa 178 (4), p. 287. DOI: 10.11646/phytotaxa.178.4.2.

**Francisco-Ortega, Javier; Fuertes-Aguilar, Javier; Gómez-Campo, C.; Santos-Guerra, Arnoldo; Jansen, Robert K. (1999):**
**Internal transcribed spacer Sequence Phylogeny of *Crambe* L. (Brassicaceae): molecular Data reveal two Old World Disjunctions.**
In: Molecular Phylogenetics and Evolution 11 (3), p. 361–380. DOI: 10.1006/mpev.1998.0592.

**Francisco-Ortega, Javier; Fuertes-Aguilar, Javier; Kim, Seung-Chul; Santos-Guerra, Arnoldo; Crawford, Daniel J.; Jansen, Robert K. (2002):**
**Phylogeny of the Macaronesian endemic *Crambe* Section *Dendrocrambe* (Brassicaceae) based on internal transcribed Spacer Sequences of nuclear ribosomal DNA.**
In: American Journal of Botany 89 (12), p. 1984–1990. DOI: 10.3732/ajb.89.12.1984.

**Franzke, Andreas; German, Dmitry A.; Al-Shehbaz, Ihsan A.; Mummenhoff, Klaus (2009):**
**Arabidopsis Family ties: molecular Phylogeny and age estimates in Brassicaceae.**
In: Taxon 58 (2), p. 425–437.

**Franzke, Andreas; Lysak, Martin A.; Al-Shehbaz, Ihsan A.; Koch, Marcus A.; Mummenhoff, Klaus (2011):**
**Cabbage Family affairs: the evolutionary History of Brassicaceae.**
In: Trends in Plant Science 16 (2), p. 108–116. DOI: 10.1016/j.tplants.2010.11.005.

**German, Dmitry A. (2016):**
**Taxonomic Notes on miscellaneous Cruciferae.**
In: Turczaninowia 19 (4), p. 129–135. DOI: 10.14258/turczaninowia.19.4.17.

**German, Dmitry A.; Al-Shehbaz, Ihsan A. (2008):**
**Five Additional Tribes (Aphragmeae, Biscutelleae, Calepineae, Conringieae, and Erysimeae) in the Brassicaceae (Cruciferae).**
In: Harvard Papers in Botany 13 (1), p. 165–170. DOI: 10.3100/1043-4534(2008)13[165:FATABC]2.0.CO;2.

**German, Dmitry A.; Friesen, Nikolai; Neuffer, Barbara; Al-Shehbaz, Ihsan A.; Hurka, Herbert (2009):**
**Contribution to ITS Phylogeny of the Brassicaceae, with special Reference to some Asian taxa.**
In: Plant Systematics and Evolution 283 (1-2), p. 33–56. DOI: 10.1007/s00606-009-0213-5.

**German, Dmitry A.; Grant, Jason R.; Lysak, Martin A.; Al-Shehbaz, Ihsan A. (2011):**
**Molecular Phylogeny and Systematics of the Tribe Chorisporeae (Brassicaceae).**
In: Plant Systematics and Evolution 294 (1-2), p. 65–86. DOI: 10.1007/s00606-011-0452-0.

**Goodson, Barbara E.; Rehman, Sumaiyah K.; Jansen, Robert K. (2011):**
**Molecular Systematics and Biogeography of *Descurainia* (Brassicaceae) based on Nuclear ITS and Non-Coding Chloroplast DNA.**
In: Systematic Botany 36 (4), p. 957–980. DOI: 10.1600/036364411X604976.

**Goodson, Barbara E.; Santos-Guerra, Arnoldo; Jansen, Robert K. (2006):**
**Molecular Systematics of *Descurainia* (Brassicaceae) in the Canary Islands: Biogeographic and taxonomic Implications.**
In: Taxon 55 (3), p. 671–682. DOI: 10.2307/25065643.

**Grundt, Hanne Hegre; Popp, Magnus; Brochmann, Christian; Oxelman, Bengt (2004):**

**Polyploid Origins in a circumpolar complex in *Draba* (Brassicaceae) inferred from cloned nuclear DNA Sequences and Fingerprints.**
In: Molecular Phylogenetics and Evolution 32 (3), p. 695–710. DOI: 10.1016/j.ympev.2004.04.006.

**Hall, Jocelyn C.; Tisdale, Tracy E.; Donohue, Kathleen; Wheeler, Andrew; Al-Yahya, Mohammed A.; Kramer, Elena M. (2011):**
**Convergent Evolution of a complex Fruit Structure in the Tribe Brassiceae (Brassicaceae).**
In: American Journal of Botany 98 (12), p. 1989–2003. DOI: 10.3732/ajb.1100203.

**Hao, Guoqian; Al-Shehbaz, Ihsan A.; Ahani, Hamid; Liang, Qianlong; Mao, Kang-Shan; Wang, Qian; Liu, Jian-Quan (2017):**
**An integrative Study of evolutionary Diversification of *Eutrema* (Eutremeae, Brassicaceae).**
In: Botanical Journal of the Linnean Society 184 (2), p. 204–223. DOI: 10.1093/botlinnean/box024.

**Heenan, Peter B.; Goeke, Dagmar F.; Houliston, Gary J.; Lysak, Martin A. (2012):**
**Phylogenetic Analyses of ITS and rbcL DNA Sequences for sixteen Genera of Australian and New Zealand Brassicaceae result in the Expansion of the Tribe Microlepidieae.**
In: Taxon 61 (5), p. 970–979.

**Heenan, Peter B.; Mitchell, Anthony D.; Koch, Marcus A. (2002):**
**Molecular Systematics of the New Zealand *Pachycladon* (Brassicaceae) complex: Generic Circumscription and Relationships to *Arabidopsis* sens. lat. and *Arabis* sens. lat.**
In: New Zealand Journal of Botany 40 (4), p. 543–562. DOI: 10.1080/0028825X.2002.9512815.

**Hu, Huan; Al-Shehbaz, Ihsan A.; Sun, Yongshuai; Hao, Guoqian; Wang, Qian; Liu, Jian-Quan (2015):**
**Species Delimitation in *Orychophragmus* (Brassicaceae) based on Chloroplast and nuclear DNA Barcodes.**
In: Taxon 64 (4), p. 714–726. DOI: 10.12705/644.4.

**Huang, Chien-Hsun; Sun, Renran; Hu, Yi; Zeng, Liping; Zhang, Ning; Cai, Liming et al. (2016):**
**Resolution of Brassicaceae Phylogeny using nuclear Genes uncovers nested Radiations and supports convergent morphological Evolution.**
In: Molecular Biology and Evolution 33 (2), p. 394–412. DOI: 10.1093/molbev/msv226.

**Ivalú Cacho, N.; Millie Burrell, A.; Pepper, Alan E.; Strauss, Sharon Y. (2014):**
**Novel nuclear Markers inform the Systematics and the Evolution of Serpentine Use in *Streptanthus* and allies (Thelypodieae, Brassicaceae).**
In: Molecular Phylogenetics and Evolution 72, p. 71–81. DOI: 10.1016/j.ympev.2013.11.018.

**Jaén-Molina, Ruth; Caujapé-Castells, Juli; Reyes-Betancort, Jorge Alfredo; Akhani, Hossein; Fernández-Palacios, Olga; Paz, Julia Pérez et al. (2009):**
**The molecular Phylogeny of *Matthiola* R. Br. (Brassicaceae) inferred from ITS Sequences, with special Emphasis on the Macaronesian Endemics.**
In: Molecular Phylogenetics and Evolution 53 (3), p. 972–981. DOI: 10.1016/j.ympev.2009.08.031.

**Joly, Simon; Heenan, Peter B.; Lockhart, Peter J. (2009):**
**A Pleistocene inter-tribal allopolyploidization event precedes the Species Radiation of *Pachycladon* (Brassicaceae) in New Zealand.**
In: Molecular Phylogenetics and Evolution 51 (2), p. 365–372. DOI: 10.1016/j.ympev.2009.02.015.

**Jordon-Thaden, Ingrid; Hase, Irina; Al-Shehbaz, Ihsan A.; Koch, Marcus A. (2010):**
**Molecular Phylogeny and Systematics of the Genus *Draba* (Brassicaceae) and Identification of its most closely related Genera.**
In: Molecular Phylogenetics and Evolution 55 (2), p. 524–540. DOI: 10.1016/j.ympev.2010.02.012.

**Karl, Robert; Koch, Marcus A. (2013):**
**A world-wide Perspective on crucifer Speciation and Evolution: Phylogenetics, Biogeography and trait Evolution in Tribe Arabideae.**
In: Annals of Botany 112 (6), p. 983–1001. DOI: 10.1093/aob/mct165.

**Khosravi, Ahmad Reza; Jacquemoud, Fernand; Mohsenzadeh, Sasan; Menke, Marck; Mummenhoff, Klaus (2009):**
**Phylogenetic Position and taxonomic Classification of *Aethionema trinervium* (Brassicaceae): A Morphologically variable Subshrub from Southwestern Asia.**
In: Annals of the Missouri Botanical Garden 96 (4), p. 564–574. DOI: 10.3417/2007004.

**Kiefer, Christiane; Dobes, Christoph; Koch, Marcus A. (2009):**
***Boechera* or not? Phylogeny and Phylogeography of eastern North American *Boechera* Species (Brassicaceae).**
In: Taxon 58 (4), p. 1109–1121.

**Kiefer, Christiane; Dobes, Christoph; Sharbel, Timothy F.; Koch, Marcus A. (2009):**
**Phylogeographic Structure of the Chloroplast DNA Gene Pool in North American *Boechera* - a Genus and continental-wide Perspective.**
In: Molecular Phylogenetics and Evolution 52 (2), p. 303–311. DOI: 10.1016/j.ympev.2009.03.016.

**Koch, Marcus A. (2012):**
**Mid-Miocene Divergence of *Ionopsidium* and *Cochlearia* and its Impact on the Systematics and Biogeography of the Tribe Cochlearieae (Brassicaceae).**
In: Taxon 61 (1), p. 76–92.

**Koch, Marcus A.; Karl, Robert; German, Dmitry A.; Al-Shehbaz, Ihsan A. (2012):**
**Systematics, Taxonomy and Biogeography of three new Asian Genera of Brassicaceae Tribe Arabideae: An ancient Distribution circle around the Asian High Mountains.**
In: Taxon 61 (5), p. 955–969.

**Koch, Marcus A.; Mummenhoff, Klaus; Hurka, Herbert (1999):**
**Molecular Phylogenetics of *Cochlearia* (Brassicaceae) and allied Genera based on nuclear ribosomal ITS DNA Sequence Analysis contradict traditional Concepts of their evolutionary Relationship.**
In: Plant Systematics and Evolution 216 (3-4), p. 207–230. DOI: 10.1007/BF01084399.

**Kučera, Jaromír; Marhold, Karol; Lihová, Judita (2010):**
***Cardamine maritima* Group (Brassicaceae) in the amphi-Adriatic Area: A Hotspot of Species Diversity revealed by DNA Sequences and morphological Variation.**
In: Taxon 59 (1), p. 148–164.

**Li, Yan; Feng, Ying; Lv, Guanghui; Liu, Bing; Qi, Aladaer (2015):**
**The Phylogeny of *Alyssum* (Brassicaceae) inferred from molecular Data.**
In: Nordic Journal of Botany 33 (6), p. 715–721. DOI: 10.1111/njb.00588.

**Lihová, Judita; Fuertes-Aguilar, Javier; Marhold, Karol; Nieto Feliner, Gonzalo (2004):**
**Origin of the disjunct tetraploid *Cardamine amporitana* (Brassicaceae) assessed with nuclear and Chloroplast DNA Sequence Data.**

In: American Journal of Botany 91 (8), p. 1231–1242. DOI: 10.3732/ajb.91.8.1231.

LIHOVÁ, JUDITA; MARHOLD, KAROL; KUDOH, HIROSHI; KOCH, MARCUS A. (2006):
**Worldwide Phylogeny and Biogeography of *Cardamine flexuosa* (Brassicaceae) and its relatives.**
In: American Journal of Botany 93 (8), p. 1206–1221. DOI: 10.3732/ajb.93.8.1206.

LIU, LEI; ZHAO, BO; TAN, DUN-YAN; WANG, JIANBO (2011):
**Phylogenetic Relationships of Brassicaceae in China: Insights from a non-coding Chloroplast, mitochondrial, and nuclear DNA Data set.**
In: Biochemical Systematics and Ecology 39 (4-6), p. 600–608. DOI: 10.1016/j.bse.2011.05.003.

MANDÁKOVÁ, TEREZIE; MUMMENHOFF, KLAUS; AL-SHEHBAZ, IHSAN A.; MUCINA, LADISLAV; LYSAK, ANDREAS A.; MÜHLHAUSEN, MARTIN (2012):
**Whole-Genome triplication and Species Radiation in the southern African Tribe Heliophileae (Brassicaceae).**
In: Taxon 61 (5), p. 989–1000.

MAYER, MICHAEL S.; BESEDA, LINDSAY (2010):
**Reconciling Taxonomy and Phylogeny in the *Streptanthus glandulosus* Complex (Brassicaceae).**
In: Annals of the Missouri Botanical Garden 97 (1), p. 106–116. DOI: 10.3417/2007010.

MAZIE, ABIGAIL R.; BAUM, DAVID A. (2016):
**Clade-specific positive selection on a developmental Gene: Branchless Trichome and the Evolution of stellate Trichomes in *Physaria* (Brassicaceae).**
In: Molecular Phylogenetics and Evolution 100, p. 31–40. DOI: 10.1016/j.ympev.2016.03.027.

MITCHELL, ANTHONY D.; HEENAN, PETER B. (2000):
**Systematic Relationships of New Zealand Endemic Brassicaceae inferred from nrDNA ITS Sequence Data.**
In: Systematic Botany 25 (1), p. 98–105. DOI: 10.2307/2666676.

MOAZZENI, HAMID; ZARRE, SHAHIN; AL-SHEHBAZ, IHSAN A.; MUMMENHOFF, KLAUS (2010):
**Phylogeny of *Isatis* (Brassicaceae) and allied Genera based on ITS Sequences of nuclear ribosomal DNA and morphological Characters.**
In: Flora - Morphology, Distribution, Functional Ecology of Plants 205 (5), p. 337–343. DOI: 10.1016/j.flora.2009.12.028.

MOAZZENI, HAMID; ZARRE, SHAHIN; PFEIL, BERNARD E.; BERTRAND, YANN J.K.; GERMAN, DMITRY A.; AL-SHEHBAZ, IHSAN A. ET AL. (2014):
**Phylogenetic Perspectives on Diversification and Character Evolution in the species-rich Genus *Erysimum* (Erysimeae; Brassicaceae) based on a densely sampled ITS approach.**
In: Botanical Journal of the Linnean Society 175 (4), p. 497–522. DOI: 10.1111/boj.12184.

MUMMENHOFF, KLAUS; BRÜGGEMANN, HOLGER; BOWMAN, JOHN L. (2001):
**Chloroplast DNA Phylogeny and Biogeography of *Lepidium* (Brassicaceae).**
In: American Journal of Botany 88 (11), p. 2051–2063. DOI: 10.2307/3558431.

MUMMENHOFF, KLAUS; LINDER, HANS PETER; FRIESEN, NIKOLAI; BOWMAN, JOHN L.; LEE, JI-YOUNG; FRANZKE, ANDREAS (2004):
**Molecular Evidence for bicontinental hybridogenous genomic Constitution in *Lepidium* sensu stricto (Brassicaceae) Species from Australia and New Zealand.**
In: American Journal of Botany 91 (2), p. 254–261. DOI: 10.3732/ajb.91.2.254.

**Nagpal, Ranjana; Dar, Tanvir H.; Raina, Soom Nath (2008):**
**Molecular Systematics of *Brassica* and allied Genera in Subtribes Brassicinae, Raphaninae, Moricandiinae, and Cakilinae (Brassicaceae, Tribe Brassiceae); the Organization and Evolution of ribosomal Gene Families.**
In: Botanical Journal of the Linnean Society 157 (3), p. 545–557. DOI: 10.1111/j.1095-8339.2008.00812.x.

**Ometto, Lino; Li, Mingai; Bresadola, Luisa; Varotto, Claudio (2012):**
**Rates of Evolution in stress-related Genes are associated with Habitat PReference in two Cardamine Lineages.**
In: BMC Evolutionary Biology 12, p. 7. DOI: 10.1186/1471-2148-12-7.

**Özüdoğru, Barış; Akaydin, Galip; Erik, Sadik; Al-Shehbaz, Ihsan A.; Mummenhoff, Klaus (2015):**
**Phylogeny, Diversification and biogeographic Implications of the eastern Mediterranean endemic Genus *Ricotia* (Brassicaceae).**
In: Taxon 64 (4), p. 727–740. DOI: 10.12705/644.5.

**Parolly, Gerald; Nordt, Birgit; Bleeker, Walter; Mummenhoff, Klaus (2010):**
***Heldreichia* Boiss. (Brassicaceae) revisited: A morphological and molecular Study.**
In: Taxon 59 (1), p. 187–202.

**Rešetnik, Ivana; Satović, Zlatko; Schneeweiss, Gerald M.; Liber, Zlatko (2013):**
**Phylogenetic Relationships in Brassicaceae Tribe Alysseae inferred from nuclear ribosomal and Chloroplast DNA Sequence Data.**
In: Molecular Phylogenetics and Evolution 69 (3), p. 772–786. DOI: 10.1016/j.ympev.2013.06.026.

**Reza Khosravi, Ahmad; Mohsenzadeh, Sasan; Mummenhoff, Klaus (2008):**
**Analysis of the phylogenetic Position of *Acanthocardamum erinaceum* (Brassicaceae) based on ITS-sequences shows that it should be transferred to *Aethionema* as *A. erinaceum*.**
In: Nordic Journal of Botany 26 (1-2), p. 25–30. DOI: 10.1111/j.0107-055X.2008.00205.x.

**Reza Khosravi, Ahmad; Mohsenzadeh, Sasan; Mummenhoff, Klaus (2008):**
**Phylogenetic Position of *Brossardia papyracea* (Brassicaceae) based on Sequences of nuclear ribosomal DNA.**
In: Feddes Repertorium 119 (1-2), p. 13–23. DOI: 10.1002/fedr.200811146.

**Salariato, Diego Leonel; Zuloaga, Fernando Omar (2017):**
**Climatic niche Evolution in the Andean Genus *Menonvillea* (Cremolobeae: Brassicaceae).**
In: Organisms Diversity and Evolution 17 (1), p. 11–28. DOI: 10.1007/s13127-016-0291-5.

**Salariato, Diego Leonel; Zuloaga, Fernando Omar; Al-Shehbaz, Ihsan A. (2014):**
**A Revision of the Genus *Menonvillea* (Cremolobeae, Brassicaceae).**
In: Phytotaxa 162 (5), p. 241. DOI: 10.11646/phytotaxa.162.5.1.

**Salariato, Diego Leonel; Zuloaga, Fernando Omar; Al-Shehbaz, Ihsan A. (2015):**
**A taxonomic Revision of the Genus *Xerodraba* (Eudemeae, Brassicaceae).**
In: Phytotaxa 207 (1), p. 39. DOI: 10.11646/phytotaxa.207.1.2.

**Salariato, Diego Leonel; Zuloaga, Fernando Omar; Al-Shehbaz, Ihsan A. (2013):**
**Molecular Phylogeny of *Menonvillea* and Recognition of the new Genus *Aimara* (Brassicaceae: Cremolobeae).**
In: Taxon 62 (6), p. 1220–1234. DOI: 10.12705/626.6.

**Salariato, Diego Leonel; Zuloaga, Fernando Omar; Al-Shehbaz, Ihsan A. (2013):**

**Revision and tribal Placement of the Argentinean Genus *Parodiodoxa* (Brassicaceae).**
In: Plant Systematics and Evolution 299 (2), p. 305–316. DOI: 10.1007/s00606-012-0722-5.

**Salariato, Diego Leonel; Zuloaga, Fernando Omar; Cano, Asunción; Al-Shehbaz, Ihsan A. (2015):**
**Molecular Phylogenetics of Tribe Eudemeae (Brassicaceae) and Implications for its Morphology and Distribution.**
In: Molecular Phylogenetics and Evolution 82 Pt A, p. 43–59. DOI: 10.1016/j.ympev.2014.09.030.

**Salariato, Diego Leonel; Zuloaga, Fernando Omar; Franzke, Andreas; Mummenhoff, Klaus; Al-Shehbaz, Ihsan A. (2016):**
**Diversification Patterns in the Ces-Clade (Brassicaceae Tribes Cremolobeae, Eudemeae, Schizopetaleae) in Andean South America.**
In: Botanical Journal of the Linnean Society 181, p. 543–566.

**Schulte, Katharina; Horres, Ralf; Zizka, Georg (2005):**
**Molecular Phylogeny of Bromelioideae and its Implications on Biogeography and the Evolution of CAM in the Family.**
In: Senckenbergiana Biologica 85, p. 1–14.

**Simón-Porcar, Violeta I.; Pérez-Collazos, Ernesto; Catalán, Pilar (2015):**
**Phylogeny and Systematics of the western Mediterranean *Vella pseudocytisus-V. aspera* complex (Brassicaceae).**
In: Turkish Journal of Botany 39, p. 472–486. DOI: 10.3906/bot-1406-87.

**Slotte, T.; Ceplitis, A.; Neuffer, Barbara; Hurka, Herbert; Lascoux, Martin (2006):**
**Intrageneric Phylogeny of *Capsella* (Brassicaceae) and the Origin of the tetraploid *C. bursa-pastoris* based on Chloroplast and nuclear DNA Sequences.**
In: American Journal of Botany 93 (11), p. 1714–1724. DOI: 10.3732/ajb.93.11.1714.

**Španiel, Stanislav; Kempa, Matúš; Salmerón-Sánchez, Esteban; Fuertes-Aguilar, Javier; Mota, Juan F.; Al-Shehbaz, Ihsan A. et al. (2015):**
**AlyBase: Database of Names, Chromosome Numbers, and ploidy Levels of Alysseae (Brassicaceae), with a new generic Concept of the Tribe.**
In: Plant Systematics and Evolution 301 (10), p. 2463–2491. DOI: 10.1007/s00606-015-1257-3.

**Španiel, Stanislav; Zozomová-Lihová, Judita; Marhold, Karol (2017):**
**Revised taxonomic Treatment of the *Alyssum montanum-A. repens* complex in the Balkans: a multivariate morphometric Analysis.**
In: Plant Systematics and Evolution 303 (10), p. 1413–1442. DOI: 10.1007/s00606-017-1468-x.

**Toro-Núñez, Oscar; Al-Shehbaz, Ihsan A.; Mort, Mark E. (2015):**
**Phylogenetic Study with nuclear and Chloroplast Data and ecological niche reveals *Atacama* (Brassicaceae), a new monotypic Genus endemic from the Andes of the Atacama Desert, Chile.**
In: Plant Systematics and Evolution 301 (5), p. 1377–1396. DOI: 10.1007/s00606-014-1157-y.

**Toro-Núñez, Oscar; Mort, Mark E.; Ruiz-Ponce, Eduardo; Al-Shehbaz, Ihsan A. (2013):**
**Phylogenetic Relationships of *Mathewsia* and *Schizopetalon* (Brassicaceae) inferred from nrDNA and cpDNA Regions: taxonomic and evolutionary Insights from an Atacama Desert endemic Lineage.**
In: Taxon 62 (2), p. 343–356. DOI: 10.12705/622.4.

**Warwick, Suzanne I.; Al-Shehbaz, Ihsan A.; Price, Robert A.; Sauder, Connie A. (2002):**

**Phylogeny of *Sisymbrium* (Brassicaceae) based on ITS Sequences of nuclear ribosomal DNA.**
In: Canadian Journal of Botany 80 (9), p. 1002–1017. DOI: 10.1139/B02-089.

WARWICK, SUZANNE I.; AL-SHEHBAZ, IHSAN A.; SAUDER, CONNIE A.; HARRIS, JAMES G.; KOCH, MARCUS A. (2004):
**Phylogeny of *Braya* and *Neotorularia* (Brassicaceae) based on nuclear ribosomal Internal Transcribed Spacer and Chloroplast trn L Intron Sequences.**
In: Canadian Journal of Botany 82 (3), p. 376–392. DOI: 10.1139/B04-012.

WARWICK, SUZANNE I.; MUMMENHOFF, KLAUS; SAUDER, CONNIE A.; KOCH, MARCUS A.; AL-SHEHBAZ, IHSAN A. (2010):
**Closing the gaps: phylogenetic Relationships in the Brassicaceae based on DNA Sequence Data of nuclear ribosomal ITS Region.**
In: Plant Systematics and Evolution 285 (3-4), p. 209–232. DOI: 10.1007/s00606-010-0271-8.

WARWICK, SUZANNE I.; SAUDER, CONNIE A. (2005):
**Phylogeny of Tribe Brassiceae (Brassicaceae) based on Chloroplast restriction site polymorphisms and nuclear ribosomal Internal Transcribed Spacer and Chloroplast trn L Intron Sequences.**
In: Canadian Journal of Botany 83 (5), p. 467–483. DOI: 10.1139/B05-021.

WARWICK, SUZANNE I.; SAUDER, CONNIE A.; AL-SHEHBAZ, IHSAN A. (2008):
**Phylogenetic Relationships in the Tribe Alysseae (Brassicaceae) based on nuclear ribosomal ITS DNA Sequences.**
In: Botany 86 (4), p. 315–336. DOI: 10.1139/B08-013.

WARWICK, SUZANNE I.; SAUDER, CONNIE A.; AL-SHEHBAZ, IHSAN A. (2011):
**Systematic Position of *Ivania, Scoliaxon,* and *Phravenia* (Brassicaceae).**
In: Taxon 60 (4), p. 1156–1164.

WARWICK, SUZANNE I.; SAUDER, CONNIE A.; AL-SHEHBAZ, IHSAN A.; JACQUEMOUD, FERNAND (2007):
**Phylogenetic Relationships in the Tribes Anchonieae, Chorisporeae, Euclidieae, and Hesperideae (Brassicaceae) based on nuclear Ribosomal ITS DNA Sequences.**
In: Annals of the Missouri Botanical Garden 94 (1), p. 56–78. DOI: 10.3417/0026-6493(2007)94[56:PRITTA]2.0.CO;2.

WARWICK, SUZANNE I.; SAUDER, CONNIE A.; MAYER, MICHAEL S.; AL-SHEHBAZ, IHSAN A. (2009):
**Phylogenetic Relationships in the Tribes Schizopetaleae and Thelypodieae (Brassicaceae) based on nuclear ribosomal ITS Region and Plastid ndh F DNA Sequences.**
In: Botany 87 (10), p. 961–985. DOI: 10.1139/B09-051.

YU, QIU-SHI; WANG, QIAN; WANG, AI-LAN; WU, GUI-LI; LIU, JIAN-QUAN (2010):
**Interspecific Delimitation and phylogenetic Origin of *Pugionium* (Brassicaceae).**
In: Journal of Systematics and Evolution 48 (3), p. 195–206. DOI: 10.1111/j.1759-6831.2010.00078.x.

YUE, JI-PEI; SUN, HANG; BAUM, DAVID A.; LI, JIAN-HUA; AL-SHEHBAZ, IHSAN A.; REE, RICHARD H. (2009):
**Molecular Phylogeny of *Solms-laubachia* (Brassicaceae) s.l., based on multiple nuclear and Plastid DNA Sequences, and its biogeographic Implications.**
In: Journal of Systematics and Evolution 47 (5), p. 402–415. DOI: 10.1111/j.1759-6831.2009.00041.x.

YUE, JI-PEI; SUN, HANG; LI, JIAN-HUA; AL-SHEHBAZ, IHSAN A. (2008):
**A Synopsis of an expanded *Solms-laubachia* (Brassicaceae), and the Description of four new Species from Western China.**
In: Annals of the Missouri Botanical Garden 95 (3), p. 520–538. DOI: 10.3417/2006214.

ZHOU, YING-YING; ZHANG, HONG-WEI; HU, JIANG-QIN; JIN, XIAO-FENG (2014):
**_Sinalliaria_, a new Genus of Brassicaceae from eastern China, based on morphological and molecular Data.**
In: Phytotaxa 186 (4), p. 188. DOI: 10.11646/phytotaxa.186.4.2.

ZIFFER-BERGER, J.; HANIN, N.; FOGEL, T.; MUMMENHOFF, KLAUS; BARAZANI, O. (2015):
**Molecular Phylogeny indicates Polyphyly in _Raphanus_ L. (Brassicaceae).**
In: Edinburgh Journal of Botany 72 (1), p. 1–11. DOI: 10.1017/S0960428614000286.

ZOZOMOVÁ-LIHOVÁ, JUDITA; MARHOLD, KAROL; ŠPANIEL, STANISLAV (2014):
**Taxonomy and evolutionary History of _Alyssum montanum_ (Brassicaceae) and related Taxa in southwestern Europe and Morocco: Diversification driven by polyploidy, geographic and ecological Isolation.**
In: Taxon 63 (3), p. 562–591. DOI: 10.12705/633.18.

### Bromeliaceae

AGUIRRE-SANTORO, JULIÁN; MICHELANGELI, FABIÁN ARMANDO; STEVENSON, DENNIS WM. (2016):
**Molecular Phylogenetics of the _Ronnbergia_ Alliance (Bromeliaceae, Bromelioideae) and Insights into their morphological Evolution.**
In: Molecular Phylogenetics and Evolution 100, p. 1–20. DOI: 10.1016/j.ympev.2016.04.007.

ALMEIDA, VALQUÍRIA REZENDE; FERREIRA DA COSTA, ANDREA; MANTOVANI, ANDRÉ; GONÇALVES-ESTEVES, VÂNIA; OLIVEIRA, ROSANI DO CARMO; FORZZA, RAFAELA CAMPOSTRINI (2009):
**Morphological Phylogenetics of _Quesnelia_ (Bromeliaceae, Bromelioideae).**
In: Systematic Botany 34 (4), p. 660–672. DOI: 10.1600/036364409790139619.

BARFUSS, MICHAEL H. J.; SAMUEL, ROSABELLE; TILL, WALTER; STUESSY, TOD F. (2005):
**Phylogenetic Relationships in Subfamily Tillandsioideae (Bromeliaceae) based on DNA Sequence Data from seven Plastid Regions.**
In: American Journal of Botany 92 (2), p. 337–351. DOI: 10.3732/ajb.92.2.337.

BARFUSS, MICHAEL H. J.; TILL, WALTER; LEME, ELTON M.C.; PINZÓN, JUAN P.; MANZANARES, JOSÉ M.; HALBRITTER, HEIDEMARIE ET AL. (2016):
**Taxonomic Revision of Bromeliaceae Subfam. Tillandsioideae based on a multi-locus DNA Sequence Phylogeny and Morphology.**
In: Phytotaxa 279 (1), p. 1. DOI: 10.11646/phytotaxa.279.1.1.

CASTELLO, LUCÍA V.; BARFUSS, MICHAEL H. J.; TILL, WALTER; GALETTO, LEONARDO; CHIAPELLA, JORGE O. (2016):
**Disentangling the _Tillandsia capillaris_ complex: phylogenetic Relationships and Taxon boundaries in Andean populations.**
In: Botanical Journal of the Linnean Society 181 (3), p. 391–414. DOI: 10.1111/boj.12400.

CHEW, TANIA; LUNA, EFRAÍN DE; GONZÁLEZ, DOLORES (2010):
**Phylogenetic Relationships of the Pseudobulbous _Tillandsia_ Species (Bromeliaceae) inferred from Cladistic Analyses of ITS 2, 5.8s Ribosomal RNA Gene, and ETS Sequences.**
In: Systematic Botany 35 (1), p. 86–95. DOI: 10.1600/036364410790862632.

COSTA, ANDREA FERREIRA DA; GOMES-DA-SILVA, JANAÍNA; WANDERLEY, MARIA DAS GRAÇAS LAPA (2015):
**_Vriesea_ (Bromeliaceae, Tillandsioideae): a cladistic Analysis of eastern Brazilian Species based on morphological Characters.**
In: Rodriguésia 66 (2), p. 429–440. DOI: 10.1590/2175-7860201566211.

**Cruz, Geyner A. S.; Zizka, Georg; Silvestro, Daniele; Leme, Elton M.C.; Schulte, Katharina; Benko-Iseppon, Ana Maria (2017):**
**Molecular Phylogeny, Character Evolution and historical Biogeography of *Cryptanthus* Otto & A. Dietr. (Bromeliaceae).**
In: Molecular Phylogenetics and Evolution 107, p. 152–165. DOI: 10.1016/j.ympev.2016.10.019.

**Donadío, Sabina; Pozner, Raúl E.; Giussani, Liliana M. (2015):**
**Phylogenetic Relationships within *Tillandsia* Subgenus *Diaphoranthema* (Bromeliaceae, Tillandsioideae) based on a comprehensive morphological Dataset.**
In: Plant Systematics and Evolution 301 (1), p. 387–410. DOI: 10.1007/s00606-014-1081-1.

**Gitaí, Jailson; Horres, Ralf; Benko-Iseppon, Ana Maria (2005):**
**Chromosomal features and Evolution of Bromeliaceae.**
In: Plant Systematics and Evolution 253 (1-4), p. 65–80. DOI: 10.1007/s00606-005-0306-8.

**Gitaí, Jailson; Paule, Juraj; Zizka, Georg; Schulte, Katharina; Benko-Iseppon, Ana Maria (2014):**
**Chromosome Numbers and DNA content in Bromeliaceae: additional Data and critical review.**
In: Botanical Journal of the Linnean Society 176 (3), p. 349–368. DOI: 10.1111/boj.12211.

**Givnish, Thomas J.; Barfuss, Michael H. J.; van Ee, Benjamin W.; Riina, Ricarda; Schulte, Katharina; Horres, Ralf et al. (2014):**
**Adaptive Radiation, correlated and contingent Evolution, and net Species Diversification in Bromeliaceae.**
In: Molecular Phylogenetics and Evolution 71, p. 55–78. DOI: 10.1016/j.ympev.2013.10.010.

**Givnish, Thomas J.; Barfuss, Michael H. J.; van Ee, Benjamin W.; Riina, Ricarda; Schulte, Katharina; Horres, Ralf et al. (2011):**
**Phylogeny, adaptive Radiation, and historical Biogeography in Bromeliaceae: Insights from an eight-locus Plastid Phylogeny.**
In: American Journal of Botany 98 (5), p. 872–895. DOI: 10.3732/ajb.1000059.

**Goetze, Márcia; Schulte, Katharina; Palma-silva, Clarisse; Zanella, Camila M.; Büttow, Miriam V.; Capra, Fernanda; Bered, Fernanda (2016):**
**Diversification of Bromelioideae (Bromeliaceae) in the Brazilian Atlantic Rainforest: A case Study in *Aechmea* Subgenus *Ortgiesia*.**
In: Molecular Phylogenetics and Evolution 98, p. 346–357. DOI: 10.1016/j.ympev.2016.03.001.

**Gomes-da-Silva, Janaína; Souza-Chies, Tatiana Teixeira de (2018):**
**What actually is *Vriesea*? A total Evidence approach in a polyphyletic Genus of Tillandsioideae (Bromeliaceae, Poales).**
In: Cladistics 34 (2), p. 181–199. DOI: 10.1111/cla.12200.

**Horres, Ralf; Schulte, Katharina; Weising, Kurt; Zizka, Georg (2007):**
**Systematics of Bromelioideae (Bromeliaceae) - Evidence from molecular and Anatomical Studies.**
In: Aliso 23 (1), p. 27–43. DOI: 10.5642/aliso.20072301.05.

**Jabaily, Rachel Schmidt; Sytsma, Kenneth J. (2010):**
**Phylogenetics of *Puya* (Bromeliaceae): Placement, major Lineages, and Evolution of Chilean Species.**
In: American Journal of Botany 97 (2), p. 337–356. DOI: 10.3732/ajb.0900107.

**Krapp, Florian; Barros Pinangé, Diego Sotero; Benko-Iseppon, Ana Maria; Leme, Elton M.C.; Weising, Kurt (2014):**

**Phylogeny and Evolution of *Dyckia* (Bromeliaceae) inferred from Chloroplast and nuclear Sequences.**
In: Plant Systematics and Evolution 300, p. 1591–1614. DOI: 10.1007/s00606-014-0985-0.

Louzada, Rafael Batista; Schulte, Katharina; Wanderley, Maria das Graças Lapa; Silvestro, Daniele; Zizka, Georg; Barfuss, Michael H. J.; Palma-silva, Clarisse (2014):
**Molecular Phylogeny of the Brazilian endemic Genus *Orthophytum* (Bromelioideae, Bromeliaceae) and its Implications on morphological Character Evolution.**
In: Molecular Phylogenetics and Evolution 77, p. 54–64. DOI: 10.1016/j.ympev.2014.03.007.

Louzada, Rafael Batista; Versieux, Leonardo M. (2010):
***Lapanthus* (Bromeliaceae, Bromelioideae): A new Genus from the Southern Espinhaço Range, Brazil.**
In: Systematic Botany 35 (3), p. 497–503. DOI: 10.1600/036364410792495908.

Louzada, Rafael Batista; Wanderley, Maria das Graças Lapa (2010):
**Revision of *Orthophytum* (Bromeliaceae): the Species with sessile Inflorescences.**
In: Phytotaxa 13, p. 1–26.

Martínez-Correa, Nancy; Espejo-Serna, Adolfo; López-Ferrari, Ana Rosa; Ramírez-Morillo, Ivón M. (2010):
**Two Novelties in *Hechtia* (Bromeliaceae, Hechtioideae) from Mexico.**
In: Systematic Botany 35 (4), p. 745–754. DOI: 10.1600/036364410X539835.

Monteiro, Raquel Fernandes; Mantovani, André; Forzza, Rafaela Campostrini (2015):
**Morphological phylogenetic Analysis of Two Early-Diverging Genera of Bromelioideae (Bromeliaceae).**
In: Rodriguésia 66 (2), p. 505–521. DOI: 10.1590/2175-7860201566218.

Pinzón, Juan P.; Ramírez-Morillo, Ivón M.; Carnevali, Germán; Barfuss, Michael H. J.; Till, Walter; Tun, Juan; Ortiz-Díaz, Juan J. (2016):
**Phylogenetics and Evolution of the *Tillandsia utriculata* complex (Bromeliaceae, Tillandsioideae) inferred from three Plastid DNA Markers and the ETS of the nuclear ribosomal DNA.**
In: Botanical Journal of the Linnean Society 181 (3), p. 362–390. DOI: 10.1111/boj.12425.

Rex, Martina; Schulte, Katharina; Zizka, Georg; Peters, Jule; Vásquez, Roberto; Ibisch, Pierre L.; Weising, Kurt (2009):
**Phylogenetic Analysis of *Fosterella* L.B.Sm. (Pitcairnioideae, Bromeliaceae) based on four Chloroplast DNA Regions.**
In: Molecular Phylogenetics and Evolution 51 (3), p. 472–485. DOI: 10.1016/j.ympev.2009.01.001.

Sass, Chodon; Specht, Chelsea D. (2010):
**Phylogenetic estimation of the core Bromelioids with an Emphasis on the Genus *Aechmea* (Bromeliaceae).**
In: Molecular Phylogenetics and Evolution 55 (2), p. 559–571. DOI: 10.1016/j.ympev.2010.01.005.

Schulte, Katharina; Barfuss, Michael H. J.; Zizka, Georg (2009):
**Phylogeny of Bromelioideae (Bromeliaceae) inferred from nuclear and Plastid DNA loci reveals the Evolution of the Tank Habit within the Subfamily.**
In: Molecular Phylogenetics and Evolution 51 (2), p. 327–339. DOI: 10.1016/j.ympev.2009.02.003.

Schulte, Katharina; Silvestro, Daniele; Kiehlmann, Elke; Vesely, Sanja; Novoa, Patricio; Zizka, Georg (2010):
**Detection of recent Hybridization between sympatric Chilean *Puya* Species (Bromeliaceae) using AFLP Markers and Reconstruction of complex Relationships.**
In: Molecular Phylogenetics and Evolution 57 (3), p. 1105–1119. DOI: 10.1016/j.ympev.2010.09.001.

**Schütz, Nicole; Krapp, Florian; Wagner, Natascha; Weising, Kurt (2016):**
**Phylogenetics of Pitcairnioideae s.s. (Bromeliaceae): Evidence from nuclear and Plastid DNA Sequence Data.**
In: Botanical Journal of the Linnean Society 181 (3), p. 323–342. DOI: 10.1111/boj.12403.

**Silvestro, Daniele; Zizka, Georg; Schulte, Katharina (2014):**
**Disentangling the effects of key Innovations on the Diversification of Bromelioideae (bromeliaceae).**
In: Evolution 68 (1), p. 163–175. DOI: 10.1111/evo.12236.

**Sousa, Leandro de Oliveira Furtado de; Wendt, Tania; Brown, Gregory K.; Tuthill, Dorothy E.; Evans, Timothy M. (2007):**
**Monophyly and phylogenetic Relationships in *Lymania* (Bromeliaceae: Bromelioideae) based on Morphology and Chloroplast DNA Sequences.**
In: Systematic Botany 32 (2), p. 264–270.

**Versieux, Leonardo M.; Barbará, Thelma; Wanderley, Maria das Graças Lapa; Calvente, Alice; Fay, Michael F.; Lexer, Christian (2012):**
**Molecular Phylogenetics of the Brazilian giant bromeliads (*Alcantarea*, Bromeliaceae): Implications for morphological Evolution and Biogeography.**
In: Molecular Phylogenetics and Evolution 64 (1), p. 177–189. DOI: 10.1016/j.ympev.2012.03.015.

### Brownlowiaceae

**Tan, H.S.; Chung, R. C.K.; Soepadmo, E. (2011):**
**A Synopsis of *Jarandersonia* (Malvaceae: Brownlowioideae).**
In: Gardens' Bulletin Singapore 63 (1-2), p. 137–144.

### Bruniaceae

**Claßen-Bockhoff, Regine; Oliver, Edward G.H.; Hall, Anthony V.; Quint, Marcus (2011):**
**A new Classification of the South African endemic Family Bruniaceae based on molecular and morphological Data.**
In: Taxon 60 (4), p. 1138–1155.

**Quint, Marcus; Claßen-Bockhoff, Regine (2006):**
**Phylogeny of Bruniaceae based on matK and ITS Sequence Data.**
In: International Journal of Plant Sciences 167 (1), p. 135–146.

**Quint, Marcus; Claßen-Bockhoff, Regine (2008):**
**Ancient or recent? Insights into the temporal Evolution of the Bruniaceae.**
In: Organisms Diversity and Evolution 8 (4), p. 293–304. DOI: 10.1016/j.ode.2008.03.001.

### Burmanniaceae

**Merckx, Vincent S.F.T.; Chatrou, Lars W.; Lemaire, Benny; Sainge, Moses N.; Huysmans, Suzy; Smets, Erik F. (2008):**
**Diversification of myco-heterotrophic Angiosperms: Evidence from Burmanniaceae.**
In: BMC Evolutionary Biology 8, p. 178. DOI: 10.1186/1471-2148-8-178.

**Merckx, Vincent S.F.T.; Schols, Peter; Maas-van de Kamer, Hiltje; Maas, Paul J.M.; Huysmans, Suzy; Smets, Erik (2006):**
**Phylogeny and Evolution of Burmanniaceae (Dioscoreales) based on Nuclear and Mitochondrial Data.**
In: American Journal of Botany 93 (11), p. 1684–1698.

**Neyland, Ray (2002):**

**A Phylogeny inferred from large-subunit (26s) ribosomal DNA Sequences suggests that Burmanniales are polyphyletic.**
In: Australian Systematic Botany 15 (1), p. 19–28. DOI: 10.1071/SB01001.

SUETSUGU, KENJI; KAWAKITA, ATSUSHI; KATO, MAKOTO (2014):
**Evidence for specificity to *Glomus* Group Ab in two Asian mycoheterotrophic *Burmannia* Species.**
In: Plant Species Biology 29 (1), p. 57–64. DOI: 10.1111/j.1442-1984.2012.00387.x.

YOKOYAMA, JUN U.N.; KOIZUMI, YAYOI; YOKOTA, MASATSUGU; TSUKAYA, HIROKAZU (2008):
**Phylogenetic Position of *Oxygyne shinzatoi* (Burmanniaceae) inferred from 18s rDNA Sequences.**
In: Journal of Plant Research 121 (1), p. 27–32. DOI: 10.1007/s10265-007-0136-6.

### Burseraceae

BECERRA, JUDITH X. (2003):
**Evolution of Mexican *Bursera* (Burseraceae) inferred from ITS, ETS, and 5s nuclear ribosomal DNA Sequences.**
In: Molecular Phylogenetics and Evolution 26, p. 300–309.

BECERRA, JUDITH X.; NOGE, KOGI; OLIVIER, SARAI; VENABLE, D. LAWRENCE (2012):
**The Monophyly of *Bursera* and its impact for Divergence times of Burseraceae.**
In: Taxon 61 (2), p. 333–343.

FEDERMAN, SARAH; DORNBURG, ALEX; DOWNIE, ALEXANDER; RICHARD, ALISON F.; DALY, DOUGLAS C.; DONOGHUE, MICHAEL J. (2015):
**The biogeographic Origin of a Radiation of Trees in Madagascar: Implications for the assembly of a tropical forest biome.**
In: BMC Evolutionary Biology 15, p. 216. DOI: 10.1186/s12862-015-0483-1.

FINE, PAUL V. A.; DALY, DOUGLAS C.; MUNOZ, GORKY VILLA; MESONES, ITALO; CAMERON, KENNETH M. (2005):
**The Contribution of edaphic Heterogeneity to the Evolution and Diversity of Burseraceae Trees in the Western Amazon.**
In: Evolution 59 (7), p. 1464–1478.

FINE, PAUL V. A.; ZAPATA, FELIPE; DALY, DOUGLAS C. (2014):
**Investigating processes of Neotropical Rain Forest Tree Diversification by Examining the Evolution and historical Biogeography of the Protieae (Burseraceae).**
In: Evolution 68 (7), p. 1988–2004. DOI: 10.1111/evo.12414.

GOSTEL, MORGAN R.; COY, KIERA A.; WEEKS, ANDREA (2015):
**Microfluidic PCR-based target enrichment: A case Study in two rapid Radiations of *Commiphora* (Burseraceae) from Madagascar.**
In: Journal of Systematics and Evolution 53 (5), p. 411–431. DOI: 10.1111/jse.12173.

GOSTEL, MORGAN R.; PHILLIPSON, PETER B.; WEEKS, ANDREA (2016):
**Phylogenetic Reconstruction of the Myrrh Genus, *Commiphora* (Burseraceae), reveals multiple Radiations in Madagascar and clarifies infrageneric Relationships.**
In: Systematic Botany 41 (1), p. 67–81. DOI: 10.1600/036364416X690598.

THULIN, MATS; BEIER, BJÖRN-AXEL; RAZAFIMANDIMBISON, SYLVAIN G.; BANKS, HANNAH I. (2008):
***Ambilobea*, a new Genus from Madagascar, the Position of *Aucoumea*, and comments on the tribal Classification of the Frankincense and Myrrh Family (Burseraceae).**

In: Nordic Journal of Botany 26 (3-4), p. 218–229. DOI: 10.1111/j.1756-1051.2008.00245.x.

**WEEKS, ANDREA; DALY, DOUGLAS C.; SIMPSON, BERYL B. (2005):**
**The phylogenetic History and Biogeography of the Frankincense and Myrrh Family (Burseraceae) based on nuclear and Chloroplast Sequence Data.**
In: Molecular Phylogenetics and Evolution 35 (1), p. 85–101. DOI: 10.1016/j.ympev.2004.12.021.

**WEEKS, ANDREA; SIMPSON, BERYL B. (2004):**
**Molecular Genetic Evidence for Interspecific Hybridization among Endemic Hispaniolan *Bursera* (Burseraceae).**
In: American Journal of Botany 91 (6), p. 976–984.

**WEEKS, ANDREA; SIMPSON, BERYL B. (2007):**
**Molecular phylogenetic Analysis of *Commiphora* (Burseraceae) yields Insight on the Evolution and historical Biogeography of an "impossible" Genus.**
In: Molecular Phylogenetics and Evolution 42 (1), p. 62–79. DOI: 10.1016/j.ympev.2006.06.015.

### Buxaceae

**BALTHAZAR, MARIA VON; ENDRESS, PETER K.; QIU, YIN-LONG (2000):**
**Phylogenetic Relationships in Buxaceae based on nuclear internal transcribed Spacers and Plastid ndhF Sequences.**
In: International Journal of Plant Sciences 161 (5), p. 785–792.

**JIAO, ZHIHUA; LI, JIAN-HUA (2009):**
**Phylogenetics and Biogeography of eastern Asian-North American disjunct Genus *Pachysandra* (Buxaceae) inferred from nucleotide Sequences.**
In: Journal of Systematics and Evolution 47 (3), p. 191–201. DOI: 10.1111/j.1759-6831.2009.00021.x.

**VAN LAERE, KATRIJN; HERMANS, DIDIER; LEUS, LEEN; VAN HUYLENBROECK, JOHAN (2011):**
**Genetic Relationships in European and Asiatic *Buxus* Species based on AFLP Markers, Genome sizes and Chromosome Numbers.**
In: Plant Systematics and Evolution 293 (1-4), p. 1–11. DOI: 10.1007/s00606-011-0422-6.

### Byblidaceae

**FUKUSHIMA, KENJI; IMAMURA, KAORI; NAGANO, KATSUYA; HOSHI, YOSHIKAZU (2011):**
**Contrasting Patterns of the 5s and 45s rDNA Evolutions in the *Byblis liniflora* complex (Byblidaceae).**
In: Journal of Plant Research 124 (2), p. 231–244. DOI: 10.1007/s10265-010-0366-x.

### Byttneriaceae

**BORRONE, JAMES W.; MEEROW, ALAN W.; KUHN, DAVID N.; WHITLOCK, BARBARA A.; SCHNELL, RAYMOND J. (2007):**
**The potential of the Wrky Gene Family for phylogenetic reconstruction: an Example from the Malvaceae.**
In: Molecular Phylogenetics and Evolution 44 (3), p. 1141–1154. DOI: 10.1016/j.ympev.2007.06.012.

**GONÇALEZ, VICTOR MARTINS; ESTEVES, GERLENI LOPES (2015):**
**Synopsis of *Melochia* L. (Byttnerioideae, Malvaceae) in southeastern Brazil.**
In: Phytotaxa 226 (3), p. 217. DOI: 10.11646/phytotaxa.226.3.2.

**RONDÓN, JOSÉ BAUDILIO (2007):**
**Estudio Taxonómico del género *Melochia* L. (Sterculiaceae) en el estado Sucre, Venezuela.**
In: Revista UDO Agrícola 7 (1), p. 122–137.

**Sousa Silva, C. R.; Figueira, A. (2005):**
**Phylogenetic Analysis of *Theobroma* (Sterculiaceae) based on Kunitz-like Trypsin inhibitor Sequences.**
In: Plant Systematics and Evolution 250 (1-2), p. 93–104. DOI: 10.1007/s00606-004-0223-2.

**Whitlock, Barbara A.; Bayer, Clemens; Baum, David A. (2001):**
**Phylogenetic Relationships and Floral Evolution of the Byttnerioideae ("Sterculiaceae" or Malvaceae s.l.) based on Sequences of the Chloroplast Gene, ndhF.**
In: Systematic Botany 26 (2), p. 420–437.

**Whitlock, Barbara A.; Hale, Amanda M. (2011):**
**The Phylogeny of *Ayenia, Byttneria,* and *Rayleya* (Malvaceae s. l.) and its Implications for the Evolution of Growth Forms.**
In: Systematic Botany 36 (1), p. 129–136. DOI: 10.1600/036364411X553216.

**Whitlock, Barbara A.; Hale, Amanda M.; Indorf, Jane L.; Wilkins, Carolyn F. (2011):**
**Polyphyly of *Rulingia* and *Commersonia* (Lasiopetaleae, Malvaceae s.l.).**
In: Australian Systematic Botany 24 (5), p. 215–225. DOI: 10.1071/SB09030.

**Wilkins, Carolyn F.; Whitlock, Barbara A. (2011):**
**A new Australian Genus, *Androcalva*, separated from *Commersonia* (Malvaceae s.l. or Byttneriaceae).**
In: Australian Systematic Botany 24 (5), p. 284–349. DOI: 10.1071/SB10032.

### Cabombaceae

**Vialette-Guiraud, Aurelie C. M.; Alaux, Michael; Legeai, Fabrice; Finet, Cedric; Chambrier, Pierre; Brown, Spencer C. et al. (2011):**
***Cabomba* as a model for studies of early Angiosperm Evolution.**
In: Annals of Botany 108 (4), p. 589–598. DOI: 10.1093/aob/mcr088.

### Cactaceae

**Applequist, Wendy L.; Wallace, Robert p. (2002):**
**Deletions in the Plastid trn T- trn L intergenic spacer define Clades within Cactaceae Subfamily Cactoideae.**
In: Plant Systematics and Evolution 231 (1-4), p. 153–162. DOI: 10.1007/s006060200017.

**Arias, Salvador; Terrazas, Teresa (2009):**
**Taxonomic Revision of *Pachycereus* (Cactaceae).**
In: Systematic Botany 34 (1), p. 68–83. DOI: 10.1600/036364409787602384.

**Arias, Salvador; Terrazas, Teresa; Arreola-Nava, Hilda J.; Vázquez-Sánchez, Monserrat; Cameron, Kenneth M. (2005):**
**Phylogenetic Relationships in *Peniocereus* (Cactaceae) inferred from Plastid DNA Sequence Data.**
In: Journal of Plant Research 118 (5), p. 317–328. DOI: 10.1007/s10265-005-0225-3.

**Baker, Marc A.; Butterworth, Charles A. (2013):**
**Geographic Distribution and taxonomic Circumscription of populations within *Coryphantha* Section *Robustispina* (Cactaceae).**
In: American Journal of Botany 100 (5), p. 984–997. DOI: 10.3732/ajb.1200619.

**Bárcenas, Rolando T. (2016):**

**A molecular phylogenetic approach to the Systematics of Cylindropuntieae (Opuntioideae, Cactaceae).**
In: Cladistics 32 (4), p. 351–359. DOI: 10.1111/cla.12135.

**BÁRCENAS, ROLANDO T.; YESSON, CHRIS; HAWKINS, JULIE A. (2011):**
**Molecular Systematics of the Cactaceae.**
In: Cladistics 27 (5), p. 470–489. DOI: 10.1111/j.1096-0031.2011.00350.x.

**BUTTERWORTH, CHARLES A.; WALLACE, ROBERT P. (2004):**
**Phylogenetic studies of *Mammillaria* (Cactaceae) - Insights from Chloroplast Sequence Variation and Hypothesis Testing using the parametric Bootstrap.**
In: American Journal of Botany 91 (7), p. 1086–1098. DOI: 10.3732/ajb.91.7.1086.

**BUTTERWORTH, CHARLES A.; WALLACE, ROBERT P. (2005):**
**Molecular Phylogenetics of the leafy Cactus Genus *Pereskia* (Cactaceae).**
In: Systematic Botany 30 (4), p. 800–808.

**CALVENTE, ALICE; ZAPPI, DANIELA C.; FOREST, FÉLIX; LOHMANN, LÚCIA GARCES (2011):**
**Molecular Phylogeny of Tribe Rhipsalideae (Cactaceae) and taxonomic Implications for *Schlumbergera* and *Hatiora*.**
In: Molecular Phylogenetics and Evolution 58 (3), p. 456–468. DOI: 10.1016/j.ympev.2011.01.001.

**CALVENTE, ALICE; ZAPPI, DANIELA C.; FOREST, FÉLIX; LOHMANN, LÚCIA GARCES (2011):**
**Molecular Phylogeny, Evolution, and Biogeography of South American epiphytic Cacti.**
In: International Journal of Plant Sciences 172 (7), p. 902–914. DOI: 10.1086/660881.

**CRUZ, MIGUEL ÁNGEL; ARIAS, SALVADOR; TERRAZAS, TERESA (2016):**
**Molecular Phylogeny and Taxonomy of the Genus *Disocactus* (Cactaceae), based on the DNA Sequences of six Chloroplast Markers.**
In: Willdenowia 46 (1), p. 145–164. DOI: 10.3372/wi.46.46112.

**DEMAIO, PABLO H.; BARFUSS, MICHAEL H. J.; KIESLING, ROBERTO; TILL, WALTER; CHIAPELLA, JORGE O. (2011):**
**Molecular Phylogeny of *Gymnocalycium* (Cactaceae): assessment of alternative infrageneric systems, a new Subgenus, and Trends in the Evolution of the Genus.**
In: American Journal of Botany 98 (11), p. 1841–1854. DOI: 10.3732/ajb.1100054.

**EDWARDS, ERIKA J.; NYFFELER, RETO; DONOGHUE, MICHAEL J. (2005):**
**Basal Cactus Phylogeny: Implications of *Pereskia* (Cactaceae) Paraphyly for the Transition to the Cactus Life Form.**
In: American Journal of Botany 92 (7), p. 1177–1188. DOI: 10.3732/ajb.92.7.1177.

**FRANCK, ALAN R.; COCHRANE, BRUCE J.; GAREY, JAMES R. (2013):**
**Relationships and Dispersal of the Caribbean Species of *Harrisia* (Sect. *Harrisia*; Cactaceae) using AFLPs and seven DNA Regions.**
In: Taxon 62 (3), p. 486–497. DOI: 10.12705/623.5.

**GRIFFITH, M. PATRICK; PORTER, J. MARK (2009):**
**Phylogeny of Opuntioideae (Cactaceae).**
In: International Journal of Plant Sciences 170 (1), p. 107–116. DOI: 10.1086/593048.

**GUERRERO, PABLO C.; ARROYO, MARY T.K.; BUSTAMANTE, RAMIRO O.; DUARTE, MILÉN; HAGEMANN, THOMAS K.; WALTER, HELMUT E. (2011):**
**Phylogenetics and predictive Distribution modeling provide Insights into the geographic Divergence of *Eriosyce* subgen. *Neoporteria* (Cactaceae).**

In: Plant Systematics and Evolution 297 (1-2), p. 113–128. DOI: 10.1007/s00606-011-0512-5.

**Harpke, Dörte; Peterson, Angela (2006):**
**Non-concerted ITS Evolution in *Mammillaria* (Cactaceae).**
In: Molecular Phylogenetics and Evolution 41 (3), p. 579–593. DOI: 10.1016/j.ympev.2006.05.036.

**Hernández-Hernández, Tania; Hernández, Héctor M.; De-Nova, José Arturo; Puente, Raul; Eguiarte, Luis E.; Magallón, Susana (2011):**
**Phylogenetic Relationships and Evolution of growth form in Cactaceae (Caryophyllales, Eudicotyledoneae).**
In: American Journal of Botany 98 (1), p. 44–61. DOI: 10.3732/ajb.1000129.

**Korotkova, Nadja; Borsch, Thomas; Quandt, Dietmar; Taylor, Nigel P.; Müller, Kai F.; Barthlott, Wilhelm (2011):**
**What does it take to resolve Relationships and to identify Species with molecular Markers? An example from the epiphytic Rhipsalideae (Cactaceae).**
In: American Journal of Botany 98 (9), p. 1549–1572. DOI: 10.3732/ajb.1000502.

**Larridon, Isabel; Walter, Helmut E.; Guerrero, Pablo C.; Duarte, Milén; Cisternas, Mauricio A.; Hernández, Carol Peña et al. (2015):**
**An integrative approach to Understanding the Evolution and Diversity of *Copiapoa* (Cactaceae), a threatened endemic Chilean Genus from the Atacama Desert.**
In: American Journal of Botany 102 (9), p. 1506–1520. DOI: 10.3732/ajb.1500168.

**Las Peñas, M. L.; Urdampilleta, J. D.; López-Carro, B.; Santiñaque, F.; Kiesling, Roberto; Bernardello, Gabriel (2014):**
**Classical and molecular cytogenetics and DNA content in *Maihuenia* and *Pereskia* (Cactaceae).**
In: Plant Systematics and Evolution 300 (3), p. 549–558. DOI: 10.1007/s00606-013-0903-x.

**Lemos, Renata C. C.; Machado, Marlon C.; Melo-De-Pinna, Gladys F. (2013):**
**Morpho-anatomical Diversity of the underground Systems of *Arrojadoa* (Cactaceae), an endemic Brazilian Genus.**
In: Botanical Journal of the Linnean Society 173 (1), p. 108–128. DOI: 10.1111/boj.12077.

**Majure, Lucas C.; Puente, Raul; Griffith, M. Patrick; Judd, Walter S.; Soltis, Pamela S.; Soltis, Douglas E. (2012):**
**Phylogeny of *Opuntia* s.s. (Cactaceae): Clade delineation, geographic Origins, and reticulate Evolution.**
In: American Journal of Botany 99 (5), p. 847–864. DOI: 10.3732/ajb.1100375.

**Martínez-González, César; Alcántara, Othón; Luna-Vega, Isolda; Garcia-Sandoval, Ricardo (2015):**
**Phylogenetic Placement and new Data on macro- and micro-Morphology of *Nopalxochia phyllanthoides* (Cactaceae), an endangered Species from Mexico.**
In: Phytotaxa 222 (4), p. 241. DOI: 10.11646/phytotaxa.222.4.1.

**Nyffeler, Reto (2002):**
**Phylogenetic Relationships in the Cactus Family (Cactaceae) based on Evidence from trnK/ matK and trnL-trnF Sequences.**
In: American Journal of Botany 89 (2), p. 312–326. DOI: 10.3732/ajb.89.2.312.

**Realini, María F.; González, Graciela E.; Font, Fabián; Picca, Pablo I.; Poggio, Lidia; Gottlieb, Alexandra M. (2015):**
**Phylogenetic Relationships in *Opuntia* (Cactaceae, Opuntioideae) from southern South America.**

In: Plant Systematics and Evolution 301 (4), p. 1123–1134. DOI: 10.1007/s00606-014-1154-1.

**RITZ, CHRISTIANE M.; FICKENSCHER, KARL; FÖLLER, JENS; HERRMANN, KATJA; MECKLENBURG, RAINER; WAHL, RAINER (2016):**

**Molecular phylogenetic Relationships of the Andean Genus *Aylostera* Speg. (Cactaceae, Trichocereeae), a new Classification and a morphological Identification Key.**

In: Plant Systematics and Evolution 302 (7), p. 763–780. DOI: 10.1007/s00606-016-1296-4.

**RITZ, CHRISTIANE M.; MARTINS, LUDWIG; MECKLENBURG, RAINER; GOREMYKIN, VADIM V.; HELLWIG, FRANK H. (2007):**

**The molecular Phylogeny of *Rebutia* (Cactaceae) and its Allies demonstrates the Influence of Paleogeography on the Evolution of South American Mountain Cacti.**

In: American Journal of Botany 94 (8), p. 1321–1332. DOI: 10.3732/ajb.94.8.1321.

**RITZ, CHRISTINE M.; REIKER, J.; CHARLES, G.; HOXEY, P.; HUNT, D.; LOWRY, M. ET AL. (2012):**

**Molecular Phylogeny and Character Evolution in terete-stemmed Andean Opuntias (Cactaceae-Opuntioideae).**

In: Molecular Phylogenetics and Evolution 65 (2), p. 668–681. DOI: 10.1016/j.ympev.2012.07.027.

**SCHLUMPBERGER, BORIS O. (2012):**

**New Combinations in the *Echinopsis* alliance.**

In: Cactaceae Systematics Initiatives 28, p. 29–31.

**SCHLUMPBERGER, BORIS O.; RENNER, SUSANNE P. (2012):**

**Molecular Phylogenetics of *Echinopsis* (Cactaceae): Polyphyly at all levels and convergent Evolution of Pollination modes and growth forms.**

In: American Journal of Botany 99 (8), p. 1335–1349. DOI: 10.3732/ajb.1100288.

### Calceolariaceae

**ANDERSSON, STEPHAN (2006):**

**On the Phylogeny of the Genus *Calceolaria* (Calceolariaceae) as inferred from ITS and Plastid matK Sequences.**

In: Taxon 55 (1), p. 125–137. DOI: 10.2307/25065534.

**COSACOV, ANDREA; SÉRSIC, ALICIA N.; SOSA, VICTORIA; DE-NOVA, JOSÉ ARTURO; NYLINDER, STEPHAN; COCUCCI, ANDREA A. (2009):**

**New Insights into the phylogenetic Relationships, Character Evolution, and phytogeographic Patterns of *Calceolaria* (Calceolariaceae).**

In: American Journal of Botany 96 (12), p. 2240–2255. DOI: 10.3732/ajb.0900165.

**EHRHART, CHRISTINE (2005):**

**The Chilean *Calceolaria integrifolia* s.l. Species Complex (Scrophulariaceae).**

In: Systematic Botany 30 (2), p. 383–411.

**NYLINDER, STEPHAN; SWENSON, ULF; PERSSON, CLAES; JANSSENS, STEVEN B.; OXELMAN, BENGT (2012):**

**A dated species-tree approach to the trans-Pacific Disjunction of the Genus *Jovellana* (Calceolariaceae, Lamiales).**

In: Taxon 61 (2), p. 381–391. DOI: 10.1002/tax.612009.

**PUPPO, PAMELA (2010):**

**A new Species of *Calceolaria* (Calceolariaceae) from disturbed Paramos in South Ecuador.**

In: Journal of the Botanical Research Institute of Texas 4 (1), p. 33–36.

**PUPPO, PAMELA (2014):**

**Revision of the *Calceolaria tripartita* s. l. Species complex (Calceolariaceae) using multivariate Analyses of morphological characters.**
In: Phytotaxa 167 (1), p. 61. DOI: 10.11646/phytotaxa.167.1.3.

### Calophyllaceae

**Zakaria, Radhiah; Choong, Chee Yen; Faridah-Hanum, Ibrahim (2007):**
**Systematic Study on Guttiferae Juss. of Peninsular Malaysia based on Plastid Sequences.**
In: Tropics 16 (2), p. 141–150. DOI: 10.3759/tropics.16.141.

### Calycanthaceae

**Zhou, Shiliang; Renner, Susanne S.; Wen, Jun (2006):**
**Molecular Phylogeny and intra- and intercontinental Biogeography of Calycanthaceae.**
In: Molecular Phylogenetics and Evolution 39 (1), p. 1–15. DOI: 10.1016/j.ympev.2006.01.015.

### Calyceraceae

**Denham, Silvia S.; Zavala-Gallo, Lucio; Johnson, Leigh A.; Pozner, Raúl E. (2016):**
**Insights into the Phylogeny and evolutionary History of Calyceraceae.**
In: Taxon 65 (6), p. 1328–1344. DOI: 10.12705/656.7.

**Pozner, Raúl E.; Zanotti, Christian; Johnson, Leigh A. (2012):**
**Evolutionary Origin of the Asteraceae Capitulum: Insights from Calyceraceae.**
In: American Journal of Botany 99 (1), p. 1–13. DOI: 10.3732/ajb.1100256.

**Zavala-Gallo, Lucio; Denham, Silvia S.; Pozner, Raúl E. (2010):**
**Revision of the Genus *Nastanthus* (Calyceraceae).**
In: Gayana Botánica 67 (2), p. 158–175. DOI: 10.4067/S0717-66432010000200002.

### Campanulaceae

**Antonelli, Alexandre (2008):**
**Higher level Phylogeny and evolutionary Trends in Campanulaceae Subfam. Lobelioideae: molecular Signal overshadows Morphology.**
In: Molecular Phylogenetics and Evolution 46 (1), p. 1–18. DOI: 10.1016/j.ympev.2007.06.015.

**Borsch, Thomas; Korotkova, Nadja; Raus, Thomas; Lobin, Wolfram; Löhne, Cornelia (2009):**
**The petD Group II Intron as a Species level marker: utility for Tree Inference and Species Identification in the diverse Genus *Campanula* (Campanulaceae).**
In: Willdenowia 39 (1), p. 7–33. DOI: 10.3372/wi.39.39101.

**Cheon, Kyeong-Sik; Yoo, Ki-Oug (2013):**
**Phylogeny of *Hanabusaya* (Campanulaceae), a Korean endemic, based on ITS Sequences of nuclear ribosomal DNA.**
In: Journal of Systematics and Evolution 51 (6), p. 704–714. DOI: 10.1111/jse.12039.

**Haberle, Rosemarie C.; Dang, Ashley; Lee, Tammy; Peñaflor, Cynthia; Cortes-Burns, Helen; Oestreich, Andrea et al. (2009):**
**Taxonomic and biogeographic Implications of a phylogenetic Analysis of the Campanulaceae based on three Chloroplast Genes.**
In: Taxon 58 (3), p. 715–734.

**Knox, Eric B.; Muasya, A. Muthama; Muchhala, Nathan (2008):**
**The Predominantly South American Clade of Lobeliaceae.**
In: Systematic Botany 33 (2), p. 462–468. DOI: 10.1600/036364408784571590.

**Koopman, Margaret M.; Ayers, Tina J. (2005):**
**Nectar Spur Evolution in the Mexican Lobelias (Campanulaceae: Lobelioideae).**
In: American Journal of Botany 92 (3), p. 558–562. DOI: 10.3732/ajb.92.3.558.

**Lagomarsino, Laura P.; Antonelli, Alexandre; Muchhala, Nathan; Timmermann, Allan; Mathews, Sarah; Davis, Charles C. (2014):**
**Phylogeny, Classification, and Fruit Evolution of the species-rich Neotropical bellflowers (Campanulaceae: Lobelioideae).**
In: American Journal of Botany 101 (12), p. 2097–2112. DOI: 10.3732/ajb.1400339.

**Lakušić, Dmitar; Liber, Zlatko; Nikolić, Toni; Surina, Boštjan; Kovačić, Sanja; Bogdanović, Sandro; Stefanović, Saša (2013):**
**Molecular Phylogeny of the *Campanula pyramidalis* Species complex (Campanulaceae) inferred from Chloroplast and nuclear non-coding Sequences and its taxonomic Implications.**
In: Taxon 62 (3), p. 505–524. DOI: 10.12705/623.1.

**Olesen, Jens M.; Alarcón, Marisa; Ehlers, Bodil K.; Aldasoro, Juan José; Roquet, Cristina (2012):**
**Pollination, Biogeography and Phylogeny of oceanic Island Bellflowers (Campanulaceae).**
In: Perspectives in Plant Ecology, Evolution and Systematics 14 (3), p. 169–182. DOI: 10.1016/j.ppees.2012.01.003.

**Park, Jeong-Mi; Kovačić, Sanja; Liber, Zlatko; Eddie, William M.M.; Schneeweiss, Gerald M. (2006):**
**Phylogeny and Biogeography of Isophyllous Species of *Campanula* (Campanulaceae) in the Mediterranean Area.**
In: Systematic Botany 31 (4), p. 862–880.

**Prebble, Jessica M.; Cupido, Christopher N.; Meudt, Heidi M.; Garnock-Jones, Philip J. (2011):**
**First phylogenetic and biogeographical Study of the southern Bluebells (*Wahlenbergia*, Campanulaceae).**
In: Molecular Phylogenetics and Evolution 59 (3), p. 636–648. DOI: 10.1016/j.ympev.2011.03.013.

**Prebble, Jessica M.; Meudt, Heidi M.; Garnock-Jones, Philip J. (2012):**
**An expanded molecular Phylogeny of the southern bluebells (*Wahlenbergia*, Campanulaceae) from Australia and New Zealand.**
In: Australian Systematic Botany 25 (1), p. 11. DOI: 10.1071/SB11023.

**Roquet, Cristina; Sáez, Llorenç; José Aldasoro, Juan; Susanna, Alfonso; Luisa Alarcón, María; Garcia-Jacas, Núria (2008):**
**Natural Delineation, molecular Phylogeny and Floral Evolution in *Campanula*.**
In: Systematic Botany 33 (1), p. 203–217. DOI: 10.1600/036364408783887465.

**Roquet, Cristina; Sanmartín, Isabel; Garcia-Jacas, Núria; Sáez, Llorenç; Susanna, Alfonso; Wikström, Niklas; Aldasoro, Juan José (2009):**
**Reconstructing the History of Campanulaceae with a Bayesian approach to molecular dating and Dispersal-vicariance Analyses.**
In: Molecular Phylogenetics and Evolution 52 (3), p. 575–587. DOI: 10.1016/j.ympev.2009.05.014.

**Schneeweiss, Gerald M.; Pachschwöll, Clemens; Tribsch, Andreas; Schönswetter, Peter; Barfuss, Michael H. J.; Esfeld, Korinna et al. (2013):**
**Molecular phylogenetic Analyses identify Alpine differentiation and dysploid Chromosome Number Changes as major forces for the Evolution of the European endemic *Phyteuma* (Campanulaceae).**

In: Molecular Phylogenetics and Evolution 69 (3), p. 634–652. DOI: 10.1016/j.ympev.2013.07.015.

Stefanović, Saša; Lakušić, Dmitar; Kuzmina, Maria; Međedović, Safer; Tan, Kit; Stevanović, Vladimir (2008):
**Molecular Phylogeny of *Edraianthus* (Grassy Bells; Campanulaceae) based on non-coding Plastid DNA Sequences.**
In: Taxon 57 (2), p. 452–475.

Wang, Qiang; Hong, De-Yuan (2015):
**Taxonomic Revision of the Genus *Pseudocodon* (Campanulaceae) based on Character Analysis and molecular Phylogeny.**
In: Phytotaxa 204 (1), p. 49. DOI: 10.11646/phytotaxa.204.1.4.

Wang, Qiang; Zhou, Shi-Liang; Hong, De-Yuan (2013):
**Molecular Phylogeny of the platycodonoid Group (Campanulaceae s.str.) with special Reference to the Circumscription of *Codonopsis*.**
In: Taxon 62 (3), p. 498–504.

Zhou, Zhuo; Hong, De-Yuan; Niu, Yang; Li, Guo-Dong; Nie, Ze-long; Wen, Jun; Sun, Hang (2013):
**Phylogenetic and biogeographic Analyses of the Sino-Himalayan endemic Genus *Cyananthus* (Campanulaceae) and Implications for the Evolution of its sexual system.**
In: Molecular Phylogenetics and Evolution 68 (3), p. 482–497. DOI: 10.1016/j.ympev.2013.04.027.

Zhou, Zhuo; Wen, Jun; Li, Guo-Dong; Sun, Hang (2012):
**Phylogenetic assessment and biogeographic Analyses of Tribe Peracarpeae (Campanulaceae).**
In: Plant Systematics and Evolution 298 (2), p. 323–336. DOI: 10.1007/s00606-011-0547-7.

### Canellaceae

Müller, Sebastian; Salomo, Karsten; Salazar, Jackeline; Naumann, Julia; Jaramillo, M. Alejandra; Neinhuis, Christoph et al. (2015):
**Intercontinental long-distance Dispersal of Canellaceae from the new to the Old World revealed by a nuclear single Copy Gene and Chloroplast Loci.**
In: Molecular Phylogenetics and Evolution 84, p. 205–219. DOI: 10.1016/j.ympev.2014.12.010.

Zimmer, Elizabeth A.; Suh, Youngbae; Kenneth, Karol G. (2012):
**Phylogenetic Placement of a recently described Taxon of the Genus *Pleodendron* (Canellaceae).**
In: Phytologia 94 (3), p. 404–412.

### Cannabaceae

Yang, Mei-Qing; van Velzen, Robin; Bakker, Freek T.; Sattarian, Ali; Li, De-Zhu; Yi, Ting-Shuang (2013):
**Molecular Phylogenetics and Character Evolution of Cannabaceae.**
In: Taxon 62 (3), p. 473–485.

Yesson, Chris; Russell, Stephen J.; Parrish, T.; Dalling, J. W.; Garwood, Nancy C. (2004):
**Phylogenetic Framework for *Trema* (Celtidaceae).**
In: Plant Systematics and Evolution 248 (1-4), p. 85–109. DOI: 10.1007/s00606-004-0186-3.

### Cannales

Barrett, Craig F.; Specht, Chelsea D.; Leebens-Mack, Jim H.; Stevenson, Dennis Wm.; Zomlefer, Wendy B.; Davis, Jerrold I. (2014):
**Resolving ancient Radiations: Can complete Plastid Gene sets elucidate deep Relationships among the tropical Gingers (Zingiberales)?**

In: Annals of Botany 113 (1), p. 119–133. DOI: 10.1093/aob/mct264.

**DENG, JIABIN; GAO, GANG; ZHANG, YAN; HE, FENGMEI; LUO, XUQIANG; ZHANG, FENGTAI ET AL. (2016):**
**Phylogenetic and ancestral Area Reconstruction of Zingiberales from Plastid Genomes.**
In: Biochemical Systematics and Ecology 66, p. 123–128. DOI: 10.1016/j.bse.2016.03.013.

### Capparaceae

**CORNEJO, XAVIER; ILTIS, HUGH H. (2006):**
**New Combinations in Capparaceae sensu stricto for Flora of Ecuador.**
In: Harvard Papers in Botany 11 (1), p. 17–18. DOI: 10.3100/1043-4534(2006)11[17:NCICSS]2.0.CO;2.

**CORNEJO, XAVIER; ILTIS, HUGH H. (2008):**
**The Reinstatement of *Capparidastrum* (Capparaceae).**
In: Harvard Papers in Botany 13 (2), p. 229–236. DOI: 10.3100/1043-4534-13.2.229.

**CORNEJO, XAVIER; ILTIS, HUGH H.; CERÓN, CARLOS E. (2014):**
***Capparidastrum tafallanum* (Capparaceae), a new Species from the Northwestern Andean Slopes of Ecuador.**
In: Harvard Papers in Botany 19 (2), p. 189–191. DOI: 10.3100/hpib.v19iss2.2014.n4.

**HALL, JOCELYN C. (2008):**
**Systematics of Capparaceae and Cleomaceae: an Evaluation of the generic Delimitations of *Capparis* and *Cleome* using Plastid DNA Sequence Data.**
In: Botany 86 (7), p. 682–696. DOI: 10.1139/B08-026.

**HALL, JOCELYN C.; SYTSMA, KENNETH J.; ILTIS, HUGH H. (2002):**
**Phylogeny of Capparaceae and Brassicaceae based on Chloroplast Sequence Data.**
In: American Journal of Botany 89 (11), p. 1826–1842. DOI: 10.3732/ajb.89.11.1826.

**ILTIS, HUGH H.; CORNEJO, XAVIER (2007):**
**Studies in the Capparaceae XXX: *Capparicordis*, a new Genus from the Neotropics.**
In: Botanical Journal of the Linnean Society 59 (3), p. 245–254. DOI: 10.1663/0007-196X(2007)59[245:SITCXC]2.0.CO;2.

**ILTIS, HUGH H.; HALL, JOCELYN C.; COCHRANE, THEODORE S.; SYTSMA, KENNETH J. (2011):**
**Studies in the Cleomaceae I. on the Separate Recognition of Capparaceae, Cleomaceae, and Brassicaceae.**
In: Annals of the Missouri Botanical Garden 98 (1), p. 28–36. DOI: 10.3417/2007017.

### Capparidales

**CARDINAL-MCTEAGUE, WARREN M.; SYTSMA, KENNETH J.; HALL, JOCELYN C. (2016):**
**Biogeography and Diversification of Brassicales: A 103-Million Year Tale.**
In: Molecular Phylogenetics and Evolution 99, p. 204–224. DOI: 10.1016/j.ympev.2016.02.021.

### Caprifoliaceae

**BELL, CHARLES D. (2010):**
**Towards a Species Level Phylogeny of *Symphoricarpos* (Caprifoliaceae), based on Nuclear and Chloroplast DNA.**
In: Systematic Botany 35 (2), p. 442–450. DOI: 10.1600/036364410791638351.

**JACOBS, BART; GEUTEN, KOEN; PYCK, NANCY; HUYSMANS, SUZY; JANSEN, STEVEN; SMETS, ERIK (2011):**
**Unraveling the Phylogeny of *Heptacodium* and *Zabelia* (Caprifoliaceae): An interdisciplinary Approach.**
In: Systematic Botany 36 (1), p. 231–252. DOI: 10.1600/036364411X553306.

**Landrein, Sven (2010):**

**_Diabelia_, a new Genus of Tribe Linnaeeae Subtribe Linnaeinae (Caprifoliaceae).**

In: Phytotaxa 3, p. 34–38.

**Landrein, Sven; Prenner, Gerhard (2013):**

**Unequal Twins? Inflorescence Evolution in the Twinflower Tribe Linnaeeae (Caprifoliaceae s.l.).**

In: International Journal of Plant Sciences 174 (2), p. 200–233. DOI: 10.1086/668251.

**Theis, Nina; Donoghue, Michael J.; Li, Jian-Hua (2008):**

**Phylogenetics of the Caprifolieae and _Lonicera_ (Dipsacales) based on Nuclear and Chloroplast DNA Sequences.**

In: Systematic Botany 33 (4), p. 776–783. DOI: 10.1600/036364408786500163.

**Wang, Qingxia; Quan, Qingmei; Zhou, Xueli; Zhu, Yunguo; Lan, Yunlong; Li, Shan et al. (2014):**

**A comparative Study of _Lonicera japonica_ with related species: morphological characteristics, ITS Sequences and active Compounds.**

In: Biochemical Systematics and Ecology 54, p. 198–207. DOI: 10.1016/j.bse.2014.02.002.

### Cardiopteridaceae

**Munzinger, Jérôme; Levionnois, Sebastien (2016):**

**Novitates Neocaledonicae III: A new Species of _Citronella_ (Cardiopteridaceae) endemic to New Caledonia.**

In: Phytotaxa 245 (3), p. 223. DOI: 10.11646/phytotaxa.245.3.5.

**Tobe, Hiroshi (2012):**

**Floral Structure of _Cardiopteris_ (Cardiopteridaceae) with special Emphasis on the Gynoecium: systematic and evolutionary Implications.**

In: Journal of Plant Research 125 (3), p. 361–369. DOI: 10.1007/s10265-011-0450-x.

### Caricaceae

**Antunes Carvalho, Fernanda; Filer, Denis; Renner, Susanne p. (2015):**

**Taxonomy in the electronic Age and an E-monograph of the Papaya Family (Caricaceae) as an Example.**

In: Cladistics 31 (3), p. 321–329. DOI: 10.1111/cla.12095.

**Carvalho, Fernanda Antunes; Renner, Susanne p. (2012):**

**A dated Phylogeny of the Papaya Family (Caricaceae) reveals the crop's closest relatives and the Family's biogeographic History.**

In: Molecular Phylogenetics and Evolution 65 (1), p. 46–53. DOI: 10.1016/j.ympev.2012.05.019.

**Kyndt, Tina; Romeijn-Peeters, Eliza; van Droogenbroeck, Bart; Romero-Motochi, José Parcemon; Gheysen, Godelieve; Goetghebeur, Paul (2005):**

**Species Relationships in the Genus _Vasconcellea_ (Caricaceae) based on molecular and morphological Evidence.**

In: American Journal of Botany 92 (6), p. 1033–1044. DOI: 10.3732/ajb.92.6.1033.

**Kyndt, Tina; van Droogenbroeck, Bart; Romeijn-Peeters, Eliza; Romero-Motochi, José Parcemon; Scheldeman, Xavier; Goetghebeur, Paul et al. (2005):**

**Molecular Phylogeny and Evolution of Caricaceae based on rDNA internal transcribed Spacers and Chloroplast Sequence Data.**

In: Molecular Phylogenetics and Evolution 37 (2), p. 442–459. DOI: 10.1016/j.ympev.2005.06.017.

### Caryophyllaceae

**Adams, Laurie G.; West, Judy G.; Cowley, Kirsten J. (2008):**
**Revision of *Spergularia* (Caryophyllaceae) in Australia.**
In: Australian Systematic Botany 21 (4), p. 251–270. DOI: 10.1071/SB08015.

**Aydin, Zeynep; Ertekin, Alaattin Selçuk; Långström, Elisabeth; Oxelman, Bengt (2014):**
**A new Section of *Silene* (Caryophyllaceae) including a new Species from South Anatolia, Turkey.**
In: Phytotaxa 178 (2), p. 98. DOI: 10.11646/phytotaxa.178.2.2.

**Bacchetta, Gianluigi; Brullo, Salvatore; Casti, Mauro; Pietro Giusso del Galdo, Gian (2010):**
**Taxonomic Revision of the *Dianthus sylvestris* Group (Caryophyllaceae) in central-southern Italy, Sicily and Sardinia.**
In: Nordic Journal of Botany 28 (2), p. 137–173. DOI: 10.1111/j.1756-1051.2009.00459.x.

**Cires, Eduardo; Fernández Prieto, José Antonio (2015):**
**Phylogenetic Relationships of *Petrocoptis* A. Braun ex Endl. (Caryophyllaceae), a discussed Genus from the Iberian Peninsula.**
In: Journal of Plant Research 128 (2), p. 223–238. DOI: 10.1007/s10265-014-0691-6.

**Dillenberger, Markus S.; Kadereit, Joachim W. (2014):**
**Maximum polyphyly: Multiple Origins and Delimitation with plesiomorphic Characters require a new Circumscription of *Minuartia* (Caryophyllaceae).**
In: Taxon 63 (1), p. 64–88. DOI: 10.12705/631.5.

**Dillenberger, Markus S.; Kadereit, Joachim W. (2017):**
**Simultaneous Speciation in the European high mountain flowering plant Genus *Facchinia* (*Minuartia* s.l., Caryophyllaceae) revealed by Genotyping-by-Sequencing.**
In: Molecular Phylogenetics and Evolution 112, p. 23–35. DOI: 10.1016/j.ympev.2017.04.016.

**Erixon, Per; Oxelman, Bengt (2008):**
**Reticulate or Tree-like Chloroplast DNA Evolution in Sileneae (Caryophyllaceae)?**
In: Molecular Phylogenetics and Evolution 48 (1), p. 313–325. DOI: 10.1016/j.ympev.2008.04.015.

**Fior, Simone; Karis, Per Ola (2007):**
**Phylogeny, Evolution and Systematics of *Moehringia* (Caryophyllaceae) as inferred from molecular and morphological Data: a case of homology Reassessment.**
In: Cladistics 23 (4), p. 362–372. DOI: 10.1111/j.1096-0031.2007.00150.x.

**Fior, Simone; Karis, Per Ola; Casazza, Gabriele; Minuto, Luigi; Sala, Francesco (2006):**
**Molecular Phylogeny of the Caryophyllaceae (Caryophyllales) inferred from Chloroplast matK and nuclear rDNA ITS Sequences.**
In: American Journal of Botany 93 (3), p. 399–411. DOI: 10.3732/ajb.93.3.399.

**Frajman, Božo; Eggens, Frida; Oxelman, Bengt (2009):**
**Hybrid Origins and homoploid reticulate Evolution within *Heliosperma* (Sileneae, Caryophyllaceae) - a multigene phylogenetic approach with relative dating.**
In: Systematic Biology 58 (3), p. 328–345. DOI: 10.1093/sysbio/syp030.

**Frajman, Božo; Heidari, Nahid; Oxelman, Bengt (2009):**
**Phylogenetic Relationships of *Atocion* and *Viscaria* (Sileneae, Caryophyllaceae) inferred from Chloroplast, nuclear ribosomal, and low-copy Gene DNA Sequences.**
In: Taxon 58 (3), p. 811–824.

**Frajman, Božo; Oxelman, Bengt (2007):**

**Reticulate Phylogenetics and phytogeographical Structure of *Heliosperma* (Sileneae, Caryophyllaceae) inferred from Chloroplast and nuclear DNA Sequences.**
In: Molecular Phylogenetics and Evolution 43 (1), p. 140–155. DOI: 10.1016/j.ympev.2006.11.003.

**GREENBERG, ANNE K.; DONOGHUE, MICHAEL J. (2011):**
**Molecular Systematics and Character Evolution in Caryophyllaceae.**
In: Taxon 60 (6), p. 1637–1652.

**HARBAUGH, DANICA T.; NEPOKROEFF, MOLLY; RABELER, RICHARD K.; MCNEILL, JOHN; ZIMMER, ELIZABETH A.; WAGNER, WARREN L. (2010):**
**A new Lineage-Based Tribal Classification of the Family Caryophyllaceae.**
In: International Journal of Plant Sciences 171 (2), p. 185–198. DOI: 10.1086/648993.

**KOOL, ANNELEEN; BENGTSON, ANNIKA; THULIN, MATS (2007):**
**Polyphyly of *Polycarpon* (Caryophyllaceae) inferred from DNA Sequence Data.**
In: Taxon 56 (3), p. 775–782. DOI: 10.2307/25065860.

**KOOL, ANNELEEN; PERRIGO, ALLISON; THULIN, MATS (2012):**
**Bristly versus juicy: phylogenetic Position and Taxonomy of *Sphaerocoma* (Caryophyllaceae).**
In: Taxon 61 (1), p. 67–75.

**LEUZINGER, MARIANNE; NACIRI, YAMAMA; DU PASQUIER, PIERRE-EMMANUEL; JEANMONOD, DANIEL (2015):**
**Molecular diversity, Phylogeography and genetic Relationships of the *Silene paradoxa* Group of Section *Siphonomorpha* (Caryophyllaceae).**
In: Plant Systematics and Evolution 301 (1), p. 265–278. DOI: 10.1007/s00606-014-1071-3.

**MONTESINOS, DANIEL BERNARDO; KOOL, ANNELEEN (2015):**
***Arenaria acaulis* (Caryophyllaceae), a new Species from South Peru.**
In: Phytotaxa 220 (1), p. 77. DOI: 10.11646/phytotaxa.220.1.7.

**MOORE, ABIGAIL J.; DILLENBERGER, MARKUS P. (2017):**
**A conspectus of the Genus *Cherleria* (*Minuartia* s.l., Caryophyllaceae ).**
In: Willdenowia 47 (1), p. 5–14. DOI: 10.3372/wi.47.47101.

**MOSYAKIN, SERGEI L.; FEDORONCHUK, MYKOLA M. (2015):**
**New Combinations for East European Species of *Sabulina* (Caryophyllaceae).**
In: Phytotaxa 231 (1), p. 95. DOI: 10.11646/phytotaxa.231.1.10.

**NIKETIC, MARJAN; SILJAK-YAKOVLEV, SONJA; FRAJMAN, BOŽO; LAZAREVIC, MAJA; STEVANOVIC, BRANCA; TOMOVIĆ, GORDANA; STEVANOVIĆ, VLADIMIR (2013):**
**Towards resolving the Systematics of *Cerastium* subsection *Cerastium* (Caryophyllaceae): a cytogenetic approach.**
In: Botanical Journal of the Linnean Society 172, p. 205–224.

**OXELMAN, BENGT; LIÉN, M.; RABELER, RICHARD K.; POPP, MAGNUS (2001):**
**A revised generic Classification of the Tribe Sileneae (Caryophyllaceae).**
In: Nordic Journal of Botany 20, p. 743–748.

**PETRI, ANNA; OXELMAN, BENGT (2011):**
**Phylogenetic Relationships within *Silene* (Caryophyllaceae) Section *Physolychnis*.**
In: Taxon 60 (4), p. 953–968.

**PIRANI, ATEFEH; ZARRE, SHAHIN; PFEIL, BERNARD E.; BERTRAND, YANN J.K.; ASSADI, MOSTAFA; OXELMAN, BENGT (2014):**

**Molecular Phylogeny of *Acanthophyllum* (Caryophyllaceae: Caryophylleae), with Emphasis on infrageneric Classification.**
In: Taxon 63 (3), p. 592–607. DOI: 10.12705/633.39.

**POPP, MAGNUS; ERIXON, PER; EGGENS, FRIDA; OXELMAN, BENGT (2005):**
**Origin and Evolution of a Circumpolar Polyploid Species Complex in *Silene* (Caryophyllaceae) inferred from Low Copy Nuclear RNA Polymerase Introns, rDNA, and Chloroplast DNA.**
In: Systematic Botany 30 (2), p. 302–313.

**POPP, MAGNUS; GIZAW, ABEL; NEMOMISSA, SILESHI; SUDA, JAN; BROCHMANN, CHRISTIAN (2008):**
**Original Article: Colonization and Diversification in the African 'sky Islands' by Eurasian *Lychnis* L. (Caryophyllaceae).**
In: Journal of Biogeography 35 (6), p. 1016–1029. DOI: 10.1111/j.1365-2699.2008.01902.x.

**POPP, MAGNUS; OXELMAN, BENGT (2004):**
**Evolution of a RNA polymerase Gene Family in *Silene* (Caryophyllaceae)-incomplete concerted Evolution and topological Congruence among Paralogues.**
In: Systematic Biology 53 (6), p. 914–932. DOI: 10.1080/10635150490888840.

**POPP, MAGNUS; OXELMAN, BENGT (2007):**
**Origin and Evolution of North American polyploid *Silene* (Caryophyllaceae).**
In: American Journal of Botany 94 (3), p. 330–349. DOI: 10.3732/ajb.94.3.330.

**RABELER, RICHARD K.; WAGNER, WARREN L. (2016):**
**New Combinations in *Odontostemma* (Caryophyllaceae).**
In: PhytoKeys (63), p. 77–97. DOI: 10.3897/phytokeys.63.8181.

**RAUTENBERG, ANJA; FILATOV, DMITRY; SVENNBLAD, BODIL; HEIDARI, NAHID; OXELMAN, BENGT (2008):**
**Conflicting phylogenetic signals in the SlX1/Y1 Gene in *Silene*.**
In: BMC Evolutionary Biology 8, p. 299. DOI: 10.1186/1471-2148-8-299.

**RAUTENBERG, ANJA; HATHAWAY, LOUISE; OXELMAN, BENGT; PRENTICE, HONOR C. (2010):**
**Geographic and phylogenetic Patterns in *Silene* Section *Melandrium* (Caryophyllaceae) as inferred from Chloroplast and nuclear DNA Sequences.**
In: Molecular Phylogenetics and Evolution 57 (3), p. 978–991. DOI: 10.1016/j.ympev.2010.08.003.

**SADEGHIAN, SOMAYEH; ZARRE, SHAHIN; RABELER, RICHARD K.; HEUBL, GÜNTHER (2015):**
**Molecular phylogenetic Analysis of *Arenaria* (Caryophyllaceae: Tribe Arenarieae) and its allies inferred from nuclear DNA Internal Transcribed Spacer and Plastid DNA rps16 Sequences.**
In: Botanical Journal of the Linnean Society 178 (4), p. 648–669. DOI: 10.1111/boj.12293.

**SCHEEN, ANNE-CATHRINE; BROCHMANN, CHRISTIAN; BRYSTING, ANNE K.; ELVEN, REIDAR; MORRIS, ASHLEY; SOLTIS, DOUGLAS E. ET AL. (2004):**
**Northern hemisphere Biogeography of *Cerastium* (Caryophyllaceae): Insights from phylogenetic Analysis of noncoding plastidnucleotide Sequences.**
In: American Journal of Botany 91 (6), p. 943–952. DOI: 10.3732/ajb.91.6.943.

**SLOAN, DANIEL B.; OXELMAN, BENGT; RAUTENBERG, ANJA; TAYLOR, DOUGLAS R. (2009):**
**Phylogenetic Analysis of mitochondrial substitution Rate Variation in the Angiosperm Tribe Sileneae.**
In: BMC Evolutionary Biology 9, p. 260. DOI: 10.1186/1471-2148-9-260.

Sloan, Daniel B.; Triant, Deborah A.; Forrester, Nicole J.; Bergner, Laura M.; Wu, Martin; Taylor, Douglas R. (2014):

**A recurring Syndrome of accelerated Plastid Genome Evolution in the Angiosperm Tribe Sileneae (Caryophyllaceae).**

In: Molecular Phylogenetics and Evolution 72, p. 82–89. DOI: 10.1016/j.ympev.2013.12.004.

Smissen, Rob D.O.B.; Clement, John C.; Garnock-Jones, Philip J.; Chambers, Geoffrey K. (2002):

**Subfamilial Relationships within Caryophyllaceae as inferred from 5' ndhF Sequences.**

In: American Journal of Botany 89 (8), p. 1336–1341. DOI: 10.3732/ajb.89.8.1336.

Vaezi, Jamil; Behroozian, Maryam; Memariani, Farshid; Joharchi, Mohammad Reza (2014):

**Dian*thus pseudocrinitus* (Caryophyllaceae), a new Species from Northeast of Iran identified by morphological and molecular Data.**

In: Phytotaxa 156 (2), p. 59. DOI: 10.11646/phytotaxa.156.2.1.

Valcárcel, Virginia; Vargas, Pablo; Feliner, Gonzalo Nieto (2006):

**Phylogenetic and Phylogeographic Analysis of the Western Mediterranean *Arenaria* Section *Plinthine* (Caryophyllaceae) based on Nuclear, Plastid, and morphological Markers.**

In: Taxon 55 (2), p. 297–312. DOI: 10.2307/25065579.

Willyard, Ann; Wallace, Lisa E.; Wagner, Warren L.; Weller, Stephen G.; Sakai, Ann K.; Nepokroeff, Molly (2011):

**Estimating the Species Tree for Hawaiian *Schiedea* (Caryophyllaceae) from multiple loci in the presence of reticulate Evolution.**

In: Molecular Phylogenetics and Evolution 60 (1), p. 29–48. DOI: 10.1016/j.ympev.2011.04.001.

### Caryophyllales

Brockington, Samuel F.; Alexandre, Roolse; Ramdial, Jeremy; Moore, Michael J.; Crawley, Sunny S.; Dhingra, Amit et al. (2009):

**Phylogeny of the Caryophyllales sensu lato: Revisiting Hypotheses on Pollination Biology and Perianth Differentiation in the Core Caryophyllales.**

In: International Journal of Plant Sciences 170 (5), p. 627–643. DOI: 10.1086/597785.

Brockington, Samuel F.; Yang, Ya; Gandia-Herrero, Fernando; Covshoff, Sarah; Hibberd, Julian M.; Sage, Rowan F. et al. (2015):

**Lineage-specific Gene Radiations underlie the Evolution of novel Betalain Pigmentation in Caryophyllales.**

In: the new Phytologist 207 (4), p. 1170–1180. DOI: 10.1111/nph.13441.

Crawley, Sunny S.; Hilu, Khidir W. (2012):

**Impact of missing Data, Gene choice, and Taxon Sampling on phylogenetic reconstruction: the Caryophyllales (Angiosperms).**

In: Plant Systematics and Evolution 298 (2), p. 297–312. DOI: 10.1007/s00606-011-0544-x.

Cuénoud, Philippe; Savolainen, Vincent; Chatrou, Lars W.; Powell, Martyn P.; Grayer, Renée J.; Chase, Mark W. (2002):

**Molecular Phylogenetics of Caryophyllales based on nuclear 18s rDNA and Plastid rbcL, atpB, and matK DNA Sequences.**

In: American Journal of Botany 89 (1), p. 132–144. DOI: 10.3732/ajb.89.1.132.

Hernández-Ledesma, Patricia; Berendsohn, Walter G.; Borsch, Thomas; Mering, Sabine; Akhani, Hossein; Arias, Salvador et al. (2015):

**A taxonomic backbone for the global synthesis of Species Diversity in the Angiosperm Order Caryophyllales.**
In: Willdenowia 45 (3), p. 281. DOI: 10.3372/wi.45.45301.

MOORE, ABIGAIL J.; VOS, JURRIAAN M. DE; HANCOCK, LILLIAN P.; GOOLSBY, ERIC; EDWARDS, ERIKA J. (2018):
**Targeted Enrichment of Large Gene Families for phylogenetic Inference: Phylogeny and molecular Evolution of Photosynthesis Genes in the *Portullugo* Clade (Caryophyllales).**
In: Systematic Biology 67 (3), p. 367–383. DOI: 10.1093/sysbio/syx078.

OCAMPO, GILBERTO; COLUMBUS, TRAVIS J. (2010):
**Molecular Phylogenetics of Suborder Cactineae (Caryophyllales), including Insights into photosynthetic Diversification and historical Biogeography.**
In: American Journal of Botany 97 (11), p. 1827–1847. DOI: 10.3732/ajb.1000227.

### Casuarinaceae

ROSE, PHILLIP F.; WILSON, KAREN L.; TELFORD, IAN R.H.; LAMONT, ROBERT W.; BRUHL, JEREMY J. (2014):
**Multiple lines of Evidence clarify limits of *Allocasuarina thalassoscopica*, *A. defungens* and *A. littoralis* (*Allocasuarina* Sect. *Cylindropitys*, Casuarinaceae).**
In: Australian Systematic Botany 27 (4), p. 257–281. DOI: 10.1071/SB14009.

SOGO, AKIKO; SETOGUCHI, HIROAKI; NOGUCHI, JUNKO; JAFFRÉ, TANGUY; TOBE, HIROSHI (2001):
**Molecular Phylogeny of Casuarinaceae based on rbcL and matK Gene Sequences.**
In: Journal of Plant Research 114 (4), p. 459–464. DOI: 10.1007/PL00014011.

STEANE, DOROTHY A.; WILSON, KAREN L.; HILL, ROBERT P. (2003):
**Using matK Sequence Data to unravel the Phylogeny of Casuarinaceae.**
In: Molecular Phylogenetics and Evolution 28 (1), p. 47–59. DOI: 10.1016/S1055-7903(03)00028-9.

### Celastraceae

BACON, CHRISTINE D.; SIMMONS, MARK P.; ARCHER, ROBERT H.; ZHAO, LIANG-CHENG; ANDRIANTIANA, JACKY L. (2016):
**Biogeography of the Malagasy Celastraceae: Multiple independent Origins followed by widespread Dispersal of Genera from Madagascar.**
In: Molecular Phylogenetics and Evolution 94 (Pt A), p. 365–382. DOI: 10.1016/j.ympev.2015.09.013.

BIRAL, LEONARDO; SMIDT, ERIC DE CAMARGO; BOLSON, MÔNICA; LOMBARDI, JULIO ANTONIO (2015):
**A new Species of *Maytenus* (Celastraceae) from the Brazilian Atlantic Forest, with Evidence of molecular Phylogeny, and two new Synonyms for *Maytenus floribunda*.**
In: Phytotaxa 231 (1), p. 53. DOI: 10.11646/phytotaxa.231.1.5.

COUGHENOUR, JENNIFER M.; SIMMONS, MARK P.; LOMBARDI, JULIO ANTONIO; YAKOBSON, KENDRA; ARCHER, ROBERT H. (2011):
**Phylogeny of Celastraceae Subfamily Hippocrateoideae inferred from morphological Characters and nuclear and Plastid Loci.**
In: Molecular Phylogenetics and Evolution 59 (2), p. 320–330. DOI: 10.1016/j.ympev.2011.02.017.

COUGHENOUR, JENNIFER M.; SIMMONS, MARK P.; LOMBARDI, JULIO ANTONIO; CAPPA, JENNIFER J. (2010):
**Phylogeny of Celastraceae Subfamily Salacioideae and Tribe Lophopetaleae inferred from morphological Characters and Nuclear and Plastid Genes.**
In: Systematic Botany 35 (2), p. 358–367. DOI: 10.1600/036364410791638289.

DARBYSHIRE, IAIN; SIMMONS, MARK P.; CAPPA, JENNIFER J.; BRETELER, FRANS J.; BUERKI, SVEN (2016):

**_Pleurostylia serrulata_ and two allied new Species from Africa are actually members of the New World _Crossopetalum_ (Celastraceae).**
In: Systematic Botany 41 (4), p. 851–864. DOI: 10.1600/036364416X693955.

Islam, Melissa Bauer; Simmons, Mark P.; Archer, Robert H. (2006):
**Phylogeny of the _Elaeodendron_ Group (Celastraceae) inferred from morphological Characters and Nuclear and Plastid Genes.**
In: Systematic Botany 31 (3), p. 512–524.

Li, Yan-Nan; Xie, Lei; Li, Jin-Yu; Zhang, Zhi-Xiang (2014):
**Phylogeny of _Euonymus_ inferred from molecular and morphological Data.**
In: Journal of Systematics and Evolution 52 (2), p. 149–160. DOI: 10.1111/jse.12068.

Mu, Xian-Yun; Zhao, Liang-Cheng; Zhang, Zhi-Xiang (2012):
**Phylogeny of _Celastrus_ L. (Celastraceae) inferred from two nuclear and three Plastid Markers.**
In: Journal of Plant Research 125 (5), p. 619–630. DOI: 10.1007/s10265-012-0484-8.

Simmons, Mark P.; Barrie, Fred R. (2014):
**_Haydenoxylon_, a Replacement Name for _Haydenia_ (Celastraceae).**
In: Novon: A Journal for Botanical Nomenclature 23 (2), p. 224–225. DOI: 10.3417/2014011.

Simmons, Mark P.; Cappa, Jennifer J.; Archer, Robert H.; Ford, Andrew J.; Eichstedt, Dedra; Clevinger, Curtis C. (2008):
**Phylogeny of the Celastreae (Celastraceae) and the Relationships of _Catha edulis_ (qat) inferred from morphological Characters and nuclear and Plastid Genes.**
In: Molecular Phylogenetics and Evolution 48 (2), p. 745–757. DOI: 10.1016/j.ympev.2008.04.039.

Simmons, Mark P.; Clevinger, Curtis C.; Savolainen, Vincent; Archer, Robert H.; Mathews, Sarah; Doyle, Jeff J. (2001):
**Phylogeny of the Celastraceae inferred from phytochrome B Gene Sequence and Morphology.**
In: American Journal of Botany 88 (2), p. 313–325. DOI: 10.2307/2657021.

Simmons, Mark P.; McKenna, Miles J.; Bacon, Christine D.; Yakobson, Kendra; Cappa, Jennifer J.; Archer, Robert H.; Ford, Andrew J. (2012):
**Phylogeny of Celastraceae Tribe Euonymeae inferred from morphological Characters and nuclear and Plastid Genes.**
In: Molecular Phylogenetics and Evolution 62 (1), p. 9–20. DOI: 10.1016/j.ympev.2011.08.022.

Simmons, Mark P.; Savolainen, Vincent; Clevinger, Curtis C.; Archer, Robert H.; Davis, J. I. (2001):
**Phylogeny of the Celastraceae inferred from 26s nuclear ribosomal DNA, Phytochrome B, rbcL, atpB, and Morphology.**
In: Molecular Phylogenetics and Evolution 19 (3), p. 353–366. DOI: 10.1006/mpev.2001.0937.

### Celastrales

Zhang, Li-Bing; Simmons, Mark P. (2006):
**Phylogeny and Delimitation of the Celastrales inferred from Nuclear and Plastid Genes.**
In: Systematic Botany 31 (1), p. 122–137.

### Centrolepidaceae

Sokoloff, Dmitry D.; Remizowa, Margarita V.; Barrett, Matthew D.; Conran, John G.; Rudall, Paula J. (2015):

**Morphological Diversity and Evolution of Centrolepidaceae (Poales), a species-poor Clade with diverse body plans and developmental Patterns.**
In: American Journal of Botany 102 (8), p. 1219–1249. DOI: 10.3732/ajb.1400434.

### Chloranthaceae

ANTONELLI, ALEXANDRE; SANMARTÍN, ISABEL (2011):
**Mass extinction, gradual Cooling, or rapid Radiation? Reconstructing the spatiotemporal Evolution of the ancient Angiosperm Genus *Hedyosmum* (Chloranthaceae) using empirical and simulated Approaches.**
In: Systematic Biology 60 (5), p. 596–615. DOI: 10.1093/sysbio/syr062.

DOYLE, JAMES A.; ENDRESS, PETER K. (2014):
**Integrating Early Cretaceous Fossils into the Phylogeny of Living Angiosperms. ANITA Lines and Relatives of Chloranthaceae.**
In: International Journal of Plant Sciences 175 (5), p. 555–600. DOI: 10.1086/675935.

KONG, HONG-ZHI; CHEN, ZHI-DUAN; LU, AN-MING (2002):
**Phylogeny of *Chloranthus* (Chloranthaceae) based on nuclear ribosomal ITS and Plastid Trnl-f Sequence Data.**
In: American Journal of Botany 89 (6), p. 940–946. DOI: 10.3732/ajb.89.6.940.

ZHANG, LI-BING; RENNER, SUSANNE P. (2003):
**The deepest splits in Chloranthaceae as resolved by Chloroplast Sequences.**
In: International Journal of Plant Sciences 164 (5), p. 383–392.

### Chrysophyllaceae

BARDON, LÉA; CHAMAGNE, JULIETTE; DEXTER, KYLE G.; SOTHERS, CYNTHIA A.; PRANCE, GHILLEAN T.; CHAVE, JÉRÔME (2013):
**Origin and Evolution of Chrysobalanaceae: Insights into the Evolution of plants in the Neotropics.**
In: Botanical Journal of the Linnean Society 171, p. 19–37.

BARDON, LÉA; SOTHERS, CYNTHIA A.; PRANCE, GHILLEAN T.; MALÉ, PIERRE-JEAN G.; XI, ZHENXIANG; DAVIS, CHARLES C. ET AL. (2016):
**Unraveling the biogeographical History of Chrysobalanaceae from Plastid Genomes.**
In: American Journal of Botany 103 (6), p. 1089–1102. DOI: 10.3732/ajb.1500463.

SOTHERS, CYNTHIA A.; PRANCE, GHILLEAN T.; BUERKI, SVEN; KOK, ROGIER P.J. DE; CHASE, MARK W. (2014):
**Taxonomic novelties in Neotropical Chrysobalanaceae: towards a monophyletic *Couepia*.**
In: Phytotaxa 172 (3), p. 176. DOI: 10.11646/phytotaxa.172.3.2.

YAKANDAWALA, DEEPTHI; MORTON, CYNTHIA M.; PRANCE, GHILLEAN T. (2010):
**Phylogenetic Relationships of the Chrysobalanaceae inferred from Chloroplast, Nuclear, and morphological Data.**
In: Annals of the Missouri Botanical Garden 97 (2), p. 259–281. DOI: 10.3417/2007175.

### Cistaceae

CIVEYREL, LAURE; LECLERCQ, JULIE; DEMOLY, JEAN-PIERRE; AGNAN, YANNICK; QUÈBRE, NICOLAS; PÉLISSIER, CÉLINE; OTTO, THIERRY (2011):
**Molecular Systematics, Character Evolution, and Pollen Morphology of *Cistus* and *Halimium* (Cistaceae).**
In: Plant Systematics and Evolution 295 (1-4), p. 23–54. DOI: 10.1007/s00606-011-0458-7.

**Guzmán, Beatriz; Vargas, Pablo (2005):**

**Systematics, Character Evolution, and Biogeography of *Cistus* L. (Cistaceae) based on ITS, trnL-trnF, and matK Sequences.**

In: Molecular Phylogenetics and Evolution 37 (3), p. 644–660. DOI: 10.1016/j.ympev.2005.04.026.

**Guzmán, Beatriz; Vargas, Pablo (2009):**

**Historical Biogeography and Character Evolution of Cistaceae (Malvales) based on Analysis of Plastid rbcL and trnL-trnF Sequences.**

In: Organisms Diversity and Evolution 9 (2), p. 83–99. DOI: 10.1016/j.ode.2009.01.001.

### Cleomaceae

**Feodorova, Tatiana A.; Voznesenskaya, Elena V.; Edwards, Gerald E.; Roalson, Eric H. (2010):**

**Biogeographic Patterns of Diversification and the Origins of C4 in *Cleome* (Cleomaceae).**

In: Systematic Botany 35 (4), p. 811–826. DOI: 10.1600/036364410X539880.

**Inda, Luis A.; Torrecilla, Pedro; Catalán, Pilar; Ruiz-Zapata, Thirza (2008):**

**Phylogeny of *Cleome* L. and its close relatives *Podandrogyne* Ducke and *Polanisia* Raf. (Cleomoideae, Cleomaceae) based on Analysis of nuclear ITS Sequences and Morphology.**

In: Plant Systematics and Evolution 274 (1-2), p. 111–126. DOI: 10.1007/s00606-008-0026-y.

**Patchell, Melanie J.; Roalson, Eric H.; Hall, Jocelyn C. (2014):**

**Resolved Phylogeny of Cleomaceae based on all three Genomes.**

In: Taxon 63 (2), p. 315–328. DOI: 10.12705/632.17.

**Riser, James P.; Cardinal-McTeague, Warren M.; Hall, Jocelyn C.; Hahn, William J.; Sytsma, Kenneth J.; Roalson, Eric H. (2013):**

**Phylogenetic Relationships among the North American cleomoids (Cleomaceae): a Test of Iltis's Reduction Series.**

In: American Journal of Botany 100 (10), p. 2102–2111. DOI: 10.3732/ajb.1300096.

**Roalson, Eric H.; Hall, Jocelyn C.; Riser II, James P.; Cardinal-McTeague, Warren M.; Cochrane, Theodore S.; Sytsma, Kenneth J. (2015):**

**A Revision of generic Boundaries and Nomenclature in the North American cleomoid Clade (Cleomaceae).**

In: Phytotaxa 205 (3), p. 129. DOI: 10.11646/phytotaxa.205.3.1.

**Tamboli, Asif Shabodin; Patil, Swapnil Mahadeo; Gholave, Avinash Ramchandra; Kadam, Suhas Kishor; Kotibhaskar, Shreya Vijaykumar; Yadav, Shrirang Ramchandra; Govindwar, Sanjay Prabhu (2016):**

**Phylogenetic Analysis, genetic Diversity and Relationships between the recently segregated Species of *Corynandra* and *Cleoserrata* from the Genus *Cleome* using DNA barcoding and molecular Markers.**

In: Comptes Rendus Biologies 339 (3-4), p. 123–132. DOI: 10.1016/j.crvi.2016.02.005.

### Clethraceae

**Anderberg, Arne A.; Zhang, Xiaoping (2002):**

**Phylogenetic Relationships of Cyrillaceae and Clethraceae (Ericales) with special Emphasis on the Genus *Purdiaea* Planch.**

In: Organisms Diversity and Evolution 2 (2), p. 127–137. DOI: 10.1078/1439-6092-00036.

**Fior, Simone; Karis, Per Ola; Anderberg, Arne A. (2003):**

**Phylogeny, Taxonomy, and Systematic Position of *Clethra* (Clethraceae, Ericales) with Notes on Biogeography: Evidence from Plastid and Nuclear DNA Sequences.**
In: International Journal of Plant Sciences 164 (6), p. 997–1006.

### Clusiaceae

CADDAH, MAYARA KRASINSKI; CAMPOS, TATIANA; ZUCCHI, MARIA IMACULADA; SOUZA, ANETE PEREIRA; BITTRICH, VOLKER; AMARAL, MARIA DO CARMO ESTANISLAU DO (2013):
**Species boundaries inferred from Microsatellite Markers in the *Kielmeyera coriacea* complex (Calophyllaceae) and Evidence of asymmetric Hybridization.**
In: Plant Systematics and Evolution 299 (4), p. 731–741. DOI: 10.1007/s00606-012-0755-9.

FURNESS, CAROL A. (2012):
**Pollen Evolution in the Clusioid Clade (Malpighiales).**
In: International Journal of Plant Sciences 173 (9), p. 1055–1082. DOI: 10.1086/667614.

GUSTAFSSON, MATS H.G.; BITTRICH, VOLKER; STEVENS, PETER F. (2002):
**Phylogeny of Clusiaceae based on rbcL Sequences.**
In: International Journal of Plant Sciences 163 (6), p. 1045–1054.

RAJKUMAR, S.; JANARTHANAM, M.K. (2007):
***Agasthiyamalaia* (Clusiaceae) A new Genus for *Poeciloneuron pauciflorum* an endemic and endangered Tree of Western Ghats, India.**
In: Journal of the Botanical Research Institute of Texas 1 (1), p. 129–133.

RUHFEL, BRADLEY R.; STEVENS, PETER F.; DAVIS, CHARLES C. (2013):
**Combined morphological and molecular Phylogeny of the Clusioid Clade (Malpighiales) and the Placement of the Ancient Rosid Macrofossil *Paleoclusia*.**
In: International Journal of Plant Sciences 174 (6), p. 910–936. DOI: 10.1086/670668.

SWEENEY, PATRICK W. (2008):
**Phylogeny and Floral Diversity in the Genus *Garcinia* (Clusiaceae) and Relatives.**
In: International Journal of Plant Sciences 169 (9), p. 1288–1303. DOI: 10.1086/591990.

### Codonaceae

WEIGEND, MAXIMILIAN (2010):
**Codonaceae - a newly required Family name in Boraginales.**
In: Phytotaxa 10, p. 26–30.

### Colchicaceae

CASE, ANDREA L.; GRAHAM, SEAN W.; MACFARLANE, TERRY D.; BARRETT, SPENCER C. H. (2008):
**A phylogenetic Study of evolutionary Transitions in Sexual Systems in Australasian *Wurmbea* (Colchicaceae).**
In: International Journal of Plant Sciences 169 (1), p. 141–156. DOI: 10.1086/523368.

DEL HOYO, ALBERTO; GARCÍA-MARÍN, JOSÉ LUIS; PEDROLA-MONFORT, JOAN (2009):
**Temporal and spatial Diversification of the African disjunct Genus *Androcymbium* (Colchicaceae).**
In: Molecular Phylogenetics and Evolution 53 (3), p. 848–861. DOI: 10.1016/j.ympev.2009.08.005.

LEKHAK, MANOJ M.; SURVESWARAN, SIDDHARTHAN; YADAV, SHRIRANG RAMCHANDRA (2016):
**Generic identity of *Camptorrhiza indica* (Colchicaceae) based on Cytogenetics and molecular Phylogenetics.**
In: Journal of Systematics and Evolution 54 (1), p. 75–82. DOI: 10.1111/jse.12165.

NGUYEN, THI P. A.; KIM, JUNG SUNG; KIM, JOO-HWAN (2013):

**Molecular phylogenetic Relationships and Implications for the Circumscription of Colchicaceae (Liliales).**
In: Botanical Journal of the Linnean Society 172 (3), p. 255–269. DOI: 10.1111/boj.12037.

TAMURA, MINORU N.; YAMADA, MASAHIRO; FUSE, SHIZUKA; HOTA, MITSURU (2013):
**Molecular Phylogeny and Taxonomy of the Genus *Disporum* (Colchicaceae).**
In: Acta Phytotaxonomica Geobotanica 64 (3), p. 137–147.

### Combretaceae

BARRETT, RUSSELL L. (2015):
**Examining range disjunctions in Australian *Terminalia* (Combretaceae) with taxonomic Revision of the *T. canescens* and *T. cunninghamii* Species complexes.**
In: Australian Systematic Botany 28 (1), p. 23–45. DOI: 10.1071/SB14026.

MAURIN, OLIVIER; CHASE, MARK W.; JORDAAN, MARIE; VAN DER BANK, MICHELLE (2010):
**Phylogenetic Relationships of Combretaceae inferred from nuclear and Plastid DNA Sequence Data: Implications for generic Classification.**
In: Botanical Journal of the Linnean Society 162 (3), p. 453–476. DOI: 10.1111/j.1095-8339.2010.01027.x.

TAN, FENGXIAO; SHI, SU-HUA; ZHONG, YANG; GONG, XUN; WANG, YUGUO (2002):
**Phylogenetic Relationships of Combretoideae (Combretaceae) inferred from plastid, nuclear Gene and spacer Sequences.**
In: Journal of Plant Research 115 (6), p. 475–481. DOI: 10.1007/s10265-002-0059-1.

### Commelinaceae

AONA, LIDYANNE Y. S.; BITTRICH, VOLKER; AMARAL, MARIA DO CARMO E. (2017):
**Taxonomic novelties in Brazilian *Dichorisandra* (Commelinaceae): *D. sagittata* sp. nov. and *D. glabrescens* stat. nov.**
In: Brittonia 69 (2), p. 209–217. DOI: 10.1007/s12228-016-9456-y.

AONA-PINHEIRO, L.Y.S.; AMARAL, MARIA DO CARMO E. (2012):
**Four new Species of *Dichorisandra* J.C.Mikan (Commelinaceae) from Southeast Brazil.**
In: Phytotaxa 48 (7), p. 22.

BERGAMO, STEPHANIE (2003):
**A phylogenetic Evaluation of *Callisia* Loefl. (Commelinaceae) based on molecular Data.**
Dissertation. University of Gergia, Athens, Georgia, USA.

BURNS, JEAN H.; FADEN, ROBERT B.; STEPPAN, SCOTT J. (2011):
**Phylogenetic Studies in the Commelinaceae Subfamily Commelinoideae inferred from Nuclear Ribosomal and Chloroplast DNA Sequences.**
In: Systematic Botany 36 (2), p. 268–276. DOI: 10.1600/036364411X569471.

EVANS, TIMOTHY M.; SYTSMA, KENNETH J.; FADEN, ROBERT B.; GIVNISH, THOMAS J. (2003):
**Phylogenetic Relationships in the Commelinaceae: II. A cladistic Analysis of rbcL Sequences and Morphology.**
In: Systematic Botany 28 (2), p. 270–292.

HARDY, CHRISTOPHER R.; FADEN, ROBERT B. (2004):
***Plowmanianthus*, a new Genus of Commelinaceae with five new Species from Tropical America.**
In: Systematic Botany 29 (2), p. 316–333. DOI: 10.1600/036364404774195511.

**Kelly, Shana M.; Evans, Timothy M. (2014):**
**Phylogenetic Analysis of the African Plant Genus *Aneilema* (family Commelinaceae) based on Chloroplast DNA Sequences.**
Honors Projects 285. Grand valley State University, Grand Valley.

### Convolvulaceae

**Breteler, Frans J. (2013):**
**Revision of *Calycobolus* (Convolvulaceae) in continental Africa.**
In: Plant Ecology and Evolution 146 (3), p. 328–350. DOI: 10.5091/plecevo.2013.856.

**Buril, Maria Teresa; Simões, Ana Rita; Carine, Mark A.; Alves, Marccus V. (2015):**
***Daustinia*, a replacement name for *Austinia* (Convolvulaceae).**
In: Phytotaxa 197 (1), p. 60. DOI: 10.11646/phytotaxa.197.1.8.

**Carine, Mark A.; Russell, Stephen J.; Santos-Guerra, Arnoldo; Francisco-Ortega, Javier (2004):**
**Relationships of the Macaronesian and Mediterranean Floras: molecular Evidence for Multiple Colonizations into Macaronesia and Back-Colonization of the Continent in *Convolvulus* (Convolvulaceae).**
In: American Journal of Botany 91 (7), p. 1070–1085.

**Costea, Mihai; Aiston, Fiona; Stefanović, Saša (2008):**
**Species Delimitation, phylogenetic Relationships, and two new Species in the *Cuscuta gracillima* complex (Convolvulaceae).**
In: Botany 86 (7), p. 670–681. DOI: 10.1139/B08-030.

**Costea, Mihai; Ruiz, Ignacio García; Stefanović, Saša (2011):**
**Systematics of "horned" dodders: phylogenetic Relationships, Taxonomy, and two new Species within the *Cuscuta chapalana* complex (Convolvulaceae).**
In: Botany 89 (10), p. 715–730. DOI: 10.1139/B11-049.

**Costea, Mihai; Stefanović, Saša (2010):**
**Evolutionary History and Taxonomy of the *Cuscuta umbellata* complex (Convolvulaceae): Evidence of extensive Hybridization from discordant nuclear and Plastid Phylogenies.**
In: Taxon 59 (6), p. 1783–1800.

**Demissew, Sebsebe (2001):**
**A Synopsis of the Genus *Merremia* (Convolvulaceae) in the Flora of Ethiopia and Eritrea.**
In: Kew Bulletin 56 (4), p. 931–943.

**Demissew, Sebsebe; Mill, Robert R. (2009):**
**Revision of the Genus *Seddera* (Convolvulaceae).**
In: Kew Bulletin 64, p. 197–233.

**Duncan, Tanya M.; Rausher, Mark D. (2013):**
**Morphological and genetic differentiation and reproductive Isolation among closely related Taxa in the *Ipomoea* series *Batatas*.**
In: American Journal of Botany 100 (11), p. 2183–2193. DOI: 10.3732/ajb.1200467.

**Eserman, Lauren A.; Tiley, George P.; Jarret, Robert L.; Leebens-Mack, Jim H.; Miller, Richard E. (2014):**
**Phylogenetics and Diversification of morning Glories (tribe Ipomoeeae, Convolvulaceae) based on whole Plastome Sequences.**
In: American Journal of Botany 101 (1), p. 92–103. DOI: 10.3732/ajb.1300207.

**García, Miguel Á.; Martín, Mariá P. (2007):**
**Phylogeny of *Cuscuta* Subgenus *Cuscuta* (Convolvulaceae) based on nrDNA ITS and Chloroplast trnL Intron Sequences.**
In: Systematic Botany 32 (4), p. 899–916.

**Jayasuriya, K. M. G. Gehan; Baskin, Jerry M.; Geneve, Robert L.; Baskin, Carol C. (2009):**
**Phylogeny of Seed Dormancy in Convolvulaceae, Subfamily Convolvuloideae (Solanales).**
In: Annals of Botany 103 (1), p. 45–63. DOI: 10.1093/aob/mcn217.

**Miller, Richard E.; McDonald, J. Andrew; Manos, Paul p. (2004):**
**Systematics of *Ipomoea* Subgenus *Quamoclit* (Convolvulaceae) based on ITS Sequence Data and A Bayesian phylogenetic Analysis.**
In: American Journal of Botany 91 (8), p. 1208–1218.

**Miller, Richard E.; Rausher, Mark D.; Manos, Paul p. (1999):**
**Phylogenetic Systematics of *Ipomoea* (Convolvulaceae) based on ITS and Waxy Sequences.**
In: Systematic Botany 24 (2), p. 209–227.

**Pastore, Mayara; Simão-Bianchini, Rosangela (2015):**
**Taxonomic novelties in *Jacquemontia* Choisy (Convolvulaceae) from Southeastern Brazil.**
In: Phytotaxa 221 (2), p. 193. DOI: 10.11646/phytotaxa.221.2.10.

**Rajapakse, Sriyani; Nilmalgoda, Sasanda D.; Molnar, Matthew; Ballard, Robert E.; Austin, Daniel F.; Bohac, Janice R. (2004):**
**Phylogenetic Relationships of the sweetpotato in *Ipomoea* series *Batatas* (Convolvulaceae) based on nuclear β-Amylase Gene Sequences.**
In: Molecular Phylogenetics and Evolution 30 (3), p. 623–632. DOI: 10.1016/S1055-7903(03)00249-5.

**Simões, Ana Rita; Culham, Alastair; Carine, Mark A. (2015):**
**Resolving the unresolved Tribe: a molecular phylogenetic Framework for the Merremieae (Convolvulaceae).**
In: Botanical Journal of the Linnean Society 179, p. 374–387.

**Simões, Ana Rita; Staples, G. W. (2017):**
**Dissolution of Convolvulaceae Tribe Merremieae and a new Classification of the constituent Genera.**
In: Botanical Journal of the Linnean Society 183, p. 561–586.

**Staples, G. W.; Buril, Maria Teresa; Simões, Ana Rita; Govaerts, Rafael (2015):**
**Nomenclatural corrections in Convolvulaceae diversae.**
In: Phytologia 97 (3), p. 219–223.

**Staples, George W. (2007):**
**A Synopsis of *Rivea* (Convolvulaceae).**
In: Edinburgh Journal of Botany 64 (2), p. 213–223. DOI: 10.1017/S0960428607000911.

**Stefanović, Saša; Austin, Daniel F.; Olmstead, Richard G. (2003):**
**Classification of Convolvulaceae: A phylogenetic Approach.**
In: Systematic Botany 28 (4), p. 791–806.

**Stefanović, Saša; Costea, Mihai (2008):**

**Reticulate Evolution in the parasitic Genus *Cuscuta* (Convolvulaceae): over and over again.**
In: Botany 86 (8), p. 791–808. DOI: 10.1139/B08-033.

**STEFANOVIĆ, SAŠA; KRUEGER, LORI; OLMSTEAD, RICHARD G. (2002):**
**Monophyly of the Convolvulaceae and Circumscription of their major Lineages based on DNA Sequences of multiple Chloroplast Loci.**
In: American Journal of Botany 89 (9), p. 1510–1522.

**STEFANOVIĆ, SAŠA; KUZMINA, MARIA (2007):**
**Delimitation of Major Lineages within Cuscuta Subgenus *Grammica* (Convolvulaceae) using Plastid and Nuclear DNA Sequences.**
In: American Journal of Botany 94 (4), p. 568–589.

**STEFANOVIĆ, SAŠA; OLMSTEAD, RICHARD G. (2004):**
**Testing the phylogenetic Position of a parasitic Plant (*Cuscuta*, Convolvulaceae, Asteridae): Bayesian Inference and the parametric Bootstrap on Data drawn from three Genomes.**
In: Systematic Biology 53 (3), p. 384–399. DOI: 10.1080/10635150490445896.

**TRAIPERM, PAWEENA; STAPLES, GEORGE W. (2013):**
**A new Combination in *Argyreia* Lour. (Convolvulaceae).**
In: Adansonia 35 (2), p. 359–363. DOI: 10.5252/a2013n2a7.

**WILKIN, PAUL (1999):**
**A morphological cladistic Analysis of the Ipomoeeae (Convolvulaceae).**
In: Kew Bulletin 54 (4), p. 853–876.

**WOOD, JOHN R. I.; CARINE, MARK A.; HARRIS, DAVID J.; WILKIN, PAUL; WILLIAMS, B.; SCOTLAND, ROBERT W. (2015):**
***Ipomoea* (Convolvulaceae) in Bolivia.**
In: Kew Bulletin 70 (3), p. 30–124. DOI: 10.1007/S12225-015-9592-7.

### Cordiaceae

**GOTTSCHLING, MARC; MILLER, JAMES P. (2006):**
**Clarification of the taxonomic Position of *Auxemma, Patagonula,* and *Saccellium* (Cordiaceae, Boraginales).**
In: Systematic Botany 31 (2), p. 361–367.

### Coriariaceae

**YOKOYAMA, JUN U.N.; SUZUKI, M.; IWATSUKI, KUNIO; HASEBE, M. (2000):**
**Molecular Phylogeny of *Coriaria*, with special Emphasis on the disjunct Distribution.**
In: Molecular Phylogenetics and Evolution 14 (1), p. 11–19. DOI: 10.1006/mpev.1999.0672.

### Cornaceae

**FAN, CHUANZHU; PURUGGANAN, MICHAEL D.; THOMAS, DAVID T.; WIEGMANN, BRIAN M.; XIANG, JENNY QIU-YUN (2004):**
**Heterogeneous Evolution of the Myc-like Anthocyanin regulatory Gene and its phylogenetic utility in *Cornus* L. (Cornaceae).**
In: Molecular Phylogenetics and Evolution 33 (3), p. 580–594. DOI: 10.1016/j.ympev.2004.08.002.

**FAN, CHUANZHU; XIANG, JENNY QIU-YUN (2001):**
**Phylogenetic Relationships within *Cornus* (Cornaceae) based on 26s rDNA Sequences.**
In: American Journal of Botany 88 (6), p. 1131–1138. DOI: 10.2307/2657096.

**THOMAS, QIU-YUN XIANG DAVID T. (2008):**

**Tracking Character Evolution and biogeographic History through time in Cornaceae - Does choice of methods matter?**
In: Journal of Systematics and Evolution 46 (3), p. 349–374.

XIANG, QIU-YUN; THOMAS, DAVID T.; ZHANG, WENHENG; MANCHESTER, STEVEN R.; MURRELL, ZACK (2006):
**Species Level Phylogeny of the Genus *Cornus* (Cornaceae) based on molecular and morphological Evidence-Implications for Taxonomy and Tertiary Intercontinental Migration.**
In: Taxon 55 (1), p. 9–30. DOI: 10.2307/25065525.

ZHANG, WENHENG; XIANG, QIU-YUN JENNY; THOMAS, DAVID T.; WIEGMANN, BRIAN M.; FROHLICH, MICHAEL W.; SOLTIS, DOUGLAS E. (2008):
**Molecular Evolution of Pistillata-like Genes in the dogwood Genus *Cornus* (Cornaceae).**
In: Molecular Phylogenetics and Evolution 47 (1), p. 175–195. DOI: 10.1016/j.ympev.2007.12.022.

### Cornales

LI, JIAN-HUA; ZHANG, ZHI-HONG (2010):
**Sequences of 72 Plastid Genes support the early Divergence of Cornales in the Asterids.**
In: Journal of Systematics and Evolution 48 (6), p. 426–434. DOI: 10.1111/j.1759-6831.2010.00098.x.

XIANG, QIU-YUN JENNY; THOMAS, DAVID T.; XIANG, QIAO PING (2011):
**Resolving and dating the Phylogeny of Cornales - Effects of Taxon Sampling, Data partitions, and fossil Calibrations.**
In: Molecular Phylogenetics and Evolution 59 (1), p. 123–138. DOI: 10.1016/j.ympev.2011.01.016.

### Corynocarpaceae

WAGSTAFF, STEVEN J.; DAWSON, MURRAY I. (2000):
**Classification, Origin and Patterns of Diversification of *Corynocarpus* (Corynocarpaceae) inferred from DNA Sequences.**
In: Systematic Botany 25 (1), p. 134–149.

### Costaceae

ANDRE, THIAGO JOSE DE CARVALHO; SPECHT, CHELSEA D.; SALZMAN, SHAYLA; PALMA-SILVA, CLARISSE; WENDT, TANIA (2015):
**Evolution of Species Diversity in the Genus *Chamaecostus* (Costaceae): molecular Phylogenetics and morphometric approaches.**
In: Phytotaxa 204 (4), p. 265. DOI: 10.11646/phytotaxa.204.4.3.

GOVAERTS, RAFAEL (2013):
***Hellenia* Retz., the correct name for *Cheilocostus* C.D.Specht (Costaceae).**
In: Phytotaxa 151 (1), p. 63. DOI: 10.11646/phytotaxa.151.1.7.

SPECHT, CHELSEA D.; STEVENSON, DENNIS WM. (2006):
**A new Phylogeny-based Generic Classification of Costaceae (Zingiberales).**
In: Taxon 55 (1), p. 153. DOI: 10.2307/25065537.

### Crassulaceae

ACEVEDO-ROSAS, RAÚL; CAMERON, KENNETH M.; SOSA, VICTORIA; PELL, SUSAN K. (2004):
**A molecular phylogenetic Study of *Graptopetalum* (Crassulaceae) based on ETS, ITS, Rpl16, and Trnl-f Nucleotide Sequences.**
In: American Journal of Botany 91 (7), p. 1099–1104.

**Acevedo-Rosas, Raúl; Sosa, Victoria; Lorea, Francisco G. (2004):**
**Phylogenetic Relationships and morphological Patterns in *Graptopetalum* (Crassulaceae).**
In: Brittonia 56 (2), p. 185–194.

**Carrillo-Reyes, Pablo; Sosa, Victoria; Mort, Mark E. (2008):**
***Thompsonella* and the "*Echeveria* Group" (Crassulaceae): phylogenetic Relationships based on molecular and morphological characters.**
In: Taxon 57 (3), p. 863–874.

**Carrillo-Reyes, Pablo; Sosa, Victoria; Mort, Mark E. (2009):**
**Molecular Phylogeny of the Acre Clade (Crassulaceae): dealing with the lack of definitions for *Echeveria* and *Sedum*.**
In: Molecular Phylogenetics and Evolution 53 (1), p. 267–276. DOI: 10.1016/j.ympev.2009.05.022.

**Fairfield, K. N.; Mort, Mark E.; Santos-Guerra, Arnoldo (2004):**
**Phylogenetics and Evolution of the Macaronesian Members of the Genus *Aichryson* (Crassulaceae) inferred from nuclear and Chloroplast Sequence Data.**
In: Plant Systematics and Evolution 248 (1-4), p. 71–83. DOI: 10.1007/s00606-004-0190-7.

**Gontcharova, Svetlana B.; Gontcharov, Andrey A. (2009):**
**Molecular Phylogeny and Systematics of flowering Plants of the Family Crassulaceae DC.**
In: Molecular Biology 43 (5), p. 794–803. DOI: 10.1134/S0026893309050112.

**Klein, Johannes T.; Kadereit, Joachim W. (2015):**
**Phylogeny, Biogeography, and Evolution of edaphic Association in the European Oreophytes *Sempervivum* and *Jovibarba* (Crassulaceae).**
In: International Journal of Plant Sciences 176 (1), p. 44–71. DOI: 10.1086/677948.

**Kozyrenko, Marina M.; Gontcharova, Svetlana B.; Gontcharov, Andrey A. (2013):**
**Phylogenetic Relationships among *Orostachys* subsection *Orostachys* Species (Crassulaceae) based on nuclear and Chloroplast DNA Data.**
In: Journal of Systematics and Evolution 51 (5), p. 578–589. DOI: 10.1111/jse.12016.

**Mort, Mark E.; Levsen, Nicholas D.; Randle, Christopher P.; van Jaarsveld, Ernst; Palmer, Annie (2005):**
**Phylogenetics and Diversification of *Cotyledon* (Crassulaceae) inferred from Nuclear and Chloroplast DNA Sequence Data.**
In: American Journal of Botany 92 (7), p. 1170–1176.

**Mort, Mark E.; Soltis, Douglas E.; Soltis, Pamela S.; Francisco-Ortega, Javier; Santos-Guerra, Arnoldo (2001):**
**Phylogenetic Relationships and Evolution of Crassulaceae inferred from matK Sequence Data.**
In: American Journal of Botany 88 (1), p. 76–91.

**Mort, Mark E.; Soltis, Douglas E.; Soltis, Pamela S.; Santos-Guerra, Arnoldo; Francisco-Ortega, Javier (2007):**
**Physiological Evolution and Association between Physiology and Growth Form in *Aeonium* (Crassulaceae).**
In: Taxon 56 (2), p. 453–464.

**Mort, Mark Eugene; Randle, Christopher P.; Burgoyne, Priscilla M.; Smith, Gideon F.; Jaarsveld, Ernst van; Hopper, Stephen D. (2009):**
**Analyses of cpDNA matK Sequence Data place *Tillaea* (Crassulaceae) within *Crassula*.**

In: Plant Systematics and Evolution 283 (3-4), p. 211–217. DOI: 10.1007/s00606-009-0227-z.

NIKULIN, ARTHUR YU.; NIKULIN, VYACHESLAV YU.; GONCTHAROVA, SVETLANA B.; GONTCHAROV, ANDREY A. (2015):
**ITS rDNA Sequence comparisons resolve phylogenetic Relationships in *Orostachys* subsection *Appendiculatae* (Crassulaceae).**
In: Plant Systematics and Evolution 301 (5), p. 1441–1453. DOI: 10.1007/s00606-014-1165-y.

NIKULIN, VYACHESLAV YU.; GONTCHAROVA, SVETLANA B.; STEPHENSON, RAY; GONTCHAROV, ANDREY A. (2016):
**Phylogenetic Relationships between *Sedum* L. and related Genera (Crassulaceae) based on ITS rDNA Sequence comparisons.**
In: Flora - Morphology, Distribution, Functional Ecology of Plants 224, p. 218–229. DOI: 10.1016/j.flora.2016.08.003.

ZHANG, JIAN-QIANG; MENG, SHI-YONG; ALLEN, GERALDINE A.; WEN, JUN; RAO, GUANG-YUAN (2014):
**Rapid Radiation and Dispersal out of the Qinghai-Tibetan Plateau of an alpine plant Lineage *Rhodiola* (Crassulaceae).**
In: Molecular Phylogenetics and Evolution 77, p. 147–158. DOI: 10.1016/j.ympev.2014.04.013.

### Crypteroniaceae

CONTI, ELENA; ERIKSSON, TORSTEN; SCHÖNENBERGER, JÜRGEN; SYTSMA, KENNETH J.; BAUM, DAVID A. (2002):
**Early Tertiary Out-of-India Dispersal of Crypteroniaceae: Evidence from Phylogeny and molecular Dating.**
In: Evolution 56 (10), p. 1931–1942. DOI: 10.1554/0014-3820(2002)056[1931:ETOOID]2.0.CO;2.

RUTSCHMANN, FRANK; ERIKSSON, TORSTEN; SCHÖNENBERGER, JÜRGEN; CONTI, ELENA (2004):
**Did Crypteroniaceae really Disperse out of India? molecular Dating Evidence from rbcL, ndhF, and rpl16 Intron Sequences.**
In: International Journal of Plant Sciences 165 (4S), S69-S83.

### Cucurbitaceae

BOER, HUGO J.; CROSS, HUGH B.; WILDE, WILLEM J.J.O.; DUYFJES-DE WILDE, BRIGITTA E.E.; GRAVENDEEL, BARBARA (2015):
**Molecular phylogenetic Analyses of Cucurbitaceae Tribe Benincaseae urge for Merging of *Pilogyne* with *Zehneria*.**
In: Phytotaxa 236 (2), p. 173. DOI: 10.11646/phytotaxa.236.2.6.

BOER, HUGO J.; SCHAEFER, HANNO; THULIN, MATS; RENNER, SUSANNE P. (2012):
**Evolution and loss of long-fringed petals: a case Study using a dated Phylogeny of the snake gourds, *Trichosanthes* (Cucurbitaceae).**
In: BMC Evolutionary Biology 12, p. 108. DOI: 10.1186/1471-2148-12-108.

CHOMICKI, GUILLAUME; RENNER, SUSANNE P. (2015):
**Watermelon Origin solved with molecular Phylogenetics including Linnaean Material: another Example of Museomics.**
In: the new Phytologist 205 (2), p. 526–532. DOI: 10.1111/nph.13163.

DUCHEN, PABLO; RENNER, SUSANNE P. (2010):
**The Evolution of *Cayaponia* (Cucurbitaceae): Repeated shifts from Bat to Bee Pollination and long-distance Dispersal to Africa 2-5 million years ago.**
In: American Journal of Botany 97 (7), p. 1129–1141. DOI: 10.3732/ajb.0900385.

GHEBRETINSAE, AMANUEL G.; THULIN, MATS; BARBER, JANET C. (2007):
**Relationships of Cucumbers and Melons unraveled: molecular Phylogenetics of *Cucumis* and related Genera (Benincaseae, Cucurbitaceae).**

In: American Journal of Botany 94 (7), p. 1256–1266. DOI: 10.3732/ajb.94.7.1256.

**Holstein, Norbert; Renner, Susanne p. (2011):**
**A dated Phylogeny and collection records reveal repeated Biome Shifts in the African Genus *Coccinia* (Cucurbitaceae).**
In: BMC Evolutionary Biology 11, p. 28. DOI: 10.1186/1471-2148-11-28.

**Kates, Heather R.; Soltis, Pamela S.; Soltis, Douglas E. (2017):**
**Evolutionary and domestication History of *Cucurbita* (pumpkin and squash) Species inferred from 44 nuclear loci.**
In: Molecular Phylogenetics and Evolution 111, p. 98–109. DOI: 10.1016/j.ympev.2017.03.002.

**Kocyan, Alexander; Zhang, Li-Bing; Schaefer, Hanno; Renner, Susanne p. (2007):**
**A multi-locus Chloroplast Phylogeny for the Cucurbitaceae and its Implications for Character Evolution and Classification.**
In: Molecular Phylogenetics and Evolution 44 (2), p. 553–577. DOI: 10.1016/j.ympev.2006.12.022.

**Li, Hong-Tao; Li, De-Zhu (2008):**
**Systematic Position of *Gomphogyne* (Cucurbitaceae) inferred from ITS, rpl16 and trnS-trnR DNA Sequences.**
In: Journal of Systematics and Evolution 46 (4), p. 595–599.

**Li, Hong-Tao; Yang, Jun-Bo; Li, De-Zhu; Möller, Michael; Shah, Amin (2010):**
**A molecular phylogenetic Study of *Hemsleya* (Cucurbitaceae) based on ITS, rpl16, trnH-psbA, and trnL DNA Sequences.**
In: Plant Systematics and Evolution 285 (1-2), p. 23–32. DOI: 10.1007/s00606-009-0252-y.

**Lira, Rafael; Sosa, Victoria; Legaspi, Talitha; Dávila, Patricia (2015):**
**Phylogenetic Relationships of *Ibervillea* and *Tumamoca* (Coniandreae, Cucurbitaceae), two Genera of the Dry Lands of North America.**
In: Phytotaxa 201 (3), p. 197. DOI: 10.11646/phytotaxa.201.3.3.

**Nee, Michael; Schaefer, Hanno; Renner, Susanne p. (2009):**
**The Relationship between *Anisosperma* and *Fevillea* (Cucurbitaceae), and a new Species of *Fevillea* from Bolivia.**
In: Systematic Botany 34 (4), p. 704–708. DOI: 10.1600/036364409790139718.

**Renner, Susanne S.; Schaefer, Hanno; Kocyan, Alexander (2007):**
**Phylogenetics of *Cucumis* (Cucurbitaceae): *Cucumber* (*C. sativus*) belongs in an Asian/Australian Clade far from Melon (*C. melo*).**
In: BMC Evolutionary Biology 7, p. 58. DOI: 10.1186/1471-2148-7-58.

**Schaefer, Hanno; Kocyan, Alexander; Renner, Susanne p. (2008):**
***Linnaeosicyos* (Cucurbitaceae): a new Genus for *Trichosanthes amara*, the Caribbean Sister Species of all Sicyeae.**
In: Systematic Botany 33 (2), p. 349–355. DOI: 10.1600/036364408784571707.

**Schaefer, Hanno; Renner, Susanne p. (2010):**
**A three-Genome Phylogeny of *Momordica* (Cucurbitaceae) suggests seven returns from Dioecy to Monoecy and recent long-distance Dispersal to Asia.**
In: Molecular Phylogenetics and Evolution 54 (2), p. 553–560. DOI: 10.1016/j.ympev.2009.08.006.

**Singh, N. P.; Matta, N. K. (2008):**
**Variation studies on Seed storage proteins and Phylogenetics of the Genus *Cucumis*.**
In: Plant Systematics and Evolution 275 (3-4), p. 209–218. DOI: 10.1007/s00606-008-0063-6.

Steele, P. Roxanne; Friar, Lauren M.; Gilbert, Lawrence E.; Jansen, Robert K. (2010):
**Molecular Systematics of the Neotropical Genus *Psiguria* (Cucurbitaceae): Implications for Phylogeny and Species Identification.**
In: American Journal of Botany 97 (1), p. 156–173. DOI: 10.3732/ajb.0900192.

Telford, Ian R.H.; Sebastian, Patrizia; Lange, Peter J.; Bruhl, Jeremy J.; Renner, Susanne p. (2012):
**Morphological and molecular Data reveal three rather than one Species of *Sicyos* (Cucurbitaceae) in Australia, New Zealand and Islands of the South West Pacific.**
In: Australian Systematic Botany 25 (3), p. 188–201. DOI: 10.1071/SB11032.

Volz, Stefanie M.; Renner, Susanne p. (2008):
**Hybridization, polyploidy, and evolutionary transitions between Monoecy and Dioecy in *Bryonia* (Cucurbitaceae).**
In: American Journal of Botany 95 (10), p. 1297–1306. DOI: 10.3732/ajb.0800187.

Volz, Stefanie M.; Renner, Susanne p. (2009):
**Phylogeography of the ancient Eurasian medicinal plant Genus *Bryonia* (Cucurbitaceae) inferred from nuclear and Chloroplast Sequences.**
In: Taxon 58 (2), p. 550–560.

### Cucurbitales

Schaefer, Hanno; Renner, Susanne p. (2011):
**Phylogenetic Relationships in the Order Cucurbitales and a new Classification of the gourd Family (Cucurbitaceae).**
In: Taxon 60 (1), p. 122–138.

Zhang, Li-Bing; Simmons, Mark P.; Kocyan, Alexander; Renner, Susanne p. (2006):
**Phylogeny of the Cucurbitales based on DNA Sequences of nine Loci from three Genomes: Implications for morphological and sexual System Evolution.**
In: Molecular Phylogenetics and Evolution 39 (2), p. 305–322. DOI: 10.1016/j.ympev.2005.10.002.

### Cunoniaceae

Bradford, Jason C. (2002):
**Molecular Phylogenetics and morphological Evolution in Cunonieae (Cunoniaceae).**
In: Annals of the Missouri Botanical Garden 89 (4), p. 491–503.

Bradford, Jason C.; Barnes, Richard W. (2001):
**Phylogenetics and Classification of Cunoniaceae (Oxalidales) using Chloroplast DNA Sequences and Morphology.**
In: Systematic Botany 26 (2), p. 354–385.

Hopkins, Helen C.F.; Rozefelds, Andrew C.; Pillon, Yohan (2013):
***Karrabina* gen. nov. (Cunoniaceae), for the Australian Species previously placed in *Geissois*, and a Synopsis of Genera in the Tribe Geissoieae.**
In: Australian Systematic Botany 26 (3), p. 167–185. DOI: 10.1071/SB12037.

Pillon, Yohan; Hopkins, Helen C.F.; Munzinger, Jérôme; Chase, Mark W. (2009):
**A molecular and morphological Survey of Generic Limits of *Acsmithia* and *Spiraeanthemum* (Cunoniaceae).**
In: Systematic Botany 34 (1), p. 141–148. DOI: 10.1600/036364409787602410.

Rozefelds, Andrew C.; Barnes, Richard W. (2002):
**The Systematic and Biogeographical Relationships of *Ceratopetalum* (Cunoniaceae) in Australia and New Guinea.**

In: International Journal of Plant Sciences 163 (4), p. 651–673.

### Cupressaceae

**Adams, Robert P.; Schwarzbach, Andrea E. (2013):**
**Phylogeny of *Juniperus* using nrDNA and four cpDNA Regions.**
In: Phytologia 95 (2), p. 179–187.

**Gadek, Paul A.; Alpers, Deryn L.; Heslewood, Margaret M.; Quinn, Christopher J. (2000):**
**Relationships within Cupressaceae sensu lato: A combined morphological and molecular Approach.**
In: American Journal of Botany 87 (7), p. 1044–1057.

**Kusumi, Junko; Tsumura, Yoshihiko; Yoshimaru, Hiroshi; Tachida, Hidenori (2000):**
**Phylogenetic Relationships in Taxodiaceae and Cupressaceae based on matK Gene, Chll Gene, Trnl-trnf Igs Region, and Trnl Intron Sequences.**
In: American Journal of Botany 87 (10), p. 1480–1488.

**Little, Damon P. (2006):**
**Evolution and Circumscription of the True Cypresses (Cupressaceae: *Cupressus*).**
In: Systematic Botany 31 (3), p. 461–480.

**Little, Damon P.; Schwarzbach, Andrea E.; Adams, Robert P.; Hsieh, Chang-Fu (2004):**
**The Circumscription and phylogenetic Relationships of *Callitropsis* and the newly described Genus *Xanthocyparis* (Cupressaceae).**
In: American Journal of Botany 91 (11), p. 1872–1881.

**Mao, Kang-Shan; Hao, Gang; Liu, Jian-Quan; Adams, Robert P.; Milne, Richard Ian (2010):**
**Diversification and Biogeography of *Juniperus* (Cupressaceae): variable Diversification rates and multiple intercontinental Dispersals.**
In: the new Phytologist 188 (1), p. 254–272. DOI: 10.1111/j.1469-8137.2010.03351.x.

**Peng, Dan; Wang, Xiao-Quan (2008):**
**Reticulate Evolution in *Thuja* inferred from multiple Gene Sequences: Implications for the Study of biogeographical disjunction between eastern Asia and North America.**
In: Molecular Phylogenetics and Evolution 47 (3), p. 1190–1202. DOI: 10.1016/j.ympev.2008.02.001.

**Piggin, Josephine; Bruhl, Jeremy J. (2010):**
**Phylogeny Reconstruction of *Callitris* Vent. (Cupressaceae) and its Allies leads to Inclusion of *Actinostrobus* within *Callitris*.**
In: Australian Systematic Botany 23 (2), p. 69–93. DOI: 10.1071/SB09044.

**Pye, M. G.; Gadek, Paul A.; Edwards, K. J. (2003):**
**Divergence, Diversity and Species of the Australasian *Callitris* (Cupressaceae) and allied Genera: Evidence from ITS Sequence Data.**
In: Australian Systematic Botany 16 (4), p. 505–514. DOI: 10.1071/SB02019.

**Schulz, Christian; Stützel, Thomas (2007):**
**Evolution of taxodiaceous Cupressaceae (Coniferopsida).**
In: Organisms Diversity and Evolution 7 (2), p. 124–135. DOI: 10.1016/j.ode.2006.03.001.

**Terry, Randall G.; Bartel, Jim A.; Adams, Robert P. (2012):**
**Phylogenetic Relationships among the New World *Cypresses* (*Hesperocyparis*; Cupressaceae): Evidence from noncoding Chloroplast DNA Sequences.**
In: Plant Systematics and Evolution 298 (10), p. 1987–2000. DOI: 10.1007/s00606-012-0696-3.

**Terry, Randall G.; Pyne, Matthew I.; Bartel, Jim A.; Adams, Robert P. (2016):**
**A molecular Biogeography of the New World Cypresses (*Callitropsis, Hesperocyparis*; Cupressaceae).**
In: Plant Systematics and Evolution 302 (7), p. 921–942. DOI: 10.1007/s00606-016-1308-4.

**Xu, Tingting; Abbott, Richard J.; Milne, Richard Ian; Mao, Kang-Shan; Du, Fang K.; Wu, Guili et al. (2010):**
**Phylogeography and allopatric Divergence of Cypress Species (*Cupressus* L.) in the Qinghai-Tibetan Plateau and adjacent Regions.**
In: BMC Evolutionary Biology 10, p. 194. DOI: 10.1186/1471-2148-10-194.

**Yang, Zu-Yu; Ran, Jin-Hua; Wang, Xiao-Quan (2012):**
**Three Genome-based Phylogeny of Cupressaceae s.l.: further Evidence for the Evolution of Gymnosperms and Southern Hemisphere Biogeography.**
In: Molecular Phylogenetics and Evolution 64 (3), p. 452–470. DOI: 10.1016/j.ympev.2012.05.004.

### Cyatheaceae

**Bystriakova, Nadia; Fls, Harald Schneider; Coomes, David (2011):**
**Evolution of the climatic Niche in scaly Tree Ferns (Cyatheaceae, Polypodiopsida).**
In: Botanical Journal of the Linnean Society 165, p. 1–19.

**Korall, Petra; Conant, David S.; Metzgar, Jordan S.; Schneider, Harald; Pryer, Kathleen M. (2007):**
**A molecular Phylogeny of Scaly Tree Ferns (Cyatheaceae).**
In: American Journal of Botany 94 (5), p. 873–886.

**Korall, Petra; Pryer, Kathleen M.; Metzgar, Jordan S.; Schneider, Harald; Conant, David p. (2006):**
**Tree Ferns: monophyletic Groups and their Relationships as revealed by four Protein-coding Plastid Loci.**
In: Molecular Phylogenetics and Evolution 39 (3), p. 830–845. DOI: 10.1016/j.ympev.2006.01.001.

**Lehnert, Marcus (2016):**
**A Synopsis of the exindusiate Species of *Cyathea* (Cyatheaceae-Polypodiopsida) with bipinnate-pinnatifid or more complex Fronds, with a Revision of the *C. lasiosora*-complex.**
In: Phytotaxa 243 (1), p. 1. DOI: 10.11646/phytotaxa.243.1.1.

### Cycadaceae

**Brookes, D. R.; Hereward, J. P.; Terry, L. I.; Walter, G. H. (2015):**
**Evolutionary dynamics of a cycad obligate Pollination Mutualism - Pattern and Process in extant Macrozamia Cycads and their specialist Thrips Pollinators.**
In: Molecular Phylogenetics and Evolution 93, p. 83–93. DOI: 10.1016/j.ympev.2015.07.003.

**Chaw, Shu-Miaw; Walters, Terrence W.; Chang, Chien-Chang; Hu, Shu-Hsuan; Chen, Shin-Hsiao (2005):**
**A Phylogeny of Cycads (Cycadales) inferred from Chloroplast matK gene, trnK Intron, and nuclear rDNA ITS Region.**
In: Molecular Phylogenetics and Evolution 37 (1), p. 214–234. DOI: 10.1016/j.ympev.2005.01.006.

**Condamine, Fabien L.; Nagalingum, Nathalie S.; Marshall, Charles R.; Morlon, Hélène (2015):**
**Origin and Diversification of living Cycads: a cautionary Tale on the Impact of the branching Process prior in Bayesian molecular Dating.**
In: BMC Evolutionary Biology 15, p. 65. DOI: 10.1186/s12862-015-0347-8.

**Lindstrom, Anders J.; Hill, K. D. (2007):**
**The Genus *Cycas* (Cycadaceae) in India.**
In: Telopea 11 (4), p. 463–488.

**Liu, Jian; Zhang, Shou-Zhou; Nagalingum, Nathalie S.; Chiang, Yu-Chung; Lindstrom, Anders J.; Gong, Xun (2018):**
**Phylogeny of the Gymnosperm Genus *Cycas* L. (Cycadaceae) as inferred from Plastid and nuclear Loci based on a large-scale Sampling: evolutionary Relationships and taxonomical Implications.**
In: Molecular Phylogenetics and Evolution 127, p. 87–97. DOI: 10.1016/j.ympev.2018.05.019.

**Nicolalde-Morejón, Fernando; Vergara-Silva, Francisco; González-Astorga, Jorge; Stevenson, Dennis Wm.; Vovides, Andrew P.; Sosa, Victoria (2011):**
**A character-based approach in the Mexican cycads supports diverse Multigene Combinations for DNA barcoding.**
In: Cladistics 27 (2), p. 150–164. DOI: 10.1111/j.1096-0031.2010.00321.x.

**Rai, Hardeep S.; O'Brien, Heath E.; Reeves, Patrick A.; Olmstead, Richard G.; Graham, Sean W. (2003):**
**Inference of higher-order Relationships in the Cycads from a large Chloroplast Data Set.**
In: Molecular Phylogenetics and Evolution 29 (2), p. 350–359. DOI: 10.1016/S1055-7903(03)00131-3.

**Salas-Leiva, Dayana E.; Meerow, Alan W.; Calonje, Michael; Griffith, M. Patrick; Francisco-Ortega, Javier; Nakamura, Kyoko et al. (2013):**
**Phylogeny of the Cycads based on multiple single-copy nuclear Genes: Congruence of concatenated Parsimony, Likelihood and Species Tree Inference Methods.**
In: Annals of Botany 112 (7), p. 1263–1278. DOI: 10.1093/aob/mct192.

**Treutlein, Jens; Wink, Michael (2002):**
**Molecular Phylogeny of Cycads inferred from rbcL Sequences.**
In: Naturwissenschaften 89 (5), p. 221–225. DOI: 10.1007/s00114-002-0308-0.

**Wei, Zhou; Meng-Meng, Guan; Xun, Gong (2015):**
***Cycas chenii* (Cycadaceae), a new Species from China, and its phylogenetic Position.**
In: Journal of Systematics and Evolution 53 (6), p. 489–498. DOI: 10.1111/jse.12153.

**Xiao, Long-Qian; Möller, Michael (2015):**
**Nuclear ribosomal ITS functional Paralogs resolve the phylogenetic Relationships of a late-Miocene Radiation Cycad *Cycas* (Cycadaceae).**
In: Public Library of Science One 10 (1), e0117971. DOI: 10.1371/journal.pone.0117971.

**Xiao, Long-Qian; Möller, Michael; Zhu, Hua (2010):**
**High nrDNA ITS Polymorphism in the ancient extant Seed Plant *Cycas*: incomplete concerted Evolution and the Origin of Pseudogenes.**
In: Molecular Phylogenetics and Evolution 55 (1), p. 168–177. DOI: 10.1016/j.ympev.2009.11.020.

### Cymodoceaceae

**Petersen, Gitte; Seberg, Ole; Short, Frederick T.; Fortes, Miguel D. (2014):**
**Complete genomic congruence but Non-Monophyly of *Cymodocea* (Cymodoceaceae), a small Group of Seagrasses.**
In: Taxon 63 (1), p. 3–8. DOI: 10.12705/631.2.

### Cynomoriaceae

**Zhang, Zhi-Hong; Li, Chun-Qi; Li, Jian-Hua (2009):**
**Phylogenetic Placement of *Cynomorium* in Rosales inferred from Sequences of the inverted repeat Region of the Chloroplast Genome.**
In: Journal of Systematics and Evolution 47 (4), p. 297–304. DOI: 10.1111/j.1759-6831.2009.00035.x.

### Cyperaceae

**Barrett, Russell L.; Wilson, Karen L. (2012):**

**A Review of the Genus *Lepidosperma* Labill. (Cyperaceae: Schoeneae).**

In: Australian Systematic Botany 25 (4), p. 225–294. DOI: 10.1071/SB11037.

**Bauters, Kenneth; Asselman, Pieter; Simpson, David A.A.; Muasya, A. Muthama; Goetghebeur, Paul; Larridon, Isabel (2016):**

**Phylogenetics, ancestral state reconstruction, and a new infrageneric Classification of *Scleria* (Cyperaceae) based on three DNA Markers.**

In: Taxon 65 (3), p. 444–466. DOI: 10.12705/653.2.

**Bauters, Kenneth; Larridon, Isabel; Reynders, Marc; Asselman, Pieter; Vrijdaghs, Alexander; Muasya, A. Muthama et al. (2014):**

**A new Classification for *Lipocarpha* and *Volkiella* as infrageneric Taxa of *Cyperus* s.l. (Cypereae, Cyperoideae, Cyperaceae): Insights from Species Tree Reconstruction supplemented with morphological and Floral developmental Data.**

In: Phytotaxa 166 (1), p. 1. DOI: 10.11646/phytotaxa.166.1.1.

**Castro, Olga; Gargiulo, Roberta; Del Guacchio, Emanuele; Caputo, Paolo; Luca, Paolo (2015):**

**A molecular survey concerning the Origin of *Cyperus esculentus* (Cyperaceae, Poales): two Sides of the same Coin (Weed vs. Crop).**

In: Annals of Botany 115 (5), p. 733–745. DOI: 10.1093/aob/mcv001.

**Chacón, Juliana; Madriñán, Santiago; Chase, Mark W.; Bruhl, Jeremy J. (2006):**

**Molecular Phylogenetics of *Oreobolus* (Cyperaceae) and the Origin and Diversification of the American Species.**

In: Taxon 55 (2), p. 359–366. DOI: 10.2307/25065583.

**da Silva, Carlos Roberto Maximiano; da Silva, Leandro Bento; Andrade, Jesus; Tardelli, Célia Guadalupe; Trevisan, Rafael; González-Elizondo, María Socorro; Vanzela, André Luís Laforga (2012):**

**Ornamentation of Achene Silica Walls and its Contribution to the Systematics of *Eleocharis* (Cyperaceae).**

In: Plant Systematics and Evolution 298 (2), p. 391–398. DOI: 10.1007/s00606-011-0552-x.

**Dragon, Julie A.; Barrington, David p. (2009):**

**Systematics of the *Carex aquatilis* and *C. lenticularis* Lineages: Geographically and ecologically divergent sister Clades of *Carex* Section *Phacocystis* (Cyperaceae).**

In: American Journal of Botany 96 (10), p. 1896–1906. DOI: 10.3732/ajb.0800404.

**Escudero, Marcial; Eaton, Deren A. R.; Hahn, Marlene; Hipp, Andrew L. (2014):**

**Genotyping-by-Sequencing as a tool to infer Phylogeny and ancestral Hybridization: a case Study in *Carex* (Cyperaceae).**

In: Molecular Phylogenetics and Evolution 79, p. 359–367. DOI: 10.1016/j.ympev.2014.06.026.

**Escudero, Marcial; Hipp, Andrew L.; Luceño, Modesto (2010):**

**Karyotype stability and predictors of Chromosome Number Variation in Sedges: a Study in *Carex* Section *Spirostachyae* (Cyperaceae).**

In: Molecular Phylogenetics and Evolution 57 (1), p. 353–363. DOI: 10.1016/j.ympev.2010.07.009.

**Escudero, Marcial; Luceño, Modesto (2009):**

**Systematics and Evolution of *Carex* sects. *Spirostachyae* and *Elatae* (Cyperaceae).**

In: Plant Systematics and Evolution 279 (1-4), p. 163–189. DOI: 10.1007/s00606-009-0156-x.

**Escudero, Marcial; Valcárcel, Virginia; Vargas, Pablo; Luceño, Modesto (2008):**

**Evolution in *Carex* L. Sect. *Spirostachyae* (Cyperaceae): A molecular and cytogenetic Approach.**
In: Organisms Diversity and Evolution 7 (4), p. 271–291. DOI: 10.1016/j.ode.2006.08.006.

**Escudero, Marcial; Valcárcel, Virginia; Vargas, Pablo; Luceño, Modesto (2009):**
**Significance of ecological Vicariance and long-distance Dispersal in the Diversification of *Carex* Sect. *Spirostachyae* (Cyperaceae).**
In: American Journal of Botany 96 (11), p. 2100–2114. DOI: 10.3732/ajb.0900134.

**Ford, Bruce A.; Iranpour, Mahmood; Naczi, Robert F. C.; Starr, Julian R.; Jerome, Cheryl A. (2006):**
**Phylogeny of *Carex* Subg. *Vignea* (Cyperaceae) based on Non-coding nrDNA Sequence Data.**
In: Systematic Botany 31 (1), p. 70–82.

**García-Madrid, Ana S.; Muasya, A. Muthama; Álvarez, Inés; Cantó, Paloma; Molina, José Antonio (2015):**
**Towards resolving phylogenetic Relationships in the *Ficinia* Clade and Description of the new Genus *Afroscirpoides* (Cyperaceae: Cypereae).**
In: Taxon 64 (4), p. 688–702. DOI: 10.12705/644.2.

**Gebauer, Sebastian; Starr, Julian R.; Hoffmann, Matthias H. (2014):**
**Parallel and convergent Diversification in two Northern Hemispheric species-rich *Carex* Lineages (Cyperaceae).**
In: Organisms Diversity and Evolution 14 (3), p. 247–258. DOI: 10.1007/s13127-014-0171-9.

**Gehrke, Berit; Martín-Bravo, Santiago; Muasya, Muthama; Luceño, Modesto (2010):**
**Monophyly, phylogenetic Position and the role of Hybridization in *Schoenoxiphium* Nees (Cariceae, Cyperaceae).**
In: Molecular Phylogenetics and Evolution 56 (1), p. 380–392. DOI: 10.1016/j.ympev.2010.03.036.

**Ghamkhar, Kioumars; Marchant, Adam D.; Wilson, Karen L.; Bruhl, Jeremy J. (2007):**
**Phylogeny of Abildgaardieae (Cyperaceae) inferred from ITS and trnL–F Data.**
In: Aliso 23 (1), p. 149–164.

**Hendrichs, M.; Michalski, S.; Begerow, D.; Oberwinkler, F.; Hellwig, Frank H. (2004):**
**Phylogenetic Relationships in *Carex*, Subgenus *Vignea* (Cyperaceae), based on ITS Sequences.**
In: Plant Systematics and Evolution 246 (1-2), p. 109–125. DOI: 10.1007/s00606-004-0127-1.

**Hendrichs, M.; Oberwinkler, F.; Begerow, D.; Bauer, R. (2004):**
***Carex*, Subgenus *Carex* (Cyperaceae) - A phylogenetic approach using ITS Sequences.**
In: Plant Systematics and Evolution 246 (1-2), p. 89–107. DOI: 10.1007/s00606-004-0128-0.

**Hinchliff, Cody E.; Lliully, Ariel Ernesto Lliully A.; Carey, Timothy; Roalson, Eric H. (2010):**
**The Origins of *Eleocharis* (Cyperaceae) and the Status of *Websteria, Egleria,* and *Chillania*.**
In: Taxon 59 (3), p. 709–719.

**Hinchliff, Cody E.; Roalson, Eric H. (2013):**
**Using Supermatrices for phylogenetic inquiry: an Example using the Sedges.**
In: Systematic Biology 62 (2), p. 205–219. DOI: 10.1093/sysbio/sys088.

**Hipp, Andrew L.; Reznicek, Anton A.; Rothrock, Paul E.; Weber, Jaime A. (2006):**
**Phylogeny and Classification of *Carex* Section *Ovales* (Cyperaceae).**
In: International Journal of Plant Sciences 167 (5), p. 1029–1048.

**Hipp, Andrew L.; Rothrock, Paul E.; Reznicek, Anton A.; Berry, Paul E. (2007):**

**Chromosome Number Changes Associated with Speciation in Sedges: a phylogenetic Study in *Carex* Section *Ovales* (Cyperaceae) using AFLP Data.**
In: Aliso 23, p. 193–203.

ITO, YU; VILJOEN, JAN-ADRIAAN; TANAKA, NORIO; YANO, OKIHITO; MUASYA, A. MUTHAMA (2016):
**Phylogeny of *Isolepis* (Cyperaceae) revisited: non-monophyletic Nature of *I. fluitans* sensu lato and Resurrection of *I. lenticularis*.**
In: Plant Systematics and Evolution 302 (2), p. 231–238. DOI: 10.1007/s00606-015-1253-7.

JIMÉNEZ-MEJÍAS, PEDRO; HAHN, MARLENE; LUEDERS, KATE; STARR, JULIAN R.; BROWN, BETHANY H.; CHOUINARD, BRIANNA N. ET AL. (2016):
**Megaphylogenetic Specimen-level Approaches to the *Carex* (Cyperaceae) Phylogeny using ITS, ETS, and matK Sequences: Implications for Classification.**
In: Systematic Botany 41 (3), p. 500–518. DOI: 10.1600/036364416X692497.

JUNG, JONGDUK; CHOI, HONG-KEUN (2013):
**Recognition of two major Clades and early diverged Groups within the Subfamily Cyperoideae (Cyperaceae) including Korean Sedges.**
In: Journal of Plant Research 126 (3), p. 335–349. DOI: 10.1007/s10265-012-0534-2.

KIM, CHANGKYUN; JUNG, JONGDUK; CHOI, HONG-KEUN (2012):
**Molecular Identification of *Schoenoplectiella* Species (Cyperaceae) by Use of Microsatellite Markers.**
In: Plant Systematics and Evolution 298 (4), p. 811–817. DOI: 10.1007/s00606-012-0592-x.

KING, MATTHEW G.; ROALSON, ERIC H. (2008):
**Exploring evolutionary Dynamics of nrDNA in *Carex* Subgenus *Vignea* (Cyperaceae).**
In: Systematic Botany 33 (3), p. 514–524. DOI: 10.1600/036364408785679860.

KOSNAR, JAN; KOSNAR, JIRÍ; HERBSTOVÁ, MIROSLAVA; MACEK, PETR; REJMÁNKOVÁ, ELISKA; STECH, MILAN (2010):
**Natural Hybridization in tropical Spikerushes of *Eleocharis* Subgenus *Limnochloa* (Cyperaceae): Evidence from Morphology and DNA Markers.**
In: American Journal of Botany 97 (7), p. 1229–1240. DOI: 10.3732/ajb.1000029.

LARRIDON, ISABEL; BAUTERS, KENNETH; REYNDERS, MARC; HUYGH, WIM; GOETGHEBEUR, PAUL (2014):
**Taxonomic Changes in C4 *Cyperus* (Cypereae, Cyperoideae, Cyperaceae): combining the Sedge Genera *Ascolepis, Kyllinga* and *Pycreus* into *Cyperus* s.l.**
In: Phytotaxa 166 (1), p. 33. DOI: 10.11646/phytotaxa.166.1.2.

LARRIDON, ISABEL; BAUTERS, KENNETH; REYNDERS, MARC; HUYGH, WIM; MUASYA, A. MUTHAMA; SIMPSON, DAVID A.A.; GOETGHEBEUR, PAUL (2013):
**Towards a new Classification of the giant paraphyletic Genus *Cyperus* (Cyperaceae): phylogenetic Relationships and generic Delimitation in C4-*Cyperus*.**
In: Botanical Journal of the Linnean Society 172 (1), p. 106–126. DOI: 10.1111/boj.12020.

LARRIDON, ISABEL; GOVAERTS, RAFAEL; BAUTERS, KENNETH; GOETGHEBEUR, PAUL (2016):
***Cyperus albescens*, a new Combination in *Cyperus* (Cyperaceae) for the common (sub)tropical African and Asian Species *Lipocarpha chinensis*.**
In: Kew Bulletin 71 (2). DOI: 10.1007/S12225-016-9642-9.

LARRIDON, ISABEL; REYNDERS, MARC; HUYGH, WIM; BAUTERS, KENNETH; VAN DE PUTTE, KOBEKE; MUASYA, A. MUTHAMA ET AL. (2011):
**Affinities in C3-*Cyperus* Lineages (Cyperaceae) revealed using molecular phylogenetic Data and Carbon Isotope Analysis.**
In: Botanical Journal of the Linnean Society 167 (1), p. 19–46. DOI: 10.1111/j.1095-8339.2011.01160.x.

LEHNEBACH, CARLOS A. (2011):
**Re-evaluating Species limits in *Uncinia angustifolia*, *U. caespitosa* s.str., *U. rupestris*, *U. viridis* and *U. zotovii* (Cyperaceae).**
In: Australian Systematic Botany 24 (6), p. 405–420. DOI: 10.1071/SB11014.

LÉVEILLÉ-BOURRET, ÉTIENNE; DONADÍO, SABINA; GILMOUR, CLAIRE N.; STARR, JULIAN R. (2015):
***Rhodoscirpus* (Cyperaceae: Scirpeae), a new South American Sedge Genus supported by molecular, morphological, anatomical and embryological Data.**
In: Taxon 64 (5), p. 931–944. DOI: 10.12705/645.4.

LÉVEILLÉ-BOURRET, ÉTIENNE; GILMOUR, CLAIRE N.; STARR, JULIAN R.; NACZI, ROBERT F. C.; SPALINK, DANIEL; SYTSMA, KENNETH J. (2014):
**Searching for the sister to Sedges (*Carex*): resolving Relationships in the Cariceae-Dulichieae-Scirpeae Clade (Cyperaceae).**
In: Botanical Journal of the Linnean Society 176 (1), p. 1–21. DOI: 10.1111/boj.12193.

LÉVEILLÉ-BOURRET, ÉTIENNE; STARR, JULIAN R.; FORD, BRUCE A. (2018):
**Why are there so many Sedges? Sumatroscirpeae, a missing piece in the evolutionary Puzzle of the giant Genus *Carex* (Cyperaceae).**
In: Molecular Phylogenetics and Evolution 119, p. 93–104. DOI: 10.1016/j.ympev.2017.10.025.

MAGUILLA, ENRIQUE; ESCUDERO, MARCIAL; WATERWAY, MARCIA J.; HIPP, ANDREW L.; LUCEÑO, MODESTO (2015):
**Phylogeny, Systematics, and trait Evolution of *Carex* Section *Glareosae*.**
In: American Journal of Botany 102 (7), p. 1128–1144. DOI: 10.3732/ajb.1500169.

MASSATTI, ROB; REZNICEK, ANTON A.; KNOWLES, L. LACEY (2016):
**Utilizing RADseq Data for phylogenetic Analysis of challenging taxonomic Groups: A case Study in *Carex* Sect. *Racemosae*.**
In: American Journal of Botany 103 (2), p. 337–347. DOI: 10.3732/ajb.1500315.

MUASYA, A. MUTHAMA; SIMPSON, DAVID A.A.; CHASE, MARK W. (2002):
**Phylogenetic Relationships in *Cyperus* L. s.l. (Cyperaceae) inferred from Plastid DNA Sequence Data.**
In: Botanical Journal of the Linnean Society 138 (2), p. 145–153. DOI: 10.1046/j.1095-8339.2002.138002145.x.

MUASYA, A. MUTHAMA; SIMPSON, DAVID A.A.; VERBOOM, GEORGE ANTHONY; GOETGHEBEUR, PAUL; NACZI, ROBERT F. C.; CHASE, MARK W.; SMETS, ERIK (2009):
**Phylogeny of Cyperaceae based on DNA Sequence Data: Current Progress and Future Prospects.**
In: the Botanical Review 75 (1), p. 2–21. DOI: 10.1007/s12229-008-9019-3.

MUASYA, A. MUTHAMA; VILJOEN, JAN-ADRIAAN; DLUDLU, MESHACK N.; DEMISSEW, SEBSEBE (2014):
**Phylogenetic Position of *Cyperus clandestinus* (Cypereae, Cyperaceae) clarified by morphological and molecular Evidence.**
In: Nordic Journal of Botany 32 (1), p. 106–114. DOI: 10.1111/j.1756-1051.2012.01700.x.

MUSILI, PAUL M.; GIBBS, ADELE K.; WILSON, KAREN L.; BRUHL, JEREMY J. (2016):
***Schoenus* (Cyperaceae) is not monophyletic based on ITS nrDNA Sequence Data.**
In: Australian Systematic Botany 29 (5), p. 265. DOI: 10.1071/SB15046.

MUTHAMA MUASYA, A.; SIMPSON, DAVID A.A.; CHASE, MARK W.; CULHAM, ALASTAIR (2001):
**A Phylogeny of *Isolepis* (Cyperaceae) inferred using Plastid rbcL and trnL-F Sequence Data.**
In: Systematic Botany 26 (2), p. 342–353.

**Poindexter, Derick B.; Naczi, Robert F. C. (2014):**
**Taxonomy and geographic Distribution of *Carex lucorum* var. *austrolucorum* (section *Acrocystis*, Cyperaceae).**
In: Brittonia 66 (4), p. 358–370. DOI: 10.1007/s12228-014-9348-y.

**Reid, Christopher S.; Carter, Richard J.; Urbatsch, Lowell E. (2014):**
**Phylogenetic Insights into New World *Cyperus* (Cyperaceae) using nuclear ITS Sequences.**
In: Brittonia 66 (3), p. 292–305. DOI: 10.1007/s12228-014-9324-6.

**Reynders, Marc; Vrijdaghs, Alexander; Larridon, Isabel; Huygh, Wim; Leroux, Olivier; Muasya, A. Muthama; Goetghebeur, Paul (2012):**
**Gynoecial Anatomy and development in Cyperoideae (Cyperaceae, Poales): congenital fusion of Carpels facilitates evolutionary Modifications in Pistil Structure.**
In: Plant Ecology and Evolution 145 (1), p. 96–125. DOI: 10.5091/plecevo.2012.675.

**Reznicek, Anton A.; Elizondo, M. Socorro González (2008):**
***Cypringlea* (Cyperaceae) revisited, a new Combination and Status.**
In: Acta Botanica Mexicana 83, p. 13–23.

**Roalson, Eric H.; Friar, Elizabeth A. (2000):**
**Infrageneric Classification of *Eleocharis* (Cyperaceae) Revisited: Evidence from the Internal Transcribed Spacer (ITS) Region of Nuclear Ribosomal DNA.**
In: Systematic Botany 25 (2), p. 323–336. DOI: 10.2307/2666645.

**Roalson, Eric H.; Friar, Elizabeth A. (2004):**
**Phylogenetic Analysis of the nuclear Alcohol Dehydrogenase (Adh) Gene Family in *Carex* Section *Acrocystis* (Cyperaceae) and combined Analyses of Adh and nuclear ribosomal ITS and ETS Sequences for inferring Species Relationships.**
In: Molecular Phylogenetics and Evolution 33 (3), p. 671–686. DOI: 10.1016/j.ympev.2004.08.005.

**Roalson, Eric H.; Friar, Elizabeth A. (2004):**
**Phylogenetic Relationships and biogeographic Patterns in North American members of *Carex* Section *Acrocystis* (Cyperaceae) using nrDNA ITS and ETS Sequence Data.**
In: Plant Systematics and Evolution 243 (3-4), p. 175–187. DOI: 10.1007/s00606-003-0089-8.

**Roalson, Eric H.; Hinchliff, Cody E.; Trevisan, Rafael; da Silva, Carlos Roberto Maximiano (2010):**
**Phylogenetic Relationships in *Eleocharis* (Cyperaceae): C4 Photosynthesis Origins and Patterns of Diversification in the Spikerushes.**
In: Systematic Botany 35 (2), p. 257–271. DOI: 10.1600/036364410791638270.

**Rye, Barbara L.; Barrett, Russell L.; Barrett, Matthew D.; Bruhl, Jeremy J.; Clarke, Kerri L.; Wilson, Karen L. (2015):**
**Five new Species and a new Combination in Cyperaceae from the Kimberley Region of Western Australia.**
In: Nuytsia 26, p. 167–184.

**Shekhovtsov, Sergei V.; Shekhovtsova, Irina N.; Peltek, Sergei E. (2012):**
**Phylogeny of Siberian Species of *Carex* Sect. *Vesicariae* based on nuclear and Plastid Markers.**
In: Nordic Journal of Botany 30 (3), p. 343–351. DOI: 10.1111/j.1756-1051.2011.01405.x.

**Simpson, David A.A.; Furness, Carol A.; Hodkinson, Trevor R.; Muasya, A. Muthama; Chase, Mark W. (2003):**

**Phylogenetic Relationships in Cyperaceae Subfamily Mapanioideae inferred from Pollen and Plastid DNA Sequence Data.**
In: American Journal of Botany 90 (7), p. 1071–1086. DOI: 10.3732/ajb.90.7.1071.

Simpson, David A.A.; Muasya, A. Muthama; Chayamarit, Kongkanda; Parnell, John A.N.; Suddee, Somran; Wilde, Bart D. E. et al. (2005):
***Khaosokia caricoides*, a new Genus and Species of Cyperaceae from Thailand.**
In: Botanical Journal of the Linnean Society 149 (3), p. 357–364. DOI: 10.1111/j.1095-8339.2005.00446.x.

Simpson, David A.A.; Muthama Muasya, A.; Alves, Marccus V.; Bruhl, Jeremy J.; Dhooge, Sandra; Chase, Mark W. et al. (2007):
**Phylogeny of Cyperaceae based on DNA Sequence Data–a new rbcL Analysis.**
In: Aliso 23 (1), p. 72–83.

Slingsby, Jasper A.; Britton, Matthew N.; Verboom, George Anthony (2014):
**Ecology limits the Diversity of the Cape Flora: Phylogenetics and Diversification of the Genus *Tetraria*.**
In: Molecular Phylogenetics and Evolution 72, p. 61–70. DOI: 10.1016/j.ympev.2013.11.017.

Starr, Julian R.; Ford, Bruce A. (2009):
**Phylogeny and Evolution in Cariceae (Cyperaceae): Current Knowledge and Future Directions.**
In: the Botanical Review 75 (1), p. 110–137. DOI: 10.1007/s12229-008-9020-x.

Starr, Julian R.; Harris, Stephen A.; Simpson, David A.A. (2003):
**Potential of the 5' and 3' Ends of the Intergenic Spacer (IGS) of rDNA in the Cyperaceae: new Sequences for Lower-level Phylogenies in Sedges with an Example from *Uncinia* Pers.**
In: International Journal of Plant Sciences 164 (2), p. 213–227.

Starr, Julian R.; Harris, Stephen A.; Simpson, David A.A. (2004):
**Phylogeny of the Unispicate Taxa in Cyperaceae Tribe Cariceae I: Generic Relationships and evolutionary Scenarios.**
In: Systematic Botany 29 (3), p. 528–544. DOI: 10.1600/0363644041744455.

Starr, Julian R.; Janzen, Francesco H.; Ford, Bruce A. (2015):
**Three new, early diverging *Carex* (Cariceae, Cyperaceae) Lineages from East and Southeast Asia with important evolutionary and biogeographic Implications.**
In: Molecular Phylogenetics and Evolution 88, p. 105–120. DOI: 10.1016/j.ympev.2015.04.001.

Thomas, William Wayt; Araújo, Ana Claudia; Alves, Marccus Vinícius (2009):
**A Preliminary molecular Phylogeny of the Rhynchosporeae (Cyperaceae).**
In: the Botanical Review 75 (1), p. 22–29. DOI: 10.1007/s12229-008-9023-7.

Verboom, George Anthony (2006):
**A Phylogeny of the Schoenoid Sedges (Cyperaceae: Schoeneae) based on Plastid DNA Sequences, with special Reference to the Genera found in Africa.**
In: Molecular Phylogenetics and Evolution 38 (1), p. 79–89. DOI: 10.1016/j.ympev.2005.05.012.

Viljoen, Jan-Adriaan; Muasya, A. Muthama; Barrett, Russell L.; Bruhl, Jeremy J.; Gibbs, Adele K.; Slingsby, Jasper A. et al. (2013):
**Radiation and repeated transoceanic Dispersal of Schoeneae (Cyperaceae) through the Southern Hemisphere.**
In: American Journal of Botany 100 (12), p. 2494–2508. DOI: 10.3732/ajb.1300105.

**Waterway, Marcia J.; Hoshino, Takuji; Masaki, Tomomi (2009):**
**Phylogeny, Species Richness, and Ecological Specialization in Cyperaceae Tribe Cariceae.**
In: the Botanical Review 75 (1), p. 138–159. DOI: 10.1007/s12229-008-9024-6.

**Waterway, Marcia J.; Martins, Kyle T.; Dabros, Anna; Prado, Alberto; Lechowicz, Martin J. (2016):**
**Ecological and evolutionary Diversification Within the Genus *Carex* (Cyperaceae): Consequences for Community Assembly in subarctic Fens.**
In: Systematic Botany 41 (3), p. 558–579. DOI: 10.1600/036364416X692514.

**Waterway, Marcia J.; Starr, Julian R. (2007):**
**Phylogenetic Relationships in Tribe Cariceae (Cyperaceae) based on nested Analyses of four molecular Data Sets.**
In: Aliso 23 (1), p. 165–192.

**Yang, Peizhi; Zheng, Hongmei; Larson, Steven; Miao, Yanjun; Hu, Tianming (2010):**
**Phylogenetic Relationships of eleven *Kobresia* accessions from the Tibetan Plateau.**
In: African Journal of Biotechnology 9 (23), p. 3359–3367.

**Yano, Okihito; Hoshino, Takuji (2005):**
**Molecular Phylogeny and Chromosomal Evolution of Japanese *Schoenoplectus* (Cyperaceae), based on ITS and ETS1f Sequences.**
In: Acta Phytotaxonomica Geobotanica 56 (2), p. 183–195.

**Yano, Okihito; Hoshino, Takuji (2006):**
**Phylogenetic Relationships *Fimbristylis* (Cyperaceae) and Chromosomal Evolution of Japanese using nrDNA ITS and ETS lf Sequence Data.**
In: Acta Phytotaxonomica Geobotanica 57 (3), p. 205–217.

**Yano, Okihito; Ikeda, Hiroshi; Jin, Xiao-Feng; Hoshino, Takuji (2014):**
**Phylogeny and chromosomal Variations in East Asian *Carex*, Siderostictae Group (Cyperaceae), based on DNA Sequences and cytological Data.**
In: Journal of Plant Research 127 (1), p. 99–107. DOI: 10.1007/s10265-013-0578-y.

**Yano, Okihito; Ikeda, Hiroshi; Watson, Mark F.; Rajbhandari, Keshab R.; Jin, Xiao-Feng; Hoshino, Takuji et al. (2012):**
**Phylogenetic Position of the Himalayan Genus *Erioscirpus* (Cyperaceae) inferred from DNA Sequence Data.**
In: Botanical Journal of the Linnean Society 170 (1), p. 1–11. DOI: 10.1111/j.1095-8339.2012.01255.x.

**Yano, Okihito; Katsuyama, Teruo; Tsubota, Hiromi; Hoshino, Takuji (2004):**
**Molecular Phylogeny of Japanese *Eleocharis* (Cyperaceae) based on ITS Sequence Data, and chromosomal Evolution.**
In: Journal of Plant Research 117 (5), p. 409–419. DOI: 10.1007/s10265-004-0173-3.

**Yen, Alan C.; Olmstead, Richard G. (2000):**
**Molecular Systematics of Cyperaceae Tribe Cariceae based on two Chloroplast DNA Regions: ndhF and trnL Intron-Intergenic Spacer.**
In: Systematic Botany 25 (3), p. 479–494. DOI: 10.2307/2666691.

**Zhang, Xiufu; Bruhl, Jeremy J.; Wilson, Karen L.; Marchant, Adam (2007):**
**Phylogeny of *Carpha* and related Genera (Schoeneae, Cyperaceae) inferred from morphological and molecular Data.**
In: Australian Systematic Botany 20 (2), p. 93–106. DOI: 10.1071/SB06023.

**Zhang, Xiufu; Marchant, Adam; Wilson, Karen L.; Bruhl, Jeremy J. (2004):**
**Phylogenetic Relationships of *Carpha* and its relatives (Schoeneae, Cyperaceae) inferred from Chloroplast trnL Intron and trnL-trnF intergenic Spacer Sequences.**
In: Molecular Phylogenetics and Evolution 31 (2), p. 647–657. DOI: 10.1016/j.ympev.2003.09.004.

**Zhang, Xiufu; Wilson, Karen L.; Bruhl, Jeremy J. (2006):**
**Species limits in *Carpha* (Schoeneae, Cyperaceae) based on phenetic Analyses.**
In: Australian Systematic Botany 19 (5), p. 437–465. DOI: 10.1071/SB06003.

### Cytinaceae

**Nickrent, Daniel Lee (2007):**
**Cytinaceae are sister to Muntingiaceae (Malvales).**
In: Taxon 56 (4), p. 1129–1135. DOI: 10.2307/25065907.

### Davalliaceae

**Tsutsumi, Chie; Chen, Cheng-Wei; Larsson, Anders; Hirayama, Yumiko; Kato, Masahiro (2016):**
**Phylogeny and Classification of Davalliaceae on the basis of Chloroplast and nuclear Markers.**
In: Taxon 65 (6), p. 1236–1248. DOI: 10.12705/656.2.

**Tsutsumi, Chie; Zhang, Xian-Chun; Kato, Masahiro (2008):**
**Molecular Phylogeny of Davalliaceae and Implications for Generic Classification.**
In: Systematic Botany 33 (1), p. 44–48.

### Dennstaedtiaceae

**Der, Joshua P.; Thomson, John A.; Stratford, Jeran K.; Wolf, Paul G. (2009):**
**Global Chloroplast Phylogeny and Biogeography of Bracken (*Pteridium*; Dennstaedtiaceae).**
In: American Journal of Botany 96 (5), p. 1041–1049.

**Luong, Thien Tam; Hovenkamp, Peter H.; Sosef, Marc p. M. (2015):**
**Revision of the Fern Genus *Orthiopteris* (Saccolomataceae) in Malesia and adjacent Regions.**
In: PhytoKeys (53), p. 39–71. DOI: 10.3897/phytokeys.53.4955.

**Perrie, Leon R.; Shepherd, Lara D.; Brownsey, Patrick J. (2015):**
**An expanded Phylogeny of the Dennstaedtiaceae Ferns: *Oenotrichia* falls within a non-monophyletic *Dennstaedtia*, and *Saccoloma* is polyphyletic.**
In: Australian Systematic Botany 28 (4), p. 256–264. DOI: 10.1071/SB15035.

**Schwartsburd, Pedro Bond; Prado, Jefferson (2016):**
**A taxonomic Revision of the South American Species of *Hypolepis* (Dennstaedtiaceae), Part II.**
In: American Fern Journal 106 (1), p. 1–53. DOI: 10.1640/0002-8444-106.1.1.

**Wolf, Paul G.; Soltis, Pamela S.; Soltis, Douglas E. (1994):**
**Phylogenetic Relationships of Dennstaedtioid Ferns: Evidence from rbcl Sequences.**
In: Molecular Phylogenetics and Evolution 3 (4), p. 383–392.

**Zhou, Shiliang; Dong, Wen-Pan; Chen, Xiaoqing; Zhang, Xian-Chun; Wen, Jun; Schneider, Harald (2014):**
**How many Species of Bracken (*Pteridium*) are there? Assessing the Chinese Brackens using molecular Evidence.**
In: Taxon 63 (3), p. 509–521. DOI: 10.12705/633.9.

### Diapensiaceae

**Higashi, Hiroyuki; Ikeda, Hajime; Setoguchi, Hiroaki (2015):**

**Molecular Phylogeny of *Shortia* sensu lato (Diapensiaceae) based on multiple nuclear Sequences.**

In: Plant Systematics and Evolution 301 (2), p. 523–529. DOI: 10.1007/s00606-014-1088-7.

### Dichapetalaceae

**Amorim, André M.; Lisboa, Décio S.; Marinho, Lucas C.; Fiaschi, Pedro (2016):**

**Novelties in *Tapura* (Dichapetalaceae) from the Brazilian Atlantic Forest.**

In: Systematic Botany 41 (3), p. 747–757. DOI: 10.1600/036364416X692523.

### Didiereaceae

**Applequist, Wendy L.; Wallace, Robert p. (2000):**

**Phylogeny of the Madagascan endemic Family Didiereaceae.**

In: Plant Systematics and Evolution 221, p. 157–166.

### Didymochlaenaceae

**Zhang, Li-Bing; Zhang, Liang (2015):**

**Didymochlaenaceae: A new Fern Family of Eupolypods I (Polypodiales).**

In: Taxon 64 (1), p. 27–38. DOI: 10.12705/641.4.

### Dilleniaceae

**Aymard, Gerardo A.C. (2017):**

**Novelties in Neotropical Dilleniaceae.**

In: Journal of the Botanical Research Institute of Texas 11 (1), p. 45–51.

**Horn, James W. (2009):**

**Phylogenetics of Dilleniaceae using Sequence Data from four Plastid Loci (RBCL, infA, rps4, rpl16 Intron).**

In: International Journal of Plant Sciences 170 (6), p. 794–813. DOI: 10.1086/599239.

### Dioncophyllaceae

**Heubl, Günther; Bringmann, Gerhard; Meimberg, Harald (2006):**

**Molecular Phylogeny and Character Evolution of carnivorous Plant Families in Caryophyllales--revisited.**

In: Plant Biology 8 (6), p. 821–830. DOI: 10.1055/s-2006-924460.

### Dioscoreaceae

**Gao, Xing; Zhu, Yu-Ping; Wu, Bao-Cheng; Zhao, Ya-Mei; Jian-Qun; Hang, Chen Yue-Yu (2008):**

**Phylogeny of *Dioscorea* Sect. *Stenophora* based on Chloroplast matK, rbcL and trnL-F Sequences.**

In: Journal of Systematics and Evolution 46 (3), p. 315–321.

**Hsu, K.-M.; Tsai, J.-L.; Chen, M.-Y.; Ku, H.-M.; Liu, S.-C. (2013):**

**Molecular Phylogeny of *Dioscorea* (Dioscoreaceae) in East and Southeast Asia.**

In: Blumea 58 (1), p. 21–27. DOI: 10.3767/000651913X669022.

**Mukherjee, Papiya; Bhat, K. V. (2013):**

**Phylogenetic Relationship of wild and cultivated yam Species (*Dioscorea* spp.) of India inferred from Pcr–rflp Analysis of two cpDNA loci.**

In: Plant Systematics and Evolution 299 (8), p. 1587–1597. DOI: 10.1007/s00606-013-0847-1.

**Wilkin, Paul; Schols, Peter; Chase, Mark W.; Chayamarit, Kongkanda; Furness, Carol A.; Huysmans, Suzy et al. (2005):**
**A Plastid Gene Phylogeny of the Yam Genus, *Dioscorea*: Roots, Fruits and Madagascar.**
In: Systematic Botany 30 (4), p. 736–749. DOI: 10.1600/036364405775097879.

### Dipsacaceae

**Avino, M.; Tortoriello, G.; Caputo, P. (2009):**
**A phylogenetic Analysis of Dipsacaceae based on four DNA Regions.**
In: Plant Systematics and Evolution 279 (1-4), p. 69–86. DOI: 10.1007/s00606-009-0147-y.

**Caputo, P.; Cozzolino, Salvatore; Moretti, A. (2004):**
**Molecular Phylogenetics of Dipsacaceae reveals parallel Trends in Seed Dispersal Syndromes.**
In: Plant Systematics and Evolution 246 (3-4). DOI: 10.1007/s00606-004-0154-y.

**Carlson, Sara E.; Howarth, Dianella G.; Donoghue, Michael J. (2011):**
**Diversification of Cycloidea-like Genes in Dipsacaceae (Dipsacales): Implications for the Evolution of Capitulum Inflorescences.**
In: BMC Evolutionary Biology 11, p. 325. DOI: 10.1186/1471-2148-11-325.

**Carlson, Sara E.; Mayer, Veronika; Donoghue, Michael J. (2009):**
**Phylogenetic Relationships, Taxonomy, and morphological Evolution in Dipsacaceae (Dipsacales) inferred by DNA Sequence Data.**
In: Taxon 58 (4), p. 1075–1091.

**Frajman, Božo; Rešetnik, Ivana; Weiss-Schneeweiss, Hanna; Ehrendorfer, Friedrich; Schönswetter, Peter (2015):**
**Cytotype Diversity and Genome Size Variation in *Knautia* (Caprifoliaceae, Dipsacoideae).**
In: BMC Evolutionary Biology 15, p. 140. DOI: 10.1186/s12862-015-0425-y.

**Mayer, Veronika; Ehrendorfer, Friedrich (2013):**
**The phylogenetic Position of *Pterocephalidium* and the new African Genus *Pterothamnus* within an improved Classification of Dipsacaceae.**
In: Taxon 62 (1), p. 112–126.

**Rešetnik, Ivana; Frajman, Božo; Bogdanović, Sandro; Ehrendorfer, Friedrich; Schönswetter, Peter (2014):**
**Disentangling Relationships among the diploid Members of the intricate Genus *Knautia* (Caprifoliaceae, Dipsacoideae).**
In: Molecular Phylogenetics and Evolution 74, p. 97–110. DOI: 10.1016/j.ympev.2014.01.028.

### Dipterocarpaceae

**Cao, Cui-Ping; Gailing, Oliver; Siregar, Iskandar; Indrioko, Sapto; Finkeldey, Reiner (2006):**
**Genetic Variation at AFLPs for the Dipterocarpaceae and its relation to molecular Phylogenies and taxonomic Subdivisions.**
In: Journal of Plant Research 119 (5), p. 553–558. DOI: 10.1007/s10265-006-0005-8.

**Choong, Chee Yen; Wickneswari, R.; Norwati, M.; Abbott, Richard J. (2008):**
**Phylogeny of *Hopea* (Dipterocarpaceae) inferred from Chloroplast DNA and nuclear PgiC Sequences.**
In: Molecular Phylogenetics and Evolution 48 (3), p. 1238–1243. DOI: 10.1016/j.ympev.2008.01.004.

**Dayanandan, Selvadurai; Ashton, Peter Shaw; Williams, Scott M.; Primack, Richard B. (1999):**

**Phylogeny of the tropical Tree Family Dipterocarpaceae based on nucleotide Sequences of the Chloroplast Rbcl gene.**
In: American Journal of Botany 86 (8), p. 1182–1190. DOI: 10.2307/2656982.

INDRIOKO, SAPTO; GAILING, OLIVER; FINKELDEY, REINER (2006):
**Molecular Phylogeny of Dipterocarpaceae in Indonesia based on Chloroplast DNA.**
In: Plant Systematics and Evolution 261 (1-4), p. 99–115. DOI: 10.1007/s00606-006-0435-8.

KAJITA, TADASHI; KAMIYA, KOICHI; NAKAMURA, K.; TACHIDA, HIDENORI; WICKNESWARI, R.; TSUMURA, YOSHIHIKO ET AL. (1998):
**Molecular Phylogeny of Dipetrocarpaceae in Southeast Asia based on Nucleotide Sequences of matK, trnL Intron, and trnL-trnF Intergenic Spacer Region in Chloroplast DNA.**
In: Molecular Phylogenetics and Evolution 10 (2), p. 202–209. DOI: 10.1006/mpev.1998.0516.

KAMIYA, KOICHI; HARADA, KO; TACHIDA, HIDENORI; ASHTON, PETER SHAW (2005):
**Phylogeny of PgiC Gene in *Shorea* and its closely related Genera (Dipterocarpaceae), the dominant Trees in Southeast Asian tropical Rain Forests.**
In: American Journal of Botany 92 (5), p. 775–788. DOI: 10.3732/ajb.92.5.775.

LI, QIAO-MING; HE, TIAN-HUA; XU, ZAI-FU (2004):
**Generic Relationships of Parash*orea chinensis* WangHsie (Dipterocarpaceae) based on cpDNA Sequences.**
In: Taxon 53 (2), p. 461–466. DOI: 10.2307/4135622.

MORTON, CYNTHIA M.; DAYANANDAN, SELVADURAI; DISSANAYAKE, D. (1999):
**Phylogeny and Biosystematics of *Pseudomonotes* (Dipterocarpaceae) based on molecular and morphological Data.**
In: Plant Systematics and Evolution 216 (3-4), p. 197–205. DOI: 10.1007/BF01084398.

TSUMURA, YOSHIHIKO; KADO, TOMOYUKI; YOSHIDA, KAZUMASA; ABE, HISASHI; OHTANI, MASATO; TAGUCHI, YURIKO ET AL. (2011):
**Molecular Database for classifying *Shorea* Species (Dipterocarpaceae) and Techniques for checking the Legitimacy of Timber and Wood Products.**
In: Journal of Plant Research 124 (1), p. 35–48. DOI: 10.1007/s10265-010-0348-z.

YULITA, KUSUMADEWI S.; BAYER, RANDALL J.; WEST, JUDITH G. (2005):
**Molecular phylogenetic Study of *Hopea* and *Shorea* (Dipterocarpaceae): Evidence from the trnL-trnF and internal transcribed Spacer Regions.**
In: Plant Species Biology 20 (3), p. 167–182. DOI: 10.1111/j.1442-1984.2005.00136.x.

### Dombeyaceae

APPLEQUIST, WENDY L. (2009):
**A Revision of the Malagasy Endemic *Helmiopsis* (Malvaceae s.l.).**
In: Annals of the Missouri Botanical Garden 96 (4), p. 521–540. DOI: 10.3417/2007184.

LE PÉCHON, TIMOTHÉE; CAO, NATHANAËL; DUBUISSON, JEAN-YVES; GIGORD, LUC D.B. (2009):
**Systematics of Dombeyoideae (Malvaceae) in the Mascarene Archipelago (Indian Ocean) inferred from Morphology.**
In: Taxon 58 (2), p. 519–531. DOI: 10.1002/tax.582016.

LE PÉCHON, TIMOTHÉE; DAI, QIANG; ZHANG, LI-BING; GAO, XIN-FEN; SAUQUET, HERVÉ (2015):
**Diversification of Dombeyoideae (Malvaceae) in the Mascarenes: Old Taxa on Young Islands?**

In: International Journal of Plant Sciences 176 (3), p. 211–221. DOI: 10.1086/679350.

**Le Péchon, Timothée; Dubuisson, Jean-Yves; Haevermans, Thomas; Cruaud, Corinne; Couloux, Arnaud; Gigord, Luc D.B. (2010):**

**Multiple colonizations from Madagascar and converged Acquisition of Dioecy in the Mascarene Dombeyoideae (Malvaceae) as inferred from Chloroplast and nuclear DNA Sequence Analyses.**

In: Annals of Botany 106 (2), p. 343–357. DOI: 10.1093/aob/mcq116.

**Skema, Cynthia (2012):**

**Toward a new Circumscription of *Dombeya* (Malvales: Dombeyaceae): A molecular phylogenetic and morphological Study of *Dombeya* of Madagascar and a new segregate Genus, *Andringitra*.**

In: Taxon 61 (3), p. 612–628. DOI: 10.1002/tax.613010.

**Wilkie, Peter (2007):**

**A new Species of *Pterospermum* (Dombeyoideae, Malvaceae/sterculiaceae) from Cambodia and Vietnam.**

In: Edinburgh Journal of Botany 64 (2), p. 179–183. DOI: 10.1017/S0960428607000881.

**Won, Hyosig (2009):**

**Phylogenetic Position of *Corchoropsis* Siebold & Zucc. (Malvaceae s.l.) inferred from Plastid DNA Sequences.**

In: Journal of Plant Biology 52 (5), p. 411–416. DOI: 10.1007/s12374-009-9052-8.

### Droseraceae

**Gonella, Paulo M.; Fleischmann, Andreas; Rivadavia, Fernando; Neill, David A.; Sano, Paulo Takeo (2016):**

**A Revision of *Drosera* (Droseraceae) from the central and northern Andes, including a new Species from the Cordillera del Cóndor (Peru and Ecuador).**

In: Plant Systematics and Evolution 302 (10), p. 1419–1432. DOI: 10.1007/s00606-016-1341-3.

**Rivadavia, Fernando; Kondo, Katsuhiko; Kato, Masahiro; Hasebe, Mitsuyasu (2003):**

**Phylogeny of the Sundews, *Drosera* (Droseraceae), based on Chloroplast rbcL and nuclear 18s ribosomal DNA Sequences.**

In: American Journal of Botany 90 (1), p. 123–130. DOI: 10.3732/ajb.90.1.123.

### Dryopteridaceae

**Canestraro, Bianca Kalinowski; Labiak, Paulo Henrique (2015):**

**The Fern Genus *Polybotrya* (Dryopteridaceae) in the Atlantic Forest of Brazil, with the Description of a new Species.**

In: Brittonia 67 (3), p. 191–215. DOI: 10.1007/s12228-015-9369-1.

**Chang, Yi-Han; Kuo, Li-Yaung; Chiou, Wen-Liang; Chen, Cheng-Wei; Chang, Ho-Ming; Amoroso, Victor B. (2013):**

***Stenolepia* Alderw. (Dryopteridaceae), a Fern Genus new to the Philippines.**

In: Philippine Journal of Science 142, p. 83–88.

**Driscoll, Heather E.; Barrington, David p. (2007):**

**Origin of Hawaiian *Polystichum* (Dryopteridaceae) in the Context of a World Phylogeny.**

In: American Journal of Botany 94 (8), p. 1413–1424.

**Geiger, Jennifer M.O.; Ranker, Tom A. (2005):**

**Molecular Phylogenetics and historical Biogeography of Hawaiian *Dryopteris* (Dryopteridaceae).**
In: Molecular Phylogenetics and Evolution 34 (2), p. 392–407. DOI: 10.1016/j.ympev.2004.11.001.

HENNEQUIN, SABINE; ROUHAN, GERMINAL; SALINO, ALEXANDRE; DUAN, YI-FAN; LEPEIGNEUX, MARIE-CAPUCINE; GUILLOU, MARGUERITE ET AL. (2017):
**Global Phylogeny and Biogeography of the Fern Genus *Ctenitis* (Dryopteridaceae), with a focus on the Indian Ocean Region.**
In: Molecular Phylogenetics and Evolution 112, p. 277–289. DOI: 10.1016/j.ympev.2017.04.012.

JUSLÉN, AINO; VÄRE, HENRY; WIKSTRÖM, NIKLAS (2011):
**Relationships and evolutionary Origins of polyploid *Dryopteris* (Dryopteridaceae) from Europe inferred using nuclear pgiC and Plastid trnL-F Sequence Data.**
In: Taxon 60 (5), p. 1284–1294.

LABIAK, PAULO HENRIQUE; SUNDUE, MICHAEL A.; ROUHAN, GERMINAL; HANKS, JUDITH GARRISON; MICKEL, JOHN T.; MORAN, ROBBIN C. (2014):
**Phylogeny and historical Biogeography of the Lastreopsid Ferns (Dryopteridaceae).**
In: American Journal of Botany 101 (7), p. 1207–1228. DOI: 10.3732/ajb.1400071.

LABIAK, PAULO HENRIQUE; SUNDUE, MICHAEL A.; ROUHAN, GERMINAL; MORAN, ROBBIN C. (2015):
**New Combinations in *Lastreopsis* and *Parapolystichum* (Dryopteridaceae).**
In: Brittonia 67 (1), p. 79–86. DOI: 10.1007/s12228-014-9351-3.

LEE, SANG-JUN; PARK, CHONG-WOOK (2013):
**Relationships and Origins of the *Dryopteris varia* (L.) Kuntze Species complex (Dryopteridaceae) in Korea inferred from nuclear and Chloroplast DNA Sequences.**
In: Biochemical Systematics and Ecology 50, p. 371–382. DOI: 10.1016/j.bse.2013.05.008.

LI, CHUN-XIANG; LU, SHU-GANG (2006):
**Phylogenetics of Chinese *Dryopteris* (Dryopteridaceae) based on the Chloroplast rps4-trnS Sequence Data.**
In: Journal of Plant Research 119 (6), p. 589–598. DOI: 10.1007/s10265-006-0003-x.

LI, CHUN-XIANG; LU, SHU-GANG; BARRINGTON, DAVID P. (2008):
**Phylogeny of Chinese *Polystichum* (Dryopteridaceae) based on Chloroplast DNA Sequence Data (trnL-F and rps4-trnS).**
In: Journal of Plant Research 121 (1), p. 19–26. DOI: 10.1007/s10265-007-0120-1.

LI, CHUN-XIANG; SHU-GANG, LU (2006):
**Phylogenetic Analysis of Dryopteridaceae based on Chloroplast rbcL Sequences.**
In: Acta Phytotaxonomica Sinica 44 (5), p. 503–515. DOI: 10.1360/aps050081.

LITTLE, DAMON P.; BARRINGTON, DAVID P. (2003):
**Major evolutionary Events in the Origin and Diversification of the Fern Genus *Polystichum* (Dryopteridaceae).**
In: American Journal of Botany 90 (3), p. 508–514.

LIU, HONG-MEI; ZHANG, XIAN-CHUN; WANG, MEI-PING; SHANG, HUI; ZHOU, SHI-LIANG; YAN, YUE-HONG ET AL. (2016):
**Phylogenetic Placement of the enigmatic Fern Genus *Trichoneuron* informs on the infra-familial Relationship of Dryopteridaceae.**
In: Plant Systematics and Evolution 302 (3), p. 319–332. DOI: 10.1007/s00606-015-1265-3.

LIU, HONG-MEI; ZHANG, XIAN-CHUN; WANG, WEI; QIU, YIN-LONG; CHEN, ZHI-DUAN (2007):

**Molecular Phylogeny of the Fern Family Dryopteridaceae inferred from Chloroplast rbc L and atp B Genes.**
In: International Journal of Plant Sciences 168 (9), p. 1311–1323. DOI: 10.1086/521710.

**Liu, Hong-Mei; Zhang, Xian-Chun; Wang, Wei; Zeng, Hui (2010):**
**Molecular Phylogeny of the endemic Fern Genera *Cyrtomidictyum* and *Cyrtogonellum* (Dryopteridaceae) from East Asia.**
In: Organisms Diversity and Evolution 10 (1), p. 57–68. DOI: 10.1007/s13127-010-0010-6.

**Lóriga, Josmaily; Schmidt, Alexander R.; Moran, Robbin C.; Feldberg, Kathrin; Schneider, Harald; Heinrichs, Jochen (2014):**
**The first fossil of a bolbitidoid Fern belongs to the early-divergent Lineages of *Elaphoglossum* (Dryopteridaceae).**
In: American Journal of Botany 101 (9), p. 1466–1475. DOI: 10.3732/ajb.1400262.

**Lóriga, Josmaily; Vasco, Alejandra; Regalado, Ledis; Heinrichs, Jochen; Moran, Robbin C. (2014):**
**Phylogeny and Classification of the Cuban Species of *Elaphoglossum* (Dryopteridaceae), with Description of *Elaphoglossum* Sect. *Wrightiana* Sect. nov.**
In: Plant Systematics and Evolution 300 (5), p. 937–951. DOI: 10.1007/s00606-013-0933-4.

**Lu, Jin-Mei; Barrington, David S.; Li, De-Zhu (2007):**
**Molecular Phylogeny of the Polystichoid Ferns in Asia based on rbcL Sequences.**
In: Systematic Botany 32 (1), p. 26–33.

**Lyons, Brendan M.; McHenry, Monique A.; Barrington, David p. (2017):**
**Insights into Evolution in Andean *Polystichum* (Dryopteridaceae) from expanded Understanding of the cytosolic Phosphoglucose Isomerase Gene.**
In: Molecular Phylogenetics and Evolution 112, p. 36–46. DOI: 10.1016/j.ympev.2017.04.010.

**McHenry, Monique A.; Barrington, David p. (2014):**
**Phylogeny and Biogeography of exindusiate Andean *Polystichum* (Dryopteridaceae).**
In: American Journal of Botany 101 (2), p. 365–375. DOI: 10.3732/ajb.1300191.

**McHenry, Monique A.; Sundue, Michael A.; Barrington, David p. (2013):**
**The Fern Genus *Adenoderris* (Family incertae sedis) is artificial.**
In: Taxon 62 (6), p. 1153–1160. DOI: 10.12705/626.13.

**Moran, Robbin C.; Hanks, Judith Garrison; Labiak, Paulo Henrique; Sundue, Michael A. (2010):**
**Perispore Morphology of Bolbitidoid Ferns (Dryopteridaceae) in Relation to Phylogeny.**
In: International Journal of Plant Sciences 171 (8), p. 872–881. DOI: 10.1086/655856.

**Moran, Robbin C.; Hanks, Judith Garrison; Rouhan, Germinal (2007):**
**Spore Morphology in Relation to Phylogeny in the Fern Genus *Elaphoglossum* (Dryopteridaceae).**
In: International Journal of Plant Sciences 168 (6), p. 905–929.

**Moran, Robbin C.; Labiak, Paulo Henrique (2015):**
**Phylogeny of the Polybotryoid Fern Clade (Dryopteridaceae).**
In: International Journal of Plant Sciences 176 (9), p. 880–891. DOI: 10.1086/683393.

**Ro, Jacobus P. (2012):**
**A Revision of the Fern Genus *Dryopteris* (Dryopteridaceae) in Sub-Saharan Africa.**
In: Phytotaxa 70, p. 1–118.

**Rouhan, Germinal; Dubuisson, Jean-Yves; Rakotondrainibe, France; Motley, Timothy J.; Mickel, John T.; Labat, Jean-Noel; Moran, Robbin C. (2004):**
**Molecular Phylogeny of the Fern Genus *Elaphoglossum* (Elaphoglossaceae) based on Chloroplast non-coding DNA Sequences: contributions of Species from the Indian Ocean area.**
In: Molecular Phylogenetics and Evolution 33 (3), p. 745–763. DOI: 10.1016/j.ympev.2004.08.006.

**Sano, R.; Takamiya, Masayuki; Ito, Motomi; Kurita, S.; Hasebe, M. (2000):**
**Phylogeny of the lady Fern Group, Tribe Physematieae (Dryopteridaceae), based on Chloroplast rbcL Gene Sequences.**
In: Molecular Phylogenetics and Evolution 15 (3), p. 403–413. DOI: 10.1006/mpev.1999.0708.

**Sessa, Emily B.; Zimmer, Elizabeth A.; Givnish, Thomas J. (2012):**
**Phylogeny, Divergence times, and historical Biogeography of New World *Dryopteris* (Dryopteridaceae).**
In: American Journal of Botany 99 (4), p. 730–750. DOI: 10.3732/ajb.1100294.

**Sessa, Emily B.; Zimmer, Elizabeth A.; Givnish, Thomas J. (2012):**
**Reticulate Evolution on a global scale: a nuclear Phylogeny for New World *Dryopteris* (Dryopteridaceae).**
In: Molecular Phylogenetics and Evolution 64 (3), p. 563–581. DOI: 10.1016/j.ympev.2012.05.009.

**Sessa, Emily B.; Zimmer, Elizabeth A.; Givnish, Thomas J. (2012):**
**Unraveling reticulate Evolution in North American *Dryopteris* (Dryopteridaceae).**
In: BMC Evolutionary Biology 12, p. 104. DOI: 10.1186/1471-2148-12-104.

**Skog, Judith E.; Mickel, John T.; Moran, Robbin C.; Volovsek, Miguel; Zimmer, Elizabeth A. (2004):**
**Molecular Studies of Representative Species in the Fern Genus *Elaphoglossum* (Dryopteridaceae) based on cpDNA Sequences rbcL, trnL-F, and rps4-trnS.**
In: International Journal of Plant Sciences 165 (6), p. 1063–1075.

**Vasco, Alejandra; Moran, Robbin C.; Rouhan, Germinal (2009):**
**Circumscription and Phylogeny of the *Elaphoglossum ciliatum* Group (*E.* Sect. *Lepidoglossa*, Dryopteridaceae) based on cpDNA Sequences.**
In: Taxon 58 (3), p. 825–834.

**Zhang, Li-Bing; He, Hai (2010):**
***Polystichum speluncicola* sp. nov. (Sect. *Haplopolystichum*, Dryopteridaceae) based on Morphological, Palynological, and molecular Evidence with Reference to the Non-Monophyly of *Cyrtogonellum*.**
In: Systematic Botany 35 (1), p. 13–19. DOI: 10.1600/036364410790862605.

**Zhang, Li-Bing; Zhang, Liang (2012):**
**The inclusion of *Acrophorus, Diacalpe, Nothoperanema,* and *Peranema* in *Dryopteris*: the molecular Phylogeny, Systematics, and Nomenclature of *Dryopteris* Subg. *Nothoperanema* (Dryopteridaceae).**
In: Taxon 61 (6), p. 1199–1216.

**Zhang, Li-Bing; Zhang, Liang; Dong, Shi-Yong; Sessa, Emily B.; Gao, Xin-Fen; Ebihara, Atsushi (2012):**
**Molecular Circumscription and major evolutionary Lineages of the Fern Genus *Dryopteris* (Dryopteridaceae).**
In: BMC Evolutionary Biology 12, p. 180. DOI: 10.1186/1471-2148-12-180.

### Ebenaceae

**Berry, Paul E.; Savolainen, Vincent; Sytsma, K. J.; Hall, J.C.; Chase, Mark W. (2001):**
**_Lissocarpa_ is sister to _Diospyros_ (Ebenaceae).**
In: Kew Bulletin 56, p. 725–729.

**Du, Xin-Yu; Zhang, Q. L.; Luo, Zheng-Rong (2009):**
**Comparison of four molecular Markers for genetic Analysis in _Diospyros_ L. (Ebenaceae).**
In: Plant Systematics and Evolution 281 (1-4), p. 171–181. DOI: 10.1007/s00606-009-0199-z.

**Duangjai, Sutee; Samuel, Rosabelle; Munzinger, Jérôme; Forest, Félix; Wallnöfer, Bruno; Barfuss, Michael H. J. et al. (2009):**
**A multi-locus Plastid phylogenetic Analysis of the pantropical Genus _Diospyros_ (Ebenaceae), with an Emphasis on the Radiation and biogeographic Origins of the New Caledonian endemic Species.**
In: Molecular Phylogenetics and Evolution 52 (3), p. 602–620. DOI: 10.1016/j.ympev.2009.04.021.

**Duangjai, Sutee; Wallnöfer, Bruno; Samuel, Rosabelle; Munzinger, Jérôme; Chase, Mark W. (2006):**
**Generic Delimitation and Relationships in Ebenaceae sensu lato: Evidence from six Plastid DNA Regions.**
In: American Journal of Botany 93 (12), p. 1808–1827. DOI: 10.3732/ajb.93.12.1808.

**Paun, Ovidiu; Turner, Barbara; Trucchi, Emiliano; Munzinger, Jérôme; Chase, Mark W.; Samuel, Rosabelle (2016):**
**Processes driving the adaptive Radiation of a tropical Tree (_Diospyros_, Ebenaceae) in New Caledonia, a Biodiversity Hotspot.**
In: Systematic Biology 65 (2), p. 212–227. DOI: 10.1093/sysbio/syv076.

**Turner, Barbara; Munzinger, Jérôme; Duangjai, Sutee; Temsch, Eva M.; Stockenhuber, Reinhold; Barfuss, Michael H. J. et al. (2013):**
**Molecular Phylogenetics of New Caledonian _Diospyros_ (Ebenaceae) using Plastid and nuclear Markers.**
In: Molecular Phylogenetics and Evolution 69 (3), p. 740–763. DOI: 10.1016/j.ympev.2013.07.002.

**Turner, Barbara; Paun, Ovidiu; Munzinger, Jérôme; Duangjai, Sutee; Chase, Mark W.; Samuel, Rosabelle (2013):**
**Analyses of amplified fragment length polymorphisms (Aflp) indicate rapid Radiation of _Diospyros_ Species (Ebenaceae) endemic to New Caledonia.**
In: BMC Evolutionary Biology 13, p. 269. DOI: 10.1186/1471-2148-13-269.

**Venkatasamy, Shadila; Khittoo, Govindranathsing; Nowbuth, Prakash; Vencatasamy, Dass Roy (2006):**
**Phylogenetic Relationships based on Morphology among the _Diospyros_ (Ebenaceae) Species endemic to the Mascarene Islands.**
In: Botanical Journal of the Linnean Society 150 (3), p. 307–313. DOI: 10.1111/j.1095-8339.2006.00474.x.

### Ecdeicoleaceae

**Marchant, Adam; Briggs, Barbara G. (2007):**
**Ecdeiocoleaceae and Joinvilleaceae, sisters of Poaceae (Poales): Evidence from rbcL and matK Data.**
In: Telopea 11 (4), p. 437–450. DOI: 10.7751/telopea20075743.

### Ehretiaceae

**Gottschling, Marc; Hilger, Hartmut H. (2004):**

**The Systematic Position of *Ehretia cortesia* nom. nov. (identical to *Cortesia cuneifolia*: Ehretiaceae, Boraginales) inferred from molecular and morphological Data.**
In: Taxon 53 (4), p. 919. DOI: 10.2307/4135559.

GOTTSCHLING, MARC; LUEBERT, FEDERICO; HILGER, HARTMUT H.; MILLER, JAMES P. (2014):
**Molecular Delimitations in the Ehretiaceae (Boraginales).**
In: Molecular Phylogenetics and Evolution 72, p. 1–6. DOI: 10.1016/j.ympev.2013.12.005.

GOTTSCHLING, MARC; MILLER, JAMES P. (2007):
**A Revision of *Bourreria* (Boraginales, Ehretiaceae) in South America.**
In: Annals of the Missouri Botanical Garden 94 (4), p. 734–744. DOI: 10.3417/0026-6493(2007)94[734:AROBBE]2.0.CO;2.

IRIMIA, RAMONA-ELENA; PÉREZ-ESCOBAR, OSCAR ALEJANDRO; GOTTSCHLING, MARC (2015):
**Strong biogeographic Signal in the phylogenetic Relationships of *Rochefortia* Sw. (Ehretiaceae, Boraginales).**
In: Plant Systematics and Evolution 301 (5), p. 1509–1516. DOI: 10.1007/s00606-014-1162-1.

### Elaeagnaceae

SON, OGYEONG; YOON, CHANG YOUNG; PARK, SEON-JOO (2014):
**Phylogenetic Relationships in Korean *Elaeagnus* L. based on nrDNA ITS Sequences.**
In: Korean Journal of Plant Resources 27 (6), p. 671–679. DOI: 10.7732/kjpr.2014.27.6.671.

SUN, K.; CHEN, X.; MA, R.; LI, C.; WANG, Q.; GE, SONG (2002):
**Molecular Phylogenetics of *Hippophae* L. (Elaeagnaceae) based on the Internal Transcribed Spacer (ITS) Sequences of nrDNA.**
In: Plant Systematics and Evolution 235 (1), p. 121–134. DOI: 10.1007/s00606-002-0206-0.

### Elaeocarpaceae

BUTCHER, RYONEN; BYRNE, MARGARET; CRAYN, DARREN M. (2007):
**Evidence for convergent Evolution among phylogenetically distant rare Species of *Tetratheca* (Elaeocarpaceae, formerly Tremandraceae) from Western Australia.**
In: Australian Systematic Botany 20 (2), p. 126–138. DOI: 10.1071/SB06017.

CRAYN, DARREN M.; ROSSETTO, MAURIZIO; MAYNARD, DAVID J. (2006):
**Molecular Phylogeny and Dating reveals an Oligo-Miocene Radiation of dry-adapted Shrubs (former Tremandraceae) from rainforest Tree Progenitors (Elaeocarpaceae) in Australia.**
In: American Journal of Botany 93 (9), p. 1328–1342. DOI: 10.3732/ajb.93.9.1328.

DOWNING, TRISHA L.; LADIGES, PAULINE Y.; DURETTO, MARCO F. (2008):
**Trichome Morphology provides phylogenetically informative Characters for *Tremandra*, *Platytheca* and *Tetratheca* (former Tremandraceae).**
In: Plant Systematics and Evolution 271 (3-4), p. 199–221. DOI: 10.1007/s00606-007-0622-2.

### Elatinaceae

CAI, LIMING; XI, ZHENXIANG; PETERSON, KYLEE; RUSHWORTH, CATHERINE A.; BEAULIEU, JEREMY M.; DAVIS, CHARLES C. (2016):
**Phylogeny of Elatinaceae and the Tropical Gondwanan Origin of the Centroplacaceae (Malpighiaceae, Elatinaceae) Clade.**
In: Public Library of Science One 11 (9), e0161881. DOI: 10.1371/journal.pone.0161881.

### Ephedraceae

**Huang, Jinling; Giannasi, David E.; Price, Robert A. (2005):**

**Phylogenetic Relationships in *Ephedra* (Ephedraceae) inferred from Chloroplast and nuclear DNA Sequences.**

In: Molecular Phylogenetics and Evolution 35 (1), p. 48–59. DOI: 10.1016/j.ympev.2004.12.020.

**Ickert-Bond, Stefanie M.; Wojciechowski, Martin F. (2004):**

**Phylogenetic Relationships in *Ephedra* (Gnetales): Evidence from Nuclear and Chloroplast DNA Sequence Data.**

In: Systematic Botany 29 (4), p. 834–849. DOI: 10.1600/0363644042451143.

**Loera, Israel; Sosa, Victoria; Ickert-Bond, Stefanie M. (2012):**

**Diversification in North American arid Lands: Niche Conservatism, Divergence and Expansion of habitat explain Speciation in the Genus *Ephedra*.**

In: Molecular Phylogenetics and Evolution 65 (2), p. 437–450. DOI: 10.1016/j.ympev.2012.06.025.

**Rydin, Catarina; Korall, Petra (2009):**

**Evolutionary Relationships in *Ephedra* (Gnetales), with Implications for Seed Plant Phylogeny.**

In: International Journal of Plant Sciences 170 (8), p. 1031–1043. DOI: 10.1086/605116.

### Equisetaceae

**Des Marais, David L.; Smith, Alan R.; Britton, Donald M.; Pryer, Kathleen M. (2003):**

**Phylogenetic Relationships and Evolution of extant Horsetails, *Equisetum*, based on Chloroplast DNA Sequence Data (rbcL and trnL-F).**

In: International Journal of Plant Sciences 164 (5), p. 737–751.

**Guillon, Jean-Michel (2004):**

**Phylogeny of Horsetails (*Equisetum*) based on the Chloroplast rps4 Gene and Adjacent Noncoding Sequences.**

In: Systematic Botany 29 (2), p. 251–259. DOI: 10.1600/036364404774195467.

**Guillon, Jean-Michel (2007):**

**Molecular Phylogeny of horsetails (*Equisetum*) including Chloroplast atpB Sequences.**

In: Journal of Plant Research 120 (4), p. 569–574. DOI: 10.1007/s10265-007-0088-x.

### Ericaceae

**Albrecht, D. E.; Owens, C. T.; Weiller, C. M.; Quinn, Christopher J. (2010):**

**Generic Concepts in Ericaceae: Styphelioideae - the *Monotoca* Group.**

In: Australian Systematic Botany 23 (5), p. 320–332. DOI: 10.1071/SB10009.

**Argent, G. (2014):**

**A Contribution to the Study of the Genus *Diplycosia* (Ericaceae) in Sulawesi, Indonesia.**

In: Edinburgh Journal of Botany 71 (1), p. 83–115. DOI: 10.1017/S0960428613000309.

**Boykin, Laura M.; Vasey, Michael C.; Parker, V. Thomas; Patterson, Robert W. (2005):**

**Two Lineages of *Arctostaphylos* (Ericaceae) identified using the Internal Transcribed Spacer (ITS) Region of the nuclear Genome.**

In: Madroño 52 (3), p. 139–147. DOI: 10.3120/0024-9637(2005)52[139:TLOAEI]2.0.CO;2.

**Brown, G. K.; Craven, L. A.; Udovicic, Frank; Ladiges, Pauline Y. (2006):**

**Phylogeny of *Rhododendron Section Vireya* (Ericaceae) based on two non-coding Regions of cpDNA.**
In: Plant Systematics and Evolution 257 (1-2), p. 57–93. DOI: 10.1007/s00606-005-0367-8.

**BROWN, GILLIAN K.; CRAVEN, LYN A.; UDOVICIC, FRANK; LADIGES, PAULINE Y. (2006):**
**Phylogenetic Relationships of *Rhododendron Section Vireya* (Ericaceae) inferred from the ITS nrDNA Region.**
In: Australian Systematic Botany 19 (4), p. 329–342. DOI: 10.1071/SB05019.

**BUSH, CATHERINE M.; JUDD, WALTER S.; FRAZIER, LOUIS; KRON, KATHLEEN A. (2010):**
**The Phylogeny of *Leucothoë* s. l. (Ericaceae: Vaccinioideae) based on morphological and molecular (ndhF, matK, and nrITS) Data.**
In: Systematic Botany 35 (1), p. 201–206. DOI: 10.1600/036364410790862443.

**BUSH, CATHERINE M.; KRON, KATHLEEN A. (2008):**
**A Phylogeny of *Bejaria* (Ericaceae: Ericoideae) based on molecular Data.**
In: Journal of the Botanical Research Institute of Texas 2 (2), p. 1193–1205.

**BUSH, CATHERINE M.; LU, LU; FRITSCH, PETER W.; LI, DE-ZHU; KRON, KATHLEEN A. (2009):**
**Phylogeny of Gaultherieae (Ericaceae: Vaccinioideae) based on DNA Sequence Data from matK, ndhF, and nrITS.**
In: International Journal of Plant Sciences 170 (3), p. 355–364. DOI: 10.1086/596330.

**BUSH, CATHERINE M.; WAGSTAFF, STEVEN J.; FRITSCH, PETER W.; KRON, KATHLEEN A. (2009):**
**The Phylogeny, Biogeography and morphological Evolution of *Gaultheria* (Ericaceae) from Australia and New Zealand.**
In: Australian Systematic Botany 22 (4), p. 229. DOI: 10.1071/SB08049.

**CHUNG, JENG-DER; LIN, TSAN-PIAO; CHEN, YU-LING; CHENG, YU-PIN; HWANG, SHIH-YING (2007):**
**Phylogeographic Study reveals the Origin and evolutionary History of a *Rhododendron* Species complex in Taiwan.**
In: Molecular Phylogenetics and Evolution 42 (1), p. 14–24. DOI: 10.1016/j.ympev.2006.06.027.

**CRAYN, DARREN M.; QUINN, CHRISTOPHER J. (2000):**
**The Evolution of the atpbeta-rbcL intergenic spacer in the Epacrids (Ericales) and its systematic and evolutionary Implications.**
In: Molecular Phylogenetics and Evolution 16 (2), p. 238–252. DOI: 10.1006/mpev.2000.0794.

**DORR, LAURENCE J. (2006):**
**New Combinations in *Erica* (Ericaceae: Ericoideae) from the High Mountains of East Africa.**
In: Novon: A Journal for Botanical Nomenclature 16 (1), p. 56–58. DOI: 10.3417/1055-3177(2006)16[56:NCIEEE]2.0.CO;2.

**FLOYD, JENNIFER WHITEHEAD (2002):**
**Phylogenetic and Biogeographic Patterns in *Gaylussacia* (Ericaceae) based on Morphological, nuclear DNA, and Chloroplast DNA Variation.**
In: Systematic Botany 27 (1), p. 99–115.

**FRITSCH, PETER W.; LU, LU; BUSH, CATHERINE M.; CRUZ, BONI C.; KRON, KATHLEEN A.; LI, DE-ZHU (2011):**
**Phylogenetic Analysis of the Wintergreen Group (Ericaceae) based on six genic Regions.**
In: Systematic Botany 36 (4), p. 990–1003. DOI: 10.1600/036364411X604994.

**GILLESPIE, EMILY; KRON, KATHLEEN A. (2010):**

**Molecular phylogenetic Relationships and a revised Classification of the Subfamily Ericoideae (Ericaceae).**
In: Molecular Phylogenetics and Evolution 56 (1), p. 343–354. DOI: 10.1016/j.ympev.2010.02.028.

**Goetsch, Loretta A.; Craven, Lyn A.; Hall, Benjamin D. (2011):**
**Major Speciation accompanied the Dispersal of *Vireya* Rhododendrons (Ericaceae, *Rhododendron* Sect. *Schistanthe*) through the Malayan archipelago: Evidence from nuclear Gene Sequences.**
In: Taxon 60 (4), p. 1015–1028.

**Goetsch, Loretta A.; Eckert, Andrew J.; Hall, Benjamin D. (2005):**
**The molecular Systematics of *Rhododendron* (Ericaceae): A Phylogeny based Upon RPB2 Gene Sequences.**
In: Systematic Botany 30 (3), p. 616–626.

**Hart, Robbie; Georgian, Elizabeth M.; Salick, Jan (2016):**
**Fast and cheap in the fall: phylogenetic determinants of late flowering phenologies in Himalayan *Rhododendron*.**
In: American Journal of Botany 103 (2), p. 198–206. DOI: 10.3732/ajb.1500440.

**Hileman, Lena C.; Vasey, Michael C.; Parker, V. Thomas (2001):**
**Phylogeny and Biogeography of the Arbutoideae (Ericaceae): Implications for the Madrean-Tethyan Hypothesis.**
In: Systematic Botany 26 (1), p. 131–143.

**Hirai, Masashi; Yoshimura, Sayuri; Ohsako, Takanori; Kubo, Nakao (2010):**
**Genetic Diversity and phylogenetic Relationships of the endangered Species** *Vaccinium sieboldii* and *Vaccinium ciliatum* **(Ericaceae).**
In: Plant Systematics and Evolution 287 (1-2), p. 75–84. DOI: 10.1007/s00606-010-0291-4.

**Hislop, Michael; Puente-Lelièvre, Caroline; Crayn, Darren M. (2012):**
***Leucopogon extremus* (Styphelieae, Styphelioideae, Ericaceae), a remarkable new Species that expands the morphological Circumscription of *Leucopogon* sens. str.**
In: Australian Systematic Botany 25 (3), p. 202–209. DOI: 10.1071/SB11033.

**Hou, Yan; Nowak, Michael D.; Mirré, Virginia; Bjorå, Charlotte Sletten; Brochmann, Christian; Popp, Magnus (2016):**
**RAD-seq Data point to a northern Origin of the arctic-alpine Genus *Cassiope* (Ericaceae).**
In: Molecular Phylogenetics and Evolution 95, p. 152–160. DOI: 10.1016/j.ympev.2015.11.009.

**Ikeda, Hajime; Yakubov, Valentin; Barkalov, Vyacheslav; Setoguchi, Hiroaki (2014):**
**Molecular Evidence for ancient Relicts of arctic-alpine plants in East Asia.**
In: the new Phytologist 203 (3), p. 980–988. DOI: 10.1111/nph.12863.

**Jin, Xiao-Feng; Ding, Bing-Yang; Jin, Shui-Hu; Qian, Li (2009):**
**Taxonomic Revision of *Rhododendron mariae* (Ericaceae) and related taxa.**
In: Nordic Journal of Botany 27 (3), p. 186–202. DOI: 10.1111/j.1756-1051.2009.00316.x.

**Johnson, Karen A.; Holland, Barbara R.; Heslewood, Margaret M.; Crayn, Darren M. (2012):**
**Supermatrices, Supertrees and serendipitous scaffolding: inferring a well-resolved, Genus-level Phylogeny of Styphelioideae (Ericaceae) despite missing Data.**
In: Molecular Phylogenetics and Evolution 62 (1), p. 146–158. DOI: 10.1016/j.ympev.2011.09.011.

**Kron, Kathleen A.; Powell, E. Ann; Luteyn, James L. (2002):**

**Phylogenetic Relationships within the Blueberry Tribe (Vaccineae, Ericaceae) based on Sequence Data from matK and nuclear ribosomal ITS Regions, with Comments on the Placements of *Satyria*.**
In: American Journal of Botany 89 (2), p. 327–336.

**Kurashige, Y.; Etoh, J.-I.; Handa, Takashi; Takayanagi, K.; Yukawa, Tomohisa (2001):**
**Sectional Relationships in the Genus *Rhododendron* (Ericaceae): Evidence from matK and trn K Intron Sequences.**
In: Plant Systematics and Evolution 228 (1-2), p. 1–14. DOI: 10.1007/s006060170033.

**Li, J. (2002):**
**Phylogenetic Relationships of Empetraceae inferred from Sequences of Chloroplast Gene matK and nuclear ribosomal DNA ITS Region.**
In: Molecular Phylogenetics and Evolution 25 (2), p. 306–315. DOI: 10.1016/S1055-7903(02)00241-5.

**Li, Mimi; Li, Jian-Hua; Jiang, Jin-Huo; Fu, Cheng-Xin; Judd, Walter p. (2009):**
**Phylogenetics and Biogeography of *Pieris* (Lyonieae, Ericaceae) inferred from Sequences of Nuclear and Chloroplast Genomes.**
In: Systematic Botany 34 (3), p. 553–560. DOI: 10.1600/036364409789271272.

**Lian-Ming, Gao; De-Zhu, Li; Chang-Qin, Zhang (2003):**
**Phylogenetic Relationships of *Rhododendron Section Azaleastrum* (Ericaceae) based on ITS Sequences.**
In: Acta Phytotaxonomica Sinica 41 (2), p. 173–179.

**Liu, Zhen-Wen; Wang, Ze-Huan; Zhou, Jing; Peng, Hua (2011):**
**Phylogeny of Pyroleae (Ericaceae): Implications for Character Evolution.**
In: Journal of Plant Research 124 (3), p. 325–337. DOI: 10.1007/s10265-010-0376-8.

**Liu, Zhen-Wen; Zhou, Jing; Liu, En-De; Peng, Hua (2010):**
**A molecular Phylogeny and a new Classification of *Pyrola* (Pyroleae, Ericaceae).**
In: Taxon 59 (6), p. 1690–1700.

**McGuire, Avery F.; Kron, Kathleen A. (2005):**
**Phylogenetic Relationships of European and African Ericas.**
In: International Journal of Plant Sciences 166 (2), p. 311–318.

**Milne, Richard Ian (2004):**
**Phylogeny and Biogeography of *Rhododendron* subsection *Pontica*, a Group with a tertiary Relict Distribution.**
In: Molecular Phylogenetics and Evolution 33 (2), p. 389–401. DOI: 10.1016/j.ympev.2004.06.009.

**Milne, Richard Ian; Davies, Chantel; Prickett, Ruby; Inns, Lucy H.; Chamberlain, David F. (2010):**
**Phylogeny of *Rhododendron* Subgenus *Hymenanthes* based on Chloroplast DNA Markers: between-lineage Hybridisation during adaptive Radiation?**
In: Plant Systematics and Evolution 285 (3-4), p. 233–244. DOI: 10.1007/s00606-010-0269-2.

**Mugrabi de Kuppler, A. L.; Fagúndez, J.; Bellstedt, Dirk U.; Oliver, Edward G.H.; Léon, J.; Pirie, Michael D. (2015):**
**Testing reticulate versus coalescent Origins of *Erica lusitanica* using a Species Phylogeny of the northern Heathers (Ericeae, Ericaceae).**
In: Molecular Phylogenetics and Evolution 88, p. 121–131. DOI: 10.1016/j.ympev.2015.04.005.

**Oliver, Edward G.H.; Oliver, Inge M. (1991):**

**Studies in the Ericoideae (Ericaceae). VIII. new Species in *Erica*, Section *Pseuderemia*, from southern Africa.**
In: Bothalia 21 (2), p. 137–142. DOI: 10.4102/abc.v21i2.873.

PEDRAZA-PEÑALOSA, PAOLA (2009):
**Insensitive blueberries: a total-Evidence Analysis of *Disterigma* s.l. (Ericaceae) exploring Transformation Costs.**
In: Cladistics 26 (1), p. 388–407. DOI: 10.1111/j.1096-0031.2009.00293.x.

PEDRAZA-PEÑALOSA, PAOLA (2009):
**Systematics of the Neotropical Blueberry Genus *Disterigma* (Ericaceae).**
In: Systematic Botany 34 (2), p. 406–413. DOI: 10.1600/036364409788606352.

PEDRAZA-PEÑALOSA, PAOLA (2016):
**Three new Species of *Psammisia* s.l. (Ericaceae: Vaccinieae), Blueberry Relatives endemic to Colombia.**
In: Brittonia 68 (1), p. 25–36. DOI: 10.1007/s12228-015-9388-y.

PEDRAZA-PEÑALOSA, PAOLA; LUTEYN, JAMES L. (2011):
**Andean *Vaccinium* (Ericaceae: Vaccinieae): Seven new Species from South America.**
In: Brittonia 63 (2), p. 257–275.

PEDRAZA-PEÑALOSA, PAOLA; SALINAS, NELSON R.; VIRNIG, ANNE LUCY S.; WHEELER, WARD C. (2015):
**Preliminary phylogenetic Analysis of the Andean Clade and the Placement of new Colombian Blueberries (Ericaceae, Vaccinieae).**
In: PhytoKeys (49), p. 13–31. DOI: 10.3897/phytokeys.49.8622.

PEDRAZA-PEÑALOSA, PAOLA; SALINAS, NELSON R.; WHEELER, WARD C. (2013):
**Venation Patterns of Neotropical Blueberries (Vaccinieae: Ericaceae) and their phylogenetic utility.**
In: Phytotaxa 96 (1), p. 1. DOI: 10.11646/phytotaxa.96.1.1.

PIRIE, MICHAEL D.; OLIVER, EDWARD G.H.; BELLSTEDT, DIRK U. (2011):
**A densely sampled ITS Phylogeny of the Cape flagship Genus *Erica* L. suggests numerous shifts in Floral Macro-Morphology.**
In: Molecular Phylogenetics and Evolution 61 (2), p. 593–601. DOI: 10.1016/j.ympev.2011.06.007.

POWELL, E. ANN; KRON, KATHLEEN A. (2002):
**Hawaiian Blueberries and their Relatives—A phylogenetic Analysis of *Vaccinium* Sections *Macropelma, Myrtillus,* and *Hemimyrtillus* (Ericaceae).**
In: Systematic Botany 27 (4), p. 768–779.

POWELL, E. ANN; KRON, KATHLEEN A. (2003):
**Molecular Systematics of the Northern Andean Blueberries (Vaccinieae, Vaccinioideae, Ericaceae).**
In: International Journal of Plant Sciences 164 (6), p. 987–995.

PUENTE-LELIÈVRE, CAROLINE; HARRINGTON, MARK G.; BROWN, ELIZABETH A.; KUZMINA, MARIA; CRAYN, DARREN M. (2013):
**Cenozoic extinction and Recolonization in the New Zealand Flora: the case of the fleshy-fruited Epacrids (Styphelieae, Styphelioideae, Ericaceae).**
In: Molecular Phylogenetics and Evolution 66 (1), p. 203–214. DOI: 10.1016/j.ympev.2012.09.027.

PUENTE-LELIÈVRE, CAROLINE; HISLOP, MICHAEL; HARRINGTON, MARK G.; BROWN, ELIZABETH A.; KUZMINA, MARIA; CRAYN, DARREN M. (2015):

**A five-marker molecular Phylogeny of the Styphelieae (Epacridoideae, Ericaceae) supports a broad Concept of *Styphelia*.**
In: Australian Systematic Botany 28 (6), p. 368–387. DOI: 10.1071/SB14041.

QUINN, CHRISTOPHER J.; CROWDEN, RONALD K.; BROWN, ELIZABETH A.; SOUTHAM, MICHAEL J.; THORNHILL, ANDREW H.; CRAYN, DARREN M. (2015):
**A reappraisal of the generic Concepts of *Epacris, Rupicola* and *Budawangia* (Ericaceae, Epacridoideae, Epacrideae) based on phylogenetic Analysis of morphological and molecular Data.**
In: Australian Systematic Botany 28 (1), p. 63–77. DOI: 10.1071/SB13009.

SCHWERY, ORLANDO; ONSTEIN, RENSKE E.; BOUCHENAK-KHELLADI, YANIS; XING, YAOWU; CARTER, RICHARD J.; LINDER, HANS PETER (2015):
**As old as the Mountains: the Radiations of the Ericaceae.**
In: the new Phytologist 207 (2), p. 355–367. DOI: 10.1111/nph.13234.

SETOGUCHI, HIROAKI; WATANABE, W.; MAEDA, Y.; PENG, CHING-I. (2008):
**Molecular Phylogeny of the Genus *Pieris* (Ericaceae) with special Reference to phylogenetic Relationships of insular Plants on the Ryukyu Islands.**
In: Plant Systematics and Evolution 270 (3-4), p. 217–230. DOI: 10.1007/s00606-007-0600-8.

TONG, YI HUA; XIA, NIAN-HE (2014):
**New Taxa of *Agapetes* (Ericaceae) from Myanmar.**
In: Phytotaxa 184 (1), p. 39. DOI: 10.11646/phytotaxa.184.1.5.

TSAI, CHI-CHU; CHEN, CHIH-HUI; CHOU, CHANG-HUNG (2012):
**DNA barcodes reveal high levels of morphological plasticity among *Rhododendron* Species (Ericaceae) in Taiwan.**
In: Biochemical Systematics and Ecology 40, p. 169–177. DOI: 10.1016/j.bse.2011.10.013.

TSUKAYA, HIROKAZU; YOKOYAMA, JUN U.N.; IMAICHI, RYOKO; OHBA, HIDEAKI (2008):
**Taxonomic Status of *Monotropastrum humile*, with special Reference to *M. humile* var. *glaberrimum* (Ericaceae, Monotropoideae).**
In: Journal of Plant Research 121 (3), p. 271–278. DOI: 10.1007/s10265-008-0157-9.

VAN DER KLOET, SAM P.; BALTZER, JENNIFER L.; APPLEBY, JULIA H.; EVANS, RODGER C.; STEWART, DONALD T. (2004):
**A re-examination of the taxonomic boundaries of *Symphysia* (Ericaceae).**
In: Taxon 53 (1), p. 91–98. DOI: 10.2307/4135492.

VANDER KLOET, P. P.; DICKINSON, TIMOTHY A. (2009):
**A subgeneric Classification of the Genus *Vaccinium* and the metamorphosis of *V.* Section *Bracteata* Nakai: more terrestrial and less epiphytic in Habit, more continental and less insular in Distribution.**
In: Journal of Plant Research 122 (3), p. 253–268. DOI: 10.1007/s10265-008-0211-7.

WAGSTAFF, STEVEN J.; DAWSON, MURRAY I.; VENTER, STEPHANUS; MUNZINGER, JÉRÔME; CRAYN, DARREN M.; STEANE, DOROTHY A.; LEMSON, KRISTINA L. (2010):
**Origin, Diversification, and Classification of the Australasian Genus *Dracophyllum* (Richeeae, Ericaceae).**
In: Annals of the Missouri Botanical Garden 97 (2), p. 235–258. DOI: 10.3417/2008130.

WASELKOV, KATHERINE; JUDD, WALTER P. (2008):
**A phylogenetic Analysis of *Leucothoe* s.l. (Ericaceae; Tribe Gaultherieae) base on phenotypic Characters.**
In: Brittonia 60 (4), 382-297.

ZHANG, MING-YING; FRITSCH, PETER W.; MA, PENG-FEI; WANG, HONG; LU, LU; LI, DE-ZHU (2017):

**Plastid phylogenomics and adaptive Evolution of *Gaultheria* series *Trichophyllae* (Ericaceae), a Clade from Sky Islands of the Himalaya-Hengduan Mountains.**

In: Molecular Phylogenetics and Evolution 110, p. 7–18. DOI: 10.1016/j.ympev.2017.01.015.

### Ericales

ROSE, JEFFREY P.; KLEIST, THOMAS J.; LÖFSTRAND, STEFAN D.; DREW, BRYAN T.; SCHÖNENBERGER, JÜRGEN; SYTSMA, KENNETH J. (2018):

**Phylogeny, historical Biogeography, and Diversification of Angiosperm Order Ericales suggest ancient Neotropical and East Asian Connections.**

In: Molecular Phylogenetics and Evolution 122, p. 59–79. DOI: 10.1016/j.ympev.2018.01.014.

### Eriocaulaceae

ANDRADE, MARIA JOSÉ GOMES DE; GIULIETTI, ANA MARIA; HARLEY, RAYMOND MERVYN; VAN DEN BERG, CÁSSIO (2011):

***Blastocaulon* (Eriocaulaceae), a synonym of *Paepalanthus*: morphological and molecular Evidence.**

In: Taxon 60 (1), p. 178–184.

ANDRADE, MARIA JOSÉ GOMES DE; GIULIETTI, ANA MARIA; RAPINI, ALESSANDRO; QUEIROZ, LUCIANO PAGANUCCI; CONCEIÇÃO, ADILVA DE SOUZA; ALMEIDA, PAULO RICARDO MACHADO DE; VAN DEN BERG, CÁSSIO (2010):

**A comprehensive phylogenetic Analysis of Eriocaulaceae: Evidence from nuclear (ITS) and Plastid (psbA-trnH and trnL-F) DNA Sequences.**

In: Taxon 59 (2), p. 379–388.

COSTA, FABIANE NEPOMUEENO (2013):

**New Circumscription of the Endemic Brazilian Genus *Actinocephalus* (Eriocaulaceae).**

In: Novon: A Journal for Botanical Nomenclature 22, p. 281–287.

DAVIES, RICHARD J.-P.; CRAIGIE, ANDREW I.; MACKAY, DUNCAN A.; WHALEN, MOLLY A.; CHEONG, JUDY P.-E.; LEACH, GREGORY J. (2007):

**Resolution of the Taxonomy of *Eriocaulon* (Eriocaulaceae) Taxa endemic to Australian mound Springs, using Morphometrics and AFLP Markers.**

In: Australian Systematic Botany 20 (5), p. 428–447. DOI: 10.1071/SB07019.

ECHTERNACHT, LIVIA; SANO, PAULO TAKEO; BONILLO, CÉLINE; CRUAUD, CORINNE; COULOUX, ARNAUD; DUBUISSON, JEAN-YVES (2014):

**Phylogeny and Taxonomy of *Syngonanthus* and *Comanthera* (Eriocaulaceae): Evidence from expanded Sampling.**

In: Taxon 63 (1), p. 47–63. DOI: 10.12705/631.36.

ECHTERNACHT, LIVIA; SANO, PAULO TAKEO; DUBUISSON, JEAN-YVES (2015):

**Taxonomic Study of *Comanthera* Subg. *Thysanocephalus* (Eriocaulaceae).**

In: Systematic Botany 40 (1), p. 136–150. DOI: 10.1600/036364415X686431.

ECHTERNACHT, LIVIA; SANO, PAULO TAKEO; TROVÓ, MARCELO; DUBUISSON, JEAN-YVES (2011):

**Phylogenetic Analysis of the Brazilian microendemic *Paepalanthus* Subgenus *Xeractis* (Eriocaulaceae) inferred from Morphology.**

In: Botanical Journal of the Linnean Society 167 (2), p. 137–152. DOI: 10.1111/j.1095-8339.2011.01170.x.

GIULIETTI, ANA MARIA; ANDRADE, MARIA JOSÉ GOMES DE; SCATENA, VERA LUCIA; TROVÓ, MARCELO; COAN, ALESSANDRA I.; SANO, PAULO TAKEO ET AL. (2012):

**Molecular Phylogeny, Morphology and their Implications for the Taxonomy of Eriocaulaceae.**

In: Rodriguésia 63 (1), p. 1–19.

**Parra, Lara Regina; Giulietti, Ana Maria; Andrade, Maria José Gomes De; van den Berg, Cássio (2010):**
**Re-establishment and new Circumscription of *Comanthera* (Eriocaulaceae).**
In: Taxon 59 (4), p. 1135–1146.

**Phillips, Sylvia M.; Mesterházy, Attila (2015):**
**Revision of small ephemeral Species of *Eriocaulon* (Eriocaulaceae) in West Africa with long involucral Bracts.**
In: Kew Bulletin 70, p. 1–17. DOI: 10.1007/S12225-014-9557-2.

**Sano, Paulo Takeo (2004):**
***Actinocephalus* (Körn.) Sano (*Paepalanthus* Sect. *Actinocephalus*), a new Genus of Eriocaulaceae, and other taxonomic and Nomenclatural Changes Involving *Paepalanthus* Mart.**
In: Taxon 53 (1), p. 99–107. DOI: 10.2307/4135493.

**Scatena, Vera Lucia; Giulietti, Ana Maria; Borba, Eduardo L.; van den Berg, Cássio (2005):**
**Anatomy of Brazilian Eriocaulaceae: correlation with Taxonomy and habitat using multivariate Analyses.**
In: Plant Systematics and Evolution 253 (1-4), p. 1–22. DOI: 10.1007/s00606-004-0295-z.

**Trovó, Marcelo; Andrade, Maria José Gomes De; Sano, Paulo Takeo; Ribeiro, Patrícia Luz; van den Berg, Cássio (2013):**
**Molecular Phylogenetics and Biogeography of Neotropical Paepalanthoideae with Emphasis on Brazilian *Paepalanthus* (Eriocaulaceae).**
In: Botanical Journal of the Linnean Society 171, p. 225–243.

**Trovó, Marcelo; Sano, Paulo Takeo (2010):**
**Taxonomic survey of *Paepalanthus Section Diphyomene* (Eriocaulaceae).**
In: Phytotaxa 14 (1), p. 49–55. DOI: 10.11646/phytotaxa.14.1.4.

**Unwin, Matthew M. (2004):**
**Molecular Systematics of the Eriocaulaceae Martinov.**
Dissertation. Miami University, Oxford, Ohio.

**Watanabe, Mauricio Takashi Coutinho; Hensold, Nancy; Sano, Paulo Takeo (2015):**
**Tidying up the mess: Lectotype Selections, Synonyms, a new Status and a new Species in *Syngonanthus* Sect. *Carphocephalus* (Eriocaulaceae).**
In: Phytotaxa 226 (2), p. 157. DOI: 10.11646/phytotaxa.226.2.5.

### Erythroxylaceae

**Emche, Stephen D.; Zhang, Dapeng; Islam, Melissa Bauer; Bailey, Bryan A.; Meinhardt, Lyndel W. (2011):**
**AFLP Phylogeny of 36 *Erythroxylum* Species.**
In: Tropical Plant Biology 4 (2), p. 126–133. DOI: 10.1007/s12042-011-9070-9.

### Escalloniaceae

**Esser, Hans-Joachim; Saw, Leng Guan (2015):**
**A new Species of *Polyosma* (Escalloniaceae / Polyosmaceae) from Thailand and new Synonyms.**
In: Phytotaxa 221 (1), p. 89. DOI: 10.11646/phytotaxa.221.1.10.

**Sede, Silvana M.; Dürnhöfer, Sofía I.; Morello, Santiago; Zapata, Felipe (2013):**
**Phylogenetics of *Escallonia* (Escalloniaceae) based on Plastid DNA Sequence Data.**
In: Botanical Journal of the Linnean Society 173 (3), p. 442–451. DOI: 10.1111/boj.12091.

**Zapata, Felipe (2013):**

**A multilocus phylogenetic Analysis of *Escallonia* (Escalloniaceae): Diversification in montane South America.**
In: American Journal of Botany 100 (3), p. 526–545. DOI: 10.3732/ajb.1200297.

### Euphorbiaceae

AUBRIOT, XAVIER; LOWRY, PORTER PRESCOTT II.; HAEVERMANS, THOMAS (2014):
**Taxonomic Revision of the Malagasy endemic and enigmatic *Euphorbia Section Pachysanthae* (Euphorbiaceae).**
In: Phytotaxa 159 (3), p. 221. DOI: 10.11646/phytotaxa.159.3.5.

BARRES, LAIA; VILATERSANA, ROSER; MOLERO, JULIÀ; SUSANNA, ALFONSO; GALBANY-CASALS, MERCÈ (2011):
**Molecular Phylogeny of *Euphorbia* Subg. *Esula* Sect. *Aphyllis* (Euphorbiaceae) inferred from nrDNA and cpDNA Markers with biogeographic Insights.**
In: Taxon 60 (3), p. 705–720. DOI: 10.1002/tax.603007.

BERRY, PAUL E.; HIPP, ANDREW L.; WURDACK, KENNETH J.J.; VAN EE, BENJAMIN W.; RIINA, RICARDA (2005):
**Molecular Phylogenetics of the giant Genus *Croton* and Tribe Crotoneae (Euphorbiaceae sensu stricto) using ITS and Trnl-trnf DNA Sequence Data.**
In: American Journal of Botany 92 (9), p. 1520–1534. DOI: 10.3732/ajb.92.9.1520.

BLATTNER, FRANK R.; WEISING, KURT; BÄNFER, G.; MASCHWITZ, ULRICH; FIALA, B. (2001):
**Molecular Analysis of phylogenetic Relationships among Myrmecophytic *Macaranga* Species (Euphorbiaceae).**
In: Molecular Phylogenetics and Evolution 19 (3), p. 331–344. DOI: 10.1006/mpev.2001.0941.

BRUYNS, PETER V.; KLAK, CORNELIA; HANÁČEK, PAVEL (2011):
**Age and Diversity in Old World succulent Species of *Euphorbia* (Euphorbiaceae).**
In: Taxon 60 (6), p. 1717–1733.

BRUYNS, PETER V.; MAPAYA, RUVIMBO J.; HEDDERSON, TERRENCE J. (2006):
**A new subgeneric Classification for *Euphorbia* (Euphorbiaceae) in southern Africa based on ITS and psbA-trnH Sequence Data.**
In: Taxon 55 (2), p. 397–420. DOI: 10.2307/25065587.

CARDINAL-MCTEAGUE, WARREN M.; GILLESPIE, LYNN J. (2016):
**Molecular Phylogeny and Pollen Evolution of Euphorbiaceae Tribe Plukenetieae.**
In: Systematic Botany 41 (2), p. 329–347. DOI: 10.1600/036364416X691759.

CARUZO, MARIA BEATRIZ R.; VAN EE, BENJAMIN W.; CORDEIRO, INÊS; BERRY, PAUL E.; RIINA, RICARDA (2011):
**Molecular Phylogenetics and Character Evolution of the "*Sacaca*" Clade: novel Relationships of *Croton Section Cleodora* (Euphorbiaceae).**
In: Molecular Phylogenetics and Evolution 60 (2), p. 193–206. DOI: 10.1016/j.ympev.2011.04.013.

CHACÓN, JULIANA; MADRIÑÁN, SANTIAGO; DEBOUCK, DANIEL; RODRIGUEZ, FAUSTO; TOHME, JOE (2008):
**Phylogenetic Patterns in the Genus *Manihot* (Euphorbiaceae) inferred from Analyses of nuclear and Chloroplast DNA Regions.**
In: Molecular Phylogenetics and Evolution 49 (1), p. 260–267. DOI: 10.1016/j.ympev.2008.07.015.

DE-NOVA, JOSÉ ARTURO; SOSA, VICTORIA (2007):
**Phylogeny and generic Delimitation of *Adelia* (Euphorbiaceae) inferred from molecular and morphological Data.**
In: Taxon 56 (4), p. 1027–1036. DOI: 10.2307/25065902.

DE-NOVA, JOSÉ ARTURO; SOSA, VICTORIA; STEINMANN, VICTOR W. (2007):
**A Synopsis of *Adelia* (Euphorbiaceae s.s.).**

In: Systematic Botany 32 (3), p. 583–595.

**De-Nova, José Arturo; Sosa, Victoria; Wurdack, Kenneth J.J. (2006):**

**Phylogenetic Relationships and the Description of a new Species of *Enriquebeltrania* (Euphorbiaceae s.s.): An Enigmatic Genus Endemic to Mexico.**

In: Systematic Botany 31 (3), p. 533–546.

**Dorsey, Brian L.; Haevermans, Thomas; Aubriot, Xavier; Morawetz, Jeffery J.; Riina, Ricarda; Steinmann, Victor W.; Berry, Paul E. (2013):**

**Phylogenetics, morphological Evolution, and Classification of *Euphorbia* Subgenus *Euphorbia*.**

In: Taxon 62 (2), p. 291–315. DOI: 10.12705/622.1.

**Fatemi, M.; Gross, C. L.; Bruhl, J. J. (2007):**

**The first phenetic Analysis of Species limits in *Bertya* (Euphorbiaceae).**

In: Australian Systematic Botany 20 (5), p. 448–463. DOI: 10.1071/SB06019.

**Frajman, Božo; Schönswetter, Peter (2011):**

**Giants and dwarfs: molecular Phylogenies reveal multiple Origins of annual Spurges within *Euphorbia* Subg. *Esula*.**

In: Molecular Phylogenetics and Evolution 61 (2), p. 413–424. DOI: 10.1016/j.ympev.2011.06.011.

**Gillespie, Lynn J. (2007):**

**A Revision of Paleotropical *Plukenetia* (Euphorbiaceae) including two new Species from Madagascar.**

In: Systematic Botany 32 (4), p. 780–802.

**Halford, David A.; Henderson, Rodney J.F. (2005):**

***Shonia* R.J.F.Hend. & Halford (Ricinocarpeae, Ricinocarpinae), a new Australian endemic Genus.**

In: Austrobaileya 7 (1), p. 215–228.

**Hand, Ralf; Hadjikyriakou, Georgios N.; Christodoulou, Charalambos S.; Frajman, Božo (2015):**

**Multiple Origins of dendroid shrubs in the eastern Mediterranean *Euphorbia hierosolymitana* Group (Euphorbiaceae) with Description of a new Species, *Euphorbia lemesiana*, from Cyprus.**

In: Botanical Journal of the Linnean Society 179 (2), p. 295–307. DOI: 10.1111/boj.12319.

**Hoffmann, Petra; McPherson, Gordon (2007):**

**Revision of *Wielandia*, Including *Blotia* and *Petalodiscus* (Phyllanthaceae; Euphorbiaceae S.l.).**

In: Annals of the Missouri Botanical Garden 94 (3), p. 519–553. DOI: 10.3417/0026-6493(2007)94[519:ROWIBA]2.0.CO;2.

**Hoffmann, Petra; Wurdack, Kenneth J.J. (2006):**

***Radcliffea*, a new Genus of Euphorbiaceae sensu stricto from Madagascar.**

In: Kew Bulletin 61, p. 193–197.

**Horn, James W.; van Ee, Benjamin W.; Morawetz, Jeffery J.; Riina, Ricarda; Steinmann, Victor W.; Berry, Paul E.; Wurdack, Kenneth J.J. (2012):**

**Phylogenetics and the Evolution of major structural Characters in the giant Genus *Euphorbia* L. (Euphorbiaceae).**

In: Molecular Phylogenetics and Evolution 63 (2), p. 305–326. DOI: 10.1016/j.ympev.2011.12.022.

**Jestrow, Brett; Rodríguez, Francisco Jiménez; Francisco-Ortega, Javier (2010):**

**Generic Delimitation in the Antillean Adelieae (Euphorbiaceae) with Description of the Hispaniolan endemic Genus *Garciadelia*.**
In: Taxon 59 (6), p. 1801–1814. DOI: 10.1002/tax.596012.

JORDAL, BJARTE H.; HEWITT, GODFREY M. (2004):
**The Origin and Radiation of Macaronesian Beetles breeding in *Euphorbia*: the relative Importance of multiple Data Partitions and Population Sampling.**
In: Systematic Biology 53 (5), p. 711–734. DOI: 10.1080/10635150490468710.

KAWAKITA, ATSUSHI; TAKIMURA, ATSUSHI; TERACHI, TORU; SOTA, TEIJI; KATO, MAKOTO (2004):
**Cospeciation Analysis of an obligate Pollination Mutualism: Have Glochidion Trees (Euphorbiaceae) and Pollinating *Epicephala* Moths (Gracillariidae) diversified in parallel?**
In: Evolution 58 (10), p. 2201–2214. DOI: 10.1554/04-187.

KULJU, KRISTO K.M.; HAM, RAYMOND W.J.M. VAN DER; BRETELER, FRANS J. (2008):
**Rediscovery and phylogenetic Position of the incertae sedis Genus *Afrotrewia* (Euphorbiaceae): morphological, Pollen and molecular Evidence.**
In: Taxon 57 (1), p. 137–143. DOI: 10.6027/9789289333788-4-en.

KULJU, KRISTO K.M.; SIERRA, SORAYA E.C.; DRAISMA, STEFANO G. A.; SAMUEL, ROSABELLE; VAN WELZEN, PETER C. (2007):
**Molecular Phylogeny of *Macaranga, Mallotus*, and related Genera (Euphorbiaceae s.s.): Insights from Plastid and nuclear DNA Sequence Data.**
In: American Journal of Botany 94 (10), p. 1726–1743. DOI: 10.3732/ajb.94.10.1726.

MELO, ANDRÉ LAURÉNIO; ESSER, HANS-JOACHIM; SALES, MARGARETH FERREIRA (2013):
**New Combinations in *Pleradenophora* (Euphorbiaceae s.s.).**
In: Phytotaxa 81 (1). DOI: 10.11646/phytotaxa.81.1.10.

OBBARD, DARREN J.; HARRIS, STEPHEN A.; BUGGS, RICHARD J. A.; PANNELL, JOHN R. (2006):
**Hybridization, Polyploidy, and the Evolution of Sexual Systems in *Mercurialis* (Euphorbiaceae).**
In: Evolution 60 (9), p. 1801–1815. DOI: 10.1554/06-104.1.

PEIRSON, JESS A.; BRUYNS, PETER V.; RIINA, RICARDA; MORAWETZ, JEFFERY J.; BERRY, PAUL E. (2013):
**A molecular Phylogeny and Classification of the largely succulent and mainly African *Euphorbia* Subg. *Athymalus* (Euphorbiaceae).**
In: Taxon 62 (6), p. 1178–1199. DOI: 10.12705/626.12.

PEIRSON, JESS A.; RIINA, RICARDA; MAYFIELD, MARK H.; FERGUSON, CAROLYN J.; URBATSCH, LOWELL E.; BERRY, PAUL E. (2014):
**Phylogenetics and Taxonomy of the New World leafy Spurges, *Euphorbia Section Tithymalus* (Euphorbiaceae).**
In: Botanical Journal of the Linnean Society 175 (2), p. 191–228. DOI: 10.1111/boj.12167.

RIINA, RICARDA; BERRY, PAUL E.; VAN EE, BENJAMIN W. (2009):
**Molecular Phylogenetics of the Dragon's Blood *Croton* Section *Cyclostigma* (Euphorbiaceae): A polyphyletic Assemblage unraveled.**
In: Systematic Botany 34 (2), p. 360–374. DOI: 10.1600/036364409788606415.

RIINA, RICARDA; PEIRSON, JESS A.; GELTMAN, DMITRY V.; MOLERO, JULIÁN; FRAJMAN, BOŽO; PAHLEVANI, AMIRHOSSEIN ET AL. (2013):

**A worldwide molecular Phylogeny and Classification of the leafy Spurges, *Euphorbia* Subgenus *Esula* (Euphorbiaceae).**
In: Taxon 62 (2), p. 316–342. DOI: 10.12705/622.3.

RITZ, CHRISTINE M.; ZIMMERMANN, N. F. A.; HELLWIG, FRANK H. (2003):
**Phylogeny of subSect. *Meleuphorbia* (A. Berger) Pax & Hoffm. (*Euphorbia* L.) reflects the climatic regime in South Africa.**
In: Plant Systematics and Evolution 241 (3-4), p. 245–259. DOI: 10.1007/s00606-003-0064-4.

SIERRA, SORAYA E.C.; KULJU, KRISTO K.M.; FIŠER, ŽIVA; APARICIO, MARCELA; VAN WELZEN, PETER C. (2010):
**The Phylogeny of *Mallotus* s.str. (Euphorbiaceae) inferred from DNA Sequence and morphological Data.**
In: Taxon 59 (1), p. 101–116. DOI: 10.1002/tax.591011.

TOKUOKA, TORU (2007):
**Molecular phylogenetic Analysis of Euphorbiaceae sensu stricto based on Plastid and nuclear DNA Sequences and Ovule and Seed Character Evolution.**
In: Journal of Plant Research 120 (4), p. 511–522. DOI: 10.1007/s10265-007-0090-3.

TURNER, BILLIE L. (2011):
**Taxonomy of the papillose *Euphorbia [Tithymalus] longecornuta* complex (Euphorbiaceae) of northern Mexico.**
In: Phytoneuron 14, p. 1–7.

VAN EE, BENJAMIN W.; BERRY, PAUL E. (2009):
**A phylogenetic and taxonomic Review of *Croton* (Euphorbiaceae s.s.) on Jamaica including the Description of *Croton jamaicensis*, a new Species of Section *Eluteria*.**
In: Systematic Botany 34 (1), p. 129–140. DOI: 10.1600/036364409787602203.

VAN EE, BENJAMIN W.; BERRY, PAUL E. (2010):
**Taxonomy and Phylogeny of *Croton* Section *Heptallon* (Euphorbiaceae).**
In: Systematic Botany 35 (1), p. 151–167. DOI: 10.1600/036364410790862461.

VAN EE, BENJAMIN W.; BERRY, PAUL E. (2011):
***Croton* Section *Pedicellati* (Euphorbiaceae), a novel New World Group, and a new Subsectional Classification of *Croton* Section *Lamprocroton*.**
In: Systematic Botany 36 (1), p. 88–98. DOI: 10.1600/036364411X553162.

VAN EE, BENJAMIN W.; FORSTER, PAUL I.; BERRY, PAUL E. (2015):
**Phylogenetic Relationships and a new Sectional Classification of *Croton* (Euphorbiaceae) in Australia.**
In: Australian Systematic Botany 28 (4), p. 219–233. DOI: 10.1071/SB15016.

VAN EE, BENJAMIN W.; RIINA, RICARDA; BERRY, PAUL E. (2011):
**A revised infrageneric Classification and molecular Phylogeny of New World *Croton* (Euphorbiaceae).**
In: Taxon 60 (3), p. 791–823. DOI: 10.1002/tax.603013.

VAN WELZEN, PETER C. (2010):
**Revision of the Asian Genus *Koilodepas* (Euphorbiaceae).**
In: Annals of the Missouri Botanical Garden 97 (2), p. 218–234. DOI: 10.3417/2007149.

WANG, SHUQIONG; CHEN, YUNYUN; YANG, YUCHEN; WU, WEI; LIU, YING; FAN, QIANG; ZHOU, REN-CHAO (2016):
**Phylogenetic Relationships and natural Hybridization in *Triadica* inferred from nuclear and Chloroplast DNA Analyses.**

In: Biochemical Systematics and Ecology 64, p. 142–148. DOI: 10.1016/j.bse.2015.11.011.

**Wurdack, Kenneth J.J.; Hoffmann, Petra; Chase, Mark W. (2005):**
**Molecular phylogenetic Analysis of uniovulate Euphorbiaceae (Euphorbiaceae sensu stricto) using Plastid RBCL and Trnl-f DNA Sequences.**
In: American Journal of Botany 92 (8), p. 1397–1420. DOI: 10.3732/ajb.92.8.1397.

**Yang, Ya; Berry, Paul E. (2011):**
**Phylogenetics of the *Chamaesyce* Clade (Euphorbia, Euphorbiaceae): reticulate Evolution and long-distance Dispersal in a prominent C4 Lineage.**
In: American Journal of Botany 98 (9), p. 1486–1503. DOI: 10.3732/ajb.1000496.

**Yang, Ya; Riina, Ricarda; Morawetz, Jeffery J.; Haevermans, Thomas; Aubriot, Xavier; Berry, Paul E. (2012):**
**Molecular Phylogenetics and Classification of *Euphorbia* Subgenus *Chamaesyce* (Euphorbiaceae).**
In: Taxon 61 (4), p. 764–789. DOI: 10.1002/tax.614005.

**Zapata, Felipe; Fine, Paul V. A. (2013):**
**Diversification of the Monoterpene Synthase Gene Family (TPSb) in *Protium*, a highly diverse Genus of tropical Trees.**
In: Molecular Phylogenetics and Evolution 68 (3), p. 432–442. DOI: 10.1016/j.ympev.2013.04.024.

**Zimmermann, N. F. A.; Ritz, Christiane M.; Hellwig, Frank H. (2010):**
**Further support for the phylogenetic Relationships within *Euphorbia* L. (Euphorbiaceae) from nrITS and trnL–trnF Igs Sequence Data.**
In: Plant Systematics and Evolution 286 (1-2), p. 39–58. DOI: 10.1007/s00606-010-0272-7.

### Fabaceae

**Achimón, Fernanda; Johnson, Leigh A.; Cocucci, Andrea A.; Sérsic, Alicia N.; Baranzelli, Matias C. (2018):**
**Species Tree Phylogeny, Character Evolution, and Biogeography of the Patagonian Genus *Anarthrophyllum* Benth. (Fabaceae).**
In: Organisms Diversity and Evolution 18 (1), p. 71–86. DOI: 10.1007/s13127-017-0355-1.

**Aïnouche, Abdelkader; Bayer, Randall J. (1999):**
**Phylogenetic Relationships in *Lupinus* (Fabaceae: Papilionoideae) based on Internal Transcribed Spacer Sequences (ITS) of nuclear ribosomal DNA.**
In: American Journal of Botany 86 (4), p. 590–607. DOI: 10.2307/2656820.

**Allan, Gerard J.; Francisco-Ortega, Javier; Santos-Guerra, Arnoldo; Boerner, Erin; Zimmer, Elizabeth A. (2004):**
**Molecular phylogenetic Evidence for the geographic Origin and Classification of Canary Island *Lotus* (Fabaceae: Loteae).**
In: Molecular Phylogenetics and Evolution 32 (1), p. 123–138. DOI: 10.1016/j.ympev.2003.11.018.

**Allan, Gerard J.; Porter, J. Mark (2000):**
**Tribal Delimitation and phylogenetic Relationships of Loteae and Coronilleae (Faboideae: Fabaceae) with special Reference to *Lotus*: Evidence from nuclear ribosomal ITS Sequences.**
In: American Journal of Botany 87 (12), p. 1871–1881. DOI: 10.2307/2656839.

**Amirahmadi, Atefe; Kazempour-Osaloo, Shahrokh; Kaveh, Akram; Maassoumi, Ali Aasghar; Naderi, Reza (2016):**
**The Phylogeny and new Classification of the Genus *Onobrychis* (Fabaceae-Hedysareae): Evidence from molecular Data.**

In: Plant Systematics and Evolution 302 (10), p. 1445–1456. DOI: 10.1007/s00606-016-1343-1.

**Amirahmadi, Atefe; Kazempour-Osaloo, Shahrokh; Moein, Fatemeh; Kaveh, Akram; Maassoumi, Ali Aasghar (2014):**
**Molecular Systematics of the Tribe Hedysareae (Fabaceae) based on nrDNA ITS and Plastid trnL-F and matK Sequences.**
In: Plant Systematics and Evolution 300 (4), p. 729–747. DOI: 10.1007/s00606-013-0916-5.

**Anouche, A.; Bayer, Randall J.; Misset, Marie-Thérèse (2004):**
**Molecular Phylogeny, Diversification and Character Evolution in *Lupinus* (Fabaceae) with special Attention to Mediterranean and African Lupines.**
In: Plant Systematics and Evolution 246 (3-4), p. 211–222. DOI: 10.1007/s00606-004-0149-8.

**Arambarri, A. (2000):**
**A Cladistic Analysis of the New World Species of *Lotus* L. (Fabaceae, Loteae).**
In: Cladistics 16 (3), p. 283–297. DOI: 10.1006/clad.2000.0133.

**Asmussen-Lange, Conny B.; Liston, Aaron (1998):**
**Chloroplast DNA Characters, Phylogeny, and Classification of *Lathyrus* (Fabaceae).**
In: American Journal of Botany 85 (3), p. 387–401. DOI: 10.2307/2446332.

**Bagheri, Ali; Maassoumi, Ali Aasghar; Ghahremaninejad, Farrokh (2011):**
**A taxonomic Revision of the Genus *Astragalus* L. (Fabaceae) in Zanjan Province, Iran and describing a new Species.**
In: Taxonomy and Biosystematics 3 (8), p. 7–16.

**Banks, Hannah I.; Forest, Félix; Lewis, Gwilym Peter (2014):**
**Evolution and Diversity of Pollen Morphology in Tribe Cercideae (Leguminosae).**
In: Taxon 63 (2), p. 299–314. DOI: 10.12705/632.37.

**Bartha, László; Dragoş, Nicolae; Molnár V., Attila; Sramkó, Gábor (2013):**
**Molecular Evidence for reticulate Speciation in *Astragalus* (Fabaceae) as revealed by a Case Study from Sect. *Dissitiflori*.**
In: Botany 91 (10), p. 702–714. DOI: 10.1139/cjb-2013-0036.

**Bena, G.; Prosperi, J. M.; Lejeune, Bernard; Olivieri, Isabelle (1998):**
**Evolution of annual Species of the Genus *Medicago*: a molecular phylogenetic Approach.**
In: Molecular Phylogenetics and Evolution 9 (3), p. 552–559. DOI: 10.1006/mpev.1998.0493.

**Bessega, C.; Fortunato, René H. (2011):**
**Section *Mimadenia*: its phylogenetic Relationships within the Genus *Mimosa* (Leguminosae, Mimosoideae) using Plastid trnL-F Sequence Data.**
In: Australian Systematic Botany 24 (2), p. 104–110. DOI: 10.1071/SB10022.

**Bessega, C.; Hopp, H. E.; Fortunato, René H. (2008):**
**Toward a Phylogeny of *Mimosa* (Leguminosae: Mimosoidae): A preliminary Analysis of Southern South American Species based on Chloroplast DNA Sequence.**
In: Annals of the Missouri Botanical Garden 95 (4), p. 567–579. DOI: 10.3417/2006012.

**Boatwright, J. S.; Tilney, Patricia M.; van Wyk, Ben-Erik (2008):**
**A taxonomic Revision of the Genus *Rothia* (Crotalarieae, Fabaceae).**
In: Australian Systematic Botany 21 (6), p. 422–430. DOI: 10.1071/SB08033.

**Boatwright, James S.; Le Roux, Marianne M.; Wink, Michael; Morozova, Tatjana; van Wyk, Ben-Erik (2008):**

**Phylogenetic Relationships of Tribe Crotalarieae (Fabaceae) inferred from DNA Sequences and Morphology.**
In: Systematic Botany 33 (4), p. 752–761. DOI: 10.1600/036364408786500271.

**Boatwright, James S.; Maurin, Olivier; van der Bank, Michelle (2015):**
**Phylogenetic Position of Madagascan Species of *Acacia* s.l. and new Combinations in *Senegalia* and *Vachellia* (Fabaceae, Mimosoideae, Acacieae).**
In: Botanical Journal of the Linnean Society 179 (2), p. 288–294. DOI: 10.1111/boj.12320.

**Boatwright, James S.; Savolainen, Vincent; van Wyk, Ben-Erik; Lise Schutte-Vlok, Anne; Forest, Félix; van der Bank, Michelle (2008):**
**Systematic Position of the Anomalous Genus *Cadia* and the Phylogeny of the Tribe Podalyrieae (Fabaceae).**
In: Systematic Botany 33 (1), p. 133–147. DOI: 10.1600/036364408783887500.

**Boatwright, James S.; Wink, Michael; van Wyk, Ben-Erik (2011):**
**The generic Concept of *Lotononis* (Crotalarieae, Fabaceae): Reinstatement of the Genera *Euchlora, Leobordea* and *Listia* and the new Genus *Ezoloba*.**
In: Taxon 60 (1), p. 161–177. DOI: 10.1002/tax.601014.

**Bogdanović, Sandro; Brullo, Cristian; Brullo, Salvatore; Ljubičić, Ivica; Giusso del Galdo, Gianpietro (2016):**
***Bituminaria plumosa* (Fabaceae), a critical Species of the Croatian Flora.**
In: Plant Ecology and Evolution 149 (3), p. 347–355. DOI: 10.5091/plecevo.2016.1211.

**Borges, Leonardo Maurici; Simon, Marcelo Fragomeni; Pirani, José Rubens (2017):**
**Less is more. Adjusting the Taxonomy of the polytypic *Mimosa setosa* (Leguminosae, Mimosoid).**
In: Rodriguésia 68 (2), p. 515–540. DOI: 10.1590/2175-7860201768215.

**Bouchenak-Khelladi, Yanis; Maurin, Olivier; Hurter, Johan; van der Bank, Michelle (2010):**
**The evolutionary History and Biogeography of Mimosoideae (Leguminosae): an Emphasis on African Acacias.**
In: Molecular Phylogenetics and Evolution 57 (2), p. 495–508. DOI: 10.1016/j.ympev.2010.07.019.

**Brouat, Carine; Gielly, Ludovic; McKey, Doyle (2001):**
**Phylogenetic Relationships in the Genus *Leonardoxa* (Leguminosae: Caesalpinioideae) inferred from Chloroplast trnL Intron and trnL-trnF intergenic spacer Sequences.**
In: American Journal of Botany 88 (1), p. 143–149. DOI: 10.2307/2657134.

**Brown, Anthony H.D.; Grace, J. P.; Doyle, J. J. (2002):**
**Molecular phylogenetic Relationships within and among diploid Races of *Glycine tomentella* (Leguminosae).**
In: Australian Systematic Botany 15 (1), p. 37–47. DOI: 10.1071/SB01003.

**Brown, Gillian K.; Ariati, Siti R.; Murphy, Daniel J.; Miller, Joseph T.H.; Ladiges, Pauline Y. (2006):**
**Bipinnate Acacias (*Acacia* Subg. *Phyllodineae* Sect. *Botrycephalae*) of eastern Australia are polyphyletic based on DNA Sequence Data.**
In: Australian Systematic Botany 19 (4), p. 315–326. DOI: 10.1071/SB05039.

**Brown, Gillian K.; Clowes, Catherine; Murphy, Daniel J.; Ladiges, Pauline Y. (2010):**

**Phylogenetic Analysis based on nuclear DNA and Morphology defines a Clade of eastern Australian Species of *Acacia* s.s. (section *Juliflorae*): the '*Acacia longifolia* Group'.**

In: Australian Systematic Botany 23 (3), p. 162–172. DOI: 10.1071/SB09037.

BROWN, GILLIAN K.; MURPHY, DANIEL J.; KIDMAN, JAMES; LADIGES, PAULINE Y. (2012):

**Phylogenetic connections of phyllodinous Species of *Acacia* outside Australia are explained by geological History and human-mediated Dispersal.**

In: Australian Systematic Botany 25 (6), p. 390–403. DOI: 10.1071/SB12027.

BROWN, GILLIAN K.; MURPHY, DANIEL J.; LADIGES, PAULINE Y. (2011):

**Relationships of the Australo-Malesian Genus *Paraserianthes* (Mimosoideae: Leguminosae) identifies the Sister Group of *Acacia* sensu stricto and two biogeographical Tracks.**

In: Cladistics 27 (4), p. 380–390. DOI: 10.1111/j.1096-0031.2011.00349.x.

BROWN, GILLIAN K.; MURPHY, DANIEL J.; MILLER, JOSEPH T.H.; LADIGES, PAULINE Y. (2008):

**Acacia s.s. and its Relationship among tropical Legumes, Tribe Ingeae (Leguminosae: Mimosoideae).**

In: Systematic Botany 33 (4), p. 739–751. DOI: 10.1600/036364408786500136.

BRULLO, SALVATORE; BRULLO, CRISTIAN; MINISSALE, PIETRO; SALMERI, CRISTINA; GALDO, GIANPIETRO GIUSSO DEL (2016):

**Taxonomic Investigations on *Psoralea palaestina* (Fabaceae), a critical Species of Mediterranean Flora.**

In: Phytotaxa 266 (2), p. 61. DOI: 10.11646/phytotaxa.266.2.1.

BRUNEAU, ANNE; FOREST, FÉLIX; HERENDEEN, PATRICK S.; KLITGAARD, BENTE B.; LEWIS, GWILYM PETER (2001):

**Phylogenetic Relationships in the Caesalpinioideae (Leguminosae) as inferred from Chloroplast trnL Intron Sequences.**

In: Systematic Botany 26 (3), p. 487–514.

BRUNEAU, ANNE; KLITGAARD, BENTE B.; PRENNER, GERHARD; FOUGÈRE-DANEZAN, MARIE; TUCKER, SHIRLEY C. (2014):

**Floral Evolution in the Detarieae (Leguminosae): phylogenetic Evidence for labile Floral Development in an early-diverging Legume Lineage.**

In: International Journal of Plant Sciences 175 (4), p. 392–417. DOI: 10.1086/675574.

BRUNEAU, ANNE; MERCURE, MARJORIE; LEWIS, GWILYM PETER; HERENDEEN, PATRICK P. (2008):

**Phylogenetic Patterns and Diversification in the Caesalpinioid Legumes.**

In: Botany 86 (7), p. 697–718. DOI: 10.1139/B08-058.

BURGHARDT, ALICIA D.; ESPERT, SHIRLEY M. (2007):

**Phylogeny of *Prosopis* (Leguminosae) as shown by morphological and biochemical Evidence.**

In: Australian Systematic Botany 20 (4), p. 332–339. DOI: 10.1071/SB06043.

CAMARGO, RODRIGO AUGUSTO; AZEVEDO TOZZI, ANA MARIA GOULART (2014):

**A Synopsis of the Genus *Deguelia* (Leguminosae, Papilionoideae, Millettieae) in Brazil.**

In: Brittonia 66 (1), p. 12–32. DOI: 10.1007/s12228-013-9302-4.

CAMARGO, RODRIGO AUGUSTO; TOZZI, ANA MARIA GOULART DE AZEVEDO (2016):

**Taxonomic Placement of *Millettia occidentalis* (Leguminosae, Papilionoideae), a rare Liana from the Amazon Basin.**

In: Phytotaxa 261 (1), p. 75. DOI: 10.11646/phytotaxa.261.1.3.

**Cardoso, Domingos; Harris, David J.; Wieringa, Jan J.; São-Mateus, Wallace M. B.; Batalha-Filho, Henrique; Torke, Benjamin M. et al. (2017):**

**A molecular-dated Phylogeny and Biogeography of the monotypic Legume Genus *Haplormosia*, a missing African branch of the otherwise American-Australian Brongniartieae Clade.**

In: Molecular Phylogenetics and Evolution 107, p. 431–442. DOI: 10.1016/j.ympev.2016.12.012.

**Cardoso, Domingos; Lima, Haroldo Cavalcante de; Rodrigues, Rodrigo Schütz; Queiroz, Luciano Paganucci; Pennington, R. Toby; Lavin, Matt (2012):**

**The *Bowdichia* Clade of Genistoid Legumes: phylogenetic Analysis of combined molecular and morphological Data and a recircumscription of *Diplotropis*.**

In: Taxon 61 (5), p. 1074–1087.

**Cardoso, Domingos; Paganucci de Queiroz, Luciano; Cavalcante de Lima, Haroldo; Suganuma, Elisa; van den Berg, Cássio; Lavin, Matt (2013):**

**A molecular Phylogeny of the vataireoid Legumes underscores Floral evolvability that is General to many early-branching Papilionoid Lineages.**

In: American Journal of Botany 100 (2), p. 403–421. DOI: 10.3732/ajb.1200276.

**Cardoso, Domingos; Queiroz, Luciano Paganucci; Pennington, R. Toby; Lima, Haroldo Cavalcante de; Fonty, Emile; Wojciechowski, Martin F.; Lavin, Matt (2012):**

**Revisiting the Phylogeny of papilionoid Legumes: new Insights from comprehensively sampled early-branching Lineages.**

In: American Journal of Botany 99 (12), p. 1991–2013. DOI: 10.3732/ajb.1200380.

**Cardoso, Domingos; São-Mateus, Wallace M. B.; da Cruz, Daiane Trabuco; Zartman, Charles E.; Komura, Dirce L.; Kite, Geoffrey C. et al. (2015):**

**Filling in the gaps of the papilionoid Legume Phylogeny: the enigmatic Amazonian Genus *Petaladenium* is a new Branch of the early-diverging Amburaneae Clade.**

In: Molecular Phylogenetics and Evolution 84, p. 112–124. DOI: 10.1016/j.ympev.2014.12.015.

**Casimiro-Soriguer, Ramón; Talavera, María; Balao, Francisco; Terrab, Anass; Herrera, Javier; Talavera, Salvador (2010):**

**Phylogeny and genetic Structure of *Erophaca* (Leguminosae), a East-West Mediterranean disjunct Genus from the Tertiary.**

In: Molecular Phylogenetics and Evolution 56 (1), p. 441–450. DOI: 10.1016/j.ympev.2010.02.025.

**Castro, Olga; Cozzolino, Salvatore; Jury, p. L.; Caputo, P. (2002):**

**Molecular Relationships in *Genista* L. Sect. *Spartocarpus* Spach (Fabaceae).**

In: Plant Systematics and Evolution 231 (1-4), p. 91–108. DOI: 10.1007/s006060200013.

**Ceolin, Guilherme B.; Miotto, Silvia T. p. (2012):**

**Combining ecological and morphometrical Approaches to increase the Resolution within the *Galactia neesii* (Leguminosae) Complex.**

In: Plant Systematics and Evolution 298 (3), p. 645–652. DOI: 10.1007/s00606-011-0573-5.

**Chandler, Gregory T.; Bayer, Randall J.; Crisp, Michael D. (2001):**

**A molecular Phylogeny of the endemic Australian Genus *Gastrolobium* (Fabaceae: Mirbelieae) and allied Genera using Chloroplast and nuclear Markers.**

In: American Journal of Botany 88 (9), p. 1675–1687. DOI: 10.2307/3558413.

**Chandler, Gregory T.; Bayer, Randall J.; Gilmore, Simon R. (2003):**

**_Oxylobium/Gastrolobium_ (Fabaceae : Mirbelieae) Conundrum: further Studies using molecular Data and a reappraisal of morphological Characters.**
In: Plant Species Biology 18 (2-3), p. 91–101. DOI: 10.1111/j.1442-1984.2003.00094.x.

**Chappill, Jennifer A.; Wilkins, Carolyn F.; Crisp, Michael D. (2008):**
**Taxonomic Revision of _Gompholobium_ (Leguminosae: Mirbelieae).**
In: Australian Systematic Botany 21 (2), p. 67–151. DOI: 10.1071/SB07030.

**Chappill, Jennifer A.; Wilkins, Carolyn F.; Crisp, Michael D. (2007):**
**Taxonomic Revision of _Jacksonia_ (Leguminosae: Mirbelieae).**
In: Australian Systematic Botany 20 (6), p. 473–623. DOI: 10.1071/SB06047.

**Chaudhary, Lal Babu; Rana, Tikam Singh; Anand, Kumar Kamal (2008):**
**Current Status of the Systematics of _Astragalus_ L. (Fabaceae) with special Reference to the Himalayan Species in India.**
In: Taiwania 55 (4), p. 338–355.

**Chennaoui-Kourda, Houda; Marghali, Sonia; Zitouna, Nadia; Trifi-Farah, Neila (2012):**
**Phylogenetic Relationships of Mediterranean _Hedysarea_ Species assessed by AFLP Markers.**
In: Plant Systematics and Evolution 298 (1), p. 51–58. DOI: 10.1007/s00606-011-0522-3.

**Choi, Byoung-Hee; Seok, Dong-Im; Endo, Yasuhiko; Ohashi, Hiroyoshi (2006):**
**Phylogenetic Significance of stylar Features in Genus _Vicia_ (Leguminosae): an Analysis with molecular Phylogeny.**
In: Journal of Plant Research 119 (5), p. 513–523. DOI: 10.1007/s10265-006-0015-6.

**Crisp, Michael D.; Appels, R.; Smith, F. M.; Keys, W. M. p. (1999):**
**Phylogenetic Evaluation of 5s ribosomal RNA Gene and Spacer in the _Callistachys_ Group (Fabaceae: Mirbelieae).**
In: Plant Systematics and Evolution 218 (1-2), p. 33–42. DOI: 10.1007/BF01087032.

**Crisp, Michael D.; Cayzer, Lindy; Chandler, Gregory T.; Cook, Lyn G. (2017):**
**A monograph of _Daviesia_ (Mirbelieae, Faboideae, Fabaceae).**
In: Phytotaxa 300 (1), p. 1. DOI: 10.11646/phytotaxa.300.1.1.

**Crisp, Michael D.; Cook, Lyn G. (2003):**
**Phylogeny and Evolution of anomalous Roots in _Daviesia_ (Fabaceae: Mirbelieae).**
In: International Journal of Plant Sciences 164 (4), p. 603–612.

**Cubas, Paloma; Pardo, Cristina; Tahiri, Hikmat (2002):**
**Molecular approach to the Phylogeny and Systematics of _Cytisus_ Leguminosae) and related Genera based on nucleotide Sequences of nrDNA (ITS Region) and cpDNA (trnL-trnF intergenic spacer).**
In: Plant Systematics and Evolution 233 (3), p. 223–242. DOI: 10.1007/s00606-002-0194-0.

**Cubas, Paloma; Pardo, Cristina; Tahiri, Hikmat (2006):**
**Morphological convergence or Lineage Sorting? The case of _Cytisus purgans_ auct. (Leguminosae).**
In: Taxon 55 (3), p. 695–704. DOI: 10.2307/25065645.

**Cubas, Paloma; Pardo, Cristina; Tahiri, Hikmat; Castroviejo, Santiago (2010):**
**Phylogeny and evolutionary Diversification of _Adenocarpus_ DC. (Leguminosae).**
In: Taxon 59 (3), p. 720–732.

**da Silva, Marcos José; Azevedo Tozzi, Ana Maria Goulart (2012):**

**Three new species, a new Combination, and a neotypification in *Dahlstedtia* (Leguminosae, Millettieae, Papilionoideae) from South America.**
In: Brittonia 64 (3), p. 268–276. DOI: 10.1007/s12228-012-9242-4.

**da Silva, Marcos José; Queiroz, Luciano Paganucci; Tozzi, Ana Maria Goulart de Azevedo; Lewis, Gwilym Peter; Sousa, Anete Pereira de (2012):**
**Phylogeny and Biogeography of *Lonchocarpus* sensu lato and its Allies in the Tribe Millettieae (Leguminosae, Papilionoideae).**
In: Taxon 61 (1), p. 93–108. DOI: 10.1002/tax.611007.

**Dastpak, Arezoo; Kazempour-Osaloo, Shahrokh; Maassoumi, Ali Aasghar; Amirahmadi, Atefe (2013):**
**Phylogenetic Analysis of *Astragalus* Sect. *Ammodendron* (Fabaceae) based on nrDNA ITS and two cpDNAs, psbA-trnH and trnT-trnY Sequences.**
In: Biochemical Systematics and Ecology 50, p. 459–466. DOI: 10.1016/j.bse.2013.06.008.

**Degtjareva, Galina V.; Kramina, Tatiana E.; Sokoloff, Dmitry D.; Samigullin, Tahir H.; Valiejo-Roman, Carmen M.; Antonov, A. p. (2006):**
**Phylogeny of the Genus *Lotus* (Leguminosae, Loteae): Evidence from nrITS Sequences and Morphology.**
In: Canadian Journal of Botany 84 (5), p. 813–830. DOI: 10.1139/B06-035.

**Degtjareva, Galina V.; Valiejo-Roman, Carmen M.; Samigullin, Tahir H.; Guara-Requena, Miguel; Sokoloff, Dmitry D. (2012):**
**Phylogenetics of *Anthyllis* (Leguminosae: Papilionoideae: Loteae): Partial Incongruence between nuclear and Plastid Markers, a long Branch Problem and Implications for morphological Evolution.**
In: Molecular Phylogenetics and Evolution 62 (2), p. 693–707. DOI: 10.1016/j.ympev.2011.11.010.

**Delgado-Salinas, Alfonso; Bibler, Ryan; Lavin, Matt (2006):**
**Phylogeny of the Genus *Phaseolus* (Leguminosae): A Recent Diversification in an ancient Landscape.**
In: Systematic Botany 31 (4), p. 779–791.

**Dizkirici, Ayten; Ekici, Murat; Kaya, Zeki (2014):**
**Comparative molecular Phylogenetics of *Astragalus* L. Sections from Turkey with New World *Astragalus* Species using nrDNA ITS Sequences.**
In: Plant Systematics and Evolution 300 (1), p. 163–175. DOI: 10.1007/s00606-013-0868-9.

**Donkpegan, Armel p. L.; Doucet, Jean-Louis; Migliore, Jérémy; Duminil, Jérôme; Daïnou, Kasso; Piñeiro, Rosalía et al. (2017):**
**Evolution in African tropical Trees displaying Ploidy-habitat Association: the Genus *Afzelia* (Leguminosae).**
In: Molecular Phylogenetics and Evolution 107, p. 270–281. DOI: 10.1016/j.ympev.2016.11.004.

**Drummond, Christopher p. (2008):**
**Diversification of *Lupinus* (Leguminosae) in the western New World: derived Evolution of perennial life History and Colonization of montane Habitats.**
In: Molecular Phylogenetics and Evolution 48 (2), p. 408–421. DOI: 10.1016/j.ympev.2008.03.009.

**Drummond, Christopher S.; Eastwood, Ruth J.; Miotto, Silvia T. S.; Hughes, Colin E. (2012):**
**Multiple continental Radiations and correlates of Diversification in *Lupinus* (Leguminosae): testing for Key Innovation with incomplete Taxon Sampling.**
In: Systematic Biology 61 (3), p. 443–460. DOI: 10.1093/sysbio/syr126.

**Duan, Lei; Wen, Jun; Yang, Xue; Liu, Pei-Liang; Arslan, Emine; Ertuğrul, Kuddisi; Chang, Zhao-Yang (2015):**

**Phylogeny of *Hedysarum* and Tribe Hedysareae (Leguminosae: Papilionoideae) inferred from Sequence Data of ITS, matK, trnL-F and psbA-trnH.**
In: Taxon 64 (1), p. 49–64. DOI: 10.12705/641.26.

DUAN, LEI; YANG, XUE; LIU, PEI-LIANG; JOHNSON, GABRIEL; WEN, JUN; CHANG, ZHAO-YANG (2016):
**A molecular Phylogeny of Caraganeae (Leguminosae, Papilionoideae) reveals Insights into new generic and infrageneric Delimitations.**
In: PhytoKeys (70), p. 111–137. DOI: 10.3897/phytokeys.70.9641.

DUMINIL, JÉRÔME; BROWN, RICHARD P.; EWÉDJÈ, EBEN-EZER B. K.; MARDULYN, PATRICK; DOUCET, JEAN-LOUIS; HARDY, OLIVIER J. (2013):
**Large-scale Pattern of genetic differentiation within African Rainforest Trees: Insights on the Roles of ecological Gradients and past Climate Changes on the Evolution of *Erythrophleum* spp (Fabaceae).**
In: BMC Evolutionary Biology 13, p. 195. DOI: 10.1186/1471-2148-13-195.

EDWARDS, DAWN; HAWKINS, JULIE A. (2007):
**Are Cape Floral Clades the same age? Contemporaneous Origins of two Lineages in the Genistoids s.l. (Fabaceae).**
In: Molecular Phylogenetics and Evolution 45 (3), p. 952–970. DOI: 10.1016/j.ympev.2007.09.014.

EDWARDS, DAWN; HORN, ANNE; TAYLOR, DIANE; SAVOLAINEN, VINCENT; HAWKINS, JULIE A. (2008):
**DNA barcoding of a large Genus, *Aspalathus* L. (Fabaceae).**
In: Taxon 57 (4), p. 1317–1327. DOI: 10.1002/tax.574021.

EGAN, ASHLEY N.; CRANDALL, KEITH A. (2008):
**Incorporating gaps as phylogenetic Characters across eight DNA Regions: Ramifications for North American Psoraleeae (Leguminosae).**
In: Molecular Phylogenetics and Evolution 46 (2), p. 532–546. DOI: 10.1016/j.ympev.2007.10.006.

EGAN, ASHLEY N.; VATANPARAST, MOHAMMAD; CAGLE, WILLIAM (2016):
**Parsing polyphyletic *Pueraria*: Delimiting distinct evolutionary Lineages through Phylogeny.**
In: Molecular Phylogenetics and Evolution 104, p. 44–59. DOI: 10.1016/j.ympev.2016.08.001.

EKICI, MURAT; AKAN, HASAN; AYTAÇ, ZEKI (2015):
**Taxonomic Revision of *Astragalus* L. Section *Onobrychoidei* DC. (Fabaceae) in Turkey.**
In: Turkish Journal of Botany 39, p. 708–745. DOI: 10.3906/bot-1405-41.

ELLISON, NICK W.; LISTON, AARON; STEINER, JEFFREY J.; WILLIAMS, WARREN M.; TAYLOR, NORMAN L. (2006):
**Molecular Phylogenetics of the clover Genus (*Trifolium* - Leguminosae).**
In: Molecular Phylogenetics and Evolution 39 (3), p. 688–705. DOI: 10.1016/j.ympev.2006.01.004.

ENDO, YASUHIKO; CHOI, BYOUNG-HEE; OHASHI, HIROYOSHI; DELGADO-SALINAS, ALFONSO (2008):
**Phylogenetic Relationships of New World *Vicia* (Leguminosae) inferred from nrDNA Internal Transcribed Spacer Sequences and Floral Characters.**
In: Systematic Botany 33 (2), p. 356–363. DOI: 10.1600/036364408784571536.

ERKUL, SEHER KARAMAN; AYTAÇ, ZEKI (2013):
**The Revision of the Genus *Oxytropis* (Leguminosae) in Turkey.**
In: Turkish Journal of Botany 37, p. 24–38.

ESPERT, SHIRLEY M.; DREWES, SUSANA I.; BURGHARDT, ALICIA D. (2007):
**Phylogeny of *Macroptilium* (Leguminosae): morphological, biochemical and molecular Evidence.**

In: Cladistics 23 (2), p. 119–129. DOI: 10.1111/j.1096-0031.2006.00140.x.

FORTUNA-PEREZ, ANA PAULA; DA SILVA, MARCOS JOSÉ; QUEIROZ, LUCIANO PAGANUCCI; LEWIS, GWILYM PETER; SIMÕES, ANDRÉ OLMOS; AZEVEDO TOZZI, ANA MARIA GOULART ET AL. (2013):
**Phylogeny and Biogeography of the Genus *Zornia* (Leguminosae: Papilionoideae: Dalbergieae).**
In: Taxon 62 (4), p. 723–732. DOI: 10.12705/624.35.

FOUGÈRE-DANEZAN, MARIE; HERENDEEN, PATRICK S.; MAUMONT, STÉPHAN; BRUNEAU, ANNE (2010):
**Morphological Evolution in the variable resin-producing Detarieae (Fabaceae): do morphological Characters retain a phylogenetic Signal?**
In: Annals of Botany 105 (2), p. 311–325. DOI: 10.1093/aob/mcp280.

FOUGÈRE-DANEZAN, MARIE; MAUMONT, STÉPHAN; BRUNEAU, ANNE (2007):
**Relationships among Resin-Producing Detarieae s.l. (Leguminosae) as inferred by molecular Data.**
In: Systematic Botany 32 (4), p. 748–761.

FRIEND, P. A.; QUANDT, DIETMAR; TALLURY, P. P.; STALKER, H. T.; HILU, KHIDIR W. (2010):
**Species, Genomes, and Section Relationships in the Genus *Arachis* (Fabaceae): a molecular Phylogeny.**
In: Plant Systematics and Evolution 290 (1-4), p. 185–199. DOI: 10.1007/s00606-010-0360-8.

FRITSCH, PETER W.; CRUZ, BONI C. (2012):
**Phylogeny of *Cercis* based on DNA Sequences of nuclear ITS and four Plastid Regions: Implications for transatlantic historical Biogeography.**
In: Molecular Phylogenetics and Evolution 62 (3), p. 816–825. DOI: 10.1016/j.ympev.2011.11.016.

GAGNON, EDELINE; HUGHES, COLIN E.; LEWIS, GWILYM PETER; BRUNEAU, ANNE (2015):
**A new cryptic Species in a new cryptic Genus in the *Caesalpinia* Group (Leguminosae) from the seasonally dry inter-Andean valleys of South America.**
In: Taxon 64 (3), p. 468–490. DOI: 10.12705/643.6.

GAGNON, EDELINE; LEWIS, GWILYM PETER; SOLANGE SOTUYO, J.; HUGHES, COLIN E.; BRUNEAU, ANNE (2013):
**A molecular Phylogeny of *Caesalpinia* sensu lato: Increased Sampling reveals new Insights and more Genera than expected.**
In: South African Journal of Botany 89, p. 111–127. DOI: 10.1016/j.sajb.2013.07.027.

GERVAIS, G. Y. F.; BRUNEAU, ANNE (2002):
**Phylogenetic Analysis of a polyphyletic African Genus of Caesalpinioideae (Leguminosae): *Monopetalanthus* Harms.**
In: Plant Systematics and Evolution 235 (1), p. 19–34. DOI: 10.1007/s00606-002-0222-0.

GOVINDARAJULU, RAJANIKANTH; HUGHES, COLIN E.; BAILEY, C. DONOVAN (2011):
**Phylogenetic and population genetic Analyses of diploid *Leucaena* (Leguminosae; Mimosoideae) reveal cryptic Species Diversity and Patterns of divergent allopatric speciation.**
In: American Journal of Botany 98 (12), p. 2049–2063. DOI: 10.3732/ajb.1100259.

GROUP, THE LEGUME PHYLOGENY WORKING (2013):
**Legume Phylogeny and Classification in the 21st century: Progress, Prospects and Lessons for other species-rich Clades.**
In: Taxon 62 (2), p. 217–248. DOI: 10.12705/622.8.

HAN, JEONG EUN; CHUNG, KOOK-HYUN; NEMOTO, TOMOYUKI; CHOI, BYOUNG-HEE (2010):

**Phylogenetic Analysis of eastern Asian and eastern North American disjunct *Lespedeza* (Fabaceae) inferred from nuclear ribosomal ITS and Plastid Region Sequences.**
In: Botanical Journal of the Linnean Society 164 (3), p. 221–235. DOI: 10.1111/j.1095-8339.2010.01084.x.

**Hardion, Laurent; Baumel, Alex; Dumas, Pierre-Jean; Duong, Nathalie; Affre, Laurence; Tatoni, Thierry (2010):**
**Phylogenetic Relationships and infrageneric Classification of *Astragalus tragacantha* L. (Fabaceae), inferred from nuclear ribosomal DNA Internal transcribed Spacers Data (nrDNA ITS).**
In: Ecologia Mediterranea 36 (1), p. 99–106.

**Haston, Elspeth M.; Lewis, Gwilym Peter; Hawkins, Julie A. (2005):**
**A phylogenetic reappraisal of the *Peltophorum* Group (Caesalpinieae: Leguminosae) based on the Chloroplast trnL-F, rbcL and rps16 Sequence Data.**
In: American Journal of Botany 92 (8), p. 1359–1371. DOI: 10.3732/ajb.92.8.1359.

**Hou, Xin; Liu, Jun-E; Zhao, Yi-Zhi (2008):**
**Molecular Phylogeny of *Caragana* (Fabaceae) in China.**
In: Journal of Systematics and Evolution 46 (4), p. 600–607.

**Hu, Jer-Ming; Lavin, Matt; Wojciechowski, Martin F.; Sanderson, Michael J. (2002):**
**Phylogenetic Analysis of Nuclear Ribosomal ITS/5.8S Sequences in the Tribe Millettieae (Fabaceae): *Poecilanthe-Cyclolobium*, the Core Millettieae, and the *Callerya* Group.**
In: Systematic Botany 27 (4), p. 722–733.

**Hu, Jer-Ming; Lavin, Matt; Wojciechowski, Martin F.; Sanderson, Michael J. (2000):**
**Phylogenetic Systematics of the Tribe Millettieae (Leguminosae) based on Chloroplast trnK/matK Sequences and its Implications for evolutionary Patterns in Papilionoideae.**
In: American Journal of Botany 87 (3), p. 418–430. DOI: 10.2307/2656638.

**Iganci, João R. V.; Soares, Marcos V.; Guerra, Ethiéne; Morim, Marli P. (2016):**
**A Preliminary molecular Phylogeny of the *Abarema* Alliance (Leguminosae) and Implications for taxonomic Rearrangement.**
In: International Journal of Plant Sciences 177 (1), p. 34–43. DOI: 10.1086/684078.

**Ireland, Helen E.; Kite, Geoffrey C.; Veitch, Nigel C.; Chase, Mark W.; Schrire, Brian D.; Lavin, Matt et al. (2010):**
**Biogeographical, ecological and morphological Structure in a phylogenetic Analysis of *Ateleia* (Swartzieae, Fabaceae) derived from combined molecular, morphological and chemical Data.**
In: Botanical Journal of the Linnean Society 162, p. 39–53.

**Jaaska, Vello (2005):**
**Isozyme Variation and phylogenetic Relationships in *Vicia* Subgenus *Cracca* (Fabaceae).**
In: Annals of Botany 96 (6), p. 1085–1096. DOI: 10.1093/aob/mci260.

**Jaaska, Vello (2015):**
**Phylogenetic Relationships among Sections *Vicia, Sepium* and *Lathyroides* of *Vicia* Subgenus *Vicia*: isozyme Evidence.**
In: Biochemical Systematics and Ecology 62, p. 186–193. DOI: 10.1016/j.bse.2015.08.002.

**Jalilian, N.; Rahiminejad, Mohammad Reza; Maassoumi, Ali Aasghar; Maroofi, Hosein (2014):**
**Taxonomic Revision of the Genus *Vicia* L. (Fabaceae) in Iran.**
In: Iranian Journal of Botany 20 (2), p. 155–164.

**Javadi, Firouzeh; Tun, Ye Tun; Kawase, Makoto; Guan, Kaiyun; Yamaguchi, Hirofumi (2011):**
**Molecular Phylogeny of the Subgenus *Ceratotropis* (genus Vigna, Leguminosae) reveals three eco-geographical Groups and Late Pliocene-Pleistocene Diversification: Evidence from four Plastid DNA Region Sequences.**
In: Annals of Botany 108 (2), p. 367–380. DOI: 10.1093/aob/mcr141.

**Javanmardi, Farahnaz; Kazempour-Osaloo, Shahrokh; Maassoumi, Ali Aasghar; Nejadsattari, Taher (2012):**
**Molecular Phylogeny of *Astragalus Section Alopecuroidei* (Fabaceae) and its Allies based on nrDNA ITS and three cpDNAs, matK, trnT-trnY and trnH-psbA Sequences.**
In: Biochemical Systematics and Ecology 45, p. 171–178. DOI: 10.1016/j.bse.2012.07.029.

**Jobson, Richard W.; Luckow, Melissa (2007):**
**Phylogenetic Study of the Genus *Piptadenia* (Mimosoideae: Leguminosae) using Plastid trnL-F and trnK / matK Sequence Data.**
In: Systematic Botany 32 (3), p. 569–575. DOI: 10.1600/036364407782250544.

**Junior, Marcus José de Azevedo Falcao; Pinto, Rafael Barbosa; Mansano, Vidal de Freitas (2016):**
**A taxonomic Revision of the Genus *Dialium* (Leguminosae: Dialiinae) in the Netotropics.**
In: Phytotaxa 283 (2), p. 123–142. DOI: 10.11646/phytotaxa.283.2.2.

**Käss, E.; Wink, Michael (1997):**
**Phylogenetic Relationships in the Papilionoideae (Family Leguminosae) based on nucleotide Sequences of cpDNA (rbcL) and ncDNA (ITS 1 and 2).**
In: Molecular Phylogenetics and Evolution 8 (1), p. 65–88. DOI: 10.1006/mpev.1997.0410.

**Kazemi, Marzie; Kazempour-Osaloo, Shahrokh; Asghar Maassoumi, Ali; Rastegar Pouyani, Eskandar (2009):**
**Molecular Phylogeny of selected Old World *Astragalus* (Fabaceae): incongruence among Chloroplast trnL-F, ndhF and nuclear ribosomal DNA ITS Sequences.**
In: Nordic Journal of Botany 27 (5), p. 425–436. DOI: 10.1111/j.1756-1051.2009.00285.x.

**Kramina, Tatiana E.; Degtjareva, Galina V.; Samigullin, Tahir H.; Valiejo-Roman, Carmen M.; Kirkbride, Joseph H.; Volis, Sergei et al. (2016):**
**Phylogeny of *Lotus* (Leguminosae: Loteae): Partial incongruence between nrITS, nrETS and Plastid Markers and biogeographic Implications.**
In: Taxon 65 (5), p. 997–1018. DOI: 10.12705/655.4.

**Kyalangalilwa, Bruce; Boatwright, James S.; Daru, Barnabas H.; Maurin, Olivier; van der Bank, Michelle (2013):**
**Phylogenetic Position and revised Classification of *Acacia* s.l. (Fabaceae: Mimosoideae) in Africa, including new Combinations in *Vachellia* and *Senegalia*.**
In: Botanical Journal of the Linnean Society 172 (4), p. 500–523. DOI: 10.1111/boj.12047.

**La Estrella, Manuel; Wieringa, Jan J.; Mackinder, Barbara; van der Burgt, Xander; Devesa, Juan A.; Bruneau, Anne (2014):**
**Phylogenetic Analysis of the African Genus *Gilbertiodendron* J. Léonard and related Genera (Leguminosae-Caesalpinioideae-Detarieae).**
In: International Journal of Plant Sciences 175 (9), p. 975–985. DOI: 10.1086/677648.

**Lavin, Matt; Pennington, R. Toby; Klitgaard, Bente B.; Sprent, Janet I.; Lima, Haroldo Cavalcante de; Gasson, Peter E. (2001):**
**The dalbergioid Legumes (Fabaceae): Delimitation of a pantropical monophyletic Clade.**
In: American Journal of Botany 88 (3), p. 503–533. DOI: 10.2307/2657116.

**Le Roux, Marianne M.; Boatwright, James S.; van Wyk, Ben-Erik (2013):**
**A global infrageneric Classification system for the Genus *Crotalaria* (Leguminosae) based on molecular and morphological Evidence.**
In: Taxon 62 (5), p. 957–971. DOI: 10.12705/625.1.

**Le Roux, Marianne M.; van Wyk, Ben-Erik; Boatwright, James S.; Tilney, Patricia M. (2011):**
**The systematic Significance of morphological and anatomical Variation in fruits of *Crotalaria* and related Genera of Tribe Crotalarieae (Fabaceae).**
In: Botanical Journal of the Linnean Society 165, p. 84–106.

**Lee, Jeongran; Hymowitz, Theodore (2001):**
**A molecular phylogenetic Study of the Subtribe Glycininae (Leguminosae) derived from the Chloroplast DNA rps 16 Intron Sequences.**
In: American Journal of Botany 88 (11), p. 2064–2073. DOI: 10.2307/3558432.

**Leht, Malle (2005):**
**Cladistic and phenetic Analyses of Relationships in *Vicia* Subgenus *Cracca* (Fabaceae) based on morphological Data.**
In: Taxon 54 (4), p. 1023–1032. DOI: 10.2307/25065486.

**Leht, Malle (2009):**
**Phylogenetics of *Vicia* (Fabaceae) based on morphological Data.**
In: Feddes Repertorium 120 (7-8), p. 379–393. DOI: 10.1002/fedr.200911117.

**Leht, Malle (2009):**
**Phylogeny of Old World *Lathyrus* L. (Fabaceae) based on morphological Data.**
In: Feddes Repertorium 120 (1-2), p. 59–74. DOI: 10.1002/fedr.200811182.

**Li, Jian-Hua; Jiang, Jin-Huo; Fu, Cheng-Xin; Tang, Shao-Qing (2014):**
**Molecular Systematics and Biogeography of *Wisteria* inferred from nucleotide Sequences of nuclear and Plastid Genes.**
In: Journal of Systematics and Evolution 52 (1), p. 40–50. DOI: 10.1111/jse.12061.

**Li, Jian-Hua; Jiang, Jin-Huo; Stel, Holly Vander; Homkes, Austin; Corajod, Jeffrey; Brown, Kenneth; Chen, Zhi-Duan (2014):**
**Phylogenetics and Biogeography of *Apios* (Fabaceae) inferred from Sequences of Nuclear and Plastid Genes.**
In: International Journal of Plant Sciences 175 (7), p. 764–780. DOI: 10.1086/676972.

**Lima, Laura Cristina Pires; Queiroz, Luciano Paganucci; Azevedon., Tozzi. Maria Goulart de; Lewis, Gwilym Peter (2014):**
**A taxonomic Revision of *Desmodium* (Leguminosae, Papilionoideae) in Brazil.**
In: Phytotaxa 169 (1), p. 1–119. DOI: 10.11646/phytotaxa.169.1.1.

**Luckow, Melissa; Fortunato, René H.; Sede, Silvana M.; Livshultz, Tatyana (2005):**
**The phylogenetic Affinities of two mysterious monotypic Mimosoids from Southern South America.**
In: Systematic Botany 30 (3), p. 585–602.

**Mahé, Frédéric; Markova, Dragomira N.; Pasquet, Rémy; Misset, Marie-Thérèse; Aïnouche, Abdelkader (2011):**
**Isolation, Phylogeny and Evolution of the SymRK Gene in the Legume Genus *Lupinus* L.**
In: Molecular Phylogenetics and Evolution 60 (1), p. 49–61. DOI: 10.1016/j.ympev.2011.04.017.

**Malaviya, D. R.; Roy, A. K.; Kaushal, P.; Kumar, B.; Tiwari, A. (2008):**
**Genetic similarity among *Trifolium* Species based on Isozyme Banding Pattern.**
In: Plant Systematics and Evolution 276 (1-2), p. 125–136. DOI: 10.1007/s00606-008-0070-7.

**Manzanilla, Vincent; Bruneau, Anne (2012):**
**Phylogeny Reconstruction in the Caesalpinieae grade (Leguminosae) based on duplicated copies of the sucrose synthase Gene and Plastid Markers.**
In: Molecular Phylogenetics and Evolution 65 (1), p. 149–162. DOI: 10.1016/j.ympev.2012.05.035.

**Marazzi, Brigitte; Endress, Peter K.; Queiroz, Luciano Paganucci; Conti, Elena (2006):**
**Phylogenetic Relationships within *Senna* (Leguminosae, Cassiinae) based on three Chloroplast DNA Regions: Patterns in the Evolution of Floral symmetry and extrafloral Nectaries.**
In: American Journal of Botany 93 (2), p. 288–303. DOI: 10.3732/ajb.93.2.288.

**Maroua, Gharbi; Nadia, Zitouna; Imen, Fadhlaoui; Neila, Trifi-Farah; Sonia, Marghali (2014):**
**Molecular characterization of *Lathyrus* Species using Chloroplast DNA trnH-psbA.**
In: Biochemical Systematics and Ecology 57, p. 439–444. DOI: 10.1016/j.bse.2014.09.002.

**Maureira-Butler, Iván J.; Pfeil, Bernard E.; Muangprom, Amorntip; Osborn, Thomas C.; Doyle, Jeff J. (2008):**
**The reticulate History of *Medicago* (Fabaceae).**
In: Systematic Biology 57 (3), p. 466–482. DOI: 10.1080/10635150802172168.

**Mayer, M. S.; Bagga, p. K. (2002):**
**The Phylogeny of *Lens* (Leguminosae): new Insight from ITS Sequence Analysis.**
In: Plant Systematics and Evolution 232 (3-4), p. 145–154. DOI: 10.1007/s006060200038.

**McMahon, Michelle; Hufford, Larry (2004):**
**Phylogeny of Amorpheae (Fabaceae: Papilionoideae).**
In: American Journal of Botany 91 (8), p. 1219–1230. DOI: 10.3732/ajb.91.8.1219.

**McMahon, Michelle; Hufford, Larry (2005):**
**Evolution and Development in the Amorphoid Clade (Amorpheae: Papilionoideae: Leguminosae): Petal Loss and Dedifferentiation.**
In: International Journal of Plant Sciences 166 (3), p. 383–396.

**Meireles, Jose Eduardo; Azevedo Tozzi, Ana Maria Goulart (2015):**
***Limadendron*: a new Genus of Leguminosae (Papilionoideae, Brongniartieae) from South America.**
In: Plant Systematics and Evolution 301 (2), p. 701–707. DOI: 10.1007/s00606-014-1108-7.

**Meng, Hong-Hu; Jacques, Frédéric M.B.; Su, Tao; Huang, Yong-Jiang; Zhang, Shi-Tao; Ma, Hong-Jie; Zhou, Zhe-Kun (2014):**
**New Biogeographic Insight into *Bauhinia* s.l. (Leguminosae): integration from fossil Records and molecular Analyses.**
In: BMC Evolutionary Biology 14, p. 181. DOI: 10.1186/s12862-014-0181-4.

**Miller, Joseph T.H.; Bayer, Randall J. (2001):**

**Molecular Phylogenetics of *Acacia* (Fabaceae: Mimosoideae) based on the Chloroplast Matk coding Sequence and Flanking Trnk Intron Spacer Regions.**
In: American Journal of Botany 88 (4), p. 697–705. DOI: 10.2307/2657071.

MILLER, JOSEPH T.H.; GRIMES, JAMES W.; MURPHY, DANIEL J.; BAYER, RANDALL J.; LADIGES, PAULINE Y. (2003):
**A phylogenetic Analysis of the Acacieae and Ingeae (Mimosoideae: Fabaceae) based on trnK, matK, psbA-trnH, and trnL/trnF Sequence Data.**
In: Systematic Botany 28 (3), p. 558–566.

MILLER, JOSEPH T.H.; SEIGLER, DAVID P. (2012):
**Evolutionary and taxonomic Relationships of *Acacia* s.l. (Leguminosae: Mimosoideae).**
In: Australian Systematic Botany 25 (3), p. 217–224. DOI: 10.1071/SB11042.

MOTEETEE, A.; VAN WYK, BEN-ERIK (2008):
**A Synopsis of the Middle-eastern and Asian Species of *Argyrolobium* (Genisteae–Fabaceae).**
In: South African Journal of Botany 74 (1), p. 10–24. DOI: 10.1016/j.sajb.2007.07.007.

MOURA, TANIA MARIA DE; LEWIS, GWILYM PETER (2014):
**Taxonomic studies in the *Mucuna poggei* complex (Leguminosae: Papilionoideae).**
In: Kew Bulletin 69 (4). DOI: 10.1007/S12225-014-9544-7.

MOURA, TANIA MARIA DE; VATANPARAST, MOHAMMAD; TOZZI, ANA MARIA GOULART DE AZEVEDO; FOREST, FÉLIX; WILMOT-DEAR, C. MELANIE; SIMON, MARCELO FRAGOMENI ET AL. (2016):
**A molecular Phylogeny and new Infrageneric Classification of *Mucuna* Adans. (Leguminosae-Papilionoideae) including Insights from Morphology and Hypotheses about Biogeography.**
In: International Journal of Plant Sciences 177 (1), p. 76–89. DOI: 10.1086/684131.

MOURA, TANIA MARIA DE; WILMOT-DEAR, C. MELANIE; VATANPARAST, MOHAMMAD; FORTUNA-PEREZ, ANA PAULA; TOZZI, ANA MARIA GOULART DE AZEVEDO; LEWIS, GWILYM PETER (2016):
**A new Infrageneric Classification of *Mucuna* (Leguminosae-Papilionoideae): Supported by Morphology, molecular Phylogeny and Biogeography.**
In: Systematic Botany 41 (3), p. 606–616. DOI: 10.1600/036364416X692532.

MURPHY, DANIEL J.; BROWN, GILLIAN K.; MILLER, JOSEPH T.H.; LADIGES, PAULINE Y. (2010):
**Molecular Phylogeny of *Acacia* Mill. (Mimosoideae: Leguminosae): Evidence for major Clades and informal Classification.**
In: Taxon 59 (1), p. 7–19. DOI: 10.1002/tax.591002.

MURPHY, DANIEL J.; UDOVICIC, FRANK; LADIGES, PAULINE Y. (2000):
**Phylogenetic Analysis of Australian *Acacia* (Leguminosae: Mimosoideae) by using Sequence Variations of an Intron and two intergenic Spacers of Chloroplast DNA.**
In: Australian Systematic Botany 13 (5), p. 745–754. DOI: 10.1071/SB99027.

NADERI SAFAR, KOSAR; KAZEMPOUR-OSALOO, SHAHROKH; MAASSOUMI, ALI AASGHAR; ZARRE, SHAHIN (2014):
**Molecular Phylogeny of *Astragalus* Section Anthylloidei (Fabaceae) inferred from nrDNA ITS and Plastid rpl32-trnL(UAG) Sequence Data.**
In: Turkish Journal of Botany 38, p. 637–652. DOI: 10.3906/bot-1308-44.

NADIA, ZITOUNA; MAROUA, GHARBI; HELA, BEN RHOUMA; HOUDA, CHENNAOUI-KOURDA; ABDELMAJID, HADDIOUI; NEILA, TRIFI-FARAH; SONIA, MARGHALI (2014):

**Evolutionary and demographic History among Maghrebian *Medicago* Species (Fabaceae) based on the nucleotide Sequences of the Chloroplast DNA barcode trnH-psbA.**
In: Biochemical Systematics and Ecology 55, p. 296–304. DOI: 10.1016/j.bse.2014.03.016.

NAGANOWSKA, B.; WOLKO, B.; ŚLIWIŃSKA, E.; KACZMAREK, Z.; SCHIFINO-WITTMANN, M. T. (2005):
**2c DNA Variation and Relationships among New World Species of the Genus *Lupinus* (Fabaceae).**
In: Plant Systematics and Evolution 256 (1-4), p. 147–157. DOI: 10.1007/s00606-005-0364-y.

NANNI, L.; FERRADINI, N.; TAFFETANI, F.; PAPA, R. (2004):
**Molecular Phylogeny of *Anthyllis* spp.**
In: Plant Biology 6 (4), p. 454–464. DOI: 10.1055/s-2004-820968.

NASSEH, Y.; JOHARCHI, MOHAMMAD REZA (2012):
**New Records and new Synonym of *Astragalus* Sections *Ammodendron* and *Carpini* (Fabaceae) from Iran.**
In: Iranian Journal of Botany 18 (1), p. 76–83.

NORES, MARÍA J.; SIMPSON, BERYL B.; HICK, PASCALE; ANTON, ANA M.; FORTUNATO, RENÉ H. (2012):
**The phylogenetic Relationships of four monospecific Caesalpinioids (Leguminosae) endemic to southern South America.**
In: Taxon 61 (4), p. 790–802.

OJEDA, ISIDRO; SANTOS-GUERRA, ARNOLDO; JAÉN-MOLINA, RUTH; OLIVA-TEJERA, FELICIA; CAUJAPÉ-CASTELLS, JULI; CRONK, QUENTIN C.B. (2012):
**The Origin of Bird Pollination in Macaronesian *Lotus* (Loteae, Leguminosae).**
In: Molecular Phylogenetics and Evolution 62 (1), p. 306–318. DOI: 10.1016/j.ympev.2011.10.001.

ORTEGA-OLIVENCIA, ANA; CATALÁN, PILAR (2009):
**Systematics and evolutionary History of the Circum-mediterranean Genus *Anagyris* L. (Fabaceae) based on morphological and molecular Data.**
In: Taxon 58 (4), p. 1290–1306. DOI: 10.1002/tax.584018.

ORTHIA, L. A.; COOK, LYN G.; CRISP, MICHAEL D. (2005):
**Generic Delimitation and phylogenetic uncertainty: an Example from a Group that has undergone an explosive Radiation.**
In: Australian Systematic Botany 18 (1), p. 41–47. DOI: 10.1071/SB04016.

OSALOO, SHAHROKH KAZEMPOUR; MAASSOUMI, ALI AASGHAR; MURAKAMI, NORIAKI (2003):
**Molecular Systematics of the Genus *Astragalus* L. (Fabaceae): phylogenetic *Analyses* of nuclear ribosomal DNA internal transcribed Spacers and Chloroplast Gene ndh F Sequences.**
In: Plant Systematics and Evolution 242 (1-4), p. 1–32. DOI: 10.1007/s00606-003-0014-1.

OSKOUEIYAN, ROGHAYEH; KAZEMPOUR-OSALOO, SHAHROKH; MAASSOUMI, ALI AASGHAR; NEJADSATTARI, TAHER; MOZAFFARIAN, VALIOLLAH (2010):
**Phylogenetic Status of *Vavilovia formosa* (Fabaceae-Fabeae) based on nrDNA ITS and cpDNA Sequences.**
In: Biochemical Systematics and Ecology 38 (3), p. 313–319. DOI: 10.1016/j.bse.2010.01.011.

ÖZTÜRK, MERYEM; DURAN, AHMET; HAKKI, ERDOGAN E. (2013):
**Cladistic and phylogenetic Analyses of the Genus *Cicer* in Turkey.**
In: Plant Systematics and Evolution 299 (10), p. 1955–1966. DOI: 10.1007/s00606-013-0850-6.

**Pardo, Cristina; Cubas, Paloma; Tahiri, Hikmat (2004):**
**Molecular Phylogeny and Systematics of *Genista* (Leguminosae) and related Genera based on nucleotide Sequences of nrDNA (ITS Region) and cpDNA (trnL-trnF intergenic spacer).**
In: Plant Systematics and Evolution 244 (1-2), p. 93–119. DOI: 10.1007/s00606-003-0091-1.

**Pooyan, P.; Ghahremaninejad, Farrokh; Assadi, Mostafa (2014):**
**A Synopsis of the Genus *Colutea* (Fabaceae) in Iran.**
In: Edinburgh Journal of Botany 71 (1), p. 35–49. DOI: 10.1017/S0960428613000280.

**Queiroz, Luciano Paganucci; Lavin, Matt (2011):**
***Coursetia* (Leguminosae) from Eastern Brazil: Nuclear Ribosomal and Chloroplast DNA Sequence Analysis reveal the Monophyly of three Caatinga-inhabiting Species.**
In: Systematic Botany 36 (1), p. 69–79. DOI: 10.1600/036364411X553144.

**Queiroz, Luciano Paganucci; Pastore, José Floriano B.; Cardoso, Domingos; Snak, Cristiane; Lima, Ana Luísa C.; Gagnon, Edeline et al. (2015):**
**A multilocus phylogenetic Analysis reveals the Monophyly of a recircumscribed papilionoid Legume Tribe Diocleae with well-supported generic Relationships.**
In: Molecular Phylogenetics and Evolution 90, p. 1–19. DOI: 10.1016/j.ympev.2015.04.016.

**Ramos, Gustavo; Lima, Haroldo Cavalcante de; Prenner, Gerhard; Queiroz, Luciano Paganucci; Zartman, Charles E.; Cardoso, Domingos (2016):**
**Molecular Systematics of the Amazonian Genus *Aldina*, a phylogenetically enigmatic ectomycorrhizal Lineage of papilionoid Legumes.**
In: Molecular Phylogenetics and Evolution 97, p. 11–18. DOI: 10.1016/j.ympev.2015.12.017.

**Rando, Juliana Gastaldello; Zuntini, Alexandre R.; Conceição, Adilva de Souza; van den Berg, Cássio; Pirani, José Rubens; Queiroz, Luciano Paganucci (2016):**
**Phylogeny of *Chamaecrista* ser. *Coriaceae* (Leguminosae) unveils a Lineage recently diversified in Brazilian Campo-Rupestre Vegetation.**
In: International Journal of Plant Sciences 177 (1), p. 3–17. DOI: 10.1086/683846.

**Ranjbar, Massoud; Hajmoradi, Fatemeh; Waycott, Michelle; van Dijk, Kor-jent (2014):**
**A Phylogeny of the Tribe Caraganeae (Fabaceae) based on DNA Sequence Data from ITS.**
In: Feddes Repertorium 125 (3-4), p. 78–84. DOI: 10.1002/fedr.201400051.

**Redden, Karen M.; Herendeen, Patrick p. (2006):**
**Morphology and phylogenetic Analysis of *Paloue* and related Genera in the *Brownea* Clade (Detarieae, Caesalpinioideae).**
In: International Journal of Plant Sciences 167 (6), p. 1229–1246.

**Redden, Karen M.; Herendeen, Patrick S.; Wurdack, Kenneth J.J.; Bruneau, Anne (2010):**
**Phylogenetic Relationships of the Northeastern South American *Brownea* Clade of Tribe Detarieae (Leguminosae: Caesalpinioideae) based on Morphology and molecular Data.**
In: Systematic Botany 35 (3), p. 524–533. DOI: 10.1600/036364410792495863.

**Riahi, Mehrshid; Zarre, Shahin; Maassoumi, Ali Aasghar; Kazempour-Osaloo, Shahrokh; Wojciechowski, Martin F. (2011):**
**Towards a Phylogeny for *Astragalus Section Caprini* (Fabaceae) and its Allies based on nuclear and Plastid DNA Sequences.**
In: Plant Systematics and Evolution 293 (1-4), p. 119–133. DOI: 10.1007/s00606-011-0417-3.

**Ribeiro, Renata Acácio; Lavin, Matt; Lemos-Filho, José Pires; Filho, Carlos Victor Mendonça; Santos, Fabrício Rodrigues dos; Lovato, Maria Bernadete (2007):**

**The Genus *Machaerium* (Leguminosae) is more closely related to *Aeschynomene* Sect. *Ochopodium* than to *Dalbergia*: Inferences from combined Sequence Data.**

In: Systematic Botany 32 (4), p. 762–771.

**Rockinger, Alexander; Flores, Andréia Silva; Renner, Susanne p. (2017):**

**Clock-dated Phylogeny for 48% of the 700 Species of *Crotalaria* (Fabaceae-Papilionoideae) resolves Sections worldwide and implies conserved flower and Leaf traits throughout its Pantropical Range.**

In: BMC Evolutionary Biology 17 (1), p. 61. DOI: 10.1186/s12862-017-0903-5.

**Rodrigues, Rodrigo Schütz; Flores, Andréia Silva (2012):**

**A new Combination in *Entada* (Leguminosae) from Roraima, Brazil.**

In: Phytotaxa 39, p. 47–50.

**Saini, Ajay; Jawali, Narendra (2009):**

**Molecular Evolution of 5s rDNA Region in *Vigna* Subgenus *Ceratotropis* and its phylogenetic Implications.**

In: Plant Systematics and Evolution 280 (3-4), p. 187–206. DOI: 10.1007/s00606-009-0178-4.

**Sakai, M.; Kanazawa, A.; Fujii, A.; Thseng, F. S.; Abe, J.; Shimamoto, Y. (2003):**

**Phylogenetic Relationships of the Chloroplast Genomes in the Genus *Glycine* inferred from four intergenic spacer Sequences.**

In: Plant Systematics and Evolution 239 (1-2), p. 29–54. DOI: 10.1007/s00606-002-0226-9.

**Samad, Farah Abdel; Baumel, Alex; Juin, Marianick; Pavon, Daniel; Siljak-Yakovlev, Sonja; Médail, Frédéric; Bou Dagher Kharrat, Magda (2014):**

**Phylogenetic Diversity and Genome sizes of *Astragalus* (Fabaceae) in the Lebanon biogeographical Crossroad.**

In: Plant Systematics and Evolution 300 (5), p. 819–830. DOI: 10.1007/s00606-013-0921-8.

**Särkinen, Tiina E.; Marcelo-Peña, José Luis; Yomona, A. Daza; Simon, Marcelo Fragomeni; Pennington, R. Toby; Hughes, Colin E. (2011):**

**Underestimated endemic Species Diversity in the dry inter-Andean valley of the Río Marañón, northern Peru: An example from *Mimosa* (Leguminosae, Mimosoideae).**

In: Taxon 60 (1), p. 139–150. DOI: 10.1002/tax.601012.

**Saslis-Lagoudakis, Charilaos; Chase, Mark W.; Robinson, Daniel N.; Russell, Stephen J.; Klitgaard, Bente B. (2008):**

**Phylogenetics of Neotropical *Platymiscium* (Leguminosae: Dalbergieae): Systematics, Divergence times, and Biogeography inferred from nuclear ribosomal and Plastid DNA Sequence Data.**

In: American Journal of Botany 95 (10), p. 1270–1286. DOI: 10.3732/ajb.0800101.

**Schaefer, Hanno; Hechenleitner, Paulina; Santos-Guerra, Arnoldo; Menezes de Sequeira, Miguel; Pennington, R. Toby; Kenicer, Gregory; Carine, Mark A. (2012):**

**Systematics, Biogeography, and Character Evolution of the Legume Tribe Fabeae with special Focus on the Middle-Atlantic Island Lineages.**

In: BMC Evolutionary Biology 12, p. 250. DOI: 10.1186/1471-2148-12-250.

**Scherson, Rosa A.; Vidal, Rodrigo; Sanderson, Michael J. (2008):**

**Phylogeny, Biogeography, and rates of Diversification of New World *Astragalus* (Leguminosae) with an Emphasis on South American Radiations.**

In: American Journal of Botany 95 (8), p. 1030–1039. DOI: 10.3732/ajb.0800017.

**Schley, Rowan J.; La Estrella, Manuel; Pérez-Escobar, Oscar Alejandro; Bruneau, Anne; Barraclough, Timothy G.; Forest, Félix; Klitgård, Bente B. (2018):**
**Is Amazonia a 'Museum' for Neotropical Trees? the Evolution of the *Brownea* Clade (Detarioideae, Leguminosae).**
In: Molecular Phylogenetics and Evolution 126, p. 279–292. DOI: 10.1016/j.ympev.2018.04.029.

**Schnabel, Andrew; McDonel, Patrick E.; Wendel, Jonathan F. (2003):**
**Phylogenetic Relationships in *Gleditsia* (Leguminosae) based on ITS Sequences.**
In: American Journal of Botany 90 (2), p. 310–320. DOI: 10.3732/ajb.90.2.310.

**Schnabel, Andrew; Wendel, Jonathan F. (1998):**
**Cladistic Biogeography of *Gleditsia* (Leguminosae) based on ndhF and rpl16 Chloroplast Gene Sequences.**
In: American Journal of Botany 85 (12), p. 1753–1765. DOI: 10.2307/2446510.

**Schrire, Brian D.; Lavin, Matt; Barker, Nigel P.; Forest, Félix (2009):**
**Phylogeny of the Tribe Indigofereae (Leguminosae-Papilionoideae): Geographically Structured more in succulent-rich and temperate Settings than in grass-rich Environments.**
In: American Journal of Botany 96 (4), p. 816–852. DOI: 10.3732/ajb.0800185.

**Schutte-Vlok, Anne Lise; van Wyk, Ben-Erik (2011):**
**A taxonomic Revision of *Podalyria* (Fabaceae).**
In: Systematic Botany 36 (3), p. 631–660. DOI: 10.1600/036364411X583628.

**Sede, Silvana M.; Tosto, D. S.; Gottlieb, Alexandra M.; Poggio, Lidia; Fortunato, René H. (2008):**
**Genetic Relationships in the *Galactia–Camptosema–Collaea* complex (Leguminosae) inferred from AFLP Markers.**
In: Plant Systematics and Evolution 276 (3-4), p. 261–270. DOI: 10.1007/s00606-008-0100-5.

**Shahi Shavvon, Robabeh; Kazempour-Osaloo, Shahrokh; Maassoumi, Ali Aasghar; Moharrek, Farideh; Karaman Erkul, Seher; Lemmon, Alan R. et al. (2017):**
**Increasing phylogenetic support for explosively radiating taxa: the promise of high-throughput Sequencing for *Oxytropis* (Fabaceae).**
In: Journal of Systematics and Evolution 55 (4), p. 385–404. DOI: 10.1111/jse.12269.

**Silva, Marcos José da; Queiroz, Luciano Paganucci; Tozzi, Ana Maria Goulart de Azevedo; Lewis, Gwilym Peter; Sousa, Anete Pereira de (2012):**
**Phylogeny and Biogeography of *Lonchocarpus* sensu lato and its Allies in the Tribe Millettieae (Leguminosae, Papilionoideae).**
In: Taxon 61 (1), p. 93–108.

**Simon, Marcelo Fragomeni; Grether, Rosaura; Queiroz, Luciano Paganucci; Särkinen, Tiina E.; Dutra, Valquíria F.; Hughes, Colin E. (2011):**
**The evolutionary History of *Mimosa* (Leguminosae): toward a Phylogeny of the sensitive Plants.**
In: American Journal of Botany 98 (7), p. 1201–1221. DOI: 10.3732/ajb.1000520.

**Simon, Marcelo Fragomeni; Pastore, José Floriano B.; Souza, Adriana F.; Borges, Leonardo Maurici; Scalon, Viviane R.; Ribeiro, Pétala G. et al. (2016):**
**Molecular Phylogeny of *Stryphnodendron* (Mimosoideae, Leguminosae) and Generic Delimitations in the *Piptadenia* Group.**
In: International Journal of Plant Sciences 177 (1), p. 44–59. DOI: 10.1086/684077.

**Simpson, Beryl B.; Larkin, Leah L.; Weeks, Andrea; McDill, Joshua R. (2006):**
**Phylogeny and Biogeography of *Pomaria* (Caesalpinioideae: Leguminosae).**
In: Systematic Botany 31 (4), p. 792–804.

**Sinou, Carole; Forest, Félix; Lewis, Gwilym Peter; Bruneau, Anne (2009):**
**The Genus *Bauhinia* s.l. (Leguminosae): a Phylogeny based on the Plastid trnL–trnF Region.**
In: Botany 87 (10), p. 947–960. DOI: 10.1139/B09-065.

**Sirichamorn, Yotsawate; Adema, Frits A.C.B.; Gravendeel, Barbara; van Welzen, Peter C. (2012):**
**Phylogeny of Palaeotropic *Derris*-like Taxa (Fabaceae) based on Chloroplast and nuclear DNA Sequences shows Reorganization of (infra)generic Classifications is needed.**
In: American Journal of Botany 99 (11), p. 1793–1808. DOI: 10.3732/ajb.1200390.

**Sirichamorn, Yotsawate; Adema, Frits A.C.B.; Roos, Marco C.; van Welzen, Peter C. (2014):**
**Molecular and morphological phylogenetic Reconstruction reveals a new generic Delimitation of Asian *Derris* (Fabaceae): Reinstatement of *Solori* and Synonymisation of *Paraderris* with *Derris*.**
In: Taxon 63 (3), p. 522–538. DOI: 10.12705/633.13.

**Smith, James F.; Zimmers., J. C. (2017):**
**New Combination in *Astragalus* (Fabaceae).**
In: Phytoneuron 38, p. 1–3.

**Snak, Cristiane; Vatanparast, Mohammad; Silva, Christian; Lewis, Gwilym Peter; Lavin, Matt; Kajita, Tadashi; Queiroz, Luciano Paganucci (2016):**
**A dated Phylogeny of the papilionoid Legume Genus *Canavalia* reveals recent Diversification by a Pantropical Liana Lineage.**
In: Molecular Phylogenetics and Evolution 98, p. 133–146. DOI: 10.1016/j.ympev.2016.02.001.

**Sotuyo, Solange; Contreras, José Luis; Gagnon, Edeline; Lewis, Gwilym Peter (2017):**
**A Synopsis of *Coulteria* (Leguminosae), including new Names and Synonyms.**
In: Phytotaxa 291 (1), p. 33. DOI: 10.11646/phytotaxa.291.1.3.

**Sousa, Filipe; Bertrand, Yann J.K.; Pfeil, Bernard E. (2016):**
**Patterns of phylogenetic Incongruence in *Medicago* found among six Loci.**
In: Plant Systematics and Evolution 302 (5), p. 493–513. DOI: 10.1007/s00606-016-1278-6.

**Sousa, Mario S.; Solange Sotuyo, J. (2012):**
**El Genero *Muellera* L.f. en Mesoamerica y Norte de Sudamerica.**
In: Acta Botanica Mexicana 100, p. 15–40.

**Sousa, Mario S.; Sotuyo, Solange; Pedraza-Ortega, Euler (2014):**
**Sistemática de *Lonchocarpus* sección *Punctati* (Fabaceae: Milletiieae), basada en Datos morfológicos y moleculares, con la Descripción de nueve Especies nuevas.**
In: Acta Botanica Mexicana 109, p. 79. DOI: 10.21829/abm109.2014.1148.

**Souza Conceição, Adilva; Paganucci de Queiroz, Luciano; Lewis, Gwilym Peter; Gomes de Andrade, Maria José; Machado de Almeida, Paulo Ricardo; Schnadelbach, Alessandra Selbach; van den Berg, Cássio (2009):**
**Phylogeny of *Chamaecrista* Moench (Leguminosae-Caesalpinioideae) based on nuclear and Chloroplast DNA Regions.**
In: Taxon 58 (4), p. 1168–1180. DOI: 10.1002/tax.584010.

**Souza, Alessandro Oliveira; Silva, Marcos José da (2015):**

**What's new in *Chamaecrista* (Fabaceae, Caesalpinioideae) from the Brazilian Cerrado?**
In: Phytotaxa 213 (3), p. 253. DOI: 10.11646/phytotaxa.213.3.5.

SOUZA, ÉLVIA RODRIGUES; LEWIS, GWILYM PETER; FOREST, FÉLIX; SCHNADELBACH, ALESSANDRA SELBACH; VAN DEN BERG, CÁSSIO; QUEIROZ, LUCIANO PAGANUCCI (2013):
**Phylogeny of *Calliandra* (Leguminosae: Mimosoideae) based on nuclear and Plastid molecular Markers.**
In: Taxon 62 (6), p. 1200–1219. DOI: 10.12705/626.2.

SOUZA, ISYS MASCARENHAS; FUNCH, LIGIA SILVEIRA; QUEIROZ, LUCIANO PAGANUCCI (2014):
**Morphological Analyses suggest a new taxonomic Circumscription for *Hymenaea courbaril* L. (Leguminosae, Caesalpinioideae).**
In: PhytoKeys (38), p. 101–118. DOI: 10.3897/phytokeys.38.7408.

STEELE, KELLY P.; ICKERT-BOND, STEFANIE M.; ZARRE, SHAHIN; WOJCIECHOWSKI, MARTIN F. (2010):
**Phylogeny and Character Evolution in *Medicago* (Leguminosae): Evidence from Analyses of Plastid trnK/matK and nuclear GA3ox1 Sequences.**
In: American Journal of Botany 97 (7), p. 1142–1155. DOI: 10.3732/ajb.1000009.

STEFANOVIĆ, SAŠA; PFEIL, BERNARD E.; PALMER, JEFFREY D.; DOYLE, JEFF J. (2009):
**Relationships among Phaseoloid Legumes based on Sequences from eight Chloroplast Regions.**
In: Systematic Botany 34 (1), p. 115–128. DOI: 10.1600/036364409787602221.

STRAUB, SHANNON C. K.; DOYLE, JEFF J. (2014):
**Molecular Phylogenetics of *Amorpha* (Fabaceae): an Evaluation of Monophyly, Species Relationships, and polyploid Origins.**
In: Molecular Phylogenetics and Evolution 76, p. 49–66. DOI: 10.1016/j.ympev.2014.02.025.

SUBRAMANIAM, SHWETA; PANDEY, ARUN KUMAR; GEETA, R.; MORT, MARK E. (2013):
**Molecular Systematics of Indian *Crotalaria* (Fabaceae) based on Analyses of nuclear ribosomal ITS DNA Sequences.**
In: Plant Systematics and Evolution 299 (6), p. 1089–1106. DOI: 10.1007/s00606-013-0781-2.

SUBRAMANIAM, SHWETA; PANDEY, ARUN KUMAR; RATHER, SHABIR A. (2015):
**A revised Circumscription of the Species in *Bracteatae* complex (section *Calycinae*) in the Genus *Crotalaria* L.: Evidence from nuclear and Chloroplast Markers.**
In: Plant Systematics and Evolution 301 (9), p. 2261–2290. DOI: 10.1007/s00606-015-1228-8.

SVEINSSON, SAEMUNDUR; CRONK, QUENTIN C.B. (2014):
**Evolutionary Origin of highly repetitive Plastid Genomes within the clover Genus (*Trifolium*).**
In: BMC Evolutionary Biology 14, p. 228. DOI: 10.1186/s12862-014-0228-6.

TEKPINAR, AYTEN; KARAMAN ERKUL, SEHER; AYTAÇ, ZEKI; KAYA, ZEKI (2016):
**Phylogenetic Relationships among native *Oxytropis* Species in Turkey Using the trnL Intron, trnL-F Igs, and trnV Intron cpDNA Regions.**
In: Turkish Journal of Botany 40, p. 472–479. DOI: 10.3906/bot-1506-45.

THOMPSON, IAN R. (2012):
**A Revision of eastern Australian *Bossiaea* (Fabaceae: Bossiaeeae).**
In: Muelleria 30 (2), p. 106–174.

TORKE, BENJAMIN M.; MANSANO, VIDAL DE FREITAS (2009):

**A phylogenetically based Sectional Classification of *Swartzia* (Leguminosae-Papilionoideae).**
In: Taxon 58 (3), p. 913–924. DOI: 10.1002/tax.583019.

TORKE, BENJAMIN M.; SCHAAL, BARBARA A. (2008):
**Molecular Phylogenetics of the species-rich Neotropical Genus *Swartzia* (Leguminosae, Papilionoideae) and related Genera of the Swartzioid Clade.**
In: American Journal of Botany 95 (2), p. 215–228. DOI: 10.3732/ajb.95.2.215.

TOSSO, FÉLICIEN; HARDY, OLIVIER J.; DOUCET, JEAN-LOUIS; DAÏNOU, KASSO; KAYMAK, ESRA; MIGLIORE, JÉRÉMY (2018):
**Evolution in the Amphi-Atlantic tropical Genus *Guibourtia* (Fabaceae, Detarioideae), combining NGS Phylogeny and Morphology.**
In: Molecular Phylogenetics and Evolution 120, p. 83–93. DOI: 10.1016/j.ympev.2017.11.026.

TRETHOWAN, LIAM; CLARK, RUTH; MACKINDER, BARBARA (2015):
**A Synopsis of the Neotropical Genus *Schnella* (Cercideae: Caesalpinioideae: Leguminosae) including 12 new Combinations.**
In: Phytotaxa 204 (4), p. 237. DOI: 10.11646/phytotaxa.204.4.1.

TURINI, FLORIAN G.; BRÄUCHLER, CHRISTIAN; HEUBL, GÜNTHER (2010):
**Phylogenetic Relationships and Evolution of morphological Characters in *Ononis* L. (Fabaceae).**
In: Taxon 59 (4), p. 1077–1090. DOI: 10.1002/tax.594008.

UYSAL, TUNA; ERTUĞRUL, KUDDISI; BOZKURT, MERYEM (2014):
**A new Genus segregated from *Thermopsis* (Fabaceae: Papilionoideae): *Vuralia*.**
In: Plant Systematics and Evolution 300 (7), p. 1627–1637. DOI: 10.1007/s00606-014-0988-x.

VAN DE WOUW, M.; MAXTED, N.; CHABANE, K.; FORD-LLOYD, B. V. (2001):
**Molecular Taxonomy of *Vicia ser. Vicia* based on amplified Fragment Length Polymorphisms.**
In: Plant Systematics and Evolution 229 (1-2), p. 91–105. DOI: 10.1007/s006060170020.

VAN DER BANK, MICHELLE; CHASE, MARK W.; VAN WYK, BEN-ERIK; FAY, MICHAEL F.; VAN DER BANK, FREDERICK H.; REEVES, GAIL; HULME, ALAN (2002):
**Systematics of the Tribe Podalyrieae (Fabaceae) based on DNA, morphological and chemical Data.**
In: Botanical Journal of the Linnean Society 139 (2), p. 159–170. DOI: 10.1046/j.1095-8339.2002.00051.x.

VAN DER MAESEN, L. J. G. (2003):
**Cajaninae of Australia (Leguminosae: Papilionoideae).**
In: Australian Systematic Botany 16 (2), p. 219–227. DOI: 10.1071/SB01047.

VAN DER WERFF, HENK (2008):
**A Synopsis of the Genus *Tachigali* (Leguminosae: Caesalpinioideae) in Northern South America.**
In: Annals of the Missouri Botanical Garden 95 (4), p. 618–661. DOI: 10.3417/2007159.

VANDER STAPPEN, J.; LAET, JAN; GAMA-LÓPEZ, S.; VAN CAMPENHOUT, S.; VOLCKAERT, G. (2002):
**Phylogenetic Analysis of *Stylosanthes* (Fabaceae) based on the Internal Transcribed Spacer Region (ITS) of nuclear ribosomal DNA.**
In: Plant Systematics and Evolution 234 (1), p. 27–51. DOI: 10.1007/s00606-002-0193-1.

VARELA, EDUARDO S.; LIMA, JOÃO P.M.S.; GALDINO, ALEXSANDRO S.; PINTO, LUCIANO DA S.; BEZERRA, WALDERLY M.; NUNES, EDSON P. ET AL. (2004):

**Relationships in Subtribe Diocleinae (Leguminosae; Papilionoideae) inferred from Internal Transcribed Spacer Sequences from nuclear ribosomal DNA.**
In: Phytochemistry 65 (1), p. 59–69. DOI: 10.1016/j.phytochem.2003.08.005.

VATANPARAST, MOHAMMAD; KLITGÅRD, BENTE B.; ADEMA, FRITS A.C.B.; PENNINGTON, R. TOBY; YAHARA, TETSUKAZU; KAJITA, TADASHI (2013):
**First molecular Phylogeny of the pantropical Genus *Dalbergia*: Implications for infrageneric Circumscription and Biogeography.**
In: South African Journal of Botany 89, p. 143–149. DOI: 10.1016/j.sajb.2013.07.001.

WAGSTAFF, STEVEN J.; HEENAN, PETER B.; SANDERSON, MICHAEL J. (1999):
**Classification, Origins, and Patterns of Diversification in New Zealand Carmichaelinae (Fabaceae).**
In: American Journal of Botany 86 (9), p. 1346–1356. DOI: 10.2307/2656781.

WANG, H. C.; SUN, H.; COMPTON, J. A.; YANG, J.-B. (2006):
**A Phylogeny of Thermopsideae (Leguminosae: Papilionoideae) inferred from nuclear ribosomal Internal Transcribed Spacer (ITS) Sequences.**
In: Botanical Journal of the Linnean Society 151 (3), p. 365–373. DOI: 10.1111/j.1095-8339.2006.00512.x.

WARWICK, M. C.; LEWIS, GWILYM PETER; LIMA, HAROLDO CAVALCANTE DE (2008):
**A reappraisal of *Barnebydendron* (Leguminosae: Caesalpinioideae: Detarieae).**
In: Kew Bulletin 63 (1), p. 143–149. DOI: 10.1007/s12225-007-9001-y.

WATSON, LINDA E.; SAYED-AHMED, H.; BADR, A. (2000):
**Molecular Phylogeny of Old World *Trifolium* (Fabaceae), based on Plastid and nuclear Markers.**
In: Plant Systematics and Evolution 224 (3-4), p. 153–171. DOI: 10.1007/BF00986340.

WILLIAMS, ANNA V.; MILLER, JOSEPH T.H.; SMALL, IAN; NEVILL, PAUL G.; BOYKIN, LAURA M. (2016):
**Integration of complete Chloroplast Genome Sequences with small Amplicon Datasets improves phylogenetic Resolution in *Acacia*.**
In: Molecular Phylogenetics and Evolution 96, p. 1–8. DOI: 10.1016/j.ympev.2015.11.021.

WOJCIECHOWSKI, MARTIN F. (2005):
***Astragalus* (Fabaceae): A molecular phylogenetic Perspective.**
In: Brittonia 57 (4), p. 382–396. DOI: 10.1663/0007-196X(2005)057[0382:AFAMPP]2.0.CO;2.

XIE, YAN-PING; MENG, YING; SUN, HANG; NIE, ZE-LONG (2016):
**Molecular Phylogeny of Gueldenstaedtia and Tibetia (Fabaceae) and their biogeographic Differentiation within Eastern Asia.**
In: Public Library of Science One 11 (9), e0162982. DOI: 10.1371/journal.pone.0162982.

XU, BO; WU, NING; GAO, XIN-FEN; ZHANG, LI-BING (2012):
**Analysis of DNA Sequences of six Chloroplast and nuclear Genes suggests Incongruence, Introgression, and incomplete Lineage Sorting in the Evolution of *Lespedeza* (Fabaceae).**
In: Molecular Phylogenetics and Evolution 62 (1), p. 346–358. DOI: 10.1016/j.ympev.2011.10.007.

YILDIZDOGAN, ZEYNEP; İKTEN, CENGIZ; MUTLU, NEDIM; TOKER, CENGIZ (2016):
**Genetic Relationships among the Genera *Cicer* L., *Lathyrus* L., *Lens* Mill., and *Vicia* L., together with similarity of *Lens* Taxa based on morphological and AFLP Markers.**
In: Turkish Journal of Botany 40, p. 566–575. DOI: 10.3906/bot-1507-9.

**Yoder, Jeremy B.; Briskine, Roman; Mudge, Joann; Farmer, Andrew; Paape, Timothy; Steele, Kelly P. et al. (2013):**
**Phylogenetic signal Variation in the Genomes of *Medicago* (Fabaceae).**
In: Systematic Biology 62 (3), p. 424–438. DOI: 10.1093/sysbio/syt009.

**Yue, Xue-Kun; Yue, Ji-Pei; Yang, Li-Er; Li, Zhi-Ming; Sun, Hang (2011):**
**Systematics of the Genus *Salweenia* (Leguminosae) from Southwest China with Discovery of a second Species.**
In: Taxon 60 (5), p. 1366–1374.

**Zhang, Ming-Li; Fritsch, Peter W.; Cruz, Boni C. (2009):**
**Phylogeny of *Caragana* (Fabaceae) based on DNA Sequence Data from rbcL, trnS-trnG, and ITS.**
In: Molecular Phylogenetics and Evolution 50 (3), p. 547–559. DOI: 10.1016/j.ympev.2008.12.001.

**Zhang, Ming-Li; Kang, Yun; Zhong, Yang; Sanderson, Stewart C. (2012):**
**Intense uplift of the Qinghai-Tibetan Plateau triggered rapid Diversification of *Phyllolobium* (Leguminosae) in the Late Cenozoic.**
In: Plant Ecology and Diversity 5 (4), p. 491–499. DOI: 10.1080/17550874.2012.727875.

**Zhang, Ming-Li; Wen, Zhi-Bin; Hao, Xiao-Li; Byalt, Vyacheslav V.; Sukhorukov, Alexander P.; Sanderson, Stewart C. (2015):**
**Taxonomy, Phylogenetics and Biogeography of *Chesneya* (Fabaceae), Evidenced from Data of three Sequences, ITS, trnS-trnG, and rbcL.**
In: Biochemical Systematics and Ecology 63, p. 80–89. DOI: 10.1016/j.bse.2015.09.017.

**Zitouna, Nadia; Marghali, Sonia; Gharbi, Maroua; Chennaoui-Kourda, Houda; Haddioui, Abdelmajid; Trifi-Farah, Neila (2013):**
**Mediterranean *Hedysarum* Phylogeny by transferable Microsatellites from *Medicago*.**
In: Biochemical Systematics and Ecology 50, p. 129–135. DOI: 10.1016/j.bse.2013.03.040.

### Fabales

**Bello, Maria Angélica; Bruneau, Anne; Forest, Félix; Hawkins, J. A. (2009):**
**Elusive Relationships within Order Fabales: phylogenetic Analyses using matK and rbcL Sequence Data.**
In: Systematic Botany 34 (1), p. 102–114. DOI: 10.1600/036364409787602348.

**Bello, Maria Angélica; Rudall, Paula J.; Hawkins, Julie A. (2012):**
**Combined phylogenetic Analyses reveal interfamilial Relationships and Patterns of Floral Evolution in the Eudicot Order Fabales.**
In: Cladistics 28 (4), p. 393–421. DOI: 10.1111/j.1096-0031.2012.00392.x.

**Ravi, V.; Khurana, Jitendra P.; Tyagi, Akhilesh K.; Khurana, Paramjit (2007):**
**Rosales Sister to Fabales: towards resolving the Rosid Puzzle.**
In: Molecular Phylogenetics and Evolution 44 (1), p. 488–493. DOI: 10.1016/j.ympev.2006.11.014.

### Fagaceae

**Chen, Li; Li, Jian-Qiang; Wang, Heng-Chang; Li, Xin-Wei; Peng, Yan-Song (2009):**
***Lithocarpus longzhouicus* comb. nov. (Fagaceae) from China: based on morphological and molecular Data.**
In: Nordic Journal of Botany 27 (2), p. 90–96. DOI: 10.1111/j.1756-1051.2008.00313.x.

**Deng, Min; Jiang, Xiao-Long; Hipp, Andrew L.; Manos, Paul S.; Hahn, Marlene (2018):**

**Phylogeny and Biogeography of East Asian evergreen oaks (*Quercus* Section *Cyclobalanopsis*; Fagaceae): Insights into the Cenozoic History of evergreen broad-leaved Forests in subtropical Asia.**
In: Molecular Phylogenetics and Evolution 119, p. 170–181. DOI: 10.1016/j.ympev.2017.11.003.

DENK, THOMAS (2003):
**Phylogeny of *Fagus* L. (Fagaceae) based on morphological Data.**
In: Plant Systematics and Evolution 240 (1-4), p. 55–81. DOI: 10.1007/s00606-003-0018-x.

DENK, THOMAS; GRIMM, GUIDO W. (2010):
**The Oaks of western Eurasia: Traditional Classifications and Evidence from two nuclear Markers.**
In: Taxon 59 (2), p. 351–366. DOI: 10.1002/tax.592002.

MANOS, PAUL S.; CANNON, CHARLES H.; OH, SANG-HUN (2008):
**Phylogenetic Relationships and taxonomic Status of the Paleoendemic Fagaceae of Western North America: Recognition of a new Genus, *Notholithocarpus*.**
In: Madroño 55 (3), p. 181–190. DOI: 10.3120/0024-9637-55.3.181.

MANOS, PAUL S.; DOYLE, J. J.; NIXON, K. C. (1999):
**Phylogeny, Biogeography, and Processes of molecular differentiation in *Quercus* Subgenus *Quercus* (Fagaceae).**
In: Molecular Phylogenetics and Evolution 12 (3), p. 333–349. DOI: 10.1006/mpev.1999.0614.

MANOS, PAUL S.; ZHOU, ZHE-KUN; CANNON, CHARLES H. (2001):
**Systematics of Fagaceae: phylogenetic Tests of reproductive Trait Evolution.**
In: International Journal of Plant Sciences 162 (6), p. 1361–1379.

OH, SANG-HUN; MANOS, PAUL P. (2008):
**Molecular Phylogenetics and Cupule Evolution in Fagaceae as inferred from nuclear CRABS CLAW Sequences.**
In: Taxon 57 (2), p. 434–451.

PEARSE, IAN S.; HIPP, ANDREW L. (2012):
**Global Patterns of Leaf Defenses in Oak Species.**
In: Evolution 66 (7), p. 2272–2286. DOI: 10.1111/j.1558-5646.2012.01591.x.

SIMEONE, MARCO C.; PIREDDA, ROBERTA; PAPINI, ALESSIO; VESSELLA, FEDERICO; SCHIRONE, BARTOLOMEO (2013):
**Application of Plastid and nuclear Markers to DNA barcoding of Euro-Mediterranean Oaks (Quercus, Fagaceae): Problems, Prospects and phylogenetic Implications.**
In: Botanical Journal of the Linnean Society 172 (4), p. 478–499. DOI: 10.1111/boj.12059.

### Flagellariaceae

WEPFER, PATRICIA H.; LINDER, HANS PETER (2014):
**The Taxonomy of *Flagellaria* (Flagellariaceae).**
In: Australian Systematic Botany 27 (3), p. 159–179. DOI: 10.1071/SB13048.

### Fouquieriaceae

DE-NOVA, JOSÉ ARTURO; SÁNCHEZ-REYES, LUNA L.; EGUIARTE, LUIS E.; MAGALLÓN, SUSANA (2018):
**Recent Radiation and Dispersal of an ancient Lineage: the case of *Fouquieria* (Fouquiericeae, Ericales) in North American Deserts.**
In: Molecular Phylogenetics and Evolution 126, p. 92–104. DOI: 10.1016/j.ympev.2018.03.026.

SCHULTHEIS, LISA M.; BALDWIN, BRUCE G. (1999):
**Molecular Phylogenetics of Fouquieriaceae: Evidence from nuclear rDNA ITS studies.**

In: American Journal of Botany 86 (4), p. 578–589. DOI: 10.2307/2656819.

### Fumariaceae

**Bertrand, Yann J.K.; Scheen, Anne-Cathrine; Marcussen, Thomas; Pfeil, Bernard E.; Sousa, Filipe; Oxelman, Bengt (2015):**

**Assignment of homoeologs to parental Genomes in Allopolyploids for Species Tree Inference, with an example from *Fumaria* (Papaveraceae).**

In: Systematic Biology 64 (3), p. 448–471. DOI: 10.1093/sysbio/syv004.

**Damerval, Catherine; Citerne, Hélène; Le Guilloux, Martine; Domenichini, Séverine; Dutheil, Justine; Ronse De Craene, Louis P.; Nadot, Sophie (2013):**

**Asymmetric morphogenetic Cues along the transverse Plane: Shift from Disymmetry to Zygomorphy in the Flower of Fumarioideae.**

In: American Journal of Botany 100 (2), p. 391–402. DOI: 10.3732/ajb.1200376.

**Pérez-Gutiérrez, Miguel A.; Romero-García, Ana T.; Fernández, M. Carmen; Blanca, Gabriel; Salinas-Bonillo, María J.; Suárez-Santiago, Víctor N. (2015):**

**Evolutionary History of fumitories (Subfamily Fumarioideae, Papaveraceae): An old Story shaped by the main geological and climatic Events in the Northern Hemisphere.**

In: Molecular Phylogenetics and Evolution 88, p. 75–92. DOI: 10.1016/j.ympev.2015.03.026.

**Pérez-Gutiérrez, Miguel A.; Romero-García, Ana T.; Salinas, María J.; Blanca, Gabriel; Fernández, M. Carmen; Suárez-Santiago, Víctor N. (2012):**

**Phylogeny of the Tribe Fumarieae (Papaveraceae s.l.) based on Chloroplast and nuclear DNA Sequences: Evolutionary and biogeographic Implications.**

In: American Journal of Botany 99 (3), p. 517–528. DOI: 10.3732/ajb.1100374.

**Salinas, María J.; Ruz-Rejn, C.; Morales, C.; Ruz-Rejn, M.; Romero, A. T.; Blanca, Gabriel et al. (2003):**

**Contribution to the Taxonomy and Phylogeny of *Sarcocapnos* DC. (Fumariaceae).**

In: Plant Systematics and Evolution 237 (3-4), p. 153–164. DOI: 10.1007/s00606-002-0260-7.

**Sauquet, Hervé; Carrive, Laetitia; Poullain, Noëlie; Sannier, Julie; Damerval, Catherine; Nadot, Sophie (2015):**

**Zygomorphy evolved from Disymmetry in Fumarioideae (Papaveraceae, Ranunculales): new Evidence from an expanded molecular phylogenetic Famework.**

In: Annals of Botany 115 (6), p. 895–914. DOI: 10.1093/aob/mcv020.

### Garryaceae

**Burge, Dylan O. (2011):**

**Molecular Phylogenetics of *Garrya* (Garryaceae).**

In: Madroño 58 (4), p. 249–255. DOI: 10.3120/0024-9637-58.4.249.

### Gelsemiaceae

**Jiao, Zhen; Jianhua, Li (2007):**

**Phylogeny of Intercontinental Disjunct Gelsemiaceae inferred from Chloroplast and Nuclear DNA Sequences.**

In: Systematic Botany 32 (3), p. 617–627.

**Struwe, Lena; Soza, Valerie L.; Manickam, Sugumaran; Olmstead, Richard G. (2014):**

**Gelsemiaceae (Gentianales) expanded to include the enigmatic Asian Genus *Pteleocarpa*.**

In: Botanical Journal of the Linnean Society 175, p. 482–496.

### Gentianaceae

**Calió, Maria Fernanda; Lepis, Katherine B.; Pirani, José Rubens; Struwe, Lena (2017):**

**Phylogeny of Helieae (Gentianaceae): Resolving taxonomic Chaos in a Neotropical Clade.**

In: Molecular Phylogenetics and Evolution 106, p. 192–208. DOI: 10.1016/j.ympev.2016.09.013.

**Calió, Maria Fernanda; Pirani, José Rubens; Struwe, Lena (2008):**

**Morphology-Based Phylogeny and Revision of *Prepusa* and *Senaea* (Gentianaceae: Helieae) —Rare Endemics from Eastern Brazil.**

In: Kew Bulletin 63 (2), p. 169–191.

**Chassot, Philippe; Nemomissa, Sileshi; Yuan, Yong-Ming; Küpfer, Philippe (2001):**

**High Paraphyly of *Swertia* L. (Gentianaceae) in the Gentianell-lineage as revealed by nuclear and Chloroplast DNA Sequence Variation.**

In: Plant Systematics and Evolution 229, p. 1–21.

**Chen, Sheng-Yun; Shi-Long, Chen; Tao, Xia; Yu-Jin, Wang (2005):**

**Phylogeny of *Metagentiana* and related Genera (Gentianaceae) inferred from nuclear ribosomal ITS Sequences.**

In: Acta Phytotaxonomica Sinica 43 (6), p. 491–502. DOI: 10.1360/aps040136.

**Chen, Sheng-Yun; Xia, Tao; Wang, Yujin; Liu, Jian-Quan; Chen, Shi-Long (2005):**

**Molecular Systematics and Biogeography of *Crawfurdia, Metagentiana* and *Tripterospermum* (Gentianaceae) based on nuclear ribosomal and Plastid DNA Sequences.**

In: Annals of Botany 96 (3), p. 413–424. DOI: 10.1093/aob/mci188.

**Christe, C.; Caetano, S.; Aeschimann, D.; Kropf, Matthias; Diadema, K.; Naciri, Y. (2014):**

**The intraspecific genetic variability of siliceous and calcareous *Gentiana* Species is shaped by contrasting demographic and Re-Colonization Processes.**

In: Molecular Phylogenetics and Evolution 70, p. 323–336. DOI: 10.1016/j.ympev.2013.09.022.

**Favre, Adrien; Matuszak, Sabine; Sun, Hang; Liu, Ende; Yuan, Yong-Ming; Müllner-Riehl, Alexandra N. (2014):**

**Two new Genera of Gentianinae (Gentianaceae): *Sinogentiana* and *Kuepferia* supported by molecular phylogenetic Evidence.**

In: Taxon 63 (2), p. 342–354. DOI: 10.12705/632.5.

**Favre, Adrien; Yuan, Yong-Ming; Küpfer, Philippe; Alvarez, Nadir (2010):**

**Phylogeny of Subtribe Gentianinae (Gentianaceae): Biogeographic Inferences despite limitations in temporal calibration Points.**

In: Taxon 59 (6), p. 1701–1711.

**Frasier, Cynthia (2008):**

**Amazonian Lowland, white Sand areas as ancestral Regions for South American Biodiversity: Biogeographic and phylogenetic Patterns in *Potalia* (Angiospermae: Gentianaceae).**

In: Organisms Diversity and Evolution 8 (1), p. 44–57. DOI: 10.1016/j.ode.2006.11.003.

**Hagen, K. Bernhard von; Kadereit, Joachim W. (2001):**

**The Phylogeny of (Gentianaceae) and its Colonization of the Southern Hemisphere as revealed by nuclear and Chloroplast DNA Sequence Variation.**

In: Organisms Diversity and Evolution 1 (1), p. 61–79. DOI: 10.1078/1439-6092-00005.

**Kissling, Jonathan; Buerki, Sven; Mansion, Guilhem (2009):**
***Klackenbergia* (Gentianaceae - Exaceae), a new endemic Genus from Madagascar.**
In: Taxon 58 (3), p. 907–912.

**Kissling, Jonathan; Yuan, Yong-Ming; Küpfer, Philippe; Mansion, Guilhem (2009):**
**The polyphyletic Genus *Sebaea* (Gentianaceae): a Step forward in Understanding the morphological and karyological Evolution of the Exaceae.**
In: Molecular Phylogenetics and Evolution 53 (3), p. 734–748. DOI: 10.1016/j.ympev.2009.07.025.

**Mansion, Guilhem (2004):**
**A new Classification of the polyphyletic Genus *Centaurium* Hill (Chironiinae, Gentianaceae): Description of the New World endemic *Zeltnera*, and Reinstatement of *Gyrandra* Griseb. and *Schenkia* Griseb.**
In: Taxon 53 (3), p. 719–740.

**Mansion, Guilhem; Struwe, Lena (2004):**
**Generic Delimitation and phylogenetic Relationships within the Subtribe Chironiinae (Chironieae: Gentianaceae), with special Reference to *Centaurium*: Evidence from nrDNA and cpDNA Sequences.**
In: Molecular Phylogenetics and Evolution 32 (3), p. 951–977. DOI: 10.1016/j.ympev.2004.03.016.

**Mansion, Guilhem; Zeltner, Louis (2004):**
**Phylogenetic Relationships within the New World endemic *Zeltnera* (Gentianaceaechironiinae) inferred from molecular and Karyological Data.**
In: American Journal of Botany 91 (12), p. 2069–2086.

**Mansion, Guilhem; Zeltner, Louis; Bretagnolle, François (2005):**
**Phylogenetic Patterns and polyploid Evolution within the Mediterranean Genus *Centaurium* (Gentianaceae - Chironieae).**
In: Taxon 54 (4), p. 931–950.

**Mathews, Katherine G.; Dunne, Niall; York, Emily; Struwe, Lena (2009):**
**A phylogenetic Analysis and taxonomic Revision of *Bartonia* (Gentianaceae: Gentianeae), based on molecular and morphological Evidence.**
In: Systematic Botany 34 (1), p. 162–172. DOI: 10.1600/036364409787602320.

**Matuszak, Sabine; Favre, Adrien; Schnitzler, Jan; Müllner-Riehl, Alexandra N. (2016):**
**Key innovations and climatic niche Divergence as drivers of Diversification in subtropical Gentianinae in southeastern and eastern Asia.**
In: American Journal of Botany 103 (5), p. 899–911. DOI: 10.3732/ajb.1500352.

**Merckx, Vincent S.F.T.; Kissling, Jonathan; Hentrich, Heiko; Janssens, Steven B.; Mennes, Constantijn B.; Specht, Chelsea D.; Smets, Erik F. (2013):**
**Phylogenetic Relationships of the mycoheterotrophic Genus *Voyria* and the Implications for the biogeographic History of Gentianaceae.**
In: American Journal of Botany 100 (4), p. 712–721. DOI: 10.3732/ajb.1200330.

**Molina, Jeanmaire E.; Struwe, Lena (2009):**
**Utility of secondary Structure in phylogenetic Reconstructions Using nrDNA ITS Sequences - an Example from Potalieae (Gentianaceae: Asteridae).**
In: Systematic Botany 34 (2), p. 414–428. DOI: 10.1600/036364409788606424.

**Struwe, Lena; Albert, Victor A.; Caliá, Fernanda M.; Frasier, Cynthia; Lepis, Katherine B.; Mathews, Katherine G.; Grant, Jason R. (2009):**

**Evolutionary Patterns in Neotropical Helieae (Gentianaceae): Evidence from Morphology, Chloroplast and nuclear DNA Sequences.**
In: Taxon 58 (2), p. 479–499.

SUGUMARAN, M.; WONG, KHOON MENG (2012):
**Studies in Malesian Gentianaceae I: *Fagraea* sensu lato-complex Genus or several Genera? A molecular phylogenetic study.**
In: Gardens' Bulletin Singapore 64 (2), p. 301–332.

SUGUMARAN, M.; WONG, KHOON MENG (2014):
**Studies in Malesian Gentianaceae VI. A Revision of *Utania* in the Malay Peninsula with two new Species.**
In: Plant Ecology and Evolution 147 (2), p. 213–223. DOI: 10.5091/plecevo.2014.971.

WONG, KHOON MENG (2012):
**Studies in Malesian Gentianaceae IV: A Revision of *Picrophloeus*.**
In: Gardens' Bulletin Singapore 64 (2), p. 511–522.

WONG, KHOON MENG; SUGUMARAN, M. (2012):
**Studies in Malesian Gentianaceae II: A taxonomic Famework for the *Fagraea*-complex, including the new Genus *Limahlania*.**
In: Gardens' Bulletin Singapore 64 (2), p. 481–495.

WONG, KHOON MENG; SUGUMARAN, M. (2012):
**Studies in Malesian Gentianaceae III: *Cyrtophyllum* reapplied to the *Fagraea fragrans* Alliance.**
In: Gardens' Bulletin Singapore 64 (2), p. 497–510.

WONG, KHOON MENG; SUGUMARAN, M.; SUGAU, J. B. (2013):
**Studies in Malesian Gentianaceae V. the *Fagraea* complex in Borneo: new generic assignments and Recombinations.**
In: Gardens' Bulletin Singapore 65 (2), p. 235–239.

YUAN, YONG-MING; WOHLHAUSER, SÉBASTIEN; MÖLLER, MICHAEL; CHASSOT, PHILIPPE; MANSION, GUILHEM; GRANT, JASON R. ET AL. (2003):
**Monophyly and Relationships of the Tribe Exaceae (Gentianaceae) inferred from nuclear ribosomal and Chloroplast DNA Sequences.**
In: Molecular Phylogenetics and Evolution 28 (3), p. 500–517. DOI: 10.1016/S1055-7903(03)00068-X.

YUAN, YONG-MING; WOHLHAUSER, SÉBASTIEN; MÖLLER, MICHAEL; KLACKENBERG, JENS; CALLMANDER, MARTIN W.; KÜPFER, PHILIPPE (2005):
**Phylogeny and Biogeography of *Exacum* (Gentianaceae): a disjunctive Distribution in the Indian Ocean Basin resulted from long Distance Dispersal and extensive Radiation.**
In: Systematic Biology 54 (1), p. 21–34. DOI: 10.1080/10635150590905867.

ZHANG, XIAO-LAN; WANG, YU-JIN; GE, XUE-JUN; YUAN, YONG-MING; YANG, HUI-LING; LIU, JIAN-QUAN (2009):
**Molecular Phylogeny and Biogeography of *Gentiana* Sect. *Cruciata* (Gentianaceae) based on four Chloroplast DNA Datasets.**
In: Taxon 58 (3), p. 862–870.

### Geraniaceae

AEDO, CARLOS; BARBERÁ, PATRICIA; BUIRA, ANTONI (2016):
**Taxonomic Revision of *Geranium* Sect. *Trilopha* (Geraniaceae).**

In: Systematic Botany 41 (2), p. 354–377. DOI: 10.1600/036364416X691812.

AEDO, CARLOS; GARCÍA, MIGUEL Á.; ALARCÓN, MARÍA LUISA; ALDASORO, JUAN JOSÉ; NAVARRO, CARMEN (2007):
**Taxonomic Revision of *Geranium* SubSect. *Mediterranea* (Geraniaceae).**
In: Systematic Botany 32 (1), p. 93–128.

BAKKER, FREEK T.; CULHAM, ALASTAIR; HETTIARACHI, PRIYANI; TOULOUMENIDOU, TASOULA; GIBBY, MARY (2004):
**Phylogeny of *Pelargonium* (Geraniaceae) based on DNA Sequences from three Genomes.**
In: Taxon 53 (1), p. 17–31. DOI: 10.2307/4135669.

BAKKER, FREEK T.; CULHAM, ALASTAIR; PANKHURST, CLIVE E.; GIBBY, MARY (2000):
**Mitochondrial and Chloroplast DNA-based Phylogeny of *Pelargonium* (Geraniaceae).**
In: American Journal of Botany 87 (5), p. 727–734. DOI: 10.2307/2656859.

FIZ, OMAR; VARGAS, PABLO; ALARCÓN, MARÍA LUISA; ALDASORO, JUAN JOSÉ (2006):
**Phylogenetic Relationships and Evolution in *Erodium* (Geraniaceae) based on trnL-trnF Sequences.**
In: Systematic Botany 41 (4), p. 739–763.

FIZ, OMAR; VARGAS, PABLO; ALARCÓN, MARISA; AEDO, CARLOS; GARCÍA, JOSÉ LUIS; ALDASORO, JUAN JOSÉ (2008):
**Phylogeny and Historical Biogeography of Geraniaceae in Relation to Climate Changes and Pollination Ecology.**
In: Systematic Botany 33 (2), p. 326–342. DOI: 10.1600/036364408784571482.

FIZ-PALACIOS, OMAR; VARGAS, PABLO; VILA, ROGER; PAPADOPULOS, ALEXANDER P. T.; ALDASORO, JUAN JOSÉ (2010):
**The uneven Phylogeny and Biogeography of *Erodium* (Geraniaceae): Radiations in the Mediterranean and recent recurrent intercontinental colonization.**
In: Annals of Botany 106 (6), p. 871–884. DOI: 10.1093/aob/mcq184.

JAMES, C. M.; GIBBY, MARY; BARRETT, J. A. (2004):
**Molecular studies in *Pelargonium* (Geraniaceae). A taxonomic Appraisal of Section *Ciconium* and the Origin of the 'Zonal' and 'Ivy-leaved' Cultivars.**
In: Plant Systematics and Evolution 243 (3-4), p. 131–146. DOI: 10.1007/s00606-003-0074-2.

JONES, CYNTHIA S.; BAKKER, FREEK T.; SCHLICHTING, CARL D.; NICOTRA, ADRIENNE B. (2009):
**Leaf shape Evolution in the South African Genus *Pelargonium* L'Her. (Geraniaceae).**
In: Evolution 63 (2), p. 479–497. DOI: 10.1111/j.1558-5646.2008.00552.x.

MARCUSSEN, THOMAS; MESEGUER, ANDREA SÁNCHEZ (2017):
**Species-level Phylogeny, Fruit Evolution and Diversification History of *Geranium* (Geraniaceae).**
In: Molecular Phylogenetics and Evolution 110, p. 134–149. DOI: 10.1016/j.ympev.2017.03.012.

MITCHELL, ANTHONY D.; HEENAN, PETER B.; PATERSON, A. M. (2009):
**Phylogenetic Relationships of *Geranium* Species indigenous to New Zealand.**
In: New Zealand Journal of Botany 47 (1), p. 21–31. DOI: 10.1080/00288250909509789.

ROESCHENBLECK, JOACHIM; ALBERS, FOCKE; MÜLLER, KAI F.; WEINL, STEFAN; KUDLA, JÖRG (2014):
**Phylogenetics, Character Evolution and a Subgeneric Revision of the Genus *Pelargonium* (Geraniaceae).**
In: Phytotaxa 159 (2), p. 31. DOI: 10.11646/phytotaxa.159.2.1.

TOULOUMENIDOU, TASOULA; BAKKER, FREEK T.; ALBERS, F. (2007):
**The Phylogeny of *Monsonia* L. (Geraniaceae).**
In: Plant Systematics and Evolution 264 (1-2), p. 1–14. DOI: 10.1007/s00606-006-0490-1.

**Weng, Mao-Lun; Ruhlman, Tracey A.; Gibby, Mary; Jansen, Robert K. (2012):**
**Phylogeny, rate Variation, and Genome size Evolution of *Pelargonium* (Geraniaceae).**
In: Molecular Phylogenetics and Evolution 64 (3), p. 654–670. DOI: 10.1016/j.ympev.2012.05.026.

### Gerrardinaceae

**Alford, Mac H. (2006):**
**Gerrardinaceae: a new Family of African Flowering Plants unresolved among Brassicales, Huerteales, Malvales, and Sapindales.**
In: Taxon 55 (4), p. 959–964. DOI: 10.2307/25065689.

### Gesneriaceae

**Anderson, Benjamin M.; Middleton, David J. (2013):**
**A Revision of *Rhynchotechum* Blume (Gesneriaceae).**
In: Edinburgh Journal of Botany 70 (01), p. 121–176. DOI: 10.1017/S0960428612000376.

**Araujo, Andréa Onofre de; Chautems, Alain (2015):**
**A new Species of *Sinningia* (Gesneriaceae) and additional floristic Data from Serra dos Carajás, Pará, Brazil.**
In: Phytotaxa 227 (2), p. 158. DOI: 10.11646/phytotaxa.227.2.5.

**Araujo, Andréa Onofre de; Chautems, Alain; Cardoso-Gustavson, Poliana; Souza, Vinicius Castro; Perret, Mathieu (2016):**
**Taxonomic Revision and phylogenetic Position of the Brazilian Endemic Genus *Sphaerorrhiza* (Sphaerorrhizinae, Gesneriaceae) including two new Species.**
In: Systematic Botany 41 (3), p. 651–664. DOI: 10.1600/036364416X692352.

**Berger, Andreas; Clark, John L.; Weber, Anton (2015):**
***Besleria macropoda* (Gesneriaceae): Lectotypification, Distribution, functional Epiphylly and discordant Fruit Morphology of a rare Costa Rican endemic.**
In: Phytotaxa 233 (2), p. 139–152. DOI: 10.11646/phytotaxa.233.2.2.

**Bransgrove, K.; Middleton, David J. (2015):**
**A Revision of *Epithema* (Gesneriaceae).**
In: Gardens' Bulletin Singapore 67 (01), p. 159. DOI: 10.3850/S2382581215000174.

**Christie, Frieda; Barber, S.; Möller, Michael (2012):**
**New Chromosome Counts in Old World Gesneriaceae: Numbers for Species hitherto regarded as *Chirita*, and their Systematic and evolutionary Significance.**
In: Edinburgh Journal of Botany 69 (02), p. 323–345. DOI: 10.1017/S0960428612000169.

**Clark, John L. (2012):**
***Gasteranthus diverticularis*, a new Species of Gesneriaceae from Southern Ecuador.**
In: Brittonia 64 (1), p. 1–5.

**Clark, John L.; Funke, Mindie M.; Duffy, Aaron M.; Smith, James F. (2012):**
**Phylogeny of a Neotropical Clade in the Gesneriaceae: More Tales of Convergent Evolution.**
In: International Journal of Plant Sciences 173 (8), p. 894–916. DOI: 10.1086/667229.

**Clark, John L.; Herendeen, Patrick S.; Skog, Laurence E.; Zimmer, Elizabeth A. (2006):**
**Phylogenetic Relationships and generic boundaries in the Episcieae (Gesneriaceae) inferred from nuclear, chloroplast, and morphological Data.**
In: Taxon 55 (2), p. 313–336. DOI: 10.2307/25065580.

**Clark, John L.; Neill, David A.; Weber, Anton; Gruhn, Jennifer A.; Katan, Tuntiak (2010):**

***Shuaria* (Gesneriaceae), an Arborescent new Genus from the Cordillera del Cóndor and Amazonian Ecuador.**
In: Systematic Botany 35 (3), p. 662–674. DOI: 10.1600/036364410792495917.

**CLARK, JOHN L.; ROALSON, ERIC H.; PRITCHARD, ROSS A.; COLEMAN, CASSANDRA L.; TEOH, VIEW-HUNE; MATOS, JESÚS (2011):**
**Independent Origin of Radial Floral Symmetry in the Gloxinieae (Gesnerioideae: Gesneriaceae) is supported by the Rediscovery of *Phinaea pulchella* in Cuba.**
In: Systematic Botany 36 (3), p. 757–767. DOI: 10.1600/036364411X583709.

**CLARK, JOHN R.; WAGNER, WARREN L.; ROALSON, ERIC H. (2009):**
**Patterns of Diversification and ancestral range Reconstruction in the southeast Asian-Pacific Angiosperm Lineage *Cyrtandra* (Gesneriaceae).**
In: Molecular Phylogenetics and Evolution 53 (3), p. 982–994. DOI: 10.1016/j.ympev.2009.09.002.

**CRONK, QUENTIN C.B.; KIEHN, MICHAEL; WAGNER, WARREN L.; SMITH, JAMES F. (2005):**
**Evolution of *Cyrtandra* (Gesneriaceae) in the Pacific Ocean: the Origin of a Supertramp Clade.**
In: American Journal of Botany 92 (6), p. 1017–1024.

**DENDUANGBORIPANT, J.; MENDUM, M.; CRONK, QUENTIN C.B. (2001):**
**Evolution in *Aeschynanthus* (Gesneriaceae) inferred from ITS Sequences.**
In: Plant Systematics and Evolution 228 (3-4), p. 181–197. DOI: 10.1007/s006060170028.

**FERREIRA, GABRIEL E.; CHAUTEMS, ALAIN; HOPKINS, MICHAEL J. G.; PERRET, MATHIEU (2016):**
**Independent Evolution of pouched flowers in the Amazon is supported by the Discovery of a new Species of *Lesia* (Gesneriaceae) from Serra do Aracá Tepui in Brazil.**
In: Plant Systematics and Evolution 302 (8), p. 1109–1119. DOI: 10.1007/s00606-016-1320-8.

**FREIBERG, MARTIN (1996):**
**The Gesneriad Flora of the Los Cedros Biological Reserve, Northwest Ecuador, Part 1: Four new Species in *Gasteranthus* (Gesneriaceae).**
In: Phyton (Horn, Austria) 36 (2), p. 303–309.

**FREIBERG, MARTIN (1997):**
**The Gesneriad Flora of the Los Cedros Biological Reserve, Northwest Ecuador, Part 2: new Species in *Alloplectus, Dalbergaria, Paradrymonia,* and *Pentadenia* (Gesneriaceae).**
In: Phyton (Horn, Austria) 37 (1), p. 133–140.

**FREIBERG, MARTIN (1998):**
**Two remarkable new Species of *Gasteranthus* (Gesneriaceae) from Central Ecuador.**
In: Phyton (Horn, Austria) 37 (2), p. 345–351.

**FREIBERG, MARTIN (2000):**
**Three new Species of *Gasteranthus* (Gesneriaceae) from Ecuador.**
In: Brittonia 52, p. 227–232.

**KOKUBUGATA, GORO; HIRAYAMA, YUMIKO; PENG, CHING-I.; YOKOTA, MASATSUGU; MÖLLER, MICHAEL (2011):**
**Phytogeographic aspects of *Lysionotus pauciflorus* sensu lato (Gesneriaceae) in the China, Japan and Taiwan Regions: phylogenetic and morphological Relationships and taxonomic Consequences.**
In: Plant Systematics and Evolution 292 (3-4), p. 177–188. DOI: 10.1007/s00606-010-0410-2.

LI, JIA-MEI; WANG, YIN-ZHENG (2007):
**Phylogenetic Reconstruction among Species of *Chiritopsis* and *Chirita* Sect. *Gibbosaccus* (Gesneriaceae) based on nrDNA ITS and cpDNA trnL-F Sequences.**
In: Systematic Botany 32 (4), p. 888–898.

LINDQVIST, CHARLOTTE; ALBERT, VICTOR A. (1999):
**Phylogeny and conservation of African violets (*Saintpaulia*: Gesneriaceae): new findings based on nuclear ribosomal 5s non-transcribed Spacer Sequences.**
In: Kew Bulletin 54, p. 363–377.

MAYER, VERONIKA; MÖLLER, MICHAEL; PERRET, MATHIEU; WEBER, ANTON (2003):
**Phylogenetic Position and generic differentiation of Epithemateae (Gesneriaceae) inferred from Plastid DNA Sequence Data.**
In: American Journal of Botany 90 (2), p. 321–329. DOI: 10.3732/ajb.90.2.321.

MIDDLETON, DAVID J.; MÖLLER, MICHAEL (2012):
***Tribounia*, a new Genus of Gesneriaceae from Thailand.**
In: Taxon 61 (6), p. 1286–1295. DOI: 10.1002/tax.616009.

MIDDLETON, DAVID J.; NISHII, KANAE; PUGLISI, CARMEN; FORREST, LAURA L.; MÖLLER, MICHAEL (2015):
***Chayamaritia* (Gesneriaceae: Didymocarpoideae), a new Genus from Southeast Asia.**
In: Plant Systematics and Evolution 301 (7), p. 1947–1966. DOI: 10.1007/s00606-015-1213-2.

MIDDLETON, DAVID J.; WEBER, ANTON; YAO, T. L.; SONTAG, SUSANNE; MÖLLER, MICHAEL (2013):
**The current Status of the Species hitherto assigned to *Henckelia* (Gesneriaceae).**
In: Edinburgh Journal of Botany 70 (03), p. 385–404. DOI: 10.1017/S0960428613000127.

MÖLLER, MICHAEL; FORREST, ALAN; WEI, YI-GANG; WEBER, ANTON (2011):
**A molecular phylogenetic assessment of the advanced Asiatic and Malesian didymocarpoid Gesneriaceae with focus on non-monophyletic and monotypic Genera.**
In: Plant Systematics and Evolution 292 (3-4), p. 223–248. DOI: 10.1007/s00606-010-0413-z.

MÖLLER, MICHAEL; KOKUBUGATA, GORO; JONG, KWITON (2008):
**Aspects of Genome Evolution in Gesneriaceae: Patterns of 45S-nrDNA Site Evolution based on Evidence from Fluorescent in Situ Hybridization (Fish).**
In: International Journal of Plant Sciences 169 (5), p. 667–676. DOI: 10.1086/533609.

MÖLLER, MICHAEL; NISHII, KANAE; ATKINS, H. J.; KONG, H. H.; KANG, MING; WEI, Y. G. ET AL. (2016):
**An Expansion of the Genus *Deinostigma* (Gesneriaceae).**
In: Gardens' Bulletin Singapore 68 (01), p. 145. DOI: 10.3850/S2382581216000119.

MÖLLER, MICHAEL; PFOSSER, MARTIN; JANG, CHANG-GEE; MAYER, VERONIKA; CLARK, ALEXANDRA; HOLLINGSWORTH, MICHELLE L. ET AL. (2009):
**A preliminary Phylogeny of the 'Didymocarpoid Gesneriaceae' based on three molecular Data sets: Incongruence with available tribal Classifications.**
In: American Journal of Botany 96 (5), p. 989–1010. DOI: 10.3732/ajb.0800291.

MORA, M. MARCELA; CLARK, JOHN L. (2016):
**Molecular Phylogeny of the Neotropical Genus *Paradrymonia* (Gesneriaceae), Reexamination of Generic Concepts and the Resurrection of *Trichodrymonia* and *Centrosolenia*.**
In: Systematic Botany 41 (1), p. 82–104. DOI: 10.1600/036364416X690561.

NING, ZU-LIN; PAN, BO; WEN, FANG; KANG, MING; ZHUANG, XUE-YING (2016):

***Primulina yingdeensis*, a new Species from Guangdong, China, and *P. rosulata*, a new Combination (Gesneriaceae), based on morphological and molecular Evidence.**
In: Willdenowia 46 (3), p. 399–409. DOI: 10.3372/wi.46.46308.

**NISHII, KANAE; HUGHES, MARK; BRIGGS, MARIE; HASTON, ELSPETH M.; CHRISTIE, FRIEDA; DEVILLIERS, MARGARET J. ET AL. (2015):**
***Streptocarpus* redefined to include all Afro-Malagasy Gesneriaceae: molecular Phylogenies prove congruent with geographical Distribution and basic Chromosome Numbers and uncover remarkable morphological Homoplasies.**
In: Taxon 64 (6), p. 1243–1274. DOI: 10.12705/646.8.

**PERRET, MATHIEU; CHAUTEMS, ALAIN; ARAUJO, ANDRÉA ONOFRE DE; SALAMIN, NICOLAS (2013):**
**Temporal and spatial Origin of Gesneriaceae in the New World inferred from Plastid DNA Sequences.**
In: Botanical Journal of the Linnean Society 171, p. 61–79.

**PERRET, MATHIEU; CHAUTEMS, ALAIN; SPICHIGER, RODOLPHE (2006):**
**Dispersal-vicariance Analyses in the Tribe Sinningieae (Gesneriaceae): A Clue To Understanding Biogeographical History of the Brazilian Atlantic Forest.**
In: Annals of the Missouri Botanical Garden 93 (2), p. 340–358. DOI: 10.3417/0026-6493(2006)93[340:DAITTS]2.0.CO;2.

**PERRET, MATHIEU; CHAUTEMS, ALAIN; SPICHIGER, RODOLPHE; BARRACLOUGH, TIMOTHY G.; SAVOLAINEN, VINCENT (2007):**
**The geographical Pattern of Speciation and Floral Diversification in the Neotropics: the Tribe Sinningieae (Gesneriaceae) as a Case Study.**
In: Evolution 61 (7), p. 1641–1660. DOI: 10.1111/j.1558-5646.2007.00136.x.

**PERRET, MATHIEU; CHAUTEMS, ALAIN; SPICHIGER, RODOLPHE; KITE, GEOFFREY C.; SAVOLAINEN, VINCENT (2003):**
**Systematics and Evolution of Tribe Sinningieae (Gesneriaceae): Evidence from phylogenetic Analyses of six Plastid DNA Regions and Nuclear NCpgs.**
In: American Journal of Botany 90 (3), p. 445–460.

**PETROVA, GALYA; MOYANKOVA, DANIELA; NISHII, KANAE; FORREST, LAURA L.; TSIRIPIDIS, IOANNIS; DROUZAS, ANDREAS D. ET AL. (2015):**
**The European Paleoendemic *Haberlea rhodopensis* (Gesneriaceae) has an Oligocene Origin and a Pleistocene Diversification and occurs in a long-persisting Refugial Area in Southeastern Europe.**
In: International Journal of Plant Sciences 176 (6), p. 499–514. DOI: 10.1086/681990.

**PILLON, YOHAN; JOHANSEN, JENNIFER B.; SAKISHIMA, TOMOKO; ROALSON, ERIC H.; PRICE, DONALD K.; STACY, ELIZABETH A. (2013):**
**Gene discordance in phylogenomics of recent plant Radiations, an example from Hawaiian *Cyrtandra* (Gesneriaceae).**
In: Molecular Phylogenetics and Evolution 69 (1), p. 293–298. DOI: 10.1016/j.ympev.2013.05.003.

**PUGLISI, CARMEN; MIDDLETON, DAVID J.; TRIBOUN, PRAMOTE; MÖLLER, MICHAEL (2011):**
**New Insights into the Relationships between *Paraboea, Trisepalum,* and *Phylloboea* (Gesneriaceae) and their taxonomic consequences.**
In: Taxon 60 (6), p. 1693–1702. DOI: 10.1002/tax.606014.

**PUGLISI, CARMEN; YAO, TZE LEONG; MILNE, RICHARD IAN; MÖLLER, MICHAEL; MIDDLETON, DAVID J. (2016):**
**Generic Recircumscription in the Loxocarpinae (Gesneriaceae), as inferred by phylogenetic and morphological Data.**

In: Taxon 65 (2), p. 277–292. DOI: 10.12705/652.5.

QIU, ZHI-JING; WANG, XIAO-LING; LIU, ZHENG-YU; YANG, JIAN-FEN (2012):
**Cytological and phylogenetic Study of *Petrocosmea hexiensis* (Gesneriaceae), a new Species from Chongqing, China.**
In: Phytotaxa 74, p. 30–38.

ROALSON, ERIC H.; BOGGAN, JOHN K.; SKOG, LAURENCE E.; ZIMMER, ELIZABETH A. (2005):
**Untangling Gloxinieae (Gesneriaceae). I. phylogenetic Patterns and generic Boundaries inferred from nuclear, chloroplast, and morphological cladistic Datasets.**
In: Taxon 54 (2), p. 389–410. DOI: 10.2307/25065368.

ROALSON, ERIC H.; SKOG, LAURENCE E.; ZIMMER, ELIZABETH A. (2003):
**Phylogenetic Relationships and the Diversification of Floral Form in *Achimenes* (Gesneriaceae).**
In: Systematic Botany 28 (3), p. 593–608.

ROALSON, ERIC H.; SKOG, LAURENCE E.; ZIMMER, ELIZABETH A. (2008):
**Untangling Gloxinieae (Gesneriaceae). II. Reconstructing Biogeographic Patterns and Estimating Divergence Times among New World Continental and Island Lineages.**
In: Systematic Botany 33 (1), p. 159–175.

SCHULTE, LACIE J.; CLARK, JOHN L.; NOVAK, STEPHEN J.; JEFFRIES, SHANDRA K.; SMITH, JAMES F. (2015):
**Speciation within *Columnea* Section *Angustiflora* (Gesneriaceae): Islands, pollinators and climate.**
In: Molecular Phylogenetics and Evolution 84, p. 125–144. DOI: 10.1016/j.ympev.2014.12.008.

SERRANO-SERRANO, MARTHA LILIANA; PERRET, MATHIEU; GUIGNARD, MAÏTÉ; CHAUTEMS, ALAIN; SILVESTRO, DANIELE; SALAMIN, NICOLAS (2015):
**Decoupled Evolution of Floral traits and climatic Preferences in a Clade of Neotropical Gesneriaceae.**
In: BMC Evolutionary Biology 15, p. 247. DOI: 10.1186/s12862-015-0527-6.

SMITH, JAMES F. (2000):
**A phylogenetic Analysis of Tribes Beslerieae and Napeantheae (Gesneriaceae) and Evolution of Fruit Types: Parsimony and Maximum Likelihood Analyses of ndhF Sequences.**
In: Systematic Botany 25 (1), p. 72. DOI: 10.2307/2666674.

SMITH, JAMES F. (2000):
**Phylogenetic Resolution within the Tribe Episcieae (Gesneriaceae): Congruence of ITS and Ndhf Sequences from Parsimony and Maximum-likelihood Analyses.**
In: American Journal of Botany 87 (6), p. 883–897.

SMITH, JAMES F.; CLARK, JOHN L. (2013):
**Molecular phylogenetic Analyses reveal undiscovered monospecific Genera in the Tribe Episcieae (Gesneriaceae).**
In: Systematic Botany 38 (2), p. 451–463. DOI: 10.1600/036364413X666723.

SMITH, JAMES F.; CLARK, JOHN L.; AMAYA-MÁRQUEZ, MARISOL; MARÍN-GÓMEZ, OSCAR H. (2017):
**Resolving incongruence: Species of hybrid Origin in *Columnea* (Gesneriaceae).**
In: Molecular Phylogenetics and Evolution 106, p. 228–240. DOI: 10.1016/j.ympev.2016.10.001.

SMITH, JAMES F.; DRAPER, SCOTTIE B.; HILEMAN, LENA C.; BAUM, DAVID A. (2004):

**A phylogenetic Analysis within Tribes Gloxinieae and Gesnerieae (Gesnerioideae: Gesneriaceae).**
In: Systematic Botany 29 (4), p. 947–958. DOI: 10.1600/0363644042451116.

SMITH, JAMES F.; FUNKE, MINDIE M.; WOO, V. L. (2006):
**A duplication of Gcyc predates Divergence within Tribe Coronanthereae (Gesneriaceae): phylogenetic Analysis and Evolution.**
In: Plant Systematics and Evolution 261 (1-4), p. 245–256. DOI: 10.1007/s00606-006-0445-6.

SMITH, JAMES F.; HILEMAN, LENA C.; POWELL, MARTYN P.; BAUM, DAVID A. (2004):
**Evolution of Gcyc, a Gesneriaceae homolog of Cycloidea, within Gesnerioideae (Gesneriaceae).**
In: Molecular Phylogenetics and Evolution 31 (2), p. 765–779. DOI: 10.1016/j.ympev.2003.09.012.

WANG, CHUN-NENG; MÖLLER, MICHAEL; CRONK, QUENTIN C.B. (2004):
**Phylogenetic Position of *Titanotrichum oldhamii* (Gesneriaceae) inferred from Four Different Gene Regions.**
In: Systematic Botany 29 (2), p. 407–418.

WANG, Y.-Z.; LI, Z.-Y. (2002):
**Inflorescence development of *Whytockia* (Epithemateae, Gesneriaceae) and phylogenetic Implications within Gesneriaceae.**
In: Plant Systematics and Evolution 236 (1), p. 45–54. DOI: 10.1007/s00606-002-0234-9.

WANG, YIN-ZHENG; LIANG, RONG-HUA; WANG, BO-HAN; LI, JIA-MEI; QIU, ZHI-JING; LI, ZHEN-YU; WEBER, ANTON (2010):
**Origin and phylogenetic Relationships of the Old World Gesneriaceae with actinomorphic flowers inferred from ITS and trnL-trnF Sequences.**
In: Taxon 59 (4), p. 1044–1052. DOI: 10.1002/tax.594005.

WANG, YIN-ZHENG; MAO, RU-BING; LIU, YAN; LI, JIA-MEI; DONG, YANG; LI, ZHEN-YU; SMITH, JAMES F. (2011):
**Phylogenetic Reconstruction of *Chirita* and Allies (Gesneriaceae) with taxonomic Treatments.**
In: Journal of Systematics and Evolution 49 (1), p. 50–64. DOI: 10.1111/j.1759-6831.2010.00113.x.

WEBER, ANTON; CLARK, JOHN L.; MÖLLER, MICHAEL (2013):
**A new formal Classification of Gesneriaceae.**
In: Selbyana 31 (2), p. 68–94.

WEBER, ANTON; MIDDLETON, DAVID J.; FORREST, ALAN; KIEW, RUTH; LIM, CHUNG LU; RAFIDAH, A. R. ET AL. (2011):
**Molecular Systematics and remodelling of *Chirita* and associated Genera (Gesneriaceae).**
In: Taxon 60 (3), p. 767–790. DOI: 10.1002/tax.603012.

YU, XUN-LIN; LI, MING; ZHOU, JIAN-JUN; LI, PENG-WEI (2015):
***Petrocodon hunanensis* (Gesneriaceae), a new Species identified by both morphological and molecular Evidence from Limestone Area in Hunan, China.**
In: Phytotaxa 195 (1), p. 65. DOI: 10.11646/phytotaxa.195.1.4.

### Gisekiaceae

BISSINGER, KERSTIN; KHOSHRAVESH, ROXANNE; KOTRADE, JAN PETER; OAKLEY, JASON; SAGE, TAMMY L.; SAGE, ROWAN F. ET AL. (2014):
***Gisekia* (Gisekiaceae): phylogenetic Relationships, Biogeography, and Ecophysiology of a poorly known $C_4$ Lineage in the Caryophyllales.**

In: American Journal of Botany 101 (3), p. 499–509. DOI: 10.3732/ajb.1300279.

### Gleicheniaceae

**Brownsey, P. J.; Ewans, R.; Rance, B.; Walls, S.; Perrie, Leon R. (2013):**

**A Review of the Fern Genus *Sticherus* (Gleicheniaceae) in New Zealand with Confirmation of two new Species Records.**

In: New Zealand Journal of Botany 51 (2), p. 104–115. DOI: 10.1080/0028825X.2013.773917.

**Gonzales, R. Jasivia; Kessler, Michael (2011):**

**A Synopsis of the Neotropical Species of *Sticherus* (Gleicheniaceae), with Descriptions of nine new Species.**

In: Phytotaxa 31, p. 1–54.

**Perrie, Leon R.; Bayly, Michael J.; Lehnebach, Carlos A.; Brownsey, Patrick J. (2007):**

**Molecular Phylogenetics and molecular Dating of the New Zealand Gleicheniaceae.**

In: Brittonia 59 (2), p. 129–141. DOI: 10.1663/0007-196X(2007)59[129:MPAMDO]2.0.CO;2.

### Gnetaceae

**Biye, Elvire H.; Balkwill, Kevin; Cron, Glynis V. (2014):**

**A clarification of *Gnetum* L. (Gnetaceae) in Africa and the Description of two new Species.**

In: Plant Systematics and Evolution 300 (2), p. 263–272. DOI: 10.1007/s00606-013-0879-6.

**Hajibabaei, Mehrdad; Xia, Junnan; Drouin, Guy (2006):**

**Seed plant Phylogeny: Gnetophytes are derived Conifers and a Sister Group to Pinaceae.**

In: Molecular Phylogenetics and Evolution 40 (1), p. 208–217. DOI: 10.1016/j.ympev.2006.03.006.

**Hou, Chen; Humphreys, Aelys M.; Thureborn, Olle; Rydin, Catarina (2015):**

**New Insights into the evolutionary History of *Gnetum* (Gnetales).**

In: Taxon 64 (2), p. 239–253. DOI: 10.12705/642.12.

**Won, Hyosig; Renner, Susanne p. (2005):**

**The Internal Transcribed Spacer of nuclear ribosomal DNA in the Gymnosperm *Gnetum*.**

In: Molecular Phylogenetics and Evolution 36 (3), p. 581–597. DOI: 10.1016/j.ympev.2005.03.011.

**Won, Hyosig; Renner, Susanne p. (2006):**

**Dating Dispersal and Radiation in the Gymnosperm *Gnetum* (Gnetales) - Clock Calibration When Outgroup Relationships are uncertain.**

In: Systematic Biology 55 (4), p. 610–622. DOI: 10.1080/10635150600812619.

**Wu, Chung-Shien; Lai, Yu-Ting; Lin, Ching-Ping; Wang, Ya-Nan; Chaw, Shu-Miaw (2009):**

**Evolution of reduced and compact Chloroplast Genomes (cpDNAs) in Gnetophytes: selection toward a lower-cost Strategy.**

In: Molecular Phylogenetics and Evolution 52 (1), p. 115–124. DOI: 10.1016/j.ympev.2008.12.026.

### Goodeniaceae

**Gardner, Andrew G.; Sessa, Emily B.; Michener, Pryce; Johnson, Eden; Shepherd, Kelly A.; Howarth, Dianella G.; Jabaily, Rachel Schmidt (2016):**

**Utilizing Next-generation Sequencing to resolve the Backbone of the core Goodeniaceae and inform future taxonomic and Floral form Studies.**

In: Molecular Phylogenetics and Evolution 94 (Pt B), p. 605–617. DOI: 10.1016/j.ympev.2015.10.003.

**Howarth, Dianella G.; Baum, David A. (2005):**

**Genealogical Evidence of homoploid Hybrid Speciation in an adaptive Radiation of *Scaevola* (Goodeniaceae) in the Hawaiian Islands.**
In: Evolution 59 (5), p. 948. DOI: 10.1554/03-752.

HOWARTH, DIANELLA G.; GUSTAFSSON, MATS H.G.; BAUM, DAVID A.; MOTLEY, TIMOTHY J. (2003):
**Phylogenetics of the Genus *Scaevola* (Goodeniaceae): Implication for Dispersal Patterns across the Pacific Basin and Colonization of the Hawaiian Islands.**
In: American Journal of Botany 90 (6), p. 915–923. DOI: 10.3732/ajb.90.6.915.

JABAILY, RACHEL SCHMIDT; SHEPHERD, KELLY A.; GUSTAFSSON, MATS H.G.; SAGE, LEIGH W.; KRAUSS, SIEGY L.; HOWARTH, DIANELLA G.; MOTLEY, TIMOTHY J. (2012):
**Systematics of the Austral-Pacific Family Goodeniaceae: Establishing a taxonomic and evolutionary Famework.**
In: Taxon 61 (2), p. 419–436.

### Grewiaceae

BRUNKEN, ULRIKE; MÜLLNER, ALEXANDRA N. (2012):
**A new tribal Classification of Grewioideae (Malvaceae) based on morphological and molecular phylogenetic Evidence.**
In: Systematic Botany 37 (3), p. 699–711. DOI: 10.1600/036364412X648670.

WHITLOCK, BARBARA A.; KAROL, KENNETH G.; ALVERSON, WILLIAM P. (2003):
**Chloroplast DNA Sequences confirm the Placement of the Enigmatic *Oceanopapaver* within *Corchorus* (Grewioideae: Malvaceae s.l., formerly Tiliaceae).**
In: International Journal of Plant Sciences 164 (1), p. 35–41.

### Grossulariaceae

SCHULTHEIS, LISA M.; DONOGHUE, MICHAEL J. (2004):
**Molecular Phylogeny and Biogeography of *Ribes* (Grossulariaceae), with an Emphasis on Gooseberries (Subg. *Grossularia*).**
In: Systematic Botany 29 (1), p. 77–96.

SENTERS, ANNE E.; SOLTIS, DOUGLAS E. (2003):
**Phylogenetic Relationships in *Ribes* (Grossulariaceae) inferred from ITS Sequence Data.**
In: Taxon 52, p. 51–66.

### Guamatelaceae

OH, SANG-HUN; POTTER, DANIEL (2006):
**Description and phylogenetic Position of a new Angiosperm Family, Guamatelaceae, inferred from Chloroplast rbcL, atpB, and matK Sequences.**
In: Systematic Botany 31 (4), p. 730–738.

### Gunneraceae

WANNTORP, LIVIA; WANNTORP, HANS-ERIK; KÄLLERSJÖ, MARI (2002):
**The identity of *Gunnera manicata* Linden ex André - resolving a Brazilian-Colombian Enigma.**
In: Taxon 51 (3), p. 493–497. DOI: 10.2307/1554861.

WANNTORP, LIVIA; WANNTORP, HANS-ERIK; OXELMAN, BENGT; KÄLLERSJÖ, MARI (2001):
**Phylogeny of *Gunnera*.**
In: Plant Systematics and Evolution 226 (1-2), p. 85–107. DOI: 10.1007/s006060170075.

### Gymnospermae

**Averyanov, Leonid V.; Nguyen, Tien Hiep; Sinh, Khang Nguyen; van Pham, The; Lamxay, Vichith; Bounphanmy, Somchanh et al. (2014):**

**Gymnosperms of Laos.**

In: Nordic Journal of Botany 32 (6), p. 765–805. DOI: 10.1111/njb.00498.

**Rai, Hardeep S.; Reeves, Patrick A.; Peakall, Rod; Olmstead, Richard G.; Graham, Sean W. (2008):**

**Inference of higher-order Conifer Relationships from a multi-locus Plastid Data Set.**

In: Botany 86 (7), p. 658–669. DOI: 10.1139/B08-062.

**Ran, Jin-Hua; Gao, Hui; Wang, Xiao-Quan (2010):**

**Fast Evolution of the retroprocessed mitochondrial RPS3 Gene in Conifer II and further Evidence for the Phylogeny of Gymnosperms.**

In: Molecular Phylogenetics and Evolution 54 (1), p. 136–149. DOI: 10.1016/j.ympev.2009.09.011.

**Wang, Xiao-Quan; Ran, Jin-Hua (2014):**

**Evolution and Biogeography of Gymnosperms.**

In: Molecular Phylogenetics and Evolution 75, p. 24–40. DOI: 10.1016/j.ympev.2014.02.005.

### Haemodoraceae

**Aerne-Hains, Layla; Simpson, Michael G. (2017):**

**Vegetative Anatomy of the Haemodoraceae and its phylogenetic Significance.**

In: International Journal of Plant Sciences 178 (2), p. 117–156. DOI: 10.1086/689199.

**Hopper, Stephen D.; Smith, Rhian J.; Fay, Michael F.; Manning, John C.; Chase, Mark W. (2009):**

**Molecular Phylogenetics of Haemodoraceae in the Greater Cape and Southwest Australian Floristic Regions.**

In: Molecular Phylogenetics and Evolution 51 (1), p. 19–30. DOI: 10.1016/j.ympev.2008.11.015.

### Haloragaceae

**Chen, Ling-Yun; Zhao, Shu-Ying; Mao, Kang-Shan; Les, Donald H.; Wang, Qing-Feng; Moody, Michael L. (2014):**

**Historical Biogeography of Haloragaceae: an out-of-Australia Hypothesis with multiple intercontinental Dispersals.**

In: Molecular Phylogenetics and Evolution 78, p. 87–95. DOI: 10.1016/j.ympev.2014.04.030.

**Moody, Michael L.; Les, Donald H. (2007):**

**Phylogenetic Systematics and Character Evolution in the Angiosperm Family Haloragaceae.**

In: American Journal of Botany 94 (12), p. 2005–2025.

**Moody, Michael L.; Les, Donald H. (2010):**

**Systematics of the aquatic Angiosperm Genus *Myriophyllum* (Haloragaceae).**

In: Systematic Botany 35 (1), p. 121–139. DOI: 10.1600/036364410790862470.

### Hamamelidaceae

**Li, Jian-Hua; Linn Bogle, A.; Klein, Anita p. (1999):**

**Phylogenetic Relationships of the Hamamelidaceae inferred from Sequences of Internal Transcribed Spacers (ITS) of Nuclear Ribosomal DNA.**

In: American Journal of Botany 86 (7), p. 1027–1037.

**Magallón, Susana (2007):**

**From Fossils to Molecules: Phylogeny and the Core Eudicot Floral Groundplan in Hamamelidoideae (Hamamelidaceae, Saxifragales).**

In: Systematic Botany 32 (2), p. 317–347.

**Qi, Zhe-Chen; Yu, Yi; Liu, Xiang; Pais, Andrew; Ranney, Thomas; Whetten, Ross; Xiang, Qiu-Yun Jenny (2015):**
**Phylogenomics of polyploid *Fothergilla* (Hamamelidaceae) by RAD-tag based GBS-Insights into Species Origin and effects of software pipelines.**
In: Journal of Systematics and Evolution 53 (5), p. 432–447. DOI: 10.1111/jse.12176.

**Xie, Lei; Yi, Ting-Shuang; Li, Rong; Li, De-Zhu; Wen, Jun (2010):**
**Evolution and biogeographic Diversification of the witch-hazel Genus (*Hamamelis* L., Hamamelidaceae) in the Northern Hemisphere.**
In: Molecular Phylogenetics and Evolution 56 (2), p. 675–689. DOI: 10.1016/j.ympev.2010.02.018.

### Hanguanaceae

**Leong-Škorničková, Jana; Niissalo, Matti A. (2017):**
**Identity and typification of *Hanguana malayana* and *H. anthelminthica* and Notes on other early Names in Hanguanaceae.**
In: Plant Systematics and Evolution 303 (9), p. 1213–1223. DOI: 10.1007/s00606-017-1432-9.

**Rahman, Siti Nurfazibt Abdul; Othman, Ahmad Sofiman; Boyce, Peter C. (2011):**
**Studies on *Hanguana* (Commelinales-Hanguanaceae) for Sunda I: *Hanguana bakoensis*, a new Forest Species from Sarawak, Malaysian Borneo, and Notes on Critical Morphologies for Elucidating *Hanguana* Taxonomy.**
In: Acta Phytotaxonomica Geobotanica 61 (3), p. 139–143.

### Heliconiaceae

**Iles, William J. D.; Sass, Chodon; Lagomarsino, Laura P.; Benson-Martin, Gracie; Driscoll, Heather E.; Specht, Chelsea D. (2017):**
**The Phylogeny of *Heliconia* (Heliconiaceae) and the Evolution of Floral presentation.**
In: Molecular Phylogenetics and Evolution 117, p. 150–167. DOI: 10.1016/j.ympev.2016.12.001.

**Isaza, L.; Marulanda, M. L.; López, A. M. (2012):**
**Genetic Diversity and molecular characterization of several *Heliconia* Species in Colombia.**
In: Genetics and molecular Research 11 (4), p. 4552–4563. DOI: 10.4238/2012.November.12.9.

**Marouelli, L. P.; Inglis, P. W.; Ferreira, M. A.; Buso, G. p. C. (2010):**
**Genetic Relationships among *Heliconia* (Heliconiaceae) Species based on RAPD Markers.**
In: Genetics and molecular Research 9 (3), p. 1377–1387. DOI: 10.4238/vol9-3gmr847.

### Heliotropiaceae

**Diane, Nadja; Förther, Harald; Hilger, Hartmut H. (2002):**
**A systematic Analysis of *Heliotropium, Tournefortia*, and allied Taxa of the Heliotropiaceae (Boraginales) based on ITS1 Sequences and morphological Data.**
In: American Journal of Botany 89 (2), p. 287–295. DOI: 10.3732/ajb.89.2.287.

**Feuillet, Christian (2016):**
**Two new Combinations in *Euploca* Nutt. (Heliotropiaceae, Boraginales) and a Conspectus of the Species of the Guiana Shield Area.**
In: PhytoKeys (61), p. 101–124. DOI: 10.3897/phytokeys.61.6260.

**Luebert, Federico; Brokamp, Grischa; Wen, Jun; Weigend, Maximilian; Hilger, Hartmut H. (2011):**
**Phylogenetic Relationships and morphological Diversity in Neotropical *Heliotropium* (Heliotropiaceae).**

In: Taxon 60 (3), p. 663–680.

**LUEBERT, FEDERICO; FROHLICH, MICHAEL W. (2016):**

**Four new Combinations in Argentinian Heliotropiaceae.**

In: Darwiniana, nueva serie 4 (2), p. 192–194. DOI: 10.14522/darwiniana.2016.42.717.

**LUEBERT, FEDERICO; HILGER, HARTMUT H.; WEIGEND, MAXIMILIAN (2011):**

**Diversification in the Andes: Age and Origins of South American *Heliotropium* Lineages (Heliotropiaceae, Boraginales).**

In: Molecular Phylogenetics and Evolution 61 (1), p. 90–102. DOI: 10.1016/j.ympev.2011.06.001.

**LUEBERT, FEDERICO; WEN, JUN (2008):**

**Phylogenetic Analysis and evolutionary Diversification of *Heliotropium* Sect. *Cochranea* (Heliotropiaceae) in the Atacama Desert.**

In: Systematic Botany 33 (2), p. 390–402. DOI: 10.1600/036364408784571635.

### Hemerocallidaceae

**FURNESS, CAROL A.; CONRAN, JOHN G.; GREGORY, THOMAS; RUDALL, PAULA J. (2013):**

**The trichotomosulcate asparagoids: Pollen Morphology of Hemerocallidaceae in Relation to Systematics and Pollination Biology.**

In: Australian Systematic Botany 26, p. 393–407.

**HUH, MAN KYU; KWON, OH SUNG; LEE, BYEONG RYONG (2013):**

**Phylogenetic Relationships of the Genus *Hemerocallis* in Korea Using rps16-trnK Sequences in Chloroplast DNA.**

In: Journal of Life Science 23 (7), p. 847–853. DOI: 10.5352/JLS.2013.23.7.847.

**WURDACK, KENNETH J.J.; DORR, LAURENCE J. (2009):**

**The South American Genera of Hemerocallidaceae (*Eccremis* and *Pasithea*): two introductions to the New World.**

In: Taxon 58 (4), p. 1122–1134. DOI: 10.1002/tax.584006.

### Herreriaceae

**LOPES, ROSANA CONRADO; ANDREATA, REGINA HELENA POTSCH (2006):**

**A new Species of *Clara* (Herreriaceae) from Brazil.**

In: Systematic Botany 31 (2), p. 298–301.

### Huerteales

**WORBERG, ANDREAS; ALFORD, MAC H.; QUANDT, DIETMAR; BORSCH, THOMAS (2009):**

**Huerteales sister to Brassicales plus Malvales, and newly circumscribed to include *Dipentodon, Gerrardina, Huertea, Perrottetia,* and *Tapiscia*.**

In: Taxon 58 (2), p. 468–478. DOI: 10.1002/tax.582012.

### Humiriaceae

**HERRERA, FABIANY; MANCHESTER, STEVEN R.; JARAMILLO, CARLOS; MACFADDEN, BRUCE; DA SILVA-CAMINHA, SILANE A. (2010):**

**Phytogeographic History and Phylogeny of the Humiriaceae.**

In: International Journal of Plant Sciences 171 (4), p. 392–408. DOI: 10.1086/651229.

### Hyacinthaceae

**ALI, SYED SHUJAIT; PFOSSER, MARTIN; WETSCHNIG, WOLFGANG; MARTÍNEZ-AZORÍN, MARIO; CRESPO, MANUEL B.; YU, YANG (2013):**

**Out of Africa: Miocene Dispersal, Vicariance, and Extinction within Hyacinthaceae Subfamily Urgineoideae.**
In: Journal of Integrative Plant Biology 55 (10), p. 950–964. DOI: 10.1111/jipb.12065.

ALI, SYED SHUJAIT; YU, YANG; PFOSSER, MARTIN; WETSCHNIG, WOLFGANG (2012):
**Inferences of biogeographical Histories within Subfamily Hyacinthoideae using S-diva and Bayesian binary MCMC Analysis implemented in RASP (Reconstruct Ancestral State in Phylogenies).**
In: Annals of Botany 109 (1), p. 95–107. DOI: 10.1093/aob/mcr274.

DESAI, NEETIN; KAWALKAR, HEMANT; DIXIT, GHANSHAM (2012):
**Biosystematics and evolutionary Studies in Indian *Drimia* Species.**
In: Journal of Systematics and Evolution 50 (6), p. 512–518. DOI: 10.1111/j.1759-6831.2012.00212.x.

GRUNDMANN, MICHAEL; RUMSEY, FRED J.; ANSELL, STEPHEN W.; RUSSELL, STEPHEN J.; DARWIN, SARAH C.; VOGEL, JOHANNES C. ET AL. (2010):
**Phylogeny and Taxonomy of the Bluebell Genus *Hyacinthoides*, Asparagaceae [Hyacinthaceae].**
In: Taxon 59 (1), p. 68–82. DOI: 10.1002/tax.591008.

KNIRSCH, WALTER; MARTÍNEZ-AZORÍN, MARIO; PFOSSER, MARTIN; WETSCHNIG, WOLFGANG (2015):
**The Reinstatement and Rediagnosis of the madagascan Genus *Rhodocodon* (Asparagaceae, Scilloideae), with validation and remarks on H. Perrier's Taxa.**
In: Phytotaxa 195 (2), p. 101. DOI: 10.11646/phytotaxa.195.2.1.

LEBATHA, PEARL; BUYS, MATT H.; STEDJE, BRITA (2006):
***Ledebouria, Resnova* and *Drimiopsis* : a Tale of three Genera.**
In: Taxon 55 (3), p. 643–652. DOI: 10.2307/25065640.

MANNING, JOHN C.; DEACON, JAMES; GOLDBLATT, PETER (2014):
**A Review of the *Schizobasis* Group of *Drimia* Jacq. (Hyacinthaceae: Urgineoideae), and the new Species *D. sigmoidea* from Western Cape, South Africa.**
In: South African Journal of Botany 94, p. 263–269. DOI: 10.1016/j.sajb.2014.07.011.

MANNING, JOHN C.; FOREST, FÉLIX; DEVEY, DION S.; FAY, MICHAEL F.; GOLDBLATT, PETER (2009):
**A molecular Phylogeny and a revised Classification of Ornithogaloideae (Hyacinthaceae) based on an Analysis of four Plastid DNA Regions.**
In: Taxon 58 (1), p. 77–107. DOI: 10.1002/tax.581011.

MANNING, JOHN C.; GOLDBLATT, PETER; FAY, MICHAEL F. (2003):
**A Revised Generic Synopsis of Hyacinthaceae in Sub-Saharan Africa, based on molecular Evidence, including new Combinations and the new Tribe Pseudoprospereae.**
In: Edinburgh Journal of Botany 60 (3). DOI: 10.1017/S0960428603000404.

MARTÍNEZ-AZORÍN, MARIO; CRESPO, MANUEL B.; DOLD, ANTHONY P.; WETSCHNIG, WOLFGANG; PINTER, MICHAEL; PFOSSER, MARTIN; VAN JAARSVELD, ERNST (2013):
***Sagittanthera* (Hyacinthaceae, Urgineoideae), a new Buzz pollinated Genus from the Eastern Cape Province of South Africa.**
In: Phytotaxa 98 (2), p. 43. DOI: 10.11646/phytotaxa.98.2.2.

MARTÍNEZ-AZORÍN, MARIO; CRESPO, MANUEL B.; JUAN, ANA; FAY, MICHAEL F. (2011):
**Molecular Phylogenetics of Subfamily Ornithogaloideae (Hyacinthaceae) based on nuclear and Plastid DNA Regions, including a new taxonomic Arrangement.**

In: Annals of Botany 107 (1), p. 1–37. DOI: 10.1093/aob/mcq207.

**Pfosser, Martin; Knirsch, Walter; Pinter, Michael; Ali, Syed Shujait; Dutta, Suchandra; Wetschnig, Wolfgang (2012):**
**Phylogenetic Relationships of Malagasy Hyacinthaceae.**
In: Plant Ecology and Evolution 145 (1), p. 65–72. DOI: 10.5091/plecevo.2012.590.

**Pfosser, Martin; Wetschnig, Wolfgang; Ungar, S.; Prenner, Gerhard (2003):**
**Phylogenetic Relationships among Genera of Massonieae (Hyacinthaceae) inferred from Plastid DNA and Seed Morphology.**
In: Journal of Plant Research 116 (2), p. 115–132. DOI: 10.1007/s10265-003-0076-8.

**Pinter, Michael; Crespo, Manuel B.; Ilg, Irmgard; Luidold, Anna K.; Martínez-Azorín, Mario; Müller-Doblies, Ute et al. (2013):**
**Mucinaea (Hyacinthaceae-Urgineoideae), a remarkable new Genus from Namaqualand (Northern Cape Province, South Africa).**
In: Phyton (Horn, Austria) 53 (2), p. 289–304.

**Stedje, Brita (1998):**
**Phylogenetic Relationships and generic Delimitation of Sub-Saharan *Scilla* (Hyacinthaceae) and allied African Genera as inferred from morphological and DNA Sequence Data.**
In: Plant Systematics and Evolution 211 (1-2), p. 1–11. DOI: 10.1007/BF00984908.

**Suárez-Santiago, Víctor N.; Salinas, María J.; Romero-García, Ana T.; Garrido-Ramos, Manuel A.; La Herrán, Roberto; Ruiz-Rejón, Carmelo et al. (2007):**
**Polyploidy, the major Speciation mechanism in *Muscari* Subgenus *Botryanthus* in the Iberian Peninsula.**
In: Taxon 56 (4), p. 1171–1184. DOI: 10.2307/25065910.

**Wetschnig, Wolfgang; Knirsch, Walter; Ali, Syed Shujait; Pfosser, Martin (2007):**
**Systematic Position of three little known an frequently misplaced Species of Hyacinthaceae from Madagascar.**
In: Phyton (Horn, Austria) 47 (1-2), p. 321–337.

**Wetschnig, Wolfgang; Pfosser, Martin (2003):**
**The *Scilla plumbea* puzzle-present Status of the Genus *Scilla* sensu lato in southern Africa and Description of *Spetaea lachenaliiflora*, a new Genus and Species of Massonieae (Hyacinthaceae).**
In: Taxon 52 (1), p. 75–92. DOI: 10.2307/3647303.

**Yildirim, Hasan; Gemící, Yusuf; Wilkin, Paul (2013):**
***Scilla vardaria* (Asparagaceae Subfamily Scilloideae): a threatened new Species of *Scilla* L. from Northeast Turkey with a Floral Corona.**
In: Phytotaxa 91 (2), p. 50. DOI: 10.11646/phytotaxa.91.2.3.

### Hydatellaceae

**Iles, William J. D.; Rudall, Paula J.; Sokoloff, Dmitry D.; Remizowa, Margarita V.; Macfarlane, Terry D.; Logacheva, Maria D.; Graham, Sean W. (2012):**
**Molecular Phylogenetics of Hydatellaceae (Nymphaeales): Sexual-system Homoplasy and a new Sectional Classification.**
In: American Journal of Botany 99 (4), p. 663–676. DOI: 10.3732/ajb.1100524.

**Saarela, Jeffery M.; Rai, Hardeep S.; Doyle, James A.; Endress, Peter K.; Mathews, Sarah; Marchant, Adam D. et al. (2007):**

**Hydatellaceae identified as a new Branch near the Base of the Angiosperm phylogenetic Tree.**
In: Nature 446 (7133), p. 312–315. DOI: 10.1038/nature05612.

SOKOLOFF, DMITRY D.; REMIZOWA, MARGARITA V.; MACFARLANE, TERRY D.; RUDALL, PAULA J. (2008):
**Classification of the early-divergent Angiosperm Family Hydatellaceae: one Genus instead of two, four new Species and sexual Dimorphism in dioecious Taxa.**
In: Taxon 57 (1), p. 179–200.

### Hydnoraceae

NICKRENT, DANIEL LEE; BLARER, ALBERT; QIU, YIN-LONG; SOLTIS, DOUGLAS E.; SOLTIS, PAMELA S.; ZANIS, MICHAEL J. (2002):
**Molecular Data place Hydnoraceae with Aristolochiaceae.**
In: American Journal of Botany 89 (11), p. 1809–1817. DOI: 10.3732/ajb.89.11.1809.

### Hydrangeaceae

GUO, YUE-LONG; PAIS, ANDREW; WEAKLEY, ALAN S.; XIANG, QIU-YUN JENNY (2013):
**Molecular phylogenetic Analysis suggests Paraphyly and early Diversification of *Philadelphus* (Hydrangeaceae) in western North America: new Insights into Affinity with *Carpenteria*.**
In: Journal of Systematics and Evolution 51 (5), p. 545–563. DOI: 10.1111/jse.12041.

KIM, CHANGKYUN; DENG, TAO; WEN, JUN; NIE, ZE-LONG; SUN, HANG (2015):
**Systematics, Biogeography, and Character Evolution of *Deutzia* (Hydrangeaceae) inferred from nuclear and Chloroplast DNA Sequences.**
In: Molecular Phylogenetics and Evolution 87, p. 91–104. DOI: 10.1016/j.ympev.2015.03.002.

SMET, YANNICK; CLERCK, OLIVIER; UEMACHI, TATSUYA; GRANADOS MENDOZA, CAROLINA; WANKE, STEFAN; GOETGHEBEUR, PAUL; SAMAIN, MARIE-STÉPHANIE (2017):
**Multilocus coalescent Species Delimitation to evaluate traditionally defined Morphotypes in *Hydrangea* Sect. *Asperae* (Hydrangeaceae).**
In: Molecular Phylogenetics and Evolution 114, p. 415–425. DOI: 10.1016/j.ympev.2017.05.021.

SMET, YANNICK; GRANADOS MENDOZA, CAROLINA; WANKE, STEFAN; GOETGHEBEUR, PAUL; SAMAIN, MARIE-STÉPHANIE (2015):
**Molecular Phylogenetics and new (infra)generic Classification to alleviate Polyphyly in Tribe Hydrangeeae (Cornales: Hydrangeaceae).**
In: Taxon 64 (4), p. 741–753. DOI: 10.12705/644.6.

### Hydrocharitaceae

CHEN, LING-YUN; CHEN, JIN-MING; GITURU, ROBERT WAHITI; WANG, QING-FENG (2012):
**Generic Phylogeny, historical Biogeography and Character Evolution of the cosmopolitan aquatic Plant Family Hydrocharitaceae.**
In: BMC Evolutionary Biology 12, p. 30. DOI: 10.1186/1471-2148-12-30.

LES, DONALD H.; JACOBS, SURREY W.L.; TIPPERY, NICHOLAS P.; CHEN, LEI; MOODY, MICHAEL L.; WILSTERMANN-HILDEBRAND, MAIKE (2008):
**Systematics of *Vallisneria* (Hydrocharitaceae).**
In: Systematic Botany 33 (1), p. 49–65. DOI: 10.1600/036364408783887483.

LES, DONALD H.; SHELDON, SALLIE P.; TIPPERY, NICHOLAS P. (2010):
**Hybridization in Hydrophiles: Natural Interspecific Hybrids in *Najas* (Hydrocharitaceae).**

In: Systematic Botany 35 (4), p. 736–744. DOI: 10.1600/036364410X539826.

**Waycott, Michelle; Freshwater, D. Wilson; York, Robert A.; Calladine, Ainsley; Kenworthy, W. Judson (2002):**

**Evolutionary Trends in the Seagrass Genus *Halophila* (Thourars): Insights from molecular Phylogeny.**

In: Bulletin of Marine Science 71 (3), p. 1299–1308.

### Hydroleaceae

**Erbar, C.; Porembski, Stefan; Leins, Peter (2005):**

**Contributions to the systematic Position of *Hydrolea* (Hydroleaceae) based on Floral development.**

In: Plant Systematics and Evolution 252 (1-2), p. 71–83. DOI: 10.1007/s00606-004-0263-7.

### Hydrophyllaceae

**Barr, Camille M. (2014):**

**Phylogenetic Relationships and crossing Data reveal a new Species of *Nemophila* (Boraginaceae).**

In: Madroño 61 (3), p. 308–315. DOI: 10.3120/0024-9637-61.3.308.

**Ferguson, Diane M. (1999):**

**Phylogenetic Analysis and Relationships in Hydrophyllaceae based on ndhF Sequence Data.**

In: Systematic Botany 23 (3), p. 253–268.

**Gilbert, Cynthia; Dempcy, John; Ganong, Constance; Patterson, Robert W.; Spicer, Greg p. (2005):**

**Phylogenetic Relationships within *Phacelia* Subgenus *Phacelia* (Hydrophyllaceae) inferred from Nuclear rDNA ITS Sequence Data.**

In: Systematic Botany 30 (3), p. 627–634.

**Hansen, Debra R.; Spicer, Greg S.; Patterson, Robert W. (2009):**

**Phylogenetic Relationships between and within *Phacelia* Sections *Whitlavia* and *Gymnobythus* (Boraginaceae).**

In: Systematic Botany 34 (4), p. 737–746. DOI: 10.1600/036364409790139637.

### Hydrostachyaceae

**Fan, Chuanzhu; Xiang, Qiu-Yun (2003):**

**Phylogenetic Analyses of Cornales based on 26s rRNA and combined 26s rDNA-MATK-RBCL Sequence Data.**

In: American Journal of Botany 90 (9), p. 1357–1372. DOI: 10.3732/ajb.90.9.1357.

### Hymenophyllaceae

**Bauret, Lucie; Grall, Aurelie; Senterre, Bruno; Rouhan, Germinal; Hennequin, Sabine; Ebihara, Atsushi; Dubuisson, Jean-Yves (2015):**

**New Circumscription of *Trichomanes cupressoides* Desvaux (Hymenophyllaceae), an endemic filmy Fern from the Seychelles (Indian Ocean), and new Insights into the Genus *Abrodictyum* C.Presl in the Western Indian Ocean.**

In: Phytotaxa 202 (1), p. 1. DOI: 10.11646/phytotaxa.202.1.1.

**Dubuisson, Jean-Yves; Hébant-Mauri, R.; Galtier, J. (1998):**

**Molecules and Morphology: Conflicts and Congruence within the Fern Genus *Trichomanes* (Hymenophyllaceae).**

In: Molecular Phylogenetics and Evolution 9 (3), p. 390–397.

**Dubuisson, Jean-Yves; Hennequin, Sabine; Douzery, Emmanuel J. P.; Cranfill, Raymond B.; Smith, Alan R.; Pryer, Kathleen M. (2003):**
**RbcL Phylogeny of the Fern Genus *Trichomanes* (Hymenophyllaceae), with Special Reference to Neotropical Taxa.**
In: International Journal of Plant Sciences 164 (5), p. 753–761.

**Dubuisson, Jean-Yves; Rouhan, Germinal; Grall, Aurelie; Hennequin, Sabine; Senterre, Bruno; Pynee, K.; Ebihara, Atsushi (2013):**
**New Insights into the Systematics and Evolution of the filmy Fern Genus *Crepidomanes* (Hymenophyllaceae) in the Mascarene Archipelago with a Focus on dwarf Species.**
In: Acta Botanica Gallica 160 (2), p. 173–194. DOI: 10.1080/12538078.2013.819294.

**Ebihara, Atsushi; Dubuisson, Jean-Yves; Iwatsuki, Kunio; Ito, Motomi (2007):**
**Systematics of *Trichomanes* (Hymenophyllaceae: Pteridophyta), Progress and Future Interests.**
In: the Fern Gazette 18 (2), p. 53–58.

**Ebihara, Atsushi; Ishikawa, Hiroshi; Matsumoto, Sadamu; Lin, Su-Juan; Iwatsuki, Kunio; Takamiya, Masayuki et al. (2005):**
**Nuclear DNA, Chloroplast DNA, and Ploidy Analysis clarified biological Complexity of the *Vandenboschia radicans* Complex (Hymenophyllaceae) in Japan and adjacent Areas.**
In: American Journal of Botany 92 (9), p. 1535–1547.

**Ebihara, Atsushi; Iwatsuki, Kunio; Ito, Motomi; Hennequin, Sabine; Dubuisson, Jean-Yves (2007):**
**A global molecular Phylogeny of the Fern Genus *Trichomanes* (Hymenophyllaceae) with special Reference to stem Anatomy.**
In: Botanical Journal of the Linnean Society 155, p. 1–27.

**Hennequin, Sabine (2003):**
**Phylogenetic Relationships within the Fern Genus *Hymenophyllum* s.l. (Hymenophyllaceae, Filicopsida): Contribution of Morphology and Cytology.**
In: Comptes Rendus Biologies 326 (6), p. 599–611. DOI: 10.1016/S1631-0691(03)00146-X.

**Hennequin, Sabine; Ebihara, Atsushi; Ito, Motomi; Iwatsuki, Kunio; Dubuisson, Jean-Yves (2003):**
**Molecular Systematics of the Fern Genus *Hymenophyllum* s.l. (Hymenophyllaceae) based on chloroplastic Coding and Noncoding Regions.**
In: Molecular Phylogenetics and Evolution 27 (2), p. 283–301. DOI: 10.1016/S1055-7903(02)00404-9.

**Hennequin, Sabine; Ebihara, Atsushi; Ito, Motomi; Iwatsuki, Kunio; Dubuisson, Jean-Yves (2006):**
**New Insights into the Phylogeny of the Genus *Hymenophyllum* s.l. (Hymenophyllaceae): Revealing the Polyphyly of *Mecodium*.**
In: Systematic Botany 31 (2), p. 271–284.

**Hennequin, Sabine; Schuettpelz, Eric; Pryer, Kathleen M.; Ebihara, Atsushi; Dubuisson, Jean-Yves (2008):**
**Divergence Times and the Evolution of Epiphytism in Filmy Ferns (Hymenophyllaceae) revisited.**
In: International Journal of Plant Sciences 169 (9), p. 1278–1287. DOI: 10.1086/591983.

**Nitta, Joel H. (2008):**
**Exploring the Utility of three Plastid Loci for Biocoding the filmy Ferns (Hymenophyllaceae) of *Moorea*.**
In: Taxon 57 (3), p. 725–736.

SCHUETTPELZ, ERIC; PRYER, KATHLEEN M. (2006):
**Reconciling extreme Branch Length Differences: decoupling Time and Rate through the evolutionary History of filmy Ferns.**
In: Systematic Biology 55 (3), p. 485–502. DOI: 10.1080/10635150600755438.

### Hypericaceae

MESEGUER, ANDREA SÁNCHEZ; ALDASORO, JUAN JOSÉ; SANMARTÍN, ISABEL (2013):
**Bayesian Inference of Phylogeny, Morphology and Range Evolution reveals a complex evolutionary History in St. John's wort (*Hypericum*).**
In: Molecular Phylogenetics and Evolution 67 (2), p. 379–403. DOI: 10.1016/j.ympev.2013.02.007.

MESEGUER, ANDREA SÁNCHEZ; SANMARTÍN, ISABEL; MARCUSSEN, THOMAS; PFEIL, BERNARD E. (2014):
**Utility of low-copy nuclear Markers in phylogenetic Reconstruction of *Hypericum* L. (Hypericaceae).**
In: Plant Systematics and Evolution 300 (6), p. 1503–1514. DOI: 10.1007/s00606-013-0977-5.

NÜRK, NICOLAI M.; BLATTNER, FRANK R. (2010):
**Cladistic Analysis of morphological Characters in *Hypericum* (Hypericaceae).**
In: Taxon 59 (5), p. 1495–1507. DOI: 10.1002/tax.595014.

NÜRK, NICOLAI M.; MADRIÑÁN, SANTIAGO; CARINE, MARK A.; CHASE, MARK W.; BLATTNER, FRANK R. (2013):
**Molecular Phylogenetics and morphological Evolution of St. John's wort (*Hypericum*; Hypericaceae).**
In: Molecular Phylogenetics and Evolution 66 (1), p. 1–16. DOI: 10.1016/j.ympev.2012.08.022.

ROBSON, NORMAN K. B. (2010):
**Studies in the Genus *Hypericum* L (Hypericaceae) 5(2). Sections 17. *Hirtella* to 19. *Coridium*.**
In: Phytotaxa 4, p. 127–258.

ROBSON, NORMAN K. B. (2010):
**Studies in the Genus *Hypericum* L. (Hypericaceae) 5(1). Sections 10. *Olympia* to 15/16. Crossophyllum.**
In: Phytotaxa 4, p. 5–126.

### Hypoxidaceae

KOCYAN, ALEXANDER; SNIJMAN, DEIRDRE A.; FOREST, FÉLIX; DEVEY, DION S.; FREUDENSTEIN, JOHN V.; WILAND-SZYMAŃSKA, JUSTYNA ET AL. (2011):
**Molecular Phylogenetics of Hypoxidaceae - Evidence from Plastid DNA Data and Inferences on Morphology and Biogeography.**
In: Molecular Phylogenetics and Evolution 60 (1), p. 122–136. DOI: 10.1016/j.ympev.2011.02.021.

KOCYAN, ALEXANDER; WILAND-SZYMAŃSKA, JUSTYNA (2016):
***Friedmannia*: a new Genus from the Seychelles and the beginning of a generic realignment of *Curculigo* (Hypoxidaceae).**
In: Phytotaxa 283 (1), p. 54. DOI: 10.11646/phytotaxa.283.1.3.

LIU, KE-WEI; XIE, GAO-CHANG; CHEN, LI-JUN; XIAO, XIN-JU; ZHENG, YU-YUN; CAI, JING ET AL. (2012):
***Sinocurculigo*, a new Genus of Hypoxidaceae from China based on molecular and morphological Evidence.**
In: Public Library of Science One 7 (6), e38880. DOI: 10.1371/journal.pone.0038880.

SNIJMAN, DEIRDRE A. (2014):
**A taxonomic Revision of the Genus *Pauridia* (Hypoxidaceae) in southern Africa.**

In: Phytotaxa 182 (1), p. 1. DOI: 10.11646/phytotaxa.182.1.1.

### Icacinaceae

**Byng, James W.; Bernardini, Benedetta; Joseph, Jeffrey A.; Chase, Mark W.; Utteridge, Timothy M.A. (2014):**
**Phylogenetic Relationships of Icacinaceae focusing on the vining Genera.**
In: Botanical Journal of the Linnean Society 176 (3), p. 277–294. DOI: 10.1111/boj.12205.

**Duno de Stefano, Rodrigo; Fernández-Concha, Germán Carnevali (2011):**
**Morphology-inferred Phylogeny and a Revision of the Genus *Emmotum* (Icacinaceae) 1.**
In: Annals of the Missouri Botanical Garden 98 (1), p. 1–27. DOI: 10.3417/2007129.

**Kårehed, Jesper (2001):**
**Multiple Origin of the tropical Forest Tree Family Icacinaceae.**
In: American Journal of Botany 88 (12), p. 2259–2274. DOI: 10.2307/3558388.

**Lens, Frederic; Kårehed, Jesper; Baas, Pieter; Jansen, Steven; Rabaey, David; Huysmans, Suzy et al. (2008):**
**The wood Anatomy of the polyphyletic Icacinaceae s.l., and their Relationships within Asterids.**
In: Taxon 57 (2), p. 525–552.

**Stull, Gregory W.; Duno de Stefano, Rodrigo; Soltis, Douglas E.; Soltis, Pamela p. (2015):**
**Resolving basal Lamiid Phylogeny and the Circumscription of Icacinaceae with a plastome-scale Data Set.**
In: American Journal of Botany 102 (11), p. 1794–1813. DOI: 10.3732/ajb.1500298.

### Iridaceae

**Alves, Tiago L.S.; Chauveau, Olivier; Eggers, Lilian; Souza-Chies, Tatiana Teixeira de (2014):**
**Species discrimination in *Sisyrinchium* (Iridaceae): assessment of DNA Barcodes in a Taxonomically challenging Genus.**
In: molecular Ecology Resources 14 (2), p. 324–335. DOI: 10.1111/1755-0998.12182.

**Chauveau, Olivier; Eggers, Lilian; Raquin, Christian; Silvério, Adriano; Brown, Spencer C.; Couloux, Arnaud et al. (2011):**
**Evolution of oil-producing Trichomes in *Sisyrinchium* (Iridaceae): Insights from the first comprehensive phylogenetic Analysis of the Genus.**
In: Annals of Botany 107 (8), p. 1287–1312. DOI: 10.1093/aob/mcr080.

**Coşkun, Fatih; Selvİ, Selami; Satil, Fatih (2010):**
**Phylogenetic Relationships of some Turkish *Crocus* (Iridaceae) Taxa based on morphological and anatomical Characters.**
In: Turkish Journal of Botany 34, p. 171–178.

**Crespo, Manuel B.; Martínez-Azorín, Mario; Mavrodiev, Evgeny V. (2015):**
**Can a Rainbow consist of a single colour? A new comprehensive generic Arrangement of the '*Iris* sensu latissimo' Clade (Iridaceae), congruent with Morphology and molecular Data.**
In: Phytotaxa 232 (1), p. 1. DOI: 10.11646/phytotaxa.232.1.1.

**Erol, Osman; Harpke, Dörte; Yildirim, Hasan (2015):**
**A new *Crocus* L. (Iridaceae) Species from SE-Turkey, based on morphological and molecular Data.**
In: Phytotaxa 239 (3), p. 223. DOI: 10.11646/phytotaxa.239.3.3.

**Goldblatt, Peter (2002):**

**Radiation in the Cape Flora and the Phylogeny of peacock Irises *Moraea* (Iridaceae) based on four Plastid DNA Regions.**
In: Molecular Phylogenetics and Evolution 25 (2), p. 341–360. DOI: 10.1016/S1055-7903(02)00235-X.

**Goldblatt, Peter; Bernhardt, Peter; Manning, John C. (2009):**
**Adaptive Radiation of the putrid Perianth: *Ferraria* (Iridaceae: Irideae) and its unusual Pollinators.**
In: Plant Systematics and Evolution 278 (1-2), p. 53–65. DOI: 10.1007/s00606-008-0132-x.

**Goldblatt, Peter; Le Thomas, Annick; Suárez-Cervera, Maria (2004):**
**Phylogeny of the Afro-Madagascan *Aristea* (Iridaceae) revisited in the Light of new Data on Pollen Morphology.**
In: Botanical Journal of the Linnean Society 144 (1), p. 41–68. DOI: 10.1111/j.0024-4074.2004.00246.x.

**Goldblatt, Peter; Manning, John C. (2011):**
**Systematics and biology of the African Genus *Ferraria* (Iridaceae: Irideae).**
In: Bothalia 41 (1), p. 1–40. DOI: 10.4102/abc.v41i1.33.

**Goldblatt, Peter; Manning, John C. (2012):**
**Systematics of the southern African Genus *Ixia* (Iridaceae: Crocoideae): 4. Revision of Sect.** *Dichone*.
In: Bothalia 42 (2), p. 87–110. DOI: 10.4102/abc.v42i2.11.

**Goldblatt, Peter; Manning, John C.; Munzinger, Jérôme; Lowry, Porter Prescott (2011):**
**A new native Family and new endemic Species for the Flora of New Caledonia: *Patersonia neocaledonica* sp. nov. (Iridaceae, Patersonioideae), from the Mount Humboldt Massif.**
In: Adansonia 33 (2), p. 201–208. DOI: 10.5252/a2011n2a4.

**Goldblatt, Peter; Manning, John C.; Schnitzler, Jan (2013):**
**A revised infrageneric Classification and Synopsis of the Afro-Eurasian Genus *Moraea* (Iridaceae: Irideae).**
In: Bothalia 43 (1), p. 29–41. DOI: 10.4102/abc.v43i1.84.

**Goldblatt, Peter; Rodriguez, Aaron; Powell, M. P.; Davies, Jonathan T.; Manning, John C.; van der Bank, Michelle; Savolainen, Vincent (2008):**
**Iridaceae 'Out of Australasia'? Phylogeny, Biogeography, and Divergence Time based on Plastid DNA Sequences.**
In: Systematic Botany 33 (3), p. 495–508. DOI: 10.1600/036364408785679806.

**Harpke, Dörte; Carta, Angelino; Tomović, Gordana; Randelović, Vladimir; Randelović, Novica; Blattner, Frank R.; Peruzzi, Lorenzo (2015):**
**Phylogeny, karyotype Evolution and Taxonomy of *Crocus* series *Verni* (Iridaceae).**
In: Plant Systematics and Evolution 301 (1), p. 309–325. DOI: 10.1007/s00606-014-1074-0.

**Harpke, Dörte; Meng, Shuchun; Rutten, Twan; Kerndorff, Helmut; Blattner, Frank R. (2013):**
**Phylogeny of *Crocus* (Iridaceae) based on one Chloroplast and two nuclear Loci: ancient Hybridization and Chromosome Number Evolution.**
In: Molecular Phylogenetics and Evolution 66 (3), p. 617–627. DOI: 10.1016/j.ympev.2012.10.007.

**Harpke, Dörte; Peruzzi, Lorenzo; Kerndorff, Helmut; Karamplianis, Theophanis; Constantinidis, Theophanis; Randelović, Vladimir et al. (2014):**
**Phylogeny, geographic Distribution, and new taxonomic Circumscription of the *Crocus reticulatus* Species Group (Iridaceae).**
In: Turkish Journal of Botany 38, p. 1182–1198. DOI: 10.3906/bot-1405-60.

**Ikinci, Nursel; Hall, Tony; Lledó, M. Dolores; Clarkson, James J.; Tillie, Nico; Seisums, Arnis et al. (2011):**
**Molecular Phylogenetics of the Juno Irises, *Iris* Subgenus *Scorpiris* (Iridaceae), based on six Plastid Markers.**
In: Botanical Journal of the Linnean Society 167 (3), p. 281–300. DOI: 10.1111/j.1095-8339.2011.01176.x.

**Kerndorff, Helmut; Pasche, Erich; Blattner, Frank R.; Harpke, Dörte (2013):**
**Fourteen new Species of *Crocus* (Liliiflorae, Iridaceae) from West, South-West and South-Central Turkey.**
In: Stapfia 99, p. 145–158.

**Kozyrenko, Marina M.; Artyukova, E. V.; Zhuravlev, Yu. N. (2009):**
**Independent Species Status of *Iris vorobievii* N.S.Pavlova, *Iris mandshurica* Maxim., and *Iris humilis* Georgi (Iridaceae): Evidence from the nuclear and Chloroplast Genomes.**
In: Russian Journal of Genetics 45 (11), p. 1394–1402. DOI: 10.1134/S1022795409110143.

**Lee, HyunJung; Park, Seon-Joo (2013):**
**A phylogenetic Study of Korean *Iris* L. based on Plastid DNA (psbA-trnH, trnL-F) Sequences.**
In: Korean Journal of Plant Taxonomy 43 (3), p. 227–235. DOI: 10.11110/kjpt.2013.43.3.227.

**Lovo, Juliana; Winkworth, Richard C.; Mello-Silva, Renato (2012):**
**New Insights into Trimezieae (Iridaceae) Phylogeny: what do molecular Data tell us?**
In: Annals of Botany 110 (3), p. 689–702. DOI: 10.1093/aob/mcs127.

**Makarevitch, Irina; Golovnina, Kseniya; Scherbik, Svetlana; Blinov, Alexander G. (2003):**
**Phylogenetic Relationships of the Siberian *Iris* Species inferred from Noncoding Chloroplast DNA Sequences.**
In: International Journal of Plant Sciences 164 (2), p. 229–237. DOI: 10.1086/346160.

**Manning, John C.; Forest, Félix; Vinnersten, Annika (2007):**
**The Genus *Colchicum* L. redefined to include *Androcymbium* Willd. based on molecular Evidence.**
In: Taxon 56 (3), p. 872–882. DOI: 10.2307/25065868.

**Manning, John C.; Goldblatt, Peter (2001):**
**A synoptic Review of *Romulea* (Iridaceae: Crocoideae) in Sub-Saharan Africa, the Arabian Peninsula and Socotra including new Species, biological Notes, and a new infrageneric Classification.**
In: Adansonia 23 (1), p. 59–108.

**Manning, John C.; Goldblatt, Peter (2011):**
**Taxonomic Revision of the Genus *Thereianthus* (Iridaceae: Crocoideae).**
In: Bothalia 41 (2), p. 239–267. DOI: 10.4102/abc.v41i2.56.

**Martínez, Jorge; Vargas, Pablo; Luceño, Modesto; Cuadrado, Ángeles (2010):**
**Evolution of *Iris* Subgenus *Xiphium* based on Chromosome Numbers, Fish of nrDNA (5s, 45s) and trnL–trnF Sequence Analysis.**
In: Plant Systematics and Evolution 289 (3-4), p. 223–235. DOI: 10.1007/s00606-010-0345-7.

**Mavrodiev, Evgeny V.; Martínez-Azorín, Mario; Dranishnikov, Peter; Crespo, Manuel B. (2014):**
**At least 23 Genera instead of one: the case of *Iris* L. s.l. (Iridaceae).**
In: Public Library of Science One 9 (8), e106459. DOI: 10.1371/journal.pone.0106459.

**Moraes, Ana Paula; Souza-Chies, Tatiana Teixeira de; Stiehl-Alves, Eudes M.; Burchardt, Paula; Eggers, Lilian; Siljak-Yakovlev, Sonja et al. (2015):**
**Evolutionary Trends in Iridaceae: new cytogenetic Findings from the New World.**
In: Botanical Journal of the Linnean Society 177, p. 27–49.

**Petersen, Gitte; Seberg, Ole; Thorsøe, Sarah; Jørgensen, Tina; Mathew, Brian (2008):**
**A Phylogeny of the Genus *Crocus* (Iridaceae) based on Sequence Data from five Plastid Regions.**
In: Taxon 57 (2), p. 487–499.

**Reeves, Gail; Chase, Mark W.; Goldblatt, Peter; Rudall, Paula J.; Fay, Michael F.; Cox, Antony V. et al. (2001):**
**Molecular Systematics of Iridaceae: Evidence from four Plastid DNA Regions.**
In: American Journal of Botany 88 (11), p. 2074–2087. DOI: 10.2307/3558433.

**Reeves, Gail; Goldblatt, Peter; Rudall, Paula J.; Chase, Mark W. (2000):**
**Molecular Systematics of Iridaceae: a combined Analysis of four Plastid DNA Sequence Matrices.**
In: Annali di Botanica 58, p. 29–42.

**Rymer, Paul D.; Manning, John C.; Goldblatt, Peter; Powell, Martyn P.; Savolainen, Vincent (2010):**
**Evidence of recent and continuous Speciation in a biodiversity Hotspot: a Population genetic approach in Southern African Gladioli (*Gladiolus*; Iridaceae).**
In: molecular Ecology 19 (21), p. 4765–4782. DOI: 10.1111/j.1365-294X.2010.04794.x.

**Souza-Chies, Tatiana Teixeira de; Santos, Eliane Kaltchuk dos; Eggers, Lilian; Flores, Alice Mainieri; Alves, Eudes M. Stiehl; Fachinetto, Juliana et al. (2012):**
**Studies on Diversity and Evolution of Iridaceae Species in Southern Brazil.**
In: Genetics and Molecular Biology 35 (4 suppl 1), p. 1027–1035. DOI: 10.1590/S1415-47572012000600018.

**Valente, Luis M.; Savolainen, Vincent; Manning, John C.; Goldblatt, Peter; Vargas, Pablo (2011):**
**Explaining disparities in Species Richness between Mediterranean floristic Regions: a case Study in *Gladiolus* (Iridaceae).**
In: Global Ecology and Biogeography 20 (6), p. 881–892. DOI: 10.1111/j.1466-8238.2010.00644.x.

**Wilson, Carol A. (2003):**
**Phylogenetic Relationships in *Iris* Series *Californicae* based on ITS Sequences of Nuclear Ribosomal DNA.**
In: Systematic Botany 28 (1), p. 39–46.

**Wilson, Carol A. (2009):**
**Phylogenetic Relationships among the recognized Series in *Iris* Section *Limniris*.**
In: Systematic Botany 34 (2), p. 277–284. DOI: 10.1600/036364409788606316.

**Wilson, Carol A. (2011):**
**Subgeneric Classification in *Iris* re-examined using Chloroplast Sequence Data.**
In: Taxon 60 (1), p. 27–35. DOI: 10.1002/tax.601004.

**Wilson, Carol A.; Padiernos, Justin; Sapir, Yuval (2016):**
**The royal Irises (*Iris* Subg. *Iris* Sect. *Oncocyclus*): Plastid and low-copy nuclear Data contribute to an Understanding of their phylogenetic Relationships.**
In: Taxon 65 (1), p. 35–46. DOI: 10.12705/651.3.

**Young, Nelson D. (1998):**
**Pacific Coast *Iris* Species Delimitation using three Species Definitions: biological, phylogenetic and genealogical.**

In: Botanical Journal of the Linnean Society 63 (1), p. 99–120. DOI: 10.1111/j.1095-8312.1998.tb01641.x.

### Iridales

**Seberg, Ole; Petersen, Gitte; Davis, Jerrold I.; Pires, J. Chris; Stevenson, Dennis Wm.; Chase, Mark W. et al. (2012):**
**Phylogeny of the Asparagales based on three Plastid and two mitochondrial Genes.**
In: American Journal of Botany 99 (5), p. 875–889. DOI: 10.3732/ajb.1100468.

### Isoetaceae

**Hoot, Sara B.; Napier, Nancy S.; Taylor, W. Carl (2004):**
**Revealing unknown or extinct Lineages within *Isoetes* (Isoetaceae) using DNA Sequences from Hybrids.**
In: American Journal of Botany 91 (6), p. 899–904.

**Hoot, Sara B.; Taylor, W. Carl; Napier, Nancy p. (2006):**
**Phylogeny and Biogeography of *Isoetes* (Isoetaceae) based on Nuclear and Chloroplast DNA Sequence Data.**
In: Systematic Botany 31 (3), p. 449–460.

**Kim, Changkyun; Shin, Hyunchur; Chang, Yung-Ta; Choi, Hong-Keun (2010):**
**Speciation pathway of *Isoetes* (Isoetaceae) in East Asia inferred from molecular phylogenetic Relationships.**
In: American Journal of Botany 97 (6), p. 958–969. DOI: 10.3732/ajb.0900162.

**Larsén, Eva; Rydin, Catarina (2016):**
**Disentangling the Phylogeny of *Isoetes* (Isoetales), using nuclear and Plastid Data.**
In: International Journal of Plant Sciences 177 (2), p. 157–174. DOI: 10.1086/684179.

**Rydin, Catarina; Wikström, Niklas (2002):**
**Phylogeny of *Isoetes* (Lycopsida): resolving basal Relationships using rbcL Sequences.**
In: Taxon 51, p. 83–89.

**Troia, Angelo; Pereira, Jovani B.; Kim, Changkyun; Taylor, W. Carl (2016):**
**The Genus *Isoetes* (Isoetaceae): a provisional Checklist of the accepted and unresolved Taxa.**
In: Phytotaxa 277 (2), p. 101. DOI: 10.11646/phytotaxa.277.2.1.

### Ixioliriaceae

**Zhao, Y.; Li, Y.; Liu, Y.; Yang, Y. F. (2015):**
**DNA barcoding for efficient Identification of *Ixiolirion* Species (Ixioliriaceae).**
In: Genetics and molecular Research 14 (1), p. 1903–1910. DOI: 10.4238/2015.March.13.19.

### Juglandaceae

**Manos, Paul S.; Soltis, Pamela S.; Soltis, Douglas E.; Manchester, Steven R.; Oh, Sang-Hun; Bell, Charles D. et al. (2007):**
**Phylogeny of extant and fossil Juglandaceae inferred from the Integration of molecular and morphological Data Sets.**
In: Systematic Biology 56 (3), p. 412–430. DOI: 10.1080/10635150701408523.

**Manos, Paul S.; Stone, Donald E. (2001):**
**Evolution, Phylogeny, and Systematics of the Juglandaceae.**
In: Annals of the Missouri Botanical Garden 88, p. 231–269.

**Stanford, Alice M.; Harden, Rachel; Parks, Clifford R. (2000):**

**Phylogeny and Biogeography of *Juglans* (Juglandaceae) based on matK and ITS Sequence Data.**
In: American Journal of Botany 87 (6), p. 872–882. DOI: 10.2307/2656895.

ZHAO, PENG; ZHOU, HUI-JUAN; POTTER, DANIEL; HU, YI-HENG; FENG, XIAO-JIA; DANG, MENG ET AL. (2018):
**Population genetics, phylogenomics and hybrid Speciation of *Juglans* in China determined from whole Chloroplast Genomes, Transcriptomes, and Genotyping-by-Sequencing (GbS).**
In: Molecular Phylogenetics and Evolution 126, p. 250–265. DOI: 10.1016/j.ympev.2018.04.014.

### Juglandales

LI, RUI-QI; CHEN, ZHI-DUAN; LU, AN-MING; SOLTIS, DOUGLAS E.; SOLTIS, PAMELA S.; MANOS, PAUL P. (2004):
**Phylogenetic Relationships in Fagales based on DNA Sequences from Three Genomes.**
In: International Journal of Plant Sciences 165 (2), p. 311–324. DOI: 10.1086/381920.

### Juncaceae

DRÁBKOVÁ, LENKA; KIRSCHNER, JAN; VLČEK, ČESTMÍR (2006):
**Phylogenetic Relationships within *Luzula* DC. and *Juncus* L. (Juncaceae): A Comparison of phylogenetic signals of trnL-trnF intergenic Spacer, trnL Intron and rbcL Plastome Sequence Data.**
In: Cladistics 22 (2), p. 132–143. DOI: 10.1111/j.1096-0031.2006.00095.x.

DRÁBKOVÁ, LENKA; VLČEK, ČESTMÍR (2007):
**The phylogenetic Position of *Oxychloe* (Juncaceae): Evidence from Morphology, nuclear and Plastid DNA Regions.**
In: Taxon 56 (1), p. 95–102.

DRBKOV, L.; KIRSCHNER, JAN; SEBERG, OLE; PETERSEN, GITTE; VLČEK, ČESTMÍR (2003):
**Phylogeny of the Juncaceae based on rbc L Sequences, with special Emphasis on *Luzula* DC. and Juncus L.**
In: Plant Systematics and Evolution 240 (1-4), p. 133–147. DOI: 10.1007/s00606-003-0001-6.

ROALSON, ERIC H. (2005):
**Phylogenetic Relationships in the Juncaceae inferred from Nuclear Ribosomal DNA Internal Transcribed Spacer Sequence Data.**
In: International Journal of Plant Sciences 166 (3), p. 397–413.

ZÁVESKÁ DRÁBKOVÁ, LENKA; VLČEK, ČESTMÍR (2009):
**DNA Variation within Juncaceae: Comparison of impact of Organelle Regions on Phylogeny.**
In: Plant Systematics and Evolution 278 (3-4), p. 169–186. DOI: 10.1007/s00606-008-0135-7.

ZÁVESKÁ DRÁBKOVÁ, LENKA; VLČEK, ČESTMÍR (2010):
**Molecular Phylogeny of the Genus *Luzula* DC. (Juncaceae, Monocotyledones) based on plastome and nuclear ribosomal Regions: a Case of Incongruence, incomplete Lineage Sorting and Hybridisation.**
In: Molecular Phylogenetics and Evolution 57 (2), p. 536–551. DOI: 10.1016/j.ympev.2010.07.022.

### Juncaginaceae

MERING, SABINE; KADEREIT, JOACHIM W. (2015):

**Phylogeny, Biogeography and Evolution of *Triglochin* L. (Juncaginaceae) - morphological Diversification is linked to Habitat Shifts rather than to genetic Diversification.**
In: Molecular Phylogenetics and Evolution 83, p. 200–212. DOI: 10.1016/j.ympev.2014.10.014.

### Kewaceae

Christenhusz, Maarten J.M.; Brockington, Samuel F.; Christin, Pascal-Antoine; Sage, Rowan F. (2014):
**On the disintegration of Molluginaceae: a new Genus and Family (*Kewa*, Kewaceae) segregated from *Hypertelis*, and Placement of *Macarthuria* in Macarthuriaceae.**
In: Phytotaxa 181 (4), p. 238. DOI: 10.11646/phytotaxa.181.4.4.

### Koeberliniaceae

Tobe, Hiroshi; Raven, Peter H. (2008):
**Embryology of *Koeberlinia* (Koeberliniaceae): Evidence for Core-Brassicalean Affinities.**
In: American Journal of Botany 95 (11), p. 1475–1486. DOI: 10.3732/ajb.0800218.

### Krameriaceae

Simpson, Beryl B.; Weeks, Andrea; Helfgott, D. Megan; Larkin, Leah L. (2004):
**Species Relationships in *Krameria* (Krameriaceae) based on ITS Sequences and Morphology: Implications for Character Utility and Biogeography.**
In: Systematic Botany 29 (1), p. 97–108. DOI: 10.1600/036364404772974013.

### Lamiaceae

Agostini, G.; Echeverrigaray, S.; Souza-Chies, Tatiana Teixeira de (2012):
**A preliminary Phylogeny of the Genus *Cunila* D. Royen ex L. (Lamiaceae) based on ITS rDNA and trnL-F Regions.**
In: Molecular Phylogenetics and Evolution 65 (2), p. 739–747. DOI: 10.1016/j.ympev.2012.07.030.

Agostini, G.; Echeverrigaray, S.; Souza-Chies, Tatiana Teixeira de (2008):
**Genetic Relationships among South American Species of *Cunila* D.Royen ex L. based on Issr.**
In: Plant Systematics and Evolution 274 (3-4), p. 135–141. DOI: 10.1007/s00606-008-0037-8.

Barber, Janet C.; Finch, Courtney C.; Francisco-Ortega, Javier; Santos-Guerra, Arnoldo; Jansen, Robert K. (2007):
**Hybridization in Macaronesian *Sideritis* (Lamiaceae): Evidence from incongruence of multiple independent Nuclear and Chloroplast Sequence Datasets.**
In: Taxon 56 (1), p. 74–88.

Barber, Janet C.; Francisco-Ortega, Javier; Santos-Guerra, Arnoldo; Turner, Kathryn G.; Jansen, Robert K. (2002):
**Origin of Macaronesian *Sideritis* L. (Lamioideae: Lamiaceae) inferred from nuclear and Chloroplast Sequence Datasets.**
In: Molecular Phylogenetics and Evolution 23, p. 293–306.

Barrabé, Laure; Karnadi-Abdelkader, Giliane; Ounemoa, Jacqueline; Kok, Rogier P.J. de; Robert, Nadia; Gateble, Gildas (2015):
**Recircumscription of *Oxera* (Lamiaceae: Ajugoideae) to include *Faradaya* based on molecular and anatomical Data.**
In: Botanical Journal of the Linnean Society 179, p. 693–711.

Bendiksby, Mika; Brysting, Anne K.; Thorbek, Lisbeth; Gussarova, Galina; Ryding, Olof (2011):

**Molecular Phylogeny and Taxonomy of the Genus *Lamium* L. (Lamiaceae): Disentangling Origins of presumed Allotetraploids.**
In: Taxon 60 (4), p. 986–1000.

BENDIKSBY, MIKA; SALMAKI, YASAMAN; BRÄUCHLER, CHRISTIAN; RYDING, OLOF (2014):
**The generic Position of *Stachys tibetica* Vatke and Amalgamation of the Genera *Eriophyton* and *Stachyopsis* (Lamiaceae Subfam. Lamioideae).**
In: Plant Systematics and Evolution 300 (5), p. 961–971. DOI: 10.1007/s00606-013-0935-2.

BENDIKSBY, MIKA; THORBEK, LISBETH; SCHEEN, ANNE-CATHRINE; LINDQVIST, CHARLOTTE; RYDING, OLOF (2011):
**An updated Phylogeny and Classification of Lamiaceae Subfamily Lamioideae.**
In: Taxon 60 (2), p. 471–484.

BRAMLEY, GEMMA L.C.; FOREST, FÉLIX; KOK, ROGIER P.J. DE (2009):
**Troublesome tropical Mints: re-examining generic Limits of *Vitex* and Relations (Lamiaceae) in South East Asia.**
In: Taxon 58 (2), p. 500–510.

BRÄUCHLER, CHRISTIAN; MEIMBERG, HARALD; ABELE, TILMAN; HEUBL, GÜNTHER (2005):
**Polyphyly of the Genus *Micromeria* (Lamiaceae) as Evidence from cpDNA Sequence Data.**
In: Taxon 54 (3), p. 639–650.

BRÄUCHLER, CHRISTIAN; MEIMBERG, HARALD; HEUBL, GÜNTHER (2010):
**Molecular Phylogeny of Menthinae (Lamiaceae, Nepetoideae, Mentheae) - Taxonomy, Biogeography and Conflicts.**
In: Molecular Phylogenetics and Evolution 55 (2), p. 501–523. DOI: 10.1016/j.ympev.2010.01.016.

CAROVIĆ-STANKO, KLAUDIJA; LIBER, ZLATKO; BESENDORFER, VIŠNJA; JAVORNIK, BRANKA; BOHANEC, BORUT; KOLAK, IVAN; SATOVIĆ, ZLATKO (2010):
**Genetic relations among Basil Taxa (*Ocimum* L.) based on molecular Markers, nuclear DNA content, and Chromosome Number.**
In: Plant Systematics and Evolution 285 (1-2), p. 13–22. DOI: 10.1007/s00606-009-0251-z.

CHEN, YA-PING; DREW, BRYAN T.; LI, BO; SOLTIS, DOUGLAS E.; SOLTIS, PAMELA S.; XIANG, CHUN-LEI (2016):
**Resolving the phylogenetic Position of *Ombrocharis* (Lamiaceae), with Reference to the molecular Phylogeny of Tribe Elsholtzieae.**
In: Taxon 65 (1), p. 123–136. DOI: 10.12705/651.8.

CHEN, YA-PING; LI, BO; OLMSTEAD, RICHARD G.; CANTINO, PHILIP D.; LIU, EN-DE; XIANG, CHUN-LEI (2014):
**Phylogenetic Placement of the enigmatic Genus *Holocheila* (Lamiaceae) inferred from Plastid DNA Sequences.**
In: Taxon 63 (2), p. 355–366. DOI: 10.12705/632.8.

CONN, BARRY J.; HENWOOD, MURRAY J.; PROFT, KIRSTIN M.; WILSON, TREVOR C. (2016):
**Molecular Phylogenetics reveals a new Species of *Prostanthera* from tropical Queensland with links to more southerly Taxa.**
In: Telopea 19, p. 13–22. DOI: 10.7751/telopea10037.

CONN, BARRY J.; HENWOOD, MURRAY J.; STREIBER, N. (2011):
**Synopsis of the Tribe Chloantheae and new nomenclatural Combinations in *Pityrodia* s.lat. (Lamiaceae).**
In: Australian Systematic Botany 24 (1), p. 1–9. DOI: 10.1071/SB10039.

CONN, BARRY J.; STREIBER, N.; BROWN, E. A.; HENWOOD, MURRAY J.; OLMSTEAD, RICHARD G. (2009):

**Infrageneric Phylogeny of Chloantheae (Lamiaceae) based on Chloroplast ndhF and nuclear ITS Sequence Data.**
In: Australian Systematic Botany 22 (4), p. 243–256. DOI: 10.1071/SB09011.

CURTO, MANUEL; SCHACHTLER, CHRISTINA; PUPPO, PAMELA; MEIMBERG, HARALD (2018):
**Using a new RAD-Sequencing approach to Study the Evolution of *Micromeria* in the Canary Islands.**
In: Molecular Phylogenetics and Evolution 119, p. 160–169. DOI: 10.1016/j.ympev.2017.11.005.

DIZKIRICI, AYTEN; CELEP, FERHAT; KANSU, CIGDEM; KAHRAMAN, AHMET; DOGAN, MUSA; KAYA, ZEKI (2015):
**A molecular Phylogeny of *Salvia euphratica* sensu lato (*Salvia* L., Lamiaceae) and its closely related Species with a Focus on the Section *Hymenosphace*.**
In: Plant Systematics and Evolution 301 (10), p. 2313–2323. DOI: 10.1007/s00606-015-1230-1.

DREW, BRYAN T.; SYTSMA, KENNETH J. (2012):
**Phylogenetics, Biogeography, and staminal Evolution in the Tribe Mentheae (Lamiaceae).**
In: American Journal of Botany 99 (5), p. 933–953. DOI: 10.3732/ajb.1100549.

DREW, BRYAN T.; SYTSMA, KENNETH J. (2013):
**The South American Radiation of *Lepechinia* (Lamiaceae): Phylogenetics, Divergence times and Evolution of Dioecy.**
In: Botanical Journal of the Linnean Society 171, p. 171–190.

EDWARDS, CHRISTINE E.; LEFKOWITZ, DAVID; SOLTIS, DOUGLAS E.; SOLTIS, PAMELA P. (2008):
**Phylogeny of *Conradina* and related Southeastern Scrub Mints (Lamiaceae) based on GapC Gene Sequences.**
In: International Journal of Plant Sciences 169 (4), p. 579–594. DOI: 10.1086/528758.

EDWARDS, CHRISTINE E.; SOLTIS, DOUGLAS E.; SOLTIS, PAMELA P. (2006):
**Molecular Phylogeny of *Conradina* and Other Scrub Mints (Lamiaceae) from the Southeastern USA: Evidence for Hybridization in Pleistocene Refugia?**
In: Systematic Botany 31 (1), p. 193–207.

FRAGOSO-MARTÍNEZ, ITZI; SALAZAR, GERARDO A.; MARTÍNEZ-GORDILLO, MARTHA; MAGALLÓN, SUSANA; SÁNCHEZ-REYES, LUNA L.; MORIARTY LEMMON, EMILY ET AL. (2017):
**A pilot Study applying the plant anchored Hybrid Enrichment Method to New World Sages (*Salvia* Subgenus *Calosphace*; Lamiaceae).**
In: Molecular Phylogenetics and Evolution 117, p. 124–134. DOI: 10.1016/j.ympev.2017.02.006.

GUERIN, G. R. (2009):
**A Revision of *Westringia* Section *Cephalowestringia* (Lamiaceae: Westringieae).**
In: Australian Systematic Botany 22 (2), p. 121–136. DOI: 10.1071/SB08030.

HARLEY, RAYMOND MERVYN (2012):
**Checklist and key of Genera and Species of the Lamiaceae of the Brazilian Amazon.**
In: Rodriguésia 63 (1), p. 129–144.

HARLEY, RAYMOND MERVYN (2014):
**Four new Taxa of *Oocephalus* (Hyptidinae: Lamiaceae) from Bahia, Brazil.**
In: Kew Bulletin 69 (4), p. 9539. DOI: 10.1007/S12225-014-9539-4.

HARLEY, RAYMOND MERVYN; PASTORE, JOSÉ FLORIANO B. (2012):
**A generic Revision and new Combinations in the Hyptidinae (Lamiaceae), based on molecular and morphological Evidence.**

In: Phytotaxa 58, p. 1–55.

HUANG, MINGJUAN; CRAWFORD, DANIEL J.; FREUDENSTEIN, JOHN V.; CANTINO, PHILIP D. (2008):
**Systematics of *Trichostema* (Lamiaceae): Evidence from ITS, ndhF, and Morphology.**
In: Systematic Botany 33 (2), p. 437–446.

JAMZAD, ZIBA; CHASE, MARK W.; INGROUILLE, MARTIN J.; SIMMONDS, MONIQUE P. J.; JALILI, ADEL (2003):
**Phylogenetic Relationships in *Nepeta* L. (Lamiaceae) and related Genera based on ITS Sequence Data.**
In: Taxon 52, p. 21–32.

JENKS, AARON A.; WALKER, JAY B.; KIM, SEUNG-CHUL (2013):
**Phylogeny of New World *Salvia* Subgenus *Calosphace* (Lamiaceae) based on cpDNA (psbA-trnH) and nrDNA (ITS) Sequence Data.**
In: Journal of Plant Research 126 (4), p. 483–496. DOI: 10.1007/s10265-012-0543-1.

KOK, ROGIER P.J. DE (2012):
**A Revision of the Genus *Gmelina* (Lamiaceae).**
In: Kew Bulletin 67, p. 293–329.

LI, BO; CANTINO, PHILIP D.; OLMSTEAD, RICHARD G.; BRAMLEY, GEMMA L.C.; XIANG, CHUN-LEI; MA, ZHONG-HUI ET AL. (2016):
**A large-scale Chloroplast Phylogeny of the Lamiaceae sheds new light on its subfamilial Classification.**
In: Scientific Reports 6, p. 34343. DOI: 10.1038/srep34343.

LI, BO; XU, WEIXIANG; TU, TIE-YAO; WANG, ZHONGSHENG; OLMSTEAD, RICHARD G.; PENG, HUA ET AL. (2012):
**Phylogenetic Position of *Wenchengia* (Lamiaceae): A Taxonomically enigmatic and critically endangered Genus.**
In: Taxon 61 (2), p. 392–401.

LI, QIAN-QUAN; LI, MIN-HUI; YUAN, QING-JUN; CUI, ZHAN-HU; HUANG, LU-QI; XIAO, PEI-GEN (2013):
**Phylogenetic Relationships of *Salvia* (Lamiaceae) in China: Evidence from DNA Sequence Datasets.**
In: Journal of Systematics and Evolution 51 (2), p. 184–195. DOI: 10.1111/j.1759-6831.2012.00232.x.

LINDQVIST, CHARLOTTE (2003):
**Cladogenesis and reticulation in the Hawaiian endemic Mints (Lamiaceae).**
In: Cladistics 19 (6), p. 480–495. DOI: 10.1016/j.cladistics.2003.09.003.

LINDQVIST, CHARLOTTE; ALBERT, VICTOR A. (2002):
**Origin of the Hawaiian endemic Mints within North American *Stachys* (Lamiaceae).**
In: American Journal of Botany 89 (10), p. 1709–1724.

MAKI, MASAYUKI; YAMASHIRO, TADASHI; DOHZONO, IKUMI; SUZUKI, KAZUO (2010):
**Molecular Phylogeny of *Isodon* (Lamiaceae) in Japan using Chloroplast DNA Sequences: recent rapid Radiations or ancient introgressive Hybridization?**
In: Plant Species Biology 25 (3), p. 240–248. DOI: 10.1111/j.1442-1984.2010.00290.x.

MATHIESEN, CECILIE; SCHEEN, ANNE-CATHRINE; LINDQVIST, CHARLOTTE (2011):
**Phylogeny and Biogeography of the Lamioid Genus *Phlomis* (Lamiaceae).**
In: Kew Bulletin 66, p. 83–99.

MEIMBERG, HARALD; ABELE, TILMAN; BRÄUCHLER, CHRISTIAN; MCKAY, JOHN K.; PÉREZ DE PAZ, PEDRO LUIS; HEUBL, GÜNTHER (2006):

**Molecular Evidence for adaptive Radiation of *Micromeria* Benth. (Lamiaceae) on the Canary Islands as inferred from Chloroplast and nuclear DNA Sequences and ISSR fingerprint Data.**
In: Molecular Phylogenetics and Evolution 41 (3), p. 566–578. DOI: 10.1016/j.ympev.2006.05.037.

**MENG, HONG-HU; ZHANG, MING-LI (2013):**
**Diversification of Plant Species in arid Northwest China: species-level phylogeographical History of *Lagochilus* Bunge ex Bentham (Lamiaceae).**
In: Molecular Phylogenetics and Evolution 68 (3), p. 398–409. DOI: 10.1016/j.ympev.2013.04.012.

**MOON, HYE-KYOUNG; FLS, SUK-PYO HONG; FLS, ERIK SMETS; HUYSMANS, SUZY (2009):**
**Phylogenetic Significance of Leaf Micromorphology and Anatomy in the Tribe Mentheae (Nepetoideae: Lamiaceae).**
In: Botanical Journal of the Linnean Society 160, p. 211–231.

**MOON, HYE-KYOUNG; SMETS, ERIK; HUYSMANS, SUZY (2010):**
**Phylogeny of Tribe Mentheae (Lamiaceae): the story of Molecules and Micromorphological characters.**
In: Taxon 59 (4), p. 1065–1076.

**MOON, HYE-KYOUNG; VINCKIER, STEFAN; WALKER, JAY B.; SMETS, ERIK; HUYSMANS, SUZY (2008):**
**A Search for phylogenetically Informative Pollen Characters in the Subtribe Salviinae (Mentheae: Lamiaceae).**
In: International Journal of Plant Sciences 169 (3), p. 455–471. DOI: 10.1086/526463.

**O'LEARY, NATALY (2015):**
**Synopsis of Subtribe Hyptidinae (Lamiaceae) in Argentina.**
In: Phytotaxa 233 (3), p. 201. DOI: 10.11646/phytotaxa.233.3.1.

**OTIENO, DONALD F.; BALKWILL, KEVIN; PATON, ALAN J.; SAVOLAINEN, VINCENT (2006):**
**A reassessment of *Hemizygia* and *Syncolostemon* (Ocimeae-Lamiaceae).**
In: Taxon 55 (4), p. 941–958.

**PAN, YUE-ZHI; FANG, LI-QIN; HAO, GANG; CAI, JIE; GONG, XUN (2009):**
**Systematic Positions of *Lamiophlomis* and *Paraphlomis* (Lamiaceae) based on nuclear and Chloroplast Sequences.**
In: Journal of Systematics and Evolution 47 (6), p. 535–542. DOI: 10.1111/j.1759-6831.2009.00050.x.

**PASTORE, JOSÉ FLORIANO B.; HARLEY, RAYMOND MERVYN; FOREST, FÉLIX; PATON, ALAN J.; VAN DEN BERG, CÁSSIO (2011):**
**Phylogeny of the Subtribe Hyptidinae (Lamiaceae Tribe Ocimeae) as inferred from nuclear and Plastid DNA.**
In: Taxon 60 (5), p. 1317–1329.

**PATON, ALAN J.; SPRINGATE, DAVID; SUDDEE, SOMRAN; OTIENO, DONALD F.; GRAYER, RENÉE J.; HARLEY, MADELINE M. ET AL. (2004):**
**Phylogeny and Evolution of Basils and Allies (Ocimeae, Labiatae) based on three Plastid DNA Regions.**
In: Molecular Phylogenetics and Evolution 31 (1), p. 277–299. DOI: 10.1016/j.ympev.2003.08.002.

**PEIRSON, JESS A.; CANTINO, PHILIP D.; JR., HARVEY E. BALLARD (2006):**
**A taxonomic Revision of *Collinsonia* (Lamiaceae) based on phenetic Analyses of morphological Variation.**
In: Systematic Botany 31 (2), p. 398–409.

**Pollard, Benedict John; Paton, Alan J. (2009):**
**The African *Plectranthus* (Lamiaceae) Expansion continues. Vale Leocus!**
In: Kew Bulletin 64, p. 259–261.

**Puppo, Pamela; Curto, Manuel; Gusmão-Guedes, Joana; Cochofel, Jaqueline; Pérez de Paz, Pedro Luis; Bräuchler, Christian; Meimberg, Harald (2015):**
**Molecular Phylogenetics of *Micromeria* (Lamiaceae) in the Canary Islands, Diversification and inter-Island Colonization Patterns inferred from nuclear Genes.**
In: Molecular Phylogenetics and Evolution 89, p. 160–170. DOI: 10.1016/j.ympev.2015.04.017.

**Roy, Tilottama; Chang, Tien-Hao; Lan, Tianying; Lindqvist, Charlotte (2013):**
**Phylogeny and Biogeography of New World Stachydeae (Lamiaceae) with Emphasis on the Origin and Diversification of Hawaiian and South American Taxa.**
In: Molecular Phylogenetics and Evolution 69 (1), p. 218–238. DOI: 10.1016/j.ympev.2013.05.023.

**Roy, Tilottama; Cole, Logan W.; Chang, Tien-Hao; Lindqvist, Charlotte (2015):**
**Untangling reticulate evolutionary Relationships among New World and Hawaiian Mints (Stachydeae, Lamiaceae).**
In: Molecular Phylogenetics and Evolution 89, p. 46–62. DOI: 10.1016/j.ympev.2015.03.023.

**Roy, Tilottama; Lindqvist, Charlotte (2015):**
**New Insights into evolutionary Relationships within the Subfamily Lamioideae (Lamiaceae) based on Pentatricopeptide Repeat (PPR) nuclear DNA Sequences.**
In: American Journal of Botany 102 (10), p. 1721–1735. DOI: 10.3732/ajb.1500233.

**Salmaki, Yasaman; Bendiksby, Mika; Heubl, Günther (2015):**
**Molecular Phylogeny confirms the Placement of enigmatic *Stachys persepolitana* in *Lamium* (Lamiaceae; Subfam. Lamioideae).**
In: Phytotaxa 192 (4), p. 254. DOI: 10.11646/phytotaxa.192.4.3.

**Salmaki, Yasaman; Kattari, Stefan; Heubl, Günther; Bräuchler, Christian (2016):**
**Phylogeny of non-monophyletic *Teucrium* (Lamiaceae: Ajugoideae): Implications for Character Evolution and Taxonomy.**
In: Taxon 65 (4), p. 805–822. DOI: 10.12705/654.8.

**Salmaki, Yasaman; Zarre, Shahin; Govaerts, Rafael; Bräuchler, Christian (2012):**
**A taxonomic Revision of the Genus *Stachys* (Lamiaceae: Lamioideae) in Iran.**
In: Botanical Journal of the Linnean Society 170, p. 573–617.

**Salmaki, Yasaman; Zarre, Shahin; Lindqvist, Charlotte; Heubl, Günther; Bräuchler, Christian (2011):**
**Comparative Leaf Anatomy of *Stachys* (Lamiaceae: Lamioideae) in Iran with a Discussion on its subgeneric Classification.**
In: Plant Systematics and Evolution 294 (1-2), p. 109–125. DOI: 10.1007/s00606-011-0450-2.

**Salmaki, Yasaman; Zarre, Shahin; Ryding, Olof; Lindqvist, Charlotte; Bräuchler, Christian; Heubl, Günther et al. (2013):**
**Molecular Phylogeny of Tribe Stachydeae (Lamiaceae Subfamily Lamioideae).**
In: Molecular Phylogenetics and Evolution 69 (3), p. 535–551. DOI: 10.1016/j.ympev.2013.07.024.

**Salmaki, Yasaman; Zarre, Shahin; Ryding, Olof; Lindqvist, Charlotte; Scheunert, Agnes; Bräuchler, Christian; Heubl, Günther (2012):**
**Phylogeny of the Tribe Phlomideae (Lamioideae: Lamiaceae) with special Focus on *Eremostachys* and *Phlomoides*: new Insights from nuclear and Chloroplast Sequences.**
In: Taxon 61 (1), p. 161–179.

**Scheen, Anne-Cathrine; Albert, Victor A. (2009):**
**Molecular Phylogenetics of the *Leucas* Group (Lamioideae; Lamiaceae).**
In: Systematic Botany 34 (1), p. 173–181. DOI: 10.1600/036364409787602366.

**Scheen, Anne-Cathrine; Bendiksby, Mika; Ryding, Olof; Mathiesen, Cecilie; Albert, Victor A.; Lindqvist, Charlotte (2010):**
**Molecular Phylogenetics, Character Evolution, and Suprageneric Classification of Lamioideae (Lamiaceae).**
In: Annals of the Missouri Botanical Garden 97 (2), p. 191–217. DOI: 10.3417/2007174.

**Scheen, Anne-Cathrine; Lindqvist, Charlotte; Fossdal, Carl G.; Albert, Victor A. (2008):**
**Molecular Phylogenetics of Tribe Synandreae, a North American Lineage of Lamioid Mints (Lamiaceae).**
In: Cladistics 24 (3), p. 299–314. DOI: 10.1111/j.1096-0031.2007.00180.x.

**Schmidt-Lebuhn, Alexander N. (2008):**
**Monophyly and phylogenetic Relationships of *Minthostachys* (Labiatae, Nepetoideae) examined using morphological and nrITS Data.**
In: Plant Systematics and Evolution 270 (1-2), p. 25–38. DOI: 10.1007/s00606-007-0598-y.

**Shepherd, Kelly A.; Perkins, Andrew J.; Collins, Joel; Byrne, Margaret; Thiele, Kevin R. (2013):**
**Morphological and molecular Evidence supports the Recognition of a new Subspecies of the critically endangered *Pityrodia scabra* (Lamiaceae).**
In: Australian Systematic Botany 26 (1), p. 1–12. DOI: 10.1071/SB12009.

**Steane, Dorothy A.; Kok, Rogier P.J. de; Olmstead, Richard G. (2004):**
**Phylogenetic Relationships between *Clerodendrum* (Lamiaceae) and other Ajugoid Genera inferred from nuclear and Chloroplast DNA Sequence Data.**
In: Molecular Phylogenetics and Evolution 32 (1), p. 39–45. DOI: 10.1016/j.ympev.2003.11.011.

**Steane, Dorothy A.; Scotland, Robert W.; Mabberley, David J.; Olmstead, Richard G. (1999):**
**Molecular Systematics of *Clerodendrum* (Lamiaceae): ITS Sequences and total Evidence.**
In: American Journal of Botany 86 (1), p. 98–107.

**Turner, Ian M. (2014):**
**A new Combination in Asian *Clinopodium* (Lamiaceae).**
In: Phytotaxa 174 (4), p. 242. DOI: 10.11646/phytotaxa.174.4.6.

**Walker, Jay B.; Sytsma, Kenneth J. (2007):**
**Staminal Evolution in the Genus *Salvia* (Lamiaceae): molecular phylogenetic Evidence for multiple Origins of the staminal Lever.**
In: Annals of Botany 100 (2), p. 375–391. DOI: 10.1093/aob/mcl176.

**Walker, Jay B.; Sytsma, Kenneth J.; Treutlein, Jens; Wink, Michael (2004):**
**Salvia (Lamiaceae) is not monophyletic: Implications for the Systematics, Radiation, and Ecological Specializations of *Salvia* and Tribe Mentheae.**
In: American Journal of Botany 91 (7), p. 1115–1125.

**Wearn, James A.; Mabberley, David J. (2011):**
***Clerodendrum* (Lamiaceae) in Borneo.**
In: Systematic Botany 36 (4), p. 1050–1061. DOI: 10.1600/036364411X605056.

**Welch, Andreanna J.; Collins, Katherine; Ratan, Aakrosh; Drautz-Moses, Daniela I.; Schuster, Stephan C.; Lindqvist, Charlotte (2016):**

**The quest to resolve recent Radiations: Plastid Phylogenomics of extinct and endangered Hawaiian endemic Mints (Lamiaceae).**
In: Molecular Phylogenetics and Evolution 99, p. 16–33. DOI: 10.1016/j.ympev.2016.02.024.

WILL, MARIA; CLAßEN-BOCKHOFF, REGINE (2014):
**Why Africa matters: Evolution of Old World *Salvia* (Lamiaceae) in Africa.**
In: Annals of Botany 114 (1), p. 61–83. DOI: 10.1093/aob/mcu081.

WILL, MARIA; CLAßEN-BOCKHOFF, REGINE (2017):
**Time to split *Salvia* s.l. (Lamiaceae) - new Insights from Old World *Salvia* Phylogeny.**
In: Molecular Phylogenetics and Evolution 109, p. 33–58. DOI: 10.1016/j.ympev.2016.12.041.

WILSON, TREVOR C.; CONN, BARRY J.; HENWOOD, MURRAY J. (2012):
**Molecular Phylogeny and Systematics of *Prostanthera* (Lamiaceae).**
In: Australian Systematic Botany 25 (5), p. 341–352. DOI: 10.1071/SB12006.

XIANG, CHUN-LEI; ZHANG, QIANG; SCHEEN, ANNE-CATHRINE; CANTINO, PHILIP D.; FUNAMOTO, TSUNEO; PENG, HUA (2013):
**Molecular Phylogenetics of *Chelonopsis* (Lamiaceae: Gomphostemmateae) as inferred from nuclear and Plastid DNA and Morphology.**
In: Taxon 62 (2), p. 375–386.

YAO, GANG; DREW, BRYAN T.; YI, TING-SHUANG; YAN, HAI-FEI; YUAN, YONG-MING; GE, XUE-JUN (2016):
**Phylogenetic Relationships, Character Evolution and biogeographic Diversification of *Pogostemon* s.l. (Lamiaceae).**
In: Molecular Phylogenetics and Evolution 98, p. 184–200. DOI: 10.1016/j.ympev.2016.01.020.

YU, XIANG-QIN; MAKI, MASAYUKI; DREW, BRYAN T.; PATON, ALAN J.; LI, HSI-WEN; ZHAO, JIAN-LI ET AL. (2014):
**Phylogeny and historical Biogeography of *Isodon* (Lamiaceae): rapid Radiation in south-west China and Miocene overland Dispersal into Africa.**
In: Molecular Phylogenetics and Evolution 77, p. 183–194. DOI: 10.1016/j.ympev.2014.04.017.

YUAN, YAO-WU; MABBERLEY, DAVID J.; STEANE, DOROTHY A.; OLMSTEAD, RICHARD G. (2010):
**Further disintegration and redefinition of *Clerodendrum* (Lamiaceae): Implications for the Understanding of the Evolution of an intriguing breeding Strategy.**
In: Taxon 59 (1), p. 125–133.

ZHONG, JIN-SHUN; LI, JIE; LI, LANG; CONRAN, JOHN G.; LI, HSI-WEN (2010):
**Phylogeny of *Isodon* (Schrad. ex Benth.) Spach (Lamiaceae) and related Genera inferred from Nuclear Ribosomal ITS, trnL–trnF Region, and rps16 Intron Sequences and Morphology.**
In: Systematic Botany 35 (1), p. 207–219. DOI: 10.1600/036364410790862614.

### Lamiales

WORTLEY, ALEXANDRA H.; RUDALL, PAULA J.; HARRIS, DAVID J.; SCOTLAND, ROBERT W. (2005):
**How much Data are needed to resolve a difficult Phylogeny?: case Study in Lamiales.**
In: Systematic Biology 54 (5), p. 697–709. DOI: 10.1080/10635150500221028.

### Lamiidae

REFULIO-RODRIGUEZ, NANCY F.; OLMSTEAD, RICHARD G. (2014):
**Phylogeny of Lamiidae.**
In: American Journal of Botany 101 (2), p. 287–299. DOI: 10.3732/ajb.1300394.

### Lardizabalaceae

TIAN, SHUANG; LEI, SHU-QING; HU, WAN; DENG, LING-LI; LI, BO; MENG, QING-LIN ET AL. (2015):

**Repeated range expansions and inter-/postglacial Recolonization routes of *Sargentodoxa cuneata* (Oliv.) Rehd. et Wils. (Lardizabalaceae) in subtropical China revealed by Chloroplast Phylogeography.**

In: Molecular Phylogenetics and Evolution 85, p. 238–246. DOI: 10.1016/j.ympev.2015.02.016.

### Lauraceae

Alves, Flavio Macedo; Souza, Vinicius Castro (2013):

**Phylogenetic Analysis of the Neotropical Genus *Mezilaurus* and Reestablishment of *Clinostemon* (Lauraceae).**

In: Taxon 62 (2), p. 281–290.

Chanderbali, André S.; van der Werff, Henk; Renner, Susanne p. (2001):

**Phylogeny and historical Biogeography of Lauraceae: Evidence from the Chloroplast and nuclear Genomes.**

In: Annals of the Missouri Botanical Garden 88, p. 104–134.

Fijridiyanto, Izu A.; Murakami, Noriaki (2009):

**Molcecular Systematics of Malesian *Litsea* Lam. and putative related Genera (Lauraceae).**

In: Acta Phytotaxonomica Geobotanica 60 (1), p. 1–18.

Fijridiyanto, Izu A.; Murakami, Noriaki (2009):

**Phylogeny of *Litsea* and related Genera (Laureae-Lauraceae) based on Analysis of rpb2 Gene Sequences.**

In: Journal of Plant Research 122 (3), p. 283–298. DOI: 10.1007/s10265-009-0218-8.

Huang, Jian-Feng; Li, Lang; Conran, John G.; Li, Jie (2016):

**Phylogenetic Utility of Leafy Gene in *Cinnamomum* (Lauraceae): Gene Duplication and Polymerase Chain Reaction-mediated recombination.**

In: Journal of Systematics and Evolution 54 (3), p. 238–249. DOI: 10.1111/jse.12189.

Huang, Jian-Feng; Li, Lang; van der Werff, Henk; Li, Hsi-Wen; Rohwer, Jens Gunter; Crayn, Darren M. et al. (2016):

**Origins and Evolution of Cinnamon and Camphor: A phylogenetic and historical biogeographical Analysis of the *Cinnamomum* Group (Lauraceae).**

In: Molecular Phylogenetics and Evolution 96, p. 33–44. DOI: 10.1016/j.ympev.2015.12.007.

Julia, S.; Soepadmo, E.; Yahud, W. (2009):

**Problem in the generic Delimitation between *Alseodaphne, Dehaasia* and *Nothaphoebe* (Lauraceae) in Borneo.**

In: Blumea 54 (1), p. 192–197. DOI: 10.3767/000651909X476148.

Kokubugata, Goro; Nakamura, Koh; Forster, Paul I.; Wilson, Gary W.; Holland, Ailsa E.; Hirayama, Yumiko; Yokota, Masatsugu (2012):

***Cassytha pubescens* and *C. glabella* (Lauraceae) are not disjunctly distributed between Australia and the Ryukyu Archipelago of Japan - Evidence from morphological and molecular Data.**

In: Australian Systematic Botany 25 (5), p. 364–373. DOI: 10.1071/SB10040.

Li, Jie; Christophel, David C. (2000):

**Systematic Relationships within the *Litsea* Complex (Lauraceae): a cladistic Analysis on the Basis of morphological and Leaf Cuticle Data.**

In: Australian Systematic Botany 13, p. 1–13.

**Li, Jie; Christophel, David C.; Conran, John G.; Li, Hsi-Wen (2004):**

**Phylogenetic Relationships within the "Core" Laureae (*Litsea* complex, Lauraceae) inferred from Sequences of the Chloroplast Gene matK and nuclear ribosomal DNA ITS Regions.**

In: Plant Systematics and Evolution 246 (1-2), p. 19–34. DOI: 10.1007/s00606-003-0113-z.

**Li, Jie; Conran, John G.; Christophel, David C.; Li, Zhi-Ming; Li, Lang; Li, Hsi-Wen (2008):**

**Phylogenetic Relationships of the *Litsea* Complex and Core Laureae (Lauraceae) using ITS and ETS Sequences and Morphology.**

In: Annals of the Missouri Botanical Garden 95 (4), p. 580–599. DOI: 10.3417/2006125.9504.

**Li, L.; Li, J.; Conran, John G.; Li, X.-W.; Li, H.-W. (2007):**

**Phylogeny of *Neolitsea* (Lauraceae) inferred from Bayesian Analysis of nrDNA ITS and ETS Sequences.**

In: Plant Systematics and Evolution 269 (3-4), p. 203–221. DOI: 10.1007/s00606-007-0580-8.

**Li, Lang; Li, Jie; Rohwer, Jens Gunter; van der Werff, Henk; Wang, Zhi-hua; Li, Hsi-Wen (2011):**

**Molecular phylogenetic Analysis of the *Persea* Group (Lauraceae) and its biogeographic Implications on the Evolution of tropical and subtropical Amphi-Pacific Disjunctions.**

In: American Journal of Botany 98 (9), p. 1520–1536. DOI: 10.3732/ajb.1100006.

**Li, Lang; Madriñán, Santiago; Li, Jie (2016):**

**Phylogeny and Biogeography of *Caryodaphnopsis* (Lauraceae) inferred from low-copy nuclear Gene and ITS Sequences.**

In: Taxon 65 (3), p. 433–443. DOI: 10.12705/653.1.

**Li, Zhi-Ming (2006):**

**Polyphyly of the Genus *Actinodaphne* (Lauraceae) inferred from the Analyses of nrDNA ITS and ETS Sequences.**

In: Acta Phytotaxonomica Sinica 44 (3), p. 272. DOI: 10.1360/aps040150.

**Nie, Z.-L.; Wen, Jun; Sun, H. (2007):**

**Phylogeny and Biogeography of *Sassafras* (Lauraceae) disjunct between Eastern Asia and Eastern North America.**

In: Plant Systematics and Evolution 267 (1-4), p. 191–203. DOI: 10.1007/s00606-007-0550-1.

**Nishida, Sachiko; van der Werff, Henk (2011):**

**An Evaluation of Classification by cuticular Characters of the Lauraceae. A Comparison to molecular Phylogeny.**

In: Annals of the Missouri Botanical Garden 98 (3), p. 348–357. DOI: 10.3417/2010054.

**Nishida, Sachiko; van der Werff, Henk (2014):**

**Do Cuticle Characters support the Recognition of *Alseodaphne, Nothaphoebe* and *Dehaasia* as distinct Genera?**

In: Reinwardtia 14 (1), p. 53–66.

**Rohwer, Jens Gunter (2000):**

**Toward a phylogenetic Classification of the Lauraceae. Evidence from matK Sequences.**

In: Systematic Botany 25 (1), p. 60. DOI: 10.2307/2666673.

**Rohwer, Jens Gunter; Li, Jie; Rudolph, Barbara; Schmidt, Sabrina A.; van der Werff, Henk; Li, Hsi-Wen (2009):**

**Is *Persea* (Lauraceae) monophyletic? Evidence from nuclear ribosomal ITS Sequence.**
In: Taxon 58 (4), p. 1153–1167.

**Rohwer, Jens Gunter; Moraes, Pedro Luis Rodrigues; Rudolph, Barbara; van der Werff, Henk (2014):**
**A phylogenetic Analysis of the *Cryptocarya* Group (Lauraceae), and Relationships of *Dahlgrenodendron, Sinopora, Triadodaphne,* and *Yasunia*.**
In: Phytotaxa 158 (2), p. 111–132. DOI: 10.11646/phytotaxa.158.2.1.

**Trofimov, Dimitrij; Rudolph, Barbara; Rohwer, Jens Gunter (2016):**
**Phylogenetic Study of the Genus *Nectandra* (Lauraceae), and Reinstatement of *Damburneya*.**
In: Taxon 65 (5), p. 980–996. DOI: 10.12705/655.3.

**van der Merwe, Marlien M.; Crayn, Darren M.; Ford, Andrew J.; Weston, Peter H.; Rossetto, Maurizio (2016):**
**Evolution of Australian *Cryptocarya* (Lauraceae) based on nuclear and Plastid phylogenetic Trees. Evidence of recent Landscape-level Disjunctions.**
In: Australian Systematic Botany 29 (2), p. 157–166. DOI: 10.1071/SB16023.

**Wang, Zhi-hua; Li, Jie; Conran, John G.; Li, Hsi-Wen (2010):**
**Phylogeny of the Southeast Asian endemic Genus *Neocinnamomum* H.Liu (Lauraceae).**
In: Plant Systematics and Evolution 290 (1-4), p. 173–184. DOI: 10.1007/s00606-010-0359-1.

### Lecythidaceae

**Barrett, Russell L. (2006):**
**A Review of *Planchonia* (Lecythidaceae) in Australia.**
In: Australian Systematic Botany 19 (2), p. 147–153. DOI: 10.1071/SB05008.

**Huang, Ya-Yi; Mori, Scott A.; Kelly, Lawrence M. (2011):**
**A morphological cladistic Analysis of Lecythidoideae with Emphasis on *Bertholletia, Corythophora, Eschweilera,* and *Lecythis*.**
In: Brittonia 63 (3), p. 396–417. DOI: 10.1007/s12228-011-9202-4.

**Huang, Ya-Yi; Mori, Scott A.; Kelly, Lawrence M. (2015):**
**Toward a phylogenetic-based Generic Classification of Neotropical Lecythidaceae - I. Status of *Bertholletia, Corythophora, Eschweilera* and *Lecythis*.**
In: Phytotaxa 203 (2), p. 85. DOI: 10.11646/phytotaxa.203.2.1.

**Huang, Ya-Yi; Mori, Scott A.; Prance, Ghillean T. (2008):**
**A Phylogeny of *Cariniana* (Lecythidaceae) based on morphological and anatomical Data.**
In: Brittonia 60 (1), p. 69–81. DOI: 10.1007/s12228-008-9014-3.

**Mori, Scott A.; Smith, Nathan P.; Huang, Ya-Yi; Prance, Ghillean T.; Kelly, Lawrence M.; Matos, Carol Carollo (2015):**
**Toward a Phylogenetic-based Generic Classification of Neotropical Lecythidaceae - II. Status of *Allantoma, Cariniana, Couratari, Couroupita, Grias* and *Gustavia*.**
In: Phytotaxa 203 (2), p. 122. DOI: 10.11646/phytotaxa.203.2.2.

**Mori, Scott A.; Tsou, Chi-Hua; Wu, Chi-Chih; Cronholm, Bodil; Anderberg, Arne A. (2007):**
**Evolution of Lecythidaceae with an Emphasis on the Circumscription of Neotropical Genera: information from combined ndhF and trnL-F Sequence Data.**
In: American Journal of Botany 94 (3), p. 289–301. DOI: 10.3732/ajb.94.3.289.

**Prance, Ghillean T.; Jongkind, Carel C. H. (2015):**
**A Revision of African Lecythidaceae.**
In: Kew Bulletin 70 (6). DOI: 10.1007/S12225-014-9547-4.

### Lentibulariaceae

**Cieslak, Thomas; Polepalli, Jai Santosh; White, Adam; Müller, Kai F.; Borsch, Thomas; Barthlott, Wilhelm et al. (2005):**
**Phylogenetic Analysis of *Pinguicula* (Lentibulariaceae): Chloroplast DNA Sequences and Morphology support several geographically distinct Adiations.**
In: American Journal of Botany 92 (10), p. 1723–1736.

**Degtjareva, Galina V.; Casper, p. J.; Hellwig, Frank H.; Schmidt, A. R.; Steiger, Jürgen; Sokoloff, Dmitry D. (2006):**
**Morphology and nrITS Phylogeny of the Genus *Pinguicula* L. (Lentibulariaceae), with special attention to Embryo Evolution.**
In: Plant Biology 8 (6), p. 778–790. DOI: 10.1055/s-2006-924560.

**Fleischmann, Andreas; Schäferhoff, Bastian; Heubl, Günther; Rivadavia, Fernando; Barthlott, Wilhelm; Müller, Kai F. (2010):**
**Phylogenetics and Character Evolution in the carnivorous Plant Genus *Genlisea* A.St.-Hil. (Lentibulariaceae).**
In: Molecular Phylogenetics and Evolution 56 (2), p. 768–783. DOI: 10.1016/j.ympev.2010.03.009.

**Kondo, Katsuhiko; Shimai, Hiro (2006):**
**Phylogenetic based Analysis of the Northern *Pinguicula* (Lentibulariaceae) on Internal Transcribed Spacer (ITS) Sequence.**
In: Acta Phytotaxonomica Geobotanica 57 (2), p. 155–164.

**Müller, Kai F.; Borsch, Thomas (2005):**
**Phylogenetics of *Utricularia* (Lentibulariaceae) and molecular Evolution of the trnK Intron in a Lineage with high substitutional Rates.**
In: Plant Systematics and Evolution 250 (1-2), p. 39–67. DOI: 10.1007/s00606-004-0224-1.

**Müller, Kai F.; Borsch, Thomas; Legendre, L.; Porembski, Stefan; Barthlott, Wilhelm (2006):**
**Recent progress in Understanding the Evolution of carnivorous Lentibulariaceae (Lamiales).**
In: Plant Biology 8 (6), p. 748–757. DOI: 10.1055/s-2006-924706.

**Reut, Markus S.; Jobson, Richard W. (2010):**
**A phylogenetic Study of Subgenus Polypompholyx: a parallel Radiation of *Utricularia* (Lentibulariaceae) throughout Australasia.**
In: Australian Systematic Botany 23 (3), p. 152–161. DOI: 10.1071/SB09054.

**Silva, Saura R.; Gibson, Robert; Adamec, Lubomír; Domínguez, Yoannis; Miranda, Vitor F. O. (2018):**
**Molecular Phylogeny of Bladderworts: A wide approach of *Utricularia* (Lentibulariaceae) Species Relationships based on six plastidial and nuclear DNA Sequences.**
In: Molecular Phylogenetics and Evolution 118, p. 244–264. DOI: 10.1016/j.ympev.2017.10.010.

### Liliaceae

**Bartha, László; Stepanov, Nikolay V.; Rukšāns, Jānis; Banciu, Horia L.; Keresztes, Lujza (2015):**
**Non-Monophyly of Siberian *Erythronium* (Liliaceae) leads to the Recognition of the formerly neglected *Erythronium sajanense*.**
In: Journal of Plant Research 128 (5), p. 721–729. DOI: 10.1007/s10265-015-0734-7.

**Booy, G.; van der Schoot, J.; Vosman, Ben (2000):**
**Heterogeneity of the Internal Transcribed Spacer 1 (ITS1) in *Tulipa* (Liliaceae).**
In: Plant Systematics and Evolution 225 (1-4), p. 29–41. DOI: 10.1007/BF00985457.

**Christenhusz, Maarten J.M.; Govaerts, Rafael; David, John C.; Hall, Tony; Borland, Katherine; Roberts, Penelope P. et al. (2013):**
**Tiptoe through the Tulips - cultural History, molecular Phylogenetics and Classification of *Tulipa* (Liliaceae).**
In: Botanical Journal of the Linnean Society 172 (3), p. 280–328. DOI: 10.1111/boj.12061.

**Clennett, John C. B.; Chase, Mark W.; Forest, Félix; Maurin, Olivier; Wilkin, Paul (2012):**
**Phylogenetic Systematics of *Erythronium* (Liliaceae): morphological and molecular Analyses.**
In: Botanical Journal of the Linnean Society 170 (4), p. 504–528. DOI: 10.1111/j.1095-8339.2012.01302.x.

**Day, Peter D.; Berger, Madeleine; Hill, Laurence; Fay, Michael F.; Leitch, Andrew R.; Leitch, Ilia J.; Kelly, Laura J. (2014):**
**Evolutionary Relationships in the medicinally important Genus *Fritillaria* L. (Liliaceae).**
In: Molecular Phylogenetics and Evolution 80, p. 11–19. DOI: 10.1016/j.ympev.2014.07.024.

**Eker, İsmaíl; Babaç, Mehmet Tekİn; Koyuncu, Mehmet (2014):**
**Revision of the Genus *Tulipa* L. (Liliaceae) in Turkey.**
In: Phytotaxa 157 (1), p. 1. DOI: 10.11646/phytotaxa.157.1.1.

**Gao, Yun-Dong; Harris, A. J.; He, Xing-Jin (2015):**
**Morphological and ecological Divergence of *Lilium* and *Nomocharis* within the Hengduan Mountains and Qinghai-Tibetan Plateau may result from Habitat Specialization and Hybridization.**
In: BMC Evolutionary Biology 15, p. 147. DOI: 10.1186/s12862-015-0405-2.

**Gao, Yun-Dong; Harris, A. J.; Zhou, Song-Dong; He, Xing-Jin (2013):**
**Evolutionary events in *Lilium* (including *Nomocharis*, Liliaceae) are temporally correlated with Orogenies of the Q-T Plateau and the Hengduan Mountains.**
In: Molecular Phylogenetics and Evolution 68 (3), p. 443–460. DOI: 10.1016/j.ympev.2013.04.026.

**Hayashi, Kazuhiko; Kawano, Shoichi (2000):**
**Molecular Systematics of *Lilium* and allied Genera (Liliaceae): phylogenetic Relationships among *Lilium* and related Genera based on the rbcL and matK Gene Sequence Data.**
In: Plant Species Biology 15 (1), p. 73–93. DOI: 10.1046/j.1442-1984.2000.00025-2.x.

**Hayashi, Kazuhiko; Yoshida, Seiji; Utech, Frederick H.; Kawano, Shoichi (2001):**
**Molecular Systematics in the Genus *Clintonia* and related Taxa based on rbcL and matK Gene Sequence Data.**
In: Plant Species Biology 16 (2), p. 119–137. DOI: 10.1046/j.1442-1984.2001.00057.x.

**Ikinci, Nursel; Oberprieler, Christoph (2010):**
**Genetic Relationships among NE-Turkish *Lilium* L. (Liliaceae) Species based on a random amplified polymorphic DNA Analysis.**
In: Plant Systematics and Evolution 284 (1-2), p. 41–48. DOI: 10.1007/s00606-009-0239-8.

**Khourang, Mahmoud; Babaei, Alireza; Sefidkon, Fatemeh; Naghavi, Mohammad Reza; Asgari, Davood; Potter, Daniel (2014):**

**Phylogenetic Relationship in *Fritillaria* spp. of Iran inferred from ribosomal ITS and Chloroplast trnL-trnF Sequence Data.**
In: Biochemical Systematics and Ecology 57, p. 451–457. DOI: 10.1016/j.bse.2014.10.001.

**Kiani, Mahnaz; Memariani, Farshid; Zarghami, Homa (2012):**
**Molecular Analysis of Species of *Tulipa* L. from Iran based on ISSR Markers.**
In: Plant Systematics and Evolution 298 (8), p. 1515–1522. DOI: 10.1007/s00606-012-0654-0.

**Mucciarelli, Marco; Fay, Michael F. (2013):**
**Plastid DNA fingerprinting of the rare *Fritillaria moggridgei* (Liliaceae) reveals Population Differentiation and genetic Isolation within the *Fritillaria tubiformis* complex.**
In: Phytotaxa 91 (1), p. 1. DOI: 10.11646/phytotaxa.91.1.1.

**Muratović, Edina; Robin, Odil; Bogunić, Faruk; Šoljan, Dubravka; Siljak-Yakovlev, Sonja (2010):**
**Karyotype Evolution and Speciation of European Lilies from *Lilium* Sect. *Liriotypus*.**
In: Taxon 59 (1), p. 165–175. DOI: 10.1002/tax.591016.

**Nishikawa, Tomotaro; Okazaki, Keiichi; Uchino, Tae; Arakawa, Katsuro; Nagamine, Tsukasa (1999):**
**A molecular Phylogeny of *Lilium* in the Internal Transcribed Spacer Region of Nuclear Ribosomal DNA.**
In: Molecular Phylogenetics and Evolution 49 (49), p. 238–249.

**Peruzzi, Lorenzo; Peterson, Angela; Tison, Jean-Marc; Harpke, Dörte (2011):**
**New light on Phylogeny and Taxonomy of the Eurasian *Gagea villosa*-*G. fragifera* Complex (Liliaceae).**
In: Nordic Journal of Botany 29 (6), p. 722–733. DOI: 10.1111/j.1756-1051.2011.01187.x.

**Peruzzi, Lorenzo; Peterson, Angela; Tison, Jean-Marc; Peterson, Jens (2008):**
**Phylogenetic Relationships of Gagea Salisb. (Liliaceae) in Italy, inferred from molecular and morphological Data matrices.**
In: Plant Systematics and Evolution 276 (3-4), p. 219–234. DOI: 10.1007/s00606-008-0081-4.

**Peruzzi, Lorenzo; Tison, Jean-Marc; Peterson, Angela; Peterson, Jens (2008):**
**On the phylogenetic Position and taxonomic value of *Gagea trinervia* (Viv.) Greuter and *Gagea* Sect. *Anthericoides* A.Terracc. (Liliaceae).**
In: Taxon 57 (4), p. 1201–1214. DOI: 10.1002/tax.574013.

**Peterson, Angela; Harpke, Dörte; Levichev, Igor G.; Beisenova, Saltanat; Schnittler, Martin; Peterson, Jens (2016):**
**Morphological and molecular Investigations of *Gagea* (Liliaceae) in Southeastern Kazakhstan with special Reference to putative altitudinal Hybrid Zones.**
In: Plant Systematics and Evolution 302 (8), p. 985–1007. DOI: 10.1007/s00606-016-1313-7.

**Peterson, Angela; John, H.; Koch, E.; Peterson, Jens (2004):**
**A molecular Phylogeny of the Genus Gagea (Liliaceae) in Germany inferred from non-coding Chloroplast and nuclear DNA Sequences.**
In: Plant Systematics and Evolution 245 (3-4), p. 145–162. DOI: 10.1007/s00606-003-0114-y.

**Peterson, Angela; Levichev, Igor G.; Peterson, Jens (2008):**
**Systematics of *Gagea* and *Lloydia* (Liliaceae) and infrageneric Classification of *Gagea* based on molecular and morphological Data.**
In: Molecular Phylogenetics and Evolution 46 (2), p. 446–465. DOI: 10.1016/j.ympev.2007.11.016.

**Peterson, Angela; Levichev, Igor G.; Peterson, Jens; Harpke, Dörte; Schnittler, Martin (2011):**

**New Insights into the Phylogeny and Taxonomy of Chinese Species of *Gagea* (Liliaceae) - Speciation through Hybridization.**
In: Organisms Diversity and Evolution 11 (5), p. 387–407. DOI: 10.1007/s13127-011-0059-x.

REŠETNIK, I.; LIBER, ZLATKO; SATOVIĆ, ZLATKO; CIGIĆ, P.; NIKOLIĆ, TONI (2007):
**Molecular Phylogeny and Systematics of the *Lilium carniolicum* Group (Liliaceae) based on nuclear ITS Sequences.**
In: Plant Systematics and Evolution 265 (1-2), p. 45–58. DOI: 10.1007/s00606-006-0513-y.

RØNSTED, NINA; LAW, STEVE; THORNTON, HANNAH; FAY, MICHAEL F.; CHASE, MARK W. (2005):
**Molecular phylogenetic Evidence for the Monophyly of *Fritillaria* and *Lilium* (Liliaceae; Liliales) and the infrageneric Classification of *Fritillaria*.**
In: Molecular Phylogenetics and Evolution 35 (3), p. 509–527. DOI: 10.1016/j.ympev.2004.12.023.

TAN, DUN-YAN; LI, XIN-RONG; HONG, DE-YUAN (2007):
***Amana kuocangshanica* (Liliaceae), a new Species from South-east China.**
In: Botanical Journal of the Linnean Society 154 (3), p. 435–442. DOI: 10.1111/j.1095-8339.2007.00660.x.

TEKŞEN, MEHTAP; KARAMAN ERKUL, SEHER (2015):
**The Synopsis of the Genus *Gagea* (Liliaceae) in Turkey.**
In: Phytotaxa 230 (2), p. 101. DOI: 10.11646/phytotaxa.230.2.1.

TURKTAS, MINE; METIN, ÖZGE KARAKAŞ; BAŞTUĞ, BERK; ERTUĞRUL, FAHRIYE; SARAÇ, YASEMIN IZGI; KAYA, ERDAL (2013):
**Molecular phylogenetic Analysis of *Tulipa* (Liliaceae) based on noncoding Plastid and nuclear DNA Sequences with an Emphasis on Turkey.**
In: Botanical Journal of the Linnean Society 172 (3), p. 270–279. DOI: 10.1111/boj.12040.

WIETSMA, WILLEM A.; DEINUM, DANIËL; TEUNISSEN, HEDWICH A. S.; VAN DEN BERG, RONALD G. (2015):
**Phylogenetic Relationships within *Fritillaria* Section *Petilium* based on AFLP fingerprints.**
In: Plant Systematics and Evolution 301 (3), p. 1043–1054. DOI: 10.1007/s00606-014-1135-4.

ZARREI, MEHDI; WILKIN, PAUL; FAY, MICHAEL F.; INGROUILLE, MARTIN J.; ZARRE, SHAHIN; CHASE, MARK W. (2009):
**Molecular Systematics of *Gagea* and *Lloydia* (Liliaceae; Liliales): Implications of Analyses of nuclear ribosomal and Plastid DNA Sequences for infrageneric Classification.**
In: Annals of Botany 104 (1), p. 125–142. DOI: 10.1093/aob/mcp103.

ZHAO, LIHUA; LIU, HUAMIN; CAI, GUANGZE; XIA, MINGZHONG (2014):
**Assessment of the genetic Diversity and genetic Relationships of *Lilium* in China using ISSR Markers.**
In: Biochemical Systematics and Ecology 55, p. 184–189. DOI: 10.1016/j.bse.2014.03.024.

ZONNEVELD, BEN J. M. (2009):
**The systematic value of nuclear Genome size for "all" Species of *Tulipa* L. (Liliaceae).**
In: Plant Systematics and Evolution 281 (1-4), p. 217–245. DOI: 10.1007/s00606-009-0203-7.

### Liliales

GIVNISH, THOMAS J.; ZULUAGA, ALEJANDRO; MARQUES, ISABEL; LAM, VIVIENNE K. Y.; GOMEZ, MARYBEL SOTO; ILES, WILLIAM J. D. ET AL. (2016):
**Phylogenomics and historical Biogeography of the Monocot Order Liliales: out of Australia and through Antarctica.**
In: Cladistics 32 (6), p. 581–605. DOI: 10.1111/cla.12153.

KIM, JUNG SUNG; HONG, JEONG-KI; CHASE, MARK W.; FAY, MICHAEL F.; KIM, JOO-HWAN (2013):
**Familial Relationships of the Monocot Order Liliales based on a molecular phylogenetic Analysis using four Plastid Loci: MatK, rbcl, atpB and atpF.**
In: Botanical Journal of the Linnean Society 172 (1), p. 5–21. DOI: 10.1111/boj.12039.

PATTERSON, THOMAS B.; GIVNISH, THOMAS J. (2002):
**Phylogeny, concerted Convergence, and phylogenetic Niche Conservatism in the Core Liliales: Insights from rbcL and ndhF Sequence Data.**
In: Evolution 56 (2), p. 233–252. DOI: 10.1554/0014-3820(2002)056[0233:PCCAPN]2.0.CO;2.

PETERSEN, GITTE; SEBERG, OLE; DAVIS, JERROLD I. (2013):
**Phylogeny of the Liliales (Monocotyledons) with special Emphasis on Data Partition congruence and RNA editing.**
In: Cladistics 29 (3), p. 274–295. DOI: 10.1111/j.1096-0031.2012.00427.x.

### Limnanthaceae

MEYERS, STEPHEN C.; LISTON, AARON; MEINKE, ROBERT (2010):
**A molecular Phylogeny of *Limnanthes* (Limnanthaceae) and Investigation of an anomalous *Limnanthes* Population from California, U.S.A.**
In: Systematic Botany 35 (3), p. 552–558. DOI: 10.1600/036364410792495854.

### Linaceae

MCDILL, JOSHUA R.; REPPLINGER, MIRIAM; SIMPSON, BERYL B.; KADEREIT, JOACHIM W. (2009):
**The Phylogeny of *Linum* and Linaceae Subfamily Linoideae, with Implications for their Systematics, Biogeography, and Evolution of Heterostyly.**
In: Systematic Botany 34 (2), p. 386–405. DOI: 10.1600/036364409788606244.

MCDILL, JOSHUA R.; SIMPSON, BERYL B. (2011):
**Molecular Phylogenetics of Linaceae with complete generic Sampling and Data from two Plastid Genes.**
In: Botanical Journal of the Linnean Society 165, p. 64–83.

SVEINSSON, SAEMUNDUR; MCDILL, JOSHUA R.; WONG, GANE KA-SHU; LI, JUANJUAN; LI, XIA; DEYHOLOS, MICHAEL K.; CRONK, QUENTIN C.B. (2014):
**Phylogenetic pinpointing of a Paleopolyploidy event within the flax Genus (*Linum*) using Transcriptomics.**
In: Annals of Botany 113 (5), p. 753–761. DOI: 10.1093/aob/mct306.

### Linderniaceae

FISCHER, EBERHARD; SCHÄFERHOFF, BASTIAN; MÜLLER, KAI F. (2013):
**The Phylogeny of Linderniaceae — the new Genus *Linderniella*, and new Combinations within *Bonnaya, Craterostigma, Lindernia, Micranthemum, Torenia* and *Vandellia*.**
In: Willdenowia 43 (2), p. 209–238. DOI: 10.3372/wi.43.43201.

LIANG, YI-SHUO; WANG, JENN-CHE (2014):
**A systematic Study of *Bonnaya* Section *Bonnaya* (Linderniaceae).**
In: Australian Systematic Botany 27 (3), p. 180–198. DOI: 10.1071/SB14002.

LIANG, YI-SHUO; WANG, JENN-CHE (2016):
**New Combinations in *Craterostigma* and *Torenia* (Linderniaceae) from Asia with key to allied Species.**
In: Phytotaxa 247 (2), p. 148. DOI: 10.11646/phytotaxa.247.2.8.

MENDONÇA DE ALMEIDA, ERTON; WANDERLEY, ARTUR MAIA; NOLLET, FELIPE; COSTA, FABIANE R.; SOUZA, LUIZ GUSTAVO R.; FELIX, LEONARDO P. (2016):
**A new Species of *Ameroglossum* (Scrophulariaceae) Growing on Inselbergs in Northeastern Brazil.**
In: Systematic Botany 41 (2), p. 423–429. DOI: 10.1600/036364416X691740.

RAHMANZADEH, R.; MÜLLER, KAI F.; FISCHER, EBERHARD; BARTELS, D.; BORSCH, THOMAS (2005):
**The Linderniaceae and Gratiolaceae are further Lineages distinct from the Scrophulariaceae (Lamiales).**
In: Plant Biology 7 (1), p. 67–78. DOI: 10.1055/s-2004-830444.

SCHÄFERHOFF, BASTIAN; FLEISCHMANN, ANDREAS; FISCHER, EBERHARD; ALBACH, DIRK C.; BORSCH, THOMAS; HEUBL, GÜNTHER; MÜLLER, KAI F. (2010):
**Towards resolving Lamiales Relationships: Insights from rapidly evolving Chloroplast Sequences.**
In: BMC Evolutionary Biology 10, p. 352. DOI: 10.1186/1471-2148-10-352.

### Lindsaeaceae

LEHTONEN, SAMULI; TUOMISTO, HANNA; ROUHAN, GERMINAL; CHRISTENHUSZ, MAARTEN J.M. (2010):
**Phylogenetics and Classification of the pantropical Fern Family Lindsaeaceae.**
In: Botanical Journal of the Linnean Society 163, p. 305–359.

PERRIE, LEON R.; SHEPHERD, LARA D.; THOUVENOT, L.; KONRAT, M. (2014):
**Chloroplast DNA Sequences support the Transfer of the New Caledonian endemic Fern *Sphenomeris alutacea* to *Odontosoria*.**
In: New Zealand Journal of Botany 52 (3), p. 310–314. DOI: 10.1080/0028825X.2014.907816.

### Linnaeaceae

JACOBS, BART; PYCK, NANCY; SMETS, ERIK (2010):
**Phylogeny of the *Linnaea* Clade: are *Abelia* and *Zabelia* closely related?**
In: Molecular Phylogenetics and Evolution 57 (2), p. 741–752. DOI: 10.1016/j.ympev.2010.08.007.

### Loasaceae

BROKAW, JOSHUA M.; HUFFORD, LARRY (2010):
**Origins and introgression of polyploid Species in *Mentzelia* Section *Trachyphytum* (Loasaceae).**
In: American Journal of Botany 97 (9), p. 1457–1473. DOI: 10.3732/ajb.0900388.

HUFFORD, LARRY; MCMAHON, MICHELLE M.; O'QUINN, ROBIN L.; POSTON, MURIEL E. (2005):
**A phylogenetic Analysis of Loasaceae Subfamily Loasoideae based on Plastid DNA Sequences.**
In: International Journal of Plant Sciences (166), p. 289–300.

HUFFORD, LARRY; MCMAHON, MICHELLE M.; SHERWOOD, ANNA M.; REEVES, GAIL; CHASE, MARK W. (2003):
**The major Clades of Loasaceae: phylogenetic Analysis using the Plastid matK and trnL-trnF Regions.**
In: American Journal of Botany 90 (8), p. 1215–1228. DOI: 10.3732/ajb.90.8.1215.

MOODY, MICHAEL L.; HUFFORD, LARRY; SOLTIS, DOUGLAS E.; SOLTIS, PAMELA P. (2001):
**Phylogenetic Relationships of Loasaceae Subfamily Gronovioideae inferred from matK and ITS Sequence Data.**
In: American Journal of Botany 88 (2), p. 326–336. DOI: 10.2307/2657022.

SCHENK, JOHN J.; HUFFORD, LARRY (2011):

**Phylogeny and Taxonomy of *Mentzelia* Section *Bartonia* (Loasaceae).**
In: Systematic Botany 36 (3), p. 711–720. DOI: 10.1600/036364411X583673.

STRELIN, MARINA MICAELA; ARROYO, JOSÉ IGNACIO; FLIESSWASSER, STELLA; ACKERMANN, MARKUS (2017):
**Diversification of *Caiophora* (Loasaceae Subfam. Loasoideae) during the Uplift of the Central Andes.**
In: Organisms Diversity and Evolution 17 (1), p. 29–41. DOI: 10.1007/s13127-016-0312-4.

WEIGEND, MAXIMILIAN (2006):
**Validating Subfamily, Genus and Species Names in Loasaceae (Cornales).**
In: Taxon 55 (2), p. 463–468.

WEIGEND, MAXIMILIAN (2007):
**Systematics of the Genus *Mentzelia* (Loasaceae) in South America.**
In: Annals of the Missouri Botanical Garden 94 (3), p. 655–689. DOI: 10.3417/0026-6493(2007)94[655:SOTGML]2.0.CO;2.

### Loganiaceae

BACKLUND, MARIA; OXELMAN, BENGT; BREMER, BIRGITTA (2000):
**Phylogenetic Relationships within the Gentianales based on Ndhf and Rbcl Sequences, with particular Reference to the Loganiaceae.**
In: American Journal of Botany 87 (7), p. 1029–1043.

FOSTER, CHARLES S.P.; CONN, BARRY J.; HENWOOD, MURRAY J.; HO, SIMON Y.W. (2014):
**Molecular Data support *Orianthera*: a new Genus of Australian Loganiaceae.**
In: Telopea 16, p. 149–158.

FOSTER, CHARLES S.P.; HO, SIMON Y.W.; CONN, BARRY J.; HENWOOD, MURRAY J. (2014):
**Molecular Systematics and Biogeography of *Logania* R.Br. (Loganiaceae).**
In: Molecular Phylogenetics and Evolution 78, p. 324–333. DOI: 10.1016/j.ympev.2014.06.001.

GIBBONS, KERRY L.; HENWOOD, MURRAY J.; CONN, BARRY J. (2012):
**Phylogenetic Relationships in Loganieae (Loganiaceae) inferred from nuclear ribosomal and Chloroplast DNA Sequence Data.**
In: Australian Systematic Botany 25 (5), p. 331–340. DOI: 10.1071/SB12002.

GRANT, JASON R. (2009):
**A Revision of Neotropical *Bonyunia* (Loganiaceae: Antonieae).**
In: Annals of the Missouri Botanical Garden 96 (4), p. 541–563. DOI: 10.3417/2006135.

### Lomariopsidaceae

CHRISTENHUSZ, MAARTEN J.M.; JONES, MIRKKA; LEHTONEN, SAMULI (2013):
**Phylogenetic Placement of the enigmatic Fern Genus *Dracoglossum*.**
In: American Fern Journal 103 (2), p. 131–138. DOI: 10.1640/0002-8444-103.2.131.

HENNEQUIN, SABINE; HOVENKAMP, PETER H.; CHRISTENHUSZ, MAARTEN J.M.; SCHNEIDER, HARALD (2010):
**Phylogenetics and Biogeography of *Nephrolepis* – a tale of old Settlers and young Tramps.**
In: Botanical Journal of the Linnean Society 164, p. 113–127.

### Loranthaceae

AMICO, GUILLERMO C.; VIDAL-RUSSELL, ROMINA (2007):
**Phylogenetic Relationships and ecological Speciation in the Mistletoe *Tristerix* (Loranthaceae): the Influence of Pollinators, Dispersers, and Hosts.**
In: American Journal of Botany 94 (4), p. 558–567.

**Caires, Claudenir Simões; Azevedo, Cecília Oliveira de (2015):**

***Peristethium phaneroneurum*** **(Loranthaceae): a new Combination expands the Distribution of the Genus from Honduras to Brazil.**

In: Rodriguésia 66 (3), p. 859–861. DOI: 10.1590/2175-7860201566313.

**Kim, Chan-Soo; Kim, Soo-Young; Sun, Byung-Yun; Yi, Jae Seon (2013):**

**A Review of the taxonomic and ecological characteristics of Korean mistletoe types (*Viscum, Korthalsella, Loranthus* and *Taxillus*).**

In: Korean Journal of Plant Taxonomy 43 (2), p. 81–89. DOI: 10.11110/kjpt.2013.43.2.81.

**Kuijt, Job (2012):**

**Reinstatement and Expansion of the Genus *Peristethium* (Loranthaceae).**

In: Annals of the Missouri Botanical Garden 98 (4), p. 542–577. DOI: 10.3417/2010121.

**Kuijt, Job (2014):**

**A Monograph of the Genus *Aetanthus* (Loranthaceae).**

In: Plant Diversity and Evolution 131 (1), p. 1–51. DOI: 10.1127/1869-6155/2014/0131-0063.

**Liu, Bing; Le, Chi-Toan; Barrett, Russell L.; Nickrent, Daniel Lee; Chen, Zhi-Duan; Lu, Limin; Vidal-Russell, Romina (2018):**

**Historical Biogeography of Loranthaceae (Santalales): Diversification agrees with Emergence of tropical Forests and Radiation of Songbirds.**

In: Molecular Phylogenetics and Evolution 124, p. 199–212. DOI: 10.1016/j.ympev.2018.03.010.

**Vidal-Russell, Romina; Nickrent, Daniel Lee (2007):**

**A molecular Phylogeny of the Feathery Mistletoe *Misodendrum*.**

In: Systematic Botany 32 (3), p. 560–568.

**Vidal-Russell, Romina; Nickrent, Daniel Lee (2008):**

**Evolutionary Relationships in the showy Mistletoe Family (Loranthaceae).**

In: American Journal of Botany 95 (8), p. 1015–1029. DOI: 10.3732/ajb.0800085.

**Wilson, Carol A. (2006):**

**An Origin of Aerial Branch Parasitism in the Mistletoe Family, Loranthaceae.**

In: American Journal of Botany 93 (5), p. 787–796.

### Lowiaceae

**Johansen, Louise B. (2005):**

**Phylogeny of *Orchidantha* (Lowiaceae) and the Zingiberales based on six DNA Regions.**

In: Systematic Botany 30 (1), p. 106–117.

### Lycopodiaceae

**Aagaard, Sunniva M.D.; Greilhuber, Johann; Zhang, Xian-Chun; Wikström, Niklas (2009):**

**Occurrence and evolutionary Origins of polyploids in the clubmoss Genus *Diphasiastrum* (Lycopodiaceae).**

In: Molecular Phylogenetics and Evolution 52 (3), p. 746–754. DOI: 10.1016/j.ympev.2009.05.004.

**Aagaard, Sunniva M.D.; Vogel, Johannes C.; Wikström, Niklas (2009):**

**Resolving maternal Relationships in the Clubmoss Genus *Diphasiastrum* (Lycopodiaceae).**

In: Taxon 58 (3), p. 835–848.

**Field, Ashley R.; Testo, Weston; Bostock, Peter D.; Holtum, Joseph A. M.; Waycott, Michelle (2016):**

**Molecular Phylogenetics and the Morphology of the Lycopodiaceae Subfamily Huperzioideae supports three Genera: *Huperzia, Phlegmariurus* and *Phylloglossum*.**
In: Molecular Phylogenetics and Evolution 94 (Pt B), p. 635–657. DOI: 10.1016/j.ympev.2015.09.024.

**Ji, Sheng-Guo; Huo, Ke-Ke; Wang, Jun; Pan, Sheng-Li (2008):**
**A molecular phylogenetic Study of Huperziaceae based on Chloroplast rbcL and psbA-trnH Sequences.**
In: Journal of Systematics and Evolution 46 (2), p. 213–219.

**Øllgaard, Benjamin (2012):**
**Nomenclatural Changes in Brazilian Lycopodiaceae.**
In: Rodriguésia 63 (2), p. 479–482.

**Øllgaard, Benjamin; Windisch, Paulo G. (2014):**
**Lycopodiaceae in Brazil. Conspectus of the Family. I. the Genera *Lycopodium, Austrolycopodium, Diphasium,* and *Diphasiastrum*.**
In: Rodriguésia 65 (2), p. 293–309.

**Wikström, Niklas; Kenrick, Paul (2001):**
**Evolution of Lycopodiaceae (Lycopsida): estimating Divergence times from rbcL Gene Sequences by use of nonparametric Rate Smoothing.**
In: Molecular Phylogenetics and Evolution 19 (2), p. 177–186. DOI: 10.1006/mpev.2001.0936.

### Lygodiaceae

**Madeira, Paul T.; Pemberton, Robert W.; Center, Ted D. (2008):**
**A molecular Phylogeny of the Genus *Lygodium* (Schizaeaceae) with special Reference to the biological Control and Host Range Testing of *Lygodium microphyllum*.**
In: Biological Control 45 (3), p. 308–318. DOI: 10.1016/j.biocontrol.2008.02.004.

### Lythraceae

**Barber, Janet C.; Ghebretinsae, Amanuel G.; Graham, Shirley A. (2010):**
**An expanded Phylogeny of *Cuphea* (Lythraceae) and a North American Monophyly.**
In: Plant Systematics and Evolution 289 (1-2), p. 35–44. DOI: 10.1007/s00606-010-0329-7.

**Graham, Shirley A. (2010):**
**Revision of the Caribbean Genus *Ginoria* (Lythraceae), including Haitia from Hispaniola.**
In: Annals of the Missouri Botanical Garden 97 (1), p. 34–90. DOI: 10.3417/2007028.

**Graham, Shirley A.; Diazgranados, Mauricio; Barber, Janet C. (2011):**
**Relationships among the confounding Genera *Ammannia, Hionanthera, Nesaea* and *Rotala* (Lythraceae).**
In: Botanical Journal of the Linnean Society 166 (1), p. 1–19. DOI: 10.1111/j.1095-8339.2011.01126.x.

**Graham, Shirley A.; Freudenstein, John V.; Luker, Melissa (2006):**
**A phylogenetic Study of *Cuphea* (Lythraceae) based on Morphology and Nuclear rDNA ITS Sequences.**
In: Systematic Botany 31 (4), p. 764–778.

**Graham, Shirley A.; Hall, Jocelyn C.; Sytsma, Kenneth J.; Shi, Su-hua (2005):**
**Phylogenetic Analysis of the Lythraceae based on four Gene Regions and Morphology.**
In: International Journal of Plant Sciences 166 (6), p. 996–1017.

**Huang, Ye-Lin; Shi, Su-hua (2002):**

**Phylogenetics of Lythraceae sensu lato: A Preliminary Analysis based on Chloroplast rbcL Gene, psaA-ycf3 Spacer, and Nuclear rDNA Internal Transcribed Spacer (ITS) Sequences.**
In: International Journal of Plant Sciences 163 (2), p. 215–225.

YANG, YU-CHEN; YANG, SHU-HUAN; FANG, LU; LI, JIAN-FANG; ZHONG, CAI-RONG; ZHOU, REN-CHAO; SHI, SU-HUA (2015):
**Phylogenetic Position of *Sonneratia griffithii* based on Sequences of the nuclear ribosomal Internal Transcribed Spacer and 13 nuclear Genes.**
In: Journal of Systematics and Evolution 53 (1), p. 47–52. DOI: 10.1111/jse.12102.

### Maesaceae

ANDERBERG, ARNE A.; STÅHL, BERTIL; KÄLLERSJÖ, MARI (2000):
**Maesaceae, a new Primuloid Family in the Order Ericales s.l.**
In: Taxon 49 (2), p. 183–187.

MA, O. P. W.; SAUNDERS, RICHARD M.K. (2003):
**Comparative Floral ontogeny of *Maesa* (Maesaceae), *Aegiceras* (Myrsinaceae) and *Embelia* (Myrsinaceae): taxonomic and phylogenetic Implications.**
In: Plant Systematics and Evolution 243 (1-2), p. 39–58. DOI: 10.1007/s00606-003-0062-6.

UTTERIDGE, TIMOTHY M.A. (2012):
**Four new Species of *Maesa* Forssk. (Primulaceae) from Malesia.**
In: Kew Bulletin 67 (3), p. 367–378. DOI: 10.1007/s12225-012-9383-3.

### Magnoliaceae

AZUMA, HIROSHI; GARCÍA-FRANCO, JOSÉ G.; RICO-GRAY, VICTOR; THIEN, LEONARD B. (2001):
**Molecular Phylogeny of the Magnoliaceae: the Biogeography of Tropical and Temerate Disjunctions.**
In: American Journal of Botany 88 (12), p. 2275–2285.

KIM, SANGTAE; PARK, CHONG-WOOK; KIM, YOUNG-DONG; SUH, YOUNGBAE (2001):
**Phylogenetic Relationships in Family Magnoliaceae inferred from Ndhf Sequences.**
In: American Journal of Botany 88 (4), p. 717–728.

KUMAR, V.SAMPATH (2006):
**New Combinatuons and new Names in Asian Magnoliaceae.**
In: Kew Bulletin 61 (2), p. 183–186.

NIE, ZE-LONG; WEN, JUN; AZUMA, HIROSHI; QIU, YIN-LONG; SUN, HANG; MENG, YING ET AL. (2008):
**Phylogenetic and biogeographic complexity of Magnoliaceae in the Northern Hemisphere inferred from three nuclear Data Sets.**
In: Molecular Phylogenetics and Evolution 48 (3), p. 1027–1040. DOI: 10.1016/j.ympev.2008.06.004.

WANG, YA-LING (2006):
**The utility of matK Gene in the phylogenetic Analysis of the Genus *Magnolia*.**
In: Acta Phytotaxonomica Sinica 44 (2), p. 135. DOI: 10.1360/aps040013.

WANG, YA-LING; EJDER, ERLAND; YANG, JIAN-FEN; LIU, R. A.O.; YE, LI-MING; HE, ZI-CAN; ZHANG, SHOU-ZHOU (2013):
***Magnolia sinostellata* and relatives (Magnoliaceae).**
In: Phytotaxa 154 (1), p. 47. DOI: 10.11646/phytotaxa.154.1.3.

### Malesherbiaceae

GENGLER-NOWAK, KARLA M. (2003):
**Molecular Phylogeny and Taxonomy of Malesherbiaceae.**

In: Systematic Botany 28 (2), p. 333–344.

**Weigend, Maximilian; Jossberger, Thomas; Beltrán, Hamilton (2015):**

**Notes on *Malesherbia* (Passifloraceae) in Peru: a new Species from southern Peru, a new record and a first report on interspecific Hybridization in *Malesherbia*.**

In: Phytotaxa 202 (4), p. 250. DOI: 10.11646/phytotaxa.202.4.2.

### Malpighiaceae

**Davis, Charles C.; Anderson, William R. (2010):**

**A complete generic Phylogeny of Malpighiaceae inferred from nucleotide Sequence Data and Morphology.**

In: American Journal of Botany 97 (12), p. 2031–2048. DOI: 10.3732/ajb.1000146.

**Davis, Charles C.; Anderson, William R.; Donoghue, Michael J. (2001):**

**Phylogeny of Malpighiaceae: Evidence from Chloroplast ndhF and trnl-F Nucleotide Sequences.**

In: American Journal of Botany 88 (10), p. 1830–1846. DOI: 10.2307/3558360.

**Davis, Charles C.; Bell, Charles D.; Fritsch, Peter W.; Mathews, Sarah (2002):**

**Phylogeny of *Acridocarpus-Brachylophon* (Malpighiaceae): Implications for Tertiary Tropical Floras and Afroasian Biogeography.**

In: Evolution 56 (12), p. 2395–2405. DOI: 10.1554/0014-3820(2002)056[2395:POABMI]2.0.CO;2.

**Davis, Charles C.; Fritsch, Peter W.; Bell, Charles D.; Mathews, Sarah (2004):**

**High-Latitude Tertiary Migrations of an exclusively tropical Clade: Evidence from Malpighiaceae.**

In: International Journal of Plant Sciences 165 (S4), S107-S121.

### Malpighiales

**Davis, Charles C.; Chase, Mark W. (2004):**

**Elatinaceae are sister to Malpighiaceae; Peridiscaceae belong to Saxifragales.**

In: American Journal of Botany 91 (2), p. 262–273. DOI: 10.3732/ajb.91.2.262.

**Korotkova, Nadja; Schneider, Julio Valentin; Quandt, Dietmar; Worberg, Andreas; Zizka, Georg; Borsch, Thomas (2009):**

**Phylogeny of the Eudicot Order Malpighiales: Analysis of a Recalcitrant Clade with Sequences of the petD Group II Intron.**

In: Plant Systematics and Evolution 282 (3-4), p. 201–228. DOI: 10.1007/s00606-008-0099-7.

**Ruhfel, Bradley R.; Bittrich, Volker; Bove, Claudia P.; Gustafsson, Mats H.G.; Philbrick, C. Thomas; Rutishauser, Rolf et al. (2011):**

**Phylogeny of the clusioid Clade (Malpighiales): Evidence from the Plastid and mitochondrial Genomes.**

In: American Journal of Botany 98 (2), p. 306–325. DOI: 10.3732/ajb.1000354.

**Wurdack, Kenneth J.J.; Davis, Charles C. (2009):**

**Malpighiales Phylogenetics: Gaining Ground on one of the most Recalcitrant Clades in the Angiosperm Tree of Life.**

In: American Journal of Botany 96 (8), p. 1551–1570. DOI: 10.3732/ajb.0800207.

**Xi, Zhenxiang; Ruhfel, Bradley R.; Schaefer, Hanno; Amorim, André M.; Sugumaran, M.; Wurdack, Kenneth J.J. et al. (2012):**

**Phylogenomics and a posteriori Data Partitioning resolve the Cretaceous Angiosperm Radiation Malpighiales.**

In: Proceedings of the National Academy of Sciences of the United States of America 109 (43), p. 17519–17524. DOI: 10.1073/pnas.1205818109.

### Malvaceae

**Aguilar, Javier Fuertes; Fryxell, Paul A.; Jansen, Robert K. (2003):**
**Phylogenetic Relationships and Classification of the Sida Generic Alliance (Malvaceae) based on nrDNA ITS Evidence.**
In: Systematic Botany 28 (2), p. 352–364.

**Álvarez, Inés; Cronn, Richard C.; Wendel, Jonathan F. (2005):**
**Phylogeny of the New World diploid Cottons (*Gossypium* L., Malvaceae) based on Sequences of three low-copy nuclear Genes.**
In: Plant Systematics and Evolution 252 (3-4), p. 199–214. DOI: 10.1007/s00606-004-0294-0.

**Andreasen, Katarina (2012):**
**Phylogeny, Hybridization, and Evolution of Habit and Breeding System in *Sidalcea* and *Eremalche* (Malvaceae).**
In: International Journal of Plant Sciences 173 (5), p. 532–548. DOI: 10.1086/665269.

**Andreasen, Katarina; Baldwin, Bruce G. (2003):**
**Reexamination of Relationships, habital Evolution, and Phylogeography of checker Mallows (*Sidalcea*; Malvaceae) based on molecular phylogenetic Data.**
In: American Journal of Botany 90 (3), p. 436–444. DOI: 10.3732/ajb.90.3.436.

**Areces-Berazain, F.; Ackerman, J. D. (2016):**
**Phylogenetics, Delimitation and historical Biogeography of the pantropical Tree Genus *Thespesia* (Malvaceae, Gossypieae).**
In: Botanical Journal of the Linnean Society 181 (2), p. 171–198. DOI: 10.1111/boj.12414.

**Blanchard, Orland J. (2012):**
**Chromosome Numbers, Phytogeography, and Evolution in *Kosteletzkya* (Malvaceae).**
In: Rhodora 114 (957), p. 37–49. DOI: 10.3119/0035-4902-114.957.37.

**Bovini, Massimo Giuseppe; Baumgratz, José Fernando A. (2016):**
**Taxonomic Revision of *Wissadula* (Malvoideae, Malvaceae) in Brazil.**
In: Phytotaxa 243 (3), p. 201. DOI: 10.11646/phytotaxa.243.3.1.

**Cronn, Richard C.; Small, Randall L.; Haselkorn, Tamara; Wendel, Jonathan F. (2002):**
**Rapid Diversification of the cotton Genus (*Gossypium*: Malvaceae) revealed by Analysis of sixteen nuclear and Chloroplast Genes.**
In: American Journal of Botany 89 (4), p. 707–725. DOI: 10.3732/ajb.89.4.707.

**Donnell, Aliya A.; Cantino, Philip D.; Ballard, Harvey Eugene (2015):**
**A phenetic Analysis of the *Bakeridesia integerrima* complex (Malvaceae).**
In: Brittonia 67 (3), p. 250–271. DOI: 10.1007/s12228-015-9362-8.

**Escobar García, Pedro; Pakravan, Maneezheh; Schönswetter, Peter; Aguilar, Javier Fuertes; Schneeweiss, Gerald M. (2012):**
**Phylogenetic Relationships in the species-rich Irano-Turanian Genus *Alcea* (Malvaceae).**
In: Taxon 61 (2), p. 324–332. DOI: 10.1002/tax.612004.

**Escobar García, Pedro; Schönswetter, Peter; Fuertes-Aguilar, Javier; Nieto Feliner, Gonzalo; Schneeweiss, Gerald M. (2009):**

**Five molecular Markers reveal extensive morphological homoplasy and reticulate Evolution in the *Malva* alliance (Malvaceae).**
In: Molecular Phylogenetics and Evolution 50 (2), p. 226–239. DOI: 10.1016/j.ympev.2008.10.015.

GROVER, CORRINNE E.; GALLAGHER, JOSEPH P.; JARECZEK, JOSEF J.; PAGE, JUSTIN T.; UDALL, JOSHUA A.; GORE, MICHAEL A.; WENDEL, JONATHAN F. (2015):
**Re-evaluating the Phylogeny of allopolyploid *Gossypium* L.**
In: Molecular Phylogenetics and Evolution 92, p. 45–52. DOI: 10.1016/j.ympev.2015.05.023.

GROVER, CORRINNE E.; GRUPP, K. K.; WANZEK, R. J.; WENDEL, JONATHAN F. (2012):
**Assessing the Monophyly of polyploid *Gossypium* Species.**
In: Plant Systematics and Evolution 298 (6), p. 1177–1183. DOI: 10.1007/s00606-012-0615-7.

HEENAN, PETER B.; DAWSON, MURRAY I.; REDMOND, D. N.; WAGSTAFF, STEVEN J. (2005):
**Relationships of the New Zealand mountain Ribbonwoods (*Hoheria glabrata* and *H. lyallii*: Malvaceae), based on molecular and morphological Data.**
In: New Zealand Journal of Botany 43 (2), p. 527–549. DOI: 10.1080/0028825X.2005.9512973.

HUERTAS, MARILÚ L.; SCHNEIDER, JULIO VALENTIN; ZIZKA, GEORG (2007):
**Phylogenetic Analysis of *Palaua* (Malveae, Malvaceae) based on Plastid and Nuclear Sequences.**
In: Systematic Botany 32 (1), p. 157–165.

KOOPMAN, MARGARET M.; BAUM, DAVID A. (2008):
**Phylogeny and Biogeography of Tribe Hibisceae (Malvaceae) on Madagascar.**
In: Systematic Botany 33 (2), p. 364–374. DOI: 10.1600/036364408784571653.

KRAPOVICKAS, ANTONIO (2015):
**Notas sobre *Urocarpidium* y *Fuertesimalva* (Malvaceae).**
In: Bonplandia 24 (1), p. 43–48.

NEUBIG, KURT M.; JR., ORLAND J. BLANCHARD; WHITTEN, WILLIAM MARK; MCDANIEL, STUART F. (2015):
**Molecular Phylogenetics of *Kosteletzkya* (Malvaceae, Hibisceae) reveals multiple independent and successive polyploid Speciation Events.**
In: Botanical Journal of the Linnean Society 179, p. 421–435. DOI: 10.1111/boj.12330.

PAKRAVAN, MANEEZHEH; GHAREMAN, A. (2003):
**Some new Combinations and Synonyms in *Alcea* (Malvaceae) from Iran.**
In: Annalen des Naturhistorischen Museums in Wien, Serie B 104, p. 713–716.

PFEIL, BERNARD E.; CRISP, MICHAEL D. (2005):
**What to do with *Hibiscus*? A proposed nomenclatural resolution for a large and well known Genus of Malvaceae and comments on Paraphyly.**
In: Australian Systematic Botany 18 (1), p. 49–60. DOI: 10.1071/SB04024.

SATYA, PRATIK; KARAN, M.; KAR, C. S.; MAHAPATRA, A. K.; MAHAPATRA, B. P. (2013):
**Assessment of molecular Diversity and evolutionary Relationship of Kenaf (*Hibiscus cannabinus* L.), roselle (*H. sabdariffa* L.) and their wild Relatives.**
In: Plant Systematics and Evolution 299 (3), p. 619–629. DOI: 10.1007/s00606-012-0748-8.

SCHNEIDER, JULIO VALENTIN; SCHULTE, KATHARINA; AGUILAR, JAVIER FUERTES; HUERTAS, MARILÚ L. (2011):
**Molecular Evidence for Hybridization and introgression in the Neotropical coastal desert-endemic *Palaua* (Malveae, Malvaceae).**
In: Molecular Phylogenetics and Evolution 60 (3), p. 373–384. DOI: 10.1016/j.ympev.2011.05.010.

SMALL, L. (2004):

**Phylogeny of *Hibiscus* Sect. *Muenchhusia* (Malvaceae) based on Chloroplast rpL16 and ndhF, and Nuclear ITS and GBSSI Sequences.**
In: Systematic Botany 29 (2), p. 385–392. DOI: 10.1600/036364404774195575.

**Tabbasam, Nabila; Zafar, Yusuf; Mehboob-Ur-Rahman (2014):**
**Pros and Cons of using genomic SSRs and EST-SSRs for resolving Phylogeny of the Genus *Gossypium*.**
In: Plant Systematics and Evolution 300 (3), p. 559–575. DOI: 10.1007/s00606-013-0891-x.

**Tate, Jennifer A. (2011):**
**The Status of *Urocarpidium* (Malvaceae): Insight from nuclear and plastid-based Phylogenies.**
In: Taxon 60 (5), p. 1330–1338. DOI: 10.1002/tax.605009.

**Tate, Jennifer A.; Fuertes-Aguilar, Javier; Wagstaff, Steven J.; La Duke, John C.; Bodo Slotta, Tracey A.; Simpson, Beryl B. (2005):**
**Phylogenetic Relationships within the Tribe Malveae (Malvaceae, Subfamily Malvoideae) as inferred from ITS Sequence Data.**
In: American Journal of Botany 92 (4), p. 584–602. DOI: 10.3732/ajb.92.4.584.

**Wagstaff, Steven J.; Tate, Jennifer A. (2011):**
**Phylogeny and Character Evolution in the New Zealand endemic Genus *Plagianthus* (Malveae, Malvaceae).**
In: Systematic Botany 36 (2), p. 405–418. DOI: 10.1600/036364411X569589.

**Zhu, Huayu; Lv, Junhong; Zhao, Liang; Tong, Xiangchao; Zhou, Baoliang; Zhang, Tianzhen; Guo, Wangzhen (2012):**
**Molecular Evolution and phylogenetic Analysis of Genes related to Cotton Fibers Development from wild and domesticated Cotton Species in *Gossypium*.**
In: Molecular Phylogenetics and Evolution 63 (3), p. 589–597. DOI: 10.1016/j.ympev.2012.01.025.

### Malvales

**Alverson, William S.; Whitlock, Barbara A.; Nyffeler, Reto; Bayer, Clemens; Baum, David A. (1999):**
**Phylogeny of the core Malvales: Evidence from ndh F Sequence Data.**
In: American Journal of Botany 86 (10), p. 1474–1486. DOI: 10.2307/2656928.

**Nyffeler, Reto; Bayer, Clemens; Alverson, William S.; Yen, Alan C.; Whitlock, Barbara A.; Chase, Mark W.; Baum, David A. (2005):**
**Phylogenetic Analysis of the *Malvadendrina* Clade (Malvaceae s.l.) based on Plastid DNA Sequences.**
In: Organisms Diversity and Evolution 5 (2), p. 109–123. DOI: 10.1016/j.ode.2004.08.001.

### Marantaceae

**Andersson, Lennart; Chase, Mark W. (2001):**
**Phylogeny and Classification of Marantaceae.**
In: Botanical Journal of the Linnean Society 135 (3), p. 275–287. DOI: 10.1111/j.1095-8339.2001.tb01097.x.

**Ley, Alexandra C.; Claßen-Bockhoff, Regine (2011):**
**Evolution in African Marantaceae - Evidence from Phylogenetic, Ecological and morphological Studies.**
In: Systematic Botany 36 (2), p. 277–290. DOI: 10.1600/036364411X569480.

**Ley, Alexandra C.; Hardy, Olivier J. (2014):**

**Contrasting Patterns of Gene flow between sister plant Species in the Understorey of African moist Forests - the Case of sympatric and parapatric Marantaceae Species.**
In: Molecular Phylogenetics and Evolution 77, p. 264–274. DOI: 10.1016/j.ympev.2014.04.026.

PISCHTSCHAN, ELKE; LEY, ALEXANDRA C.; CLAßEN-BOCKHOFF, REGINE (2010):
**Ontogenetic and phylogenetic Diversification of the hooded staminode in Marantaceae.**
In: Taxon 59 (4), p. 1111–1125.

PRINCE, LINDA M.; KRESS, W. JOHN (2006):
**Phylogenetic Relationships and Classification in Marantaceae: Insights from Plastid DNA Sequence Data.**
In: Taxon 55 (2), p. 281. DOI: 10.2307/25065578.

SUKSATHAN, PIYAKASET; GUSTAFSSON, MATS H.G.; BORCHSENIUS, FINN (2009):
**Phylogeny and generic Delimitation of Asian Marantaceae.**
In: Botanical Journal of the Linnean Society 159 (3), p. 381–395. DOI: 10.1111/j.1095-8339.2009.00949.x.

VIEIRA, SILVANA; MAAS, PAUL J.M.; BORCHSENIUS, FINN (2012):
**Taxonomic Revision of *Myrosma* (Marantaceae).**
In: Blumea 57 (2), p. 125–130. DOI: 10.3767/000651912X654218.

VIEIRA, SILVANA; SOUZA, VINICIUS CASTRO (2008):
**Four new Species of *Maranta* L. (Marantaceae) from Brazil.**
In: Botanical Journal of the Linnean Society 158 (1), p. 131–139. DOI: 10.1111/j.1095-8339.2008.00852.x.

### Marattiaceae

CHRISTENHUSZ, MAARTEN J.M. (2010):
***Danaea* (Marattiaceae) revisited: Biodiversity, a new Classification and ten new Species of a Neotropical Fern Genus.**
In: Botanical Journal of the Linnean Society 163, p. 360–385.

CHRISTENHUSZ, MAARTEN J.M. (2010):
**Revision of the Neotropical Fern Genus *Eupodium* (Marattiaceae).**
In: Kew Bulletin 65, p. 115–121.

CHRISTENHUSZ, MAARTEN J.M.; TUOMISTO, HANNA; METZGAR, JORDAN S.; PRYER, KATHLEEN M. (2008):
**Evolutionary Relationships within the Neotropical, Eusporangiate Fern Genus *Danaea* (Marattiaceae).**
In: Molecular Phylogenetics and Evolution 46 (1), p. 34–48. DOI: 10.1016/j.ympev.2007.09.015.

MURDOCK, ANDREW G. (2008):
**A taxonomic Revision of the Eusporangiate Fern Family Marattiaceae, with Description of a new Genus *Ptisana*.**
In: Taxon 57 (3), p. 737–755.

MURDOCK, ANDREW G. (2008):
**Phylogeny of Marattioid Ferns (Marattiaceae): inferring a Root in the Absence of a closely related Outgroup.**
In: American Journal of Botany 95 (5), p. 626–641. DOI: 10.3732/ajb.2007308.

### Marcgraviaceae

WARD, N. MISA; PRICE, ROBERT A. (2002):
**Phylogenetic Relationships of Marcgraviaceae: Insights from three Chloroplast Genes.**

In: Systematic Botany 27 (1), p. 149–160.

### Marsileaceae

NAGALINGUM, NATHALIE S.; SCHNEIDER, HARALD; PRYER, KATHLEEN M. (2007):
**Molecular phylogenetic Relationships and morphological Evolution in the Heterosporous Fern Genus *Marsilea*.**
In: Systematic Botany 32 (1), p. 16–25. DOI: 10.1600/036364407780360256.

### Martyniaceae

GUTIERREZ, RAUL (2008):
**Preliminary Chloroplast DNA Studies in the Flowering Plant Family Martyniaceae (Order Lamiales).**
In: Journal of the Arizona-Nevada Academy of Science 40 (1), p. 105–110. DOI: 10.2181/1533-6085(2008)40[105:PCDSIT]2.0.CO;2.

### Maundiaceae

SOKOLOFF, DMITRY D.; MERING, SABINE; JACOBS, SURREY W.L.; REMIZOWA, MARGARITA V. (2013):
**Morphology of Maundia supports its isolated phylogenetic Position in the early-divergent Monocot Order Alismatales.**
In: Botanical Journal of the Linnean Society 173 (1), p. 12–45. DOI: 10.1111/boj.12068.

### Melanthiaceae

FARMER, SUSAN N.; SCHILLING, EDWARD E. (2002):
**Phylogenetic Analyses of Trilliaceae based on morphological and molecular Data.**
In: Systematic Botany 27 (4), p. 674–692.

FUSE, SHIZUKA; SOOK LEE, NAM N.; TAMURA, MINORU N. (2004):
**Biosystematic studies on the Genus *Heloniopsis* (Melanthiaceae) II. Two new Species from Korea based on morphological and molecular Evidence.**
In: Taxon 53 (4), p. 949–958. DOI: 10.2307/4135562.

HSU, TSAI-WEN; KONO, YOSHIKO; CHIANG, TZEN-YUH; PENG, CHING-I. (2011):
***Ypsilandra* (Melanthiaceae; Liliaceae sensu lato), a new generic record for Taiwan.**
In: Botanical Studies 52, p. 99–104.

JI, YUNHENG; FRITSCH, PETER W.; LI, HENG; XIAO, TIAOJIANG; ZHOU, ZHEKUN (2006):
**Phylogeny and Classification of *Paris* (Melanthiaceae) inferred from DNA Sequence Data.**
In: Annals of Botany 98 (1), p. 245–256. DOI: 10.1093/aob/mcl095.

KIM, JIN OHK; TAMURA, MINORU N.; FUSE, SHIZUKA; LEE, NAM SOOK (2014):
**Taxonomic Status and Phylogeny of *Veratrum* Section *Veratrum* (Melanthiaceae) in Korea and Japan based on Chloroplast and nuclear Sequence Data.**
In: Plant Systematics and Evolution 300 (1), p. 75–89. DOI: 10.1007/s00606-013-0861-3.

KIM, JUNG SUNG; BODIN, SARAH S.; KIM, JOO-HWAN (2015):
**Complete Chloroplast Genome of *Chionographis japonica* (Willd.) Maxim. (Melanthiaceae): Comparative Genomics and Evaluation of Universal Primers for Liliales.**
In: Plant Molecular Biology Reporter. DOI: 10.13140/2.1.4598.6249.

KIM, SANG-CHUL; KIM, JUNG SUNG; CHASE, MARK W.; FAY, MICHAEL F.; KIM, JOO-HWAN (2016):

**Molecular phylogenetic Relationships of Melanthiaceae (Liliales) based on Plastid DNA Sequences.**
In: Botanical Journal of the Linnean Society 181 (4), p. 567–584. DOI: 10.1111/boj.12405.

**Liao, Wan-Jin; Yuan, Yong-Ming; Zhang, D.-Y. (2007):**
**Biogeography and Evolution of flower color in *Veratrum* (Melanthiaceae) through Inference of a Phylogeny based on multiple DNA Markers.**
In: Plant Systematics and Evolution 267 (1-4), p. 177–190. DOI: 10.1007/s00606-007-0528-z.

**Osaloo, Shahrokh Kazempour; Kawano, Shoichi (1999):**
**Molecular Systematics of Trilliaceae II. phylogenetic Analyses of *Trillium* and its allies using Sequences of rbcL and matK Genes of cp DNA and internal transcribed Spacers of 18s-26s nrDNA.**
In: Plant Species Biology 14 (1), p. 75–94. DOI: 10.1046/j.1442-1984.1999.00009.x.

**Pellicer, Jaume; Kelly, Laura J.; Leitch, Ilia J.; Zomlefer, Wendy B.; Fay, Michael F. (2014):**
**A universe of dwarfs and giants: Genome size and Chromosome Evolution in the Monocot Family Melanthiaceae.**
In: the new Phytologist 201 (4), p. 1484–1497. DOI: 10.1111/nph.12617.

**Schilling, Edward E.; Floden, Aaron; Farmer, Susan B. (2013):**
**A new sessile-flowered *Trillium* Species from Tennessee.**
In: Castanea 78 (2), p. 140–147. DOI: 10.2179/12-043.

**Tanaka, Noriyuki (2017):**
**A Synopsis of the Genus *Chamaelirium* (Melanthiaceae) with a new infrageneric Classification including *Chionographis*.**
In: Taiwania 62 (2), p. 157–167.

**Zomlefer, Wendy B.; Judd, Walter p. (2008):**
**Two new Species of *Schoenocaulon* (Liliales: Melanthiaceae) from Mexico supported by ITS Sequence Data.**
In: Systematic Botany 33 (1), p. 117–124. DOI: 10.1600/036364408783887474.

**Zomlefer, Wendy B.; Whitten, William Mark; Williams, Norris H.; Judd, Walter p. (2003):**
**An Overview of *Veratrum* s.l. (Liliales: Melanthiaceae) and an Infrageneric Phylogeny based on ITS Sequence Data.**
In: Systematic Botany 28 (2), p. 250–269.

**Zomlefer, Wendy B.; Whitten, William Mark; Williams, Norris H.; Judd, Walter p. (2006):**
**Infrageneric Phylogeny of *Schoenocaulon* (Liliales: Melanthiaceae) with Clarification of cryptic Species based on ITS Sequence Data and geographical Distribution.**
In: American Journal of Botany 93 (8), p. 1178–1192. DOI: 10.3732/ajb.93.8.1178.

**Zomlefer, Wendy B.; Williams, Norris H.; Whitten, William Mark; Judd, Walter p. (2001):**
**Generic Circumscription and Relationships in the Tribe Melanthieae (Liliales, Melanthiaceae), with Emphasis on *Zigadenus* : Evidence from ITS and trnL-F Sequence Data.**
In: American Journal of Botany 88 (9), p. 1657–1669. DOI: 10.2307/3558411.

### Melastomataceae

**Almeda, Frank (1993):**
***Stanmarkia*, a new Genus of Melastomataceae from the Volcanic Highlands of Western Guatemala and Adjacent Mexico.**

In: Brittonia 45 (3), p. 187–203. DOI: 10.2307/2807100.

**AMORIM, ANDRÉ M.; GOLDENBERG, RENATO; MICHELANGELI, FABIÁN ARMANDO (2009):**
**A new Species of *Physeterostemon* (Melastomataceae) from Bahia, Brazil, with Notes on the Phylogeny of the Genus.**
In: Systematic Botany 34 (2), p. 324–329. DOI: 10.1600/036364409788606389.

**CÁMARA-LERET, R.; RIDDER-NUMAN, J.W.A.; VELDKAMP, J. F. (2013):**
**Revision of *Heteroblemma* gen. nov. (Dissochaeteae – Melastomataceae ) from Malesia and Vietnam.**
In: Blumea 58 (3), p. 229–240. DOI: 10.3767/000651913X674945.

**CÁMARA-LERET, R.; VELDKAMP, J. F. (2011):**
**A remarkable new *Medinilla* (Melastomataceae) from Celebes (Sulawesi), Indonesia.**
In: Gardens' Bulletin Singapore 62 (2), p. 213–221.

**CELLINESE, NICOLETTA; RENNER, SUSANNE P. (1997):**
**New Species and new Combinations in *Sonerila* and *Phyllagathis* (Melastomataceae) from Thailand.**
In: Novon: A Journal for Botanical Nomenclature 7, p. 106–112.

**CLAUSING, G. (2001):**
**Evolution of Growth form in epiphytic Dissochaeteae (Melastomataceae).**
In: Organisms Diversity and Evolution 1 (1), p. 45–60. DOI: 10.1078/1439-6092-00004.

**CLAUSING, G.; RENNER, SUSANNE P. (2001):**
**Molecular Phylogenetics of Melastomataceae and Memecylaceae: Implications for Character Evolution.**
In: American Journal of Botany 88 (3), p. 486–498. DOI: 10.2307/2657114.

**FIDANZA, KARINA; ALMEDA, FRANK (2011):**
**New Species of the Brazilian Endemic Genus *Cambessedesia* (Melastomataceae).**
In: Harvard Papers in Botany 16 (1), p. 57–63. DOI: 10.3100/025.016.0109.

**FRITSCH, PETER W.; ALMEDA, FRANK; RENNER, SUSANNE S.; MARTINS, ANGELA B.; CRUZ, BONI C. (2004):**
**Phylogeny and Circumscription of the near-endemic Brazilian Tribe Microlicieae (Melastomataceae).**
In: American Journal of Botany 91 (7), p. 1105–1114. DOI: 10.3732/ajb.91.7.1105.

**GOLDENBERG, RENATO; ALMEDA, FRANK; CADDAH, MAYARA KRASINSKI; MARTINS, ANGELA B.; MEIRELLES, JULIA; MICHELANGELI, FABIÁN ARMANDO; WEISS, MARKUS (2013):**
**Nomenclator Botanicus for the Neotropical Genus *Miconia* (Melastomataceae: Miconieae).**
In: Phytotaxa 106 (1), p. 1. DOI: 10.11646/phytotaxa.106.1.1.

**GOLDENBERG, RENATO; AMORIM, ANDRÉ M. (2006):**
***Physeterostemon* (Melastomataceae): a new Genus and two new Species from the Bahian Atlantic Forest, Brazil.**
In: Taxon 55 (4), p. 965–972. DOI: 10.2307/25065690.

**GOLDENBERG, RENATO; FRAGA, CLAUDIO N.; FONTANA, ANDRÉ P.; NICOLAS, ANTOINE N.; MICHELANGELI, FABIÁN ARMANDO (2012):**
**Taxonomy and Phylogeny of *Merianthera* (Melastomataceae).**
In: Taxon 61 (5), p. 1040–1056. DOI: 10.1002/tax.615010.

**Goldenberg, Renato; Penneys, Darin S.; Almeda, Frank; Judd, Walter S.; Michelangeli, Fabián Armando (2008):**
**Phylogeny of *Miconia* (Melastomataceae): Patterns of Stamen Diversification in a Megadiverse Neotropical Genus.**
In: International Journal of Plant Sciences 169 (7), p. 963–979. DOI: 10.1086/589697.

**Ionta, Gretchen M.; Judd, Walter S.; Williams, Norris H.; Whitten, William Mark (2007):**
**Phylogenetic Relationships in *Rhexia* (Melastomataceae): Evidence from DNA Sequence Data and Morphology.**
In: International Journal of Plant Sciences 168 (7), p. 1055–1066.

**Jr., Jacinto C. Regalado (1990):**
**Revision of *Medinilla* (Melastomataceae) of Borneo.**
In: Blumea 35, p. 5–70.

**Kartonegoro, A.; Veldkamp, J. F. (2013):**
**Revision of *Creochiton* (Melastomataceae ).**
In: Blumea 58 (3), p. 217–227. DOI: 10.3767/000651913X674134.

**Kriebel, Ricardo (2016):**
**Phylogenetic Placement of the monotypic Genus *Schwackaea* (Melastomeae: Melastomataceae) and the Evolution of its unique Fruit.**
In: International Journal of Plant Sciences 177 (5), p. 440–448. DOI: 10.1086/685689.

**Kriebel, Ricardo; Michelangeli, Fabián Armando; Kelly, Lawrence M. (2015):**
**Discovery of unusual anatomical and continuous Characters in the evolutionary History of *Conostegia* (Miconieae: Melastomataceae).**
In: Molecular Phylogenetics and Evolution 82, p. 289–313. DOI: 10.1016/j.ympev.2014.09.021.

**Majure, Lucas C.; Neubig, Kurt M.; Skean, J. Dan; Bécquer, Eldis R.; Judd, Walter p. (2015):**
**Evolution of the Sandpaper Clade (Miconieae, Melastomataceae).**
In: International Journal of Plant Sciences 176 (7), p. 607–626. DOI: 10.1086/682148.

**Martin, Claire V.; Little, Damon P.; Goldenberg, Renato; Michelangeli, Fabián Armando (2008):**
**A phylogenetic Evaluation of *Leandra* (Miconieae, Melastomataceae): a polyphyletic Genus where the Seeds tell the Story, not the Petals.**
In: Cladistics 24 (3), p. 315–327. DOI: 10.1111/j.1096-0031.2007.00185.x.

**Meyer, Karsten (2001):**
**Revision of the Southeast Asian Genus *Melastoma* (Melastomataceae).**
In: Blumea 46, p. 351–398.

**Michelangeli, Fabián Armando; Carmenate Reyes, Wilder; Sosa, Karla (2015):**
**A Revision of *Meriania* (Melastomataceae) in the Greater Antilles with Emphasis on the Status of the Cuban Species.**
In: Brittonia 67 (2), p. 118–137. DOI: 10.1007/s12228-015-9366-4.

**Michelangeli, Fabián Armando; Guimaraes, Paulo J. F.; Penneys, Darin S.; Almeda, Frank; Kriebel, Ricardo (2013):**
**Phylogenetic Relationships and Distribution of New World Melastomeae (Melastomataceae).**
In: Botanical Journal of the Linnean Society 171, p. 38–60.

**Michelangeli, Fabián Armando; Nicolas, Antoine N.; Morales-P, María Eugenia; David, Heriberto (2011):**

**Phylogenetic Relationships of *Allomaieta, Alloneuron, Cyphostyla,* and *Wurdastom* (Melastomataceae) and the Resurrection of the Tribe Cyphostyleae.**
In: International Journal of Plant Sciences 172 (9), p. 1165–1178. DOI: 10.1086/662032.

**Michelangeli, Fabián Armando; Penneys, Darin S.; Giza, Joanna; Soltis, Douglas E.; Hils, Mathew H.; Skean, J. Dan (2004):**
**A preliminary Phylogeny of the Tribe Miconieae (Melastomataceae) based on nrITS Sequence Data and its Implications on Inflorescence Position.**
In: Taxon 53 (2), p. 279–290. DOI: 10.2307/4135608.

**Morley, Robert J.; Dick, Christopher W. (2003):**
**Missing fossils, molecular Clocks, and the Origin of the Melastomataceae.**
In: American Journal of Botany 90 (11), p. 1638–1644. DOI: 10.3732/ajb.90.11.1638.

**Oliveira da Silva, Marcus Felippe; Fernandes Guimarães, Paulo José; Michelangeli, Fabián Armando (2014):**
**Nomenclatural and taxonomic Novelties in the Tribe Melastomeae (Melastomataceae).**
In: Phytotaxa 186 (4), p. 222. DOI: 10.11646/phytotaxa.186.4.6.

**Penneys, Darin S.; Judd, Walter p. (2005):**
**A Systematic Revision and Cladistic Analysis of *Charianthus* (Melastomataceae) using morphological and molecular Characters.**
In: Systematic Botany 30 (3), p. 559–584.

**Penneys, Darin S.; Judd, Walter p. (2011):**
**Phylogenetics and Morphology in the Blakeeae (Melastomataceae).**
In: International Journal of Plant Sciences 172 (1), p. 78–106. DOI: 10.1086/657284.

**Penneys, Darin S.; Judd, Walter p. (2013):**
**A revised Circumscription for the Blakeeae (Melastomataceae) with associated nomenclatural Adjustments.**
In: PhytoKeys (20), p. 17–32. DOI: 10.3897/phytokeys.20.4344.

**Penneys, Darin S.; Judd, Walter p. (2013):**
**Combined molecular and morphological phylogenetic Analyses of the Blakeeae (Melastomataceae).**
In: International Journal of Plant Sciences 174 (5), p. 802–817. DOI: 10.1086/670011.

**Penneys, Darin S.; Michelangeli, Fabián Armando; Judd, Walter S.; Almeda, Frank (2010):**
**Henrietteeae (Melastomataceae): A new Neotropical Berry-Fruited Tribe.**
In: Systematic Botany 35 (4), p. 783–800. DOI: 10.1600/036364410X539862.

**Pennys, Darin p. (2013):**
**Preliminary Phylogeny of the Astronieae (Melastomataceae) based on Nuclear and Plastid DNA Sequence Data, with Comments on the Philippine endemic Genus, *Astrocalyx*.**
In: Philippine Journal of Science 142, p. 159–168.

**Reginato, Marcelo; Michelangeli, Fabián Armando (2016):**
**Untangling the Phylogeny of *Leandra* s.str. (Melastomataceae, Miconieae).**
In: Molecular Phylogenetics and Evolution 96, p. 17–32. DOI: 10.1016/j.ympev.2015.11.015.

**Reginato, Marcelo; Michelangeli, Fabián Armando; Goldenberg, Renato (2010):**
**Phylogeny of *Pleiochiton* (Melastomataceae, Miconieae): total Evidence.**
In: Botanical Journal of the Linnean Society 162, p. 423–434.

**Renner, Susanne p. (2004):**
**Bayesian Analysis of combined Chloroplast loci, using multiple Calibrations, supports the recent Arrival of Melastomataceae in Africa and Madagascar.**
In: American Journal of Botany 91 (9), p. 1427–1435. DOI: 10.3732/ajb.91.9.1427.

**Rocha, Maria José R.; Batista, João Aguiar Nogueira; Guimaraes, Paulo J. F.; Michelangeli, Fabián Armando (2016):**
**Phylogenetic Relationships in the *Marcetia* alliance (Melastomeae, Melastomataceae) and Implications for generic Circumscription.**
In: Botanical Journal of the Linnean Society 181 (4), p. 585–609. DOI: 10.1111/boj.12429.

**Romero, Rosana; Versiane, Ana Flávia Alves (2014):**
**Taxonomic Novelty and Typifications in *Microlepis* (Melastomataceae).**
In: Novon: A Journal for Botanical Nomenclature 23 (2), p. 217–223. DOI: 10.3417/2011077.

**Veranso-Libalah, Marie Claire; Stone, Robert Douglas; Fongod, Augustina G.N.; Couvreur, Thomas L.P.; Kadereit, Gudrun (2017):**
**Phylogeny and Systematics of African Melastomateae (Melastomataceae).**
In: Taxon 66 (3), p. 584–614. DOI: 10.12705/663.5.

### Meliaceae

**Clarkson, James J.; Pennington, Terence D.; Chase, Mark W.; Haynes, Gwilym; Engstrand, Rachel; Kaye, Maria et al. (2016):**
**Phylogenetic Relationships in *Trichilia* (Meliaceae) based on Ribosomal ITS Sequences.**
In: Phytotaxa 259 (1), p. 6. DOI: 10.11646/phytotaxa.259.1.4.

**Duminil, Jérôme; Kenfack, David; Viscosi, Vincenzo; Grumiau, Laurent; Hardy, Olivier J. (2012):**
**Testing Species Delimitation in sympatric Species complexes: the case of an African tropical Tree, *Carapa* spp. (Meliaceae).**
In: Molecular Phylogenetics and Evolution 62 (1), p. 275–285. DOI: 10.1016/j.ympev.2011.09.020.

**Fukuda, Tatsuya; Yokoyama, Jun U.N.; Tsukaya, Hirokazu (2003):**
**Phylogenetic Relationships among Species in the Genera *Chisocheton* and *Guarea* that have unique indeterminate Leaves as inferred from Sequences of Chloroplast DNA.**
In: International Journal of Plant Sciences 164 (1), p. 13–24.

**Grudinski, Melanie; Pannell, Caroline M.; Chase, Mark W.; Ahmad, Joffre A.; Müllner-Riehl, Alexandra N. (2014):**
**An Evaluation of taxonomic Concepts of the widespread plant Genus *Aglaia* and its allies across Wallace's Line (tribe Aglaieae, Meliaceae).**
In: Molecular Phylogenetics and Evolution 73, p. 65–76. DOI: 10.1016/j.ympev.2014.01.025.

**Koecke, A. Valerie; Müllner-Riehl, Alexandra N.; Pennington, Terence D.; Schorr, Gertrud; Schnitzler, Jan (2013):**
**Niche Evolution through time and across Continents: the story of Neotropical *Cedrela* (Meliaceae).**
In: American Journal of Botany 100 (9), p. 1800–1810. DOI: 10.3732/ajb.1300059.

**Koenen, Erik J. M.; Clarkson, James J.; Pennington, Terence D.; Chatrou, Lars W. (2015):**
**Recently evolved Diversity and convergent Radiations of rainforest Mahoganies (Meliaceae) shed new Light on the Origins of Rainforest Hyperdiversity.**
In: the new Phytologist 207 (2), p. 327–339. DOI: 10.1111/nph.13490.

**Müllner, Alexandra N.; Pennington, Terence D.; Chase, Mark W. (2009):**

**Molecular Phylogenetics of Neotropical Cedreleae (Mahogany Family, Meliaceae) based on nuclear and Plastid DNA Sequences reveal multiple Origins of "*Cedrela odorata*".**

In: Molecular Phylogenetics and Evolution 52 (2), p. 461–469. DOI: 10.1016/j.ympev.2009.03.025.

**Müllner, Alexandra N.; Samuel, Rosabelle; Chase, Mark W.; Coleman, Annette; Stuessy, Tod F. (2008):**

**An Evaluation of Tribes and generic Relationships in Melioideae (Meliaceae) based on nuclear ITS ribosomal DNA.**

In: Taxon 57 (1), p. 98–108.

**Müllner, Alexandra N.; Samuel, Rosabelle; Chase, Mark W.; Pannell, Caroline M.; Greger, Harald (2005):**

***Aglaia* (Meliaceae): An Evaluation of taxonomic Concepts based on DNA Data and secondary Metabolites.**

In: American Journal of Botany 92 (3), p. 534–543.

**Müllner, Alexandra N.; Samuel, Rosabelle; Johnson, Sheila A.; Cheek, Martin R.; Pennington, Terence D.; Chase, Mark W. (2003):**

**Molecular Phylogenetics of Meliaceae (Sapindales) based on Nuclear and Plastid DNA Sequences.**

In: American Journal of Botany 90 (3), p. 471–480.

**Müllner, Alexandra N.; Savolainen, Vincent; Samuel, Rosabelle; Chase, Mark W. (2006):**

**The Mahogany Family "out-of-Africa": Divergence time Estimation, global Biogeographic Patterns inferred from Plastid rbcL DNA Sequences, extant, and fossil Distribution of Diversity.**

In: Molecular Phylogenetics and Evolution 40 (1), p. 236–250. DOI: 10.1016/j.ympev.2006.03.001.

**Pennington, Terence D.; Clarkson, James J. (2013):**

**A Revision of *Guarea* (Meliaceae).**

In: Edinburgh Journal of Botany 70 (02), p. 179–362. DOI: 10.1017/S0960428613000036.

### Melianthaceae

**Linder, Hans Peter; Dlamini, Titus; Henning, Jack; Verboom, George Anthony (2006):**

**The evolutionary History of *Melianthus* (Melianthaceae).**

In: American Journal of Botany 93 (7), p. 1052–1064. DOI: 10.3732/ajb.93.7.1052.

### Memecylaceae

**Stone, Robert Douglas (2006):**

**Phylogeny of Major Lineages in Melastomataceae, Subfamily Olisbeoideae: Utility of Nuclear Glyceraldehyde 3-Phosphate Dehydrogenase (GapC) Gene Sequences.**

In: Systematic Botany 31 (1), p. 107–121.

**Stone, Robert Douglas (2014):**

**The species-rich, paleotropical Genus *Memecylon* (Melastomataceae): Molecular Phylogenetics and revised infrageneric Classification of the African Species.**

In: Taxon 63 (3), p. 539–561. DOI: 10.12705/633.10.

**Stone, Robert Douglas; Andreasen, Katarina (2010):**

**The Afro-Madagascan Genus *Warneckea* (Melastomataceae): Molecular Systematics and revised infrageneric Classification.**

In: Taxon 59 (1), p. 83–92. DOI: 10.1002/tax.591009.

### Menispermaceae

**Ahmed, P. M.; Verma, V.; Qazi, P. H.; Ganaie, M. M.; Bakshi, P. K.; Qazi, G. N. (2005):**
**Molecular Phylogeny in Indian *Tinospora* Species by DNA based molecular Markers.**
In: Plant Systematics and Evolution 256 (1-4), p. 75–87. DOI: 10.1007/s00606-004-0293-1.

**Hoot, Sara B.; Zautke, Herbert; Harris, David J.; Crane, Peter R.; Neves, Susana P. (2009):**
**Phylogenetic Patterns in Menispermaceae based on multiple Chloroplast Sequence Data.**
In: Systematic Botany 34 (1), p. 44–56. DOI: 10.1600/036364409787602339.

**Jacques, Frédéric M.B.; Bertolino, P. (2008):**
**Molecular and morphological Phylogeny of Menispermaceae (Ranunculales).**
In: Plant Systematics and Evolution 274 (1-2), p. 83–97. DOI: 10.1007/s00606-008-0038-7.

**Ortiz, Rosa del C.; Kellogg, Elizabeth A. (2007):**
**Molecular Phylogeny of the Moonseed Family (Menispermaceae): Implications for morphological Diversification.**
In: American Journal of Botany 94 (8), p. 1425–1438.

**Ortiz, Rosa del C.; Wang, Wei; Jacques, Frédéric M.B.; Chen, Zhi-Duan (2016):**
**Phylogeny and a revised tribal Classification of Menispermaceae (Moonseed Family) based on molecular and morphological Data.**
In: Taxon 65 (6), p. 1288–1312. DOI: 10.12705/656.5.

**Wang, Wei; Ortiz, Rosa del C.; Jacques, Frédéric M.B.; Chung, Shih-Wen; Liu, Yang; Xiang, Xiao-Guo; Chen, Zhi-Duan (2017):**
**New Insights into the Phylogeny of Burasaieae (Menispermaceae) with the Recognition of a new Genus and Emphasis on the southern Taiwanese and mainland Chinese Disjunction.**
In: Molecular Phylogenetics and Evolution 109, p. 11–20. DOI: 10.1016/j.ympev.2016.12.038.

**Wang, Wei; Ortiz, Rosa del C.; Jacques, Frédéric M.B.; Xiang, Xiao-Guo; Li, Hong-Lei; Lin, Li et al. (2012):**
**Menispermaceae and the Diversification of tropical rainforests near the Cretaceous-Paleogene Boundary.**
In: the new Phytologist 195 (2), p. 470–478. DOI: 10.1111/j.1469-8137.2012.04158.x.

**Wang, Wei; Wang, Heng-Chang; Chen, Zhi-Duan (2007):**
**Phylogeny and morphological Evolution of Tribe Menispermeae (Menispermaceae) inferred from Chloroplast and nuclear Sequences.**
In: Perspectives in Plant Ecology, Evolution and Systematics 8 (3), p. 141–154. DOI: 10.1016/j.ppees.2006.12.001.

**Wefferling, Keir M.; Hoot, Sara B.; Neves, Susana P. (2013):**
**Phylogeny and Fruit Evolution in Menispermaceae.**
In: American Journal of Botany 100 (5), p. 883–905. DOI: 10.3732/ajb.1200556.

**Xie, Daotao; He, Jiayong; Huang, Jianming; Xie, Hui; Wang, Yaqin; Kang, Yun et al. (2015):**
**Molecular Phylogeny of Chinese *Stephania* (Menispermaceae) and Reassessment of the Subgeneric and Sectional Classifications.**
In: Australian Systematic Botany 28 (4), p. 246–255. DOI: 10.1071/SB14023.

### Menyanthaceae

**Tippery, Nicholas P.; Les, Donald H. (2008):**
**Phylogenetic Analysis of the Internal Transcribed Spacer (ITS) Region in Menyanthaceae using predicted secondary Structure.**

In: Molecular Phylogenetics and Evolution 49 (2), p. 526–537. DOI: 10.1016/j.ympev.2008.07.019.

**Tippery, Nicholas P.; Les, Donald H. (2011):**
**Phylogenetic Relationships and morphological Evolution in *Nymphoides* (Menyanthaceae).**
In: Systematic Botany 36 (4), p. 1101–1113. DOI: 10.1600/036364411X605092.

**Tippery, Nicholas P.; Les, Donald H.; Padgett, Donald J.; Jacobs, Surrey W.L. (2008):**
**Generic Circumscription in Menyanthaceae: A phylogenetic Evaluation.**
In: Systematic Botany 33 (3), p. 598–612. DOI: 10.1600/036364408785679851.

### Metteniusaceae

**González, Favio; Betancur, Julio; Maurin, Olivier; Freudenstein, John V.; Chase, Mark W. (2007):**
**Metteniusaceae, an early-diverging Family in the Lamiid Clade.**
In: Taxon 56 (3), p. 795–800. DOI: 10.2307/25065861.

### Molluginaceae

**Brockington, Samuel F.; Dos Santos, Patricia; Glover, Beverley; Craene, Louis Ronse (2013):**
**Androecial Evolution in Caryophyllales in light of a paraphyletic Molluginaceae.**
In: American Journal of Botany 100 (9), p. 1757–1778. DOI: 10.3732/ajb.1300083.

**Christin, Pascal-Antoine; Sage, Tammy L.; Edwards, Erika J.; Ogburn, R. Matthew; Khoshravesh, Roxanne; Sage, Rowan F. (2011):**
**Complex evolutionary transitions and the Significance of C3-C4 intermediate Forms of Photosynthesis in Molluginaceae.**
In: Evolution 65 (3), p. 643–660. DOI: 10.1111/j.1558-5646.2010.01168.x.

**Thulin, Mats; Moore, Abigail J.; El-Seedi, Hesham; Larsson, Anders; Christin, Pascal-Antoine; Edwards, Erika J. (2016):**
**Phylogeny and generic Delimitation in Molluginaceae, new Pigment Data in Caryophyllales, and the new Family Corbichoniaceae.**
In: Taxon 65 (4), p. 775–793. DOI: 10.12705/654.6.

### Monimiaceae

**Lirio, Elton John; Peixoto, Ariane Luna (2015):**
**A new Species of *Mollinedia* (Monimiaceae, Mollinedioideae, Mollinedieae) from Atlantic Rainforest, Brazil.**
In: Phytotaxa 239 (1), p. 89. DOI: 10.11646/phytotaxa.239.1.9.

**Lirio, Elton John; Peixoto, Ariane Luna; Siqueira, Marinez Ferreira (2015):**
**Taxonomy, conservation, geographic and potential Distribution of *Macrotorus* Perkins (Mollinedioideae, Monimiaceae), and a key to the Neotropical Genera of Monimiaceae.**
In: Phytotaxa 234 (3), p. 201. DOI: 10.11646/phytotaxa.234.3.1.

**Renner, Susanne S.; Takeuchi, Wayne N. (2009):**
**A Phylogeny and revised Circumscription for *Kairoa* (Monimiaceae), with the Description of a new Species from Papua New Guinea.**
In: Harvard Papers in Botany 14 (1), p. 71–81. DOI: 10.3100/025.014.0111.

### Monocotyledoneae

**Givnish, Thomas J.; Pires, J. Chris; Graham, Sean W.; McPherson, Marc; Prince, Linda M.; Patterson, Thomas B. et al. (2006):**

**Phylogenetic Relationships of Monocots based on the Highly Informative Plastid Gene ndhF.**
In: Aliso 22 (1), p. 28–51. DOI: 10.5642/aliso.20062201.04.

### Montiaceae

APPLEQUIST, WENDY L.; WAGNER, WARREN L.; ZIMMER, ELIZABETH A.; NEPOKROEFF, MOLLY (2006):
**Molecular Evidence Resolving the Systematic Position of *Hectorella* (Portulacaceae).**
In: Systematic Botany 31 (2), p. 310–319. DOI: 10.1600/036364406777585900.

MATTHEW OGBURN, R.; EDWARDS, ERIKA J. (2015):
**Life History Lability underlies rapid Climate Niche Evolution in the Angiosperm Clade Montiaceae.**
In: Molecular Phylogenetics and Evolution 92, p. 181–192. DOI: 10.1016/j.ympev.2015.06.006.

O'QUINN, ROBIN; HUFFORD, LARRY (2005):
**Molecular Systematics of Montieae (Portulacaceae): Implications for Taxonomy, Biogeography and Ecology.**
In: Systematic Botany 30 (2), p. 314–331.

STOUGHTON, THOMAS R.; JOLLES, DIANA D.; O'QUINN, ROBIN L. (2017):
**The Western Spring Beauties, *Claytonia lanceolata* (Montiaceae): A Review and Revised Taxonomy for California.**
In: Systematic Botany 42 (2), p. 283–300. DOI: 10.1600/036364417X695475.

### Moraceae

BRUUN-LUND, SAM; CLEMENT, WENDY L.; KJELLBERG, FINN; RØNSTED, NINA (2017):
**First Plastid phylogenomic Study reveals potential cyto-nuclear Discordance in the evolutionary History of *Ficus* L. (Moraceae).**
In: Molecular Phylogenetics and Evolution 109, p. 93–104. DOI: 10.1016/j.ympev.2016.12.031.

CRUAUD, ASTRID; RØNSTED, NINA; CHANTARASUWAN, BHANUMAS; CHOU, LIEN SIANG; CLEMENT, WENDY L.; COULOUX, ARNAUD ET AL. (2012):
**An extreme Case of Plant-Insect Codiversification: Figs and Fig-pollinating Wasps.**
In: Systematic Biology 61 (6), p. 1029–1047. DOI: 10.1093/sysbio/sys068.

DATWYLER, SHANNON L.; WEIBLEN, GEORGE D. (2004):
**On the Origin of the Fig: phylogenetic Relationships of Moraceae from Ndhf Sequences.**
In: American Journal of Botany 91 (5), p. 767–777.

GARDNER, ELLIOT M.; SARRAF, PAYA; WILLIAMS, EVELYN W.; ZEREGA, NYREE J.C. (2017):
**Phylogeny and Biogeography of *Maclura* (Moraceae) and the Origin of an anachronistic fruit.**
In: Molecular Phylogenetics and Evolution 117, p. 49–59. DOI: 10.1016/j.ympev.2017.06.021.

LI, HONG-QING; WANG, SHUANG; CHEN, JI-YUN; GUI, PING (2012):
**Molecular Phylogeny of *Ficus* Section *Ficus* in China based on four DNA Regions.**
In: Journal of Systematics and Evolution 50 (5), p. 422–432. DOI: 10.1111/j.1759-6831.2012.00221.x.

MACHADO, ANDERSON FERREIRA PINTO; RØNSTED, NINA; BRUUN-LUND, SAM; PEREIRA, RODRIGO AUGUSTO SANTINELO; PAGANUCCI DE QUEIROZ, LUCIANO (2018):
**Atlantic forests to the all Americas: Biogeographical History and Divergence times of Neotropical *Ficus* (Moraceae).**
In: Molecular Phylogenetics and Evolution 122, p. 46–58. DOI: 10.1016/j.ympev.2018.01.015.

**Misiewicz, T. M.; Zerega, Nyree J.C. (2012):**
**Phylogeny, Biogeography and Character Evolution of Dorstenia (Moraceae).**
In: Edinburgh Journal of Botany 69 (3), p. 413–440. DOI: 10.1017/S096042861200025X.

**Renoult, Julien P.; Kjellberg, Finn; Grout, Cinderella; Santoni, Sylvain; Khadari, Bouchaïb (2009):**
**Cyto-nuclear Discordance in the Phylogeny of *Ficus* Section *Galoglychia* and Host Shifts in Plant-pollinator associations.**
In: BMC Evolutionary Biology 9, p. 248. DOI: 10.1186/1471-2148-9-248.

**Rønsted, Nina; Salvo, Gabriele; Savolainen, Vincent (2007):**
**Biogeographical and phylogenetic Origins of African Fig Species (*Ficus* Section *Galoglychia*).**
In: Molecular Phylogenetics and Evolution 43 (1), p. 190–201. DOI: 10.1016/j.ympev.2006.12.010.

**Rønsted, Nina; Weiblen, George D.; Savolainen, Vincent; Cook, James M. (2008):**
**Phylogeny, Biogeography, and Ecology of *Ficus* Section *Malvanthera* (Moraceae).**
In: Molecular Phylogenetics and Evolution 48 (1), p. 12–22. DOI: 10.1016/j.ympev.2008.04.005.

**Vijayan, Kunjupillai (2004):**
**Genetic Relationships of Japanese and Indian mulberry (*Morus* spp.) Genotypes revealed by DNA Fingerprinting.**
In: Plant Systematics and Evolution 243 (3-4), p. 221–232. DOI: 10.1007/s00606-003-0078-y.

**Xu, Lei; Harrison, Rhett D.; Yang, Pei; Yang, Da-Rong (2011):**
**New Insight into the phylogenetic and biogeographic History of Genus *Ficus*: Vicariance played a relatively minor Role compared with ecological Opportunity and Dispersal.**
In: Journal of Systematics and Evolution 49 (6), p. 546–557. DOI: 10.1111/j.1759-6831.2011.00155.x.

**Yang, Li-Yuan; Machado, Carlos A.; Dang, Xiao-Dong; Peng, Yan-Qiong; Yang, Da-Rong; Zhang, Da-Yong; Liao, Wan-Jin (2015):**
**The incidence and Pattern of Copollinator Diversification in dioecious and monoecious Figs.**
In: Evolution 69 (2), p. 294–304. DOI: 10.1111/evo.12584.

**Zerega, Nyree J.C.; Clement, Wendy L.; Datwyler, Shannon L.; Weiblen, George D. (2005):**
**Biogeography and Divergence Times in the Mulberry Family (Moraceae).**
In: Molecular Phylogenetics and Evolution 37 (2), p. 402–416. DOI: 10.1016/j.ympev.2005.07.004.

**Zerega, Nyree J.C.; Nur Supardi, M. N.; Motley, Timothy J. (2010):**
**Phylogeny and Recircumscription of Artocarpeae (Moraceae) with a Focus on *Artocarpus*.**
In: Systematic Botany 35 (4), p. 766–782. DOI: 10.1600/036364410X539853.

**Zerega, Nyree J.C.; Ragone, Diane; Motley, Timothy J. (2005):**
**Systematics and Species limits of Breadfruit (*Artocarpus*, Moraceae).**
In: Systematic Botany 30 (3), p. 603–615.

### Morinaceae

**Bell, C. (2003):**
**Phylogeny and Biogeography of Morinaceae (Dipsacales) based on nuclear and Chloroplast DNA Sequences.**
In: Organisms Diversity and Evolution 3 (3), p. 227–237. DOI: 10.1078/1439-6092-00077.

### Musaceae

**Christelová, Pavla; Valárik, Miroslav; Hřibová, Eva; Langhe, Edmond de; Doležel, Jaroslav (2011):**
**A multi Gene Sequence-based Phylogeny of the Musaceae (Banana) Family.**
In: BMC Evolutionary Biology 11, p. 103. DOI: 10.1186/1471-2148-11-103.

**Hareesh, V. S.; Joe, A.; Sreejith, P. E.; Sabu, M. (2017):**
***Musa markkuana* stat. nov. (Musaceae) - A reassessment of *Musa velutina* subsp. *markkuana*.**
In: Phytotaxa 303 (3), p. 279. DOI: 10.11646/phytotaxa.303.3.8.

**Langhe, Edmond de; Pillay, M.; Tenkouano, A.; Swennen, R. (2005):**
**Integrating morphological and molecular Taxonomy in *Musa*: the African plantains (*Musa* spp. Aab Group).**
In: Plant Systematics and Evolution 255 (3-4), p. 225–236. DOI: 10.1007/s00606-005-0346-0.

**Li, Lin-Feng; Häkkinen, Markku; Yuan, Yong-Ming; Hao, Gang; Ge, Xue-Jun (2010):**
**Molecular Phylogeny and Systematics of the Banana Family (Musaceae) inferred from multiple nuclear and Chloroplast DNA Fragments, with a special Reference to the Genus *Musa*.**
In: Molecular Phylogenetics and Evolution 57 (1), p. 1–10. DOI: 10.1016/j.ympev.2010.06.021.

**Liu, Ai-Zhong; Kress, W. John; Li, De-Zhu (2010):**
**Phylogenetic Analyses of the Banana Family (Musaceae) based on nuclear ribosomal (ITS) and Chloroplast (trnL-F) Evidence.**
In: Taxon 59 (1), p. 20–28.

### Myricaceae

**Herbert, Jane; Chase, Mark W.; Möller, Michael; Abbott, Richard J. (2006):**
**Nuclear and Plastid DNA Sequences confirm the Placement of the enigmatic *Canacomyrica monticola* in Myricaceae.**
In: Taxon 55 (2), p. 349–357. DOI: 10.2307/25065582.

**Huguet, Valérie; Gouy, Manolo; Normand, Philippe; Zimpfer, Jeff F.; Fernandez, Maria P. (2005):**
**Molecular Phylogeny of Myricaceae: a Reexamination of host-symbiont Specificity.**
In: Molecular Phylogenetics and Evolution 34 (3), p. 557–568. DOI: 10.1016/j.ympev.2004.11.018.

**Liu, Luxian; Jin, Xin-Jie; Chen, Nan; Li, Xian; Li, Pan; Fu, Cheng-Xin (2015):**
**Phylogeny of *Morella rubra* and its Relatives (Myricaceae) and Genetic Resources of Chinese Bayberry using RAD Sequencing.**
In: Public Library of Science One 10 (10), e0139840. DOI: 10.1371/journal.pone.0139840.

### Myristiacaceae

**Sauquet, Hervé; Doyle, James A.; Scharaschkin, Tanya; Borsch, Thomas; Hilu, Khidir W.; Chatrou, Lars W.; Le Thomas, Annick (2003):**
**Phylogenetic Analysis of Magnoliales and Myristicaceae based on multiple Data Sets: Implications for Character Evolution.**
In: Botanical Journal of the Linnean Society 142 (2), p. 125–186. DOI: 10.1046/j.1095-8339.2003.00171.x.

**Sauquet, Hervé; Le Thomas, Annick (2003):**
**Pollen Diversity and Evolution in Myristicaceae (Magnoliales).**
In: International Journal of Plant Sciences 164 (4), p. 613–628.

**Sheeja, T. E.; Sabeesh, C.; Shabna, O. V.; Shalini, R. S.; Krishnamoorthy, B. (2013):**
**Genetic Diversity Analysis of *Myristica* and related Genera using Rapd and ISSR Markers.**

In: Journal of Spices and Aromatic Crops 22 (1), p. 38–46.

**Steeves, Royce Allan David (2011):**
**An Intrageneric Study of morphological and Genetic Variation in the Neotropical Compsoneura and *Virola* (Myristiaceae).**
University of Guelph, Ontario, Canada. Guelph, Ontario, Canada.

### Myrsinaceae

**Anderberg, Arne A. (1994):**
**Phylogeny and Subgeneric Classification of *Cyclamen* L. (Primulaceae).**
In: Kew Bulletin 49 (3), p. 455–467.

**Anderberg, Arne A.; Manns, Ulrika; Källersjö, Mari (2007):**
**Phylogeny and Floral Evolution of the Lysimachieae (Ericales, Myrsinaceae): Evidence from ndhF Sequence Data.**
In: Willdenowia 37 (2), p. 407–421. DOI: 10.3372/wi.37.37202.

**Anderberg, Arne A.; Trift, Ida; Kllersj, Mari (2000):**
**Phylogeny of *Cyclamen* L. (Primulaceae): Evidence from Morphology and Sequence Data from the internal transcribed Spacers of nuclear ribosomal DNA.**
In: Plant Systematics and Evolution 220 (3-4), p. 147–160. DOI: 10.1007/BF00985043.

**Bone, Ruth E.; Strijk, Joeri S.; Fritsch, Peter W.; Buerki, Sven; Strasberg, Dominique; Thébaud, Christophe; Hodkinson, Trevor R. (2012):**
**Phylogenetic Inference of *Badula* (Primulaceae), a rare and threatened Genus endemic to the Mascarene Archipelago.**
In: Botanical Journal of the Linnean Society 169 (2), p. 284–296. DOI: 10.1111/j.1095-8339.2012.01221.x.

**Dubéarnès, Anne; Julius, Avelinah; Utteridge, Timothy M.A. (2015):**
**A Synopsis of the Genus *Embelia* in Peninsular Malaysia and Singapore. Studies in Malaysian Myrsinaceae III.**
In: Kew Bulletin 70 (2), p. 1–33. DOI: 10.1007/S12225-015-9570-0.

**Gielly, L.; Debussche, M.; Thompson, J. D. (2001):**
**Geographic Isolation and Evolution of Mediterranean endemic *Cyclamen* : Insights from Chloroplast trnL (Uaa) Intron Sequence Variation.**
In: Plant Systematics and Evolution 230 (1-2), p. 75–88. DOI: 10.1007/s006060170005.

**Hao, Gang; Yuan, Yong-Ming; Hu, Chi-Ming; Ge, Xue-Jun; Zhao, Nan-Xian (2004):**
**Molecular Phylogeny of *Lysimachia* (Myrsinaceae) based on Chloroplast trnL–F and nuclear ribosomal ITS Sequences.**
In: Molecular Phylogenetics and Evolution 31 (1), p. 323–339. DOI: 10.1016/S1055-7903(03)00286-0.

**Källersjö, Mari; Bergqvist, Gullevi; Anderberg, Arne A. (2000):**
**Generic Realignment in primuloid Families of the Ericales s.l.: a phylogenetic Analysis based on DNA Sequences from three Chloroplast Genes and Morphology.**
In: American Journal of Botany 87 (9), p. 1325–1341. DOI: 10.2307/2656725.

**Lemaire, Benny; Smets, Erik; Dessein, Steven (2011):**
**Bacterial Leaf symbiosis in *Ardisia* (Myrsinoideae, Primulaceae): molecular Evidence for Host Specificity.**
In: Research in Microbiology 162 (5), p. 528–534. DOI: 10.1016/j.resmic.2011.04.003.

**Manns, Ulrika; Anderberg, Arne A. (2005):**

**Molecular Phylogeny of *Anagallis* (Myrsinaceae) based on ITS, trnL-F, and ndhF Sequence Data.**
In: International Journal of Plant Sciences 166 (6), p. 1019–1028.

**MANNS, ULRIKA; ANDERBERG, ARNE A. (2007):**
**Character Evolution in *Anagallis* (Myrsinaceae) inferred from morphological and molecular Data.**
In: Systematic Botany 32 (1), p. 166–179.

**MARTINS, L.; OBERPRIELER, CHRISTOPH; HELLWIG, FRANK H. (2003):**
**A phylogenetic Analysis of Primulaceae s.l. based on Internal Transcribed Spacer (ITS) DNA Sequence Data.**
In: Plant Systematics and Evolution 237 (1-2), p. 75–85. DOI: 10.1007/s00606-002-0258-1.

**OH, IL-CHAN; SCHÖNENBERGER, JÜRGEN; MOTLEY, TIMOTHY J.; MYRENÅS, MATTIAS; ANDERBERG, ARNE A. (2013):**
**Phylogenetic Relationships among endemic Hawaiian *Lysimachia* (Ericales: Primulaceae): Insights from Nuclear and Chloroplast DNA Sequence Data.**
In: Pacific Science 67 (2), p. 237–251. DOI: 10.2984/67.2.7.

**PIPOLY, JOHN J.; RICKETSON, JON M. (1998):**
**A Revision of the Genus *Ardisia* Subgenus *Graphardisia* (Myrsinaceae).**
In: Sida, Contributions to Botany 18 (2), p. 433–472.

**RICKETSON, JON M.; PIPOLY, JOHN J. (2010):**
**A Synopsis of Neotropical *Stylogyne* (Myrsinaceae).**
In: Novon: A Journal for Botanical Nomenclature 20 (4), p. 437–447. DOI: 10.3417/2010027.

**SCHMID, MAURICE (2009):**
**Contribution à la Connaissance des Primulaceae (ex Myrsinaceae) de Nouvelle-Calédonie. II. Le genre *Rapanea* Aubl.**
In: Adansonia 31 (2), p. 341–395. DOI: 10.5252/a2009n2a8.

**SCHMID, MAURICE (2012):**
**Contribution à la connaissance des Primulaceae (ex Myrsinaceae) de Nouvelle-Calédonie. III. Les genres *Tapeinosperma* Hook.f. et Mangenotiella gen. nov.**
In: Adansonia 34 (2), p. 279–341. DOI: 10.5252/a2012n2a7.

**SLEUMER, H. (1988):**
**A Revision of the Genus *Ardisia* Sw. (Myrsinaceae) in New Guinea.**
In: Blumea 33, p. 115–140.

**SUNARNO, BAMBANG (2002):**
**New Species of *Labisia* (Myrsinaceae) from Sumatra.**
In: Reinwardtia 12 (1), p. 121–124.

**SUNARNO, BAMBANG (2005):**
**Revision of the Genus *Labisia* (Myrsinaceae).**
In: Blumea 50 (3), p. 579–597. DOI: 10.3767/000651905X622879.

**WANNTORP, LIVIA; RONSE DE CRAENE, LOUIS P.; PENG, CHING-I.; ANDERBERG, ARNE A. (2012):**
**Floral Ontogeny and Morphology of *Stimpsonia* and *Ardisiandra*, two aberrant Genera of the Primuloid Clade of Ericales.**
In: International Journal of Plant Sciences 173 (9), p. 1023–1035. DOI: 10.1086/667607.

**YESSON, CHRIS; CULHAM, ALASTAIR (2006):**
**A phyloclimatic Study of *Cyclamen*.**

In: BMC Evolutionary Biology 6, p. 72. DOI: 10.1186/1471-2148-6-72.

ZHANG, CAI-YUN; WANG, FENG-YING; YAN, HAI-FEI; HAO, GANG; HU, CHI-MING; GE, XUE-JUN (2012):
**Testing DNA barcoding in closely related Groups of *Lysimachia* L. (Myrsinaceae).**
In: molecular Ecology Resources 12 (1), p. 98–108. DOI: 10.1111/j.1755-0998.2011.03076.x.

### Myrtaceae

BAYLY, MICHAEL J.; LADIGES, PAULINE Y. (2007):
**Divergent paralogues of ribosomal DNA in Eucalypts (Myrtaceae).**
In: Molecular Phylogenetics and Evolution 44 (1), p. 346–356. DOI: 10.1016/j.ympev.2006.10.027.

BAYLY, MICHAEL J.; RIGAULT, PHILIPPE; SPOKEVICIUS, ANTANAS; LADIGES, PAULINE Y.; ADES, PETER K.; ANDERSON, CHARLOTTE ET AL. (2013):
**Chloroplast Genome Analysis of Australian Eucalypts - *Eucalyptus, Corymbia, Angophora, Allosyncarpia* and *Stockwellia* (Myrtaceae).**
In: Molecular Phylogenetics and Evolution 69 (3), p. 704–716. DOI: 10.1016/j.ympev.2013.07.006.

BAYLY, MICHAEL J.; UDOVICIC, FRANK; GIBBS, ADELE K.; PARRA-O., CARLOS; LADIGES, PAULINE Y. (2008):
**Ribosomal DNA pseudogenes are widespread in the Eucalypt Group (Myrtaceae): Implications for phylogenetic Analysis.**
In: Cladistics 24 (2), p. 131–146. DOI: 10.1111/j.1096-0031.2007.00175.x.

BIFFIN, ED; CRAVEN, LYN A.; CRISP, MICHAEL D.; GADEK, PAUL A. (2006):
**Molecular Systematics of *Syzygium* and allied Genera (Myrtaceae): Evidence from the Chloroplast Genome.**
In: Taxon 55 (1), p. 79–94. DOI: 10.2307/25065530.

BIFFIN, ED; HARRINGTON, MARK G.; CRISP, MICHAEL D.; CRAVEN, L. A.; GADEK, PAUL A. (2007):
**Structural Partitioning, paired-sites Models and Evolution of the ITS Transcript in *Syzygium* and Myrtaceae.**
In: Molecular Phylogenetics and Evolution 43 (1), p. 124–139. DOI: 10.1016/j.ympev.2006.08.013.

BOHTE, A.; DRINNAN, ANDREW N. (2005):
**Floral development and systematic Position of *Arillastrum, Allosyncarpia, Stockwellia* and *Eucalyptopsis* (Myrtaceae).**
In: Plant Systematics and Evolution 251 (1), p. 53–70. DOI: 10.1007/s00606-004-0161-z.

BROOKER, M. I.H. (2000):
**A new Classification of the Genus *Eucalyptus* L'Hér. (Myrtaceae).**
In: Australian Systematic Botany 13 (1), p. 79–148. DOI: 10.1071/SB98008.

BROOKER, M. I.H.; HOOPER, S.D. (2002):
**Taxonomy of Species deriving from the Publication of *Eucalyptus* subseries *Cornutae* (Myrtaceae).**
In: Nuytsia 14 (3), p. 325–360.

BROWN, GILLIAN K.; UDOVICIC, FRANK; LADIGES, PAULINE Y. (2001):
**Molecular Phylogeny and Biogeography of *Melaleuca, Callistemon* and related Genera (Myrtaceae).**
In: Australian Systematic Botany 14 (4), p. 565–585. DOI: 10.1071/SB00029.

COPELAND, LACHLAN M.; CRAVEN, LYN A.; BRUHL, JEREMY J. (2011):
**A taxonomic Review of *Homoranthus* (Myrtaceae: Chamelaucieae).**
In: Australian Systematic Botany 24 (6), p. 351–374. DOI: 10.1071/SB11015.

CRAVEN, L. A.; BIFFIN, ED (2010):

**An infrageneric Classification of *Syzygium* (Myrtaceae).**
In: Blumea 55 (1), p. 94–99. DOI: 10.3767/000651910X499303.

**Craven, Lyn A.; Edwards, Robert D.; Cowley, Kirsten J. (2014):**
**New Combinations and names in *Melaleuca* (Myrtaceae).**
In: Taxon 63 (3), p. 663–670. DOI: 10.12705/633.38.

**Edwards, Robert D.; Craven, Lyn A.; Crisp, Michael D.; Cook, Lyn G. (2010):**
***Melaleuca* revisited: cpDNA and morphological Data confirm that *Melaleuca* L. (Myrtaceae) is not monophyletic.**
In: Taxon 59 (3), p. 744–754. DOI: 10.1002/tax.593007.

**Flores-Rentería, Lluvia; Rymer, Paul D.; Riegler, Markus (2017):**
**Unpacking boxes: Integration of molecular, morphological and ecological Approaches reveals extensive Patterns of reticulate Evolution in box Eucalypts.**
In: Molecular Phylogenetics and Evolution 108, p. 70–87. DOI: 10.1016/j.ympev.2017.01.019.

**Gibbs, Adele K.; Udovicic, Frank; Drinnan, Andrew N.; Ladiges, Pauline Y. (2009):**
**Phylogeny and Classification of *Eucalyptus* Subgenus *Eudesmia* (Myrtaceae) based on nuclear ribosomal DNA, Chloroplast DNA and Morphology.**
In: Australian Systematic Botany 22 (3), p. 158–179. DOI: 10.1071/SB08043.

**Harrington, Mark G.; Gadek, Paul A. (2004):**
**Molecular Systematics of the *Acmena* alliance (Myrtaceae): phylogenetic Analyses and evolutionary Implications with Reference to Australian taxa.**
In: Australian Systematic Botany 17 (1), p. 63–72. DOI: 10.1071/SB03009.

**Harrington, Mark G.; Jackes, Betsy R.; Barrett, Matthew D.; Craven, Lyn A.; Barrett, Russell L. (2012):**
**Phylogenetic Revision of Backhousieae (Myrtaceae): Neogene Divergence, a revised Circumscription of *Backhousia* and two new Species.**
In: Australian Systematic Botany 25 (6), p. 404–417. DOI: 10.1071/SB12015.

**Jones, Rebecca C.; Nicolle, Dean; Steane, Dorothy A.; Vaillancourt, René E.; Potts, Bradley M. (2016):**
**High density, Genome-wide Markers and intra-specific replication yield an unprecedented phylogenetic Reconstruction of a globally significant, speciose Lineage of *Eucalyptus*.**
In: Molecular Phylogenetics and Evolution 105, p. 63–85. DOI: 10.1016/j.ympev.2016.08.009.

**Lam, N.; Wilson, Paul G.; Heslewood, Margaret M.; Quinn, Christopher J. (2002):**
**A phylogenetic Analysis of the *Chamelaucium* alliance (Myrtaceae).**
In: Australian Systematic Botany 15 (4), p. 535–543. DOI: 10.1071/SB01039.

**Lange, Peter J.; Smissen, Rob D.O.B.; Wagstaff, Steven J.; Keeling, D. Jeannette; Murray, B. G.; Toelken, H. R. (2010):**
**A molecular Phylogeny and infrageneric Classification for *Kunzea* (Myrtaceae) inferred from rDNA ITS and ETS Sequences.**
In: Australian Systematic Botany 23 (5), p. 309–319. DOI: 10.1071/SB10019.

**Lucas, Eve J.; Harris, Stephen A.; Mazine, Fiorella Fernanda; Belsham, Stephen R.; Nic Lughadha, Eimear M.; Telford, Annika et al. (2007):**
**Suprageneric Phylogenetics of Myrteae, the generically richest Tribe in Myrtaceae (Myrtales).**
In: Taxon 56 (4), p. 1105–1128. DOI: 10.2307/25065906.

**Lucas, Eve J.; Matsumoto, Kazue; Harris, Stephen A.; Nic Lughadha, Eimear M.; Benardini, Benedetta; Chase, Mark W. (2011):**
**Phylogenetics, Morphology, and Evolution of the large Genus *Myrcia* s.l. (Myrtaceae).**
In: International Journal of Plant Sciences 172 (7), p. 915–934. DOI: 10.1086/660913.

**Mazine, Fiorella Fernanda; Souza, Vinicius Castro; Sobral, Marcos; Forest, Félix; Lucas, Eve J. (2014):**
**A preliminary phylogenetic Analysis of *Eugenia* (Myrtaceae: Myrteae), with a focus on Neotropical Species.**
In: Kew Bulletin 69 (2), p. 9497. DOI: 10.1007/S12225-014-9497-X.

**McKinnon, Gay E.; Vaillancourt, René E.; Steane, Dorothy A.; Potts, Bradley M. (2008):**
**An AFLP marker approach to lower-level Systematics in *Eucalyptus* (Myrtaceae).**
In: American Journal of Botany 95 (3), p. 368–380. DOI: 10.3732/ajb.95.3.368.

**Mills, Penelope J.; Cook, Lyn G. (2014):**
**Rapid chromosomal Evolution in a morphologically cryptic Radiation.**
In: Molecular Phylogenetics and Evolution 77, p. 126–135. DOI: 10.1016/j.ympev.2014.03.015.

**Murillo-A, José; Ruiz-Ponce, Eduardo; Landrum, Leslie R.; Stuessy, Tod F.; Barfuss, Michael H. J. (2012):**
**Phylogenetic Relationships in *Myrceugenia* (Myrtaceae) based on Plastid and nuclear DNA Sequences.**
In: Molecular Phylogenetics and Evolution 62 (2), p. 764–776. DOI: 10.1016/j.ympev.2011.11.021.

**Murillo-A, José; Stuessy, Tod F.; Ruiz, Eduardo (2013):**
**Phylogenetic Relationships among *Myrceugenia, Blepharocalyx,* and *Luma* (Myrtaceae) based on paired-sites Models and the secondary Structures of ITS and ETS Sequences.**
In: Plant Systematics and Evolution 299 (4), p. 713–729. DOI: 10.1007/s00606-012-0754-x.

**Nicolle, Dean; Whalen, Molly A.; Mackay, Duncan A. (2006):**
**Morphological Variation and phylogenetic Relationships within *Eucalyptus* series *Subulatae* (Myrtaceae) of southern Australia.**
In: Australian Systematic Botany 19 (1), p. 59–86. DOI: 10.1071/SB04036.

**Ochieng, Joel W.; Henry, Robert J.; Baverstock, Peter R.; Steane, Dorothy A.; Shepherd, Mervyn (2007):**
**Nuclear ribosomal pseudogenes resolve a corroborated Monophyly of the Eucalypt Genus *Corymbia* despite misleading Hypotheses at functional ITS Paralogs.**
In: Molecular Phylogenetics and Evolution 44 (2), p. 752–764. DOI: 10.1016/j.ympev.2007.04.017.

**Parra-O., Carlos; Bayly, Michael J.; Drinnan, Andrew N.; Udovicic, Frank; Ladiges, Pauline Y. (2010):**
**Corrigendum to: Phylogeny, major Clades and infrageneric Classification of *Corymbia* (Myrtaceae), based on nuclear ribosomal DNA and Morphology.**
In: Australian Systematic Botany 23 (2). DOI: 10.1071/SB09028_CO.

**Parra-O., Carlos; Bayly, Michael J.; Drinnan, Andrew N.; Udovicic, Frank; Ladiges, Pauline Y. (2009):**
**Phylogeny, major Clades and infrageneric Classification of *Corymbia* (Myrtaceae), based on nuclear ribosomal DNA and Morphology.**
In: Australian Systematic Botany 22 (5), p. 384–399. DOI: 10.1071/SB09028.

**Parra-O., Carlos; Bayly, Michael J.; Udovicic, Frank; Ladiges, Pauline Y. (2006):**
**Ets Sequences support the Monophyly of the Eucalypt Genus *Corymbia* (Myrtaceae).**
In: Taxon 55 (3), p. 653–663. DOI: 10.2307/25065641.

**Poke, Fiona S.; Martin, Darren P.; Steane, Dorothy A.; Vaillancourt, René E.; Reid, James B. (2006):**

**The impact of intragenic recombination on phylogenetic Reconstruction at the Sectional level in *Eucalyptus* when using a single copy nuclear Gene (Cinnamoyl CoA Reductase).**
In: Molecular Phylogenetics and Evolution 39 (1), p. 160–170. DOI: 10.1016/j.ympev.2005.11.016.

PROENÇA, CAROLYN E.B.; LUGHADHA, EIMEAR M. NIC; LUCAS, EVE J.; WOODGYER, ELIZABETH M. (2006):
**_Algrizea_ (Myrteae, Myrtaceae): A new Genus from the Highlands of Brazil.**
In: Systematic Botany 31 (2), p. 320–326.

RYE, BARBARA L. (2013):
**A Revision of the south-western Australian Genus *Astartea* (Myrtaceae: Chamelaucieae).**
In: Nuytsia 23, p. 189–269.

RYE, BARBARA L. (2015):
**A Revision of the south-western Australian Genus *Babingtonia* (Myrtaceae: Chamelaucieae).**
In: Nuytsia 25, p. 219–250.

RYE, BARBARA L. (2015):
**Reinstatement of *Ericomyrtus* (Myrtaceae: Chamaelaucieae) with three new Combinations.**
In: Nuytsia 25, p. 131–143.

RYE, BARBARA L.; TRUDGEN, MALCOLM E. (2012):
**Seven new Combinations for Western Australian members of Myrtaceae Tribe Chamelaucieae.**
In: Nuytsia 22 (6), p. 393–398.

SANTOS, MATHEUS FORTES; LUCAS, EVE J.; SANO, PAULO TAKEO; BUERKI, SVEN; STAGGEMEIER, VANESSA GRAZIELE; FOREST, FÉLIX (2017):
**Biogeographical Patterns of *Myrcia* s.l. (Myrtaceae) and their correlation with geological and climatic History in the Neotropics.**
In: Molecular Phylogenetics and Evolution 108, p. 34–48. DOI: 10.1016/j.ympev.2017.01.012.

SANTOS, MATHEUS FORTES; SANO, PAULO TAKEO; FOREST, FÉLIX; LUCAS, EVE J. (2016):
**Phylogeny, Morphology and Circumscription of *Myrcia* Sect. *Sympodiomyrcia* (*Myrcia* s.l., Myrtaceae).**
In: Taxon 65 (4), p. 759–774. DOI: 10.12705/654.5.

SNOW, NEIL (2009):
**_Kanakomyrtus_ (Myrtaceae): A new Endemic Genus from New Caledonia with Linear Stigma Lobes and Baccate Fruits.**
In: Systematic Botany 34 (2), p. 330–344. DOI: 10.1600/036364409788606253.

SNOW, NEIL; DAWSON, JOHN W.; CALLMANDER, MARTIN W.; GANDHI, KANCHI; MUNZINGER, JÉRÔME (2016):
**New Species, new Combinations, and Lectotypifications in New Caledonian *Eugenia* L. (Myrtaceae).**
In: Candollea 71 (1), p. 67–81. DOI: 10.15553/c2016v711a9.

SNOW, NEIL; MCFADDEN, JESSIE; EVANS, TIMOTHY M.; SALYWON SALYWON, ANDREW M.; WOJCIECHOWSKI, MARTIN F.; WILSON, PETER G. (2011):
**Morphological and molecular Evidence of Polyphyly in *Rhodomyrtus* (Myrtaceae: Myrteae).**

In: Systematic Botany 36 (2), p. 390–404. DOI: 10.1600/036364411X569570.

**Sobral, Marcos; Costa, Idimá G.; Souza, Marcelo C.; Zorzanelli, Joao Paulo F. (2017):**
**Five new Species and one new Combination in Brazilian Myrtaceae.**
In: Phytotaxa 307 (4), p. 233. DOI: 10.11646/phytotaxa.307.4.1.

**Soh, Wuu Kuang; Parnell, John A.N. (2011):**
**Comparative Leaf Anatomy and Phylogeny of *Syzygium* Gaertn.**
In: Plant Systematics and Evolution 297 (1-2), p. 1–32. DOI: 10.1007/s00606-011-0495-2.

**Staggemeier, Vanessa Graziele; Lucas, Eve J. (2014):**
**Morphological diagnosis of a new Species in *Myrcia* sensu lato (Myrtaceae) from Bahia, Brazil, with molecular highlights.**
In: Phytotaxa 181 (4), p. 229. DOI: 10.11646/phytotaxa.181.4.3.

**Steane, Dorothy A.; McKinnon, Gay E.; Vaillancourt, René E.; Potts, Bradley M. (1999):**
**ITS Sequence Data resolve higher level Relationships among the Eucalypts.**
In: Molecular Phylogenetics and Evolution 12 (2), p. 215–223. DOI: 10.1006/mpev.1999.0612.

**Steane, Dorothy A.; Nicolle, Dean; McKinnon, Gay E.; Vaillancourt, René E.; Potts, Bradley M. (2002):**
**Higher-level Relationships among the Eucalypts are resolved by ITS-sequence Data.**
In: Australian Systematic Botany 15 (1), p. 49–62. DOI: 10.1071/SB00039.

**Steane, Dorothy A.; Nicolle, Dean; Potts, Bradley M. (2007):**
**Phylogenetic Positioning of anomalous Eucalypts by using ITS Sequence Data.**
In: Australian Systematic Botany 20 (5), p. 402–408. DOI: 10.1071/SB07013.

**Steane, Dorothy A.; Nicolle, Dean; Sansaloni, Carolina P.; Petroli, César D.; Carling, Jason; Kilian, Andrzej et al. (2011):**
**Population genetic Analysis and Phylogeny Reconstruction in *Eucalyptus* (Myrtaceae) using high-throughput, Genome-wide Genotyping.**
In: Molecular Phylogenetics and Evolution 59 (1), p. 206–224. DOI: 10.1016/j.ympev.2011.02.003.

**Thornhill, Andrew H.; Ho, Simon Y.W.; Külheim, Carsten; Crisp, Michael D. (2015):**
**Interpreting the modern Distribution of Myrtaceae using a dated molecular Phylogeny.**
In: Molecular Phylogenetics and Evolution 93, p. 29–43. DOI: 10.1016/j.ympev.2015.07.007.

**Trudgen, Malcolm E.; Rye, Barbara L. (2005):**
***Astus*, a new Western Australian Genus of Myrtaceae with heterocarpidic Fruits.**
In: Nuytsia 15 (3), p. 495–512.

**Trudgen, Malcolm E.; Rye, Barbara L. (2010):**
***Enekbatus*, a new Western Australian Genus of Myrtaceae with a multi-locular indehiscent Fruit.**
In: Nuytsia 20, p. 229–259.

**Udovicic, Frank; Ladiges, Pauline Y. (2000):**
**Informativeness of Nuclear and Chloroplast DNA Regions and the Phylogeny of the Eucalypts and related Genera (Myrtaceae).**
In: Kew Bulletin 55 (3), p. 633–645.

**Udovicic, Frank; Spencer, Roger D. (2012):**
**New Combinations in *Callistemon* (Myrtaceae).**
In: Muelleria 30 (1), p. 23–25.

**van der Merwe, Marlien M.; van Wyk, Abraham E.; Botha, A. M. (2005):**

**Molecular phylogenetic Analysis of *Eugenia* L. (Myrtaceae), with Emphasis on southern African Taxa.**
In: Plant Systematics and Evolution 251 (1), p. 21–34. DOI: 10.1007/s00606-004-0160-0.

VASCONCELOS, THAÍS N. C.; PROENÇA, CAROLYN E.B.; AHMAD, BERHAMAN; AGUILAR, DANIEL S.; AGUILAR, REINALDO; AMORIM, BRUNO P. ET AL. (2017):
**Myrteae Phylogeny, Calibration, Biogeography and Diversification Patterns: Increased Understanding in the most Species rich Tribe of Myrtaceae.**
In: Molecular Phylogenetics and Evolution 109, p. 113–137. DOI: 10.1016/j.ympev.2017.01.002.

WILSON, CHRISTINE E.; FOREST, FÉLIX; DEVEY, DION S.; LUCAS, EVE J. (2016):
**Phylogenetic Relationships in *Calyptranthes* (Myrtaceae) with Particular Emphasis on its Monophyly relative to *Myrcia* s. l.**
In: Systematic Botany 41 (2), p. 378–386. DOI: 10.1600/036364416X691786.

WILSON, PAUL G.; O'BRIEN, M. M.; HESLEWOOD, MARGARET M.; QUINN, CHRISTOPHER J. (2005):
**Relationships within Myrtaceae sensu lato based on a matK Phylogeny.**
In: Plant Systematics and Evolution 251 (1), p. 3–19. DOI: 10.1007/s00606-004-0162-y.

WILSON, PETER G.; HESLEWOOD, MARGARET M. (2014):
**An expanded phylogenetic Analysis of *Sannantha* (Myrtaceae) and Description of a new Species.**
In: Australian Systematic Botany 27 (1), p. 78–84. DOI: 10.1071/SB14011.

WILSON, PETER G.; HESLEWOOD, MARGARET M. (2016):
**Phylogenetic Position of *Meteoromyrtus* (Myrtaceae).**
In: Telopea 19, p. 45–55. DOI: 10.7751/telopea10389.

WILSON, PETER G.; HESLEWOOD, MARGARET M.; QUINN, CHRISTOPHER J. (2007):
**Re-evaluation of the Genus *Babingtonia* (Myrtaceae) in eastern Australia and New Caledonia.**
In: Australian Systematic Botany 20 (4), p. 302–318. DOI: 10.1071/SB06032.

WRIGHT, SHANE D.; KEELING, D. JEANNETTE; ASHTON, FERN G.; DAWSON, JOHN W.; GARDNER, RICHARD C. (2000):
**Phylogenetic Analyses of New Caledonian *Metrosideros* and *Carpolepis* (Myrtaceae) from nrDNA (ITS) Sequences.**
In: Australian Systematic Botany 13 (6), p. 919–926. DOI: 10.1071/SB00009.

### Myrtales

BERGER, BRENT A.; KRIEBEL, RICARDO; SPALINK, DANIEL; SYTSMA, KENNETH J. (2016):
**Divergence times, historical Biogeography, and shifts in Speciation Rates of Myrtales.**
In: Molecular Phylogenetics and Evolution 95, p. 116–136. DOI: 10.1016/j.ympev.2015.10.001.

VASCONCELOS, THAÍS N. C.; PROENÇA, CAROLYN E.B. (2015):
**Floral cost vs. Floral display: Insights from the megadiverse Myrtales suggest that energetically expensive Floral Parts are less phylogenetically constrained.**
In: American Journal of Botany 102 (6), p. 900–909. DOI: 10.3732/ajb.1400509.

### Nartheciaceae

FUSE, SHIZUKA; LEE, NAM SOOK; TAMURA, MINORU N. (2012):
**Biosystematic studies on the Family Nartheciaceae (Dioscoreales) I. phylogenetic Relationships, Character Evolution and taxonomic Re-examination.**
In: Plant Systematics and Evolution 298 (8), p. 1575–1584. DOI: 10.1007/s00606-012-0660-2.

ZHAO, YI-MIN; WANG, WEI; ZHANG, SHU-REN (2012):

**Delimitation and Phylogeny of *Aletris* (Nartheciaceae) with Implications for Perianth Evolution.**

In: Journal of Systematics and Evolution 50 (2), p. 135–145. DOI: 10.1111/j.1759-6831.2011.00172.x.

### Nelumbonaceae

**XUE, JIAN-HUA; DONG, WEN-PAN; CHENG, TAO; ZHOU, SHI-LIANG (2012):**

**Nelumbonaceae: Systematic Position and Species Diversification revealed by the complete Chloroplast Genome.**

In: Journal of Systematics and Evolution 50 (6), p. 477–487. DOI: 10.1111/j.1759-6831.2012.00224.x.

### Nepenthaceae

**ALAMSYAH, FIRMAN; ITO, MOTOMI (2013):**

**Phylogenetic Analysis of Nepenthaceae, based on Internal Transcribed Spacer Nuclear Ribosomal DNA Sequences.**

In: Acta Phytotaxonomica Geobotanica 64 (3), p. 113–126.

**MEIMBERG, HARALD; HEUBL, GÜNTHER (2006):**

**Introduction of a nuclear Marker for phylogenetic Analysis of Nepenthaceae.**

In: Plant Biology 8 (6), p. 831–840. DOI: 10.1055/s-2006-924676.

**MEIMBERG, HARALD; THALHAMMER, STEFAN; BRACHMANN, ANDREAS; HEUBL, GÜNTHER (2006):**

**Comparative Analysis of a translocated Copy of the trnK Intron in carnivorous Family Nepenthaceae.**

In: Molecular Phylogenetics and Evolution 39 (2), p. 478–490. DOI: 10.1016/j.ympev.2005.11.023.

**MEIMBERG, HARALD; WISTUBA, A.; DITTRICH, P.; HEUBL, GÜNTHER (2001):**

**Molecular Phylogeny of Nepenthaceae based on Cladistic Analysis of Plastid trnK Intron Sequence Data.**

In: Plant Biology 3 (2), p. 164–175. DOI: 10.1055/s-2001-12897.

### Nitrariaceae

**TEMIRBAYEVA, KAMSHAT; ZHANG, MING-LI (2015):**

**Molecular phylogenetic and biogeographical Analysis of *Nitraria* based on nuclear and Chloroplast DNA Sequences.**

In: Plant Systematics and Evolution 301 (7), p. 1897–1906. DOI: 10.1007/s00606-015-1202-5.

### Nothofagaceae

**ACOSTA, M. CRISTINA; PREMOLI, ANDREA C. (2010):**

**Evidence of Chloroplast capture in South American *Nothofagus* (Subgenus *Nothofagus*, Nothofagaceae).**

In: Molecular Phylogenetics and Evolution 54 (1), p. 235–242. DOI: 10.1016/j.ympev.2009.08.008.

**HEENAN, PETER B.; SMISSEN, ROB D.O.B. (2013):**

**Revised Circumscription of *Nothofagus* and Recognition of the segregate Genera *Fuscospora, Lophozonia,* and *Trisyngyne* (Nothofagaceae).**

In: Phytotaxa 146 (1), p. 1. DOI: 10.11646/phytotaxa.146.1.1.

**SAUQUET, HERVÉ; HO, SIMON Y.W.; GANDOLFO, MARIA A.; JORDAN, GREGORY J.; WILF, PETER; CANTRILL, DAVID J. ET AL. (2012):**

**Testing the impact of calibration on molecular Divergence times using a fossil-rich Group: the case of *Nothofagus* (Fagales).**

In: Systematic Biology 61 (2), p. 289–313. DOI: 10.1093/sysbio/syr116.

Smissen, Rob D.O.B.; Richardson, p. J.; Morse, C. W.; Heenan, Peter B. (2014):

**Relationships, Gene flow and Species boundaries among New Zealand *Fuscospora* (Nothofagaceae: Southern Beech).**

In: New Zealand Journal of Botany 52 (4), p. 389–406. DOI: 10.1080/0028825X.2014.960946.

### Nyctaginaceae

Douglas, Norman A.; Manos, Paul p. (2007):

**Molecular Phylogeny of Nyctaginaceae: Taxonomy, Biogeography, and Characters associated with a Radiation of xerophytic Genera in North America.**

In: American Journal of Botany 94 (5), p. 856–872. DOI: 10.3732/ajb.94.5.856.

Douglas, Norman A.; Spellenberg, Richard (2010):

**A new tribal Classification of Nyctaginaceae.**

In: Taxon 59 (3), p. 905–910.

Hernández-Ledesma, Patricia; Olvera, Hilda Flores; Ochoterena, Helga (2010):

**Cladistic Analysis and taxonomic Synopsis of *Anulocaulis* (Nyctaginaceae) based on morphological Data.**

In: Systematic Botany 35 (4), p. 858–876. DOI: 10.1600/036364410X539916.

Levin, Rachel A. (2000):

**Phylogenetic Relationships within Nyctaginaceae Tribe Nyctagineae: Evidence from Nuclear and Chloroplast Genomes.**

In: Systematic Botany 25 (4), p. 738. DOI: 10.2307/2666731.

Levin, Rachel A.; McDade, Lucinda A.; Raguso, Robert A. (2003):

**The Systematic Utility of Floral and vegetative Fragrance in two Genera of Nyctaginaceae.**

In: Systematic Biology 52 (3), p. 334–351. DOI: 10.1080/10635150390196975.

Struwig, Madeleen; Klaassen, Esmerialda S.; Kwembeya, Ezekiel G. (2015):

**Nyctaginaceae: A taxonomic Treatment for the Flora of Namibia.**

In: Phytotaxa 238 (2), p. 101. DOI: 10.11646/phytotaxa.238.2.1.

### Nymphaeaceae

Borsch, Thomas; Hilu, Khidir W.; Wiersema, John H.; Löhne, Cornelia; Barthlott, Wilhelm; Wilde, Volker (2007):

**Phylogeny of *Nymphaea* (Nymphaeaceae): Evidence from Substitutions and Microstructural Changes in the Chloroplast trnT-trnF Region.**

In: International Journal of Plant Sciences 168 (5), p. 639–671.

Borsch, Thomas; Löhne, Cornelia; Mbaye, Mame Samba; Wiersema, John H. (2011):

**Towards a complete Species Tree of *Nymphaea*: shedding further Light on Subg. *Brachyceras* and its Relationships to the Australian water-lilies.**

In: Telopea 13 (1-2), p. 193–217.

Borsch, Thomas; Wiersema, John H.; Hellquist, C. Barre; Löhne, Cornelia; Govers, Kim (2014):

**Speciation in North American water lilies: Evidence for the hybrid Origin of the newly discovered Canadian endemic *Nymphaea loriana* sp. nov. (Nymphaeaceae) in a past Contact Zone.**

In: Botany 92 (12), p. 867–882. DOI: 10.1139/cjb-2014-0060.

Dkhar, Jeremy; Kumaria, Suman; Rao, Satyawada Rama; Tandon, Pramod (2012):

**Sequence characteristics and phylogenetic Implications of the nrDNA internal transcribed Spacers (ITS) in the Genus *Nymphaea* with focus on some Indian Representatives.**
In: Plant Systematics and Evolution 298 (1), p. 93–108. DOI: 10.1007/s00606-011-0526-z.

LÖHNE, CORNELIA; BORSCH, THOMAS; JACOBS, SURREY W.L.; HELLQUIST, C. BARRE; WIERSEMA, JOHN H. (2008):
**Nuclear and Plastid DNA Sequences reveal complex reticulate Patterns in Australian water-Lilies (*Nymphaea* Subgenus *Anecphya*, Nymphaeaceae).**
In: Australian Systematic Botany 21 (4), p. 229–250. DOI: 10.1071/SB07010.

PADGETT, DONALD J.; LES, DONALD H.; CROW, GARRETT E. (1999):
**Phylogenetic Relationships in *Nuphar* (Nymphaeaceae): Evidence from Morphology, Chloroplast DNA, and Nuclear Ribosomal DNA.**
In: American Journal of Botany 86 (9), p. 1316–1324.

TAYLOR, DAVID WINSHIP (2008):
**Phylogenetic Analysis of Cabombaceae and Nymphaeaceae based on vegetative and Leaf architectural Characters.**
In: Taxon 57 (4), p. 1082–1095. DOI: 10.1002/tax.574005.

### Nymphaeales

BORSCH, THOMAS; LÖHNE, CORNELIA; WIERSEMA, JOHN H. (2008):
**Phylogeny and evolutionary Patterns in Nymphaeales: integrating genes, Genomes and Morphology.**
In: Taxon 57 (4), p. 1052–1081. DOI: 10.1002/tax.574004.

LÖHNE, CORNELIA; BORSCH, THOMAS; WIERSEMA, JOHN H. (2007):
**Phylogenetic Analysis of Nymphaeales using fast-evolving and noncoding Chloroplast Markers.**
In: Botanical Journal of the Linnean Society 154 (2), p. 141–163. DOI: 10.1111/j.1095-8339.2007.00659.x.

YOO, MI-JEONG; BELL, CHARLES D.; SOLTIS, PAMELA S.; SOLTIS, DOUGLAS E. (2005):
**Divergence Times and Historical Biogeography of Nymphaeales.**
In: Systematic Botany 30 (4), p. 693–704.

### Nyssaceae


Wang, Nian; Milne, Richard Ian; Jacques, Frédéric M.B.; Sun, Bao-Ling; Zhang, Chang-Qin; Yang, Jun-Bo (2012):

**Phylogeny and a revised Classification of the Chinese Species of *Nyssa* (Nyssaceae) based on morphological and molecular Data.**

In: Taxon 61 (2), p. 344–354.

Xiang, Qiu-Yun; Moody, Michael L.; Soltis, Douglas E.; Fan, Chuanzhu; Soltis, Pamela p. (2002):

**Relationships within Cornales and Circumscription of Cornaceae - matK and rbcL Sequence Data and effects of outgroups and long Branches.**

In: Molecular Phylogenetics and Evolution 24 (1), p. 35–57. DOI: 10.1016/S1055-7903(02)00267-1.

Zhou, Wenbin; Ji, Xiang; Obata, Shihori; Pais, Andrew; Dong, Yibo; Peet, Robert; Xiang, Qiu-Yun Jenny (2018):

**Resolving Relationships and phylogeographic History of the *Nyssa sylvatica* complex using Data from RAD-seq and Species Distribution Modeling.**

In: Molecular Phylogenetics and Evolution 126, p. 1–16. DOI: 10.1016/j.ympev.2018.04.001.


### Ochnaceae


Sastre, Claude; Offroy, Bérangère (2016):

**Révision Nomenclaturale des Binômes du Genre Néotropical *Ouratea* Aublet (Ochnaceae) décrits par Van Tieghem.**

In: Adansonia 38 (1), p. 55–98. DOI: 10.5252/a2016n1a5.

Schneider, Julio Valentin; Bissiengou, Pulcherie; Amaral, Maria do Carmo E.; Tahir, Ali; Fay, Michael F.; Thines, Marco et al. (2014):

**Phylogenetics, ancestral state reconstruction, and a new infrafamilial Classification of the pantropical Ochnaceae (Medusagynaceae, Ochnaceae s.str., Quiinaceae) based on five DNA Regions.**

In: Molecular Phylogenetics and Evolution 78, p. 199–214. DOI: 10.1016/j.ympev.2014.05.018.


### Octoknemaceae


Gosline, George; Malécot, Valéry (2011):

**A monograph of *Octoknema* (Octoknemaceae - Olacaceae s.l.).**

In: Kew Bulletin 66, p. 367–404.


### Olacaceae


Malécot, Valéry; Nickrent, Daniel Lee (2008):

**Molecular phylogenetic Relationships of Olacaceae and related Santalales.**

In: Systematic Botany 33 (1), p. 97–106. DOI: 10.1600/036364408783887384.


### Oleaceae


Baldoni, L.; Guerrero, C.; Sossey-Aloui, K.; Abott, A. G.; Angiolillo, A.; Lumaret R. (2002):

**Phylogenetic Relationships among *Olea* Species, based on Nucleotide Variation at a Non-Coding Chloroplast DNA Region.**

In: Plant Biology 4, p. 346–351.

Besnard, Guillaume; Rubio de Casas, Rafael; Christin, Pascal-Antoine; Vargas, Pablo (2009):

**Phylogenetics of *Olea* (Oleaceae) based on Plastid and nuclear ribosomal DNA Sequences: Tertiary climatic Shifts and Lineage differentiation Times.**

In: Annals of Botany 104 (1), p. 143–160. DOI: 10.1093/aob/mcp105.

Gu, Jing; Su, Jun-Xia; Lin, Ruo-Zhu; Li, Rui-Qi; Xiao, Pei-Gen (2011):

**Testing four proposed barcoding Markers for the Identification of Species within *Ligustrum* L. (Oleaceae).**
In: Journal of Systematics and Evolution 49 (3), p. 213–224. DOI: 10.1111/j.1759-6831.2011.00136.x.

**Guo, Shi-Quan; Xiong, Min; Ji, Chun-Feng; Zhang, Zhi-Rong; Li, De-Zhu; Zhang, Zhi-Yong (2011):**
**Molecular phylogenetic Reconstruction of *Osmanthus* Lour. (Oleaceae) and related Genera based on three Chloroplast intergenic spacers.**
In: Plant Systematics and Evolution 294 (1-2), p. 57–64. DOI: 10.1007/s00606-011-0445-z.

**Hinsinger, Damien D.; Gaudeul, Myriam; Couloux, Arnaud; Bousquet, Jean; Frascaria-Lacoste, Nathalie (2014):**
**The Phylogeography of Eurasian *Fraxinus* Species reveals ancient transcontinental Reticulation.**
In: Molecular Phylogenetics and Evolution 77, p. 223–237. DOI: 10.1016/j.ympev.2014.04.021.

**Hong-Wa, Cynthia; Besnard, Guillaume (2013):**
**Intricate Patterns of phylogenetic Relationships in the Olive Family as inferred from multi-locus Plastid and nuclear DNA Sequence Analyses: a close-up on *Chionanthus* and *Noronhia* (Oleaceae).**
In: Molecular Phylogenetics and Evolution 67 (2), p. 367–378. DOI: 10.1016/j.ympev.2013.02.003.

**Jeandroz, Sylvain; Roy, Alice; Bousquet, Jean (1997):**
**Phylogeny and Phylogeography of the Circumpolar Genus *Fraxinus* (Oleaceae) based on Internal Transcribed Spacer Sequences of Nuclear Ribosomal DNA.**
In: Molecular Phylogenetics and Evolution 7 (2), p. 241–251.

**Kim, Dong-Kap; Kim, Joo-Hwan (2011):**
**Molecular Phylogeny of Tribe Forsythieae (Oleaceae) based on nuclear ribosomal DNA internal transcribed Spacers and Plastid DNA trnL-F and matK Gene Sequences.**
In: Journal of Plant Research 124 (3), p. 339–347. DOI: 10.1007/s10265-010-0383-9.

**Kim, Ki-joong (1999):**
**Molecular Phylogeny of *Forsythia* (Oleaceae) based on Chloroplast DNA Variation.**
In: Plant Systematics and Evolution 218, p. 113–123.

**Kim, Ki-joong; Jansen, Robert K. (1998):**
**A Chloroplast DNA Phylogeny of Lilacs (*Syringa*, Oleaceae): Plastome Groups show a strong Correlation with crossing Groups.**
In: American Journal of Botany 85 (9), p. 1338–1351.

**Lowry, Porter Prescott; Hong-Wa, Cynthia (Hg.) (2016):**
**A taxonomic Revision of the Genus *Noronhia* Stadtm. ex Thouars (Oleaceae) in Madagascar and the Comoro Islands.**
Genève: Conservatoire et jardin botaniques de la Ville de Genève (Boissiera, vol. 70).

**Nesom, Guy L. (2014):**
**Phylogeny of *Fraxinus* Sect. *Melioides* (Oleaceae): Review and an alternative Hypothesis.**
In: Phytoneuron 95, p. 1–9.

**Wallander, Eva (2008):**
**Systematics of *Fraxinus* (Oleaceae) and Evolution of Dioecy.**
In: Plant Systematics and Evolution 273 (1-2), p. 25–49. DOI: 10.1007/s00606-008-0005-3.

**WALLANDER, EVA; ALBERT, VICTOR A. (2000):**
**Phylogeny and Classification of Oleaceae based on Rps16 and Trnl-f Sequence Data.**
In: American Journal of Botany 87 (12), p. 1827–1841.

**YUAN, WANG-JUN; ZHANG, WEI-RUI; HAN, YUAN-JI; DONG, MEI-FANG; SHANG, FU-DE (2010):**
**Molecular Phylogeny of *Osmanthus* (Oleaceae) based on non-coding Chloroplast and nuclear ribosomal Internal Transcribed Spacer Regions.**
In: Journal of Systematics and Evolution 48 (6), p. 482–489. DOI: 10.1111/j.1759-6831.2010.00099.x.


### Oliniaceae

**SEBOLA, R. J.; BALKWILL, KEVIN (2013):**
**A monographic Study of the Oliniaceae.**
In: Kew Bulletin 68 (3), p. 419–456. DOI: 10.1007/S12225-013-9465-X.


### Onagraceae

**BEDOYA, ANA M.; MADRIÑÁN, SANTIAGO (2015):**
**Evolution of the aquatic habit in *Ludwigia* (Onagraceae): Morpho-anatomical adaptive Strategies in the Neotropics.**
In: Aquatic Botany 120, p. 352–362. DOI: 10.1016/j.aquabot.2014.10.005.

**BERRY, PAUL E.; HAHN, WILLIAM J.; SYTSMA, KENNETH J.; HALL, JOCELYN C.; MAST, AUSTIN R. (2004):**
**Phylogenetic Relationships and Biogeography of *Fuchsia* (Onagraceae) based on noncoding nuclear and Chloroplast DNA Data.**
In: American Journal of Botany 91 (4), p. 601–614. DOI: 10.3732/ajb.91.4.601.

**FORD, V. S.; GOTTLIEB, L. D. (2003):**
**Reassessment of phylogenetic Relationships in *Clarkia* Sect. *Sympherica*.**
In: American Journal of Botany 90 (2), p. 284–292. DOI: 10.3732/ajb.90.2.284.

**FORD, V. S.; GOTTLIEB, L. D. (2007):**
**Tribal Relationships within Onagraceae inferred from PgiC Sequences.**
In: Systematic Botany 32 (2), p. 348–356. DOI: 10.1600/036364407781179725.

**GRÍMSSON, FRIÐGEIR; ZETTER, REINHARD; LENG, QIN (2012):**
**Diverse fossil Onagraceae Pollen from a Miocene Palynoflora of North-east China: early Steps in Resolving the Phytogeographic History of the Family.**
In: Plant Systematics and Evolution 298 (3), p. 671–687. DOI: 10.1007/s00606-011-0578-0.

**HOGGARD, GLORIA D.; KORES, PAUL J.; MOLVRAY, MIA; HOGGARD, RONALD K. (2004):**
**The Phylogeny of *Gaura* (Onagraceae) based on ITS, ETS, and trnL-F Sequence Data.**
In: American Journal of Botany 91 (1), p. 139–148. DOI: 10.3732/ajb.91.1.139.

**HUNG, KUO-HSIANG; SCHAAL, BARBARA A.; HSU, TSAI-WEN; CHIANG, YU-CHUNG; PENG, CHING-I.; CHIANG, TZEN-YUH (2009):**
**Phylogenetic Relationships of diploid and polyploid Species in *Ludwigia* Sect. *Isnardia* (Onagraceae) based on Chloroplast and nuclear DNAs.**
In: Taxon 58 (4), p. 1216–1226. DOI: 10.1002/tax.584013.

**KRAKOS, KYRA N.; REECE, JOSHUA S.; RAVEN, PETER H. (2014):**
**Molecular Phylogenetics and Reproductive Biology of *Oenothera* Section *Kneiffia* (Onagraceae).**
In: Systematic Botany 39 (2), p. 523–532. DOI: 10.1600/036364414X680744.

**PENG, CHING-I.; SCHMIDT, CLIFFORD L.; HOCH, PETER C.; RAVEN, PETER H. (2005):**
**Systematics and Evolution of *Ludwigia* Section *Dantia* (Onagraceae).**

In: Annals of the Missouri Botanical Garden 92 (3), p. 307–359.

**Xie, Lei; Wagner, Warren L.; Ree, Richard H.; Berry, Paul E.; Wen, Jun (2009):**
**Molecular Phylogeny, Divergence time estimates, and historical Biogeography of *Circaea* (Onagraceae) in the Northern Hemisphere.**
In: Molecular Phylogenetics and Evolution 53 (3), p. 995–1009. DOI: 10.1016/j.ympev.2009.09.009.

### Ophioglossaceae

**Dauphin, Benjamin; Vieu, Julien; Grant, Jason R. (2014):**
**Molecular Phylogenetics supports widespread cryptic Species in Moonworts (*Botrychium* s.s., Ophioglossaceae).**
In: American Journal of Botany 101 (1), p. 128–140. DOI: 10.3732/ajb.1300154.

**Hauk, Warren D.; Parks, Clifford R.; Chase, Mark W. (2003):**
**Phylogenetic studies of Ophioglossaceae: Evidence from rbcL and trnL-F Plastid DNA Sequences and Morphology.**
In: Molecular Phylogenetics and Evolution 28 (1), p. 131–151. DOI: 10.1016/S1055-7903(03)00032-0.

**Small, Randall L.; Lickey, Edgar B.; Shaw, Joey; Hauk, Warren D. (2005):**
**Amplification of noncoding Chloroplast DNA for phylogenetic studies in Lycophytes and Monilophytes with a comparative example of relative phylogenetic Utility from Ophioglossaceae.**
In: Molecular Phylogenetics and Evolution 36 (3), p. 509–522. DOI: 10.1016/j.ympev.2005.04.018.

**Sun, Byung-Yun; Kim, Moon Hong; Kim, Chul Hwan; Park, Chong-Wook (2001):**
***Mankyua* (Ophioglossaceae): a new Fern Genus from Cheju Island, Korea.**
In: Taxon 50 (4), p. 1019–1024.

**Williams, Evelyn W.; Waller, Donald M. (2012):**
**Phylogenetic Placement of Species within the Genus *Botrychium* s.s. (Ophioglossaceae) on the Basis of Plastid Sequences, amplified Fragment Length Polymorphisms, and flow Cytometry.**
In: International Journal of Plant Sciences 173 (5), p. 516–531. DOI: 10.1086/664711.

### Opiliaceae

**Le, Chi-Toan; Liu, Bing; Barrett, Russell L.; Lu, Li-Min; Wen, Jun; Chen, Zhi-Duan (2018):**
**Phylogeny and a new tribal Classification of Opiliaceae (Santalales) based on molecular and morphological Evidence.**
In: Journal of Systematics and Evolution 56 (1), p. 56–66. DOI: 10.1111/jse.12295.

**Yang, Liu; Yang, Guan-Song; Ma, Hai-Ying; Wang, Yue-Hua; Shen, Shi-Kang (2018):**
**Phylogenetic Placement of *Yunnanopilia* (Opiliaceae) inferred from molecular and morphological Data.**
In: Journal of Systematics and Evolution 56 (1), p. 48–55. DOI: 10.1111/jse.12285.

### Orchidaceae

**Aceto, S.; Caputo, P.; Cozzolino, Salvatore; Gaudio, L.; Moretti, A. (1999):**
**Phylogeny and Evolution of *Orchis* and allied Genera based on ITS DNA Variation: morphological Gaps and molecular Continuity.**
In: Molecular Phylogenetics and Evolution 13 (1), p. 67–76. DOI: 10.1006/mpev.1999.0628.

**Adams, Peter B. (2011):**

**Systematics of Dendrobiinae (Orchidaceae), with special Reference to Australian Taxa.**
In: Botanical Journal of the Linnean Society 166 (2), p. 105–126. DOI: 10.1111/j.1095-8339.2011.01141.x.

**Alvarez-Molina, Aída; Cameron, Kenneth M. (2009):**
**Molecular Phylogenetics of Prescottiinae s.l. and their close allies (Orchidaceae, Cranichideae) inferred from Plastid and nuclear ribosomal DNA Sequences.**
In: American Journal of Botany 96 (5), p. 1020–1040. DOI: 10.3732/ajb.0800219.

**Andriananjamanantsoa, Herinandrianina N.; Engberg, Shannon; Louis, Edward E.; Brouillet, Luc (2016):**
**Diversification of *Angraecum* (Orchidaceae, Vandeae) in Madagascar: Revised Phylogeny reveals Species Accumulation through time rather than rapid Radiation.**
In: Public Library of Science One 11 (9), e0163194. DOI: 10.1371/journal.pone.0163194.

**Arévalo, Rafael; Cameron, Kenneth M. (2013):**
**Molecular Phylogenetics of *Mormolyca* (Orchidaceae: Maxillariinae) based on combined molecular Data Sets.**
In: Lankesteriana 13 (1-2), p. 1–11. DOI: 10.15517/lank.v0i0.11528.

**Averyanov, Leonid V. (2009):**
***Hayata glandulifera* (Orchidaceae), new Genus and Species from Northern Vietnam.**
In: Taiwania 54 (4), p. 311–316.

**Averyanov, Leonid V.; Kuznetsov, Andrey N.; Kuznetsova, Svetlana P. (2013):**
***Vietorchis furcata* (Orchidaceae, VietOrchidinae) - a new Species from Southern Vietnam.**
In: Taiwania 58 (4), p. 251–256.

**Azevedo, Cecília Oliveira de; van den Berg, Cássio; Barros, Fábio (2014):**
**A Revision of *Prescottia* (Orchidaceae: Orchidoideae, Cranichideae).**
In: Phytotaxa 178 (4), p. 233. DOI: 10.11646/phytotaxa.178.4.1.

**Baranow, Przemysław; Szlachetko, Dariusz L. (2016):**
**The taxonomic Revision of *Sobralia* Ruiz & Pav. (Orchidaceae) in the Guyanas (Guyana, Suriname, French Guiana).**
In: Plant Systematics and Evolution 302 (3), p. 333–355. DOI: 10.1007/s00606-015-1266-2.

**Baranow, Przemysław; Szlachetko, Dariusz L.; Dudek, Magdalena (2014):**
**New Species of *Sobralia* Section *Abbreviatae* Brieger (Orchidaceae) from Colombia. A morphological and molecular Evidence.**
In: Plant Systematics and Evolution 300 (7), p. 1663–1670. DOI: 10.1007/s00606-014-0991-2.

**Barrett, Craig F.; Freudenstein, John V. (2008):**
**Molecular Evolution of rbcL in the mycoheterotrophic Coralroot Orchids (*Corallorhiza* Gagnebin, Orchidaceae).**
In: Molecular Phylogenetics and Evolution 47 (2), p. 665–679. DOI: 10.1016/j.ympev.2008.02.014.

**Barrett, Craig F.; Freudenstein, John V.; Taylor, D. Lee; Kõljalg, Urmas (2010):**
**Rangewide Analysis of fungal associations in the fully mycoheterotrophic *Corallorhiza striata* complex (Orchidaceae) reveals extreme Specificity on ectomycorrhizal *Tomentella* (Thelephoraceae) across North America.**
In: American Journal of Botany 97 (4), p. 628–643. DOI: 10.3732/ajb.0900230.

**Bateman, Richard M.; Hollingsworth, Peter M.; Preston, Jillian; Yi-Bo, L. U.O.; Pridgeon, Alec M.; Chase, Mark W. (2003):**

**Molecular Phylogenetics and Evolution of Orchidinae and selected Habenariinae (Orchidaceae).**
In: Botanical Journal of the Linnean Society 142 (1), p. 1–40. DOI: 10.1046/j.1095-8339.2003.00157.x.

**Bateman, Richard M.; Rudall, Paula J.; James, Karen E. (2006):**
**Phylogenetic Context, generic Affinities and evolutionary Origin of the enigmatic Balkan Orchid *Gymnadenia frivaldii* Hampe ex Griseb.**
In: Taxon 55 (1), p. 107–118. DOI: 10.2307/25065532.

**Batista, João Aguiar Nogueira; Borges, Karina S.; Faria, Marina W. F.; Proite, Karina; Ramalho, Aline Joseph; Salazar, Gerardo A.; van den Berg, Cássio (2013):**
**Molecular Phylogenetics of the species-rich Genus *Habenaria* (Orchidaceae) in the New World based on nuclear and Plastid DNA Sequences.**
In: Molecular Phylogenetics and Evolution 67 (1), p. 95–109. DOI: 10.1016/j.ympev.2013.01.008.

**Batista, João Aguiar Nogueira; Meneguzzo, Thiago E.C.; Salazar, Gerardo A.; Ramalho, Aline Joseph; Bem Bianchetti, Luciano (2011):**
**Phylogenetic placement, taxonomic Revision and a new Species of *Nothostele* (Orchidaceae), an enigmatic Genus endemic to the Cerrado of Central Brazil.**
In: Botanical Journal of the Linnean Society 165 (4), p. 348–363. DOI: 10.1111/j.1095-8339.2011.01113.x.

**Batista, João Aguiar Nogueira; Mota, A. Carolina M.N.A.; Proite, Karina; Bianchetti, Luciano De Bem; Romero-González, Gustavo A.; Huerta, Héctor; Salazar, Gerardo A. (2014):**
**Molecular Phylogenetics of Neotropical *Cyanaeorchis* (Cymbidieae, Epidendroideae, Orchidaceae): geographical rather than morphological Similarities plus a new Species.**
In: Phytotaxa 156 (5), p. 251. DOI: 10.11646/phytotaxa.156.5.1.

**Bellstedt, Dirk U.; Linder, Hans Peter; Harley, Eric H. (2001):**
**Phylogenetic Relationships in *Disa* based on non-coding trnL-trnF Chloroplast Sequences: Evidence of numerous repeat Regions.**
In: American Journal of Botany 88 (11), p. 2088–2100. DOI: 10.2307/3558434.

**Bellusci, F.; Pellegrino, G.; Palermo, A. M.; Musacchio, A. (2008):**
**Phylogenetic Relationships in the Orchid Genus *Serapias* L. based on noncoding Regions of the Chloroplast Genome.**
In: Molecular Phylogenetics and Evolution 47 (3), p. 986–991. DOI: 10.1016/j.ympev.2008.03.019.

**Bernardos, Sonia; Santos, María A.; Tyteca, Daniel; Amich, Francisco (2006):**
**Phylogenetic Relationships of Mediterranean Neottieae and Orchideae (Orchidaceae) inferred from nuclear ribosomal ITS Sequences.**
In: Acta Botanica Gallica 153 (2), p. 153–165. DOI: 10.1080/12538078.2006.10515534.

**Blanco, Mario A.; Carnevali, Germán; Witten, W. Mark; Singer, Rodrigo B.; Koehler, Samantha; Williams, Norris H. et al. (2011):**
**Generic Realignments in Maxillariinae (Orchidaceae).**
In: Lankesteriana 7 (3), p. 515–537. DOI: 10.15517/lank.v0i0.7935.

**Bogarín, Diego; Karremans, Adam P.; Rincón, Rafael; Gravendeel, Barbara (2013):**
**A new *Specklinia* (Orchidaceae: Pleurothallidinae) from Costa Rica and Panama.**
In: Phytotaxa 115 (2), p. 31. DOI: 10.11646/phytotaxa.115.2.1.

**Bone, Ruth E.; Cribb, Phillip J.; Buerki, Sven (2015):**
**Phylogenetics of Eulophiinae (Orchidaceae: Epidendroideae): evolutionary Patterns and Implications for generic Delimitation.**

In: Botanical Journal of the Linnean Society 179 (1), p. 43–56. DOI: 10.1111/boj.12299.

**Borba, Eduardo L.; Salazar, Gerardo A.; Mazzoni-Viveiros, Solange; Batista, João Aguiar Nogueira (2014):**
**Phylogenetic Position and Floral Morphology of the Brazilian endemic, monospecific Genus *Cotylolabium* : a sister Group for the remaining Spiranthinae (Orchidaceae).**
In: Botanical Journal of the Linnean Society 175 (1), p. 29–46. DOI: 10.1111/boj.12136.

**Bouetard, Anthony; Lefeuvre, Pierre; Gigant, Rodolphe; Bory, Séverine; Pignal, Marc; Besse, Pascale; Grisoni, Michel (2010):**
**Evidence of transoceanic Dispersion of the Genus *Vanilla* based on Plastid DNA phylogenetic Analysis.**
In: Molecular Phylogenetics and Evolution 55 (2), p. 621–630. DOI: 10.1016/j.ympev.2010.01.021.

**Breitkopf, Hendrik; Onstein, Renske E.; Cafasso, Donata; Schlüter, Philipp M.; Cozzolino, Salvatore (2015):**
**Multiple shifts to different pollinators fuelled rapid Diversification in sexually deceptive *Ophrys* Orchids.**
In: the new Phytologist 207 (2), p. 377–389. DOI: 10.1111/nph.13219.

**Burke, Jacinta M.; Bayly, Michael J.; Adams, Peter B.; Ladiges, Pauline Y. (2008):**
**Molecular phylogenetic Analysis of *Dendrobium* (Orchidaceae), with Emphasis on the Australian Section *Dendrocoryne*, and Implications for generic Classification.**
In: Australian Systematic Botany 21 (1), p. 1–14. DOI: 10.1071/SB07038.

**Bytebier, Benny; Bellstedt, Dirk U.; Linder, Hans Peter (2007):**
**A molecular Phylogeny for the large African Orchid Genus *Disa*.**
In: Molecular Phylogenetics and Evolution 43 (1), p. 75–90. DOI: 10.1016/j.ympev.2006.08.014.

**Bytebier, Benny; Bellstedt, Dirk U.; Linder, Hans Peter (2008):**
**A new Phylogeny-based Sectional Classification for the large African Orchid Genus *Disa*.**
In: Taxon 57 (4), p. 1233–1251. DOI: 10.1002/tax.574015.

**Cameron, Kenneth M. (2004):**
**Utility of Plastid psaB Gene Sequences for investigating intrafamilial Relationships within Orchidaceae.**
In: Molecular Phylogenetics and Evolution 31 (3), p. 1157–1180. DOI: 10.1016/j.ympev.2003.10.010.

**Cameron, Kenneth M. (2005):**
**Leave it to the leaves: a molecular phylogenetic Study of Malaxideae (Epidendroideae, Orchidaceae).**
In: American Journal of Botany 92 (6), p. 1025–1032. DOI: 10.3732/ajb.92.6.1025.

**Cameron, Kenneth M. (2009):**
**On the Value of nuclear and mitochondrial Gene Sequences for Reconstructing the Phylogeny of Vanilloid Orchids (Vanilloideae, Orchidaceae).**
In: Annals of Botany 104 (3), p. 377–385. DOI: 10.1093/aob/mcp024.

**Cameron, Kenneth M.; Chase, Mark W. (1999):**
**Phylogenetic Relationships of Pogoniinae (Vanilloideae, Orchidaceae): An Herbaceous Example of the Eastern North America-Eastern Asia Phytogeographic Disjunction.**
In: Journal of Plant Research 112 (3), p. 317–329. DOI: 10.1007/PL00013873.

**Carlsward, Barbara S.; Stern, William Louis (2009):**
**Vegetative Anatomy and Systematics of Triphorinae (Orchidaceae).**

In: Botanical Journal of the Linnean Society 159 (2), p. 203–210. DOI: 10.1111/j.1095-8339.2008.00930.x.

**Carlsward, Barbara S.; Whitten, William Mark; Williams, Norris H. (2003):**
**Molecular Phylogenetics of Neotropical Leafless Angraecinae (Orchidaceae): Reevaluation of Generic Concepts.**
In: International Journal of Plant Sciences 164 (1), p. 43–51.

**Carlsward, Barbara S.; Whitten, William Mark; Williams, Norris H. (2006):**
**Molecular Phylogenetics of Vandeae (Orchidaceae) and the Evolution of Leaflessness.**
In: American Journal of Botany 93 (5), p. 770–786.

**Carnevali, Germán; Cetzal, William; Whitten, William Mark (2012):**
***Cryptocentrum beckendorfii* (Orchidaceae: Maxillariinae), an extraordinary new Species from Andean Peru.**
In: Phytotaxa 68, p. 45–51.

**Chase, Mark W.; Cameron, Kenneth M.; Freudenstein, John V.; Pridgeon, Alec M.; Salazar, Gerardo A.; van den Berg, Cássio; Schuiteman, André (2015):**
**An updated Classification of Orchidaceae.**
In: Botanical Journal of the Linnean Society 177 (2), p. 151–174. DOI: 10.1111/boj.12234.

**Chase, Mark W.; Whitten, William Mark (2011):**
**Further taxonomic Transfers in Oncidiinae (Orchidaceae).**
In: Phytotaxa 20 (1), p. 26–32. DOI: 10.11646/phytotaxa.20.1.2.

**Chemisquy, Maria Amelia; Morrone, Osvaldo (2010):**
**Phylogenetic Analysis of the Subtribe Chloraeinae (Orchidaceae): a preliminary Approach based on three Chloroplast Markers.**
In: Australian Systematic Botany 23 (1), p. 38–46. DOI: 10.1071/SB09026.

**Chemisquy, Maria Amelia; Morrone, Osvaldo (2012):**
**Molecular Phylogeny of *Gavilea* (Chloraeinae: Orchidaceae) using Plastid and nuclear Markers.**
In: Molecular Phylogenetics and Evolution 62 (3), p. 889–897. DOI: 10.1016/j.ympev.2011.11.026.

**Chemisquy, Maria Amelia; Prevosti, Francisco J.; Morrone, Osvaldo (2009):**
**Seed Morphology in the Tribe Chloraeeae (Orchidaceae): combining traditional and geometric morphometrics.**
In: Botanical Journal of the Linnean Society 160 (2), p. 171–183. DOI: 10.1111/j.1095-8339.2009.00968.x.

**Chiron, G. R.U.Y.; Karremans, Adam P.; van den Berg, Cássio (2016):**
**Nomenclatural Notes in the Pleurothallidinae (Orchidaceae): *Phloeophila*.**
In: Phytotaxa 270 (1), p. 56. DOI: 10.11646/phytotaxa.270.1.6.

**Chiron, Guy R.; Guiard, Josiane; van den Berg, Cássio (2012):**
**Phylogenetic Relationships in Brazilian *Pleurothallis* sensu lato (Pleurothallidinae, Orchidaceae): Evidence from nuclear ITS rDNA Sequences.**
In: Phytotaxa 46, p. 34–58.

**Chiron, Guy R.; Oliveira, Reyjane Patrícia; Santos, Tarciso M.; Bellvert, Florian; Bertrand, Cédric; van den Berg, Cássio (2009):**
**Phylogeny and Evolution of *Baptistonia* (Orchidaceae, Oncidiinae) based on molecular Analyses, Morphology and Floral Oil Evidences.**
In: Plant Systematics and Evolution 281 (1-4), p. 35–49. DOI: 10.1007/s00606-009-0181-9.

**Chochai, Araya; Leitch, Ilia J.; Ingrouille, Martin J.; Fay, Michael F. (2012):**

**Molecular Phylogenetics of *Paphiopedilum* (Cypripedioideae; Orchidaceae) based on nuclear ribosomal ITS and Plastid Sequences.**
In: Botanical Journal of the Linnean Society 170 (2), p. 176–196. DOI: 10.1111/j.1095-8339.2012.01293.x.

**Cisternas, Mauricio A.; Salazar, Gerardo A.; Verdugo, Gabriela; Novoa, Patricio; Calderón, Ximena; Negritto, María A. (2012):**
**Phylogenetic Analysis of Chloraeinae (Orchidaceae) based on Plastid and nuclear DNA Sequences.**
In: Botanical Journal of the Linnean Society 168 (3), p. 258–277. DOI: 10.1111/j.1095-8339.2011.01200.x.

**Clements, Mark A. (2003):**
**Molecular phylogenetic Systematics in the Dendrobiinae (Orchidaceae), with Emphasis on *Dendrobium* Sect. *Pedilonum*.**
In: Telopea 10 (1), p. 247–298. DOI: 10.7751/telopea20035619.

**Clements, Mark A.; Howard, Christopher G.; Miller, Joseph T.H. (2015):**
***Caladenia* revisited: Results of molecular phylogenetic Analyses of Caladeniinae Plastid and nuclear loci.**
In: American Journal of Botany 102 (4), p. 581–597. DOI: 10.3732/ajb.1500021.

**Clements, Mark A.; Tupac Otero, J.; Miller, Joseph T.H. (2011):**
**Phylogenetic Relationships in Pterostylidinae (Cranichideae: Orchidaceae): combined Evidence from nuclear ribomsomal and Plastid DNA Sequences.**
In: Australian Journal of Botany 59 (2), p. 99–117. DOI: 10.1071/BT10190.

**Cox, Antony V.; Pridgeon, Alec M.; Albert, Victor A.; Chase, Mark W. (1997):**
**Phylogenetics of the slipper Orchids (Cypripedioideae, Orchidaceae): Nuclear rDNA ITS Sequences.**
In: Plant Systematics and Evolution 208 (3-4), p. 197–223. DOI: 10.1007/BF00985442.

**Cozzolino, Salvatore; Widmer, Alex (2005):**
**The evolutionary basis of reproductive isolation in Mediterranean Orchids.**
In: Taxon 54 (4), p. 977–985. DOI: 10.2307/25065482.

**Cribb, Phillip J.; Fischer, Eberhard; Killmann, Dorothee (2010):**
**A Revision of *Gastrodia* (Orchidaceae: Epidendroideae, Gastrodieae) in tropical Africa.**
In: Kew Bulletin 65 (2), p. 315–321. DOI: 10.1007/s12225-010-9193-4.

**Deng, H. U.A.; Zhang, Guo-Qiang; Liu, Zhong-Jian; Wang, Y. A.N. (2015):**
**A new Species and a new Combination of *Phalaenopsis* (Orchidaceae: Epidendroideae: Aeridinae): Evidence from morphological and DNA Analysis.**
In: Phytotaxa 238 (3), p. 243. DOI: 10.11646/phytotaxa.238.3.3.

**Devey, Dion S.; Bateman, Richard M.; Fay, Michael F.; Hawkins, Julie A. (2008):**
**Friends or relatives? Phylogenetics and Species Delimitation in the controversial European Orchid Genus *Ophrys*.**
In: Annals of Botany 101 (3), p. 385–402. DOI: 10.1093/aob/mcm299.

**Devos, Nicolas; Raspé, Olivier; Oh, Sang-Hun; Tyteca, Daniel; Jacquemart, Anne-Laure (2006):**
**The Evolution of *Dactylorhiza* (Orchidaceae) allotetraploid Complex: Insights from nrDNA Sequences and cpDNA PCR-rflp Data.**
In: Molecular Phylogenetics and Evolution 38 (3), p. 767–778. DOI: 10.1016/j.ympev.2005.11.013.

**Doucette, Alfonso; Timyan, Joel; Henrys, Ingrid; Cameron, Kenneth M. (2016):**

**A tiny new Species of *Specklinia* from Haiti's Parc National Naturel Macaya and new Combinations in *Acianthera* (Pleurothallidinae, Epidendreae, Epidendroideae, Orchidaceae).**
In: Phytotaxa 275 (3), p. 263. DOI: 10.11646/phytotaxa.275.3.4.

**Dressler, Robert L.; Whitten, William Mark; Williams, Norris H. (2004):**
**Phylogenetic Relationships of *Scaphyglottis* and related Genera (Laeliinae: Orchidaceae) based on nrDNA ITS Sequence Data.**
In: Brittonia 56 (1), p. 58–66. DOI: 10.1663/0007-196X(2004)056[0058:PROSAR]2.0.CO;2.

**Droissart, Vincent; Sonké, Bonaventure; Nguembou K., Charlemagne; Djuikouo, Marie-Noël K.; Parmentier, Ingrid; Stévart, Tariq (2009):**
**Synopsis of the Genus *Chamaeangis* (Orchidaceae), including two new Taxa.**
In: Systematic Botany 34 (2), p. 285–296. DOI: 10.1600/036364409788606361.

**Endara, Lorena A.; Williams, Norris H.; Whitten, William Mark (2011):**
**Filogenia molecular preliminar de *Scaphosepalum* (Orchidaceae: Pleurothallidinae).**
In: Lankesteriana 11 (3), p. 245–252.

**Fan, Jie; Qin, Hai-Ning; Li, De-Zhu; Jin, Xiao-Hua (2009):**
**Molecular Phylogeny and Biogeography of *Holcoglossum* (Orchidaceae: Aeridinae) based on nuclear ITS, and Chloroplast trnL-F and matK.**
In: Taxon 58 (3), p. 849–861. DOI: 10.1002/tax.583013.

**Fischer, Gunter Alexander; Gravendeel, Barbara; Sieder, Anton; Andriantiana, Jacky L.; Heiselmayer, Paul; Cribb, Phillip J. et al. (2007):**
**Evolution of Resupination in Malagasy Species of *Bulbophyllum* (Orchidaceae).**
In: Molecular Phylogenetics and Evolution 45 (1), p. 358–376. DOI: 10.1016/j.ympev.2007.06.023.

**Fraga, Claudio N.; Meneguzzo, Thiago E.C.; Saddi, Eduardo M. (2015):**
**The true identity of *Serapias nitida*, a Species of *Buchtienia* (Orchidaceae) from Brazilian Atlantic Forest.**
In: Taxon 64 (2), p. 355–361. DOI: 10.12705/642.27.

**Freudenstein, John V.; Chase, Mark W. (2015):**
**Phylogenetic Relationships in Epidendroideae (Orchidaceae), one of the great Flowering Plant Radiations: progressive Specialization and Diversification.**
In: Annals of Botany 115 (4), p. 665–681. DOI: 10.1093/aob/mcu253.

**Freudenstein, John V.; Senyo, Diana M. (2008):**
**Relationships and Evolution of matK in a Group of leafless Orchids (*Corallorhiza* and Corallorhizinae; Orchidaceae: Epidendroideae).**
In: American Journal of Botany 95 (4), p. 498–505. DOI: 10.3732/ajb.95.4.498.

**Freudenstein, John V.; van den Berg, Cássio; Goldman, Douglas H.; Kores, Paul J.; Molvray, Mia; Chase, Mark W. (2004):**
**An expanded Plastid DNA Phylogeny of Orchidaceae and Analysis of Jackknife Branch Support Strategy.**
In: American Journal of Botany 91 (1), p. 149–157. DOI: 10.3732/ajb.91.1.149.

**Gamisch, Alexander; Fischer, Gunter Alexander; Comes, Hans Peter (2016):**
**Frequent but asymmetric Niche Shifts in *Bulbophyllum* Orchids support Environmental and climatic Instability in Madagascar over Quaternary Time Scales.**
In: BMC Evolutionary Biology 16, p. 14. DOI: 10.1186/s12862-016-0586-3.

**Gamisch, Alexander; Fischer, Gunter Alexander; Comes, Hans Peter (2015):**
**Multiple independent Origins of auto-Pollination in tropical Orchids (*Bulbophyllum*) in light of the Hypothesis of selfing as an evolutionary dead end.**
In: BMC Evolutionary Biology 15, p. 192. DOI: 10.1186/s12862-015-0471-5.

**Garcia-Cruz, Javier; Sosa, Victoria (2005):**
**Phylogenetic Relationships and Character Evolution in *Govenia* (Orchidaceae).**
In: Canadian Journal of Botany 83 (10), p. 1329–1339. DOI: 10.1139/b05-098.

**Gardiner, Lauren Maria (2012):**
**New Combinations in the Genus *Vanda* (Orchidaceae).**
In: Phytotaxa 61, p. 47–54.

**Gardiner, Lauren Maria; Kocyan, Alexander; Motes, Martin; Roberts, David L.; Emerson, Brent C. (2013):**
**Molecular Phylogenetics of *Vanda* and related Genera (Orchidaceae).**
In: Botanical Journal of the Linnean Society 173 (4), p. 549–572. DOI: 10.1111/boj.12102.

**Goldman, Douglas H.; Freudenstein, John V.; Kores, Paul J.; Molvray, Mia; Jarrell, David C.; Whitten, William Mark et al. (2001):**
**Phylogenetics of Arethuseae (Orchidaceae) based on Plastid matK and rbcL Sequences.**
In: Systematic Botany 26 (3), p. 670–695.

**Goldman, Douglas H.; Jansen, Robert K.; van den Berg, Cássio; Leitch, Ilia J.; Fay, Michael F.; Chase, Mark W. (2004):**
**Molecular and cytological examination of *Calopogon* (Orchidaceae, Epidendroideae): Circumscription, Phylogeny, Polyploidy, and possible hybrid Speciation.**
In: American Journal of Botany 91 (5), p. 707–723. DOI: 10.3732/ajb.91.5.707.

**Górniak, Marcin; Paun, Ovidiu; Chase, Mark W. (2010):**
**Phylogenetic Relationships within Orchidaceae based on a low-copy nuclear coding Gene, XDH: Congruence with organellar and nuclear ribosomal DNA Results.**
In: Molecular Phylogenetics and Evolution 56 (2), p. 784–795. DOI: 10.1016/j.ympev.2010.03.003.

**Gravendeel, Barbara; Chase, Mark W.; Vogel, Eduard F.; Roos, Marco C.; Mes, Ted H. M.; Bachmann, Konrad (2001):**
**Molecular Phylogeny of *Coelogyne* (Epidendroideae; Orchidaceae) based on Plastid Rflps, matK, and nuclear ribosomal ITS Sequences: Evidence for Polyphyly.**
In: American Journal of Botany 88 (10), p. 1915–1927. DOI: 10.2307/3558367.

**Gustafsson, A.; Lovisa S.; Verola, Christiano Franco; Antonelli, Alexandre (2010):**
**Reassessing the temporal Evolution of Orchids with new Fossils and a Bayesian relaxed Clock, with Implications for the Diversification of the rare South American Genus *Hoffmannseggella* (Orchidaceae: Epidendroideae).**
In: BMC Evolutionary Biology 10, p. 177. DOI: 10.1186/1471-2148-10-177.

**Heads, Michael (2008):**
**Biological Disjunction along the West Caledonian fault, New Caledonia: a Synthesis of molecular Phylogenetics and Panbiogeography.**
In: Botanical Journal of the Linnean Society 158 (3), p. 470–488. DOI: 10.1111/j.1095-8339.2008.00866.x.

**Hermans, Johan; Andriantiana, Jacky L.; Sieder, Anton; Kiehn, Michael; Cribb, Phillip J.; Rajavelona, Landy; Gardiner, Lauren Maria (2017):**
**New Species and nomenclatural Changes in *Cynorkis* (Orchidaceae: Orchidoideae) from Madagascar and the Mascarenes.**

In: Kew Bulletin 72 (3), p. 144. DOI: 10.1007/S12225-017-9715-4.

**Hidayata, Topik; Weston, Peter H.; Yukawa, Tomohisa; Ito, Motomi; Rice, Rod (2012):**
**Phylogeny of Subtribe Aeridinae (Orchidaceae) inferred from DNA Sequences Data: Advanced Analyses including Australasian Genera.**
In: Jurnal Teknologi 59, p. 87–95.

**Hills, Harold G. (2012):**
**Taxonomic Revision of *Dressleria* (Orchidaceae, Catasetinae).**
In: Phytoneuron 48, p. 1–28.

**Hills, Harold G.; Weber, Michael H. (2012):**
***Dressleria morenoi* (Orchidaceae, Catasetinae): a new Species from Colombia.**
In: Phytoneuron 103, p. 1–5.

**Hopper, Stephen D. (2009):**
**Taxonomic turmoil Down-under: recent Developments in Australian Orchid Systematics.**
In: Annals of Botany 104 (3), p. 447–455. DOI: 10.1093/aob/mcp090.

**Hopper, Stephen D.; Brown, Andrew P. (2004):**
**Robert Brown's *Caladenia* revisited, including a Revision of its Sister Genera *Cyanicula*, *Ericksonella* and *Pheladenia* (Caladeniinae: Orchidaceae).**
In: Australian Systematic Botany 17 (2), p. 171–240. DOI: 10.1071/SB03002.

**Hopper, Stephen D.; Brown, Andrew P. (2007):**
**A Revision of Australia' s hammer Orchids (*Drakaea*: Orchidaceae), with some field Data on species-specific sexually deceived Wasp Pollinators.**
In: Australian Systematic Botany 20 (3), p. 252–285. DOI: 10.1071/SB06033.

**Hosseini, Shahla; Dadkhah, Kourosh; Go, Rusea (2016):**
**Molecular Systematics of Genus *Bulbophyllum* (Orchidaceae) in Peninsular Malaysia based on combined nuclear and Plastid DNA Sequences.**
In: Biochemical Systematics and Ecology 65, p. 40–48. DOI: 10.1016/j.bse.2016.01.003.

**Hsu, Tien-Chuan; Chung, Shih-Wen (2008):**
***Oberonia segawae* (Orchidaceae): A new Orchid Species in Taiwan.**
In: Taiwania 53 (2), p. 165–169.

**Inda, Luis A.; Pimentel, Manuel; Chase, Mark W. (2010):**
**Contribution of mitochondrial cox1 Intron Sequences to the Phylogenetics of Tribe Orchideae (Orchidaceae): Do the Distribution and Sequence of this Intron in Orchids also tell us something about its Evolution?**
In: Taxon 59 (4), p. 1053–1064. DOI: 10.1002/tax.594006.

**Inda, Luis A.; Pimentel, Manuel; Chase, Mark W. (2012):**
**Phylogenetics of Tribe Orchideae (Orchidaceae: Orchidoideae) based on combined DNA Matrices: Inferences regarding Timing of Diversification and Evolution of Pollination Syndromes.**
In: Annals of Botany 110 (1), p. 71–90. DOI: 10.1093/aob/mcs083.

**Indsto, James O.; Weston, Peter H.; Clements, Mark A. (2009):**
**A molecular phylogenetic Analysis of *Diuris* (Orchidaceae) based on AFLP and ITS reveals three major Clades and a basal Species.**
In: Australian Systematic Botany 22 (1), p. 1–15. DOI: 10.1071/SB08029.

**Janes, Jasmine K.; Duretto, Marco F. (2010):**
**A new Classification for Subtribe Pterostylidinae (Orchidaceae), Reaffirming *Pterostylis* in the broad sense.**
In: Australian Systematic Botany 23 (4), p. 260–269. DOI: 10.1071/SB09052.

**Janes, Jasmine K.; Steane, Dorothy A.; Vaillancourt, René E.; Duretto, Marco F. (2010):**
**A molecular Phylogeny of the Subtribe Pterostylidinae (Orchidaceae): resolving the taxonomic Confusion.**
In: Australian Systematic Botany 23 (4), p. 248–259. DOI: 10.1071/SB10006.

**Jersáková, Jana; Trávníček, Pavel; Kubátová, Barbora; Krejčíková, Jana; Urfus, Tomáš; Liu, Zhong-Jian et al. (2013):**
**Genome size Variation in Orchidaceae Subfamily Apostasioideae: filling the phylogenetic Gap.**
In: Botanical Journal of the Linnean Society 172 (1), p. 95–105. DOI: 10.1111/boj.12027.

**Jin, Wei-Tao; Jin, Xiao-Hua; Schuiteman, André; Li, De-Zhu; Xiang, Xiao-Guo; Huang, Wei-Chang et al. (2014):**
**Molecular Systematics of Subtribe Orchidinae and Asian Taxa of Habenariinae (Orchideae, Orchidaceae) based on Plastid matK, rbcL and nuclear ITS.**
In: Molecular Phylogenetics and Evolution 77, p. 41–53. DOI: 10.1016/j.ympev.2014.04.004.

**Jin, Wei-Tao; Xiang, Xiao-Guo; Jin, Xiao-Hua (2015):**
**Generic Delimitation of Orchidaceae from China: current Situation and Perspective.**
In: Biodiversity Science 23 (2), p. 237–242. DOI: 10.17520/biods.2014268.

**Jin, Xiao-Hua; Li, De-Zhu; Xiang, Xiao-Guo; Lai, Yang-Jun; Shi, Xiao-Chun (2012):**
***Nujiangia* (Orchidaceae: Orchideae): A new Genus from the Himalayas.**
In: Journal of Systematics and Evolution 50 (1), p. 64–71. DOI: 10.1111/j.1759-6831.2011.00167.x.

**Karremans, Adam P. (2014):**
***Lankesteriana*, a new Genus in the Pleurothallidinae (Orchidaceae).**
In: Lankesteriana 13 (3), p. 319–332.

**Karremans, Adam P. (2015):**
**Nomenclatural Notes in the Pleurothallidinae (Orchidaceae): *Stelis*.**
In: Phytotaxa 203 (3), p. 292. DOI: 10.11646/phytotaxa.203.3.9.

**Karremans, Adam P.; Albertazzi, Federico J.; Bakker, Freek T.; Bogarín, Diego; Eurlings, Marcel C.M.; Pridgeon, Alec M. et al. (2016):**
**Phylogenetic reassessment of *Specklinia* and its allied Genera in the Pleurothallidinae (Orchidaceae).**
In: Phytotaxa 272 (1), p. 1. DOI: 10.11646/phytotaxa.272.1.1.

**Karremans, Adam P.; Bakker, Freek T.; Pupulin, Franco; Solano-Gómez, Rodolfo; Smulders, Marinus J. M. (2013):**
**Phylogenetics of *Stelis* and closely related Genera (Orchidaceae: Pleurothallidinae).**
In: Plant Systematics and Evolution 299 (1), p. 151–176. DOI: 10.1007/s00606-012-0712-7.

**Karremans, Adam P.; Bogarín, Diego; Pupulin, Franco; Luer, Carlyle A.; Gravendeel, Barbara (2015):**
**The glandulous *Specklinia*: morphological Convergence versus phylogenetic Divergence.**
In: Phytotaxa 218 (2), p. 101. DOI: 10.11646/phytotaxa.218.2.1.

**Karremans, Adam P.; Pupulin, Franco; Gravendeel, Barbara (2015):**

**_Specklinia dunstervillei_, a new Species long confused with _Specklinia endotrachys_ (Orchidaceae: Pleurothallidinae).**
In: Public Library of Science One 10 (7), e0131971. DOI: 10.1371/journal.pone.0131971.

**Karremans, Adam P.; Rincón-González, Milton (2015):**
**Nomenclatural Notes in the Pleurothallidinae (Orchidaceae): _Apoda-prorepentia_.**
In: Phytotaxa 238 (2), p. 174. DOI: 10.11646/phytotaxa.238.2.5.

**Kennedy, Aaron H.; Watson, Linda E. (2010):**
**Species Delimitations and phylogenetic Relationships within the Fully Myco-heterotrophic _Hexalectris_ (Orchidaceae).**
In: Systematic Botany 35 (1), p. 64–76.

**Kita, Koichi; Handa, Takashi; Hidayata, Topik; Weston, Peter H.; Yukawa, Tomohisa; Ricef, Rod; Ito, Motomi (2005):**
**Molecular Phylogenetics of _Phalaenopsis_ (Orchidaceae) Genera: Re-evaluation of Generic Concepts.**
In: Acta Phytotaxonomica Geobotanica 56 (2), p. 141–161.

**Kocyan, Alexander; Qiu, Y.-L.; Endress, Peter K.; Conti, Elena (2004):**
**A phylogenetic Analysis of Apostasioideae (Orchidaceae) based on ITS, trnL-F and matK Sequences.**
In: Plant Systematics and Evolution 247 (3-4), p. 203–213. DOI: 10.1007/s00606-004-0133-3.

**Kocyan, Alexander; Schuiteman, André (2014):**
**New Combinations in Aeridinae (Orchidaceae).**
In: Phytotaxa 161 (1), p. 61. DOI: 10.11646/phytotaxa.161.1.3.

**Kocyan, Alexander; Vogel, Eduard F.; Conti, Elena; Gravendeel, Barbara (2008):**
**Molecular Phylogeny of _Aerides_ (Orchidaceae) based on one nuclear and two Plastid Markers: a step forward in Understanding the Evolution of the Aeridinae.**
In: Molecular Phylogenetics and Evolution 48 (2), p. 422–443. DOI: 10.1016/j.ympev.2008.02.017.

**Koehler, Samantha; Cabral, Juliano S.; Whitten, William Mark; Williams, Norris H.; Singer, Rodrigo B.; Neubig, Kurt M. et al. (2008):**
**Molecular Phylogeny of the Neotropical Genus _Christensonella_ (Orchidaceae, Maxillariinae): Species Delimitation and Insights into Chromosome Evolution.**
In: Annals of Botany 102 (4), p. 491–507. DOI: 10.1093/aob/mcn128.

**Koehler, Samantha; Williams, Norris H.; Whitten, William Mark; Amaral, Maria do Carmo E. (2002):**
**Phylogeny of the _Bifrenaria_ (Orchidaceae) Complex based on Morphology and Sequence Data from Nuclear rDNA Internal Transcribed Spacers (ITS) and Chloroplast trnL - trnF Region.**
In: International Journal of Plant Sciences 163 (6), p. 1055–1066. DOI: 10.1086/342035.

**Kolanowska, Marta; Szlachetko, Dariusz L. (2015):**
**Notes on Pachyphyllinae (Vandoideae, Orchidaceae) with a Description of a new Genus.**
In: Plant Systematics and Evolution 301 (1), p. 95–111. DOI: 10.1007/s00606-014-1057-1.

**Kolanowska, Marta; Szlachetko, Dariusz L. (2015):**
**Overview of _Cranichis_ (Orchidaceae, Cranichidinae) and allied Genera with Notes on their Colombian Representatives.**
In: Plant Systematics and Evolution 301 (2), p. 709–724. DOI: 10.1007/s00606-014-1110-0.

**Kolanowska, Marta; Szlachetko, Dariusz L. (2016):**
**Problems with generic Delimitation in the *Odontoglossum* complex (Orchidaceae, Oncidiinae) and an Attempt for a Solution.**
In: Plant Systematics and Evolution 302 (2), p. 203–217. DOI: 10.1007/s00606-015-1254-6.

**Kollmann, Ludovic Jean Charles (2010):**
**New Combinations and Description of two new Species in *Pabstiella* Brieger & Senghas (Orchidaceae) from Brazil.**
In: Candollea 65 (1), p. 95–100. DOI: 10.15553/c2010v651a8.

**Kores, Paul J.; Molvray, Mia; Weston, Peter H.; Hopper, Stephen D.; Brown, Andrew P.; Cameron, Kenneth M.; Chase, Mark W. (2001):**
**A phylogenetic Analysis of Diurideae (Orchidaceae) based on Plastid DNA Sequence Data.**
In: American Journal of Botany 88 (10), p. 1903–1914. DOI: 10.2307/3558366.

**Kras, Marta; Szlachetko, Dariusz L. (2013):**
**The Genus *Bilabrella* Lindl. (Orchidaceae, Habenariinae): General Characteristic and Research History of the Genus.**
In: Biodiversity: Research and Conservation 32 (1), p. 9–24. DOI: 10.2478/biorc-2013-0012.

**Larsen, Michael W.; Peter, Craig I.; Johnson, Steven D.; Olesen, Jens M. (2008):**
**Comparative biology of Pollination Systems in the African-Malagasy Genus *Brownleea* (Brownleeinae: Orchidaceae).**
In: Botanical Journal of the Linnean Society 156 (1), p. 65–78. DOI: 10.1111/j.1095-8339.2007.00725.x.

**Leopardi, Carlos; Carnevali, Germán; Romero-González, Gustavo A. (2012):**
***Amoana* (Orchidaceae, Laeliinae), a new Genus and Species from Mexico.**
In: Phytotaxa 65, p. 23–35.

**Leopardi, Carlos; Cumana, Luis (2015):**
**Listado de Especies de la Familia Orchidaceae para el Estado Sucre, Venezuela.**
In: Lankesteriana 8 (3). DOI: 10.15517/lank.v8i3.18323.

**Leopardi-Verde, Carlos L.; Carnevali, Germán; Romero-González, Gustavo A. (2017):**
**A Phylogeny of the Genus *Encyclia* (Orchidaceae: Laeliinae), with Emphasis on the Species of the Northern Hemisphere.**
In: Journal of Systematics and Evolution 55 (2), p. 110–123. DOI: 10.1111/jse.12225.

**Li, Ji-hong; Liu, Zhong-Jian; Salazar, Gerardo A.; Bernhardt, Peter; Perner, Holger; Tomohisa, Yukawa et al. (2011):**
**Molecular Phylogeny of *Cypripedium* (Orchidaceae: Cypripedioideae) inferred from multiple nuclear and Chloroplast Regions.**
In: Molecular Phylogenetics and Evolution 61 (2), p. 308–320. DOI: 10.1016/j.ympev.2011.06.006.

**Li, Lin; Yan, Haifei (2013):**
**A remarkable new Species of *Liparis* (Orchidaceae) from China and its phylogenetic Implications.**
In: Public Library of Science One 8 (11), e78112. DOI: 10.1371/journal.pone.0078112.

**Li, Lin; Yan, Hai-Fei; Niu, Miao; Tu, Tie-Yao; Li, Shi-Jin; Xing, Fu-Wu (2014):**
**Re-establishment of the Genus *Ania* Lindl. (Orchidaceae).**
In: Public Library of Science One 9 (7), e103129. DOI: 10.1371/journal.pone.0103129.

**Li, Lin; Ye, De-Ping; Niu, Miao; Yan, Hai-Fei; Wen, Tie-Long; Li, Shi-Jin (2015):**

***Thuniopsis*: A new Orchid Genus and Phylogeny of the Tribe Arethuseae (Orchidaceae).**
In: Public Library of Science One 10 (8), e0132777. DOI: 10.1371/journal.pone.0132777.

LI, MING-HE; ZHANG, GUO-QIANG; LAN, SI-REN; LIU, ZHONG-JIAN (2016):
**A molecular Phylogeny of Chinese Orchids.**
In: Journal of Systematics and Evolution 54 (4), p. 349–362. DOI: 10.1111/jse.12187.

LI, MING-HE; ZHANG, GUO-QIANG; LIU, ZHONG-JIAN; LAN, SI-REN (2014):
**Revision of *Hygrochilus* (Orchidaceae: Epidendroideae: Aeridinae) and a molecular phylogenetic Analysis.**
In: Phytotaxa 159 (4), p. 256. DOI: 10.11646/phytotaxa.159.4.2.

LIN, WEI-MIN; LING-LONG; HUANG, KUO; LIN, TSAN-PIAO (2006):
**Newly Discovered native Orchids of Taiwan.**
In: Taiwania 51 (3), p. 162–169.

LUER, CARLYLE A.; TOSCANO DE BRITO, A. L. V. (2011):
**Miscellaneous new Species and Combinations in the Pleurothallidinae (Orchidaceae) from Brazil and Argentina.**
In: Harvard Papers in Botany 16 (2), p. 361–382. DOI: 10.3100/0.25.016.0207.

LUO, JING; HOU, BEI-WEI; NIU, ZHI-TAO; LIU, WEI; XUE, QING-YUN; DING, XIAO-YU (2014):
**Comparative Chloroplast Genomes of photosynthetic Orchids: Insights into Evolution of the Orchidaceae and development of molecular Markers for phylogenetic Applications.**
In: Public Library of Science One 9 (6), e99016. DOI: 10.1371/journal.pone.0099016.

LUO, YI-BO; ZHU, GUANG-HUA; KURZWEIL, HUBERT (2005):
**The Gynostemium of *Hemipiliopsis purpureopunctata* and *Senghasiella glaucifolia*, two Taxonomically disputed Species of Habenariinae (Orchidaceae).**
In: Botanical Journal of the Linnean Society 147 (2), p. 191–196. DOI: 10.1111/j.1095-8339.2005.00367.x.

MARGONSKA, HANNA B.; SZLACHETKO, DARIUSZ L. (2001):
***Alatiliparis* (Orchidaceae, Malaxidinae), a new Orchid Genus with two new Species from Sumatra.**
In: Ann Bot Fennici 38, p. 77–81.

MARGONSKA, HANNA B.; SZLACHETKO, DARIUSZ L. (2012):
**Two new Combinations and a new Subsection in *Crepidium* Bl. emend. Szlach. (Orchidaceae, Malaxidinae).**
In: Annalen des Naturhistorischen Museums in Wien, Serie B 111, p. 175–180.

MARTOS, FLORENT; JOHNSON, STEVEN D.; PETER, CRAIG I.; BYTEBIER, BENNY (2014):
**A molecular Phylogeny reveals Paraphyly of the large Genus *Eulophia* (Orchidaceae): A case for the Reinstatement of *Orthochilus*.**
In: Taxon 63 (1), p. 9–23. DOI: 10.12705/631.6.

MCMURTRY, DOUGLAS; BYTEBIER, BENNY (2015):
***Disa staerkeriana* (Orchidaceae): a new Species from Mpumalanga, South Africa.**
In: Phytotaxa 203 (2), p. 192. DOI: 10.11646/phytotaxa.203.2.9.

MICHENEAU, CLAIRE; CARLSWARD, BARBARA S.; FAY, MICHAEL F.; BYTEBIER, BENNY; PAILLER, THIERRY; CHASE, MARK W. (2008):

**Phylogenetics and Biogeography of Mascarene Angraecoid Orchids (Vandeae, Orchidaceae).**
In: Molecular Phylogenetics and Evolution 46 (3), p. 908–922. DOI: 10.1016/j.ympev.2007.12.001.

**Miller, Joseph T.H.; Clements, Mark A. (2014):**
**Molecular phylogenetic Analyses of Drakaeinae: Diurideae (Orchidaceae) based on DNA Sequences of the Internal Transcribed Spacer Region.**
In: Australian Systematic Botany 27 (1), p. 3–22. DOI: 10.1071/SB13036.

**Molinari-Novoa, Eduardo Antonio (2015):**
**Homage to Christenson: Combinations under *Maxillaria*.**
In: Richardiana 15, p. 291–305.

**Monteiro, Silvana Helena N.; Selbach-Schnadelbach, Alessandra; Oliveira, Reyjane Patrícia; van den Berg, Cássio (2010):**
**Molecular Phylogenetics of *Galeandra* (Orchidaceae: Catasetinae) based on Plastid and Nuclear DNA Sequences.**
In: Systematic Botany 35 (3), p. 476–486. DOI: 10.1600/036364410792495944.

**Mytnik-Ejsmont, Joanna; Baranow, Przemysław (2010):**
**Taxonomic Study of *Polystachya* Hook. (Orchidaceae) from Asia.**
In: Plant Systematics and Evolution 290 (1-4), p. 57–63. DOI: 10.1007/s00606-010-0348-4.

**Mytnik-Ejsmont, Joanna; Szlachetko, Dariusz L.; Baranow, Przemysław; Górniak, Marcin (2014):**
**A phylogenetic and morphological Study of *Polystachya* Sect. *Superpositae* (Orchidaceae) with Description of a new Species from Cameroon.**
In: Plant Systematics and Evolution 300 (1), p. 19–28. DOI: 10.1007/s00606-013-0856-0.

**Mytnik-Ejsmont, Joanna; Szlachetko, Dariusz L.; Baranow, Przemysław; Jolliffe, Kevin; Górniak, Marcin (2015):**
**Phylogenetic Placement and Taxonomy of the Genus *Hederorkis* (Orchidaceae).**
In: Public Library of Science One 10 (4), e0122306. DOI: 10.1371/journal.pone.0122306.

**Neubig, Kurt M.; Whitten, William Mark; Blanco, Mario A.; Endara, Lorena A.; Williams, Norris H.; Koehler, Samantha (2015):**
**Preliminary molecular Phylogenetics of *Sobralia* and relatives (Orchidaceae: Sobralieae).**
In: Lankesteriana 11 (3), p. 307–317. DOI: 10.15517/lank.v11i3.18286.

**Neubig, Kurt M.; Whitten, William Mark; Fls, Norris H. Williams; Blanco, Mario A.; Endara, Lorena A.; Burleigh, John Gordon et al. (2012):**
**Generic recircumscriptions of Oncidiinae (Orchidaceae: Cymbidieae) based on maximum likelihood Analysis of combined DNA Datasets.**
In: Botanical Journal of the Linnean Society 168, p. 117–146.

**Neubig, Kurt M.; Williams, Norris H.; Whitten, William Mark; Pupulin, Franco (2009):**
**Molecular Phylogenetics and the Evolution of Fruit and Leaf Morphology of *Dichaea* (Orchidaceae: Zygopetalinae).**
In: Annals of Botany 104 (3), p. 457–467. DOI: 10.1093/aob/mcp004.

**Noguera-Savelli, Eliana J.; Germán Carnevali, F.-C.; Romero-González, Gustavo A. (2008):**
**Description of a new Species and Notes on *Crossoglossa* (Orchidaceae: Epidendroideae: Malaxideae) from the eastern Andes in Colombia and Venezuela.**
In: Brittonia 60 (3), p. 240–244. DOI: 10.1007/s12228-008-9020-5.

**Nowak, Sławomir; Szlachetko, Dariusz L.; Mytnik-Ejsmont, Joanna; Cleef, Antoine M. (2015):**

**Eight new Species of *Gomphichis* (Orchidaceae, Spiranthoideae) from Colombia.**
In: Plant Systematics and Evolution 301 (1), p. 61–76. DOI: 10.1007/s00606-014-1054-4.

**OJEDA, ISIDRO; FERNÁNDEZ-CONCHA, GERMÁN CARNEVALI; ROMERO-GONZÁLEZ, GUSTAVO A. (2009):**
***Nitidobulbon*, a new Genus of Maxillariinae (Orchidaceae).**
In: Novon: A Journal for Botanical Nomenclature 19 (1), p. 96–101. DOI: 10.3417/2007039.

**PANSARIN, EMERSON R.; SALATINO, ANTONIO; SALATINO, MARIA LUIZA FARIA (2008):**
**Phylogeny of South American Pogonieae (Orchidaceae, Vanilloideae) based on Sequences of nuclear ribosomal (ITS) and Chloroplast (psaB, rbcL, rps16, and trnL-F) DNA, with Emphasis on *Cleistes* and Discussion of biogeographic Implications.**
In: Organisms Diversity and Evolution 8 (3), p. 171–181. DOI: 10.1016/j.ode.2007.09.003.

**PANSARIN, LUDMILA M.; PANSARIN, EMERSON R.; SAZIMA, MARLIES (2014):**
**Osmophore Structure and Phylogeny of *Cirrhaea* (Orchidaceae, Stanhopeinae).**
In: Botanical Journal of the Linnean Society 176, p. 369–383.

**PEDERSEN, HENRIK Æ.; WATTHANA, SANTI; ROY, MÉLANIE; SUDDEE, SOMRAN; SELOSSE, MARC-ANDRÉ (2009):**
***Cephalanthera exigua* rediscovered: new Insights in the Taxonomy, Habitat Requirements and breeding System of a rare mycoheterotrophic Orchid.**
In: Nordic Journal of Botany 27 (6), p. 460–468. DOI: 10.1111/j.1756-1051.2009.00465.x.

**PEDRON, MARCELO; BUZATTO, CRISTIANO ROBERTO; RAMALHO, ALINE JOSEPH; CARVALHO, BRUNO M.; RADINS, JOSÉ A.; SINGER, RODRIGO B.; BATISTA, JOÃO AGUIAR NOGUEIRA (2014):**
**Molecular Phylogenetics and taxonomic Revision of *Habenaria* Section *Pentadactylae* (Orchidaceae, Orchidinae).**
In: Botanical Journal of the Linnean Society 175 (1), p. 47–73. DOI: 10.1111/boj.12161.

**PERAZA-FLORES, LIZANDRON.; CARNEVALI, GERMÁN; VAN DEN BERG, CÁSSIO (2016):**
**A molecular Phylogeny of the *Laelia* alliance (Orchidaceae) and a reassessment of *Laelia* and *Schomburgkia*.**
In: Taxon 65 (6), p. 1249–1262. DOI: 10.12705/656.3.

**PÉREZ-ESCOBAR, OSCAR ALEJANDRO; GOTTSCHLING, MARC; WHITTEN, WILLIAM MARK; SALAZAR, GERARDO A.; GERLACH, GÜNTER (2016):**
**Sex and the Catasetinae (Darwin's favourite Orchids).**
In: Molecular Phylogenetics and Evolution 97, p. 1–10. DOI: 10.1016/j.ympev.2015.11.019.

**PÉREZ-ESCOBAR, OSCAR ALEJANDRO; KOLANOWSKA, MARTA; RINCÓN-USECHE, CRISTIAN (2013):**
**A new Species of *Lepanthes* (Pleurothallidinae, Orchidaceae) from Colombia.**
In: Systematic Botany 38 (2), p. 316–319. DOI: 10.1600/036364413X666822.

**PESSOA, EDLLEY MAX; ALVES, MARCCUS V.; ALVES-ARAÚJO, ANDERSON; PALMA-SILVA, CLARISSE; PINHEIRO, FÁBIO (2012):**
**Integrating different Tools to disentangle Species Complexes: A case Study in *Epidendrum* (Orchidaceae).**
In: Taxon 61 (4), p. 721–734. DOI: 10.1002/tax.614002.

**PILLON, YOHAN; FAY, MICHAEL F.; HEDRÉN, MIKAEL; BATEMAN, RICHARD M.; DEVEY, DION S.; SHIPUNOV, ALEXEY B. ET AL. (2007):**
**Evolution and temporal Diversification of western European polyploid Species Complexes in *Dactylorhiza* (Orchidaceae).**
In: Taxon 56 (4), p. 1185–1208. DOI: 10.2307/25065911.

**PINHEIRO, FÁBIO; COZZOLINO, SALVATORE (2013):**

**_Epidendrum_ (Orchidaceae) as a Model System for ecological and evolutionary Studies in the Neotropics.**
In: Taxon 62 (1), p. 77–78.

Pinheiro, Fábio; Koehler, Samantha; Corrêa, Andréa Macêdo; Salatino, Maria Luiza Faria; Salatino, Antonio; Barros, Fábio (2009):
**Phylogenetic Relationships and infrageneric Classification of _Epidendrum_ Subgenus _Amphiglottium_ (Laeliinae, Orchidaceae).**
In: Plant Systematics and Evolution 283 (3-4), p. 165–177. DOI: 10.1007/s00606-009-0224-2.

Ponsie, M. E.; Mitchell, Anthony D.; Edwards, Trevor J.; Johnson, Steven D. (2007):
**Phylogeny of _Bonatea_ (Orchidaceae: Habenariinae) based on molecular and morphological Data.**
In: Plant Systematics and Evolution 263 (3-4), p. 253–268. DOI: 10.1007/s00606-006-0487-9.

Pridgeon, Alec M.; Solano, Rodolfo; Chase, Mark W. (2001):
**Phylogenetic Relationships in Pleurothallidinae (Orchidaceae): combined Evidence from nuclear and Plastid DNA Sequences.**
In: American Journal of Botany 88 (12), p. 2286–2308. DOI: 10.2307/3558390.

Pupulin, Franco; Bogarín, Diego (2007):
**A second Species of _Restrepiella_ (Orchidaceae: Pleurothallidinae).**
In: Willdenowia 37 (1), p. 323–329. DOI: 10.3372/wi.37.37122.

Pupulin, Franco; Karremans, Adam P.; Belfort Oconitrillo, Noelia (2017):
**Two new Species of _Echinosepala_ (Orchidaceae: Pleurothallidinae).**
In: Lankesteriana 17 (2). DOI: 10.15517/lank.v17i2.30205.

Pupulin, Franco; Karremans, Adam P.; Gravendeel, Barbara (2012):
**A Reconsideration of the empusellous Species of _Specklinia_ (Orchidaceae: Pleurothallidinae) in Costa Rica.**
In: Phytotaxa 63, p. 1–20.

Quintanilla-Quintero, Sonia; Ortiz, Pedro L.; Bernal, Jaime E.; Gómez, Alberto (2011):
**Phylogenetic Relationships among Genera of the Subtribe Oncidiinae Epidendroideae: Orchidaceae and a new Genus: _Santanderella_.**
In: Phytologia 93 (3), p. 388–406.

Rakotoarivelo, Fanny P.; Razafimandimbison, Sylvain G.; Mallet, Bertrand; Faliniaina, Lucien; Pailler, Thierry (2012):
**Molecular Systematics and evolutionary Trends and Relationships in the Genus _Jumellea_ (Orchidaceae): Implications for its Species limits.**
In: Taxon 61 (3), p. 534–544. DOI: 10.1002/tax.613004.

Raskoti, Bhakta Bahadur; Jin, Wei-Tao; Xiang, Xiao-Guo; Schuiteman, André; Li, De-Zhu; Li, Jian-Wu et al. (2016):
**A phylogenetic Analysis of molecular and morphological Characters of _Herminium_ (Orchidaceae, Orchideae): evolutionary Relationships, Taxonomy, and Patterns of Character Evolution.**
In: Cladistics 32 (2), p. 198–210. DOI: 10.1111/cla.12125.

Romero-González, Gustavo A.; Batista, João Aguiar Nogueira; Bem Bianchetti, Luciano (2008):
**A Synopsis of the Genus _Cyrtopodium_ (Catasetinae: Orchidaceae).**
In: Harvard Papers in Botany 13 (1), p. 189–206. DOI: 10.3100/1043-4534(2008)13[189:ASOTGC]2.0.CO;2.

**Romero-González, Gustavo A.; Fernández-Concha, Germán Carnevali; Brito, A. L. V. Toscano de (2016):**
**Novelties in the Orchid Flora of Venezuela IX. Subtribe Pleurothallidinae. New Combinations in *Anathallis* and a new Report for the Orchid Flora of Colombia.**
In: Harvard Papers in Botany 21 (1), p. 23–29. DOI: 10.3100/hpib.v21iss1.2016.n3.

**Russell, Anton; Samuel, Rosabelle; Bogarín, Diego; Fernando, Suranjan; Wijesundera, Siril; Klejna, Verena; Chase, Mark W. (2011):**
**Genetic Variation and phylogenetic Relationships of a pantropical Species Group in *Polystachya* (Orchidaceae).**
In: Botanical Journal of the Linnean Society 165 (3), p. 235–250. DOI: 10.1111/j.1095-8339.2010.01108.x.

**Russell, Anton; Samuel, Rosabelle; Rupp, Barbara; Barfuss, Michael H. J.; Šafran, Marko; Besendorfer, Višnja; Chase, Mark W. (2010):**
**Phylogenetics and Cytology of a pantropical Orchid Genus *Polystachya* (Polystachyinae, Vandeae, Orchidaceae): Evidence from Plastid DNA Sequence Data.**
In: Taxon 59 (2), p. 389–404. DOI: 10.1002/tax.592005.

**Saeki, Ikuyo; Kitazawa, Asako; Abe, Atsushi; Minemoto, Koya; Koike, Fumito (2014):**
**Phylogeography of a rare Orchid, *Vexillabium yakushimense*: Comparison of Populations in central Honshu and the Nansei Island Chain, Japan.**
In: Plant Systematics and Evolution 300 (1), p. 1–12. DOI: 10.1007/s00606-013-0854-2.

**Salazar, Gerardo A.; Ballesteros-Barrera, Claudia (2010):**
***Sotoa*, a new Genus of Spiranthinae (Orchidaceae) from Mexico and the Southern United States.**
In: Lankesteriana 9 (3), p. 491–504.

**Salazar, Gerardo A.; Cabrera, Lidia I.; Figueroa, Coyolxauhqui (2011):**
**Molecular Phylogenetics, Floral convergence and Systematics of *Dichromanthus* and *Stenorrhynchos* (Orchidaceae: Spiranthinae).**
In: Botanical Journal of the Linnean Society 167 (1), p. 1–18. DOI: 10.1111/j.1095-8339.2011.01161.x.

**Salazar, Gerardo A.; Cabrera, Lidia I.; Madriñán, Santiago; Chase, Mark W. (2009):**
**Phylogenetic Relationships of Cranichidinae and Prescottiinae (Orchidaceae, Cranichideae) inferred from Plastid and nuclear DNA Sequences.**
In: Annals of Botany 104 (3), p. 403–416. DOI: 10.1093/aob/mcn257.

**Salazar, Gerardo A.; Chase, Mark W.; Soto-Arenas, Miguel A.; Ingrouille, Martin J. (2003):**
**Phylogenetics of Cranichideae with Emphasis on Spiranthinae (Orchidaceae, Orchidoideae): Evidence from Plastid and nuclear DNA Sequences.**
In: American Journal of Botany 90 (5), p. 777–795. DOI: 10.3732/ajb.90.5.777.

**Salazar, Gerardo A.; van den Berg, Cássio; Popovkin, Alex (2014):**
**Phylogenetic Relationships of *Discyphus scopulariae* (Orchidaceae, Cranichideae) inferred from Plastid and nuclear DNA Sequences: Evidence supporting Recognition of a new Subtribe, Discyphinae.**
In: Phytotaxa 173 (2), p. 127. DOI: 10.11646/phytotaxa.173.2.3.

**Sardaro, Maria Luisa Savo; Atallah, Maroun; Picarella, Maurizio Enea; Aracri, Benedetto; Pagnotta, Mario A. (2012):**
**Genetic Diversity, Population Structure and phylogenetic Inference among Italian Orchids of the *Serapias* Genus assessed by AFLP molecular Markers.**
In: Plant Systematics and Evolution 298 (9), p. 1701–1710. DOI: 10.1007/s00606-012-0671-z.

**Schuiteman, André; Bonnet, Pierre; Svengsuksa, Bouakhaykhone; Barthélémy, Daniel (2008):**

**An annotated Checklist of the Orchidaceae of Laos.**
In: Nordic Journal of Botany 26 (5-6), p. 257–316. DOI: 10.1111/j.1756-1051.2008.00265.x.

SCHUITEMAN, ANDRÉ; CHASE, MARK W. (2015):
**A reappraisal of *Maxillaria* (Orchidaceae).**
In: Phytotaxa 225 (1), p. 1. DOI: 10.11646/phytotaxa.225.1.1.

SCHUITEMAN, ANDRÉ; VOGEL, EDUARD F. (2003):
**New Names and Combinations in Orchidaceae from the Philippines and New Guinea.**
In: Blumea 48 (3), p. 507–514. DOI: 10.3767/000651903X489500.

SHIPUNOV, ALEXEY B.; FAY, MICHAEL F.; PILLON, YOHAN; BATEMAN, RICHARD M.; CHASE, MARK W. (2004):
***Dactylorhiza* (Orchidaceae) in European Russia: Combined molecular and morphological Analysis.**
In: American Journal of Botany 91 (9), p. 1419–1426.

SIMO-DROISSART, MURIELLE; MICHENEAU, CLAIRE; SONKÉ, BONAVENTURE; DROISSART, VINCENT; PLUNKETT, GREGORY M.; LOWRY, PORTER PRESCOTT II. ET AL. (2013):
**Morphometrics and molecular Phylogenetics of the continental African Species of *Angraecum* Section *Pectinaria* (Orchidaceae).**
In: Plant Ecology and Evolution 146 (3), p. 295–309. DOI: 10.5091/plecevo.2013.900.

SIMO-DROISSART, MURIELLE; PLUNKETT, GREGORY M.; DROISSART, VINCENT; EDWARDS, MOLLY B.; FARMINHÃO, JOÃO N. M.; JEČMENICA, VLADIMIR ET AL. (2018):
**New phylogenetic Insights toward Developing a natural generic Classification of African Angraecoid Orchids (Vandeae, Orchidaceae).**
In: Molecular Phylogenetics and Evolution 126, p. 241–249. DOI: 10.1016/j.ympev.2018.04.021.

SIMO-DROISSART, MURIELLE; SONKÉ, BONAVENTURE; DROISSART, VINCENT; MICHENEAU, CLAIRE; LOWRY, PORTER PRESCOTT; HARDY, OLIVIER J. ET AL. (2016):
**Morphometrics and molecular Phylogenetics of *Angraecum* Section *Dolabrifolia* (Orchidaceae, Angraecinae).**
In: Plant Systematics and Evolution 302 (8), p. 1027–1045. DOI: 10.1007/s00606-016-1315-5.

SMIDT, ERIC C.; BORBA, EDUARDO L.; GRAVENDEEL, BARBARA; FISCHER, GUNTER ALEXANDER; VAN DEN BERG, CÁSSIO (2011):
**Molecular Phylogeny of the Neotropical Sections of *Bulbophyllum* (Orchidaceae) using Nuclear and Plastid Spacers.**
In: Taxon 60 (4), p. 1050–1064.

SOLIVA, M.; KOCYAN, ALEXANDER; WIDMER, ALEX (2001):
**Molecular Phylogenetics of the sexually deceptive Orchid Genus *Ophrys* (Orchidaceae) based on nuclear and Chloroplast DNA Sequences.**
In: Molecular Phylogenetics and Evolution 20 (1), p. 78–88. DOI: 10.1006/mpev.2001.0953.

SOSA, VICTORIA (2007):
**A molecular and morphological phylogenetic Study of Subtribe Bletiinae (Epidendreae, Orchidaceae).**
In: Systematic Botany 32 (1), p. 34–42.

SRAMKÓ, GÁBOR; ATTILA, MOLNÁR V.; HAWKINS, JULIE A.; BATEMAN, RICHARD M. (2014):
**Molecular Phylogeny and evolutionary History of the Eurasiatic Orchid Genus *Himantoglossum* s.l. (Orchidaceae).**
In: Annals of Botany 114 (8), p. 1609–1626. DOI: 10.1093/aob/mcu179.

SRIKANTH, KRISHNAMOORTHY; KOO, JA CHOON; WHANG, SUNG SOO (2013):

**Genetic Relationships between Korean *Calanthe* Species and some naturally occurring Mutants based on multiple DNA Markers.**
In: Nordic Journal of Botany 31 (6), p. 757–766. DOI: 10.1111/j.1756-1051.2013.00152.x.

**Surveswaran, Siddharthan; Gowda, Vinita; Sun, Mei (2018):**
**Using an integrated approach to identify cryptic species, Divergence Patterns and hybrid Species in Asian ladies' tresses Orchids (*Spiranthes*, Orchidaceae).**
In: Molecular Phylogenetics and Evolution 124, p. 106–121. DOI: 10.1016/j.ympev.2018.02.025.

**Szlachetko, Dariusz L.; Górniak, Marcin; Kolanowska, Marta; Mytnik-Ejsmont, Joanna; Kowalkowska, Agnieszka K.; Rutkowski, Piotr; Koliński, Tomasz (2014):**
**Taxonomic Position and Phylogeny of the Genus *Vargasiella* (Orchidaceae, Vandoideae) based on molecular and morphological Evidence.**
In: Public Library of Science One 9 (6), e98472. DOI: 10.1371/journal.pone.0098472.

**Szlachetko, Dariusz L.; Kolanowska, Marta (2013):**
**Three new Species of *Ponthieva* (Orchidaceae, Spiranthoideae) from Colombia and Venezuela.**
In: Plant Systematics and Evolution 299 (9), p. 1671–1678. DOI: 10.1007/s00606-013-0824-8.

**Szlachetko, Dariusz L.; Kolanowska, Marta (2014):**
**A new Species of *Scaphyglottis* (Orchidaceae, Epidendroideae) from Colombia.**
In: Plant Systematics and Evolution 300 (5), p. 1031–1034. DOI: 10.1007/s00606-013-0941-4.

**Szlachetko, Dariusz L.; Kolanowska, Marta (2015):**
**Reconsideration of *Heteranthocidium* (Oncidiinae, Orchidaceae): new Species and taxonomic Transfers.**
In: Plant Systematics and Evolution 301 (7), p. 1793–1805. DOI: 10.1007/s00606-014-1189-3.

**Szlachetko, Dariusz L.; Nowak, Sławomir (2014):**
**Two new Species of *Aa* (Orchidaceae, Spiranthoideae) from Colombia.**
In: Plant Systematics and Evolution 300 (8), p. 1833–1841. DOI: 10.1007/s00606-014-1010-3.

**Szlachetko, Dariusz L.; Sitko, Magdalena; Tukałło, Piotr; Mytnik-Ejsmont, Joanna (2012):**
**Taxonomy of the Subtribe Maxillariinae (Orchidaceae, Vandoideae) revised.**
In: Biodiversity: Research and Conservation 25 (1), p. 13–38. DOI: 10.2478/v10119-012-0017-2.

**Takamiya, Tomoko; Wongsawad, Pheravut; Sathapattayanon, Apirada; Tajima, Natsuko; Suzuki, Shunichiro; Kitamura, Saki et al. (2014):**
**Molecular Phylogenetics and Character Evolution of morphologically diverse Groups, *Dendrobium* Section *Dendrobium* and Allies.**
In: AoB PLANTS 6. DOI: 10.1093/aobpla/plu045.

**Tang, Ying; Yukawa, Tomohisa; Bateman, Richard M.; Jiang, Hong; Peng, Hua (2015):**
**Phylogeny and Classification of the East Asian *Amitostigma* alliance (Orchidaceae: Orchideae) based on six DNA Markers.**
In: BMC Evolutionary Biology 15, p. 96. DOI: 10.1186/s12862-015-0376-3.

**Topik, Hidayat; Yukawa, Tomohisa; Ito, Motomi (2005):**
**Molecular Phylogenetics of Subtribe Aeridinae (Orchidaceae): Insights from Plastid matK and nuclear ribosomal ITS Sequences.**
In: Journal of Plant Research 118 (4), p. 271–284. DOI: 10.1007/s10265-005-0217-3.

**Tsai, Chi-Chu; Chiang, Yu-Chung; Huang, S.C.; Chen, C. H.; Chou, Chang-Hung (2010):**

**Molecular Phylogeny of *Phalaenopsis* Blume (Orchidaceae) on the basis of Plastid and nuclear DNA.**
In: Plant Systematics and Evolution 288 (1-2), p. 77–98. DOI: 10.1007/s00606-010-0314-1.

TSAI, CHI-CHU; HUANG, S.C.; CHOU, CHANG-HUNG (2005):
**Molecular Phylogeny of *Phalaenopsis* Blume (Orchidaceae) based on the Internal Transcribed Spacer of the nuclear ribosomal DNA.**
In: Plant Systematics and Evolution 256 (1-4), p. 1–16. DOI: 10.1007/s00606-005-0356-y.

TSUKAYA, HIROKAZU; NAKAJIMA, MUTSUKO; OKADA, HIROSHI (2011):
***Kalimantanorchis*: a new Genus of Mycotrophic Orchid from West Kalimantan, Borneo.**
In: Systematic Botany 36 (1), p. 49–52. DOI: 10.1600/036364411X553117.

TSUTSUMI, CHIE; YUKAWA, TOMOHISA; LEE, NAM SOOK; LEE, CHANG-SHOOK; KATO, MASAHIRO (2007):
**Phylogeny and comparative Seed Morphology of epiphytic and terrestrial Species of *Liparis* (Orchidaceae) in Japan.**
In: Journal of Plant Research 120 (3), p. 405–412. DOI: 10.1007/s10265-007-0077-0.

VAN DEN BERG, CÁSSIO (2014):
**Reaching a compromise between conflicting nuclear and Plastid phylogenetic Trees: a new Classification for the Genus *Cattleya* (Epidendreae; Epidendroideae; Orchidaceae).**
In: Phytotaxa 186 (2), p. 75. DOI: 10.11646/phytotaxa.186.2.2.

VAN DEN BERG, CÁSSIO (2015):
**Nomenclatural Notes in *Guarianthe* (Orchidaceae: Laeliinae): Clarification of *Guarianthe ×deckeri, G. ×guatemalensis and G. patinii*.**
In: Phytotaxa 239 (1), p. 65. DOI: 10.11646/phytotaxa.239.1.6.

VAN DEN BERG, CÁSSIO; GOLDMAN, DOUGLAS H.; FREUDENSTEIN, JOHN V.; PRIDGEON, ALEC M.; CAMERON, KENNETH M.; CHASE, MARK W. (2005):
**An overview of the phylogenetic Relationships within Epidendroideae inferred from multiple DNA Regions and Recircumscription of Epidendreae and Arethuseae (Orchidaceae).**
In: American Journal of Botany 92 (4), p. 613–624. DOI: 10.3732/ajb.92.4.613.

VAN DEN BERG, CÁSSIO; HIGGINS, WESLEY E.; DRESSLER, ROBERT L.; WHITTEN, WILLIAM MARK; SOTO-ARENAS, MIGUEL A.; CHASE, MARK W. (2009):
**A phylogenetic Study of Laeliinae (Orchidaceae) based on combined nuclear and Plastid DNA Sequences.**
In: Annals of Botany 104 (3), p. 417–430. DOI: 10.1093/aob/mcp101.

VAN DER NIET, TIMOTHEÜS; JÜRGENS, A.; JOHNSON, STEVEN D. (2010):
**Pollinators, Floral Morphology and scent Chemistry in the southern African Orchid Genus *Schizochilus*.**
In: South African Journal of Botany 76 (4), p. 726–738. DOI: 10.1016/j.sajb.2010.07.004.

VAN DER NIET, TIMOTHEÜS; LILTVED, WILLIAM R.; JOHNSON, STEVEN D. (2011):
**More than meets the Eye: a morphological and phylogenetic Comparison of long-spurred, white-flowered *Satyrium* Species (Orchidaceae) in South Africa.**
In: Botanical Journal of the Linnean Society 166 (4), p. 417–430. DOI: 10.1111/j.1095-8339.2011.01143.x.

VAN DER NIET, TIMOTHEÜS; LINDER, HANS PETER; BYTEBIER, BENNY; BELLSTEDT, DIRK U. (2005):

**Molecular Markers Reject Monophyly of the Subgenera of *Satyrium* (Orchidaceae).**
In: Systematic Botany 30 (2), p. 263–274.

VAN DER NIET, TIMOTHEÜS; PETER LINDER, H. (2008):
**Dealing with Incongruence in the Quest for the Species Tree: a case Study from the Orchid Genus *Satyrium*.**
In: Molecular Phylogenetics and Evolution 47 (1), p. 154–174. DOI: 10.1016/j.ympev.2007.12.008.

VÁZQUEZ, FRANCISCO M.; JOSÉ BLANCO, DAVID GARCÍA; MÁRQUEZ, FRANCISCO; GUERRA, MARÍA JOSÉ (2015):
**Review of *Anacamptis* Sect. *Morianthus* Taxa from SW-Iberian Peninsula.**
In: Journal Europäischer Orchideen 47 (2-4), p. 338–364.

VERMA, DURGESH; SINGH, P. K.; MAO, A. A. (2013):
**Notes on Diversity and Distribution of Tribe Orchideae in *Meghalaya*.**
In: Keanean Journal of Science 2, p. 83–92.

VERMEULEN, JAAP J. (2008):
**New Species of *Bulbophyllum* from eastern Malesia (Orchidaceae).**
In: Nordic Journal of Botany 26 (3-4), p. 129–195. DOI: 10.1111/j.1756-1051.2008.00220.x.

VERMEULEN, JAAP J.; SCHUITEMAN, ANDRÉ; VOGEL, EDUARD F. (2014):
**Nomenclatural Changes in *Bulbophyllum* (Orchidaceae; Epidendroideae).**
In: Phytotaxa 166 (2), p. 101. DOI: 10.11646/phytotaxa.166.2.1.

VERMEULEN, JAAP J.; VOGEL, EDUARD F.; VOGEL, A.P.T.M. (2010):
**A new Species of *Bulbophyllum* Section *Epicrianthes* (Orchidaceae) from Papua Province, Indonesia.**
In: Blumea 55 (3), p. 276–277. DOI: 10.3767/000651910X547101.

WATERMAN, RICHARD J.; PAUW, ANTON; BARRACLOUGH, TIMOTHY G.; SAVOLAINEN, VINCENT (2009):
**Pollinators underestimated: a molecular Phylogeny reveals widespread Floral Convergence in oil-secreting Orchids (Sub-tribe Coryciinae) of the Cape of South Africa.**
In: Molecular Phylogenetics and Evolution 51 (1), p. 100–110. DOI: 10.1016/j.ympev.2008.05.020.

WHITTEN, WILLIAM MARK; BLANCO, MARIO A.; WILLIAMS, NORRIS H.; KOEHLER, SAMANTHA; CARNEVALI, GERMÁN; SINGER, RODRIGO B. ET AL. (2007):
**Molecular Phylogenetics of *Maxillaria* and related Genera (Orchidaceae: Cymbidieae) based on combined molecular Data Sets.**
In: American Journal of Botany 94 (11), p. 1860–1889. DOI: 10.3732/ajb.94.11.1860.

WHITTEN, WILLIAM MARK; NEUBIG, KURT M.; WILLIAMS, NORRIS H. (2014):
**Generic and Subtribal Relationships in Neotropical Cymbidieae (Orchidaceae) based on matK/ycf1 Plastid Data.**
In: Lankesteriana 13 (3), p. 375–392. DOI: 10.15517/lank.v13i3.14425.

WHITTEN, WILLIAM MARK; WILLIAMS, NORRIS H.; CHASE, MARK W. (2000):
**Subtribal and generic Relationships of Maxillarieae (Orchidaceae) with Emphasis on Stanhopeinae: combined molecular Evidence.**
In: American Journal of Botany 87 (12), p. 1842–1856. DOI: 10.2307/2656837.

WHITTEN, WILLIAM MARK; WILLIAMS, NORRIS H.; DRESSLER, ROBERT L.; GERLACH, GÜNTER; PUPULIN, FRANCO (2005):
**Generic Relationships of Zygopetalinae (Orchidaceae: Cymbidieae): combined molecular Evidence.**
In: Lankesteriana 5 (2), p. 87–107.

**Williams, Norris H.; Whitten, William Mark; Dressler, Robert L. (2005):**

**Molecular Systematics of *Telipogon* (Orchidaceae: Oncidiinae) and ITS Allies: Nuclear and Plastid DNA Sequence Data.**

In: Lankesteriana 5 (3), p. 163–184.

**Wilson, Mark; Frank, Graham S.; Jost, Lou; Pridgeon, Alec M.; Vieira-Uribe, Sebastian; Karremans, Adam P. (2017):**

**Phylogenetic Analysis of *Andinia* (Pleurothallidinae; Orchidaceae) and a systematic Re-Circumscription of the Genus.**

In: Phytotaxa 295 (2), p. 101. DOI: 10.11646/phytotaxa.295.2.1.

**Wonnapinij, Passorn; Sriboonlert, Ajaraporn (2015):**

**Molecular Phylogenetics of Species of *Bulbophyllum* Sect. *Trias* (Orchidaceae; Epidendroideae; Malaxidae) based on nrITS and Plastid rbcL and matK.**

In: Phytotaxa 226 (1), p. 1–17. DOI: 10.11646/phytotaxa.226.1.1.

**Xiang, Xiao-Guo; Jin, Wei-Tao; Li, De-Zhu; Schuiteman, André; Huang, Wei-Chang; Li, Jian-Wu et al. (2014):**

**Phylogenetics of Tribe Collabieae (Orchidaceae, Epidendroideae) based on four Chloroplast Genes with morphological Appraisal.**

In: Public Library of Science One 9 (1), e87625. DOI: 10.1371/journal.pone.0087625.

**Xiang, Xiao-Guo; Li, De-Zhu; Jin, Wei-Tao; Zhou, Hai-Lang; Li, Jian-Wu; Jin, Xiao-Hua (2012):**

**Phylogenetic Placement of the enigmatic Orchid Genera *Thaia* and *Tangtsinia*: Evidence from molecular and morphological Characters.**

In: Taxon 61 (1), p. 45–54. DOI: 10.1002/tax.611003.

**Xiang, Xiao-Guo; Schuiteman, André; Li, De-Zhu; Huang, Wei-Chang; Chung, Shih-Wen; Li, Jian-Wu et al. (2013):**

**Molecular Systematics of *Dendrobium* (Orchidaceae, Dendrobieae) from mainland Asia based on Plastid and nuclear Sequences.**

In: Molecular Phylogenetics and Evolution 69 (3), p. 950–960. DOI: 10.1016/j.ympev.2013.06.009.

**Yang, Jun-Bo; Tang, Min; Li, Hong-Tao; Zhang, Zhi-Rong; Li, De-Zhu (2013):**

**Complete Chloroplast Genome of the Genus *Cymbidium*: lights into the Species Identification, phylogenetic Implications and Population genetic Analyses.**

In: BMC Evolutionary Biology 13, p. 84. DOI: 10.1186/1471-2148-13-84.

**Yukawa, Tomohisa; Cribb, Phillip J. (2014):**

**Nomenclatural Changes in the Genus *Calanthe* (Orchidaceae).**

In: Bulletin of the National Museum of Nature and Science, Tokyo, Series Botany 40 (4), p. 145–151.

**Yukawa, Tomohisa; Ohba, Hideaki; Cameron, Kenneth M.; Chase, Mark W. (1996):**

**Chloroplast DNA Phylogeny of Subtribe Dendrobiinae (Orchidaceae): Insights from a combined Analysis based on RbcL Sequences and Restriction Site Variation.**

In: Journal of Plant Research 109 (2), p. 169–176. DOI: 10.1007/BF02344542.

**Zhai, Jun-Wen; Zhang, Guo-Qiang; Chen, Li-Jun; Xiao, Xin-Ju; Liu, Ke-Wei; Tsai, Wen-Chieh et al. (2013):**

**A new Orchid Genus, *Danxiaorchis*, and phylogenetic Analysis of the Tribe Calypsoeae.**

In: Public Library of Science One 8 (4), e60371. DOI: 10.1371/journal.pone.0060371.

**Zhai, Jun-Wen; Zhang, Guo-Qiang; Li, Lin; Wang, Meina; Chen, Li-Jun; Chung, Shih-Wen et al. (2014):**

**A new phylogenetic Analysis sheds new Light on the Relationships in the *Calanthe* alliance (Orchidaceae) in China.**

In: Molecular Phylogenetics and Evolution 77, p. 216–222. DOI: 10.1016/j.ympev.2014.04.005.

Zhang, Guo-Qiang; Liu, Ke-Wei; Chen, Li-Jun; Xiao, Xin-Ju; Zhai, Jun-Wen; Li, Li-Qiang et al. (2013):
**A new molecular Phylogeny and a new Genus, *Pendulorchis*, of the *Aerides-Vanda* alliance (Orchidaceae: Epidendroideae).**
In: Public Library of Science One 8 (4), e60097. DOI: 10.1371/journal.pone.0060097.

Zou, Long-Hai; Huang, Jiu-Xiang; Zhang, Guo-Qiang; Liu, Zhong-Jian; Zhuang, Xue-Ying (2015):
**A molecular Phylogeny of Aeridinae (Orchidaceae: Epidendroideae) inferred from multiple nuclear and Chloroplast Regions.**
In: Molecular Phylogenetics and Evolution 85, p. 247–254. DOI: 10.1016/j.ympev.2015.02.014.

### Orobanchaceae

Dong, Li-Na; Wang, Hong; Wortley, Alexandra H.; Lu, Lu; Li, De-Zhu (2013):
**Phylogenetic Relationships in the *Pterygiella* complex (Orobanchaceae) inferred from molecular and morphological Evidence.**
In: Botanical Journal of the Linnean Society 171, p. 491–507.

Eaton, Deren A. R.; Ree, Richard H. (2013):
**Inferring Phylogeny and Introgression using RADseq Data: an Example from Flowering Plants (*Pedicularis*: Orobanchaceae).**
In: Systematic Biology 62 (5), p. 689–706. DOI: 10.1093/sysbio/syt032.

Fischer, Eberhard; Schäferhoff, Bastian; Müller, Kai F. (2012):
**The new monotypic Genus *Bardotia* (Orobanchaceae) from Madagascar and Remarks on the phylogenetic Relationships of the African and Madagascan Genera *Parastriga*, *Radamaea, Rhamphicarpa* and *Sieversandreas*.**
In: Phytotaxa 46, p. 19–33.

Frajman, Božo; Carlón, Luis; Kosachev, Petr A.; Sánchez Pedraja, Óscar; Schneeweiss, Gerald M.; Schönswetter, Peter (2013):
**Phylogenetic Position and Taxonomy of the enigmatic *Orobanche krylowii* (Orobanchaceae), a predominatly Asian Species newly found in Albania (SE Europe).**
In: Phytotaxa 137 (1), p. 1. DOI: 10.11646/phytotaxa.137.1.1.

Garg, Arti (2009):
**Critical taxonomic Appraisal of some Taxa of *Pedicularis* from Indian Himalayas belonging to Section *Siphonanthae*.**
In: Taiwania 54 (2), p. 122–133.

Gaudeul, Myriam; Véla, Errol; Rouhan, Germinal (2016):
**Eastward Colonization of the Mediterranean Basin by two geographically Structured Clades: the Case of *Odontites* Ludw. (Orobanchaceae).**
In: Molecular Phylogenetics and Evolution 96, p. 140–149. DOI: 10.1016/j.ympev.2015.12.008.

Gussarova, Galina (2017):
**New Combinations in Nearctic endemic *Euphrasia* (Orobanchaceae).**
In: Journal of the Botanical Research Institute of Texas 11 (2), p. 289–290.

Gussarova, Galina; Popp, Magnus; Vitek, Ernst; Brochmann, Christian (2008):
**Molecular Phylogeny and Biogeography of the bipolar *Euphrasia* (Orobanchaceae): recent Radiations in an old Genus.**
In: Molecular Phylogenetics and Evolution 48 (2), p. 444–460. DOI: 10.1016/j.ympev.2008.05.002.

Liu, Min-lu; Yu, Wen-Bin; Kuss, Patrick; Li, De-Zhu; Wang, Hong (2015):
**Floral nectary Morphology and Evolution in *Pedicularis* (Orobanchaceae).**

In: Botanical Journal of the Linnean Society 178, 592-507.

**Manen, Jean-François; Habashi, Christine; Jeanmonod, Daniel; Park, Jeong-Mi; Schneeweiss, Gerald M. (2004):**
**Phylogeny and intraspecific variability of holoparasitic *Orobanche* (Orobanchaceae) inferred from Plastid rbcL Sequences.**
In: Molecular Phylogenetics and Evolution 33 (2), p. 482–500. DOI: 10.1016/j.ympev.2004.06.010.

**McNeal, Joel R.; Bennett, Jonathan R.; Wolfe, Andrea D.; Mathews, Sarah (2013):**
**Phylogeny and Origins of Holoparasitism in Orobanchaceae.**
In: American Journal of Botany 100 (5), p. 971–983. DOI: 10.3732/ajb.1200448.

**Morawetz, Jeffery J.; Randle, Christopher P.; Wolfe, Andrea D. (2010):**
**Phylogenetic Relationships within the tropical Clade of Orobanchaceae.**
In: Taxon 59 (2), p. 416–426.

**Morawetz, Jeffery J.; Wolfe, Andrea D. (2009):**
**Assessing the Monophyly of *Alectra* and its Relationship to *Melasma* (Orobanchaceae).**
In: Systematic Botany 34 (3), p. 561–569. DOI: 10.1600/036364409789271281.

**Oja, Tatjana; Talve, Tiina (2012):**
**Genetic Diversity and Differentiation in six Species of the Genus *Rhinanthus* (Orobanchaceae).**
In: Plant Systematics and Evolution 298 (5), p. 901–911. DOI: 10.1007/s00606-012-0599-3.

**Park, Jeong-Mi; Manen, Jean-François; Colwell, Alison E. L.; Schneeweiss, Gerald M. (2008):**
**A Plastid Gene Phylogeny of the non-photosynthetic parasitic *Orobanche* (Orobanchaceae) and related Genera.**
In: Journal of Plant Research 121 (4), p. 365–376. DOI: 10.1007/s10265-008-0169-5.

**Park, Jeong-Mi; Manen, Jean-François; Schneeweiss, Gerald M. (2007):**
**Horizontal Gene transfer of a Plastid Gene in the non-photosynthetic Flowering Plants *Orobanche* and *Phelipanche* (Orobanchaceae).**
In: Molecular Phylogenetics and Evolution 43 (3), p. 974–985. DOI: 10.1016/j.ympev.2006.10.011.

**Pettengill, James B.; Neel, Maile C. (2008):**
**Phylogenetic Patterns and conservation among North American members of the Genus *Agalinis* (Orobanchaceae).**
In: BMC Evolutionary Biology 8, p. 264. DOI: 10.1186/1471-2148-8-264.

**Pinto-Carrasco, Daniel; Scheunert, Agnes; Heubl, Günther; Rico, Enrique; Martínez-Ortega, M. Montserrat (2017):**
**Unravelling the Phylogeny of the root-hemiparasitic Genus *Odontites* (Tribe Rhinantheae, Orobanchaceae): Evidence for five main Lineages.**
In: Taxon 66 (4), p. 886–908. DOI: 10.12705/664.6.

**Pradeep, Ayilliath K.; Pramod, Choorakkay (2013):**
***Parasopubia hofmannii* Pradeep & Pramod and *Parasopubia hofmannii* var. *albiflora* Pradeep & Pramod (Orobanchaceae), two new Taxa from India.**
In: Candollea 68 (1), p. 115–122. DOI: 10.15553/c2013v681a16.

**Randle, Christopher P.; Wolfe, Andrea D. (2005):**
**The Evolution and Expression of Rbcl in Holoparasitic Sister-genera *Harveya* and *Hyobanche* (Orobanchaceae)1.**

In: American Journal of Botany 92 (9), p. 1575–1585.

**Ree, Richard H. (2005):**
**Phylogeny and the Evolution of Floral Diversity in *Pedicularis* (Orobanchaceae).**
In: International Journal of Plant Sciences 166 (4), p. 595–613.

**Rodrigues, Anuar G.; Colwell, Alison E. L.; Stefanović, Saša (2011):**
**Molecular Systematics of the parasitic Genus *Conopholis* (Orobanchaceae) inferred from Plastid and nuclear Sequences.**
In: American Journal of Botany 98 (5), p. 896–908. DOI: 10.3732/ajb.1000375.

**Scheunert, Agnes; Fleischmann, Andreas; Olano-Marín, Catalina; Bräuchler, Christian; Heubl, Günther (2012):**
**Phylogeny of Tribe Rhinantheae (Orobanchaceae) with a Focus on Biogeography, Cytology and Re-examination of generic Concepts.**
In: Taxon 61 (6), p. 1269–1285.

**Schneeweiss, Gerald M.; Colwell, Alison E. L.; Park, Jeong-Mi; Jang, Chang-Gee; Stuessy, Tod F. (2004):**
**Phylogeny of holoparasitic *Orobanche* (Orobanchaceae) inferred from nuclear ITS Sequences.**
In: Molecular Phylogenetics and Evolution 30 (2), p. 465–478. DOI: 10.1016/S1055-7903(03)00210-0.

**Tank, David C.; Egger, J. Mark; Olmstead, Richard G. (2009):**
**Phylogenetic Classification of Subtribe Castillejinae (Orobanchaceae).**
In: Systematic Botany 34, p. 182–197.

**Tank, David C.; Olmstead, Richard G. (2008):**
**From Annuals to Perennials: Phylogeny of Subtribe Castillejinae (Orobanchaceae).**
In: American Journal of Botany 95 (5), p. 608–625. DOI: 10.3732/ajb.2007346.

**Tank, David C.; Olmstead, Richard G. (2009):**
**The evolutionary Origin of a second Radiation of Annual *Castilleja* (Orobanchaceae) Species in South America: the role of long Distance Dispersal and Allopolyploidy.**
In: American Journal of Botany 96 (10), p. 1907–1921. DOI: 10.3732/ajb.0800416.

**Tkach, Natalia V.; Ree, Richard H.; Kuss, Patrick; Röser, Martin; Hoffmann, Matthias H. (2014):**
**High mountain Origin, Phylogenetics, Evolution, and Niche Conservatism of arctic Lineages in the hemiparasitic Genus *Pedicularis* (Orobanchaceae).**
In: Molecular Phylogenetics and Evolution 76, p. 75–92. DOI: 10.1016/j.ympev.2014.03.004.

**Uribe-Convers, Simon; Tank, David C. (2016):**
**Phylogenetic Revision of the Genus *Bartsia* (Orobanchaceae): Disjunct Distributions correlate to independent Lineages.**
In: Systematic Botany 41 (3), p. 672–684. DOI: 10.1600/036364416X692299.

**Wang, H.; Mill, Robert R.; Blackmore, p. (2003):**
**Pollen Morphology and infra-generic evolutionary Relationships in some Chinese Species of *Pedicularis* (Scrophulariaceae).**
In: Plant Systematics and Evolution 237 (1-2), p. 1–17. DOI: 10.1007/s00606-002-0188-y.

**Wang, Hong-Juan; Li, Wei-Tao; Liu, Ya-Nan; Yang, Fu-Sheng; Wang, Xiao-Quan (2015):**
**Range-wide multilocus phylogenetic Analyses of *Pedicularis* Sect. *Cyathophora* (Orobanchaceae): Implications for Species Delimitation and Speciation.**
In: Taxon 64 (5), p. 959–974. DOI: 10.12705/645.6.

**Wang, Hong-Juan; Li, Wei-Tao; Liu, Ya-Nan; Yang, Fu-Sheng; Wang, Xiao-Quan (2017):**

**Resolving interspecific Relationships within Evolutionarily young Lineages using RNA-Seq Data: An Example from *Pedicularis* Section *Cyathophora* (Orobanchaceae).**
In: Molecular Phylogenetics and Evolution 107, p. 345–355. DOI: 10.1016/j.ympev.2016.11.018.

WEISS-SCHNEEWEISS, HANNA; GREILHUBER, JOHANN; SCHNEEWEISS, GERALD M. (2006):
**Genome Size Evolution in Holoparasitic *Orobanche* (Orobanchaceae) and related Genera.**
In: American Journal of Botany 93 (1), p. 148–156.

WU, MING-JOU; HUANG, TSENG-CHIENG (2004):
**Taxonomy of the *Euphrasia transmorrisonensis* (Orobanchaceae) Complex in Taiwan based on nrITS.**
In: Taxon 53 (4), p. 911–918.

YANG, F.-S.; WANG, X.-Q. (2007):
**Extensive Length Variation in the cpDNA trnT-trnF Region of hemiparasitic *Pedicularis* and its phylogenetic Implications.**
In: Plant Systematics and Evolution 264 (3-4), p. 251–264. DOI: 10.1007/s00606-006-0510-1.

YU, WEN-BIN; HUANG, PAN-HUI; LI, DE-ZHU; WANG, HONG (2013):
**Incongruence between nuclear and Chloroplast DNA Phylogenies in *Pedicularis* Section *Cyathophora* (Orobanchaceae).**
In: Public Library of Science One 8 (9), e74828. DOI: 10.1371/journal.pone.0074828.

### Osmundaceae

BOMFLEUR, BENJAMIN; GRIMM, GUIDO W.; MCLOUGHLIN, STEPHEN (2015):
***Osmunda pulchella* sp. nov. from the Jurassic of Sweden - reconciling molecular and fossil Evidence in the Phylogeny of modern Royal Ferns (Osmundaceae).**
In: BMC Evolutionary Biology 15, p. 126. DOI: 10.1186/s12862-015-0400-7.

METZGAR, JORDAN S.; SKOG, JUDITH E.; ZIMMER, ELIZABETH A.; PRYER, KATHLEEN M. (2008):
**The Paraphyly of *Osmunda* is confirmed by phylogenetic Analyses of seven Plastid Loci.**
In: Systematic Botany 33 (1), p. 31–36.

### Oxalidaceae

HEIBL, CHRISTOPH; RENNER, SUSANNE P. (2012):
**Distribution models and a dated Phylogeny for Chilean Oxalis Species reveal occupation of new habitats by different Lineages, not rapid adaptive Radiation.**
In: Systematic Biology 61 (5), p. 823–834. DOI: 10.1093/sysbio/sys034.

JOOSTE, MICHELLE; DREYER, LÉANNE L.; OBERLANDER, KENNETH C. (2016):
**The phylogenetic Significance of Leaf anatomical Traits of southern African *Oxalis*.**
In: BMC Evolutionary Biology 16 (1), p. 225. DOI: 10.1186/s12862-016-0792-z.

OBERLANDER, KENNETH C.; DREYER, LÉANNE L.; BELLSTEDT, DIRK U. (2011):
**Molecular Phylogenetics and Origins of southern African *Oxalis*.**
In: Taxon 60 (6), p. 1667–1677. DOI: 10.1002/tax.606011.

OBERLANDER, KENNETH C.; DREYER, LÉANNE L.; BELLSTEDT, DIRK U.; REEVES, GAIL (2004):
**Systematic Relationships in southern African *Oxalis* L. (Oxalidaceae): Congruence between palynological and Plastid trnL-F Evidence.**
In: Taxon 53 (4), p. 977–985. DOI: 10.2307/4135564.

**Oberlander, Kenneth C.; Emshwiller, Eve; Bellstedt, Dirk U.; Dreyer, L. L. (2009):**
**A model of Bulb Evolution in the Eudicot Genus *Oxalis* (Oxalidaceae).**
In: Molecular Phylogenetics and Evolution 51 (1), p. 54–63. DOI: 10.1016/j.ympev.2008.11.022.

**Vaio, Magdalena; Gardner, Andrew; Emshwiller, Eve; Guerra, Marcelo (2013):**
**Molecular Phylogeny and Chromosome Evolution among the creeping herbaceous *Oxalis* Species of Sections *Corniculatae* and Ripariae (Oxalidaceae).**
In: Molecular Phylogenetics and Evolution 68 (2), p. 199–211. DOI: 10.1016/j.ympev.2013.03.019.

**Vaio, Magdalena; Gardner, Andrew; Speranza, Pablo R.; Emshwiller, Eve; Guerra, Marcelo (2016):**
**Phylogenetic and cytogenetic Relationships among Species of *Oxalis* Section *Articulatae* (Oxalidaceae).**
In: Plant Systematics and Evolution 302 (9), p. 1253–1265. DOI: 10.1007/s00606-016-1330-6.

### Paeoniaceae

**Hong, De-Yuan; Zhang, Da-Ming; Wang, Xiao-Quan; Koruklu, Selcuk Tugrul; Tzanoudakis, Dimitris (2008):**
**Relationships and Taxonomy of *Paeonia arietina* G. Anderson complex (Paeoniaceae) and its allies.**
In: Taxon 57 (3), p. 922–932.

**Pan, Jin; Zhang, Da-Ming; Sang, Tao (2007):**
**Molecular phylogenetic Evidence for the Origin of a diploid Hybrid of *Paeonia* (Paeoniaceae).**
In: American Journal of Botany 94 (3), p. 400–408.

**Tank, David C.; Sang, Tao (2001):**
**Phylogenetic utility of the Glycerol-3-Phosphate Acyltransferase Gene: Evolution and Implications in *Paeonia* (Paeoniaceae).**
In: Molecular Phylogenetics and Evolution 19 (3), p. 421–429. DOI: 10.1006/mpev.2001.0931.

**Zhang, Jin-Mei; López-Pujol, Jordi; Gong, Xun; Wang, Hua-Feng; Vilatersana, Roser; Zhou, Shi-Liang (2018):**
**Population genetic dynamics of Himalayan-Hengduan Tree Paeonies, *Paeonia* Subsect. *Delavayanae*.**
In: Molecular Phylogenetics and Evolution 125, p. 62–77. DOI: 10.1016/j.ympev.2018.03.003.

**Zhao, Xuan; Zhou, Zhi-Qin; Qi-Bing; Pan, Lin Kai-Yu; Li, Ming-Yang (2008):**
**Phylogenetic Analysis of *Paeonia* Sect. *Moutan* (Paeoniaceae) based on multiple DNA fragments and morphological Data.**
In: Journal of Systematics and Evolution 46 (4), p. 563–572.

### Pandanaceae

**Buerki, Sven; Callmander, Martin W.; Devey, Dion S.; Chappell, Lauren; Gallaher, Timothy; Munzinger, Jérôme et al. (2012):**
**Straightening out the screw-pines: A first step in Understanding phylogenetic Relationships within Pandanaceae.**
In: Taxon 61 (5), p. 1010–1020.

**Callmander, Martin W.; Booth, Thomas J.; Beentje, Henk; Buerki, Sven (2013):**
**Update on the Systematics of *Benstonea* (Pandanaceae): When a visionary Taxonomist foresees phylogenetic Relationships.**
In: Phytotaxa 112 (2), p. 57. DOI: 10.11646/phytotaxa.112.2.4.

**Callmander, Martin W.; Buerki, Sven (2016):**

**Two new threatened Species of *Benstonea* Callm. & Buerki (Pandanaceae) from Sabah (Borneo, Malaysia).**

In: Candollea 71 (2), p. 257–263. DOI: 10.15553/c2016v712a10.

CALLMANDER, MARTIN W.; BUERKI, SVEN; KEIM, ARY P.R.Y.; PHILLIPSON, PETER B. (2014):

**Notes on *Benstonea* (Pandanaceae) from the Islands of Halmahera, New Guinea and Sulawesi.**

In: Phytotaxa 175 (3), p. 161. DOI: 10.11646/phytotaxa.175.3.6.

CALLMANDER, MARTIN W.; LOWRY, PORTER PRESCOTT; FOREST, FÉLIX; DEVEY, DION S.; BEENTJE, HENK; BUERKI, SVEN (2012):

***Benstonea* Callm. & Buerki (Pandanaceae): Characterization, Circumscription, and Distribution of a new Genus of Screw-Pines, with a Synopsis of accepted Species.**

In: Candollea 67 (2), p. 323–345. DOI: 10.15553/c2012v672a12.

GALLAHER, TIMOTHY; CALLMANDER, MARTIN W.; BUERKI, SVEN; KEELEY, STERLING C. (2015):

**A long distance Dispersal Hypothesis for the Pandanaceae and the Origins of the *Pandanus tectorius* Complex.**

In: Molecular Phylogenetics and Evolution 83, p. 20–32. DOI: 10.1016/j.ympev.2014.11.002.

### Pandanales

CADDICK, LIZABETH R.; RUDALL, PAULA J.; WILKIN, PAUL; HEDDERSON, TERRY A.; CHASE, MARK W. (2002):

**Phylogenetics of Dioscoreales based on combined Analyses of morphological and molecular Data.**

In: Botanical Journal of the Linnean Society 138 (2), p. 123–144. DOI: 10.1046/j.1095-8339.2002.138002123.x.

CADDICK, LIZABETH R.; WILKIN, PAUL; RUDALL, PAULA J.; HEDDERSON, TERRY A.; CHASE, MARK W. (2002):

**Yams reclassified: a Recircumscription of Dioscoreaceae and Dioscoreales.**

In: Taxon 51 (1), p. 103–114. DOI: 10.2307/1554967.

### Papaveraceae

CAROLAN, JAMES C.; HOOK, INGRID L. I.; CHASE, MARK W.; KADEREIT, JOACHIM W.; HODKINSON, TREVOR R. (2006):

**Phylogenetics of Papaver and related Genera based on DNA Sequences from ITS nuclear ribosomal DNA and Plastid trnL Intron and trnL-F intergenic Spacers.**

In: Annals of Botany 98 (1), p. 141–155. DOI: 10.1093/aob/mcl079.

GÜRKÖK, TUGBA; KAYMAK, ELIF; BOZTEPE, GÜLSEN; KOYUNCU, MESUT; PARMAKSIZ, ISKENDER (2013):

**Molecular characterization of the Genus *Papaver* Gülsen Section *Oxytona* using ISSR Markers.**

In: Turkish Journal of Botany 37, p. 644–650. DOI: 10.3906/bot-1208-16.

LI, JIAN-HUA; MURRAY, K. GREGORY; LI, PAN; BROWN, KENNETH (2018):

**Differential Diversifications of South American and Eastern Asian disjunct Genera *Bocconia* and *Macleaya* (Papaveraceae).**

In: Journal of Systematics and Evolution 56 (1), p. 25–34. DOI: 10.1111/jse.12286.

ZHANG, ZHONG-XIN; WANG, DONG; YANG, XUE (2016):

**The taxonomic Position of *Corydalis parviflora* Su & Lidén (Papaveraceae), a genetically distinct species: Evidence from cpDNA and nDNA Sequences.**

In: Biochemical Systematics and Ecology 67, p. 134–141. DOI: 10.1016/j.bse.2016.06.003.

### Passifloraceae

HEARN, DAVID J. (2006):

**_Adenia_ (Passifloraceae) and its Adaptive Radiation: Phylogeny and Growth Form Diversification.**
In: Systematic Botany 31 (4), p. 805–821.

**KROSNICK, SHAWN E.; FORD, ANDREW J.; FREUDENSTEIN, JOHN V. (2009):**
**Taxonomic Revision of _Passiflora_ Subgenus _Tetrapathea_ Including the Monotypic Genera _Hollrungia_ and _Tetrapathea_ (Passifloraceae), and a new Species of _Passiflora_.**
In: Systematic Botany 34 (2), p. 375–385. DOI: 10.1600/036364409788606343.

**MILWARD-DE-AZEVEDO, MICHAELE ALVIM; BAUMGRATZ, JOSÉ FERNANDO A.; GONÇALVES-ESTEVES, VÂNIA (2012):**
**A taxonomic Revision of _Passiflora_ Subgenus _Decaloba_ (Passifloraceae) in Brazil.**
In: Phytotaxa 53, p. 1–68.

**MUSCHNER, VALÉRIA C.; LORENZ, ALINE P.; CERVI, ARMANDO C.; BONATTO, SANDRO LUIS; SOUZA-CHIES, TATIANA TEIXEIRA DE; SALZANO, FRANCISCO M.; FREITAS, LORETA BRANDÃO (2003):**
**A first molecular phylogenetic Analysis of _Passiflora_ (Passifloraceae).**
In: American Journal of Botany 90 (8), p. 1229–1238.

**PECCOUD, JEAN; PIATSCHECK, FINN; YOCKTENG, ROXANA; GARCIA, MARJORIE; SAUVE, MATHIEU; DJIÉTO-LORDON, CHAMPLAIN ET AL. (2013):**
**Multi-locus Phylogenies of the Genus _Barteria_ (Passifloraceae) portray complex Patterns in the Evolution of Myrmecophytism.**
In: Molecular Phylogenetics and Evolution 66 (3), p. 824–832. DOI: 10.1016/j.ympev.2012.11.006.

**PIGNAL, MARC; YOCKTENG, ROXANA; HEARN, DAVID J.; LABAT, JEAN-NOEL (2013):**
**_Adenia barthelatii_ (Passifloraceae), a new endemic Species of Mayotte and its phylogenetic Status within the Genus _Adenia_.**
In: Phytotaxa 99 (1), p. 40. DOI: 10.11646/phytotaxa.99.1.3.

**TOKUOKA, TORU (2012):**
**Molecular phylogenetic Analysis of Passifloraceae sensu lato (Malpighiales) based on Plastid and nuclear DNA Sequences.**
In: Journal of Plant Research 125 (4), p. 489–497. DOI: 10.1007/s10265-011-0472-4.

**YOCKTENG, ROXANA; NADOT, SOPHIE (2004):**
**Infrageneric Phylogenies: a Comparison of chloroplast-expressed glutamine Synthetase, cytosol-expressed Glutamine Synthetase and cpDNA Maturase K in _Passiflora_.**
In: Molecular Phylogenetics and Evolution 31 (1), p. 397–402. DOI: 10.1016/S1055-7903(03)00276-8.

**YOCKTENG, ROXANA; NADOT, SOPHIE (2004):**
**Phylogenetic Relationships among _Passiflora_ Species based on the Glutamine Synthetase nuclear Gene expressed in Chloroplast (ncpGS).**
In: Molecular Phylogenetics and Evolution 31 (1), p. 379–396. DOI: 10.1016/S1055-7903(03)00277-X.

### Paulowniaceae

**ZHOU, QING-MEI; JENSEN, SØREN ROSENDAL; LIU, GUO-LI; WANG, SHUANG; LI, HONG-QING (2014):**
**Familial Placement of _Wightia_ (Lamiales).**
In: Plant Systematics and Evolution 300 (9), p. 2009–2017. DOI: 10.1007/s00606-014-1029-5.

### Pedaliaceae

**GORMLEY, INGEBORG C.; BEDIGIAN, DOROTHEA; OLMSTEAD, RICHARD G. (2015):**

**Phylogeny of Pedaliaceae and Martyniaceae and the Placement of *Trapella* in Plantaginaceae s. l.**
In: Systematic Botany 40 (1), p. 259–268. DOI: 10.1600/036364415X686558.

### Penaeaceae

ONSTEIN, RENSKE E.; CARTER, RICHARD J.; XING, YAOWU; LINDER, HANS PETER (2014):
**Diversification Rate Shifts in the Cape Floristic Region: the right Traits in the right Place at the right Time.**
In: Perspectives in Plant Ecology, Evolution and Systematics 16 (6), p. 331–340. DOI: 10.1016/j.ppees.2014.08.002.

SCHÖNENBERGER, JÜRGEN; CONTI, ELENA (2003):
**Molecular Phylogeny and Floral Evolution of Penaeaceae, Oliniaceae, Rhynchocalycaceae, and Alzateaceae (Myrtales).**
In: American Journal of Botany 90 (2), p. 293–309. DOI: 10.3732/ajb.90.2.293.

### Peridiscaceae

SOLTIS, DOUGLAS E.; CLAYTON, JOSHUA W.; DAVIS, CHARLES C.; GITZENDANNER, MATTHEW A.; CHEEK, MARTIN R.; SAVOLAINEN, VINCENT ET AL. (2007):
**Monophyly and Relationships of the enigmatic Family Peridiscaceae.**
In: Taxon 56 (1), p. 65–73.

### Petenaeaceae

CHRISTENHUSZ, MAARTEN J.M.; FAY, MICHAEL F.; CLARKSON, JAMES J.; GASSON, PETER E.; MORALES CAN, JULIO; JIMÉNEZ BARRIOS, JORGE B.; CHASE, MARK W. (2010):
**Petenaeaceae, a new Angiosperm Family in Huerteales with a distant Relationship to *Gerrardina* (Gerrardinaceae).**
In: Botanical Journal of the Linnean Society 164 (1), p. 16–25. DOI: 10.1111/j.1095-8339.2010.01074.x.

### Philydraceae

SAARELA, JEFFERY M.; PRENTIS, PETER J.; RAI, HARDEEP S.; GRAHAM, SEAN W. (2008):
**Phylogenetic Relationships in the Monocot Order Commelinales, with a focus on Philydraceae.**
In: Botany 86 (7), p. 719–731. DOI: 10.1139/B08-063.

### Phrymaceae

BARKER, WILLIAM R.; NESOM, GUY L.; BEARDSLEY, PAUL M.; FRAGA., N. P. (2012):
**A taxonomic Conspectus of Phrymaceae: A narrowed Circumscriptions for *Mimulus*, new and resurrected Genera, and new Names and Combinations.**
In: Phytoneuron 39, p. 1–60.

BEARDSLEY, PAUL M.; BAKER, WILLIAM R. (2005):
**Patterns of Evolution in Australian *Mimulus* and related Genera (Phrymaceae~Scrophulariaceae): a molecular Phylogeny using Chloroplast and nuclear Sequence Data.**
In: Australian Systematic Botany 18, p. 61–73.

BEARDSLEY, PAUL M.; SCHOENIG, STEVE E.; WHITTALL, JUSTEN B.; OLMSTEAD, RICHARD G. (2004):
**Patterns of Evolution in Western North American *Mimulus* (Phrymaceae).**
In: American Journal of Botany 91 (3), p. 474–489.

LES, DONALD H.; CAPERS, ROBERT P. (2006):

**Introduction of *Glossostigma* (Phrymaceae) to North America: A taxonomic and Ecological Overview.**
In: American Journal of Botany 93 (6), p. 927–939.

NESOM, GUY L. (2014):
**Updated Classification and hypothetical Phylogeny of *Erythranthe* Sect. *Simiola* (Phrymaceae).**
In: Phytoneuron 81, p. 1–6.

NIE, ZE-LONG; SUN, HANG; BEARDSLEY, PAUL M.; OLMSTEAD, RICHARD G. (2006):
**Evolution of Biogeographic Disjunction between Eastern Asia and Eastern North America in *Phryma* (Phrymaceae).**
In: American Journal of Botany 93 (9), p. 1343–1356.


### Phyllanthaceae

CHALLEN, GILL; VORONTSOVA, MARIA S.; SCHNEIDER, HARALD; CHEEK, MARTIN R. (2011):
**Phylogenetically distinct and critically Endangered new Tree Species of *Phyllanthus* from Cameroon (Phyllanthaceae, Euphorbiaceae s. l.).**
In: Systematic Botany 36 (4), p. 933–938. DOI: 10.1600/036364411X604949.

KATHRIARACHCHI, HASHENDRA; HOFFMANN, PETRA; SAMUEL, ROSABELLE; WURDACK, KENNETH J.J.; CHASE, MARK W. (2005):
**Molecular Phylogenetics of Phyllanthaceae inferred from five Genes (plastid atpB, matK, 3'ndhF, rbcL, and nuclear Phyc).**
In: Molecular Phylogenetics and Evolution 36 (1), p. 112–134. DOI: 10.1016/j.ympev.2004.12.002.

KATHRIARACHCHI, HASHENDRA; SAMUEL, ROSABELLE; HOFFMANN, PETRA; MLINAREC, JELENA; WURDACK, KENNETH J.J.; RALIMANANA, HÉLÈNE ET AL. (2006):
**Phylogenetics of Tribe Phyllantheae (Phyllanthaceae; Euphorbiaceae sensu lato) based on nrITS and Plastid matK DNA Sequence Data.**
In: American Journal of Botany 93 (4), p. 637–655. DOI: 10.3732/ajb.93.4.637.

MANISSORN, JUTHATIP; SUKRONG, SUCHADA; RUANGRUNGSI, NIJSIRI; MIZUKAMI, HAJIME (2010):
**Molecular phylogenetic Analysis of *Phyllanthus* Species in Thailand and the Application of Polymerase Chain Reaction-Restriction Fragment Length Polymorphism for *Phyllanthus amarus* Identification.**
In: Biological and Pharmaceutical Bulletin 33 (10), p. 1723–1727. DOI: 10.1248/bpb.33.1723.

PRUESAPAN, KANCHANA; TELFORD, IAN R.H.; BRUHL, JEREMY J.; DRAISMA, STEFANO G. A.; VAN WELZEN, PETER C. (2008):
**Delimitation of *Sauropus* (Phyllanthaceae) based on Plastid matK and nuclear ribosomal ITS DNA Sequence Data.**
In: Annals of Botany 102 (6), p. 1007–1018. DOI: 10.1093/aob/mcn193.

PRUESAPAN, KANCHANA; TELFORD, IAN R.H.; BRUHL, JEREMY J.; VAN WELZEN, PETER C. (2012):
**Phylogeny and proposed Circumscription of *Breynia, Sauropus* and *Synostemon* (Phyllanthaceae), based on Chloroplast and nuclear DNA Sequences.**
In: Australian Systematic Botany 25 (5), p. 313–330. DOI: 10.1071/SB11005.

SAMUEL, ROSABELLE; KATHRIARACHCHI, HASHENDRA; HOFFMANN, PETRA; BARFUSS, MICHAEL H. J.; WURDACK, KENNETH J.J.; DAVIS, CHARLES C.; CHASE, MARK W. (2005):
**Molecular Phylogenetics of Phyllanthaceae: Evidence from Plastid matK and Nuclear Phyc Sequences.**
In: American Journal of Botany 92 (1), p. 132–141.

**Telford, Ian R.H.; Pruesapan, Kanchana; van Welzen, Peter C.; Bruhl, Jeremy J. (2014):**
**Molecular Data consistently recover a 'Queensland Clade' of *Synostemon* Phyllanthaceae, Phyllantheae) with distinctive Floral Morphology.**
In: Australian Systematic Botany 27 (6), p. 450–461. DOI: 10.1071/SB14034.

**Telford, Ian R.H.; Pruesapan, Kanchana; van Welzen, Peter C.; Bruhl, Jeremy J. (2016):**
**Morphological and molecular Data show *Synostemon trachyspermus* (Phyllanthaceae, Phyllantheae) to be a heterogeneous Species Assemblage.**
In: Australian Systematic Botany 29 (3), p. 218–234. DOI: 10.1071/SB16008.

**van Welzen, Peter C.; Pruesapan, Kanchana; Telford, Ian R.H.; Esser, Hans-Joachim; Bruhl, J. J. (2014):**
**Phylogenetic Reconstruction prompts taxonomic Changes in *Sauropus, Synostemon* and *Breynia* (Phyllanthaceae Tribe Phyllantheae).**
In: Blumea 59 (2), p. 77–94. DOI: 10.3767/000651914X684484.

**Vorontsova, Maria S.; Hoffmann, Petra (2008):**
**A phylogenetic Classification of Tribe Poranthereae (Phyllanthaceae, Euphorbiaceae sensu lato).**
In: Kew Bulletin 63 (1), p. 41–59. DOI: 10.1007/s12225-007-9012-8.

**Vorontsova, Maria S.; Hoffmann, Petra; Maurin, Olivier; Chase, Mark W. (2007):**
**Molecular Phylogenetics of Tribe Poranthereae (Phyllanthaceae; Euphorbiaceae sensu lato).**
In: American Journal of Botany 94 (12), p. 2026–2040. DOI: 10.3732/ajb.94.12.2026.

**Webster, Grady L.; Carpenter, Kevin J. (2008):**
**Pollen Morphology and Systematics of palaeotropical *Phyllanthus* and related Genera of Subtribe Phyllanthinae (Euphorbiaceae).**
In: Botanical Journal of the Linnean Society 157 (4), p. 591–608. DOI: 10.1111/j.1095-8339.2008.00781.x.

**Wurdack, Kenneth J.J.; Hoffmann, Petra; Samuel, Rosabelle; Bruijn, Anette D.E.; van der Bank, Michelle; Chase, Mark W. (2004):**
**Molecular phylogenetic Analysis of Phyllanthaceae (Phyllanthoideae pro parte, Euphorbiaceae sensu lato) using Plastid Rbcl DNA Sequences.**
In: American Journal of Botany 91 (11), p. 1882–1900.

**Yao, Gang; Zhang, Dian-Xiang (2016):**
**Pollen Morphology of Chinese *Glochidion* (Phyllanthaceae) and its taxonomic Implications.**
In: Nordic Journal of Botany 34 (1), p. 102–110. DOI: 10.1111/njb.00912.

### Phytolaccaceae

**Ali, M. Ajmal; Lee, Joongku; Kim, Soo-Young; Park, Sang-Hong; Al-Hemaid, Fahad M.A. (2015):**
**Molecular phylogenetic Analyses if Internal Transcribed Spacer (ITS) Sequences of nuclear ribosomal DNA indicate Monophyly of the Genus Phytolacca L. (Phytolaccaceae).**
In: Bangladesh Journal of Plant Taxononmy 22 (1), p. 1–8.

**Lee, J.; Kim, p. Y.; Park, Sang-Hong; Ali, M. Ajmal (2013):**
**Molecular phylogenetic Relationships among Members of the Family Phytolaccaceae sensu lato inferred from Internal Transcribed Spacer Sequences of nuclear ribosomal DNA.**
In: Genetics and molecular Research 12 (4), p. 4515–4525. DOI: 10.4238/2013.February.28.15.

**Meirelles, Julia (2016):**

**Flora das cangas da Serra dos Carajás, Pará, Brasil: Phytolaccaceae.**

In: Rodriguésia 67 (5spe), p. 1443–1445. DOI: 10.1590/2175-7860201667545.

**Schäferhoff, Bastian; Müller, Kai F.; Borsch, Thomas (2009):**

**Caryophyllales Phylogenetics: disentangling Phytolaccaceae and Molluginaceae and Description of Microteaceae as a new isolated Family.**

In: Willdenowia 39 (2), p. 209–228. DOI: 10.3372/wi.39.39201.


### Picramniales

**Logacheva, Maria D.; Shipunov, Alexey B. (2017):**

**Phylogenomic Analysis of *Picramnia, Alvaradoa,* and *Leitneria* supports the independent Picramniales.**

In: Journal of Systematics and Evolution 55 (3), p. 171–176. DOI: 10.1111/jse.12246.


### Picrodendraceae

**Sutter, D. Merino; Forster, Paul I.; Endress, Peter K. (2006):**

**Female flowers and systematic Position of Picrodendraceae (Euphorbiaceae s.l., Malpighiales).**

In: Plant Systematics and Evolution 261 (1-4), p. 187–215. DOI: 10.1007/s00606-006-0414-0.


### Pinaceae

**Aguirre-Planter, Erika; Jaramillo-Correa, Juan P.; Gómez-Acevedo, Sandra; Khasa, Damase P.; Bousquet, Jean; Eguiarte, Luis E. (2012):**

**Phylogeny, Diversification rates and Species boundaries of Mesoamerican Firs (*Abies*, Pinaceae) in a genus-wide Context.**

In: Molecular Phylogenetics and Evolution 62 (1), p. 263–274. DOI: 10.1016/j.ympev.2011.09.021.

**Campbell, Christopher S.; Wright, Wesley A.; Cox, Margaret; Vining, Thomas F.; Major, C. Smoot; Arsenault, Matthew P. (2005):**

**Nuclear ribosomal DNA Internal Transcribed Spacer 1 (ITS1) in *Picea* (Pinaceae): Sequence Divergence and Structure.**

In: Molecular Phylogenetics and Evolution 35 (1), p. 165–185. DOI: 10.1016/j.ympev.2004.11.010.

**Eckert, Andrew J.; Hall, Benjamin D. (2006):**

**Phylogeny, historical Biogeography, and Patterns of Diversification for *Pinus* (Pinaceae): phylogenetic Tests of fossil-based Hypotheses.**

In: Molecular Phylogenetics and Evolution 40 (1), p. 166–182. DOI: 10.1016/j.ympev.2006.03.009.

**Flores-Rentería, Lluvia; Wegier, Ana; Ortega Del Vecchyo, Diego; Ortíz-Medrano, Alejandra; Piñero, Daniel; Whipple, Amy V. et al. (2013):**

**Genetic, morphological, geographical and ecological approaches reveal phylogenetic Relationships in complex Groups, an Example of recently diverged Pinyon Pine Species (Subsection *Cembroides*).**

In: Molecular Phylogenetics and Evolution 69 (3), p. 940–949. DOI: 10.1016/j.ympev.2013.06.010.

**Gernandt, David S.; Hernández-León, Sergio; Salgado-Hernández, Esmeralda; La Pérez de Rosa, Jorge A. (2009):**

**Phylogenetic Relationships of *Pinus* Subsection *Ponderosae* inferred from Rapidly Evolving cpDNA Regions.**

In: Systematic Botany 34 (3), p. 481–491. DOI: 10.1600/036364409789271290.

**Gernandt, David S.; Liston, Aaron; Piñero, Daniel (2001):**

**Variation in the nrDNA ITS of *Pinus* subsection *Cembroides*: Implications for molecular systematic studies of pine Species Complexes.**
In: Molecular Phylogenetics and Evolution 21 (3), p. 449–467. DOI: 10.1006/mpev.2001.1026.

Gernandt, David S.; López, Gretel Geada; García, Sol Ortiz; Liston, Aaron (2005):
**Phylogeny and Classification of *Pinus*.**
In: Taxon 54 (1), p. 29–42. DOI: 10.2307/25065300.

Grivet, Delphine; Climent, José; Zabal-Aguirre, Mario; Neale, David B.; Vendramin, Giovanni G.; González-Martínez, Santiago C. (2013):
**Adaptive Evolution of Mediterranean Pines.**
In: Molecular Phylogenetics and Evolution 68 (3), p. 555–566. DOI: 10.1016/j.ympev.2013.03.032.

Gugerli, Felix; Sperisen, C.; Büchler, U.; Brunner, I.; Brodbeck, Sabine; Palmer, Jeffrey D.; Qiu, Y.-L. (2001):
**The evolutionary split of Pinaceae from other conifers: Evidence from an Intron Loss and a multigene Phylogeny.**
In: Molecular Phylogenetics and Evolution 21 (2), p. 167–175. DOI: 10.1006/mpev.2001.1004.

Hao, Zhen-Zhen; Liu, Yan-Yan; Nazaire, Mare; Wei, Xiao-Xin; Wang, Xiao-Quan (2015):
**Molecular Phylogenetics and evolutionary History of Sect. *Quinquefoliae* (Pinus): Implications for Northern Hemisphere Biogeography.**
In: Molecular Phylogenetics and Evolution 87, p. 65–79. DOI: 10.1016/j.ympev.2015.03.013.

Havill, Nathan P.; Campbell, Christopher S.; Vining, Thomas F.; LePage, Ben; Bayer, Randall J.; Donoghue, Michael J. (2008):
**Phylogeny and Biogeography of *Tsuga* (Pinaceae) inferred from Nuclear Ribosomal ITS and Chloroplast DNA Sequence Data.**
In: Systematic Botany 33 (3), p. 478–489.

Kan, Xian-Zhao; Wang, Shan-Shan; Ding, Xin; Wang, Xiao-Quan (2007):
**Structural Evolution of nrDNA ITS in Pinaceae and its phylogenetic Implications.**
In: Molecular Phylogenetics and Evolution 44 (2), p. 765–777. DOI: 10.1016/j.ympev.2007.05.004.

Kaundun, Shiv Shankhar; Lebreton, Philippe (2010):
**Taxonomy and Systematics of the Genus *Pinus* based on morphological, biogeographical and biochemical Characters.**
In: Plant Systematics and Evolution 284 (1-2), p. 1–15. DOI: 10.1007/s00606-009-0228-y.

Lockwood, Jared D.; Aleksić, Jelena M.; Zou, Jiabin; Wang, Jing; Liu, Jian-Quan; Renner, Susanne p. (2013):
**A new Phylogeny for the Genus *Picea* from plastid, mitochondrial, and nuclear Sequences.**
In: Molecular Phylogenetics and Evolution 69 (3), p. 717–727. DOI: 10.1016/j.ympev.2013.07.004.

Parks, Matthew; Cronn, Richard C.; Liston, Aaron (2012):
**Separating the Wheat from the Chaff: Mitigating the Effects of Noise in a Plastome phylogenomic Data Set from *Pinus* L. (Pinaceae).**
In: BMC Evolutionary Biology 12, p. 100. DOI: 10.1186/1471-2148-12-100.

Potenko, V. V. (2007):
**Relationships among Spruces (*Picea* A.Dietr., Pinaceae) of the Russian Far East.**
In: Plant Systematics and Evolution 268 (1-4), p. 1–13. DOI: 10.1007/s00606-007-0551-0.

Qiao, Cai-Yuan; Ran, Jin-Hua; Li, Yan; Wang, Xiao-Quan (2007):
**Phylogeny and Biogeography of *Cedrus* (Pinaceae) inferred from Sequences of seven paternal Chloroplast and maternal mitochondrial DNA Regions.**

In: Annals of Botany 100 (3), p. 573–580. DOI: 10.1093/aob/mcm134.

**Ran, Jin-Hua; Wei, Xiao-Xin; Wang, Xiao-Quan (2006):**
**Molecular Phylogeny and Biogeography of *Picea* (Pinaceae): Implications for phylogeographical studies using cytoplasmic Haplotypes.**
In: Molecular Phylogenetics and Evolution 41 (2), p. 405–419. DOI: 10.1016/j.ympev.2006.05.039.

**Semerikov, Vladimir L.; Zhang, Hengqing; Sun, Mei; Lascoux, Martin (2003):**
**Conflicting Phylogenies of *Larix* (Pinaceae) based on cytoplasmic and nuclear DNA.**
In: Molecular Phylogenetics and Evolution 27 (2), p. 173–184. DOI: 10.1016/S1055-7903(02)00447-5.

**Semerikova, p. A.; Semerikov, Vladimir L. (2014):**
**Molecular phylogenetic Analysis of the Genus *Abies* (Pinaceae) based on the nucleotide Sequence of Chloroplast DNA.**
In: Russian Journal of Genetics 50 (1), p. 7–19. DOI: 10.1134/S1022795414010104.

**Suyama, Y.; Yoshimaru, Hiroshi; Tsumura, Yoshihiko (2000):**
**Molecular phylogenetic Position of Japanese *Abies* (Pinaceae) based on Chloroplast DNA Sequences.**
In: Molecular Phylogenetics and Evolution 16 (2), p. 271–277. DOI: 10.1006/mpev.2000.0795.

**Syring, John; Willyard, Ann; Cronn, Richard C.; Liston, Aaron (2005):**
**Evolutionary Relationships among *Pinus* (Pinaceae) Subsections inferred from Multiple Low-copy Nuclear Loci.**
In: American Journal of Botany 92 (12), p. 2086–2100.

**Wang, Baosheng; Wang, Xiao-Ru (2014):**
**Mitochondrial DNA capture and Divergence in *Pinus* provide new Insights into the Evolution of the Genus.**
In: Molecular Phylogenetics and Evolution 80, p. 20–30. DOI: 10.1016/j.ympev.2014.07.014.

**Wang, Xiao-Quan; Han, Ying; Hong, De-Yuan (1998):**
**A molecular Systematic Study of *Cathaya*, a relic Genus of the Pinaceae in China.**
In: Plant Systematics and Evolution 213, p. 165–172.

**Wang, Xiao-Ru; Tsumura, Yoshihiko; Yoshimaru, Hiroshi; Nagasaka, Kazutoshi; Schmidt, Alfred E. (1999):**
**Phylogenetic Relationships of Eurasian Pines (*Pinus*, Pinaceae) based on Chloroplast Rbcl, Matk, Rpl20-rps18 Spacer, and Trnv Intron Sequences.**
In: American Journal of Botany 86 (12), p. 1742–1753.

**Wei, X.-X.; Wang, X.-Q. (2003):**
**Phylogenetic split of *Larix* : Evidence from paternally inherited cpDNA trnT-trnF Region.**
In: Plant Systematics and Evolution 239 (1-2), p. 67–77. DOI: 10.1007/s00606-002-0264-3.

**Wei, Xiao-Xin; Wang, Xiao-Quan (2004):**
**Evolution of 4-coumarate: Coenzyme A Ligase (4cl) Gene and Divergence of *Larix* (Pinaceae).**
In: Molecular Phylogenetics and Evolution 31 (2), p. 542–553. DOI: 10.1016/j.ympev.2003.08.015.

**Wei, Xiao-Xin; Yang, Zu-Yu; Li, Yan; Wang, Xiao-Quan (2010):**
**Molecular Phylogeny and Biogeography of *Pseudotsuga* (Pinaceae): Insights into the floristic Relationship between Taiwan and its adjacent Areas.**
In: Molecular Phylogenetics and Evolution 55 (3), p. 776–785. DOI: 10.1016/j.ympev.2010.03.007.

**Xiang, Qiao-Ping; Wei, Ran; Shao, Yi-Zhen; Yang, Zu-Yu; Wang, Xiao-Quan; Zhang, Xian-Chun (2015):**

**Phylogenetic Relationships, possible ancient Hybridization, and biogeographic History of *Abies* (Pinaceae) based on Data from nuclear, plastid, and mitochondrial Genomes.**
In: Molecular Phylogenetics and Evolution 82 Pt A, p. 1–14. DOI: 10.1016/j.ympev.2014.10.008.

XIANG, QIAO-PING; XIANG, QIU-YUN; GUO, YAN-YAN; ZHANG, XIAN-CHUN (2009):
**Phylogeny of *Abies* (Pinaceae) inferred from nrITS Sequence Data.**
In: Taxon 58 (1), p. 141–152.

XIANG, QIAO-PING; XIANG, QIU-YUN; LISTON, AARON; ZHANG, XIAN-CHUN (2004):
**Phylogenetic Relationships in *Abies* (Pinaceae): Evidence from Pcr-rflp of the nuclear ribosomal DNA Internal Transcribed Spacer Region.**
In: Botanical Journal of the Linnean Society 145, p. 425–435.

XIE, QING; LIU, ZHI-HONG; WANG, SHU-HUI; LI, ZHOU-QI (2015):
**Genetic Diversity and phylogenetic Relationships among five endemic *Pinus* Taxa (Pinaceae) of China as revealed by Srap Markers.**
In: Biochemical Systematics and Ecology 62, p. 115–120. DOI: 10.1016/j.bse.2015.08.005.

ZHANG, HONG-XIANG; ZHANG, MING-LI; WILLIAMS, DAVID M. (2014):
**Genetic Evidence and Species Distribution modelling reveal the response of *Larix sibirica* and its related Species to Quaternary climatic and ancient historical Events.**
In: Biochemical Systematics and Ecology 54, p. 316–325. DOI: 10.1016/j.bse.2014.02.017.

### Piperaceae

HORNER, HARRY T.; WANKE, STEFAN; SAMAIN, MARIE-STÉPHANIE (2009):
**Evolution and Systematic Value of Leaf Crystal Macropatterns in the Genus *Peperomia* (Piperaceae).**
In: International Journal of Plant Sciences 170 (3), p. 343–354. DOI: 10.1086/596338.

JARAMILLO, M. ALEJANDRA; CALLEJAS, RICARDO; DAVIDSON, CHRISTOPHER; SMITH, JAMES F.; STEVENS, ANGELA C.; TEPE, ERIC J. (2008):
**A Phylogeny of the Tropical Genus Piper using ITS and the Chloroplast Intron psbJ–petA.**
In: Systematic Botany 33 (4), p. 647–660. DOI: 10.1600/036364408786500244.

JARAMILLO, M. ALEJANDRA; MANOS, PAUL P. (2001):
**Phylogeny and Patterns of Floral Diversity in the Genus *Piper* (Piperaceae).**
In: American Journal of Botany 88 (4), p. 706–716. DOI: 10.2307/2657072.

MAI, PATRICIA; ROSSADO, ANDRÉS; BONIFACINO, JOSÉ MAURICIO; WAECHTER, JORGE LUIZ (2016):
**Taxonomic Revision of *Peperomia* (Piperaceae) from Uruguay.**
In: Phytotaxa 244 (2), p. 125. DOI: 10.11646/phytotaxa.244.2.2.

MOLINA-HENAO, Y. FRANCHESCO; GUERRERO-CHACÓN, ADRIANA L.; JARAMILLO, M. ALEJANDRA (2016):
**Ecological and Geographic Dimensions of Diversification in *Piper* Subgenus *Ottonia*: A Lineage of Neotropical Rainforest Shrubs.**
In: Systematic Botany 41 (2), p. 253–262. DOI: 10.1600/036364416X691777.

NAUMANN, JULIA; SYMMANK, LARS; SAMAIN, MARIE-STÉPHANIE; MÜLLER, KAI F.; NEINHUIS, CHRISTOPH; DEPAMPHILIS, CLAUDE W.; WANKE, STEFAN (2011):
**Chasing the Hare - Evaluating the phylogenetic Utility of a nuclear Single copy Gene Region at and below Species level within the Species rich Group *Peperomia* (Piperaceae).**

In: BMC Evolutionary Biology 11, p. 357. DOI: 10.1186/1471-2148-11-357.

SAMAIN, MARIE-STÉPHANIE; MATHIEU, GUIDO; WANKE, STEFAN; NEINHUIS, CHRISTOPH; GOETGHEBEUR, PAUL (2008):
**_Verhuellia_ revisited - unravelling its intricate taxonomic History and a new subfamilial Classification of Piperaceae.**
In: Taxon 57 (2), p. 583–587.

SAMAIN, MARIE-STÉPHANIE; VRIJDAGHS, ALEXANDER; HESSE, MICHAEL; GOETGHEBEUR, PAUL; JIMÉNEZ RODRÍGUEZ, FRANCISCO; STOLL, ALEXANDRA ET AL. (2010):
**_Verhuellia_ is a segregate Lineage in Piperaceae: more Evidence from Flower, Fruit and Pollen Morphology, Anatomy and Development.**
In: Annals of Botany 105 (5), p. 677–688. DOI: 10.1093/aob/mcq031.

WANKE, STEFAN; SAMAIN, MARIE-STÉPHANIE; VANDERSCHAEVE, L.; MATHIEU, GUIDO; GOETGHEBEUR, PAUL; NEINHUIS, C. (2006):
**Phylogeny of the Genus _Peperomia_ (Piperaceae) inferred from the trnK/matK Region (cpDNA).**
In: Plant Biology 8 (1), p. 93–102. DOI: 10.1055/s-2005-873060.

### Piperales

JARAMILLO, M. ALEJANDRA; KRAMER, ELENA M. (2007):
**Molecular Evolution of the Petal and Stamen Identity Genes, Apetala3 and Pistillata, after Petal Loss in the Piperales.**
In: Molecular Phylogenetics and Evolution 44 (2), p. 598–609. DOI: 10.1016/j.ympev.2007.03.015.

TRUEBA, SANTIAGO; ROWE, NICK P.; NEINHUIS, CHRISTOPH; WANKE, STEFAN; WAGNER, SARAH T.; ISNARD, SANDRINE (2015):
**Stem Anatomy and the Evolution of Woodiness in Piperales.**
In: International Journal of Plant Sciences 176 (5), p. 468–485. DOI: 10.1086/680595.

WANKE, STEFAN; JARAMILLO, M. ALEJANDRA; BORSCH, THOMAS; SAMAIN, MARIE-STÉPHANIE; QUANDT, DIETMAR; NEINHUIS, CHRISTOPH (2007):
**Evolution of Piperales - matK Gene and trnK Intron Sequence Data reveal Lineage specific Resolution Contrast.**
In: Molecular Phylogenetics and Evolution 42 (2), p. 477–497. DOI: 10.1016/j.ympev.2006.07.007.

### Pittosporaceae

BACON, CHRISTINE D.; ALLAN, GERARD J.; ZIMMER, ELIZABETH A.; WAGNER, WARREN L. (2011):
**Genome scans reveal high levels of Gene flow in Hawaiian _Pittosporum_.**
In: Taxon 60 (3), p. 733–741.

CAYZER, LINDY W.; CRISP, MICHAEL D.; DONALDSON, STUART (2007):
**_Cheiranthera_ (Pittosporaceae).**
In: Australian Systematic Botany 20 (4), p. 340. DOI: 10.1071/SB06045.

CAYZER, LINDY W.; CRISP, MICHAEL D.; TELFORD, IAN R.H. (2004):
**Cladistic Analysis and Revision of _Billardiera_ (Pittosporaceae).**
In: Australian Systematic Botany 17 (1), p. 83. DOI: 10.1071/SB03028.

CHANDLER, GREGORY T.; PLUNKETT, GREGORY M.; PINNEY, STEVEN M.; CAYZER, LINDY W.; GEMMILL, CHRISSEN E.C. (2007):
**Molecular and morphological Agreement in Pittosporaceae: phylogenetic Analysis with nuclear ITS and Plastid trn L– trn F Sequence Data.**
In: Australian Systematic Botany 20 (5), p. 390. DOI: 10.1071/SB07004.

GEMMILL, CHRISSEN E.C.; ALLAN, GERARD J.; WAGNER, WARREN L.; ZIMMER, ELIZABETH A. (2002):
**Evolution of insular Pacific *Pittosporum* (Pittosporaceae): Origin of the Hawaiian Radiation.**
In: Molecular Phylogenetics and Evolution 22 (1), p. 31–42. DOI: 10.1006/mpev.2001.1019.

### Plantaginaceae

ALBACH, DIRK C.; BRIGGS, BARBARA G. (2012):
**Phylogenetic Analysis of Australian Species of *Veronica* (V. Section *Labiatoides*; Plantaginaceae).**
In: Australian Systematic Botany 25 (5), p. 353–363. DOI: 10.1071/SB12014.

ALBACH, DIRK C.; CHASE, MARK W. (2004):
**Incongruence in Veroniceae (Plantaginaceae): Evidence from two Plastid and a nuclear ribosomal DNA Region.**
In: Molecular Phylogenetics and Evolution 32 (1), p. 183–197. DOI: 10.1016/j.ympev.2003.12.001.

ALBACH, DIRK C.; MEUDT, HEIDI M. (2010):
**Phylogeny of *Veronica* in the Southern and Northern Hemispheres based on plastid, nuclear ribosomal and nuclear low-copy DNA.**
In: Molecular Phylogenetics and Evolution 54 (2), p. 457–471. DOI: 10.1016/j.ympev.2009.09.030.

ALBACH, DIRK C.; MEUDT, HEIDI M.; OXELMAN, BENGT (2005):
**Piecing together the ''new'' Plantaginaceae.**
In: American Journal of Botany 92 (2), p. 297–315.

ALBACH, DIRK C.; STERNBURG, MAXIMILIAN VON; SCALONE, ROMAIN; BARDY, KATHARINA E. (2009):
**Phylogenetic Analysis and differentiation of *Veronica* Subgenus *Stenocarpon* in the Balkan Peninsula.**
In: Botanical Journal of the Linnean Society 159, p. 616–636.

ALBACH, DIRK C.; UTTERIDGE, TIMOTHY M.A.; WAGSTAFF, STEVEN J. (2005):
**Origin of Veroniceae (Plantaginaceae, formerly Scrophulariaceae) on New Guinea.**
In: Systematic Botany 30 (2), p. 412–423.

BALDWIN, BRUCE G.; KALISZ, SUSAN; ARMBRUSTER, W. SCOTT (2011):
**Phylogenetic Perspectives on Diversification, Biogeography, and Floral Evolution of *Collinsia* and *Tonella* (Plantaginaceae).**
In: American Journal of Botany 98 (4), p. 731–753.

BUHK, NIKLAS; ZHAO, LIANG; LI, HONG-QING; ALBACH, DIRK C. (2015):
**Molecular Systematics and Morphometrics in *Veronica* Subsect. Canae (Plantaginaceae).**
In: Plant Systematics and Evolution 301 (8), p. 1967–1979. DOI: 10.1007/s00606-015-1214-1.

DUNBAR-CO, STEPHANIE; WIECZOREK, ANIA M.; MORDEN, CLIFFORD W. (2008):
**Molecular Phylogeny and adaptive Radiation of the endemic Hawaiian *Plantago* Species (Plantaginaceae).**
In: American Journal of Botany 95 (9), p. 1177–1188. DOI: 10.3732/ajb.0800132.

EKER, İSMAÍL; YÜCESAN, BUHARA; SAMEEULLAH, MUHAMMAD; WEIß, WALTER; MÜLLER-URI, FRIEDER; GÜREL, EKREM; KREIS, WOLFGANG (2016):
**Phylogeny of Anatolian (Turkey) Species in the *Digitalis* Sect. *Globiflorae* (Plantaginaceae).**
In: Phytotaxa 244 (3), p. 263–282.

**Estes, Dwayne; Small, Randall L. (2008):**
**Phylogenetic Relationships of the Monotypic Genus *Amphianthus* (Plantaginaceae Tribe Gratioleae) inferred from Chloroplast DNA Sequences.**
In: Systematic Botany 33 (1), p. 176–182.

**Fernández-Mazuecos, Mario; Blanco-Pastor, José Luis; Vargas, Pablo (2013):**
**A Phylogeny of Toadflaxes (*Linaria* Mill.) based on Nuclear Internal Transcribed Spacer Sequences: Systematic and evolutionary Consequences.**
In: International Journal of Plant Sciences 174 (2), p. 234–249. DOI: 10.1086/668790.

**Hoggard, Ronald K.; Kores, Paul J.; Molvray, Mia; Hoggard, Gloria D.; Broughton, David A. (2003):**
**Molecular Systematics and Biogeography of the Amphibious Genus *Littorella* (Plantaginaceae).**
In: American Journal of Botany 90 (3), p. 429–435.

**Ishikawa, Naoko; Yokoyama, Jun U.N.; Tsukaya, Hirokazu (2009):**
**Molecular Evidence of reticulate Evolution in the Subgenus Plantago (Plantaginaceae).**
In: American Journal of Botany 96 (9), p. 1627–1635. DOI: 10.3732/ajb.0800400.

**Kelly, Laura J.; Culham, Alastair (2008):**
**Phylogenetic utility of more Axillary Growth4 (MAX4)-like Genes: a case Study in *Digitalis/Isoplexis* (Plantaginaceae).**
In: Plant Systematics and Evolution 273 (3-4), p. 133–149. DOI: 10.1007/s00606-008-0008-0.

**Kosachev, Petr A.; Behçet, Lütfi; Mayland-Quellhorst, Eike; Albach, Dirk C. (2016):**
**Analyzing reticulate Relationships using CpDNA and Pyrosequenced ITS1 as exemplified by *Veronica* Subgen. *Pseudolysimachium* (Plantaginaceae).**
In: Systematic Botany 41 (1), p. 105–119. DOI: 10.1600/036364416X690697.

**Li, Guo-Dong; Kim, Changkyun; Zha, Hong-Guang; Zhou, Zhuo; Nie, Ze-long; Sun, Hang (2014):**
**Molecular Phylogeny and Biogeography of the arctic-alpine Genus *Lagotis* (Plantaginaceae).**
In: Taxon 63 (1), p. 103–115. DOI: 10.12705/631.47.

**Martins, Aline C.; Scherz, Mark D.; Renner, Susanne p. (2014):**
**Several Origins of Floral Oil in the Angelonieae, a Southern Hemisphere disjunct Clade of Plantaginaceae.**
In: American Journal of Botany 101 (12), p. 2113–2120. DOI: 10.3732/ajb.1400470.

**Meudt, Heidi M. (2008):**
**Taxonomic Revision of Australasian snow Hebes (*Veronica*, Plantaginaceae).**
In: Australian Systematic Botany 21 (6), p. 387–421. DOI: 10.1071/SB08034.

**Meudt, Heidi M. (2011):**
**Amplified Fragment Length Polymorphism Data reveal a History of Auto- and Allopolyploidy in New Zealand endemic Species of *Plantago* (Plantaginaceae): new Perspectives on a taxonomically challenging Group.**
In: International Journal of Plant Sciences 172 (2), p. 220–237. DOI: 10.1086/657657.

**Meudt, Heidi M.; Bayly, Michael J. (2008):**
**Phylogeographic Patterns in the Australasian Genus *Chionohebe* (*Veronica* s.l., Plantaginaceae) based on AFLP and Chloroplast DNA Sequences.**
In: Molecular Phylogenetics and Evolution 47 (1), p. 319–338. DOI: 10.1016/j.ympev.2007.12.019.

**Meudt, Heidi M.; Simpson, Beryl B. (2007):**

**Pphylogenetic Analysis of morphological Characters in *Ourisia* (Plantaginaceae): taxonomic and evolutionary Implications.**

In: Annals of the Missouri Botanical Garden 94 (3), p. 554–570. DOI: 10.3417/0026-6493(2007)94[554:PAOMCI]2.0.CO;2.

**Müller, Kai F.; Albach, Dirk C. (2010):**

**Evolutionary rates in *Veronica* L. (Plantaginaceae): disentangling the influence of life History and Breeding System.**

In: Journal of molecular Evolution 70 (1), p. 44–56. DOI: 10.1007/s00239-009-9307-5.

**Muñoz-Centeno, Luz M.; Albach, Dirk C.; Sánchez-Agudo, Jose A.; Martínez-Ortega, M. Montserrat (2006):**

**Systematic Significance of Seed Morphology in *Veronica* (Plantaginaceae): a phylogenetic Perspective.**

In: Annals of Botany 98 (2), p. 335–350. DOI: 10.1093/aob/mcl120.

**Ogutcen, Ezgi; Vamosi, Jana C. (2016):**

**A phylogenetic Study of the Tribe Antirrhineae: Genome duplications and long-distance Dispersals from the Old World to the New World.**

In: American Journal of Botany 103 (6), p. 1071–1081. DOI: 10.3732/ajb.1500464.

**Oyama, Ryan K.; Baum, David A. (2004):**

**Phylogenetic Relationships of North American *Antirrhinum* (Veronicaceae).**

In: American Journal of Botany 91 (6), p. 918–925.

**Padilla-García, Nélida; Rojas-Andrés, Blanca M.; López-González, Noemí; Castro, Mariana; Castro, Sílvia; Loureiro, João et al. (2018):**

**The challenge of Species Delimitation in the diploid-polyploid Complex *Veronica* subsection *Pentasepalae*.**

In: Molecular Phylogenetics and Evolution 119, p. 196–209. DOI: 10.1016/j.ympev.2017.11.007.

**Philbrick, C. Thomas; Les, Donald H. (2000):**

**Phylogenetic studies in *Callitriche*: Implications for interpretation of ecological, karyological and Pollination System Evolution.**

In: Aquatic Botany 68, p. 123–141.

**Rojas-Andrés, Blanca M.; Martínez-Ortega, M. Montserrat (2016):**

**Taxonomic Revision of *Veronica* subsection *Pentasepalae* (*Veronica*, Plantaginaceae sensu APG III).**

In: Phytotaxa 285 (1), p. 1. DOI: 10.11646/phytotaxa.285.1.1.

**Rønsted, Nina; Chase, Mark W.; Albach, Dirk C.; Bello, Maria Angélica (2002):**

**Phylogenetic Relationships within *Plantago* (Plantaginaceae): Evidence from nuclear ribosomal ITS and Plastid trnL-F Sequence Data.**

In: Botanical Journal of the Linnean Society 139, p. 323–338.

**Saeidi-Mehrvarz, Shahryar; Zarre, Shahin (2004):**

**A cladistic Analysis of the Iranian Species of the Genus *Veronica* (Scrophulariaceae) with Emphasis on the Patterns of Homoplasy.**

In: Feddes Repertorium 115 (7-8), p. 519–529. DOI: 10.1002/fedr.200311051.

**Sánchez Agudo, José Ángel; Rico, Enrique; Sánchez Sánchez, José; Montserrat Martínez-Ortega, Maria (2009):**

**Pollen Morphology in the Genus *Veronica* L. (Plantaginaceae) and its systematic Significance.**

In: Grana 48 (4), p. 239–257. DOI: 10.1080/00173130903364723.

**Scalone, Romain; Kolf, Markus; Albach, Dirk C. (2013):**
**Mating System Variation in *Veronica* (Plantaginaceae): Inferences from Pollen/Ovule Ratios and other reproductive Traits.**
In: Nordic Journal of Botany 31 (3), p. 372–384. DOI: 10.1111/j.1756-1051.2012.01706.x.

**Sosa, Maria De Las Mercedes; Dematteis, Massimiliano (2013):**
**Taxonomic Position and identity of *Stemodia scoparioides* (Gratiolae, Plantaginaceae).**
In: Phytotaxa 135 (1), p. 35. DOI: 10.11646/phytotaxa.135.1.5.

**Surina, Boštjan; Pfanzelt, Simon; Einzmann, Helena J.R.; Albach, Dirk C. (2014):**
**Bridging the Alps and the Middle East: Evolution, Phylogeny and Systematics of the Genus *Wulfenia* (Plantaginaceae).**
In: Taxon 63 (4), p. 843–858. DOI: 10.12705/634.18.

**Tay, Mei Lin; Meudt, Heidi M.; Garnock-Jones, Philip J.; Ritchie, Peter A. (2010):**
**Erratum to: DNA Sequences from three Genomes reveal multiple long-distance Dispersals and non-Monophyly of Sections in Australasian *Plantago* (Plantaginaceae).**
In: Australian Systematic Botany 23 (4), p. 306–307. DOI: 10.1071/SB09040_ER.

**Tay, Mei Lin; Meudt, Heidi M.; Garnock-Jones, Philip J.; Ritchie, Peter A. (2010):**
**Testing Species Limits of New Zealand *Plantago* (Plantaginaceae) using Internal Transcribed Spacer (ITS) DNA Sequences.**
In: New Zealand Journal of Botany 48 (3-4), p. 205–224. DOI: 10.1080/0028825X.2010.518318.

**Thiv, Mike; Thulin, Mats; Hjertson, Mats; Kropf, Matthias; Linder, Hans Peter (2010):**
**Evidence for a vicariant Origin of Macaronesian–Eritreo/Arabian disjunctions in *Campylanthus* Roth (Plantaginaceae).**
In: Molecular Phylogenetics and Evolution 54 (2), p. 607–616. DOI: 10.1016/j.ympev.2009.10.009.

**Vargas, Pablo; Rosselló, J. A.; Oyama, Ryan K.; Gemes, J. (2004):**
**Molecular Evidence for Naturalness of Genera in the Tribe Antirrhineae (Scrophulariaceae) and three independent evolutionary Lineages from the New World and the Old World.**
In: Plant Systematics and Evolution 249 (3-4), p. 151–172. DOI: 10.1007/s00606-004-0216-1.

**Wagstaff, Steven J.; Garnock-Jones, Philip J. (1998):**
**Evolution and Biogeography of the *Hebe* complex (Scrophulariaceae) inferred from ITS Sequences.**
In: New Zealand Journal of Botany 36 (3), p. 425–437. DOI: 10.1080/0028825X.1998.9512581.

**Wagstaff, Steven J.; Garnock-Jones, Philip J. (2000):**
**Patterns of Diversification in *Chionohebe* and *Parahebe* (Scrophulariaceae) inferred from ITS Sequences.**
In: New Zealand Journal of Botany 38 (3), p. 389–407. DOI: 10.1080/0028825X.2000.9512691.

**Wessinger, Carolyn A.; Freeman, Craig C.; Mort, Mark E.; Rausher, Mark D.; Hileman, Lena C. (2016):**
**Multiplexed shotgun Genotyping resolves Species Relationships within the North American Genus *Penstemon*.**
In: American Journal of Botany 103 (5), p. 912–922. DOI: 10.3732/ajb.1500519.

**Wolfe, Andrea D.; Datwyler, Shannon L.; Randle, Christopher P. (2002):**

**A phylogenetic and Biogeographic Analysis of the Cheloneae (Scrophulariaceae) based on ITS and matK Sequence Data.**
In: Systematic Botany 27 (1), p. 138–148.

**Wolfe, Andrea D.; Randle, Christopher P.; Datwyler, Shannon L.; Morawetz, Jeffery J.; Arguedas, Nidia (2006):**
**Phylogeny, taxonomic Affinities, and Biogeography of *Penstemon* (Plantaginaceae) based on ITS and cpDNA Sequence Data.**
In: American Journal of Botany 93 (11), p. 1699–1713.

**Yousefi, Nafiseh; Zarre, Shahin; Heubl, Günther (2016):**
**Molecular Phylogeny of the mainly Mediterranean Genera *Chaenorhinum, Kickxia* and *Nanorrhinum* (Plantaginaceae, Tribe Antirrhineae), with Focus on Taxa in the Flora Iranica Region.**
In: Nordic Journal of Botany 34 (4), p. 455–463. DOI: 10.1111/njb.01000.

### Platanaceae

**Feng, Yun; Oh, Sang-Hun; Manos, Paul p. (2005):**
**Phylogeny and Historical Biogeography of the Genus *Platanus* as inferred from Nuclear and Chloroplast DNA.**
In: Systematic Botany 30 (4), p. 786–799.

**Grimm, Guido W.; Denk, Thomas (2008):**
**ITS Evolution in *Platanus* (Platanaceae): homoeologues, pseudogenes and ancient Hybridization.**
In: Annals of Botany 101 (3), p. 403–419. DOI: 10.1093/aob/mcm305.

**Grimm, Guido W.; Denk, Thomas (2010):**
**The reticulate Origin of modern Plane Trees (*Platanus*, Platanaceae): A nuclear Marker Puzzle.**
In: Taxon 59 (1), p. 134–147.

### Plumbaginaceae

**Akhani, Hossein; Malekmohammadi, Maryam; Mahdavi, Parastoo; Gharibiyan, Arootin; Chase, Mark W. (2013):**
**Phylogenetics of the Irano-Turanian Taxa of *Limonium* (Plumbaginaceae) based on ITS nrDNA Sequences and Leaf Anatomy provides Evidence for Species Delimitation and Relationships of Lineages.**
In: Botanical Journal of the Linnean Society 171 (3), p. 519–550. DOI: 10.1111/boj.12015.

**Brullo, Salvatore; Erben, Matthias (2016):**
**The Genus *Limonium* (Plumbaginaceae) in Greece.**
In: Phytotaxa 240 (1), p. 1. DOI: 10.11646/phytotaxa.240.1.1.

**Fuertes-Aguilar, Javier; Nieto Feliner, Gonzalo (2003):**
**Additive Polymorphisms and Reticulation in an ITS Phylogeny of Thrifts (*Armeria*, Plumbaginaceae).**
In: Molecular Phylogenetics and Evolution 28 (3), p. 430–447. DOI: 10.1016/S1055-7903(02)00301-9.

**Lledó, M. Dolores (2000):**
**Polyphyly of *Limoniastrum* (Plumbaginaceae): Evidence from DNA Sequences of Plastid rbc L, trn L Intron and trnL-F intergene Spacer.**
In: Botanical Journal of the Linnean Society 132 (2), p. 175–191. DOI: 10.1006/bojl.1999.0304.

**Lledó, M. Dolores; Crespo, Manuel B.; Fay, Michael F.; Chase, Mark W. (2005):**
**Molecular Phylogenetics of *Limonium* and related Genera (Plumbaginaceae): biogeographical and systematic Implications.**
In: American Journal of Botany 92 (7), p. 1189–1198. DOI: 10.3732/ajb.92.7.1189.

**Lledó, M. Dolores; Karis, Per Ola; Crespo, Manuel B.; Fay, Michael F.; Chase, Mark W. (2001):**
**Phylogenetic Position and taxonomic Status of the Genus *Aegialitis* and subfamilies Staticoideae and Plumbaginoideae (Plumbaginaceae): Evidence from Plastid DNA Sequences and Morphology.**
In: Plant Systematics and Evolution 229 (1-2), p. 107–124. DOI: 10.1007/s006060170021.

**Moharrek, Farideh; Kazempour-Osaloo, Shahrokh; Assadi, Mostafa (2014):**
**Molecular Phylogeny of Plumbaginaceae with Emphasis on *Acantholimon* Boiss. based on nuclear and Plastid DNA Sequences in Iran.**
In: Biochemical Systematics and Ecology 57, p. 117–127. DOI: 10.1016/j.bse.2014.07.023.

**Palacios, C.; Rosselló, J. A.; González-Candelas, F. (2000):**
**Study of the evolutionary Relationships among *Limonium* Species (Plumbaginaceae) using nuclear and cytoplasmic molecular Markers.**
In: Molecular Phylogenetics and Evolution 14 (2), p. 232–249. DOI: 10.1006/mpev.1999.0690.

### Poaceae

**Acosta, Juan M.; Scataglini, Maria Amalia; Reinheimer, R.; Zuloaga, Fernando Omar (2014):**
**A phylogenetic Study of Subtribe Otachyriinae (Poaceae, Panicoideae, Paspaleae).**
In: Plant Systematics and Evolution 300 (10), p. 2155–2166. DOI: 10.1007/s00606-014-1034-8.

**Adoukonou-Sagbadja, Hubert; Wagner, Carola; Ordon, Frank; Friedt, Wolfgang (2010):**
**Reproductive System and molecular phylogenetic Relationships of Fonio Millets (*Digitaria* spp., Poaceae) with some polyploid wild Relatives.**
In: Tropical Plant Biology 3 (4), p. 240–251. DOI: 10.1007/s12042-010-9063-0.

**Akiyama, Yukio; Goel, Shailendra; Conner, Joann A.; Hanna, Wayne W.; Yamada-Akiyama, Hitomi; Ozias-Akins, Peggy (2011):**
**Evolution of the Apomixis transmitting Chromosome in *Pennisetum*.**
In: BMC Evolutionary Biology 11, p. 289.

**Aliscioni, Sandra S.; Giussani, Liliana M.; Zuloaga, Fernando Omar; Kellogg, Elizabeth A. (2003):**
**A molecular Phylogeny of *Panicum* (Poaceae: Paniceae): tests of Monophyly and phylogenetic Placement within the Panicoideae.**
In: American Journal of Botany 90 (5), p. 796–821. DOI: 10.3732/ajb.90.5.796.

**Allende, Jose Ramon Grande (2014):**
**Novitates Agrostologicae IV. Additional Segregates from *Panicum* Incertae Sedis.**
In: Phytoneuron 22, p. 1–6.

**Amarilla, Leonardo D.; Chiapella, Jorge O.; Nagahama, Nicolás; Anton, Ana M. (2013):**
**Inclusion of *Dasyochloa* in the amphitropical Genus *Munroa* (Poaceae, Chloridoideae) based on morphological Evidence.**
In: Darwiniana, nueva serie 1 (2), p. 241–252. DOI: 10.14522/darwiniana.2013.12.502.

**Anderson, Benjamin M.; Barrett, Matthew D.; Krauss, Siegfried L.; Thiele, Kevin R. (2016):**
**Untangling a Species complex of arid zone Grasses (*Triodia*) reveals Patterns congruent with co-occurring Animals.**
In: Molecular Phylogenetics and Evolution 101, p. 142–162. DOI: 10.1016/j.ympev.2016.05.014.

**Arthan, Watchara; Traiperm, Paweena; Gale, Stephan W.; Norsaengsri, Monthon; Kethirun, Lalita (2016):**
**Re-evaluation of the taxonomic Status of *Hackelochloa* (Poaceae) based on anatomical and phenetic Analyses.**
In: Botanical Journal of the Linnean Society 181 (2), p. 224–245. DOI: 10.1111/boj.12411.

**Attigala, Lakshmi; Triplett, Jimmy K.; Kathriarachchi, Hashendra-Suvini; Clark, Lynn G. (2014):**
**A new Genus and a major temperate bamboo Lineage of the Arundinarieae (Poaceae: Bambusoideae) from Sri Lanka based on a multi-locus Plastid Phylogeny.**
In: Phytotaxa 174 (4), p. 187. DOI: 10.11646/phytotaxa.174.4.1.

**Attigala, Lakshmi; Wysocki, William P.; Duvall, Melvin R.; Clark, Lynn G. (2016):**
**Phylogenetic estimation and morphological Evolution of Arundinarieae (Bambusoideae: Poaceae) based on plastome phylogenomic Analysis.**
In: Molecular Phylogenetics and Evolution 101, p. 111–121. DOI: 10.1016/j.ympev.2016.05.008.

**Banfi, Enrico; Galasso, Gabriele; Foggi, Bruno; Kopecký, David; Ardenghi, Nicola M.G. (2017):**
**From *Schedonorus* and *Micropyropsis* to *Lolium* (Poaceae: Loliinae): new Combinations and typifications.**
In: Taxon 66 (3), p. 708–717. DOI: 10.12705/663.11.

**Barber, Janet C.; Hames, Kasey A.; Cialdella, Ana María; Giussani, Liliana M.; Morrone, Osvaldo (2009):**
**Phylogenetic Relationships of *Piptochaetium* Presl (Poaceae: Stipeae) and related Genera reconstructed from nuclear and Chloroplast Sequence Datasets.**
In: Taxon 58 (2), p. 375–380. DOI: 10.1002/tax.582005.

**Barker, Nigel P.; Galley, Chloé; Verboom, George Anthony; Mafa, P.; Gilbert, M.; Linder, Hans Peter (2007):**
**The Phylogeny of the Austral grass Subfamily Danthonioideae: Evidence from multiple Data sets.**
In: Plant Systematics and Evolution 264 (3-4), p. 135–156. DOI: 10.1007/s00606-006-0479-9.

**Besnard, Guillaume; Christin, Pascal-Antoine; Malé, Pierre-Jean G.; Coissac, Eric; Ralimanana, Hélène; Vorontsova, Maria p. (2013):**
**Phylogenomics and Taxonomy of Lecomtelleae (Poaceae), an isolated panicoid Lineage from Madagascar.**
In: Annals of Botany 112 (6), p. 1057–1066. DOI: 10.1093/aob/mct174.

**Bess, Emilie C.; Doust, Andrew N.; Kellogg, Elizabeth A. (2005):**
**A naked Grass in the "Bristle Clade": A phylogenetic and Developmental Study of *Panicum* Section *Bulbosa* (Paniceae: Poaceae).**
In: International Journal of Plant Sciences 166 (3), p. 371–381.

**Bieniek, Wojciech; Mizianty, Marta; Szklarczyk, Marek (2015):**
**Sequence Variation at the three Chloroplast loci (matK, rbcL, trnH-psbA) in the Triticeae Tribe (Poaceae): Comments on the Relationships and Utility in DNA Barcoding of selected Species.**
In: Plant Systematics and Evolution 301 (4), p. 1275–1286. DOI: 10.1007/s00606-014-1138-1.

**Birch, Joanne L.; Berwick, Freya M.; Walsh, Neville G.; Cantrill, David J.; Murphy, Daniel J. (2014):**
**Distribution of morphological Diversity within widespread Australian Species of Poa (Poaceae, Tribe Poeae) and Implications for Taxonomy of the Genus.**
In: Australian Systematic Botany 27 (6), p. 333–354. DOI: 10.1071/SB14028.

**Birch, Joanne L.; Cantrill, David J.; Walsh, Neville G.; Murphy, Daniel J. (2014):**

**Phylogenetic Investigation and Divergence dating of *Poa* (Poaceae, Tribe Poeae) in the Australasian Region.**
In: Botanical Journal of the Linnean Society 175 (4), p. 523–552. DOI: 10.1111/boj.12185.

BOUCHENAK-KHELLADI, YANIS; SALAMIN, NICOLAS; SAVOLAINEN, VINCENT; FOREST, FÉLIX; VAN DER BANK, MICHELLE; CHASE, MARK W.; HODKINSON, TREVOR R. (2008):
**Large multi-gene phylogenetic Trees of the Grasses (Poaceae): Progress towards complete tribal and generic level Sampling.**
In: Molecular Phylogenetics and Evolution 47 (2), p. 488–505. DOI: 10.1016/j.ympev.2008.01.035.

BRASSAC, JONATHAN; BLATTNER, FRANK R. (2015):
**Species-Level Phylogeny and Polyploid Relationships in *Hordeum* (Poaceae) inferred by Next-Generation Sequencing and in Silico Cloning of Multiple Nuclear Loci.**
In: Systematic Biology 64 (5), p. 792–808. DOI: 10.1093/sysbio/syv035.

BROWN, AUSTIN J. (2015):
**A morphological search for *Lachnagrostis* among the South African *Agrostis* and *Polypogon* (Poacaeae).**
In: Muelleria 34, p. 23–46.

BURKE, SEAN V.; GRENNAN, COLIN P.; DUVALL, MELVIN R. (2012):
**Plastome Sequences of two New World bamboos - *Arundinaria gigantea* and *Cryptochloa strictiflora* (Poaceae) - extend phylogenomic Understanding of Bambusoideae.**
In: American Journal of Botany 99 (12), p. 1951–1961. DOI: 10.3732/ajb.1200365.

BUSTOS, ALFREDO DE; JOUVE, NICOLAS (2002):
**Phylogenetic Relationships of the Genus *Secale* based on the Characterisation of rDNA ITS Sequences.**
In: Plant Systematics and Evolution 235 (1), p. 147–154. DOI: 10.1007/s00606-002-0215-z.

BUSTOS, ALFREDO DE; JOUVE, NICOLAS (2006):
**Characterisation and phylogenetic Analysis of the Genes coding for high molecular Weigth Glutenin Subunits in the diploid Species of *Aegilops*.**
In: International Journal of Plant Sciences 167 (2), p. 359–366.

CATALÁN, PILAR; OLMSTEAD, RICHARD G. (2000):
**Phylogenetic Reconstruction of the Genus *Brachypodium* P.Beauv. (Poaceae) from combined Sequences of Chloroplast ndhF Gene and nuclear ITS.**
In: Plant Systematics and Evolution 220, p. 1–19.

CATALÁN, PILAR; TORRECILLA, PEDRO; LÓPEZ RODRÍGUEZ, JOSE ANGEL; OLMSTEAD, RICHARD G. (2004):
**Phylogeny of the festucoid Grasses of Subtribe Loliinae and Allies (Poeae, Pooideae) inferred from ITS and trnL-F Sequences.**
In: Molecular Phylogenetics and Evolution 31 (2), p. 517–541. DOI: 10.1016/j.ympev.2003.08.025.

CATALÁN, PILAR; TORRECILLA, PEDRO; LÓPEZ-RODRÍGUEZ, JOSÉ A.; MÜLLER, JOCHEN; STACE, CLIVE A. (2007):
**A Systematic Approach to Subtribe Loliinae (Poaceae: Pooideae) based on phylogenetic Evidence.**
In: Aliso 23 (1), p. 380–405.

CERROS-TLATILPA, ROSA; COLUMBUS, TRAVIS J.; BARKER, NIGEL P. (2011):
**Phylogenetic Relationships of *Aristida* and Relatives (Poaceae, Aristidoideae) based on noncoding Chloroplast (trnL-F, rpl16) and nuclear (ITS) DNA Sequences.**

In: American Journal of Botany 98 (11), p. 1868–1886. DOI: 10.3732/ajb.1100103.

**Che, Yonghe; Yang, Yanping; Yang, Xinming; Li, Xiuquan; Li, Lihui (2015):**

**Phylogenetic Relationship and Diversity among *Agropyron* Gaertn. Germplasm using SSRs Markers.**

In: Plant Systematics and Evolution 301 (1), p. 163–170. DOI: 10.1007/s00606-014-1062-4.

**Chemisquy, Maria Amelia; Giussani, Liliana M.; Scataglini, Maria Amalia; Kellogg, Elizabeth A.; Morrone, Osvaldo (2010):**

**Phylogenetic studies favour the Unification of *Pennisetum, Cenchrus* and *Odontelytrum* (Poaceae): a combined nuclear, Plastid and morphological Analysis, and nomenclatural Combinations in *Cenchrus*.**

In: Annals of Botany 106 (1), p. 107–130. DOI: 10.1093/aob/mcq090.

**Chiapella, Jorge O. (2007):**

**A molecular phylogenetic Study of *Deschampsia* (Poaceae: Aveneae) inferred from nuclear ITS and Plastid trnL Sequence Data: support for the Recognition of *Avenella* and *Vahlodea*.**

In: Taxon 56 (1), p. 55–64.

**Christin, Pascal-Antoine; Besnard, Guillaume (2009):**

**Two independent C4 Origins in Aristidoideae (Poaceae) revealed by the Recruitment of distinct Phosphoenolpyruvate Carboxylase Genes.**

In: American Journal of Botany 96 (12), p. 2234–2239. DOI: 10.3732/ajb.0900111.

**Cialdella, Ana María; Giussani, Liliana M.; Aagesen, Lone; Zuloaga, Fernando Omar; Morrone, Osvaldo (2007):**

**A Phylogeny of *Piptochaetium* (Poaceae: Pooideae: Stipeae) and related Genera based on a combined Analysis including trnL-F, rpl16, and Morphology.**

In: Systematic Botany 32 (3), p. 545–559.

**Cialdella, Ana María; Salariato, Diego Leonel; Aagesen, Lone; Giussani, Liliana M.; Zuloaga, Fernando Omar; Morrone, Osvaldo (2010):**

**Phylogeny of New World Stipeae (Poaceae): an Evaluation of the Monophyly of *Aciachne* and *Amelichloa*.**

In: Cladistics 26 (6), p. 563–578. DOI: 10.1111/j.1096-0031.2010.00310.x.

**Cialdella, Ana María; Zuloaga, Fernando Omar (2011):**

**Taxonomic Study of *Gymnopogon* (Poaceae, Chloridoideae, Cynodonteae).**

In: Annals of the Missouri Botanical Garden 98 (3), p. 301–330. DOI: 10.3417/2009071.

**Clark, Lynn G.; Dransfield, Soejatmi; Triplett, Jimmy K.; Sánchez-Ken, J. Gabriel (2007):**

**Phylogenetic Relationships among the One-Flowered, Determinate Genera of Bambuseae (Poaceae: Bambusoideae).**

In: Aliso 23 (1), p. 315–332.

**Columbus, Travis J.; Cerros-Tlatilpa, Rosa; Kinney, Michael S.; Siqueiros-Delgado, Maria Elena; Bell, Hester L.; Griffith, M. Patrick; Refulio-Rodriguez, Nancy F. (2007):**

**Phylogenetics of Chloridoideae (Gramineae): a preliminary Study based on Nuclear Ribosomal Internal Transcribed Spacer and Chloroplast trnL-F Sequences.**

In: Aliso 23 (1), p. 565–579.

**Consaul, Laurie L.; Gillespie, Lynn J.; Waterway, Marcia J. (2008):**

**Systematics of North American arctic diploid *Puccinellia* (Poaceae): Morphology, DNA Content, and AFLP Markers.**

In: Systematic Botany 33 (2), p. 251–261. DOI: 10.1600/036364408784571662.

**Consaul, Laurie L.; Gillespie, Lynn J.; Waterway, Marcia J. (2010):**
**Evolution and polyploid Origins in North American arctic *Puccinellia* (Poaceae) based on nuclear ribosomal spacer and Chloroplast DNA Sequences.**
In: American Journal of Botany 97 (2), p. 324–336. DOI: 10.3732/ajb.0900180.

**Davidse, Gerrit; Soreng, Robert J.; Peterson, Paul M. (2009):**
***Agrostopoa* (Poaceae, Pooideae, Poeae, Poinae), a new Genus with three Species from Colombia.**
In: Novon: A Journal for Botanical Nomenclature 19 (1), p. 32–40. DOI: 10.3417/2007132.

**Davis, Jerrold I.; Soreng, Robert J. (2007):**
**A preliminary phylogenetic Analysis of the Grass Subfamily Pooideae (Poaceae), with Attention to structural Features of the Plastid and Nuclear Genomes, including an Intron Loss in GBSSI.**
In: Aliso 23 (1), p. 335–348.

**Davis, Jerrold I.; Soreng, Robert J. (2010):**
**Migration of Endpoints of two Genes relative to Boundaries between Regions of the Plastid Genome in the Grass Family (Poaceae).**
In: American Journal of Botany 97 (5), p. 874–892. DOI: 10.3732/ajb.0900228.

**Denham, Silvia S.; Zuloaga, Fernando Omar (2007):**
**Phylogenetic Relationships of the Decumbentes Group of *Paspalum, Thrasya,* and *Thrasyopsis* (Poaceae: Panicoideae: Paniceae).**
In: Aliso 23 (1), p. 545–562.

**Díaz-Pérez, A. J.; Sharifi-Tehrani, M.; Inda, Luis A.; Catalán, Pilar (2014):**
**Polyphyly, Gene-duplication and extensive Allopolyploidy framed the Evolution of the ephemeral *Vulpia* Grasses and other fine-leaved Loliinae (Poaceae).**
In: Molecular Phylogenetics and Evolution 79, p. 92–105. DOI: 10.1016/j.ympev.2014.06.009.

**Dillon, p. L.; Lawrence, P. K.; Henry, Robert J. (2001):**
**The Use of ribosomal ITS to determine phylogenetic Relationships within *Sorghum*.**
In: Plant Systematics and Evolution 230 (1-2), p. 97–110. DOI: 10.1007/s006060170007.

**Dillon, p. L.; Lawrence, P. K.; Henry, Robert J.; Price, H. J. (2007):**
***Sorghum* resolved as a distinct Genus based on combined ITS1, ndhF and Adh1 Analyses.**
In: Plant Systematics and Evolution 268 (1-4), p. 29–43. DOI: 10.1007/s00606-007-0571-9.

**Donadío, Sabina; Giussani, Liliana M.; Kellogg, Elizabeth A.; Zuolaga, Fernando O.; Morrone, Osvaldo (2009):**
**A preliminary molecular Phylogeny of *Pennisetum* and *Cenchrus* (Poaceae-Paniceae) based on the trnL-F, rpl16 Chloroplast Markers.**
In: Taxon 58 (2), p. 392–404. DOI: 10.1002/tax.582007.

**Dong, Zhen-Zhen; Fan, Xing; Sha, Li-Na; Zeng, Jian; Wang, Yi; Chen, Qian et al. (2013):**
**Phylogeny and molecular Evolution of the rbcL Gene of St Genome in *Elymus* sensu lato (Poaceae: Triticeae).**
In: Biochemical Systematics and Ecology 50, p. 322–330. DOI: 10.1016/j.bse.2013.05.005.

**Döring, Elke; Schneider, Julia; Hilu, Khidir W.; Röser, Martin (2007):**
**Phylogenetic Relationships in the Aveneae/Poeae complex (Pooideae, Paceae).**

In: Kew Bulletin 62, p. 407–424.

**Doust, Andrew N.; Penly, Anya M.; Jacobs, Surrey W.L.; Kellogg, Elizabeth A. (2007):**
**Congruence, Conflict, and Polyploidization shown by Nuclear and Chloroplast Markers in the Monophyletic "Bristle Clade" (Paniceae, Panicoideae, Poaceae).**
In: Systematic Botany 32 (3), p. 531–544.

**Duvall, Melvin R.; Davis, Jerrold; Clark, Lynn G.; Noll, Jeffrey; Goldman, Douglas H.; Sánchez-Ken, J. Gabriel (2007):**
**Phylogeny of the Grasses (Poaceae) Revisited.**
In: Aliso 23 (1), p. 237–247. DOI: 10.5642/aliso.20072301.18.

**Duvall, Melvin R.; Fisher, Amanda E.; Columbus, Travis J.; Ingram, Amanda L.; Wysocki, William P.; Burke, Sean V. et al. (2016):**
**Phylogenomics and Plastome Evolution of the Chloridoid Grasses (Chloridoideae: Poaceae).**
In: International Journal of Plant Sciences 177 (3), p. 235–246. DOI: 10.1086/684526.

**Escobar, Juan S.; Scornavacca, Céline; Cenci, Alberto; Guilhaumon, Claire; Santoni, Sylvain; Douzery, Emmanuel J. P. et al. (2011):**
**Multigenic Phylogeny and Analysis of Tree Incongruences in Triticeae (Poaceae).**
In: BMC Evolutionary Biology 11, p. 181. DOI: 10.1186/1471-2148-11-181.

**Essi, Liliana; Longhi-Wagner, Hilda Maria; Souza-Chies, Tatiana Teixeira de (2008):**
**Phylogenetic Analysis of the *Briza*-complex (Poaceae).**
In: Molecular Phylogenetics and Evolution 47 (3), p. 1018–1029. DOI: 10.1016/j.ympev.2008.03.007.

**Essi, Liliana; Wagner, Hilda Maria Longhi; Souza-Chies, Tatiana Teixeira de (2014):**
**ISSR as a Tool to support taxonomic Decisions: a first Approach for *Chascolytrum* Species complexes (Poaceae).**
In: Journal of Biotechnology and Biodiversity 5 (2).

**Essi, Liliana; Wagner, Hilda Maria Longhi; Souza-Chies, Tatiana Teixeira de (2011):**
**New Combinations within the *Briza* Complex (Poaceae, Pooideae, Poeae).**
In: Novon: A Journal for Botanical Nomenclature 21 (3), p. 326–330. DOI: 10.3417/2010026.

**Fan, Xing; Liu, Jing; Sha, Li-Na; Sun, Gen-Lou; Hu, Zhi-Qin; Zeng, Jian et al. (2014):**
**Evolutionary Pattern of rDNA following Polyploidy in *Leymus* (Triticeae: Poaceae).**
In: Molecular Phylogenetics and Evolution 77, p. 296–306. DOI: 10.1016/j.ympev.2014.04.016.

**Fan, Xing; Sha, Li-Na; Dong, Zhen-Zhen; Zhang, Hai-Qin; Kang, Hou-Yang; Wang, Yi et al. (2013):**
**Phylogenetic Relationships and Y Genome Origin in *Elymus* L. sensu lato (Triticeae; Poaceae) based on single-copy nuclear Acc1 and Pgk1 Gene Sequences.**
In: Molecular Phylogenetics and Evolution 69 (3), p. 919–928. DOI: 10.1016/j.ympev.2013.06.012.

**Fan, Xing; Sha, Li-Na; Yang, Rui-Wu; Zhang, Hai-Qin; Kang, Hou-Yang; Ding, Chun-Bang et al. (2009):**
**Phylogeny and evolutionary History of *Leymus* (Triticeae; Poaceae) based on a single-copy nuclear Gene Encoding Plastid Acetyl-CoA Carboxylase.**
In: BMC Evolutionary Biology 9, p. 247. DOI: 10.1186/1471-2148-9-247.

**Fan, Xing; Sha, Li-Na; Yu, Shuang-Bin; Wu, Dan-Dan; Chen, Xiao-Hong; Zhuo, Xiao-Feng et al. (2013):**
**Phylogenetic Reconstruction and Diversification of the Triticeae (Poaceae) based on single-copy nuclear Acc1 and Pgk1 Gene Data.**
In: Biochemical Systematics and Ecology 50, p. 346–360. DOI: 10.1016/j.bse.2013.05.010.

**Fisher, Amanda E.; Hasenstab-Lehman, Kristen M.; Bell, Hester L.; Blaine, Ellen; Ingram, Amanda L.; Columbus, Travis J. (2016):**

**Evolutionary History of chloridoid Grasses estimated from 122 nuclear Loci.**
In: Molecular Phylogenetics and Evolution 105, p. 1–14. DOI: 10.1016/j.ympev.2016.08.011.

FISHER, AMANDA E.; TRIPLETT, JAMES K.; HO, CHI-SING; SCHILLER, ALICIA D.; OLTROGGE, KIMBERLY A.; SCHRODER, ERIC P. ET AL. (2009):
**Paraphyly in the Bamboo Subtribe Chusqueinae (Poaceae: Bambusoideae) and a revised infrageneric Classification for *Chusquea*.**
In: Systematic Botany 34 (4), p. 673–683. DOI: 10.1600/036364409790139790.

FORTUNE, PHILIPPE M.; POURTAU, NATHALIE; VIRON, NICOLAS; AÏNOUCHE, MALIKA L. (2008):
**Molecular Phylogeny and reticulate Origins of the polyploid Bromus Species from Section *Genea* (Poaceae).**
In: American Journal of Botany 95 (4), p. 454–464. DOI: 10.3732/ajb.95.4.454.

GALLEY, CHLOÉ; LINDER, HANS PETER (2007):
**The Phylogeny of the Pentaschistis Clade (Danthonioideae, Poaceae) based on Chloroplast DNA, and the Evolution and loss of complex Characters.**
In: Evolution 61 (4), p. 864–884. DOI: 10.1111/j.1558-5646.2007.00067.x.

GAMACHE, JACQUES; SUN, GEN-LOU (2015):
**Phylogenetic Analysis of the Genus *Pseudoroegneria* and the Triticeae Tribe using the RbcL Gene.**
In: Biochemical Systematics and Ecology 62, p. 73–81. DOI: 10.1016/j.bse.2015.07.038.

GAO, GANG; TANG, ZILIN; WANG, QIAN; GOU, XUEMEI; DING, CHUN-BANG; ZHANG, LI ET AL. (2014):
**Phylogeny and maternal donor of *Kengyilia* (Triticeae: Poaceae) based on Chloroplast trnT–trnL Sequences.**
In: Biochemical Systematics and Ecology 57, p. 102–107. DOI: 10.1016/j.bse.2014.07.004.

GE, YA-LONG GUO AND SONG (2005):
**Molecular Phylogeny of Oryzeae (Poaceae) based on DNA Sequences from Chloroplast, Mitochontrial and Nuclear Genomes.**
In: American Journal of Botany 92 (9), p. 1548–1558.

GILLESPIE, LYNN J.; ARCHAMBAULT, ANNIE; SORENG, ROBERT J. (2007):
**Phylogeny of *Poa* (Poaceae) based on trnT–trnF Sequence Data: Major Clades and basal Relationships.**
In: Aliso 23 (1), p. 420–434.

GILLESPIE, LYNN J.; SORENG, ROBERT J. (2005):
**A phylogenetic Analysis of the Bluegrass Genus Poa based on cpDNA Restriction Site Data.**
In: Systematic Botany 30 (1), p. 84–105.

GILLESPIE, LYNN J.; SORENG, ROBERT J.; BULL, ROGER D.; JACOBS, SURREY W.L.; REFULIO-RODRIGUEZ, NANCY F. (2008):
**Phylogenetic Relationships in Subtribe Poinae (Poaceae, Poeae) based on nuclear ITS and Plastid trnT– trnL– trnF Sequences.**
In: Botany 86 (8), p. 938–967. DOI: 10.1139/B08-076.

GILLESPIE, LYNN J.; SORENG, ROBERT J.; JACOBS, SURREY W.L. (2009):
**Phylogenetic Relationships of Australian *Poa* (Poaceae: Poinae), including molecular Evidence for two new Genera, *Saxipoa* and *Sylvipoa*.**
In: Australian Systematic Botany 22 (6), p. 413–436. DOI: 10.1071/SB09016.

**GIUSSANI, LILIANA M.; ZULOAGA, FERNANDO OMAR; QUARÍN, CAMILO L.; COTA-SÁNCHEZ, J. HUGO; UBAYASENA, KUMARY; MORRONE, OSVALDO (2009):**

**Phylogenetic Relationships in the Genus *Paspalum* (Poaceae: Panicoideae: Paniceae): An Assessment of the *Quadrifaria* and *Virgata* Informal Groups.**

In: Systematic Botany 34 (1), p. 32–43. DOI: 10.1600/036364409787602258.

**GOH, W. L.; CHANDRAN, S.; FRANKLIN, D. C.; ISAGI, YUJI; KOSHY, K. C.; SUNGKAEW, P. ET AL. (2013):**

**Multi-gene Region phylogenetic Analyses suggest reticulate Evolution and a Clade of Australian Origin among paleotropical woody Bamboos (Poaceae: Bambusoideae: Bambuseae).**

In: Plant Systematics and Evolution 299 (1), p. 239–257. DOI: 10.1007/s00606-012-0718-1.

**GOH, W. L.; CHANDRAN, S.; LIN, R.-S.; XIA, NIAN-HE; WONG, KHOON MENG (2010):**

**Phylogenetic Relationships among Southeast Asian climbing Bamboos (Poaceae: Bambusoideae) and the *Bambusa* Complex.**

In: Biochemical Systematics and Ecology 38 (4), p. 764–773. DOI: 10.1016/j.bse.2010.07.006.

**GOLOVNINA, K. A.; GLUSHKOV, P. A.; BLINOV, ALEXANDER G.; MAYOROV, V. I.; ADKISON, L. R.; GONCHAROV, N. P. (2007):**

**Molecular Phylogeny of the Genus *Triticum* L.**

In: Plant Systematics and Evolution 264 (3-4), p. 195–216. DOI: 10.1007/s00606-006-0478-x.

**GUO, GUO-YE; YANG, RUI-WU; DING, CHUN-BANG; FAN, XING; ZHANG, LI; ZHOU, YONG-HONG (2014):**

**Phylogenetic Relationships among *Leymus* and related diploid Genera (Triticeae: Poaceae) based on Chloroplast trnQ-rps16 Sequences.**

In: Nordic Journal of Botany 32 (5), p. 658–666. DOI: 10.1111/j.1756-1051.2013.00233.x.

**GUO, JUN; YU, XIAOCHENG; YIN, HUAYAN; LIU, GUOJUAN; LI, ANFEI; WANG, HONGWEI; KONG, LINGRANG (2016):**

**Phylogenetic Relationships of *Thinopyrum* and *Triticum* Species revealed by SCoT and Cddp Markers.**

In: Plant Systematics and Evolution 302 (9), p. 1301–1309. DOI: 10.1007/s00606-016-1332-4.

**HAMASHA, HASSAN R.; HAGEN, K. BERNHARD VON; RÖSER, MARTIN (2012):**

***Stipa* (Poaceae) and allies in the Old World: Molecular Phylogenetics realigns Genus Circumscription and gives Evidence on the Origin of American and Australian Lineages.**

In: Plant Systematics and Evolution 298 (2), p. 351–367. DOI: 10.1007/s00606-011-0549-5.

**HARDION, LAURENT; VERLAQUE, RÉGINE; BAUMEL, ALEX; JUIN, MARIANICK; VILA, BRUNO (2012):**

**Revised Systematics of Mediterranean *Arundo* (Poaceae) based on AFLP Fingerprints and Morphology.**

In: Taxon 61 (6), p. 1217–1226.

**HEMPEL, WERNER (2011):**

**Revision und Phylogenie der Arten der Gattung *Melica* L. (Poaceae) in Eurasien und Nordafrika.**

In: Feddes Repertorium 122 (1-2), p. 1–253. DOI: 10.1002/fedr.201100029.

**HOCHBACH, ANNE; SCHNEIDER, JULIA; RÖSER, MARTIN (2015):**

**A multi-locus Analysis of phylogenetic Relationships within grass Subfamily Pooideae (Poaceae) inferred from Sequences of nuclear single copy Gene Regions compared with Plastid DNA.**

In: Molecular Phylogenetics and Evolution 87, p. 14–27. DOI: 10.1016/j.ympev.2015.03.010.

**Hodge, Curtis D.; Wang, Huan; Sun, Gen-Lou (2010):**

**Phylogenetic Analysis of the maternal Genome of tetraploid StStYY *Elymus* (Triticeae: Poaceae) Species and the monogenomic Triticeae based on rps16 Sequence Data.**

In: Plant Science 178 (5), p. 463–468. DOI: 10.1016/j.plantsci.2010.03.002.

**Hodge, John G.; Kellogg, Elizabeth A. (2014):**

**Patterns of Inflorescence Development of three Prairie Grasses (Andropogoneae, Poaceae).**

In: International Journal of Plant Sciences 175 (9), p. 963–974. DOI: 10.1086/677652.

**Hodkinson, Trevor R.; Chase, Mark W.; Lledó, M. Dolores; Salamin, Nicolas; Renvoize, Stephen A. (2002):**

**Phylogenetics of *Miscanthus, Saccharum* and related Genera (Saccharinae, Andropogoneae, Poaceae) based on DNA Sequences from ITS nuclear ribosomal DNA and Plastid trnL-Intron and trnL-F intergenic spacers.**

In: Journal of Plant Research 115 (5), p. 381–392. DOI: 10.1007/s10265-002-0049-3.

**Hodkinson, Trevor R.; Chase, Mark W.; Takahashi, Chigusa; Leitch, Ilia J.; Bennett, Michael D.; Renvoize, Stephen A. (2002):**

**The use of DNA Sequencing (Its and trnL-F), AFLP, and fluorescent in situ Hybridization to Study allopolyploid *Miscanthus* (Poaceae).**

In: American Journal of Botany 89 (2), p. 279–286. DOI: 10.3732/ajb.89.2.279.

**Hodkinson, Trevor R.; Salamin, Nicolas; Chase, Mark W. (2007):**

**Large Trees, Supertrees, and Diversification of the Grass Family.**

In: Aliso 23 (1), p. 248–258. DOI: 10.5642/aliso.20072301.19.

**Hsiao, C.; Jacobs, Surrey W.L.; Barker, Nigel P.; Chatterton, N. J. (1998):**

**A molecular Phylogeny of the Subfamily Arundinoideae (Poaceae) based on Sequences of rDNA.**

In: Australian Systematic Botany 11 (1), p. 41–52. DOI: 10.1071/SB97001.

**Hunter, Angela M.; Orlovich, David A.; Lloyd, Kelvin M.; Lee, William G.; Murphy, Daniel J. (2004):**

**The generic Position of *Austrofestuca littoralis* and the Reinstatement of *Hookerochloa* and *Festucella* (Poaceae) based on Evidence from nuclear (ITS) and Chloroplast (trnL-trnF) DNA Sequences.**

In: New Zealand Journal of Botany 42 (2), p. 253–262. DOI: 10.1080/0028825X.2004.9512902.

**Ingram, Amanda L.; Doyle, Jeff J. (2007):**

***Eragrostis* (Poaceae): Monophyly and Infrageneric Classification.**

In: Aliso 23 (1), p. 595–604.

**Jacobs, Surrey W.L. (2001):**

**The Genus *Lachnagrostis* (Gramineae) in Australia.**

In: Telopea 9 (3), p. 439–448.

**Jacobs, Surrey W.L.; Bayer, Randall J.; Everett, Joy; Arriaga, Mirta; Barkworth, Mary E.; Sabin-Badereau, Alexandru et al. (2007):**

**Systematics of the Tribe Stipeae (Gramineae) using molecular Data.**

In: Aliso 23 (1), p. 349–361. DOI: 10.5642/aliso.20072301.28.

**Jones, Samuel S.; Burke, Sean V.; Duvall, Melvin R. (2014):**

**Phylogenomics, molecular Evolution, and estimated Ages of Lineages from the deep Phylogeny of Poaceae.**

In: Plant Systematics and Evolution 300 (6), p. 1421–1436. DOI: 10.1007/s00606-013-0971-y.

**Kelchner, Scot A. (2013):**
**Higher level phylogenetic Relationships within the Bamboos (Poaceae: Bambusoideae) based on five Plastid Markers.**
In: Molecular Phylogenetics and Evolution 67 (2), p. 404–413. DOI: 10.1016/j.ympev.2013.02.005.

**Kellogg, Elizabeth A. (2009):**
**The evolutionary History of Ehrhartoideae, Oryzeae, and *Oryza*.**
In: Rice 2 (1), p. 1–14. DOI: 10.1007/s12284-009-9022-2.

**Kellogg, Elizabeth A.; Hiser, Kenneth M.; Doust, Andrew N. (2004):**
**Taxonomy, Phylogeny, and Inflorescence Development of the Genus *Ixophorus* (Panicoideae: Poaceae).**
In: International Journal of Plant Sciences 165 (6), p. 1089–1105.

**Kim, Seonhee; Kim, Chang-Seok; Lee, Jeongran; Lee, In-Yong; Chung, Young-Jae; Cho, Myong-Suk; Kim, Seung-Chul (2015):**
**Phylogenetic Relationships among Species of *Setaria* (Paniceae; Panicoideae; Poaceae) in Korea: Insights from nuclear (Its and kn1) and Chloroplast DNA Sequence Data.**
In: Plant Systematics and Evolution 301 (2), p. 725–736. DOI: 10.1007/s00606-014-1111-z.

**Kinney, Michael S.; Columbus, Travis J.; Friar, Elizabeth A. (2007):**
**Dicliny in *Bouteloua* (Poaceae: Chloridoideae): Implications for the Evolution of Dioecy.**
In: Aliso 23 (1), p. 605–614.

**Kobayashi, Mikio (2015):**
**Phylogeny, Speciation, and Distribution of the Japanese dwarf Bamboos of Genus *Sasa* and allies.**
In: Plant Species Biology 30 (1), p. 45–62. DOI: 10.1111/1442-1984.12058.

**Kristiansen, Kim A.; Cilieborg, Malene; Drábková, Lenka; Jørgensen, Tina; Petersen, Gitte; Seberg, Ole (2005):**
**DNA Taxonomy - the Riddle of Oxychloe (Juncaceae).**
In: Systematic Botany 30 (2), p. 284–289.

**Lambertini, C.; Gustafsson, Mats H.G.; Frydenberg, J.; Lissner, J.; Speranza, M.; Brix, H. (2006):**
**A phylogeographic Study of the cosmopolitan Genus *Phragmites* (Poaceae) based on AFLPs.**
In: Plant Systematics and Evolution 258 (3-4), p. 161–182. DOI: 10.1007/s00606-006-0412-2.

**Lei, Ying-Xia; Liu, Jia; Fan, Xing; Sha, Li-Na; Wang, Yi; Kang, Hou-Yang et al. (2018):**
**Phylogeny and maternal Donor of *Roegneria* and its affinitive Genera (Poaceae: Triticeae) based on Sequence Data for two Chloroplast DNA Regions (ndhF and trnH-psbA ).**
In: Journal of Systematics and Evolution 56 (2), p. 105–119. DOI: 10.1111/jse.12291.

**Liao, Jin-Qiu; Ross, Loretta; Fan, Xing; Sha, Li-Na; Kang, Hou-Yang; Zhang, Hai-Qin et al. (2013):**
**Phylogeny and maternal Donors of the tetraploid Species with St Genome (Poaceae: Triticeae) inferred from CoxII and ITS Sequences.**
In: Biochemical Systematics and Ecology 50, p. 277–285. DOI: 10.1016/j.bse.2013.05.004.

**Linder, Hans Peter; Baeza, Marcelo; Barker, Nigel P.; Galley, Chloé; Humphreys, Aelys M.; Lloyd, Kelvin M. et al. (2010):**
**A generic Classification of the Danthonioideae (Poaceae).**
In: Annals of the Missouri Botanical Garden 97 (3), p. 306–364. DOI: 10.3417/2009006.

LIU, Q.; ZHANG, D. X.; PETERSON, PAUL M. (2010):
**Lemma micromorphological Characters in the Chloridoideae (Poaceae) optimized on a molecular Phylogeny.**
In: South African Journal of Botany 76 (2), p. 196–209. DOI: 10.1016/j.sajb.2009.10.006.

LIU, QUANLAN; GE, SONG; TANG, HAIBAO; ZHANG, XIANGLIN; ZHU, GENFENG; LU, BAO-RONG (2006):
**Phylogenetic Relationships in *Elymus* (Poaceae: Triticeae) based on the nuclear ribosomal Internal Transcribed Spacer and Chloroplast trnL-F Sequences.**
In: the new Phytologist 170 (2), p. 411–420. DOI: 10.1111/j.1469-8137.2006.01665.x.

LIZARAZU, MABEL A.; NICOLA, MARCELA V.; SALARIATO, DIEGO LEONEL (2014):
**Taxonomic re-evaluation of *Panicum* Sections *Tuerckheimiana* and *Valida* (Poaceae: Panicoideae) Using morphological and molecular Data.**
In: Taxon 63 (2), p. 265–274. DOI: 10.12705/632.34.

LLOYD, KELVIN M.; HUNTER, ANGELA M.; ORLOVICH, DAVID A.; DRAFFIN, SUZANNE J.; STEWART, ALAN V.; LEE, WILLIAM G. (2007):
**Phylogeny and Biogeography of Endemic *Festuca* (Poaceae) from New Zealand based on Nuclear (ITS) and Chloroplast (trnL–trnF) Nucleotide Sequences.**
In: Aliso 23 (1), p. 406–419.

MA, HONG-ZHENG; CAI, ZHE; ZHANG, FU-MIN; ZHANG, HUI; GE, SONG; DAI, SI-LAN; CHEN, WEN-LI (2015):
**Taxonomic Evaluation of *Miscanthus nudipes* (Poaceae) based on morphological and molecular Evidence.**
In: Phytotaxa 205 (1), p. 1. DOI: 10.11646/phytotaxa.205.1.1.

MARKOVA, DRAGOMIRA N.; MASON-GAMER, ROBERTA J. (2015):
**Diversity, abundance, and evolutionary dynamics of Pong-like transposable elements in Triticeae.**
In: Molecular Phylogenetics and Evolution 93, p. 318–330. DOI: 10.1016/j.ympev.2015.07.008.

MARTÍNEZ, ANA MARÍA SORIANO; SALAZAR, GERARDO A.; ARANDA, PATRICIA DÁVILA (2007):
**Phylogenetic Relationships of *Zeugites* (Poaceae: Centothecoideae) inferred from Plastid and Nuclear DNA Sequences and Morphology.**
In: Systematic Botany 32 (4), p. 722–730.

MASON-GAMER, ROBERTA J. (2005):
**The {beta}-Amylase Genes of Grasses and a phylogenetic Analysis of the Triticeae (Poaceae).**
In: American Journal of Botany 92 (6), p. 1045–1058. DOI: 10.3732/ajb.92.6.1045.

MATHEWS, SARAH; TSAI, ROCKY C.; KELLOGG, ELIZABETH A. (2000):
**Phylogenetic Structure in the grass Family (Poaceae): Evidence from the nuclear Gene Phytochrome B.**
In: American Journal of Botany 87 (1), p. 96–107. DOI: 10.2307/2656688.

MEIMBERG, HARALD; RICE, KEVIN J.; MILAN, NEIL F.; NJOKU, COLLINS C.; MCKAY, JOHN K. (2009):
**Multiple Origins promote the ecological Amplitude of allopolyploid *Aegilops* (Poaceae).**
In: American Journal of Botany 96 (7), p. 1262–1273. DOI: 10.3732/ajb.0800345.

MORRIS, LEAH M.; DUVALL, MELVIN R. (2010):
**The Chloroplast Genome of *Anomochloa marantoidea* (Anomochlooideae; Poaceae) comprises a mixture of grass-like and unique Features.**

In: American Journal of Botany 97 (4), p. 620–627. DOI: 10.3732/ajb.0900226.

**Morrone, Osvaldo; Aagesen, Lone; Scataglini, Maria Amalia; Salariato, Diego Leonel; Denham, Silvia S.; Chemisquy, Maria Amelia et al. (2012):**
**Phylogeny of the Paniceae (Poaceae: Panicoideae): integrating Plastid DNA Sequences and Morphology into a new Classification.**
In: Cladistics 28 (4), p. 333–356. DOI: 10.1111/j.1096-0031.2011.00384.x.

**Morrone, Osvaldo; Denham, Silvia S.; Aliscioni, Sandra S.; Zuloaga, Fernando Omar (2008):**
***Parodiophyllochloa*, a new Genus segregated from *Panicum* (Paniceae, Poaceae) based on morphological and molecular Data.**
In: Systematic Botany 33 (1), p. 66–76. DOI: 10.1600/036364408783887393.

**Morrone, Osvaldo; Scataglini, Maria Amalia; Zuloaga, Fernando Omar (2007):**
***Cyphonanthus*, a new Genus segregated from *Panicum* (Poaceae: Panicoideae: Paniceae) based on morphological, anatomical and molecular Data.**
In: Taxon 56 (2), p. 521–532.

**Naghavi, Mohammad Reza; Rad, Marjan Behzadii; Riahi, Mehrshid; Taleie, Alireza (2013):**
**Phylogenetic Analysis in some *Hordeum* Species (Triticeae; Poaceae) based on two single-copy nuclear Genes encoding Acetyl-CoA Carboxylase.**
In: Biochemical Systematics and Ecology 47, p. 148–155. DOI: 10.1016/j.bse.2012.10.007.

**Nasernakhaei, Fatemeh; Rahiminejad, Mohammad Reza; Saeidi, Hojjatollah; Tavassoli, Manoochehr (2015):**
**Phylogenetic Relationships among the Iranian *Triticum* diploid Gene Pool as inferred from the loci Acc1 and Pgk1.**
In: Phytotaxa 201 (2), p. 111. DOI: 10.11646/phytotaxa.201.2.1.

**Neves, Susana S.; Swire-Clark, Ginger; Hilu, Khidir W.; Baird, Wm Vance (2005):**
**Phylogeny of *Eleusine* (Poaceae: Chloridoideae) based on nuclear ITS and Plastid trnT-trnF Sequences.**
In: Molecular Phylogenetics and Evolution 35 (2), p. 395–419. DOI: 10.1016/j.ympev.2004.12.005.

**Ng'uni, Dickson; Geleta, Mulatu; Fatih, Moneim; Bryngelsson, Tomas (2010):**
**Phylogenetic Analysis of the Genus *Sorghum* based on combined Sequence Data from cpDNA Regions and ITS Generate well-supported Trees with two major Lineages.**
In: Annals of Botany 105 (3), p. 471–480. DOI: 10.1093/aob/mcp305.

**Nicola, Marcela V.; Lizarazu, Mabel A.; Scataglini, Maria Amalia (2015):**
**Phylogenetic Analysis and taxonomic Position of Section *Verrucosa* of *Panicum* and its Relationship with Taxa of the *Sacciolepis–Trichanthecium* Clade (Poaceae: Panicoideae: Paniceae).**
In: Plant Systematics and Evolution 301 (9), p. 2247–2260. DOI: 10.1007/s00606-015-1227-9.

**Oliveira, Reyjane Patrícia; Clark, Lynn G.; Schnadelbach, Alessandra Selbach; Monteiro, Silvana Helena N.; Borba, Eduardo L.; Longhi-Wagner, Hilda Maria; van den Berg, Cássio (2014):**
**A molecular Phylogeny of *Raddia* and its allies within the Tribe Olyreae (Poaceae, Bambusoideae) based on noncoding Plastid and nuclear Spacers.**
In: Molecular Phylogenetics and Evolution 78, p. 105–117. DOI: 10.1016/j.ympev.2014.04.012.

**Pattanaik, Swapnendu; Hall, John B. (2011):**
**Molecular Evidence for Polyphyly in the woody Bamboo Genus *Dendrocalamus* (Subtribe Bambusinae).**

In: Plant Systematics and Evolution 291 (1-2), p. 59–67. DOI: 10.1007/s00606-010-0380-4.

**Peng, Sheng; Yang, Han-Qi; Li, De-Zhu (2008):**
**Highly heterogeneous generic Delimitation within the temperate Bamboo Clade (Poaceae: Bambusoideae): Evidence from GBSSI and ITS Sequences.**
In: Taxon 57 (3), p. 799–810. DOI: 10.1002/tax.573011.

**Peng, Yuan-Ying; Wei, Yu-Ming; Baum, Bernard R.; Jiang, Qian-Tao; Lan, Xiu-Jin; Dai, Shou-Fen; Zheng, You-Liang (2010):**
**Phylogenetic Investigation of *Avena* diploid Species and the maternal Genome Donor of *Avena* Polyploids.**
In: Taxon 59 (5), p. 1472–1482.

**Persson, Nannie; Rydin, Catarina (2016):**
**Phylogenetic Relationships of the '*Briza* complex' to other members of the Subfamily Pooideae (Poaceae).**
In: Plant Ecology and Evolution 149 (2), p. 216–227. DOI: 10.5091/plecevo.2016.1194.

**Peter Linder, H.; Bouchenak-Khelladi, Yanis (2017):**
**Adaptive Radiations should not be simplified: the case of the danthonioid Grasses.**
In: Molecular Phylogenetics and Evolution 117, p. 179–190. DOI: 10.1016/j.ympev.2017.10.003.

**Petersen, Gitte; Seberg, Ole; Baden, C. (2004):**
**A phylogenetic Analysis of the Genus *Psathyrostachys* (Poaceae) based on one nuclear Gene, three Plastid Genes, and Morphology.**
In: Plant Systematics and Evolution 249 (1-2), p. 99–110. DOI: 10.1007/s00606-004-0196-1.

**Peterson, Paul M.; Romaschenko, Konstantin; Arrieta, Yolanda Herrera (2014):**
**A molecular Phylogeny and Classification of the Cteniinae, Farragininae, Gouiniinae, Gymnopogoninae, Perotidinae, and Trichoneurinae (Poaceae: Chloridoideae: Cynodonteae).**
In: Taxon 63 (2), p. 275–286. DOI: 10.12705/632.35.

**Peterson, Paul M.; Romaschenko, Konstantin; Arrieta, Yolanda Herrera (2015):**
**A molecular Phylogeny and Classification of the Eleusininae with a new Genus, *Micrachne* (Poaceae: Chloridoideae: Cynodonteae).**
In: Taxon 64 (3), p. 445–467. DOI: 10.12705/643.5.

**Peterson, Paul M.; Romaschenko, Konstantin; Arrieta, Yolanda Herrera; Saarela, Jeffery M. (2014):**
**A molecular Phylogeny and new subgeneric Classification of *Sporobolus* (Poaceae: Chloridoideae: Sporobolinae).**
In: Taxon 63 (6), p. 1212–1243. DOI: 10.12705/636.19.

**Peterson, Paul M.; Romaschenko, Konstantin; Arrieta, Yolanda Herrera (2015):**
**Phylogeny and subgeneric Classification of *Bouteloua* with a new species, *B. herrera-arrietae* (Poaceae: Chloridoideae: Cynodonteae: Boutelouinae).**
In: Journal of Systematics and Evolution 53 (4), p. 351–366. DOI: 10.1111/jse.12159.

**Peterson, Paul M.; Romaschenko, Konstantin; Barker, Nigel P.; Linder, Hans Peter (2011):**
**Centropodieae and *Ellisochloa*, a new Tribe and Genus in Chloridoideae (Poaceae).**
In: Taxon 60 (4), p. 1113–1122.

**Peterson, Paul M.; Romaschenko, Konstantin; Herrera Arrieta, Yolanda (2016):**

**A molecular Phylogeny and Classification of the Cynodonteae (Poaceae: Chloridoideae) with four new Genera: *Orthacanthus, Triplasiella, Tripogonella,* and *Zaqiqah*; three new Subtribes: Dactylocteniinae, Orininae, and Zaqiqahinae; and a subgeneric Classification of *Distichlis*.**
In: Taxon 65 (6), p. 1263–1287. DOI: 10.12705/656.4.

PETERSON, PAUL M.; ROMASCHENKO, KONSTANTIN; JOHNSON, GABRIEL (2010):
**A Classification of the Chloridoideae (Poaceae) based on multi-gene phylogenetic Trees.**
In: Molecular Phylogenetics and Evolution 55 (2), p. 580–598. DOI: 10.1016/j.ympev.2010.01.018.

PETERSON, PAUL M.; ROMASCHENKO, KONSTANTIN; JOHNSON, GABRIEL (2010):
**A Phylogeny and Classification of the Muhlenbergiinae (Poaceae: Chloridoideae: Cynodonteae) based on Plastid and nuclear DNA Sequences.**
In: American Journal of Botany 97 (9), p. 1532–1554. DOI: 10.3732/ajb.0900359.

PETERSON, PAUL M.; ROMASCHENKO, KONSTANTIN; SNOW, NEIL; JOHNSON, GABRIEL (2012):
**A molecular Phylogeny and Classification of *Leptochloa* (Poaceae: Chloridoideae: Chlorideae) sensu lato and related Genera.**
In: Annals of Botany 109 (7), p. 1317–1330. DOI: 10.1093/aob/mcs077.

PETERSON, PAUL M.; VEGA, ISIDORO SÁNCHEZ (2007):
**Eragrostis (Poaceae: Chloridoideae: Eragrostideae: Eragrostidinae) of Peru.**
In: Annals of the Missouri Botanical Garden 94 (4), p. 745–790. DOI: 10.3417/0026-6493(2007)94[745:EPCEEO]2.0.CO;2.

PIMENTEL, MANUEL; SAHUQUILLO, ELVIRA; CATALÁN, PILAR (2007):
**Genetic Diversity and spatial correlation Patterns unravel the biogeographical History of the European sweet vernal Grasses (*Anthoxanthum* L., Poaceae).**
In: Molecular Phylogenetics and Evolution 44 (2), p. 667–684. DOI: 10.1016/j.ympev.2007.04.006.

PIRIE, MICHAEL D.; HUMPHREYS, AELYS M.; BARKER, NIGEL P.; LINDER, HANS PETER (2009):
**Reticulation, Data Combination, and inferring evolutionary History: an Example from Danthonioideae (Poaceae).**
In: Systematic Biology 58 (6), p. 612–628. DOI: 10.1093/sysbio/syp068.

PLEINES, THEKLA; BLATTNER, FRANK R. (2008):
**Phylogeographic Implications of an AFLP Phylogeny of the American diploid *Hordeum* Species (Poaceae: Triticeae).**
In: Taxon 57 (3), p. 875–881. DOI: 10.1002/tax.573016.

QUINTANAR, ALEJANDRO; CASTROVIEJO, SANTIAGO; CATALÁN, PILAR (2007):
**Phylogeny of the Tribe Aveneae (Pooideae, Poaceae) inferred from Plastid trnT-F and nuclear ITS Sequences.**
In: American Journal of Botany 94 (9), p. 1554–1569. DOI: 10.3732/ajb.94.9.1554.

RAHMANIAN, S.; SAEIDI, HOJJATOLLAH; ASSADI, MOSTAFA; RAHIMINEJAD, MOHAMMAD REZA (2014):
**A taxonomic Revision of the Genus *Eremopoa* Roschv. (Poaceae, Poeae) in Iran.**
In: Iranian Journal of Botany 20 (1), p. 8–15.

REINHEIMER, R.; VEGETTI, A. C. (2008):
**Inflorescence Diversity and Evolution in the PCK Clade (Poaceae: Panicoideae: Paniceae).**
In: Plant Systematics and Evolution 275 (3-4), p. 133–167. DOI: 10.1007/s00606-008-0057-4.

**Romaschenko, Konstantin; Peterson, Paul M.; Soreng, Robert J.; Futorna, Oksana; Susanna, Alfonso (2011):**

**Phylogenetics of *Piptatherum* s.l. (Poaceae: Stipeae): Evidence for a new Genus, *Piptatheropsis*, and Resurrection of *Patis*.**

In: Taxon 60 (6), p. 1703–1716.

**Romaschenko, Konstantin; Peterson, Paul M.; Soreng, Robert J.; Garcia-Jacas, Núria; Futorna, Oksana; Susanna, Alfonso (2012):**

**Systematics and Evolution of the needle Grasses (Poaceae: Pooideae: Stipeae) based on Analysis of multiple Chloroplast loci, ITS, and Lemma Micromorphology.**

In: Taxon 61 (1), p. 18–44.

**Rousseau, Hélène; Rousseau-Gueutin, Mathieu; Dauvergne, Xavier; Boutte, Julien; Simon, Gaëlle; Marnet, Nathalie et al. (2017):**

**Evolution of DMSP (Dimethylsulfoniopropionate) Biosynthesis Pathway: Origin and phylogenetic Distribution in polyploid *Spartina* (Poaceae, Chloridoideae).**

In: Molecular Phylogenetics and Evolution 114, p. 401–414. DOI: 10.1016/j.ympev.2017.07.003.

**Rousseau-Gueutin, Mathieu; Bellot, Sidonie; Martin, G. E.; Boutte, J.; Chelaifa, H.; Lima, O. et al. (2015):**

**The Chloroplast Genome of the hexaploid *Spartina maritima* (Poaceae, Chloridoideae): Comparative Analyses and molecular Dating.**

In: Molecular Phylogenetics and Evolution 93, p. 5–16. DOI: 10.1016/j.ympev.2015.06.013.

**Rua, Gabriel H.; Speranza, Pablo R.; Vaio, Magdalena; Arakaki, Mónica (2010):**

**A phylogenetic Analysis of the Genus *Paspalum* (Poaceae) based on cpDNA and Morphology.**

In: Plant Systematics and Evolution 288 (3-4), p. 227–243. DOI: 10.1007/s00606-010-0327-9.

**Ruiz-Sanchez, Eduardo (2015):**

**Parametric and non-parametric Species Delimitation methods Result in the Recognition of two new Neotropical woody Bamboo Species.**

In: Molecular Phylogenetics and Evolution 93, p. 261–273. DOI: 10.1016/j.ympev.2015.08.004.

**Ruiz-Sanchez, Eduardo; Sosa, Victoria (2015):**

**Origin and Evolution of fleshy Fruit in woody Bamboos.**

In: Molecular Phylogenetics and Evolution 91, p. 123–134. DOI: 10.1016/j.ympev.2015.05.020.

**Ruiz-Sanchez, Eduardo; Sosa, Victoria; Mejía-Saules, M. Teresa (2011):**

**Molecular Phylogenetics of the Mesoamerican Bamboo *Olmeca* (Poaceae, Bambuseae): Implications for Taxonomy.**

In: Taxon 60 (1), p. 89–98.

**Ruiz-Sanchez, Eduardo; Sosa, Victoria; Mejía-Saules, M. Teresa (2008):**

**Phylogenetics of *Otatea* inferred from Morphology and Chloroplast DNA Sequence Data, and Recircumscription of Guaduinae (Poaceae: Bambusoideae).**

In: Systematic Botany 33 (2), p. 277–283. DOI: 10.1600/036364408784571644.

**Saarela, Jeffery M.; Bull, Roger D.; Paradis, Michel J.; Ebata, Sharon N.; Paul, M. Peterson; Soreng, Robert J.; Paszko, Beata (2017):**

**Molecular Phylogenetics of cool-season Grasses in the Subtribes Agrostidinae, Anthoxanthinae, Aveninae, Brizinae, Calothecinae, Koeleriinae and Phalaridinae (Poaceae, Pooideae, Poeae, Poeae Chloroplast Group 1).**

In: PhytoKeys (87), p. 1–139. DOI: 10.3897/phytokeys.87.12774.

**Saarela, Jeffery M.; Graham, Sean W. (2010):**

**Inference of phylogenetic Relationships among the subfamilies of Grasses (Poaceae: Poales) using meso-scale exemplar-based Sampling of the Plastid Genome.**
In: Botany 88 (1), p. 65–84. DOI: 10.1139/B09-093.

**Saarela, Jeffery M.; Peterson, Paul M.; Keane, Ryan M.; Cayouette, Jacques; Graham, Sean W. (2007):**
**Molecular Phylogenetics of *Bromus* (Poaceae: Pooideae) based on Chloroplast and Nuclear DNA Sequence Data.**
In: Aliso 23 (1), p. 450–467.

**Saarela, Jeffery M.; Peterson, Paul M.; Valdés-Reyna, Jesus (2014):**
**A taxonomic Revision of *Bromus* (Poaceae: Pooideae: Bromeae) in México and Central America.**
In: Phytotaxa 185 (1), p. 1. DOI: 10.11646/phytotaxa.185.1.1.

**Salamin, Nicolas; Hodkinson, Trevor R.; Savolainen, Vincent (2002):**
**Building Supertrees: an Empirical Assessment using the Grass Family (Poaceae).**
In: Systematic Biology 51 (1), p. 136–150.

**Salariato, Diego Leonel; Giussani, Liliana M.; Morrone, Osvaldo; Zuloaga, Fernando Omar (2009):**
**Rupichloa, a new Genus segregated from *Urochloa* (Poaceae) based on morphological and molecular Data.**
In: Taxon 58 (2), p. 381–391. DOI: 10.1002/tax.582006.

**Salariato, Diego Leonel; Morrone, Osvaldo; Zuloaga, Fernando Omar (2012):**
***Mayariochloa*, a new monotypic Genus segregated from *Scutachne* (Poaceae, Panicoideae, Paniceae).**
In: Systematic Botany 37 (1), p. 105–116. DOI: 10.1600/036364412X616684.

**Sánchez-Ken, J. Gabriel; Clark, Lynn G. (2007):**
**Phylogenetic Relationships within the Centothecoideae + Panicoideae Clade (Poaceae) based on ndhF and rpl16 Intron Sequences and Structural Data.**
In: Aliso 23 (1), p. 487–501.

**Sánchez-Ken, J. Gabriel; Clark, Lynn G. (2010):**
**Phylogeny and a new tribal Classification of the Panicoideae s.l. (Poaceae) based on Plastid and nuclear Sequence Data and structural Data.**
In: American Journal of Botany 97 (10), p. 1732–1748. DOI: 10.3732/ajb.1000024.

**Sánchez-Ken, J. Gabriel; Clark, Lynn G.; Kellogg, Elizabeth A.; Kay, Elma E. (2007):**
**Reinstatement and Emendation of Subfamily Micrairoideae (Poaceae).**
In: Systematic Botany 32 (1), p. 71–80.

**Scataglini, Maria Amalia; Zuloaga, Fernando Omar; Giussani, Liliana M.; Denham, Silvia S.; Morrone, Osvaldo (2014):**
**Phylogeny of New World *Paspalum* (Poaceae, Panicoideae, Paspaleae) based on Plastid and nuclear Markers.**
In: Plant Systematics and Evolution 300 (5), p. 1051–1070. DOI: 10.1007/s00606-013-0944-1.

**Schneider, Julia; Döring, Elke; Hilu, Khidir W.; Röser, Martin (2009):**
**Phylogenetic Structure of the Grass Subfamily Pooideae based on Comparison of Plastid matK gene–3'trnK Exon and nuclear ITS Sequences.**
In: Taxon 58 (2), p. 405–424.

**Schneider, Julia; Winterfeld, Grit; Röser, Martin (2012):**

**Polyphyly of the grass Tribe Hainardieae (Poaceae: Pooideae): Identification of its different Lineages based on molecular Phylogenetics, including morphological and cytogenetic Characteristics.**
In: Organisms Diversity and Evolution 12 (2), p. 113–132. DOI: 10.1007/s13127-012-0077-3.

**Sclovich, Sergio E.; Giussani, Liliana M.; Cialdella, Ana María; Sede, Silvana M. (2015):**
**Phylogenetic Analysis of *Jarava* (Poaceae, Pooideae, Stipeae) and related Genera: testing the value of the awn Indumentum in the Circumscription of *Jarava*.**
In: Plant Systematics and Evolution 301 (6), p. 1625–1641. DOI: 10.1007/s00606-014-1175-9.

**Seberg, Ole; Petersen, Gitte (2007):**
**Phylogeny of Triticeae (Poaceae) based on three Organelle Genes, two Single-Copy nuclear Genes, and Morphology.**
In: Aliso 23 (1), p. 362–371.

**Sede, Silvana M.; Morrone, Osvaldo; Aliscioni, Sandra S.; Giussani, Liliana M.; Zuloaga, Fernando Omar (2009):**
***Oncorachis* and *Sclerochlamys*, two new segregated Genera from *Streptostachys* (Poaceae, Panicoideae, Paniceae): a Revision based on molecular, morphological and anatomical Characters.**
In: Taxon 58 (2), p. 365–374.

**Sede, Silvana M.; Morrone, Osvaldo; Giussani, Liliana M.; Zuloaga, Fernando Omar (2008):**
**Phylogenetic Studies in the Paniceae (Poaceae): a Realignment of Section *Lorea* of *Panicum*.**
In: Systematic Botany 33 (2), p. 284–300. DOI: 10.1600/036364408784571626.

**Sede, Silvana M.; Zuloaga, Fernando Omar; Morrone, Osvaldo (2009):**
**Phylogenetic Studies in the Paniceae (Poaceae-Panicoideae): *Ocellochloa*, a new Genus from the New World.**
In: Systematic Botany 34 (4), p. 684–692. DOI: 10.1600/036364409790139655.

**Sha, Li-Na; Fan, Xing; Li, Jun; Liao, Jin-Qiu; Zeng, Jian; Wang, Yi et al. (2017):**
**Contrasting evolutionary Patterns of multiple Loci uncover new Aspects in the Genome Origin and evolutionary History of *Leymus* (Triticeae; Poaceae).**
In: Molecular Phylogenetics and Evolution 114, p. 175–188. DOI: 10.1016/j.ympev.2017.05.015.

**Sha, Li-Na; Fan, Xing; Zhang, Hai-Qin; Kang, Hou-Yang; Wang, Yi; Wang, Xiao-Li et al. (2016):**
**Phylogeny and molecular Evolution of the Dmc1 Gene in the polyploid Genus *Leymus* (Triticeae: Poaceae) and its diploid Relatives.**
In: Journal of Systematics and Evolution 54 (3), p. 250–263. DOI: 10.1111/jse.12188.

**Silva, Christian; Snak, Cristiane; Schnadelbach, Alessandra Selbach; van den Berg, Cássio; Oliveira, Reyjane Patrícia (2015):**
**Phylogenetic Relationships of *Echinolaena* and *Ichnanthus* within Panicoideae (Poaceae) reveal two new Genera of tropical grasses.**
In: Molecular Phylogenetics and Evolution 93, p. 212–233. DOI: 10.1016/j.ympev.2015.07.015.

**Skendzic, Elizabeth; Columbus, Travis J.; Cerros-Tlatilpa, Rosa (2007):**
**Phylogenetics of Andropogoneae (Poaceae: Panicoideae) based on Nuclear Ribosomal Internal Transcribed Spacer and Chloroplast trnL–F Sequences.**
In: Aliso 23 (1), p. 530–544. DOI: 10.5642/aliso.20072301.40.

**Snow, Neil; Peterson, Paul M. (2012):**

**Nomenclatural Notes on *Dinebra, Diplachne, Disakisperma* and *Leptochloa* (Poaceae: Chloridoidea).**
In: Phytoneuron 71, p. 1–2.

SORENG, ROBERT J.; GILLESPIE, LYNN J.; JACOBS, SURREY W.L. (2009):
***Saxipoa* and *Sylvipoa* - two new Genera and a new Classification for Australian *Poa* (Poaceae: Poinae).**
In: Australian Systematic Botany 22 (6), p. 401–412. DOI: 10.1071/SB09003.

SORENG, ROBERT J.; PETERSON, PAUL M.; ROMASCHENKO, KONSTANTIN; DAVIDSE, GERRIT; TEISHER, JORDAN K.; CLARK, LYNN G. ET AL. (2017):
**A worldwide phylogenetic Classification of the Poaceae (Gramineae) II: An Update and a Comparison of two 2015 Classifications.**
In: Journal of Systematics and Evolution 55 (4), p. 259–290. DOI: 10.1111/jse.12262.

SORENG, ROBERT J.; PETERSON, PAUL M.; ROMASCHENKO, KONSTANTIN; DAVIDSE, GERRIT; ZULOAGA, FERNANDO OMAR; JUDZIEWICZ, EMMET J. ET AL. (2015):
**A worldwide phylogenetic Classification of the Poaceae (Gramineae).**
In: Journal of Systematics and Evolution 53 (2), p. 117–137. DOI: 10.1111/jse.12150.

SOUTO, D. P. FERNÁNDEZ; CATALANO, SANTIAGO A.; TOSTO, D. S.; BERNASCONI, P.; SALA, A.; WAGNER, M.; CORACH, D. (2006):
**Phylogenetic Relationships of *Deschampsia antarctica* (Poaceae): Insights from nuclear ribosomal ITS.**
In: Plant Systematics and Evolution 261 (1-4), p. 1–9. DOI: 10.1007/s00606-006-0425-x.

SUN, GEN-LOU; POURKHEIRANDISH, MOHAMMAD; KOMATSUDA, TAKAO (2009):
**Molecular Evolution and Phylogeny of the Rpb2 Gene in the Genus *Hordeum*.**
In: Annals of Botany 103 (6), p. 975–983. DOI: 10.1093/aob/mcp020.

SUN, QIAN; LIN, Q. I.; YI, ZI-LI; YANG, ZHI-RONG; ZHOU, FA-SONG (2010):
**A taxonomic Revision of *Miscanthus* s.l. (Poaceae) from China.**
In: Botanical Journal of the Linnean Society 164 (2), p. 178–220. DOI: 10.1111/j.1095-8339.2010.01082.x.

SUN, YE; XIA, NIAN-HE; LIN, RUSHUN (2005):
**Phylogenetic Analysis of *Bambusa* (Poaceae: Bambusoideae) based on Internal Transcribed Spacer Sequences of nuclear ribosomal DNA.**
In: Biochemical Genetics 43 (11-12), p. 603–612. DOI: 10.1007/s10528-005-9117-4.

SYME, ANNA E.; MURPHY, DANIEL J.; HOLMES, GARETH D.; GARDNER, STUART; FOWLER, RACHAEL; CANTRILL, DAVID J. (2012):
**An expanded phylogenetic Analysis of *Austrostipa* (Poaceae: Stipeae) to test infrageneric Relationships.**
In: Australian Systematic Botany 25 (1), p. 1–10. DOI: 10.1071/SB10049.

TANG, LIANG; ZOU, XIN-HUI; ZHANG, LIN-BIN; GE, SONG (2015):
**Multilocus Species Tree Analyses resolve the ancient Radiation of the Subtribe Zizaniinae (Poaceae).**
In: Molecular Phylogenetics and Evolution 84, p. 232–239. DOI: 10.1016/j.ympev.2015.01.011.

TEERAWATANANON, ATCHARA; JACOBS, SURREY W.L.; HODKINSON, TREVOR R. (2009):
**Phylogenetics of Panicoideae (Poaceae) based on Chloroplast and nuclear DNA Sequences.**
In: Telopea 13 (1-2), p. 115–142.

TORRECILLA, PEDRO; LÓPEZ RODRÍGUEZ, JOSE ANGEL; STANCIK, D.; CATALÁN, PILAR (2003):

**Systematics of *Festuca* L. Sects. *Eskia* Willk., *Pseudatropis* Kriv., *Amphigenes* (Janka) Tzvel., *Pseudoscariosa* Kriv. and *Scariosae* Hack. based on Analysis of morphological Characters and DNA Sequences.**
In: Plant Systematics and Evolution 239 (1-2), p. 113–139. DOI: 10.1007/s00606-002-0265-2.

**Triplett, Jimmy K.; Clark, Lynn G. (2010):**
**Phylogeny of the Temperate Bamboos (Poaceae: Bambusoideae: Bambuseae) with an Emphasis on *Arundinaria* and Allies.**
In: Systematic Botany 35 (1), p. 102–120. DOI: 10.1600/036364410790862678.

**Tyrrell, Christopher D.; Santos-Gonçalves, Ana Paula; Londoño, Ximena; Clark, Lynn G. (2012):**
**Molecular Phylogeny of the arthrostylidioid Bamboos (Poaceae: Bambusoideae: Bambuseae: Arthrostylidiinae) and new Genus *Didymogonyx*.**
In: Molecular Phylogenetics and Evolution 65 (1), p. 136–148. DOI: 10.1016/j.ympev.2012.05.033.

**Verboom, George Anthony; Ntsohi, Refiloe; Barker, Nigel P. (2006):**
**Molecular Phylogeny of African *Rytidosperma* –affiliated danthonioid Grasses reveals generic Polyphyly and convergent Evolution in Spikelet Morphology.**
In: Taxon 55 (2), p. 337–348. DOI: 10.2307/25065581.

**Vorontsova, Maria S.; Simon, Bryan K. (2012):**
**Updating Classifications to reflect Monophyly: 10 to 20 percent of Species names Change in Poaceae.**
In: Taxon 51 (4), p. 735–746.

**Voshell, Stephanie M.; Baldini, Riccardo M.; Kumar, Rohit; Tatalovich, Nicholas; Hilu, Khidir W. (2011):**
**Canary Grasses (*Phalaris*, Poaceae): molecular Phylogenetics, polyploidy and floret Evolution.**
In: Taxon 60 (5), p. 1306–1316.

**Washburn, Jacob D.; Schnable, James C.; Davidse, Gerrit; Pires, J. Chris (2015):**
**Phylogeny and photosynthesis of the Grass Tribe Paniceae.**
In: American Journal of Botany 102 (9), p. 1493–1505. DOI: 10.3732/ajb.1500222.

**Welker, Cassiano A. D.; Souza-Chies, Tatiana Teixeira de; Longhi-Wagner, Hilda Maria; Peichoto, Myriam Carolina; McKain, Michael R.; Kellogg, Elizabeth A. (2016):**
**Multilocus Phylogeny and phylogenomics of *Eriochrysis* P. Beauv. (Poaceae-Andropogoneae): taxonomic Implications and Evidence of interspecific Hybridization.**
In: Molecular Phylogenetics and Evolution 99, p. 155–167. DOI: 10.1016/j.ympev.2016.02.022.

**Welker, Cassiano A. D.; Souza-Chies, Tatiana Teixeira de; Longhi-Wagner, Hilda Maria; Peichoto, Myriam Carolina; McKain, Michael R.; Kellogg, Elizabeth A. (2015):**
**Phylogenetic Analysis of *Saccharum* s.l. (Poaceae; Andropogoneae), with Emphasis on the Circumscription of the South American Species.**
In: American Journal of Botany 102 (2), p. 248–263. DOI: 10.3732/ajb.1400397.

**Whipple, Ian G.; Barkworth, Mary E.; Bushman, Bradley p. (2007):**
**Molecular Insights into the Taxonomy of *Glyceria* (Poaceae: Meliceae) in North America.**
In: American Journal of Botany 94 (4), p. 551–557. DOI: 10.3732/ajb.94.4.551.

**Winterfeld, Grit; Schneider, Julia; Becher, Hannes; Dickie, John; Röser, Martin (2015):**

**Karyosystematics of the Australasian stipoid grass *Austrostipa* and related Genera: Chromosome sizes, Ploidy, Chromosome Base Numbers and Phylogeny.**
In: Australian Systematic Botany 28 (3), p. 145–159. DOI: 10.1071/SB14029.

WÖLK, ALEXANDRA; RÖSER, MARTIN (2013):
**The new Genus *Trisetopsis* and new Combinations in oat-like Grasses (Poaceae).**
In: Schlechtendalia 25, p. 57–61.

WÖLK, ALEXANDRA; RÖSER, MARTIN (2014):
**Polyploid Evolution, intercontinental biogeographical Relationships and Morphology of the recently described African Oat Genus *Trisetopsis* (Poaceae).**
In: Taxon 63 (4), p. 773–788. DOI: 10.12705/634.1.

WYSOCKI, WILLIAM P.; CLARK, LYNN G.; ATTIGALA, LAKSHMI; RUIZ-SANCHEZ, EDUARDO; DUVALL, MELVIN R. (2015):
**Evolution of the Bamboos (Bambusoideae; Poaceae): a full plastome phylogenomic Analysis.**
In: BMC Evolutionary Biology 15, p. 50. DOI: 10.1186/s12862-015-0321-5.

XU, CHONG-MEI; QU, CHANG-YOU; YU, WEN-GUANG; ZHANG, XUE-JIE; LI, FA-ZENG (2009):
**Phylogenetic Origin of *Beckmannia* (Poaceae) inferred from molecular Evidence.**
In: Journal of Systematics and Evolution 47 (4), p. 305–310. DOI: 10.1111/j.1759-6831.2009.00032.x.

YAN, CHI; SUN, GEN-LOU (2011):
**Nucleotide Divergence and genetic Relationships of *Pseudoroegneria* Species.**
In: Biochemical Systematics and Ecology 39 (4-6), p. 309–319. DOI: 10.1016/j.bse.2011.08.009.

YANG, HAN-QI; PENG, SHENG; LI, DE-ZHU (2007):
**Generic Delimitations of *Schizostachyum* and its Allies (Gramineae: Bambusoideae) inferred from GBSSI and trnL-F Sequence Phylogenies.**
In: Taxon 56 (1), p. 45–54.

YANG, HAN-QI; YANG, JUN-BO; PENG, ZHEN-HUA; GAO, JIAN; YANG, YU-MING; PENG, SHENG; LI, DE-ZHU (2008):
**A molecular phylogenetic and Fruit evolutionary Analysis of the major Groups of the paleotropical woody Bamboos (Gramineae: Bambusoideae) based on nuclear ITS, GBSSI Gene and Plastid trnL-F DNA Sequences.**
In: Molecular Phylogenetics and Evolution 48 (3), p. 809–824. DOI: 10.1016/j.ympev.2008.06.001.

YANG, JUN-BO; YANG, HAN-QI; LI, DE-ZHU; WONG, KHOON MENG; YANG, YU-MING (2010):
**Phylogeny of *Bambusa* and its allies (Poaceae: Bambusoideae) inferred from nuclear GBSSI Gene and Plastid psbA-trnH, rpl32-trnL and rps16 Intron DNA Sequences.**
In: Taxon 59 (4), p. 1102–1110.

ZENG, CHUN-XIA; ZHANG, YU-XIAO; TRIPLETT, JIMMY K.; YANG, JUN-BO; LI, DE-ZHU (2010):
**Large multi-locus Plastid Phylogeny of the Tribe Arundinarieae (Poaceae: Bambusoideae) reveals ten major Lineages and low Rate of molecular Divergence.**
In: Molecular Phylogenetics and Evolution 56 (2), p. 821–839. DOI: 10.1016/j.ympev.2010.03.041.

ZHANG, H-Q; FAN, X.; SHA, L-N; ZHANG, C.; YANG, R-W; ZHOU, Y-H (2008):
**Phylogeny of *Hystrix* and related Genera (Poaceae: Triticeae) based on nuclear rDNA ITS Sequences.**
In: Plant Biology 10 (5), p. 635–642. DOI: 10.1111/j.1438-8677.2008.00065.x.

ZHANG, XIAN-ZHI; ZENG, CHUN-XIA; MA, PENG-FEI; HAEVERMANS, THOMAS; ZHANG, YU-XIAO; ZHANG, LI-NA ET AL. (2016):

**Multi-locus Plastid phylogenetic Biogeography supports the Asian Hypothesis of the temperate woody Bamboos (Poaceae: Bambusoideae).**
In: Molecular Phylogenetics and Evolution 96, p. 118–129. DOI: 10.1016/j.ympev.2015.11.025.

**Zhang, Yu-Xiao; Zeng, Chun-Xia; Li, De-Zhu (2012):**
**Complex Evolution in Arundinarieae (Poaceae: Bambusoideae): incongruence between Plastid and nuclear GBSSI Gene Phylogenies.**
In: Molecular Phylogenetics and Evolution 63 (3), p. 777–797. DOI: 10.1016/j.ympev.2012.02.023.

**Zheng, Chaohan; Xia, Nian-He; Deng, Yun-Fei (2013):**
***Ampelocalamus stoloniformis*, a new Combination for Chinese Bambusoideae (Poaceae).**
In: Phytotaxa 135 (1), p. 62. DOI: 10.11646/phytotaxa.135.1.7.

**Zimmermann, Tanja; Bocksberger, Gaëlle; Brüggemann, Wolfgang; Berberich, Thomas (2013):**
**Phylogenetic Relationship and molecular Taxonomy of African Grasses of the Genus *Panicum* inferred from four Chloroplast DNA-Barcodes and nuclear Gene Sequences.**
In: Journal of Plant Research 126 (3), p. 363–371. DOI: 10.1007/s10265-012-0538-y.

**Zuloaga, Fernando Omar; Morrone, Osvaldo; Davidse, Gerrit; Pennington, Susan J. (2007):**
**Classification and Biogeography of Panicoideae (Poaceae) in the New World.**
In: Aliso 23 (1), p. 503–529.

**Zuloaga, Fernando Omar; Salariato, Diego Leonel; Scataglini, Maria Amalia (2018):**
**Molecular Phylogeny of *Panicum* s. str. (Poaceae, Panicoideae, Paniceae) and Insights into its Biogeography and Evolution.**
In: Public Library of Science One 13 (2), e0191529. DOI: 10.1371/journal.pone.0191529.

**Zuloaga, Fernando Omar; Salomón, Luciana; Scataglini, Maria Amalia (2015):**
**Phylogeny of Sections *Clavelligerae* and *Pectinatae* of *Panicum* (Poaceae, Panicoideae, Paniceae): establishment of the new Subtribe Dichantheliinae and the Genus *Adenochloa*.**
In: Plant Systematics and Evolution 301 (6), p. 1693–1711. DOI: 10.1007/s00606-014-1186-6.

**Zuloaga, Fernando Omar; Scataglini, Maria Amalia; Morrone, Osvaldo (2010):**
**A phylogenetic Evaluation of *Panicum* Sects. *Agrostoidea, Megista, Prionitia* and *Tenera* (Panicoideae, Poaceae): Two new Genera, *Stephostachys* and *Sorengia*.**
In: Taxon 59 (5), p. 1535–1546.

### Podocarpaceae

**Herbert, Jane; Hollingsworth, Peter M.; Gardner, Martin F.; Mill, Robert R.; Thomas, Philip I.; Jaffré, Tanguy (2002):**
**Conservation genetics and Phylogenetics of New Caledonian *Retrophyllum* (Podocarpaceae) Species.**
In: New Zealand Journal of Botany 40 (2), p. 175–188. DOI: 10.1080/0028825X.2002.9512781.

**Kelch, Dean G. (1998):**
**Phylogeny of Podocarpaceae: Comparison of Evidence from Morphology and 18s Rdna1.**
In: American Journal of Botany 85 (7), p. 986–996.

**Kelch, Dean G. (2002):**

**Phylogenetic assessment of the monotypic Genera *Sundacarpus* and *Manoao* (Coniferales: Podocarpaceae) utilising Evidence from 18s rDNA Sequences.**
In: Australian Systematic Botany 15 (1), p. 29–35. DOI: 10.1071/SB01002.

**KNOPF, PATRICK; SCHULZ, CHRISTIAN; LITTLE, DAMON P.; STÜTZEL, THOMAS; STEVENSON, DENNIS WM. (2012):**
**Relationships within Podocarpaceae based on DNA Sequence, anatomical, morphological, and biogeographical Data.**
In: Cladistics 28 (3), p. 271–299. DOI: 10.1111/j.1096-0031.2011.00381.x.

**MILL, ROBERT R. (2015):**
**A Monographic Revision of the Genus *Podocarpus* (Podocarpaceae): II. the Species of the Caribbean Bioregion.**
In: Edinburgh Journal of Botany 72 (1), p. 61–185. DOI: 10.1017/S0960428614000328.

**SINCLAIR, W. T.; MILL, ROBERT R.; GARDNER, MARTIN F.; WOLTZ, P.; JAFFRÉ, TANGUY; PRESTON, JILLIAN ET AL. (2002):**
**Evolutionary Relationships of the New Caledonian heterotrophic Conifer, *Parasitaxus usta* (Podocarpaceae), inferred from Chloroplast trn L-f Intron/spacer and nuclear rDNA ITS2 Sequences.**
In: Plant Systematics and Evolution 233 (1-2), p. 79–104. DOI: 10.1007/s00606-002-0199-8.

### Podostemaceae

**FUJINAMI, RIEKO; IMAICHI, RYOKO (2015):**
**Developmental Morphology of flattened shoots in *Dalzellia ubonensis* and *Indodalzellia* gracilis with Implications for the Evolution of diversified shoot morphologies in the Subfamily Tristichoideae (Podostemaceae).**
In: American Journal of Botany 102 (6), p. 848–859. DOI: 10.3732/ajb.1500206.

**KATO, MASAHIRO (2016):**
**Multidisciplinary studies of the Diversity and Evolution in River-weeds.**
In: Journal of Plant Research 129 (3), p. 397–410. DOI: 10.1007/s10265-016-0801-8.

**KATO, MASAHIRO; KITA, YOKO; KOI, SATOSHI (2003):**
**Molecular Phylogeny, Taxonomy and Biogeography of *Malaccotristicha australis* comb. nov. (syn. *Tristicha australis*) (Podostemaceae).**
In: Australian Systematic Botany 16 (2), p. 177–183. DOI: 10.1071/SB02020.

**KELLY, LAURA J.; AMEKA, GABRIEL K.; CHASE, MARK W. (2010):**
**DNA barcoding of African Podostemaceae (river-weeds): A test of proposed Barcode Regions.**
In: Taxon 59 (1), p. 251–260. DOI: 10.1002/tax.591023.

**KHANDURI, PRIYANKA; TANDON, RAJESH; UNIYAL, PREM LAL; BHAT, VISHNU; PANDEY, ARUN KUMAR (2015):**
**Comparative Morphology and molecular Systematics of Indian Podostemaceae.**
In: Plant Systematics and Evolution 301 (3), p. 861–882. DOI: 10.1007/s00606-014-1121-x.

**KITA, YOKO; KATO, M. (2001):**
**Infrafamilial Phylogeny of the aquatic Angiosperm Podostemaceae inferred from the Nucleotide Sequences of the matK Gene.**
In: Plant Biology 3 (2), p. 156–163. DOI: 10.1055/s-2001-12895.

**KOI, SATOSHI; KATO, MASAHIRO (2010):**
**Developmental Morphology of Seedling and Shoot and phylogenetic Relationship of *Diplobryum koyamae* (Podostemaceae).**
In: American Journal of Botany 97 (3), p. 373–387. DOI: 10.3732/ajb.0900157.

**Koi, Satoshi; Kato, Masahiro (2012):**
**A taxonomic Study of Podostemaceae Subfamily Podostemoideae of Laos with phylogenetic Analyses of *Cladopus, Paracladopus* and *Polypleurum*.**
In: Kew Bulletin 67 (3), p. 331–365. DOI: 10.1007/s12225-012-9399-8.

**Koi, Satoshi; Kita, Yoko; Hirayama, Yumiko; Rutishauser, Rolf; Huber, Konrad A.; Kato, Masahiro (2012):**
**Molecular phylogenetic Analysis of Podostemaceae: Implications for Taxonomy of major Groups.**
In: Botanical Journal of the Linnean Society 169 (3), p. 461–492. DOI: 10.1111/j.1095-8339.2012.01258.x.

**Koi, Satoshi; Kita, Yoko; Kato, Masahiro (2008):**
***Paracladopus chanthaburiensis*, a new Species of Podostemaceae from Thailand, with Notes on its Morphology, Phylogeny and Distribution.**
In: Taxon 57 (1), p. 201–210. DOI: 10.6027/9789289333788-4-en.

**Koi, Satoshi; Rutishauser, Rolf; Kato, Masahiro (2009):**
**Phylogenetic Relationship and Morphology of *Dalzellia gracilis* (Podostemaceae, Subfamily Tristichoideae) with Proposal of a new Genus.**
In: International Journal of Plant Sciences 170 (2), p. 237–246. DOI: 10.1086/595292.

**Moline, Philip M.; Thiv, Mike; Ameka, G. K.; Ghogue, Jean-Paul; Pfeifer, Evelin; Rutishauser, Rolf (2007):**
**Comparative Morphology and molecular Systematics of African Podostemaceae-Podostemoideae, with Emphasis on *Dicraeanthus* and *Ledermanniella* from Cameroon.**
In: International Journal of Plant Sciences 168 (2), p. 159–180.

**Pfeifer, Evelin; Grob, Valentin; Thiv, Mike; Rutishauser, Rolf (2009):**
***Stonesia ghoguei*, peculiar Morphology of a new Cameroonian Species (Podostemaceae, Podostemoideae).**
In: Novon: A Journal for Botanical Nomenclature 19 (1), p. 102–116. DOI: 10.3417/2007080.

**Philbrick, C. Thomas; Bove, Claudia P.; Edson Jr., Thomas C. (2009):**
**Monograph of *Castelnavia* (Podostemaceae).**
In: Systematic Botany 34 (4), p. 715–729. DOI: 10.1600/036364409790139781.

**Philbrick, C. Thomas; Bove, Claudia P.; Stevens, Hannah I. (2010):**
**Endemism in Neotropical Podostemaceae.**
In: Annals of the Missouri Botanical Garden 97 (3), p. 425–456. DOI: 10.3417/2008087.

**Soltis, Douglas E.; Mort, Mark E.; Soltis, Pamela S.; Hibsch-Jetter, Carola; Zimmer, Elizabeth A.; Morgan, D. R. (1999):**
**Phylogenetic Relationships of the enigmatic Angiosperm Family Podostemaceae inferred from 18s rDNA and rbcL Sequence Data.**
In: Molecular Phylogenetics and Evolution 11 (2), p. 261–272. DOI: 10.1006/mpev.1998.0577.

**Thiv, Mike; Ghogue, Jean-Paul; Grob, Valentin; Huber, Konrad A.; Pfeifer, Evelin; Rutishauser, Rolf (2009):**
**How to get off the Mismatch at the generic Rank in African Podostemaceae?**
In: Plant Systematics and Evolution 283 (1-2), p. 57–77. DOI: 10.1007/s00606-009-0214-4.

**Tippery, Nicholas P.; Philbrick, C. Thomas; Bove, Claudia P.; Les, Donald H. (2011):**
**Systematics and Phylogeny of Neotropical Riverweeds (Podostemaceae: Podostemoideae).**
In: Systematic Botany 36 (1), p. 105–118. DOI: 10.1600/036364411X553180.

### Polemoniaceae

**Bell, Charles D.; Patterson, Robert W. (2000):**
**Molecular Phylogeny and Biogeography of *Linanthus* (Polemoniaceae).**
In: American Journal of Botany 87 (12), p. 1857–1870. DOI: 10.2307/2656838.

**Ferguson, Carolyn J.; Jansen, Robert K. (2002):**
**A Chloroplast DNA Phylogeny of eastern *Phlox* (Polemoniaceae): Implications of Congruence and Incongruence with the ITS Phylogeny.**
In: American Journal of Botany 89 (8), p. 1324–1335. DOI: 10.3732/ajb.89.8.1324.

**Goodwillie, Carol (1999):**
**Multiple Origins of Self-compatibility in *Linanthus* Section *Leptosiphon* (Polemoniaceae): phylogenetic Evidence from Internal-transcribed-spacer Sequence Data.**
In: Evolution 53 (5), p. 1387–1395.

**Johnson, Leigh A.; Chan, Lauren M.; Weese, Terri L.; Busby, Lisa D.; McMurry, Samuel (2008):**
**Nuclear and cpDNA Sequences combined provide strong Inference of higher phylogenetic Relationships in the Phlox Family (Polemoniaceae).**
In: Molecular Phylogenetics and Evolution 48 (3), p. 997–1012. DOI: 10.1016/j.ympev.2008.05.036.

**Johnson, Leigh A.; Soltis, Douglas E. (1995):**
**Phylogenetic Inference in Saxifragaceae sensu stricto and *Gilia* (Polemoniaceae) using matK Sequences.**
In: Annals of the Missouri Botanical Garden 82 (2), p. 149–175.

**Monfils, Anna K.; Prather, L. Alan (2010):**
**Phylogeny of *Cantua* (Polemoniaceae): Evidence from Chloroplast and Nuclear DNA Sequence Data.**
In: Systematic Botany 35 (4), p. 877–884. DOI: 10.1600/036364410X539925.

**Porter, J. Mark; Johnson, Leigh A.; Wilken, Dieter H. (2010):**
**Phylogenetic Systematics of *Ipomopsis* (Polemoniaceae): Relationships and Divergence Times estimated from Chloroplast and Nuclear DNA Sequences.**
In: Systematic Botany 35 (1), p. 181–200. DOI: 10.1600/036364410790862542.

**Prather, L. Alan; Ferguson, Carolyn J.; Jansen, Robert K. (2000):**
**Polemoniaceae Phylogeny and Classification: Implications of Sequence Data from the Chloroplast Gene ndhF.**
In: American Journal of Botany 87 (9), p. 1300–1308. DOI: 10.2307/2656723.

**Weese, Terri L.; Johnson, Leigh A. (2005):**
**Utility of NADP-dependent Isocitrate Dehydrogenase for species-level evolutionary Inference in Angiosperm Phylogeny: a case Study in *Saltugilia*.**
In: Molecular Phylogenetics and Evolution 36 (1), p. 24–41. DOI: 10.1016/j.ympev.2005.03.004.

**Worley, Anne C.; Ghazvini, Habibollah; Schemske, Douglas W. (2009):**
**A Phylogeny of the Genus *Polemonium* based on Amplified Fragment Length Polymorphism (Aflp) Markers.**
In: Systematic Botany 34 (1), p. 149–161. DOI: 10.1600/036364409787602267.

### Polygalaceae

**Abbott, Richard J.; Pastore, José Floriano B. (2015):**
**Preliminary Synopsis of the Genus *Hebecarpa* (Polygalaceae).**
In: Kew Bulletin 70 (3), p. 125. DOI: 10.1007/S12225-015-9589-2.

**Forest, Félix; Manning, John C. (2006):**
**Evidence for Inclusion of South African Endemic *Nylandtia* in *Muraltia* (Polygalaceae).**
In: Systematic Botany 31 (3), p. 525–532.

**Forest, Félix; Nänni, Ingrid; Chase, Mark W.; Crane, Peter R.; Hawkins, Julie A. (2007):**
**Diversification of a large Genus in a continental Biodiversity Hotspot: temporal and spatial Origin of *Muraltia* (Polygalaceae) in the Cape of South Africa.**
In: Molecular Phylogenetics and Evolution 43 (1), p. 60–74. DOI: 10.1016/j.ympev.2006.08.017.

**Kerrigan, Raelee A. (2012):**
**A Treatment for *Polygala* of northern Australia.**
In: Australian Systematic Botany 25 (2), p. 83–137. DOI: 10.1071/SB08032.

**Mennes, Constantijn B.; Moerland, Michelangelo S.; Rath, Magnus; Smets, Erik F.; Merckx, Vincent S.F.T. (2015):**
**Evolution of Mycoheterotrophy in Polygalaceae: the Case of *Epirixanthes*.**
In: American Journal of Botany 102 (4), p. 598–608. DOI: 10.3732/ajb.1400549.

**Pastore, José Floriano B. (2012):**
***Caamembeca*: generic Status and new name for *Polygala* Subgenus *Ligustrina* (Polygalaceae).**
In: Kew Bulletin 67 (3), p. 435–442. DOI: 10.1007/s12225-012-9360-x.

**Pastore, José Floriano B. (2016):**
***Acanthocladus dichromus*, a new Combination for a South American Polygalaceae.**
In: Phytotaxa 286 (1), p. 54. DOI: 10.11646/phytotaxa.286.1.7.

**Pastore, José Floriano B.; Abbott, Richard J. (2012):**
**Taxonomic Notes and new Combinations for *Asemeia* (Polygalaceae).**
In: Kew Bulletin 67 (4), p. 801–813. DOI: 10.1007/s12225-012-9397-x.

**Pastore, José Floriano B.; Abbott, Richard J.; Neubig, Kurt M.; Whitten, William Mark; Mascarenhas, Renata B.; Mota, Michelle Christine Almeida; van den Berg, Cássio (2017):**
**A molecular Phylogeny and taxonomic Notes in *Caamembeca* (Polygalaceae).**
In: Systematic Botany 42 (1), p. 54–62. DOI: 10.1600/036364417X694935.

### Polygonaceae

**Burke, Janelle M.; Sanchez, Adriana (2011):**
**Revised Subfamily Classification for Polygonaceae, with a tribal Classification for Eriogonoideae.**
In: Brittonia 63 (4), p. 510–520. DOI: 10.1007/s12228-011-9197-x.

**Burke, Janelle M.; Sanchez, Adriana; Kron, Kathleen A.; Luckow, Melissa (2010):**
**Placing the woody tropical Genera of Polygonaceae: A Hypothesis of Character Evolution and Phylogeny.**
In: American Journal of Botany 97 (8), p. 1377–1390. DOI: 10.3732/ajb.1000022.

**Fan, Deng-Mei; Chen, Jia-Hui; Meng, Ying; Wen, Jun; Huang, Jin-Ling; Yang, Yong-Ping (2013):**
**Molecular Phylogeny of *Koenigia* L. (Polygonaceae: Persicarieae): Implications for Classification, Character Evolution and Biogeography.**
In: Molecular Phylogenetics and Evolution 69 (3), p. 1093–1100. DOI: 10.1016/j.ympev.2013.08.018.

**Galasso, Gabriele; Banfi, Enrico; Mattia, Fabrizio de; Grassi, Fabrizio; Sgorbati, Sergio; Labra, Massimo (2009):**

**Molecular Phylogeny of *Polygonum* L. s.l. (Polygonoideae, Polygonaceae), focusing on European taxa: preliminary Results and systematic Considerations based on rbcL plastidial Sequence Data.**
In: Atti della Societa Italiana di Scienze Naturali e del Museo Civico di Storia Naturale di Milano 150 (1), p. 113–148.

**Kempton, Elizabeth A. (2012):**
**Systematics of Eriogonoideae s. s. (Polygonaceae).**
In: Systematic Botany 37 (3), p. 723–737. DOI: 10.1600/036364412X648698.

**Kim, Sang-Tae; Donoghue, Michael J. (2008):**
**Incongruence between cpDNA and nrITS Trees indicates extensive Hybridization within *Eupersicaria* (Polygonaceae).**
In: American Journal of Botany 95 (9), p. 1122–1135. DOI: 10.3732/ajb.0700008.

**Kim, Sang-Tae; Donoghue, Michael J. (2008):**
**Molecular Phylogeny of *Persicaria* (Persicarieae, Polygonaceae).**
In: Systematic Botany 33 (1), p. 77–86. DOI: 10.1600/036364408783887302.

**Li, Fen-Long; Zeller, Friedrich J.; Huang, Kai-Feng; Shi, Tao-Xiong; Chen, Qing-Fu (2013):**
**Improvement of fluorescent Chromosome in situ PCR and its application in the Phylogeny of the Genus *Fagopyrum* Mill. using nuclear Genes of Chloroplast Origin (cpDNA).**
In: Plant Systematics and Evolution 299 (9), p. 1679–1691. DOI: 10.1007/s00606-013-0825-7.

**Ohsako, Takanori; Ohnishi, Ohmi (2000):**
**Intra- and interspecific Phylogeny of wild *Fagopyrum* (Polygonaceae) Species based on nucleotide Sequences of noncoding Regions in Chloroplast DNA.**
In: American Journal of Botany 87 (4), p. 573–582. DOI: 10.2307/2656601.

**Sanchez, Adriana; Kron, Kathleen A. (2008):**
**Phylogenetics of Polygonaceae with an Emphasis on the Evolution of Eriogonoideae.**
In: Systematic Botany 33 (1), p. 87–96. DOI: 10.1600/036364408783887456.

**Sanchez, Adriana; Kron, Kathleen A. (2009):**
**Phylogenetic Relationships of *Afrobrunnichia* Hutch. & Dalziel (Polygonaceae) based on three Chloroplast Genes and ITS.**
In: Taxon 58 (3), p. 781–792.

**Sanchez, Adriana; Kron, Kathleen A. (2011):**
**Phylogenetic Relationships of *Triplaris* and *Ruprechtia*: Re-Delimitation of the recognized Genera and two new Genera for Tribe Triplarideae (Polygonaceae).**
In: Systematic Botany 36 (3), p. 702–710. DOI: 10.1600/036364411X583664.

**Sanchez, Adriana; Schuster, Tanja M.; Burke, Janelle M.; Kron, Kathleen A. (2011):**
**Taxonomy of Polygonoideae (Polygonaceae): A new tribal Classification.**
In: Taxon 60 (1), p. 151–160.

**Sanchez, Adriana; Schuster, Tanja M.; Kron, Kathleen A. (2009):**
**A Large-Scale Phylogeny of Polygonaceae based on molecular Data.**
In: International Journal of Plant Sciences 170 (8), p. 1044–1055. DOI: 10.1086/605121.

**Schuster, Tanja M.; Reveal, James L.; Bayly, Michael J.; Kron, Kathleen A. (2015):**
**An updated molecular Phylogeny of Polygonoideae (Polygonaceae): Relationships of *Oxygonum, Pteroxygonum and Rumex*, and a new Circumscription of *Koenigia*.**

In: Taxon 64 (6), p. 1188–1208. DOI: 10.12705/646.5.

SCHUSTER, TANJA M.; REVEAL, JAMES L.; KRON, KATHLEEN A. (2011):
**Phylogeny of Polygoneae (Polygonaceae: Polygonoideae).**
In: Taxon 60 (6), p. 1653–1666.

SCHUSTER, TANJA M.; WILSON, KAREN L.; KRON, KATHLEEN A. (2011):
**Phylogenetic Relationships of *Muehlenbeckia, Fallopia,* and *Reynoutria* (Polygonaceae) Investigated with Chloroplast and Nuclear Sequence Data.**
In: International Journal of Plant Sciences 172 (8), p. 1053–1066. DOI: 10.1086/661293.

SUN, WEI; ZHONG-ZE; MING-ZHEN, ZHOU; HE-WEN, LIU; DONG, WAN XIANG (2008):
**Reappraisal of the generic Status of *Pteroxygonum* (Polygonaceae) on the Basis of Morphology, Anatomy and nrDNA ITS Sequence Analysis.**
In: Journal of Systematics and Evolution 46 (1), p. 73–79.

SUN, YONGSHUAI; WANG, AILAN; WAN, DONGSHI; WANG, QIAN; LIU, JIAN-QUAN (2012):
**Rapid Radiation of *Rheum* (Polygonaceae) and parallel Evolution of morphological Traits.**
In: Molecular Phylogenetics and Evolution 63 (1), p. 150–158. DOI: 10.1016/j.ympev.2012.01.002.

TAVAKKOLI, SOLMAZ; KAZEMPOUR-OSALOO, SHAHROKH; MOZAFFARIAN, VALIOLLAH; MAASSOUMI, ALI AASGHAR (2015):
**Molecular Phylogeny of *Atraphaxis* and the woody *Polygonum* Species (Polygonaceae): Taxonomic Implications based on molecular and morphological Evidence.**
In: Plant Systematics and Evolution 301 (4), p. 1157–1170. DOI: 10.1007/s00606-014-1140-7.

TAVAKKOLI, SOLMAZ; OSALOO, SHAHROKH KAZEMPOUR; MAASSOUMI, ALI AASGHAR (2010):
**The Phylogeny of *Calligonum* and *Pteropyrum* (Polygonaceae) based on nuclear ribosomal DNA ITS and Chloroplast trnL-F Sequences.**
In: Iranian Journal of Biotechnology 8 (1), p. 7–15.

TIAN, XINMIN; LUO, JIAN; WANG, AILAN; MAO, KANG-SHAN; LIU, JIAN-QUAN (2011):
**On the Origin of the woody buckwheat *Fagopyrum tibeticum* (=*Parapteropyrum tibeticum*) in the Qinghai-Tibetan Plateau.**
In: Molecular Phylogenetics and Evolution 61 (2), p. 515–520. DOI: 10.1016/j.ympev.2011.07.001.

VYSOCHINA, G. (2014):
**Phenolic compounds in Systematics and Phylogeny of the Family Polygonaceae Juss. Vi. Genus *Knorringia* (Chukav.) Tzvel.**
In: Turczaninowia 17 (1), p. 33–41. DOI: 10.14258/turczaninowia.17.1.4.

WANG, AILAN; YANG, MEIHUA; LIU, JIAN-QUAN (2005):
**Molecular Phylogeny, recent Radiation and Evolution of gross Morphology of the rhubarb Genus *Rheum* (Polygonaceae) inferred from Chloroplast DNA trnL-F Sequences.**
In: Annals of Botany 96 (3), p. 489–498. DOI: 10.1093/aob/mci201.

ZHOU, MEI-LIANG; WANG, CHENG-LONG; WANG, DE-ZHOU; ZHENG, YA-DI; LI, FA-LIANG; ZHU, XUE-MEI ET AL. (2014):
**Phylogenetic Relationship of four new Species related to southwestern Sichuan *Fagopyrum* based on morphological and molecular Characterization.**
In: Biochemical Systematics and Ecology 57, p. 403–409. DOI: 10.1016/j.bse.2014.09.024.

### Polypodiaceae

**Bauret, Lucie; Gaudeul, Myriam; Sundue, Michael A.; Parris, Barbara S.; Ranker, Tom A.; Rakotondrainibe, France et al. (2017):**
**Madagascar sheds new light on the molecular Systematics and Biogeography of Grammitid Ferns: new unexpected Lineages and numerous long-distance Dispersal Events.**
In: Molecular Phylogenetics and Evolution 111, p. 1–17. DOI: 10.1016/j.ympev.2017.03.005.

**Brownsey, Patrick J.; Perrie, Leon R. (Decmber 2014):**
**Polypodiaceae (excluding *Notogrammitis*).**
In: Breitwieser I., Heenan P., Wilton A. (eds) Flora of New Zealand – Seed Plants. Fascicle 1. Manaaki Whenua Press, Lincoln. With contribution of Landcare Research NZ Ltd.

**Fls, Harald Schneider; Kreier, Hans-Peter; Hovenkamp, Peter H.; Janssen, Thomas (2008):**
**Phylogenetic Relationships of the Fern Genus *Christiopteris* shed new light onto the Classification and Biogeography of Drynarioid Ferns.**
In: Botanical Journal of the Linnean Society 157, p. 645–656.

**Hirai, Regina Y.; Rouhan, Germinal; Labiak, Paulo Henrique; Ranker, Tom A.; Prado, Jefferson (2011):**
**Moranopteris: A new Neotropical Genus of Grammitid Ferns (Polypodiaceae) segregated from Asian *Micropolypodium*.**
In: Taxon 60 (4), p. 1123–1137.

**Kim, Changkyun; Zha, Hong-Guang; Deng, Tao; Sun, Hang; Wu, Su-Gong (2013):**
**Phylogenetic Position of *Kontumia* (Polypodiaceae) inferred from four Chloroplast DNA Regions.**
In: Journal of Systematics and Evolution 51 (2), p. 154–163. DOI: 10.1111/j.1759-6831.2012.00230.x.

**Kreier, Hans-Peter; Rex, Martina; Weising, Kurt; Kessler, Michael; Smith, Alan R.; Schneider, Harald (2008):**
**Inferring the Diversification of the epiphytic Fern Genus *Serpocaulon* (Polypodiaceae) in South America using Chloroplast Sequences and Amplified Fragment Length Polymorphisms.**
In: Plant Systematics and Evolution 274 (1-2), p. 1–16. DOI: 10.1007/s00606-008-0021-3.

**Kreier, Hans-Peter; Schneider, Harald (2006):**
**Phylogeny and Biogeography of the Staghorn Fern Genus *Platycerium* (Polypodiaceae, Polypodiidae).**
In: American Journal of Botany 93 (2), p. 217–225.

**Kreier, Hans-Peter; Schneider, Harald (2006):**
**Reinstatement of *Loxogramme dictyopteris*, based on phylogenetic Evidence, for the New Zealand endemic Fern, *Anarthropteris lanceolata* (Polypodiaceae, Polypodiidae).**
In: Australian Systematic Botany 19 (4), p. 309–314. DOI: 10.1071/SB05033.

**Kreier, Hans-Peter; Zhang, Xian-Chun; Muth, Heiko; Schneider, Harald (2008):**
**The Microsoroid Ferns: Inferring the Relationships of a highly diverse Lineage of Paleotropical epiphytic Ferns (Polypodiaceae, Polypodiopsida).**
In: Molecular Phylogenetics and Evolution 48 (3), p. 1155–1167. DOI: 10.1016/j.ympev.2008.05.001.

**Labiak, Paulo Henrique; Rouhan, Germinal; Sundue, Michael A. (2010):**
**Phylogeny and Taxonomy of *Leucotrichum* (Polypodiaceae): A new Genus of Grammitid Ferns from the Neotropics.**
In: Taxon 59 (3), p. 911–921.

LABIAK, PAULO HENRIQUE; SUNDUE, MICHAEL A.; ROUHAN, GERMINAL (2010):
**Molecular Phylogeny, Character Evolution, and Biogeography of the Grammitid Fern Genus *Lellingeria* (Polypodiaceae).**
In: American Journal of Botany 97 (8), p. 1354–1364. DOI: 10.3732/ajb.0900393.

LEHNERT, MARCUS; KESSLER, MICHAEL; SCHMIDT-LEBUHN, ALEXANDER N.; KLIMAS, SUSAN A.; FEHLBERG, SHANNON D.; RANKER, TOM A. (2009):
**Phylogeny of the Fern Genus *Melpomene* (Polypodiaceae) inferred from Morphology and Chloroplast DNA Analysis.**
In: Systematic Botany 34 (1), p. 17–27. DOI: 10.1600/036364409787602276.

LINDSAY, STUART; MIDDLETON, DAVID J. (2009):
**New Combinations in the Ferns of Thailand.**
In: Edinburgh Journal of Botany 66 (2), p. 355–361. DOI: 10.1017/S096042860900568X.

OTTO, ELISABETH M.; JANSSEN, THOMAS; KREIER, HANS-PETER; SCHNEIDER, HARALD (2009):
**New Insights into the Phylogeny of *Pleopeltis* and related Neotropical Genera (Polypodiaceae, Polypodiopsida).**
In: Molecular Phylogenetics and Evolution 53 (1), p. 190–201. DOI: 10.1016/j.ympev.2009.05.001.

PERRIE, LEON R.; PARRIS, BARBARA P. (2012):
**Chloroplast DNA Sequences indicate the Grammitid Ferns (Polypodiaceae) in New Zealand belong to a single Clade, *Notogrammitis* gen. nov.**
In: New Zealand Journal of Botany 50 (4), p. 457–472. DOI: 10.1080/0028825X.2012.735247.

RANKER, TOM A.; GEIGER, JENNIFER M.O.; KENNEDY, S.C; SMITH, ALAN R.; HAUFLER, CHRISTOPHER H.; PARRIS, BARBARA P. (2003):
**Molecular Phylogenetics and Evolution of the endemic Hawaiian Genus *Adenophorus* (Grammitidaceae).**
In: Molecular Phylogenetics and Evolution 26 (3), p. 337–347. DOI: 10.1016/S1055-7903(02)00234-8.

RANKER, TOM A.; SMITH, ALAN R.; PARRIS, BARBARA S.; GEIGER, JENNIFER M.O.; HAUFLER, CHRISTOPHER H.; STRAUB, SHANNON C. K.; SCHNEIDER, HARALD (2004):
**Phylogeny and Evolution of Grammitid Ferns (Grammitidaceae): a case of rampant morphological Homoplasy.**
In: Taxon 53 (2), p. 415–428.

SALINO, ALEXANDRE; ALMEIDA, THAÍS ELIAS; SMITH, ALAN R.; GÓMEZ, ADRIANNA NAVARRO; KREIER, HANS-PETER; SCHNEIDER, HARALD (2008):
**A new Species of *Microgramma* (Polypodiaceae) from Brazil and Recircumscription of the Genus based on phylogenetic Evidence.**
In: Systematic Botany 33 (4), p. 630–635. DOI: 10.1600/036364408786500208.

SCHNEIDER, HARALD; JANSSEN, THOMAS; HOVENKAMP, PETER H.; SMITH, ALAN R.; CRANFILL, RAYMOND B.; HAUFLER, CHRISTOPHER H.; RANKER, TOM A. (2004):
**Phylogenetic Relationships of the Enigmatic Malesian Fern *Thylacopteris* (Polypodiaceae, Polypodiidae).**
In: International Journal of Plant Sciences 165 (6), p. 1077–1087.

SCHNEIDER, HARALD; KREIER, HANS-PETER; PERRIE, LEON R.; BROWNSEY, PATRICK J. (2006):
**The Relationships of *Microsorum* (Polypodiaceae) Species occurring in New Zealand.**
In: New Zealand Journal of Botany 44 (2), p. 121–127. DOI: 10.1080/0028825X.2006.9513012.

SCHNEIDER, HARALD; KREIER, HANS-PETER; WILSON, ROSEMARY; SMITH, ALAN R. (2006):

**The *Synammia* Enigma: Evidence for a Temperate Lineage of Polygrammoid Ferns (Polypodiaceae, Polypodiidae) in Southern South America.**
In: Systematic Botany 31 (1), p. 31–41.

SCHNEIDER, HARALD; SMITH, ALAN R.; CRANFILL, RAYMOND B.; HILDEBRAND, TERRI J.; HAUFLER, CHRISTOPHER H.; RANKER, TOM A. (2004):
**Unraveling the Phylogeny of Polygrammoid Ferns (Polypodiaceae and Grammitidaceae): exploring Aspects of the Diversification of epiphytic Plants.**
In: Molecular Phylogenetics and Evolution 31 (3), p. 1041–1063. DOI: 10.1016/j.ympev.2003.09.018.

SHAO, WEN; LU, SHU-GANG (2011):
***Himalayopteris*, a new Fern Genus from India and the adjacent Himalayas (Polypodiaceae, Polypodioideae).**
In: Novon: A Journal for Botanical Nomenclature 21 (1), p. 90–93. DOI: 10.3417/2009115.

SMITH, ALAN R.; KREIER, HANS-PETER; HAUFLER, CHRISTOPHER H.; RANKER, TOM A.; SCHNEIDER, HARALD (2006):
***Serpocaulon* (Polypodiaceae), a new Genus segregated from *Polypodium*.**
In: Taxon 55 (4), p. 919–930.

SPRUNT, SUSAN V.; SCHNEIDER, HARALD; WATSON, LINDA E.; RUSSELL, STEPHEN J.; NAVARRO-GOMEZ, ADRIANA; JAMES HICKEY, R. (2011):
**Exploring the molecular Phylogeny and Biogeography of *Pleopeltis polypodioides* (Polypodiaceae, Polypodiales) inferred from Plastid DNA Sequences.**
In: Systematic Botany 36 (4), p. 862–869. DOI: 10.1600/036364411X604886.

SUNDUE, MICHAEL A. (2010):
**A morphological Cladistic Analysis of *Terpsichore* (Polypodiaceae).**
In: Systematic Botany 35 (4), p. 716–729. DOI: 10.1600/036364410X539808.

SUNDUE, MICHAEL A.; ISLAM, MELISSA BAUER; RANKER, TOM A. (2010):
**Systematics of Grammitid Ferns (Polypodiaceae): using Morphology and Plastid Sequence Data to resolve the Circumscriptions of *Melpomene* and the Polyphyletic Genera *Lellingeria* and *Terpsichore*.**
In: Systematic Botany 35 (4), p. 701–715. DOI: 10.1600/036364410X539790.

SUNDUE, MICHAEL A.; PARRIS, BARBARA S.; RANKER, TOM A.; SMITH, ALAN R.; FUJIMOTO, ERIN L.; ZAMORA-CROSBY, DELIA ET AL. (2014):
**Global Phylogeny and Biogeography of Grammitid Ferns (Polypodiaceae).**
In: Molecular Phylogenetics and Evolution 81, p. 195–206. DOI: 10.1016/j.ympev.2014.08.017.

TESTO, WESTON; SUNDUE, MICHAEL A. (2014):
**Primary Hemiepiphytism in *Colysis ampla* (Polypodiaceae) provides new Insight into the Evolution of Growth Habit in Ferns.**
In: International Journal of Plant Sciences 175 (5), p. 526–536. DOI: 10.1086/675934.

WANG, FA-GUO; BARRATT, P. A.M.; WILFREDO, FALCÓN; FAY, MICHAEL F.; LEHTONEN, SAMULI; TUOMISTO, HANNA ET AL. (2014):
**On the Monophyly of Subfamily Tectarioideae (Polypodiaceae) and the phylogenetic Placement of some associated Fern Genera.**
In: Phytotaxa 164 (1), p. 1. DOI: 10.11646/phytotaxa.164.1.1.

WANG, LI; QI, XIN-PING; XIANG, QIAO-PING; HEINRICHS, JOCHEN; SCHNEIDER, HARALD; ZHANG, XIAN-CHUN (2010):
**Phylogeny of the paleotropical Fern Genus *Lepisorus* (Polypodiaceae, Polypodiopsida) inferred from four Chloroplast DNA Regions.**
In: Molecular Phylogenetics and Evolution 54 (1), p. 211–225. DOI: 10.1016/j.ympev.2009.08.032.

**Wang, Li; Wu, Zhi-qiang; Xiang, Qiao-Ping; Heinrichs, Jochen; Fls, Harald Schneider; Zhang, Xian-Chun (2010):**

**A molecular Phylogeny and a revised Classification of Tribe Lepisoreae (Polypodiaceae) based on an Analysis of four Plastid DNA Regions.**

In: Botanical Journal of the Linnean Society 162, p. 28–38.

**Xiao-Dong; Shu-Gang, Dong; Li, Lu Chun-Xiang (2008):**

**Molecular Phylogeny of *Colysis* (Polypodiaceae) based on Chloroplast rbcL and rps4-trnS Sequences.**

In: Journal of Systematics and Evolution 46 (5), p. 658–666.

**Zhou, Xin-Mao; Zhang, Liang; Chen, Cheng-Wei; Li, Chun-Xiang; Huang, Yao-Moan; Chen, De-Kui et al. (2017):**

**A Plastid Phylogeny and Character Evolution of the Old World Fern Genus *Pyrrosia* (Polypodiaceae) with the Description of a new Genus: *Hovenkampia* (Polypodiaceae).**

In: Molecular Phylogenetics and Evolution 114, p. 271–294. DOI: 10.1016/j.ympev.2017.06.020.

### Polypodiales

**Rothfels, Carl J.; Sundue, Michael A.; Kuo, Li-Yaung; Larsson, Anders; Kato, Masahiro; Schuettpelz, Eric; Pryer, Kathleen M. (2012):**

**A revised Family-level Classification for Eupolypod II Ferns (Polypodiidae: Polypodiales).**

In: Taxon 61 (3), p. 515–533.

### Pontederiaceae

**Ness, Rob W.; Graham, Sean W.; Barrett, Spencer C. H. (2011):**

**Reconciling Gene and Genome duplication events: using multiple nuclear Gene Families to infer the Phylogeny of the aquatic Plant Family Pontederiaceae.**

In: Molecular Biology and Evolution 28 (11), p. 3009–3018. DOI: 10.1093/molbev/msr119.

**Oliveira Pellegrini, Marco Octávio (2017):**

**Two new Synonyms in *Heteranthera* (Pontederiaceae, Commelinales).**

In: Nordic Journal of Botany 35 (1), p. 124–128. DOI: 10.1111/njb.01152.

### Portulacaceae

**Applequist, Wendy L.; Wallace, Robert p. (2001):**

**Phylogeny of the Portulacaceous Cohort based on ndhF Sequence Data.**

In: Systematic Botany 26 (2), p. 406–419.

**Hershkovitz, Mark A.; Zimmer, Elizabeth A. (2000):**

**Ribosomal DNA Evidence and Disjunctions of western American Portulacaceae.**

In: Molecular Phylogenetics and Evolution 15 (3), p. 419–439. DOI: 10.1006/mpev.1999.0720.

**Ocampo, Gilberto (2015):**

**Systematic Implications of Seed morphological Diversity in Portulacaceae (Caryophyllales).**

In: Plant Systematics and Evolution 301 (4), p. 1215–1226. DOI: 10.1007/s00606-014-1146-1.

**Ocampo, Gilberto; Columbus, Travis J. (2012):**

**Molecular Phylogenetics, historical Biogeography, and Chromosome Number Evolution of *Portulaca* (Portulacaceae).**

In: Molecular Phylogenetics and Evolution 63 (1), p. 97–112. DOI: 10.1016/j.ympev.2011.12.017.

### Posidoniaceae

AIRES, T.; MARBÀ, N.; CUNHA, R. L.; KENDRICK, G. A.; DI WALKER; SERRÃO, E. A. ET AL. (2011):
**Evolutionary History of the seagrass Genus *Posidonia*.**
In: Marine Ecology Progress Series 421, p. 117–130. DOI: 10.3354/meps08879.

### Potamogetonaceae

ITO, YU; TANAKA, NORIO; GARCÍA-MURILLO, PABLO; MUASYA, A. MUTHAMA (2016):
**A new Delimitation of the Afro-Eurasian plant Genus *Althenia* to include its Australasian relative, *Lepilaena* (Potamogetonaceae) - Evidence from DNA and morphological Data.**
In: Molecular Phylogenetics and Evolution 98, p. 261–270. DOI: 10.1016/j.ympev.2016.02.008.

LINDQVIST, CHARLOTTE; LAET, JAN; HAYNES, ROBERT R.; AAGESEN, LONE; KEENER, BRIAN R.; ALBERT, VICTOR A. (2006):
**Molecular Phylogenetics of an aquatic plant Lineage, Potamogetonaceae.**
In: Cladistics 22 (6), p. 568–588. DOI: 10.1111/j.1096-0031.2006.00124.x.

WANG, Q. D.; ZHANG, TIANZHEN; WANG, J. B. (2007):
**Phylogenetic Relationships and hybrid Origin of *Potamogeton* Species (Potamogetonaceae) distributed in China: Insights from the nuclear ribosomal Internal Transcribed Spacer Sequence (ITS).**
In: Plant Systematics and Evolution 267 (1-4), p. 65–78. DOI: 10.1007/s00606-006-0499-5.

### Primulaceae

BELLINO, ALESSANDRO; BELLINO, LEONARDO; BALDANTONI, DANIELA; SARACINO, ANTONIO (2015):
**Evolution, Ecology and Systematics of *Soldanella* (Primulaceae) in the southern Apennines (Italy).**
In: BMC Evolutionary Biology 15, p. 158. DOI: 10.1186/s12862-015-0433-y.

BOUCHER, FLORIAN C.; CASAZZA, GABRIELE; SZÖVÉNYI, P.; CONTI, ELENA (2016):
**Sequence capture using RAD probes clarifies phylogenetic Relationships and Species boundaries in *Primula* Sect. *Auricula*.**
In: Molecular Phylogenetics and Evolution 104, p. 60–72. DOI: 10.1016/j.ympev.2016.08.003.

BOUCHER, FLORIAN C.; THUILLER, WILFRIED; ROQUET, CRISTINA; DOUZET, ROLLAND; AUBERT, SERGE; ALVAREZ, NADIR; LAVERGNE, SÉBASTIEN (2012):
**Reconstructing the Origins of high-alpine Niches and Cushion Life form in the Genus *Androsace* s.l. (Primulaceae).**
In: Evolution 66 (4), p. 1255–1268. DOI: 10.1111/j.1558-5646.2011.01483.x.

GUGGISBERG, ALESSIA; MANSION, GUILHEM; CONTI, ELENA (2009):
**Disentangling reticulate Evolution in an Arctic-alpine polyploid Complex.**
In: Systematic Biology 58 (1), p. 55–73. DOI: 10.1093/sysbio/syp010.

GÜLTEPE, MUTLU; UZUNER, UĞUR; COŞKUNÇELEBI, KAMIL; BELDÜZ, ALI OSMAN; TERZÍOĞLU, SALIH (2010):
**Internal transcribed spacer (ITS) Polymorphism in the wild *Primula* (Primulaceae) Taxa of Turkey.**
In: Turkish Journal of Botany 34, p. 147–157.

KELSO, SYLVIA; BEARDSLEY, PAUL M.; WEITEMIER, KEVIN (2009):
**Phylogeny and Biogeography of *Primula* Sect. *Parryi* (Primulaceae).**
In: International Journal of Plant Sciences 170 (1), p. 93–106. DOI: 10.1086/593041.

KOVTONYUK, N. K.; GONCHAROV, A. A. (2009):

**Phylogenetic Relationships in the Genus *Primula* L. (Primulaceae) inferred from the ITS Region Sequences of nuclear rDNA.**
In: Russian Journal of Genetics 45 (6), p. 663–670. DOI: 10.1134/S1022795409060052.

LIU, YUN-JIAO; LIU, JUAN; HU, CHI-MING; HAO, GANG (2015):
**Non-Monophyly of *Primula* subgenera *Auganthus* and *Carolinella* (Primlaceae) as confirmed by the nuclear DNA Sequence Variation.**
In: Plant Systematics and Evolution 301 (8), p. 2057–2071. DOI: 10.1007/s00606-015-1207-0.

MAST, AUSTIN R.; FELLER, DANIELLE M.S.; KELSO, SYLVIA; CONTI, ELENA (2004):
**Buzz-pollinated *Dodecatheon* originated from within the heterostylous *Primula* Subgenus *Auriculastrum* (Primulaceae): A Seven-region CpDNA Phylogeny and ITS Implications for Floral Evolution.**
In: American Journal of Botany 91 (6), p. 926–942.

MAST, AUSTIN R.; KELSO, SYLVIA; JOHN RICHARDS, A.; LANG, DANIELA J.; FELLER, DANIELLE M.S.; CONTI, ELENA (2001):
**Phylogenetic Relationships in *Primula* L. and related Genera (Primulaceae) based on Noncoding Chloroplast DNA.**
In: International Journal of Plant Sciences 162 (6), p. 1381–1400.

MAST, AUSTIN R.; REVEAL, JAMES L. (2007):
**Transfer of *Dodecatheon* to *Primula* (Primulaceae).**
In: Brittonia 59 (1), p. 7982. DOI: 10.1663/0007-196X(2007)59[79:TODTPP]2.0.CO;2.

REN, GUANGPENG; CONTI, ELENA; SALAMIN, NICOLAS (2015):
**Phylogeny and Biogeography of *Primula* Sect. *Armerina*: Implications for plant Evolution under climate Change and the Uplift of the Qinghai-Tibet Plateau.**
In: BMC Evolutionary Biology 15, p. 161. DOI: 10.1186/s12862-015-0445-7.

SCHMIDT-LEBUHN, ALEXANDER N.; VOS, JURRIAAN M. DE; KELLER, BARBARA; CONTI, ELENA (2012):
**Phylogenetic Analysis of *Primula* Section *Primula* reveals rampant Non-Monophyly among morphologically distinct Species.**
In: Molecular Phylogenetics and Evolution 65 (1), p. 23–34. DOI: 10.1016/j.ympev.2012.05.015.

SCHNEEWEISS, GERALD M.; SCHÖNSWETTER, PETER; KELSO, SYLVIA; NIKLFELD, HARALD (2004):
**Complex biogeographic Patterns in *Androsace* (Primulaceae) and related Genera: Evidence from phylogenetic Analyses of nuclear Internal Transcribed Spacer and Plastid trnL-F Sequences.**
In: Systematic Biology 53 (6), p. 856–876. DOI: 10.1080/10635150490522566.

TRIFT, IDA; LIDÉN, MAGNUS; ANDERBERG, ARNE A. (2004):
**Phylogeny and Biogeography of *Dionysia* (Primulaceae).**
In: International Journal of Plant Sciences 165 (5), p. 845–860.

ZHANG, LI-BING; COMES, HANS PETER; KADEREIT, JOACHIM W. (2001):
**Phylogeny and quaternary History of the European montane/alpine endemic *Soldanella* (Primulaceae) based on ITS and AFLP Variation.**
In: American Journal of Botany 88 (12), p. 2331–2345. DOI: 10.2307/3558393.

ZHANG, LI-BING; COMES, HANS PETER; KADEREIT, JOACHIM W. (2004):
**The temporal Course of Quaternary Diversification in the European High Mountain Endemic *Primula* Sect. *Auricula* (Primulaceae).**
In: International Journal of Plant Sciences 165 (1), p. 191–207.

ZHANG, LI-BING; KADEREIT, JOACHIM W. (2004):

**Classification of *Primula* Sect. *Auricula* (Primulaceae) based on two molecular Data sets (ITS, AFLPs), Morphology and geographical Distribution.**
In: Botanical Journal of the Linnean Society 146 (1), p. 1–26. DOI: 10.1111/j.1095-8339.2004.00301.x.

ZHANG, LI-BING; KADEREIT, JOACHIM W. (2004):
**Nomenclature of *Soldanella* L. (Primulaceae).**
In: Taxon 53 (3), p. 741–752. DOI: 10.2307/4135448.

### Proteaceae

BARKER, NIGEL P.; VANDERPOORTEN, ALAIN; MORTON, CYNTHIA M.; ROURKE, JOHN P. (2004):
**Phylogeny, Biogeography, and the Evolution of Life-history Traits in *Leucadendron* (Proteaceae).**
In: Molecular Phylogenetics and Evolution 33 (3), p. 845–860. DOI: 10.1016/j.ympev.2004.07.007.

BUTCHER, RYONEN; THIELE, KEVIN R. (2014):
**An Investigation of Taxon Boundaries in rare and range-restricted *Synaphea* (Proteaceae: Conosperminae) Species from South-west Western Australia.**
In: Australian Systematic Botany 27 (2), p. 119–144. DOI: 10.1071/SB14015.

CARDILLO, MARCEL; PRATT, RENAE (2013):
**Evolution of a Hotspot Genus: geographic Variation in Speciation and extinction rates in *Banksia* (Proteaceae).**
In: BMC Evolutionary Biology 13, p. 155. DOI: 10.1186/1471-2148-13-155.

DUCHENE, DAVID; BROMHAM, LINDELL (2013):
**Rates of molecular Evolution and Diversification in plants: Chloroplast Substitution Rates correlate with Species-richness in the Proteaceae.**
In: BMC Evolutionary Biology 13, p. 65. DOI: 10.1186/1471-2148-13-65.

HOLMES, GARETH D.; DOWNING, TRISHA L.; JAMES, ELIZABETH A.; BLACKET, MARK J.; HOFFMANN, ARY A.; BAYLY, MICHAEL J. (2014):
**Phylogeny of the holly Grevilleas (Proteaceae) based on nuclear ribosomal and Chloroplast DNA.**
In: Australian Systematic Botany 27 (1), p. 56–77. DOI: 10.1071/SB13045.

MAST, AUSTIN R.; GIVNISH, THOMAS J. (2002):
**Historical Biogeography and the Origin of stomatal Distributions in *Banksia* and *Dryandra* (Proteaceae) based on their CpDNA Phylogeny.**
In: American Journal of Botany 89 (8), p. 1311–1323.

MAST, AUSTIN R.; JONES, ERIC H.; HAVERY, SHAWN P. (2005):
**An assessment of old and new DNA Sequence Evidence for the Paraphyly of *Banksia* with respect to *Dryandra* (Proteaceae).**
In: Australian Systematic Botany 18 (1), p. 75–88. DOI: 10.1071/SB04015.

MAST, AUSTIN R.; MILTON, ETHAN F.; JONES, ERIC H.; BARKER, ROBYN M.; BARKER, WILLIAM R.; WESTON, PETER H. (2012):
**Time-calibrated Phylogeny of the woody Australian Genus *Hakea* (Proteaceae) supports multiple Origins of insect-Pollination among bird-pollinated Ancestors.**
In: American Journal of Botany 99 (3), p. 472–487. DOI: 10.3732/ajb.1100420.

MAST, AUSTIN R.; OLDE, PETER M.; MAKINSON, ROBERT O.; JONES, ERIC H.; KUBES, AMANDA; MILLER, ELIOT T.; WESTON, PETER H. (2015):

**Paraphyly changes Understanding of Timing and Tempo of Diversification in Subtribe Hakeinae (Proteaceae), a giant Australian plant Radiation.**
In: American Journal of Botany 102 (10), p. 1634–1646. DOI: 10.3732/ajb.1500195.

**Mast, Austin R.; Thiele, Kevin R. (2007):**
**The transfer of *Dryandra* R.Br. to *Banksia* L.f. (Proteaceae).**
In: Australian Systematic Botany 20 (1), p. 63–71. DOI: 10.1071/SB06016.

**Mast, Austin R.; Willis, Crystal L.; Jones, Eric H.; Downs, Katherine M.; Weston, Peter H. (2008):**
**A smaller *Macadamia* from a more vagile Tribe: Inference of phylogenetic Relationships, Divergence times, and diaspore Evolution in *Macadamia* and relatives (tribe Macadamieae; Proteaceae).**
In: American Journal of Botany 95 (7), p. 843–870. DOI: 10.3732/ajb.0700006.

**Milner, Melita L.; McIntosh, Emma J.; Crisp, Michael D.; Weston, Peter H.; Rossetto, Maurizio (2013):**
**Microsatellite Variation for phylogenetic, phylogeographic and population-genetic studies in *Lomatia* (Proteaceae).**
In: Australian Systematic Botany 26 (3), p. 186–195. DOI: 10.1071/SB13002.

**Nash, Chloe M. (2017):**
**Digest: Shifting biomes: Insight into Patterns of plant Radiation and Dispersal.**
In: Evolution 71 (8), p. 2130–2131. DOI: 10.1111/evo.13300.

**Pharmawati, M.; Yan, G.; McFarlane, I. J. (2004):**
**Application of Rapd and ISSR Markers to analyse molecular Relationships in *Grevillea* (Proteaceae).**
In: Australian Systematic Botany 17 (1), p. 49–61. DOI: 10.1071/SB03016.

**Reyes, Elisabeth; Morlon, Hélène; Sauquet, Hervé (2015):**
**Presence in Mediterranean Hotspots and Floral Symmetry affect Speciation and Extinction Rates in Proteaceae.**
In: the new Phytologist 207 (2), p. 401–410. DOI: 10.1111/nph.13244.

**Sauquet, Hervé; Weston, Peter H.; Barker, Nigel P.; Anderson, Cajsa Lisa; Cantrill, David J.; Savolainen, Vincent (2009):**
**Using fossils and molecular Data to reveal the Origins of the Cape Proteas (Subfamily Proteoideae).**
In: Molecular Phylogenetics and Evolution 51 (1), p. 31–43. DOI: 10.1016/j.ympev.2008.12.013.

**Tonnabel, Jeanne; Mignot, Agnès; Douzery, Emmanuel J. P.; Rebelo, Anthony G.; Schurr, Frank M.; Midgley, Jeremy J. et al. (2014):**
**Convergent and correlated Evolution of major Life-history Traits in the Angiosperm Genus *Leucadendron* (Proteaceae).**
In: Evolution 68 (10), p. 2775–2792. DOI: 10.1111/evo.12480.

**Tonnabel, Jeanne; Olivieri, Isabelle; Mignot, Agnès; Rebelo, Anthony G.; Justy, Fabienne; Santoni, Sylvain et al. (2014):**
**Developing nuclear DNA phylogenetic Markers in the Angiosperm Genus *Leucadendron* (Proteaceae): a Next-generation Sequencing transcriptomic Approach.**
In: Molecular Phylogenetics and Evolution 70, p. 37–46. DOI: 10.1016/j.ympev.2013.07.027.

**Welsford, Megan R.; Hobbhahn, Nina; Midgley, Jeremy J.; Johnson, Steven D. (2016):**

**Floral trait Evolution associated with shifts between Insect and Wind Pollination in the dioecious Genus *Leucadendron* (Proteaceae).**
In: Evolution 70 (1), p. 126–139. DOI: 10.1111/evo.12821.

**Weston, Peter H.; Kooyman, Robert M. (2002):**
**Systematics of *Eidothea* (Proteaceae), with the Description of a new Species, *E. hardeniana*, from the Nightcap Range, north-eastern New South Wales.**
In: Telopea 9 (4), p. 821–832.

### Pteridaceae

**Bouma, Whitney L. M.; Ritchie, Peter A.; Perrie, Leon R. (2010):**
**Phylogeny and generic Taxonomy of the New Zealand Pteridaceae Ferns from Chloroplast rbcL DNA Sequences.**
In: Australian Systematic Botany 23 (3), p. 143–151. DOI: 10.1071/SB09047.

**Chao, Yi-Shan; Rouhan, Germinal; Amoroso, Victor B.; Chiou, Wen-Liang (2014):**
**Molecular Phylogeny and Biogeography of the Fern Genus *Pteris* (Pteridaceae).**
In: Annals of Botany 114 (1), p. 109–124. DOI: 10.1093/aob/mcu086.

**Cochran, Alyssa T.; Prado, Jefferson; Schuettpelz, Eric (2014):**
**Tryonia, a new Taenitidoid Fern Genus segregated from *Jamesonia* and *Eriosorus* (Pteridaceae).**
In: PhytoKeys (35), p. 23–43. DOI: 10.3897/phytokeys.35.6886.

**Eiserhardt, Wolf L.; Rohwer, Jens Gunter; Russell, Stephen J.; Yesilyurt, Jovita C.; Schneider, Harald (2011):**
**Evidence for Radiations of Cheilanthoid Ferns in the Greater Cape Floristic Region.**
In: Taxon 60 (5), p. 1269–1283.

**Grusz, Amanda L.; Windham, Michael D.; Pryer, Kathleen M. (2009):**
**Deciphering the Origins of apomictic Polyploids in the *Cheilanthes yavapensis* complex (Pteridaceae).**
In: American Journal of Botany 96 (9), p. 1636–1645. DOI: 10.3732/ajb.0900019.

**Hirai, Regina Y.; Schuettpelz, Eric; Huiet, Layne; Pryer, Kathleen M.; Smith, Alan R.; Prado, Jefferson (2016):**
**Phylogeny and Relationships of the Neotropical *Adiantum raddianum* Group (Pteridaceae).**
In: Taxon 65 (6), p. 1225–1235. DOI: 10.12705/656.1.

**Kirkpatrick, Ruth E. B. (2007):**
**Investigating the Monophyly of *Pellaea* (Pteridaceae) in the Context of a phylogenetic Analysis of Cheilanthoid Ferns.**
In: Systematic Botany 32 (3), p. 504–518.

**Li, Fay-Wei; Pryer, Kathleen M.; Windham, Michael D. (2012):**
***Gaga*, a new Fern Genus Segregated from Cheilanthes (Pteridaceae).**
In: Systematic Botany 37 (4), p. 845–860. DOI: 10.1600/036364412X656626.

**Lu, Jin-Mei; Wen, Jun; Lutz, Sue; Wang, Yi-Ping; Li, De-Zhu (2012):**
**Phylogenetic Relationships of Chinese *Adiantum* based on five Plastid Markers.**
In: Journal of Plant Research 125 (2), p. 237–249. DOI: 10.1007/s10265-011-0441-y.

**Metzgar, Jordan S.; Alverson, Edward R.; Chen, Sinian; Vaganov, Alexey V.; Ickert-Bond, Stefanie M. (2013):**
**Diversification and reticulation in the circumboreal Fern Genus *Cryptogramma*.**
In: Molecular Phylogenetics and Evolution 67 (3), p. 589–599. DOI: 10.1016/j.ympev.2013.02.020.

**NAKAZATO, TAKUYA; GASTONY, GERALD J. (2003):**

**Molecular Phylogenetics of *Anogramma* Species and related Genera (Pteridaceae: Taenitidoideae).**

In: Systematic Botany 28 (3), p. 490–502.

**PABÓN-MORA, NATALIA; GONZÁLEZ, FAVIO (2016):**

***Nephopteris* out of the Clouds: molecular Evidence places the enigmatic *N. maxonii* (Pteridaceae) within the *Jamesonia* Clade.**

In: Brittonia 68 (1), p. 83–92. DOI: 10.1007/s12228-015-9394-0.

**PONCE, MONICA; SCATAGLINI, MARIA AMALIA (2016):**

**Do two South American Species of *Cheilanthes* (Pteridaceae) traditionally linked to the *Cheilanthes marginata* Group, belong to *Gaga*?**

In: Phytotaxa 257 (2), p. 149. DOI: 10.11646/phytotaxa.257.2.4.

**PRADO, JEFFERSON; RODRIGUES, CRISTIANE DEL NERO; SALATINO, ANTONIO; SALATINO, MARIA LUIZA FARIA (2007):**

**Phylogenetic Relationships among Pteridaceae, including Brazilian species, inferred from rbcL Sequences.**

In: Taxon 56 (2), p. 355–368.

**PRYER, KATHLEEN M.; HUIET, LAYNE; LI, FAY-WEI; ROTHFELS, CARL J.; SCHUETTPELZ, ERIC (2016):**

**Maidenhair Ferns, *Adiantum*, are indeed monophyletic and Sister to Shoestring Ferns, Vittarioids (Pteridaceae).**

In: Systematic Botany 41 (1), p. 17–23. DOI: 10.1600/036364416X690660.

**ROTHFELS, CARL J.; SCHUETTPELZ, ERIC (2014):**

**Accelerated rate of molecular Evolution for Vittarioid Ferns is strong and not driven by Selection.**

In: Systematic Biology 63 (1), p. 31–54. DOI: 10.1093/sysbio/syt058.

**ROTHFELS, CARL J.; WINDHAM, MICHAEL D.; GRUSZ, AMANDA L.; GASTONY, GERALD J.; PRYER, KATHLEEN M. (2008):**

**Toward a monophyletic *Notholaena* (Pteridaceae): resolving Patterns of evolutionary Convergence in xeric-adapted Ferns.**

In: Taxon 57 (3), p. 712–724.

**RUHFEL, BRADLEY R.; LINDSAY, STUART; DAVIS, CHARLES C. (2008):**

**Phylogenetic Placement of *Rheopteris* and the Polyphyly of *Monogramma* (Pteridaceae s.l.): Evidence from rbcL Sequence Data.**

In: Systematic Botany 33 (1), p. 37–43. DOI: 10.1600/036364408783887410.

**SÁNCHEZ-BARACALDO, PATRICIA (2004):**

**Phylogenetic Relationships of the Subfamily Taenitidoideae, Pteridaceae.**

In: American Fern Journal 94 (3), p. 126–142. DOI: 10.1640/0002-8444(2004)094[0126:PROTST]2.0.CO;2.

**SÁNCHEZ-BARACALDO, PATRICIA (2004):**

**Phylogenetics and Biogeography of the Neotropical Fern Genera *Jamesonia* and *Eriosorus* (Pteridaceae).**

In: American Journal of Botany 91 (2), p. 274–284.

**SCATAGLINI, MARIA AMALIA; AMALIA, M. (2012):**

**Nuevas Combinaciones en *Adiantopsis* (Pteridaceae, Cheilantheae) de América del Sur.**

In: Novon: A Journal for Botanical Nomenclature 22 (1), p. 62–66. DOI: 10.3417/2010059.

**SCHNEIDER, HARALD; HE, LI-JUAN; HENNEQUIN, SABINE; ZHANG, XIAN-CHUN (2013):**

**Towards a natural Classification of Pteridaceae: inferring the Relationships of enigmatic Pteridoid Fern Species occurring in the Sino-Himalaya and Afro-Madagascar.**
In: Phytotaxa 77 (4). DOI: 10.11646/phytotaxa.77.4.1.

SCHUETTPELZ, ERIC; CHEN, CHENG-WEI; KESSLER, MICHAEL; PINSON, JERALD B.; JOHNSON, GABRIEL; DAVILA, ALEX ET AL. (2016):
**A revised generic Classification of Vittarioid Ferns (Pteridaceae) based on molecular, micromorphological, and geographic Data.**
In: Taxon 65 (4), p. 708–722. DOI: 10.12705/654.2.

SCHUETTPELZ, ERIC; PRYER, KATHLEEN M.; WINDHAM, MICHAEL D. (2015):
**A unified Approach to taxonomic Delimitation in the Fern Genus *Pentagramma* (Pteridaceae).**
In: Systematic Botany 40 (3), p. 629–644. DOI: 10.1600/036364415X689366.

SCHUETTPELZ, ERIC; SCHNEIDER, HARALD; HUIET, LAYNE; WINDHAM, MICHAEL D.; PRYER, KATHLEEN M. (2007):
**A molecular Phylogeny of the Fern Family Pteridaceae: assessing overall Relationships and the Affinities of previously unsampled Genera.**
In: Molecular Phylogenetics and Evolution 44 (3), p. 1172–1185. DOI: 10.1016/j.ympev.2007.04.011.

SIGEL, ERIN M.; WINDHAM, MICHAEL D.; HUIET, LAYNE; YATSKIEVYCH, GEORGE; PRYER, KATHLEEN M. (2011):
**Species Relationships and Farina Evolution in the Cheilanthoid Fern Genus *Argyrochosma* (Pteridaceae).**
In: Systematic Botany 36 (3), p. 554–564. DOI: 10.1600/036364411X583547.

WANG, W. A.N.; YANG, WENLI; MAO, XINGXING; ZHAO, RANRAN; DOU, PING; ZHANG, GANGMIN (2015):
**The phylogenetic affinities of *Pellaea connectens*, a rare endemic Chinese Fern.**
In: Phytotaxa 220 (1), p. 30. DOI: 10.11646/phytotaxa.220.1.2.

YESILYURT, JOVITA C.; SCHNEIDER, HARALD (2010):
**The new Fern Genus *Calciphilopteris* (Pteridaceae).**
In: Phytotaxa 7, p. 52–59.

YESILYURT, JOVITA CISLINSKI; BARBARÁ, THELMA; SCHNEIDER, HARALD; RUSSELL, STEPHEN J.; CULHAM, ALASTAIR; GIBBY, MARY (2015):
**Identifying the generic Limits of the Cheilanthoid Genus *Doryopteris*.**
In: Phytotaxa 221 (2), p. 101. DOI: 10.11646/phytotaxa.221.2.1.

ZHANG, GANGMIN; ZHANG, XIAN-CHUN; CHEN, ZHI-DUAN; LIU, HONG-MEI; YANG, WENLI (2007):
**First Insights in the Phylogeny of Asian Cheilanthoid Ferns based on Sequences of two Chloroplast Markers.**
In: Taxon 56 (2), p. 369–378.

ZHANG, LIANG; ROTHFELS, CARL J.; EBIHARA, ATSUSHI; SCHUETTPELZ, ERIC; LE PÉCHON, TIMOTHÉE; KAMAU, PERIS ET AL. (2015):
**A global Plastid Phylogeny of the Brake Fern Genus *Pteris* (Pteridaceae) and related Genera in the Pteridoideae.**
In: Cladistics 31 (4), p. 406–423. DOI: 10.1111/cla.12094.

ZHANG, LIANG; ZHANG, LI-BING (2018):
**Phylogeny and Systematics of the brake Fern Genus *Pteris* (Pteridaceae) based on molecular (plastid and nuclear) and morphological Evidence.**
In: Molecular Phylogenetics and Evolution 118, p. 265–285. DOI: 10.1016/j.ympev.2017.09.011.

**Zhang, Liang; Zhou, Xin-Mao; Lu, Ngan Thi; Zhang, Li-Bing (2017):**
**Phylogeny of the Fern Subfamily Pteridoideae (Pteridaceae; Pteridophyta), with the Description of a new Genus: *Gastoniella*.**
In: Molecular Phylogenetics and Evolution 109, p. 59–72. DOI: 10.1016/j.ympev.2016.12.037.

### Pteridophyta

**Christenhusz, Maarten J.M.; Zhang, Xian-Chun; Schneider, Harald (2011):**
**A linear Sequence of extant families and Genera of Lycophytes and Ferns.**
In: Phytotaxa 19, p. 7–54.

**Ebihara, Atsushi (2011):**
**RbcL Phylogeny of Japanese Pteridophyte Flora and Implications on Infrafamilial Systematics.**
In: Bulletin of the National Museum of Nature and Science, Tokyo, Series Botany 37 (2), p. 63–74.

**Knie, Nils; Fischer, Simon; Grewe, Felix; Polsakiewicz, Monika; Knoop, Volker (2015):**
**Horsetails are the sister Group to all other Monilophytes and Marattiales are sister to Leptosporangiate Ferns.**
In: Molecular Phylogenetics and Evolution 90, p. 140–149. DOI: 10.1016/j.ympev.2015.05.008.

**Kuo, Li-Yaung; Li, Fay-Wei; Chiou, Wen-Liang; Wang, Chun-Neng (2011):**
**First Insights into Fern matK Phylogeny.**
In: Molecular Phylogenetics and Evolution 59 (3), p. 556–566. DOI: 10.1016/j.ympev.2011.03.010.

**Lehtonen, Samuli (2011):**
**Towards resolving the complete Fern Tree of Life.**
In: Public Library of Science One 6 (10), e24851. DOI: 10.1371/journal.pone.0024851.

**Lu, Jin-Mei; Zhang, Ning; Du, Xin-Yu; Wen, Jun; Li, De-Zhu (2015):**
**Chloroplast phylogenomics resolves Key Relationships in Ferns.**
In: Journal of Systematics and Evolution 53 (5), p. 448–457. DOI: 10.1111/jse.12180.

**Pryer, Kathleen M.; Schuettpelz, Eric; Wolf, Paul G.; Schneider, Harald; Smith, Alan R.; Cranfill, Raymond B. (2004):**
**Phylogeny and Evolution of Ferns (Monilophytes) with a Focus on the aarly Leptosporangiate Divergences.**
In: American Journal of Botany 91 (10), p. 1582–1598.

**Rai, Hardeep S.; Graham, Sean W. (2010):**
**Utility of a large, multigene Plastid Data set in inferring higher-order Relationships in Ferns and relatives (Monilophytes).**
In: American Journal of Botany 97 (9), p. 1444–1456. DOI: 10.3732/ajb.0900305.

**Rothfels, Carl J.; Larsson, Anders; Kuo, Li-Yaung; Korall, Petra; Chiou, Wen-Liang; Pryer, Kathleen M. (2012):**
**Overcoming deep Roots, fast Rates, and short Internodes to resolve the ancient rapid Radiation of Eupolypod II Ferns.**
In: Systematic Biology 61 (3), p. 490–509. DOI: 10.1093/sysbio/sys001.

**Rothfels, Carl J.; Li, Fay-Wei; Sigel, Erin M.; Huiet, Layne; Larsson, Anders; Burge, Dylan O. et al. (2015):**
**The evolutionary History of Ferns inferred from 25 low-copy nuclear Genes.**
In: American Journal of Botany 102 (7), p. 1089–1107. DOI: 10.3732/ajb.1500089.

**Schuettpelz, Eric; Pryer, Kathleen M. (2007):**
**Fern Phylogeny inferred from 400 Leptosporangiate Species and three Plastid Genes.**

In: Taxon 56 (4), p. 1037–1050.

**Tanaka, Takayuki; Isaka, Yuichi; Hattori, Mitsuru; Sato, Toshiyuki (2014):**
**Ecological and phylogenetic Approaches for Diversification of apogamous Ferns in Japan.**
In: Plant Systematics and Evolution 300 (9), p. 2041–2050. DOI: 10.1007/s00606-014-1036-6.

**Testo, Weston; Sundue, Michael A. (2016):**
**A 4000-species Dataset provides new Insight into the Evolution of Ferns.**
In: Molecular Phylogenetics and Evolution 105, p. 200–211. DOI: 10.1016/j.ympev.2016.09.003.

**Xianchun, Zhang; Ran, Wei; Hongmei, Liu; Lijuan, He; Li, Wang; Gangmin, Zhang (2013):**
**Phylogeny and Classification of the extant Lycophytes and Ferns from China.**
In: Chinese Bulletin of Botany 48 (2), p. 119–137. DOI: 10.3724/SP.J.1259.2013.00119.

### Quiinaceae

**Schneider, Julio Valentin; Swenson, Ulf; Samuel, L.; Stuessy, Tod F.; Zizka, Georg (2006):**
**Phylogenetics of Quiinaceae (Malpighiales): Evidence from trnL-trnF Sequence Data and Morphology.**
In: Plant Systematics and Evolution 257 (3-4), p. 189–203. DOI: 10.1007/s00606-005-0386-5.

### Quillajaceae

**Luebert, Federico (2013):**
**Taxonomy and Distribution of the Genus *Quillaja* Molina (Quillajaceae).**
In: Feddes Repertorium 124 (4), p. 157–162. DOI: 10.1002/fedr.201400029.

### Rafflesiaceae

**Bendiksby, Mika; Schumacher, Trond; Gussarova, Galina; Nais, Jamili; Mat-Salleh, Kamarudin; Sofiyanti, Nery et al. (2010):**
**Elucidating the evolutionary History of the Southeast Asian, holoparasitic, giant-flowered Rafflesiaceae: pliocene vicariance, morphological convergence and Character Displacement.**
In: Molecular Phylogenetics and Evolution 57 (2), p. 620–633. DOI: 10.1016/j.ympev.2010.08.005.

**Nickrent, Daniel Lee; Blarer, Albert; Qiu, Yin-Long; Vidal-Russell, Romina; Anderson, Frank E. (2004):**
**Phylogenetic Inference in Rafflesiales: the Influence of Rate Heterogeneity and horizontal Gene Transfer.**
In: BMC Evolutionary Biology 4, p. 40. DOI: 10.1186/1471-2148-4-40.

### Ranunculaceae

**Baltisberger, Matthias; Hörandl, Elvira (2016):**
**Karyotype Evolution supports the molecular Phylogeny in the Genus *Ranunculus* (Ranunculaceae).**
In: Perspectives in Plant Ecology, Evolution and Systematics 18, p. 1–14. DOI: 10.1016/j.ppees.2015.11.001.

**Bastida, Jesús M.; Alcántara, Julio M.; Rey, Pedro J.; Vargas, Pablo; Herrera, Carlos M. (2010):**
**Extended Phylogeny of *Aquilegia*: the biogeographical and ecological Patterns of two simultaneous but contrasting Radiations.**
In: Plant Systematics and Evolution 284 (3-4), p. 171–185. DOI: 10.1007/s00606-009-0243-z.

**Bittkau, Christiane; Comes, Hans Peter (2009):**
**Molecular Inference of a late Pleistocene Diversification Shift in *Nigella* s. lat. (Ranunculaceae) resulting from increased Speciation in the Aegean Archipelago.**
In: Journal of Biogeography 36 (7), p. 1346–1360. DOI: 10.1111/j.1365-2699.2008.02003.x.

**Chartier, Marion; Dressler, Stefan; Schönenberger, Jürgen; Mora, Alfonso Rojas; Sarthou, Corinne; Wang, Wei; Jabbour, Florian (2016):**

**The Evolution of afro-montane *Delphinium* (Ranunculaceae): Morphospecies, Phylogenetics and Biogeography.**

In: Taxon 65 (6), p. 1313–1327. DOI: 10.12705/656.6.

**Cheng, Jin; Xie, Lei (2014):**

**Molecular Phylogeny and historical Biogeography of *Caltha* (Ranunculaceae) based on Analyses of multiple nuclear and Plastid Sequences.**

In: Journal of Systematics and Evolution 52 (1), p. 51–67. DOI: 10.1111/jse.12051.

**Cires, Eduardo; Baltisberger, Matthias; Cuesta, Candela; Vargas, Pablo; Prieto, José Antonio Fernández (2014):**

**Allopolyploid Origin of the Balkan endemic *Ranunculus wettsteinii* (Ranunculaceae) inferred from nuclear and Plastid DNA Sequences.**

In: Organisms Diversity and Evolution 14 (1), p. 1–10. DOI: 10.1007/s13127-013-0150-6.

**Compton, J. A.; Culham, Alastair; Gibbings, J. G.; Jury, p. L. (1998):**

**Phylogeny of *Actaea* including *Cimicifuga* (Ranunculaceae) inferred from nrDNA ITS Sequence Variation.**

In: Biochemical Systematics and Ecology 26, p. 185–197.

**Cossard, Guillaume; Sannier, Julie; Sauquet, Hervé; Damerval, Catherine; Craene, Louis Ronse; Jabbour, Florian; Nadot, Sophie (2016):**

**Subfamilial and tribal Relationships of Ranunculaceae: Evidence from eight molecular Markers.**

In: Plant Systematics and Evolution 302 (4), p. 419–431. DOI: 10.1007/s00606-015-1270-6.

**Ehrendorfer, Friedrich; Ziman, Svetlana N.; König, Christiane; Keener, Carl S.; Dutton, Bryan E.; Tsarenko, Olga N. et al. (2009):**

**Taxonomic Revision, Phylogenetics and transcontinental Distribution of *Anemone* Section *Anemone* (Ranunculaceae).**

In: Botanical Journal of the Linnean Society 160, p. 312–354.

**Emadzade, Khatere; Gehrke, Berit; Linder, Hans Peter; Hörandl, Elvira (2011):**

**The biogeographical History of the cosmopolitan Genus *Ranunculus* L. (Ranunculaceae) in the temperate to meridional Zones.**

In: Molecular Phylogenetics and Evolution 58 (1), p. 4–21. DOI: 10.1016/j.ympev.2010.11.002.

**Emadzade, Khatere; Lebmann, Markus J.; Hoffmann, Matthias H.; Tkach, Natalia V.; Lone, Fayaz A.; Hörandl, Elvira (2015):**

**Phylogenetic Relationships and Evolution of high mountain Buttercups (*Ranunculus*) in North America and Central Asia.**

In: Perspectives in Plant Ecology, Evolution and Systematics 17 (2), p. 131–141. DOI: 10.1016/j.ppees.2015.02.001.

**Emadzade, Khatere; Lehnebach, Carlos A.; Lockhart, Peter J.; Hörandl, Elvira (2010):**

**A molecular Phylogeny, Morphology and Classification of Genera of Ranunculeae (Ranunculaceae).**

In: Taxon 59 (3), p. 809–828.

**He, Yang; Hou, Pei; Fan, Gang; Arain, Saima; Peng, Cheng (2014):**

**Comprehensive Analyses of molecular Phylogeny and main Alkaloids for *Coptis* (Ranunculaceae) Species Identification.**

In: Biochemical Systematics and Ecology 56, p. 88–94. DOI: 10.1016/j.bse.2014.05.002.

**Heiss, Andreas G.; Kropf, Matthias; Sontag, Susanne; Weber, Anton (2011):**
**Seed Morphology of *Nigella* s.l. (Ranunculaceae): Identification, diagnostic Traits, and their potential phylogenetic Relevance.**
In: International Journal of Plant Sciences 172 (2), p. 267–284. DOI: 10.1086/657676.

**Hoffmann, Matthias H. (1999):**
**The Phylogeny of *Actaea* (Ranunculaceae): a biogeographical Approach.**
In: Plant Systematics and Evolution 216, p. 251–263.

**Hoffmann, Matthias H.; Hagen, K. Bernhard von; Hörandl, Elvira; Röser, Martin; Tkach, Natalia V. (2010):**
**Sources of the Arctic Flora: Origins of Arctic Species in *Ranunculus* and related Genera.**
In: International Journal of Plant Sciences 171 (1), p. 90–106. DOI: 10.1086/647918.

**Hong, Yu; Luo, Yan; Gao, Qi; Ren, Chen; Yuan, Qiong; Yang, Qin-Er (2017):**
**Phylogeny and reclassification of *Aconitum* Subgenus *Lycoctonum* (Ranunculaceae).**
In: Public Library of Science One 12 (1), e0171038. DOI: 10.1371/journal.pone.0171038.

**Hoot, Sara B.; Kramer, Jenny; Arroyo, Mary T.K. (2008):**
**Phylogenetic Position of the South American dioecious Genus *Hamadryas* and related Ranunculeae (Ranunculaceae).**
In: International Journal of Plant Sciences 169 (3), p. 433–443. DOI: 10.1086/526460.

**Hörandl, Elvira; Emadzade, Khatere (2011):**
**The Evolution and Biogeography of alpine Species in *Ranunculus* (Ranunculaceae): A global Comparison.**
In: Taxon 60 (2), p. 415–426.

**Hörandl, Elvira; Greilhuber, Johann; Klímová, Katarina; Paun, Ovidiu; Temsch, Eva M.; Emadzade, Khatere (2009):**
**Reticulate Evolution and taxonomic Concepts in the *Ranunculus auricomus* complex (Ranunculaceae): Insights from Analysis of morphological, karyological and molecular Data.**
In: Taxon 58 (4), p. 1194–1215.

**Hörandl, Elvira; Paun, Ovidiu; Johansson, Jan Thomas; Lehnebach, Carlos A.; Armstrong, Tristan; Chen, Lixue; Lockhart, Peter J. (2005):**
**Phylogenetic Relationships and evolutionary traits in *Ranunculus* s.l. (Ranunculaceae) inferred from ITS Sequence Analysis.**
In: Molecular Phylogenetics and Evolution 36 (2), p. 305–327. DOI: 10.1016/j.ympev.2005.02.009.

**Jabbour, Florian; Renner, Susanne p. (2011):**
***Consolida* and *Aconitella* are an annual Clade of *Delphinium* (Ranunculaceae) that diversified in the Mediterranean basin and the Irano-Turanian Region.**
In: Taxon 60 (4), p. 1029–1040.

**Jabbour, Florian; Renner, Susanne p. (2012):**
**A Phylogeny of Delphinieae (Ranunculaceae) shows that *Aconitum* is nested within *Delphinium* and that Late Miocene Transitions to long Life Cycles in the Himalayas and Southwest China coincide with bursts in Diversification.**
In: Molecular Phylogenetics and Evolution 62 (3), p. 928–942. DOI: 10.1016/j.ympev.2011.12.005.

**Jabbour, Florian; Renner, Susanne p. (2012):**

**Spurs in a Spur: Perianth Evolution in the Delphinieae (Ranunculaceae).**
In: International Journal of Plant Sciences 173 (9), p. 1036–1054. DOI: 10.1086/667613.

**Jabbour, Florian; Ronse De Craene, Louis P.; Nadot, Sophie; Damerval, Catherine (2009):**
**Establishment of Zygomorphy on an ontogenic spiral and Evolution of Perianth in the Tribe Delphinieae (Ranunculaceae).**
In: Annals of Botany 104 (5), p. 809–822. DOI: 10.1093/aob/mcp162.

**Johansson, Jan Thomas (1999):**
**There large inversions in the Chloroplast Genomes and one loss of the Chloroplast Gene rpsl6 suggest an early evolutionary Split in the Genus *Adonis* (Ranunculaceae).**
In: Plant Systematics and Evolution 218, p. 133–143.

**Lehnebach, Carlos A.; Cano, A.; Monsalve, C.; McLenachan, P.; Hörandl, Elvira; Lockhart, Peter J. (2007):**
**Phylogenetic Relationships of the monotypic Peruvian Genus *Laccopetalum* (Ranunculaceae).**
In: Plant Systematics and Evolution 264 (1-2), p. 109–116. DOI: 10.1007/s00606-006-0488-8.

**Luo, Jiang-Ping; Hong, Yu; Ren, Chen; Yang, Qin-Er; Yuan, Qiong (2016):**
**Reinstatement of the Chinese Species *Cimicifuga lancifoliolata* (Ranunculaceae).**
In: Nordic Journal of Botany 34 (5), p. 627–640. DOI: 10.1111/njb.01092.

**Luo, Yan (2005):**
**Taxonomic Revision of *Aconitum* (Ranunculaceae) from Sichuan, China.**
In: Acta Phytotaxonomica Sinica 43 (4), p. 289–386. DOI: 10.1360/aps040102.

**Luo, Yan; Zhang, Fu-Min; Yang, Qin-Er (2005):**
**Phylogeny of *Aconitum* Subgenus *Aconitum* (Ranunculaceae) inferred from ITS Sequences.**
In: Plant Systematics and Evolution 252 (1-2), p. 11–25. DOI: 10.1007/s00606-004-0257-5.

**Manning, John C.; Goldblatt, Peter (2013):**
**The native and naturalised Species of *Peltocalathos* and *Ranunculus* (Ranunculaceae: Ranunculeae) in southern Africa.**
In: Bothalia 43 (2), p. 179–195.

**Meyer, Kyle M.; Hoot, Sara B.; Arroyo, Mary T.K. (2010):**
**Phylogenetic Affinities of South American *Anemone* (Ranunculaceae), including the endemic segregate Genera, *Barneoudia* and *Oreithales*.**
In: International Journal of Plant Sciences 171 (3), p. 323–331. DOI: 10.1086/650153.

**Miikeda, Osamu; Kita, Koichi; Handa, Takashi; Tomohisa, Yukawa (2006):**
**Phylogenetic Relationships of *Clematis* (Ranunculaceae) based on Chloroplast and nuclear DNA Sequences.**
In: Botanical Journal of the Linnean Society 152, p. 153–168.

**Mlinarec, Jelena; Franjevic, Damjan; Bockor, Luka; Besendorfer, Višnja (2016):**
**Diverse evolutionary Pathways shaped 5s rDNA of Species of Tribe Anemoneae (Ranunculaceae) and reveal phylogenetic Signal.**
In: Botanical Journal of the Linnean Society 182, p. 80–99.

**Mlinarec, Jelena; Satović, Zlatko; Mihelj, D.; Malenica, N.; Besendorfer, Višnja (2012):**
**Cytogenetic and phylogenetic Studies of diploid and polyploid Members of Tribe Anemoninae (Ranunculaceae).**

In: Plant Biology 14 (3), p. 525–536. DOI: 10.1111/j.1438-8677.2011.00519.x.

**Paun, Ovidiu; Lehnebach, Carlos A.; Johansson, Jan Thomas; Lockhart, Peter J.; Hörandl, Elvira (2005):**
**Phylogenetic Relationships and Biogeography of *Ranunculus* and allied Genera (Ranunculaceae) in the Mediterranean Region and in the European Alpine System.**
In: Taxon 54 (4), p. 911–930.

**Schuettpelz, Eric; Hoot, Sara B. (2004):**
**Phylogeny and Biogeography of *Caltha* (Ranunculaceae) based on Chloroplast and nuclear DNA Sequences.**
In: American Journal of Botany 91 (2), p. 247–253.

**Soza, Valerie L.; Brunet, Johanne; Liston, Aaron; Smith, Patricia Salles; Di Stilio, Verónica p. (2012):**
**Phylogenetic Insights into the Correlates of Dioecy in Meadow-rues (*Thalictrum*, Ranunculaceae).**
In: Molecular Phylogenetics and Evolution 63 (1), p. 180–192. DOI: 10.1016/j.ympev.2012.01.009.

**Soza, Valerie L.; Haworth, Kendall L.; Di Stilio, Verónica p. (2013):**
**Timing and Consequences of recurrent Polyploidy in Meadow-rues (Thalictrum, Ranunculaceae).**
In: Molecular Biology and Evolution 30 (8), p. 1940–1954. DOI: 10.1093/molbev/mst101.

**Sun, Hang; McLewin, Will; Fay, Michael F. (2001):**
**Molecular Phylogeny of *Helleborus* (Ranunculaceae), with an Emphasis on the East Asian-Mediterranean Disjunction.**
In: Taxon 50 (4), p. 1001–1018.

**Wang, W.; Li, R.-Q.; Chen, Z.-D. (2005):**
**Systematic Position of *Asteropyrum* (Ranunculaceae) inferred from Chloroplast and nuclear Sequences.**
In: Plant Systematics and Evolution 255 (1-2), p. 41–54. DOI: 10.1007/s00606-005-0339-z.

**Wang, Wei; Chen, Zhi-Duan (2007):**
**Generic level Phylogeny of Thalictroideae (Ranunculaceae) as Implications for the taxonomic Status of *Paropyrum* and petal Evolution.**
In: Taxon 56 (3), p. 811–821.

**Wang, Wei; Hu, Hao; Xiang, Xiao-Guo; Yu, Sheng-Xiang; Chen, Zhi-Duan (2010):**
**Phylogenetic Placements of *Calathodes* and *Megaleranthis* (Ranunculaceae): Evidence from molecular and morphological Data.**
In: Taxon 59 (6), p. 1712–1720.

**Wang, Wei; Li, Hong-Lei; Xiang, Xiao-Guo; Chen, Zhi-Duan (2014):**
**Revisiting the Phylogeny of Ranunculeae: Implications for Divergence time Estimation and historical Biogeography.**
In: Journal of Systematics and Evolution 52 (5), p. 551–565. DOI: 10.1111/jse.12101.

**Wang, Wei; Liu, Yang; Yu, Sheng-Xiang; Gao, Tian-Gang; Chen, Zhi-Duan (2013):**
***Gymnaconitum*, a new Genus of Ranunculaceae endemic to the Qinghai-Tibetan Plateau.**
In: Taxon 62 (4), p. 713–722. DOI: 10.12705/624.10.

**Wang, Wen-Tsai (2007):**
**A Revision of *Clematis* Sect. *Tubulosae* (Ranunculaceae).**
In: Acta Phytotaxonomica Sinica 45 (4), p. 425–457. DOI: 10.1360/aps06114.

**Xiang, Kun-Li; Erst, Andrey S.; Xiang, Xiao-Guo; Jabbour, Florian; Wang, Wei (2018):**
**Biogeography of *Coptis* Salisb. (Ranunculales, Ranunculaceae, Coptidoideae), an Eastern Asian and North American Genus.**
In: BMC Evolutionary Biology 18 (1), p. 74. DOI: 10.1186/s12862-018-1195-0.

**Xiang, Kun-Li; Wu, Sheng-Dan; Yu, Sheng-Xiang; Liu, Yang; Jabbour, Florian; Erst, Andrey p. et al. (2016):**
**The First Comprehensive Phylogeny of *Coptis* (Ranunculaceae) and its Implications for Character Evolution and Classification.**
In: Public Library of Science One 11 (4), e0153127. DOI: 10.1371/journal.pone.0153127.

**Xiang, Kun-Li; Zhao, Liang; Erst, Andrey S.; Yu, Sheng-Xiang; Jabbour, Florian; Wang, Wei (2017):**
**A molecular Phylogeny of *Dichocarpum* (Ranunculaceae): Implications for eastern Asian Biogeography.**
In: Molecular Phylogenetics and Evolution 107, p. 594–604. DOI: 10.1016/j.ympev.2016.12.026.

**Xie, Lei; Li, Liang-Qian (2012):**
**Variation of Pollen Morphology, and its Implications in the Phylogeny of *Clematis* (Ranunculaceae).**
In: Plant Systematics and Evolution 298 (8), p. 1437–1453. DOI: 10.1007/s00606-012-0648-y.

**Xie, Lei; Wen, Jun; Li, Liang-Qian (2011):**
**Phylogenetic Analyses of *Clematis* (Ranunculaceae) based on Sequences of Nuclear Ribosomal ITS and three Plastid Regions.**
In: Systematic Botany 36 (4), p. 907–921. DOI: 10.1600/036364411X604921.

**Zhang, Yu; Hong, Yu; Ren, Chen; Tang, Ming; Hoot, Sara B.; Yang, Qin-Er (2015):**
**Palynology, Cytology, and molecular Systematics of *Anemone* Section *Begoniifolia* (Ranunculaceae).**
In: Plant Systematics and Evolution 301 (1), p. 411–424. DOI: 10.1007/s00606-014-1082-0.

**Zhang, Yu; Kong, Hang-Hui; Yang, Qin-Er (2015):**
**Phylogenetic Relationships and taxonomic Status of the monotypic Chinese Genus *Anemoclema* (Ranunculaceae).**
In: Plant Systematics and Evolution 301 (5), p. 1335–1344. DOI: 10.1007/s00606-014-1160-3.

### Ranunculales

**Wang, Wei; Lu, An-Ming; Ren, Yi; Endress, Mary E.; Chen, Zhi-Duan (2009):**
**Phylogeny and Classification of Ranunculales: Evidence from four molecular Loci and morphological Data.**
In: Perspectives in Plant Ecology, Evolution and Systematics 11 (2), p. 81–110. DOI: 10.1016/j.ppees.2009.01.001.

### Rapateaceae

**Givnish, Thomas J.; Evans, Timothy M.; Zjhra, Michelle L.; Patterson, T. B.; Berry, Paul E.; Sytsma, K. J. (2000):**
**Molecular Evolution, Adaptive Radiation, and Geographic Diversification in the Amphiatlantic Family Rapateaceae: Evidence from ndhF Sequences and Morphology.**
In: Evolution 54 (6), p. 1915–1937. DOI: 10.1111/j.0014-3820.2000.tb01237.x.

**Givnish, Thomas J.; Millam, Kendra C.; Evans, Timothy M.; Hall, Jocelyn C.; Chris Pires, J.; Berry, Paul E.; Sytsma, Kenneth J. (2004):**

**Ancient Vicariance or Recent Long-Distance Dispersal? Inferences about Phylogeny and South American–African Disjunctions in Rapateaceae and Bromeliaceae based on ndh F Sequence Data.**
In: International Journal of Plant Sciences 165 (S4), S35-S54. DOI: 10.1086/421067.

**Oriani, Aline; Scatena, Vera Lucia (2013):**
**The taxonomic value of Floral Characters in Rapateaceae (Poales-Monocotyledons).**
In: Plant Systematics and Evolution 299 (2), p. 291–303. DOI: 10.1007/s00606-012-0721-6.

### Rehmanniaceae

**Albach, Dirk C.; Yan, Kun; Jensen, Søren Rosendal; Li, Hong-Qing (2009):**
**Phylogenetic Placement of *Triaenophora* (formerly Scrophulariaceae) with some Implications for the Phylogeny of Lamiales.**
In: Taxon 58 (3), p. 749–756.

**Liu, Chun-yan; Yuan, Chao; Li, Xue-tong; Li, Zhong-Hu; Zhao, Peng; Qian, Zeng-Qiang et al. (2015):**
**Development of Microsatellites from *Rehmannia glutinosa* Transcriptome and Evaluation of genetic Diversity among *Rehmannia* Species.**
In: Biochemical Systematics and Ecology 59, p. 177–182. DOI: 10.1016/j.bse.2015.01.022.

### Resedaceae

**Martín-Bravo, Santiago; Meimberg, Harald; Luceño, Modesto; Märkl, Wolfgang; Valcárcel, Virginia; Bräuchler, Christian et al. (2007):**
**Molecular Systematics and Biogeography of Resedaceae based on ITS and trnL-F Sequences.**
In: Molecular Phylogenetics and Evolution 44 (3), p. 1105–1120. DOI: 10.1016/j.ympev.2006.12.016.

### Restionaceae

**Briggs, Barbara G. (2012):**
**Chromosome Numbers in some Australian Restionaceae (Poales): new Counts and an inferred Base Number for Leptocarpoideae.**
In: Telopea 14, p. 37–42. DOI: 10.7751/telopea2012006.

**Briggs, Barbara G. (2014):**
***Desmocladus* (Restionaceae) enlarged to include the Western Australian Genera *Harperia*, *Kulinia* and *Onychosepalum*.**
In: Telopea 17, p. 29–33. DOI: 10.7751/telopea20147402.

**Briggs, Barbara G.; Dixon, K. W.; Johnson, Lawrence A.S. (2012):**
**New Species and Variation in *Lepidobolus* (Restionaceae) from Western Australia.**
In: Telopea 14, p. 29–36. DOI: 10.7751/telopea2012005.

**Briggs, Barbara G.; Johnson, Lawrence A.S. (2004):**
**New Combinations in *Chordifex* (Restionaceae) from eastern Australia and new Species from Western Australia.**
In: Telopea 10 (3), p. 683–700.

**Briggs, Barbara G.; Johnson, Lawrence A.S. (2014):**
***Leptocarpus* (Restionaceae) enlarged to include *Meeboldina* and *Stenotalis*, with new Western Australian Species and Subgenera.**
In: Telopea 16, p. 19–41. DOI: 10.7751/telopea20147400.

**Briggs, Barbara G.; Linder, Hans Peter (2009):**
**A new subfamilial and tribal Classification of Restionaceae (Poales).**

In: Telopea 12 (3), p. 333–345.

**Briggs, Barbara G.; Marchant, Adam D.; Perkins, Andrew J. (2014):**
**Phylogeny of the Restiid Clade (Poales) and Implications for the Classification of Anarthriaceae, Centrolepidaceae and Australian Restionaceae.**
In: Taxon 63 (1), p. 24–46. DOI: 10.12705/631.1.

**Eldenäs, Pia K.; Linder, Hans Peter (2000):**
**Congruence and Complementarity of morphological and trnL-trnF Sequence Data and the Phylogeny of the African Restionaceae.**
In: Systematic Botany 25 (4), p. 692–707. DOI: 10.2307/2666728.

**Hardy, Christopher R.; Linder, Hans Peter (2007):**
**Phylogeny and Historical Ecology of *Rhodocoma* (Restionaceae) from the Cape Floristic Region.**
In: Aliso 23, p. 213–226.

**Linder, Hans Peter; Eldenäs, Pia K.; Briggs, Barbara G. (2003):**
**Contrasting Patterns of Radiation in African and Australian Restionaceae.**
In: Evolution 57 (12), p. 2688–2702. DOI: 10.1554/03-097.

**Linder, Hans Peter; Hardy, Christopher R. (2010):**
**A generic Classification of the Restioneae (Restionaceae), southern Africa.**
In: Bothalia 40 (1), p. 1–35. DOI: 10.4102/abc.v40i1.178.

**Linder, Hans Peter; Hardy, Christopher R.; Rutschmann, Frank (2005):**
**Taxon Sampling effects in molecular Clock Dating: an Example from the African Restionaceae.**
In: Molecular Phylogenetics and Evolution 35 (3), p. 569–582. DOI: 10.1016/j.ympev.2004.12.006.

**Linder, Hans Peter; Mann, D. M. (1998):**
**The Phylogeny and Biogeography of *Thamnochortus* (Restionaceae).**
In: Botanical Journal of the Linnean Society 128 (4), p. 319–357. DOI: 10.1111/j.1095-8339.1998.tb02125.x.

**Moline, Philip M.; Linder, Hans Peter (2005):**
**Molecular Phylogeny and Generic Delimitation in the *Elegia* Group (Restionaceae, South Africa) based on a complete Taxon Sampling and four Chloroplast DNA Regions.**
In: Systematic Botany 30 (4), p. 759–772.

**Moline, Philip M.; Linder, Hans Peter (2006):**
**Input Data, analytical Methods and Biogeography of *Elegia* (Restionaceae).**
In: Journal of Biogeography 33 (1), p. 47–62. DOI: 10.1111/j.1365-2699.2005.01369.x.

**Wagstaff, Steven J.; Clarkson, Beverley R. (2012):**
**Systematics and Ecology of the Australasian Genus *Empodisma* (Restionaceae) and Description of a new Species from Peatlands in northern New Zealand.**
In: PhytoKeys (13), p. 39–79. DOI: 10.3897/phytokeys.13.3259.

### Rhamnaceae

**Aagesen, Lone; Medan, D.; Kellermann, Jürgen; Hilger, Hartmut H. (2005):**
**Phylogeny of the Tribe Colletieae (Rhamnaceae)? A sensitivity Analysis of the Plastid Region trnL-trnF combined with Morphology.**
In: Plant Systematics and Evolution 250 (3-4), p. 197–214. DOI: 10.1007/s00606-004-0204-5.

**Bolmgren, Kjell; Oxelman, Bengt (2004):**

**Generic limits in *Rhamnus* L. s.l. (Rhamnaceae) inferred from nuclear and Chloroplast DNA Sequence Phylogenies.**
In: Taxon 53 (2), p. 383–390.

BUERKI, SVEN; PHILLIPSON, PETER B.; CALLMANDER, MARTIN W. (2011):
**A taxonomic Revision of *Gouania* (Rhamnaceae) in Madagascar and the other Islands of the Western Indian Ocean (the Comoro and Mascarene Islands, and the Seychelles).**
In: Annals of the Missouri Botanical Garden 98 (2), p. 157–195. DOI: 10.3417/2007075.

BURGE, DYLAN O.; ERWIN, DIANE M.; ISLAM, MELISSA BAUER; KELLERMANN, JÜRGEN; KEMBEL, STEVEN W.; WILKEN, DIETER H.; MANOS, PAUL P. (2011):
**Diversification of *Ceanothus* (Rhamnaceae) in the California Floristic Province.**
In: International Journal of Plant Sciences 172 (9), p. 1137–1164. DOI: 10.1086/662028.

FAY, MICHAEL F.; LLED, MA D.; RICHARDSON, JAMES E.; RYE, BARBARA L.; HOPPER, STEPHEN D. (2001):
**Molecular Data confirm the Affinities of the South-west Australian endemic Granitites with *Alphitonia* (Rhamnaceae).**
In: Kew Bulletin 56, p. 669–675.

HARDIG, TERRY M.; SOLTIS, PAMELA S.; SOLTIS, DOUGLAS E. (2000):
**Diversification of the North American Shrub Genus *Ceanothus* (Rhamnaceae): Conflicting Phylogenies from Nuclear Ribosomal DNA and Chloroplast DNA.**
In: American Journal of Botany 87 (1), p. 108–123.

HAUENSCHILD, FRANK (2016):
**Corrigendum to Hauenschild, F. & al., Analysis of the cosmopolitan Buckthorn Genera *Frangula* and *Rhamnus* s.l. supports the Description of a new Genus.**
In: Taxon 65 (4), p. 926–927. DOI: 10.12705/654.49.

HAUENSCHILD, FRANK; FAVRE, ADRIEN; SALAZAR, GERARDO A.; MÜLLNER-RIEHL, ALEXANDRA N. (2016):
**Analysis of the cosmopolitan Buckthorn Genera *Frangula* and *Rhamnus* s.l. supports the Description of a new Genus, *Ventia*.**
In: Taxon 65 (1), p. 65–78. DOI: 10.12705/651.5.

HAUENSCHILD, FRANK; MATUSZAK, SABINE; MÜLLNER-RIEHL, ALEXANDRA N.; FAVRE, ADRIEN (2016):
**Phylogenetic Relationships within the cosmopolitan Buckthorn Family (Rhamnaceae) support the Resurrection of *Sarcomphalus* and the Description of *Pseudoziziphus* gen. nov.**
In: Taxon 65 (1), p. 47–64. DOI: 10.12705/651.4.

ISLAM, MELISSA BAUER; GURALNICK, ROBERT P. (2015):
**Generic Placement of the Former *Condaliopsis* (Rhamnaceae) Species.**
In: Phytotaxa 236 (1), p. 25. DOI: 10.11646/phytotaxa.236.1.2.

KELLERMANN, JÜRGEN; MEDAN, D.; AAGESEN, LONE; HILGER, HARTMUT H. (2005):
**Rehabilitation of the South American Genus *Ochetophila* Poepp. ex Endl. (Rhamnaceae: Colletieae).**
In: New Zealand Journal of Botany 43 (4), p. 865–869. DOI: 10.1080/0028825X.2005.9512996.

KELLERMANN, JÜRGEN; RYE, BARBARA L.; THIELE, KEVIN R. (2006):
***Polianthion*, a new Genus of Rhamnaceae (Pomaderreae) from Western Australia and Queensland.**
In: Australian Systematic Botany 19 (2), p. 169. DOI: 10.1071/SB05027.

KELLERMANN, JÜRGEN; UDOVICIC, FRANK; LADIGES, PAULINE Y. (2005):
**Phylogenetic Analysis and generic Limits of the Tribe Pomaderreae (Rhamnaceae) using Internal Transcribed Spacer DNA Sequences.**
In: Taxon 54 (3), p. 619–631.

ONSTEIN, RENSKE E.; CARTER, RICHARD J.; XING, YAOWU; RICHARDSON, JAMES E.; LINDER, HANS PETER (2015):
**Do Mediterranean-type Ecosystems have a common History? - Insights from the Buckthorn Family (Rhamnaceae).**
In: Evolution 69 (3), p. 756–771. DOI: 10.1111/evo.12605.

POOL, AMY (2013):
**New Species, Combinations, and Lectotypifications in Neotropical and Northern Mexican *Frangula* (Rhamnaceae).**
In: Novon: A Journal for Botanical Nomenclature 22 (4), p. 447–467. DOI: 10.3417/2013009.

RICHARDSON, JAMES E.; FAY, MICHAEL F.; CRONK, QUENTIN C.B.; BOWMAN, DIANE; CHASE, MARK W. (2000):
**A phylogenetic Analysis of Rhamnaceae using Rbcl and Trnl-f Plastid DNA Sequences.**
In: American Journal of Botany 87 (9), p. 1309–1324.

RICHARDSON, JAMES E.; WEITZ, FRANS M.; FAY, MICHAEL F.; CRONK, QUENTIN C.B.; LINDER, HANS PETER; REEVES, GAIL; CHASE, MARK W. (2001):
**Phylogenetic Analysis of *Phylica* L. (Rhamnaceae) with an Emphasis on Island species: Evidence from Plastid trnL-F and nuclear Internal Transcribed Spacer (ribosomal) DNA Sequences.**
In: Taxon 50 (2), p. 405–427.

### Rhizophoraceae

SCHWARZBACH, ANDREA E.; RICKLEFS, ROBERT E. (2000):
**Systematic affinities of Rhizophoraceae and Anisophylleaceae, and intergeneric Relationships within Rhizophoraceae, based on Chloroplast DNA, nuclear ribosomal DNA, and Morphology.**
In: American Journal of Botany 87 (4), p. 547–564. DOI: 10.2307/2656599.

SETOGUCHI, HIROAKI; KOSUGE, KEIKO; TOBE, HIROSHI (1999):
**Molecular Phylogeny of Rhizophoraceae based on rbcL Gene Sequences.**
In: Journal of Plant Research 112 (4), p. 443–455. DOI: 10.1007/PL00013899.

TSAI, CHI-CHU; LI, SHU-JU; SU, YU-YEN; YONG, JEAN W.H.; SAENGER, PETER; CHESSON, PETER ET AL. (2012):
**Molecular Phylogeny and Evidence for natural Hybridization and historical introgression between *Ceriops* Species (Rhizophoraceae).**
In: Biochemical Systematics and Ecology 43, p. 178–191. DOI: 10.1016/j.bse.2012.03.009.

### Rosaceae

ACOSTA, JUAN M.; SALARIATO, DIEGO LEONEL; CIALDELLA, ANA MARÍA (2016):
**Molecular Phylogeny and morphological Analysis of *Tetraglochin* (Rosaceae: Rosoideae: Sanguisorbeae) and Recognition of the new Species *T. andina*.**
In: Systematic Botany 41 (4), p. 839–850. DOI: 10.1600/036364416X693946.

ALICE, LAWRENCE A.; CAMPBELL, CHRISTOPHER P. (1999):
**Phylogeny of *Rubus* (Rosaceae) based on Nuclear Ribosomal DNA Internal Transcribed Spacer Region Sequences.**
In: American Journal of Botany 86 (1), p. 81–97.

**Bortiri, E.; Heuvel, B. Vanden; Potter, Daniel (2006):**
**Phylogenetic Analysis of Morphology in *Prunus* reveals extensive Homoplasy.**
In: Plant Systematics and Evolution 259 (1), p. 53–71. DOI: 10.1007/s00606-006-0427-8.

**Bruneau, Anne; Starr, Julian R.; Joly, Simon (2007):**
**Phylogenetic Relationships in the Genus *Rosa*: new Evidence from Chloroplast DNA Sequences and an appraisal of current Knowledge.**
In: Systematic Botany 32 (2), p. 366–378. DOI: 10.1600/036364407781179653.

**Burgess, Michael B.; Cushman, Kevin R.; Doucette, Eric T.; Frye, Christopher T.; Campbell, Christopher p. (2015):**
**Understanding diploid Diversity: A first Step in Unraveling polyploid, apomictic Complexity in *Amelanchier*.**
In: American Journal of Botany 102 (12), p. 2041–2057. DOI: 10.3732/ajb.1500330.

**Campbell, Christopher S.; Evans, Rodger C.; Morgan, D. R.; Dickinson, Timothy A.; Arsenault, Matthew P. (2007):**
**Phylogeny of Subtribe Pyrinae (formerly the Maloideae, Rosaceae): Limited Resolution of a complex evolutionary History.**
In: Plant Systematics and Evolution 266 (1-2), p. 119–145. DOI: 10.1007/s00606-007-0545-y.

**Chin, Siew-wai; Shaw, Joey; Haberle, Rosemarie C.; Wen, Jun; Potter, Daniel (2014):**
**Diversification of Almonds, Peaches, Plums and Cherries - molecular Systematics and biogeographic History of *Prunus* (Rosaceae).**
In: Molecular Phylogenetics and Evolution 76, p. 34–48. DOI: 10.1016/j.ympev.2014.02.024.

**Chin, Siew-wai; Wen, Jun; Johnson, Gabriel; Potter, Daniel (2010):**
**Merging *Maddenia* with the morphologically diverse *Prunus* (Rosaceae).**
In: Botanical Journal of the Linnean Society 164, p. 236–245.

**Chung, Kyong-Sook; Elisens, Wayne J.; Skvarla, John J. (2010):**
**Pollen Morphology and its phylogenetic Significance in Tribe Sanguisorbeae (Rosaceae).**
In: Plant Systematics and Evolution 285 (3-4), p. 139–148. DOI: 10.1007/s00606-009-0262-9.

**Danet, Frédéric (2016):**
**Le genre *Argentina* Hill (Rosaceae) en Nouvelle-Guinée: une Espèce et une Combinaison nouvelles.**
In: Adansonia 38 (2), p. 233–239. DOI: 10.5252/a2016n2a7.

**Depypere, Leander; Chaerle, Peter; Breyne, Peter; Vander Mijnsbrugge, Kristine; Goetghebeur, Paul (2009):**
**A combined morphometric and AFLP based Diversity Study challenges the Taxonomy of the European Members of the complex *Prunus* L. Section *Prunus*.**
In: Plant Systematics and Evolution 279 (1-4), p. 219–231. DOI: 10.1007/s00606-009-0158-8.

**Dobes, Christoph; Paule, Juraj (2010):**
**A comprehensive Chloroplast DNA-based Phylogeny of the Genus *Potentilla* (Rosaceae): Implications for its geographic Origin, Phylogeography and generic Circumscription.**
In: Molecular Phylogenetics and Evolution 56 (1), p. 156–175. DOI: 10.1016/j.ympev.2010.03.005.

**Eriksson, Torsten; Lundberg, Magnus; Töpel, Mats; Östensson, Pia; Smedmark, Jenny E.E. (2015):**
***Sibbaldia*: a molecular phylogenetic Study of a remarkably polyphyletic Genus in Rosaceae.**
In: Plant Systematics and Evolution 301 (1), p. 171–184. DOI: 10.1007/s00606-014-1063-3.

**Evans, Rodger C.; Campbell, Christopher p. (2002):**
**The Origin of the Apple Subfamily (Maloideae; Rosaceae) is clarified by DNA Sequence Data from duplicated GBSSI Genes.**
In: American Journal of Botany 89 (9), p. 1478–1484.

**Faghir, Marzieh Beygom; Attar, Farideh; Farazmand, Ali; Kazempour-Osaloo, Shahrokh (2014):**
**Phylogeny of the Genus *Potentilla* (Rosaceae) in Iran based on nrDNA ITS and cpDNA trnL-F Sequences with a Focus on Leaf and Style characters' Evolution.**
In: Turkish Journal of Botany 38, p. 417–429. DOI: 10.3906/bot-1303-67.

**Faghir, Marzieh Beygom; Attar, Farideh; Shavvon, Robabeh Shahi; Mehrmanesh, Atefeh (2015):**
**Pollen Morphology of the Genus *Alchemilla* L. (Rosaceae) in Iran.**
In: Turkish Journal of Botany 39, p. 267–279. DOI: 10.3906/bot-1406-23.

**Feng, Tao; Moore, Michael J.; Sun, Yanxia; Meng, Ai-Ping; Chu, Haijia; Li, Jian-Qiang; Wang, Heng-Chang (2015):**
**A new Species of *Argentina* (Rosaceae, Potentilleae) from Southeast Tibet, with Reference to the taxonomic Status of the Genus.**
In: Plant Systematics and Evolution 301 (3), p. 911–921. DOI: 10.1007/s00606-014-1125-6.

**Feng, Tao; Moore, Michael J.; Yan, Min-Hui; Sun, Yan-Xia; Zhang, Hua-Jie; Meng, Ai-Ping et al. (2017):**
**Phylogenetic Study of the Tribe Potentilleae (Rosaceae), with further Insight into the disintegration of *Sibbaldia*.**
In: Journal of Systematics and Evolution 55 (3), p. 177–191. DOI: 10.1111/jse.12243.

**Fougère-Danezan, Marie; Joly, Simon; Bruneau, Anne; Gao, Xin-Fen; Zhang, Li-Bing (2015):**
**Phylogeny and Biogeography of wild roses with specific attention to polyploids.**
In: Annals of Botany 115 (2), p. 275–291. DOI: 10.1093/aob/mcu245.

**Gehrke, Berit; Bräuchler, Christian; Romoleroux, K.; Lundberg, Magnus; Heubl, Günther; Eriksson, Torsten (2008):**
**Molecular Phylogenetics of *Alchemilla, Aphanes* and *Lachemilla* (Rosaceae) inferred from Plastid and nuclear Intron and spacer DNA Sequences, with Comments on generic Classification.**
In: Molecular Phylogenetics and Evolution 47 (3), p. 1030–1044. DOI: 10.1016/j.ympev.2008.03.004.

**Guo, Wei; Yu, Yi; Shen, Ru-Jiang; Liao, Wen-bo; Chin, Siew-wai; Potter, Daniel (2011):**
**A Phylogeny of *Photinia* sensu lato (Rosaceae) and related Genera based on nrITS and cpDNA Analysis.**
In: Plant Systematics and Evolution 291 (1-2), p. 91–102. DOI: 10.1007/s00606-010-0368-0.

**Joly, Simon; Bruneau, Anne (2006):**
**Incorporating Allelic Variation for Reconstructing the evolutionary History of Organisms from Multiple Genes: An Example from *Rosa* in North America.**
In: Systematic Biology 55 (4), p. 623–636. DOI: 10.1080/10635150600863109.

**Joly, Simon; Bruneau, Anne (2007):**
**Delimiting Species Boundaries in *Rosa* Sect. *Cinnamomeae* (Rosaceae) in Eastern North America.**
In: Systematic Botany 32 (4), p. 819–836.

**Kessler, Michael; Schmidt-Lebuhn, Alexander N. (2006):**
**Taxonomical and Distributional Notes on *Polylepis* (Rosaceae).**
In: Organisms Diversity and Evolution 6 (1), p. 67–69. DOI: 10.1016/j.ode.2005.04.001.

**Khan, Gulzar; Zhang, Fa-Qi; Gao, Qing-Bo; Fu, Peng-Cheng; Xing, Rui; Wang, Jiu-Li et al. (2016):**
**Phylogenetic Analyses of *Spiraea* (Rosaceae) distributed in the Qinghai-Tibetan Plateau and adjacent Regions: Insights from molecular Data.**
In: Plant Systematics and Evolution 302 (1), p. 11–21. DOI: 10.1007/s00606-015-1238-6.

**Koopman, Wim J. M.; Wissemann, Volker; Cock, Katrien; van Huylenbroeck, Johan; Riek, Jan de; Sabatino, Gerda J. H. et al. (2008):**
**AFLP Markers as a Tool to reconstruct complex Relationships: A Case Study in *Rosa* (Rosaceae).**
In: American Journal of Botany 95 (3), p. 353–366.

**Korotkova, Nadja; Parolly, Gerald; Khachatryan, Anahit; Ghulikyan, Lusine; Sargsyan, Harutyun; Akopian, Janna et al. (2018):**
**Towards resolving the evolutionary History of Caucasian Pears (*Pyrus*, Rosaceae)- Phylogenetic Relationships, Divergence Times and Leaf Trait Evolution.**
In: Journal of Systematics and Evolution 56 (1), p. 35–47. DOI: 10.1111/jse.12276.

**Lee, Chunghee; Hong, Suk-Pyo (2011):**
**Phylogenetic Relationships of the rare Korean monotypic endemic Genus *Pentactina* Nakai in the Tribe Spiraeeae (Rosaceae) based on molecular Data.**
In: Plant Systematics and Evolution 294 (3-4), p. 159–166. DOI: 10.1007/s00606-011-0457-8.

**Lee, Sangtae; Heo, Kyoung-In; Cho, Juhee; Lee, Chunghee; Chen, Wen-Li; Kim, Seung-Chul (2011):**
**New Insights into Pollen Morphology and its Implications in the Phylogeny of *Sanguisorba* L. (Rosaceae; Sanguisorbeae).**
In: Plant Systematics and Evolution 291 (3-4), p. 227–242. DOI: 10.1007/s00606-010-0384-0.

**Lee, William G.; Macmillan, B. H.; Partridge, T. R.; Lister, R.; Lloyd, Kelvin M. (2001):**
**Fruit features in relation to the Ecology and Distribution of *Acaena* (Rosaceae) Species in New Zealand.**
In: New Zealand Journal of Ecology 25 (1), p. 17–27.

**Li, Feifei; Fan, Qiang; Li, Qingyan; Chen, Sufang; Guo, Wei; Cui, Dafang; Liao, Wen-bo (2014):**
**Molecular Phylogeny of *Cotoneaster* (Rosaceae) inferred from nuclear ITS and multiple Chloroplast Sequences.**
In: Plant Systematics and Evolution 300 (6), p. 1533–1546. DOI: 10.1007/s00606-014-0980-5.

**Li, Meng; Ohi-Toma, Tetsuo; Gao, Yun-Dong; Xu, Bo; Zhu, Zhang-Ming; Ju, Wen-Bin; Gao, Xin-Fen (2017):**
**Molecular Phylogenetics and historical Biogeography of *Sorbus* sensu stricto (Rosaceae).**
In: Molecular Phylogenetics and Evolution 111, p. 76–86. DOI: 10.1016/j.ympev.2017.03.018.

**Liu, Xiao-Lin; Wen, Jun; Nie, Ze-long; Johnson, Gabriel; Liang, Zong-Suo; Chang, Zhao-Yang (2013):**
**Polyphyly of the *Padus* Group of *Prunus* (Rosaceae) and the Evolution of biogeographic Disjunctions between Eastern Asia and Eastern North America.**
In: Journal of Plant Research 126 (3), p. 351–361. DOI: 10.1007/s10265-012-0535-1.

**Lo Eugenia, Y.Y.; Stefanović, Saša; Dickinson, Timothy A. (2007):**
**Molecular Reappraisal of Relationships between *Crataegus* and *Mespilus* (Rosaceae, Pyreae)- Two Genera or One?**
In: Systematic Botany 32 (3), p. 596–616.

**Lo, Eugenia Y.Y.; Donoghue, Michael J. (2012):**

**Expanded phylogenetic and dating Analyses of the Apples and their Relatives (Pyreae, Rosaceae).**
In: Molecular Phylogenetics and Evolution 63 (2), p. 230–243. DOI: 10.1016/j.ympev.2011.10.005.

LO, EUGENIA Y.Y.; STEFANOVIĆ, SAŠA; CHRISTENSEN, KNUD IB; DICKINSON, TIMOTHY A. (2009):
**Evidence for genetic association between East Asian and western North American *Crataegus* L. (Rosaceae) and rapid Divergence of the eastern North American Lineages based on multiple DNA Sequences.**
In: Molecular Phylogenetics and Evolution 51 (2), p. 157–168. DOI: 10.1016/j.ympev.2009.01.018.

LUNDBERG, MAGNUS; TÖPEL, MATS; ERIKSEN, BENTE; NYLANDER, JOHAN A.A.; ERIKSSON, TORSTEN (2009):
**Allopolyploidy in Fragariinae (Rosaceae): comparing four DNA Sequence Regions, with comments on Classification.**
In: Molecular Phylogenetics and Evolution 51 (2), p. 269–280. DOI: 10.1016/j.ympev.2009.02.020.

MARTICORENA, ALICIA (2006):
**Revisión del Género *Acaena* (Rosaceae) en Chile.**
In: Annals of the Missouri Botanical Garden 93 (3), p. 412–454. DOI: 10.3417/0026-6493(2007)93[412:RDGARE]2.0.CO;2.

NJUGUNA, WAMBUI; LISTON, AARON; CRONN, RICHARD C.; ASHMAN, TIA-LYNN; BASSIL, NAHLA (2013):
**Insights into Phylogeny, sex Function and Age of *Fragaria* based on whole Chloroplast Genome Sequencing.**
In: Molecular Phylogenetics and Evolution 66 (1), p. 17–29. DOI: 10.1016/j.ympev.2012.08.026.

OH, SANG-HUN; POTTER, DANIEL (2003):
**Phylogenetic utility of the second Intron of Leafy in *Neillia* and *Stephanandra* (Rosaceae) and Implications for the Origin of *Stephanandra*.**
In: Molecular Phylogenetics and Evolution 29 (2), p. 203–215. DOI: 10.1016/S1055-7903(03)00093-9.

OH, SANG-HUN; POTTER, DANIEL (2005):
**Molecular phylogenetic Systematics and Biogeography of Tribe Neillieae (Rosaceae) using DNA Sequences of CpDNA, RDNA, and Leafy.**
In: Systematic Botany 92 (1), p. 179–192.

OHTA, S.; YAMAMOTO, T.; NISHITANI, C.; KATSUKI, T.; IKETANI, H.; OMURA, M. (2007):
**Phylogenetic Relationships among Japanese flowering Cherries (*Prunus* Subgenus *Cerasus*) based on nucleotide Sequences of Chloroplast DNA.**
In: Plant Systematics and Evolution 263 (3-4), p. 209–225. DOI: 10.1007/s00606-006-0474-1.

POTTER, DANIEL; ERIKSSON, TORSTEN; EVANS, RODGER C.; OH, SANG-HUN; SMEDMARK, JENNY E.E.; MORGAN, D. R. ET AL. (2007):
**Phylogeny and Classification of Rosaceae.**
In: Plant Systematics and Evolution 266 (1-2), p. 5–43. DOI: 10.1007/s00606-007-0539-9.

POTTER, DANIEL; STILL, P. M.; GREBENC, T.; BALLIAN, D.; BOŽIČ, G.; FRANJIÆ, J.; KRAIGHER, H. (2007):
**Phylogenetic Relationships in Tribe Spiraeeae (Rosaceae) inferred from nucleotide Sequence Data.**
In: Plant Systematics and Evolution 266 (1-2), p. 105–118. DOI: 10.1007/s00606-007-0544-z.

RIEK, JAN DE; COCK, KATRIEN; SMULDERS, MARINUS J. M.; NYBOM, HILDE (2013):
**AFLP-based population Structure Analysis as a Means to validate the complex Taxonomy of Dog-Roses (*Rosa* Section *Caninae*).**
In: Molecular Phylogenetics and Evolution 67 (3), p. 547–559. DOI: 10.1016/j.ympev.2013.02.024.

**Rousseau-Gueutin, Mathieu; Gaston, A.; Aïnouche, Abdelkader; Aïnouche, Malika L.; Olbricht, K.; Staudt, G. et al. (2009):**
**Tracking the evolutionary History of Polyploidy in *Fragaria* L. (Strawberry): new Insights from phylogenetic Analyses of low-copy nuclear Genes.**
In: Molecular Phylogenetics and Evolution 51 (3), p. 515–530. DOI: 10.1016/j.ympev.2008.12.024.

**Schmidt-Lebuhn, Alexander N.; Kessler, Michael; Kumar, Mahendra (2006):**
**Promiscuity in the Andes: Species Relationships in *Polylepis* (Rosaceae, Sanguisorbeae) based on AFLP and Morphology.**
In: Systematic Botany 31 (3), p. 547–559.

**Scott, Jun Wen; Berggren, T.; Lee, Chung-Hee; Ickert-Bond, Stefanie M.; Yi, Ting-Shuang; Yoo, Ki-Oug et al. (2008):**
**Phylogenetic Inferences in *Prunus* (Rosaceae) using Chloroplast ndhF and nuclear ribosomal ITS Sequences.**
In: Journal of Systematics and Evolution 46 (3), p. 322–332.

**Sennikov, Alexander N.; Kurtto, Arto (2017):**
**A phylogenetic checklist of *Sorbus* s.l. (Rosaceae) in Europe.**
In: Memoranda Societatis pro Fauna et Flora Fennica 93, p. 1–78.

**Sepp, Silvia; Bobrova, Vera K.; Troitsky, Alex K.; Glazunova, Klavdiya P. (2000):**
**Genetic polymorphism detected with Rapd Analysis and morphological Variability in some Microspecies of apomictic *Alchemilla*.**
In: Ann Bot Fennici 37, p. 105–123.

**Shaw, Joey; Small, Randall L. (2005):**
**Chloroplast DNA Phylogeny and Phylogeography of the North American Plums (*Prunus* Subgenus *Prunus* Section *Prunocerasus*, Rosaceae).**
In: American Journal of Botany 92 (12), p. 2011–2030.

**Shi, Shuo; Li, Jinlu; Sun, Jiahui; Yu, Jing; Zhou, Shiliang (2013):**
**Phylogeny and Classification of *Prunus* sensu lato (Rosaceae).**
In: Journal of Integrative Plant Biology 55 (11), p. 1069–1079. DOI: 10.1111/jipb.12095.

**Smedmark, Jenny E.E.; Eriksson, Torsten (2002):**
**Phylogenetic Relationships of *Geum* (Rosaceae) and Relatives inferred from the nrITS and trnL-trnF Regions.**
In: Systematic Botany 27 (2), p. 303–317.

**Smedmark, Jenny E.E.; Eriksson, Torsten; Bremer, Birgitta (2005):**
**Allopolyploid Evolution in Geinae (Colurieae: Rosaceae) – building reticulate Species Trees from bifurcating Gene Trees.**
In: Organisms Diversity and Evolution 5 (4), p. 275–283. DOI: 10.1016/j.ode.2004.12.003.

**Sochor, Michal; Vašut, Radim J.; Sharbel, Timothy F.; Trávníček, Bohumil (2015):**
**How just a few makes a lot: Speciation via Reticulation and Apomixis on Example of European Brambles (*Rubus* subgen. *Rubus*, Rosaceae).**
In: Molecular Phylogenetics and Evolution 89, p. 13–27. DOI: 10.1016/j.ympev.2015.04.007.

**Soják, Jiří (2008):**
**Notes on *Potentilla* XXI. A new division of the Tribe Potentilleae (Rosaceae) and Notes on generic Delimitations.**
In: Botanische Jahrbücher für Systematik, Pflanzengeschichte und Pflanzengeographie 127, p. 349–358.

**Töpel, Mats; Antonelli, Alexandre; Yesson, Chris; Eriksen, Bente (2012):**
**Past climate Change and plant Evolution in Western North America: a case Study in Rosaceae.**
In: Public Library of Science One 7 (12), e50358. DOI: 10.1371/journal.pone.0050358.

**Vieira, Jorge; Ferreira, Pedro G.; Aguiar, Bruno; Fonseca, Nuno A.; Vieira, Cristina P. (2010):**
**Evolutionary Patterns at the RNase based gametophytic Self-incompatibility System in two divergent Rosaceae Groups (Maloideae and Prunus).**
In: BMC Evolutionary Biology 10, p. 200. DOI: 10.1186/1471-2148-10-200.

**Wissemann, Volker; Ritz, Christine M. (2005):**
**The Genus *Rosa* (Rosoideae, Rosaceae) revisited: molecular Analysis of nrITS-1 and atpB-rbc L intergenic spacer (Igs) versus conventional Taxonomy.**
In: Botanical Journal of the Linnean Society 147, p. 275–290.

**Wissemann, Volker; Ritz, Christine M. (2007):**
**Evolutionary Patterns and processes in the Genus *Rosa* (Rosaceae) and their Implications for host-parasite Co-Evolution.**
In: Plant Systematics and Evolution 266 (1-2), p. 79–89. DOI: 10.1007/s00606-007-0542-1.

**Xu, Qiang; Wen, Xiaopeng; Deng, Xiuxin (2007):**
**Phylogenetic and evolutionary Analysis of NBS-encoding Genes in Rosaceae Fruit Crops.**
In: Molecular Phylogenetics and Evolution 44 (1), p. 315–324. DOI: 10.1016/j.ympev.2006.12.029.

**Yang, Ji Young; Pak, Jae-Hong (2006):**
**Phylogeny of Korean *Rubus* (Rosaceae) based on its (nrDNA) and trnL/F intergenic Region (cpDNA).**
In: Journal of Plant Biology 49 (1), p. 44–54.

**Yazbek, M.; Oh, Sang-Hun (2013):**
**Peaches and Almonds: Phylogeny of *Prunus* Subg. *Amygdalus* (Rosaceae) based on DNA Sequences and Morphology.**
In: Plant Systematics and Evolution 299 (8), p. 1403–1418. DOI: 10.1007/s00606-013-0802-1.

**Yi-Hua, Tong; Nian-HeI.A., X. (2016):**
**New Combinations for Chinese *Argentina* Hill (Rosaceae).**
In: Journal of Tropical and Subtropical Botany 24 (4), p. 426–428.

**Zamani, Asghar; Attar, Farideh; Maroofi, Hosein (2012):**
**A Synopsis of the Genus *Pyrus* (Rosaceae) in Iran.**
In: Nordic Journal of Botany 30, p. 310–332.

**Zhang, Zhaoyang; Fan, Liming; Yang, Junbo; Hao, Xiaojiang; Gu, Zhijian (2006):**
**Alkaloid Polymorphism and ITS Sequence Variation in the *Spiraea japonica* Complex (Rosaceae) in China: Traces of the Biological Effects of the Himalaya-Tibet Plateau Uplift.**
In: American Journal of Botany 93 (5), p. 762–769.

**Zheng, Xiaoyan; Cai, Danying; Potter, Daniel; Postman, Joseph; Liu, Jing; Teng, Yuanwen (2014):**
**Phylogeny and evolutionary histories of *Pyrus* L. revealed by phylogenetic Trees and networks based on Data from multiple DNA Sequences.**
In: Molecular Phylogenetics and Evolution 80, p. 54–65. DOI: 10.1016/j.ympev.2014.07.009.

**Zheng, Xiaoyan; Hu, Chunyun; Spooner, David M.; Liu, Jing; Cao, Jiashu; Teng, Yuanwen (2011):**

**Molecular Evolution of Adh and Leafy and the phylogenetic Utility of their Introns in *Pyrus* (Rosaceae).**
In: BMC Evolutionary Biology 11, p. 255. DOI: 10.1186/1471-2148-11-255.

ZHU, ZHANG-MING; GAO, XIN-FEN; FOUGÈRE-DANEZAN, MARIE (2015):
**Phylogeny of *Rosa* Sections *Chinenses* and *Synstylae* (Rosaceae) based on Chloroplast and nuclear Markers.**
In: Molecular Phylogenetics and Evolution 87, p. 50–64. DOI: 10.1016/j.ympev.2015.03.014.

### Rousseaceae

PILLON, YOHAN; HOPKINS, HELEN C.F.; BARRABÉ, LAURE; STACY, E. A. (2014):
**A new record for *Carpodetus* (Rousseaceae) in Vanuatu.**
In: New Zealand Journal of Botany 52 (4), p. 449–452. DOI: 10.1080/0028825X.2014.928332.

### Rubiaceae

ACHILLE, FRÉDÉRIC; MOTLEY, TIMOTHY J.; LOWRY, PORTER PRESCOTT; JÉRÉMIE, JOËL (2006):
**Polyphyly in *Guettarda* L. (Rubiaceae, Guettardeae) based on nrDNA ITS Sequence Data.**
In: Annals of the Missouri Botanical Garden 93 (1), p. 103–121. DOI: 10.3417/0026-6493(2006)93[103:PIGLRG]2.0.CO;2.

ALEJANDRO, GRECEBIO JONATHAN D.; MEVE, ULRICH; MOULY, ARNAUD; THIV, MIKE; LIEDE-SCHUMANN, SIGRID (2011):
**Molecular Phylogeny and taxonomic Revision of the Philippine endemic *Villaria* Rolfe (Rubiaceae).**
In: Plant Systematics and Evolution 296 (1-2), p. 1–20. DOI: 10.1007/s00606-011-0472-9.

ALEJANDRO, GRECEBIO JONATHAN D.; MEVE, ULRICH; UY, MILLARD; MOULY, ARNAUD; THIV, MIKE; LIEDE-SCHUMANN, SIGRID (2010):
**Molecular support of the Classification of *Greeniopsis* Merr. in Aleisanthieae (Rubiaceae), with a Revision of the Genus.**
In: Taxon 59 (5), p. 1547–1564.

ALEJANDRO, GRECEBIO JONATHAN D.; RAZAFIMANDIMBISON, SYLVAIN G.; LIEDE-SCHUMANN, SIGRID (2005):
**Polyphyly of *Mussaenda* inferred from ITS and trnT-F Data and ITS Implication for Generic Limits in Mussaendeae (Rubiaceae).**
In: American Journal of Botany 92 (3), p. 544–557.

ANDERSON, CAJSA LISA; ROVA, JOHAN H.E.; ANDERSSON, LENNART (2001):
**Molecular Phylogeny of the Tribe Anthospermeae (Rubiaceae): Systematic and biogeographic Implications.**
In: Australian Systematic Botany 14 (2), p. 231–244. DOI: 10.1071/SB00021.

ANDERSSON, LENNART (2002):
**Relationships and Generic Circumscriptions in the *Psychotria* Complex (Rubiaceae, Psychotrieae).**
In: Systematics and Geography of Plants 72 (1-2), p. 167–202.

ANDERSSON, LENNART; ANTONELLI, ALEXANDRE (2005):
**Phylogeny of the Tribe Cinchoneae (Rubiaceae), its Position in Cinchonoideae, and Description of a new Genus, *Ciliosemina*.**
In: Taxon 54 (1), p. 17–28.

ANDREASEN, KATARINA; BREMER, BIRGITTA (2000):

**Combined phylogenetic Analysis in the Rubiaceae-Ixoroideae: Morphology, Nuclear and Chloroplast DNA Data.**
In: American Journal of Botany 87 (11), p. 1731–1748.

Backlund, Maria; Bremer, Birgitta; Thulin, Mats (2007):
**Paraphyly of Paederieae, Recognition of Putorieae and Expansion of *Plocama* (Rubiaceae-Rubioideae).**
In: Taxon 56 (2), p. 315–328.

Barrabé, Laure; Buerki, Sven; Mouly, Arnaud; Davis, Aaron P.; Munzinger, Jérôme; Maggia, Laurent (2012):
**Delimitation of the Genus *Margaritopsis* (Rubiaceae) in the Asian, Australasian and Pacific Region, based on molecular phylogenetic Inference and Morphology.**
In: Taxon 61 (6), p. 1251–1268.

Barrabé, Laure; Maggia, Laurent; Pillon, Yohan; Rigault, Frédéric; Mouly, Arnaud; Davis, Aaron P.; Buerki, Sven (2014):
**New Caledonian Lineages of *Psychotria* (Rubiaceae) reveal different evolutionary Histories and the largest documented Plant Radiation for the Archipelago.**
In: Molecular Phylogenetics and Evolution 71, p. 15–35. DOI: 10.1016/j.ympev.2013.10.020.

Barrabé, Laure; Mouly, Arnaud; Lowry, Porter Prescott; Munzinger, Jérôme (2011):
**Reinstatement of the endemic New Caledonian Genus *Thiollierea* Montrouz. (Rubiaceae) necessitated by the Polyphyly of *Bikkia* Reinw. as currently circumscribed.**
In: Adansonia 33 (1), p. 115–134. DOI: 10.5252/a2011n1a8.

Block, Petra J.; Razafimandimbison, Sylvain G.; Janssens, Steven B.; Ochoterena, Helga; Robbrecht, Elmar; Bremer, Birgitta (2015):
**Molecular Phylogenetics and generic assessment in the Tribe Pavetteae (Rubiaceae).**
In: Taxon 64 (1), p. 79–95. DOI: 10.12705/641.19.

Borhidi, A.; Darók, J.; Stranczinger, Szilvia (2011):
***Donnellyanthus* (Rubiaceae, Rondeletieae), a new Genus in the Flora of Mexico and Meso-America.**
In: Acta Botanica Hungarica 53 (3-4), p. 273–281. DOI: 10.1556/ABot.53.2011.3–4.9.

Borhidi, A.; Lozada-Pérez, L. (2011):
***Martensianthus* nomen novum to replace *Neomartensia* Borhidi et Lozada-Pérez 2010 (Rubiaceae) non Yoshida et Mikami 1996 (Delesseriaceae).**
In: Acta Botanica Hungarica 53 (1-2), p. 37–42.

Bremer, Birgitta (2009):
**A Review of molecular phylogenetic Studies of Rubiaceae.**
In: Annals of the Missouri Botanical Garden 96 (1), p. 4–26. DOI: 10.3417/2006197.

Bremer, Birgitta; Eriksson, Torsten (2009):
**Time Tree of Rubiaceae: Phylogeny and Dating the Family, Subfamilies, and Tribes.**
In: International Journal of Plant Sciences 170 (6), p. 766–793. DOI: 10.1086/599077.

Cabaña Fader, Andrea A.; Salas, Roberto Manuel; Dessein, Steven; Cabral, Elsa L. (2016):
**Synopsis of *Hexasepalum* (Rubiaceae), the Priority Name for *Diodella* and a new Species from Brazil.**
In: Systematic Botany 41 (2), p. 408–422. DOI: 10.1600/036364416X691803.

Church, Sheri A. (2003):

**Molecular Phylogenetics of *Houstonia* (Rubiaceae): descending aneuploidy and breeding system Evolution in the Radiation of the Lineage across North America.**
In: Molecular Phylogenetics and Evolution 27 (2), p. 223–238. DOI: 10.1016/S1055-7903(02)00446-3.

CHURCH, SHERI A.; TAYLOR, DOUGLAS R. (2005):
**Speciation and Hybridization among *Houstonia* (Rubiaceae) Species: the Influence of Polyploidy on Reticulate Evolution.**
In: American Journal of Botany 92 (8), p. 1372–1380.

CORTÉS-B., ROCIO; DELPRETE, PIERO G.; MOTLEY, TIMOTHY J. (2009):
**Phylogenetic Placement of the Tribe Retiniphylleae among the Subfamily Ixoroideae (Rubiaceae).**
In: Annals of the Missouri Botanical Garden 96 (1), p. 61–67. DOI: 10.3417/2006198.

CORTÉS-B., ROCIO; MOTLEY, TIMOTHY J. (2015):
**Phylogeny of the Henriquezieae-Posoquerieae-Sipaneeae, a Guayanan-centered Clade of Rubiaceae: Implications for morphological Evolution.**
In: Phytotaxa 206 (1), p. 90. DOI: 10.11646/phytotaxa.206.1.12.

DAVIS, AARON P.; CHESTER, MICHAEL; MAURIN, OLIVIER (2007):
**Searching for the relatives of *Coffea* (Rubiaceae, Ixoroideae): the Circumscription and Phylogeny of Coffeeae based on Plastid Sequence Data and Morphology.**
In: American Journal of Botany 94 (3), p. 313–329.

DELPRETE, PIERO G.; CORTÉS-B., ROCIO (2004):
**A phylogenetic Study of the Tribe Sipaneeae (Rubiaceae, Ixoroideae), using trnL-F and ITS Sequence Data.**
In: Taxon 53 (2), p. 347–356.

DELPRETE, PIERO G.; PERSSON, CLAES (2012):
***Octavia sessiliflora* DC. and *Mussaenda glomerulata* Lam. ex Poir., two obscure Taxa from French Guiana synonymous with Members of the *Alibertia* Group (Rubiaceae, Gardenieae).**
In: Adansonia 34 (2), p. 353–363. DOI: 10.5252/a2012n2a9.

DENG, YUN-FEI (2007):
***Fleroya*, a substitute name for *Hallea* J.-f. Leroy (Rubiaceae).**
In: Taxon 56 (1), p. 247–248.

DUAN, TINGTING; DENG, XIAOFANG; CHEN, SHI; LUO, ZHONGLAI; ZHAO, ZHONGTAO; TU, TIE-YAO ET AL. (2018):
**Evolution of sexual systems and growth habit in *Mussaenda* (Rubiaceae): Insights into the evolutionary pathways of Dioecy.**
In: Molecular Phylogenetics and Evolution 123, p. 113–122. DOI: 10.1016/j.ympev.2018.02.015.

GARGIULO, ROBERTA; GUACCHIO, EMANUELE DEL; CAPUTO, PAOLO (2015):
**Phylogenetic Reconstruction of *Asperula* Sect. *Cynanchicae* (Rubiaceae) reveals a Mosaic of evolutionary Histories.**
In: Taxon 64 (4), p. 754–769. DOI: 10.12705/644.7.

GINTER, ANNA; RAZAFIMANDIMBISON, SYLVAIN G.; BREMER, BIRGITTA (2015):
**Phylogenetic affinities of *Myrioneuron* and *Cyanoneuron*, generic Limits of the Tribe Argostemmateae and Description of a new Asian Tribe, Cyanoneuroneae (Rubiaceae).**
In: Taxon 64 (2), p. 286–298. DOI: 10.12705/642.2.

**Groeninckx, Inge; Dessein, Steven; Ochoterena, Helga; Persson, Claes; Motley, Timothy J.; Kårehed, Jesper et al. (2009):**

**Phylogeny of the Herbaceous Tribe Spermacoceae (Rubiaceae) based on Plastid DNA Data.**

In: Annals of the Missouri Botanical Garden 96 (1), p. 109–132. DOI: 10.3417/2006201.

**Groeninckx, Inge; Ochoterena, Helga; Smets, Erik; Dessein, Steven (2010):**

**Molecular phylogenetic and morphological Study of *Kohautia* (Spermacoceae, Rubiaceae), with the Recognition of the new Genus *Cordylostigma*.**

In: Taxon 59 (5), p. 1457–1471.

**Guo, Xing; Wang, Rui-Jiang; Simmons, Mark P.; But, Paul Pui-Hay; Yu, Jing (2013):**

**Phylogeny of the Asian *Hedyotis-Oldenlandia* complex (Spermacoceae, Rubiaceae): Evidence for high Levels of Polyphyly and the parallel Evolution of diplophragmous Capsules.**

In: Molecular Phylogenetics and Evolution 67 (1), p. 110–122. DOI: 10.1016/j.ympev.2013.01.006.

**Gustafsson, Claes; Persson, Claes (2002):**

**Phylogenetic Relationships among Species of the Neotropical Genus *Randia* (Rubiaceae, Gardenieae) inferred from molecular and morphological Data.**

In: Taxon 51, p. 661–674.

**Hamon, Perla; Grover, Corrinne E.; Davis, Aaron P.; Rakotomalala, Jean-Jacques; Raharimalala, Nathalie E.; Albert, Victor A. et al. (2017):**

**Genotyping-by-Sequencing provides the first well-resolved Phylogeny for Coffee (*Coffea*) and Insights into the Evolution of caffeine Content in its species: GBS Coffee Phylogeny.**

In: Molecular Phylogenetics and Evolution 109, p. 351–361. DOI: 10.1016/j.ympev.2017.02.009.

**Huang, Wei-Ping; Sun, Hang; Deng, Tao; Razafimandimbison, Sylvain G.; Nie, Ze-long; Wen, Jun (2013):**

**Molecular Phylogenetics and Biogeography of the eastern Asian–eastern North American disjunct *Mitchella* and its close relative *Damnacanthus* (Rubiaceae, Mitchelleae).**

In: Botanical Journal of the Linnean Society 171, p. 395–412.

**Jalaluddin, Susan; Bruhl, Jeremy J. (2008):**

**Testing Species limits in *Rennellia* (Prismatomerideae, Rubioideae, Rubiaceae).**

In: Taxon 57 (1), p. 43–52.

**Janssens, Steven B.; Groeninckx, Inge; Block, Petra J.; Verstraete, Brecht; Smets, Erik F.; Dessein, Steven (2016):**

**Dispersing towards Madagascar: Biogeography and Evolution of the Madagascan endemics of the Spermacoceae Tribe (Rubiaceae).**

In: Molecular Phylogenetics and Evolution 95, p. 58–66. DOI: 10.1016/j.ympev.2015.10.024.

**Kainulainen, Kent; Bremer, Birgitta (2014):**

**Phylogeny of *Euclinia* and allied Genera of Gardenieae (Rubiaceae), and Description of *Melanoxerus*, an endemic Genus of Madagascar.**

In: Taxon 63 (4), p. 819–830. DOI: 10.12705/634.2.

**Kainulainen, Kent; Mouly, Arnaud; Khodabandeh, Anbar; Bremer, Birgitta (2009):**

**Molecular phylogenetic Analysis of the Tribe Alberteae (Rubiaceae), with Description of a new Genus, *Razafimandimbisonia*.**

In: Taxon 58 (3), p. 757–768.

**Kainulainen, Kent; Persson, Claes; Eriksson, Torsten; Bremer, Birgitta (2010):**
**Molecular Systematics and morphological Character Evolution of the Condamineeae (Rubiaceae).**
In: American Journal of Botany 97 (12), p. 1961–1981. DOI: 10.3732/ajb.1000090.

**Kainulainen, Kent; Razafimandimbison, Sylvain G. (2016):**
**New taxonomic Combinations in West Indian Ocean Vanguerieae (Rubiaceae).**
In: Phytotaxa 282 (4), p. 267. DOI: 10.11646/phytotaxa.282.4.3.

**Kainulainen, Kent; Razafimandimbison, Sylvain G.; Bremer, Birgitta (2013):**
**Phylogenetic Relationships and new tribal Delimitations in Subfamily Ixoroideae (Rubiaceae).**
In: Botanical Journal of the Linnean Society 173, p. 387–406.

**Kårehed, Jesper; Bremer, Birgitta (2007):**
**The Systematics of Knoxieae (Rubiaceae) - molecular Data and their taxonomic Consequences.**
In: Taxon 56 (4), p. 1051–1076.

**Kårehed, Jesper; Groeninckx, Inge; Dessein, Steven; Motley, Timothy J.; Bremer, Birgitta (2008):**
**The phylogenetic utility of Chloroplast and nuclear DNA Markers and the Phylogeny of the Rubiaceae Tribe Spermacoceae.**
In: Molecular Phylogenetics and Evolution 49 (3), p. 843–866. DOI: 10.1016/j.ympev.2008.09.025.

**Khan, Saleh Ahammad; Razafimandimbison, Sylvain G.; Bremer, Birgitta; Liede-Schumann, Sigrid (2008):**
**Phylogeny and Biogeography of the African Genus *Virectaria* Bremek. (Sabiceeae s.l., Ixoroideae, Rubiaceae).**
In: Plant Systematics and Evolution 275 (1-2), p. 43–58. DOI: 10.1007/s00606-008-0053-8.

**Khan, Saleh Ahammad; Razafimandimbison, Sylvain G.; Bremer, Birgitta; Liede-Schumann, Sigrid (2008):**
**Sabiceeae and Virectarieae (Rubiaceae, Ixoroideae): one or two Tribes? New tribal and generic Circumscriptions of Sabiceeae and Biogeography of Sabicea s.l.**
In: Taxon 57 (1), p. 7–23.

**Kirkbride, Joseph H.; Delprete, Piero G., JR. (2015):**
**New Combinations in *Hexasepalum* (Rubiaceae: Spermacoceae).**
In: Journal of the Botanical Research Institute of Texas 9 (1), p. 103–106.

**Krüger, Åsa; Razafimandimbison, Sylvain G.; Bremer, Birgitta (2012):**
**Molecular Phylogeny of the Tribe Danaideae (Rubiaceae: Rubioideae): Another Example of out-of-Madagascar Dispersal.**
In: Taxon 61 (3), p. 629–636.

**Lantz, Henrik; Bremer, Birgitta (2004):**
**Phylogeny inferred from Morphology and DNA Data: Characterizing well-supported Groups in Vanguerieae (Rubiaceae).**
In: Botanical Journal of the Linnean Society 146, p. 257–283.

**Lantz, Henrik; Bremer, Birgitta (2005):**
**Phylogeny of the complex Vanguerieae (Rubiaceae) Genera *Fadogia, Rytigynia,* and *Vangueria* with close relatives and a new Circumscription of *Vangueria*.**
In: Plant Systematics and Evolution 253 (1-4), p. 159–183. DOI: 10.1007/s00606-005-0313-9.

**Löfstrand, Stefan D.; Krüger, Åsa; Razafimandimbison, Sylvain G.; Bremer, Birgitta (2014):**

**Phylogeny and Generic Delimitations in the Sister Tribes Hymenodictyeae and Naucleeae (Rubiaceae).**
In: Systematic Botany 39 (1), p. 304–315. DOI: 10.1600/036364414X678116.

MALCOMBER, SIMON T.; TAYLOR, CHARLOTTE M. (2009):
**A Systematic Revision of *Gaertnera* (Rubiaceae, Gaertnereae).**
In: Annals of the Missouri Botanical Garden 96 (4), p. 575–671. DOI: 10.3417/2002161.

MANNS, ULRIKA; WIKSTRÖM, NIKLAS; TAYLOR, CHARLOTTE M.; BREMER, BIRGITTA (2012):
**Historical Biogeography of the predominantly Neotropical Subfamily Cinchonoideae (Rubiaceae): into or out of America?**
In: International Journal of Plant Sciences 173 (3), p. 261–286. DOI: 10.1086/663971.

MARTINEZ-CAMILO, RUBEN; MARTÍNEZ-MELENDEZ, NAYELY; PÉREZ-FARRERA, MIGUEL ANGEL; LORENCE, DAVID H. (2011):
**Una nueva Especie de *Pseudomiltemia* (Rubiaceae) de Chiapas, México.**
In: Brittonia 63 (2), p. 197–202.

MAURIN, OLIVIER; DAVIS, AARON P.; CHESTER, MICHAEL; MVUNGI, ESTHER F.; JAUFEERALLY-FAKIM, YASMINA; FAY, MICHAEL F. (2007):
**Towards a Phylogeny for *Coffea* (Rubiaceae): identifying well-supported Lineages based on nuclear and Plastid DNA Sequences.**
In: Annals of Botany 100 (7), p. 1565–1583. DOI: 10.1093/aob/mcm257.

MCDOWELL, TIM; VOLOVSEK, MIGUEL; MANOS, PAUL P. (2003):
**Biogeography of *Exostema* (Rubiaceae) in the Caribbean Region in Light of molecular phylogenetic Analyses.**
In: Systematic Botany 28 (2), p. 431–441.

MOTLEY, TIMOTHY J.; WURDACK, KENNETH J.J.; DELPRETE, PIERO G. (2005):
**Molecular Systematics of the Catesbaeeae-Chiococceae Complex (Rubiaceae): Flower and Fruit Evolution and Biogeographic Implications.**
In: American Journal of Botany 92 (2), p. 316–329.

MOULY, ARNAUD; KAINULAINEN, KENT; PERSSON, CLAES; DAVIS, AARON P.; WONG, KHOON MENG; RAZAFIMANDIMBISON, SYLVAIN G.; BREMER, BIRGITTA (2014):
**Phylogenetic Structure and Clade Circumscriptions in the Gardenieae complex (Rubiaceae).**
In: Taxon 63 (4), p. 801–818. DOI: 10.12705/634.4.

MOULY, ARNAUD; RAZAFIMANDIMBISON, SYLVAIN G.; FLORENCE, JACQUES; JÉRÉMIE, JOËL; BREMER, BIRGITTA (2009):
**Paraphyly of *Ixora* and new Tribal Delimitation of Ixoreae (Rubiaceae): Inference from Combined Chloroplast (RPS16, RBCL, and trnT-F) Sequence Data.**
In: Annals of the Missouri Botanical Garden 96 (1), p. 146–160. DOI: 10.3417/2006194.

MOULY, ARNAUD; RAZAFIMANDIMBISON, SYLVAIN G.; KHODABANDEH, ANBAR; BREMER, BIRGITTA (2009):
**Phylogeny and Classification of the species-rich pantropical showy Genus *Ixora* (Rubiaceae-Ixoreae) with Indications of geographical monophyletic Units and Hybrids.**
In: American Journal of Botany 96 (3), p. 686–706. DOI: 10.3732/ajb.0800235.

NAKAMURA, KOH; CHUNG, SHIH-WEN; KOKUBUGATA, GORO; DENDA, TETSUO; YOKOTA, MASATSUGU (2006):
**Phylogenetic Systematics of the monotypic Genus *Hayataella* (Rubiaceae) endemic to Taiwan.**
In: Journal of Plant Research 119 (6), p. 657–661. DOI: 10.1007/s10265-006-0017-4.

NEGRÓN-ORTIZ, V.; WATSON, LINDA E. (2002):
**Molecular Phylogeny and Biogeography of *Erithalis* (Rubiaceae), an Endemic of the Caribbean Basin.**
In: Plant Systematics and Evolution 234 (1), p. 71–83. DOI: 10.1007/s00606-002-0192-2.

NEPOKROEFF, MOLLY; BREMER, BIRGITTA; SYTSMA, KENNETH J. (1999):
**Reorganization of the Genus *Psychotria* and Tribe Psychotrieae (Rubiaceae) inferred from ITS and rbcL Sequence Data.**
In: Systematic Botany 24 (1), p. 5–27.

NEPOKROEFF, MOLLY; SYTSMA, KENNETH J.; WAGNER, WARREN L.; ZIMMER, ELIZABETH A. (2003):
**Reconstructing Ancestral Patterns of Colonization and Dispersal in the Hawaiian Understory Tree Genus *Psychotria* (Rubiaceae): A Comparison of Parsimony and Likelihood Approaches.**
In: Systematic Biology 52 (6), p. 820–838. DOI: 10.1080/10635150390251072.

NEUPANE, SUMAN; DESSEIN, STEVEN; WIKSTRÖM, NIKLAS; LEWIS, PAUL O.; LONG, CHUNLIN; BREMER, BIRGITTA; MOTLEY, TIMOTHY J. (2015):
**The *Hedyotis-Oldenlandia* complex (Rubiaceae: Spermacoceae) in Asia and the Pacific: Phylogeny revisited with new generic Delimitations.**
In: Taxon 64 (2), p. 299–322. DOI: 10.12705/642.8.

OGURI, EMIKO; SUGAWARA, TAKASHI; PENG, CHING-I.; YANG, T. Y. ALECK; MURAKAMI, NORIAKI (2013):
**Geographical Origin and sexual-system Evolution of the androdioecious Plant *Gynochthodes boninensis* (Rubiaceae), endemic to the Bonin Islands, Japan.**
In: Molecular Phylogenetics and Evolution 68 (3), p. 699–708. DOI: 10.1016/j.ympev.2013.04.019.

OLIVEIRA, CAETANO TRONCOSO; GIACOMIN, LEANDRO L.; ZAPPI, DANIELA C. (2011):
***Kerianthera longiflora* (Rubiaceae), a remarkable new Species from eastern Brazil, with some observations on *K. preclara*.**
In: Kew Bulletin 66, p. 143–148.

PERSSON, CLAES (2000):
**Phylogeny of the Neotropical *Alibertia* Group (Rubiaceae), with Emphasis on the Genus *Alibertia*, inferred from ITS and 5s Ribosomal DNA Sequences.**
In: American Journal of Botany 87 (7), p. 1018–1028.

RAZAFIMANDIMBISON, SYLVAIN G.; KAINULAINEN, KENT; WONG, KHOON MENG; BEAVER, KATY; BREMER, BIRGITTA (2011):
**Molecular support for a basal Grade of morphologically distinct, monotypic Genera in the species-rich Vanguerieae alliance (Rubiaceae, Ixoroideae): Its systematic and conservation Implications.**
In: Taxon 60 (4), p. 941–952.

RAZAFIMANDIMBISON, SYLVAIN G.; KELLOGG, ELIZABETH A.; BREMER, BIRGITTA (2004):
**Recent Origin and phylogenetic utility of divergent ITS putative Pseudogenes: a case Study from Naucleeae (Rubiaceae).**
In: Systematic Biology 53 (2), p. 177–192. DOI: 10.1080/10635150490423278.

RAZAFIMANDIMBISON, SYLVAIN G.; LANTZ, HENRIK; MOULY, ARNAUD; BREMER, BIRGITTA (2009):
**Evolutionary Trends, major Lineages, and new generic Limits in the dioecious Group of the Tribe Vanguerieae (Rubiaceae): Insights into the Evolution of functional Dioecy.**
In: Annals of the Missouri Botanical Garden 96 (1), p. 161–181. DOI: 10.3417/2006191.

**Razafimandimbison, Sylvain G.; McDowell, Timothy D.; Halford, David A.; Bremer, Birgitta (2009):**
**Molecular Phylogenetics and generic assessment in the Tribe Morindeae (Rubiaceae-Rubioideae): how to circumscribe *Morinda* L. to be monophyletic?**
In: Molecular Phylogenetics and Evolution 52 (3), p. 879–886. DOI: 10.1016/j.ympev.2009.04.007.

**Razafimandimbison, Sylvain G.; Moog, Joachim; Lantz, Henrik; Maschwitz, Ulrich; Bremer, Birgitta (2005):**
**Re-assessment of Monophyly, Evolution of Myrmecophytism, and rapid Radiation in *Neonauclea* s.s. (Rubiaceae).**
In: Molecular Phylogenetics and Evolution 34 (2), p. 334–354. DOI: 10.1016/j.ympev.2004.10.005.

**Razafimandimbison, Sylvain G.; Rydin, Catarina; Bremer, Birgitta (2008):**
**Evolution and Trends in the Psychotrieae alliance (Rubiaceae)- a rarely reported evolutionary Change of many-seeded Carpels from one-seeded Carpels.**
In: Molecular Phylogenetics and Evolution 48 (1), p. 207–223. DOI: 10.1016/j.ympev.2008.03.034.

**Razafimandimbison, Sylvain G.; Taylor, Charlotte M.; Wikström, Niklas; Pailler, Thierry; Khodabandeh, Anbar; Bremer, Birgitta (2014):**
**Phylogeny and generic limits in the sister Tribes Psychotrieae and Palicoureeae (Rubiaceae): Evolution of Schizocarps in *Psychotria* and Origins of bacterial Leaf Nodules of the Malagasy Species.**
In: American Journal of Botany 101 (7), p. 1102–1126. DOI: 10.3732/ajb.1400076.

**Rova, Johan H.E.; Delprete, Piero G.; Bremer, Birgitta (2009):**
**The *Rondeletia* Complex (Rubiaceae): An attempt to use ITS, rps16, and trnL-F Sequence Data to delimit Guettardeae, Rondeletieae, and Sections within *Rondeletia*.**
In: Annals of the Missouri Botanical Garden 96 (1), p. 182–193. DOI: 10.3417/2006179.

**Rydin, Catarina; Razafimandimbison, Sylvain G.; Bremer, Birgitta (2008):**
**Rare and enigmatic Genera (*Dunnia, Schizocolea, Colletoecema*), sisters to species-rich Clades: Phylogeny and aspects of Conservation Biology in the Coffee Family.**
In: Molecular Phylogenetics and Evolution 48 (1), p. 74–83. DOI: 10.1016/j.ympev.2008.04.006.

**Rydin, Catarina; Razafimandimbison, Sylvain G.; Khodabandeh, Anbar; Bremer, Birgitta (2009):**
**Evolutionary Relationships in the Spermacoceae alliance (Rubiaceae) using information from six molecular loci: Insights into systematic affinities of *Neohymenopogon* and *Mouretia*.**
In: Taxon 58 (3), p. 793–810.

**Salas, Roberto Manuel; Cabral, Elsa L. (2011):**
***Staelia culcita* (Rubiaceae), a new Species from Minas Gerais, Brazil.**
In: Plant Ecology and Evolution 144 (3), p. 372–376. DOI: 10.5091/plecevo.2011.620.

**Salas, Roberto Manuel; Viana, Pedro L.; Cabral, Elsa L.; Dessein, Steven; Janssens, Steven B. (2015):**
***Carajasia* (Rubiaceae), a new and endangered Genus from Carajás Mountain Range, Pará, Brazil.**
In: Phytotaxa 206 (1), p. 14. DOI: 10.11646/phytotaxa.206.1.4.

**Smedmark, Jenny E.E.; Bremer, Birgitta (2011):**
**Molecular Systematics and incongruent Gene Trees of Urophylleae (Rubiaceae).**
In: Taxon 60 (5), p. 1397–1406.

**Smedmark, Jenny E.E.; Razafimandimbison, Sylvain G.; Wikström, Niklas; Bremer, Birgitta (2014):**

**Inferring geographic range Evolution of a pantropical Tribe in the Coffee Family (Lasiantheae, Rubiaceae) in the face of topological Uncertainty.**
In: Molecular Phylogenetics and Evolution 70, p. 182–194. DOI: 10.1016/j.ympev.2013.09.007.

SMEDMARK, JENNY E.E.; RYDIN, CATARINA; RAZAFIMANDIMBISON, SYLVAIN G.; KHAN, SALEH AHAMMAD; LIEDE-SCHUMANN, SIGRID; BREMER, BIRGITTA (2008):
**A Phylogeny of Urophylleae (Rubiaceae) based on rps16 Intron Data.**
In: Taxon 57 (1), p. 24–32.

SOZA, VALERIE L.; OLMSTEAD, RICHARD G. (2010):
**Evolution of breeding systems and Fruits in New World *Galium* and Relatives (Rubiaceae).**
In: American Journal of Botany 97 (10), p. 1630–1646. DOI: 10.3732/ajb.1000130.

SOZA, VALERIE L.; OLMSTEAD, RICHARD G. (2010):
**Molecular Systematics of Tribe Rubieae (Rubiaceae): Evolution of major Clades, development of leaf-like Whorls, and Biogeography.**
In: Taxon 59 (3), p. 755–771.

STRANCZINGER, SZILVIA; GALAMBOS, ANIKO; SZENASY, DALMA; SZALONTAI, BALINT (2014):
**Phylogenetic Relationships in the Neotropical Tribe Hamelieae (Rubiaceae, Cinchonoideae) and Comments on its generic Limits.**
In: Journal of Systematics and Evolution 52 (5), p. 643–650. DOI: 10.1111/jse.12103.

TAYLOR, CHARLOTTE M. (2004):
**The Neotropical Genus *Ronabea* (Rubiaceae, Lasiantheae).**
In: Systematics and Geography of Plants 74 (1), p. 35–42.

TAYLOR, CHARLOTTE M. (2014):
**Rubiacearum Americanarum Magna Hama Pars XXXII. New Species and a new Combination in *Coussarea* (Coussareeae) from Western South America.**
In: Brittonia 66 (3), p. 256–268. DOI: 10.1007/s12228-014-9328-2.

TAYLOR, CHARLOTTE M.; GEREAU, ROY E. (2013):
**The Genus *Carapichea* (Rubiaceae, Psychotrieae).**
In: Annals of the Missouri Botanical Garden 99 (1), p. 100–127.

TAYLOR, CHARLOTTE M.; NEILL, DAVID A.; GEREAU, ROY E. (2011):
**Rubiacearum Americanarum Magna Hama Pars XXIX: Overview of the Neotropical Genus *Schizocalyx* (Condamineeae) and Description of two new Species.**
In: Novon: A Journal for Botanical Nomenclature 21 (4), p. 496–507. DOI: 10.3417/2008095.

TERRELL, EDWARD E.; ROBINSON, HAROLD ERNEST (2009):
**A new Genus, *Mexotis*, for five Mexican Species of Hedyotideae (Rubiaceae).**
In: Journal of the Botanical Research Institute of Texas 3 (1), p. 59–70.

THULIN, MATS; BREMER, BIRGITTA (2004):
**Studies in the Tribe Spermacoceae (Rubiaceae-Rubioideae): the Circumscriptions of *Amphiasma* and *Pentanopsis* and the Affinities of *Phylohydrax*.**
In: Plant Systematics and Evolution 247 (3-4). DOI: 10.1007/s00606-004-0155-x.

TOSH, JAMES; DAVIS, AARON P.; DESSEIN, STEVEN; BLOCK, PETRA J.; HUYSMANS, SUZY; FAY, MIKE F. ET AL. (2009):
**Phylogeny of *Tricalysia* (Rubiaceae) and its Relationships with allied Genera based on Plastid DNA Data: Resurrection of the Genus *Empogona*.**
In: Annals of the Missouri Botanical Garden 96 (1), p. 194–213. DOI: 10.3417/2006202.

**Verstraete, Brecht; Janssens, Steven B.; Lemaire, Benny; Smets, Erik; Dessein, Steven (2013):**
**Phylogenetic Lineages in Vanguerieae (Rubiaceae) associated with *Burkholderia* Bacteria in Sub-Saharan Africa.**
In: American Journal of Botany 100 (12), p. 2380–2387. DOI: 10.3732/ajb.1300303.

**Verstraete, Brecht; Janssens, Steven B.; Rønsted, Nina (2017):**
**Non-nodulated bacterial Leaf Symbiosis promotes the evolutionary Success of its Host Plants in the Coffee Family (Rubiaceae).**
In: Molecular Phylogenetics and Evolution 113, p. 161–168. DOI: 10.1016/j.ympev.2017.05.022.

**Verstraete, Brecht; Lachenaud, Olivier; Smets, Erik; Dessein, Steven; Bonaventure, Sonké (2013):**
**Taxonomy and Phylogenetics of *Cuviera* (RubiaceaeVanguerieae) and Reinstatement of *Globulostylis* with the Description of three new Species.**
In: Botanical Journal of the Linnean Society 173, p. 407–441.

**Wen, Hai-Zhen; Wang, Rui-Jiang (2012):**
***Foonchewia guangdongensis* gen. et sp. nov. (Rubioideae: Rubiaceae) and its systematic Position inferred from Chloroplast Sequences and Morphology.**
In: Journal of Systematics and Evolution 50 (5), p. 467–476. DOI: 10.1111/j.1759-6831.2012.00196.x.

**Wikström, Niklas; Neupane, Suman; Kårehed, Jesper; Motley, Timothy J.; Bremer, Birgitta (2013):**
**Phylogeny of *Hedyotis* L. (Rubiaceae: Spermacoceae): Redefining a complex Asian-Pacific Assemblage.**
In: Taxon 62 (2), p. 357–374.

**Xie, Peiwu; Tu, Tie-Yao; Razafimandimbison, Sylvain G.; Zhu, Chengjie; Zhang, Dian-Xiang (2014):**
**Phylogenetic Position of *Guihaiothamnus* (Rubiaceae): its evolutionary and ecological Implications.**
In: Molecular Phylogenetics and Evolution 78, p. 375–385. DOI: 10.1016/j.ympev.2014.05.022.

**Yang, Li-E; Meng, Ying; Peng, De-Li; Nie, Ze-long; Sun, Hang (2018):**
**Molecular Phylogeny of *Galium* L. of the Tribe Rubieae (Rubiaceae) - Emphasis on Chinese Species and Recognition of a new Genus *Pseudogalium*.**
In: Molecular Phylogenetics and Evolution 126, p. 221–232. DOI: 10.1016/j.ympev.2018.04.004.

**Yang, Li-E; Sun, Hang; Ehrendorfer, Friedrich; Nie, Ze-long (2016):**
**Molecular Phylogeny of Chinese *Rubia* (Rubiaceae: Rubieae) inferred from nuclear and Plastid DNA Sequences.**
In: Journal of Systematics and Evolution 54 (1), p. 37–47. DOI: 10.1111/jse.12157.

### Rubiales

**Yang, Lei-Lei; Li, Hong-Lei; Wei, Lei; Yang, Tuo; Kuang, Dai-Yong; Li, Ming-Hong et al. (2016):**
**A Supermatrix Approach provides a comprehensive Genus-level Phylogeny for Gentianales.**
In: Journal of Systematics and Evolution 54 (4), p. 400–415. DOI: 10.1111/jse.12192.

### Ruppiaceae

**Ito, Yu; Ohi-Toma, Tetsuo; Murata, Jin; Tanaka, Norio (2010):**
**Hybridization and Polyploidy of an aquatic Plant, *Ruppia* (Ruppiaceae), inferred from Plastid and nuclear DNA Phylogenies.**
In: American Journal of Botany 97 (7), p. 1156–1167. DOI: 10.3732/ajb.0900168.

### Ruscaceae

**Kim, Joo-Hwan; Kim, Dong-Kap; Forest, Félix; Fay, Michael F.; Chase, Mark W. (2010):**

**Molecular Phylogenetics of Ruscaceae sensu lato and related families (Asparagales) based on Plastid and nuclear DNA Sequences.**
In: Annals of Botany 106 (5), p. 775–790. DOI: 10.1093/aob/mcq167.

KIM, S.-C.; LEE, NAM SOOK (2007):
**Generic Delimitation and Biogeography of *Maianthemum* and *Smilacina* (Ruscaceae sensu lato): preliminary Results based on partial 3′ matK Gene and trnK 3′ Intron Sequences of cpDNA.**
In: Plant Systematics and Evolution 265 (1-2), p. 1–12. DOI: 10.1007/s00606-007-0517-2.

MENG, YING; NIE, ZE-LONG; DENG, TAO; WEN, JUN; YANG, YONG-PING (2014):
**Phylogenetics and Evolution of Phyllotaxy in the Solomon's seal Genus *Polygonatum* (Asparagaceae: Polygonateae).**
In: Botanical Journal of the Linnean Society 176 (4), p. 435–451. DOI: 10.1111/boj.12218.

MENG, YING; WEN, JUN; NIE, ZE-LONG; SUN, HANG; YANG, YONG-PING (2008):
**Phylogeny and biogeographic Diversification of *Maianthemum* (Ruscaceae: Polygonatae).**
In: Molecular Phylogenetics and Evolution 49 (2), p. 424–434. DOI: 10.1016/j.ympev.2008.07.017.

ROJAS-PIÑA, VANESSA; OLSON, MARK E.; ALVARADO-CÁRDENAS, LEONARDO O.; EGUIARTE, LUIS E. (2014):
**Molecular Phylogenetics and Morphology of *Beaucarnea* (Ruscaceae) as distinct from *Nolina*, and the Submersion of *Calibanus* into *Beaucarnea*.**
In: Taxon 63 (6), p. 1193–1211. DOI: 10.12705/636.31.

TILLICH, H.-J. (2005):
**A key for *Aspidistra* (Ruscaceae), including fifteen new Species from Vietnam.**
In: Feddes Repertorium 116 (5-6), p. 313–338. DOI: 10.1002/fedr.200511076.

TILLICH, H.-J.; AVERYANOV, LEONID V.; DZU, N. V. (2007):
**Six new Species of *Aspidistra* (Ruscaceae) from northern Vietnam.**
In: Blumea 52 (2), p. 335–344. DOI: 10.3767/000651907X609070.

### Rutaceae

APPELHANS, MARC S.; KROHM, SABRINA; MANAFZADEH, SARA; WEN, JUN (2016):
**Phylogenetic Placement of *Psilopeganum*, a rare monotypic Genus of Rutaceae (the Citrus Family) endemic to China.**
In: Journal of Systematics and Evolution 54 (5), p. 535–544. DOI: 10.1111/jse.12208.

APPELHANS, MARC S.; REICHELT, NIKLAS; GROPPO, MILTON; PAETZOLD, CLAUDIA; WEN, JUN (2018):
**Phylogeny and Biogeography of the pantropical Genus *Zanthoxylum* and its closest relatives in the Proto-Rutaceae Group (Rutaceae).**
In: Molecular Phylogenetics and Evolution 126, p. 31–44. DOI: 10.1016/j.ympev.2018.04.013.

APPELHANS, MARC S.; SMETS, ERIK; RAZAFIMANDIMBISON, SYLVAIN G.; HAEVERMANS, THOMAS; VAN MARLE, E. J.; COULOUX, ARNAUD ET AL. (2011):
**Phylogeny, evolutionary Trends and Classification of the *Spathelia-Ptaeroxylon* Clade: morphological and molecular Insights.**
In: Annals of Botany 107 (8), p. 1259–1277. DOI: 10.1093/aob/mcr076.

APPELHANS, MARC S.; WAGNER, WARREN L.; WOOD, KENNETH R. (2014):
***Melicope balgooyi* Appelhans, W.I. Wagner & K.R.Wood, a new Species and new Record in *Melicope* Section *Melicope* (Rutaceae) for the Austral Islands.**
In: PhytoKeys (39), p. 77–86. DOI: 10.3897/phytokeys.39.7691.

APPELHANS, MARC S.; WEN, JUN; WAGNER, WARREN L. (2014):
**A molecular Phylogeny of *Acronychia, Euodia, Melicope* and Relatives (Rutaceae) reveals polyphyletic Genera and Key Innovations for Species richness.**
In: Molecular Phylogenetics and Evolution 79, p. 54–68. DOI: 10.1016/j.ympev.2014.06.014.

APPELHANS, MARC S.; WEN, JUN; WOOD, KENNETH R.; ALLAN, GERARD J.; ZIMMER, ELIZABETH A.; WAGNER, WARREN L. (2014):
**Molecular phylogenetic Analysis of Hawaiian Rutaceae (*Melicope, Platydesma* and *Zanthoxylum*) and their different Colonization Patterns.**
In: Botanical Journal of the Linnean Society 174, p. 425–448.

ARAÚJO, EDSON FREITAS DE; QUEIROZ, LUCIANO PAGANUCCI; MACHADO, MARCOS ANTÔNIO (2003):
**What is *Citrus*? Taxonomic Implications from a Study of cp-DNA Evolution in the Tribe Citreae (Rutaceae Subfamily Aurantioideae).**
In: Organisms Diversity and Evolution 3, p. 55–62.

ARMSTRONG, JIM A. (2002):
**The Genus *Zieria* (Rutaceae): a systematic and evolutionary Study.**
In: Australian Systematic Botany 15 (3), p. 277–463. DOI: 10.1071/SB00040.

BARRETT, ROSEMARY A.; BAYLY, MICHAEL J.; DURETTO, MARCO F.; FORSTER, PAUL I.; LADIGES, PAULINE Y.; CANTRILL, DAVID J. (2014):
**A Chloroplast Phylogeny of *Zieria* (Rutaceae) in Australia and New Caledonia shows widespread Incongruence with Species-level Taxonomy.**
In: Australian Systematic Botany 27 (6), p. 427–449. DOI: 10.1071/SB14033.

BAYER, RANDALL J.; MABBERLEY, DAVID J.; MORTON, CYNTHIA M.; MILLER, CATHY H.; SHARMA, ISH K.; PFEIL, BERNARD E. ET AL. (2009):
**A molecular Phylogeny of the Orange Subfamily (Rutaceae: Aurantioideae) using nine cpDNA Sequences.**
In: American Journal of Botany 96 (3), p. 668–685. DOI: 10.3732/ajb.0800341.

BAYLY, MICHAEL J.; DURETTO, MARCO F.; HOLMES, GARETH D.; FORSTER, PAUL I.; CANTRILL, DAVID J.; LADIGES, PAULINE Y. (2015):
**Transfer of the New Caledonian Genus *Boronella* to *Boronia* (Rutaceae) based on Analyses of cpDNA and nrDNA.**
In: Australian Systematic Botany 28 (3), p. 111–123. DOI: 10.1071/SB15008.

BAYLY, MICHAEL J.; HOLMES, GARETH D.; FORSTER, PAUL I.; MUNZINGER, JÉRÔME; CANTRILL, DAVID J.; LADIGES, PAULINE Y. (2016):
**Phylogeny, Classification and Biogeography of *Halfordia* (Rutaceae) in Australia and New Caledonia.**
In: Plant Systematics and Evolution 302 (10), p. 1457–1470. DOI: 10.1007/s00606-016-1344-0.

BRUNIERA, CARLA POLESELLI; KALLUNKI, JACQUELYN A.; GROPPO, MILTON (2015):
***Almeidea* A.St.-Hil. belongs to *Conchocarpus* J.C. Mikan (Galipeinae, Rutaceae): Evidence from morphological and molecular Data, with a first Analysis of Subtribe Galipeinae.**
In: Public Library of Science One 10 (5), e0125650. DOI: 10.1371/journal.pone.0125650.

BUT, PAUL PUI-HAY; POON, ALICE WING-SEM; SHAW, PANG-CHUI; SIMMONS, MARK P.; GREGER, HARALD (2009):
**Contribution of molecular Cladistics to the Taxonomy of Rutaceae in China.**
In: Journal of Systematics and Evolution 47 (2), p. 144–150. DOI: 10.1111/j.1759-6831.2009.00013.x.

CHOI, BO-KYUNG; DURETTO, MARCO F.; HONG, SUK-PYO (2012):

**Comparative Seed Morphology of *Boronia* and related Genera (Boroniinae: Rutaceae) and its systematic Implications.**
In: Nordic Journal of Botany 30 (2), p. 241–256. DOI: 10.1111/j.1756-1051.2011.01251.x.

**DIAS, PEDRO; UDULUTSCH, RENATA GIASSI; PIRANI, JOSÉ RUBENS (2015):**
**Molecular Phylogeny and Biogeography of the South-American Genus *Metrodorea* (Rutaceae).**
In: Turkish Journal of Botany 39, p. 825–834. DOI: 10.3906/bot-1410-49.

**FRENCH, PHILIPPA A.; BROWN, GILLIAN K.; BAYLY, MICHAEL J. (2016):**
**Incongruent Patterns of nuclear and Chloroplast Variation in *Correa* (Rutaceae): Introgression and Biogeography in South-eastern Australia.**
In: Plant Systematics and Evolution 302 (4), p. 447–468. DOI: 10.1007/s00606-016-1277-7.

**GROPPO, MILTON; PIRANI, JOSÉ RUBENS; SALATINO, MARIA LUIZA FARIA; BLANCO, SILVIA R.; KALLUNKI, JACQUELYN A. (2008):**
**Phylogeny of Rutaceae based on two non-coding Regions from cpDNA.**
In: American Journal of Botany 95 (8), p. 985–1005. DOI: 10.3732/ajb.2007313.

**MOLE, B. J.; UDOVICIC, FRANK; LADIGES, PAULINE Y.; DURETTO, MARCO F. (2004):**
**Molecular Phylogeny of *Phebalium* (Rutaceae: Boronieae) and related Genera based on the nrDNA Regions ITS 1+2.**
In: Plant Systematics and Evolution 249 (3-4), p. 197–212. DOI: 10.1007/s00606-004-0218-z.

**MORTON, CYNTHIA M. (2009):**
**Phylogenetic Relationships of the Aurantioideae (Rutaceae) based on the nuclear ribosomal DNA ITS Region and three noncoding Chloroplast DNA Regions, atpB-rbcL spacer, rps16, and trnL-trnF.**
In: Organisms Diversity and Evolution 9 (1), p. 52–68. DOI: 10.1016/j.ode.2008.11.001.

**OTHMAN, RAJA NUR ATEEKA; JORDAN, GREGORY J.; WORTH, JAMES R. P.; STEANE, DOROTHY A.; DURETTO, MARCO F. (2010):**
**Phylogeny and infrageneric Classification of *Correa* Andrews (Rutaceae) on the Basis of nuclear and Chloroplast DNA.**
In: Plant Systematics and Evolution 288 (3-4), p. 127–138. DOI: 10.1007/s00606-010-0315-0.

**POON, ALICE WING-SEM; SHAW, PANG-CHUI; SIMMONS, MARK P.; BUT, PAUL PUI-HAY (2007):**
**Congruence of Molecular, Morphological, and Biochemical Profiles in Rutaceae: a Cladistic Analysis of the Subfamilies Rutoideae and Toddalioideae.**
In: Systematic Botany 32 (4), p. 837–846.

**RAZAFIMANDIMBISON, SYLVAIN G.; APPELHANS, MARC S.; RABARISON, HARISON; HAEVERMANS, THOMAS; RAKOTONDRAFARA, ANDRIARIMALALA; RAKOTONANDRASANA, STEPHAN R. ET AL. (2010):**
**Implications of a molecular phylogenetic Study of the Malagasy Genus *Cedrelopsis* and its Relatives (Ptaeroxylaceae).**
In: Molecular Phylogenetics and Evolution 57 (1), p. 258–265. DOI: 10.1016/j.ympev.2010.06.023.

**SALVO, GABRIELE; BACCHETTA, GIANLUIGI; GHAHREMANINEJAD, FARROKH; CONTI, ELENA (2008):**
**Phylogenetic Relationships of Ruteae (Rutaceae): new Evidence from the Chloroplast Genome and comparisons with non-molecular Data.**
In: Molecular Phylogenetics and Evolution 49 (3), p. 736–748. DOI: 10.1016/j.ympev.2008.09.004.

**SALVO, GABRIELE; HO, SIMON Y.W.; ROSENBAUM, GIDEON; REE, RICHARD H.; CONTI, ELENA (2010):**

**Tracing the temporal and spatial Origins of Island Endemics in the Mediterranean Region: a case Study from the Citrus-family (*Ruta* L., Rutaceae).**
In: Systematic Biology 59 (6), p. 705–722. DOI: 10.1093/sysbio/syq046.

SALVO, GABRIELE; MANAFZADEH, SARA; GHAHREMANINEJAD, FARROKH; TOJIBAEV, KOMILJON; ZELTNER, LOUIS; CONTI, ELENA (2011):
**Phylogeny, Morphology, and Biogeography of *Haplophyllum* (Rutaceae), a species-rich Genus of the Irano-Turanian floristic Region.**
In: Taxon 60 (2), p. 513–527.

SAMUEL, L.; EHRENDORFER, FRIEDRICH; CHASE, MARK W.; GREGER, HARALD (2001):
**Phylogenetic Analyses of Aurantioideae (Rutaceae) based on Non-Coding Plastid DNA Sequences and Phytochemical Features.**
In: Plant Biology 3, p. 77–87.

SHIVAKUMAR, VIKRAM S.; APPELHANS, MARC S.; JOHNSON, GABRIEL; CARLSEN, MÓNICA M.; ZIMMER, ELIZABETH A. (2017):
**Analysis of whole Chloroplast Genomes from the Genera of the Clauseneae, the Curry Tribe (Rutaceae, Citrus-Family).**
In: Molecular Phylogenetics and Evolution 117, p. 135–140. DOI: 10.1016/j.ympev.2016.12.015.

THIV, MIKE; VAN DER NIET, TIMOTHEÜS; RUTSCHMANN, FRANK; THULIN, MATS; BRUNE, THOMAS; LINDER, HANS PETER (2011):
**Old-New World and Trans-African Disjunctions of *Thamnosma* (Rutaceae): intercontinental long-distance Dispersal and local Differentiation in the succulent Biome.**
In: American Journal of Botany 98 (1), p. 76–87. DOI: 10.3732/ajb.1000339.

TRINDER-SMITH, TERRY H.; LINDER, HANS PETER; VAN DER NIET, TIMOTHEÜS; VERBOOM, GEORGE ANTHONY; NOWELL, TRACEY L. (2007):
**Plastid DNA Sequences Reveal Generic Paraphyly within Diosmeae (Rutoideae, Rutaceae).**
In: Systematic Botany 32 (4), p. 847–855.

WEGE, JULIET A. (2017):
**Taxonomic Notes on *Asterolasia* (Rutaceae) in Western Australia to inform Conservation.**
In: Nuytsia 28, p. 141–146.

XIE, RANG-JIN; ZHI-QIN; DENG, ZHOU LIE (2008):
**Taxonomic and phylogenetic Relationships among the Genera of the True Citrus-Fruit Trees Group (Aurantioideae, Rutaceae) based on AFLP Markers.**
In: Journal of Systematics and Evolution 46 (5), p. 682–691.

YAN, G.; SHAN, F.; PLUMMER, J. A. (2002):
**Genomic Relationships within *Boronia* (Rutaceae) as revealed by karyotype Analysis and Rapd molecular Markers.**
In: Plant Systematics and Evolution 233 (3), p. 147–161. DOI: 10.1007/s00606-002-0202-4.


### Sabiaceae


MORALES, J. FRANCISCO (2013):
**Sinopsis del Género *Meliosma* (Sabiaceae) en México y Centroamérica.**
In: Phytoneuron 82, p. 1–86.

Yang, Tuo; Lu, Li-Min; Wang, Wei; Li, Jian-Hua; Manchester, Steven R.; Wen, Jun; Chen, Zhi-Duan (2018):
**Boreotropical Range Expansion and long-distance Dispersal explain two Amphi-Pacific tropical Disjunctions in Sabiaceae.**
In: Molecular Phylogenetics and Evolution 124, p. 181–191. DOI: 10.1016/j.ympev.2018.03.005.

Zúñiga, Jose Daniel (2015):
**Phylogenetics of Sabiaceae with Emphasis on *Meliosma* based on Nuclear and Chloroplast Data.**
In: Systematic Botany 40 (3), p. 761–775. DOI: 10.1600/036364415X689221.

### Salicaceae

Abdollahzadeh, A.; Osaloo, p. Kazempour Osaloo; Maassoumi, Ali Aasghar (2011):
**Molecular Phylogeny of the Genus *Salix* (Salicaceae) with an Emphasize to its Species in Iran.**
In: Iranian Journal of Botany 17 (2), p. 244–253.

Alford, Mac H. (2006):
**A taxonomic Revision of the Andean Genus *Pineda* (Salicaceae).**
In: Kew Bulletin 61, p. 205–214.

Alford, Mac H. (2008):
**Revision of *Neosprucea* (Salicaceae).**
In: Systematic Botany Monographs 85, p. 1–62.

Applequist, Wendy L.; Schatz, George E. (2016):
**A synoptic Revision of the Malagasy Species of *Scolopia* Schreb. (Salicaceae, Scolopieae).**
In: Adansonia 38 (1), p. 99–115. DOI: 10.5252/a2016n1a6.

Argus, George W. (2007):
**Salix (Salicaceae) Distribution Maps and a Synopsis of their Classification in North America, North of Mexico.**
In: Harvard Papers in Botany 12 (2), p. 335–368. DOI: 10.3100/1043-4534(2007)12[335:SSDMAA]2.0.CO;2.

Azuma, Takayuki; Kajita, Tadashi; Yokoyama, Jun U.N.; Ohashi, Hiroyoshi (2000):
**Phylogenetic Relationships of *Salix* (Salicaceae) based on rbcL Sequence Data.**
In: American Journal of Botany 87 (1), p. 67–75. DOI: 10.2307/2656686.

Chen, Jia-Hui; Sun, Hang; Wen, Jun; Yang, Yong-Ping (2010):
**Molecular Phylogeny of *Salix* L. (Salicaceae) inferred from three Chloroplast Datasets and its systematic Implications.**
In: Taxon 59 (1), p. 29–37. DOI: 10.1002/tax.591004.

Hamzeh, Mona; Dayanandan, Selvadurai (2004):
**Phylogeny of *Populus* (Salicaceae) based on nucleotide Sequences of Chloroplast Trnt-trnf Region and nuclear rDNA.**
In: American Journal of Botany 91 (9), p. 1398–1408. DOI: 10.3732/ajb.91.9.1398.

Hardig, Terry M.; Anttila, C. K.; Brunsfeld, p. J. (2010):
**A phylogenetic Analysis of *Salix* (Salicaceae) based on matK and Ribosomal DNA Sequence Data.**
In: Journal of Botany 2010 (8), p. 1–12. DOI: 10.1155/2010/197696.

Huang, Daisie I.; Hefer, Charles A.; Kolosova, Natalia; Douglas, Carl J.; Cronk, Quentin C.B. (2014):

**Whole plastome Sequencing reveals deep Plastid Divergence and cytonuclear Discordance between closely related Balsam Poplars, *Populus balsamifera* and *P. trichocarpa* (Salicaceae).**
In: the new Phytologist 204 (3), p. 693–703. DOI: 10.1111/nph.12956.

INGVARSSON, PÄR K. (2008):
**Molecular Evolution of synonymous Codon usage in *Populus*.**
In: BMC Evolutionary Biology 8, p. 307. DOI: 10.1186/1471-2148-8-307.

LEVSEN, NICHOLAS D.; TIFFIN, PETER; OLSON, MATTHEW P. (2012):
**Pleistocene Speciation in the Genus *Populus* (Salicaceae).**
In: Systematic Biology 61 (3), p. 401–412. DOI: 10.1093/sysbio/syr120.

LIU, XIA; WANG, ZHAOSHAN; WANG, DONGSHENG; ZHANG, JIANGUO (2016):
**Phylogeny of *Populus-Salix* (Salicaceae) and their relative Genera using molecular Datasets.**
In: Biochemical Systematics and Ecology 68, p. 210–215. DOI: 10.1016/j.bse.2016.07.016.

MARQUETE, RONALDO; MANSANO, VIDAL DE FREITAS (2012):
**Taxonomic Revision of the *Casearia ulmifolia* Complex (Salicaceae).**
In: Novon: A Journal for Botanical Nomenclature 22 (2), p. 196–206. DOI: 10.3417/2011011.

WAN, XUE QIN; ZHANG, FAN; ZHONG, YU; DING, YUN HAI; WANG, CHANG LIANG; HU, TING XIN (2013):
**Study of genetic Relationships and Phylogeny of the native *Populus* in Southwest China based on nucleotide Sequences of Chloroplast trnT–trnF and nuclear DNA.**
In: Plant Systematics and Evolution 299 (1), p. 57–65. DOI: 10.1007/s00606-012-0702-9.

WANG, DONGSHENG; WANG, ZHAOSHAN; DU, SHUHUI; ZHANG, JIANGUO (2015):
**Phylogeny of Section *Leuce* (*Populus*, Salicaceae) inferred from 34 Chloroplast DNA fragments.**
In: Biochemical Systematics and Ecology 63, p. 212–217. DOI: 10.1016/j.bse.2015.09.020.

WANG, ZHAOSHAN; DU, SHUHUI; DAYANANDAN, SELVADURAI; WANG, DONGSHENG; ZENG, YANFEI; ZHANG, JIANGUO (2014):
**Phylogeny Reconstruction and hybrid Analysis of *Populus* (Salicaceae) based on nucleotide Sequences of multiple single-copy nuclear Genes and Plastid Fragments.**
In: Public Library of Science One 9 (8), e103645. DOI: 10.1371/journal.pone.0103645.

WU, JIE; NYMAN, TOMMI; WANG, DONG-CHAO; ARGUS, GEORGE W.; YANG, YONG-PING; CHEN, JIA-HUI (2015):
**Phylogeny of *Salix* Subgenus *Salix* s.l. (Salicaceae): Delimitation, Biogeography, and reticulate Evolution.**
In: BMC Evolutionary Biology 15, p. 31. DOI: 10.1186/s12862-015-0311-7.

### Salviniaceae

METZGAR, JORDAN S.; SCHNEIDER, HARALD; PRYER, KATHLEEN M. (2007):
**Phylogeny and Divergence Time Estimates for the Fern Genus *Azolla* (Salviniaceae).**
In: International Journal of Plant Sciences 168 (7), p. 1045–1953.

NAGALINGUM, NATHALIE S.; NOWAK, MICHAEL D.; PRYER, KATHLEEN M. (2008):
**Assessing phylogenetic Relationships in extant heterosporous Ferns (Salviniales), with a focus on *Pilularia* and *Salvinia*.**
In: Botanical Journal of the Linnean Society 157, p. 673–685.

PEREIRA, ANA L.; MARTINS, MADALENA; OLIVEIRA, M. MARGARIDA; CARRAPIÇO, FRANCISCO (2011):

**Morphological and genetic Diversity of the Family Azollaceae inferred from vegetative Characters and Rapd Markers.**
In: Plant Systematics and Evolution 297 (3-4), p. 213–226. DOI: 10.1007/s00606-011-0509-0.

REID, JILL D.; PLUNKETT, GREGORY M.; PETERS, GERALD A. (2006):
**Phylogenetic Relationships in the heterosporous Fern Genus *Azolla* (Azollaceae) based on DNA Sequence Data from three noncoding Regions.**
In: International Journal of Plant Sciences 167 (3), p. 529–538.

### Samolaceae

JONES, KEITH; ANDERBERG, ARNE A.; CRAENE, L. P. RONSE; WANNTORP, LIVIA (2012):
**Origin, Diversification, and Evolution of *Samolus valerandi* (Samolaceae, Ericales).**
In: Plant Systematics and Evolution 298 (8), p. 1523–1531. DOI: 10.1007/s00606-012-0655-z.

WANNTORP, LIVIA; ANDERBERG, ARNE A. (2011):
**Evolution and Diversification of Brook Weeds (*Samolus*, Samolaceae, Ericales).**
In: International Journal of Plant Sciences 172 (2), p. 250–266. DOI: 10.1086/657647.

### Santalaceae

DER, JOSHUA P.; NICKRENT, DANIEL LEE (2008):
**A molecular Phylogeny of Santalaceae (Santalales).**
In: Systematic Botany 33 (1), p. 107–116.

HARBAUGH, DANICA T. (2008):
**Polyploid and Hybrid Origins of Pacific Island Sandalwoods (*Santalum*, Santalaceae) inferred from Low-Copy Nuclear and Flow Cytometry Data.**
In: International Journal of Plant Sciences 169 (5), p. 677–685. DOI: 10.1086/533610.

HARBAUGH, DANICA T.; BALDWIN, BRUCE G. (2007):
**Phylogeny and Biogeography of the Sandalwoods (*Santalum*, Santalaceae): Repeated Dispersals throughout the Pacific.**
In: American Journal of Botany 94 (6), p. 1028–1040.

ROGERS, ZACHARY S.; NICKRENT, DANIEL LEE; MALÉCOT, VALÉRY (2008):
***Staufferia* and *Pilgerina*: Two new endemic monotypic arborescent Genera of Santalaceae from Madagascar.**
In: Annals of the Missouri Botanical Garden 95 (2), p. 391–404. DOI: 10.3417/2006148.

VIDAL-RUSSELL, ROMINA; NICKRENT, DANIEL LEE (2008):
**The first Mistletoes: Origins of aerial Parasitism in Santalales.**
In: Molecular Phylogenetics and Evolution 47 (2), p. 523–537. DOI: 10.1016/j.ympev.2008.01.016.

### Santalales

NICKRENT, DANIEL LEE; MALÉCOT, VALÉRY; VIDAL-RUSSELL, ROMINA; DER, JOSHUA P. (2010):
**A revised Classification of Santalales.**
In: Taxon 59 (2), p. 538–558.

### Sapindaceae

BUERKI, SVEN; DAVIDSON, CHRISTOPHER; PEREIRA, JOAN T.; CALLMANDER, MARTIN W. (2013):
**A new endemic Species of *Trigonachras* (Sapindaceae) from Sabah, Malaysia (Borneo).**
In: Phytotaxa 88 (2), p. 19–24. DOI: 10.11646/phytotaxa.88.2.1.

**Buerki, Sven; Forest, Félix; Acevedo-Rodríguez, Pedro; Callmander, Martin W.; Nylander, Johan A.A.; Harrington, Mark G. et al. (2009):**

**Plastid and nuclear DNA Markers reveal intricate Relationships at subfamilial and tribal levels in the Soapberry Family (Sapindaceae).**

In: Molecular Phylogenetics and Evolution 51 (2), p. 238–258. DOI: 10.1016/j.ympev.2009.01.012.

**Buerki, Sven; Forest, Félix; Callmander, Martin W.; Lowry, Porter Prescott II.; Devey, Dion S.; Munzinger, Jérôme (2012):**

**Phylogenetic Inference of New Caledonian Lineages of Sapindaceae: molecular Evidence requires a Reassessment of generic Circumscriptions.**

In: Taxon 61 (1), p. 109–119.

**Buerki, Sven; Forest, Félix; Salamin, Nicolas; Alvarez, Nadir (2011):**

**Comparative performance of Supertree Algorithms in large Data sets using the Soapberry Family (Sapindaceae) as a case Study.**

In: Systematic Biology 60 (1), p. 32–44. DOI: 10.1093/sysbio/syq057.

**Buerki, Sven; Lowry, Porter Prescott II.; Alvarez, Nadir; Razafimandimbison, Sylvain G.; Küpfer, Philippe; Callmander, Martin W. (2010):**

**Phylogeny and Circumscription of Sapindaceae revisited: molecular Sequence Data, Morphology and Biogeography support Recognition of a new Family, Xanthoceraceae.**

In: Plant Ecology and Evolution 143 (2), p. 148–159. DOI: 10.5091/plecevo.2010.437.

**Buerki, Sven; Lowry, Porter Prescott II.; Andriambololonera, Sylvie; Phillipson, Peter B.; Vary, Laura; Callmander, Martin W. (2011):**

**How to kill two Genera with one Tree: clarifying generic Circumscriptions in an endemic Malagasy Clade of Sapindaceae.**

In: Botanical Journal of the Linnean Society 165, p. 223–234.

**Buerki, Sven; Lowry, Porter Prescott; Phillipson, Peter B.; Callmander, Martin W. (2010):**

**Molecular phylogenetic and morphological Evidence supports Recognition of *Gereaua*, a new Endemic Genus of Sapindaceae from Madagascar.**

In: Systematic Botany 35 (1), p. 172–180. DOI: 10.1600/036364410790862669.

**Buijsen, J. R. M.; van Welzen, Peter C.; Ham, Raymond W.J.M. van der (2003):**

**A phylogenetic Analysis of *Harpullia* (Sapindaceae) with Notes on Historical Biogeography.**

In: Systematic Botany 28 (1), p. 106–117.

**Coulleri, J. P.; Dematteis, Massimiliano; Ferrucci, M. p. (2012):**

**A new Insight into *Serjania* Mill. (Sapindaceae, Paullinieae) infrageneric Classification: a cytogenetic approach.**

In: Plant Systematics and Evolution 298 (9), p. 1743–1753. DOI: 10.1007/s00606-012-0675-8.

**Edwards, K. J.; Gadek, Paul A. (2001):**

**Evolution and Biogeography of *Alectryon* (Sapindaceae).**

In: Molecular Phylogenetics and Evolution 20 (1), p. 14–26. DOI: 10.1006/mpev.2001.0952.

**Forest, Félix; Drouin, Josée Nadia; Charest, René; Brouillet, Luc; Bruneau, Anne (2001):**

**A morphological phylogenetic Analysis of *Aesculus* L. and *Billia* Peyr. (Sapindaceae).**

In: Canadian Journal of Botany 79 (2), p. 154–169. DOI: 10.1139/cjb-79-2-154.

**Grimm, Guido W.; Denk, Thomas; Hemleben, V. (2007):**

**Evolutionary History and Systematics of *Acer* Section *Acer* – a case Study of low-level Phylogenetics.**
In: Plant Systematics and Evolution 267 (1-4), p. 215–253. DOI: 10.1007/s00606-007-0572-8.

HARRINGTON, MARK G.; BIFFIN, ED; GADEK, PAUL A. (2009):
**Comparative Study of the Evolution of nuclear ribosomal Spacers Incorporating secondary Structure Analyzes within Dodonaeoideae, Hippocastanoideae and Xanthoceroideae (Sapindaceae).**
In: Molecular Phylogenetics and Evolution 50 (2), p. 364–375. DOI: 10.1016/j.ympev.2008.11.010.

HARRINGTON, MARK G.; EDWARDS, KAREN J.; JOHNSON, SHEILA A.; CHASE, MARK W.; GADEK, PAUL A. (2005):
**Phylogenetic Inference in Sapindaceae sensu lato using Plastid matK and rbcL DNA Sequences.**
In: Systematic Botany 30 (2), p. 366–382.

HARRINGTON, MARK G.; GADEK, PAUL A. (2010):
**Phylogenetics of Hopbushes and Pepperflowers (*Dodonaea, Diplopeltis* - Sapindaceae), based on nuclear ribosomal ITS and partial ETS Sequences incorporating secondary-Structure Models.**
In: Australian Systematic Botany 23 (6), p. 431–442. DOI: 10.1071/SB10002.

HARRIS, A. J.; FU, CHENG-XIN; XIANG, QIU-YUN JENNY; HOLLAND, LARINDA; WEN, JUN (2016):
**Testing the Monophyly of *Aesculus* L. and *Billia* Peyr., woody Genera of Tribe Hippocastaneae of the Sapindaceae.**
In: Molecular Phylogenetics and Evolution 102, p. 145–151. DOI: 10.1016/j.ympev.2016.06.001.

HARRIS, A. J.; XIANG, QIU-YUN; THOMAS, DAVID T. (2009):
**Phylogeny, Origin, and biogeographic History of Aesculus L. (Sapindales) - an Update from combined Analysis of DNA Sequences, Morphology, and Fossils.**
In: Taxon 58 (1), p. 108–126.

HOPKINS, HELEN C.F. (2013):
**A second Species in the little known African Genus *Blighiopsis* (Sapindaceae).**
In: Kew Bulletin 68 (2), p. 345–353.

LI, JIAN-HUA (2011):
**Phylogenetic Evaluation of Series Delimitations in Section *Palmata* (*Acer*, Aceroideae, Sapindaceae) based on Sequences of Nuclear and Chloroplast Genes.**
In: Aliso 29 (1), p. 43–49.

MUNZINGER, JÉRÔME; LOWRY, PORTER PRESCOTT; BUERKI, SVEN; CALLMANDER, MARTIN W. (2016):
**A taxonomic Revision of the Endemic New Caledonian Genus *Storthocalyx* (Sapindaceae).**
In: Systematic Botany 41 (2), p. 387–400. DOI: 10.1600/036364416X691902.

RENNER, SUSANNE S.; GRIMM, GUIDO W.; SCHNEEWEISS, GERALD M.; STUESSY, TOD F.; RICKLEFS, ROBERT E. (2008):
**Rooting and dating Maples (*Acer*) with an uncorrelated-rates molecular clock: Implications for north American/Asian disjunctions.**
In: Systematic Biology 57 (5), p. 795–808. DOI: 10.1080/10635150802422282.

URDAMPILLETA, J. D.; COULLERI, J. P.; FERRUCCI, M. S.; FORNI-MARTINS, E. R. (2013):
**Karyotype Evolution and phylogenetic Analyses in the Genus *Cardiospermum* L. (Paullinieae, Sapindaceae).**
In: Plant Biology 15 (5), p. 868–881. DOI: 10.1111/j.1438-8677.2012.00679.x.

**Weckerle, Caroline S.; Rutishauser, Rolf (2003):**

**Comparative Morphology and Systematic Position of *Averrhoidium* within Sapindaceae.**

In: International Journal of Plant Sciences 164 (5), p. 775–792.

**Zhang, Zhihong; Li, Chunqi; Li, Jian-Hua (2010):**

**Conflicting Phylogenies of Section *Macrantha* (Acer Aceroideae, Sapindaceae) based on Chloroplast and Nuclear DNA.**

In: Systematic Botany 35 (4), p. 801–810. DOI: 10.1600/036364410X539871.


### Sapindales

**Müllner-Riehl, Alexandra N.; Weeks, Andrea; Clayton, Joshua W.; Buerki, Sven; Nauheimer, Lars; Chiang, Yu-Chung et al. (2016):**

**Molecular Phylogenetics and molecular Clock Dating of Sapindales based on Plastid rbcL, atpB and trnL-trnF DNA Sequences.**

In: Taxon 65 (5), p. 1019–1036. DOI: 10.12705/655.5.


### Sapotaceae

**Anderberg, Arne A.; Swenson, Ulf (2003):**

**Evolutionary Lineages in Sapotaceae (Ericales): A Cladistic Analysis based on ndhF Sequence Data.**

In: International Journal of Plant Sciences 164 (5), p. 763–773.

**Armstrong, K. E. (2013):**

**A Revision of the Asian-pacific Species of *Manilkara* (Sapotaceae).**

In: Edinburgh Journal of Botany 70 (1), p. 7–56. DOI: 10.1017/S0960428612000327.

**Bartish, Igor V.; Swenson, Ulf; Munzinger, Jérôme; Anderberg, Arne A. (2005):**

**Phylogenetic Relationships among New Caledonian Sapotaceae (Ericales): molecular Evidence for generic Polyphyly and repeated Dispersal.**

In: American Journal of Botany 92 (4), p. 667–673. DOI: 10.3732/ajb.92.4.667.

**Gautier, Laurent; Naciri, Yamama; Anderberg, Arne A.; Smedmark, Jenny E.E.; Randrianaivo, Richard; Swenson, Ulf (2013):**

**A new species, Genus and Tribe of Sapotaceae, endemic to Madagascar.**

In: Taxon 62 (5), p. 972–983. DOI: 10.12705/625.17.

**Smedmark, Jenny E.E.; Anderberg, Arne A. (2007):**

**Boreotropical migration explains Hybridization between geographically distant Lineages in the pantropical Clade Sideroxyleae (Sapotaceae).**

In: American Journal of Botany 94 (9), p. 1491–1505. DOI: 10.3732/ajb.94.9.1491.

**Smedmark, Jenny E.E.; Swenson, Ulf; Anderberg, Arne A. (2006):**

**Accounting for Variation of Substitution Rates through Time in Bayesian Phylogeny Reconstruction of Sapotoideae (Sapotaceae).**

In: Molecular Phylogenetics and Evolution 39 (3), p. 706–721. DOI: 10.1016/j.ympev.2006.01.018.

**Swenson, Ulf; Anderberg, Arne A. (2005):**

**Phylogeny, Character Evolution, and Classification of Sapotaceae (Ericales).**

In: Cladistics 21 (2), p. 101–130. DOI: 10.1111/j.1096-0031.2005.00056.x.

**Swenson, Ulf; Bartish, Igor V.; Munzinger, Jérôme (2007):**

**Phylogeny, diagnostic Characters and generic limitation of Australasian Chrysophylloideae (Sapotaceae, Ericales): Evidence from ITS Sequence Data and Morphology.**
In: Cladistics 23 (3), p. 201–228. DOI: 10.1111/j.1096-0031.2006.00141.x.

SWENSON, ULF; LOWRY, PORTER PRESCOTT; MUNZINGER, JÉRÔME; RYDIN, CATARINA; BARTISH, IGOR V. (2008):
**Phylogeny and generic limits in the *Niemeyera complex* of New Caledonian Sapotaceae: Evidence of multiple Origins of the anisomerous Flower.**
In: Molecular Phylogenetics and Evolution 49 (3), p. 909–929. DOI: 10.1016/j.ympev.2008.09.022.

SWENSON, ULF; MUNZINGER, JÉRÔME (2010):
**Taxonomic Revision of *Pycnandra* Subgenus *Trouettia* (Sapotaceae), with six new Species from New Caledonia.**
In: Australian Systematic Botany 23 (5), p. 333–370. DOI: 10.1071/SB10025.

SWENSON, ULF; MUNZINGER, JÉRÔME (2012):
**Revision of *Pichonia* (Sapotaceae) in New Caledonia.**
In: Australian Systematic Botany 25 (1), p. 31–48. DOI: 10.1071/SB11027.

SWENSON, ULF; MUNZINGER, JÉRÔME; BARTISH, IGOR V. (2007):
**Molecular Phylogeny of *Planchonella* (Sapotaceae) and eight new Species from New Caledonia.**
In: Taxon 56 (2), p. 329–354. DOI: 10.1002/tax.562007.

SWENSON, ULF; NYLINDER, STEPHAN; MUNZINGER, JÉRÔME (2013):
**Towards a natural Classification of Sapotaceae Subfamily Chrysophylloideae in *Oceania* and Southeast Asia based on nuclear Sequence Data.**
In: Taxon 62 (4), p. 746–770. DOI: 10.12705/624.11.

SWENSON, ULF; RICHARDSON, JAMES E.; BARTISH, IGOR V. (2008):
**Multi-gene Phylogeny of the pantropical Subfamily Chrysophylloideae (Sapotaceae): Evidence of generic Polyphyly and extensive morphological Homoplasy.**
In: Cladistics 24 (6), p. 1006–1031. DOI: 10.1111/j.1096-0031.2008.00235.x.

TERRA-ARAUJO, MÁRIO H.; FARIA, APARECIDA D.; VICENTINI, ALBERTO; NYLINDER, STEPHAN; SWENSON, ULF (2015):
**Species Tree Phylogeny and Biogeography of the Neotropical Genus *Pradosia* (Sapotaceae, Chrysophylloideae).**
In: Molecular Phylogenetics and Evolution 87, p. 1–13. DOI: 10.1016/j.ympev.2015.03.007.

TRIONO, TEGUH; BROWN, ANTHONY H.D.; WEST, JUDY G.; CRISP, MICHAEL D. (2007):
**A Phylogeny of *Pouteria* (Sapotaceae) from Malesia and Australasia.**
In: Australian Systematic Botany 20 (2), p. 107–118. DOI: 10.1071/SB06011.

### Sarcolaenaceae

HONG-WA, CYNTHIA (2009):
**Endemic families of Madagascar. XII. Resurrection and taxonomic Revision of the Genera *Mediusella* (Cavaco) Hutchinson and *Xerochlamys* Baker (Sarcolaenaceae).**
In: Adansonia 31 (2), p. 311–339. DOI: 10.5252/a2009n2a7.

LOWRY, PORTER PRESCOTT II.; HAEVERMANS, THOMAS; LABAT, JEAN-NOEL; SCHATZ, GEORGE E.; LEROY, JEAN-FRANCOIS; WOLF, ANNE-ELIZABETH (2000):
**Endemic families of Madagascar. V. A synoptic Revision of *Eremolaena, Pentachlaena* and *Perrierodendron* (Sarcolaenaceae).**
In: Adansonia 22 (1), p. 11–31.

**Lowry, Porter Prescott II.; Schatz, George E.; Wolf, Anne-Elizabeth (2002):**
**Endemic families of Madagascar. VIII. A synoptic Revision of *Xyloolaena* Baill. (Sarcolaenaceae).**
In: Adansonia 24 (1), p. 7–19.

**Schatz, George E.; Lowry, Porter Prescott II.; Wolf, Anne-Elizabeth (2000):**
**Endemic families of Madagascar. VI. A synoptic Revision of *Rhodolaena* (Sarcolaenaceae).**
In: Adansonia 22 (2), p. 239–252.

**Schatz, George E.; Lowry, Porter Prescott II.; Wolf, Anne-Elizabeth (2001):**
**Endemic families of Madagascar. VII. A synoptic Revision of *Leptolaena* Thouars sensu stricto (Sarcolaenaceae).**
In: Adansonia 23 (2), p. 171–189.

### Sarraceniaceae

**Ellison, Aaron M.; Butler, Elena D.; Hicks, Emily Jean; Naczi, Robert F. C.; Calie, Patrick J.; Bell, Charles D.; Davis, Charles C. (2012):**
**Phylogeny and Biogeography of the carnivorous Plant Family Sarraceniaceae.**
In: Public Library of Science One 7 (6), e39291. DOI: 10.1371/journal.pone.0039291.

**Stephens, Jessica D.; Rogers, Willie L.; Heyduk, Karolina; Cruse-Sanders, Jennifer M.; Determann, Ron O.; Glenn, Travis C.; Malmberg, Russell L. (2015):**
**Resolving phylogenetic Relationships of the recently radiated carnivorous Plant Genus *Sarracenia* using target Enrichment.**
In: Molecular Phylogenetics and Evolution 85, p. 76–87. DOI: 10.1016/j.ympev.2015.01.015.

### Saururaceae

**Meng, Shao-Wu; Chen, Zhi-Duan; Li, De-Zhu; Liang, Han-Xing (2002):**
**Phylogeny of Saururaceae based on mitochondrial matR Gene Sequence Data.**
In: Journal of Plant Research 115, p. 71–76. DOI: 10.1007/s102650200011.

### Saxifragaceae

**Deng, Jia-bin; Drew, Bryan T.; Mavrodiev, Evgeny V.; Gitzendanner, Matthew A.; Soltis, Pamela S.; Soltis, Douglas E. (2015):**
**Phylogeny, Divergence times, and historical Biogeography of the Angiosperm Family Saxifragaceae.**
In: Molecular Phylogenetics and Evolution 83, p. 86–98. DOI: 10.1016/j.ympev.2014.11.011.

**Ebersbach, Jana; Müllner-Riehl, Alexandra N.; Michalak, Ingo; Tkach, Natalia V.; Hoffmann, Matthias H.; Röser, Martin et al. (2017):**
**In and out of the Qinghai-Tibet Plateau: Divergence time Estimation and historical Biogeography of the large arctic-alpine Genus *Saxifraga* L.**
In: Journal of Biogeography 44 (4), p. 900–910. DOI: 10.1111/jbi.12899.

**Gao, Qing-Bo; Li, Yin-Hu; Gornall, Richard J.; Zhang, Zhuo-Xin; Zhang, Fa-Qi; Xing, Rui et al. (2015):**
**Phylogeny and Speciation in *Saxifraga* Sect. *Ciliatae* (Saxifragaceae): Evidence from psbA-trnH, trnL-F and ITS Sequences.**
In: Taxon 64 (4), p. 703–713. DOI: 10.12705/644.3.

**Mas de Xaxars, Gemma; García-Fernández, Alfredo; Barnola, Pere; Martín, Joan; Mercadé, Arnau; Vallès, Joan et al. (2015):**
**Phylogenetic and cytogenetic Studies reveal hybrid Speciation in *Saxifraga* Subsect. *Triplinervium* (Saxifragaceae).**

In: Journal of Systematics and Evolution 53 (1), p. 53–62. DOI: 10.1111/jse.12105.

**Okuyama, Yudai; Pellmyr, Olle; Kato, Makoto (2008):**
**Parallel Floral Adaptations to Pollination by Fungus Gnats within the Genus *Mitella* (Saxifragaceae).**
In: Molecular Phylogenetics and Evolution 46 (2), p. 560–575. DOI: 10.1016/j.ympev.2007.09.020.

**Soltis, Douglas E.; Tago-Nakazawa, Miyuki; Xiang, Qiu-Yun; Kawano, Shoichi; Murata, Jin; Wakabayashi, Michio; Hibsch-Jetter, Carola (2001):**
**Phylogenetic Relationships and Evolution in *Chrysosplenium* (Saxifragaceae) based on matK Sequence Data.**
In: American Journal of Botany 88 (5), p. 883–893.

**Tkach, Natalia V.; Röser, Martin; Hoffmann, Matthias H. (2015):**
**Molecular Phylogenetics, Character Evolution and Systematics of the Genus *Micranthes* (Saxifragaceae).**
In: Botanical Journal of the Linnean Society 178, p. 47–66.

**Tkach, Natalia V.; Röser, Martin; Miehe, Georg; Müllner-Riehl, Alexandra N.; Ebersbach, Jana; Favre, Adrien; Hoffmann, Matthias H. (2015):**
**Molecular Phylogenetics, Morphology and a revised Classification of the complex Genus *Saxifraga* (Saxifragaceae).**
In: Taxon 64 (6), p. 1159–1187. DOI: 10.12705/646.4.

**Vargas, Pablo; Morton, Cynthia M.; Jury, Stephen L. (1999):**
**Biogeographic Patterns in Mediterranean and Macaronesian Species of *Saxifraga* (Saxifragaceae) inferred from phylogenetic Analyses of ITS Sequences.**
In: American Journal of Botany 86 (5), p. 724–734.

**Winkler, Manuela; Tribsch, Andreas; Schneeweiss, Gerald M.; Brodbeck, Sabine; Gugerli, Felix; Holderegger, Rolf; Schönswetter, Peter (2013):**
**Strong nuclear differentiation contrasts with widespread sharing of Plastid DNA Haplotypes across Taxa in European purple Saxifrages (*Saxifraga* Section *Porphyrion* subsection *Oppositifoliae*).**
In: Botanical Journal of the Linnean Society 173, p. 622–636.

**Xiang, Chun-Lei; Gitzendanner, Matthew A.; Soltis, Douglas E.; Peng, Hua; Lei, Li-Gong (2012):**
**Phylogenetic Placement of the enigmatic and critically endangered Genus *Saniculiphyllum* (Saxifragaceae) inferred from combined Analysis of Plastid and nuclear DNA Sequences.**
In: Molecular Phylogenetics and Evolution 64 (2), p. 357–367. DOI: 10.1016/j.ympev.2012.04.010.

**Zhang, De-Jun; Chen, Sheng-Yun; Gao, Qing-Bo (2008):**
**Circumscription and Phylogeny of *Saxifraga* Sect. *Ciliatae*: Evidence from nrDNA ITS Sequences.**
In: Journal of Systematics and Evolution 46 (5), p. 667–675.

**Zhu, Wei-Dong; Nie, Ze-long; Wen, Jun; Sun, Hang (2013):**
**Molecular Phylogeny and Biogeography of *Astilbe* (Saxifragaceae) in Asia and Eastern North America.**
In: Botanical Journal of the Linnean Society 171, p. 377–394.

### Saxifragales

**Jian, Shu-Guang; Soltis, Pamela S.; Gitzendanner, Matthew A.; Moore, Michael J.; Li, Rui-Qi; Hendry, Tory A. et al. (2008):**

**Resolving an ancient, rapid Radiation in Saxifragales.**

In: Systematic Biology 57 (1), p. 38–57. DOI: 10.1080/10635150801888871.

**Soltis, Douglas E.; Mort, Mark E.; Latvis, Maribeth; Mavrodiev, Evgeny V.; O'Meara, Brian C.; Soltis, Pamela p. et al. (2013):**

**Phylogenetic Relationships and Character Evolution Analysis of Saxifragales using a Supermatrix Approach.**

In: American Journal of Botany 100 (5), p. 916–929. DOI: 10.3732/ajb.1300044.

### Schisandraceae

**Denk, Thomas; Oh, I.-C. (2005):**

**Phylogeny of Schisandraceae based on morphological Data: Evidence from modern Plants and the fossil Record.**

In: Plant Systematics and Evolution 256 (1-4), p. 113–145. DOI: 10.1007/s00606-005-0327-3.

**Hao, Gang (2001):**

**A phylogenetic Analysis of the Schisandraceae based on Morphology and nuclear ribosomal ITS Sequences.**

In: Botanical Journal of the Linnean Society 135 (4), p. 401–411. DOI: 10.1006/bojl.2000.0420.

**Hao, Gang; Saunders, Richard M.K.; Chye, Mee-Len (2000):**

**A phylogenetic Analysis of the Illiciaceae based on Sequences of internal transcribed Spacers (ITS) of nuclear ribosomal DNA.**

In: Plant Systematics and Evolution 223, p. 81–90.

**Liu, Zhong; Hao, Gang; Luo, Yi-bo; Thien, Leonard B.; Rosso, Samuel W.; Lu, An-Ming; Chen, Zhi-Duan (2006):**

**Phylogeny and Androecial Evolution in Schisandraceae, inferred from Sequences of Nuclear Ribosomal DNA ITS and Chloroplast DNA trnL-F Regions.**

In: International Journal of Plant Sciences 167 (3), p. 539–550.

### Schizaeaceae

**Wikström, Niklas; Kenrick, Paul; Vogel, Johannes C. (2002):**

**Schizaeaceae: a phylogenetic approach.**

In: Review Palaobotany and Palynology 119, p. 35–50.

### Schlegeliaceae

**Barringer, Kerry (2004):**

**A Revision of *Gibsoniothamnus* L.O.Williams (Schlegeliaceae).**

In: Brittonia 56 (3), p. 213–237.

### Scrophulariaceae

**Archibald, Jenny K.; Mort, Mark E.; Wolfe, Andrea D. (2005):**

**Phylogenetic Relationships within *Zaluzianskya* (Scrophulariaceae s. s., Tribe Manuleeae): Classification based on DNA Sequences from Multiple Genomes and Implications for Character Evolution and Biogeography.**

In: Systematic Botany 30 (1), p. 196–215.

**Gándara, Etelvina; Sosa, Victoria (2013):**

**Testing the Monophyly and Position of the North American shrubby Desert Genus *Leucophyllum* (Scrophulariaceae: Leucophylleae).**

In: Botanical Journal of the Linnean Society 171, p. 508–518.

**Gándara, Etelvina; Sosa, Victoria (2014):**
**Spatio-temporal Evolution of *Leucophyllum pringlei* and allies (Scrophulariaceae): a Group endemic to North American xeric Regions.**
In: Molecular Phylogenetics and Evolution 76, p. 93–101. DOI: 10.1016/j.ympev.2014.02.027.

**Ghahremaninejad, Farrokh; Riahi, Mehrshid; Babaei, Melina; Attar, Farideh; Behçet, Lütfi; Sonboli, Ali (2014):**
**Monophyly of *Verbascum* (Scrophularieae : Scrophulariaceae): Evidence from nuclear and Plastid phylogenetic Analyses.**
In: Australian Journal of Botany 62 (8), p. 638–646. DOI: 10.1071/BT14159.

**Karaveliogullari, Faik Ahmet; Aytaç, Zeki (2008):**
**Revision of the Genus *Verbascum* L. (Group A) in Turkey.**
In: Botany Research Journal 1 (1), p. 9–32.

**Kornhall, Per (2004):**
**New Circumscription of the Tribe Limoselleae (Scrophulariaceae) that includes the Taxa of the Tribe Manuleeae.**
In: Botanical Journal of the Linnean Society 146, p. 453–467.

**Kornhall, Per; Heidari, Nahid; Bremer, Birgitta (2001):**
**Selagineae and Manuleeae, two Tribes or one? Phylogenetic studies in the Scrophulariaceae.**
In: Plant Systematics and Evolution 228, p. 199–218.

**Mosyakin, Sergei L.; Tsymbalyuk, Zoya M. (2015):**
**Pollen Morphology of Tribes Aptosimeae and Myoporeae supports the phylogenetic Pattern in early-branching Scrophulariaceae revealed by molecular Studies.**
In: Willdenowia 45 (2), p. 209–222. DOI: 10.3372/wi.45.45207.

**Navarro-Pérez, María L.; López, Josefa; Fernández-Mazuecos, Mario; Rodríguez-Riaño, Tomás; Vargas, Pablo; Ortega-Olivencia, Ana (2013):**
**The role of Birds and Insects in Pollination Shifts of *Scrophularia* (Scrophulariaceae).**
In: Molecular Phylogenetics and Evolution 69 (1), p. 239–254. DOI: 10.1016/j.ympev.2013.05.027.

**Oxelman, Bengt; Kornhall, Per; Olmstead, Richard G.; Bremer, Birgitta (2005):**
**Further disintegration of Scrophulariaceae.**
In: Taxon 54 (2), p. 411–425.

**Scheunert, Agnes; Heubl, Günther (2011):**
**Phylogenetic Relationships among New World *Scrophularia* L. (Scrophulariaceae): new Insights inferred from DNA Sequence Data.**
In: Plant Systematics and Evolution 291 (1-2), p. 69–89. DOI: 10.1007/s00606-010-0369-z.

**Scheunert, Agnes; Heubl, Günther (2014):**
**Diversification of *Scrophularia* (Scrophulariaceae) in the Western Mediterranean and Macaronesia - phylogenetic Relationships, reticulate Evolution and biogeographic Patterns.**
In: Molecular Phylogenetics and Evolution 70, p. 296–313. DOI: 10.1016/j.ympev.2013.09.023.

**Tank, David C.; Beardsley, Paul M.; Kelchner, Scot A.; Olmstead, Richard G. (2006):**
**Review of the Systematics of Scrophulariaceae s.l. and their current Disposition.**
In: Australian Systematic Botany 19 (4), p. 289–307. DOI: 10.1071/SB05009.

Verboom, George Anthony; Herron, Margaret L.; Moncrieff, Glenn R.; Slingsby, Jasper A. (2016):
**Maintenance of Species integrity in the context of a recent Radiation: the case of *Jamesbrittenia* (Scrophulariaceae: Limoselleae) in southern Africa.**
In: Botanical Journal of the Linnean Society 182, p. 115–139.

### Selaginellaceae

Arrigo, Nils; Therrien, James; Anderson, Cajsa Lisa; Windham, Michael D.; Haufler, Christopher H.; Barker, Michael p. (2013):
**A total Evidence approach to Understanding phylogenetic Relationships and ecological Diversity in *Selaginella* Subg. *Tetragonostachys*.**
In: American Journal of Botany 100 (8), p. 1672–1682. DOI: 10.3732/ajb.1200426.

Korall, Petra; Kenrick, Paul (2004):
**The phylogenetic History of Selaginellaceae based on DNA Sequences from the Plastid and Nucleus: extreme Substitution Rates and Rate Heterogeneity.**
In: Molecular Phylogenetics and Evolution 31 (3), p. 852–864. DOI: 10.1016/j.ympev.2003.10.014.

Weststrand, Stina; Korall, Petra (2016):
**A subgeneric Classification of *Selaginella* (Selaginellaceae).**
In: American Journal of Botany 103 (12), p. 2160–2169. DOI: 10.3732/ajb.1600288.

Weststrand, Stina; Korall, Petra (2016):
**Phylogeny of Selaginellaceae: There is Value in Morphology after all!**
In: American Journal of Botany 103 (12), p. 2136–2159. DOI: 10.3732/ajb.1600156.

Zhou, Xin-Mao; Jiang, Li-Jü; Zhang, Liang; Gao, Xin-Fen; He, Zhao-Rong; Zhang, Li-Bing (2015):
**Spore Morphology of *Selaginella* (Selaginellaceae) from China and its systematic Significance.**
In: Phytotaxa 237 (1), p. 1. DOI: 10.11646/phytotaxa.237.1.1.

Zhou, Xin-Mao; Rothfels, Carl J.; Zhang, Liang; He, Zhao-Rong; Le Péchon, Timothée; He, Hai et al. (2016):
**A large-scale Phylogeny of the Lycophyte Genus *Selaginella* (Selaginellaceae: Lycopodiopsida) based on Plastid and nuclear Loci.**
In: Cladistics 32 (4), p. 360–389. DOI: 10.1111/cla.12136.

Zhou, Xin-Mao; Zhang, Li-Bing (2015):
**A Classification of *Selaginella* (Selaginellaceae) based on molecular (chloroplast and nuclear), macromorphological, and Spore Features.**
In: Taxon 64 (6), p. 1117–1140. DOI: 10.12705/646.2.

### Setchellanthaceae

Hernández-Hernández, Tania; Colorado, Wendy B.; Sosa, Victoria (2013):
**Molecular Evidence for the Origin and evolutionary History of the rare American Desert monotypic Family Setchellanthaceae.**
In: Organisms Diversity and Evolution 13 (4), p. 485–496. DOI: 10.1007/s13127-013-0136-4.

### Simaroubaceae

Clayton, Joshua W.; Fernando, Edwino S.; Soltis, Pamela S.; Soltis, Douglas E. (2007):
**Molecular Phylogeny of the Tree-of-Heaven Family (Simaroubaceae) based on Chloroplast and Nuclear Markers.**
In: International Journal of Plant Sciences 168 (9), p. 1325–1339. DOI: 10.1086/521796.

Clayton, Joshua W.; Soltis, Pamela S.; Soltis, Douglas E. (2009):

**Recent long-distance Dispersal overshadows ancient biogeographical Patterns in a pantropical Angiosperm Family (Simaroubaceae, Sapindales).**
In: Systematic Biology 58 (4), p. 395–410. DOI: 10.1093/sysbio/syp041.

**Devecchi, Marcelo Fernando; Thomas, William Wayt; Plunkett, Gregory M.; Pirani, José Rubens (2018):**
**Testing the Monophyly of *Simaba* (Simaroubaceae): Evidence from five molecular Regions and Morphology.**
In: Molecular Phylogenetics and Evolution 120, p. 63–82. DOI: 10.1016/j.ympev.2017.11.024.

### Siparunaceae

**Renner, Susanne S.; Hausner, Gerlinde (2000):**
**New Species of *Siparuna* (Siparunaceae) III. Three new Species and one newly ranked Entity from Colombia, Ecuador, and Peru.**
In: Novon: A Journal for Botanical Nomenclature 10, p. 134–143.

**Renner, Susanne S.; Hausner, Gerlinde (2005):**
**New Species of *Siparuna* (Siparunaceae) IV. A new Subcanopy Tree from White-Sand Areas in Brazil and Venezuela.**
In: Novon: A Journal for Botanical Nomenclature 15, p. 202–206.

### Smilacaceae

**Chen, Chen; Qi, Zhe-Chen; Xu, Xi-Hui; Comes, Hans Peter; Koch, Marcus A.; Jin, Xin-Jie et al. (2014):**
**Understanding the formation of Mediterranean-African-Asian disjunctions: Evidence for Miocene climate-driven Vicariance and recent long-distance Dispersal in the Tertiary relict *Smilax aspera* (Smilacaceae).**
In: the new Phytologist 204 (1), p. 243–255. DOI: 10.1111/nph.12910.

**Chen, Shi-Chao (2006):**
**A phylogenetic Analysis of the Smilacaceae based on morphological Data.**
In: Acta Phytotaxonomica Sinica 44 (2), p. 113–125. DOI: 10.1360/aps050065.

**Fu, Cheng-Xin; Kong, Hanghui; Qiu, Ying-Xiong; Cameron, Kenneth M. (2005):**
**Molecular Phylogeny of the East Asian–North American disjunct *Smilax* Sect. *Nemexia* (Smilacaceae).**
In: International Journal of Plant Sciences 166 (2), p. 301–309.

**Li, Pan; Li, Mimi; Shi, Yin; Zhao, Yunpeng; Wan, Ying; Fu, Cheng-Xin; Cameron, Kenneth M. (2013):**
**Phylogeography of North American herbaceous *Smilax* (Smilacaceae): Combined AFLP and cpDNA Data support a northern Refugium in the Driftless Area.**
In: American Journal of Botany 100 (4), p. 801–814. DOI: 10.3732/ajb.1200250.

**Qi, Zhe-Chen; Cameron, Kenneth M.; Li, Pan; Zhao, Yunpeng; Chen, Shi-Chao; Chen, Guangcun; Fu, Cheng-Xin (2013):**
**Phylogenetics, Character Evolution, and Distribution Patterns of the Greenbriers, Smilacaceae (Liliales), a near-cosmopolitan Family of Monocots.**
In: Botanical Journal of the Linnean Society 173 (4), p. 535–548. DOI: 10.1111/boj.12096.

**Qi, Zhe-Chen; Li, P. A.N.; Fu, Cheng-Xin (2013):**
**New Combinations and a new Name in *Smilax* for Species of *Heterosmilax* in Eastern and Southeast Asian Smilacaceae (Liliales).**
In: Phytotaxa 117 (2), p. 58. DOI: 10.11646/phytotaxa.117.2.4.

**Zhao, Yunpeng; Qi, Zhe-Chen; Ma, Weiwei; Dai, Qiongyan; Li, Pan; Cameron, Kenneth M. et al. (2013):**

**Comparative Phylogeography of the *Smilax hispida* Group (Smilacaceae) in Eastern Asia and North America - Implications for allopatric Speciation, causes of Diversity Disparity, and Origins of temperate Elements in Mexico.**
In: Molecular Phylogenetics and Evolution 68 (2), p. 300–311. DOI: 10.1016/j.ympev.2013.03.025.

### Solanaceae

**Ames, Mercedes; Spooner, David M. (2010):**
**Phylogeny of *Solanum* Series *Piurana* and related Species in *Solanum* Section *Petota* based on five conserved ortholog Sequences.**
In: Taxon 59 (4), p. 1091–1101.

**Ando, Toshio; Kokubun, Hisashi; Watanabe, Hitoshi; Tanaka, Norio; Yukawa, Tomohisa; Hashimoto, Goro et al. (2005):**
**Phylogenetic Analysis of *Petunia* sensu Jussieu (Solanaceae) using Chloroplast DNA Rflp.**
In: Annals of Botany 96 (2), p. 289–297. DOI: 10.1093/aob/mci177.

**Aoki, S.; Ito, Motomi (2000):**
**Molecular Phylogeny of *Nicotiana* (Solanaceae) based on the Nucleotide Sequence of the matK Gene.**
In: Plant Biology 2, p. 316–324.

**Aubriot, Xavier; Fls, Paramjit Singh; Knapp, Sandra (2016):**
**Tropical Asian Species show that the Old World Clade of 'spiny Solanums' (*Solanum* Subgenus *Leptostemonum* pro parte: Solanaceae) is not monophyletic.**
In: Botanical Journal of the Linnean Society 181, p. 199–223.

**Averett, John E. (2009):**
***Schraderanthus*, a new Genus of Solanaceae.**
In: Phytologia 91 (1), p. 54–61.

**Bohs, Lynn (2007):**
**Phylogeny of the *Cyphomandra* Clade of the Genus *Solanum* (Solanaceae) based on ITS Sequence Data.**
In: Taxon 56 (4), p. 1012–1026.

**Carrizo García, Carolina; Wahlert, Gregory A.; Orozco, Clara Inés; Barboza, Gloria Estela; Bohs, Lynn (2015):**
**Phylogeny of the Andean Genus *Deprea* (Physalideae, Solanaceae): testing the generic Circumscription.**
In: Phytotaxa 238 (1), p. 71. DOI: 10.11646/phytotaxa.238.1.3.

**Chase, Mark W.; Knapp, Sandra; Cox, Antony V.; Clarkson, James J.; Butsko, Yelena; Joseph, Jeffrey A. et al. (2003):**
**Molecular Systematics, Gish and the Origin of hybrid Taxa in *Nicotiana* (Solanaceae).**
In: Annals of Botany 92 (1), p. 107–127. DOI: 10.1093/aob/mcg087.

**Chase, Sandra Knapp: Mark W.; Clarkson, James J. (2004):**
**Nomenclatural Changes and a new sectional Classification in *Nicotiana* (Solanaceae).**
In: Taxon 53 (1), p. 73–82.

**Chen, Sumei; Matsubara, Kiyoshi; Omori, Takahiro; Kokubun, Hisashi; Kodama, Hiroaki; Watanabe, Hitoshi et al. (2007):**

**Phylogenetic Analysis of the Genus *Petunia* (Solanaceae) based on the Sequence of the Hf1 gene.**
In: Journal of Plant Research 120 (3), p. 385–397. DOI: 10.1007/s10265-006-0070-z.

CLARKSON, JAMES J.; KELLY, LAURA J.; LEITCH, ANDREW R.; KNAPP, SANDRA; CHASE, MARK W. (2010):
**Nuclear Glutamine Synthetase Evolution in *Nicotiana*: Phylogenetics and the Origins of allotetraploid and homoploid (diploid) Hybrids.**
In: Molecular Phylogenetics and Evolution 55 (1), p. 99–112. DOI: 10.1016/j.ympev.2009.10.003.

CLARKSON, JAMES J.; KNAPP, SANDRA; GARCIA, VICENTE F.; OLMSTEAD, RICHARD G.; LEITCH, ANDREW R.; CHASE, MARK W. (2004):
**Phylogenetic Relationships in *Nicotiana* (Solanaceae) inferred from multiple Plastid DNA Regions.**
In: Molecular Phylogenetics and Evolution 33 (1), p. 75–90. DOI: 10.1016/j.ympev.2004.05.002.

DEANNA, ROCÍO; BARBOZA, GLORIA ESTELA; CARRIZO GARCÍA, CAROLINA (2018):
**Phylogenetic Relationships of *Deprea*: new Insights into the evolutionary History of physaloid Groups.**
In: Molecular Phylogenetics and Evolution 119, p. 71–80. DOI: 10.1016/j.ympev.2017.11.001.

DILLON, MICHAEL O.; TU, TIE-YAO; SOEJIMA, AKIKO; YI, TING-SHUANG; NIE, ZE-LONG; TYE, ALAN; WEN, JUN (2007):
**Phylogeny of *Nolana* (Nolaneae, Solanoideae, Solanaceae) as inferred from granule-bound Starch Synthase I (Gbssi) Sequences.**
In: Taxon 56 (4), p. 1000–1011.

FAJARDO, DIEGO; SPOONER, DAVID M. (2011):
**Phylogenetic Relationships of *Solanum* Series *Conicibaccata* and related Species in *Solanum* Section *Petota* inferred from five conserved ortholog Sequences.**
In: Systematic Botany 36 (1), p. 163–170. DOI: 10.1600/036364411x553252.

FILIPOWICZ, NATALIA; RENNER, SUSANNE P. (2012):
***Brunfelsia* (Solanaceae): a Genus evenly divided between South America and Radiations on Cuba and other Antillean Islands.**
In: Molecular Phylogenetics and Evolution 64 (1), p. 1–11. DOI: 10.1016/j.ympev.2012.02.026.

FLS, MARIA. P. VORONTSOVA; STERN, STEPHEN R.; BOHS, LYNN; KNAPP, SANDRA (2013):
**African spiny *Solanum* (Subgenus *Leptostemonum*, Solanaceae): a thorny phylogenetic Tangle.**
In: Botanical Journal of the Linnean Society 173, p. 176–193.

FREGONEZI, JEFFERSON NUNES; FREITAS, LORETA BRANDÃO; BONATTO, SANDRO LUIS; SEMIR, JOÃO; STEHMANN, RENATO (2012):
**Infrageneric Classification of *Calibrachoa* (Solanaceae) based on morphological and molecular Evidence.**
In: Taxon 61 (1), p. 120–130.

FUKUDA, TATSUYA; YOKOYAMA, JUN U.N.; OHASHI, HIROYOSHI (2001):
**Phylogeny and Biogeography of the Genus *Lycium* (Solanaceae): Inferences from Chloroplast DNA Sequences.**
In: Molecular Phylogenetics and Evolution 19 (2), p. 246–258. DOI: 10.1006/mpev.2001.0921.

GARCIA, VICENTE F.; OLMSTEAD, RICHARD G. (2003):
**Phylogenetics of Tribe Anthocercideae (Solanaceae) based on ndhF and trnL/F Sequence Data.**

In: Systematic Botany 28 (3), p. 609–615.

**Gates, Daniel J.; Pilson, Diana; Smith, Stacey Dewitt (2018):**
**Filtering of target Sequence capture Individuals facilitates Species Tree Construction in the plant Subtribe Iochrominae (Solanaceae).**
In: Molecular Phylogenetics and Evolution 123, p. 26–34. DOI: 10.1016/j.ympev.2018.02.002.

**Jamil, Ishrat; Qamarunnisa, Syeda; Azhar, Abid; Shinwari, Zabta K.; Ali, Syed Irtifaq; Qaiser, Muhammad (2014):**
**Subfamilial Relationships within Solanaceae as inferred from atpβ-rbcL Intergenic Spacer.**
In: Pakistan Journal of Botany 46 (2), p. 585–590.

**Kelly, Laura J.; Leitch, Andrew R.; Clarkson, James J.; Hunter, Robin B.; Knapp, Sandra; Chase, Mark W. (2010):**
**Intragenic Recombination Events and Evidence for hybrid Speciation in *Nicotiana* (Solanaceae).**
In: Molecular Biology and Evolution 27 (4), p. 781–799. DOI: 10.1093/molbev/msp267.

**Kelly, Laura J.; Leitch, Andrew R.; Clarkson, James J.; Knapp, Sandra; Chase, Mark W. (2013):**
**Reconstructing the complex evolutionary Origin of wild allopolyploid Tobaccos (*Nicotiana* Section *suaveolentes*).**
In: Evolution 67 (1), p. 80–94. DOI: 10.1111/j.1558-5646.2012.01748.x.

**Knapp, Sandra; Vorontsova, Maria p. (2016):**
**A Revision of the "African Non-Spiny" Clade of *Solanum* L. (*Solanum* Sections *Afrosolanum* Bitter, *Benderianum* Bitter, *Lemurisolanum* Bitter, *Lyciosolanum* Bitter, *Macronesiotes* Bitter, and *Quadrangulare* Bitter: Solanaceae).**
In: PhytoKeys (66), p. 1–142. DOI: 10.3897/phytokeys.66.8457.

**Levin, Rachel A.; Bernardello, Gabriel; Whiting, Carolyn; Miller, Jill p. (2011):**
**A new generic Circumscription in Tribe Lycieae (Solanaceae).**
In: Taxon 60 (3), p. 681–690.

**Levin, Rachel A.; Blanton, Jessica; Miller, Jill p. (2009):**
**Phylogenetic utility of nuclear nitrate reductase: a multi-locus Comparison of nuclear and Chloroplast Sequence Data for Inference of Relationships among American Lycieae (Solanaceae).**
In: Molecular Phylogenetics and Evolution 50 (3), p. 608–617. DOI: 10.1016/j.ympev.2008.12.005.

**Levin, Rachel A.; Keyes, Edmund M.; Miller, Jill p. (2015):**
**Evolutionary Relationships, Gynodioecy, and Polyploidy in the Galápagos Endemic *Lycium minimum* (Solanaceae).**
In: International Journal of Plant Sciences 176 (2), p. 197–210. DOI: 10.1086/679492.

**Levin, Rachel A.; Miller, Jill p. (2005):**
**Relationships within Tribe Lycieae (Solanaceae): Paraphyly of *Lycium* and multiple Origins of Gender Dimorphism.**
In: American Journal of Botany 92 (12), p. 2044–2053.

**Levin, Rachel A.; Myers, Nicole R.; Bohs, Lynn (2006):**
**Phylogenetic Relationships among the ''spiny *Solanums*'' (*Solanum* Subgenus *Leptostemonum*, Solanaceae).**
In: American Journal of Botany 93 (1), p. 157–169.

**Levin, Rachel A.; Watson, Kimberly; Bohs, Lynn (2005):**

**A Four-gene Study of evolutionary Relationships in *Solanum* Section *Acanthophora*.**

In: American Journal of Botany 92 (4), p. 603–612.

**Levin, Rachel A.; Whelan, Andrew; Miller, Jill p. (2009):**

**The utility of nuclear conserved Ortholog Set II (Cosii) genomic Regions for species-level phylogenetic Inference in *Lycium* (Solanaceae).**

In: Molecular Phylogenetics and Evolution 53 (3), p. 881–890. DOI: 10.1016/j.ympev.2009.08.016.

**Li, Hong-Qing; Gui, Ping; Xiong, Shen-Zhan; Averett, John E. (2013):**

**The generic Position of two Species of Tribe Physaleae (Solanaceae) inferred from three DNA Sequences: A case Study on *Physaliastrum* and *Archiphysalis*.**

In: Biochemical Systematics and Ecology 50, p. 82–89. DOI: 10.1016/j.bse.2013.03.038.

**Marks, Claire E.; Newbigin, Ed; Ladiges, Pauline Y. (2011):**

**Comparative Morphology and Phylogeny of *Nicotiana* Section *Suaveolentes* (Solanaceae) in Australia and the South Pacific.**

In: Australian Systematic Botany 24 (2), p. 61. DOI: 10.1071/SB11006.

**Martine, Christopher T.; Anderson, Gregory J.; Les, Donald H. (2009):**

**Gender-bending Aubergines: molecular Phylogenetics of cryptically dioecious *Solanum* in Australia.**

In: Australian Systematic Botany 22 (2), p. 107–120. DOI: 10.1071/SB07039.

**Martine, Christopher T.; Vanderpool, Dan; Anderson, Gregory J.; Les, Donald H. (2006):**

**Phylogenetic Relationships of Andromonoecious and Dioecious Australian Species of *Solanum* Subgenus *Leptostemonum* Section *Melongena*: Inferences from ITS Sequence Data.**

In: Systematic Botany 31 (2), p. 410–420.

**Martins, Talline R.; Barkman, Todd J. (2005):**

**Reconstruction of Solanaceae Phylogeny using the Nuclear Gene SAMT.**

In: Systematic Botany 30 (2), p. 435–447.

**Miller, Ryan J.; Mione, Thomas; Phan, Hanh-La; Olmstead, Richard G. (2011):**

**Color by Numbers: Nuclear Gene Phylogeny of *Jaltomata* (Solanaceae), Sister Genus to *Solanum*, supports three Clades differing in Fruit Color.**

In: Systematic Botany 36 (1), p. 153–162. DOI: 10.1600/036364411X553243.

**Montero-Castro, Juan Carlos; Delgado-Salinas, Alfonso; Luna, Efraín de; Eguiarte, Luis E. (2006):**

**Phylogenetic Analysis of *Cestrum* Section *Habrothamnus* (Solanaceae) based on Plastid and Nuclear DNA Sequences.**

In: Systematic Botany 31 (4), p. 843–850.

**Moré, Marcela; Cocucci, Andrea A.; Sérsic, Alicia N.; Barboza, Gloria Estela (2015):**

**Phylogeny and Floral trait Evolution in *Jaborosa* (Solanaceae).**

In: Taxon 64 (3), p. 523–534. DOI: 10.12705/643.8.

**Muchhala, Nathan; Johnsen, Sönke; Smith, Stacey Dewitt (2014):**

**Competition for Hummingbird Pollination shapes Flower Color Variation in Andean Solanaceae.**

In: Evolution 68 (8), p. 2275–2286. DOI: 10.1111/evo.12441.

**Olmstead, Richard G.; Bohs, Lynn; Migid, Hala Abdel; Santiago-Valentin, Eugenio; Garcia, Vicente F.; Collier, Sarah M. (2008):**

**A molecular Phylogeny of the Solanaceae.**
In: Taxon 57 (4), p. 1159–1181.

OREJUELA, ANDRÉS; WAHLERT, GREGORY A.; OROZCO, CLARA INÉS; BARBOZA, GLORIA ESTELA; BOHS, LYNN (2017):
**Phylogeny of the Tribes Juanulloeae and Solandreae (Solanaceae).**
In: Taxon 66 (2), p. 379–392. DOI: 10.12705/662.6.

POCZAI, P.; TALLER, J.; SZABÓ, I. (2008):
**Analysis of phylogenetic Relationships in the Genus *Solanum* (Solanaceae) as revealed by Rapd Markers.**
In: Plant Systematics and Evolution 275 (1-2), p. 59–67. DOI: 10.1007/s00606-008-0051-x.

RAMOS-FREGONEZI, ALINE M. C.; FREGONEZI, JEFFERSON NUNES; CYBIS, GABRIELA B.; FAGUNDES, NELSON J. R.; BONATTO, SANDRO LUIS; FREITAS, LORETA BRANDÃO (2015):
**Were Sea Level Changes during the Pleistocene in the South Atlantic Coastal Plain a driver of Speciation in *Petunia* (Solanaceae)?**
In: BMC Evolutionary Biology 15, p. 92. DOI: 10.1186/s12862-015-0363-8.

RECK-KORTMANN, MAIKEL; SILVA-ARIAS, GUSTAVO ADOLFO; SEGATTO, ANA LÚCIA ANVERSA; MÄDER, GERALDO; BONATTO, SANDRO LUIS; FREITAS, LORETA BRANDÃO (2014):
**Multilocus Phylogeny Reconstruction: new Insights into the evolutionary History of the Genus *Petunia*.**
In: Molecular Phylogenetics and Evolution 81, p. 19–28. DOI: 10.1016/j.ympev.2014.08.022.

RECK-KORTMANN, MAIKEL; SILVA-ARIAS, GUSTAVO ADOLFO; STEHMANN, JOÃO R.; GREPPI, JULIÁN A.; FREITAS, LORETA BRANDÃO (2015):
**Phylogenetic Relationships of *Petunia patagonica* (Solanaceae) revealed by molecular and biogeographical Evidence.**
In: Phytotaxa 222 (1), p. 17. DOI: 10.11646/phytotaxa.222.1.2.

SÄRKINEN, TIINA E.; BARBOZA, GLORIA ESTELA; KNAPP, SANDRA (2015):
**True Black Nightshades: Phylogeny and Delimitation of the Morelloid Clade of *Solanum*.**
In: Taxon 64 (5), p. 945–958. DOI: 10.12705/645.5.

SÄRKINEN, TIINA E.; BOHS, LYNN; OLMSTEAD, RICHARD G.; KNAPP, SANDRA (2013):
**A phylogenetic Famework for evolutionary Study of the nightshades (Solanaceae): a dated 1000-tip Tree.**
In: BMC Evolutionary Biology 13, p. 214. DOI: 10.1186/1471-2148-13-214.

SMITH, STACEY DEWITT; BAUM, DAVID A. (2006):
**Phylogenetics of the Florally diverse Andean Clade Iochrominae (Solanaceae).**
In: American Journal of Botany 93 (8), p. 1140–1153.

STERN, STEPHEN R.; AGRA, FÁTIMA DE MARIA; BOHS, LYNN (2011):
**Molecular Delimitation of Clades within New World Species of the "spiny Solanums" (*Solanum* Subg. *Leptostemonum*).**
In: Taxon 60 (5), p. 1429–1441.

STERN, STEPHEN R.; BOHS, LYNN (2016):
**An eight Marker Phylogeny of *Solanum* Sect. *Micracantha* (Solanaceae).**
In: Systematic Botany 41 (1), p. 120–127. DOI: 10.1600/036364416X690589.

STERN, STEPHEN R.; WEESE, TERRI L.; BOHS, LYNN (2010):
**Phylogenetic Relationships in *Solanum* Section *Androceras* (Solanaceae).**
In: Systematic Botany 35 (4), p. 885–893. DOI: 10.1600/036364410X539934.

**Tate, Jennifer A.; Acosta, M. Cristina; McDill, Joshua R.; Moscone, Eduardo A.; Simpson, Beryl B.; Cocucci, Andrea A. (2009):**
**Phylogeny and Character Evolution in *Nierembergia* (Solanaceae): Molecular, Morphological, and Cytogenetic Evidence.**
In: Systematic Botany 34 (1), p. 198–206.

**Tepe, Eric J.; Bohs, Lynn (2010):**
**A molecular Phylogeny of *Solanum* Sect. *Pteroidea* (Solanaceae) and the Utility of Cosii Markers in resolving Relationships among closely related Species.**
In: Taxon 59 (3), p. 733–743.

**Tepe, Eric J.; Bohs, Lynn (2011):**
**A Revision of *Solanum* Section *Herpystichum*.**
In: Systematic Botany 36 (4), p. 1068–1087. DOI: 10.1600/036364411X605074.

**Tepe, Eric J.; Farruggia, Frank T.; Bohs, Lynn (2011):**
**A 10-gene Phylogeny of *Solanum* Section *Herpystichum* (Solanaceae) and a Comparison of phylogenetic Methods.**
In: American Journal of Botany 98 (8), p. 1356–1365. DOI: 10.3732/ajb.1000516.

**Tu, Tie-Yao; Dillon, Michael O.; Sun, Hang; Wen, Jun (2008):**
**Phylogeny of *Nolana* (Solanaceae) of the Atacama and Peruvian Deserts inferred from Sequences of four Plastid Markers and the nuclear Leafy second Intron.**
In: Molecular Phylogenetics and Evolution 49 (2), p. 561–573. DOI: 10.1016/j.ympev.2008.07.018.

**Tu, Tie-Yao; Volis, Sergei; Dillon, Michael O.; Sun, Hang; Wen, Jun (2010):**
**Dispersals of Hyoscyameae and Mandragoreae (Solanaceae) from the New World to Eurasia in the early Miocene and their biogeographic Diversification within Eurasia.**
In: Molecular Phylogenetics and Evolution 57 (3), p. 1226–1237. DOI: 10.1016/j.ympev.2010.09.007.

**Volkov, Roman A.; Komarova, Nataliya Y.; Panchuk, Irina I.; Hemleben, Vera (2003):**
**Molecular Evolution of rDNA external transcribed spacer and Phylogeny of Sect. *Petota* (genus *Solanum*).**
In: Molecular Phylogenetics and Evolution 29 (2), p. 187–202. DOI: 10.1016/S1055-7903(03)00092-7.

**Walsh, Brian M.; Hoot, Sara B. (2001):**
**Phylogenetic Relationships of *Capsicum* (Solanaceae) using DNA Sequences from Two Noncoding Regions: the Chloroplast atpB-rbcL Spacer Region and Nuclear waxy Introns.**
In: International Journal of Plant Sciences 162 (6), p. 1409–1418.

**Weese, Terri L.; Bohs, Lynn (2007):**
**A Three-Gene Phylogeny of the Genus *Solanum* (Solanaceae).**
In: Systematic Botany 32 (2), p. 445–463.

**Weese, Terri L.; Bohs, Lynn (2010):**
**Eggplant Origins: Out of Africa, into the Orient.**
In: Taxon 59 (1), p. 49–56.

**Whitson, Maggie; Manos, Paul p. (2005):**
**Untangling *Physalis* (Solanaceae) from the Physaloids: A Two-Gene Phylogeny of the Physalinae.**
In: Systematic Botany 30 (1), p. 216–230.

**Zamberlan, Priscilla M.; Rodrigues, Izabella M. C.; Geraldo Mäder, Luana Castro, João R. Stehmann; Bonatto, Sandro Luis; Freitas, Loreta Brandão (2015):**

**Re-evaluation of the generic Status of *Athenaea* and *Aureliana* (Withaniinae, Solanaceae) based on molecular Phylogeny and Morphology of the Calyx.**

In: Botanical Journal of the Linnean Society 177, p. 322–334.

**Zamora-Tavares, María del Pilar; Martínez, Mahinda; Magallón, Susana; Guzmán-Dávalos, Laura; Vargas-Ponce, Ofelia (2016):**

***Physalis* and physaloids: A recent and complex evolutionary History.**

In: Molecular Phylogenetics and Evolution 100, p. 41–50. DOI: 10.1016/j.ympev.2016.03.032.

### Stachyuraceae

**Zhu, Yu-Ping; Wen, Jun; Zhang, Zhi-Yun; Chen, Zhi-Duan (2006):**

**Evolutionary Relationships and Diversification of Stachyuraceae based on Sequences of four Chloroplast Markers and the nuclear ribosomal ITS Region.**

In: Taxon 55 (4), p. 931–940. DOI: 10.2307/25065687.

### Staphyleaceae

**Harris, A. J.; Chen, Ping-Ting; Xu, Xin-Wei; Zhang, Jian-Qiang; Yang, Xue; Wen, Jun (2017):**

**A molecular Phylogeny of Staphyleaceae: Implications for generic Delimitation and classical biogeographic Disjunctions in the Family.**

In: Journal of Systematics and Evolution 55 (2), p. 124–141. DOI: 10.1111/jse.12236.

### Stemonaceae

**Fan, Lan-Lan; Zhu, Shu; Chen, Hu-Biao; Yang, Dong-Hui; Cai, Shao-Qing; Komatsu, Katsuko (2009):**

**Molecular Analysis of *Stemona* Plants in China based on Sequences of four Chloroplast DNA Regions.**

In: Biological and Pharmaceutical Bulletin 32 (8), p. 1439–1446. DOI: 10.1248/bpb.32.1439.

**Li, En-Xiang; Yi, Sun; Qiu, Ying-Xiong; Guo, Jiang-Tao; Comes, Hans Peter; Fu, Cheng-Xin (2008):**

**Phylogeography of two East Asian Species in *Croomia* (Stemonaceae) inferred from Chloroplast DNA and Issr Fingerprinting Variation.**

In: Molecular Phylogenetics and Evolution 49 (3), p. 702–714. DOI: 10.1016/j.ympev.2008.09.012.

### Stemonuraceae

**Munzinger, Jérôme; McPherson, Gordon; Lowry, Porter Prescott II. (2008):**

**A second Species in the endemic New Caledonian Genus *Gastrolepis* (Stemonuraceae) and its Implications for the conservation Status of high-altitude Maquis Vegetation: coherent application of the IUCN Red List Criteria is urgently needed in New Caledonia.**

In: Botanical Journal of the Linnean Society 157, p. 775–783.

**Schori, Melanie (2010):**

**A Systematic Revision of *Gomphandra* (Stemonuraceae).**

Dissertation. Ohio University, Faculty of the College of Arts and Sciences of Ohio University.

**Schori, Melanie; Utteridge, Timothy M.A. (2010):**

**Three new Species and a new name in Southeast Asian *Gomphandra* (Stemonuraceae / Icacinaceae s.l.).**

In: Blumea 55 (2), p. 189–195. DOI: 10.3767/000651910X527699.

**Utteridge, Timothy M.A. (2011):**

**A Revision of the Genus *Medusanthera* (Stemonuraceae, Icacinaceae s.l.).**

In: Kew Bulletin 66, p. 49–81.

### Sterculiaceae

DORR, L.J.; CHEEK, MARTIN R. (2011):

**The identity and typification of *Pimia* Seem. (Sterculiaceae).**

In: Kew Bulletin 66, p. 629–631.

WILKIE, PETER; CLARK, ALEXANDRA; PENNINGTON, R. TOBY; CHEEK, MARTIN R.; BAYER, CLEMENS; WILCOCK, CHRIS C. (2006):

**Phylogenetic Relationships within the Subfamily Sterculioideae (Malvaceae/Sterculiaceae-Sterculieae) using the Chloroplast Gene ndhF.**

In: Systematic Botany 31 (1), p. 160–170.

### Strelitziaceae

CRON, GLYNIS V.; PIRONE, CARY; BARTLETT, MADELAINE; KRESS, W. JOHN; SPECHT, CHELSEA D. (2012):

**Phylogenetic Relationships and Evolution in the Strelitziaceae (Zingiberales).**

In: Systematic Botany 37 (3), p. 606–619. DOI: 10.1600/036364412X648562.

### Stylidiaceae

GLENNY, DAVID P. (2009):

**A Revision of the Genus *Forstera* (Stylidiaceae) in New Zealand.**

In: New Zealand Journal of Botany 47 (3), p. 285–315. DOI: 10.1080/00288250909509811.

LAURENT, NADINA; BREMER, BIRGITTA; BREMER, KARE (1998):

**Phylogeny and Generic Interrelationships of the Stylidiaceae (Asterales), with a possible extreme Case of Floral Paedomorphosis.**

In: Systematic Botany 23 (3), p. 289. DOI: 10.2307/2419506.

WEGE, JULIET A. (2012):

**Navigating the Floral Milky Way: the Taxonomy of the microgeophytic Triggerplants (*Stylidium petiolare* and allies: Stylidiaceae).**

In: Australian Systematic Botany 25 (2), p. 138. DOI: 10.1071/SB12001.

### Styracaceae

FRITSCH, PETER W. (2001):

**Phylogeny and Biogeography of the flowering plant Genus *Styrax* (Styracaceae) based on Chloroplast DNA Restriction Sites and DNA Sequences of the Internal Transcribed Spacer Region.**

In: Molecular Phylogenetics and Evolution 19 (3), p. 387–408. DOI: 10.1006/mpev.2001.0933.

FRITSCH, PETER W. (2004):

**New Species and taxonomic Changes in *Styrax* (Styracaceae) from South America.**

In: Novon: A Journal for Botanical Nomenclature 14 (1), p. 43–57.

FRITSCH, PETER W.; MORTON, CYNTHIA M.; CHEN, TAO; MELDRUM, CANDICE (2001):

**Phylogeny and Biogeography of the Styracaceae.**

In: International Journal of Plant Sciences 162 (S6), S95-S116. DOI: 10.1086/323418.

YAO, XIAOHONG; YE, QIGANG; FRITSCH, PETER W.; CRUZ, BONI C.; HUANG, HONGWEN (2008):

**Phylogeny of *Sinojackia* (Styracaceae) based on DNA Sequence and Microsatellite Data: Implications for Taxonomy and Conservation.**

In: Annals of Botany 101 (5), p. 651–659. DOI: 10.1093/aob/mcm332.

### Symplocaceae

**Fritsch, Peter W.; Al, Frank (2015):**
**A taxonomic Revision of Antillean *Symplocos* (Symplocaceae).**
In: Phytotaxa 194 (1), p. 1. DOI: 10.11646/phytotaxa.194.1.1.

**Fritsch, Peter W.; Cruz, Boni C.; Almeda, Frank; Wang, Yuguo; Shi, Su-hua (2006):**
**Phylogeny of *Symplocos* based on DNA Sequences of the Chloroplast trnC-trnD Intergenic Region.**
In: Systematic Botany 31 (1), p. 181–192.

**Fritsch, Peter W.; Kelly, Lawrence M.; Wang, Yuguo; Almeda, Frank; Kriebel, Ricardo (2008):**
**Revised infrafamilial Classification of Symplocaceae based on phylogenetic Data from DNA Sequences and Morphology.**
In: Taxon 57 (3), p. 823–852. DOI: 10.1002/tax.573013.

**Soejima, Akiko; Nagamasu, Hidetoshi (2004):**
**Phylogenetic Analysis of Asian *Symplocos* (Symplocaceae) based on nuclear and Chloroplast DNA Sequences.**
In: Journal of Plant Research 117 (3), p. 199–207. DOI: 10.1007/s10265-004-0151-9.

**Wang, Yuguo; Fritsch, Peter W.; Shi, Su-hua; Almeda, Frank; Cruz, Boni C.; Kelly, Lawrence M. (2004):**
**Phylogeny and infrageneric Classification of *Symplocos* (Symplocaceae) inferred from DNA Sequence Data.**
In: American Journal of Botany 91 (11), p. 1901–1914. DOI: 10.3732/ajb.91.11.1901.

### Talinaceae

**Nyffeler, Reto; Eggli, Urs (2010):**
**Disintegrating Portulacaceae: A new familial Classification of the Suborder Portulacineae (Caryophyllales) based on molecular and morphological Data.**
In: Taxon 59 (1), p. 227–240.

### Tamaricaceae

**Arianmanesh, Reza; Mehregan, Iraj; Nejadsattari, Taher; Assadi, Mostafa (2015):**
**Molecular Phylogeny of *Tamarix* (Tamaricaceae) Species from Iran based on ITS Sequence Data.**
In: European Journal of Experimental Biology 5 (6), p. 44–50.

**Chen, Youhua (2013):**
**A Revisited Study on Phylogeography and phylogenetic Diversity of *Myricaria* (Tamaricaceae).**
In: Journal of Ecosystem & Ecography 3 (132). DOI: 10.4172/2157-7625.1000132.

**Gaskin, John F.; Ghahremaninejad, Farrokh; Zhang, Dao-Yuan; Londo, Jason P. (2004):**
**A systematic overviewn of Frankeniaceae and Tamaricaceae from nuclear rDNA and Plastid Sequence Data.**
In: Annals of the Missouri Botanical Garden 91, p. 401–409.

**Gaskin, John F.; Schaal, Barbara A. (2003):**
**Molecular phylogenetic Investigation of U.S. Invasive *Tamarix*.**
In: Systematic Botany 28 (1), p. 86–95.

**Sun, Likun; Yang, Ruiqi; Zhang, Baogui; Zhang, Gaosen; Wu, Xiukun; Zhang, Wei et al. (2016):**
**Phylogenetic Relationships among Species of *Tamarix* (Tamaricaceae) in China.**
In: Biochemical Systematics and Ecology 69, p. 213–221. DOI: 10.1016/j.bse.2016.10.003.

**Wang, Yong; Liu, Yifei; Liu, Songbai; Huang, Hongwen (2009):**

**Molecular Phylogeny of *Myricaria* (Tamaricaceae): Implications for Taxonomy and Conservation in China.**
In: Botanical Studies 50, p. 343–352.

ZHANG, MING-LI; HAO, XIAO-LI; SANDERSON, STEWART C.; VYACHESLAV, BYALT V.; SUKHORUKOV, ALEXANDER P.; ZHANG, X. I.A. (2014):
**Spatiotemporal Evolution of *Reaumuria* (Tamaricaceae) in Central Asia: Insights from molecular Biogeography.**
In: Phytotaxa 167 (1), p. 89. DOI: 10.11646/phytotaxa.167.1.5.

### Taxaceae

CHENG, Y.; NICOLSON, R. G.; TRIPP, K.; CHAW, SHU-MIAW (2000):
**Phylogeny of Taxaceae and Cephalotaxaceae Genera inferred from Chloroplast matK Gene and nuclear rDNA ITS Region.**
In: Molecular Phylogenetics and Evolution 14 (3), p. 353–365. DOI: 10.1006/mpev.1999.0710.

COLLINS, DENNIS; MILL, ROBERT R.; MÖLLER, MICHAEL (2003):
**Species Separation of *Taxus baccata, T. canadensis,* and *T. cuspidata* (Taxaceae) and Origins of their reputed Hybrids inferred from Rapd and Cpdna Data.**
In: American Journal of Botany 90 (2), p. 175–182.

GHIMIRE, BALKRISHNA; HEO, KWEON (2014):
**Cladistic Analysis of Taxaceae s.l.**
In: Plant Systematics and Evolution 300 (2), p. 217–223. DOI: 10.1007/s00606-013-0874-y.

LI, JIAN-HUA; DAVIS, CHARLES C.; TREDICI, PETER DEL; DONOGHUE, MICHAEL J. (2001):
**Phylogeny and Biogeography of *Taxus* (Taxaceae) inferred from Sequences of the Internal Transcribed Spacer Region of nuclear ribosomal DNA.**
In: Harvard Papers in Botany 6 (1), p. 267–274.

### Tectariaceae

DING, HUI-HUI; CHAO, YI-SHAN; CALLADO, JOHN REY; DONG, SHI-YONG (2014):
**Phylogeny and Character Evolution of the Fern Genus *Tectaria* (Tectariaceae) in the Old World inferred from Chloroplast DNA Sequences.**
In: Molecular Phylogenetics and Evolution 80, p. 66–78. DOI: 10.1016/j.ympev.2014.06.004.

LIU, HONG-MEI; HE, LI-JUAN; SCHNEIDER, HARALD (2014):
**Towards the natural Classification of Tectarioid Ferns: Confirming the phylogenetic Relationships of *Pleocnemia* and *Pteridrys* (Eupolypods I).**
In: Journal of Systematics and Evolution 52 (2), p. 161–174. DOI: 10.1111/jse.12073.

ZHANG, LIANG; SCHUETTPELZ, ERIC; ROTHFELS, CARL J.; ZHOU, XIN-MAO; GAO, XIN-FEN; ZHANG, LI-BING (2016):
**Circumscription and Phylogeny of the Fern Family Tectariaceae based on Plastid and nuclear Markers, with the Description of two new Genera: *Draconopteris* and *Malaifilix* (Tectariaceae).**
In: Taxon 65 (4), p. 723–738. DOI: 10.12705/654.3.

ZHANG, LIANG; ZHOU, XIN-MAO; CHEN, DE-KUI; SCHUETTPELZ, ERIC; KNAPP, RALF; LU, NGAN THI ET AL. (2017):
**A global Phylogeny of the Fern Genus *Tectaria* (Tectariaceae: Polypodiales) based on Plastid and nuclear Markers Identifies major evolutionary Lineages and suggests repeated Evolution of free Venation from Anastomosing Venation.**
In: Molecular Phylogenetics and Evolution 114, p. 295–333. DOI: 10.1016/j.ympev.2017.05.020.

### Ternstroemiaceae

**Su, Yingjuan; Liao, Wen-bo; Wang, Ting; Sun, Yufei; Wei, Qiang; Chang, Hungta (2011):**
**Phylogeny and evolutionary Divergence times in *Apterosperma* and *Euryodendron*: Evidence of a Tertiary Origin in South China.**
In: Biochemical Systematics and Ecology 39 (4-6), p. 769–777. DOI: 10.1016/j.bse.2011.07.004.

**Tsou, Chi-Hua; Li, Lu; Vijayan, Kunjupillai (2016):**
**The Intra-familial Relationships of Pentaphylacaceae s.l. as revealed by DNA Sequence Analysis.**
In: Biochemical Genetics 54 (3), p. 270–282. DOI: 10.1007/s10528-016-9717-1.

**Wu, Chi-Chih; Hsu, Zhi-Fu; Tsou, Chi-Hua (2007):**
**Phylogeny and Taxonomy of *Eurya* (Ternstroemiaceae) from Taiwan, as inferred from ITS Sequence Data.**
In: Botanical Studies 48, p. 97–116.

### Theaceae

**Heo, Kyeong-In; Lee, Sangtae; Lee, Chunghee; Kim, Seung-Chul (2011):**
**Generic Delimitation and infrageneric Classification of *Stewartia* and *Hartia* (Theaceae; Stewartieae): Insight from Pollen Morphology.**
In: Plant Systematics and Evolution 297 (1-2), p. 33–50. DOI: 10.1007/s00606-011-0497-0.

**Li, Mi-Mi; Li, Jian-Hua; Del Tredici, Peter; Corajod, Jeffrey; Fu, Cheng-Xin (2013):**
**Phylogenetics and Biogeography of Theaceae based on Sequences of Plastid Genes.**
In: Journal of Systematics and Evolution 51 (4), p. 396–404. DOI: 10.1111/jse.12017.

**Li, Rong; Yang, Jun-Bo; Yang, Shi-Xiong; Li, De-Zhu (2011):**
**Phylogeny and Taxonomy of the *Pyrenaria* complex (Theaceae) based on nuclear ribosomal ITS Sequences.**
In: Nordic Journal of Botany 29 (6), p. 780–787. DOI: 10.1111/j.1756-1051.2011.01175.x.

**Luna, Isolda; Ochoterena, Helga (2004):**
**Phylogenetic Relationships of the Genera of Theaceae based on Morphology.**
In: Cladistics 20 (3), p. 223–270. DOI: 10.1111/j.1096-0031.2004.00024.x.

**Prince, Linda M.; Parks, Clifford R. (2001):**
**Phylogenetic Relationships of Theaceae inferred from Chloroplast DNA Sequence Data.**
In: American Journal of Botany 88 (12), p. 2309–2320. DOI: 10.2307/3558391.

**Vijayan, Kunjupillai; Zhang, Wen-Ju; Tsou, Chi-Hua (2009):**
**Molecular Taxonomy of *Camellia* (Theaceae) inferred from nrITS Sequences.**
In: American Journal of Botany 96 (7), p. 1348–1360. DOI: 10.3732/ajb.0800205.

**Wang, Y. H.; He, H.; Min, T. L.; Zhou, L. H.; Fritsch, Peter W. (2006):**
**The phylogenetic Position of *Apterosperma* (Theaceae) based on morphological and karyotype Characters.**
In: Plant Systematics and Evolution 260 (1), p. 39–52. DOI: 10.1007/s00606-006-0426-9.

**Yang, S.-X.; Yang, J.-B.; Lei, Li-Gong; Li, D.-Z.; Yoshino, H.; Ikeda, T. (2004):**
**Reassessing the Relationships between *Gordonia* and *Polyspora* (Theaceae) based on the combined Analyses of molecular Data from the nuclear, plastid and mitochondrial Genomes.**
In: Plant Systematics and Evolution 248 (1-4), p. 45–55. DOI: 10.1007/s00606-004-0178-3.

YU, XIANG-QIN; DREW, BRYAN T.; YANG, JUN-BO; GAO, LIAN-MING; LI, DE-ZHU (2017):
**Comparative Chloroplast Genomes of eleven *Schima* (Theaceae) species: Insights into DNA barcoding and Phylogeny.**
In: Public Library of Science One 12 (6), e0178026. DOI: 10.1371/journal.pone.0178026.

### Thecophilaeaceae

BRUMMITT, R.K.; BANKS, HANNAH I.; JOHNSON, MARGARET A.T.; DOCHERTY, KATHERINE A.; JONES, KEITH; CHASE, MARK W.; RUDALL, PAULA J. (1998):
**Taxonomy of Cyanastroideae (Tecophilaeaceae): a multidisciplinary Approach.**
In: Kew Bulletin 53 (4), p. 769–803.

MANNING, JOHN C.; FOREST, FÉLIX; MANNHEIMER, C. A. (2005):
***Eremiolirion*, a new Genus of southern African Tecophilaeaceae, and taxonomic Notes on *Cyanella alba*.**
In: Bothalia 35 (2), p. 115–120. DOI: 10.4102/abc.v35i2.386.

MANNING, JOHN C.; GOLDBLATT, PETER (2012):
**A Revision of Tecophilaeaceae Subfam. Tecophilaeoideae in Africa.**
In: Bothalia 42 (1), p. 21–41. DOI: 10.4102/abc.v42i1.6.

### Thelypteridaceae

ALMEIDA, THAÍS ELIAS; HENNEQUIN, SABINE; SCHNEIDER, HARALD; SMITH, ALAN R.; BATISTA, JOÃO AGUIAR NOGUEIRA; RAMALHO, ALINE JOSEPH ET AL. (2016):
**Towards a phylogenetic generic Classification of Thelypteridaceae: Additional Sampling suggests alterations of Neotropical Taxa and further Study of paleotropical Genera.**
In: Molecular Phylogenetics and Evolution 94 (Pt B), p. 688–700. DOI: 10.1016/j.ympev.2015.09.009.

FERNANDES, ROZIJANE SANTOS; YESILYURT, JOVITA CISLINSKI; SALINO, ALEXANDRE (2014):
**New Species and Combinations in *Meniscium* (Thelypteridaceae).**
In: Phytotaxa 184 (1), p. 1. DOI: 10.11646/phytotaxa.184.1.1.

HE, LI-JUAN; ZHANG, XIAN-CHUN (2012):
**Exploring generic Delimitation within the Fern Family Thelypteridaceae.**
In: Molecular Phylogenetics and Evolution 65 (2), p. 757–764. DOI: 10.1016/j.ympev.2012.07.021.

LI, ZHONG-YANG; HE, ZHAO-RONG; ZHANG, XIAN-CHUN (2013):
**A taxonomic Revision of *Cyclosorus* Subgenus *Cyclosoriopsis* (Thelypteridaceae) from China.**
In: Journal of Systematics and Evolution 51 (5), p. 609–638. DOI: 10.1111/jse.12013.

SMITH, ALAN R.; CRANFILL, RAYMOND B. (2002):
**Intrafamilial Relationships of the Thelypteroid Ferns (Thelypteridaceae).**
In: American Fern Journal 92 (2), p. 131. DOI: 10.1640/0002-8444(2002)092[0131:IROTTF]2.0.CO;2.

WANG, REN-XIANG; SHAO, WEN; LIU, LING; LIU, JING; DENG, XI-CHAO; LU, SHU-GANG (2015):
**A Systematic Study of the Fern Genus *Mesopteris* Ching (Thelypteridaceae).**
In: American Fern Journal 105 (1), p. 11–19. DOI: 10.1640/0002-8444-105.1.11.

### Themidaceae

PIRES, J. CHRIS; FAY, MICHAEL F.; DAVIS, WARREN S.; HUFFORD, LARRY; ROVA, JOHAN H.E.; CHASE, MARK W.; SYTSMA, KENNETH J. (2001):
**Molecular and morphological phylogenetic Analyses of Themidaceae (Asparagales).**
In: Kew Bulletin 56, p. 601–626.

**Pires, J. Chris; Sytsma, Kenneth J. (2002):**

**A phylogenetic Evaluation of a biosystematic Famework: *Brodiaea* and related petaloid Monocots (Themidaceae).**

In: American Journal of Botany 89 (8), p. 1342–1359. DOI: 10.3732/ajb.89.8.1342.

### Theophrastaceae

**Källersjö, Mari; Ståhl, Bertil (2003):**

**Phylogeny of Theophrastaceae (Ericales s. lat.).**

In: International Journal of Plant Sciences 164 (4), p. 579–591.

**Ståhl, Bertil; Källersjö, Mari (2004):**

**Reinstatement of *Bonellia* (Theophrastaceae).**

In: Novon: A Journal for Botanical Nomenclature 14 (1), p. 115–118.

### Thesiaceae

**Moore, Timothy E.; Verboom, George Anthony; Forest, Félix (2010):**

**Phylogenetics and Biogeography of the parasitic Genus *Thesium* L. (Santalaceae), with an Emphasis on the Cape of South Africa.**

In: Botanical Journal of the Linnean Society 162, p. 435–454.

**Nickrent, Daniel Lee; García, Miguel Angel (2015):**

***Lacomucinaea*, a new monotypic Genus in Thesiaceae (Santalales).**

In: Phytotaxa 224 (2), p. 173. DOI: 10.11646/phytotaxa.224.2.4.

### Thismiaceae

**Merckx, Vincent S.F.T.; Bakker, Freek T.; Huysmans, Suzy; Smets, Erik (2009):**

**Bias and conflict in phylogenetic Inference of myco-heterotrophic Plants: a case Study in Thismiaceae.**

In: Cladistics 25 (1), p. 64–77. DOI: 10.1111/j.1096-0031.2008.00241.x.

**Yahara, Tetsukazu; Tsukaya, Hirokazu (2008):**

***Oxygyne yamashitae*, a new Species of Thismiaceae from Yaku Island, Japan.**

In: Acta Phytotaxonomica Geobotanica 59 (2), p. 97–104.

### Thomandersiaceae

**Wortley, Alexandra H.; Harris, David J.; Scotland, Robert W. (2007):**

**On the Taxonomy and phylogenetic Position of *Thomandersia*.**

In: Systematic Botany 32 (2), p. 415–444.

### Thymelaeaceae

**Beaumont, Angela J.; Edwards, Trevor J.; Manning, John C.; Maurin, Olivier; Rautenbach, Marline; Motsi, Moleboheng C. et al. (2009):**

***Gnidia* (Thymelaeaceae) is not monophyletic: taxonomic Implications for Thymelaeoideae and a partial new generic Taxonomy for *Gnidia*.**

In: Botanical Journal of the Linnean Society 160 (4), p. 402–417. DOI: 10.1111/j.1095-8339.2009.00988.x.

**Bredenkamp, C. L.; van Wyk, Abraham E. (2003):**

**Taxonomy of the Genus *Passerina* (Thymelaeaceae).**

In: Bothalia 33 (1), p. 59–98. DOI: 10.4102/abc.v33i1.433.

**Eurlings, Marcel C.M.; Gravendeel, Barbara (2005):**

**TrnL-trnF Sequence Data imply Paraphyly of *Aquilaria* and *Gyrinops* (Thymelaeaceae) and provide new Perspectives for Agarwood Identification.**

In: Plant Systematics and Evolution 254 (1-2), p. 1–12. DOI: 10.1007/s00606-005-0312-x.

**Floden, Aaron J.; Mayfield, Mark H.; Ferguson, Carolyn J. (2009):**
**A new narrowly endemic Species of *Dirca* (Thymelaeaceae) from Kansas and Arkansas, with a phylogenetic Overview and taxonomic Synopsis of the Genus.**
In: Journal of the Botanical Research Institute of Texas 3 (2), p. 485–499.

**Foster, Charles S.P.; Cantrill, David J.; James, Elizabeth A.; Syme, Anna E.; Jordan, Rebecca; Douglas, Rachel et al. (2016):**
**Molecular Phylogenetics provides new Insights into the Systematics of *Pimelea* and *Thecanthes* (Thymelaeaceae).**
In: Australian Systematic Botany 29 (3), p. 185–196. DOI: 10.1071/SB16013.

**Galicia-Herbada, D. (2006):**
**Origin and Diversification of *Thymelaea* (Thymelaeaceae): Inferences from a phylogenetic Study based on ITS (rDNA) Sequences.**
In: Plant Systematics and Evolution 257 (3-4), p. 159–187. DOI: 10.1007/s00606-005-0371-z.

**Rogers, Zachary p. (2009):**
**A Revision of Malagasy *Gnidia* (Thymelaeaceae, Thymelaeoideae).**
In: Annals of the Missouri Botanical Garden 96 (2), p. 324–369. DOI: 10.3417/2006114.

**Rogers, Zachary p. (2010):**
**Nomenclatural Notes on American Thymelaeaceae.**
In: Novon: A Journal for Botanical Nomenclature 20 (4), p. 448–462. DOI: 10.3417/2009069.

**van der Bank, Michelle; Fay, Michael F.; Chase, Mark W. (2002):**
**Molecular Phylogenetics of Thymelaeaceae with particular Reference to African and Australian Genera.**
In: Taxon 51, p. 329–339.

**Zhang, Yong-Hong; Volis, Sergei; Sun, Hang (2010):**
**Chloroplast Phylogeny and Phylogeography of *Stellera chamaejasme* on the Qinghai-Tibet Plateau and in adjacent Regions.**
In: Molecular Phylogenetics and Evolution 57 (3), p. 1162–1172. DOI: 10.1016/j.ympev.2010.08.033.

### Tiliaceae

**Cai, Jie; Ma, Peng-Fei; Li, Hong-Tao; Li, De-Zhu (2015):**
**Complete Plastid Genome Sequencing of four *Tilia* Species (Malvaceae): A comparative Analysis and phylogenetic Implications.**
In: Public Library of Science One 10 (11), e0142705. DOI: 10.1371/journal.pone.0142705.

**Yousefzadeh, Hamed; Hosseinzadeh Colagar, Abasalt; Tabari, Masoud; Sattarian, Ali; Assadi, Mostafa (2012):**
**Utility of ITS Region Sequence and Structure for molecular Identification of *Tilia* Species from Hyrcanian forests, Iran.**
In: Plant Systematics and Evolution 298 (5), p. 947–961. DOI: 10.1007/s00606-012-0604-x.

### Tofieldiaceae

**Azuma, Hiroshi; Tobe, Hiroshi (2011):**
**Molecular phylogenetic Analyses of Tofieldiaceae (Alismatales): Family Circumscription and intergeneric Relationships.**
In: Journal of Plant Research 124 (3), p. 349–357. DOI: 10.1007/s10265-010-0387-5.

**Campbell, Lisa M.; Dorr, Laurence J. (2013):**
**A Synopsis of *Harperocallis* (Tofieldiaceae, Alismatales) with ten new Combinations.**

In: PhytoKeys (21), p. 37–52. DOI: 10.3897/phytokeys.21.4859.

**Remizowa, Margarita V.; Sokoloff, Dmitry D.; Campbell, Lisa M.; Stevenson, Dennis Wm.; Rudall, Paula J. (2011):**
**_Harperocallis_ is congeneric with _Isidrogalvia_ (Tofieldiaceae, Alismatales): Evidence from comparative Floral Morphology.**
In: Taxon 60 (4), p. 1076–1094.

**Tamura, Minoru N.; Azuma, Hiroshi; Yamashita, Jun; Fuse, Shizuka; Ishii, Takaaki (2010):**
**Biosystematic studies on the Family Tofieldiaceae II. Phylogeny of Species of _Tofieldia_ and _Triantha_ inferred from Plastid and Nuclear DNA Sequences.**
In: Acta Phytotaxonomica Geobotanica 60 (3), p. 131–140.

**Tamura, Minoru N.; Fuse, Shizuka; Azuma, H.; Hasebe, M. (2004):**
**Biosystematic Studies on the Family Tofieldiaceae I. Phylogeny and Circumscription of the Family inferred from DNA Sequences of matK and rbcL.**
In: Plant Biology 6 (5), p. 562–567. DOI: 10.1055/s-2004-821278.

**Tamura, Minoru N.; Fuse, Shizuka; Lee, Nam Sook; Kim, Jin Ohk; Yamashita, Jun; Ishii, Takaaki (2011):**
**Biosystematic studies on the Family Tofieldiaceae III. Classification of _Tofieldia nuda_ into three Species and three Varieties.**
In: Taxon 60 (5), p. 1339–1348.

### Tracheophyta

**Cantino, Philip D.; Doyle, James A.; Graham, Sean W.; Judd, Walter S.; Olmstead, Richard G.; Soltis, Douglas E. et al. (2007):**
**Towards a phylogenetic Nomenclature of Tracheophyta.**
In: Taxon 56 (3), p. 1–44.

### Triuridaceae

**Mennes, Constantijn B.; Smets, Erik F.; Moses, Sainge N.; Merckx, Vincent S.F.T. (2013):**
**New Insights in the long-debated evolutionary History of Triuridaceae (Pandanales).**
In: Molecular Phylogenetics and Evolution 69 (3), p. 994–1004. DOI: 10.1016/j.ympev.2013.05.031.

### Trochodendraceae

**Ren, Yi; Chen, L.; Tian, X. H.; Zhang, X. H.; Lu, A. M. (2007):**
**Discovery of Vessels in _Tetracentron_ (Trochodendraceae) and its systematic Significance.**
In: Plant Systematics and Evolution 267 (1-4), p. 155–161. DOI: 10.1007/s00606-007-0563-9.

### Tropaeolaceae

**Andersson, Lennart; Andersson, Stephan (2000):**
**A molecular Phylogeny of Tropaeolaceae and its Systematic Implications.**
In: Taxon 49 (2), p. 721–736.

**Hershkovitz, Mark A.; Hernández-Pellicer, C. C.; Arroyo, Mary T.K. (2006):**
**Ribosomal DNA Evidence for the Diversification of _Tropaeolum_ Sect. _Chilensia_ (Tropaeolaceae).**
In: Plant Systematics and Evolution 260 (1), p. 1–24. DOI: 10.1007/s00606-006-0428-7.

### Turneraceae

**Arbo, María M.; Gonzalez, Ana M.; Sede, Silvana M. (2015):**

**Phylogenetic Relationships within Turneraceae based on morphological Characters with Emphasis on Seed micromorphology.**
In: Plant Systematics and Evolution 301 (7), p. 1907–1926. DOI: 10.1007/s00606-015-1204-3.

LÓPEZ, ALICIA; FERNÁNDEZ, AVELIANO; SHORE, JOEL P. (2013):
**Inferences on the Origins of polyploid *Turnera* Species (Passifloraceae) based on molecular Data.**
In: Botany 91 (3), p. 167–175. DOI: 10.1139/cjb-2012-0135.

MERCEDES ARBO, MARIA; ESPERT, SHIRLEY M. (2009):
**Morphology, Phylogeny and Biogeography of *Turnera* L. (Turneraceae).**
In: Taxon 58 (2), p. 457–467. DOI: 10.1002/tax.582011.

THULIN, MATS; RAZAFIMANDIMBISON, SYLVAIN G.; CHAFE, PAUL; HEIDARI, NAHID; KOOL, ANNELEEN; SHORE, JOEL P. (2012):
**Phylogeny of the Turneraceae Clade (Passifloraceae s.l.): Trans-Atlantic Disjunctions and two new Genera in Africa.**
In: Taxon 61 (2), p. 308–323. DOI: 10.1002/tax.612003.

TRUYENS, SIMON; ARBO, MARÍA M.; SHORE, JOEL P. (2005):
**Phylogenetic Relationships, Chromosome and breeding System Evolution in *Turnera* (Turneraceae): Inferences from its Sequence Data.**
In: American Journal of Botany 92 (10), p. 1749–1758. DOI: 10.3732/ajb.92.10.1749.

### Typhaceae

ITO, YU; TANAKA, NORIO; KIM, CHANGKYUN; KAUL, ROBERT B.; ALBACH, DIRK C. (2016):
**Phylogeny of *Sparganium* (Typhaceae) revisited: non-monophyletic Nature of *S. emersum* sensu lato and Resurrection of *S. acaule*.**
In: Plant Systematics and Evolution 302 (1), p. 129–135. DOI: 10.1007/s00606-015-1245-7.

KIM, CHANGKYUN; CHOI, HONG-KEUN (2011):
**Molecular Systematics and Character Evolution of *Typha* (Typhaceae) inferred from nuclear and Plastid DNA Sequence Data.**
In: Taxon 60 (5), p. 1417–1428.

SULMAN, JOSHUA D.; DREW, BRYAN T.; DRUMMOND, CHLOE; HAYASAKA, EISUKE; SYTSMA, KENNETH J. (2013):
**Systematics, Biogeography, and Character Evolution of *Sparganium* (Typhaceae): Diversification of a widespread, aquatic Lineage.**
In: American Journal of Botany 100 (10), p. 2023–2039. DOI: 10.3732/ajb.1300048.

### Ulmaceae

DENK, THOMAS; GRIMM, GUIDO W. (2005):
**Phylogeny and Biogeography of *Zelkova* (Ulmaceae sensu stricto) as inferred from Leaf Morphology, ITS Sequence Data and the fossil Record.**
In: Botanical Journal of the Linnean Society 147, p. 129–157.

### Urticaceae

GROSSE-VELDMANN, BERNADETTE; NÜRK, NICOLAI M.; SMISSEN, ROB D.O.B.; BREITWIESER, ILSE; QUANDT, DIETMAR; WEIGEND, MAXIMILIAN (2016):
**Pulling the sting out of Nettle Systematics - A comprehensive Phylogeny of the Genus *Urtica* L. (Urticaceae).**
In: Molecular Phylogenetics and Evolution 102, p. 9–19. DOI: 10.1016/j.ympev.2016.05.019.

GROSSE-VELDMANN, BERNADETTE; WEIGEND, MAXIMILIAN (2018):

**The geometry of gender: Hyper-Diversification of sexual systems in *Urtica* L. (Urticaceae).**
In: Cladistics 34 (2), p. 131–150. DOI: 10.1111/cla.12193.

GUTIÉRREZ-VALENCIA, JUANITA; CHOMICKI, GUILLAUME; RENNER, SUSANNE P. (2017):
**Recurrent Breakdowns of Mutualisms with Ants in the Neotropical Ant-plant Genus *Cecropia* (Urticaceae).**
In: Molecular Phylogenetics and Evolution 111, p. 196–205. DOI: 10.1016/j.ympev.2017.04.009.

HADIAH, JULISASI T.; CONN, BARRY J.; QUINN, CHRISTOPHER J. (2008):
**Infra-familial Phylogeny of Urticaceae, using Chloroplast Sequence Data.**
In: Australian Systematic Botany 21 (5), p. 375–385. DOI: 10.1071/SB08041.

HADIAH, JULISASI T.; QUINN, CHRISTOPHER J.; CONN, BARRY J. (2003):
**Phylogeny of *Elatostema* (Urticaceae) using Chloroplast DNA Data.**
In: Telopea 10 (1), p. 235–246.

JESTROW, BRETT; VALDÉS, JAMES J.; RODRÍGUEZ, FRANCISCO JIMÉNEZ; FRANCISCO-ORTEGA, JAVIER (2012):
**Phylogenetic Placement of the Dominican Republic endemic Genus *Sarcopilea* (Urticaceae).**
In: Taxon 61 (3), p. 592–600.

KIM, CHANGKYUN; DENG, TAO; CHASE, MARK W.; ZHANG, DAI-GUI; NIE, ZE-LONG; SUN, HANG (2015):
**Generic Phylogeny and Character Evolution in Urticeae (Urticaceae) inferred from nuclear and Plastid DNA Regions.**
In: Taxon 64 (1), p. 65–78. DOI: 10.12705/641.20.

MONRO, ALEX K. (2006):
**The Revision of Species-rich Genera: A phylogenetic Framework for the Strategic Revision of *Pilea* (Urticaceae) based on cpDNA, nrDNA, and Morphology.**
In: American Journal of Botany 93 (3), p. 426–441.

TREIBER, ERIN L.; GAGLIOTI, ANDRÉ LUIZ; ROMANIUC-NETO, SERGIO; MADRIÑÁN, SANTIAGO; WEIBLEN, GEORGE D. (2016):
**Phylogeny of the Cecropieae (Urticaceae) and the Evolution of an Ant-Plant Mutualism.**
In: Systematic Botany 41 (1), p. 56–66. DOI: 10.1600/036364416X690633.

WU, ZENG-YUAN; MONRO, ALEX K.; MILNE, RICHARD IAN; WANG, HONG; YI, TING-SHUANG; LIU, JIE; LI, DE-ZHU (2013):
**Molecular Phylogeny of the nettle Family (Urticaceae) inferred from multiple Loci of three Genomes and extensive generic Sampling.**
In: Molecular Phylogenetics and Evolution 69 (3), p. 814–827. DOI: 10.1016/j.ympev.2013.06.022.

### Valerianaceae

BELL, C. (2005):
**Phylogeny and Biogeography of Valerianaceae (Dipsacales) with special Reference to the South American Valerians.**
In: Organisms Diversity and Evolution 5 (2), p. 147–159. DOI: 10.1016/j.ode.2004.10.014.

BELL, CHARLES D. (2004):
**Preliminary Phylogeny of Valerianaceae (Dipsacales) inferred from nuclear and Chloroplast DNA Sequence Data.**
In: Molecular Phylogenetics and Evolution 31 (1), p. 340–350. DOI: 10.1016/j.ympev.2003.07.006.

**Bell, Charles D. (2007):**
**Phylogenetic Placement and Biogeography of the North American Species of *Valerianella* (Valerianaceae: Dipsacales) based on Chloroplast and nuclear DNA.**
In: Molecular Phylogenetics and Evolution 44 (3), p. 929–941. DOI: 10.1016/j.ympev.2007.03.013.

**Bell, Charles D.; Gonzalez, Lauren A. (2018):**
**Exploring the utility of "Next-generation" Sequence Data on Inferring the Phylogeny of the South American *Valeriana* (Valerianaceae).**
In: Molecular Phylogenetics and Evolution 123, p. 44–49. DOI: 10.1016/j.ympev.2018.02.014.

**Bell, Charles D.; Kutschker, Adriana; Arroyo, Mary T.K. (2012):**
**Phylogeny and Diversification of Valerianaceae (Dipsacales) in the Southern Andes.**
In: Molecular Phylogenetics and Evolution 63 (3), p. 724–737. DOI: 10.1016/j.ympev.2012.02.015.

**Hidalgo, Oriane; Garnatje, Teresa; Susanna, Alfonso; Mathez, Joel (2004):**
**Phylogeny of Valerianaceae based on matK and ITS Markers, with Reference to matK individual Polymorphism.**
In: Annals of Botany 93 (3), p. 283–293. DOI: 10.1093/aob/mch042.

**Jacobs, Bart; Bell, Charles D.; Smets, Erik (2010):**
**Fruits and Seeds of the *Valeriana* Clade (Dipsacales): Diversity and Evolution.**
In: International Journal of Plant Sciences 171 (4), p. 421–434. DOI: 10.1086/651243.

**Pyck, Nancy; van Lysebetten, A.; Stessens, J.; Smets, Erik (2002):**
**The Phylogeny of Patrinieae sensu Graebner (Valerianaceae) revisited: additional Evidence from ndh F Sequence Data.**
In: Plant Systematics and Evolution 233 (1-2), p. 29–46. DOI: 10.1007/s006060200053.

### Velloziaceae

**Behnke, H.-Dietmar; Hummel, Eric; Hillmer, Stefan; Sauer-Gürth, Hedwig; Gonzalez, Javier; Wink, Michael (2013):**
**A Revision of African Velloziaceae based on Leaf Anatomy Characters and rbcL Nucleotide Sequences.**
In: Botanical Journal of the Linnean Society 172 (1), p. 22–94. DOI: 10.1111/boj.12018.

**Mello-Silva, Renato (2005):**
**Morphological Analysis, Phylogenies and Classification in Velloziaceae.**
In: Botanical Journal of the Linnean Society 148 (2), p. 157–173. DOI: 10.1111/j.1095-8339.2005.00399.x.

**Mello-Silva, Renato; Montserrat, Laura (2015):**
**Depicting *Barbacenia flava* and *Vellozia intermedia* with a short History of illustrated Velloziaceae.**
In: Kew Bulletin 70 (2), p. 22. DOI: 10.1007/S12225-015-9575-8.

**Menezes, Nanuza L.; Mello-Silva, Renato; Mayo, Simon J. (1993):**
**A Cladistic Analysis of the Velloziaceae.**
In: Kew Bulletin 49 (1), p. 71–92. DOI: 10.2307/4110200.

### Verbenaceae

**Lu-Irving, Patricia; O'Leary, Nataly; O'Brien, Anna; Olmstead, Richard G. (2014):**
**Resolving the Genera *Aloysia* and *Acantholippia* within Tribe Lantaneae (Verbenaceae), using Chloroplast and Nuclear Sequences.**
In: Systematic Botany 39 (2), p. 644–655. DOI: 10.1600/036364414X680816.

**Lu-Irving, Patricia; Olmstead, Richard G. (2013):**

**Investigating the Evolution of Lantaneae (Verbenaceae) using multiple Loci.**
In: Botanical Journal of the Linnean Society 171, p. 103–119.

MARX, HANNAH E.; O'LEARY, NATALY; YUAN, YAO-WU; LU-IRVING, PATRICIA; TANK, DAVID C.; MÚLGURA, MARÍA EMA; OLMSTEAD, RICHARD G. (2010):
**A molecular Phylogeny and Classification of Verbenaceae.**
In: American Journal of Botany 97 (10), p. 1647–1663. DOI: 10.3732/ajb.1000144.

NESOM, GUY L. (2010):
**Infrageneric Classification of *Verbena* (Verbenaceae).**
In: Phytoneuron 11, p. 1–15.

O'LEARY, NATALY; CALVIÑO, CAROLINA ISABEL; MARTÍNEZ, SUSANA G.; LU-IRVING, PATRICIA; OLMSTEAD, RICHARD G.; MÚLGURA, MARÍA EMA (2012):
**Evolution of morphological Traits in Verbenaceae.**
In: American Journal of Botany 99 (11), p. 1778–1792. DOI: 10.3732/ajb.1200123.

O'LEARY, NATALY; YUAN, YAO-WU; CHEMISQUY, MARIA AMELIA; OLMSTEAD, RICHARD G. (2009):
**Reassignment of Species of paraphyletic *Junellia* s.l. to the new Genus *Mulguraea* (Verbenaceae) and new Circumscription of Genus *Junellia* : molecular and morphological Congruence.**
In: Systematic Botany 34 (4), p. 777–786. DOI: 10.1600/036364409790139691.

PERALTA, PAOLA; MÚLGURA DE ROMERO, MARÍA E.; DENHAM, SILVIA S.; BOTTA, SILVIA M. (2008):
**Revisión del Género *Junellia* (Verbenaceae).**
In: Annals of the Missouri Botanical Garden 95 (2), p. 338–390. DOI: 10.3417/2004167.

YUAN, YAO-WU; LIU, CHANG; MARX, HANNAH E.; OLMSTEAD, RICHARD G. (2010):
**An empirical Demonstration of using Pentatricopeptide Repeat (PPR) Genes as plant phylogenetic tools: Phylogeny of Verbenaceae and the *Verbena*-complex.**
In: Molecular Phylogenetics and Evolution 54 (1), p. 23–35. DOI: 10.1016/j.ympev.2009.08.029.

YUAN, YAO-WU; OLMSTEAD, RICHARD G. (2008):
**A species-level phylogenetic Study of the *Verbena* complex (Verbenaceae) indicates two independent intergeneric Chloroplast Transfers.**
In: Molecular Phylogenetics and Evolution 48 (1), p. 23–33. DOI: 10.1016/j.ympev.2008.04.004.

YUAN, YAO-WU; OLMSTEAD, RICHARD G. (2008):
**Evolution and phylogenetic utility of the Phot Gene duplicates in the *Verbena* complex (Verbenaceae): dramatic Intron size Variation and Footprint of ancestral Recombination.**
In: American Journal of Botany 95 (9), p. 1166–1176. DOI: 10.3732/ajb.0800133.

### Viburnaceae

CHATELET, DAVID S.; CLEMENT, WENDY L.; SACK, LAWREN; DONOGHUE, MICHAEL J.; EDWARDS, ERIKA J. (2013):
**The Evolution of Photosynthetic Anatomy in *Viburnum* (Adoxaceae).**
In: International Journal of Plant Sciences 174 (9), p. 1277–1291. DOI: 10.1086/673241.

CLEMENT, WENDY L.; ARAKAKI, MÓNICA; SWEENEY, PATRICK W.; EDWARDS, ERIKA J.; DONOGHUE, MICHAEL J. (2014):
**A Chloroplast Tree for *Viburnum* (Adoxaceae) and its Implications for phylogenetic Classification and Character Evolution.**
In: American Journal of Botany 101 (6), p. 1029–1049. DOI: 10.3732/ajb.1400015.

CLEMENT, WENDY L.; DONOGHUE, MICHAEL J. (2011):

**Dissolution of *Viburnum* Section *Megalotinus* (Adoxaceae) of Southeast Asia and its Implications for morphological Evolution and Biogeography.**
In: International Journal of Plant Sciences 172 (4), p. 559–573. DOI: 10.1086/658927.

CLEMENT, WENDY L.; DONOGHUE, MICHAEL J. (2012):
**Barcoding Success as a function of phylogenetic Relatedness in *Viburnum*, a Clade of woody Angiosperms.**
In: BMC Evolutionary Biology 12, p. 73. DOI: 10.1186/1471-2148-12-73.

SPRIGGS, ELIZABETH L.; CLEMENT, WENDY L.; SWEENEY, PATRICK W.; MADRIÑÁN, SANTIAGO; EDWARDS, ERIKA J.; DONOGHUE, MICHAEL J. (2015):
**Temperate Radiations and dying embers of a tropical past: the Diversification of *Viburnum*.**
In: the new Phytologist 207 (2), p. 340–354. DOI: 10.1111/nph.13305.

WINKWORTH, RICHARD C.; DONOGHUE, MICHAEL J. (2004):
***Viburnum* Phylogeny: Evidence from the duplicated nuclear Gene GBSSI.**
In: Molecular Phylogenetics and Evolution 33 (1), p. 109–126. DOI: 10.1016/j.ympev.2004.05.006.

WINKWORTH, RICHARD C.; DONOGHUE, MICHAEL J. (2005):
***Viburnum* Phylogeny based on combined molecular Data: Implications for Taxonomy and Biogeography.**
In: American Journal of Botany 92 (4), p. 653–666. DOI: 10.3732/ajb.92.4.653.

### Violaceae

BALLARD, HARVEY EUGENE; SYTSMA, KENNETH J. (2000):
**Evolution and Biogeography of the Woody Hawaiian Violets (*Viola*, Violaceae): Arctic Origins, herbaceous Ancestry and Bird Dispersal.**
In: Evolution 54 (5), p. 1521–1532. DOI: 10.1554/0014-3820(2000)054[1521:EABOTW]2.0.CO;2.

CONESA, M. À.; MUS, M.; ROSSELLÓ, J. A. (2008):
**Hybridization between insular endemic and widespread Species of *Viola* in non-disturbed Environments assessed by nuclear ribosomal and cpDNA Sequences.**
In: Plant Systematics and Evolution 273 (3-4), p. 169–177. DOI: 10.1007/s00606-008-0006-2.

HAVRAN, J. CHRISTOPHER; SYTSMA, KENNETH J.; BALLARD, HARVEY EUGENE (2009):
**Evolutionary Relationships, inter-Island Biogeography, and molecular Evolution in the Hawaiian violets (*Viola*: Violaceae).**
In: American Journal of Botany 96 (11), p. 2087–2099. DOI: 10.3732/ajb.0900021.

LIANGGUAN-XIN; XINGFU-WU (2010):
**Infrageneric Phylogeny of the Genus *Viola* (Violaceae) based on trnL trnF, psbA-trnH, rpL16,ITS Sequences, cytological and morphological Data.**
In: Acta Botanica Yunnanica 32 (6), p. 477–488.

MARCUSSEN, THOMAS; HEIER, LISE; BRYSTING, ANNE K.; OXELMAN, BENGT; JAKOBSEN, KJETILL P. (2015):
**From Gene Trees to a dated allopolyploid Network: Insights from the Angiosperm Genus Viola (Violaceae).**
In: Systematic Biology 64 (1), p. 84–101. DOI: 10.1093/sysbio/syu071.

MARCUSSEN, THOMAS; JAKOBSEN, KJETILL S.; DANIHELKA, JIRÍ; BALLARD, HARVEY EUGENE; BLAXLAND, KIM; BRYSTING, ANNE K.; OXELMAN, BENGT (2012):
**Inferring Species Networks from Gene Trees in high-polyploid North American and Hawaiian Violets (*Viola*, Violaceae).**

In: Systematic Biology 61 (1), p. 107–126. DOI: 10.1093/sysbio/syr096.

**Marcussen, Thomas; Oxelman, Bengt; Skog, Anna; Jakobsen, Kjetill p. (2010):**
**Evolution of plant RNA Polymerase IV/V genes: Evidence of Subneofunctionalization of duplicated NRPD2/NRPE2-like paralogs in *Viola* (Violaceae).**
In: BMC Evolutionary Biology 10, p. 45. DOI: 10.1186/1471-2148-10-45.

**Mitchell, Anthony D.; Heenan, Peter B.; Murray, B. G.; Molloy, B. P. J.; Lange, Peter J. (2009):**
**Evolution of the south-western Pacific Genus *Melicytus* (Violaceae): Evidence from DNA Sequence Data, Cytology and Sex Expression.**
In: Australian Systematic Botany 22 (3), p. 143–157. DOI: 10.1071/SB08042.

**Nakamura, Koh; Denda, Tetsuo; Kokubugata, Goro; Huang, Chiun-Jr; Peng, Ching-I.; Yokota, Masatsugu (2015):**
**Phylogeny and Biogeography of the *Viola iwagawae-tashiroi* Species complex (Violaceae, Section *Plagiostigma*) endemic to the Ryukyu Archipelago, Japan.**
In: Plant Systematics and Evolution 301 (1), p. 337–351. DOI: 10.1007/s00606-014-1076-y.

**Paula-Souza, Juliana; Ballard, Harvey Eugene (2014):**
**Re-establishment of the name *Pombalia*, and new Combinations from the polyphyletic *Hybanthus* (Violaceae).**
In: Phytotaxa 183 (1), p. 1. DOI: 10.11646/phytotaxa.183.1.1.

**Stajsic, V.; Walsh, Neville G.; Douglas, R.; Messina, Andre; Molloy, B. P. J. (2014):**
**A Revision of *Melicytus* (Violaceae) in mainland Australia and Tasmania.**
In: Australian Systematic Botany 27 (4), p. 305–323. DOI: 10.1071/SB14022.

**Tokuoka, Toru (2008):**
**Molecular phylogenetic Analysis of Violaceae (Malpighiales) based on Plastid and nuclear DNA Sequences.**
In: Journal of Plant Research 121 (3), p. 253–260. DOI: 10.1007/s10265-008-0153-0.

**Wahlert, Gregory A.; Marcussen, Thomas; Paula-Souza, Juliana; Feng, Min; Ballard, Harvey Eugene (2014):**
**A Phylogeny of the Violaceae (Malpighiales) inferred from Plastid DNA Sequences: Implications for Generic Diversity and Intrafamilial Classification.**
In: Systematic Botany 39 (1), p. 239–252. DOI: 10.1600/036364414X678008.

**Yoo, Ki-Oug; Jang, Su-Kil (2010):**
**Infrageneric Relationships of Korean *Viola* based on eight Chloroplast Markers.**
In: Journal of Systematics and Evolution 48 (6), p. 474–481. DOI: 10.1111/j.1759-6831.2010.00102.x.

### Viscaceae

**Molvray, Mia; Kores, Paul J.; Chase, Mark W. (1999):**
**Phylogenetic Relationships within *Korthalsella* (Viscaceae) based on Nuclear ITS and Plastid Trnl-f Sequence Data.**
In: American Journal of Botany 86 (2), p. 249–260.

**Nickrent, Daniel Lee; García, Miguel Á.; Martín, Mariá P.; Mathiasen, Robert L. (2004):**
**A Phylogeny of all Species of *Arceuthobium* (Viscaceae) using Nuclear and Chloroplast DNA Sequences.**
In: American Journal of Botany 91 (1), p. 125–138.

### Vitaceae

**Chen, Ping-Ting; Chen, Long-Qing; Wen, Jun (2011):**

**The first phylogenetic Analysis of *Tetrastigma* (Miq.) Planch., the host of Rafflesiaceae.**
In: Taxon 60 (2), p. 499–512.

INGROUILLE, MARTIN J.; CHASE, MARK W.; FAY, MICHAEL F.; BOWMAN, DIANE; VAN DER BANK, MICHELLE; BRUIJN, ANETTE D.E. (2002):
**Systematics of Vitaceae from the Viewpoint of Plastid rbcL DNA Sequence Data.**
In: Botanical Journal of the Linnean Society 138 (421-432).

JANSEN, ROBERT K.; KAITTANIS, CHARALAMBOS; SASKI, CHRISTOPHER; LEE, SEUNG-BUM; TOMKINS, JEFFREY; ALVERSON, ANDREW J.; DANIELL, HENRY (2006):
**Phylogenetic Analyses of *Vitis* (Vitaceae) based on complete Chloroplast Genome Sequences: effects of Taxon Sampling and phylogenetic methods on resolving Relationships among Rosids.**
In: BMC Evolutionary Biology 6, p. 32. DOI: 10.1186/1471-2148-6-32.

LIU, XIU-QUN; ICKERT-BOND, STEFANIE M.; CHEN, LONG-QING; WEN, JUN (2013):
**Molecular Phylogeny of *Cissus* L. of Vitaceae (the grape Family) and Evolution of its pantropical intercontinental Disjunctions.**
In: Molecular Phylogenetics and Evolution 66 (1), p. 43–53. DOI: 10.1016/j.ympev.2012.09.003.

LU, LIMIN; COX, CYMON J.; MATHEWS, SARAH; WANG, WEI; WEN, JUN; CHEN, ZHI-DUAN (2018):
**Optimal Data Partitioning, multispecies Coalescent and Bayesian Concordance Analyses resolve early Divergences of the grape Family (Vitaceae).**
In: Cladistics 34 (1), p. 57–77. DOI: 10.1111/cla.12191.

LU, LIMIN; WANG, WEI; CHEN, ZHI-DUAN; WEN, JUN (2013):
**Phylogeny of the non-monophyletic *Cayratia* Juss. (Vitaceae) and Implications for Character Evolution and Biogeography.**
In: Molecular Phylogenetics and Evolution 68 (3), p. 502–515. DOI: 10.1016/j.ympev.2013.04.023.

LU, LIMIN; WEN, JUN; CHEN, ZHI-DUAN (2012):
**A combined morphological and molecular phylogenetic Analysis of *Parthenocissus* (Vitaceae) and taxonomic Implications.**
In: Botanical Journal of the Linnean Society 168, p. 43–63.

MOLINA, JEANMAIRE E.; WEN, JUN; STRUWE, LENA (2013):
**Systematics and Biogeography of the non-viny Grape relative *Leea* (Vitaceae).**
In: Botanical Journal of the Linnean Society 171, p. 354–376.

NIE, ZE-LONG; SUN, HANG; CHEN, ZHI-DUAN; MENG, YING; MANCHESTER, STEVEN R.; WEN, JUN (2010):
**Molecular Phylogeny and biogeographic Diversification of *Parthenocissus* (Vitaceae) disjunct between Asia and North America.**
In: American Journal of Botany 97 (8), p. 1342–1353. DOI: 10.3732/ajb.1000085.

NIE, ZE-LONG; SUN, HANG; MANCHESTER, STEVEN R.; MENG, YING; LUKE, QUENTIN; WEN, JUN (2012):
**Evolution of the intercontinental disjunctions in six Continents in the *Ampelopsis* Clade of the Grape Family (Vitaceae).**
In: BMC Evolutionary Biology 12, p. 17. DOI: 10.1186/1471-2148-12-17.

PELSER, PIETER B.; NICKRENT, DANIEL LEE; BARCELONA, JULIE F. (2016):
**Untangling a Vine and its Parasite: Host specificity of Philippine *Rafflesia* (Rafflesiaceae).**
In: Taxon 65 (4), p. 739–758. DOI: 10.12705/654.4.

Ren, Hui; Lu, Li-Min; Soejima, Akiko; Luke, Quentin; Zhang, Dian-Xiang; Chen, Zhi-Duan; Wen, Jun (2011):
**Phylogenetic Analysis of the grape Family (Vitaceae) based on the noncoding Plastid trnC-petN, trnH-psbA, and trnL-F Sequences.**
In: Taxon 60 (3), p. 629–637.

Rodrigues, Jacqueline Gomes; Lombardi, Julio Antonio; Lovato, Maria Bernadete (2014):
**Phylogeny of *Cissus* (Vitaceae) focusing on South American Species.**
In: Taxon 63 (2), p. 287–298. DOI: 10.12705/632.33.

Rossetto, Maurizio; Crayn, Darren M.; Jackes, Betsy R.; Porter, Carolyn (2007):
**An updated estimate of intergeneric phylogenetic Relationships in the Australian Vitaceae.**
In: Canadian Journal of Botany 85 (8), p. 722–730. DOI: 10.1139/B07-022.

Soejima, Akiko; Wen, Jun (2006):
**Phylogenetic Analysis of the Grape Family (Vitaceae) based on three Chloroplast Markers.**
In: American Journal of Botany 93 (2), p. 278–287.

Trias-Blasi, Anna; Chayamarit, Kongkanda; Teerawatananon, Atchara; Parnell, John A.N. (2014):
**A taxonomic Revision of *Pterisanthes* (Vitaceae) in Thailand and a new Thai Record for *Pterisanthes cissioides*.**
In: Phytotaxa 159 (2), p. 95. DOI: 10.11646/phytotaxa.159.2.3.

Tröndle, Dorothee; Schröder, Stephan; Kassemeyer, Hanns-Heinz; Kiefer, Christiane; Koch, Marcus A.; Nick, Peter (2010):
**Molecular Phylogeny of the Genus *Vitis* (Vitaceae) based on Plastid Markers.**
In: American Journal of Botany 97 (7), p. 1168–1178. DOI: 10.3732/ajb.0900218.

Wan, Yizhen; Schwaninger, Heidi R.; Baldo, Angela M.; Labate, Joanne A.; Zhong, Gan-Yuan; Simon, Charles J. (2013):
**A phylogenetic Analysis of the grape Genus (*Vitis* L.) reveals broad Reticulation and concurrent Diversification during Neogene and Quaternary Climate Change.**
In: BMC Evolutionary Biology 13, p. 141. DOI: 10.1186/1471-2148-13-141.

Wen, Jun; Nie, Ze-long; Soejima, Akiko; Meng, Ying (2007):
**Phylogeny of Vitaceae based on the nuclear Gai1 Gene Sequences.**
In: Canadian Journal of Botany 85 (8), p. 731–745. DOI: 10.1139/B07-071.

Zecca, Giovanni; Abbott, Richard J.; Sun, Wei-Bang; Spada, Alberto; Sala, Francesco; Grassi, Fabrizio (2012):
**The timing and the Mode of Evolution of wild Grapes (Vitis).**
In: Molecular Phylogenetics and Evolution 62 (2), p. 736–747. DOI: 10.1016/j.ympev.2011.11.015.

Zhang, Ning; Wen, Jun; Zimmer, Elizabeth A. (2015):
**Expression Patterns of AP1, FUL, FT and leafy Orthologs in Vitaceae support the Homology of Tendrils and Inflorescences throughout the Grape Family.**
In: Journal of Systematics and Evolution 53 (5), p. 469–476. DOI: 10.1111/jse.12138.

Zhang, Ning; Wen, Jun; Zimmer, Elizabeth A. (2016):
**Another Look at the phylogenetic Position of the Grape Order Vitales: Chloroplast phylogenomics with an expanded Sampling of Key Lineages.**
In: Molecular Phylogenetics and Evolution 101, p. 216–223. DOI: 10.1016/j.ympev.2016.04.034.

### Vochysiaceae

**Sytsma, Kenneth J.; Litt, Amy; Zjhra, Michelle L.; Chris Pires, J.; Nepokroeff, Molly; Conti, Elena et al. (2004):**
**Clades, Clocks, and Continents: Historical and Biogeographical Analysis of Myrtaceae, Vochysiaceae, and Relatives in the Southern Hemisphere.**
In: International Journal of Plant Sciences 165 (S4), S85-S105. DOI: 10.1086/421066.

### Winteraceae

**Marquínez, Xavier; Lohmann, Lúcia Garces; Salatino, Maria Luiza Faria; Salatino, Antonio; González, Favio (2009):**
**Generic Relationships and dating of Lineages in Winteraceae based on nuclear (ITS) and Plastid (rpS16 and psbA-trnH) Sequence Data.**
In: Molecular Phylogenetics and Evolution 53 (2), p. 435–449. DOI: 10.1016/j.ympev.2009.07.001.

### Woodsiaceae

**Chunxiang, Li; Shugang, Lu; Xiaoyan, Sun; Qun, Yang (2011):**
**Phylogenetic Positions of the Enigmatic Asiatic Fern Genera *Diplaziopsis* and *Rhachidosorus* from Analyses of Four Plastid Genes.**
In: American Fern Journal 101 (3), p. 142–155. DOI: 10.1640/0002-8444-101.3.142.

**Shao, Yi-Zhen; Wei, Ran; Zhang, Xian-Chun; Xiang, Qiao Ping (2015):**
**Molecular Phylogeny of the Cliff Ferns (Woodsiaceae: Polypodiales) with a proposed Infrageneric Classification.**
In: Public Library of Science One 10 (9), e0136318. DOI: 10.1371/journal.pone.0136318.

**Wei, Ran; Zhang, Xian-Chun (2014):**
**Rediscovery of *Cystoathyrium chinense* Ching (Cystopteridaceae): phylogenetic Placement of the critically endangered Fern Species endemic to China.**
In: Journal of Systematics and Evolution 52 (4), p. 450–457. DOI: 10.1111/jse.12075.

### Xyridaceae

**Campbell, Lisa M. (2012):**
**Pollen Morphology of Xyridaceae Systematic (Poales) and its Potential.**
In: the Botanical Review 78 (4), p. 428–439. DOI: 10.1007/s12229-012-9110-7.

**Oriani, Aline; Scatena, Vera Lucia (2014):**
**Ovule, Fruit and Seed development in *Abolboda* (Xyridaceae, Poales): Implications for Taxonomy and Phylogeny.**
In: Botanical Journal of the Linnean Society 175 (1), p. 144–154. DOI: 10.1111/boj.12152.

### Zamiaceae

**González, Dolores; Vovides, Andrew P.; Bárcenas, Cristina (2008):**
**Phylogenetic Relationships of the Neotropical Genus *Dioon* (Cycadales, Zamiaceae) based on Nuclear and Chloroplast DNA Sequence Data.**
In: Systematic Botany 33 (2), p. 229–236.

**Nicolalde-Morejón, Fernando; Vovides, Andrew P.; Stevenson, Dennis Wm. (2009):**
**Taxonomic Revision of *Zamia* in Mega-Mexico.**
In: Brittonia 61 (4), p. 301–335.

**Nolasco-Soto, Janet; González-Astorga, Jorge; Nicolalde-Morejón, Fernando; Vergara-Silva, Francisco; los Monteros, Alejandro Espinosa; Medina-Villarreal, Anwar (2015):**
**Phylogeography and demographic History of *Zamia paucijuga* Wieland (Zamiaceae), a Cycad Species from the Mexican Pacific Slope.**

In: Plant Systematics and Evolution 301 (2), p. 623–637. DOI: 10.1007/s00606-014-1101-1.

**Treutlein, Jens; Vorster, P.; Wink, Michael (2005):**

**Molecular Relationships in *Encephalartos* (Zamiaceae, Cycadales) based on Nucleotide Sequences of nuclear ITS 1&2, rbcL, and genomic Issr Fingerprinting.**

In: Plant Biology 7 (1), p. 79–90. DOI: 10.1055/s-2004-830478.

**Vovides, Andrew P.; González, Dolores; Pérez-Farrera, Miguel Angel; Avendaño, Sergio; Bárcenas, Cristina (2004):**

**A Review of research on the *Cycad* Genus *Ceratozamia* Brongn. (Zamiaceae) in Mexico.**

In: Taxon 53 (2), p. 291–297.

### Zingiberaceae

**Benedict, John C.; Smith, Selena Y.; Collinson, Margaret E.; Leong-Škorničková, Jana; Specht, Chelsea D.; Marone, Federica et al. (2015):**

**Seed Morphology and Anatomy and its utility in recognizing Subfamilies and Tribes of Zingiberaceae.**

In: American Journal of Botany 102 (11), p. 1814–1841. DOI: 10.3732/ajb.1500300.

**Droop, A. J.; Newman, Mark F. (2014):**

**A Revision of *Amomum* (Zingiberaceae) in Sumatra.**

In: Edinburgh Journal of Botany 71 (02), p. 193–258. DOI: 10.1017/S0960428614000043.

**Funakoshi, Hidenobu; Kress, W. John; Skornickova, Jana; Liu, Ai-Zhong; Inoue, Ken (2005):**

**Return from the Lost: Rediscovery of the Presumed extinct *Leptosolen* (Zingiberaceae) of the Philippines and its phylogenetic Placement in Gingers.**

In: Acta Phytotaxonomica Geobotanica 56 (1), p. 41–53.

**Gao, Li-Xia; Hu, Xiu; Liu, Nian; Bang-Hai; Li, Huang Zheng-Jun; Li, Yan (2008):**

**Cluster Analysis of Chinese *Hedychium* based on SRAP Markers.**

In: Journal of Systematics and Evolution 46 (6), p. 899–905.

**Julius, Avelinah; Suleiman, Monica; Takano, Atsuko; Nagamasu, Hidetoshi (2008):**

**Preliminary molecular Phylogeny of Bornean *Plagiostachys* (Zingiberaceae) based on DNA Sequence Data of Internal Transcribed Spacer (ITS).**

In: Journal of Tropical Biology and Conservation 4 (1), p. 67–80.

**Kaewsri, Wittaya; Paisooksantivatana, Yingyong; Veesommai, Uamporn; Eiadthong, Wichan; Vajrodaya, Srunya (2007):**

**Phylogenetic Analysis of Thai *Amomum* (Alpinioideae: Zingiberaceae) using AFLP Markers.**

In: Kasetsart Journal - Natural Science 41, p. 213–226.

**Kress, W. John; Liu, Ai-Zhong; Newman, Mark F.; Li, Qing-Jun (2005):**

**The molecular Phylogeny of *Alpinia* (Zingiberaceae): a complex and polyphyletic Genus of Gingers.**

In: American Journal of Botany 92 (1), p. 167–178. DOI: 10.3732/ajb.92.1.167.

**Kress, W. John; Prince, Linda M.; Williams, Kyle J. (2002):**

**The Phylogeny and a new Classification of the Gingers (Zingiberaceae): Evidence from molecular Data.**

In: American Journal of Botany 89 (10), p. 1682–1696. DOI: 10.3732/ajb.89.10.1682.

**Lamxay, Vichith; Newman, Mark F. (2012):**

**A Revision of *Amomum* (Zingiberaceae) in Cambodia, Laos and Vietnam.**
In: Edinburgh Journal of Botany 69 (01), p. 99–206. DOI: 10.1017/S0960428611000436.

LEONG-ŠKORNIČKOVÁ, JANA; ŠÍDA, OTAKAR; ZÁVESKÁ, ELIŠKA; MARHOLD, KAROL (2015):
**History of infrageneric Classification, typification of supraspecific Names and outstanding Transfers in *Curcuma* (Zingiberaceae).**
In: Taxon 64 (2), p. 362–373. DOI: 10.12705/642.11.

PEDERSEN, L. B. (2004):
**Phylogenetic Analysis of the Subfamily Alpinioideae (Zingiberaceae), particularly *Etlingera* Giseke, based on nuclear and Plastid DNA.**
In: Plant Systematics and Evolution 245 (3-4), p. 239–258. DOI: 10.1007/s00606-004-0126-2.

SÄRKINEN, TIINA E.; NEWMAN, MARK F.; MAAS, PAUL J.M.; MAAS, HILTJE; POULSEN, AXEL D.; HARRIS, DAVID J. ET AL. (2007):
**Recent oceanic long-distance Dispersal and Divergence in the amphi-Atlantic Rain Forest Genus *Renealmia* L.f. (Zingiberaceae).**
In: Molecular Phylogenetics and Evolution 44 (3), p. 968–980. DOI: 10.1016/j.ympev.2007.06.007.

TAKANO, ATSUKO; NAGAMASU, HIDETOSHI (2007):
***Myxochlamys* (Zingiberaceae), a new Genus from Borneo.**
In: Acta Phytotaxonomica Geobotanica 58 (1), p. 19–32.

TAKANO, ATSUKO; OKADA, HIROSHI (2002):
**Multiple occurrences of triploid Formation in *Globba* (Zingiberaceae) from molecular Evidence.**
In: Plant Systematics and Evolution 230 (3-4), p. 143–159. DOI: 10.1007/s006060200001.

TECHAPRASAN, JIRANAN; LEONG-ŠKORNIČKOVÁ, JANA (2011):
**Transfer of *Kaempferia candida* to *Curcuma* (Zingiberaceae) based on morphological and molecular Data.**
In: Nordic Journal of Botany 29 (6), p. 773–779. DOI: 10.1111/j.1756-1051.2011.00970.x.

VALDERRAMA, EUGENIO; RICHARDSON, JAMES E.; KIDNER, CATHERINE ANNE; MADRIÑÁN, SANTIAGO; STONE, GRAHAM N. (2018):
**Transcriptome mining for phylogenetic Markers in a recently radiated Genus of tropical plants (*Renealmia* L.f., Zingiberaceae).**
In: Molecular Phylogenetics and Evolution 119, p. 13–24. DOI: 10.1016/j.ympev.2017.10.001.

WILLIAMS, KYLE J.; KRESS, W. JOHN; MANOS, PAUL P. (2004):
**The Phylogeny, Evolution, and Classification of the Genus *Globba* and Tribe Globbeae (Zingiberaceae): Appendages do matter.**
In: American Journal of Botany 91 (1), p. 100–114. DOI: 10.3732/ajb.91.1.100.

WOOD, T. H.; WHITTEN, WILLIAM MARK; WILLIAMS, NORRIS H. (2000):
**Phylogeny of *Hedychium* and related Genera (Zingiberaceae) based on ITS Sequence Data.**
In: Edinburgh Journal of Botany 57 (2), p. 261–270. DOI: 10.1017/S0960428600000196.

XIA, YONG-MEI; KRESS, W. JOHN; PRINCE, LINDA M. (2004):
**Phylogenetic Analyses of *Amomum* (Alpinioideae: Zingiberaceae) using ITS and matK DNA Sequence Data.**
In: Systematic Botany 29 (2), p. 334–344.

ZÁVESKÁ, ELIŠKA; FÉR, TOMÁŠ; ŠÍDA, OTAKAR; KRAK, KAROL; MARHOLD, KAROL; LEONG-ŠKORNIČKOVÁ, JANA (2012):

**Phylogeny of *Curcuma* (Zingiberaceae) based on Plastid and nuclear Sequences: Proposal of the new Subgenus *Ecomata*.**
In: Taxon 61 (4), 747-743.

ZÁVESKÁ, ELIŠKA; FÉR, TOMÁŠ; ŠÍDA, OTAKAR; MARHOLD, KAROL; LEONG-ŠKORNIČKOVÁ, JANA (2016):
**Hybridization among distantly related species: Examples from the polyploid Genus *Curcuma* (Zingiberaceae).**
In: Molecular Phylogenetics and Evolution 100, p. 303–321. DOI: 10.1016/j.ympev.2016.04.017.

ZHAO, JIAN-LI; ZHONG, JIN-SHUN; FAN, YONG-LI; XIA, YONG-MEI; LI, QING-JUN (2017):
**A preliminary species-level Phylogeny of the alpine Ginger *Roscoea* : Implications for Speciation.**
In: Journal of Systematics and Evolution 55 (3), p. 215–224. DOI: 10.1111/jse.12247.

ZOU, P. U.; YE, YU-SHI; LIAO, JING-PING (2016):
***Alpinia austrosinense* (Zingiberaceae), a new Combination from China and its Relationship with *A. pumila*.**
In: Phytotaxa 255 (2), p. 175. DOI: 10.11646/phytotaxa.255.2.8.

### Zosteraceae

TANAKA, NORIO; KUO, JOHN; OMORI, YUJI; NAKAOKA, MASAHIRO; AIOI, KEIKO (2003):
**Phylogenetic Relationships in the Genera *Zostera* and *Heterozostera* (Zosteraceae) based on matK Sequence Data.**
In: Journal of Plant Research 116 (4), p. 273–279. DOI: 10.1007/s10265-003-0090-x.

### Zygophyllaceae

BEIER, BJÖRN-AXEL; CHASE, MARK W.; THULIN, MATS (2003):
**Phylogenetic Relationships and Taxonomy of Subfamily Zygophylloideae (Zygophyllaceae) based on molecular and morphological Data.**
In: Plant Systematics and Evolution 240 (1-4), p. 11–39. DOI: 10.1007/s00606-003-0007-0.

BEIER, BJÖRN-AXEL; NYLANDER, JOHAN A.A.; CHASE, MARK W.; THULIN, MATS (2004):
**Phylogenetic Relationships and Biogeography of the desert plant Genus *Fagonia* (Zygophyllaceae), inferred by Parsimony and Bayesian Model Averaging.**
In: Molecular Phylogenetics and Evolution 33 (1), p. 91–108. DOI: 10.1016/j.ympev.2004.05.010.

BELLSTEDT, DIRK U.; VAN ZYL, L.; MARAIS, E. M.; BYTEBIER, BENNY; VILLIERS, C. A.; MAKWARELA, A. M.; DREYER, L. L. (2008):
**Phylogenetic Relationships, Character Evolution and Biogeography of southern African members of *Zygophyllum* (Zygophyllaceae) based on three Plastid Regions.**
In: Molecular Phylogenetics and Evolution 47 (3), p. 932–949. DOI: 10.1016/j.ympev.2008.02.019.

LAUTERBACH, MAXIMILIAN; VAN DER MERWE, PIETER DE WET; KEẞLER, LISA; PIRIE, MICHAEL D.; BELLSTEDT, DIRK U.; KADEREIT, GUDRUN (2016):
**Evolution of Leaf Anatomy in arid Environments - A case Study in southern African *Tetraena* and *Roepera* (Zygophyllaceae).**
In: Molecular Phylogenetics and Evolution 97, p. 129–144. DOI: 10.1016/j.ympev.2016.01.002.

LIA, V. V.; CONFALONIERI, V. A.; COMAS, C. I.; HUNZIKER, J. H. (2001):
**Molecular Phylogeny of *Larrea* and its allies (Zygophyllaceae): reticulate Evolution and the probable time of Creosote bush Arrival to North America.**
In: Molecular Phylogenetics and Evolution 21 (2), p. 309–320. DOI: 10.1006/mpev.2001.1025.
