## Supplementary file 4 - Literature used to compile LCVP ordered by plant families 2 for "The Leipzig Catalogue of Vascular Plants (LCVP) – An improved taxonomic reference list for all known vascular plants"

TY  - JOUR
AB  - Nucleotide sequences of Rubisco Large Subunit (rbcL) and the internal transcribed spacers (ITS) of nrDNA were obtained for the five species of Larrea and one species each of Bulnesia (ITS only) and Plectrocarpa (rbcL only). Parsimony analyses were conducted, including sequences from seven genera of Zygophyllaceae reported by other authors-Kallstroemia, Zygophyllum, Augea, Fagonia, Pintoa, Guaiacum, and Porlieria. The main conclusions of the present study are (1) the Argentine endemic Plectrocarpa tetracantha belongs to the subfamily Larreoideae (New World Clade); (2) all three phylogenies obtained from rbcL, ITS, and combined data sets show a close relationship between the tetraploid L. cuneifolia (sect. Bifolium) and the diploid multifoliolate pair L. nitida-L. ameghinoi (sect. Larrea), which could result from a possible intersectional hybrid origin of the tetraploid; (3) L. divaricata (sect. Bifolium) and L. tridentata (sect. Bifolium) form a highly supported monophyletic group, which agrees with previous cytogenetic and molecular evidence; and (4) the rate of nucleotide substitution of rbcL was estimated based on geological and fossil records. Under the molecular clock hypothesis, nucleotide sequence divergence between L. divaricata and L. tridentata suggests a Late Neogene (8.4 to 4.2 mybp) time of arrival of the diploid ancestors of L. tridentata to North American deserts.
AD  - Departamento de Cs. Biológicas, Universidad de Buenos Aires, 1428 Buenos Aires, Argentina. M4 - Citavi
AU  - Lia, V. V.
AU  - Confalonieri, V. A.
AU  - Comas, C. I.
AU  - Hunziker, J. H.
DO  - 10.1006/mpev.2001.1025
IS  - 2
KW  - Zygophyllaceae
LA  - eng
N1  - Journal Article Research Support, Non-U.S. Gov't Journal Article Research Support, Non-U.S. Gov't Journal Article Research Support, Non-U.S. Gov't Journal Article Research Support, Non-U.S. Gov't
PY  - 2001
SN  - 1055-7903
SP  - 309-320
ST  - Molecular phylogeny of Larrea and its allies (Zygophyllaceae): reticulate evolution and the probable time of creosote bush arrival to North America
T2  - Molecular phylogenetics and evolution
TI  - Molecular phylogeny of Larrea and its allies (Zygophyllaceae): reticulate evolution and the probable time of creosote bush arrival to North America
VL  - 21
ID  - 4249
ER  - 


TY  - JOUR
AB  - The dry biomes of southern Africa (Desert, Nama Karoo and Succulent Karoo) are home to a rich and diverse xerophytic flora. This flora includes two morphologically diverse clades of Zygophyllaceae, Tetraena and Roepera (Zygophylloideae), which inhabit some of the most arid habitats in the region. Using a plastid phylogeny of Zygophylloideae we assess whether the evolution of putatively adaptive traits (leaf shape, vasculature, mode of water storage and photosynthetic type: C3 versus C4) coincides with the successful colonisation of environments with different drought regimes within southern Africa. Our results show general niche conservatism within arid habitats in Tetraena, but niche shifts from arid to more mesic biomes with longer and/or cooler growing season (Fynbos and Thicket) in Roepera. However, these distinct broad-scale biogeographical patterns are not reflected in leaf anatomy, which seems to vary at more local scales. We observed considerable variability and multiple convergences to similar leaf anatomies in both genera, including shifts between "all cell succulence" leaf types and leaf types with distinct chlorenchyma and hydrenchyma. Our survey of C4 photosynthesis in the Zygophylloideae showed that the C4 pathway is restricted to Tetraena simplex, which also having an annual life history and a widespread distribution, is rather atypical for this group.
AD  - Institut für Allgemeine und Spezielle Botanik, Johannes Gutenberg-Universität Mainz, 55099 Mainz, Germany. Electronic address:. Department of Biochemistry, University of Stellenbosch, Stellenbosch, South Africa. Institut für Allgemeine und Spezielle Botanik, Johannes Gutenberg-Universität Mainz, 55099 Mainz, Germany. Institut für Allgemeine und Spezielle Botanik, Johannes Gutenberg-Universität Mainz, 55099 Mainz, Germany; Department of Biochemistry, University of Stellenbosch, Stellenbosch, South Africa. Department of Biochemistry, University of Stellenbosch, Stellenbosch, South Africa. Institut für Allgemeine und Spezielle Botanik, Johannes Gutenberg-Universität Mainz, 55099 Mainz, Germany. M4 - Citavi
AU  - Lauterbach, Maximilian
AU  - van der Merwe, Pieter de Wet
AU  - Keßler, Lisa
AU  - Pirie, Michael D.
AU  - Bellstedt, Dirk U.
AU  - Kadereit, Gudrun
DO  - 10.1016/j.ympev.2016.01.002
KW  - Zygophyllaceae
LA  - eng
N1  - Journal Article Research Support, Non-U.S. Gov't Journal Article Research Support, Non-U.S. Gov't Journal Article Research Support, Non-U.S. Gov't Journal Article Research Support, Non-U.S. Gov't
PY  - 2016
SN  - 1055-7903
SP  - 129-144
ST  - Evolution of leaf anatomy in arid environments - A case study in southern African Tetraena and Roepera (Zygophyllaceae)
T2  - Molecular phylogenetics and evolution
TI  - Evolution of leaf anatomy in arid environments - A case study in southern African Tetraena and Roepera (Zygophyllaceae)
VL  - 97
ID  - 275
ER  - 


TY  - JOUR
AB  - The plastid coding rbcL and non-coding trnLF regions of 53 of 55 southern African Zygophyllum species were sequenced and used to evaluate the phylogenetic relationships within the southern African representatives of the genus. Published sequences of the same gene regions of Australian, Asian and North African Zygophyllum species were included to assess the relationships of the species from these regions to the southern African species. The addition of Z. stapffii from Namibia, found to be conspecific with Z. orbiculatum from Angola, lead to a greatly resolved tree. The molecular results were largely congruent with a recent sectional classification of the southern African species and supported their subdivision into subgenera Agrophyllum and Zygophyllum. Reconstruction of the character evolution of capsule dehiscence, seed attachment and seed mucilage showed that these characters allowed a division of southern African species into the two subgenera but that this could not be applied to species occurring elsewhere. Other morphological characters were found to vary and unique character combinations, rather than unique characters, were found to be of systematic value in sectional delimitation. The study suggests that repeated radiations from the horn of Africa to southern Africa and Asia and back lead to the present distribution of the taxa in the subfamily Zygophylloideae. Although this study supports some of the recent taxonomic changes in the group, the unresolved relationships between the proposed genera Tetraena and Roepera and those retained as Zygophyllum species suggest that changes to the taxonomy may have been premature.
AD  - Department of Biochemistry, University of Stellenbosch, Stellenbosch, Private Bag X1, Matieland 7602, South Africa. M4 - Citavi
AU  - Bellstedt, Dirk U.
AU  - van Zyl, L.
AU  - Marais, E. M.
AU  - Bytebier, Benny
AU  - Villiers, C. A.
AU  - Makwarela, A. M.
AU  - Dreyer, L. L.
DO  - 10.1016/j.ympev.2008.02.019
IS  - 3
KW  - Zygophyllaceae
LA  - eng
N1  - Journal Article Research Support, Non-U.S. Gov't Journal Article Research Support, Non-U.S. Gov't Journal Article Research Support, Non-U.S. Gov't Journal Article Research Support, Non-U.S. Gov't
PY  - 2008
SN  - 1055-7903
SP  - 932-949
ST  - Phylogenetic relationships, character evolution and biogeography of southern African members of Zygophyllum (Zygophyllaceae) based on three plastid regions
T2  - Molecular phylogenetics and evolution
TI  - Phylogenetic relationships, character evolution and biogeography of southern African members of Zygophyllum (Zygophyllaceae) based on three plastid regions
VL  - 47
ID  - 2747
ER  - 


TY  - JOUR
AB  - Phylogenetic relationships within Fagonia were inferred from analyses of plastid trnL intron and nuclear ribosomal ITS DNA sequences. Sampling of the genus was nearly complete, including 32 of 34 species. Phylogenetic analysis was carried out using parsimony, and Bayesian model averaging. The latter method allows model-based inference while accounting for model-selection uncertainty, and is here used for the first time in phylogenetic analyses. All species of Fagonia in the Old World, except F. cretica, form a weakly supported clade, and all Fagonia species of the New World, except F. scoparia, are well supported as sister to the Old World clade. Fagonia scoparia, from Mexico, and F. cretica, from Northern Africa, are well supported as sisters to all other Fagonia species. Vicariance-dispersal analysis, using DIVA, indicated that the occurrences of Fagonia in South America and southern Africa are due to dispersals, and also, that the ancestor of Fagonia had a distribution compatible with the boreotropics hypothesis.
AD  - Department of Systematic Botany, Evolutionary Biology Centre, Uppsala University, Norbyvägen 18D, 752 36 Uppsala, Sweden. M4 - Citavi
AU  - Beier, Björn-Axel
AU  - Nylander, J. A. A.
AU  - Chase, Mark W.
AU  - Thulin, M.
DO  - 10.1016/j.ympev.2004.05.010
IS  - 1
KW  - Zygophyllaceae
LA  - eng
N1  - Comparative Study Journal Article Research Support, Non-U.S. Gov't Comparative Study Journal Article Research Support, Non-U.S. Gov't Comparative Study Journal Article Research Support, Non-U.S. Gov't Comparative Study Journal Article Research Support, Non-U.S. Gov't
PY  - 2004
SN  - 1055-7903
SP  - 91-108
ST  - Phylogenetic relationships and biogeography of the desert plant genus Fagonia (Zygophyllaceae), inferred by parsimony and Bayesian model averaging
T2  - Molecular phylogenetics and evolution
TI  - Phylogenetic relationships and biogeography of the desert plant genus Fagonia (Zygophyllaceae), inferred by parsimony and Bayesian model averaging
VL  - 33
ID  - 3754
ER  - 


TY  - JOUR
AU  - Beier, Björn-Axel
AU  - Chase, Mark W.
AU  - Thulin, M.
DO  - 10.1007/s00606-003-0007-0 M4 - Citavi
IS  - 1-4
KW  - Zygophyllaceae
N1  - PII: 6XFDGHCX9KXKV9EM PII: 6XFDGHCX9KXKV9EM PII: 6XFDGHCX9KXKV9EM PII: 6XFDGHCX9KXKV9EM
PY  - 2003
SN  - 0378-2697
SP  - 11-39
ST  - Phylogenetic relationships and taxonomy of subfamily Zygophylloideae (Zygophyllaceae) based on molecular and morphological data
T2  - Plant Systematics and Evolution
TI  - Phylogenetic relationships and taxonomy of subfamily Zygophylloideae (Zygophyllaceae) based on molecular and morphological data
VL  - 240
ID  - 3958
ER  - 


TY  - JOUR
AB  - Phylogenetic analysis of the plastid (chloroplast) DNA matK gene of Zosteraceae species was undertaken. A molecular phylogenetic tree based on matK sequence data showed the monophyly of Heterozostera tasmanica and subgenus Zosterella and did not support the separation of Heterozostera from the genus Zostera. The tree based on matK supported the monophyly of the subgenus Zostera, and showed that Zosteraceae consist of three main groups: Phyllospadix, which is clearly defined by being dioecious; the subgenus Zosterella and Heterozostera; and the subgenus Zostera. Character-state reconstruction of chromosome number and geographic distribution for our molecular phylogenetic tree showed that 2 n=12 is a plesiomorphic character for Zostera and Heterozostera, that the chromosome number was doubled or tripled in two lineages, and that the initial speciation of Zostera and Heterozostera occurred in the Northern Hemisphere. The matK tree showed the close affinity of Z. noltii and Z. japonica, which have disjunct distributions. Zostera marina, which is the only widely distributed species in the subgenus Zostera, also occurring in the northern Atlantic, was shown to be embedded within other subgenus Zostera species.
AD  - Tsukuba Botanical Garden, National Science Museum, 4-1-1 Amakubo, Tsukuba 305-0005 Japan. M4 - Citavi
AU  - Tanaka, Norio
AU  - Kuo, John
AU  - Omori, Yuji
AU  - Nakaoka, Masahiro
AU  - Aioi, Keiko
DO  - 10.1007/s10265-003-0090-x
IS  - 4
KW  - Zosteraceae
LA  - eng
N1  - Journal Article Research Support, Non-U.S. Gov't Journal Article Research Support, Non-U.S. Gov't Journal Article Research Support, Non-U.S. Gov't Journal Article Research Support, Non-U.S. Gov't
PY  - 2003
SP  - 273-279
ST  - Phylogenetic relationships in the genera Zostera and Heterozostera (Zosteraceae) based on matK sequence data
T2  - Journal of Plant Research
TI  - Phylogenetic relationships in the genera Zostera and Heterozostera (Zosteraceae) based on matK sequence data
VL  - 116
ID  - 4057
ER  - 


TY  - JOUR
AU  - Zou, P. U.
AU  - Ye, Yu-shi
AU  - Liao, Jing-ping
DO  - 10.11646/phytotaxa.255.2.8 M4 - Citavi
IS  - 2
KW  - Zingiberaceae
PY  - 2016
SN  - 1179-3163
SP  - 175
ST  - Alpinia austrosinense (Zingiberaceae), a new combination from China and its relationship with A. pumila
T2  - Phytotaxa
TI  - Alpinia austrosinense (Zingiberaceae), a new combination from China and its relationship with A. pumila
VL  - 255
ID  - 424
ER  - 


TY  - JOUR
AD  - Laboratory of Ecology and Evolutionary Biology, State Key Laboratory for Conservation and Utilization of Bio-Resources in Yunnan; Yunnan University; Kunming 650091 China Xishuangbanna Tropical Botanical Garden; Chinese Academy of Sciences; Mengla 666303 China Department of Plant Biology; University of Vermont; 63 Carrigan Drive Burlington, VT 05405 USA China Forest Exploration and Design Institute of Kunming; Kunming 650216 China M4 - Citavi
AU  - Zhao, Jian-Li
AU  - Zhong, Jinshun
AU  - Fan, Yong-Li
AU  - Xia, Yong-Mei
AU  - Li, Qing-Jun
DO  - 10.1111/jse.12247
IS  - 3
KW  - Zingiberaceae
PY  - 2017
SN  - 16744918
SP  - 215-224
ST  - A preliminary species-level phylogeny of the alpine ginger Roscoea : Implications for speciation
T2  - Journal of Systematics and Evolution
TI  - A preliminary species-level phylogeny of the alpine ginger Roscoea : Implications for speciation
VL  - 55
ID  - 144
ER  - 


TY  - JOUR
AB  - Discerning relationships among species evolved by reticulate and/or polyploid evolution is not an easy task, although it is widely discussed. The economically important genus Curcuma (ca. 120 spp.; Zingiberaceae), broadly distributed in tropical SE Asia, is a particularly interesting example of a group of palaeopolyploid origin whose evolution is driven mainly by hybridization and polyploidization. Although a phylogeny and a new infrageneric classification of Curcuma, based on commonly used molecular markers (ITS and cpDNA), have recently been proposed, significant evolutionary questions remain unresolved. We applied a multilocus approach and a combination of modern analytical methods to this genus to distinguish causes of gene tree incongruence and to identify hybrids and their parental species. Five independent regions of nuclear DNA (DCS, GAPDH, GLOBOSA3, LEAFY, ITS) and four non-coding cpDNA regions (trnL-trnF, trnT-trnL, psbA-trnH and matK), analysed as a single locus, were employed to construct a species tree and hybrid species trees using (*)BEAST and STEM-hy. Detection of hybridogenous species in the dataset was also conducted using the posterior predictive checking approach as implemented in JML. The resulting species tree outlines the relationships among major evolutionary lineages within Curcuma, which were previously unresolved or which conflicted depending upon whether they were based on ITS or cpDNA markers. Moreover, by using the additional markers in tests of plausible topologies of hybrid species trees for C. vamana, C. candida, C. roscoeana and C. myanmarensis suggested by previous molecular and morphological evidence, we found strong evidence that all the species except C. candida are of subgeneric hybrid origin.
AD  - Department of Botany, Faculty of Science, Charles University, Prague, Czech Republic. Electronic address:. Department of Botany, Faculty of Science, Charles University, Prague, Czech Republic. Department of Botany, National Museum in Prague, Prague, Czech Republic. Department of Botany, Faculty of Science, Charles University, Prague, Czech Republic; Institute of Botany, Slovak Academy of Sciences, Bratislava, Slovak Republic. The Herbarium, Singapore Botanic Gardens, Singapore. M4 - Citavi
AU  - Záveská, Eliška
AU  - Fér, Tomáš
AU  - Šída, Otakar
AU  - Marhold, Karol
AU  - Leong-Škorničková, Jana
DO  - 10.1016/j.ympev.2016.04.017
KW  - Zingiberaceae
LA  - eng
N1  - Journal Article Journal Article Journal Article Journal Article
PY  - 2016
SN  - 1055-7903
SP  - 303-321
ST  - Hybridization among distantly related species: Examples from the polyploid genus Curcuma (Zingiberaceae)
T2  - Molecular phylogenetics and evolution
TI  - Hybridization among distantly related species: Examples from the polyploid genus Curcuma (Zingiberaceae)
VL  - 100
ID  - 417
ER  - 


TY  - JOUR
AU  - Záveská, Eliška
AU  - Fér, Tomáš
AU  - Šída, Otakar
AU  - Krak, Karol
AU  - Marhold, Karol
AU  - Leong-Škorničková, Jana
IS  - 4
KW  - Zingiberaceae
PY  - 2012
SN  - 00400262
SP  - 747-743
ST  - Phylogeny of Curcuma (Zingiberaceae) based on plastid and nuclear sequences: Proposal of the new subgenus Ecomata
T2  - Taxon
TI  - Phylogeny of Curcuma (Zingiberaceae) based on plastid and nuclear sequences: Proposal of the new subgenus Ecomata
VL  - 61
ID  - 1782
ER  - 


TY  - JOUR
AU  - Xia, Yong-Mei
AU  - Kress, W. John
AU  - Prince, Linda M.
IS  - 2
KW  - Zingiberaceae
PY  - 2004
SN  - 03636445
SP  - 334-344
ST  - Phylogenetic Analyses of Amomum (Alpinioideae: Zingiberaceae) Using ITS and matK DNA Sequence Data
T2  - Systematic Botany
TI  - Phylogenetic Analyses of Amomum (Alpinioideae: Zingiberaceae) Using ITS and matK DNA Sequence Data
VL  - 29
ID  - 3933
ER  - 


TY  - JOUR
AB  - Edinburgh Journal of Botany
AU  - Wood, T. H.
AU  - Whitten, William Mark
AU  - Williams, Norris H.
DO  - 10.1017/S0960428600000196 Y3 - 23.01.2019 M4 - Citavi
IS  - 2
KW  - Zingiberaceae
N1  - PII: S0960428600000196 PII: S0960428600000196 PII: S0960428600000196 PII: S0960428600000196
PY  - 2000
SN  - 0960-4286
SP  - 261-270
ST  - Phylogeny of Hedychium And Related Genera (Zingiberaceae) based on ITS Sequence Data
T2  - Edinburgh Journal of Botany
TI  - Phylogeny of Hedychium And Related Genera (Zingiberaceae) based on ITS Sequence Data
VL  - 57
ID  - 4393
ER  - 


TY  - JOUR
AB  - The genus Globba (100 species) is one of the largest genera in the primarily tropical Zingiberaceae. Globba along with the small genera Gagnepainia, Hemiorchis, and Mantisia comprise the Globbeae, one of the two tribes of subfamily Zingiberoideae. Traditional infrageneric classification in Globba has focused on the number of anther appendages: zero, two, or four. Parsimony and Bayesian analyses were conducted on nuclear internal transcribed spacer (ITS) and plastid trnK-matK data from a broad sampling of Globba and related genera. Results show Mantisia to be monophyletic but nested within Globba, while Hemiorchis and Gagnepainia are monophyletic genera that are sister to each other. Anther appendage number and shape, along with inflorescence and fruit morphology, are the most important characters for understanding evolutionary relationships in Globba. A new infrageneric classification system for Globba, recognizing three subgenera and seven sections is presented. The four species of Mantisia are formally transferred into Globba but retained as a distinct section. Within Globba, a notable biogeographic boundary is seen at the Isthmus of Kra in southern Thailand.
AD  - Department of Biology, Box 90338, Duke University, Durham, North Carolina 27708 USA; M4 - Citavi
AU  - Williams, Kyle J.
AU  - Kress, W. John
AU  - Manos, Paul S.
DO  - 10.3732/ajb.91.1.100
IS  - 1
KW  - Zingiberaceae
LA  - eng
N1  - Journal Article Journal Article Journal Article Journal Article
PY  - 2004
SN  - 0002-9122
SP  - 100-114
ST  - The phylogeny, evolution, and classification of the genus Globba and tribe Globbeae (Zingiberaceae): appendages do matter
T2  - American journal of botany
TI  - The phylogeny, evolution, and classification of the genus Globba and tribe Globbeae (Zingiberaceae): appendages do matter
VL  - 91
ID  - 3929
ER  - 


TY  - JOUR
AB  - The reconstruction of relationships within species-rich groups that have recently evolved in biodiversity hotspots is hampered by a lack of phylogenetically informative markers. It is also made difficult by the lack of sampling necessary to reconstruct a species-level phylogeny. We use transcriptome mining to search for markers to reconstruct a phylogeny of the amphi-Atlantic genus Renealmia L. f. (Zingiberaceae). We recover seven introns from single copy genes and use them to reconstruct the phylogeny of the genus together with a commonly used phylogenetic marker, internal transcribed spacers of ribosomal DNA (ITS) that has previously been used to reconstruct the phylogeny of the genus. We targeted genes with low numbers of base pairs that improves sequencing success using highly degraded DNA from herbarium specimens. The use of herbarium specimens greatly increased the number of species in the study as these were readily available in historical collections. Data were obtained for 14 of the 17 African species and 54 of the 65 Neotropical species. The phylogeny was well-supported for a number of Renealmia subgroups although relationships among those clades remained poorly supported.
AD  - Royal Botanic Garden Edinburgh, Edinburgh, Scotland, United Kingdom; Institute of Evolutionary Biology, University of Edinburgh, Edinburgh, Scotland, United Kingdom; Laboratorio de Análisis del Comportamiento Humano, Facultad de Psicología y Grupo de Investigación de Biología (GRIB), Universidad El Bosque, Bogotá, Colombia. Electronic address:. Royal Botanic Garden Edinburgh, Edinburgh, Scotland, United Kingdom; Programa de Biología, Universidad del Rosario, Bogotá, Colombia. Electronic address:. Royal Botanic Garden Edinburgh, Edinburgh, Scotland, United Kingdom; Institute of Molecular Plant Sciences, University of Edinburgh, Edinburgh, Scotland, United Kingdom. Electronic address:. Laboratorio de Botánica y Sistemática, Universidad de Los Andes, Bogotá, Colombia. Electronic address:. Royal Botanic Garden Edinburgh, Edinburgh, Scotland, United Kingdom; Institute of Evolutionary Biology, University of Edinburgh, Edinburgh, Scotland, United Kingdom. Electronic address:. M4 - Citavi
AU  - Valderrama, Eugenio
AU  - Richardson, James E.
AU  - Kidner, Catherine Anne
AU  - Madriñán, Santiago
AU  - Stone, Graham N.
DO  - 10.1016/j.ympev.2017.10.001
KW  - Zingiberaceae
LA  - eng
N1  - Journal Article Research Support, Non-U.S. Gov't Journal Article Research Support, Non-U.S. Gov't Journal Article Research Support, Non-U.S. Gov't Journal Article Research Support, Non-U.S. Gov't
PY  - 2018
SN  - 1055-7903
SP  - 13-24
ST  - Transcriptome mining for phylogenetic markers in a recently radiated genus of tropical plants (Renealmia L.f., Zingiberaceae)
T2  - Molecular phylogenetics and evolution
TI  - Transcriptome mining for phylogenetic markers in a recently radiated genus of tropical plants (Renealmia L.f., Zingiberaceae)
VL  - 119
ID  - 37
ER  - 


TY  - JOUR
AU  - Techaprasan, Jiranan
AU  - Leong-Škorničková, Jana
DO  - 10.1111/j.1756-1051.2011.00970.x M4 - Citavi
IS  - 6
KW  - Zingiberaceae
PY  - 2011
SN  - 0107055X
SP  - 773-779
ST  - Transfer of Kaempferia candida to Curcuma (Zingiberaceae) based on morphological and molecular data
T2  - Nordic Journal of Botany
TI  - Transfer of Kaempferia candida to Curcuma (Zingiberaceae) based on morphological and molecular data
VL  - 29
ID  - 2053
ER  - 


TY  - JOUR
AU  - Takano, A.
AU  - Okada, H.
DO  - 10.1007/s006060200001 Y3 - 23.01.2019 M4 - Citavi
IS  - 3-4
KW  - Zingiberaceae
N1  - PII: 59K4UQKVDVV70XKT PII: 59K4UQKVDVV70XKT PII: 59K4UQKVDVV70XKT PII: 59K4UQKVDVV70XKT
PY  - 2002
SN  - 0378-2697
SP  - 143-159
ST  - Multiple occurrences of triploid formation in Globba (Zingiberaceae) from molecular evidence
T2  - Plant Systematics and Evolution
TI  - Multiple occurrences of triploid formation in Globba (Zingiberaceae) from molecular evidence
VL  - 230
ID  - 4176
ER  - 


TY  - JOUR
AU  - Takano, Atsuko
AU  - Nagamasu, Hidetoshi
IS  - 1
KW  - Zingiberaceae
PY  - 2007
SP  - 19-32
ST  - Myxochlamys (Zingiberaceae), a new genus from Borneo
T2  - Acta Phytotaxonomica Geobotanica
TI  - Myxochlamys (Zingiberaceae), a new genus from Borneo
VL  - 58
ID  - 3274
ER  - 


TY  - JOUR
AB  - Renealmia L.f. (Zingiberaceae) is one of the few tropical plant genera with numerous species in both Africa and South America but not in Asia. Based on phylogenetic analysis of nuclear ribosomal internal transcribed spacer (ITS) and chloroplast trnL-F DNA, Renealmia is shown to be monophyletic with high branch support. Low sequence divergence found in the two genome regions (ITS: 0-2.4%; trnL-F: 0-1.9%) suggests recent diversification within the genus. Molecular divergence age estimates give further support to the recent origin of the genus and show that Renealmia has attained its amphi-Atlantic distribution by an oceanic long-distance dispersal event from Africa to South America during the Miocene or Pliocene (15.8-2.7 My ago). Some support is found for the hypothesis that speciation in neotropical Renealmia was influenced by the Andean orogeny. Speciation has been approximately simultaneous on both sides of the Atlantic, but increased taxon sampling is required to compare the speciation rates between the New World and Old World tropics.
AD  - Royal Botanic Garden Edinburgh, 20A Inverleith Row, EH3 5LR Edinburgh, UK. M4 - Citavi
AU  - Särkinen, Tiina E.
AU  - Newman, Mark F.
AU  - Maas, Paul J. M.
AU  - Maas, Hiltje
AU  - Poulsen, Axel D.
AU  - Harris, David J.
AU  - Richardson, James E.
AU  - Clark, Alexandra
AU  - Hollingsworth, Michelle L.
AU  - Pennington, R. Toby
DO  - 10.1016/j.ympev.2007.06.007
IS  - 3
KW  - Zingiberaceae
LA  - eng
N1  - Journal Article Research Support, Non-U.S. Gov't Journal Article Research Support, Non-U.S. Gov't Journal Article Research Support, Non-U.S. Gov't Journal Article Research Support, Non-U.S. Gov't
PY  - 2007
SN  - 1055-7903
SP  - 968-980
ST  - Recent oceanic long-distance dispersal and divergence in the amphi-Atlantic rain forest genus Renealmia L.f. (Zingiberaceae)
T2  - Molecular phylogenetics and evolution
TI  - Recent oceanic long-distance dispersal and divergence in the amphi-Atlantic rain forest genus Renealmia L.f. (Zingiberaceae)
VL  - 44
ID  - 3254
ER  - 


TY  - JOUR
AU  - Pedersen, L. B.
DO  - 10.1007/s00606-004-0126-2 M4 - Citavi
IS  - 3-4
KW  - Zingiberaceae
N1  - PII: QV31XAACK79074GB PII: QV31XAACK79074GB PII: QV31XAACK79074GB PII: QV31XAACK79074GB
PY  - 2004
SN  - 0378-2697
SP  - 239-258
ST  - Phylogenetic analysis of the subfamily Alpinioideae (Zingiberaceae), particularly Etlingera Giseke, based on nuclear and plastid DNA
T2  - Plant Systematics and Evolution
TI  - Phylogenetic analysis of the subfamily Alpinioideae (Zingiberaceae), particularly Etlingera Giseke, based on nuclear and plastid DNA
VL  - 245
ID  - 3857
ER  - 


TY  - JOUR
AD  - The Herbarium, Singapore Botanic Gardens, 1 Cluny Road, 259569, Singapore;, Department of Botany, National Museum in Prague, Cirkusová 1840, 193 00 Praha 9, Czech Republic Department of Botany, Faculty of Science, Charles University, Benátská 2, 128 01 Praha 2, Czech Republic Department of Botany, Faculty of Science, Charles University, Benátská 2, 128 01 Praha 2, Czech Republic, Institute of Botany, Slovak Academy of Sciences, Dúbravská cesta 9, 845 23 Bratislava, Slovak Republic M4 - Citavi
AU  - Leong-Škorničková, Jana
AU  - Šída, Otakar
AU  - Záveská, Eliška
AU  - Marhold, Karol
DO  - 10.12705/642.11
IS  - 2
KW  - Zingiberaceae
PY  - 2015
SN  - 00400262
SP  - 362-373
ST  - History of infrageneric classification, typification of supraspecific names and outstanding transfers in Curcuma (Zingiberaceae)
T2  - Taxon
TI  - History of infrageneric classification, typification of supraspecific names and outstanding transfers in Curcuma (Zingiberaceae)
VL  - 64
ID  - 582
ER  - 


TY  - JOUR
AU  - Lamxay, Vichith
AU  - Newman, M. F.
DO  - 10.1017/S0960428611000436 M4 - Citavi
IS  - 01
KW  - Zingiberaceae
N1  - PII: S0960428611000436 PII: S0960428611000436 PII: S0960428611000436 PII: S0960428611000436
PY  - 2012
SN  - 0960-4286
SP  - 99-206
ST  - A Revision of Amomum (Zingiberaceae) in Cambodia, Laos and Vietnam
T2  - Edinburgh Journal of Botany
TI  - A Revision of Amomum (Zingiberaceae) in Cambodia, Laos and Vietnam
VL  - 69
ID  - 1603
ER  - 


TY  - JOUR
AB  - The pantropical Zingiberaceae is the largest family in the order Zingiberales with 53 genera and over 1200 species. Classifications of the family first proposed in 1889 and refined by others since that time recognize four tribes (Globbeae, Hedychieae, Alpinieae, and Zingibereae) based on morphological features, such as number of locules and placentation in the ovary, development of staminodia, modifications of the fertile anther, and rhizome-shoot-leaf orientation. New phylogenetic analyses based on DNA sequences of the nuclear internal transcribed spacer (ITS) and plastid matK regions suggest that at least some of these morphological traits are homoplasious and three of the tribes are paraphyletic. The African genus Siphonochilus and Bornean genus Tamijia are basal clades. The former Alpinieae and Hedychieae for the most part are monophyletic taxa with the Globbeae and Zingibereae included within the latter. The results of these phylogenetic investigations are used to propose a new classification of the Zingiberaceae that recognizes four subfamilies and four tribes: Siphonochiloideae (Siphonochileae), Tamijioideae (Tamijieae), Alpinioideae (Alpinieae, Riedelieae), and Zingiberoideae (Zingibereae, Globbeae). Morphological features congruent with this classification and the taxonomic status of various monotypic genera are discussed.
AD  - Botany, MRC-166, United States National Herbarium, National Museum of Natural History, Smithsonian Institution, P.O. Box 37012, Washington, D.C. 20013-7012 USA; M4 - Citavi
AU  - Kress, W. John
AU  - Prince, Linda M.
AU  - Williams, Kyle J.
DO  - 10.3732/ajb.89.10.1682
IS  - 10
KW  - Zingiberaceae
LA  - eng
N1  - Journal Article Journal Article Journal Article Journal Article
PY  - 2002
SN  - 0002-9122
SP  - 1682-1696
ST  - The phylogeny and a new classification of the gingers (Zingiberaceae): evidence from molecular data
T2  - American journal of botany
TI  - The phylogeny and a new classification of the gingers (Zingiberaceae): evidence from molecular data
VL  - 89
ID  - 4127
ER  - 


TY  - JOUR
AB  - Alpinia is the largest, most widespread, and most taxonomically complex genus in the Zingiberaceae with 230 species occurring throughout tropical and subtropical Asia. Species of Alpinia often predominate in the understory of forests, while others are important ornamentals and medicinals. Investigations of the evolutionary relationships of a subset of species of Alpinia using DNA sequence-based methods specifically test the monophyly of the genus and the validity of the previous classifications. Seventy-two species of Alpinia, 27 non-Alpinia species in the subfamily Alpinioideae, eight species in the subfamily Zingiberoideae, one species in the subfamily Tamijioideae, and three species in the outgroup genus Siphonochilus (Siphonochiloideae) were sequenced for the plastid matK region and the nuclear internal transcribed spacer (ITS) loci. Parsimony analyses of both individual and combined data sets identified six polyphyletic clades containing species of Alpinia distributed across the tribe Alpinieae. These results were supported by a Bayesian analysis of the combined data set. Except in a few specific cases, these monophyletic groupings of species do not correspond with either Schumann's (1904) or Smith's (1990) classification of the genus. Here we build on previous molecular analyses of the Alpinioideae and propose the next steps necessary to recognize new generic boundaries in the Alpinieae.
AD  - Department of Botany, MRC-166, United States National Herbarium, National Museum of Natural History, Smithsonian Institution, PO Box 37012, Washington, D.C. 20013-7012 USA; M4 - Citavi
AU  - Kress, W. John
AU  - Liu, Ai-Zhong
AU  - Newman, Mark
AU  - Li, Qing-Jun
DO  - 10.3732/ajb.92.1.167
IS  - 1
KW  - Zingiberaceae
LA  - eng
N1  - Journal Article Journal Article Journal Article Journal Article
PY  - 2005
SN  - 0002-9122
SP  - 167-178
ST  - The molecular phylogeny of Alpinia (Zingiberaceae): a complex and polyphyletic genus of gingers
T2  - American journal of botany
TI  - The molecular phylogeny of Alpinia (Zingiberaceae): a complex and polyphyletic genus of gingers
VL  - 92
ID  - 3619
ER  - 


TY  - JOUR
AU  - Kaewsri, Wittaya
AU  - Paisooksantivatana, Yingyong
AU  - Veesommai, Uamporn
AU  - Eiadthong, Wichan
AU  - Vajrodaya, Srunya
KW  - Zingiberaceae
PY  - 2007
SP  - 213-226
ST  - Phylogenetic Analysis of Thai Amomum (Alpinioideae: Zingiberaceae) Using AFLP Markers
T2  - Kasetsart J. (Nat. Sci.)
TI  - Phylogenetic Analysis of Thai Amomum (Alpinioideae: Zingiberaceae) Using AFLP Markers
VL  - 41
ID  - 3153
ER  - 


TY  - JOUR
AU  - Julius, Avelinah
AU  - Suleiman, Monica
AU  - Takano, Atsuko
AU  - Nagamasu, Hidetoshi
IS  - 1
KW  - Zingiberaceae
PY  - 2008
SP  - 67-80
ST  - Preliminary molecular phylogeny of Bornean Plagiostachys (Zingiberaceae) based on DNA sequence data of internal transcribed spacer (ITS)
T2  - Journal of Tropical Biology and Conservation
TI  - Preliminary molecular phylogeny of Bornean Plagiostachys (Zingiberaceae) based on DNA sequence data of internal transcribed spacer (ITS)
VL  - 4
ID  - 2847
ER  - 


TY  - JOUR
AU  - Gao, Li-Xia
AU  - Hu, Xiu
AU  - Liu, Nian
AU  - Bang, Hai
AU  - Li, Huang Zheng-Jun
AU  - Li, Yan
IS  - 6
KW  - Zingiberaceae
PY  - 2008
SN  - 16744918
SP  - 899-905
ST  - Cluster analysis of Chinese Hedychium based on SRAP markers
T2  - Journal of Systematics and Evolution
TI  - Cluster analysis of Chinese Hedychium based on SRAP markers
VL  - 46
ID  - 2808
ER  - 


TY  - JOUR
AU  - Funakoshi, Hidenobu
AU  - Kress, W. John
AU  - Skornickova, Jana
AU  - Liu, Ai-Zhong
AU  - Inoue, Ken
IS  - 1
KW  - Zingiberaceae
PY  - 2005
SP  - 41-53
ST  - Return from the Lost: Rediscovery of the Presumed extinct Leptosolen (Zingiberaceae) of the Philippines and its phylogenetic Placement in Gingers
T2  - Acta Phytotaxonomica Geobotanica
TI  - Return from the Lost: Rediscovery of the Presumed extinct Leptosolen (Zingiberaceae) of the Philippines and its phylogenetic Placement in Gingers
VL  - 56
ID  - 3589
ER  - 


TY  - JOUR
AU  - Droop, A. J.
AU  - Newman, M. F.
DO  - 10.1017/S0960428614000043 M4 - Citavi
IS  - 02
KW  - Zingiberaceae
PY  - 2014
SN  - 0960-4286
SP  - 193-258
ST  - A Revision of Amomum (Zingiberaceae) In Sumatra
T2  - Edinburgh Journal of Botany
TI  - A Revision of Amomum (Zingiberaceae) In Sumatra
VL  - 71
ID  - 884
ER  - 


TY  - JOUR
AB  - PREMISE OF THE STUDY Recent phylogenetic analyses based on molecular data suggested that the monocot family Zingiberaceae be separated into four subfamilies and four tribes. Robust morphological characters to support these clades are lacking. Seeds were analyzed in a phylogenetic context to test independently the circumscription of clades and to better understand evolution of seed characters within Zingiberaceae. METHODS Seventy-five species from three of the four subfamilies were analyzed using synchrotron based x-ray tomographic microscopy (SRXTM) and scored for 39 morphoanatomical characters. KEY RESULTS Zingiberaceae seeds are some of the most structurally complex seeds in angiosperms. No single seed character was found to distinguish each subfamily, but combinations of characters were found to differentiate between the subfamilies. Recognition of the tribes based on seeds was possible for Globbeae, but not for Alpinieae, Riedelieae, or Zingibereae, due to considerable variation. CONCLUSIONS SRXTM is an excellent, nondestructive tool to capture morphoanatomical variation of seeds and allows for the study of taxa with limited material available. Alpinioideae, Siphonochiloideae, Tamijioideae, and Zingiberoideae are well supported based on both molecular and morphological data, including multiple seed characters. Globbeae are well supported as a distinctive tribe within the Zingiberoideae, but no other tribe could be differentiated using seeds due to considerable homoplasy when compared with currently accepted relationships based on molecular data. Novel seed characters suggest tribal affinities for two currently unplaced Zingiberaceae taxa: Siliquamomum may be related to Riedelieae and Monolophus to Zingibereae, but further work is needed before formal revision of the family.
AD  - Department of Earth and Environmental Sciences, University of Michigan, Ann Arbor, Michigan 48109-1005 USA. Department of Earth and Environmental Sciences, University of Michigan, Ann Arbor, Michigan 48109-1005 USA Museum of Paleontology, University of Michigan, Ann Arbor, Michigan 48109-1079 USA. Department of Earth Sciences, Royal Holloway University of London, London, UK, TW20 0EX. Herbarium, Singapore Botanic Gardens, National Parks Board, Singapore 259569. Department of Plant and Microbial Biology, Integrative Biology and the University and Jepson Herbaria, University of California, Berkeley, California 94750-2465 USA. Swiss Light Source, Paul Scherrer Institut, 5232 Villigen PSI, Switzerland. Advanced Photon Source, Argonne National Laboratories, Argonne, Illinois 60439 USA. Advanced Light Source, Lawrence Berkeley National Laboratories, Berkeley, California 94720 USA. M4 - Citavi
AU  - Benedict, John C.
AU  - Smith, Selena Y.
AU  - Collinson, Margaret E.
AU  - Leong-Škorničková, Jana
AU  - Specht, Chelsea D.
AU  - Marone, Federica
AU  - Xiao, Xianghui
AU  - Parkinson, Dilworth Y.
DO  - 10.3732/ajb.1500300
IS  - 11
KW  - Zingiberaceae
LA  - eng
N1  - Journal Article Research Support, Non-U.S. Gov't Research Support, U.S. Gov't, Non-P.H.S. Journal Article Research Support, Non-U.S. Gov't Research Support, U.S. Gov't, Non-P.H.S. Journal Article Research Support, Non-U.S. Gov't Research Support, U.S. Gov't, Non-P.H.S. Journal Article Research Support, Non-U.S. Gov't Research Support, U.S. Gov't, Non-P.H.S.
PY  - 2015
SN  - 0002-9122
SP  - 1814-1841
ST  - Seed morphology and anatomy and its utility in recognizing subfamilies and tribes of Zingiberaceae
T2  - American journal of botany
TI  - Seed morphology and anatomy and its utility in recognizing subfamilies and tribes of Zingiberaceae
VL  - 102
ID  - 443
ER  - 


TY  - JOUR
AU  - Vovides, Andrew P.
AU  - González, Dolores
AU  - Pérez-Farrera, Miguel Angel
AU  - Avendaño, Sergio
AU  - Bárcenas, Cristina
IS  - 2
KW  - Zamiaceae
PY  - 2004
SN  - 00400262
SP  - 291-297
ST  - A review of research on the Cycad genus Ceratozamia Brongn. (Zamiaceae) in Mexico
T2  - Taxon
TI  - A review of research on the Cycad genus Ceratozamia Brongn. (Zamiaceae) in Mexico
VL  - 53
ID  - 3921
ER  - 


TY  - JOUR
AB  - The cycad genus Encephalartos is restricted to Africa and is threatened with extinction in most of its range. Total DNA was extracted from 51, i.e., 78 %, of the described species of Encephalartos. The accessions were sampled from the furthest western occurrence of the genus in Nigeria, via Sudan and Uganda, to southern South Africa. The sequences of nuclear ribosomal internal transcribed spacer regions 1 and 2 (ITS 1&2), the chloroplast encoded rbcL gene, and ISSR genomic fingerprinting were employed to resolve the molecular history and the relationships within the genus. Sequence alignment, as well as ISSR fingerprinting, data show low genetic variation among all analysed accessions, indicating diversification within the Pliocene/Pleistocene. ITS 1&2 data agree well with morphological and geographical characters and resolved three major genetic clusters with overlapping distribution ranges in eastern South Africa. This area, that contains the largest diversity of genotypes of Encephalartos, may have served as a Pliocene/Pleistocene refugium.
AD  - Institut für Pharmazie und Molekulare Biotechnologie, Universität Heidelberg, Im Neuenheimer Feld 364, 69120 Heidelberg, Germany. M4 - Citavi
AU  - Treutlein, J.
AU  - Vorster, P.
AU  - Wink, M.
DO  - 10.1055/s-2004-830478
IS  - 1
KW  - Zamiaceae
LA  - eng
N1  - Journal Article Journal Article Journal Article Journal Article
PY  - 2005
SN  - 1435-8603
SP  - 79-90
ST  - Molecular relationships in Encephalartos (Zamiaceae, Cycadales) based on nucleotide sequences of nuclear ITS 1&2, rbcL, and genomic Issr fingerprinting
T2  - Plant Biology
TI  - Molecular relationships in Encephalartos (Zamiaceae, Cycadales) based on nucleotide sequences of nuclear ITS 1&2, rbcL, and genomic Issr fingerprinting
VL  - 7
ID  - 3710
ER  - 


TY  - JOUR
AU  - Nolasco-Soto, Janet
AU  - González-Astorga, Jorge
AU  - Nicolalde-Morejón, Fernando
AU  - Vergara-Silva, Francisco
AU  - los Monteros, Alejandro Espinosa
AU  - Medina-Villarreal, Anwar
DO  - 10.1007/s00606-014-1101-1 M4 - Citavi
IS  - 2
KW  - Zamiaceae
N1  - PII: 1101 PII: 1101 PII: 1101 PII: 1101
PY  - 2015
SN  - 0378-2697
SP  - 623-637
ST  - Phylogeography and demographic history of Zamia paucijuga Wieland (Zamiaceae), a cycad species from the Mexican Pacific slope
T2  - Plant Systematics and Evolution
TI  - Phylogeography and demographic history of Zamia paucijuga Wieland (Zamiaceae), a cycad species from the Mexican Pacific slope
VL  - 301
ID  - 653
ER  - 


TY  - JOUR
AU  - Nicolalde-Morejón, Fernando
AU  - Vovides, Andrew P.
AU  - Stevenson, Dennis W.
IS  - 4
KW  - Zamiaceae
PY  - 2009
SN  - 0007-196X
SP  - 301-335
ST  - Taxonomic revision of Zamia in Mega-Mexico
T2  - Brittonia
TI  - Taxonomic revision of Zamia in Mega-Mexico
VL  - 61
ID  - 2619
ER  - 


TY  - JOUR
AU  - González, Dolores
AU  - Vovides, Andrew P.
AU  - Bárcenas, Cristina
IS  - 2
KW  - Zamiaceae
PY  - 2008
SN  - 03636445
SP  - 229-236
ST  - Phylogenetic Relationships of the Neotropical Genus Dioon (Cycadales, Zamiaceae) Based on Nuclear and Chloroplast DNA Sequence Data
T2  - Systematic Botany
TI  - Phylogenetic Relationships of the Neotropical Genus Dioon (Cycadales, Zamiaceae) Based on Nuclear and Chloroplast DNA Sequence Data
VL  - 33
ID  - 2816
ER  - 


TY  - JOUR
AD  - Departamento de Botânica; Instituto de Biociências; Universidade Estadual Paulista; C. Postal 199 13506900 Rio Claro SP Brazil M4 - Citavi
AU  - Oriani, Aline
AU  - Scatena, Vera L.
DO  - 10.1111/boj.12152 Y3 - 10.02.2019
IS  - 1
KW  - Xyridaceae
PY  - 2014
SP  - 144-154
ST  - Ovule, fruit and seed development in Abolboda (Xyridaceae, Poales): implications for taxonomy and phylogeny
T2  - Botanical Journal of the Linnean Society
TI  - Ovule, fruit and seed development in Abolboda (Xyridaceae, Poales): implications for taxonomy and phylogeny
VL  - 175
ID  - 1011
ER  - 


TY  - JOUR
AU  - Campbell, Lisa M.
DO  - 10.1007/s12229-012-9110-7 M4 - Citavi
IS  - 4
KW  - Xyridaceae
N1  - PII: 9110 PII: 9110 PII: 9110 PII: 9110
PY  - 2012
SN  - 0006-8101
SP  - 428-439
ST  - Pollen Morphology of Xyridaceae Systematic (Poales) and its Potential
T2  - The Botanical Review
TI  - Pollen Morphology of Xyridaceae Systematic (Poales) and its Potential
VL  - 78
ID  - 1487
ER  - 


TY  - JOUR
AD  - State Key Laboratory of Systematic and Evolutionary Botany, Institute of Botany; Chinese Academy of Sciences; Beijing 100093 China University of Chinese Academy of Sciences; Beijing 100049 China M4 - Citavi
AU  - Wei, Ran
AU  - Zhang, Xian-Chun
DO  - 10.1111/jse.12075
IS  - 4
KW  - Woodsiaceae
PY  - 2014
SN  - 16744918
SP  - 450-457
ST  - Rediscovery of Cystoathyrium chinense Ching (Cystopteridaceae): Phylogenetic placement of the critically endangered fern species endemic to China
T2  - Journal of Systematics and Evolution
TI  - Rediscovery of Cystoathyrium chinense Ching (Cystopteridaceae): Phylogenetic placement of the critically endangered fern species endemic to China
VL  - 52
ID  - 1126
ER  - 


TY  - JOUR
AB  - The cliff fern family Woodsiaceae has experienced frequent taxonomic changes at the familial and generic ranks since its establishment. The bulk of its species were placed in Woodsia, while Cheilanthopsis, Hymenocystis, Physematium, and Protowoodsia are segregates recognized by some authors. Phylogenetic relationships among the genera of Woodsiaceae remain unclear because of the extreme morphological diversity and inadequate taxon sampling in phylogenetic studies to date. In this study, we carry out comprehensive phylogenetic analyses of Woodsiaceae using molecular evidence from four chloroplast DNA markers (atpA, matK, rbcL and trnL-F) and covering over half the currently recognized species. Our results show three main clades in Woodsiaceae corresponding to Physematium (clade I), Cheilanthopsis-Protowoodsia (clade II) and Woodsia s.s. (clade III). In the interest of preserving monophyly and taxonomic stability, a broadly defined Woodsia including the other segregates is proposed, which is characterized by the distinctive indument and inferior indusia. Therefore, we present a new subgeneric classification of the redefined Woodsia based on phylogenetic and ancestral state reconstructions to better reflect the morphological variation, geographic distribution pattern, and evolutionary history of the genus. Our analyses of the cytological character evolution support multiple aneuploidy events that have resulted in the reduction of chromosome base number from 41 to 33, 37, 38, 39 and 40 during the evolutionary history of the cliff ferns.
AD  - State Key Laboratory of Systematic and Evolutionary Botany, Institute of Botany, Chinese Academy of Sciences, Beijing, 100093, China; University of the Chinese Academy of Sciences, Beijing, 100049, China. State Key Laboratory of Systematic and Evolutionary Botany, Institute of Botany, Chinese Academy of Sciences, Beijing, 100093, China. State Key Laboratory of Systematic and Evolutionary Botany, Institute of Botany, Chinese Academy of Sciences, Beijing, 100093, China. State Key Laboratory of Systematic and Evolutionary Botany, Institute of Botany, Chinese Academy of Sciences, Beijing, 100093, China. M4 - Citavi
AU  - Shao, Yizhen
AU  - Wei, Ran
AU  - Zhang, Xianchun
AU  - Xiang, Qiaoping
DO  - 10.1371/journal.pone.0136318
IS  - 9
KW  - Woodsiaceae
LA  - eng
N1  - Journal Article Research Support, Non-U.S. Gov't Journal Article Research Support, Non-U.S. Gov't Journal Article Research Support, Non-U.S. Gov't Journal Article Research Support, Non-U.S. Gov't
PY  - 2015
SP  - e0136318
ST  - Molecular Phylogeny of the Cliff Ferns (Woodsiaceae: Polypodiales) with a proposed Infrageneric Classification
T2  - PloS one
TI  - Molecular Phylogeny of the Cliff Ferns (Woodsiaceae: Polypodiales) with a proposed Infrageneric Classification
UR  - https://www.ncbi.nlm.nih.gov/pmc/articles/PMC4562699
VL  - 10
ID  - 713
ER  - 


TY  - JOUR
AU  - Chunxiang, Li
AU  - Shugang, Lu
AU  - Xiaoyan, Sun
AU  - Qun, Yang
DO  - 10.1640/0002-8444-101.3.142 M4 - Citavi
IS  - 3
KW  - Woodsiaceae
PY  - 2011
SN  - 0002-8444
SP  - 142-155
ST  - Phylogenetic Positions of the Enigmatic Asiatic Fern Genera Diplaziopsis and Rhachidosorus from Analyses of Four Plastid Genes
T2  - American Fern Journal
TI  - Phylogenetic Positions of the Enigmatic Asiatic Fern Genera Diplaziopsis and Rhachidosorus from Analyses of Four Plastid Genes
VL  - 101
ID  - 1855
ER  - 


TY  - JOUR
AB  - Phylogenetic analyses of representative species from the five genera of Winteraceae (Drimys, Pseudowintera, Takhtajania, Tasmannia, and Zygogynum s.l.) were performed using ITS nuclear sequences and a combined data-set of ITS+psbA-trnH+rpS16 sequences (sampling of 30 and 15 species, respectively). Indel informativity using simple gap coding or gaps as a fifth character was examined in both data-sets. Parsimony and Bayesian analyses support the monophyly of Drimys, Tasmannia, and Zygogynum s.l., but do not support the monophyly of Belliolum, Zygogynum s.s., and Bubbia. Within Drimys, the combined data-set recovers two subclades. Divergence time estimates suggest that the splitting between Drimys and its sister clade (Pseudowintera+Zygogynum s.l.) occurred around the end of the Cretaceous; in contrast, the divergence between the two subclades within Drimys is more recent (15.5-18.5MY) and coincides in time with the Andean uplift. Estimates suggest that the earliest divergences within Winteraceae could have predated the first events of Gondwana fragmentation.
AU  - Marquínez, Xavier
AU  - Lohmann, Lúcia Garces
AU  - Salatino, Maria L. Faria
AU  - Salatino, Antonio
AU  - González, Favio
DO  - 10.1016/j.ympev.2009.07.001
IS  - 2
KW  - Winteraceae
LA  - eng
N1  - Journal Article Research Support, Non-U.S. Gov't Journal Article Research Support, Non-U.S. Gov't Journal Article Research Support, Non-U.S. Gov't Journal Article Research Support, Non-U.S. Gov't
PY  - 2009
SN  - 1055-7903
SP  - 435-449
ST  - Generic relationships and dating of lineages in Winteraceae based on nuclear (Its) and plastid (rpS16 and psbA-trnH) sequence data
T2  - Molecular phylogenetics and evolution
TI  - Generic relationships and dating of lineages in Winteraceae based on nuclear (Its) and plastid (rpS16 and psbA-trnH) sequence data
VL  - 53
ID  - 2594
ER  - 


TY  - JOUR
AU  - Sytsma, Kenneth J.
AU  - Litt, Amy
AU  - Zjhra, Michelle L.
AU  - Chris Pires, J.
AU  - Nepokroeff, Molly
AU  - Conti, Elena
AU  - Walker, Jay
AU  - Wilson, Peter G.
DO  - 10.1086/421066 Y3 - 21.02.2019 M4 - Citavi
IS  - S4
KW  - Vochysiaceae
PY  - 2004
SN  - 1058-5893
SP  - S85-S105
ST  - Clades, Clocks, and Continents: Historical and Biogeographical Analysis of Myrtaceae, Vochysiaceae, and Relatives in the Southern Hemisphere
T2  - International Journal of Plant Sciences
TI  - Clades, Clocks, and Continents: Historical and Biogeographical Analysis of Myrtaceae, Vochysiaceae, and Relatives in the Southern Hemisphere
VL  - 165
ID  - 3909
ER  - 


TY  - JOUR
AB  - Vitales is well-known for including the economically important fruit crop, the wine grape (Vitis vinifera). However, the position of the Vitales in the higher eudicots has been a subject of much debate. It has been variously reported to be sister to the Saxifragales and together as sister to the rest of rosids, or sister to the fabids-malvids clade, or sister to the Santalales, or sister to the fabids-malvids-Saxifragales clade. However, in all of these scenarios, the support values for the relationship of Vitales on the phylogenies were only low to moderate. Additionally, all previous studies sampled only Vitis vinifera as the representative of the Vitales. We herein expanded the sampling of the Vitales to include representatives of all major clades of the order, as well as representatives of other key lineages including Saxifragales, Dilleniales, and Santalales. Extensive likelihood and Bayesian analyses were conducted to test the position of Vitales, using different numbers of genes, a variety of partitioning strategies, and both nucleotide and amino acid sequences. With the expanded sampling strategy, almost all analyses supported the relationship of Vitales as sister to Saxifragales. This relationship was supported in a 72-gene data set with a bootstrap value of 91%, the highest support value reported to date. Based on this topology, we discuss possible morphological synapomorphies shared between Vitales and Saxifragales. Furthermore, a hypothesis of reticulate evolution was postulated as an explanation for the incongruence of Vitales' position when comparing plastid and nuclear gene phylogenies.
AD  - Department of Botany, National Museum of Natural History, MRC 166, Smithsonian Institution, Washington, DC 20013-7012, USA. Department of Botany, National Museum of Natural History, MRC 166, Smithsonian Institution, Washington, DC 20013-7012, USA. Electronic address:. Department of Botany, National Museum of Natural History, MRC 166, Smithsonian Institution, Washington, DC 20013-7012, USA. Electronic address:. M4 - Citavi
AU  - Zhang, Ning
AU  - Wen, Jun
AU  - Zimmer, Elizabeth A.
DO  - 10.1016/j.ympev.2016.04.034
KW  - Vitaceae
LA  - eng
N1  - Journal Article Journal Article Journal Article Journal Article
PY  - 2016
SN  - 1055-7903
SP  - 216-223
ST  - Another look at the phylogenetic position of the grape order Vitales: Chloroplast phylogenomics with an expanded sampling of key lineages
T2  - Molecular phylogenetics and evolution
TI  - Another look at the phylogenetic position of the grape order Vitales: Chloroplast phylogenomics with an expanded sampling of key lineages
VL  - 101
ID  - 419
ER  - 


TY  - JOUR
AD  - Department of Botany; National Museum of Natural History; MRC166, Smithsonian Institution Washington DC 20013-7012 USA M4 - Citavi
AU  - Zhang, Ning
AU  - Wen, Jun
AU  - Zimmer, Elizabeth A.
DO  - 10.1111/jse.12138
IS  - 5
KW  - Vitaceae
PY  - 2015
SN  - 16744918
SP  - 469-476
ST  - Expression patterns of AP1 , FUL , FT and Leafy orthologs in Vitaceae support the homology of tendrils and inflorescences throughout the grape family
T2  - Journal of Systematics and Evolution
TI  - Expression patterns of AP1 , FUL , FT and Leafy orthologs in Vitaceae support the homology of tendrils and inflorescences throughout the grape family
VL  - 53
ID  - 799
ER  - 


TY  - JOUR
AB  - Wild grapes are woody climbers, found mostly in temperate regions of the northern hemisphere, comprising the genus Vitis. Despite its importance, the evolutionary history of Vitis is still contentious. Past studies have led to conflicting hypotheses about the phylogeny, speciation events, and biogeographic history of the genus. Here we investigate the evolutionary history of Vitis using data from four chloroplast spacers (trnH-psbA, trnK-rps16, trnF-nahJ, and rpl32-trnL) and the nuclear gene RPB2-I, and we explore mechanisms that could have shaped the observed distribution of current species. Maximum likelihood and Bayesian analyses provided similar results, strongly supporting the presence of two subgenera and suggesting a species clustering within subgenus Vitis that mainly mirrors the disjunction between the Old and New World. Vitis vinifera subsp. sylvestris was found to be sister to the Asian species while three major clades were found in the American species. A network approach confirmed the main geographic groups and highlighted different chloroplast haplotype patterns between Asian and American species. Molecular dating analysis provided the time boundaries to discuss our results. Our study shows wild grape diversification to be a continuous and complex process that concerned the Tertiary as well as the Quaternary, most likely involving both geographical and climatic forces. Local variations in extent and timing of these forces were discussed based on observed differences between groups. In the context of the Tertiary-Quaternary debate, we provide evidence in favor of the "continuous hypothesis" to explain present diversity. Finally, two directions for future research are highlighted: (i) was the earliest grape American or Asian? and (ii) are all modern grape species real?
AD  - Botanical Garden, Department of Biology, Università degli Studi di Milano, Via Celoria 26, 20133 Milan, Italy. M4 - Citavi
AU  - Zecca, Giovanni
AU  - Abbott, Richard J.
AU  - Sun, Wei-Bang
AU  - Spada, Alberto
AU  - Sala, Francesco
AU  - Grassi, Fabrizio
DO  - 10.1016/j.ympev.2011.11.015
IS  - 2
KW  - Vitaceae
LA  - eng
N1  - Journal Article Research Support, Non-U.S. Gov't Journal Article Research Support, Non-U.S. Gov't Journal Article Research Support, Non-U.S. Gov't Journal Article Research Support, Non-U.S. Gov't
PY  - 2012
SN  - 1055-7903
SP  - 736-747
ST  - The timing and the mode of evolution of wild grapes (Vitis)
T2  - Molecular phylogenetics and evolution
TI  - The timing and the mode of evolution of wild grapes (Vitis)
VL  - 62
ID  - 1783
ER  - 


TY  - JOUR
AD  - Department of Botany, MRC 166, National Museum of Natural History, Smithsonian Institution, Washington, DC 20013-7012, USA. Laboratory of Systematic and Evolutionary Botany, Institute of Botany, Chinese Academy of Sciences, Nanxincun 20, Xiangshan, Beijing 100093, P. R. China. Laboratory of Biodiversity and Biogeography, Kunming Institute of Botany, Chinese Academy of Sciences, Kunming, Yunnan 650204, P. R. China. Graduate School of Chinese Academy of Sciences, Beijing 100039, P. R. China. Department of Biological Science, School of Science, Osaka Prefecture University, Sakai, Osaka 599-8531, Japan. M4 - Citavi
AU  - Wen, Jun
AU  - Nie, Ze-long
AU  - Soejima, Akiko
AU  - Meng, Ying
DO  - 10.1139/B07-071
IS  - 8
KW  - Vitaceae
PY  - 2007
SN  - 0008-4026
SP  - 731-745
ST  - Phylogeny of Vitaceae based on the nuclear Gai1 gene sequencesThis article is one of a selection of papers presented at the symposium on Vitis at the XVII International Botanical Congress held in Vienna, Austria, in 2005
T2  - Canadian Journal of Botany
TI  - Phylogeny of Vitaceae based on the nuclear Gai1 gene sequencesThis article is one of a selection of papers presented at the symposium on Vitis at the XVII International Botanical Congress held in Vienna, Austria, in 2005
VL  - 85
ID  - 3302
ER  - 


TY  - JOUR
AB  - BACKGROUND Grapes are one of the most economically important fruit crops. There are about 60 species in the genus Vitis. The phylogenetic relationships among these species are of keen interest for the conservation and use of this germplasm. We selected 309 accessions from 48 Vitis species,varieties, and outgroups, examined ~11 kb (~3.4 Mb total) of aligned nuclear DNA sequences from 27 unlinked genes in a phylogenetic context, and estimated divergence times based on fossil calibrations. RESULTS Vitis formed a strongly supported clade. There was substantial support for species and less for the higher-level groupings (series). As estimated from extant taxa, the crown age of Vitis was 28 Ma and the divergence of subgenera (Vitis and Muscadinia) occurred at ~18 Ma. Higher clades in subgenus Vitis diverged 16 - 5 Ma with overlapping confidence intervals, and ongoing divergence formed extant species at 12 - 1.3 Ma. Several species had species-specific SNPs. NeighborNet analysis showed extensive reticulation at the core of subgenus Vitis representing the deeper nodes, with extensive reticulation radiating outward. Fitch Parsimony identified North America as the origin of the most recent common ancestor of extant Vitis species. CONCLUSIONS Phylogenetic patterns suggested origination of the genus in North America, fragmentation of an ancestral range during the Miocene, formation of extant species in the late Miocene-Pleistocene, and differentiation of species in the context of Pliocene-Quaternary tectonic and climatic change. Nuclear SNPs effectively resolved relationships at and below the species level in grapes and rectified several misclassifications of accessions in the repositories. Our results challenge current higher-level classifications, reveal the abundance of genetic diversity in the genus that is potentially available for crop improvement, and provide a valuable resource for species delineation, germplasm conservation and use.
AD  - College of Horticulture, Northwest A&F University, Yangling, Shaanxi 712100, People's Republic of China. M4 - Citavi
AU  - Wan, Yizhen
AU  - Schwaninger, Heidi R.
AU  - Baldo, Angela M.
AU  - Labate, Joanne A.
AU  - Zhong, Gan-Yuan
AU  - Simon, Charles J.
DO  - 10.1186/1471-2148-13-141
KW  - Vitaceae
LA  - eng
N1  - Journal Article Research Support, Non-U.S. Gov't Research Support, U.S. Gov't, Non-P.H.S. Journal Article Research Support, Non-U.S. Gov't Research Support, U.S. Gov't, Non-P.H.S. Journal Article Research Support, Non-U.S. Gov't Research Support, U.S. Gov't, Non-P.H.S. Journal Article Research Support, Non-U.S. Gov't Research Support, U.S. Gov't, Non-P.H.S.
PY  - 2013
SP  - 141
ST  - A phylogenetic analysis of the grape genus (Vitis L.) reveals broad reticulation and concurrent diversification during neogene and quaternary climate change
T2  - BMC evolutionary biology
TI  - A phylogenetic analysis of the grape genus (Vitis L.) reveals broad reticulation and concurrent diversification during neogene and quaternary climate change
UR  - https://www.ncbi.nlm.nih.gov/pmc/articles/PMC3750556
VL  - 13
ID  - 1421
ER  - 


TY  - JOUR
AB  - UNLABELLED PREMISE OF THE STUDY This work represents the first molecular phylogeny of the economically important genus Vitis, an important genetic resource for breeding in grapevine, Vitis vinifera. • METHODS A molecular phylogeny of Vitis using a combined data set of three noncoding regions of the plastid DNA genome was constructed from 47 accessions covering 30 species of Vitis. The data for the trnL-F marker were combined with previously published data across the Vitaceae. • KEY RESULTS The molecular phylogeny demonstrated monophyly of the genus Vitis. Based on the combined analysis of three genes, Vitis is split into three clades that mirror the continental distribution of these accessions. The diversity is highest in the Asian clade, but the general genetic distances across taxa from different continents are relatively small. • CONCLUSIONS The findings support a relatively recent and intense gene flow between East Asia and North America and the possible impact of hybridization on the evolution of the genus Vitis. Taxon identity in important stock collections should be screened carefully because roughly 10% of the accessions analyzed in the present study had been misidentified.
AD  - Botanical Institute, Molecular Cell Biology, Karlsruhe Institute of Technology, Kaiserstrasse 2 76128 Karlsruhe, Germany. M4 - Citavi
AU  - Tröndle, Dorothee
AU  - Schröder, Stephan
AU  - Kassemeyer, Hanns-Heinz
AU  - Kiefer, Christiane
AU  - Koch, Marcus A.
AU  - Nick, Peter
DO  - 10.3732/ajb.0900218
IS  - 7
KW  - Vitaceae
LA  - eng
N1  - Journal Article Journal Article Journal Article Journal Article
PY  - 2010
SN  - 0002-9122
SP  - 1168-1178
ST  - Molecular phylogeny of the genus Vitis (Vitaceae) based on plastid markers
T2  - American journal of botany
TI  - Molecular phylogeny of the genus Vitis (Vitaceae) based on plastid markers
VL  - 97
ID  - 2373
ER  - 


TY  - JOUR
AU  - Trias-Blasi, Anna
AU  - Chayamarit, Kongkanda
AU  - Teerawatananon, Atchara
AU  - Parnell, John A. N.
DO  - 10.11646/phytotaxa.159.2.3 M4 - Citavi
IS  - 2
KW  - Vitaceae
PY  - 2014
SN  - 1179-3163
SP  - 95
ST  - A taxonomic revision of Pterisanthes (Vitaceae) in Thailand and a new Thai record for Pterisanthes cissioides
T2  - Phytotaxa
TI  - A taxonomic revision of Pterisanthes (Vitaceae) in Thailand and a new Thai record for Pterisanthes cissioides
VL  - 159
ID  - 1104
ER  - 


TY  - JOUR
AU  - Soejima, Akiko
AU  - Wen, Jun
IS  - 2
KW  - Vitaceae
PY  - 2006
SN  - 0002-9122
SP  - 278-287
ST  - Phylogenetic Analysis of the Grape Family (Vitaceae) based on three Chloroplast Markers
T2  - American journal of botany
TI  - Phylogenetic Analysis of the Grape Family (Vitaceae) based on three Chloroplast Markers
VL  - 93
ID  - 3496
ER  - 


TY  - JOUR
AU  - Rossetto, Maurizio
AU  - Crayn, Darren M.
AU  - Jackes, Betsy R.
AU  - Porter, Carolyn
DO  - 10.1139/B07-022 M4 - Citavi
IS  - 8
KW  - Vitaceae
PY  - 2007
SN  - 0008-4026
SP  - 722-730
ST  - An updated estimate of intergeneric phylogenetic relationships in the Australian Vitaceae
T2  - Canadian Journal of Botany
TI  - An updated estimate of intergeneric phylogenetic relationships in the Australian Vitaceae
VL  - 85
ID  - 3246
ER  - 


TY  - JOUR
AU  - Rodrigues, Jacqueline Gomes
AU  - Lombardi, Julio Antonio
AU  - Lovato, Maria Bernadete
DO  - 10.12705/632.33 M4 - Citavi
IS  - 2
KW  - Vitaceae
PY  - 2014
SN  - 00400262
SP  - 287-298
ST  - Phylogeny of <I>Cissus</I> (Vitaceae) focusing on South American species
T2  - Taxon
TI  - Phylogeny of <I>Cissus</I> (Vitaceae) focusing on South American species
VL  - 63
ID  - 1039
ER  - 


TY  - JOUR
AU  - Ren, Hui
AU  - Lu, Li-Min
AU  - Soejima, Akiko
AU  - Luke, Quentin
AU  - Zhang, Dian-Xiang
AU  - Chen, Zhi‐Duan
AU  - Wen, Jun
IS  - 3
KW  - Vitaceae
PY  - 2011
SN  - 00400262
SP  - 629-637
ST  - Phylogenetic analysis of the grape family (Vitaceae) based on the noncoding plastid trnC-petN, trnH-psbA, and trnL-F sequences
T2  - Taxon
TI  - Phylogenetic analysis of the grape family (Vitaceae) based on the noncoding plastid trnC-petN, trnH-psbA, and trnL-F sequences
VL  - 60
ID  - 2004
ER  - 


TY  - JOUR
AU  - Pelser, Pieter B.
AU  - Nickrent, Daniel L.
AU  - Barcelona, Julie F.
DO  - 10.12705/654.4 M4 - Citavi
IS  - 4
KW  - Vitaceae
PY  - 2016
SN  - 00400262
SP  - 739-758
ST  - Untangling a vine and its parasite: Host specificity of Philippine Rafflesia (Rafflesiaceae)
T2  - Taxon
TI  - Untangling a vine and its parasite: Host specificity of Philippine Rafflesia (Rafflesiaceae)
VL  - 65
ID  - 325
ER  - 


TY  - JOUR
AB  - BACKGROUND The Ampelopsis clade (Ampelopsis and its close allies) of the grape family Vitaceae contains ca. 43 species disjunctly distributed in Asia, Europe, North America, South America, Africa, and Australia, and is a rare example to study both the Northern and the Southern Hemisphere intercontinental disjunctions. We reconstruct the temporal and spatial diversification of the Ampelopsis clade to explore the evolutionary processes that have resulted in their intercontinental disjunctions in six continents. RESULTS The Bayesian molecular clock dating and the likelihood ancestral area analyses suggest that the Ampelopsis clade most likely originated in North America with its crown group dated at 41.2 Ma (95% HPD 23.4-61.0 Ma) in the middle Eocene. Two independent Laurasian migrations into Eurasia are inferred to have occurred in the early Miocene via the North Atlantic land bridges. The ancestor of the Southern Hemisphere lineage migrated from North America to South America in the early Oligocene. The Gondwanan-like pattern of intercontinental disjunction is best explained by two long-distance dispersals: once from South America to Africa estimated at 30.5 Ma (95% HPD 16.9-45.9 Ma), and the other from South America to Australia dated to 19.2 Ma (95% HPD 6.7-22.3 Ma). CONCLUSIONS The global disjunctions in the Ampelopsis clade are best explained by a diversification model of North American origin, two Laurasian migrations, one migration into South America, and two post-Gondwanan long-distance dispersals. These findings highlight the importance of both vicariance and long distance dispersal in shaping intercontinental disjunctions of flowering plants.
AD  - Key Laboratory of Biodiversity and Biogeography, Kunming Institute of Botany, Chinese Academy of Sciences, Kunming, Yunnan 650204, China. M4 - Citavi
AU  - Nie, Ze-long
AU  - Sun, Hang
AU  - Manchester, Steven R.
AU  - Meng, Ying
AU  - Luke, Quentin
AU  - Wen, Jun
DO  - 10.1186/1471-2148-12-17
KW  - Vitaceae
LA  - eng
N1  - Journal Article Research Support, Non-U.S. Gov't Research Support, U.S. Gov't, Non-P.H.S. Journal Article Research Support, Non-U.S. Gov't Research Support, U.S. Gov't, Non-P.H.S. Journal Article Research Support, Non-U.S. Gov't Research Support, U.S. Gov't, Non-P.H.S. Journal Article Research Support, Non-U.S. Gov't Research Support, U.S. Gov't, Non-P.H.S.
PY  - 2012
SP  - 17
ST  - Evolution of the intercontinental disjunctions in six continents in the Ampelopsis clade of the grape family (Vitaceae)
T2  - BMC evolutionary biology
TI  - Evolution of the intercontinental disjunctions in six continents in the Ampelopsis clade of the grape family (Vitaceae)
UR  - https://www.ncbi.nlm.nih.gov/pmc/articles/PMC3299610
VL  - 12
ID  - 1651
ER  - 


TY  - JOUR
AB  - UNLABELLED PREMISE OF THE STUDY Parthenocissus is a genus of the grape family Vitaceae and has a disjunct distribution in Asia and North America with members in both tropical and temperate regions. The monophyly of Parthenocissus has not yet been tested, and the species relationships and the evolution of its intercontinental disjunction have not been investigated with extensive sampling and molecular phylogenetic methods. • METHODS Plastid (trnL-F, rps16, and atpB-rbcL) and nuclear GAI1 sequences of 56 accessions representing all 12 Parthenocissus species were analyzed with parsimony, likelihood, and Bayesian inference. Divergence times of disjunct lineages were estimated with relaxed Bayesian dating. Evolution of the leaflet number was assessed by tracing this character onto Bayesian trees using the Trace Character Over Trees option in the program Mesquite. • KEY RESULTS Parthenocissus is monophyletic and sister to the newly described segregate genus Yua. Two major clades within Parthenocissus are recognizable corresponding to their distribution in Asia and North America. The disjunction between the two continents is estimated to be at 21.64 (95% higher posterior densities 10.23-34.89) million years ago. • CONCLUSIONS Parthenocissus is likely to have derived from the Eocene boreotropical element. Its current Asian-North American disjunction is dated to the early Miocene, congruent with fossil and paleoclimatic evidence. The tropical species is nested within the temperate clade and is inferred to have dispersed from the adjacent temperate regions. Parthenocissus and Yua are best treated as distinct genera. Leaflet number in this genus has a complex history and cannot be used as a character for infrageneric classification.
AD  - Key Laboratory of Biodiversity and Biogeography, Kunming Institute of Botany, Chinese Academy of Sciences, Kunming, Yunnan 650204, P. R. China. M4 - Citavi
AU  - Nie, Ze-long
AU  - Sun, Hang
AU  - Chen, Zhi‐Duan
AU  - Meng, Ying
AU  - Manchester, Steven R.
AU  - Wen, Jun
DO  - 10.3732/ajb.1000085
IS  - 8
KW  - Vitaceae
LA  - eng
N1  - Journal Article Journal Article Journal Article Journal Article
PY  - 2010
SN  - 0002-9122
SP  - 1342-1353
ST  - Molecular phylogeny and biogeographic diversification of Parthenocissus (Vitaceae) disjunct between Asia and North America
T2  - American journal of botany
TI  - Molecular phylogeny and biogeographic diversification of Parthenocissus (Vitaceae) disjunct between Asia and North America
VL  - 97
ID  - 2297
ER  - 


TY  - JOUR
AU  - Molina, Jeanmaire E.
AU  - Wen, Jun
AU  - Struwe, Lena
KW  - Vitaceae
PY  - 2013
SP  - 354-376
ST  - Systematics and biogeography of the non-viny Grape relative Leea (Vitaceae)
T2  - Botanical Journal of the Linnean Society
TI  - Systematics and biogeography of the non-viny Grape relative Leea (Vitaceae)
VL  - 171
ID  - 1319
ER  - 


TY  - JOUR
AU  - Lu, Limin
AU  - Wen, Jun
AU  - Chen, Zhi‐Duan
KW  - Vitaceae
PY  - 2012
SP  - 43-63
ST  - A combined morphological and molecular phylogenetic analysis of Parthenocissus (Vitaceae) and taxonomic implications
T2  - Botanical Journal of the Linnean Society
TI  - A combined morphological and molecular phylogenetic analysis of Parthenocissus (Vitaceae) and taxonomic implications
VL  - 168
ID  - 1618
ER  - 


TY  - JOUR
AB  - Cayratia consists of ca. 60 species primarily distributed in the tropical and subtropical regions of Asia, Australia, and Africa. It is an excellent candidate for exploring the evolution of intercontinental disjunct distributions in the Old World. Previous phylogenetic work of Vitaceae with a few species of Cayratia sampled showed that Cayratia was not monophyletic and was closely related to Cyphostemma and Tetrastigma. We herein expanded taxon sampling of Cayratia (25/60 species) with its allied genera Cyphostemma (39/150 species), Tetrastigma (27/95 species), and other related genera from Vitaceae represented, employing five plastid markers (atpB-rbcL, rps16, trnC-petN, trnH-psbA, and trnL-F), to investigate the phylogeny, character evolution and biogeography of Cayratia. The phylogenetic analyses have confirmed the monophyly of the Cayratia-Cyphostemma-Tetrastigma (CCT) clade and resolved Cayratia into three lineages: the African Cayratia clade, subg. Cayratia, and subg. Discypharia. The African Cayratia was supported as the first diverging lineage within the CCT clade and Tetrastigma is resolved as sister to subg. Discypharia. Character optimizations suggest that the presence/absence of a membrane enclosing the ventral infolds in seeds is an important character for the taxonomy of Cayratia. The presence of bracts on the lower part of the inflorescence axis is inferred to have arisen only once in Cayratia, but this character evolved several times in Tetrastigma. Both the branching pattern of tendrils and the leaf architecture are suggested as important infrageneric characters, but should be used cautiously because some states evolved multiple times. Ancestral area reconstruction and molecular dating suggest that the CCT clade originated from continental Africa in the late Cretaceous, and it then reached Asia twice independently in the late Cretaceous and late Oligocene, respectively. Several dispersals are inferred from Asia to Australia since the Eocene.
AD  - State Key Laboratory of Systematic and Evolutionary Botany, Institute of Botany, Chinese Academy of Sciences, Beijing 100093, China. M4 - Citavi
AU  - Lu, Limin
AU  - Wang, Wei
AU  - Chen, Zhi‐Duan
AU  - Wen, Jun
DO  - 10.1016/j.ympev.2013.04.023
IS  - 3
KW  - Vitaceae
LA  - eng
N1  - Journal Article Research Support, U.S. Gov't, Non-P.H.S. Journal Article Research Support, U.S. Gov't, Non-P.H.S. Journal Article Research Support, U.S. Gov't, Non-P.H.S. Journal Article Research Support, U.S. Gov't, Non-P.H.S.
PY  - 2013
SN  - 1055-7903
SP  - 502-515
ST  - Phylogeny of the non-monophyletic Cayratia Juss. (Vitaceae) and implications for character evolution and biogeography
T2  - Molecular phylogenetics and evolution
TI  - Phylogeny of the non-monophyletic Cayratia Juss. (Vitaceae) and implications for character evolution and biogeography
VL  - 68
ID  - 1300
ER  - 


TY  - JOUR
AD  - State Key Laboratory of Systematic and Evolutionary Botany; Institute of Botany; Chinese Academy of Sciences; Beijing 100093 China Centro de Ciências do Mar; Universidade do Algarve; Gambelas Faro 8005-319 Portugal CSIRO National Research Collections; Australian National Herbarium; Canberra ACT 2601 Australia Department of Botany; National Museum of Natural History; MRC166, Smithsonian Institution; Washington DC 20013-7012 USA M4 - Citavi
AU  - Lu, Limin
AU  - Cox, Cymon J.
AU  - Mathews, Sarah
AU  - Wang, Wei
AU  - Wen, Jun
AU  - Chen, Zhi‐Duan
DO  - 10.1111/cla.12191
IS  - 1
KW  - Vitaceae
PY  - 2018
SN  - 07483007
SP  - 57-77
ST  - Optimal data partitioning, multispecies coalescent and Bayesian concordance analyses resolve early divergences of the grape family (Vitaceae)
T2  - Cladistics
TI  - Optimal data partitioning, multispecies coalescent and Bayesian concordance analyses resolve early divergences of the grape family (Vitaceae)
VL  - 34
ID  - 24
ER  - 


TY  - JOUR
AB  - Pantropical intercontinental disjunct distribution is a major biogeographic pattern in plants, and has been explained mainly by boreotropical migration via the North Atlantic land bridges (NALB) and transoceanic long-distance dispersal (LDD), and sometimes by vicariance. However, well-resolved phylogenies of pantropical clades are still relatively few. Cissus is the largest genus of the grape family Vitaceae and shows a pantropical intercontinental disjunction with its 300 species distributed in all major tropical regions. This study constructed the phylogenetic relationships and biogeographic diversification history of Cissus, employing five plastid markers (rps16, trnL-F, atpB-rbcL, trnH-psbA and trnC-petN). The results confirmed that Cissus polyphyletic, consisting of three main clades: the core Cissus, the Cissus striata complex, and the Australian-Neotropical disjunct Cissus antarctica -C. trianae clade. The latter two clades need to be removed from Cissus to maintain the monophyly of the genus. The core Cissus is inferred to have originated in Africa and is estimated to have diverged from its relatives in Vitaceae in the late Cretaceous. It diversified in Africa into several main lineages in the late Paleocene to the early Eocene, colonized Asia at least three times in the Miocene, and the Neotropics in the middle Eocene. The NALB seems the most plausible route for the core Cissus migration from Africa to the Neotropics in the middle Eocene. Three African-Asian and two Neotropical-Australian disjunctions in Cissus s.l. are estimated to have originated in the Miocene and may be best explained by LDD.
AD  - Key Laboratory of Horticultural Plant Biology, Ministry of Education, College of Horticulture and Forestry Science, Huazhong Agricultural University, Wuhan 430070, PR China. M4 - Citavi
AU  - Liu, Xiu-Qun
AU  - Ickert-Bond, Stefanie M.
AU  - Chen, Long-Qing
AU  - Wen, Jun
DO  - 10.1016/j.ympev.2012.09.003
IS  - 1
KW  - Vitaceae
LA  - eng
N1  - Journal Article Research Support, Non-U.S. Gov't Journal Article Research Support, Non-U.S. Gov't Journal Article Research Support, Non-U.S. Gov't Journal Article Research Support, Non-U.S. Gov't
PY  - 2013
SN  - 1055-7903
SP  - 43-53
ST  - Molecular phylogeny of Cissus L. of Vitaceae (the grape family) and evolution of its pantropical intercontinental disjunctions
T2  - Molecular phylogenetics and evolution
TI  - Molecular phylogeny of Cissus L. of Vitaceae (the grape family) and evolution of its pantropical intercontinental disjunctions
VL  - 66
ID  - 1295
ER  - 


TY  - JOUR
AB  - BACKGROUND The Vitaceae (grape) is an economically important family of angiosperms whose phylogenetic placement is currently unresolved. Recent phylogenetic analyses based on one to several genes have suggested several alternative placements of this family, including sister to Caryophyllales, asterids, Saxifragales, Dilleniaceae or to rest of rosids, though support for these different results has been weak. There has been a recent interest in using complete chloroplast genome sequences for resolving phylogenetic relationships among angiosperms. These studies have clarified relationships among several major lineages but they have also emphasized the importance of taxon sampling and the effects of different phylogenetic methods for obtaining accurate phylogenies. We sequenced the complete chloroplast genome of Vitis vinifera and used these data to assess relationships among 27 angiosperms, including nine taxa of rosids. RESULTS The Vitis vinifera chloroplast genome is 160,928 bp in length, including a pair of inverted repeats of 26,358 bp that are separated by small and large single copy regions of 19,065 bp and 89,147 bp, respectively. The gene content and order of Vitis is identical to many other unrearranged angiosperm chloroplast genomes, including tobacco. Phylogenetic analyses using maximum parsimony and maximum likelihood were performed on DNA sequences of 61 protein-coding genes for two datasets with 28 or 29 taxa, including eight or nine taxa from four of the seven currently recognized major clades of rosids. Parsimony and likelihood phylogenies of both data sets provide strong support for the placement of Vitaceae as sister to the remaining rosids. However, the position of the Myrtales and support for the monophyly of the eurosid I clade differs between the two data sets and the two methods of analysis. In parsimony analyses, the inclusion of Gossypium is necessary to obtain trees that support the monophyly of the eurosid I clade. However, maximum likelihood analyses place Cucumis as sister to the Myrtales and therefore do not support the monophyly of the eurosid I clade. CONCLUSION Phylogenies based on DNA sequences from complete chloroplast genome sequences provide strong support for the position of the Vitaceae as the earliest diverging lineage of rosids. Our phylogenetic analyses support recent assertions that inadequate taxon sampling and incorrect model specification for concatenated multi-gene data sets can mislead phylogenetic inferences when using whole chloroplast genomes for phylogeny reconstruction.
AD  - Section of Integrative Biology and Institute of Cellular and Molecular Biology, Patterson Laboratories 141, University of Texas, Austin, TX 78712, USA. M4 - Citavi
AU  - Jansen, Robert K.
AU  - Kaittanis, Charalambos
AU  - Saski, Christopher
AU  - Lee, Seung-Bum
AU  - Tomkins, Jeffrey
AU  - Alverson, Andrew J.
AU  - Daniell, Henry
DO  - 10.1186/1471-2148-6-32
KW  - Vitaceae
LA  - eng
N1  - Journal Article Research Support, N.I.H., Extramural Research Support, U.S. Gov't, Non-P.H.S. Journal Article Research Support, N.I.H., Extramural Research Support, U.S. Gov't, Non-P.H.S. Journal Article Research Support, N.I.H., Extramural Research Support, U.S. Gov't, Non-P.H.S. Journal Article Research Support, N.I.H., Extramural Research Support, U.S. Gov't, Non-P.H.S.
PY  - 2006
SP  - 32
ST  - Phylogenetic analyses of Vitis (Vitaceae) based on complete chloroplast genome sequences: effects of taxon sampling and phylogenetic methods on resolving relationships among rosids
T2  - BMC evolutionary biology
TI  - Phylogenetic analyses of Vitis (Vitaceae) based on complete chloroplast genome sequences: effects of taxon sampling and phylogenetic methods on resolving relationships among rosids
UR  - https://www.ncbi.nlm.nih.gov/pmc/articles/PMC1479384
VL  - 6
ID  - 3400
ER  - 


TY  - JOUR
AU  - Ingrouille, Martin J.
AU  - Chase, Mark W.
AU  - Fay, Michael F.
AU  - Bowman, Diane
AU  - van der Bank, Michelle
AU  - Bruijn, Anette D. E.
IS  - 421-432
KW  - Vitaceae
PY  - 2002
ST  - Systematics of Vitaceae from the viewpoint of plastid rbcL DNA sequence data
T2  - Botanical Journal of the Linnean Society
TI  - Systematics of Vitaceae from the viewpoint of plastid rbcL DNA sequence data
VL  - 138
ID  - 4123
ER  - 


TY  - JOUR
AU  - Chen, Ping-Ting
AU  - Chen, Long-Qing
AU  - Wen, Jun
IS  - 2
KW  - Vitaceae
PY  - 2011
SN  - 00400262
SP  - 499-512
ST  - The first phylogenetic analysis of Tetrastigma (Miq.) Planch., the host of Rafflesiaceae
T2  - Taxon
TI  - The first phylogenetic analysis of Tetrastigma (Miq.) Planch., the host of Rafflesiaceae
VL  - 60
ID  - 1851
ER  - 


TY  - JOUR
AU  - Nickrent, Daniel L.
AU  - García, Miguel Á
AU  - Martín, Maria P.
AU  - Mathiasen, Robert L.
IS  - 1
KW  - Viscaceae
PY  - 2004
SN  - 0002-9122
SP  - 125-138
ST  - A Phylogeny Of All Species Of Arceuthobium (Viscaceae) Using Nuclear And Chloroplast DNA Sequences
T2  - American journal of botany
TI  - A Phylogeny Of All Species Of Arceuthobium (Viscaceae) Using Nuclear And Chloroplast DNA Sequences
VL  - 91
ID  - 3850
ER  - 


TY  - JOUR
AU  - Molvray, Mia
AU  - Kores, Paul J.
AU  - Chase, Mark W.
IS  - 2
KW  - Viscaceae
PY  - 1999
SN  - 0002-9122
SP  - 249-260
ST  - Phylogenetic Relationships Within Korthalsella (Viscaceae) Based On Nuclear ITS And Plastid Trnl-f Sequence Data
T2  - American journal of botany
TI  - Phylogenetic Relationships Within Korthalsella (Viscaceae) Based On Nuclear ITS And Plastid Trnl-f Sequence Data
VL  - 86
ID  - 4422
ER  - 


TY  - JOUR
AU  - Yoo, Ki-Oug
AU  - Jang, Su-Kil
DO  - 10.1111/j.1759-6831.2010.00102.x M4 - Citavi
IS  - 6
KW  - Violaceae
PY  - 2010
SN  - 16744918
SP  - 474-481
ST  - Infrageneric relationships of Korean Viola based on eight chloroplast markers
T2  - Journal of Systematics and Evolution
TI  - Infrageneric relationships of Korean Viola based on eight chloroplast markers
VL  - 48
ID  - 2407
ER  - 


TY  - JOUR
AU  - Wahlert, Gregory A.
AU  - Marcussen, Thomas
AU  - Paula-Souza, Juliana
AU  - Feng, Min
AU  - Ballard, Harvey Eugene
DO  - 10.1600/036364414X678008 M4 - Citavi
IS  - 1
KW  - Violaceae
PY  - 2014
SN  - 03636445
SP  - 239-252
ST  - A Phylogeny of the Violaceae (Malpighiales) Inferred from Plastid DNA Sequences: Implications for Generic Diversity and Intrafamilial Classification
T2  - Systematic Botany
TI  - A Phylogeny of the Violaceae (Malpighiales) Inferred from Plastid DNA Sequences: Implications for Generic Diversity and Intrafamilial Classification
VL  - 39
ID  - 1116
ER  - 


TY  - JOUR
AB  - A phylogenetic analysis of Violaceae is presented using sequences from rbcL, atpB, matK and 18S rDNA from 39 species and 19 genera. The combined analysis of four molecular markers resulted in only one most parsimonious tree, and 33 of all 38 nodes within Violaceae are supported by a bootstrap proportion of more than 50%. Fusispermum is in a basal-most position and Rinorea, Decorsella, Rinoreocarpus and the other Violaceae are successively diverged. The monogeneric subfamily Fusispermoideae is supported, and it shares a number of plesiomorphies with Passifloraceae (a convolute petal aestivation, actinomorphic flowers and connate filaments). The other monogeneric subfamily Leonioideae is sunken within the subfamily Violoideae and is sister to Gloeospermum, sharing some seed morphological characteristics. The present molecular phylogenetic analysis suggests that the convolute, apotact and quincuncial petal aestivation is successively derived within the family. The evolutionary trends of the other morphological characteristics, such as a filament connation, the number of carpels and floral symmetry, are discussed.
AU  - Tokuoka, Toru
DO  - 10.1007/s10265-008-0153-0
IS  - 3
KW  - Violaceae
LA  - eng
PY  - 2008
SP  - 253-260
ST  - Molecular phylogenetic analysis of Violaceae (Malpighiales) based on plastid and nuclear DNA sequences
T2  - Journal of Plant Research
TI  - Molecular phylogenetic analysis of Violaceae (Malpighiales) based on plastid and nuclear DNA sequences
VL  - 121
ID  - 2998
ER  - 


TY  - JOUR
AU  - Stajsic, V.
AU  - Walsh, N. G.
AU  - Douglas, R.
AU  - Messina, A.
AU  - Molloy, B. P. J.
DO  - 10.1071/SB14022 M4 - Citavi
IS  - 4
KW  - Violaceae
N1  - PII: SB14022 PII: SB14022 PII: SB14022 PII: SB14022
PY  - 2014
SN  - 1030-1887
SP  - 305-323
ST  - A revision of Melicytus (Violaceae) in mainland Australia and Tasmania
T2  - Australian Systematic Botany
TI  - A revision of Melicytus (Violaceae) in mainland Australia and Tasmania
VL  - 27
ID  - 1076
ER  - 


TY  - JOUR
AU  - Paula-Souza, Juliana
AU  - Ballard, Harvey Eugene
DO  - 10.11646/phytotaxa.183.1.1 M4 - Citavi
IS  - 1
KW  - Violaceae
PY  - 2014
SN  - 1179-3163
SP  - 1
ST  - Re-establishment of the name Pombalia, and new combinations from the polyphyletic Hybanthus (Violaceae)
T2  - Phytotaxa
TI  - Re-establishment of the name Pombalia, and new combinations from the polyphyletic Hybanthus (Violaceae)
VL  - 183
ID  - 1016
ER  - 


TY  - JOUR
AU  - Nakamura, Koh
AU  - Denda, Tetsuo
AU  - Kokubugata, Goro
AU  - Huang, Chiun, Jr.
AU  - Peng, Ching- I.
AU  - Yokota, Masatsugu
DO  - 10.1007/s00606-014-1076-y M4 - Citavi
IS  - 1
KW  - Violaceae
N1  - PII: 1076 PII: 1076 PII: 1076 PII: 1076
PY  - 2015
SN  - 0378-2697
SP  - 337-351
ST  - Phylogeny and biogeography of the Viola iwagawae-tashiroi species complex (Violaceae, section Plagiostigma) endemic to the Ryukyu Archipelago, Japan
T2  - Plant Systematics and Evolution
TI  - Phylogeny and biogeography of the Viola iwagawae-tashiroi species complex (Violaceae, section Plagiostigma) endemic to the Ryukyu Archipelago, Japan
VL  - 301
ID  - 644
ER  - 


TY  - JOUR
AU  - Mitchell, A. D.
AU  - Heenan, Peter B.
AU  - Murray, B. G.
AU  - Molloy, B. P. J.
AU  - Lange, Peter J.
DO  - 10.1071/SB08042 M4 - Citavi
IS  - 3
KW  - Violaceae
N1  - PII: SB08042 PII: SB08042 PII: SB08042 PII: SB08042
PY  - 2009
SN  - 1030-1887
SP  - 143-157
ST  - Evolution of the south-western Pacific genus Melicytus (Violaceae): evidence from DNA sequence data, Cytology and sex Expression
T2  - Australian Systematic Botany
TI  - Evolution of the south-western Pacific genus Melicytus (Violaceae): evidence from DNA sequence data, Cytology and sex Expression
VL  - 22
ID  - 2602
ER  - 


TY  - JOUR
AB  - BACKGROUND DNA-dependent RNA polymerase IV and V (Pol IV and V) are multi-subunit enzymes occurring in plants. The origin of Pol V, specific to angiosperms, from Pol IV, which is present in all land plants, is linked to the duplication of the gene encoding the largest subunit and the subsequent subneofunctionalization of the two paralogs (NRPD1 and NRPE1). Additional duplication of the second-largest subunit, NRPD2/NRPE2, has happened independently in at least some eudicot lineages, but its paralogs are often subject to concerted evolution and gene death and little is known about their evolution nor their affinity with Pol IV and Pol V. RESULTS We sequenced a approximately 1500 bp NRPD2/E2-like fragment from 18 Viola species, mostly paleopolyploids, and 6 non-Viola Violaceae species. Incongruence between the NRPD2/E2-like gene phylogeny and species phylogeny indicates a first duplication of NRPD2 relatively basally in Violaceae, with subsequent sorting of paralogs in the descendants, followed by a second duplication in the common ancestor of Viola and Allexis. In Viola, the mutation pattern suggested (sub-) neofunctionalization of the two NRPD2/E2-like paralogs, NRPD2/E2-a and NRPD2/E2-b. The dN/dS ratios indicated that a 54 bp region exerted strong positive selection for both paralogs immediately following duplication. This 54 bp region encodes a domain that is involved in the binding of the Nrpd2 subunit with other Pol IV/V subunits, and may be important for correct recognition of subunits specific to Pol IV and Pol V. Across all Viola taxa 73 NRPD2/E2-like sequences were obtained, of which 23 (32%) were putative pseudogenes - all occurring in polyploids. The NRPD2 duplication was conserved in all lineages except the diploid MELVIO clade, in which NRPD2/E2-b was lost, and its allopolyploid derivates from hybridization with the CHAM clade, section Viola and section Melanium, in which NRPD2/E2-a occurred in multiple copies while NRPD2/E2-b paralogs were either absent or pseudogenized. CONCLUSIONS Following the relatively recent split of Pol IV and Pol V, our data indicate that these two multi-subunit enzymes are still in the process of specialization and each acquiring fully subfunctionalized copies of their subunit genes. Even after specialization, the NRPD2/E2-like paralogs are prone to pseudogenization and gene conversion and NRPD2 and NRPE2 copy number is a highly dynamic process modulated by allopolyploidy and gene death.
AD  - Centre for Ecological and Evolutionary Synthesis (CEES), Department of Biology, University of Oslo, 0316 Oslo, Norway. M4 - Citavi
AU  - Marcussen, Thomas
AU  - Oxelman, Bengt
AU  - Skog, Anna
AU  - Jakobsen, Kjetill S.
DO  - 10.1186/1471-2148-10-45
KW  - Violaceae
LA  - eng
N1  - Journal Article Research Support, Non-U.S. Gov't Journal Article Research Support, Non-U.S. Gov't Journal Article Research Support, Non-U.S. Gov't Journal Article Research Support, Non-U.S. Gov't
PY  - 2010
SP  - 45
ST  - Evolution of plant RNA polymerase IV/V genes: evidence of subneofunctionalization of duplicated NRPD2/NRPE2-like paralogs in Viola (Violaceae)
T2  - BMC evolutionary biology
TI  - Evolution of plant RNA polymerase IV/V genes: evidence of subneofunctionalization of duplicated NRPD2/NRPE2-like paralogs in Viola (Violaceae)
UR  - https://www.ncbi.nlm.nih.gov/pmc/articles/PMC2834690
VL  - 10
ID  - 2269
ER  - 


TY  - JOUR
AB  - The phylogenies of allopolyploids take the shape of networks and cannot be adequately represented as bifurcating trees. Especially for high polyploids (i.e., organisms with more than six sets of nuclear chromosomes), the signatures of gene homoeolog loss, deep coalescence, and polyploidy may become confounded, with the result that gene trees may be congruent with more than one species network. Herein, we obtained the most parsimonious species network by objective comparison of competing scenarios involving polyploidization and homoeolog loss in a high-polyploid lineage of violets (Viola, Violaceae) mostly or entirely restricted to North America, Central America, or Hawaii. We amplified homoeologs of the low-copy nuclear gene, glucose-6-phosphate isomerase (GPI), by single-molecule polymerase chain reaction (PCR) and the chloroplast trnL-F region by conventional PCR for 51 species and subspecies. Topological incongruence among GPI homoeolog subclades, owing to deep coalescence and two instances of putative loss (or lack of detection) of homoeologs, were reconciled by applying the maximum tree topology for each subclade. The most parsimonious species network and the fossil-based calibration of the homoeolog tree favored monophyly of the high polyploids, which has resulted from allodecaploidization 9-14 Ma, involving sympatric ancestors from the extant Viola sections Chamaemelanium (diploid), Plagiostigma (paleotetraploid), and Viola (paleotetraploid). Although two of the high-polyploid lineages (Boreali-Americanae, Pedatae) remained decaploid, recurrent polyploidization with tetraploids of section Plagiostigma within the last 5 Ma has resulted in two 14-ploid lineages (Mexicanae, Nosphinium) and one 18-ploid lineage (Langsdorffianae). This implies a more complex phylogenetic and biogeographic origin of the Hawaiian violets (Nosphinium) than that previously inferred from rDNA data and illustrates the necessity of considering polyploidy in phylogenetic and biogeographic reconstruction.
AU  - Marcussen, Thomas
AU  - Jakobsen, Kjetill S.
AU  - Danihelka, Jirí
AU  - Ballard, Harvey Eugene
AU  - Blaxland, Kim
AU  - Brysting, Anne K.
AU  - Oxelman, Bengt
DO  - 10.1093/sysbio/syr096
IS  - 1
KW  - Violaceae
LA  - eng
PY  - 2012
SP  - 107-126
ST  - Inferring species networks from gene trees in high-polyploid North American and Hawaiian violets (Viola, Violaceae)
T2  - Systematic Biology
TI  - Inferring species networks from gene trees in high-polyploid North American and Hawaiian violets (Viola, Violaceae)
UR  - https://www.ncbi.nlm.nih.gov/pmc/articles/PMC3243738
VL  - 61
ID  - 1626
ER  - 


TY  - JOUR
AB  - Allopolyploidization accounts for a significant fraction of speciation events in many eukaryotic lineages. However, existing phylogenetic and dating methods require tree-like topologies and are unable to handle the network-like phylogenetic relationships of lineages containing allopolyploids. No explicit framework has so far been established for evaluating competing network topologies, and few attempts have been made to date phylogenetic networks. We used a four-step approach to generate a dated polyploid species network for the cosmopolitan angiosperm genus Viola L. (Violaceae Batch.). The genus contains ca 600 species and both recent (neo-) and more ancient (meso-) polyploid lineages distributed over 16 sections. First, we obtained DNA sequences of three low-copy nuclear genes and one chloroplast region, from 42 species representing all 16 sections. Second, we obtained fossil-calibrated chronograms for each nuclear gene marker. Third, we determined the most parsimonious multilabeled genome tree and its corresponding network, resolved at the section (not the species) level. Reconstructing the "correct" network for a set of polyploids depends on recovering all homoeologs, i.e., all subgenomes, in these polyploids. Assuming the presence of Viola subgenome lineages that were not detected by the nuclear gene phylogenies ("ghost subgenome lineages") significantly reduced the number of inferred polyploidization events. We identified the most parsimonious network topology from a set of five competing scenarios differing in the interpretation of homoeolog extinctions and lineage sorting, based on (i) fewest possible ghost subgenome lineages, (ii) fewest possible polyploidization events, and (iii) least possible deviation from expected ploidy as inferred from available chromosome counts of the involved polyploid taxa. Finally, we estimated the homoploid and polyploid speciation times of the most parsimonious network. Homoploid speciation times were estimated by coalescent analysis of gene tree node ages. Polyploid speciation times were estimated by comparing branch lengths and speciation rates of lineages with and without ploidy shifts. Our analyses recognize Viola as an old genus (crown age 31 Ma) whose evolutionary history has been profoundly affected by allopolyploidy. Between 16 and 21 allopolyploidizations are necessary to explain the diversification of the 16 major lineages (sections) of Viola, suggesting that allopolyploidy has accounted for a high percentage-between 67% and 88%-of the speciation events at this level. The theoretical and methodological approaches presented here for (i) constructing networks and (ii) dating speciation events within a network, have general applicability for phylogenetic studies of groups where allopolyploidization has occurred. They make explicit use of a hitherto underexplored source of ploidy information from chromosome counts to help resolve phylogenetic cases where incomplete sequence data hampers network inference. Importantly, the coalescent-based method used herein circumvents the assumption of tree-like evolution required by most techniques for dating speciation events.
AD  - Department of Biosciences, Centre for Ecological and Evolutionary Synthesis (CEES), University of Oslo, PO Box 1066 Blindern, NO-0316 Oslo, Norway and Department of Biological and Environmental Sciences, University of Gothenburg, PO Box 461, 405 30 Gothenburg, Sweden Department of Biosciences, Centre for Ecological and Evolutionary Synthesis (CEES), University of Oslo, PO Box 1066 Blindern, NO-0316 Oslo, Norway and Department of Biological and Environmental Sciences, University of Gothenburg, PO Box 461, 405 30 Gothenburg, Sweden. Department of Biosciences, Centre for Ecological and Evolutionary Synthesis (CEES), University of Oslo, PO Box 1066 Blindern, NO-0316 Oslo, Norway and Department of Biological and Environmental Sciences, University of Gothenburg, PO Box 461, 405 30 Gothenburg, Sweden. Department of Biosciences, Centre for Ecological and Evolutionary Synthesis (CEES), University of Oslo, PO Box 1066 Blindern, NO-0316 Oslo, Norway and Department of Biological and Environmental Sciences, University of Gothenburg, PO Box 461, 405 30 Gothenburg, Sweden. Department of Biosciences, Centre for Ecological and Evolutionary Synthesis (CEES), University of Oslo, PO Box 1066 Blindern, NO-0316 Oslo, Norway and Department of Biological and Environmental Sciences, University of Gothenburg, PO Box 461, 405 30 Gothenburg, Sweden. Department of Biosciences, Centre for Ecological and Evolutionary Synthesis (CEES), University of Oslo, PO Box 1066 Blindern, NO-0316 Oslo, Norway and Department of Biological and Environmental Sciences, University of Gothenburg, PO Box 461, 405 30 Gothenburg, Sweden. M4 - Citavi
AU  - Marcussen, Thomas
AU  - Heier, Lise
AU  - Brysting, Anne K.
AU  - Oxelman, Bengt
AU  - Jakobsen, Kjetill S.
DO  - 10.1093/sysbio/syu071
IS  - 1
KW  - Violaceae
LA  - eng
N1  - Journal Article Research Support, Non-U.S. Gov't Journal Article Research Support, Non-U.S. Gov't Journal Article Research Support, Non-U.S. Gov't Journal Article Research Support, Non-U.S. Gov't
PY  - 2015
SP  - 84-101
ST  - From gene trees to a dated allopolyploid network: insights from the angiosperm genus Viola (Violaceae)
T2  - Systematic Biology
TI  - From gene trees to a dated allopolyploid network: insights from the angiosperm genus Viola (Violaceae)
UR  - https://www.ncbi.nlm.nih.gov/pmc/articles/PMC4265142
VL  - 64
ID  - 610
ER  - 


TY  - JOUR
AU  - Liangguan, xin
AU  - Xingfu, wu
IS  - 6
KW  - Violaceae
PY  - 2010
SP  - 477-488
ST  - Infrageneric phylogeny of the Genus Viola (Violaceae) based on trnL trnF, psbA-trnH, rpL16,ITS sequences, cytological and morphological Data
T2  - Acta Botanica Yunnanica
TI  - Infrageneric phylogeny of the Genus Viola (Violaceae) based on trnL trnF, psbA-trnH, rpL16,ITS sequences, cytological and morphological Data
VL  - 32
ID  - 2248
ER  - 


TY  - JOUR
AB  - The endemic Hawaiian flora offers remarkable opportunities to study the patterns of plant morphological and molecular evolution. The Hawaiian violets are a monophyletic lineage of nine taxa distributed across six main islands of the Hawaiian archipelago. To describe the evolutionary relationships, biogeography, and molecular evolution rates of the Hawaiian violets, we conducted a phylogenetic study using nuclear rDNA internal transcribed spacer sequences from specimens of each species. Parsimony, maximum likelihood (ML), and Bayesian inference reconstructions of island colonization and radiation strongly suggest that the Hawaiian violets first colonized the Maui Nui Complex, quickly radiated to Kaua'i and O'ahu, and recently dispersed to Hawai'i. The lineage consists of "wet" and "dry" clades restricted to distinct precipitation regimes. The ML and Bayesian inference reconstructions of shifts in habitat, habit, and leaf shape indicate that ecologically analogous taxa have undergone parallel evolution in leaf morphology and habit. This parallel evolution correlates with shifts to specialized habitats. Relative rate tests showed that woody and herbaceous sister species possess equal molecular evolution rates. The incongruity of molecular evolution rates in taxa on younger islands suggests that these rates may not be determined by growth form (or lifespan) alone, but may be influenced by complex dispersal events.
AD  - Department of Biology, Campbell University, Buies Creek, North Carolina 27506 USA. M4 - Citavi
AU  - Havran, J. Christopher
AU  - Sytsma, Kenneth J.
AU  - Ballard, Harvey Eugene
DO  - 10.3732/ajb.0900021
IS  - 11
KW  - Violaceae
LA  - eng
N1  - Journal Article Journal Article Journal Article Journal Article
PY  - 2009
SN  - 0002-9122
SP  - 2087-2099
ST  - Evolutionary relationships, interisland biogeography, and molecular evolution in the Hawaiian violets (Viola: Violaceae)
T2  - American journal of botany
TI  - Evolutionary relationships, interisland biogeography, and molecular evolution in the Hawaiian violets (Viola: Violaceae)
VL  - 96
ID  - 2540
ER  - 


TY  - JOUR
AU  - Conesa, M. À
AU  - Mus, M.
AU  - Rosselló, J. A.
DO  - 10.1007/s00606-008-0006-2 M4 - Citavi
IS  - 3-4
KW  - Violaceae
N1  - PII: 6 PII: 6 PII: 6 PII: 6
PY  - 2008
SN  - 0378-2697
SP  - 169-177
ST  - Hybridization between insular endemic and widespread species of Viola in non-disturbed environments assessed by nuclear ribosomal and cpDNA sequences
T2  - Plant Systematics and Evolution
TI  - Hybridization between insular endemic and widespread species of Viola in non-disturbed environments assessed by nuclear ribosomal and cpDNA sequences
VL  - 273
ID  - 2775
ER  - 


TY  - JOUR
AU  - Ballard, Harvey Eugene
AU  - Sytsma, Kenneth J.
DO  - 10.1554/0014-3820(2000)054[1521:EABOTW]2.0.CO;2 Y3 - 18.02.2019 M4 - Citavi
IS  - 5
KW  - Violaceae
PY  - 2000
SN  - 0014-3820
SP  - 1521-1532
ST  - Evolution And Biogeography Of The Woody Hawaiian Violets (Viola, Violaceae): Arctic Origins, Herbaceous Ancestry And Bird Dispersal
T2  - Evolution
TI  - Evolution And Biogeography Of The Woody Hawaiian Violets (Viola, Violaceae): Arctic Origins, Herbaceous Ancestry And Bird Dispersal
VL  - 54
ID  - 4318
ER  - 


TY  - JOUR
AB  - We investigated Viburnum phylogeny using separate and combined analyses of DNA sequence data from two chloroplast and three nuclear loci. Separate analyses of nuclear and chloroplast data sets resulted in gene trees that were generally congruent with one another and with trees from two previous analyses. Our gene trees do differ in the position of section Pseudotinus, as well as in species relationships within sections Pseudotinus and Lentago. However, tests for incongruence indicate that differences between the nuclear and chloroplast data are not significant. Furthermore, gene trees from combined analyses were highly similar to those found in separate analyses, suggesting that these localized differences do not affect other parts of the tree. Our analyses provide convincing support for numerous relationships, although there is still uncertainty at the base of the tree. To facilitate future study, we propose informal names for 12 well-supported species groups, as well as for several higher-level clades. We also discuss the biogeographic implications of our phylogeny, focusing on repeated, although apparently temporally incongruent, patterns of disjunction between the Old and New Worlds.
AD  - Department of Ecology and Evolutionary Biology, Yale University, P.O. Box 208106, New Haven, Connecticut 06520-8106 USA. M4 - Citavi
AU  - Winkworth, Richard C.
AU  - Donoghue, Michael J.
DO  - 10.3732/ajb.92.4.653
IS  - 4
KW  - Viburnaceae
LA  - eng
N1  - Journal Article Journal Article Journal Article Journal Article
PY  - 2005
SN  - 0002-9122
SP  - 653-666
ST  - Viburnum phylogeny based on combined molecular data: implications for taxonomy and biogeography
T2  - American journal of botany
TI  - Viburnum phylogeny based on combined molecular data: implications for taxonomy and biogeography
VL  - 92
ID  - 3729
ER  - 


TY  - JOUR
AB  - DNA sequencing studies of the granule-bound starch synthase gene (GBSSI) indicate the presence of two loci in Viburnum. Gene trees from separate and combined phylogenetic analyses of the GBSSI paralogues are generally congruent with each other and with trees from previous analyses, especially those of Donoghue et al. [Syst. Bot. 29 (2004) 188] based on nuclear ribosomal ITS and chloroplast trnK intron DNA sequences. Specifically, our GBSSI trees confirm (i) the monophyly of some and non-monophyly of other traditionally recognized taxonomic sections, (ii) the presence of three major supra-sectional lineages within Viburnum, and (iii) the resolution of many species relationships within the section-level clades. Analyses of GBSSI also provide greater resolution of relationships within the largest supra-sectional lineage. Relationships at the base of the Viburnum phylogeny remain uncertain; in particular, the position of the root, relationships among the supra-sectional clades, and the exact placement of several smaller groups (e.g., Viburnum clemensiae, Viburnum urceolatum, and section Pseudotinus). In two lineages each GBSSI paralogue is represented by two distinct sequences. The presence of additional copies appears to be correlated with polyploidy in these clades. Placement of the homoeologues in our gene trees suggests the possibility of a hybrid origin for these polyploids.
AD  - Department of Ecology and Evolutionary Biology, Yale University, P.O. Box 208105, New Haven, CT 06520-8105, USA. M4 - Citavi
AU  - Winkworth, Richard C.
AU  - Donoghue, Michael J.
DO  - 10.1016/j.ympev.2004.05.006
IS  - 1
KW  - Viburnaceae
LA  - eng
N1  - Comparative Study Journal Article Comparative Study Journal Article Comparative Study Journal Article Comparative Study Journal Article
PY  - 2004
SN  - 1055-7903
SP  - 109-126
ST  - Viburnum phylogeny: evidence from the duplicated nuclear gene GBSSI
T2  - Molecular phylogenetics and evolution
TI  - Viburnum phylogeny: evidence from the duplicated nuclear gene GBSSI
VL  - 33
ID  - 3930
ER  - 


TY  - JOUR
AB  - We used a near-complete phylogeny for the angiosperm clade Viburnum to assess lineage diversification rates, and to examine possible morphological and ecological factors driving radiations. Maximum-likelihood and Bayesian approaches identified shifts in diversification rate and possible links to character evolution. We inferred the ancestral environment for Viburnum and changes in diversification dynamics associated with subsequent biome shifts. Viburnum probably diversified in tropical forests of Southeast Asia in the Eocene, with three subsequent radiations in temperate clades during the Miocene. Four traits (purple fruits, extrafloral nectaries, bud scales and toothed leaves) were statistically associated with higher rates of diversification. However, we argue that these traits are unlikely to be driving diversification directly. Instead, two radiations were associated with the occupation of mountainous regions and a third with repeated shifts between colder and warmer temperate forests. Early-branching depauperate lineages imply that the rare lowland tropical species are 'dying embers' of once more diverse lineages; net diversification rates in Viburnum likely decreased in these tropical environments after the Oligocene. We suggest that 'taxon pulse' dynamics might characterize other temperate plant lineages.
AD  - Department of Ecology and Evolutionary Biology, Yale University, PO Box 208106, New Haven, CT, 06520, USA. Department of Biology, The College of New Jersey, 2000 Pennington Rd, Ewing, NJ, 08628, USA. Division of Botany, Peabody Museum of Natural History, Yale University, PO Box 208118, New Haven, CT, 06520, USA. Laboratorio de Botánica y Sistemática, Departamento de Ciencias Biológicas, Universidad de los Andes, Apartado Aéreo 4976, Bogotá, Colombia. Department of Ecology and Evolutionary Biology, Brown University, Box G-W, 80 Waterman Street, Providence, RI, 02912, USA. Department of Ecology and Evolutionary Biology, Yale University, PO Box 208106, New Haven, CT, 06520, USA. M4 - Citavi
AU  - Spriggs, Elizabeth L.
AU  - Clement, Wendy L.
AU  - Sweeney, Patrick W.
AU  - Madriñán, Santiago
AU  - Edwards, Erika J.
AU  - Donoghue, Michael J.
DO  - 10.1111/nph.13305
IS  - 2
KW  - Viburnaceae
LA  - eng
N1  - Journal Article Research Support, Non-U.S. Gov't Research Support, U.S. Gov't, Non-P.H.S. Journal Article Research Support, Non-U.S. Gov't Research Support, U.S. Gov't, Non-P.H.S. Journal Article Research Support, Non-U.S. Gov't Research Support, U.S. Gov't, Non-P.H.S. Journal Article Research Support, Non-U.S. Gov't Research Support, U.S. Gov't, Non-P.H.S.
PY  - 2015
SP  - 340-354
ST  - Temperate radiations and dying embers of a tropical past: the diversification of Viburnum
T2  - The New phytologist
TI  - Temperate radiations and dying embers of a tropical past: the diversification of Viburnum
VL  - 207
ID  - 728
ER  - 


TY  - JOUR
AB  - BACKGROUND The chloroplast genes matK and rbcL have been proposed as a "core" DNA barcode for identifying plant species. Published estimates of successful species identification using these loci (70-80%) may be inflated because they may have involved comparisons among distantly related species within target genera. To assess the ability of the proposed two-locus barcode to discriminate closely related species, we carried out a hierarchically structured set of comparisons within Viburnum, a clade of woody angiosperms containing ca. 170 species (some 70 of which are currently used in horticulture). For 112 Viburnum species, we evaluated rbcL + matK, as well as the chloroplast regions rpl32-trnL, trnH-psbA, trnK, and the nuclear ribosomal internal transcribed spacer region (nrITS). RESULTS At most, rbcL + matK could discriminate 53% of all Viburnum species, with only 18% of the comparisons having genetic distances >1%. When comparisons were progressively restricted to species within major Viburnum subclades, there was a significant decrease in both the discriminatory power and the genetic distances. trnH-psbA and nrITS show much higher levels of variation and potential discriminatory power, and their use in plant barcoding should be reconsidered. As barcoding has often been used to discriminate species within local areas, we also compared Viburnum species within two regions, Japan and Mexico and Central America. Greater success in discriminating among the Japanese species reflects the deeper evolutionary history of Viburnum in that area, as compared to the recent radiation of a single clade into the mountains of Latin America. CONCLUSIONS We found very low levels of discrimination among closely related species of Viburnum, and low levels of variation in the proposed barcoding loci may limit success within other clades of long-lived woody plants. Inclusion of the supplementary barcodes trnH-psbA and nrITS increased discrimination rates but were often more effective alone rather than in combination with rbcL + matK. We surmise that the efficacy of barcoding in plants has often been overestimated because of the lack of comparisons among closely related species. Phylogenetic information must be incorporated to properly evaluate relatedness in assessing the utility of barcoding loci.
AD  - Department of Ecology and Evolutionary Biology, Yale University, New Haven, CT 06520, USA. M4 - Citavi
AU  - Clement, Wendy L.
AU  - Donoghue, Michael J.
DO  - 10.1186/1471-2148-12-73
KW  - Viburnaceae
LA  - eng
N1  - Comparative Study Journal Article Research Support, Non-U.S. Gov't Research Support, U.S. Gov't, Non-P.H.S. Comparative Study Journal Article Research Support, Non-U.S. Gov't Research Support, U.S. Gov't, Non-P.H.S. Comparative Study Journal Article Research Support, Non-U.S. Gov't Research Support, U.S. Gov't, Non-P.H.S. Comparative Study Journal Article Research Support, Non-U.S. Gov't Research Support, U.S. Gov't, Non-P.H.S.
PY  - 2012
SP  - 73
ST  - Barcoding success as a function of phylogenetic relatedness in Viburnum, a clade of woody angiosperms
T2  - BMC evolutionary biology
TI  - Barcoding success as a function of phylogenetic relatedness in Viburnum, a clade of woody angiosperms
UR  - https://www.ncbi.nlm.nih.gov/pmc/articles/PMC3433329
VL  - 12
ID  - 1511
ER  - 


TY  - JOUR
AU  - Clement, Wendy L.
AU  - Donoghue, Michael J.
DO  - 10.1086/658927 M4 - Citavi
IS  - 4
KW  - Viburnaceae
PY  - 2011
SN  - 1058-5893
SP  - 559-573
ST  - Dissolution of Viburnum Section Megalotinus (Adoxaceae) of Southeast Asia and its Implications for Morphological Evolution and Biogeography
T2  - International Journal of Plant Sciences
TI  - Dissolution of Viburnum Section Megalotinus (Adoxaceae) of Southeast Asia and its Implications for Morphological Evolution and Biogeography
VL  - 172
ID  - 1860
ER  - 


TY  - JOUR
AB  - • Premise of the study: Despite recent progress, significant uncertainties remain concerning relationships among early-branching lineages within Viburnum (Adoxaceae), prohibiting a new classification and hindering studies of character evolution and the increasing use of Viburnum in addressing a wide range of ecological and evolutionary questions. We hoped to resolve these issues by sequencing whole plastid genomes for representative species and combining these with molecular data previously obtained from an expanded taxon sample.• Methods: We performed paired-end Illumina sequencing of plastid genomes of 22 Viburnum species and combined these data with a 10-gene data set to infer phylogenetic relationships for 113 species. We used the results to devise a comprehensive phylogenetic classification and to analyze the evolution of eight morphological characters that vary among early-branching lineages.• Key results: With greatly increased levels of confidence in most of the early branches, we propose a phylogenetic classification of Viburnum, providing formal phylogenetic definitions for 30 clades, including 13 with names recognized under the International Code of Nomenclature for Algae, Fungi, and Plants, eight with previously proposed informal names, and nine newly proposed names for major branches. Our parsimony reconstructions of bud structure, leaf margins, inflorescence form, ruminate endosperm, extrafloral nectaries, glandular trichomes, palisade anatomy, and pollen exine showed varying levels of homoplasy, but collectively provided morphological support for some, though not all, of the major clades.• Conclusions: Our study demonstrates the value of next-generation plastid sequencing, the ease of creating a formal phylogenetic classification, and the utility of such a system in describing patterns of character evolution.
AU  - Clement, Wendy L.
AU  - Arakaki, Mónica
AU  - Sweeney, Patrick W.
AU  - Edwards, Erika J.
AU  - Donoghue, Michael J.
DO  - 10.3732/ajb.1400015
IS  - 6
KW  - Viburnaceae
LA  - eng
PY  - 2014
SN  - 0002-9122
SP  - 1029-1049
ST  - A chloroplast tree for Viburnum (Adoxaceae) and its implications for phylogenetic classification and character evolution
T2  - American journal of botany
TI  - A chloroplast tree for Viburnum (Adoxaceae) and its implications for phylogenetic classification and character evolution
VL  - 101
ID  - 867
ER  - 


TY  - JOUR
AU  - Chatelet, David S.
AU  - Clement, Wendy L.
AU  - Sack, Lawren
AU  - Donoghue, Michael J.
AU  - Edwards, Erika J.
DO  - 10.1086/673241 M4 - Citavi
IS  - 9
KW  - Viburnaceae
PY  - 2013
SN  - 1058-5893
SP  - 1277-1291
ST  - The Evolution of Photosynthetic Anatomy in Viburnum (Adoxaceae)
T2  - International Journal of Plant Sciences
TI  - The Evolution of Photosynthetic Anatomy in Viburnum (Adoxaceae)
VL  - 174
ID  - 1191
ER  - 


TY  - JOUR
AB  - A well-resolved species level phylogeny is critically important in studying organismal evolution (e.g., hybridization, polyploidization, adaptive speciation). Lack of appropriate molecular markers that give sufficient resolution to gene trees is one of the major impediments to inferring species level phylogenies. In addition, sampling multiple independent loci is essential to overcome the lineage sorting problem. The availability of nuclear loci has often been a limiting factor in plant species-level phylogenetic studies. Here the two PHOT loci were developed as new sources of nuclear gene trees. The PHOT1 and PHOT2 gene trees of the Verbena complex (Verbenaceae) are well resolved and have good clade support. These gene trees are consistent with each other and previously generated chloroplast and nuclear waxy gene trees in most of the phylogenetic backbone as well as some terminal relationships, but are incongruent in some other relationships. Locus-specific primers were optimized for amplifying and sequencing these two loci in all Lamiales. Comparing intron size in the context of the gene trees shows dramatic variation within the Verbena complex, particularly at the PHOT1 locus. These variations are largely caused by invasions of short transposable elements and frequent long deletions and insertions of unknown causes. In addition, inspection of DNA sequences and phylogenetic analyses unmask a clear footprint of ancestral recombination in one species.
AD  - Department of Biology, University of Washington, Seattle, Washington 98195 USA. M4 - Citavi
AU  - Yuan, Yao-Wu
AU  - Olmstead, Richard G.
DO  - 10.3732/ajb.0800133
IS  - 9
KW  - Verbenaceae
LA  - eng
N1  - Journal Article Journal Article Journal Article Journal Article
PY  - 2008
SN  - 0002-9122
SP  - 1166-1176
ST  - Evolution and phylogenetic utility of the Phot gene duplicates in the Verbena complex (Verbenaceae): dramatic intron size variation and footprint of ancestral recombination
T2  - American journal of botany
TI  - Evolution and phylogenetic utility of the Phot gene duplicates in the Verbena complex (Verbenaceae): dramatic intron size variation and footprint of ancestral recombination
VL  - 95
ID  - 3032
ER  - 


TY  - JOUR
AB  - Two major impediments to infer plant phylogenies at inter- or intra- species level include the lack of appropriate molecular markers and the gene tree/species tree discordance. Both of these problems require more extensive investigations. One of the foci of this study is examining the phylogenetic utility of a combined chloroplast DNA dataset (>5.0kb) of seven non-coding regions, in comparison with that of a large fragment (ca. 3.0kb) of a low-copy nuclear gene (waxy), in a recent, rapidly diversifying group, the Verbena complex. The complex includes three very closely related genera, Verbena (base chromosome number x=7), Glandularia (x=5), and Junellia (x=10), comprising some 150 species distributed predominantly in South and North America. Our results confirm the inadequacy of non-coding cpDNA in resolving relationships among closely related species due to lack of variation, and the great potential of low-copy nuclear gene as source of variation. However, this study suggests that when both cpDNA and nuclear DNA are employed in low-level phylogenetic studies, cpDNA might be very useful to infer organelle evolutionary history (e.g., chloroplast transfer) and more comprehensively understand the evolutionary history of organisms. The phylogenetic framework of the Verbena complex resulted from this study suggests that Junellia is paraphyletic and most ancestral among the three genera; both Glandularia and Verbena are monophyletic and have been derived from within Junellia. Implications of this phylogenetic framework to understand chromosome number evolution and biogeography are discussed. Most interestingly, the comparison of the cpDNA and nuclear DNA phylogenies indicates two independent intergeneric chloroplast transfers, both from Verbena to Glandularia. One is from a diploid North American Verbena species to a polyploid North American Glandularia species. The other is more ancient, from the South American Verbena group to the common ancestor of a major Glandularia lineage, which has radiated subsequently in both South and North America. The commonly assumed introgressive hybridization may not explain the chloroplast transfers reported here. The underlying mechanism remains uncertain.
AD  - Department of Biology, University of Washington, PO Box 355325, Seattle, WA 98195-5325, USA. M4 - Citavi
AU  - Yuan, Yao-Wu
AU  - Olmstead, Richard G.
DO  - 10.1016/j.ympev.2008.04.004
IS  - 1
KW  - Verbenaceae
LA  - eng
N1  - Journal Article Research Support, U.S. Gov't, Non-P.H.S. Journal Article Research Support, U.S. Gov't, Non-P.H.S. Journal Article Research Support, U.S. Gov't, Non-P.H.S. Journal Article Research Support, U.S. Gov't, Non-P.H.S.
PY  - 2008
SN  - 1055-7903
SP  - 23-33
ST  - A species-level phylogenetic study of the Verbena complex (Verbenaceae) indicates two independent intergeneric chloroplast transfers
T2  - Molecular phylogenetics and evolution
TI  - A species-level phylogenetic study of the Verbena complex (Verbenaceae) indicates two independent intergeneric chloroplast transfers
VL  - 48
ID  - 3031
ER  - 


TY  - JOUR
AB  - The pentatricopeptide repeat (PPR) gene family, with hundreds of members in land plant genomes, has been recognized as a tremendous resource for plant phylogenetic studies based on publicly available genomic data from model organisms. However, whether this appealing nuclear gene marker system can be readily applied to non-model organisms remains questionable, particularly given the potential uncertainties in designing specific primers to only amplify the locus of interest from the sea of PPR genes. Here we demonstrate empirically the use of PPR genes in the family Verbenaceae and the Verbena complex. We also lay out a general scheme to design locus-specific primers to amplify and sequence PPR genes in non-model organisms. Intergeneric relationships within the family Verbenaceae were fully resolved with strong support. Relationships among the closely related genera within the Verbena complex and among some species groups within each genus were also well resolved, but resolution among very closely related species was limited. Our results suggest that PPR genes can be readily employed in non-model organisms. They may be best used to resolve relationships in a spectrum from among distantly related genera to among not-so-closely related congeneric species, but may have limited use among very closely related species.
AD  - Department of Biology, University of Washington, Seattle, WA 98195, USA. M4 - Citavi
AU  - Yuan, Yao-Wu
AU  - Liu, Chang
AU  - Marx, Hannah E.
AU  - Olmstead, Richard G.
DO  - 10.1016/j.ympev.2009.08.029
IS  - 1
KW  - Verbenaceae
LA  - eng
N1  - Journal Article Research Support, U.S. Gov't, Non-P.H.S. Journal Article Research Support, U.S. Gov't, Non-P.H.S. Journal Article Research Support, U.S. Gov't, Non-P.H.S. Journal Article Research Support, U.S. Gov't, Non-P.H.S.
PY  - 2010
SN  - 1055-7903
SP  - 23-35
ST  - An empirical demonstration of using pentatricopeptide repeat (PPR) genes as plant phylogenetic tools: phylogeny of Verbenaceae and the Verbena-complex
T2  - Molecular phylogenetics and evolution
TI  - An empirical demonstration of using pentatricopeptide repeat (PPR) genes as plant phylogenetic tools: phylogeny of Verbenaceae and the Verbena-complex
VL  - 54
ID  - 2410
ER  - 


TY  - JOUR
AU  - Peralta, Paola
AU  - Múlgura de Romero, María E.
AU  - Denham, Silvia S.
AU  - Botta, Silvia M.
DO  - 10.3417/2004167 M4 - Citavi
IS  - 2
KW  - Verbenaceae
PY  - 2008
SP  - 338-390
ST  - Revisión del Género Junellia (Verbenaceae)
T2  - Annals of the Missouri Botanical Garden
TI  - Revisión del Género Junellia (Verbenaceae)
VL  - 95
ID  - 2927
ER  - 


TY  - JOUR
AU  - O'Leary, Nataly
AU  - Yuan, Yao-Wu
AU  - Chemisquy, Maria Amelia
AU  - Olmstead, Richard G.
DO  - 10.1600/036364409790139691 M4 - Citavi
IS  - 4
KW  - Verbenaceae
PY  - 2009
SN  - 03636445
SP  - 777-786
ST  - Reassignment of Species of Paraphyletic Junellia s.l. to the new Genus Mulguraea (Verbenaceae) and New Circumscription of Genus Junellia : Molecular and Morphological Congruence
T2  - Systematic Botany
TI  - Reassignment of Species of Paraphyletic Junellia s.l. to the new Genus Mulguraea (Verbenaceae) and New Circumscription of Genus Junellia : Molecular and Morphological Congruence
VL  - 34
ID  - 2624
ER  - 


TY  - JOUR
AB  - PREMISE OF THE STUDY A new infrafamilial circumscription of the Verbenaceae with eight tribes: Casselieae, Citharexyleae, Duranteae, Lantaneae, Neospartoneae, Petreeae, Priveae, and Verbeneae, has been recently proposed, on the basis of molecular phylogenetic studies. Two genera, Dipyrena and Rhaphithamnus, remain unplaced. The aim of this work is to reconstruct the evolutionary history of morphological characters traditionally employed in the classification of the Verbenaceae, with special attention to tribes Verbeneae and Lantaneae. METHODS Twenty-one characters, related to habit and vegetative morphology, inflorescence and floral morphology, ovary and fruit morphology, as well as chromosome number, were optimized over a molecular phylogeny of Verbenaceae. KEY RESULTS All tribes are supported by at least one morphological trait except tribes Duranteae and Citharexyleae. Suffrutescent habit, sessile flowers, and four cluses are synapomorphies for tribe Verbeneae. Gynoecium with short style and entire stigma are synapomorphic traits for tribe Lantaneae. Sessile flowers and unicarpellate ovaries are morphological synapomorphies for the new tribe Neospartoneae. Suffrutescent habit is a synapomorphic trait for tribe Priveae. Homothetic pleiobotrya and absence of the adaxial staminode are synapomorphic traits for tribe Casselieae. Undivided fleshy fruits are probably a synapomorphic trait for tribe Petreeae. Putative plesiomorphies for the ancestor of the Verbenaceae are discussed as well as synapomorphic traits within other Verbenaceae clades. CONCLUSIONS Many of the characters traditionally employed in classification have proven to be very homoplastic, or have been shown not to support relationships within the family. Moreover, traditional assumptions concerning character polarity have in some cases been shown to be incorrect.
AD  - Instituto de Botánica Darwinion, Labardén 200, CC 22, San Isidro, Buenos Aires, Argentina. M4 - Citavi
AU  - O'Leary, Nataly
AU  - Calviño, Carolina Isabel
AU  - Martínez, Susana
AU  - Lu-Irving, Patricia
AU  - Olmstead, Richard G.
AU  - Múlgura, Maria Ema
DO  - 10.3732/ajb.1200123
IS  - 11
KW  - Verbenaceae
LA  - eng
N1  - Journal Article Research Support, Non-U.S. Gov't Research Support, U.S. Gov't, Non-P.H.S. Journal Article Research Support, Non-U.S. Gov't Research Support, U.S. Gov't, Non-P.H.S. Journal Article Research Support, Non-U.S. Gov't Research Support, U.S. Gov't, Non-P.H.S. Journal Article Research Support, Non-U.S. Gov't Research Support, U.S. Gov't, Non-P.H.S.
PY  - 2012
SN  - 0002-9122
SP  - 1778-1792
ST  - Evolution of morphological traits in Verbenaceae
T2  - American journal of botany
TI  - Evolution of morphological traits in Verbenaceae
VL  - 99
ID  - 1658
ER  - 


TY  - JOUR
AU  - Nesom, Guy L.
KW  - Verbenaceae
PY  - 2010
SP  - 1-15
ST  - infrageneric Classification of Verbena (Verbenaceae)
T2  - Phytoneuron
TI  - infrageneric Classification of Verbena (Verbenaceae)
VL  - 11
ID  - 2294
ER  - 


TY  - JOUR
AB  - UNLABELLED PREMISE OF THE STUDY Verbenaceae consist of trees, shrubs, lianas, and herbs distributed primarily in Latin America, where they occur in a wide array of ecosystems. A second center of diversity exists in Africa. Competing morphology-based classifications that rely on different traits conflict in significant ways. A broad phylogenetic study was undertaken to assess those classifications and to examine the historical geography of the family. • METHODS Analysis of seven chloroplast DNA regions for 109 species, representing all genera except one monotypic genus, provide inference into evolutionary relationships in Verbenaceae. • KEY RESULTS The phylogeny shows that none of the traditional classifications reflect phylogenetic relationships very well. Eight clades are recognized as tribes (Casselieae, Citharexyleae, Duranteae, Lantaneae, Neospartoneae trib. nov., Petreeae, Priveae, and Verbeneae). Two genera, Dipyrena and Rhaphithamnus, remain unplaced in these larger clades. Petreeae, which consist of Neotropical lianas, are sister to the rest of the family. Lantaneae and Verbeneae together form a derived clade that comprises approximately two-thirds of the species in Verbenaceae. • CONCLUSIONS We present a new tribal classification, including one new tribe, Neospartoneae trib. nov., to accommodate three small genera of Argentine species (Diostea, Neosparton, and Lampaya). Phylogenetic inference suggests a South American origin for Verbenaceae, with approximately six colonization events having given rise to the Old World species.
AD  - Department of Biology and Burke Museum, University of Washington, Seattle, Washington 98195, USA. M4 - Citavi
AU  - Marx, Hannah E.
AU  - O'Leary, Nataly
AU  - Yuan, Yao-Wu
AU  - Lu-Irving, Patricia
AU  - Tank, David C.
AU  - Múlgura, María E.
AU  - Olmstead, Richard G.
DO  - 10.3732/ajb.1000144
IS  - 10
KW  - Verbenaceae
LA  - eng
N1  - Journal Article Journal Article Journal Article Journal Article
PY  - 2010
SN  - 0002-9122
SP  - 1647-1663
ST  - A molecular phylogeny and classification of Verbenaceae
T2  - American journal of botany
TI  - A molecular phylogeny and classification of Verbenaceae
VL  - 97
ID  - 2273
ER  - 


TY  - JOUR
AU  - Lu-Irving, Patricia
AU  - Olmstead, Richard G.
KW  - Verbenaceae
PY  - 2013
SP  - 103-119
ST  - Investigating the evolution of Lantaneae (Verbenaceae) using multiple loci
T2  - Botanical Journal of the Linnean Society
TI  - Investigating the evolution of Lantaneae (Verbenaceae) using multiple loci
VL  - 171
ID  - 1302
ER  - 


TY  - JOUR
AU  - Lu-Irving, Patricia
AU  - O'Leary, Nataly
AU  - O'Brien, Anna
AU  - Olmstead, Richard G.
DO  - 10.1600/036364414X680816 M4 - Citavi
IS  - 2
KW  - Verbenaceae
PY  - 2014
SN  - 03636445
SP  - 644-655
ST  - Resolving the Genera Aloysia and Acantholippia within tribe Lantaneae (Verbenaceae), using Chloroplast and Nuclear Sequences
T2  - Systematic Botany
TI  - Resolving the Genera Aloysia and Acantholippia within tribe Lantaneae (Verbenaceae), using Chloroplast and Nuclear Sequences
VL  - 39
ID  - 971
ER  - 


TY  - JOUR
AU  - Menezes, Nanuza L.
AU  - Mello-Silva, Renato
AU  - Mayo, Simon J.
DO  - 10.2307/4110200 Y3 - 22.02.2019 M4 - Citavi
IS  - 1
KW  - Velloziaceae
PY  - 1993
SN  - 0075-5974
SP  - 71-92
ST  - A Cladistic Analysis of the Velloziaceae
T2  - Kew Bulletin
TI  - A Cladistic Analysis of the Velloziaceae
VL  - 49
ID  - 4477
ER  - 


TY  - JOUR
AU  - Mello-Silva, Renato
AU  - Montserrat, Laura
DO  - 10.1007/S12225-015-9575-8 M4 - Citavi
IS  - 2
KW  - Velloziaceae
N1  - PII: 9575 PII: 9575 PII: 9575 PII: 9575
PY  - 2015
SN  - 0075-5974
SP  - 22
ST  - Depicting Barbacenia flava and Vellozia intermedia with a short history of illustrated Velloziaceae
T2  - Kew Bulletin
TI  - Depicting Barbacenia flava and Vellozia intermedia with a short history of illustrated Velloziaceae
VL  - 70
ID  - 622
ER  - 


TY  - JOUR
AU  - Mello-Silva, Renato
DO  - 10.1111/j.1095-8339.2005.00399.x Y3 - 22.02.2019 M4 - Citavi
IS  - 2
KW  - Velloziaceae
PY  - 2005
SP  - 157-173
ST  - Morphological analysis, phylogenies and classification in Velloziaceae
T2  - Botanical Journal of the Linnean Society
TI  - Morphological analysis, phylogenies and classification in Velloziaceae
VL  - 148
ID  - 3654
ER  - 


TY  - JOUR
AD  - Pflanzliche Zellbiologie; Centre for Organismal Studies; Ruprecht-Karls-Universität Heidelberg; Im Neuenheimer Feld 230; 69120; Heidelberg; Germany Institut für Pharmazie und Molekulare Biotechnologie; Ruprecht-Karls-Universität Heidelberg; Im Neuenheimer Feld 364; 69120; Heidelberg; Germany M4 - Citavi
AU  - Behnke, H. Dietmar
AU  - Hummel, Eric
AU  - Hillmer, Stefan
AU  - Sauer-Gürth, Hedwig
AU  - Gonzalez, Javier
AU  - Wink, Michael
DO  - 10.1111/boj.12018 Y3 - 22.02.2019
IS  - 1
KW  - Velloziaceae
PY  - 2013
SP  - 22-94
ST  - A revision of African Velloziaceae based on leaf anatomy characters and rbcL nucleotide sequences
T2  - Botanical Journal of the Linnean Society
TI  - A revision of African Velloziaceae based on leaf anatomy characters and rbcL nucleotide sequences
VL  - 172
ID  - 1173
ER  - 


TY  - JOUR
AU  - Pyck, N.
AU  - van Lysebetten, A.
AU  - Stessens, J.
AU  - Smets, E.
DO  - 10.1007/s006060200053 Y3 - 10.02.2019 M4 - Citavi
IS  - 1-2
KW  - Valerianaceae
N1  - PII: 19YV3PX3CC7NP6WR PII: 19YV3PX3CC7NP6WR PII: 19YV3PX3CC7NP6WR PII: 19YV3PX3CC7NP6WR
PY  - 2002
SN  - 0378-2697
SP  - 29-46
ST  - The phylogeny of Patrinieae sensu Graebner (Valerianaceae) revisited: additional evidence from ndh F sequence data
T2  - Plant Systematics and Evolution
TI  - The phylogeny of Patrinieae sensu Graebner (Valerianaceae) revisited: additional evidence from ndh F sequence data
VL  - 233
ID  - 4157
ER  - 


TY  - JOUR
AU  - Jacobs, Bart
AU  - Bell, Charles D.
AU  - Smets, Erik
DO  - 10.1086/651243 M4 - Citavi
IS  - 4
KW  - Valerianaceae
PY  - 2010
SN  - 1058-5893
SP  - 421-434
ST  - Fruits and Seeds of the Valeriana Clade (Dipsacales): Diversity and Evolution
T2  - International Journal of Plant Sciences
TI  - Fruits and Seeds of the Valeriana Clade (Dipsacales): Diversity and Evolution
VL  - 171
ID  - 2214
ER  - 


TY  - JOUR
AB  - BACKGROUND AND AIMS The monophyly of Valerianaceae and the precise delimitation of the family are not totally resolved. Our knowledge on the phylogeny of the group is only partial: on a morphological basis, some contradicting taxonomic proposals have been published, which demonstrates the difficulties in establishing a natural classification of the family and especially in proposing a relevant treatment of the large genus Valeriana. The aims of this study are to contribute to the phylogeny and generic delineation of the Valerianaceae on the basis of molecular data. METHODS A cladistic analysis of the sequences of one plastid (matK) and one nuclear (ITS) molecular marker was carried out, both individually and in combination. KEY RESULTS The results of the analyses of both regions confirm that the family is monophyletic, with the exclusion of Triplostegia. The tribe Patrinieae is monophyletic, and the tribe Valerianeae is also a natural group. Two of the subtribes of Valerianeae, Fediinae and Centranthinae, are also monophyletic, with the exclusion of the genus Plectritis from Fediinae. The subtribe Valerianinae, on the other hand, is paraphyletic. CONCLUSIONS Our results confirm, for the first time on a molecular basis, the suggested paraphyly of Valeriana in its present circumscription, with profound nomenclatural and taxonomic implications. The correlation between molecular phylogeny and biogeography is close. In the course of the plastid DNA sequencing, a polymorphism concerning the matK gene was found, a fact that should be carefully evaluated in phylogenetic analyses.
AU  - Hidalgo, Oriane
AU  - Garnatje, Teresa
AU  - Susanna, Alfonso
AU  - Mathez, Joel
DO  - 10.1093/aob/mch042
IS  - 3
KW  - Valerianaceae
LA  - eng
PY  - 2004
SN  - 0305-7364
SP  - 283-293
ST  - Phylogeny of Valerianaceae based on matK and ITS markers, with reference to matK individual polymorphism
T2  - Annals of Botany
TI  - Phylogeny of Valerianaceae based on matK and ITS markers, with reference to matK individual polymorphism
UR  - https://www.ncbi.nlm.nih.gov/pmc/articles/PMC4242203
VL  - 93
ID  - 3800
ER  - 


TY  - JOUR
AB  - The southern Andean clade of Valeriana provides an excellent model for the study of biogeography. Here we provide new data to help clarify phylogenetic relationships among the South American valerians, with special focus on taxa found in the southern Andes. We found that the southern Andean taxa formed a clade in maximum likelihood and maximum parsimony analyses, and used a Bayesian relaxed clock method to estimate divergence times within Valerianaceae. Our temporal results were similar to other studies, but we found greater variance in our estimates, suggesting that the species of Valeriana have been on the South American continent for some time, and have been successful at exploiting new niche opportunities that reflects the contemporary radiation. Regardless of the time frame for the radiation of the clade, the uptick in the rate of diversification in Valerianaceae appears correlated with a dispersal event from Central to South America. The appearance of Valeriana in the southern Andes (13.7 Ma) corresponds with the transition from closed forest on the western side of the Andes in central Chile to a more open Mediterranean woodland environment. This would suggest that the high species richness of Valerianaceae in South America is the result of multiple, smaller radiations such as the one in the southern Andes, that may or may not be geographically isolated. These smaller radiations may also be driven by species moving into new biomes (migration from a temperate to a more Mediterranean-type climate and into alpine). The degree to which different ecological and geological factors interact to drive diversification is difficult to ascertain. Likewise, without a better-resolved phylogeny it is impossible to determine the directionality of dispersal in this group; did they colonize the southern Andes first, then move northward as the central Andean alpine habitat became more widely available or vice versa?
AD  - Department of Biological Sciences, University of New Orleans, 2000 Lakeshore Drive, New Orleans, LA 70148, USA. M4 - Citavi
AU  - Bell, Charles D.
AU  - Kutschker, Adriana
AU  - Arroyo, Mary T. K.
DO  - 10.1016/j.ympev.2012.02.015
IS  - 3
KW  - Valerianaceae
LA  - eng
N1  - Journal Article Research Support, Non-U.S. Gov't Research Support, U.S. Gov't, Non-P.H.S. Journal Article Research Support, Non-U.S. Gov't Research Support, U.S. Gov't, Non-P.H.S. Journal Article Research Support, Non-U.S. Gov't Research Support, U.S. Gov't, Non-P.H.S. Journal Article Research Support, Non-U.S. Gov't Research Support, U.S. Gov't, Non-P.H.S.
PY  - 2012
SN  - 1055-7903
SP  - 724-737
ST  - Phylogeny and diversification of Valerianaceae (Dipsacales) in the southern Andes
T2  - Molecular phylogenetics and evolution
TI  - Phylogeny and diversification of Valerianaceae (Dipsacales) in the southern Andes
VL  - 63
ID  - 1470
ER  - 


TY  - JOUR
AB  - This study aimed to investigate the phylogenetic utility of genotyping-by-sequencing (GBS) data in the southern South American subclade of Valerianaceae (Dipsacales). The variety of forms that has arisen in this clade, presumably over the past 5-10 million years, has all the signatures of an adaptive and rapid radiation. While the phylogeny of Valerianaceae has received a great deal of attention in the last decade, species relationships have been hard to resolve using traditional phylogenetic markers. Here, we collected high-throughput genomic sequence data from reduced-representation libraries obtained through GBS protocols. Putative orthologs were identified using within- and among-sample clustering using the computer software pyRAD. We recovered over 3000 loci for 14 species of southern South AmericanValeriana,with 140 loci present across all samples.We analyzed a set of phylogenetic trees generated from each locus using maximum likelihood methods, as well as multispecies coalescent (∗BEAST) methods. For comparative purposes, we also used a supermatrix approach to infer the phylogeny for these taxa. Across different methods and data sets, we recovered consistent relationships for the southern South American valerians that we sampled with varying degrees of support.
AD  - Department of Biological Sciences, University of New Orleans, 2000 Lakeshore Drive, New Orleans, LA 70148, USA. Electronic address:. Department of Biological Sciences, University of New Orleans, 2000 Lakeshore Drive, New Orleans, LA 70148, USA. M4 - Citavi
AU  - Bell, Charles D.
AU  - Gonzalez, Lauren A.
DO  - 10.1016/j.ympev.2018.02.014
KW  - Valerianaceae
LA  - eng
N1  - Journal Article Journal Article Journal Article Journal Article
PY  - 2018
SN  - 1055-7903
SP  - 44-49
ST  - Exploring the utility of "next-generation" sequence data on inferring the phylogeny of the South American Valeriana (Valerianaceae)
T2  - Molecular phylogenetics and evolution
TI  - Exploring the utility of "next-generation" sequence data on inferring the phylogeny of the South American Valeriana (Valerianaceae)
VL  - 123
ID  - 3
ER  - 


TY  - JOUR
AB  - Although Valerianaceae is a relatively small group of angiosperms (ca. 350 species), sample sizes of previous phylogenetic studies have been limited and taxonomic sampling has been usually geographically biased to species from Europe and/or South America. One group that has never been included in any analyses to date is the North American representatives of Valerianella. In this study I sampled 17 additional accessions from Valerianaceae, including six of the 12 described North American Valerianella species and four additional samples of European Valerianella species. Phylogenies based on parsimony and Bayesian methods show strong support for placing the North American Valerianella nested within the European species. These analyses also found Fedia to be nested within Valerianella, making Valerianella paraphyletic, a result consistent with several previously published molecular phylogenies. Divergence times estimated from the molecular data using recently proposed Bayesian 'relaxed' clock methods, suggest Valerianella was in North America by the middle Miocene (ca. 16-14 million years ago). These dates would suggest the North American species of Valerianella arrived in the New World just prior to the North Atlantic land bridge (NALB) no longer being a viable conduit for migration. Given the large amount of uncertainty in the divergence time estimates, dispersal across the North Atlantic may not be the only viable alternative to explain the current distribution of Valerianella. However, biogeographic analyses using an explicit model based approach, strongly favor the NALB as the route of dispersal for Valerianella.
AD  - M4 - Citavi
AU  - Bell, Charles D.
DO  - 10.1016/j.ympev.2007.03.013
IS  - 3
KW  - Valerianaceae
LA  - eng
N1  - Journal Article Research Support, Non-U.S. Gov't Research Support, U.S. Gov't, Non-P.H.S. Journal Article Research Support, Non-U.S. Gov't Research Support, U.S. Gov't, Non-P.H.S. Journal Article Research Support, Non-U.S. Gov't Research Support, U.S. Gov't, Non-P.H.S. Journal Article Research Support, Non-U.S. Gov't Research Support, U.S. Gov't, Non-P.H.S.
PY  - 2007
SN  - 1055-7903
SP  - 929-941
ST  - Phylogenetic placement and biogeography of the North American species of Valerianella (Valerianaceae: Dipsacales) based on chloroplast and nuclear DNA
T2  - Molecular phylogenetics and evolution
TI  - Phylogenetic placement and biogeography of the North American species of Valerianella (Valerianaceae: Dipsacales) based on chloroplast and nuclear DNA
VL  - 44
ID  - 3053
ER  - 


TY  - JOUR
AB  - Valerianaceae is a relatively small (ca. 350 species), but morphologically diverse angiosperm clade. Sequence data from the entire ndhF gene, the trnL-F intergenic spacer region, the trnL intron, the matK region, the rbcL-atpB intergenic spacer region and internal transcribed spacer (ITS) region of nuclear ribosomal DNA were collected for 21 taxa within Dipsacaceae and Valerianaceae (1 and 20, respectively). These data were included in several phylogenetic analyses with previously published sequences from Dipsacales. Results from these analyses (maximum parsimony, maximum likelihood, and Bayesian analysis) are in strong agreement with many of the conclusions from previous studies, most importantly: (1) Valerianaceae is sister to Dipsacaceae; (2) Triplostegia is more closely related to species of Dipsacaceae than to Valerianaceae; and (3) Valeriana appears not to be monophyletic, with Valeriana celtica falling outside the remainder of the species of Valeriana sampled here (with very strong support). With the exception of V. celtica, these data support two major clades within Valeriana; one that is exclusively New World and another that is distributed in both the Old and New World. Although the species of Valerianaceae and its sister group Dipsacaceae plus Triplostegia, are widely distributed in the Northern Hemisphere, and the data imply that Valerianaceae diversified initially in Asia (the Himalayan Patrinia and Nardostachys falling at the base of the clade), the center of modern species diversity for the group is in the Andes of South America with as many as 175 species restricted to that region. Although the exclusively South American taxa form a clade in the chloroplast and combined ITS and chloroplast analyses, support values tend to be low. Future studies will need to include additional data, in the form of both characters and taxa, before any strong conclusions about the character evolution, diversification, and biogeography of the South American valerians can be made.
AD  - Department of Ecology and Evolutionary Biology, Yale University, New Haven, CT 06520, USA. M4 - Citavi
AU  - Bell, Charles D.
DO  - 10.1016/j.ympev.2003.07.006
IS  - 1
KW  - Valerianaceae
LA  - eng
N1  - Journal Article Research Support, U.S. Gov't, Non-P.H.S. Journal Article Research Support, U.S. Gov't, Non-P.H.S. Journal Article Research Support, U.S. Gov't, Non-P.H.S. Journal Article Research Support, U.S. Gov't, Non-P.H.S.
PY  - 2004
SN  - 1055-7903
SP  - 340-350
ST  - Preliminary phylogeny of Valerianaceae (Dipsacales) inferred from nuclear and chloroplast DNA sequence data
T2  - Molecular phylogenetics and evolution
TI  - Preliminary phylogeny of Valerianaceae (Dipsacales) inferred from nuclear and chloroplast DNA sequence data
VL  - 31
ID  - 3755
ER  - 


TY  - JOUR
AU  - Bell, C.
DO  - 10.1016/j.ode.2004.10.014 M4 - Citavi
IS  - 2
KW  - Valerianaceae
N1  - PII: S1439609205000292 PII: S1439609205000292 PII: S1439609205000292 PII: S1439609205000292
PY  - 2005
SN  - 1439-6092
SP  - 147-159
ST  - Phylogeny and biogeography of Valerianaceae (Dipsacales) with special reference to the South American valerians
T2  - Organisms Diversity & Evolution
TI  - Phylogeny and biogeography of Valerianaceae (Dipsacales) with special reference to the South American valerians
VL  - 5
ID  - 3557
ER  - 


TY  - JOUR
AB  - Urticaceae is one of the larger Angiosperm families, but relationships within it remain poorly known. This study presents the first densely sampled molecular phylogeny of Urticaceae, using maximum likelihood (ML), maximum parsimony (MP) and Bayesian inference (BI) to analyze the DNA sequence data from two nuclear (ITS and 18S), four chloroplast (matK, rbcL, rpll4-rps8-infA-rpl36, trnL-trnF) and one mitochondrial (matR) loci. We sampled 169 accessions representing 122 species, representing 47 of the 54 recognized genera within Urticaceae, including four of the six sometimes separated as Cecropiaceae. Major results included: (1) Urticaceae including Cecropiaceae was monophyletic; (2) Cecropiaceae was biphyletic, with both lineages nested within Urticaceae; (3) Urticaceae can be divided into four well-supported clades; (4) previously erected tribes or subfamilies were broadly supported, with some additions and alterations; (5) the monophyly of many genera was supported, whereas Boehmeria, Pellionia, Pouzolzia and Urera were clearly polyphyletic, while Urtica and Pilea each had a small genus nested within them; (6) relationships between genera were clarified, mostly with substantial support. These results clarify that some morphological characters have been overstated and others understated in previous classifications of the family, and provide a strong foundation for future studies on biogeography, character evolution, and circumscription of difficult genera.
AD  - Key Laboratory of Biodiversity and Biogeography, Kunming Institute of Botany, Chinese Academy of Sciences, Kunming 650201, China; Germplasm Bank of Wild Species, Kunming Institute of Botany, Chinese Academy of Sciences, Kunming 650201, China; University of Chinese Academy of Sciences, Beijing 100049, China. M4 - Citavi
AU  - Wu, Zeng-Yuan
AU  - Monro, Alex K.
AU  - Milne, Richard I.
AU  - Wang, Hong
AU  - Yi, Ting-Shuang
AU  - Liu, Jie
AU  - Li, De-Zhu
DO  - 10.1016/j.ympev.2013.06.022
IS  - 3
KW  - Urticaceae
LA  - eng
N1  - Journal Article Research Support, Non-U.S. Gov't Journal Article Research Support, Non-U.S. Gov't Journal Article Research Support, Non-U.S. Gov't Journal Article Research Support, Non-U.S. Gov't
PY  - 2013
SN  - 1055-7903
SP  - 814-827
ST  - Molecular phylogeny of the nettle family (Urticaceae) inferred from multiple loci of three genomes and extensive generic sampling
T2  - Molecular phylogenetics and evolution
TI  - Molecular phylogeny of the nettle family (Urticaceae) inferred from multiple loci of three genomes and extensive generic sampling
VL  - 69
ID  - 1435
ER  - 


TY  - JOUR
AU  - Treiber, Erin L.
AU  - Gaglioti, André Luiz
AU  - Romaniuc-Neto, Sergio
AU  - Madriñán, Santiago
AU  - Weiblen, George D.
DO  - 10.1600/036364416X690633 M4 - Citavi
IS  - 1
KW  - Urticaceae
PY  - 2016
SN  - 03636445
SP  - 56-66
ST  - Phylogeny of the Cecropieae (Urticaceae) and the Evolution of an Ant-Plant Mutualism
T2  - Systematic Botany
TI  - Phylogeny of the Cecropieae (Urticaceae) and the Evolution of an Ant-Plant Mutualism
VL  - 41
ID  - 380
ER  - 


TY  - JOUR
AU  - Monro, Alex K.
IS  - 3
KW  - Urticaceae
PY  - 2006
SN  - 0002-9122
SP  - 426-441
ST  - The Revision of Species-rich Genera: A Phylogenetic Framework for the Strategic Revision Of Pilea (Urticaceae) based on cpDNA, nrDNA, and Morphology
T2  - American journal of botany
TI  - The Revision of Species-rich Genera: A Phylogenetic Framework for the Strategic Revision Of Pilea (Urticaceae) based on cpDNA, nrDNA, and Morphology
VL  - 93
ID  - 3452
ER  - 


TY  - JOUR
AD  - Key Laboratory for Plant Diversity and Biogeography of East Asia, Kunming Institute of Botany, Chinese Academy of Sciences, Kunming 650204, Yunnan, P.R. China Key Laboratory for Plant Diversity and Biogeography of East Asia, Kunming Institute of Botany, Chinese Academy of Sciences, Kunming 650204, Yunnan, P.R. China, School of Life Science, Yunnan University, Kunming 650091, P.R China, University of Chinese Academy of Sciences, Beijing 100039, P.R. China Jodrell Laboratory of Royal Botanic Gardens, Kew, Richmond, Surrey TW9 3AB, U.K. Key Laboratory of Plant Resources Conservation and Utilization, Jishou University, College of Hunan Province, Jishou 416000, Hunan, P.R. China Key Laboratory for Plant Diversity and Biogeography of East Asia, Kunming Institute of Botany, Chinese Academy of Sciences, Kunming 650204, Yunnan, P.R. China;, M4 - Citavi
AU  - Kim, Changkyun
AU  - Deng, Tao
AU  - Chase, Mark W.
AU  - Zhang, Dai-Gui
AU  - Nie, Ze-long
AU  - Sun, Hang
DO  - 10.12705/641.20
IS  - 1
KW  - Urticaceae
PY  - 2015
SN  - 00400262
SP  - 65-78
ST  - Generic phylogeny and character evolution in Urticeae (Urticaceae) inferred from nuclear and plastid DNA regions
T2  - Taxon
TI  - Generic phylogeny and character evolution in Urticeae (Urticaceae) inferred from nuclear and plastid DNA regions
VL  - 64
ID  - 559
ER  - 


TY  - JOUR
AU  - Jestrow, Brett
AU  - Valdés, James J.
AU  - Rodríguez, Francisco Jiménez
AU  - Francisco-Ortega, Javier
IS  - 3
KW  - Urticaceae
PY  - 2012
SN  - 00400262
SP  - 592-600
ST  - Phylogenetic placement of the Dominican Republic endemic genus Sarcopilea (Urticaceae)
T2  - Taxon
TI  - Phylogenetic placement of the Dominican Republic endemic genus Sarcopilea (Urticaceae)
VL  - 61
ID  - 1578
ER  - 


TY  - JOUR
AU  - Hadiah, Julisasi T.
AU  - Quinn, Christopher J.
AU  - Conn, Barry J.
IS  - 1
KW  - Urticaceae
PY  - 2003
SN  - 03129764
SP  - 235-246
ST  - Phylogeny of Elatostema (Urticaceae) using chloroplast DNA data
T2  - Telopea
TI  - Phylogeny of Elatostema (Urticaceae) using chloroplast DNA data
VL  - 10
ID  - 3986
ER  - 


TY  - JOUR
AU  - Hadiah, Julisasi T.
AU  - Conn, Barry J.
AU  - Quinn, Christopher J.
DO  - 10.1071/SB08041 M4 - Citavi
IS  - 5
KW  - Urticaceae
N1  - PII: SB08041 PII: SB08041 PII: SB08041 PII: SB08041
PY  - 2008
SN  - 1030-1887
SP  - 375-385
ST  - Infra-familial phylogeny of Urticaceae, using chloroplast sequence data
T2  - Australian Systematic Botany
TI  - Infra-familial phylogeny of Urticaceae, using chloroplast sequence data
VL  - 21
ID  - 2823
ER  - 


TY  - JOUR
AB  - Mutualisms could be evolutionarily unstable, with changes in partner abundances or in the spatial context of interactions potentially promoting their dissolution. We test this prediction using the defense mutualisms between species of the Neotropical genus Cecropia and Azteca ants. A new, multigene phylogeny with representatives of all five genera of Cecropieae (most of them from the Neotropics) and half of the 61 species of Cecropia shows the West African endemic Musanga (2spp.) as sister to Cecropia, implying dispersal from the Neotropics to Africa, with a molecular clock suggesting that this occurred about 23Mya. Cecropia, a genus of neotropical pioneer trees, started diversifying ca. 8Mya. We infer a single origin of specialized symbiosis with Azteca within Cecropia, eight complete losses of this symbiosis, and a potential partner shift involving the replacement of Azteca by Neoponera luteola ants. Niche space modeling based on geo-referenced occurrences of over 9000 collections representing 58 of the 61 species of Cecropia, together with several comparative analyses, implies that mutualism loss is concentrated at high altitudes and on Caribbean islands, with the surprisingly frequent breakdowns potentially facilitated by low species-specificity of interacting Cecropia and Azteca mutualists.
AD  - Systematic Botany and Mycology, University of Munich (LMU), Menzinger Str. 67, 80638 Munich, Germany. Systematic Botany and Mycology, University of Munich (LMU), Menzinger Str. 67, 80638 Munich, Germany. Electronic address:. Systematic Botany and Mycology, University of Munich (LMU), Menzinger Str. 67, 80638 Munich, Germany. M4 - Citavi
AU  - Gutiérrez-Valencia, Juanita
AU  - Chomicki, Guillaume
AU  - Renner, Susanne S.
DO  - 10.1016/j.ympev.2017.04.009
KW  - Urticaceae
LA  - eng
N1  - Journal Article Journal Article Journal Article Journal Article
PY  - 2017
SN  - 1055-7903
SP  - 196-205
ST  - Recurrent breakdowns of mutualisms with ants in the neotropical ant-plant genus Cecropia (Urticaceae)
T2  - Molecular phylogenetics and evolution
TI  - Recurrent breakdowns of mutualisms with ants in the neotropical ant-plant genus Cecropia (Urticaceae)
VL  - 111
ID  - 75
ER  - 


TY  - JOUR
AD  - Nees-Institut für Biodiversität der Pflanzen; Rheinische Friedrich-Wilhelms-Universität; Meckenheimer Allee 170 D-53115 Bonn Germany M4 - Citavi
AU  - Grosse-Veldmann, Bernadette
AU  - Weigend, Maximilian
DO  - 10.1111/cla.12193
IS  - 2
KW  - Urticaceae
PY  - 2018
SN  - 07483007
SP  - 131-150
ST  - The geometry of gender: hyper-diversification of sexual systems in Urtica L. (Urticaceae)
T2  - Cladistics
TI  - The geometry of gender: hyper-diversification of sexual systems in Urtica L. (Urticaceae)
VL  - 34
ID  - 14
ER  - 


TY  - JOUR
AB  - The genus Urtica L. is subcosmopolitan, found on all continents (except Antarctica) and most extratropical islands and ranges from Alaska to Patagonia, Spitzbergen to the Cape and Camtschatka to the subantarctic islands. However, throughout its geographical range morphologically nearly indistinguishable species are found alongside morphologically quite disparate species, with the overall diversity of morphological characters extremely limited. The systematics of Urtica have puzzled scientists for the past 200years and no single comprehensive attempt at understanding infrageneric relationships has been published in the past, nor are species delimitations unequivocally established. We here provide the first comprehensive phylogeny of the genus including 61 of the 63 species recognized, represented by 144 ingroup accessions and 14 outgroup taxa. The markers ITS1-5.8S-ITS2, psbA-trnH intergenic spacer, trnL-trnF and trnS-trnG are used. The phylogeny is well resolved. The eastern Asian Zhengyia shennongensis T. Deng, D.G. Zhang & H. Sun is retrieved as sister to Urtica. Within Urtica, a clade comprising the western Eurasian species U. pilulifera L. and U. neubaueri Chrtek is sister to all other species of the genus. The phylogenetic analyses retrieve numerous well-supported clades, suggesting previously unsuspected relationships and implying that classically used taxonomic characters such as leaf morphology and growth habit are highly homoplasious. Species delimitation is problematical, and several accessions assigned to Urtica dioica L. (as subspecies) are retrieved in widely different places in the phylogeny. The genus seems to have undergone numerous dispersal-establishment events both between continents and onto different islands. Three recent species radiations are inferred, one in America centered in the Andes, one in New Zealand, and one in northern Eurasia which includes Urtica dioica s.str. sensu Henning et al. (2014). The present study provides the basis of a critical re-examination of species limits and taxonomy, but also of the dispersal ecology of this widespread plant group and an in-depth study of the three clades with recent radiations.
AD  - Nees-Institut für Biodiversität der Pflanzen, Rheinische Friedrich-Wilhelms-Universität, Meckenheimer Allee 170, D-53115 Bonn, Germany. Centre for Organismal Studies Heidelberg, Ruprecht-Karls-Universität, Im Neuenheimer Feld 345, D-69120 Heidelberg, Germany. Allan Herbarium, Landcare Research, PO Box 69040, Lincoln 7640, New Zealand. Allan Herbarium, Landcare Research, PO Box 69040, Lincoln 7640, New Zealand. Nees-Institut für Biodiversität der Pflanzen, Rheinische Friedrich-Wilhelms-Universität, Meckenheimer Allee 170, D-53115 Bonn, Germany. Nees-Institut für Biodiversität der Pflanzen, Rheinische Friedrich-Wilhelms-Universität, Meckenheimer Allee 170, D-53115 Bonn, Germany. Electronic address:. M4 - Citavi
AU  - Grosse-Veldmann, Bernadette
AU  - Nürk, Nicolai M.
AU  - Smissen, Rob
AU  - Breitwieser, Ilse
AU  - Quandt, Dietmar
AU  - Weigend, Maximilian
DO  - 10.1016/j.ympev.2016.05.019
KW  - Urticaceae
LA  - eng
N1  - Journal Article Journal Article Journal Article Journal Article
PY  - 2016
SN  - 1055-7903
SP  - 9-19
ST  - Pulling the sting out of nettle systematics - A comprehensive phylogeny of the genus Urtica L. (Urticaceae)
T2  - Molecular phylogenetics and evolution
TI  - Pulling the sting out of nettle systematics - A comprehensive phylogeny of the genus Urtica L. (Urticaceae)
VL  - 102
ID  - 231
ER  - 


TY  - JOUR
AU  - Denk, Thomas
AU  - Grimm, Guido W.
KW  - Ulmaceae
PY  - 2005
SP  - 129-157
ST  - Phylogeny and biogeography of Zelkova (Ulmaceae sensu stricto) as inferred from leaf morphology, ITS sequence data and the fossil record
T2  - Botanical Journal of the Linnean Society
TI  - Phylogeny and biogeography of Zelkova (Ulmaceae sensu stricto) as inferred from leaf morphology, ITS sequence data and the fossil record
VL  - 147
ID  - 3577
ER  - 


TY  - JOUR
AB  - UNLABELLED PREMISE OF THE STUDY Sparganium (Typhaceae) is a genus of aquatic monocots containing ±14 species, with flowers aggregated in unisexual, spherical heads, and habit ranging from floating to emergent. Sparganium presents an opportunity to investigate diversification, character evolution, and biogeographical relationships in a widespread temperate genus of aquatic monocots. We present a fossil-calibrated, molecular phylogeny of Sparganium based on analysis of two chloroplast and two nuclear markers. Within this framework, we examine character evolution in both habit and stigma number and infer the ancestral area and biogeographic history of the genus. • METHODS Sequence data from two cpDNA and two nDNA markers were analyzed using maximum parsimony, maximum likelihood, and Bayesian inference. We used the program BEAST to simultaneously estimate phylogeny and divergence times, S-DIVA and Lagrange for biogeographical reconstruction, and BayesTraits to examine locule number and habit evolution. • KEY RESULTS Two major clades were recovered with strong support: one composed of S. erectum and S. eurycarpum; and the other containing all remaining Sparganium. We realigned the subgenera to conform to these clades. Divergence time analysis suggests a Miocene crown origin but Pliocene diversification. Importantly, the floating-leaved habit has arisen multiple times in the genus, from emergent ancestors-contrary to past hypotheses. • CONCLUSIONS Cooling trends during the Tertiary are correlated with the isolation of temperate Eurasian and North American taxa. Vicariance, long-distance dispersal, and habitat specialization are proposed as mechanisms for Sparganium diversification.
AD  - Department of Botany, University of Wisconsin, Madison, Wisconsin 53706 USA. M4 - Citavi
AU  - Sulman, Joshua D.
AU  - Drew, Bryan T.
AU  - Drummond, Chloe
AU  - Hayasaka, Eisuke
AU  - Sytsma, Kenneth J.
DO  - 10.3732/ajb.1300048
IS  - 10
KW  - Typhaceae
LA  - eng
N1  - Journal Article Research Support, Non-U.S. Gov't Journal Article Research Support, Non-U.S. Gov't Journal Article Research Support, Non-U.S. Gov't Journal Article Research Support, Non-U.S. Gov't
PY  - 2013
SN  - 0002-9122
SP  - 2023-2039
ST  - Systematics, biogeography, and character evolution of Sparganium (Typhaceae): diversification of a widespread, aquatic lineage
T2  - American journal of botany
TI  - Systematics, biogeography, and character evolution of Sparganium (Typhaceae): diversification of a widespread, aquatic lineage
VL  - 100
ID  - 1399
ER  - 


TY  - JOUR
AU  - Kim, Changkyun
AU  - Choi, Hong-Keun
IS  - 5
KW  - Typhaceae
PY  - 2011
SN  - 00400262
SP  - 1417-1428
ST  - Molecular systematics and character evolution of Typha (Typhaceae) inferred from nuclear and plastid DNA sequence data
T2  - Taxon
TI  - Molecular systematics and character evolution of Typha (Typhaceae) inferred from nuclear and plastid DNA sequence data
VL  - 60
ID  - 1932
ER  - 


TY  - JOUR
AU  - Ito, Yu
AU  - Tanaka, Norio
AU  - Kim, Changkyun
AU  - Kaul, Robert B.
AU  - Albach, Dirk C.
DO  - 10.1007/s00606-015-1245-7 M4 - Citavi
IS  - 1
KW  - Typhaceae
N1  - PII: 1245 PII: 1245 PII: 1245 PII: 1245
PY  - 2016
SN  - 0378-2697
SP  - 129-135
ST  - Phylogeny of Sparganium (Typhaceae) revisited: non-monophyletic nature of S. emersum sensu lato and resurrection of S. acaule
T2  - Plant Systematics and Evolution
TI  - Phylogeny of Sparganium (Typhaceae) revisited: non-monophyletic nature of S. emersum sensu lato and resurrection of S. acaule
VL  - 302
ID  - 250
ER  - 


TY  - JOUR
AB  - Turnera provides a useful system for exploring two significant evolutionary phenomena-shifts in breeding system (distyly vs. homostyly) and the evolution of polyploids. To explore these, the first molecular phylogeny of Turnera was constructed using sequences of the internal transcribed spacer region (ITS) of nuclear ribosomal DNA for 37 taxa. We attempted to resolve the origins of allopolyploid species using single-strand conformation polymorphism and sequencing of homeologous copies of ITS. Two allohexaploid species possessed putative ITS homeologues (T. velutina and T. orientalis). A phylogenetic analysis to identify progenitors contributing to the origins of these polyploids was unsuccessful, possibly as a result of concerted evolution of ITS. Breeding system evolution was mapped onto the phylogeny assuming distyly to be ancestral in Turnera. Self-compatible homostyly appears to have arisen independently at least three times in Turnera; however, we were not able to determine whether there have been independent origins of homostyly among hexaploid species in series Turnera. Our phylogenetic analyses suggest that series Turnera is monophyletic. Neither series Microphyllae nor Anomalae, however, appear to be monophyletic. Future taxonomic revisions may require new circumscriptions of these latter series.
AD  - Department of Biology, York University, 4700 Keele Street, Toronto, Ontario M3J 1P3 Canada; M4 - Citavi
AU  - Truyens, Simon
AU  - Arbo, María M.
AU  - Shore, Joel S.
DO  - 10.3732/ajb.92.10.1749
IS  - 10
KW  - Turneraceae
LA  - eng
N1  - Journal Article Journal Article Journal Article Journal Article
PY  - 2005
SN  - 0002-9122
SP  - 1749-1758
ST  - Phylogenetic relationships, chromosome and breeding system evolution in Turnera (Turneraceae): inferences from its sequence data
T2  - American journal of botany
TI  - Phylogenetic relationships, chromosome and breeding system evolution in Turnera (Turneraceae): inferences from its sequence data
VL  - 92
ID  - 3713
ER  - 


TY  - JOUR
AD  - Department of Systematic Biology, EBC; Uppsala University; Norbyvägen 18D 752 36 Uppsala Sweden Bergius Foundation; The Royal Swedish Academy of Sciences and Botany Department, Stockholm University; 106 91 Stockholm Sweden M4 - Citavi
AU  - Thulin, Mats
AU  - Razafimandimbison, Sylvain G.
AU  - Chafe, Paul
AU  - Heidari, Nahid
AU  - Kool, Anneleen
AU  - Shore, Joel S.
DO  - 10.1002/tax.612003 Y3 - 18.02.2019
IS  - 2
KW  - Turneraceae
PY  - 2012
SN  - 00400262
SP  - 308-323
ST  - Phylogeny of the Turneraceae clade (Passifloraceae s.l.): Trans-Atlantic disjunctions and two new genera in Africa
T2  - Taxon
TI  - Phylogeny of the Turneraceae clade (Passifloraceae s.l.): Trans-Atlantic disjunctions and two new genera in Africa
VL  - 61
ID  - 1737
ER  - 


TY  - JOUR
AD  - Instituto de Botánica del Nordeste (UNNE-CONICET); C.C. 209 Corrientes 3400 Argentina Facultad de Ciencias Exactas y Naturales; Universidad de Buenos Aires; Argentina M4 - Citavi
AU  - Mercedes Arbo, Maria
AU  - Espert, Shirley M.
DO  - 10.1002/tax.582011 Y3 - 18.02.2019
IS  - 2
KW  - Turneraceae
PY  - 2009
SN  - 00400262
SP  - 457-467
ST  - Morphology, phylogeny and biogeography of Turnera L. (Turneraceae)
T2  - Taxon
TI  - Morphology, phylogeny and biogeography of Turnera L. (Turneraceae)
VL  - 58
ID  - 2599
ER  - 


TY  - JOUR
AD  - Instituto de Botánica Darwinion (CONICET – ANCEFN), Labardén 200, CC 22, B1642HYD, San Isidro, Buenos Aires, Argentina. Instituto de Botánica del Nordeste (UNNE – CONICET), Sargento Cabral 2131, CC 209, 3400, Corrientes, Argentina. Department of Biology, York University, 4700 Keele Street, Toronto, ON M3J 1P3, Canada. M4 - Citavi
AU  - López, Alicia
AU  - Fernández, Aveliano
AU  - Shore, Joel S.
DO  - 10.1139/cjb-2012-0135
IS  - 3
KW  - Turneraceae
PY  - 2013
SN  - 1916-2790
SP  - 167-175
ST  - Inferences on the origins of polyploid Turnera species (Passifloraceae) based on molecular data
T2  - Botany
TI  - Inferences on the origins of polyploid Turnera species (Passifloraceae) based on molecular data
VL  - 91
ID  - 1299
ER  - 


TY  - JOUR
AU  - Arbo, María M.
AU  - Gonzalez, Ana M.
AU  - Sede, Silvana M.
DO  - 10.1007/s00606-015-1204-3 M4 - Citavi
IS  - 7
KW  - Turneraceae
N1  - PII: 1204 PII: 1204 PII: 1204 PII: 1204
PY  - 2015
SN  - 0378-2697
SP  - 1907-1926
ST  - Phylogenetic relationships within Turneraceae based on morphological characters with emphasis on seed micromorphology
T2  - Plant Systematics and Evolution
TI  - Phylogenetic relationships within Turneraceae based on morphological characters with emphasis on seed micromorphology
VL  - 301
ID  - 431
ER  - 


TY  - JOUR
AU  - Hershkovitz, Mark A.
AU  - Hernández-Pellicer, C. C.
AU  - Arroyo, Mary T. K.
DO  - 10.1007/s00606-006-0428-7 M4 - Citavi
IS  - 1
KW  - Tropaeolaceae
N1  - PII: 428 PII: 428 PII: 428 PII: 428
PY  - 2006
SN  - 0378-2697
SP  - 1-24
ST  - Ribosomal DNA evidence for the diversification of Tropaeolum sect. Chilensia (Tropaeolaceae)
T2  - Plant Systematics and Evolution
TI  - Ribosomal DNA evidence for the diversification of Tropaeolum sect. Chilensia (Tropaeolaceae)
VL  - 260
ID  - 3390
ER  - 


TY  - JOUR
AU  - Andersson, Lennart
AU  - Andersson, Stephan
IS  - 2
KW  - Tropaeolaceae
PY  - 2000
SN  - 00400262
SP  - 721-736
ST  - A Molecular Phylogeny of Tropaeolaceae and its Systematic Implications
T2  - Taxon
TI  - A Molecular Phylogeny of Tropaeolaceae and its Systematic Implications
VL  - 49
ID  - 4308
ER  - 


TY  - JOUR
AU  - Ren, Y.
AU  - Chen, L.
AU  - Tian, X. H.
AU  - Zhang, X. H.
AU  - Lu, A. M.
DO  - 10.1007/s00606-007-0563-9 M4 - Citavi
IS  - 1-4
KW  - Trochodendraceae
N1  - PII: 563 PII: 563 PII: 563 PII: 563
PY  - 2007
SN  - 0378-2697
SP  - 155-161
ST  - Discovery of vessels in Tetracentron (Trochodendraceae) and its systematic significance
T2  - Plant Systematics and Evolution
TI  - Discovery of vessels in Tetracentron (Trochodendraceae) and its systematic significance
VL  - 267
ID  - 3238
ER  - 


TY  - JOUR
AB  - The mycoheterotrophic plant family Triuridaceae (Pandanales) is hypothesized to be an old family, mainly based on its pantropical distribution. The existence of fossils from the Upper Cretaceous, assigned to Triuridaceae may form additional support for a great age of the family, although the affinity of these fossils to Triuridaceae is questioned. Although the circumscription of Triuridaceae has never been problematic, probably due to its distinct morphological characters, its systematic relationship has been under debate since the family was described around 1840. The lack of synapomorphies suitable for resolving higher taxonomic relationships is a function of the family's reduced vegetative growth and the highly modified floral structures. Molecular studies have assigned Triuridaceae to Pandanales, but its exact phylogenetic position remains unknown. In the present study the phylogeny of the Pandanales was reconstructed using four molecular markers and the divergence age estimates were obtained with a relaxed molecular clock method. We found that Triuridaceae are monophyletic and most likely descent form the second major split in Pandanales. The relationships between the other Pandanales families (Cyclanthaceae, Pandanaceae, Stemonaceae and Velloziaceae) are otherwise in accordance with earlier studies. Velloziaceae are sister to the rest of the Pandanales, Stemonaceae are most likely sister to a clade consisting of Pandanaceae and Cyclanthaceae, and the latter two families are sister to each other. All currently recognized tribes within Triuridaceae are also monophyletic at current taxon sampling. We estimate that the family has a Cretaceous (or Lower Paleocene) stem age, which is in accordance with earlier predictions. This old age, along with elevated mutation rates indicated by long branch lengths and the family's mycoheterotrophic lifestyle, might account for the substantial morphological differences between Triuridaceae and its closest relatives.
AD  - Naturalis Biodiversity Center, Leiden University, PO Box 9517, Leiden, The Netherlands. Electronic address:. M4 - Citavi
AU  - Mennes, Constantijn B.
AU  - Smets, Erik F.
AU  - Moses, Sainge N.
AU  - Merckx, Vincent S. F. T.
DO  - 10.1016/j.ympev.2013.05.031
IS  - 3
KW  - Triuridaceae
LA  - eng
N1  - Journal Article Journal Article Journal Article Journal Article
PY  - 2013
SN  - 1055-7903
SP  - 994-1004
ST  - New insights in the long-debated evolutionary history of Triuridaceae (Pandanales)
T2  - Molecular phylogenetics and evolution
TI  - New insights in the long-debated evolutionary history of Triuridaceae (Pandanales)
VL  - 69
ID  - 1312
ER  - 


TY  - JOUR
AU  - Cantino, Philip D.
AU  - Doyle, James A.
AU  - W.Graham, Sean
AU  - S.Judd, Walter
AU  - Olmstead, Richard G.
AU  - Soltis, Douglas E.
AU  - S.Soltis, Pamela
AU  - Donoghue, Michael J.
IS  - 3
KW  - Tracheophyta
PY  - 2007
SN  - 00400262
SP  - 1-44
ST  - Towards a phylogenetic nomenclature of Tracheophyta
T2  - Taxon
TI  - Towards a phylogenetic nomenclature of Tracheophyta
VL  - 56
ID  - 3064
ER  - 


TY  - JOUR
AU  - Tamura, Minoru N.
AU  - Fuse, Shizuka
AU  - Lee, Nam Sook
AU  - Kim, Jin Ohk
AU  - Yamashita, Jun
AU  - Ishii, Takaaki
IS  - 5
KW  - Tofieldiaceae
PY  - 2011
SN  - 00400262
SP  - 1339-1348
ST  - Biosystematic studies on the family Tofieldiaceae III. Classification of Tofieldia nuda into three species and three varieties
T2  - Taxon
TI  - Biosystematic studies on the family Tofieldiaceae III. Classification of Tofieldia nuda into three species and three varieties
VL  - 60
ID  - 2049
ER  - 


TY  - JOUR
AB  - In order to specify phylogenetic positions of the genera of Tofieldiaceae (Tofieldia, Triantha, Pleea, Harperocallis, Isidrogalvia), and to suggest reasonable circumscription of the family and genera of Tofieldiaceae, we determined DNA sequences of matK and rbcL for each genus of the family, and analyzed them phylogenetically with the 45 families and 113 genera of the monocots other than Tofieldiaceae, whose matK and rbcL sequences have already been reported. We found that Tofieldia, Triantha, Pleea, and Harperocallis form the same clade, which receives 100% bootstrap support. This clade can be regarded as corresponding to Tofieldiaceae, and is embedded in the clade of Alismatales (98%). On the other hand, Isidrogalvia is not included in this Tofieldiaceae clade, and positioned as sister to Narthecium (100%), embedded in the clade of Nartheciaceae (Dioscoreales) (100%). In the Tofieldiaceae, Pleea first diverges from the remaining three genera (100%), and then, Harperocallis diverges from the Tofieldia- Triantha complex (100%). In the Tofieldia- Triantha complex, five Tofieldia species form the same clade (100%), and two Triantha species form another clade (100%). Thus, Isidrogalvia should be transferred from Tofieldiaceae to Nartheciaceae. As Isidrogalvia, as well as the Nartheciaceae, have the carpels that are fully connate into a single style, Isidrogalvia fits the Nartheciaceae well with respect to carpel connation. After this transfer, the Tofieldiaceae correspond mainly to plants with almost free carpels and three styles. Pleea is better treated as an independent genus than included in Tofieldia. Triantha can be treated either as an independent genus or as congeneric with Tofieldia.
AD  - Botanical Gardens, Graduate School of Science, Osaka City University, 2000 Kisaichi, Katano-shi, Osaka 576-0004, Japan. M4 - Citavi
AU  - Tamura, M. N.
AU  - Fuse, Shizuka
AU  - Azuma, H.
AU  - Hasebe, M.
DO  - 10.1055/s-2004-821278
IS  - 5
KW  - Tofieldiaceae
LA  - eng
N1  - Journal Article Research Support, Non-U.S. Gov't Journal Article Research Support, Non-U.S. Gov't Journal Article Research Support, Non-U.S. Gov't Journal Article Research Support, Non-U.S. Gov't
PY  - 2004
SN  - 1435-8603
SP  - 562-567
ST  - Biosystematic studies on the family Tofieldiaceae I. Phylogeny and circumscription of the family inferred from DNA sequences of matK and rbcL
T2  - Plant Biology
TI  - Biosystematic studies on the family Tofieldiaceae I. Phylogeny and circumscription of the family inferred from DNA sequences of matK and rbcL
VL  - 6
ID  - 3911
ER  - 


TY  - JOUR
AU  - Tamura, Minoru N.
AU  - Azuma, Hiroshi
AU  - Yamashita, Jun
AU  - Fuse, Shizuka
AU  - Ishii, Takaaki
IS  - 3
KW  - Tofieldiaceae
PY  - 2010
SP  - 131-140
ST  - Biosystematic studies on the Family Tofieldiaceae II. Phylogeny of Species of Tofieldia and Triantha inferred from PIastid and Nuclear DNA Sequences
T2  - Acta Phytotaxonomica Geobotanica
TI  - Biosystematic studies on the Family Tofieldiaceae II. Phylogeny of Species of Tofieldia and Triantha inferred from PIastid and Nuclear DNA Sequences
VL  - 60
ID  - 2364
ER  - 


TY  - JOUR
AU  - Remizowa, Margarita V.
AU  - Sokoloff, Dmitry D.
AU  - Campbell, Lisa M.
AU  - Stevenson, Dennis W.
AU  - Rudall, Paula J.
IS  - 4
KW  - Tofieldiaceae
PY  - 2011
SN  - 00400262
SP  - 1076-1094
ST  - Harperocallis is congeneric with Isidrogalvia (Tofieldiaceae, Alismatales): Evidence from comparative floral morphology
T2  - Taxon
TI  - Harperocallis is congeneric with Isidrogalvia (Tofieldiaceae, Alismatales): Evidence from comparative floral morphology
VL  - 60
ID  - 2003
ER  - 


TY  - JOUR
AB  - Ten new combinations from Asagraea, Isidrogaliva, and Tofieldia are proposed in the previously monospecific genus Harperocallis (Tofieldiaceae, Alismatales). As circumscribed here, the genus is restricted to the Americas. The majority of species occur in the Andes or the Guayana region of northern South America; more than half have restricted distributions, and Harperocallis flava is narrowly endemic in the Coastal Plain of the southeastern United States. A key to species, synonymies, distributions, representative specimens, and salient notes are presented. Populations of the species are mapped and Harperocallis robustior is illustrated. A neotype is selected for Tofieldia frigida, here considered a synonym of Harperocallis falcata. Several recent records of Harperocallis longiflora, previously known only from the type collected in 1902, are reported. ResúmenSe proponen diez nuevas combinaciones de Asagraea, Isidrogaliva, y Tofieldia en Harperocallis (Tofieldiaceae, Alismatales), un género previamente monoespecífico. Como se circunscribe aquí, el género se restringe a las Américas. La mayoría de las especies se encuentran en los Andes o en la región de Guayana en el norte de América del Sur. Más de la mitad tienen distribuciones restringidas, y Harperocallis flava es estrictamente endémica en la llanura costera del sudeste de los Estados Unidos. Se presenta una clave para distinguir las especies, sinonimias, distribuciones, muestras representativas y notas salientes. Se mapearon poblaciones de las especies y se ilustra Harperocallis robustior. Se seleccionó un neotipo para Tofieldia falcata, considerado aquí como un sinónimo de Harperocallis falcata. Se reportan algunos registros adicionales de Harperocallis longiflora, especie anteriormente conocida sólo con el tipo colectado en 1902.
AD  - The New York Botanical Garden, Bronx, New York 10458, USA. M4 - Citavi
AU  - Campbell, Lisa M.
AU  - Dorr, Laurence J.
DO  - 10.3897/phytokeys.21.4859
IS  - 21
KW  - Tofieldiaceae
LA  - eng
N1  - Journal Article Journal Article Journal Article Journal Article
PY  - 2013
SN  - 1314-2011
SP  - 37-52
ST  - A synopsis of Harperocallis (Tofieldiaceae, Alismatales) with ten new combinations
T2  - PhytoKeys
TI  - A synopsis of Harperocallis (Tofieldiaceae, Alismatales) with ten new combinations
UR  - https://www.ncbi.nlm.nih.gov/pmc/articles/PMC3689118
ID  - 1185
ER  - 


TY  - JOUR
AB  - Tofieldiaceae are a small monocot family comprising about 20 species, mostly distributed in the Northern Hemisphere, with some in northern South America. To clarify the family circumscription, the number of distinguishable genera in the family, and relationships among the genera, we conducted molecular analyses of cpDNA (matK and non-coding trnL-trnL-trnF region) sequences of 17 associated species of Tofieldiaceae, along with 14 species of Acorales, Alismatales, Dioscoreales, Pandanales, and Liliales. Results showed that Tofieldiaceae are monophyletic, comprising all the species assignable to Harperocallis, Isidrogalvia, Pleea, Tofieldia, and Triantha, thus supporting the original family circumscription. Within the family, Pleea is sister to the rest of the family, in which Isidrogalvia is sister to Harperocallis, and Tofieldia to Triantha. Morphological characters supporting the relationships among the genera were briefly discussed.
AD  - Department of Botany, Graduate School of Science, Kyoto University, Kyoto, 606-8502, Japan. M4 - Citavi
AU  - Azuma, Hiroshi
AU  - Tobe, Hiroshi
DO  - 10.1007/s10265-010-0387-5
IS  - 3
KW  - Tofieldiaceae
LA  - eng
N1  - Journal Article Research Support, Non-U.S. Gov't Journal Article Research Support, Non-U.S. Gov't Journal Article Research Support, Non-U.S. Gov't Journal Article Research Support, Non-U.S. Gov't
PY  - 2011
SP  - 349-357
ST  - Molecular phylogenetic analyses of Tofieldiaceae (Alismatales): family circumscription and intergeneric relationships
T2  - Journal of Plant Research
TI  - Molecular phylogenetic analyses of Tofieldiaceae (Alismatales): family circumscription and intergeneric relationships
VL  - 124
ID  - 1805
ER  - 


TY  - JOUR
AU  - Yousefzadeh, Hamed
AU  - Hosseinzadeh Colagar, Abasalt
AU  - Tabari, Masoud
AU  - Sattarian, Ali
AU  - Assadi, Mostafa
DO  - 10.1007/s00606-012-0604-x M4 - Citavi
IS  - 5
KW  - Tiliaceae
N1  - PII: 604 PII: 604 PII: 604 PII: 604
PY  - 2012
SN  - 0378-2697
SP  - 947-961
ST  - Utility of ITS region sequence and structure for molecular identification of Tilia species from Hyrcanian forests, Iran
T2  - Plant Systematics and Evolution
TI  - Utility of ITS region sequence and structure for molecular identification of Tilia species from Hyrcanian forests, Iran
VL  - 298
ID  - 1779
ER  - 


TY  - JOUR
AB  - Tilia is an ecologically and economically important genus in the family Malvaceae. However, there is no complete plastid genome of Tilia sequenced to date, and the taxonomy of Tilia is difficult owing to frequent hybridization and polyploidization. A well-supported interspecific relationships of this genus is not available due to limited informative sites from the commonly used molecular markers. We report here the complete plastid genome sequences of four Tilia species determined by the Illumina technology. The Tilia plastid genome is 162,653 bp to 162,796 bp in length, encoding 113 unique genes and a total number of 130 genes. The gene order and organization of the Tilia plastid genome exhibits the general structure of angiosperms and is very similar to other published plastid genomes of Malvaceae. As other long-lived tree genera, the sequence divergence among the four Tilia plastid genomes is very low. And we analyzed the nucleotide substitution patterns and the evolution of insertions and deletions in the Tilia plastid genomes. Finally, we build a phylogeny of the four sampled Tilia species with high supports using plastid phylogenomics, suggesting that it is an efficient way to resolve the phylogenetic relationships of this genus.
AD  - Germplasm Bank of Wild Species, Kunming Institute of Botany, Chinese Academy of Sciences, Kunming 650201, Yunnan, China. School of Life Science, Yunnan University, Kunming 650091, Yunnan, China. Kunming College of Life Science, University of Chinese Academy of Sciences, Kunming 650204, China. Germplasm Bank of Wild Species, Kunming Institute of Botany, Chinese Academy of Sciences, Kunming 650201, Yunnan, China. Germplasm Bank of Wild Species, Kunming Institute of Botany, Chinese Academy of Sciences, Kunming 650201, Yunnan, China. Germplasm Bank of Wild Species, Kunming Institute of Botany, Chinese Academy of Sciences, Kunming 650201, Yunnan, China. M4 - Citavi
AU  - Cai, Jie
AU  - Ma, Peng-Fei
AU  - Li, Hong-Tao
AU  - Li, De-Zhu
DO  - 10.1371/journal.pone.0142705
IS  - 11
KW  - Tiliaceae
LA  - eng
N1  - Comparative Study Journal Article Research Support, Non-U.S. Gov't Comparative Study Journal Article Research Support, Non-U.S. Gov't Comparative Study Journal Article Research Support, Non-U.S. Gov't Comparative Study Journal Article Research Support, Non-U.S. Gov't
PY  - 2015
SP  - e0142705
ST  - Complete Plastid Genome Sequencing of four Tilia Species (Malvaceae): A comparative Analysis and phylogenetic Implications
T2  - PloS one
TI  - Complete Plastid Genome Sequencing of four Tilia Species (Malvaceae): A comparative Analysis and phylogenetic Implications
UR  - https://www.ncbi.nlm.nih.gov/pmc/articles/PMC4643919
VL  - 10
ID  - 469
ER  - 


TY  - JOUR
AB  - Historic events such as the uplift of Qinghai-Tibet Plateau (Q-T Plateau) and climatic oscillations in the Quaternary period greatly affected the evolution and modern distribution of Sino-Tibetan flora. Stellera chamaejasme, a perennial herb with flower color polymorphism that is distributed from the mountainous southeastern Q-T Plateau (Hengduan Mountains, H-D Mountains) to the vast platform of the Q-T Plateau and the adjacent plain of northern China, provides an excellent model to explore the effects of historic events on the origination and variation of species. In this study, we conducted a phylogenetic and phylogeographical study using three chloroplast sequences (trnT-L, trnL-F and rpL16) in 26 populations of S. chamaejasme and 12 outgroups from the Thymeleaceae. Phylogenetic analysis and molecular clock estimation revealed that the monophyletic origin of S. chamaejasme occurred ca. 6.5892 Ma, which is consistent with the radical environment changes caused by the rapid uplift of the Q-T Plateau ca. 7 Ma. Intra-specific differentiation of S. chamaejasme is estimated to have occurred after ca. 2.1 Ma. Twelve haplotypes were revealed from combined trnL-F and rpL16 sequences. High genetic diversity (h(T)=0.834) and population differentiation (N(ST)=0.997 and G(ST)=0.982) imply restricted gene flow among populations and significant geographical or environmental isolation. All populations from the vast plain of northern China were dominated by one haplotype (H1), and the same haplotype was fixed in most populations from the high elevation platform of the western and northern Q-T Plateau. In contrast, the majority of the haplotypes were found in the relatively narrow area of the H-D Mountains, in the southeastern distribution of S. chamaejasme. The contrasting haplotype distribution patterns suggested that the H-D Mountains were either a refugium for S. chamaejasme during the Quaternary climatic oscillations or a diversification center of this species. The present wide distribution of this species on the Q-T Plateau platform and in northern China is likely to have resulted from a rapid post-glacial population expansion from the southeastern refugium involving founder effects, facilitated by the adjacent geographic range with a similar grassland habitat.
AD  - Key Laboratory of Biodiversity and Biogeography, Kunming Institute of Botany, Chinese Academy of Sciences, Kunming, Yunnan 650204, People's Republic of China. M4 - Citavi
AU  - Zhang, Yong-Hong
AU  - Volis, Sergei
AU  - Sun, Hang
DO  - 10.1016/j.ympev.2010.08.033
IS  - 3
KW  - Thymelaeaceae
LA  - eng
N1  - Journal Article Research Support, Non-U.S. Gov't Journal Article Research Support, Non-U.S. Gov't Journal Article Research Support, Non-U.S. Gov't Journal Article Research Support, Non-U.S. Gov't
PY  - 2010
SN  - 1055-7903
SP  - 1162-1172
ST  - Chloroplast phylogeny and phylogeography of Stellera chamaejasme on the Qinghai-Tibet Plateau and in adjacent regions
T2  - Molecular phylogenetics and evolution
TI  - Chloroplast phylogeny and phylogeography of Stellera chamaejasme on the Qinghai-Tibet Plateau and in adjacent regions
VL  - 57
ID  - 2418
ER  - 


TY  - JOUR
AU  - van der Bank, Michelle
AU  - Fay, Michael F.
AU  - Chase, Mark W.
KW  - Thymelaeaceae
PY  - 2002
SN  - 00400262
SP  - 329-339
ST  - Molecular phylogenetics of Thymelaeaceae with particular reference to African and Australian genera
T2  - Taxon
TI  - Molecular phylogenetics of Thymelaeaceae with particular reference to African and Australian genera
VL  - 51
ID  - 4181
ER  - 


TY  - JOUR
AU  - Rogers, Zachary S.
DO  - 10.3417/2009069 M4 - Citavi
IS  - 4
KW  - Thymelaeaceae
PY  - 2010
SN  - 1055-3177
SP  - 448-462
ST  - Nomenclatural Notes on American Thymelaeaceae
T2  - Novon: A Journal for Botanical Nomenclature
TI  - Nomenclatural Notes on American Thymelaeaceae
VL  - 20
ID  - 2330
ER  - 


TY  - JOUR
AU  - Rogers, Zachary S.
DO  - 10.3417/2006114 M4 - Citavi
IS  - 2
KW  - Thymelaeaceae
PY  - 2009
SP  - 324-369
ST  - A Revision of Malagasy Gnidia (Thymelaeaceae, Thymelaeoideae)
T2  - Annals of the Missouri Botanical Garden
TI  - A Revision of Malagasy Gnidia (Thymelaeaceae, Thymelaeoideae)
VL  - 96
ID  - 2650
ER  - 


TY  - JOUR
AU  - Galicia-Herbada, D.
DO  - 10.1007/s00606-005-0371-z M4 - Citavi
IS  - 3-4
KW  - Thymelaeaceae
N1  - PII: 371 PII: 371 PII: 371 PII: 371
PY  - 2006
SN  - 0378-2697
SP  - 159-187
ST  - Origin and diversification of Thymelaea (Thymelaeaceae): inferences from a phylogenetic study based on ITS (rDNA) sequences
T2  - Plant Systematics and Evolution
TI  - Origin and diversification of Thymelaea (Thymelaeaceae): inferences from a phylogenetic study based on ITS (rDNA) sequences
VL  - 257
ID  - 3375
ER  - 


TY  - JOUR
AU  - Foster, Charles S. P.
AU  - Cantrill, David J.
AU  - James, Elizabeth A.
AU  - Syme, Anna E.
AU  - Jordan, Rebecca
AU  - Douglas, Rachel
AU  - Ho, Simon Y. W.
AU  - Henwood, Murray J.
DO  - 10.1071/SB16013 M4 - Citavi
IS  - 3
KW  - Thymelaeaceae
N1  - PII: SB16013 PII: SB16013 PII: SB16013 PII: SB16013
PY  - 2016
SN  - 1030-1887
SP  - 185-196
ST  - Molecular phylogenetics provides new insights into the systematics of Pimelea and Thecanthes (Thymelaeaceae)
T2  - Australian Systematic Botany
TI  - Molecular phylogenetics provides new insights into the systematics of Pimelea and Thecanthes (Thymelaeaceae)
VL  - 29
ID  - 217
ER  - 


TY  - JOUR
AU  - Floden, Aaron J.
AU  - Mayfield, Mark H.
AU  - Ferguson, Carolyn J.
IS  - 2
KW  - Thymelaeaceae
PY  - 2009
SP  - 485-499
ST  - A new narrowly endemic Species of Dirca (Thymelaeaceae) from Kansas and Arkansas, with a phylogenetic Overview and taxonomic Synopsis of the Genus
T2  - Journal of the Botanical Research Institute of Texas
TI  - A new narrowly endemic Species of Dirca (Thymelaeaceae) from Kansas and Arkansas, with a phylogenetic Overview and taxonomic Synopsis of the Genus
VL  - 3
ID  - 2510
ER  - 


TY  - JOUR
AU  - Eurlings, Marcel C. M.
AU  - Gravendeel, Barbara
DO  - 10.1007/s00606-005-0312-x M4 - Citavi
IS  - 1-2
KW  - Thymelaeaceae
N1  - PII: 312 PII: 312 PII: 312 PII: 312
PY  - 2005
SN  - 0378-2697
SP  - 1-12
ST  - TrnL-trnF sequence data imply paraphyly of Aquilaria and Gyrinops (Thymelaeaceae) and provide new perspectives for agarwood identification
T2  - Plant Systematics and Evolution
TI  - TrnL-trnF sequence data imply paraphyly of Aquilaria and Gyrinops (Thymelaeaceae) and provide new perspectives for agarwood identification
VL  - 254
ID  - 3584
ER  - 


TY  - JOUR
AU  - Bredenkamp, C. L.
AU  - van Wyk, A. E.
DO  - 10.4102/abc.v33i1.433 Y3 - 20.02.2019 M4 - Citavi
IS  - 1
KW  - Thymelaeaceae
PY  - 2003
SN  - 0006-8241
SP  - 59-98
ST  - Taxonomy of the genus Passerina (Thymelaeaceae)
T2  - Bothalia
TI  - Taxonomy of the genus Passerina (Thymelaeaceae)
VL  - 33
ID  - 3961
ER  - 


TY  - JOUR
AU  - Beaumont, Angela J.
AU  - Edwards, Trevor J.
AU  - Manning, John C.
AU  - Maurin, Olivier
AU  - Rautenbach, Marline
AU  - Motsi, Moleboheng C.
AU  - Fay, Michael F.
AU  - Chase, Mark W.
AU  - van der Bank, Michelle
DO  - 10.1111/j.1095-8339.2009.00988.x Y3 - 20.02.2019 M4 - Citavi
IS  - 4
KW  - Thymelaeaceae
PY  - 2009
SP  - 402-417
ST  - Gnidia (Thymelaeaceae) is not monophyletic: taxonomic implications for Thymelaeoideae and a partial new generic taxonomy for Gnidia
T2  - Botanical Journal of the Linnean Society
TI  - Gnidia (Thymelaeaceae) is not monophyletic: taxonomic implications for Thymelaeoideae and a partial new generic taxonomy for Gnidia
VL  - 160
ID  - 2442
ER  - 


TY  - JOUR
AU  - Wortley, A. H.
AU  - Harris, David J.
AU  - Scotland, R. W.
IS  - 2
KW  - Thomandersiaceae
PY  - 2007
SN  - 03636445
SP  - 415-444
ST  - On the Taxonomy and Phylogenetic Position of Thomandersia
T2  - Systematic Botany
TI  - On the Taxonomy and Phylogenetic Position of Thomandersia
VL  - 32
ID  - 3309
ER  - 


TY  - JOUR
AU  - Yahara, Tetsukazu
AU  - Tsukaya, Hirokazu
IS  - 2
KW  - Thismiaceae
PY  - 2008
SP  - 97-104
ST  - Oxygyne yamashitae, a new Species of Thismiaceae from Yaku Island, Japan
T2  - Acta Phytotaxonomica Geobotanica
TI  - Oxygyne yamashitae, a new Species of Thismiaceae from Yaku Island, Japan
VL  - 59
ID  - 3026
ER  - 


TY  - JOUR
AU  - Merckx, Vincent
AU  - Bakker, Freek T.
AU  - Huysmans, Suzy
AU  - Smets, Erik
DO  - 10.1111/j.1096-0031.2008.00241.x M4 - Citavi
IS  - 1
KW  - Thismiaceae
PY  - 2009
SN  - 07483007
SP  - 64-77
ST  - Bias and conflict in phylogenetic inference of myco-heterotrophic plants: a case study in Thismiaceae
T2  - Cladistics
TI  - Bias and conflict in phylogenetic inference of myco-heterotrophic plants: a case study in Thismiaceae
VL  - 25
ID  - 2600
ER  - 


TY  - JOUR
AU  - Nickrent, Daniel Lee
AU  - García, Miguel Angel
DO  - 10.11646/phytotaxa.224.2.4 M4 - Citavi
IS  - 2
KW  - Thesiaceae
PY  - 2015
SN  - 1179-3163
SP  - 173
ST  - Lacomucinaea, a new monotypic genus in Thesiaceae (Santalales)
T2  - Phytotaxa
TI  - Lacomucinaea, a new monotypic genus in Thesiaceae (Santalales)
VL  - 224
ID  - 649
ER  - 


TY  - JOUR
AU  - Moore, Timothy E.
AU  - Verboom, G. Anthony
AU  - Forest, Félix
KW  - Thesiaceae
PY  - 2010
SP  - 435-454
ST  - Phylogenetics and biogeography of the parasitic genus Thesium L. (Santalaceae), with an emphasis on the Cape of South Africa
T2  - Botanical Journal of the Linnean Society
TI  - Phylogenetics and biogeography of the parasitic genus Thesium L. (Santalaceae), with an emphasis on the Cape of South Africa
VL  - 162
ID  - 2286
ER  - 


TY  - JOUR
AU  - Ståhl, Bertil
AU  - Källersjö, Mari
IS  - 1
KW  - Theophrastaceae
PY  - 2004
SN  - 1055-3177
SP  - 115-118
ST  - Reinstatement of Bonellia (Theophrastaceae)
T2  - Novon: A Journal for Botanical Nomenclature
TI  - Reinstatement of Bonellia (Theophrastaceae)
VL  - 14
ID  - 3904
ER  - 


TY  - JOUR
AU  - Källersjö, Mari
AU  - Ståhl, Bertil
IS  - 4
KW  - Theophrastaceae
PY  - 2003
SN  - 1058-5893
SP  - 579-591
ST  - Phylogeny of Theophrastaceae (Ericales s. lat.)
T2  - International Journal of Plant Sciences
TI  - Phylogeny of Theophrastaceae (Ericales s. lat.)
VL  - 164
ID  - 3996
ER  - 


TY  - JOUR
AB  - Phylogenetic analyses of plastid DNA sequences of ndhF, trnL-F intron and spacer regions, and rpl16 are presented separately and combined for 41 taxa from all 12 genera of the Themidaceae and for 20 taxa from nine related families in the higher Asparagales. The results from the combined analysis are the most resolved and provide a high level of support for the monophyly of Themidaceae. Within Themidaceae, the Milla complex of Mexico is supported as monophyletic within a paraphyletic Brodiaea complex of western North America. Four major clades are identified in each of the individual and combined analyses: (1) the Milla complex; (2) Brodiaea, Dichelostemma, and Triteleiopsis; (3) Triteleia, Bloomeria, and Muilla clevelandii; and (4) Androstephium and the other species of Muilla. These well-defined clades suggest that morphological characters (e.g., an extended perianth tube) that have been traditionally used to circumscribe the genera within the Brodiaea complex have evolved independently at least twice. In addition, common biogeographic distribution patterns (e.g., Brodiaea and Triteleia having centers of diversity in northern California and the Pacific Northwest) appear to be the result of separate evolutionary radiations.
AD  - Department of Botany, University of Wisconsin, Madison, Wisconsin 53706-1381 USA. M4 - Citavi
AU  - Pires, J. Chris
AU  - Sytsma, Kenneth J.
DO  - 10.3732/ajb.89.8.1342
IS  - 8
KW  - Themidaceae
LA  - eng
N1  - Journal Article Journal Article Journal Article Journal Article
PY  - 2002
SN  - 0002-9122
SP  - 1342-1359
ST  - A phylogenetic evaluation of a biosystematic framework: Brodiaea and related petaloid monocots (Themidaceae)
T2  - American journal of botany
TI  - A phylogenetic evaluation of a biosystematic framework: Brodiaea and related petaloid monocots (Themidaceae)
VL  - 89
ID  - 4155
ER  - 


TY  - JOUR
AU  - Pires, J. Chris
AU  - Fay, Michael F.
AU  - Davis, Warren S.
AU  - Hufford, Larry
AU  - Rova, Johan
AU  - Chase, Mark W.
AU  - Sytsma, Kenneth J.
KW  - Themidaceae
PY  - 2001
SN  - 0075-5974
SP  - 601-626
ST  - Molecular and Morphological Phylogenetic Analyses of Themidaceae (Asparagales)
T2  - Kew Bulletin
TI  - Molecular and Morphological Phylogenetic Analyses of Themidaceae (Asparagales)
VL  - 56
ID  - 4269
ER  - 


TY  - JOUR
AU  - Wang, Ren-Xiang
AU  - Shao, Wen
AU  - Liu, Ling
AU  - Liu, Jing
AU  - Deng, Xi-Chao
AU  - Lu, Shu-Gang
DO  - 10.1640/0002-8444-105.1.11 M4 - Citavi
IS  - 1
KW  - Thelypteridaceae
PY  - 2015
SN  - 0002-8444
SP  - 11-19
ST  - A Systematic Study of the Fern Genus Mesopteris Ching (Thelypteridaceae)
T2  - American Fern Journal
TI  - A Systematic Study of the Fern Genus Mesopteris Ching (Thelypteridaceae)
VL  - 105
ID  - 770
ER  - 


TY  - JOUR
AU  - Smith, Alan R.
AU  - Cranfill, Raymond B.
DO  - 10.1640/0002-8444(2002)092[0131:IROTTF]2.0.CO;2 M4 - Citavi
IS  - 2
KW  - Thelypteridaceae
PY  - 2002
SN  - 0002-8444
SP  - 131
ST  - Intrafamilial Relationships of the Thelypteroid Ferns (Thelypteridaceae)
T2  - American Fern Journal
TI  - Intrafamilial Relationships of the Thelypteroid Ferns (Thelypteridaceae)
VL  - 92
ID  - 4171
ER  - 


TY  - JOUR
AU  - Li, Zhong-Yang
AU  - He, Zhao-Rong
AU  - Zhang, Xian-Chun
DO  - 10.1111/jse.12013 M4 - Citavi
IS  - 5
KW  - Thelypteridaceae
PY  - 2013
SN  - 16744918
SP  - 609-638
ST  - A taxonomic revision of Cyclosorus subgenus Cyclosoriopsis (Thelypteridaceae) from China
T2  - Journal of Systematics and Evolution
TI  - A taxonomic revision of Cyclosorus subgenus Cyclosoriopsis (Thelypteridaceae) from China
VL  - 51
ID  - 1288
ER  - 


TY  - JOUR
AB  - Thelypteridaceae is one of the largest families of polypodioid ferns. The generic classification of the family is still controversial because of high levels of convergent or parallel evolution of morphological characters and a lack of molecular phylogenetic studies. In the present study, phylogenetic analyses of three chloroplast regions (rbcL, rps4 and trnL-trnF intergenic spacer region) for 115 taxa, representing 27 recognized segregates in the family, were conducted to explore infrafamilial relationships and gain further understanding of generic boundaries. The phylogenetic reconstructions resolved six distinct clades (Clade I-VI) with strong support. Seven genera: Cyclogramma, Macrothelypteris, Oreopteris, Phegopteris, Pseudophegopteris, Stegnogramma, and Thelypteris are recognized from Clades I, II, IV, and V. In Clade III, Metathelypteris was supported as monophyletic, but the other segregates Amauropelta, Coryphopteris, and Parathelypteris were polyphyletic or paraphyletic, preventing clear recognition of generic boundaries within this clade without additional sampling. Considering great morphological homoplasy within Clade VI, a large genus Cyclosorus is recognized to comprise several small recognized segregates. Within this clade, Pronephrium, and Christella were revealed to be polyphyletic, but several Asian-endemic segregates, such as Glaphyropteridopsis, Mesopteris, and Pseudocyclosorus were strongly supported as monophyletic. Analyses of the evolution of morphological character states on the molecular phylogeny showed extremely high levels of homoplastic evolution for many diagnostic characters.
AD  - State Key Laboratory of Systematic and Evolutionary Botany, Institute of Botany, Chinese Academy of Sciences, Beijing 100093, China. M4 - Citavi
AU  - He, Li-Juan
AU  - Zhang, Xian-Chun
DO  - 10.1016/j.ympev.2012.07.021
IS  - 2
KW  - Thelypteridaceae
LA  - eng
N1  - Journal Article Research Support, Non-U.S. Gov't Journal Article Research Support, Non-U.S. Gov't Journal Article Research Support, Non-U.S. Gov't Journal Article Research Support, Non-U.S. Gov't
PY  - 2012
SN  - 1055-7903
SP  - 757-764
ST  - Exploring generic delimitation within the fern family Thelypteridaceae
T2  - Molecular phylogenetics and evolution
TI  - Exploring generic delimitation within the fern family Thelypteridaceae
VL  - 65
ID  - 1558
ER  - 


TY  - JOUR
AU  - Fernandes, Rozijane Santos
AU  - Yesilyurt, Jovita Cislinski
AU  - Salino, Alexandre
DO  - 10.11646/phytotaxa.184.1.1 M4 - Citavi
IS  - 1
KW  - Thelypteridaceae
PY  - 2014
SN  - 1179-3163
SP  - 1
ST  - New species and combinations in Meniscium (Thelypteridaceae)
T2  - Phytotaxa
TI  - New species and combinations in Meniscium (Thelypteridaceae)
VL  - 184
ID  - 895
ER  - 


TY  - JOUR
AB  - Thelypteridaceae is one of the largest fern families, having about 950 species and a cosmopolitan distribution but with most species occurring in tropical and subtropical regions. Its generic classification remains controversial, with different authors recognizing from one up to 32 genera. Phylogenetic relationships within the family have not been exhaustively studied, but previous studies have confirmed the monophyly of the lineage. Thus far, sampling has been inadequate for establishing a robust hypothesis of infrafamilial relationships within the family. In order to understand phylogenetic relationships within Thelypteridaceae and thus to improve generic reclassification, we expand the molecular sampling, including new samples of Old World taxa and, especially, many additional neotropical representatives. We also explore the monophyly of exclusively or mostly neotropical genera Amauropelta, Goniopteris, Meniscium, and Steiropteris. Our sampling includes 68 taxa and 134 newly generated sequences from two plastid genomic regions (rps4-trnS and trnL-trnF), plus 73 rps4 and 72 trnL-trnF sequences from GenBank. These data resulted in a concatenated matrix of 1980 molecular characters for 149 taxa. The combined data set was analyzed using maximum parsimony and bayesian inference of phylogeny. Our results are consistent with the general topological structure found in previous studies, including two main lineages within the family: phegopteroid and thelypteroid. The thelypteroid lineage comprises two clades; one of these included the segregates Metathelypteris, Coryphopteris, and Amauropelta (including part of Parathelypteris), whereas the other comprises all segregates of Cyclosorus s.l., such as Goniopteris, Meniscium, and Steiropteris (including Thelypteris polypodioides, previously incertae sedis). The three mainly neotropical segregates were found to be monophyletic but nested in a broadly defined Cyclosorus. The fourth mainly neotropical segregate, Amauropelta, was found to include species considered to be part of Parathelypteris. In Old World thelypteroids, which correspond to nearly half the diversity in the family, an increase in sampling is still needed to resolve relationships and circumscription of genera, particularly in the christelloid clade (i.e., Amphineuron, Chingia, Christella, Pneumatopteris, Pronephrium, and Sphaerostephanos). Based on currently available knowledge, we propose the recognition of 16 genera in the family.
AD  - Departamento de Botânica, Universidade Federal de Minas Gerais, Av. Antônio Carlos, 6627 - Belo Horizonte, P.O. Box 486, Minas Gerais 30123-970, Brazil. Electronic address:. UMR 7205, CNRS, MNHN, UPMC, EPHE Institut de Systématique, Evolution, Biodiversité, 43 rue de Buffon, 75005 Paris, France; UMR 7207, CNRS, MNHN, UPMC, 'Centre de recherche sur la Paléobiodiversité et les Paléoenvironnements', 43 rue de Buffon, 75005 Paris, France. Department of Life Sciences, Natural History Museum, Cromwell Road, London SW7 5BD, England, United Kingdom. The University Herbarium, University of California, Berkeley, 1001 Valley Life Sciences Bldg. #2465, Berkeley, CA 94720-2465, USA. Departamento de Botânica, Universidade Federal de Minas Gerais, Av. Antônio Carlos, 6627 - Belo Horizonte, P.O. Box 486, Minas Gerais 30123-970, Brazil. Departamento de Botânica, Universidade Federal de Minas Gerais, Av. Antônio Carlos, 6627 - Belo Horizonte, P.O. Box 486, Minas Gerais 30123-970, Brazil. Departamento de Botânica, Universidade Federal de Minas Gerais, Av. Antônio Carlos, 6627 - Belo Horizonte, P.O. Box 486, Minas Gerais 30123-970, Brazil. Departamento de Botânica, Universidade Federal de Minas Gerais, Av. Antônio Carlos, 6627 - Belo Horizonte, P.O. Box 486, Minas Gerais 30123-970, Brazil. M4 - Citavi
AU  - Almeida, Thaís Elias
AU  - Hennequin, Sabine
AU  - Schneider, Harald
AU  - Smith, Alan R.
AU  - Batista, João Aguiar Nogueira
AU  - Ramalho, Aline Joseph
AU  - Proite, Karina
AU  - Salino, Alexandre
DO  - 10.1016/j.ympev.2015.09.009
IS  - Pt B
KW  - Thelypteridaceae
LA  - eng
N1  - Journal Article Research Support, Non-U.S. Gov't Journal Article Research Support, Non-U.S. Gov't Journal Article Research Support, Non-U.S. Gov't Journal Article Research Support, Non-U.S. Gov't
PY  - 2016
SN  - 1055-7903
SP  - 688-700
ST  - Towards a phylogenetic generic classification of Thelypteridaceae: Additional sampling suggests alterations of neotropical taxa and further study of paleotropical genera
T2  - Molecular phylogenetics and evolution
TI  - Towards a phylogenetic generic classification of Thelypteridaceae: Additional sampling suggests alterations of neotropical taxa and further study of paleotropical genera
VL  - 94
ID  - 151
ER  - 


TY  - JOUR
AU  - R.K.Brummitt
AU  - Banks, Hannah I.
AU  - Johnson, Margaret A. T.
AU  - Docherty, Katherine A.
AU  - Jones, Keith
AU  - Chase, Mark W.
AU  - Rudall, Paula J.
IS  - 4
KW  - Thecophilaeaceae
PY  - 1998
SN  - 0075-5974
SP  - 769-803
ST  - Taxonomy of Cyanastroideae (Tecophilaeaceae): a multidisciplinary approach
T2  - Kew Bulletin
TI  - Taxonomy of Cyanastroideae (Tecophilaeaceae): a multidisciplinary approach
VL  - 53
ID  - 4459
ER  - 


TY  - JOUR
AU  - Manning, John C.
AU  - Goldblatt, Peter
DO  - 10.4102/abc.v42i1.6 Y3 - 15.02.2019 M4 - Citavi
IS  - 1
KW  - Thecophilaeaceae
PY  - 2012
SN  - 0006-8241
SP  - 21-41
ST  - A revision of Tecophilaeaceae subfam. Tecophilaeoideae in Africa
T2  - Bothalia
TI  - A revision of Tecophilaeaceae subfam. Tecophilaeoideae in Africa
VL  - 42
ID  - 1622
ER  - 


TY  - JOUR
AU  - Manning, John C.
AU  - Forest, Félix
AU  - Mannheimer, C. A.
DO  - 10.4102/abc.v35i2.386 Y3 - 15.02.2019 M4 - Citavi
IS  - 2
KW  - Thecophilaeaceae
PY  - 2005
SN  - 0006-8241
SP  - 115-120
ST  - Eremiolirion, a new genus of southern African Tecophilaeaceae, and taxonomic notes on Cyanella alba
T2  - Bothalia
TI  - Eremiolirion, a new genus of southern African Tecophilaeaceae, and taxonomic notes on Cyanella alba
VL  - 35
ID  - 3641
ER  - 


TY  - JOUR
AB  - Schima is an ecologically and economically important woody genus in tea family (Theaceae). Unresolved species delimitations and phylogenetic relationships within Schima limit our understanding of the genus and hinder utilization of the genus for economic purposes. In the present study, we conducted comparative analysis among the complete chloroplast (cp) genomes of 11 Schima species. Our results indicate that Schima cp genomes possess a typical quadripartite structure, with conserved genomic structure and gene order. The size of the Schima cp genome is about 157 kilo base pairs (kb). They consistently encode 114 unique genes, including 80 protein-coding genes, 30 tRNAs, and 4 rRNAs, with 17 duplicated in the inverted repeat (IR). These cp genomes are highly conserved and do not show obvious expansion or contraction of the IR region. The percent variability of the 68 coding and 93 noncoding (>150 bp) fragments is consistently less than 3%. The seven most widely touted DNA barcode regions as well as one promising barcode candidate showed low sequence divergence. Eight mutational hotspots were identified from the 11 cp genomes. These hotspots may potentially be useful as specific DNA barcodes for species identification of Schima. The 58 cpSSR loci reported here are complementary to the microsatellite markers identified from the nuclear genome, and will be leveraged for further population-level studies. Phylogenetic relationships among the 11 Schima species were resolved with strong support based on the cp genome data set, which corresponds well with the species distribution pattern. The data presented here will serve as a foundation to facilitate species identification, DNA barcoding and phylogenetic reconstructions for future exploration of Schima.
AD  - Germplasm Bank of Wild Species in Southwest China, Kunming Institute of Botany, Chinese Academy of Sciences, Kunming, Yunnan, China. Key Laboratory for Plant Diversity and Biogeography of East Asia, Kunming Institute of Botany, Chinese Academy of Sciences, Kunming, Yunnan, China. Department of Biology, University of Nebraska, Kearney, NE, United States of America. Germplasm Bank of Wild Species in Southwest China, Kunming Institute of Botany, Chinese Academy of Sciences, Kunming, Yunnan, China. Key Laboratory for Plant Diversity and Biogeography of East Asia, Kunming Institute of Botany, Chinese Academy of Sciences, Kunming, Yunnan, China. Germplasm Bank of Wild Species in Southwest China, Kunming Institute of Botany, Chinese Academy of Sciences, Kunming, Yunnan, China. M4 - Citavi
AU  - Yu, Xiang-Qin
AU  - Drew, Bryan T.
AU  - Yang, Jun-Bo
AU  - Gao, Lian-Ming
AU  - Li, De-Zhu
DO  - 10.1371/journal.pone.0178026
IS  - 6
KW  - Theaceae
LA  - eng
N1  - Comparative Study Journal Article Comparative Study Journal Article Comparative Study Journal Article Comparative Study Journal Article
PY  - 2017
SP  - e0178026
ST  - Comparative chloroplast genomes of eleven Schima (Theaceae) species: Insights into DNA barcoding and phylogeny
T2  - PloS one
TI  - Comparative chloroplast genomes of eleven Schima (Theaceae) species: Insights into DNA barcoding and phylogeny
UR  - https://www.ncbi.nlm.nih.gov/pmc/articles/PMC5456055
VL  - 12
ID  - 140
ER  - 


TY  - JOUR
AU  - Yang, S. X.
AU  - Yang, J. B.
AU  - Lei, Li-Gong
AU  - Li, D. Z.
AU  - Yoshino, H.
AU  - Ikeda, T.
DO  - 10.1007/s00606-004-0178-3 M4 - Citavi
IS  - 1-4
KW  - Theaceae
N1  - PII: 178 PII: 178 PII: 178 PII: 178
PY  - 2004
SN  - 0378-2697
SP  - 45-55
ST  - Reassessing the relationships between Gordonia and Polyspora (Theaceae) based on the combined analyses of molecular data from the nuclear, plastid and mitochondrial genomes
T2  - Plant Systematics and Evolution
TI  - Reassessing the relationships between Gordonia and Polyspora (Theaceae) based on the combined analyses of molecular data from the nuclear, plastid and mitochondrial genomes
VL  - 248
ID  - 3936
ER  - 


TY  - JOUR
AU  - Wang, Y. H.
AU  - He, H.
AU  - Min, T. L.
AU  - Zhou, L. H.
AU  - Fritsch, Peter W.
DO  - 10.1007/s00606-006-0426-9 M4 - Citavi
IS  - 1
KW  - Theaceae
N1  - PII: 426 PII: 426 PII: 426 PII: 426
PY  - 2006
SN  - 0378-2697
SP  - 39-52
ST  - The phylogenetic position of Apterosperma (Theaceae) based on morphological and karyotype characters
T2  - Plant Systematics and Evolution
TI  - The phylogenetic position of Apterosperma (Theaceae) based on morphological and karyotype characters
VL  - 260
ID  - 3517
ER  - 


TY  - JOUR
AB  - Camellia, comprising more than 200 species, is the type genus of the family Theaceae. Currently, the interspecies relationship of the economically important genus is still a matter of great debate and controversy. In an attempt to help settle this dispute using molecular phylogeny, we analyzed ITS sequences of 112 species of Camellia. The maximum parsimony and Bayesian trees grouped these species into eight major clades and four isolates. The current study supported the monophyly of sections Thea and Furfuracea, a merged section of Theopsis and Eriandra and the formation of section Oleifera by H, -t. Chang (Flora of Reipulicae Popularis Sinicae. Tomus 49 (3), Science Press, China). The study suggested the polyphyletic nature of the sections Camellia, Paracamellia, Pseudocamellia, and Tuberculata and the paraphyletic nature of the section Chrysantha but did not support the sectional status of the three small sections, Archecamellia, Piquetia, and Sterocarpus. We also discuss the results in terms of morphology, geographic distribution and the results from an earlier molecular phylogeny analysis.
AD  - Institute of Plant and Microbial Biology, Academia Sinica, Taipei, Taiwan-115, ROC. M4 - Citavi
AU  - Vijayan, Kunjupillai
AU  - Zhang, Wen-Ju
AU  - Tsou, Chih-Hua
DO  - 10.3732/ajb.0800205
IS  - 7
KW  - Theaceae
LA  - eng
N1  - Journal Article Journal Article Journal Article Journal Article
PY  - 2009
SN  - 0002-9122
SP  - 1348-1360
ST  - Molecular taxonomy of Camellia (Theaceae) inferred from nrITS sequences
T2  - American journal of botany
TI  - Molecular taxonomy of Camellia (Theaceae) inferred from nrITS sequences
VL  - 96
ID  - 2707
ER  - 


TY  - JOUR
AU  - Prince, Linda M.
AU  - Parks, Clifford R.
DO  - 10.2307/3558391 Y3 - 11.02.2019 M4 - Citavi
IS  - 12
KW  - Theaceae
PY  - 2001
SN  - 0002-9122
SP  - 2309-2320
ST  - Phylogenetic relationships of Theaceae inferred from chloroplast DNA sequence data
T2  - American journal of botany
TI  - Phylogenetic relationships of Theaceae inferred from chloroplast DNA sequence data
VL  - 88
ID  - 4272
ER  - 


TY  - JOUR
AU  - Luna, Isolda
AU  - Ochoterena, Helga
DO  - 10.1111/j.1096-0031.2004.00024.x Y3 - 11.02.2019 M4 - Citavi
IS  - 3
KW  - Theaceae
PY  - 2004
SN  - 07483007
SP  - 223-270
ST  - Phylogenetic relationships of the genera of Theaceae based on morphology
T2  - Cladistics
TI  - Phylogenetic relationships of the genera of Theaceae based on morphology
VL  - 20
ID  - 3830
ER  - 


TY  - JOUR
AU  - Li, Rong
AU  - Yang, Jun-Bo
AU  - Yang, Shi-Xiong
AU  - Li, De-Zhu
DO  - 10.1111/j.1756-1051.2011.01175.x M4 - Citavi
IS  - 6
KW  - Theaceae
PY  - 2011
SN  - 0107055X
SP  - 780-787
ST  - Phylogeny and taxonomy of the Pyrenaria complex (Theaceae) based on nuclear ribosomal ITS sequences
T2  - Nordic Journal of Botany
TI  - Phylogeny and taxonomy of the Pyrenaria complex (Theaceae) based on nuclear ribosomal ITS sequences
VL  - 29
ID  - 1950
ER  - 


TY  - JOUR
AD  - College of Life Sciences; Zhejiang University; Hangzhou 310058 China Jiangsu Provincial Key Laboratory for Plant Ex Situ Conservation; Institute of Botany, Jiangsu Province and Chinese Academy of Sciences; Nanjing 210014 China Department of Biology; Hope College; MI 49423 USA Arnold Arboretum; Harvard University; Jamaica Plain MA 02130 USA M4 - Citavi
AU  - Li, Mi-Mi
AU  - Li, Jian-Hua
AU  - Del Tredici, Peter
AU  - Corajod, Jeffrey
AU  - Fu, Cheng-Xin
DO  - 10.1111/jse.12017
IS  - 4
KW  - Theaceae
PY  - 2013
SN  - 16744918
SP  - 396-404
ST  - Phylogenetics and biogeography of Theaceae based on sequences of plastid genes
T2  - Journal of Systematics and Evolution
TI  - Phylogenetics and biogeography of Theaceae based on sequences of plastid genes
VL  - 51
ID  - 1285
ER  - 


TY  - JOUR
AU  - Heo, Kyeong-In
AU  - Lee, Sangtae
AU  - Lee, Chunghee
AU  - Kim, Seung-Chul
DO  - 10.1007/s00606-011-0497-0 M4 - Citavi
IS  - 1-2
KW  - Theaceae
N1  - PII: 497 PII: 497 PII: 497 PII: 497
PY  - 2011
SN  - 0378-2697
SP  - 33-50
ST  - Generic delimitation and infrageneric classification of Stewartia and Hartia (Theaceae; Stewartieae): insight from pollen morphology
T2  - Plant Systematics and Evolution
TI  - Generic delimitation and infrageneric classification of Stewartia and Hartia (Theaceae; Stewartieae): insight from pollen morphology
VL  - 297
ID  - 1915
ER  - 


TY  - JOUR
AU  - Wu, Chi-Chih
AU  - Hsu, Zhi-Fu
AU  - Tsou, Chih-Hua
KW  - Ternstroemiaceae
PY  - 2007
SP  - 97-116
ST  - Phylogeny and Taxonomy of Eurya (Ternstroemiaceae) from Taiwan, as inferred from ITS sequence data
T2  - Botanical Studies
TI  - Phylogeny and Taxonomy of Eurya (Ternstroemiaceae) from Taiwan, as inferred from ITS sequence data
VL  - 48
ID  - 3310
ER  - 


TY  - JOUR
AB  - During the past three decades, molecular taxonomy has made considerable changes in the systematic delimitations of several families in the order Ericales which were formed earlier based on morphology. For instance, the Pentaphylacaceae s.l. has been treated differently by both modern and traditional taxonomists. Modern molecular taxonomists constituted this family by combining the traditionally defined Pentaphylacaceae s.s. (Pentaphylax), Sladeniaceae s.s. (Sladenia), the subfamily Ternstroemioideae with 11 genera of Theaceae s.l. and the genus Ficalhoa. There are also treatments placing the genus Pentaphylax with Ternstroemioideae in Pentaphylacaceae and Ficalhoa with Sladenia in Sladeniaceae. Because most of these genera are poorly studied, investigations on all aspects are important to understand the phylogeny to settle the issues surrounding the treatment of the 14 genera in this family. In the present study, DNA sequences of nrITS and trnL-F genes from species of 11 genera from these 14 genera were generated and analyzed together with sequences from other closely related members of Ericales. The results suggested existence of four distinct lineages viz., Sladenia, Pentaphylax, and tribes Frezierieae (9 genera) and Ternstroemieae (2 genera). Further, it demonstrated that within the biggest lineage, Frezierieae, the Visnea remained sister to the clades Adinandra+Cleyera, Euryodendron+Symplococarpon, Freziera, and finally, Eurya. Based on the evidence, it can be concluded that Sladeniaceae and Pentaphylacaceae are very close to each other and the proposal of merging them into a mega family Pentaphylacaceae s.l. with four tribes, i.e., Sladenieae, Pentaphylaceae, Ternstroemieae, and Freziereae should be considered seriously.
AD  - Institute of Plant and Microbial Biology, Academia Sinica, Taipei, Taiwan, ROC. Research Institute of Tropical Plants, Xishuanbana, Yunnan, People's Republic of China. Institute of Plant and Microbial Biology, Academia Sinica, Taipei, Taiwan, ROC.. Central Silk Board, BTM Layout, Madiwala, Bangalore, 560068, India.. M4 - Citavi
AU  - Tsou, Chih-Hua
AU  - Li, Lu
AU  - Vijayan, Kunjupillai
DO  - 10.1007/s10528-016-9717-1
IS  - 3
KW  - Ternstroemiaceae
LA  - eng
N1  - Journal Article Journal Article Journal Article Journal Article
PY  - 2016
SN  - 0006-2928
SP  - 270-282
ST  - The Intra-familial Relationships of Pentaphylacaceae s.l. as revealed by DNA Sequence Analysis
T2  - Biochemical genetics
TI  - The Intra-familial Relationships of Pentaphylacaceae s.l. as revealed by DNA Sequence Analysis
VL  - 54
ID  - 383
ER  - 


TY  - JOUR
AU  - Su, Yingjuan
AU  - Liao, Wen-bo
AU  - Wang, Ting
AU  - Sun, Yufei
AU  - Wei, Qiang
AU  - Chang, Hungta
DO  - 10.1016/j.bse.2011.07.004 M4 - Citavi
IS  - 4-6
KW  - Ternstroemiaceae
N1  - PII: S0305197811001475 PII: S0305197811001475 PII: S0305197811001475 PII: S0305197811001475
PY  - 2011
SN  - 03051978
SP  - 769-777
ST  - Phylogeny and evolutionary divergence times in Apterosperma and Euryodendron: Evidence of a Tertiary origin in south China
T2  - Biochemical Systematics and Ecology
TI  - Phylogeny and evolutionary divergence times in Apterosperma and Euryodendron: Evidence of a Tertiary origin in south China
VL  - 39
ID  - 2046
ER  - 


TY  - JOUR
AB  - Tectaria (Tectariaceae) is one of the most confusing fern genera in terms of its circumscription and phylogeny. Since its original description, a number of genera had been moved into or related with this genus, while others had been segregated from it. Tectaria is also among the largest fern genera, comprising 150-210 mostly tropical species. Previous molecular studies have been far from comprehensive (sampling no more than 76 accessions from 52 species), limited in geographic scope (mainly restricted to Asia), and based exclusively on plastid markers. In the present study, DNA sequences of eight plastid and one nuclear marker of 360 accessions representing ca. 130 species of Tectaria, ca. 36 species of six non-Tectaria genera in Tectariaceae, 12 species of Davalliaceae, Oleandraceae, and Polypodiaceae, and 13 species of Lomariopsidaceae were used to infer a phylogeny with maximum likelihood, Bayesian inference, and maximum parsimony approaches. Our major results include: (1) the most recently proposed circumscription of Tectaria is strongly supported as monophyletic; (2) the genera Lenda, Microbrochis, Phlebiogonium, and Sagenia, sampled here for the first time, are resolved as part of Tectaria; (3) four superclades representing early splits in Tectaria are identified, with the Old World species being sister to the New World species; (4) 12 well-supported major clades in Tectaria are revealed, differing from one another in molecular, morphological, and geographical features; (5) evolution of 13 morphological characters is inferred in a phylogenetic context and morphological synapomorphies of various clades are identified; and in particular (6) free venation in Tectaria is inferred to be repeatedly derived from anastomosing venation, an evolutionary phenomenon not documented previously in vascular plants in a phylogenetic context based on both plastid and nuclear evidence.
AD  - CAS Key Laboratory of Mountain Ecological Restoration and Bioresource Utilization, Chengdu Institute of Biology, Chinese Academy of Sciences, P.O. Box 416, Chengdu, Sichuan 610041, China; Key Laboratory for Plant Diversity and Biogeography of East Asia, Kunming Institute of Botany, Chinese Academy of Sciences, Kunming, Yunnan 650201, China; Missouri Botanical Garden, P.O. Box 299, St. Louis, MO 63166-0299, USA. CAS Key Laboratory of Mountain Ecological Restoration and Bioresource Utilization, Chengdu Institute of Biology, Chinese Academy of Sciences, P.O. Box 416, Chengdu, Sichuan 610041, China; Missouri Botanical Garden, P.O. Box 299, St. Louis, MO 63166-0299, USA. College of Life Sciences, Chongqing Normal University, Shapingba, Chongqing 400047, China. Department of Botany (MRC 166), National Museum of Natural History, Smithsonian Institution, PO Box 37012, Washington, DC 20013-7012, USA. Correspondent of the Muséum national d'Histoire naturelle (MNHN, Paris, France), Steigestrasse 78, 69412 Eberbach, Germany. CAS Key Laboratory of Mountain Ecological Restoration and Bioresource Utilization, Chengdu Institute of Biology, Chinese Academy of Sciences, P.O. Box 416, Chengdu, Sichuan 610041, China; Department of Biology, Vietnam National Museum of Nature, Vietnam Academy of Science and Technology, 18 Hoang Quoc Viet, Ha Noi, Viet Nam; University of Chinese Academy of Sciences, Beijing 100049, China. Department of Ecology - Evolutionary Biology, Vietnam National University Ho Chi Minh City (VNUHCM) - University of Science, 227 Nguyen Van Cu Street, District 5, Ho Chi Minh City, Viet Nam. Southern Institute of Ecology, Vietnam Academy of Science and Technology, 1 Mạc Đĩnh Chi, District 1, Ho Chi Minh City, Viet Nam. Co-Innovation Center for Sustainable Forestry in Southern China, Nanjing Forestry University and International Cultivar Registration Center for Osmanthus, College of Biology and the Environment, Nanjing Forestry University, Nanjing, Jiangsu 210037, China. College of Life Sciences, Chongqing Normal University, Shapingba, Chongqing 400047, China. CAS Key Laboratory of Mountain Ecological Restoration and Bioresource Utilization, Chengdu Institute of Biology, Chinese Academy of Sciences, P.O. Box 416, Chengdu, Sichuan 610041, China. Electronic address:. CAS Key Laboratory of Mountain Ecological Restoration and Bioresource Utilization, Chengdu Institute of Biology, Chinese Academy of Sciences, P.O. Box 416, Chengdu, Sichuan 610041, China; Missouri Botanical Garden, P.O. Box 299, St. Louis, MO 63166-0299, USA. Electronic address:. M4 - Citavi
AU  - Zhang, Liang
AU  - Zhou, Xin-Mao
AU  - Chen, De-Kui
AU  - Schuettpelz, Eric
AU  - Knapp, Ralf
AU  - Lu, Ngan Thi
AU  - Luong, Thien Tam
AU  - Dang, Minh Tri
AU  - Duan, Yi-Fan
AU  - He, Hai
AU  - Gao, Xin-Fen
AU  - Zhang, Li-Bing
DO  - 10.1016/j.ympev.2017.05.020
KW  - Tectariaceae
LA  - eng
N1  - Journal Article Journal Article Journal Article Journal Article
PY  - 2017
SN  - 1055-7903
SP  - 295-333
ST  - A global phylogeny of the fern genus Tectaria (Tectariaceae: Polypodiales) based on plastid and nuclear markers identifies major evolutionary lineages and suggests repeated evolution of free venation from anastomosing venation
T2  - Molecular phylogenetics and evolution
TI  - A global phylogeny of the fern genus Tectaria (Tectariaceae: Polypodiales) based on plastid and nuclear markers identifies major evolutionary lineages and suggests repeated evolution of free venation from anastomosing venation
VL  - 114
ID  - 141
ER  - 


TY  - JOUR
AU  - Zhang, Liang
AU  - Schuettpelz, Eric
AU  - Rothfels, Carl J.
AU  - Zhou, Xin-Mao
AU  - Gao, Xin-Fen
AU  - Zhang, Li-Bing
DO  - 10.12705/654.3 M4 - Citavi
IS  - 4
KW  - Tectariaceae
PY  - 2016
SN  - 00400262
SP  - 723-738
ST  - Circumscription and phylogeny of the fern family Tectariaceae based on plastid and nuclear markers, with the description of two new genera: Draconopteris and Malaifilix (Tectariaceae)
T2  - Taxon
TI  - Circumscription and phylogeny of the fern family Tectariaceae based on plastid and nuclear markers, with the description of two new genera: Draconopteris and Malaifilix (Tectariaceae)
VL  - 65
ID  - 418
ER  - 


TY  - JOUR
AD  - Shenzhen Laboratory of Southern Subtropical Plant Diversity, Fairylake Botanical Garden; Shenzhen & Chinese Academy of Sciences; Shenzhen 518004 China State Key Laboratory of Systematic and Evolutionary Botany, Institute of Botany; Chinese Academy of Sciences; Beijing 100093 China Department of Life Sciences; Natural History Museum; London SW7 5BD UK M4 - Citavi
AU  - Liu, Hong-Mei
AU  - He, Li-Juan
AU  - Schneider, Harald
DO  - 10.1111/jse.12073
IS  - 2
KW  - Tectariaceae
PY  - 2014
SN  - 16744918
SP  - 161-174
ST  - Towards the natural classification of tectarioid ferns: Confirming the phylogenetic relationships of Pleocnemia and Pteridrys (Eupolypods I)
T2  - Journal of Systematics and Evolution
TI  - Towards the natural classification of tectarioid ferns: Confirming the phylogenetic relationships of Pleocnemia and Pteridrys (Eupolypods I)
VL  - 52
ID  - 964
ER  - 


TY  - JOUR
AB  - In this study we provide a phylogeny for the pantropical fern genus Tectaria, with emphasis on the Old World species, based on sequences of five plastid regions (atpB, ndhF plus ndhF-trnL, rbcL, rps16-matK plus matK, and trnL-F). Maximum parsimony, maximum likelihood, and Bayesian inference are used to analyze 115 individuals, representing ca. 56 species of Tectaria s.l. and 36 species of ten related genera. The results strongly support the monophyly of Tectaria in a broad sense, in which Ctenitopsis, Hemigramma, Heterogonium, Psomiocarpa, Quercifilix, Stenosemia, and Tectaridium should be submerged. Such broadly circumscribed Tectaria is supported by the arising pattern of veinlets and the base chromosome number (x=40). Four primary clades are well resolved within Tectaria, one from the Neotropic (T. trifoliata clade) and three from the Old World (T. subtriphylla clade, Ctenitopsis clade, and T. crenata clade). Tectaria crenata clade is the largest one including six subclades. Of the genera previously recognized as tectarioid ferns, Ctenitis, Lastreopsis, and Pleocnemia, are confirmed to be members in Dryopteridaceae; while Pteridrys and Triplophyllum are supported in Tectariaceae. To infer morphological evolution, 13 commonly used characters are optimized on the resulting phylogenetic trees and in result, are all homoplastic in Tectaria.
AU  - Ding, Hui-Hui
AU  - Chao, Yi-Shan
AU  - Callado, John Rey
AU  - Dong, Shi-Yong
DO  - 10.1016/j.ympev.2014.06.004
KW  - Tectariaceae
LA  - eng
PY  - 2014
SN  - 1055-7903
SP  - 66-78
ST  - Phylogeny and character evolution of the fern genus Tectaria (Tectariaceae) in the Old World inferred from chloroplast DNA sequences
T2  - Molecular phylogenetics and evolution
TI  - Phylogeny and character evolution of the fern genus Tectaria (Tectariaceae) in the Old World inferred from chloroplast DNA sequences
VL  - 80
ID  - 878
ER  - 


TY  - JOUR
AU  - Li, Jian-Hua
AU  - Davis, Charles C.
AU  - Tredici, Peter Del
AU  - Donoghue, Michael J.
IS  - 1
KW  - Taxaceae
PY  - 2001
SN  - 1043-4534
SP  - 267-274
ST  - Phylogeny and biogeography of Taxus (Taxaceae) inferred from sequences of the internal transcribed spacer region of nuclear ribosomal DNA
T2  - Harvard Papers in Botany
TI  - Phylogeny and biogeography of Taxus (Taxaceae) inferred from sequences of the internal transcribed spacer region of nuclear ribosomal DNA
VL  - 6
ID  - 4248
ER  - 


TY  - JOUR
AU  - Ghimire, Balkrishna
AU  - Heo, Kweon
DO  - 10.1007/s00606-013-0874-y M4 - Citavi
IS  - 2
KW  - Taxaceae
N1  - PII: 874 PII: 874 PII: 874 PII: 874
PY  - 2014
SN  - 0378-2697
SP  - 217-223
ST  - Cladistic analysis of Taxaceae s. l
T2  - Plant Systematics and Evolution
TI  - Cladistic analysis of Taxaceae s. l
VL  - 300
ID  - 910
ER  - 


TY  - JOUR
AU  - Collins, Dennis
AU  - Mill, Robert R.
AU  - Möller, Michael
IS  - 2
KW  - Taxaceae
PY  - 2003
SN  - 0002-9122
SP  - 175-182
ST  - Species Separation of Taxus baccata, T. canadensis, and T. cuspidata (Taxaceae) And Origins Of Their Reputed Hybrids Inferred From Rapd And Cpdna Data
T2  - American journal of botany
TI  - Species Separation of Taxus baccata, T. canadensis, and T. cuspidata (Taxaceae) And Origins Of Their Reputed Hybrids Inferred From Rapd And Cpdna Data
VL  - 90
ID  - 3969
ER  - 


TY  - JOUR
AB  - Phylogeny of the Taxaceae genera and the monotypic family Cephalotaxaceae has been extraordinarily controversial. In this paper chloroplast matK genes and nuclear ITS sequences were determined for all six genera of the two families and representatives of other conifer families. Analysis using either the nonsynonymous sites or the deduced amino acid sequences of matK genes strongly indicates that taxad genera and Cephalotaxaceae are monophyletic, with the Taxodiaceae/Cupressaceae clade as their sister group. Cephalotaxus is basal to the taxad genera, among which two clades, Torreya/Amentotaxus and Taxus/Pseudotaxus/Austrotaxus, are resolved. They correspond to Janchen's two tribes, Torreyeae and Taxeae. In Taxeae, Austrotaxus is the first to branch off. Analyses of the nuclear ITS sequence data corroborated the topology of the matK gene tree. These results refute the views that Cephalotaxaceae has no alliance with Taxaceae and that Austrotaxus and Amentotaxus should be excluded from the Taxaceae. We estimated the divergence time between the Taxodiaceae/Cupressaceae and the Cephalotaxaceae/Taxaceae clades to be 192-230 Myr ago and the divergence time between taxads and Cephalotaxus to be 149-179 Myr ago. Soon after the latter divergence event, within 6-8 Myr, the two taxad tribes originated. In conclusion, our data do not support Florin's claim that taxads could be traced to Devonian psilophytes (359-395 Myr ago).
AD  - Institute of Botany, Academia Sinica, Taipei, 11529, Taiwan. M4 - Citavi
AU  - Cheng, Y.
AU  - Nicolson, R. G.
AU  - Tripp, K.
AU  - Chaw, Shu-Miaw
DO  - 10.1006/mpev.1999.0710
IS  - 3
KW  - Taxaceae
LA  - eng
N1  - Journal Article Research Support, Non-U.S. Gov't Journal Article Research Support, Non-U.S. Gov't Journal Article Research Support, Non-U.S. Gov't Journal Article Research Support, Non-U.S. Gov't
PY  - 2000
SN  - 1055-7903
SP  - 353-365
ST  - Phylogeny of Taxaceae and Cephalotaxaceae genera inferred from chloroplast matK gene and nuclear rDNA ITS region
T2  - Molecular phylogenetics and evolution
TI  - Phylogeny of Taxaceae and Cephalotaxaceae genera inferred from chloroplast matK gene and nuclear rDNA ITS region
VL  - 14
ID  - 4326
ER  - 


TY  - JOUR
AU  - Zhang, Mingli
AU  - Hao, Xiao-Li
AU  - Sanderson, Stewart C.
AU  - Vyacheslav, Byalt V.
AU  - Sukhorukov, Alexander P.
AU  - Zhang, X. I. A.
DO  - 10.11646/phytotaxa.167.1.5 M4 - Citavi
IS  - 1
KW  - Tamaricaceae
PY  - 2014
SN  - 1179-3163
SP  - 89
ST  - Spatiotemporal evolution of Reaumuria (Tamaricaceae) in Central Asia: insights from molecular biogeography
T2  - Phytotaxa
TI  - Spatiotemporal evolution of Reaumuria (Tamaricaceae) in Central Asia: insights from molecular biogeography
VL  - 167
ID  - 1144
ER  - 


TY  - JOUR
AU  - Wang, Yong
AU  - Liu, Yifei
AU  - Liu, Songbai
AU  - Huang, Hongwen
KW  - Tamaricaceae
PY  - 2009
SP  - 343-352
ST  - Molecular phylogeny of Myricaria (Tamaricaceae): implications for taxonomy and conservation in China
T2  - Botanical Studies
TI  - Molecular phylogeny of Myricaria (Tamaricaceae): implications for taxonomy and conservation in China
VL  - 50
ID  - 2710
ER  - 


TY  - JOUR
AU  - Sun, Likun
AU  - Yang, Ruiqi
AU  - Zhang, Baogui
AU  - Zhang, Gaosen
AU  - Wu, Xiukun
AU  - Zhang, Wei
AU  - Zhang, Binlin
AU  - Chen, Tuo
AU  - Liu, Guangxiu
DO  - 10.1016/j.bse.2016.10.003 M4 - Citavi
KW  - Tamaricaceae
N1  - PII: S030519781630223X PII: S030519781630223X PII: S030519781630223X PII: S030519781630223X
PY  - 2016
SN  - 03051978
SP  - 213-221
ST  - Phylogenetic relationships among species of Tamarix (Tamaricaceae) in China
T2  - Biochemical Systematics and Ecology
TI  - Phylogenetic relationships among species of Tamarix (Tamaricaceae) in China
VL  - 69
ID  - 370
ER  - 


TY  - JOUR
AU  - Gaskin, John F.
AU  - Schaal, Barbara A.
IS  - 1
KW  - Tamaricaceae
PY  - 2003
SN  - 03636445
SP  - 86-95
ST  - Molecular Phylogenetic Investigation of U.S. Invasive Tamarix
T2  - Systematic Botany
TI  - Molecular Phylogenetic Investigation of U.S. Invasive Tamarix
VL  - 28
ID  - 3983
ER  - 


TY  - JOUR
AU  - Gaskin, John F.
AU  - Ghahremaninejad, Farrokh
AU  - Zhang, Dao-Yuan
AU  - Londo, Jason P.
KW  - Tamaricaceae
PY  - 2004
SP  - 401-409
ST  - A systematic overviewn of Frankeniaceae and Tamaricaceae from nuclear rDNA and plastid sequence data
T2  - Annals of the Missouri Botanical Garden
TI  - A systematic overviewn of Frankeniaceae and Tamaricaceae from nuclear rDNA and plastid sequence data
VL  - 91
ID  - 3784
ER  - 


TY  - JOUR
AU  - Chen, Youhua
DO  - 10.4172/2157-7625.1000132 M4 - Citavi
IS  - 132
KW  - Tamaricaceae
PY  - 2013
ST  - A Revisited Study On Phylogeography And Phylogenetic Diversity Of Myricaria (Tamaricaceae)
T2  - Journal of Ecosystem & Ecography
TI  - A Revisited Study On Phylogeography And Phylogenetic Diversity Of Myricaria (Tamaricaceae)
VL  - 3
ID  - 1193
ER  - 


TY  - JOUR
AU  - Arianmanesh, Reza
AU  - Mehregan, Iraj
AU  - Nejadsatari, Taher
AU  - Assadi, Mostafa
IS  - 6
KW  - Tamaricaceae
PY  - 2015
SP  - 44-50
ST  - Molecular phylogeny of Tamarix (Tamaricaceae) species from Iran based on ITS sequence data
T2  - Euro. J. Exp. Bio.
TI  - Molecular phylogeny of Tamarix (Tamaricaceae) species from Iran based on ITS sequence data
VL  - 5
ID  - 433
ER  - 


TY  - JOUR
AU  - Nyffeler, Reto
AU  - Eggli, Urs
IS  - 1
KW  - Talinaceae
PY  - 2010
SN  - 00400262
SP  - 227-240
ST  - Disintegrating Portulacaceae: A new familial classification of the suborder Portulacineae (Caryophyllales) based on molecular and morphological data
T2  - Taxon
TI  - Disintegrating Portulacaceae: A new familial classification of the suborder Portulacineae (Caryophyllales) based on molecular and morphological data
VL  - 59
ID  - 2300
ER  - 


TY  - JOUR
AB  - Symplocos comprises ∼300 species of woody flowering plants with a disjunct distribution between the warm-temperate to tropical regions of eastern Asia and the Americas. Phylogenetic analyses of 111 species of Symplocos based on the nuclear ribosomal internal transcribed spacer (ITS) region and the chloroplast genes rpl16, matK, and trnL-trnF yielded topologies in which only one of the four traditionally recognized subgenera (Epigenia; Neotropics) is monophyletic. Section Cordyloblaste (subgenus Symplocos; eastern Asia) is monophyletic and sister to a group comprising all other samples of Symplocos. Section Palura (subgenus Hopea; eastern Asia) is sister to a group comprising all other samples of Symplocos except those of section Cordyloblaste. Symplocos wikstroemiifolia (eastern Asia) and S. tinctoria (southeastern United States), both of subgenus Hopea, form a clade that groups with S. longipes (tropical North America) and the species of subgenus Epigenia. The remaining samples of subgenus Hopea (eastern Asia) form a clade. Section Neosymplocos (subgenus Microsymplocos; Neotropics) is well nested within a clade otherwise comprising the samples of section Symplocastrum (subgenus Symplocos; Neotropics). Section Urbaniocharis (subgenus Microsymplocos; Antilles) groups as sister to the clade comprising Symplocastrum and Neosymplocos. The data support the independent evolution of deciduousness among section Palura and S. tinctoria. The early initial divergence of sections Cordyloblaste and Palura from the main group warrants their recognition at taxonomic levels higher than those at which they are currently placed. An inferred eastern Asian origin for Symplocos with subsequent dispersal to the Americas is consistent with patterns from other phylogenetic studies of eastern Asian-American disjunct plant groups but contrary to a North American origin inferred from the earliest fossil occurrences of the genus.
AD  - State Key Laboratory, School of Life Sciences, Zhongshan University, Guangzhou 510275, People's Republic of China; M4 - Citavi
AU  - Wang, Yuguo
AU  - Fritsch, Peter W.
AU  - Shi, Suhua
AU  - Almeda, Frank
AU  - Cruz, Boni C.
AU  - Kelly, Lawrence M.
DO  - 10.3732/ajb.91.11.1901
IS  - 11
KW  - Symplocaceae
LA  - eng
N1  - Journal Article Journal Article Journal Article Journal Article
PY  - 2004
SN  - 0002-9122
SP  - 1901-1914
ST  - Phylogeny and infrageneric classification of Symplocos (Symplocaceae) inferred from DNA sequence data
T2  - American journal of botany
TI  - Phylogeny and infrageneric classification of Symplocos (Symplocaceae) inferred from DNA sequence data
VL  - 91
ID  - 3925
ER  - 


TY  - JOUR
AB  - Thirty species and one variety of Symplocos (Symplocaceae), including all taxa distributed in Japan, were phylogenetically analyzed with DNA sequence data. The evolution of morphological characters is discussed on the basis of the phylogenetic relationships obtained. All species were Asian, except one, S. austromexicana, from Mexico. The nuclear ribosomal internal transcribed spacer (ITS) region and two intergenic spacer regions, between trnL and trnF, and between trnH and psbA of chloroplast DNA were used. The topologies of trees obtained from ITS and the chloroplast intergenic spacers are largely congruent. S. sonoharae, the representative of the subgenus Symplocos in Asia, was sister to all other species. The position of the American species, S. austromexicana, which also belongs to the subgenus Symplocos, was not well resolved. The phylogenetic tree based on combined sequence data largely supports the monophyletic origin of the infrageneric sections proposed earlier. However, the phylogenetic relationship between them is not well resolved, probably due to rapid diversification. The section Palura, a deciduous group, is well defined in the DNA analysis, suggesting its independent status in the genus Symplocos. In spite of their morphological divergence, the three endemic species of the Bonin Islands are monophyletic. The occurrence of curved seeds seems to be homoplastic, scattered over the phylogenetic tree without showing a particular infrageneric relationship.
AD  - College of Integrated Arts and Sciences, Osaka Prefecture University, Osaka 599-8531, Japan. M4 - Citavi
AU  - Soejima, Akiko
AU  - Nagamasu, Hidetoshi
DO  - 10.1007/s10265-004-0151-9
IS  - 3
KW  - Symplocaceae
LA  - eng
N1  - Comparative Study Journal Article Research Support, Non-U.S. Gov't Comparative Study Journal Article Research Support, Non-U.S. Gov't Comparative Study Journal Article Research Support, Non-U.S. Gov't Comparative Study Journal Article Research Support, Non-U.S. Gov't
PY  - 2004
SP  - 199-207
ST  - Phylogenetic analysis of Asian Symplocos (Symplocaceae) based on nuclear and chloroplast DNA sequences
T2  - Journal of Plant Research
TI  - Phylogenetic analysis of Asian Symplocos (Symplocaceae) based on nuclear and chloroplast DNA sequences
VL  - 117
ID  - 3902
ER  - 


TY  - JOUR
AD  - Department of Botany; California Academy of Sciences; 55 Music Concourse Drive , San Francisco, California 94118 U.S.A. The New York Botanical Garden; 200th St. & Southern Blvd. Bronx, New York 10458 U.S.A. Ministry of Education Key Laboratory for Biodiversity Science and Ecological Engineering; Institute of Biodiversity Science; Fudan University; Shanghai 200433 China Department of Biology; San Francisco State University,; 1600 Holloway Ave. San Francisco, California 94132 U.S.A. M4 - Citavi
AU  - Fritsch, Peter W.
AU  - Kelly, Lawrence M.
AU  - Wang, Yuguo
AU  - Almeda, Frank
AU  - Kriebel, Ricardo
DO  - 10.1002/tax.573013 Y3 - 11.02.2019
IS  - 3
KW  - Symplocaceae
PY  - 2008
SN  - 00400262
SP  - 823-852
ST  - Revised infrafamilial classification of Symplocaceae based on phylogenetic data from DNA sequences and morphology
T2  - Taxon
TI  - Revised infrafamilial classification of Symplocaceae based on phylogenetic data from DNA sequences and morphology
VL  - 57
ID  - 2805
ER  - 


TY  - JOUR
AU  - Fritsch, Peter W.
AU  - Cruz, Boni C.
AU  - Almeda, Frank
AU  - Wang, Yuguo
AU  - Shi, Su-hua
IS  - 1
KW  - Symplocaceae
PY  - 2006
SN  - 03636445
SP  - 181-192
ST  - Phylogeny of Symplocos Based on DNA Sequences of the Chloroplast trnC-trnD Intergenic Region
T2  - Systematic Botany
TI  - Phylogeny of Symplocos Based on DNA Sequences of the Chloroplast trnC-trnD Intergenic Region
VL  - 31
ID  - 3374
ER  - 


TY  - JOUR
AU  - Fritsch, Peter W.
AU  - Al, Frank
DO  - 10.11646/phytotaxa.194.1.1 M4 - Citavi
IS  - 1
KW  - Symplocaceae
PY  - 2015
SN  - 1179-3163
SP  - 1
ST  - A taxonomic revision of Antillean Symplocos (Symplocaceae)
T2  - Phytotaxa
TI  - A taxonomic revision of Antillean Symplocos (Symplocaceae)
VL  - 194
ID  - 518
ER  - 


TY  - JOUR
AB  - BACKGROUND AND AIMS The genus Sinojackia consists of eight species, all endemic to China. All species of Sinojackia are endangered or threatened owing to poor recruitment within populations. Information on molecular phylogenetics is critical for developing successful conservation strategies for this genus. METHODS Combined DNA sequence data from the nuclear ribosomal internal transcribed spacer regions and plastid psbA-trnH intergenic spacer and microsatellite data were used to infer a phylogeny of the genus. KEY RESULTS Parsimony analysis of the combined sequence data and multivariate analysis based on fruit characters indicated that Sinojackia dolichocarpa is monophyletic and genetically well separated from the other Sinojackia species, thus supporting its rank at the generic level as Changiostyrax. Phylogenetic relationships within Sinojackia sensu stricto are unresolved from the combined sequence data. A UPGMA dendrogram based on seven microsatellite loci of 96 individual plants yielded a first-diverging cluster of all individuals of S. microcarpa. The remaining species form another cluster without any definitive patterns corresponding to current species circumscriptions, suggesting either extensive hybridization or incipient speciation. CONCLUSIONS The results suggest that there are too many species recognized within Sinojackia sensu stricto, but this must be further assessed with comprehensive morphological and taxonomic revisionary work. The implications of the phylogenetic data for conservation are discussed.
AD  - Wuhan Botanical Garden, The Chinese Academy of Sciences, Moshan, Wuhan 430074, China. M4 - Citavi
AU  - Yao, Xiaohong
AU  - Ye, Qigang
AU  - Fritsch, Peter W.
AU  - Cruz, Boni C.
AU  - Huang, Hongwen
DO  - 10.1093/aob/mcm332
IS  - 5
KW  - Styracaceae
LA  - eng
N1  - Journal Article Research Support, Non-U.S. Gov't Journal Article Research Support, Non-U.S. Gov't Journal Article Research Support, Non-U.S. Gov't Journal Article Research Support, Non-U.S. Gov't
PY  - 2008
SN  - 0305-7364
SP  - 651-659
ST  - Phylogeny of Sinojackia (Styracaceae) based on DNA sequence and microsatellite data: implications for taxonomy and conservation
T2  - Annals of Botany
TI  - Phylogeny of Sinojackia (Styracaceae) based on DNA sequence and microsatellite data: implications for taxonomy and conservation
UR  - https://www.ncbi.nlm.nih.gov/pmc/articles/PMC2710174
VL  - 101
ID  - 3028
ER  - 


TY  - JOUR
AU  - Fritsch, Peter W.
AU  - Morton, Cynthia M.
AU  - Chen, Tao
AU  - Meldrum, Candice
DO  - 10.1086/323418 Y3 - 11.02.2019 M4 - Citavi
IS  - S6
KW  - Styracaceae
PY  - 2001
SN  - 1058-5893
SP  - S95-S116
ST  - Phylogeny and Biogeography of the Styracaceae
T2  - International Journal of Plant Sciences
TI  - Phylogeny and Biogeography of the Styracaceae
VL  - 162
ID  - 4224
ER  - 


TY  - JOUR
AU  - Fritsch, Peter W.
IS  - 1
KW  - Styracaceae
PY  - 2004
SN  - 1055-3177
SP  - 43-57
ST  - New Species and Taxonomic Changes in Styrax (Styracaceae) from South America
T2  - Novon: A Journal for Botanical Nomenclature
TI  - New Species and Taxonomic Changes in Styrax (Styracaceae) from South America
VL  - 14
ID  - 3780
ER  - 


TY  - JOUR
AB  - Phylogenetic relationships within the flowering plant genus Styrax were investigated with DNA sequence data from the internal transcribed spacer (ITS) region of nuclear ribosomal DNA (nrDNA) and with chloroplast DNA restriction site data from the genes trnK, rpoC1, and rpoC2. The data sets from each genome were analyzed separately and in combination with parsimony methods. The results strongly support the monophyly of each of the four series of the genus but provide little phylogenetic resolution among them. Reticulate evolution may at least partly explain discordance between the molecular phylogenetic estimates and a prior morphological estimate within series Cyrta. The historical biogeography of the genus was inferred with unweighted parsimony character optimization of trees recovered from a combined ITS and morphological data set, after a series of combinability tests for data set congruence was conducted. The results are consistent with the fossil record in supporting a Eurasian origin of Styrax. The nested phylogenetic position of the South American members of the genus within those from southern North America and Eurasia suggests that the boreotropics hypothesis best explains the amphi-Pacific tropical disjunct distribution occurring within section Valvatae. The pattern of relationship recovered among the species of section Styrax ((western North America + western Eurasia) (eastern North America + eastern Eurasia)) is rare among north-temperate Tertiary forest relicts. The monophyly of the group of species from western North America and western Eurasia provides qualified support for the Madrean-Tethyan hypothesis, which posits a Tertiary floristic connection among the semiarid regions in which these taxa occur. A single vicariance event between eastern Asia and eastern North America accounts for the pattern of relationship among intercontinental disjuncts in series Cyrta.
AD  - Department of Botany and Osher Foundation Laboratory for Molecular Systematics, California Academy of Sciences, Golden Gate Park, San Francisco, California 94118, USA. M4 - Citavi
AU  - Fritsch, Peter W.
DO  - 10.1006/mpev.2001.0933
IS  - 3
KW  - Styracaceae
LA  - eng
N1  - Journal Article Research Support, Non-U.S. Gov't Research Support, U.S. Gov't, Non-P.H.S. Journal Article Research Support, Non-U.S. Gov't Research Support, U.S. Gov't, Non-P.H.S. Journal Article Research Support, Non-U.S. Gov't Research Support, U.S. Gov't, Non-P.H.S. Journal Article Research Support, Non-U.S. Gov't Research Support, U.S. Gov't, Non-P.H.S.
PY  - 2001
SN  - 1055-7903
SP  - 387-408
ST  - Phylogeny and biogeography of the flowering plant genus Styrax (Styracaceae) based on chloroplast DNA restriction sites and DNA sequences of the internal transcribed spacer region
T2  - Molecular phylogenetics and evolution
TI  - Phylogeny and biogeography of the flowering plant genus Styrax (Styracaceae) based on chloroplast DNA restriction sites and DNA sequences of the internal transcribed spacer region
VL  - 19
ID  - 4223
ER  - 


TY  - JOUR
AU  - Wege, Juliet A.
DO  - 10.1071/SB12001 M4 - Citavi
IS  - 2
KW  - Stylidiaceae
N1  - PII: SB12001 PII: SB12001 PII: SB12001 PII: SB12001
PY  - 2012
SN  - 1030-1887
SP  - 138
ST  - Navigating the floral Milky Way: the taxonomy of the microgeophytic triggerplants (Stylidium petiolare and allies: Stylidiaceae)
T2  - Australian Systematic Botany
TI  - Navigating the floral Milky Way: the taxonomy of the microgeophytic triggerplants (Stylidium petiolare and allies: Stylidiaceae)
VL  - 25
ID  - 1759
ER  - 


TY  - JOUR
AU  - Laurent, Nadina
AU  - Bremer, Birgitta
AU  - Bremer, Kare
DO  - 10.2307/2419506 M4 - Citavi
IS  - 3
KW  - Stylidiaceae
PY  - 1998
SN  - 03636445
SP  - 289
ST  - Phylogeny and Generic Interrelationships of the Stylidiaceae (Asterales), with a Possible Extreme Case of Floral Paedomorphosis
T2  - Systematic Botany
TI  - Phylogeny and Generic Interrelationships of the Stylidiaceae (Asterales), with a Possible Extreme Case of Floral Paedomorphosis
VL  - 23
ID  - 4455
ER  - 


TY  - JOUR
AU  - Glenny, David S.
DO  - 10.1080/00288250909509811 M4 - Citavi
IS  - 3
KW  - Stylidiaceae
PY  - 2009
SN  - 0028-825X
SP  - 285-315
ST  - A revision of the genus Forstern (Stylidiaceae) in New Zealand
T2  - New Zealand Journal of Botany
TI  - A revision of the genus Forstern (Stylidiaceae) in New Zealand
VL  - 47
ID  - 2525
ER  - 


TY  - JOUR
AU  - Cron, Glynis V.
AU  - Pirone, Cary
AU  - Bartlett, Madelaine
AU  - Kress, W. John
AU  - Specht, Chelsea D.
DO  - 10.1600/036364412X648562 M4 - Citavi
IS  - 3
KW  - Strelitziaceae
PY  - 2012
SN  - 03636445
SP  - 606-619
ST  - Phylogenetic Relationships and Evolution in the Strelitziaceae (Zingiberales)
T2  - Systematic Botany
TI  - Phylogenetic Relationships and Evolution in the Strelitziaceae (Zingiberales)
VL  - 37
ID  - 1515
ER  - 


TY  - JOUR
AU  - Wilkie, Peter
AU  - Clark, Alexandra
AU  - Pennington, R. Toby
AU  - Cheek, Martin R.
AU  - Bayer, Clemens
AU  - Wilcock, Chris C.
IS  - 1
KW  - Sterculiaceae
PY  - 2006
SN  - 03636445
SP  - 160-170
ST  - Phylogenetic Relationships within the Subfamily Sterculioideae (Malvaceae/Sterculiaceae-Sterculieae) Using the Chloroplast Gene ndhF
T2  - Systematic Botany
TI  - Phylogenetic Relationships within the Subfamily Sterculioideae (Malvaceae/Sterculiaceae-Sterculieae) Using the Chloroplast Gene ndhF
VL  - 31
ID  - 3527
ER  - 


TY  - JOUR
AU  - Dorr, L. J.
AU  - Cheek, Martin R.
KW  - Sterculiaceae
PY  - 2011
SN  - 0075-5974
SP  - 629-631
ST  - The identity and typification of Pimia Seem. (Sterculiaceae)
T2  - Kew Bulletin
TI  - The identity and typification of Pimia Seem. (Sterculiaceae)
VL  - 66
ID  - 1869
ER  - 


TY  - JOUR
AU  - Utteridge, Timothy M. A.
KW  - Stemonuraceae
PY  - 2011
SN  - 0075-5974
SP  - 49-81
ST  - A revision of the genus Medusanthera (Stemonuraceae, Icacinaceae s.l.)
T2  - Kew Bulletin
TI  - A revision of the genus Medusanthera (Stemonuraceae, Icacinaceae s.l.)
VL  - 66
ID  - 2064
ER  - 


TY  - JOUR
AU  - Schori, M.
AU  - Utteridge, T. M. A.
DO  - 10.3767/000651910X527699 M4 - Citavi
IS  - 2
KW  - Stemonuraceae
PY  - 2010
SP  - 189-195
ST  - Three new species and a new name in Southeast Asian Gomphandra (Stemonuraceae / Icacinaceae s.l.)
T2  - Blumea
TI  - Three new species and a new name in Southeast Asian Gomphandra (Stemonuraceae / Icacinaceae s.l.)
VL  - 55
ID  - 2345
ER  - 


TY  - THES
AU  - Schori, Melanie
CY  - Faculty of the College of Arts and Sciences of Ohio University
DA  - June 2010
KW  - Stemonuraceae
M3  - Dissertation
PB  - Ohio University
ST  - A Systematic Revision of Gomphandra (Stemonuraceae)
TI  - A Systematic Revision of Gomphandra (Stemonuraceae)
ID  - 2099
ER  - 


TY  - JOUR
AU  - Munzinger, Jérôme
AU  - McPherson, Gordon
AU  - Lowry, Porter Prescott, II
KW  - Stemonuraceae
PY  - 2008
SP  - 775-783
ST  - A second species in the endemic New Caledonian genus Gastrolepis (Stemonuraceae) and its implications for the conservation status of high-altitude maquis vegetation: coherent application of the IUCN Red List criteria is urgently needed in New Caledonia
T2  - Botanical Journal of the Linnean Society
TI  - A second species in the endemic New Caledonian genus Gastrolepis (Stemonuraceae) and its implications for the conservation status of high-altitude maquis vegetation: coherent application of the IUCN Red List criteria is urgently needed in New Caledonia
VL  - 157
ID  - 2908
ER  - 


TY  - JOUR
AB  - The genus Croomia (Stemonaceae) comprises three herbaceous perennial species that are distributed in temperate-deciduous forests in Southeastern North America (C.pauciflora) and East Asia (C. japonica, C. heterosepala). The two Asian species have abutting ranges in South Japan, but C. japonica also occurs disjunctively on the adjacent Asiatic mainland in East China. In our phylogenetic analysis of Croomia, based on chloroplast (cp) DNA sequence variation of the trnL-F region, and rooted with Stemona spp., the two Asian species are identified as sister that likely diverged in the Mid-to-Late Pleistocene (0.84-0.13 mya), whereas the divergence of C. pauciflora dates back to the Late Plio-/Pleistocene (<2.6 mya). Phylogeographical analysis of the two East Asian species detected seven cpDNA (trnL-F) haplotypes across 16 populations surveyed, and all of those were fixed for a particular cpDNA haplotype (H(E)=0.0, G(ST)=1). A survey of inter-simple sequence repeats (ISSRs) markers also detected remarkably low levels of within-population diversity (C. japonica: H(E)=0.085; C. heterosepala: H(E)=0.125), and high levels of inter-population differentiation (C. japonica: Phi(ST)=0.736; C. heterosepala: Phi(ST)=0.550), at least partly due to pronounced regional genetic substructure within both species. Non-overlapping distributions of cpDNA haplotypes and strong genetic (cpDNA/ISSR) differentiation among populations and/or regions accord with findings of a nested clade analysis, which inferred allopatric fragmentation as the major process influencing the spatial haplotype distribution of the two species. Based on mismatch distribution analysis and neutrality tests, we do not find evidence of population expansion in both species. Overall, we conclude that components of temperate-deciduous forest types in South Japan and East China are particularly sensitive to range fragmentation, isolation, and enhanced (incipient) species formation through climate-induced expansions of other forest types over glacial and interglacial periods of the (Late) Quaternary.
AD  - Laboratory of Systematic and Evolutionary Botany, Department of Biology, College of Life Science, Zhejiang University, Hangzhou 310058, PR China. M4 - Citavi
AU  - Li, En-Xiang
AU  - Yi, Sun
AU  - Qiu, Ying-Xiong
AU  - Guo, Jiang-Tao
AU  - Comes, Hans Peter
AU  - Fu, Cheng-Xin
DO  - 10.1016/j.ympev.2008.09.012
IS  - 3
KW  - Stemonaceae
LA  - eng
N1  - Journal Article Research Support, Non-U.S. Gov't Journal Article Research Support, Non-U.S. Gov't Journal Article Research Support, Non-U.S. Gov't Journal Article Research Support, Non-U.S. Gov't
PY  - 2008
SN  - 1055-7903
SP  - 702-714
ST  - Phylogeography of two East Asian species in Croomia (Stemonaceae) inferred from chloroplast DNA and Issr fingerprinting variation
T2  - Molecular phylogenetics and evolution
TI  - Phylogeography of two East Asian species in Croomia (Stemonaceae) inferred from chloroplast DNA and Issr fingerprinting variation
VL  - 49
ID  - 2875
ER  - 


TY  - JOUR
AU  - Fan, Lan-Lan
AU  - Zhu, Shu
AU  - Chen, Hu-Biao
AU  - Yang, Dong-Hui
AU  - Cai, Shao-Qing
AU  - Komatsu, Katsuko
DO  - 10.1248/bpb.32.1439 Y3 - 22.02.2019 M4 - Citavi
IS  - 8
KW  - Stemonaceae
PY  - 2009
SN  - 0918-6158
SP  - 1439-1446
ST  - Molecular Analysis of Stemona Plants in China based on Sequences of four Chloroplast DNA Regions
T2  - Biological & Pharmaceutical Bulletin
TI  - Molecular Analysis of Stemona Plants in China based on Sequences of four Chloroplast DNA Regions
VL  - 32
ID  - 2503
ER  - 


TY  - JOUR
AD  - Department of Botany; National Museum of Natural History; MRC-166, Smithsonian Institution Washington, DC 20013-7012 USA Institute of Agricultural Economy and Information; Hubei Academy of Agricultural Sciences; Wuhan 430064 China National Field Station of Freshwater Ecosystem of Liangzi Lake; College of Life Sciences, Wuhan University; Wuhan 430072 China College of Life Sciences; Shaanxi Normal University; Xi'an 710119 China Agriculture School; Kunming University; Kunming 650214 China M4 - Citavi
AU  - Harris, A. J.
AU  - Chen, Ping-Ting
AU  - Xu, Xin-Wei
AU  - Zhang, Jian-Qiang
AU  - Yang, Xue
AU  - Wen, Jun
DO  - 10.1111/jse.12236
IS  - 2
KW  - Staphyleaceae
PY  - 2017
SN  - 16744918
SP  - 124-141
ST  - A molecular phylogeny of Staphyleaceae: Implications for generic delimitation and classical biogeographic disjunctions in the family
T2  - Journal of Systematics and Evolution
TI  - A molecular phylogeny of Staphyleaceae: Implications for generic delimitation and classical biogeographic disjunctions in the family
VL  - 55
ID  - 79
ER  - 


TY  - JOUR
AD  - State Key Laboratory of Systematic and Evolutionary Botany; Institute of Botany; Chinese Academy of Sciences; 20 Nanxincun Xiangshan Beijing 100093 China Graduate School of the Chinese Academy of Sciences; 19 Yuquanlu Beijing 100049 China College of Life Sciences; Nanjing University; 22 Hankoulu Nanjing Jiangsu Province 210093 China Department of Botany; National Museum of Natural History; Smithsonian Institution; Washington MRC166 DC 20013-7012 U.S.A. M4 - Citavi
AU  - Zhu, Yu-Ping
AU  - Wen, Jun
AU  - Zhang, Zhi-Yun
AU  - Chen, Zhi‐Duan
DO  - 10.2307/25065687 Y3 - 11.02.2019
IS  - 4
KW  - Stachyuraceae
PY  - 2006
SN  - 00400262
SP  - 931-940
ST  - Evolutionary relationships and diversification of Stachyuraceae based on sequences of four chloroplast markers and the nuclear ribosomal ITS region
T2  - Taxon
TI  - Evolutionary relationships and diversification of Stachyuraceae based on sequences of four chloroplast markers and the nuclear ribosomal ITS region
VL  - 55
ID  - 3540
ER  - 


TY  - JOUR
AB  - The complex evolutionary history of the subtribe Physalinae is reflected in the poor resolution of the relationships of Physalis and the physaloid genera. We hypothesize that this low resolution is caused by recent evolutionary history in a complex geographic setting. The aims of this study were twofold: (1) To determine the phylogenetic relationships of the current genera recognized in Physalinae in order to identify monophyletic groups and resolve the physaloid grade; and (2) to determine the probable causes of the recent divergence in Physalinae. We conducted phylogenetic analyses with maximum likelihood (ML) and Bayesian inference with 50 Physalinae species and 19 others as outgroups, using morphological and molecular data from five plastid and two nuclear regions. A relaxed molecular clock was obtained from the ML topology and ancestral area reconstruction was conducted using the DEC model. The genera Chamaesaracha, Leucophysalis, and Physalis subgenus Rydbergis were recovered as monophyletic. Three clades, Alkekengi-Calliphysalis, Schraderanthus-Tzeltalia, and Witheringia-Brachistus, also received good support. However, even with morphological data and that of the DNA of seven regions, the tree was not completely resolved and many clades remained unsupported. Physalinae diverged at the end of the Miocene (∼9.22Mya) with one trend indicating that the greatest diversification within the subtribe occurred during the last 5My. The Neotropical region presented the highest probability (45%) of being the ancestral area of Physalinae followed by the Mexican Transition Zone (35%). During the Pliocene and Pleistocene, the geographical areas where species were found experienced significant geological and climatic changes, giving rise to rapid and relatively recent diversification events in Physalinae. Thus, recent origin, high diversification, and morphological complexity have contributed, at least with the currently available methods, to the inability to completely disentangle the phylogenetic relationships of Physalinae.
AD  - Instituto de Botánica, Departamento de Botánica y Zoología, Centro Universitario de Ciencias Biológicas y Agropecuarias, Universidad de Guadalajara, Apartado postal 1-139, CP 45101 Zapopan, Jalisco, Mexico. Electronic address:. Facultad de Ciencias Naturales, Universidad Autónoma de Querétaro, Av. De las Ciencias s/n, Juriquilla, CP 76230, Del. Sta. Rosa Jáuregui, Querétaro, Mexico. Electronic address:. Instituto de Biología, Universidad Nacional Autónoma de México, Ciudad de México, Mexico. Electronic address:. Instituto de Botánica, Departamento de Botánica y Zoología, Centro Universitario de Ciencias Biológicas y Agropecuarias, Universidad de Guadalajara, Apartado postal 1-139, CP 45101 Zapopan, Jalisco, Mexico. Electronic address:. Instituto de Botánica, Departamento de Botánica y Zoología, Centro Universitario de Ciencias Biológicas y Agropecuarias, Universidad de Guadalajara, Apartado postal 1-139, CP 45101 Zapopan, Jalisco, Mexico. Electronic address:. M4 - Citavi
AU  - Zamora-Tavares, María Del Pilar
AU  - Martínez, Mahinda
AU  - Magallón, Susana
AU  - Guzmán-Dávalos, Laura
AU  - Vargas-Ponce, Ofelia
DO  - 10.1016/j.ympev.2016.03.032
KW  - Solanaceae
LA  - eng
N1  - Journal Article Journal Article Journal Article Journal Article
PY  - 2016
SN  - 1055-7903
SP  - 41-50
ST  - Physalis and physaloids: A recent and complex evolutionary history
T2  - Molecular phylogenetics and evolution
TI  - Physalis and physaloids: A recent and complex evolutionary history
VL  - 100
ID  - 416
ER  - 


TY  - JOUR
AU  - Zamberlan, Priscilla M.
AU  - Rodrigues, Izabella M. C.
AU  - Geraldo Mäder, Luana Castro João R. Stehmann
AU  - Bonatto, Sandro Luis
AU  - Freitas, Loreta Brandão
KW  - Solanaceae
PY  - 2015
SP  - 322-334
ST  - Re-evaluation of the generic status of Athenaea and Aureliana (Withaniinae, Solanaceae) based on molecular phylogeny and morphology of the calyx
T2  - Botanical Journal of the Linnean Society
TI  - Re-evaluation of the generic status of Athenaea and Aureliana (Withaniinae, Solanaceae) based on molecular phylogeny and morphology of the calyx
VL  - 177
ID  - 795
ER  - 


TY  - JOUR
AU  - Whitson, Maggie
AU  - Manos, Paul S.
IS  - 1
KW  - Solanaceae
PY  - 2005
SN  - 03636445
SP  - 216-230
ST  - Untangling Physalis (Solanaceae) from the Physaloids: A Two-Gene Phylogeny of the Physalinae
T2  - Systematic Botany
TI  - Untangling Physalis (Solanaceae) from the Physaloids: A Two-Gene Phylogeny of the Physalinae
VL  - 30
ID  - 3724
ER  - 


TY  - JOUR
AU  - Weese, Terri L.
AU  - Bohs, Lynn
IS  - 1
KW  - Solanaceae
PY  - 2010
SN  - 00400262
SP  - 49-56
ST  - Eggplant origins: Out of Africa, into the Orient
T2  - Taxon
TI  - Eggplant origins: Out of Africa, into the Orient
VL  - 59
ID  - 2394
ER  - 


TY  - JOUR
AU  - Weese, Terri L.
AU  - Bohs, Lynn
IS  - 2
KW  - Solanaceae
PY  - 2007
SN  - 03636445
SP  - 445-463
ST  - A Three-Gene Phylogeny of the Genus Solanum (Solanaceae)
T2  - Systematic Botany
TI  - A Three-Gene Phylogeny of the Genus Solanum (Solanaceae)
VL  - 32
ID  - 3300
ER  - 


TY  - JOUR
AB  - International Journal of Plant Sciences 2001.162:1409-1418
AU  - Walsh, Brian M.
AU  - Hoot, Sara B.
IS  - 6
KW  - Solanaceae
PY  - 2001
SN  - 1058-5893
SP  - 1409-1418
ST  - Phylogenetic Relationships of Capsicum (Solanaceae) Using DNA Sequences from Two Noncoding Regions: The Chloroplast atpB‐rbcL Spacer Region and Nuclear waxy Introns
T2  - International Journal of Plant Sciences
TI  - Phylogenetic Relationships of Capsicum (Solanaceae) Using DNA Sequences from Two Noncoding Regions: The Chloroplast atpB‐rbcL Spacer Region and Nuclear waxy Introns
VL  - 162
ID  - 4295
ER  - 


TY  - JOUR
AU  - Volkov, Roman A.
AU  - Komarova, Nataliya Y.
AU  - Panchuk, Irina I.
AU  - Hemleben, Vera
DO  - 10.1016/S1055-7903(03)00092-7 M4 - Citavi
IS  - 2
KW  - Solanaceae
N1  - PII: S1055790303000927 PII: S1055790303000927 PII: S1055790303000927 PII: S1055790303000927
PY  - 2003
SN  - 1055-7903
SP  - 187-202
ST  - Molecular evolution of rDNA external transcribed spacer and phylogeny of sect. Petota (genus Solanum)
T2  - Molecular phylogenetics and evolution
TI  - Molecular evolution of rDNA external transcribed spacer and phylogeny of sect. Petota (genus Solanum)
VL  - 29
ID  - 4064
ER  - 


TY  - JOUR
AB  - The cosmopolitan Solanaceae contains 21 tribes and has the greatest diversity in South America. Hyoscyameae and Mandragoreae are the only tribes of this family distributed exclusively in Eurasia with two centers of diversity: the Mediterranean-Turanian (MT) region and the Tibetan Plateau (TP). In this study, we examined the origins and biogeographical diversifications of the two tribes based on the phylogenetic framework and chronogram inferred from a combined data set of six plastid DNA regions (the atpB gene, the ndhF gene, the rps16-trnK intergenic spacer, the rbcL gene, the trnC-psbM region and the psbA-trnH intergenic spacer) with two fossil calibration points. Our data suggest that Hyoscyameae and Mandragoreae each forms a monophyletic group independently derived from different New World lineages in the early Miocene. Phylogenetic relationships within both tribes are generally well resolved. All genera of Hyoscyameae are found to be monophyletic and they diversified in middle to late Miocene. At nearly the same time, Mandragoreae split into two clades, corresponding to the MT region and the TP region, respectively. Both the phylogenetic relationships and the estimated ages of Hyoscyameae and Mandragoreae support two independent dispersal events of their ancestors from the New World into Eurasia. After their arrivals in Eurasia, the two tribes diversified primarily in the MT region and in the TP region via multiple biogeographic processes including vicariance, dispersal, recolonization or being preserved as relicts, from the mid Miocene to the late Quaternary.
AD  - Key Laboratory of Biodiversity and Biogeography, Kunming Institute of Botany, Chinese Academic of Sciences, Kunming 650204, PR China. M4 - Citavi
AU  - Tu, Tieyao
AU  - Volis, Sergei
AU  - Dillon, Michael O.
AU  - Sun, Hang
AU  - Wen, Jun
DO  - 10.1016/j.ympev.2010.09.007
IS  - 3
KW  - Solanaceae
LA  - eng
N1  - Journal Article Research Support, Non-U.S. Gov't Research Support, U.S. Gov't, Non-P.H.S. Journal Article Research Support, Non-U.S. Gov't Research Support, U.S. Gov't, Non-P.H.S. Journal Article Research Support, Non-U.S. Gov't Research Support, U.S. Gov't, Non-P.H.S. Journal Article Research Support, Non-U.S. Gov't Research Support, U.S. Gov't, Non-P.H.S.
PY  - 2010
SN  - 1055-7903
SP  - 1226-1237
ST  - Dispersals of Hyoscyameae and Mandragoreae (Solanaceae) from the New World to Eurasia in the early Miocene and their biogeographic diversification within Eurasia
T2  - Molecular phylogenetics and evolution
TI  - Dispersals of Hyoscyameae and Mandragoreae (Solanaceae) from the New World to Eurasia in the early Miocene and their biogeographic diversification within Eurasia
VL  - 57
ID  - 2377
ER  - 


TY  - JOUR
AB  - The phylogeny of Nolana (Solanaceae), a genus primarily distributed in the coastal Atacama and Peruvian deserts with a few species in the Andes and one species endemic to the Galápagos Islands, was reconstructed using sequences of four plastid regions (ndhF, psbA-trnH, rps16-trnK and trnC-psbM) and the nuclear LEAFY second intron. The monophyly of Nolana was strongly supported by all molecular data. The LEAFY data suggested that the Chilean species, including Nolana sessiliflora, the N. acuminata group and at least some members of the Alona group, are basally diverged, supporting the Chilean origin of the genus. Three well-supported clades in the LEAFY tree were corroborated by the SINE (short interspersed elements) or SINE-like insertions. Taxa from Peru are grouped roughly into two clades. Nolana galapagensis from the Galápagos Island is most likely to have derived from a Peruvian ancestor. The monophyly of the morphologically well-diagnosed Nolana acuminata group (N. acuminata, N. baccata, N. paradoxa, N. parviflora, N. pterocarpa, N. rupicola and N. elegans) was supported by both plastid and LEAFY data. Incongruence between the plastid and the LEAFY data was detected concerning primarily the positions of N. sessiliflora, N. galapagensis, taxa of the Alona group and the two Peruvian clades. Such incongruence may be due to reticulate evolution or in some cases lineage sorting of plastid DNA. Incongruence between our previous GBSSI trees and the plastid-LEAFY trees was also detected concerning two well-supported major clades in the GBSSI tree. Duplication of the GBSSI gene may have contributed to this incongruence.
AD  - Key Laboratory of Biodiversity and Biogeography, Kunming Institute of Botany, Chinese Academy of Sciences, Kunming, Yunnan 650204, PR China. M4 - Citavi
AU  - Tu, Tieyao
AU  - Dillon, Michael O.
AU  - Sun, Hang
AU  - Wen, Jun
DO  - 10.1016/j.ympev.2008.07.018
IS  - 2
KW  - Solanaceae
LA  - eng
N1  - Journal Article Research Support, Non-U.S. Gov't Journal Article Research Support, Non-U.S. Gov't Journal Article Research Support, Non-U.S. Gov't Journal Article Research Support, Non-U.S. Gov't
PY  - 2008
SN  - 1055-7903
SP  - 561-573
ST  - Phylogeny of Nolana (Solanaceae) of the Atacama and Peruvian deserts inferred from sequences of four plastid markers and the nuclear Leafy second intron
T2  - Molecular phylogenetics and evolution
TI  - Phylogeny of Nolana (Solanaceae) of the Atacama and Peruvian deserts inferred from sequences of four plastid markers and the nuclear Leafy second intron
VL  - 49
ID  - 3003
ER  - 


TY  - JOUR
AB  - PREMISE OF THE STUDY Solanum section Herpystichum is a lineage that comprises both widespread and very narrowly distributed species. This study investigates the phylogenetic relationships of sect. Herpystichum and evaluates several phylogenetic methods for analysis of multiple sequences. METHODS Sequence data from seven nuclear (ITS, GBSSI, and five COSII) and three plastid (psbA-trnH, trnT-trnF, and trnS-trnG) regions were concatenated and analyzed under maximum parsimony and Bayesian criteria. In addition, we used two analytical methods that take into account differences in topologies resulting from the analyses of the individual markers: Bayesian Estimation of Species Trees (BEST) and supertree analysis. KEY RESULTS The monophyletic Solanum sect. Herpystichum was resolved with moderate support in the concatenated maximum parsimony and Bayesian analyses and the supertree analysis, and relationships within the section were well-resolved and strongly supported. The BEST topology, however, was poorly resolved. Also, because of how BEST deals with missing sequences, >25% of our accessions, including two species, had to be excluded from the analyses. Our results indicate a progenitor-descendent relationship with two species nested within the widespread S. evolvulifolium. CONCLUSIONS Analytical methods that consider individual topologies are important for studies based on multiple molecular markers. On the basis of analyses in this study, BEST had the serious shortcoming that taxa with missing sequences must be removed from the analysis or they can produce spurious topologies. Supertree analysis provided a good alternative for our data by allowing the inclusion of all 10 species of sect. Herpystichum.
AD  - Department of Biology, 257 South 1400 East, University of Utah, Salt Lake City, Utah 84112, USA. M4 - Citavi
AU  - Tepe, Eric J.
AU  - Farruggia, Frank T.
AU  - Bohs, Lynn
DO  - 10.3732/ajb.1000516
IS  - 8
KW  - Solanaceae
LA  - eng
N1  - Comparative Study Journal Article Research Support, U.S. Gov't, Non-P.H.S. Comparative Study Journal Article Research Support, U.S. Gov't, Non-P.H.S. Comparative Study Journal Article Research Support, U.S. Gov't, Non-P.H.S. Comparative Study Journal Article Research Support, U.S. Gov't, Non-P.H.S.
PY  - 2011
SN  - 0002-9122
SP  - 1356-1365
ST  - A 10-gene phylogeny of Solanum section Herpystichum (Solanaceae) and a comparison of phylogenetic methods
T2  - American journal of botany
TI  - A 10-gene phylogeny of Solanum section Herpystichum (Solanaceae) and a comparison of phylogenetic methods
VL  - 98
ID  - 2055
ER  - 


TY  - JOUR
AU  - Tepe, Eric J.
AU  - Bohs, Lynn
DO  - 10.1600/036364411X605074 M4 - Citavi
IS  - 4
KW  - Solanaceae
PY  - 2011
SN  - 03636445
SP  - 1068-1087
ST  - A Revision of Solanum Section Herpystichum
T2  - Systematic Botany
TI  - A Revision of Solanum Section Herpystichum
VL  - 36
ID  - 2054
ER  - 


TY  - JOUR
AU  - Tepe, Eric J.
AU  - Bohs, Lynn
IS  - 3
KW  - Solanaceae
PY  - 2010
SN  - 00400262
SP  - 733-743
ST  - A molecular phylogeny of Solanum sect. Pteroidea (Solanaceae) and the utility of Cosii markers in resolving relationships among closely related species
T2  - Taxon
TI  - A molecular phylogeny of Solanum sect. Pteroidea (Solanaceae) and the utility of Cosii markers in resolving relationships among closely related species
VL  - 59
ID  - 2369
ER  - 


TY  - JOUR
AU  - Tate, Jennifer A.
AU  - Acosta, M. Cristina
AU  - McDill, Joshua
AU  - Moscone, Eduardo A.
AU  - Simpson, Beryl B.
AU  - Cocucci, Andrea A.
IS  - 1
KW  - Solanaceae
PY  - 2009
SN  - 03636445
SP  - 198-206
ST  - Phylogeny and Character Evolution in Nierembergia (Solanaceae): Molecular, Morphological, and Cytogenetic Evidence
T2  - Systematic Botany
TI  - Phylogeny and Character Evolution in Nierembergia (Solanaceae): Molecular, Morphological, and Cytogenetic Evidence
VL  - 34
ID  - 2692
ER  - 


TY  - JOUR
AU  - Stern, Stephen R.
AU  - Weese, Terri
AU  - Bohs, Lynn
DO  - 10.1600/036364410X539934 M4 - Citavi
IS  - 4
KW  - Solanaceae
PY  - 2010
SN  - 03636445
SP  - 885-893
ST  - Phylogenetic Relationships in Solanum Section Androceras (Solanaceae)
T2  - Systematic Botany
TI  - Phylogenetic Relationships in Solanum Section Androceras (Solanaceae)
VL  - 35
ID  - 2355
ER  - 


TY  - JOUR
AU  - Stern, Stephen
AU  - Bohs, Lynn
DO  - 10.1600/036364416X690589 M4 - Citavi
IS  - 1
KW  - Solanaceae
PY  - 2016
SN  - 03636445
SP  - 120-127
ST  - An eight Marker Phylogeny of Solanum sect. Micracantha (Solanaceae)
T2  - Systematic Botany
TI  - An eight Marker Phylogeny of Solanum sect. Micracantha (Solanaceae)
VL  - 41
ID  - 369
ER  - 


TY  - JOUR
AU  - Stern, Stephen
AU  - Agra, Fátima de Maria
AU  - Bohs, Lynn
IS  - 5
KW  - Solanaceae
PY  - 2011
SN  - 00400262
SP  - 1429-1441
ST  - Molecular delimitation of clades within New World species of the "spiny solanums" (Solanum subg. Leptostemonum)
T2  - Taxon
TI  - Molecular delimitation of clades within New World species of the "spiny solanums" (Solanum subg. Leptostemonum)
VL  - 60
ID  - 2045
ER  - 


TY  - JOUR
AU  - Smith, Stacey Dewitt
AU  - Baum, David A.
IS  - 8
KW  - Solanaceae
PY  - 2006
SN  - 0002-9122
SP  - 1140-1153
ST  - Phylogenetics of the florally diverse Andean Clade Iochrominae (Solanaceae)1
T2  - American journal of botany
TI  - Phylogenetics of the florally diverse Andean Clade Iochrominae (Solanaceae)1
VL  - 93
ID  - 3495
ER  - 


TY  - JOUR
AB  - BACKGROUND The Solanaceae is a plant family of great economic importance. Despite a wealth of phylogenetic work on individual clades and a deep knowledge of particular cultivated species such as tomato and potato, a robust evolutionary framework with a dated molecular phylogeny for the family is still lacking. Here we investigate molecular divergence times for Solanaceae using a densely-sampled species-level phylogeny. We also review the fossil record of the family to derive robust calibration points, and estimate a chronogram using an uncorrelated relaxed molecular clock. RESULTS Our densely-sampled phylogeny shows strong support for all previously identified clades of Solanaceae and strongly supported relationships between the major clades, particularly within Solanum. The Tomato clade is shown to be sister to section Petota, and the Regmandra clade is the first branching member of the Potato clade. The minimum age estimates for major splits within the family provided here correspond well with results from previous studies, indicating splits between tomato and potato around 8 Million years ago (Ma) with a 95% highest posterior density (HPD) 7-10 Ma, Solanum and Capsicum c. 19 Ma (95% HPD 17-21), and Solanum and Nicotiana c. 24 Ma (95% HPD 23-26). CONCLUSIONS Our large time-calibrated phylogeny provides a significant step towards completing a fully sampled species-level phylogeny for Solanaceae, and provides age estimates for the whole family. The chronogram now includes 40% of known species and all but two monotypic genera, and is one of the best sampled angiosperm family phylogenies both in terms of taxon sampling and resolution published thus far. The increased resolution in the chronogram combined with the large increase in species sampling will provide much needed data for the examination of many biological questions using Solanaceae as a model system.
AU  - Särkinen, Tiina
AU  - Bohs, Lynn
AU  - Olmstead, Richard G.
AU  - Knapp, Sandra
DO  - 10.1186/1471-2148-13-214
KW  - Solanaceae
LA  - eng
N1  - Journal Article Research Support, U.S. Gov't, Non-P.H.S. Journal Article Research Support, U.S. Gov't, Non-P.H.S. Journal Article Research Support, U.S. Gov't, Non-P.H.S. Journal Article Research Support, U.S. Gov't, Non-P.H.S.
PY  - 2013
SP  - 214
ST  - A phylogenetic framework for evolutionary study of the nightshades (Solanaceae): a dated 1000-tip tree
T2  - BMC evolutionary biology
TI  - A phylogenetic framework for evolutionary study of the nightshades (Solanaceae): a dated 1000-tip tree
UR  - https://www.ncbi.nlm.nih.gov/pmc/articles/PMC3850475
VL  - 13
ID  - 1373
ER  - 


TY  - JOUR
AU  - Särkinen, Tiina
AU  - Barboza, Gloria Estela
AU  - Knapp, Sandra
DO  - 10.12705/645.5 M4 - Citavi
IS  - 5
KW  - Solanaceae
PY  - 2015
SN  - 00400262
SP  - 945-958
ST  - True Black nightshades: Phylogeny and delimitation of the Morelloid clade of Solanum
T2  - Taxon
TI  - True Black nightshades: Phylogeny and delimitation of the Morelloid clade of Solanum
VL  - 64
ID  - 699
ER  - 


TY  - JOUR
AU  - Reck-Kortmann, Maikel
AU  - Silva-Arias, Gustavo A.
AU  - Stehmann, João R.
AU  - Greppi, Julián A.
AU  - Freitas, Loreta Brandão
DO  - 10.11646/phytotaxa.222.1.2 M4 - Citavi
IS  - 1
KW  - Solanaceae
PY  - 2015
SN  - 1179-3163
SP  - 17
ST  - Phylogenetic relationships of Petunia patagonica (Solanaceae) revealed by molecular and biogeographical evidence
T2  - Phytotaxa
TI  - Phylogenetic relationships of Petunia patagonica (Solanaceae) revealed by molecular and biogeographical evidence
VL  - 222
ID  - 681
ER  - 


TY  - JOUR
AB  - The phylogeny of Petunia species has been difficult to resolve, primarily due to the recent diversification of the genus. Several studies have included molecular data in phylogenetic reconstructions of this genus, but all of them have failed to include all taxa and/or analyzed few genetic markers. In the present study, we employed the most inclusive genetic and taxonomic datasets for the genus, aiming to reconstruct the evolutionary history of Petunia based on molecular phylogeny, biogeographic distribution, and character evolution. We included all 20 Petunia morphological species or subspecies in these analyses. Based on nine nuclear and five plastid DNA markers, our phylogenetic analysis reinforces the monophyly of the genus Petunia and supports the hypothesis that the basal divergence is more related to the differentiation of corolla tube length, whereas the geographic distribution of species is more related to divergences within these main clades. Ancestral area reconstructions suggest the Pampas region as the area of origin and earliest divergence in Petunia. The state reconstructions suggest that the ancestor of Petunia might have had a short corolla tube and a bee pollination floral syndrome.
AU  - Reck-Kortmann, Maikel
AU  - Silva-Arias, Gustavo Adolfo
AU  - Segatto, Ana Lúcia Anversa
AU  - Mäder, Geraldo
AU  - Bonatto, Sandro Luis
AU  - Freitas, Loreta Brandão
DO  - 10.1016/j.ympev.2014.08.022
KW  - Solanaceae
LA  - eng
PY  - 2014
SN  - 1055-7903
SP  - 19-28
ST  - Multilocus phylogeny reconstruction: new insights into the evolutionary history of the genus Petunia
T2  - Molecular phylogenetics and evolution
TI  - Multilocus phylogeny reconstruction: new insights into the evolutionary history of the genus Petunia
VL  - 81
ID  - 1034
ER  - 


TY  - JOUR
AB  - BACKGROUND Quaternary climatic changes led to variations in sea level and these variations played a significant role in the generation of marine terrace deposits in the South Atlantic Coastal Plain. The main consequence of the increase in sea level was local extinction or population displacement, such that coastal species would be found around the new coastline. Our main goal was to investigate the effects of sea level changes on the geographical structure and variability of genetic lineages from a Petunia species endemic to the South Atlantic Coastal Plain. We employed a phylogeographic approach based on plastid sequences obtained from individuals collected from the complete geographic distribution of Petunia integrifolia ssp. depauperata and its sister group. We used population genetics tests to evaluate the degree of genetic variation and structure among and within populations, and we used haplotype network analysis and Bayesian phylogenetic methods to estimate divergence times and population growth. RESULTS We observed three major genetic lineages whose geographical distribution may be related to different transgression/regression events that occurred in this region during the Pleistocene. The divergence time between the monophyletic group P. integrifolia ssp. depauperata and its sister group (P. integrifolia ssp. integrifolia) was compatible with geological estimates of the availability of the coastal plain. Similarly, the origin of each genetic lineage is congruent with geological estimates of habitat availability. CONCLUSIONS Diversification of P. integrifolia ssp. depauperata possibly occurred as a consequence of the marine transgression/regression cycles during the Pleistocene. In periods of high sea level, plants were most likely restricted to a refuge area corresponding to fossil dunes and granitic hills, from which they colonized the coast once the sea level came down. The modern pattern of lineage geographical distribution and population variation was established by a range expansion with serial founder effects conditioned on soil availability.
AD  - Laboratory of Molecular Evolution, Department of Genetics, Universidade Federal do Rio Grande do Sul, P.O. Box 15053, Porto Alegre, Brazil.. Laboratory of Molecular Evolution, Department of Genetics, Universidade Federal do Rio Grande do Sul, P.O. Box 15053, Porto Alegre, Brazil.. Department of Statistics, Universidade Federal do Rio Grande do Sul, P.O. Box 15080, Porto Alegre, Brazil.. Laboratory of Molecular Evolution, Department of Genetics, Universidade Federal do Rio Grande do Sul, P.O. Box 15053, Porto Alegre, Brazil.. Genomic and Molecular Biology Laboratory, Pontifícia Universidade Católica do Rio Grande do Sul, Ipiranga 6681, 90610 001, Porto Alegre, RS, Brazil.. Laboratory of Molecular Evolution, Department of Genetics, Universidade Federal do Rio Grande do Sul, P.O. Box 15053, Porto Alegre, Brazil.. M4 - Citavi
AU  - Ramos-Fregonezi, Aline M. C.
AU  - Fregonezi, Jefferson Nunes
AU  - Cybis, Gabriela B.
AU  - Fagundes, Nelson J. R.
AU  - Bonatto, Sandro Luis
AU  - Freitas, Loreta Brandão
DO  - 10.1186/s12862-015-0363-8
KW  - Solanaceae
LA  - eng
N1  - Journal Article Research Support, Non-U.S. Gov't Journal Article Research Support, Non-U.S. Gov't Journal Article Research Support, Non-U.S. Gov't Journal Article Research Support, Non-U.S. Gov't
PY  - 2015
SP  - 92
ST  - Were sea level changes during the Pleistocene in the South Atlantic Coastal Plain a driver of speciation in Petunia (Solanaceae)?
T2  - BMC evolutionary biology
TI  - Were sea level changes during the Pleistocene in the South Atlantic Coastal Plain a driver of speciation in Petunia (Solanaceae)?
UR  - https://www.ncbi.nlm.nih.gov/pmc/articles/PMC4438590
VL  - 15
ID  - 678
ER  - 


TY  - JOUR
AU  - Poczai, P.
AU  - Taller, J.
AU  - Szabó, I.
DO  - 10.1007/s00606-008-0051-x M4 - Citavi
IS  - 1-2
KW  - Solanaceae
N1  - PII: 51 PII: 51 PII: 51 PII: 51
PY  - 2008
SN  - 0378-2697
SP  - 59-67
ST  - Analysis of phylogenetic relationships in the genus Solanum (Solanaceae) as revealed by Rapd markers
T2  - Plant Systematics and Evolution
TI  - Analysis of phylogenetic relationships in the genus Solanum (Solanaceae) as revealed by Rapd markers
VL  - 275
ID  - 2934
ER  - 


TY  - JOUR
AD  - Instituto de Ciencias Naturales, Universidad Nacional de Colombia, Carrera 30 45-03, edificio 425, Bogotá, Colombia;, Department of Biology, University of Utah, Salt Lake City, Utah 84112, U.S.A. Instituto de Ciencias Naturales, Universidad Nacional de Colombia, Carrera 30 45-03, edificio 425, Bogotá, Colombia Instituto Multidisciplinario de Biología Vegetal, Museo Botánico de Córdoba, CONICET-UNC, Córdoba, 5000 Argentina M4 - Citavi
AU  - Orejuela, Andrés
AU  - Wahlert, GregoryA
AU  - Orozco, Clara Inés
AU  - Barboza, Gloria Estela
AU  - Bohs, Lynn
DO  - 10.12705/662.6
IS  - 2
KW  - Solanaceae
PY  - 2017
SN  - 00400262
SP  - 379-392
ST  - Phylogeny of the tribes Juanulloeae and Solandreae (Solanaceae)
T2  - Taxon
TI  - Phylogeny of the tribes Juanulloeae and Solandreae (Solanaceae)
VL  - 66
ID  - 101
ER  - 


TY  - JOUR
AU  - Olmstead, Richard G.
AU  - Bohs, Lynn
AU  - Migid, Hala Abdel
AU  - Santiago-Valentin, Eugenio
AU  - Garcia, Vicente F.
AU  - Collier, Sarah M.
IS  - 4
KW  - Solanaceae
PY  - 2008
SN  - 00400262
SP  - 1159-1181
ST  - A molecular phylogeny of the Solanaceae
T2  - Taxon
TI  - A molecular phylogeny of the Solanaceae
VL  - 57
ID  - 2919
ER  - 


TY  - JOUR
AB  - One classic explanation for the remarkable diversity of flower colors across angiosperms involves evolutionary shifts among different types of pollinators with different color preferences. However, the pollinator shift model fails to account for the many examples of color variation within clades that share the same pollination system. An alternate explanation is the competition model, which suggests that color divergence evolves in response to interspecific competition for pollinators, as a means to decrease interspecific pollinator movements. This model predicts color overdispersion within communities relative to null assemblages. Here, we combine morphometric analyses, field surveys, and models of pollinator vision with a species-level phylogeny to test the competition model in the primarily hummingbird-pollinated clade Iochrominae (Solanaceae). Results show that flower color as perceived by pollinators is significantly overdispersed within sites. This pattern is not simply due to phylogenetic history: phylogenetic community structure does not deviate from random expectations, and flower color lacks phylogenetic signal. Moreover, taxa that occur in sympatry occupy a significantly larger volume of color space than those in allopatry, supporting the hypothesis that competition in sympatry drove the evolution of novel colors. We suggest that competition among close relatives may commonly underlie floral divergence, especially in species-rich habitats where congeners frequently co-occur.
AU  - Muchhala, Nathan
AU  - Johnsen, Sönke
AU  - Smith, Stacey Dewitt
DO  - 10.1111/evo.12441
IS  - 8
KW  - Solanaceae
LA  - eng
PY  - 2014
SN  - 0014-3820
SP  - 2275-2286
ST  - Competition for hummingbird pollination shapes flower color variation in Andean solanaceae
T2  - Evolution
TI  - Competition for hummingbird pollination shapes flower color variation in Andean solanaceae
VL  - 68
ID  - 995
ER  - 


TY  - JOUR
AU  - Moré, Marcela
AU  - Cocucci, Andrea A.
AU  - Sérsic, Alicia N.
AU  - Barboza, Gloria Estela
DO  - 10.12705/643.8 M4 - Citavi
IS  - 3
KW  - Solanaceae
PY  - 2015
SN  - 00400262
SP  - 523-534
ST  - Phylogeny and floral trait evolution in Jaborosa (Solanaceae)
T2  - Taxon
TI  - Phylogeny and floral trait evolution in Jaborosa (Solanaceae)
VL  - 64
ID  - 636
ER  - 


TY  - JOUR
AU  - Montero-Castro, Juan Carlos
AU  - Delgado-Salinas, Alfonso
AU  - Luna, Efraín de
AU  - Eguiarte, Luis E.
IS  - 4
KW  - Solanaceae
PY  - 2006
SN  - 03636445
SP  - 843-850
ST  - Phylogenetic Analysis of Cestrum Section Habrothamnus (Solanaceae) Based on Plastid and Nuclear DNA Sequences
T2  - Systematic Botany
TI  - Phylogenetic Analysis of Cestrum Section Habrothamnus (Solanaceae) Based on Plastid and Nuclear DNA Sequences
VL  - 31
ID  - 3453
ER  - 


TY  - JOUR
AU  - Miller, Ryan J.
AU  - Mione, Thomas
AU  - Phan, Hanh-La
AU  - Olmstead, Richard G.
DO  - 10.1600/036364411X553243 M4 - Citavi
IS  - 1
KW  - Solanaceae
PY  - 2011
SN  - 03636445
SP  - 153-162
ST  - Color by Numbers: Nuclear Gene Phylogeny of Jaltomata (Solanaceae), Sister Genus to Solanum, supports three Clades differing in Fruit Color
T2  - Systematic Botany
TI  - Color by Numbers: Nuclear Gene Phylogeny of Jaltomata (Solanaceae), Sister Genus to Solanum, supports three Clades differing in Fruit Color
VL  - 36
ID  - 1973
ER  - 


TY  - JOUR
AU  - Martins, Talline R.
AU  - Barkman, Todd J.
IS  - 2
KW  - Solanaceae
PY  - 2005
SN  - 03636445
SP  - 435-447
ST  - Reconstruction of Solanaceae Phylogeny Using the Nuclear Gene SAMT
T2  - Systematic Botany
TI  - Reconstruction of Solanaceae Phylogeny Using the Nuclear Gene SAMT
VL  - 30
ID  - 3646
ER  - 


TY  - JOUR
AU  - Martine, Christopher T.
AU  - Vanderpool, Dan
AU  - Anderson, Gregory J.
AU  - Les, Donald H.
IS  - 2
KW  - Solanaceae
PY  - 2006
SN  - 03636445
SP  - 410-420
ST  - Phylogenetic Relationships of Andromonoecious and Dioecious Australian Species of <i>Solanum</i> subgenus <i>Leptostemonum</i> section <i>Melongena</i>: Inferences from ITS Sequence Data
T2  - Systematic Botany
TI  - Phylogenetic Relationships of Andromonoecious and Dioecious Australian Species of <i>Solanum</i> subgenus <i>Leptostemonum</i> section <i>Melongena</i>: Inferences from ITS Sequence Data
VL  - 31
ID  - 3441
ER  - 


TY  - JOUR
AU  - Martine, Christopher T.
AU  - Anderson, Gregory J.
AU  - Les, Donald H.
DO  - 10.1071/SB07039 M4 - Citavi
IS  - 2
KW  - Solanaceae
N1  - PII: SB07039 PII: SB07039 PII: SB07039 PII: SB07039
PY  - 2009
SN  - 1030-1887
SP  - 107-120
ST  - Gender-bending aubergines: molecular phylogenetics of cryptically dioecious Solanum in Australia
T2  - Australian Systematic Botany
TI  - Gender-bending aubergines: molecular phylogenetics of cryptically dioecious Solanum in Australia
VL  - 22
ID  - 2595
ER  - 


TY  - JOUR
AU  - Marks, Claire E.
AU  - Newbigin, Ed
AU  - Ladiges, Pauline Y.
DO  - 10.1071/SB11006 M4 - Citavi
IS  - 2
KW  - Solanaceae
N1  - PII: SB11006 PII: SB11006 PII: SB11006 PII: SB11006
PY  - 2011
SN  - 1030-1887
SP  - 61
ST  - Comparative morphology and phylogeny of Nicotiana section Suaveolentes (Solanaceae) in Australia and the South Pacific
T2  - Australian Systematic Botany
TI  - Comparative morphology and phylogeny of Nicotiana section Suaveolentes (Solanaceae) in Australia and the South Pacific
VL  - 24
ID  - 1966
ER  - 


TY  - JOUR
AU  - Li, Hong-Qing
AU  - Gui, Ping
AU  - Xiong, Shen-Zhan
AU  - Averett, John E.
DO  - 10.1016/j.bse.2013.03.038 M4 - Citavi
KW  - Solanaceae
N1  - PII: S0305197813000835 PII: S0305197813000835 PII: S0305197813000835 PII: S0305197813000835
PY  - 2013
SN  - 03051978
SP  - 82-89
ST  - The generic position of two species of tribe Physaleae (Solanaceae) inferred from three DNA sequences: A case study on Physaliastrum and Archiphysalis
T2  - Biochemical Systematics and Ecology
TI  - The generic position of two species of tribe Physaleae (Solanaceae) inferred from three DNA sequences: A case study on Physaliastrum and Archiphysalis
VL  - 50
ID  - 1283
ER  - 


TY  - JOUR
AB  - The identification of genomic regions with sufficient variation to elucidate fine-scale relationships among closely related species is a major goal of phylogenetic systematics. However, the accumulation of such multi-locus data sets brings its own challenges, given that gene trees do not necessarily represent the true species tree. Using genomic tools developed for Solanum (Solanaceae), we have evaluated the utility of nuclear conserved ortholog set II (COSII) regions for phylogenetic inference in tribe Lycieae (Solanaceae). Five COSII regions, with intronic contents ranging from 68% to 91%, were sequenced in 10 species. Their phylogenetic utility was assessed and compared with data from more commonly used nuclear (GBSSI, nrITS) and cpDNA spacer data. We compared the effectiveness of a traditional total evidence concatenation approach versus the recently developed Bayesian estimation of species trees (BEST) method to infer species trees given multiple independent gene trees. All of the sampled COSII regions had high numbers of parsimony-informative (PI) characters, and two of the COSII regions had more PI characters than the GBSSI, ITS, and cpDNA spacer data sets combined. COSII markers are a promising new tool for phylogenetic inference in Solanaceae, and should be explored in related groups. Both the concatenation and BEST approaches yielded similar topologies; however, when multiple individuals with polyphyletic alleles were included, BEST was clearly the more robust approach for inferring species trees in the presence of gene tree incongruence.
AD  - Department of Biology, McGuire Life Sciences Building, Amherst College, Amherst, MA 01002, USA. M4 - Citavi
AU  - Levin, Rachel A.
AU  - Whelan, Andrew
AU  - Miller, Jill S.
DO  - 10.1016/j.ympev.2009.08.016
IS  - 3
KW  - Solanaceae
LA  - eng
N1  - Journal Article Research Support, Non-U.S. Gov't Research Support, U.S. Gov't, Non-P.H.S. Journal Article Research Support, Non-U.S. Gov't Research Support, U.S. Gov't, Non-P.H.S. Journal Article Research Support, Non-U.S. Gov't Research Support, U.S. Gov't, Non-P.H.S. Journal Article Research Support, Non-U.S. Gov't Research Support, U.S. Gov't, Non-P.H.S.
PY  - 2009
SN  - 1055-7903
SP  - 881-890
ST  - The utility of nuclear conserved ortholog set II (Cosii) genomic regions for species-level phylogenetic inference in Lycium (Solanaceae)
T2  - Molecular phylogenetics and evolution
TI  - The utility of nuclear conserved ortholog set II (Cosii) genomic regions for species-level phylogenetic inference in Lycium (Solanaceae)
VL  - 53
ID  - 2582
ER  - 


TY  - JOUR
AU  - Levin, Rachel A.
AU  - Watson, Kimberly
AU  - Bohs, Lynn
IS  - 4
KW  - Solanaceae
PY  - 2005
SN  - 0002-9122
SP  - 603-612
ST  - A Four-gene Study of Evolutionary Relationships in Solanum Section Acanthophora
T2  - American journal of botany
TI  - A Four-gene Study of Evolutionary Relationships in Solanum Section Acanthophora
VL  - 92
ID  - 3630
ER  - 


TY  - JOUR
AU  - Levin, Rachel A.
AU  - Myers, Nicole R.
AU  - Bohs, Lynn
IS  - 1
KW  - Solanaceae
PY  - 2006
SN  - 0002-9122
SP  - 157-169
ST  - Phylogenetic Relationships Among The ‘‘spiny Solanums’’ (Solanum Subgenus Leptostemonum, Solanaceae)
T2  - American journal of botany
TI  - Phylogenetic Relationships Among The ‘‘spiny Solanums’’ (Solanum Subgenus Leptostemonum, Solanaceae)
VL  - 93
ID  - 3424
ER  - 


TY  - JOUR
AU  - Levin, Rachel A.
AU  - Miller, Jill S.
IS  - 12
KW  - Solanaceae
PY  - 2005
SN  - 0002-9122
SP  - 2044-2053
ST  - Relationships within Tribe Lycieae (Solanaceae): Paraphyly Of Lycium And Multiple Origins Of Gender Dimorphism
T2  - American journal of botany
TI  - Relationships within Tribe Lycieae (Solanaceae): Paraphyly Of Lycium And Multiple Origins Of Gender Dimorphism
VL  - 92
ID  - 3629
ER  - 


TY  - JOUR
AU  - Levin, Rachel A.
AU  - Keyes, Edmund M.
AU  - Miller, Jill S.
DO  - 10.1086/679492 M4 - Citavi
IS  - 2
KW  - Solanaceae
PY  - 2015
SN  - 1058-5893
SP  - 197-210
ST  - Evolutionary Relationships, Gynodioecy, and Polyploidy in the Galápagos Endemic Lycium minimum (Solanaceae)
T2  - International Journal of Plant Sciences
TI  - Evolutionary Relationships, Gynodioecy, and Polyploidy in the Galápagos Endemic Lycium minimum (Solanaceae)
VL  - 176
ID  - 586
ER  - 


TY  - JOUR
AB  - There has been considerable interest and research into the benefits of multiple low-copy nuclear regions for phylogenetic studies at low taxonomic levels. In this study, the phylogenetic utility of DNA sequence data from two low-copy nuclear genes, nitrate reductase (NIA) and granule-bound starch synthase I (GBSSI), was compared with data from nuclear ribosomal ITS and the cpDNA spacers trnT-trnF and trnD-trnT across 33 closely related taxa in tribe Lycieae (Solanaceae). The NIA data (introns 1-2) had the most parsimony-informative characters, with over twice the number provided by GBSSI, but NIA also had greater homoplasy. Although gene trees were generally concordant across the four regions, there were some notable areas of incongruence, suggesting both incomplete lineage sorting as well as possible reticulate origins.
AD  - Department of Biology, McGuire Life Sciences Building, Amherst College, Amherst, MA 01002, USA. M4 - Citavi
AU  - Levin, Rachel A.
AU  - Blanton, Jessica
AU  - Miller, Jill S.
DO  - 10.1016/j.ympev.2008.12.005
IS  - 3
KW  - Solanaceae
LA  - eng
N1  - Journal Article Research Support, Non-U.S. Gov't Research Support, U.S. Gov't, Non-P.H.S. Journal Article Research Support, Non-U.S. Gov't Research Support, U.S. Gov't, Non-P.H.S. Journal Article Research Support, Non-U.S. Gov't Research Support, U.S. Gov't, Non-P.H.S. Journal Article Research Support, Non-U.S. Gov't Research Support, U.S. Gov't, Non-P.H.S.
PY  - 2009
SN  - 1055-7903
SP  - 608-617
ST  - Phylogenetic utility of nuclear nitrate reductase: a multi-locus comparison of nuclear and chloroplast sequence data for inference of relationships among American Lycieae (Solanaceae)
T2  - Molecular phylogenetics and evolution
TI  - Phylogenetic utility of nuclear nitrate reductase: a multi-locus comparison of nuclear and chloroplast sequence data for inference of relationships among American Lycieae (Solanaceae)
VL  - 50
ID  - 2581
ER  - 


TY  - JOUR
AU  - Levin, Rachel A.
AU  - Bernardello, Gabriel
AU  - Whiting, Carolyn
AU  - Miller, Jill S.
IS  - 3
KW  - Solanaceae
PY  - 2011
SN  - 00400262
SP  - 681-690
ST  - A new generic circumscription in tribe Lycieae (Solanaceae)
T2  - Taxon
TI  - A new generic circumscription in tribe Lycieae (Solanaceae)
VL  - 60
ID  - 1945
ER  - 


TY  - JOUR
AB  - The African Non-Spiny (ANS) clade contains 14 species of mostly large canopy lianas or scandent shrubs confined to Madagascar (10) and continental Africa (4, with with one species reaching the southern Arabian peninsula). Members of the clade were previously classified in sections Afrosolanum Bitter, Benderianum Bitter, Lemurisolanum Bitter, Macronesiotes Bitter and Quadrangulare Bitter, and were throught to be related to a variety of New World groups. The group is an early-branching lineage of non-spiny solanums and characters shared with other vining New World solanums are homoplastic. The 14 species of the group occupy a wide range of habitats, from wet forests in western Africa to savanna and dry forests of southern Madagascar and dune habitats in South Africa. Many members of the group are highly variable morphologically, and habit can vary between shrub and canopy vine in a single locality. We here review the taxonomic history, morphology, potential relationships and ecology of these species; we provide keys for their identification, descriptions, full synonymy (including designations of lectotypes and neotypes) and nomenclatural notes. Illustrations, distribution maps and preliminary conservation assessments are provided for all species.
AD  - Department of Life Sciences, Natural History Museum, Cromwell Road, London SW7 5BD, United Kingdom. Comparative Plant and Fungal Biology, Royal Botanic Gardens, Kew, Richmond, Surrey TW7 3AB, United Kingdom. M4 - Citavi
AU  - Knapp, Sandra
AU  - Vorontsova, Maria S.
DO  - 10.3897/phytokeys.66.8457
IS  - 66
KW  - Solanaceae
LA  - eng
N1  - Journal Article Journal Article Journal Article Journal Article
PY  - 2016
SN  - 1314-2011
SP  - 1-142
ST  - A revision of the "African Non-Spiny" Clade of Solanum L. (Solanum sections Afrosolanum Bitter, Benderianum Bitter, Lemurisolanum Bitter, Lyciosolanum Bitter, Macronesiotes Bitter, and Quadrangulare Bitter: Solanaceae)
T2  - PhytoKeys
TI  - A revision of the "African Non-Spiny" Clade of Solanum L. (Solanum sections Afrosolanum Bitter, Benderianum Bitter, Lemurisolanum Bitter, Lyciosolanum Bitter, Macronesiotes Bitter, and Quadrangulare Bitter: Solanaceae)
UR  - https://www.ncbi.nlm.nih.gov/pmc/articles/PMC4957028
ID  - 264
ER  - 


TY  - JOUR
AB  - Nicotiana (Solanaceae) provides an ideal system for understanding polyploidization, a pervasive and powerful evolutionary force in plants, as this genus contains several groups of allotetraploids that formed at different times from different diploid progenitors. However, the parental lineages of the largest group of allotetraploids, Nicotiana section Suaveolentes, have been problematic to identify. Using data from four regions of three low-copy nuclear genes, nuclear ribosomal DNA, and regions of the plastid genome, we have reconstructed the evolutionary origin of sect. Suaveolentes and identified the most likely diploid progenitors by using a combination of gene trees and network approaches to uncover the most strongly supported evidence of species relationships. Our analyses best support a scenario where a member of the sect. Sylvestres lineage acted as the paternal progenitor and a member of either sect. Petunioides or sect. Noctiflorae that also contained introgressed DNA from the other, or a hypothetical hybrid species between these two sections, was the maternal progenitor. Nicotiana exemplifies many of the factors that can complicate the reconstruction of polyploid evolutionary history and highlights how reticulate evolution at the diploid level can add even greater complexity to allopolyploid genomes.
AD  - School of Biological and Chemical Sciences, Queen Mary University of London, E1 4NS, London, United Kingdom. M4 - Citavi
AU  - Kelly, Laura J.
AU  - Leitch, Andrew R.
AU  - Clarkson, James J.
AU  - Knapp, Sandra
AU  - Chase, Mark W.
DO  - 10.1111/j.1558-5646.2012.01748.x
IS  - 1
KW  - Solanaceae
LA  - eng
N1  - Journal Article Research Support, Non-U.S. Gov't Journal Article Research Support, Non-U.S. Gov't Journal Article Research Support, Non-U.S. Gov't Journal Article Research Support, Non-U.S. Gov't
PY  - 2013
SN  - 0014-3820
SP  - 80-94
ST  - Reconstructing the complex evolutionary origin of wild allopolyploid tobaccos (Nicotiana section suaveolentes)
T2  - Evolution
TI  - Reconstructing the complex evolutionary origin of wild allopolyploid tobaccos (Nicotiana section suaveolentes)
VL  - 67
ID  - 1262
ER  - 


TY  - JOUR
AB  - Reticulate evolution may function both at the species level, through homoploid and polyploid hybridization, and below the species level, through inter and intragenic recombination. These processes represent challenges for the reconstruction of evolutionary relationships between species, because they cannot be represented adequately with bifurcating trees. We use data from low-copy nuclear genes to evaluate long-standing hypotheses of homoploid (interspecific) hybrid speciation in Nicotiana (Solanaceae) and reconstruct a complex series of reticulation events that have been important in the evolutionary history of this genus. Hybrid origins for three diploid species (Nicotiana glauca, N. linearis, and N. spegazzinii) are inferred on the basis of gene tree incongruence, evidence for interallelic recombination between likely parental alleles, and support for incompatible splits in Lento plots. Phylogenetic analysis of recombinant gene sequences illustrates that recombinants may be resolved with one of their progenitor lineages with a high posterior probability under Bayesian inference, and thus there is no indication of the conflict between phylogenetic signals that results from reticulation. Our results illustrate the importance of hybridization in shaping evolution in Nicotiana and also show that intragenic recombination may be relatively common. This finding demonstrates that it is important to investigate the possibility of recombination when aiming to detect hybrids from DNA-sequence data and reconstruct patterns of reticulate evolution between species.
AD  - Jodrell Laboratory, Royal Botanic Gardens Kew, Richmond, Surrey, United Kingdom. M4 - Citavi
AU  - Kelly, Laura J.
AU  - Leitch, Andrew R.
AU  - Clarkson, James J.
AU  - Hunter, Robin B.
AU  - Knapp, Sandra
AU  - Chase, Mark W.
DO  - 10.1093/molbev/msp267
IS  - 4
KW  - Solanaceae
LA  - eng
N1  - Journal Article Research Support, Non-U.S. Gov't Journal Article Research Support, Non-U.S. Gov't Journal Article Research Support, Non-U.S. Gov't Journal Article Research Support, Non-U.S. Gov't
PY  - 2010
SP  - 781-799
ST  - Intragenic recombination events and evidence for hybrid speciation in Nicotiana (Solanaceae)
T2  - Molecular biology and evolution
TI  - Intragenic recombination events and evidence for hybrid speciation in Nicotiana (Solanaceae)
VL  - 27
ID  - 2227
ER  - 


TY  - JOUR
AU  - Jamil, Ishrat
AU  - Qamarunnisa, Syeda
AU  - Azhar, Abid
AU  - Shinwari, Zabta K.
AU  - Ali, Syed Irtifaq
AU  - Qaiser, Muhammad
IS  - 2
KW  - Solanaceae
PY  - 2014
SP  - 585-590
ST  - Subfamilial Relationships within Solanaceae As Inferred From atpβ-rbcL Intergenic Spacer
T2  - Pakistan Journal of Botany
TI  - Subfamilial Relationships within Solanaceae As Inferred From atpβ-rbcL Intergenic Spacer
VL  - 46
ID  - 931
ER  - 


TY  - JOUR
AB  - Advances in sequencing technology have made it possible to produce large multi-locus datasets required for species tree analyses. One challenge with constructing high throughput sequencing datasets, however, is that missing information is propagated at different steps in the sequence preparation process. To date, species tree studies have focused on filtering and removing errors that occur at particular loci. Given the way that high throughput sequencing datasets are constructed, however, large amounts of error or ambiguity may also manifest across individuals. Here we use a novel tree-based multivariate clustering method to identify and remove individuals with low phylogenetic signal in a nuclear sequence capture dataset for the Iochrominae clade (Solanaceae). Our results suggest that the low quality tips are the result of the library preparation process (e.g. unequal pooling) rather than poor capture due to phylogenetic distance from the reference species. After implementing the clustering approach and removing low quality tips, we construct an Iochrominae species tree that resolves a number of unknown relationships. We propose this pipeline as a valuable tool for species tree reconstruction with phylogenomic datasets containing variable levels of missing data.
AD  - Department of Plant Sciences, University of California, Davis, CA, USA. Electronic address:. School of Biological Sciences, University of Nebraska, Lincoln, NE, USA. Department of Ecology and Evolutionary Biology, University of Colorado, Boulder, CO, USA. M4 - Citavi
AU  - Gates, Daniel J.
AU  - Pilson, Diana
AU  - Smith, Stacey D.
DO  - 10.1016/j.ympev.2018.02.002
KW  - Solanaceae
LA  - eng
N1  - Journal Article Research Support, U.S. Gov't, Non-P.H.S. Journal Article Research Support, U.S. Gov't, Non-P.H.S. Journal Article Research Support, U.S. Gov't, Non-P.H.S. Journal Article Research Support, U.S. Gov't, Non-P.H.S.
PY  - 2018
SN  - 1055-7903
SP  - 26-34
ST  - Filtering of target sequence capture individuals facilitates species tree construction in the plant subtribe Iochrominae (Solanaceae)
T2  - Molecular phylogenetics and evolution
TI  - Filtering of target sequence capture individuals facilitates species tree construction in the plant subtribe Iochrominae (Solanaceae)
VL  - 123
ID  - 12
ER  - 


TY  - JOUR
AU  - Garcia, Vicente F.
AU  - Olmstead, Richard G.
IS  - 3
KW  - Solanaceae
PY  - 2003
SN  - 03636445
SP  - 609-615
ST  - Phylogenetics of Tribe Anthocercideae (Solanaceae) Based on ndhF and trnL/F Sequence Data
T2  - Systematic Botany
TI  - Phylogenetics of Tribe Anthocercideae (Solanaceae) Based on ndhF and trnL/F Sequence Data
VL  - 28
ID  - 3982
ER  - 


TY  - JOUR
AB  - Lycium comprises approximately 70 species and is disjunctly distributed in temperate to subtropical regions in South America, North America, southern Africa, Eurasia, and Australia. Among them, only Lycium sandwicense A. Gray sporadically occurs widely on oceanic islands in the Pacific Ocean. To investigate phylogenetic and biogeographic relationships of the genus with emphasis on L. sandwicense, the coding region of matK, the two intergenic spacers trnT (UGU)-trnL (UAA) and trnL (UAA)-trnF (GAA), and the trnL (UAA) intron of chloroplast DNA (cpDNA) were sequenced. A strict consensus tree resulting from the phylogenetic analysis indicates the following: (1) New World species comprise a potentially paraphyletic assemblage; (2) southern African, Australian, and Eurasian species together are monophyletic; (3) southern African species are a paraphyletic assemblage; and (4) L. sandwicense is in a clade with certain New World species. The estimated biogeographic events based on the cpDNA analysis indicate that (1) Lycium originated in the New World, (2) all southern African, Australian, and Eurasian species have a common ancestor from the New World, (3) Australian and Eurasian species originated once from a southern African progenitor, and (4) L. sandwicense differentiated from the New World species.
AD  - Biological Institute, Tohoku University, Sendai, 980-8578, Japan. M4 - Citavi
AU  - Fukuda, Tatsuya
AU  - Yokoyama, J.
AU  - Ohashi, H.
DO  - 10.1006/mpev.2001.0921
IS  - 2
KW  - Solanaceae
LA  - eng
N1  - Journal Article Journal Article Journal Article Journal Article
PY  - 2001
SN  - 1055-7903
SP  - 246-258
ST  - Phylogeny and biogeography of the genus Lycium (Solanaceae): inferences from chloroplast DNA sequences
T2  - Molecular phylogenetics and evolution
TI  - Phylogeny and biogeography of the genus Lycium (Solanaceae): inferences from chloroplast DNA sequences
VL  - 19
ID  - 4225
ER  - 


TY  - JOUR
AU  - Fregonezi, Jefferson Nunes
AU  - Freitas, Loreta Brandão
AU  - Bonatto, Sandro Luis
AU  - Semir, Joáo
AU  - Stehmann, Renato
IS  - 1
KW  - Solanaceae
PY  - 2012
SN  - 00400262
SP  - 120-130
ST  - Infrageneric classification of <i>Calibrachoa</i> (Solanaceae) based on morphological and molecular evidence
T2  - Taxon
TI  - Infrageneric classification of <i>Calibrachoa</i> (Solanaceae) based on morphological and molecular evidence
VL  - 61
ID  - 1535
ER  - 


TY  - JOUR
AU  - Fls, Maria S. Vorontsova
AU  - Stern, Stephen
AU  - Bohs, Lynn
AU  - Knapp, Sandra
KW  - Solanaceae
PY  - 2013
SP  - 176-193
ST  - African spiny Solanum (subgenus Leptostemonum, Solanaceae): a thorny phylogenetic tangle
T2  - Botanical Journal of the Linnean Society
TI  - African spiny Solanum (subgenus Leptostemonum, Solanaceae): a thorny phylogenetic tangle
VL  - 173
ID  - 1219
ER  - 


TY  - JOUR
AB  - Hallucinogenic or toxic species of Brunfelsia (Solanaceae: Petunieae) are important in native cultures throughout South America, and the genus also contains several horticulturally important species. An earlier morphological revision of the c. 50 species recognized three main groups, one consisting of the 23 Antillean species, another of southern South American and Andean species, and a third of species from the Amazon Basin and Guiana Shield. Based on plastid and nuclear DNA sequences from up to 65 accessions representing 80% of the species, we generated a phylogeny and a calibrated chronogram for Brunfelsia to infer clade expansion and shifts in pollinators and fruit types. Brunfelsia flowers offer nectar, and attract lepidoptera, hummingbirds, or bees; the fruits are dry or fleshy. Our results imply that Brunfelsia is 16-21 Myr old and entered the Antilles from South America early during its history, with subsequent expansion along the island arc. The ancestor of the Antillean clade was hawk-moth-pollinated and had fleshy capsules, perhaps facilitating dispersal by birds. The only shift to hummingbird pollination occurred on Cuba, which also harbors the largest single radiation, with 11 species (10 included in our study) that apparently arose over the past 4 Myr. Jamaica, Hispaniola, and Puerto Rico each sustained smaller radiations. The data also reveal at least one new species.
AD  - Systematic Botany and Mycology, University of Munich (LMU), Munich, Germany. M4 - Citavi
AU  - Filipowicz, Natalia
AU  - Renner, Susanne S.
DO  - 10.1016/j.ympev.2012.02.026
IS  - 1
KW  - Solanaceae
LA  - eng
N1  - Journal Article Journal Article Journal Article Journal Article
PY  - 2012
SN  - 1055-7903
SP  - 1-11
ST  - Brunfelsia (Solanaceae): a genus evenly divided between South America and radiations on Cuba and other Antillean islands
T2  - Molecular phylogenetics and evolution
TI  - Brunfelsia (Solanaceae): a genus evenly divided between South America and radiations on Cuba and other Antillean islands
VL  - 64
ID  - 1533
ER  - 


TY  - JOUR
AU  - Fajardo, Diego
AU  - Spooner, David M.
DO  - 10.1600/036364411x553252 M4 - Citavi
IS  - 1
KW  - Solanaceae
PY  - 2011
SN  - 03636445
SP  - 163-170
ST  - Phylogenetic Relationships of Solanum Series Conicibaccata and related Species in Solanum Section Petota inferred from five conserved ortholog Sequences
T2  - Systematic Botany
TI  - Phylogenetic Relationships of Solanum Series Conicibaccata and related Species in Solanum Section Petota inferred from five conserved ortholog Sequences
VL  - 36
ID  - 1881
ER  - 


TY  - JOUR
AU  - Dillon, Michael O.
AU  - Tu, Tieyao
AU  - Soejima, Akiko
AU  - Yi, Tingshuang
AU  - Nie, Zelong
AU  - Tye, Alan
AU  - Wen, Jun
IS  - 4
KW  - Solanaceae
PY  - 2007
SN  - 00400262
SP  - 1000-1011
ST  - Phylogeny of Nolana (Nolaneae, Solanoideae, Solanaceae) as inferred from granule-bound starch synthase I (Gbssi) sequences
T2  - Taxon
TI  - Phylogeny of Nolana (Nolaneae, Solanoideae, Solanaceae) as inferred from granule-bound starch synthase I (Gbssi) sequences
VL  - 56
ID  - 3085
ER  - 


TY  - JOUR
AB  - Deprea is the genus with the second highest species richness in tribe Physalideae (Solanaceae) and comprises 50 species that are mainly distributed in the Andes of South America. The taxonomy of Deprea has been unstable after controversial hypotheses about its position and circumscription. Additionally, biogeographical inferences are only based on observations of the restricted area of distribution of some species and no ancestral area estimation have been performed. Here, we present a phylogenetic analysis and an ancestral area reconstruction of Deprea in order to establish its circumscription, resolve its position within Physalideae, and reconstruct its biogeographical history. Phylogenetic analyses were conducted using Maximum Likelihood and Bayesian approaches. Forty-three Deprea species and 26 related taxa were sampled for three DNA markers (psbA-trnH, ITS, and waxy). A Bayesian binary MCMC model was applied in order to infer ancestral areas. Deprea is resolved as a strongly supported monophyletic group according to its current circumscription and is placed within subtribe Withaninae of Physalideae. The phylogenetic relationships enabled us to solve taxonomic problems including the rejection and acceptance of previous synonyms. The most probable ancestral area for Deprea is the Northern Andes of South America and the Amotape-Huancabamba zone. Our phylogeny provides increased resolution and support for the current position and circumscription of Deprea. Better resolution of interspecific relationships was also obtained, although some affinities remain unclear. The phylogenetic and ancestral area reconstructions provide a framework for addressing taxonomic problems and investigating new evolutionary questions.
AD  - Instituto Multidisciplinario de Biología Vegetal (IMBIV), CONICET and Universidad Nacional de Córdoba, CC 495, CP 5000 Córdoba, Argentina; Facultad de Ciencias Químicas, Universidad Nacional de Córdoba, Haya de la Torre y Medina Allende, Córdoba, Argentina. Electronic address:. Instituto Multidisciplinario de Biología Vegetal (IMBIV), CONICET and Universidad Nacional de Córdoba, CC 495, CP 5000 Córdoba, Argentina; Facultad de Ciencias Químicas, Universidad Nacional de Córdoba, Haya de la Torre y Medina Allende, Córdoba, Argentina. Instituto Multidisciplinario de Biología Vegetal (IMBIV), CONICET and Universidad Nacional de Córdoba, CC 495, CP 5000 Córdoba, Argentina. M4 - Citavi
AU  - Deanna, Rocío
AU  - Barboza, Gloria Estela
AU  - Carrizo García, Carolina
DO  - 10.1016/j.ympev.2017.11.001
KW  - Solanaceae
LA  - eng
N1  - Journal Article Research Support, Non-U.S. Gov't Journal Article Research Support, Non-U.S. Gov't Journal Article Research Support, Non-U.S. Gov't Journal Article Research Support, Non-U.S. Gov't
PY  - 2018
SN  - 1055-7903
SP  - 71-80
ST  - Phylogenetic relationships of Deprea: New insights into the evolutionary history of physaloid groups
T2  - Molecular phylogenetics and evolution
TI  - Phylogenetic relationships of Deprea: New insights into the evolutionary history of physaloid groups
VL  - 119
ID  - 6
ER  - 


TY  - JOUR
AB  - For Nicotiana, with 75 naturally occurring species (40 diploids and 35 allopolyploids), we produced 4656bp of plastid DNA sequence for 87 accessions and various outgroups. The loci sequenced were trnL intron and trnL-F spacer, trnS-G spacer and two genes, ndhF and matK. Parsimony and Bayesian analyses yielded identical relationships for the diploids, and these are consistent with other data, producing the best-supported phylogenetic assessment currently available for the genus. For the allopolyploids, the line of maternal inheritance is traced via the plastid tree. Nicotiana and the Australian endemic tribe Anthocercideae form a sister pair. Symonanthus is sister to the rest of Anthocercideae. Nicotiana sect. Tomentosae is sister to the rest of the genus. The maternal parent of the allopolyploid species of N. sect. Polydicliae were ancestors of the same species, but the allopolyploids were produced at different times, thus making such sections paraphyletic to their extant diploid relatives. Nicotiana is likely to have evolved in southern South America east of the Andes and later dispersed to Africa, Australia, and southwestern North America.
AD  - Jodrell Laboratory, Royal Botanic Gardens, Kew, Richmond, Surrey TW9 3DS, UK. M4 - Citavi
AU  - Clarkson, James J.
AU  - Knapp, Sandra
AU  - Garcia, Vicente F.
AU  - Olmstead, Richard G.
AU  - Leitch, Andrew R.
AU  - Chase, Mark W.
DO  - 10.1016/j.ympev.2004.05.002
IS  - 1
KW  - Solanaceae
LA  - eng
N1  - Comparative Study Journal Article Research Support, Non-U.S. Gov't Comparative Study Journal Article Research Support, Non-U.S. Gov't Comparative Study Journal Article Research Support, Non-U.S. Gov't Comparative Study Journal Article Research Support, Non-U.S. Gov't
PY  - 2004
SN  - 1055-7903
SP  - 75-90
ST  - Phylogenetic relationships in Nicotiana (Solanaceae) inferred from multiple plastid DNA regions
T2  - Molecular phylogenetics and evolution
TI  - Phylogenetic relationships in Nicotiana (Solanaceae) inferred from multiple plastid DNA regions
VL  - 33
ID  - 3770
ER  - 


TY  - JOUR
AB  - Interspecies relationships in Nicotiana (Solanaceae) are complex because 40 species are diploid (two sets of chromosomes) and 35 species are allotetraploid (four sets of chromosomes, two from each progenitor diploid species). We sequenced a fragment (containing four introns) of the nuclear gene 'chloroplast-expressed glutamine synthetase' (ncpGS) in 65 species of Nicotiana. Here we present the first phylogenetic analysis based on a low-copy nuclear gene for this well studied and important genus. Diploid species have a single-copy of ncpGS, and allotetraploids as expected have two homeologous copies, each derived from their progenitor diploid. Results were particularly useful for determining the paternal lineage of previously enigmatic taxa (for which our previous analyses had revealed only the maternal progenitors). In particular, we were able to shed light on the origins of the two oldest and largest allotetraploid sections, N. sects. Suaveolentes and Repandae. All homeologues have an intact reading frame and apparently similar rates of divergence, suggesting both remain functional. Difficulties in fitting certain diploid species into the sectional classification of Nicotiana on morphological grounds, coupled with discordance between the ncpGS data and previous trees (i.e. plastid, nuclear ribosomal DNA), indicate a number of homoploid (diploid) hybrids in the genus. We have evidence for Nicotiana glutinosa and Nicotiana linearis being of hybrid origin and patterns of intra-allelic recombination also indicate the possibility of reticulate origins for other diploid species.
AD  - Jodrell Laboratory, Royal Botanic Gardens, Kew, Richmond, Surrey TW9 3DS, UK. M4 - Citavi
AU  - Clarkson, James J.
AU  - Kelly, Laura J.
AU  - Leitch, Andrew R.
AU  - Knapp, Sandra
AU  - Chase, Mark W.
DO  - 10.1016/j.ympev.2009.10.003
IS  - 1
KW  - Solanaceae
LA  - eng
N1  - Journal Article Research Support, Non-U.S. Gov't Journal Article Research Support, Non-U.S. Gov't Journal Article Research Support, Non-U.S. Gov't Journal Article Research Support, Non-U.S. Gov't
PY  - 2010
SN  - 1055-7903
SP  - 99-112
ST  - Nuclear glutamine synthetase evolution in Nicotiana: phylogenetics and the origins of allotetraploid and homoploid (diploid) hybrids
T2  - Molecular phylogenetics and evolution
TI  - Nuclear glutamine synthetase evolution in Nicotiana: phylogenetics and the origins of allotetraploid and homoploid (diploid) hybrids
VL  - 55
ID  - 2147
ER  - 


TY  - JOUR
AB  - Polymerase chain reaction fragment length polymorphisms and nucleotide sequences for a cytochrome P450 gene encoding flavonoid-3',5'-hydroxylase, Hf1, were studied in 19 natural taxa of Petunia. Natural Petunia taxa were classified into six groups based on major insertion or deletion events that occurred only in intron II of the locus. The maximum parsimony method was used to calculate strict consensus trees based on nucleotide sequences in selected regions of the Hf1 locus. Petunia taxa were divided into two major clades in the phylogenetic trees. Petunia axillaris (including three subspecies), P. exserta, and P. occidentalis formed a clade with 100% bootstrap support. This clade is associated with a consistently inflexed pedicel, self-compatibility in most taxa, and geographical distribution in southern and western portions of the genus range. The other clade, which comprised the remainder of the genus is, however, less supported (up to 71% bootstrap); it is characterized by a deflexed pedicel in the fruiting state (except P. inflata), self-incompatibility, and a northeastern distribution. A nuclear gene, Hf1, seems to be a useful molecular marker for elucidating the phylogeny of the genus Petunia when compared with the nucleotide sequence of trnK intron of chloroplast DNA.
AD  - Graduate School of Science and Technology, Chiba University, Chiba, Japan. M4 - Citavi
AU  - Chen, Sumei
AU  - Matsubara, Kiyoshi
AU  - Omori, Takahiro
AU  - Kokubun, Hisashi
AU  - Kodama, Hiroaki
AU  - Watanabe, Hitoshi
AU  - Hashimoto, Goro
AU  - Marchesi, Eduardo
AU  - Bullrich, Laura
AU  - Ando, Toshio
DO  - 10.1007/s10265-006-0070-z
IS  - 3
KW  - Solanaceae
LA  - eng
N1  - Journal Article Journal Article Journal Article Journal Article
PY  - 2007
SP  - 385-397
ST  - Phylogenetic analysis of the genus Petunia (Solanaceae) based on the sequence of the Hf1 gene
T2  - Journal of Plant Research
TI  - Phylogenetic analysis of the genus Petunia (Solanaceae) based on the sequence of the Hf1 gene
VL  - 120
ID  - 3069
ER  - 


TY  - JOUR
AU  - Chase, Sandra Knapp Mark W.
AU  - Clarkson, James J.
IS  - 1
KW  - Solanaceae
PY  - 2004
SN  - 00400262
SP  - 73-82
ST  - Nomenclatural changes and a new sectional classification in Nicotiana (Solanaceae)
T2  - Taxon
TI  - Nomenclatural changes and a new sectional classification in Nicotiana (Solanaceae)
VL  - 53
ID  - 3767
ER  - 


TY  - JOUR
AB  - Phylogenetic relationships in the genus Nicotiana were investigated using parsimony analyses of the internal transcribed spacer (ITS) regions of nuclear ribosomal DNA (nrDNA). In addition, origins of some amphidiploid taxa in Nicotiana were investigated using the techniques of genomic in situ hybridization (GISH), and the results of both sets of analyses were used to evaluate previous hypotheses about the origins of these taxa. Phylogenetic analyses of the ITS nrDNA data were performed on the entire genus (66 of 77 naturally occurring species, plus three artificial hybrids), comprising both diploid and polyploid taxa, and on the diploid taxa only (35 species) to examine the effects of amphidiploids on estimates of relationships. All taxa, regardless of ploidy, produced clean, single copies of the ITS region, even though some taxa are hybrids. Results are compared with a published plastid (matK) phylogeny using fewer, but many of the same, taxa. The patterns of relationships in Nicotiana, as seen in both analyses, are largely congruent with each other and previous evolutionary ideas based on morphology and cytology, but some important differences are apparent. None of the currently recognized subgenera of Nicotiana is monophyletic and, although most of the currently recognized sections are coherent, others are clearly polyphyletic. Relying solely upon ITS nrDNA analysis to reveal phylogenetic patterns in a complex genus such as Nicotiana is insufficient, and it is clear that conventional analysis of single data sets, such as ITS, is likely to be misleading in at least some respects about evolutionary history. ITS sequences of natural and well-documented amphidiploids are similar or identical to one of their two parents-usually, but not always, the maternal parent-and are not in any sense themselves 'hybrid'. Knowing how ITS evolves in artificial amphidiploids gives insight into what ITS analysis might reveal about naturally occurring amphidiploids of unknown origin, and it is in this perspective that analysis of ITS sequences is highly informative.
AD  - Molecular Systematics Section, Jodrell Laboratory, Royal Botanic Gardens, Kew, Richmond, Surrey TW9 3DS, UK. M4 - Citavi
AU  - Chase, Mark W.
AU  - Knapp, Sandra
AU  - Cox, Antony V.
AU  - Clarkson, James J.
AU  - Butsko, Yelena
AU  - Joseph, Jeffrey A.
AU  - Savolainen, Vincent
AU  - Parokonny, Alex S.
DO  - 10.1093/aob/mcg087
IS  - 1
KW  - Solanaceae
LA  - eng
N1  - Journal Article Research Support, Non-U.S. Gov't Journal Article Research Support, Non-U.S. Gov't Journal Article Research Support, Non-U.S. Gov't Journal Article Research Support, Non-U.S. Gov't
PY  - 2003
SN  - 0305-7364
SP  - 107-127
ST  - Molecular systematics, Gish and the origin of hybrid taxa in Nicotiana (Solanaceae)
T2  - Annals of Botany
TI  - Molecular systematics, Gish and the origin of hybrid taxa in Nicotiana (Solanaceae)
UR  - https://www.ncbi.nlm.nih.gov/pmc/articles/PMC4243627
VL  - 92
ID  - 3965
ER  - 


TY  - JOUR
AU  - Carrizo García, Carolina
AU  - Wahlert, Gregory
AU  - Orozco, Clara Inés
AU  - Barboza, Gloria Estela
AU  - Bohs, Lynn
DO  - 10.11646/phytotaxa.238.1.3 M4 - Citavi
IS  - 1
KW  - Solanaceae
PY  - 2015
SN  - 1179-3163
SP  - 71
ST  - Phylogeny of the Andean genus Deprea (Physalideae, Solanaceae): testing the generic circumscription
T2  - Phytotaxa
TI  - Phylogeny of the Andean genus Deprea (Physalideae, Solanaceae): testing the generic circumscription
VL  - 238
ID  - 474
ER  - 


TY  - JOUR
AU  - Bohs, Lynn
IS  - 4
KW  - Solanaceae
PY  - 2007
SN  - 00400262
SP  - 1012-1026
ST  - Phylogeny of the Cyphomandra clade of the genus Solanum (Solanaceae) based on ITS sequence data
T2  - Taxon
TI  - Phylogeny of the Cyphomandra clade of the genus Solanum (Solanaceae) based on ITS sequence data
VL  - 56
ID  - 3055
ER  - 


TY  - JOUR
AU  - Averett, John E.
IS  - 1
KW  - Solanaceae
PY  - 2009
SP  - 54-61
ST  - Schraderanthus, a new Genus of Solanaceae
T2  - Phytologia
TI  - Schraderanthus, a new Genus of Solanaceae
VL  - 91
ID  - 2435
ER  - 


TY  - JOUR
AU  - Aubriot, Xavier
AU  - Fls, Paramjit Singh
AU  - Knapp, Sandra
KW  - Solanaceae
PY  - 2016
SP  - 199-223
ST  - Tropical Asian species show that the Old World clade of ‘spiny solanums’ (Solanum subgenus Leptostemonum pro parte: Solanaceae) is not monophyletic
T2  - Botanical Journal of the Linnean Society
TI  - Tropical Asian species show that the Old World clade of ‘spiny solanums’ (Solanum subgenus Leptostemonum pro parte: Solanaceae) is not monophyletic
VL  - 181
ID  - 163
ER  - 


TY  - JOUR
AU  - Aoki, S.
AU  - Ito, Motomi
KW  - Solanaceae
PY  - 2000
SN  - 1435-8603
SP  - 316-324
ST  - Molecular Phylogeny of Nicotiana (Solanaceae) Based on the Nucleotide Sequence of the matK Gene
T2  - Plant Biology
TI  - Molecular Phylogeny of Nicotiana (Solanaceae) Based on the Nucleotide Sequence of the matK Gene
VL  - 2
ID  - 4310
ER  - 


TY  - JOUR
AB  - BACKGROUND AND AIMS The phylogenetic relationships of Petunia sensu Jussieu (Petunia sensu Wijsman plus Calibrachoa) are unclear. This study aimed to resolve this uncertainty using molecular evidence. METHODS Phylogenetic trees of 52 taxa of Petunia sensu Jussieu were constructed using restriction fragment length polymorphism (RFLP) of chloroplast DNA digested with 19 restriction enzymes and hybridized with 12 cloned Nicotiana chloroplast DNA fragments as probes. KEY RESULTS In all, 89 phylogenetically informative RFLPs were detected and one 50 % majority consensus tree was obtained, using the maximum parsimony method, and one distance matrix tree, using the neighbour joining method. Petunia sensu Wijsman and Calibrachoa were monophyletic sister clades in both trees. Calibrachoa parviflora and C. pygmaea, previously thought to differ from the other species in terms of their cross-compatibility, seed morphology, and nuclear DNA content, formed a basal clade that was sister to the remainder of Calibrachoa. Several clades found in the phylogenetic trees corresponded to their distribution ranges, suggesting that recent speciation in the genus Petunia sensu Jussieu occurred independently in several different regions. CONCLUSIONS The separation of Petunia sensu Wijsman and Calibrachoa was supported by chloroplast DNA analysis. Two groups in the Calibrachoa were also recognized with a high degree of confidence.
AD  - Faculty of Horticulture, Chiba University, 648 Matsudo, Matsudo, Chiba 271-8510, Japan. M4 - Citavi
AU  - Ando, Toshio
AU  - Kokubun, Hisashi
AU  - Watanabe, Hitoshi
AU  - Tanaka, Norio
AU  - Yukawa, Tomohisa
AU  - Hashimoto, Goro
AU  - Marchesi, Eduardo
AU  - Suárez, Enrique
AU  - Basualdo, Isabel L.
DO  - 10.1093/aob/mci177
IS  - 2
KW  - Solanaceae
LA  - eng
N1  - Comparative Study Journal Article Research Support, Non-U.S. Gov't Comparative Study Journal Article Research Support, Non-U.S. Gov't Comparative Study Journal Article Research Support, Non-U.S. Gov't Comparative Study Journal Article Research Support, Non-U.S. Gov't
PY  - 2005
SN  - 0305-7364
SP  - 289-297
ST  - Phylogenetic analysis of Petunia sensu Jussieu (Solanaceae) using chloroplast DNA Rflp
T2  - Annals of Botany
TI  - Phylogenetic analysis of Petunia sensu Jussieu (Solanaceae) using chloroplast DNA Rflp
UR  - https://www.ncbi.nlm.nih.gov/pmc/articles/PMC4246877
VL  - 96
ID  - 3550
ER  - 


TY  - JOUR
AU  - Ames, Mercedes
AU  - Spooner, David M.
IS  - 4
KW  - Solanaceae
PY  - 2010
SN  - 00400262
SP  - 1091-1101
ST  - Phylogeny of Solanum series Piurana and related species in Solanum section Petota based on five conserved ortholog sequences
T2  - Taxon
TI  - Phylogeny of Solanum series Piurana and related species in Solanum section Petota based on five conserved ortholog sequences
VL  - 59
ID  - 2108
ER  - 


TY  - JOUR
AB  - The Smilax hispida group (Smilacaceae) exhibits a discontinuous distribution in eastern Asia, eastern and western United States, and Mexico. A broad scale phylogeographic analysis was conducted for this group to evaluate the hypotheses of accelerated allopatric divergence in eastern Asia and a northern origin of the temperate elements in Mexico. Phylogeny was inferred using seven plastid and nuclear DNA sequences. Species delineation was assessed using genealogical sorting indices (GSI). Lineage divergence time, haplotype diversification rates, and ancestral distributions were estimated using Bayesian methods. Phylogeographic patterns in eastern Asia and North America were compared by analyzing 539 individuals from 64 populations to assess allopatric diversification. Results strongly supported delineation of six allopatric species, the origin of this group from a Mexican ancestor around 11.42mya, and Mexican origins of the temperate species in Mexico. Significant geographic structure of haplotypes was found in eastern Asia, and greater haplotype diversification rate was observed for the North American lineage. Our data support allopatric speciation in eastern Asia but do not find evidence of an elevated diversification rate. Greater species diversity of the study system in eastern Asia may be due to a longer evolutionary history. Our results do not support northern origins of the Mexican temperate species.
AD  - The Key Laboratory of Conservation Biology for Endangered Wildlife of the Ministry of Education, College of Life Sciences, Zhejiang University, Hangzhou 310058, PR China. M4 - Citavi
AU  - Zhao, Yunpeng
AU  - Qi, Zhechen
AU  - Ma, Weiwei
AU  - Dai, Qiongyan
AU  - Li, Pan
AU  - Cameron, Kenneth M.
AU  - Lee, Joongku
AU  - Xiang, Qiu-Yun Jenny
AU  - Fu, Cheng-Xin
DO  - 10.1016/j.ympev.2013.03.025
IS  - 2
KW  - Smilacaceae
LA  - eng
N1  - Comparative Study Journal Article Research Support, Non-U.S. Gov't Comparative Study Journal Article Research Support, Non-U.S. Gov't Comparative Study Journal Article Research Support, Non-U.S. Gov't Comparative Study Journal Article Research Support, Non-U.S. Gov't
PY  - 2013
SN  - 1055-7903
SP  - 300-311
ST  - Comparative phylogeography of the Smilax hispida group (Smilacaceae) in eastern Asia and North America - implications for allopatric speciation, causes of diversity disparity, and origins of temperate elements in Mexico
T2  - Molecular phylogenetics and evolution
TI  - Comparative phylogeography of the Smilax hispida group (Smilacaceae) in eastern Asia and North America - implications for allopatric speciation, causes of diversity disparity, and origins of temperate elements in Mexico
VL  - 68
ID  - 1449
ER  - 


TY  - JOUR
AU  - Qi, Zhechen
AU  - Li, P. A. N.
AU  - Fu, Cheng-Xin
DO  - 10.11646/phytotaxa.117.2.4 M4 - Citavi
IS  - 2
KW  - Smilacaceae
PY  - 2013
SN  - 1179-3163
SP  - 58
ST  - New combinations and a new name in Smilax for species of Heterosmilax in Eastern and Southeast Asian Smilacaceae (Liliales)
T2  - Phytotaxa
TI  - New combinations and a new name in Smilax for species of Heterosmilax in Eastern and Southeast Asian Smilacaceae (Liliales)
VL  - 117
ID  - 1358
ER  - 


TY  - JOUR
AD  - The Key Laboratory of Conservation Biology for Endangered Wildlife of the Ministry of Education; College of Life Sciences; Zhejiang University; Hangzhou 310058 China Laboratory of Systematic and Evolutionary Botany; Institute of Plant Sciences and Conservation Center for Gene Resources of Endangered Wildlife; Zhejiang University; Hangzhou 310058 China Wisconsin State Herbarium; Department of Botany; University of Wisconsin; Madison WI 53706 USA Research and Development Centre; Firmenich Aromatics (China); Shanghai 201108 China College of Life Sciences and Technology; Tongji University; Shanghai 200092 China M4 - Citavi
AU  - Qi, Zhechen
AU  - Cameron, Kenneth M.
AU  - Li, Pan
AU  - Zhao, Yunpeng
AU  - Chen, Shi-Chao
AU  - Chen, Guangcun
AU  - Fu, Cheng-Xin
DO  - 10.1111/boj.12096 Y3 - 16.02.2019
IS  - 4
KW  - Smilacaceae
PY  - 2013
SP  - 535-548
ST  - Phylogenetics, character evolution, and distribution patterns of the greenbriers, Smilacaceae (Liliales), a near-cosmopolitan family of monocots
T2  - Botanical Journal of the Linnean Society
TI  - Phylogenetics, character evolution, and distribution patterns of the greenbriers, Smilacaceae (Liliales), a near-cosmopolitan family of monocots
VL  - 173
ID  - 1357
ER  - 


TY  - JOUR
AB  - PREMISE OF THE STUDY The genetic structure of North American herbaceous Smilax, a clade of closely related understory herbs that has a wide and disjunct distribution, was investigated to test the hypothesis of whether a northern refugium in the upper Midwest may have existed for these plants during the last glaciation. METHODS We analyzed 33 populations sampled from California and throughout the eastern United States using AFLP and chloroplast DNA sequence data. KEY RESULTS Individuals of S. jamesii from northern California formed a clade sister to the eastern North American species, and they likely diverged from each other during the Pleistocene. Among the eastern species, two lineages were found on opposite sides of the Appalachian Mountains. The populations near the "Driftless Area" contained most of the chlorotypes found in the Midwest, including a unique one endemic to this area, and the AFLP data indicated that one population from this area has the highest frequency-down-weighted-marker value. CONCLUSIONS This study, and others that have focused on mammals, amphibians, and woody plants, provides further evidence for the debate over whether the upper Midwest's "Driftless Area" may have hosted a biologically diverse northern glacial refugium. Herbaceous species of Smilax from eastern North American exhibit an Appalachian discontinuity. They appear to have persisted in multiple refugia both east and west of the Appalachians, with several populations persisting in situ through the Last Glacial Maximum (LGM). We suggest that they experienced localized expansion after the LGM, possibly followed by subsequent contact between the Midwest and the East Coastal lineages.
AD  - The Key Laboratory of Conservation Biology for Endangered Wildlife of the Ministry of Education, College of Life Sciences, Zhejiang University, Hangzhou 310058, China. M4 - Citavi
AU  - Li, Pan
AU  - Li, Mimi
AU  - Shi, Yin
AU  - Zhao, Yunpeng
AU  - Wan, Ying
AU  - Fu, Cheng-Xin
AU  - Cameron, Kenneth M.
DO  - 10.3732/ajb.1200250
IS  - 4
KW  - Smilacaceae
LA  - eng
N1  - Journal Article Research Support, Non-U.S. Gov't Journal Article Research Support, Non-U.S. Gov't Journal Article Research Support, Non-U.S. Gov't Journal Article Research Support, Non-U.S. Gov't
PY  - 2013
SN  - 0002-9122
SP  - 801-814
ST  - Phylogeography of North American herbaceous Smilax (Smilacaceae): Combined AFLP and cpDNA data support a northern refugium in the Driftless Area
T2  - American journal of botany
TI  - Phylogeography of North American herbaceous Smilax (Smilacaceae): Combined AFLP and cpDNA data support a northern refugium in the Driftless Area
VL  - 100
ID  - 1286
ER  - 


TY  - JOUR
AB  - International Journal of Plant Sciences 2005.166:301-309
AU  - Fu, Cheng-Xin
AU  - Kong, Hanghui
AU  - Qiu, Yingxiong
AU  - Cameron, Kenneth M.
IS  - 2
KW  - Smilacaceae
PY  - 2005
SN  - 1058-5893
SP  - 301-309
ST  - Molecular Phylogeny of the East Asian–North American Disjunct Smilax Sect. Nemexia (Smilacaceae)
T2  - International Journal of Plant Sciences
TI  - Molecular Phylogeny of the East Asian–North American Disjunct Smilax Sect. Nemexia (Smilacaceae)
VL  - 166
ID  - 3587
ER  - 


TY  - JOUR
AU  - Chen, Shi-Chao
DO  - 10.1360/aps050065 M4 - Citavi
IS  - 2
KW  - Smilacaceae
PY  - 2006
SN  - 0529-1526
SP  - 113-125
ST  - A phylogenetic analysis of the Smilacaceae based on morphological data
T2  - Acta Phytotaxonomica Sinica
TI  - A phylogenetic analysis of the Smilacaceae based on morphological data
VL  - 44
ID  - 3351
ER  - 


TY  - JOUR
AB  - Tethyan plant disjunctions, including Mediterranean-African-Asian disjunctions, are thought to be vicariant, but their temporal origin and underlying causes remain largely unknown. To address this issue, we reconstructed the evolutionary history of Smilax aspera, a hypothesized component of the European Tertiary laurel forest flora. Thirty-eight populations and herbarium specimens representing 57 locations across the species range were sequenced at seven plastid regions and the nuclear ribosomal internal transcribed spacer region. Time-calibrated phylogenetic and phylogeographic inferences were used to trace ancestral areas and biogeographical events. The deep intraspecific split between Mediterranean and African-Asian lineages is attributable to range fragmentation of a southern Tethyan ancestor, as colder and more arid climates developed shortly after the mid-Miocene. In the Mediterranean, climate-induced vicariance has shaped regional population structure since the Late Miocene/Early Pliocene. At around the same time, East African and South Asian lineages split by vicariance, with one shared haplotype reflecting long-distance dispersal. Our results support the idea that geographic range formation and divergence of Tertiary relict species are more or less gradual (mostly vicariant) processes over long time spans, rather than point events in history. They also highlight the importance of the Mediterranean Basin as a centre of intraspecific divergence for Tertiary relict plants.
AU  - Chen, Chen
AU  - Qi, Zhe-Chen
AU  - Xu, Xi-Hui
AU  - Comes, Hans Peter
AU  - Koch, Marcus A.
AU  - Jin, Xin-Jie
AU  - Fu, Cheng-Xin
AU  - Qiu, Ying-Xiong
DO  - 10.1111/nph.12910
IS  - 1
KW  - Smilacaceae
LA  - eng
PY  - 2014
SP  - 243-255
ST  - Understanding the formation of Mediterranean-African-Asian disjunctions: evidence for Miocene climate-driven vicariance and recent long-distance dispersal in the Tertiary relict Smilax aspera (Smilacaceae)
T2  - The New phytologist
TI  - Understanding the formation of Mediterranean-African-Asian disjunctions: evidence for Miocene climate-driven vicariance and recent long-distance dispersal in the Tertiary relict Smilax aspera (Smilacaceae)
VL  - 204
ID  - 859
ER  - 


TY  - JOUR
AU  - Renner, Susanne S.
AU  - Hausner, Gerlinde
KW  - Siparunaceae
PY  - 2005
SN  - 1055-3177
SP  - 202-206
ST  - New Species of Siparuna (Siparunaceae) Iv. A new Subcanopy Tree from White-Sand Areas in Brazil and Venezuela
T2  - Novon: A Journal for Botanical Nomenclature
TI  - New Species of Siparuna (Siparunaceae) Iv. A new Subcanopy Tree from White-Sand Areas in Brazil and Venezuela
VL  - 15
ID  - 3682
ER  - 


TY  - JOUR
AU  - Renner, Susanne S.
AU  - Hausner, Gerlinde
KW  - Siparunaceae
PY  - 2000
SN  - 1055-3177
SP  - 134-143
ST  - New Species of Siparuna (Siparunaceae) III. Three New Species and One Newly Ranked Entity from Colombia, Ecuador, and Peru
T2  - Novon: A Journal for Botanical Nomenclature
TI  - New Species of Siparuna (Siparunaceae) III. Three New Species and One Newly Ranked Entity from Colombia, Ecuador, and Peru
VL  - 10
ID  - 4367
ER  - 


TY  - JOUR
AB  - Generic circumscriptions in the mostly pantropical family Simaroubaceae are somewhat controversial. Simaba is the largest genus, currently defined as exclusively neotropical, with around 25 species of trees and shrubs, but both its limits and infrageneric classification have been a matter of discussion and divergence. Traditionally, species of the genus have been treated in three sections: Simaba sect. Tenuiflorae, S. sect. Floribundae and S. sect. Grandiflorae, but a phylogenetic analysis suggested that the latter two may not be monophyletic. To test the monophyly of Simaba and its infrageneric classification, we used a molecular approach based on DNA sequence data from two nuclear ribosomal spacer regions (ITS and ETS) and three plastid regions (rps16 intron, and intergenic spacers psbA-trnH and trnL-trnF), including a comprehensive sampling of species from Simaba and closely related genera. We also performed ancestral character reconstructions to identify morphological characters that could serve as synapomorphies for major clades and to explore patterns of homoplasy in the morphological dataset. Our results show Simaba as traditionally circumscribed is not monophyletic, with taxa segregated into two strongly supported but distinct clades, one of which is more closely related to Simarouba. The three main clades that emerged in the phylogeny include a mostly Amazonian Simaba clade (which includes the type species of Simaba and the remaining species of S. sect. Tenuiflorae, here proposed to be recognized as Simaba sensu stricto), a mostly extra-Amazonian Simaba clade (a distinct lineage that will be recognized as Homalolepis, a genus currently treated in synonymy and equivalent to Simaba sections Grandiflorae and Floribundae), and the Simarouba clade (including all of its current species). These three clades are characterized by a combination of morphological characters, described in detail herein, some of which are novel features for Simaba not previously reported in the literature. Mapping character-states on the phylogenetic tree provides tests for evolutionary hypotheses. For example, our reconstruction of habit and geographic distribution suggests that the diversification of several shrubby species within the extra-Amazonian lineage in the South American cerrados probably occurred from ancestors inhabiting tropical forests, involving transitions in morphological and ecological traits.
AD  - Departamento de Botânica, Universidade de São Paulo, Herbário SPF, Rua do Matão 277, São Paulo 05508-900, Brazil. Electronic address:. Institute of Systematic Botany, New York Botanical Garden, Bronx, NY 10458-5126, USA. Cullman Program for Molecular Systematics, New York Botanical Garden, Bronx, NY 10458-5126, USA. Departamento de Botânica, Universidade de São Paulo, Herbário SPF, Rua do Matão 277, São Paulo 05508-900, Brazil. M4 - Citavi
AU  - Devecchi, Marcelo Fernando
AU  - Thomas, William Wayt
AU  - Plunkett, Gregory M.
AU  - Pirani, José Rubens
DO  - 10.1016/j.ympev.2017.11.024
KW  - Simaroubaceae
LA  - eng
N1  - Journal Article Research Support, Non-U.S. Gov't Journal Article Research Support, Non-U.S. Gov't Journal Article Research Support, Non-U.S. Gov't Journal Article Research Support, Non-U.S. Gov't
PY  - 2018
SN  - 1055-7903
SP  - 63-82
ST  - Testing the monophyly of Simaba (Simaroubaceae): Evidence from five molecular regions and morphology
T2  - Molecular phylogenetics and evolution
TI  - Testing the monophyly of Simaba (Simaroubaceae): Evidence from five molecular regions and morphology
VL  - 120
ID  - 9
ER  - 


TY  - JOUR
AB  - Detailed biogeographic studies of pantropical clades are still relatively few, and those conducted to date typically use parsimony or event-based methods to reconstruct ancestral areas. In this study, a recently developed likelihood method for reconstructing ancestral areas (the dispersal-extinction cladogenesis [DEC] model) is applied to the angiosperm family Simaroubaceae, a geographically widespread and ecologically diverse clade of pantropical and temperate trees and shrubs. To estimate divergence dates in the family, Bayesian uncorrelated rates analyses and robust fossil calibrations are applied to the well-sampled and strongly supported phylogeny. For biogeographic analyses, the effects of parameter configurations in the DEC model are assessed for different possible ancestral ranges, and the likelihood method is compared with dispersal-vicariance analysis (DIVA). Regardless of the parameters used, likelihood analyses show a common pattern of multiple recent range shifts that overshadow reconstruction of events deeper in the family's history. DIVA produced results similar to the DEC model when ancestral ranges were restricted to two areas, but some improbable ancestral ranges were also observed. Simaroubaceae exhibit an early history of range expansion between major continental areas in the Northern Hemisphere, but reconstruction of ancestral areas for lineages diverging in the early Tertiary are sensitive to the parameters of the model used. A North American origin is suggested for the family, with migration via Beringia by ancestral taxa. In contrast to traditional views, long-distance dispersal events are common, particularly in the Late Oligocene and later. Notable dispersals are inferred to have occurred across the Atlantic Ocean in both directions, as well as between Africa and Asia, and around the Indian Ocean basin and Pacific islands.
AD  - Department of Botany, University of Florida, Gainesville, FL 32611-7800, USA. M4 - Citavi
AU  - Clayton, Joshua W.
AU  - Soltis, Pamela S.
AU  - Soltis, Douglas E.
DO  - 10.1093/sysbio/syp041
IS  - 4
KW  - Simaroubaceae
LA  - eng
N1  - Journal Article Research Support, Non-U.S. Gov't Research Support, U.S. Gov't, Non-P.H.S. Journal Article Research Support, Non-U.S. Gov't Research Support, U.S. Gov't, Non-P.H.S. Journal Article Research Support, Non-U.S. Gov't Research Support, U.S. Gov't, Non-P.H.S. Journal Article Research Support, Non-U.S. Gov't Research Support, U.S. Gov't, Non-P.H.S.
PY  - 2009
SP  - 395-410
ST  - Recent long-distance dispersal overshadows ancient biogeographical patterns in a pantropical angiosperm family (Simaroubaceae, Sapindales)
T2  - Systematic Biology
TI  - Recent long-distance dispersal overshadows ancient biogeographical patterns in a pantropical angiosperm family (Simaroubaceae, Sapindales)
VL  - 58
ID  - 2477
ER  - 


TY  - JOUR
AU  - Clayton, Joshua W.
AU  - Fernando, Edwino S.
AU  - Soltis, Pamela S.
AU  - Soltis, Douglas E.
DO  - 10.1086/521796 M4 - Citavi
IS  - 9
KW  - Simaroubaceae
PY  - 2007
SN  - 1058-5893
SP  - 1325-1339
ST  - Molecular Phylogeny of the Tree‐of‐Heaven Family (Simaroubaceae) Based on Chloroplast and Nuclear Markers
T2  - International Journal of Plant Sciences
TI  - Molecular Phylogeny of the Tree‐of‐Heaven Family (Simaroubaceae) Based on Chloroplast and Nuclear Markers
VL  - 168
ID  - 3075
ER  - 


TY  - JOUR
AU  - Hernández-Hernández, Tania
AU  - Colorado, Wendy B.
AU  - Sosa, Victoria
DO  - 10.1007/s13127-013-0136-4 M4 - Citavi
IS  - 4
KW  - Setchellanthaceae
N1  - PII: 136 PII: 136 PII: 136 PII: 136
PY  - 2013
SN  - 1439-6092
SP  - 485-496
ST  - Molecular evidence for the origin and evolutionary history of the rare American desert monotypic family Setchellanthaceae
T2  - Organisms Diversity & Evolution
TI  - Molecular evidence for the origin and evolutionary history of the rare American desert monotypic family Setchellanthaceae
VL  - 13
ID  - 1244
ER  - 


TY  - JOUR
AD  - Key Laboratory of Mountain Ecological Restoration and Bioresource Utilization, Chengdu Institute of Biology, Chinese Academy of Sciences, P.O. Box 416, Chengdu, Sichuan 610041, China, School of Life Sciences, Sichuan University, Chengdu, Sichuan 610064, China Missouri Botanical Garden, P.O. Box 299, St. Louis, Missouri 63166-0299, U.S.A.;, M4 - Citavi
AU  - Zhou, Xin-Mao
AU  - Zhang, Li-Bing
DO  - 10.12705/646.2
IS  - 6
KW  - Selaginellaceae
PY  - 2015
SN  - 00400262
SP  - 1117-1140
ST  - A classification of Selaginella (Selaginellaceae) based on molecular (chloroplast and nuclear), macromorphological, and spore features
T2  - Taxon
TI  - A classification of Selaginella (Selaginellaceae) based on molecular (chloroplast and nuclear), macromorphological, and spore features
VL  - 64
ID  - 805
ER  - 


TY  - JOUR
AD  - Chengdu Institute of Biology; Chinese Academy of Sciences; PO Box 416 Chengdu Sichuan 610041 China School of Life Sciences; Sichuan University; Chengdu Sichuan 610064 China Department of Zoology; University of British Columbia; #4200-6270 University Blvd Vancouver BC V6T 1Z4 Canada University Herbarium and Department of Integrative Biology; University of California; Berkeley CA 94720-2465 USA School of Life Science; Yunnan University; Kunming Yunnan 650091 China School of Life Sciences; University of KwaZulu-Natal; Private Bag X01 Scottsville Pietermaritzburg 3209 South Africa Department of Biology; Chongqing Normal University; Shapingba, Chongqing 400047 China Department of Botany, Vietnam National Museum of Nature; Vietnam Academy of Science and Technology; 18th Hoang Quoc Viet Road Cau Giay, Ha Noi Viet Nam Correspondent of the Muséum national d'Histoire naturelle (MNHN, Paris, France); Steigestrasse 78 69412 Eberbach Germany National Tropical Botanical Garden; 3530 Papalina Road Kalaheo HI 96741 USA Missouri Botanical Garden; PO Box 299 St Louis MO 63166-0299 USA M4 - Citavi
AU  - Zhou, Xin-Mao
AU  - Rothfels, Carl J.
AU  - Zhang, Liang
AU  - He, Zhao-Rong
AU  - Le Péchon, Timothée
AU  - He, Hai
AU  - Lu, Ngan Thi
AU  - Knapp, Ralf
AU  - Lorence, David H.
AU  - He, Xing-Jin
AU  - Gao, Xin-Fen
AU  - Zhang, Li-Bing
DO  - 10.1111/cla.12136
IS  - 4
KW  - Selaginellaceae
PY  - 2016
SN  - 07483007
SP  - 360-389
ST  - A large-scale phylogeny of the lycophyte genus Selaginella (Selaginellaceae: Lycopodiopsida) based on plastid and nuclear loci
T2  - Cladistics
TI  - A large-scale phylogeny of the lycophyte genus Selaginella (Selaginellaceae: Lycopodiopsida) based on plastid and nuclear loci
VL  - 32
ID  - 423
ER  - 


TY  - JOUR
AU  - Zhou, Xin-Mao
AU  - Jiang, Li-Jü
AU  - Zhang, Liang
AU  - Gao, Xin-Fen
AU  - He, Zhao-Rong
AU  - Zhang, Li-Bing
DO  - 10.11646/phytotaxa.237.1.1 M4 - Citavi
IS  - 1
KW  - Selaginellaceae
PY  - 2015
SN  - 1179-3163
SP  - 1
ST  - Spore morphology of Selaginella (Selaginellaceae) from China and its systematic significance
T2  - Phytotaxa
TI  - Spore morphology of Selaginella (Selaginellaceae) from China and its systematic significance
VL  - 237
ID  - 804
ER  - 


TY  - JOUR
AB  - PREMISE OF THE STUDY The cosmopolitan lycophyte family Selaginellaceae, dating back to the Late Devonian-Early Carboniferous, is notorious for its many species with a seemingly undifferentiated gross morphology. This morphological stasis has for a long time hampered our understanding of the evolutionary history of the single genus Selaginella. Here we present a large-scale phylogenetic analysis of Selaginella, and based on the resulting phylogeny, we discuss morphological evolution in the group. METHODS We sampled about one-third of the approximately 750 recognized Selaginella species. Evolutionary relationships were inferred from both chloroplast (rbcL) and single-copy nuclear gene data (pgiC and SQD1) using a Bayesian inference approach. The morphology of the group was studied and important features mapped onto the phylogeny. KEY RESULTS We present an overall well-supported phylogeny of Selaginella, and the phylogenetic positions of some previously problematic taxa (i.e., S. sinensis and allies) are now resolved with strong support. We show that even though the evolution of most morphological characters involves reversals and/or parallelisms, several characters are phylogenetically informative. Seven major clades are identified, which each can be uniquely diagnosed by a suite of morphological features. There is value in morphology after all! CONCLUSIONS Our hypothesis of the evolutionary relationships of Selaginella is well founded based on DNA sequence data, as well as morphology, and is in line with previous findings. It will serve as a firm basis for further studies on Selaginella with respect to, e.g., the poorly known alpha taxonomy, as well as evolutionary questions such as historical biogeographic reconstructions.
AD  - Systematic Biology, Department of Organismal Biology, Evolutionary Biology Centre, Uppsala University, Norbyvägen 18D, SE-752 36 Uppsala, Sweden. Systematic Biology, Department of Organismal Biology, Evolutionary Biology Centre, Uppsala University, Norbyvägen 18D, SE-752 36 Uppsala, Sweden. M4 - Citavi
AU  - Weststrand, Stina
AU  - Korall, Petra
DO  - 10.3732/ajb.1600156
IS  - 12
KW  - Selaginellaceae
LA  - eng
N1  - Journal Article Journal Article Journal Article Journal Article
PY  - 2016
SN  - 0002-9122
SP  - 2136-2159
ST  - Phylogeny of Selaginellaceae: There is value in morphology after all!
T2  - American journal of botany
TI  - Phylogeny of Selaginellaceae: There is value in morphology after all!
VL  - 103
ID  - 397
ER  - 


TY  - JOUR
AB  - PREMISE OF THE STUDY The lycophyte family Selaginellaceae includes approximately 750 herbaceous species worldwide, with the main species richness in the tropics and subtropics. We recently presented a phylogenetic analysis of Selaginellaceae based on DNA sequence data and, with the phylogeny as a framework, the study discussed the character evolution of the group focusing on gross morphology. Here we translate these findings into a new classification. METHODS To present a robust and useful classification, we identified well-supported monophyletic groups from our previous phylogenetic analysis of 223 species, which together represent the diversity of the family with respect to morphology, taxonomy, and geographical distribution. Care was taken to choose groups with supporting morphology. KEY RESULTS In this classification, we recognize a single genus Selaginella and seven subgenera: Selaginella, Rupestrae, Lepidophyllae, Gymnogynum, Exaltatae, Ericetorum, and Stachygynandrum. The subgenera are all well supported based on analysis of DNA sequence data and morphology. A key to the subgenera is presented. CONCLUSIONS Our new classification is based on a well-founded hypothesis of the evolutionary relationships of Selaginella, and each subgenus can be identified by a suite of morphological features, most of them possible to study in the field. Our intention is that the classification will be useful not only to experts in the field, but also to a broader audience.
AD  - Systematic Biology, Department of Organismal Biology, Evolutionary Biology Centre, Uppsala University, Norbyvägen 18D, SE-752 36 Uppsala, Sweden. Systematic Biology, Department of Organismal Biology, Evolutionary Biology Centre, Uppsala University, Norbyvägen 18D, SE-752 36 Uppsala, Sweden. M4 - Citavi
AU  - Weststrand, Stina
AU  - Korall, Petra
DO  - 10.3732/ajb.1600288
IS  - 12
KW  - Selaginellaceae
LA  - eng
N1  - Journal Article Journal Article Journal Article Journal Article
PY  - 2016
SN  - 0002-9122
SP  - 2160-2169
ST  - A subgeneric classification of Selaginella (Selaginellaceae)
T2  - American journal of botany
TI  - A subgeneric classification of Selaginella (Selaginellaceae)
VL  - 103
ID  - 396
ER  - 


TY  - JOUR
AB  - Molecular phylogenetic research on Selaginellaceae has focused on the plastid gene rbcL, which in this family has unusually high substitution rates. Here we develop a molecular data set from the nuclear 26S ribosomal DNA gene with the aim of evaluating and extending the results of previous phylogenetic research. The 26S rDNA and the rbcL regions were sequenced for a sample of 23 species, which represent the main elements of species diversity in the family. The data were analysed independently and in combination using both maximum parsimony and Bayesian inference. Although several between genome differences were found, the general pattern of relationships uncovered by all analyses was very similar. Results corroborate the previous study supporting new groupings not previously recognised on morphological grounds. Substitution rates in the 26S rDNA were also found to be high (26% informative) for the region analysed, but lower than for rbcL (37% informative). These data indicate that high substitution rates might be widespread in all three genomes (i.e., plastid, mitochondrion, and nucleus).
AD  - Department of Botany, Stockholm University, SE-106 91 Stockholm, Sweden. M4 - Citavi
AU  - Korall, Petra
AU  - Kenrick, Paul
DO  - 10.1016/j.ympev.2003.10.014
IS  - 3
KW  - Selaginellaceae
LA  - eng
N1  - Journal Article Research Support, Non-U.S. Gov't Journal Article Research Support, Non-U.S. Gov't Journal Article Research Support, Non-U.S. Gov't Journal Article Research Support, Non-U.S. Gov't
PY  - 2004
SN  - 1055-7903
SP  - 852-864
ST  - The phylogenetic history of Selaginellaceae based on DNA sequences from the plastid and nucleus: extreme substitution rates and rate heterogeneity
T2  - Molecular phylogenetics and evolution
TI  - The phylogenetic history of Selaginellaceae based on DNA sequences from the plastid and nucleus: extreme substitution rates and rate heterogeneity
VL  - 31
ID  - 3819
ER  - 


TY  - JOUR
AB  - PREMISE OF THE STUDY Several members of Selaginella are renowned for their ability to survive extreme drought and "resurrect" when conditions improve. Many of these belong to subgenus Tetragonostachys, a group of ∼45 species primarily found in North and Central America, with substantial diversity in the Sonoran and Chihuahuan Deserts. We evaluated the monophyly and the age of subgenus Tetragonostachys and assess how drought tolerance contributed to the evolution of this clade. METHODS Our study included most Tetragonostachys species, using plastid and nuclear sequences, fossil and herbarium records, and climate variables to describe the species diversity, phylogenetic relationships, divergence times, and climatic niche evolution in the subgenus. KEY RESULTS We found that subgenus Tetragonostachys forms a monophyletic group sister to Selaginella lepidophylla and may have diverged from other Selaginella because of a Gondwanan-Laurasian vicariance event ca. 240 mya. The North American radiation of Tetragonostachys appears to be much more recent and to have occurred during the Early Cretaceous-late Paleocene interval. We identified two significant and nested ecological niche shifts during the evolution of Tetragonostachys associated with extreme drought tolerance and a more recent shift to cold climates. Our analyses suggest that drought tolerance evolved in the warm deserts of southwest North America and may have been advantageous for colonization of cold and dry boreal climates. CONCLUSIONS Our investigation provides a foundation for future research addressing the genomics of ecological niche evolution and the potential role of reticulate evolution in Selaginella subgenus Tetragonostachys.
AU  - Arrigo, Nils
AU  - Therrien, James
AU  - Anderson, Cajsa Lisa
AU  - Windham, Michael D.
AU  - Haufler, Christopher H.
AU  - Barker, Michael S.
DO  - 10.3732/ajb.1200426
IS  - 8
KW  - Selaginellaceae
LA  - eng
PY  - 2013
SN  - 0002-9122
SP  - 1672-1682
ST  - A total evidence approach to understanding phylogenetic relationships and ecological diversity in Selaginella subg. Tetragonostachys
T2  - American journal of botany
TI  - A total evidence approach to understanding phylogenetic relationships and ecological diversity in Selaginella subg. Tetragonostachys
VL  - 100
ID  - 1163
ER  - 


TY  - JOUR
AU  - Verboom, George Anthony
AU  - Herron, Margaret L.
AU  - Moncrieff, Glenn R.
AU  - Slingsby, Jasper A.
KW  - Scrophulariaceae
PY  - 2016
SP  - 115-139
ST  - Maintenance of species integrity in the context of a recent radiation: the case of Jamesbrittenia (Scrophulariaceae: Limoselleae) in southern Africa
T2  - Botanical Journal of the Linnean Society
TI  - Maintenance of species integrity in the context of a recent radiation: the case of Jamesbrittenia (Scrophulariaceae: Limoselleae) in southern Africa
VL  - 182
ID  - 389
ER  - 


TY  - JOUR
AU  - Tank, David C.
AU  - Beardsley, Paul M.
AU  - Kelchner, Scot A.
AU  - Olmstead, Richard G.
DO  - 10.1071/SB05009 M4 - Citavi
IS  - 4
KW  - Scrophulariaceae
N1  - PII: SB05009 PII: SB05009 PII: SB05009 PII: SB05009
PY  - 2006
SN  - 1030-1887
SP  - 289-307
ST  - Review of the systematics of Scrophulariaceae s.l. and their current disposition
T2  - Australian Systematic Botany
TI  - Review of the systematics of Scrophulariaceae s.l. and their current disposition
VL  - 19
ID  - 3504
ER  - 


TY  - JOUR
AB  - The flora of the Mediterranean region and Macaronesia is characterized by high levels of species diversity and endemism. We examined phylogenetic relationships of Scrophularia within one of its secondary centers of diversity located in the Iberian Peninsula and adjacent Macaronesia. In total, 65 ingroup accessions from 45 species, representing an almost complete sampling of the region, were analyzed using sequences from the internal transcribed spacer region (ITS) and the plastid trnQ-rps16 intergenic spacer. Phylogenetic relationships were inferred using Bayesian inference, maximum likelihood and statistical parsimony networking. Incongruence between datasets was assessed with statistical tests and displayed by split networks. Biogeographic inferences incorporating information from both markers (despite low resolution in some parts of the trees) and all incongruent taxa were accomplished with a novel combination of methods, using trees generated with the taxon duplication approach as input for Bayesian binary MCMC (BBM) analysis as implemented in RASP. Nuclear and chloroplast markers support a clade which comprises the majority of Iberian and Macaronesian species and consists of three subclades. Analyses of the substantial incongruence observed among markers indicate reticulate evolution and suggest that Scrophularia species diversity in this region is largely attributable to hybridization; a combination of both polyploidy and dysploidy in the karyotypic evolution of Western Mediterranean Scrophularia taxa is proposed. Our results provide support for an ancient hybridization event between two widespread lineages, which resulted in an allopolyploid ancestor of the Iberian - Macaronesian group with 2n=58 chromosomes. The ancestor then diverged into the three main lineages present in the Iberian Peninsula, Northern Africa and Macaronesia today. Subsequent interspecific hybridizations at different ploidy levels additionally generated new species. Presumably, hybridization and diversification within the genus in the Western Mediterranean have not been restricted to one particular event, but occurred repeatedly. It can be assumed that the topographical complexity found in the Iberian Peninsula has promoted diversification and hybrid speciation processes in Scrophularia, and that isolation in glacial refugia has preserved recent and ancient lineages. For the Macaronesian taxa, biogeographic analyses support several origins, by colonizations from at least four distinct lineages.
AU  - Scheunert, Agnes
AU  - Heubl, Günther
DO  - 10.1016/j.ympev.2013.09.023
KW  - Scrophulariaceae
LA  - eng
PY  - 2014
SN  - 1055-7903
SP  - 296-313
ST  - Diversification of Scrophularia (Scrophulariaceae) in the Western Mediterranean and Macaronesia - phylogenetic relationships, reticulate evolution and biogeographic patterns
T2  - Molecular phylogenetics and evolution
TI  - Diversification of Scrophularia (Scrophulariaceae) in the Western Mediterranean and Macaronesia - phylogenetic relationships, reticulate evolution and biogeographic patterns
VL  - 70
ID  - 1058
ER  - 


TY  - JOUR
AU  - Scheunert, Agnes
AU  - Heubl, Günther
DO  - 10.1007/s00606-010-0369-z M4 - Citavi
IS  - 1-2
KW  - Scrophulariaceae
N1  - PII: 369 PII: 369 PII: 369 PII: 369
PY  - 2011
SN  - 0378-2697
SP  - 69-89
ST  - Phylogenetic relationships among New World Scrophularia L. (Scrophulariaceae): new insights inferred from DNA sequence data
T2  - Plant Systematics and Evolution
TI  - Phylogenetic relationships among New World Scrophularia L. (Scrophulariaceae): new insights inferred from DNA sequence data
VL  - 291
ID  - 2026
ER  - 


TY  - JOUR
AU  - P.Kornhall
AU  - N.Heidari
AU  - Bremer, Birgitta
KW  - Scrophulariaceae
PY  - 2001
SN  - 0378-2697
SP  - 199-218
ST  - Selagineae and Manuleeae, two tribes or one? Phylogenetic studies in the Scrophulariaceae
T2  - Plant Systematics and Evolution
TI  - Selagineae and Manuleeae, two tribes or one? Phylogenetic studies in the Scrophulariaceae
VL  - 228
ID  - 4268
ER  - 


TY  - JOUR
AU  - Oxelman, Bengt
AU  - Kornhall, Per
AU  - Olmstead, Richard G.
AU  - Bremer, Birgitta
IS  - 2
KW  - Scrophulariaceae
PY  - 2005
SN  - 00400262
SP  - 411-425
ST  - Further disintegration of Scrophulariaceae
T2  - Taxon
TI  - Further disintegration of Scrophulariaceae
VL  - 54
ID  - 3670
ER  - 


TY  - JOUR
AB  - The mixed vertebrate-insect pollination system is rare in Holarctic plants. Phylogenetic relationships of 116 Scrophularia taxa were investigated based on two plastid (ndhF and trnL-trnF) and one nuclear (ITS) DNA regions. A wider time-calibrated analysis of ndhF sequences of the Lamiales revealed that Scrophularia diverged as early as in the Miocene (<22 Ma). Results of maximum-likelihood optimizations supported wasp pollination as the ancestral pollination system from which other systems derived (hoverfly, mixed vertebrate-insect and bird systems). Four origins for a mixed vertebrate-insect (MVI) pollination system were inferred, in which two western Mediterranean species (S. sambucifolia and S. grandiflora) and two island species (the Tirrenian S. trifoliata and the Canarian S. calliantha) were involved. S. calliantha is the only species in which a more complex MVI system, including pollination by the lizard Gallotia stehlini, has evolved. In addition, bird (hummingbird) floral traits found in the New Mexican S. macrantha appear to have been independently acquired. In contrast, we failed to find evidence for an ancient role of hummingbirds in the evolution of European Scrophularia. Indeed, paleontological data revealed that extinction of European hummingbirds (30-32 Ma) occurred earlier than the divergence of European MVI lineages of Scrophularia. In conclusion, our results showed that a role of birds in pollination of Scrophularia may not have been effective in the Miocene-Pliocene, but bird pollination that shows its origin in the Pliocene-Pleistocene is still operating independently in different islands and continents.
AD  - Área de Botánica, Facultad de Ciencias, Universidad de Extremadura, Avda. de Elvas s.n., 06006 Badajoz, Spain. M4 - Citavi
AU  - Navarro-Pérez, María L.
AU  - López, Josefa
AU  - Fernández-Mazuecos, Mario
AU  - Rodríguez-Riaño, Tomás
AU  - Vargas, Pablo
AU  - Ortega-Olivencia, Ana
DO  - 10.1016/j.ympev.2013.05.027
IS  - 1
KW  - Scrophulariaceae
LA  - eng
N1  - Journal Article Research Support, Non-U.S. Gov't Journal Article Research Support, Non-U.S. Gov't Journal Article Research Support, Non-U.S. Gov't Journal Article Research Support, Non-U.S. Gov't
PY  - 2013
SN  - 1055-7903
SP  - 239-254
ST  - The role of birds and insects in pollination shifts of Scrophularia (Scrophulariaceae)
T2  - Molecular phylogenetics and evolution
TI  - The role of birds and insects in pollination shifts of Scrophularia (Scrophulariaceae)
VL  - 69
ID  - 1326
ER  - 


TY  - JOUR
AU  - Mosyakin, Sergei L.
AU  - Tsymbalyuk, Zoya M.
DO  - 10.3372/wi.45.45207 M4 - Citavi
IS  - 2
KW  - Scrophulariaceae
PY  - 2015
SN  - 0511-9618
SP  - 209-222
ST  - Pollen morphology of tribes Aptosimeae and Myoporeae supports the phylogenetic pattern in early-branching Scrophulariaceae revealed by molecular studies
T2  - Willdenowia
TI  - Pollen morphology of tribes Aptosimeae and Myoporeae supports the phylogenetic pattern in early-branching Scrophulariaceae revealed by molecular studies
VL  - 45
ID  - 639
ER  - 


TY  - JOUR
AU  - Kornhall, Per
KW  - Scrophulariaceae
PY  - 2004
SP  - 453-467
ST  - New circumscription of the tribe Limoselleae (Scrophulariaceae) that includes the taxa of the tribe Manuleeae
T2  - Botanical Journal of the Linnean Society
TI  - New circumscription of the tribe Limoselleae (Scrophulariaceae) that includes the taxa of the tribe Manuleeae
VL  - 146
ID  - 3820
ER  - 


TY  - JOUR
AU  - Karaveliogullari, Faik Ahmet
AU  - Aytaç, Zeki
IS  - 1
KW  - Scrophulariaceae
PY  - 2008
SP  - 9-32
ST  - Revision of the Genus Verbascum L. (Group A) in Turkey
T2  - Botany Research Journal
TI  - Revision of the Genus Verbascum L. (Group A) in Turkey
VL  - 1
ID  - 2849
ER  - 


TY  - JOUR
AU  - Ghahremaninejad, Farrokh
AU  - Riahi, Mehrshid
AU  - Babaei, Melina
AU  - Attar, Farideh
AU  - Behçet, Lütfi
AU  - Sonboli, Ali
DO  - 10.1071/BT14159 M4 - Citavi
IS  - 8
KW  - Scrophulariaceae
N1  - PII: BT14159 PII: BT14159 PII: BT14159 PII: BT14159
PY  - 2014
SN  - 0067-1924
SP  - 638-646
ST  - Monophyly of Verbascum (Scrophularieae : Scrophulariaceae): evidence from nuclear and plastid phylogenetic analyses
T2  - Australian Journal of Botany
TI  - Monophyly of Verbascum (Scrophularieae : Scrophulariaceae): evidence from nuclear and plastid phylogenetic analyses
VL  - 62
ID  - 909
ER  - 


TY  - JOUR
AB  - The taxa of the Leucophyllum pringlei clade were used to understand the influence of the Neogene orogenesis and the Quaternary climate cycles on the diversification of the flora of the xeric regions of North America. This clade includes the five southernmost species of the genus: L. ambiguum, L. flyrii, L. pruinosum and L. ultramonticola, which are distributed throughout the Chihuahuan Desert north of the Trans-Mexican Volcanic Belt, and L. pringlei in the region of Tehuacán-Cuicatlán south of this mountain range. Here we test whether these species diverged during the pluvial periods of the Pleistocene, and whether L. pringlei diverged earlier from the other species during the uplift of the Trans-Mexican Volcanic Belt. Using three plastid regions (psbA-trnH, psbK-psbI, trnL-F) and a nuclear (ITS) marker, phylogenetic analyses were carried out, along with a reconstruction of their ancestral area. Trees retrieved the five species in a monophyletic group with the most recent common ancestor distributed in the Sinaloan dry forest during the Late Miocene (8.08Ma), from where it dispersed to the Chihuahuan Desert during the Late Miocene (6.35Ma). The secondary uplift of the Sierra Madre Occidental during the Late Miocene to Early Pliocene influenced a vicariance event. Divergence between L. pringlei and the species from north of the Trans-Mexican Volcanic Belt occurred during the second volcanic episode in the Late Miocene (7.5-3Ma). The most recent common ancestor of L. ambiguum, L. pruinosum and L. ultramonticola was widely distributed in the southern part of the Chihuahuan Desert during the Early to Late Pliocene (3.50Ma). The diversification of these three species occurred in the Middle Pleistocene (0.9Ma) during the pluvial and inter-pluvial cycles.
AD  - Biología Evolutiva, Instituto de Ecología, A.C., Carretera antigua a Coatepec 351, El Haya, 91070 Xalapa, Veracruz, Mexico. Electronic address:. Biología Evolutiva, Instituto de Ecología, A.C., Carretera antigua a Coatepec 351, El Haya, 91070 Xalapa, Veracruz, Mexico. M4 - Citavi
AU  - Gándara, Etelvina
AU  - Sosa, Victoria
DO  - 10.1016/j.ympev.2014.02.027
KW  - Scrophulariaceae
LA  - eng
N1  - Journal Article Research Support, Non-U.S. Gov't Journal Article Research Support, Non-U.S. Gov't Journal Article Research Support, Non-U.S. Gov't Journal Article Research Support, Non-U.S. Gov't
PY  - 2014
SN  - 1055-7903
SP  - 93-101
ST  - Spatio-temporal evolution of Leucophyllum pringlei and allies (Scrophulariaceae): a group endemic to North American xeric regions
T2  - Molecular phylogenetics and evolution
TI  - Spatio-temporal evolution of Leucophyllum pringlei and allies (Scrophulariaceae): a group endemic to North American xeric regions
VL  - 76
ID  - 903
ER  - 


TY  - JOUR
AU  - Gándara, Etelvina
AU  - Sosa, Victoria
KW  - Scrophulariaceae
PY  - 2013
SP  - 508-518
ST  - Testing the monophyly and position of the North American shrubby desert genus Leucophyllum (Scrophulariaceae: Leucophylleae)
T2  - Botanical Journal of the Linnean Society
TI  - Testing the monophyly and position of the North American shrubby desert genus Leucophyllum (Scrophulariaceae: Leucophylleae)
VL  - 171
ID  - 1225
ER  - 


TY  - JOUR
AU  - Archibald, Jenny K.
AU  - Mort, Mark E.
AU  - Wolfe, Andrea D.
IS  - 1
KW  - Scrophulariaceae
PY  - 2005
SN  - 03636445
SP  - 196-215
ST  - Phylogenetic Relationships within Zaluzianskya (Scrophulariaceae s. s., tribe Manuleeae): Classification Based on DNA Sequences from Multiple Genomes and Implications for Character Evolution and Biogeography
T2  - Systematic Botany
TI  - Phylogenetic Relationships within Zaluzianskya (Scrophulariaceae s. s., tribe Manuleeae): Classification Based on DNA Sequences from Multiple Genomes and Implications for Character Evolution and Biogeography
VL  - 30
ID  - 3552
ER  - 


TY  - JOUR
AU  - Barringer, Kerry
IS  - 3
KW  - Schlegeliaceae
PY  - 2004
SN  - 0007-196X
SP  - 213-237
ST  - A revision of Gibsoniothamnus L.O.Williams (Schlegeliaceae)
T2  - Brittonia
TI  - A revision of Gibsoniothamnus L.O.Williams (Schlegeliaceae)
VL  - 56
ID  - 3750
ER  - 


TY  - JOUR
AU  - Wikström, Niklas
AU  - Kenrick, Paul
AU  - Vogel, Johannes C.
KW  - Schizaeaceae
PY  - 2002
SP  - 35-50
ST  - Schizaeaceae: a phylogenetic approach
T2  - Rev. Palaobotany and Palynology
TI  - Schizaeaceae: a phylogenetic approach
VL  - 119
ID  - 4190
ER  - 


TY  - JOUR
AB  - International Journal of Plant Sciences 2006.167:539-550
AU  - Liu, Zhong
AU  - Hao, Gang
AU  - Luo, Yi‐bo
AU  - Thien, Leonard B.
AU  - Rosso, Samuel W.
AU  - Lu, An-Ming
AU  - Chen, Zhi‐Duan
IS  - 3
KW  - Schisandraceae
PY  - 2006
SN  - 1058-5893
SP  - 539-550
ST  - Phylogeny and Androecial Evolution in Schisandraceae, Inferred from Sequences of Nuclear Ribosomal DNA ITS and Chloroplast DNA trnL‐F Regions
T2  - International Journal of Plant Sciences
TI  - Phylogeny and Androecial Evolution in Schisandraceae, Inferred from Sequences of Nuclear Ribosomal DNA ITS and Chloroplast DNA trnL‐F Regions
VL  - 167
ID  - 3435
ER  - 


TY  - JOUR
AU  - Hao, Gang
AU  - Saunders, Richard M. K.
AU  - Chye, Mee-Len
KW  - Schisandraceae
PY  - 2000
SN  - 0378-2697
SP  - 81-90
ST  - A phylogenetic analysis of the Illiciaceae based on sequences of internal transcribed spacers (ITS) of nuclear ribosomal DNA
T2  - Plant Systematics and Evolution
TI  - A phylogenetic analysis of the Illiciaceae based on sequences of internal transcribed spacers (ITS) of nuclear ribosomal DNA
VL  - 223
ID  - 4340
ER  - 


TY  - JOUR
AU  - Hao, Gang
DO  - 10.1006/bojl.2000.0420 M4 - Citavi
IS  - 4
KW  - Schisandraceae
N1  - PII: S0024-4074(00)90420-0 PII: S0024-4074(00)90420-0 PII: S0024-4074(00)90420-0 PII: S0024-4074(00)90420-0
PY  - 2001
SP  - 401-411
ST  - A phylogenetic analysis of the Schisandraceae based on morphology and nuclear ribosomal ITS sequences
T2  - Botanical Journal of the Linnean Society
TI  - A phylogenetic analysis of the Schisandraceae based on morphology and nuclear ribosomal ITS sequences
VL  - 135
ID  - 4233
ER  - 


TY  - JOUR
AU  - Denk, Thomas
AU  - Oh, I. C.
DO  - 10.1007/s00606-005-0327-3 M4 - Citavi
IS  - 1-4
KW  - Schisandraceae
N1  - PII: 327 PII: 327 PII: 327 PII: 327
PY  - 2005
SN  - 0378-2697
SP  - 113-145
ST  - Phylogeny of Schisandraceae based on morphological data: evidence from modern plants and the fossil record
T2  - Plant Systematics and Evolution
TI  - Phylogeny of Schisandraceae based on morphological data: evidence from modern plants and the fossil record
VL  - 256
ID  - 3578
ER  - 


TY  - JOUR
AB  - UNLABELLED PREMISE OF THE STUDY We sought novel evolutionary insights for the highly diverse Saxifragales by constructing a large phylogenetic tree encompassing 36.8% of the species-level biodiversity. • METHODS We built a phylogenetic tree for 909 species of Saxifragales and used this hypothesis to examine character evolution for annual or perennial habit, woody or herbaceous habit, ovary position, petal number, carpel number, and stamen to petal ratio. We employed likelihood approaches to investigate the effect of habit and life history on speciation and extinction within this clade. • KEY RESULTS Two major shifts occurred from a woody ancestor to the herbaceous habit, with multiple secondary changes from herbaceous to woody. Transitions among superior, subinferior, and inferior ovaries appear equiprobable. A major increase in petal number is correlated with a large increase in carpel number; these increases have co-occurred multiple times in Crassulaceae. Perennial or woody lineages have higher rates of speciation than annual or herbaceous ones, but higher probabilities of extinction offset these differences. Hence, net diversification rates are highest for annual, herbaceous lineages and lowest for woody perennials. The shift from annuality to perenniality in herbaceous taxa is frequent. Conversely, woody perennial lineages to woody annual transitions are infrequent; if they occur, the woody annual state is left immediately. • CONCLUSIONS The large tree provides new insights into character evolution that are not obvious with smaller trees. Our results indicate that in some cases the evolution of angiosperms might be conditioned by constraints that have been so far overlooked.
AD  - Department of Biology, University of Florida, Gainesville, FL 32611, USA. M4 - Citavi
AU  - Soltis, Douglas E.
AU  - Mort, Mark E.
AU  - Latvis, Maribeth
AU  - Mavrodiev, Evgeny V.
AU  - O'Meara, Brian C.
AU  - Soltis, Pamela S.
AU  - Burleigh, John Gordon
AU  - Rubio de Casas, Rafael
DO  - 10.3732/ajb.1300044
IS  - 5
KW  - Saxifragales
LA  - eng
N1  - Journal Article Research Support, Non-U.S. Gov't Research Support, U.S. Gov't, P.H.S. Journal Article Research Support, Non-U.S. Gov't Research Support, U.S. Gov't, P.H.S. Journal Article Research Support, Non-U.S. Gov't Research Support, U.S. Gov't, P.H.S. Journal Article Research Support, Non-U.S. Gov't Research Support, U.S. Gov't, P.H.S.
PY  - 2013
SN  - 0002-9122
SP  - 916-929
ST  - Phylogenetic relationships and character evolution analysis of Saxifragales using a supermatrix approach
T2  - American journal of botany
TI  - Phylogenetic relationships and character evolution analysis of Saxifragales using a supermatrix approach
VL  - 100
ID  - 1391
ER  - 


TY  - JOUR
AB  - Despite the prior use of approximately 9000 bp, deep-level relationships within the angiosperm clade, Saxifragales remain enigmatic, due to an ancient, rapid radiation (89.5 to 110 Ma based on the fossil record). To resolve these deep relationships, we constructed several new data sets: (1) 16 genes representing the three genomic compartments within plant cells (2 nuclear, 10 plastid, 4 mitochondrial; aligned, analyzed length = 21,460 bp) for 28 taxa; (2) the entire plastid inverted repeat (IR; 26,625 bp) for 17 taxa; (3) "total evidence" (50,845 bp) for both 17 and 28 taxa (the latter missing the IR). Bayesian and ML methods yielded identical topologies across partitions with most clades receiving high posterior probability (pp = 1.0) and bootstrap (95% to 100%) values, suggesting that with sufficient data, rapid radiations can be resolved. In contrast, parsimony analyses of different partitions yielded conflicting topologies, particularly with respect to the placement of Paeoniaceae, a clade characterized by a long branch. In agreement with published simulations, the addition of characters increased bootstrap support for the putatively erroneous placement of Paeoniaceae. Although having far fewer parsimony-informative sites, slowly evolving plastid genes provided higher resolution and support for deep-level relationships than rapidly evolving plastid genes, yielding a topology close to the Bayesian and ML total evidence tree. The plastid IR region may be an ideal source of slowly evolving genes for resolution of deep-level angiosperm divergences that date to 90 My or more. Rapidly evolving genes provided support for tip relationships not recovered with slowly evolving genes, indicating some complementarity. Age estimates using penalized likelihood with and without age constraints for the 28-taxon, total evidence data set are comparable to fossil dates, whereas estimates based on the 17-taxon data are much older than implied by the fossil record. Hence, sufficient taxon density, and not simply numerous base pairs, is important in reliably estimating ages. Age estimates indicate that the early diversification of Saxifragales occurred rapidly, over a time span as short as 6 million years. Between 25,000 and 50,000 bp were needed to resolve this radiation with high support values. Extrapolating from Saxifragales, a similar number of base pairs may be needed to resolve the many other deep-level radiations of comparable age in angiosperms.
AD  - South China Botanical Garden, the Chinese Academy of Sciences, Guangzhou, China. M4 - Citavi
AU  - Jian, Shu-Guang
AU  - Soltis, Pamela S.
AU  - Gitzendanner, Matthew A.
AU  - Moore, Michael J.
AU  - Li, Rui-Qi
AU  - Hendry, Tory A.
AU  - Qiu, Yin-Long
AU  - Dhingra, Amit
AU  - Bell, Charles D.
AU  - Soltis, Douglas E.
DO  - 10.1080/10635150801888871
IS  - 1
KW  - Saxifragales
LA  - eng
N1  - Journal Article Research Support, U.S. Gov't, Non-P.H.S. Journal Article Research Support, U.S. Gov't, Non-P.H.S. Journal Article Research Support, U.S. Gov't, Non-P.H.S. Journal Article Research Support, U.S. Gov't, Non-P.H.S.
PY  - 2008
SP  - 38-57
ST  - Resolving an ancient, rapid radiation in Saxifragales
T2  - Systematic Biology
TI  - Resolving an ancient, rapid radiation in Saxifragales
VL  - 57
ID  - 2845
ER  - 


TY  - JOUR
AU  - Zhu, Wei-dong
AU  - Nie, Ze-long
AU  - Wen, Jun
AU  - Sun, Hang
KW  - Saxifragaceae
PY  - 2013
SP  - 377-394
ST  - Molecular phylogeny and biogeography of Astilbe (Saxifragaceae) in Asia and eastern North America
T2  - Botanical Journal of the Linnean Society
TI  - Molecular phylogeny and biogeography of Astilbe (Saxifragaceae) in Asia and eastern North America
VL  - 171
ID  - 1452
ER  - 


TY  - JOUR
AU  - Zhang, De-Jun
AU  - Chen, Sheng-Yun
AU  - Gao, Qing-Bo
IS  - 5
KW  - Saxifragaceae
PY  - 2008
SN  - 16744918
SP  - 667-675
ST  - Circumscription and phylogeny of Saxifraga sect. Ciliatae: Evidence from nrDNA ITS sequences
T2  - Journal of Systematics and Evolution
TI  - Circumscription and phylogeny of Saxifraga sect. Ciliatae: Evidence from nrDNA ITS sequences
VL  - 46
ID  - 3034
ER  - 


TY  - JOUR
AB  - Saniculiphyllum, a monotypic genus distributed in Southwest China, was thought to be extinct before our recent rediscovery. The taxonomic position of this genus has been enigmatic ever since its publication. It was originally treated as the only member of a distinct tribe Saniculiphylleae in the family Saxifragaceae. Some proposed a new family, Saniculophyllaceae, to accommodate this genus, although its affinities are clearly with members of Saxifragaceae. Here we analyzed six DNA regions, the nuclear ribosomal ITS and 26S rDNA and the plastid rbcL, matK, trnL-trnF, psbA-trnH genes, spacers, and intron to explore the phylogenetic position of Saniculiphyllum within Saxifragaceae. The combined nuclear and chloroplast dataset includes 63 ingroup species, representing all genera but Hieronymusia in the family. Results from likelihood, parsimony and Bayesian phylogenetic methods corroborate earlier results. Two clades of Saxifragaceae, the Heucheroid and Saxifragoid clades, were recovered. The topologies obtained from different analyses confirm the placement of Saniculiphyllum in Saxifragaceae, but our analyses reveal that Saniculiphyllum is embedded within the large Heucheroid clade. However, the closest relatives of Saniculiphyllum within the Heucheroid clade remain unclear. Combined with morphological data, our results suggest that Saniculiphyllum should best be regarded as a highly distinctive lineage within the Heucheroid clade of Saxifragaceae. Morphological novelties and conservation status of Saniculiphyllum are also presented.
AD  - Key Laboratory of Biodiversity and Biogeography, Kunming Institute of Botany, Chinese Academy of Sciences, No. 132, Lanhei Road, Kunming, Yunnan 650201, PR China. M4 - Citavi
AU  - Xiang, Chun-Lei
AU  - Gitzendanner, Matthew A.
AU  - Soltis, Douglas E.
AU  - Peng, Hua
AU  - Lei, Li-Gong
DO  - 10.1016/j.ympev.2012.04.010
IS  - 2
KW  - Saxifragaceae
LA  - eng
N1  - Journal Article Research Support, Non-U.S. Gov't Journal Article Research Support, Non-U.S. Gov't Journal Article Research Support, Non-U.S. Gov't Journal Article Research Support, Non-U.S. Gov't
PY  - 2012
SN  - 1055-7903
SP  - 357-367
ST  - Phylogenetic placement of the enigmatic and critically endangered genus Saniculiphyllum (Saxifragaceae) inferred from combined analysis of plastid and nuclear DNA sequences
T2  - Molecular phylogenetics and evolution
TI  - Phylogenetic placement of the enigmatic and critically endangered genus Saniculiphyllum (Saxifragaceae) inferred from combined analysis of plastid and nuclear DNA sequences
VL  - 64
ID  - 1769
ER  - 


TY  - JOUR
AU  - Winkler, Manuela
AU  - Tribsch, Andreas
AU  - Schneeweiss, Gerald M.
AU  - Brodbeck, Sabine
AU  - Gugerli, Felix
AU  - Holderegger, Rolf
AU  - Schönswetter, Peter
KW  - Saxifragaceae
PY  - 2013
SP  - 622-636
ST  - Strong nuclear differentiation contrasts with widespread sharing of plastid DNA haplotypes across taxa in European purple saxifrages (Saxifraga section Porphyrion subsection Oppositifoliae)
T2  - Botanical Journal of the Linnean Society
TI  - Strong nuclear differentiation contrasts with widespread sharing of plastid DNA haplotypes across taxa in European purple saxifrages (Saxifraga section Porphyrion subsection Oppositifoliae)
VL  - 173
ID  - 1432
ER  - 


TY  - JOUR
AU  - Vargas, Pablo
AU  - Morton, Cynthia M.
AU  - Jury, Stephen L.
IS  - 5
KW  - Saxifragaceae
PY  - 1999
SN  - 0002-9122
SP  - 724-734
ST  - Biogeographic Patterns In Mediterranean And Macaronesian Species Of Saxifraga (Saxifragaceae) Inferred From Phylogenetic Analyses Of ITS Sequences
T2  - American journal of botany
TI  - Biogeographic Patterns In Mediterranean And Macaronesian Species Of Saxifraga (Saxifragaceae) Inferred From Phylogenetic Analyses Of ITS Sequences
VL  - 86
ID  - 4437
ER  - 


TY  - JOUR
AD  - Martin Luther University Halle-Wittenberg, Institute of Biology, Geobotany and Botanical Garden, Neuwerk 21, 06108 Halle, Germany;, Martin Luther University Halle-Wittenberg, Institute of Biology, Geobotany and Botanical Garden, Neuwerk 21, 06108 Halle, Germany;, Philipps-University Marburg, Department of Geography, Deutschhausstraße 10, 35032 Marburg, Germany University of Leipzig, Institute of Biology, Department of Molecular Evolution and Plant Systematics, Herbarium LZ, Johannisallee 21–23, 04103 Leipzig, Germany, German Centre for Integrative Biodiversity Research (iDiv) Halle-Jena-Leipzig, Deutscher Platz 5e, 04103 Leipzig, Germany University of Leipzig, Institute of Biology, Department of Molecular Evolution and Plant Systematics, Herbarium LZ, Johannisallee 21–23, 04103 Leipzig, Germany Martin Luther University Halle-Wittenberg, Institute of Biology, Geobotany and Botanical Garden, Neuwerk 21, 06108 Halle, Germany;, M4 - Citavi
AU  - Tkach, Natalia
AU  - Röser, Martin
AU  - Miehe, Georg
AU  - Muellner-Riehl, Alexandra N.
AU  - Ebersbach, Jana
AU  - Favre, Adrien
AU  - Hoffmann, Matthias H.
DO  - 10.12705/646.4
IS  - 6
KW  - Saxifragaceae
PY  - 2015
SN  - 00400262
SP  - 1159-1187
ST  - Molecular phylogenetics, morphology and a revised classification of the complex genus Saxifraga (Saxifragaceae)
T2  - Taxon
TI  - Molecular phylogenetics, morphology and a revised classification of the complex genus Saxifraga (Saxifragaceae)
VL  - 64
ID  - 754
ER  - 


TY  - JOUR
AU  - Tkach, Natalia
AU  - Röser, Martin
AU  - Hoffmann, Matthias H.
KW  - Saxifragaceae
PY  - 2015
SP  - 47-66
ST  - Molecular phylogenetics, character evolution and systematics of the genus Micranthes (Saxifragaceae)
T2  - Botanical Journal of the Linnean Society
TI  - Molecular phylogenetics, character evolution and systematics of the genus Micranthes (Saxifragaceae)
VL  - 178
ID  - 753
ER  - 


TY  - JOUR
AU  - Soltis, Douglas E.
AU  - Tago-Nakazawa, Miyuki
AU  - Xiang, Qiu-Yun
AU  - Kawano, Shoichi
AU  - Murata, Jin
AU  - Wakabayashi, Michio
AU  - Hibsch-Jetter, Carola
IS  - 5
KW  - Saxifragaceae
PY  - 2001
SN  - 0002-9122
SP  - 883-893
ST  - Phylogenetic Relationships And Evolution In Chrysosplenium (Saxifragaceae) Based On Matk Sequence Data
T2  - American journal of botany
TI  - Phylogenetic Relationships And Evolution In Chrysosplenium (Saxifragaceae) Based On Matk Sequence Data
VL  - 88
ID  - 4288
ER  - 


TY  - JOUR
AB  - The widespread pattern of parallel flower evolution as an adaptation for particular pollinator agents, known as "pollination syndromes", has long drawn attention from evolutionary biologists. Here, we report parallel evolution of saucer-shaped flowers and an associated unusual pollination system within the lineage Heucherina, a group of saxifragaceous genera. Field observations reveal that 18 of 28 plant species studied are pollinated almost exclusively by fungus gnats (Mycetophilidae). Among the 18 species with a fungus-gnat pollination system, 13 have characteristic saucer-shaped flowers and are pollinated mainly by several unspecialized mycetophilid genera with short mouthparts. We performed phylogenetic analyses using nucleotide sequences of external and internal transcribed spacers of nuclear ribosomal DNA and reconstructed ancestral floral morphologies with an establishment of the model of floral character evolution under a maximum-likelihood framework. Our analysis indicates that there is significant directionality in the evolutionary shifts of floral forms in the Heucherina. The inferred phylogeny further supports four origins of saucer-shaped flowers, which is shared among 14 species that are traditionally classified into the genus Mitella. In addition, our analysis indicates the extensive polyphyly of genus Mitella, as also suggested previously. The results suggest that the flower-visiting fungus gnats have caused convergent selection for the saucer-shaped flower repeatedly evolved within Heucherina.
AD  - Graduate School of Human and Environmental Studies, Kyoto University, Yoshida Nihonmatsu-cho, Sakyo-ku, Kyoto 606-8501, Japan. M4 - Citavi
AU  - Okuyama, Yudai
AU  - Pellmyr, Olle
AU  - Kato, Makoto
DO  - 10.1016/j.ympev.2007.09.020
IS  - 2
KW  - Saxifragaceae
LA  - eng
N1  - Journal Article Research Support, Non-U.S. Gov't Research Support, U.S. Gov't, Non-P.H.S. Journal Article Research Support, Non-U.S. Gov't Research Support, U.S. Gov't, Non-P.H.S. Journal Article Research Support, Non-U.S. Gov't Research Support, U.S. Gov't, Non-P.H.S. Journal Article Research Support, Non-U.S. Gov't Research Support, U.S. Gov't, Non-P.H.S.
PY  - 2008
SN  - 1055-7903
SP  - 560-575
ST  - Parallel floral adaptations to pollination by fungus gnats within the genus Mitella (Saxifragaceae)
T2  - Molecular phylogenetics and evolution
TI  - Parallel floral adaptations to pollination by fungus gnats within the genus Mitella (Saxifragaceae)
VL  - 46
ID  - 2918
ER  - 


TY  - JOUR
AU  - Mas De Xaxars, Gemma
AU  - García-Fernández, Alfredo
AU  - Barnola, Pere
AU  - Martín, Joan
AU  - MercadÉ, Arnau
AU  - Vallès, Joan
AU  - Vargas, Pablo
AU  - Vigo, Jospe
AU  - Garnatje, Teresa
DO  - 10.1111/jse.12105 M4 - Citavi
IS  - 1
KW  - Saxifragaceae
PY  - 2015
SN  - 16744918
SP  - 53-62
ST  - Phylogenetic and cytogenetic studies reveal hybrid speciation in Saxifraga subsect. Triplinervium (Saxifragaceae)
T2  - Journal of Systematics and Evolution
TI  - Phylogenetic and cytogenetic studies reveal hybrid speciation in Saxifraga subsect. Triplinervium (Saxifragaceae)
VL  - 53
ID  - 613
ER  - 


TY  - JOUR
AD  - Key Laboratory of Adaption and Evolution of Plateau Biota, Northwest Institute of Plateau Biology, Chinese Academy of Sciences, Xining 810001, Qinghai, P. R. China Key Laboratory of Adaption and Evolution of Plateau Biota, Northwest Institute of Plateau Biology, Chinese Academy of Sciences, Xining 810001, Qinghai, P. R. China, University of Chinese Academy of Sciences, Beijing 100039, P. R. China Department of Biology, University of Leicester, Leicester LE1 7RH, U.K. Key Laboratory of Plant Resources and Sustainable Utilization, South China Botanical Garden, Chinese Academy of Sciences, Guangzhou 510650, Guangdong, P. R. China Key Laboratory of Adaption and Evolution of Plateau Biota, Northwest Institute of Plateau Biology, Chinese Academy of Sciences, Xining 810001, Qinghai, P. R. China;, M4 - Citavi
AU  - Gao, Qing-Bo
AU  - Li, Yin-Hu
AU  - Gornall, Richard J.
AU  - Zhang, Zhuo-Xin
AU  - Zhang, Fa-Qi
AU  - Xing, Rui
AU  - Fu, Peng-Cheng
AU  - Wang, Jiu-Li
AU  - Liu, Hai-Rui
AU  - Tian, Zun-Zhe
AU  - Chen, Shi-Long
DO  - 10.12705/644.3
IS  - 4
KW  - Saxifragaceae
PY  - 2015
SN  - 00400262
SP  - 703-713
